# Discovery, diversity and functional associations of crAss-like phages in human gut metagenomes from four Dutch cohorts

**DOI:** 10.1101/2021.06.01.446427

**Authors:** Anastasia Gulyaeva, Sanzhima Garmaeva, Renate A.A.A. Ruigrok, Daoming Wang, Niels P. Riksen, Mihai G. Netea, Cisca Wijmenga, Rinse K. Weersma, Jingyuan Fu, Arnau Vich Vila, Alexander Kurilshikov, Alexandra Zhernakova

**Affiliations:** Department of Genetics, University of Groningen, University Medical Center Groningen, the Netherlands; Department of Gastroenterology and Hepatology, University Medical Center Groningen, the Netherlands; Department of Internal Medicine, Radboud University Medical Center, Nijmegen, the Netherlands; Department of Pediatrics, University of Groningen, University Medical Center Groningen, the Netherlands

**Keywords:** CrAss-like phages, Human gut virome, Metagenomics, Population cohorts, IBD

## Abstract

The crAss-like phages are a diverse group of related viruses that includes one of the most abundant viruses of the human gut. To explore their diversity and functional role in human population and clinical cohorts, we analyzed gut metagenomic data collected from more than 2000 individuals from the Netherlands. We discovered 125 novel species-level and 32 novel genus-level clusters of crAss-like phages, all belonging to five previously recognized groups associated with the human gut. Analysis of their genomic features revealed that closely related crAss-like phages can possess strikingly divergent regions responsible for transcription, presumably acquired through recombination. Prediction of crAss-like phage hosts pointed primarily to bacteria of the phylum Bacteroidetes, consistent with previous reports. Finally, we explored the temporal stability of crAss-like phages over a 4-year period and identified associations between the abundance of crAss-like phages and several human phenotypes, including depletion of crAss-like phages in inflammatory bowel disease patients.

**Highlights:**

- 125 tentative new species of crAss-like phages were discovered
- Closely related crAss-like phages often possess highly divergent transcription gene modules possibly acquired via recombination
- CrAss-like fraction of the human gut virome remains relatively stable over the period of 4 years
- Prevalence of crAss-like phages in the human gut is associated with several metabolic, dietary and health phenotypes
- Gut crAss-like phages are depleted in inflammatory bowel disease patients

## Introduction

Our planet harbors an enormous diversity of viruses. There are millions of distinct species of cellular organisms, and all can likely be infected by viruses. The diversity of hosts, high mutation rate and propensity to exchange genetic material with hosts and other viruses co-infecting a host cell all contribute to the unparalleled diversity of virus genomes. Remarkably, viruses employ a limited number of key conserved proteins responsible for genome replication and virion formation, allowing to comprehend and classify the global virome (Koonin et al., 2020).

The advent of metagenomics led to a rapid expansion of the known viral sequence space, although it is still far from saturation. Due to the medical importance of the human gut ecosystem, its virome has become a major focus of metagenomics research, resulting in the discovery of multiple novel viruses including the crAss-like phages. The first representative of these phages, the crAssphage, was discovered in 2014 and is believed to be one of the most abundant viruses in the human gut (Dutilh et al., 2014). This initial discovery was followed by the identification of five groups of crAss-like phages associated with the human gut – alpha/gamma, beta, delta, epsilon and zeta – as well as diverse crAss-like phages associated with other environments (Guerin et al., 2018; Yutin et al., 2021; Yutin et al., 2018).

The crAss-like phages are tailed phages with up to 192 kb circular (or terminally redundant) double-stranded DNA genomes. According to phylogenetic analysis of their conserved proteins, crAss-like phages form a monophyletic clade related to the prokaryotic virus order *Caudovirales*. There are three major functional regions that can be distinguished in the crAss-like phage genome: a replication gene module, a transcription gene module and a capsid gene module (Koonin and Yutin, 2020; Yutin *et al.*, 2018). Transcription of the three modules is initiated early (by a virion-packaged RNA polymerase (Holmfeldt et al., 2013; Shkoporov et al., 2018)), in the middle and late in the infection, respectively (Drobysheva et al., 2021; Shkoporov et al., 2021). The genomes of various crAss-like phages bear unique features, including reassignment of the stop codons TAG and TGA for amino acids and a high density of introns and inteins (Yutin *et al.*, 2021).

Numerous crAss-like phages have been predicted to infect bacteria of the phylum Bacteroidetes (Dutilh *et al.*, 2014; Yutin *et al.*, 2021), and accordingly, several crAss-like phages have been isolated in cultures of Bacteroidetes hosts or enriched in the presence of Bacteroidetes hosts (Fitzgerald et al., 2021; Guerin et al., 2021; Hryckowian et al., 2020; Shkoporov *et al.*, 2018), although there is also evidence that some crAss-like phages may infect hosts from different phyla (Yutin *et al.*, 2021). Apparently, crAss-like phages may coexist with their hosts relatively peacefully, as the same crAss-like phages were often recurrently detected in metagenomic samples of longitudinal study participants (Edwards et al., 2019; Shkoporov et al., 2019; Siranosian et al., 2020) and four crAss-like phages were shown to persist in the cultures of their hosts for a long time without a significant reduction in the host population (Guerin *et al.*, 2021; Shkoporov *et al.*, 2018; Shkoporov *et al.*, 2021). Interestingly, hosts were shown to develop resistance to crAss-like phage infection via the phase variation mechanism: inversions in the bacterial genome loci responsible for cell surface coating biosynthesis were linked to altered cell surface architecture and a reduced ability of the bacteria to absorb phage (Shkoporov *et al.*, 2021). The crAss-like phages are globally distributed in the human population, although the abundance of specific crAss-like phage taxa may vary depending on the geographic location (Camarillo-Guerrero et al., 2021; Edwards *et al.*, 2019; Guerin *et al.*, 2018). The abundance of certain crAss-like phages in the human gut metagenome have been shown to correlate with several health conditions, including advanced stages of HIV infection (Oude Munnink et al., 2014), inflammatory bowel disease (IBD) (Clooney et al., 2019), Parkinson’s disease, liver cirrhosis and ankylosing spondylitis (Tisza and Buck, 2020).

To further our understanding of the diversity, abundance and functional role of the crAss-like phages in the human gut microbial community, we identified and analyzed crAss-like phages present in 2291 metagenomic samples from four cohorts collected in the Netherlands. These include 1135 baseline and 338 follow-up samples from the population cohorts Lifelines-DEEP (LLD) and LLD follow-up (Chen et al., 2021; Tigchelaar et al., 2015; Zhernakova et al., 2016), 298 samples from the 300-Obesity cohort (300OB) (Ter Horst et al., 2020), and 520 samples from a cohort of patients with IBD (Imhann et al., 2019; Vich Vila et al., 2018). Analysis of these samples led to the discovery of crAss-like phage genomes prototyping 125 novel species. We predicted hosts of crAss-like phages and found associations between the abundance of crAss-like phages and structural variations in bacterial genomes. Finally, we characterized the temporal dynamics and stability of crAss-like phages in longitudinal samples and explored links between the abundance of crAss-like phages and human metabolic, environmental and health phenotypes.

## Results

### Assembly and identification of crAss-like phage genomes

To identify crAss-like phage genomes in our gut metagenomic data, we assembled contigs for 2291 metagenomic samples from four cohorts: 1135 LLD, 338 LLD follow-up, 298 300OB and 520 IBD samples. Contigs of 10 kilobases (kb) or longer were retained for further analysis. The number of these contigs ranged from 0 to 3036 per sample and was proportional to the number of sequencing reads available for each sample (Figure S1A-B).

To identify crAss-like phage genomes, we searched for contigs encoding three conserved capsid and genome-packaging proteins of crAss-like phages: terminase large subunit (TerL), portal protein and major capsid protein (MCP). To enable a high sensitivity search and account for the stop-codon reassignments observed in some crAss-like phages (Yutin *et al.*, 2021), we used both the protein sequences predicted under the standard genetic code and the full-length contig translations in six frames as search subjects. A contig was considered as a potential crAss-like phage genome fragment if either of the two types of search subjects was hit (HMMER, E-value < 0.001) by profiles (Yutin *et al.*, 2021) of the three crAss-like phage proteins.

To verify the validity of our approach, we first applied it to four publicly available control datasets: a database of diverse manually annotated viral genomes (Viral RefSeq 202) (Brister et al., 2015), a database of 249 crAss-like phage genomes (DB249) (Guerin *et al.*, 2018), a database of 673 crAss-like phage genomes (DB673) (Yutin *et al.*, 2021) and a database of 146 nucleotide sequences (DB146) recognized as crAss-like by the NCBI Taxonomy resource (Schoch et al., 2020). As expected, only a few Viral RefSeq genomes were recognized as crAss-like phages (Table S1): three established crAss-like phages (Dutilh *et al.*, 2014; Oude Munnink *et al.*, 2014; Shkoporov *et al.*, 2018), five bacteriophages of the fish pathogen *Flavobacterium psychrophilum* (phylum Bacteroidetes) and seven bacteriophages of the aquatic bacterium *Cellulophaga baltica* (phylum Bacteroidetes). The latter 12 bacteriophages were classified as belonging to the family *Podoviridae* at the time of discovery (Castillo and Middelboe, 2016; Holmfeldt *et al.*, 2013). Also as expected, almost all entries of the three crAss-specific databases (DB249, DB673 and DB146) were recognized as crAss-like phages. The only exceptions (Table S1) were 13 unrecognized entries from DB249 and DB673, all represented by partial genomes lacking the complete capsid gene module, and three unrecognized entries of DB146 representing two marine phages, Cellulophaga phage phi38:1 and phi40:1 (Holmfeldt *et al.*, 2013), that likely form a lineage divergent from known crAss-like phages.

Following successful validation of our crAss-like phage detection approach in the control datasets, we applied it to the contigs of 10 kb or longer, which were assembled for each of the individual metagenomic samples from our cohorts. The crAss-like contigs were detected in 43% of LLD, 58% of LLD follow-up, 61% of 300OB and 29% of IBD cohort samples, and their number ranged from one to six per sample. The correlation between the number of crAss-like contigs detected and the number of sequencing reads per sample was relatively weak (r_Spearman_ = 0.182, P-value = 1.47e-18; Figure S1C). In total, we detected 1556 crAss-like contigs. Notably, the sets of crAss-like contigs identified based on hits to individual protein queries and based on hits to contig translations in six frames were almost identical: 1553 of the 1556 crAss-like contigs detected were detected by both methods. The lengths of the crAss-like contigs ranged from 10 to 196 kb, mean length 59 kb. A considerable fraction of these contigs (255, 16%) had a 5’-end sequence of 10 or more nucleotides that perfectly matched the 3’-end sequence, tentatively indicating that these are fully sequenced circular or terminally redundant linear genomes (Table S2, Material S1).

In order to assign the identified contigs to species-level virus operational taxonomic units (vOTUs), the 1556 newly detected crAss-like contigs were pooled together with the DB249, DB673 and DB146 genomes and clustered under the 95% average nucleotide identity (ANI) over the 85% alignment fraction (AF) threshold (Roux et al., 2019). As a result, we delineated 378 vOTUs (Table S3), 102 of these vOTUs included both genomes identified in this study and genomes from the databases, whereas 125 were composed entirely of the crAss-like phage genomes identified in this study (Table S4). 217 vOTUs contained tentatively complete genomes, which were selected to represent these vOTUs in subsequent analyses. The remaining 161 vOTUs were represented by the longest available genome fragments.

Next, we explored the genome organization of the 125 newly discovered vOTUs of crAss-like phages. Their representative genomes and genome fragments possess characteristics similar to those of the previously discovered crAss-like phages, including the identifiable replication, transcription and capsid gene modules, and the presence and order of the key conserved genes (Material S2). Notably, of the 125 representatives of the newly discovered vOTUs, 49 and 2 apparently utilize the TAG and the TGA stop codon reassignment for an amino acid, respectively (Material S2). Interestingly, all complete and nearly complete crAss-like phage genomes representing vOTUs possess a similar GC skew pattern, which was first reported for the ΦCrAss001 phage (Shkoporov *et al.*, 2018): each strand of the genome is characterized by high GC skew values in coding region and low GC skew values in the rest of the genome (Material S2). A similar but slightly less pronounced pattern was observed for the AT skew in most crAss-like phages outside of the alpha/gamma group (Material S2).

The 378 genomes representing species-level vOTUs were further clustered into 182 genus-level clusters under the 50% AF threshold (Adriaenssens and Brister, 2017). 60 of these clusters included both genomes identified in this study and genomes from the databases, while 32 clusters were composed entirely of crAss-like phage genome sequences identified in this study (Table S4). On TerL-based and portal-based phylogenetic trees, all of the vOTUs containing crAss-like phages identified in this study fell into the five known groups associated with the human gut: alpha/gamma, beta, delta, epsilon and zeta (Figure 1, Table S4, Material S3-S4). The names used to designate genus-level clusters in this study are composed of the name of the group to which a cluster belongs and a number, e.g. delta27.

**Figure 1.**
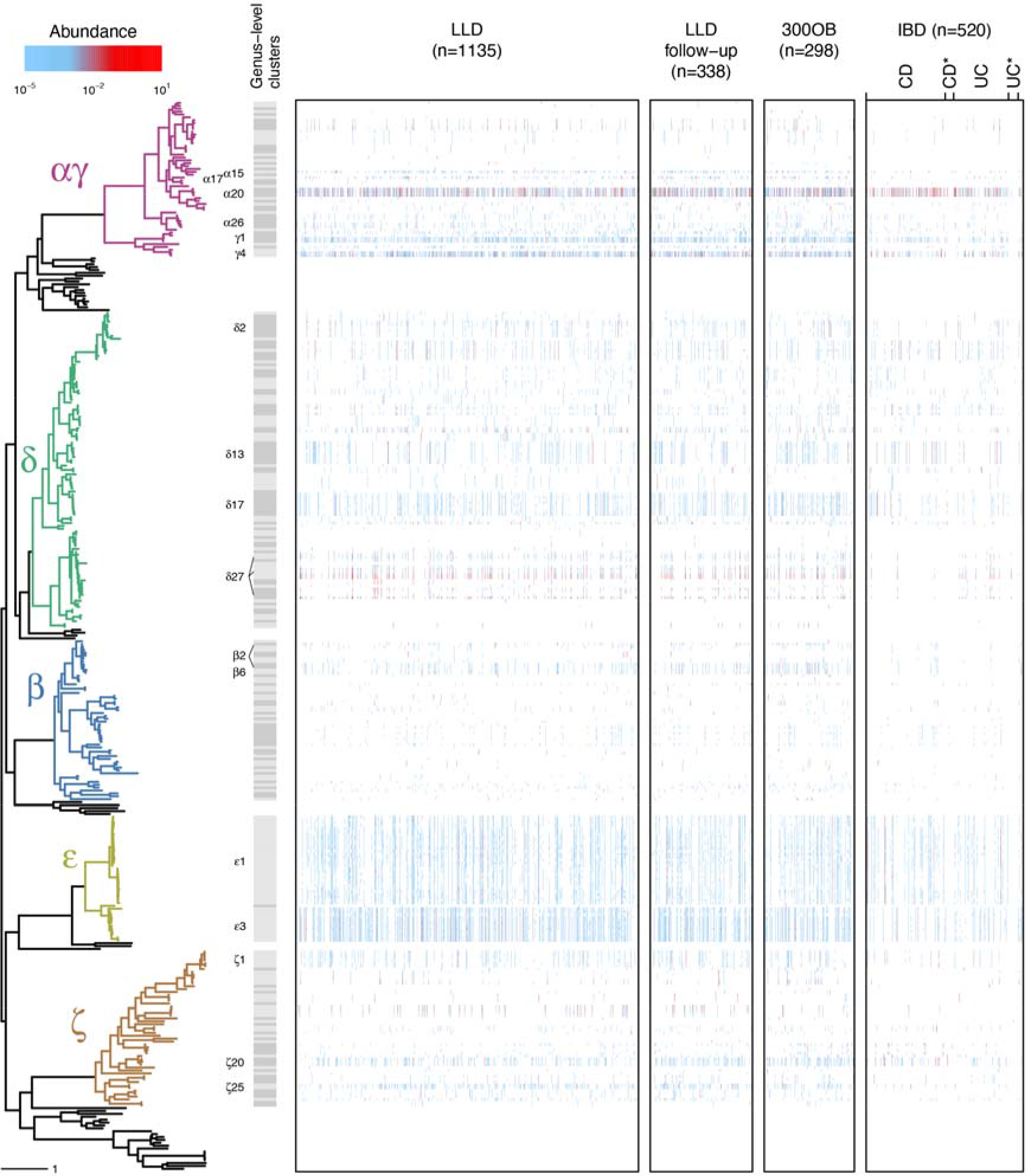
Abundance of crAss-like phages in the four cohorts. A TerL-based phylogenetic tree is juxtaposed with four abundance matrices representing the four cohorts. The tree is midpoint pseudo-rooted, and tree tips correspond to crAss-like phages representing vOTUs. Seven vOTUs represented by partial genomes lacking a TerL gene are not shown. The five clades of crAss-like phages associated with the human gut are designated by the color of the branches. Genus-level clusters of the crAss-like phages belonging to the five groups associated with the human gut are depicted by the grey bars next to the tree, with names indicated for genus-level clusters present in > 10% of samples in any cohort. See Material S3 for the detailed annotation of the tree. Rows of the matrices correspond to crAss-like phage vOTUs and are ordered according to their position on the phylogenetic tree. Columns of the matrices correspond to samples and are ordered according to the age of the individuals (LLD and LLD follow-up cohorts), body mass index (BMI) (300OB cohort) and diagnosis (IBD cohort). The color of a matrix element indicates estimated vOTU abundance in a sample. CD, Crohn’s disease; UC, ulcerative colitis; asterisk next to the diagnosis abbreviation indicates that the individual has a stoma.

### Closely related crAss-like phages can possess highly divergent genome regions responsible for transcription

Following the delineation of vOTUs, we mapped the sequencing reads from each sample to the 378 crAss-like phage genomes representing vOTUs. When analyzing the coverage of crAss-like phage genomes by sequencing reads, we noticed a common anomaly: the transcription gene module often received either zero coverage or, conversely, higher coverage than the rest of the genome in a fraction of analyzed samples (Figure 2A-B, Material S2, S5). While this anomaly was associated with multiple diverse crAss-like phage genomes, we selected one random genome, OLNE01000081 belonging to the delta27 genus-level cluster, to analyze the anomaly in-depth. The region of OLNE01000081 encoding RNA polymerase (RNAP) and a protein homologous to VP02740, an uncharacterized transcription-related protein (Yutin *et al.*, 2021), was not covered by reads in a fraction of samples, while in other samples coverage of this region was comparable to the rest of the genome (Figure 2A-B, Material S5).

**Figure 2.**
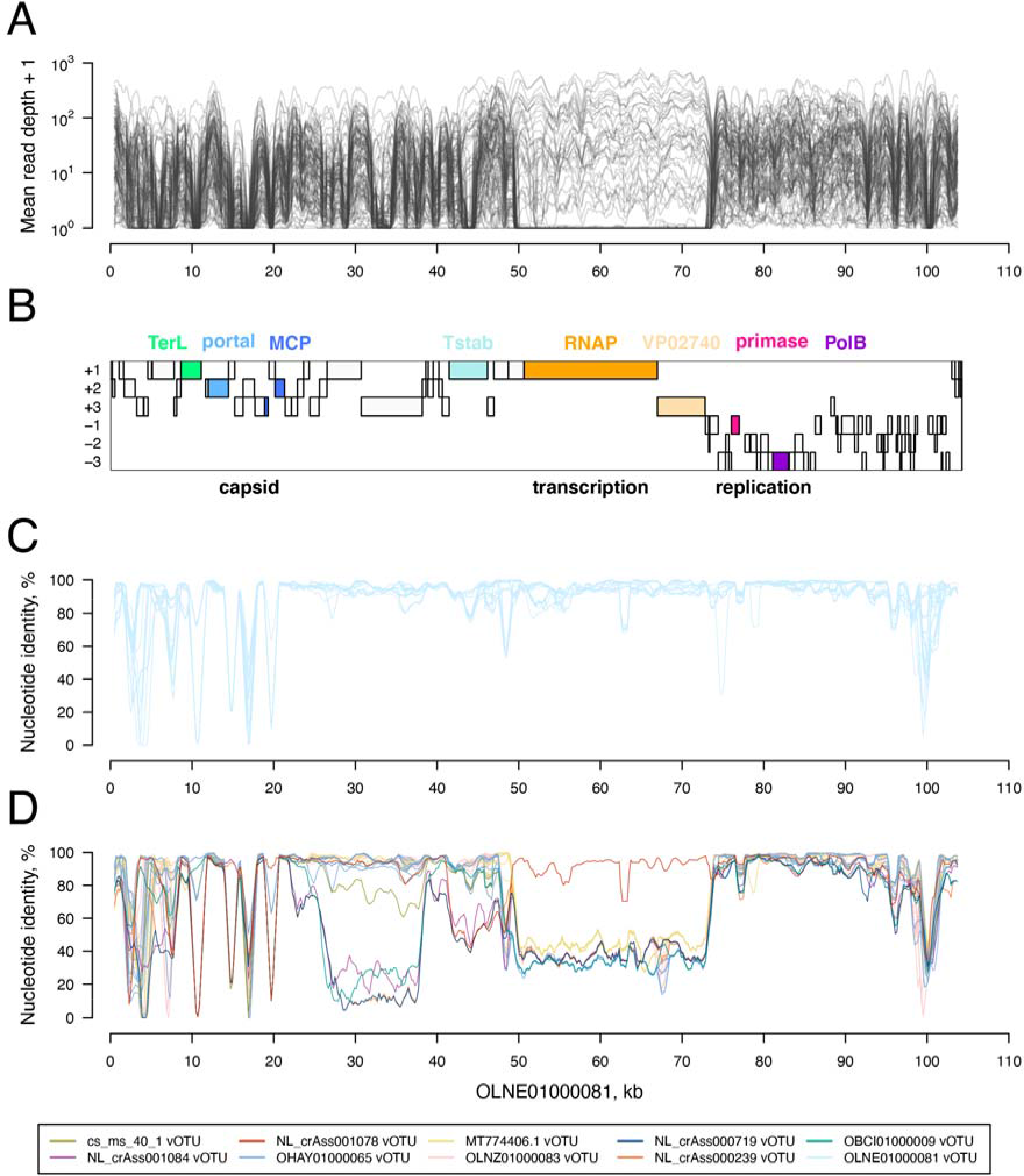
OLNE01000081 genome properties. (**A**) Depth of the OLNE01000081 genome coverage by sequencing reads from the LLD cohort. Each transparent grey line corresponds to a sample and represents mean depth in a 1001 nt sliding window. (**B**) OLNE01000081 genome organization. A genome is presented as a black rectangular contour, and three forward and three reverse frames of the genome are indicated. Open reading frames (ORFs) translated under the TAG stop codon reassignment for glutamine are presented as bars. Products of the ORFs shown in color are indicated above the genome map: TerL, terminase large subunit; MCP, major capsid protein; Tstab, tail stabilization protein; RNAP, RNA polymerase; VP02740, uncharacterized transcription-related protein (Yutin *et al.*, 2021); PolB, DNA polymerase family B. Functional regions of the genome are indicated below the genome map. (**C**) Nucleotide identity between the OLNE01000081 and other tentatively complete genomes belonging to the same vOTU. (**D**) Nucleotide identity between the OLNE01000081 genome and (nearly) complete genomes belonging to other vOTUs of the delta27 genus-level cluster. Data for genomes belonging to different vOTUs are distinguished by color (see inset). Nucleotide identity presented on panels C and D was calculated using a 1001 nt sliding window.

To investigate the cause of this coverage anomaly, we compared the OLNE01000081 genome to complete genomes belonging to the same vOTU (Figure 2C) and to complete and nearly-complete genomes belonging to other delta27 vOTUs (Figure 2D). While genomes belonging to the same vOTU were almost identical to the OLNE01000081 genome in all regions except the region encoding capsid proteins, where islands of high conservation are interspersed with low conservation stretches (Figure 2C), genomes belonging to other delta27 vOTUs demonstrated a notable divergence from OLNE01000081 in some regions and very strong similarity in others. Most strikingly, nucleotide identity between the OLNE01000081 genome and genomes from all other delta27 vOTUs (except NL_crAss001078) abruptly dropped to ~30% at the 5’-end of the RNAP gene and was abruptly restored to ~100% at the 3’-end of the VP02740 gene (Figure 2D). This pattern involves exactly the same region as the coverage anomaly described above and is consistent with the substitution of the genome region in question via recombination (Lole et al., 1999). A phylogenetic tree reconstructed based on the transcription gene module shows that there are four distinct types of this region within the delta27 genus-level cluster (Figure 3). The origin of the type present in OLNE01000081 remains unknown, as no close (> 20% gene length coverage by BLASTN hits with E-value < 0.001) non-delta27 homologs of either RNAP or VP02740 genes were detected in crAss-like phage genomes and GenBank entries (Sayers et al., 2021).

**Figure 3.**
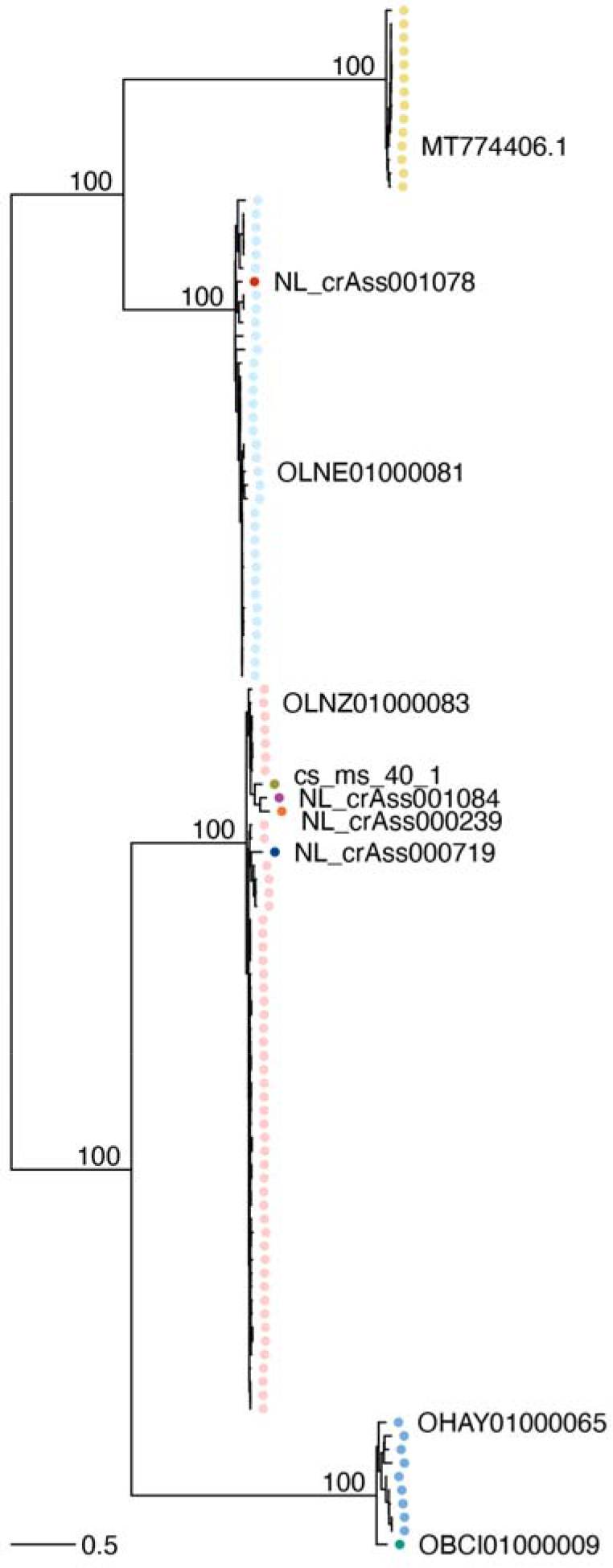
Phylogenetic tree based on the transcription gene module of the delta27 crAss-like phages. A total of 114 delta27 crAss-like phages with (nearly) complete genome sequences are included. The vOTU of each tree tip is indicated by color. Tree tips corresponding to genomes representing vOTUs are labeled. Bootstrap support is specified for long branches.

The fact that besides the delta group, the coverage anomaly associated with the transcription gene module was also observed in alpha/gamma, beta and zeta groups, indicates that the presence of highly divergent transcription gene modules in closely related crAss-like phages may be a common phenomenon. We further confirmed the link between the coverage anomaly and nucleotide sequence divergence in the genus-level clusters alpha6, beta8 and zeta9 belonging to the three latter groups (Material S6).

### Prevalence and temporal stability of crAss-like phages

Mapping sequencing reads to crAss-like phage genomes representing vOTUs allowed us to estimate the abundance of crAss-like phages in metagenomic samples. The percentage of filtered and quality-trimmed sequencing reads mapping to the crAss-like phage genomes varied from 0% to 20.7% per sample, with a mean of 0.4% in each of the four cohorts (Figure S2).

A vOTU was considered to be detected in a sample if 10% of the length of its representative sequence was covered by reads. Based on this criterion, all but 17 vOTUs belonging to the five groups associated with the human gut were detected in at least one sample in one of the four cohorts, and no vOTUs outside of these five groups were detected in any sample (Figure 1). The correlation between the number of detected vOTUs and the number of sequencing reads per sample was relatively weak (r_Spearman_ = 0.213, P-value = 6.28e-25; Figure S1D).

Almost universally, there was a correlation between detection of vOTUs belonging to the same genus-level cluster (Figure 1). This pattern is likely explained by mis-alignment of reads to reference genomes representing vOTUs closely related to, but different from, the vOTU of their origin. As it complicates the vOTU-level analysis of the abundance of crAss-like phages, we decided to conduct all the subsequent analyses at the genus and higher taxonomic levels.

A total of 118 of the 132 crAss-like phage genus-level clusters associated with the human gut were detected in at least one sample, and 1998 of the 2291 samples contained at least one crAss-like phage. On average, 4.3, 4.7, 5.3 and 1.9 crAss-like phage genus-level clusters were detected in LLD, LLD follow-up, 300OB and IBD cohort samples, respectively. The most abundant clusters were alpha20, which contains the first discovered crAssphage (Dutilh *et al.*, 2014), and delta27 (Figure S3). The most frequently detected clusters were alpha20, gamma1, gamma4, epsilon1 and epsilon3 (detected in > 20% of samples in at least one cohort, Figure S3).

To study the temporal stability of the crAss-like phages, we used the data from the LLD (Zhernakova *et al.*, 2016) and LLD follow-up (Chen *et al.*, 2021) cohorts: samples collected from the same 338 individuals at two timepoints approximately 4-years apart. First, we assessed the stability of individual genus-level clusters of crAss-like phages. Only clusters detected in > 10% of LLD or LLD follow-up samples were considered. Among the individuals positive for a genus-level cluster in at least one timepoint, 41%, on average, were positive at both timepoints (Figure 4A). This number was above 50% for two genus-level clusters: alpha20 and gamma4. The overall composition of the crAss-like fraction of the virome was also relatively stable. The dissimilarity between samples collected from the same individual at the two timepoints, quantified using Bray-Curtis (BC) distances, was smaller than the dissimilarity between pairs of samples collected from different individuals at the same timepoint (mean intra-individual BC distance 0.72, mean inter-individual BC distance in LLD cohort 0.94, Wilcoxon signed-rank test P-value 2.06e-90) (Figure 4B). Thus, we observed that, on average, people are more similar to themselves over time than they are to unrelated individuals at the same timepoint.

**Figure 4.**
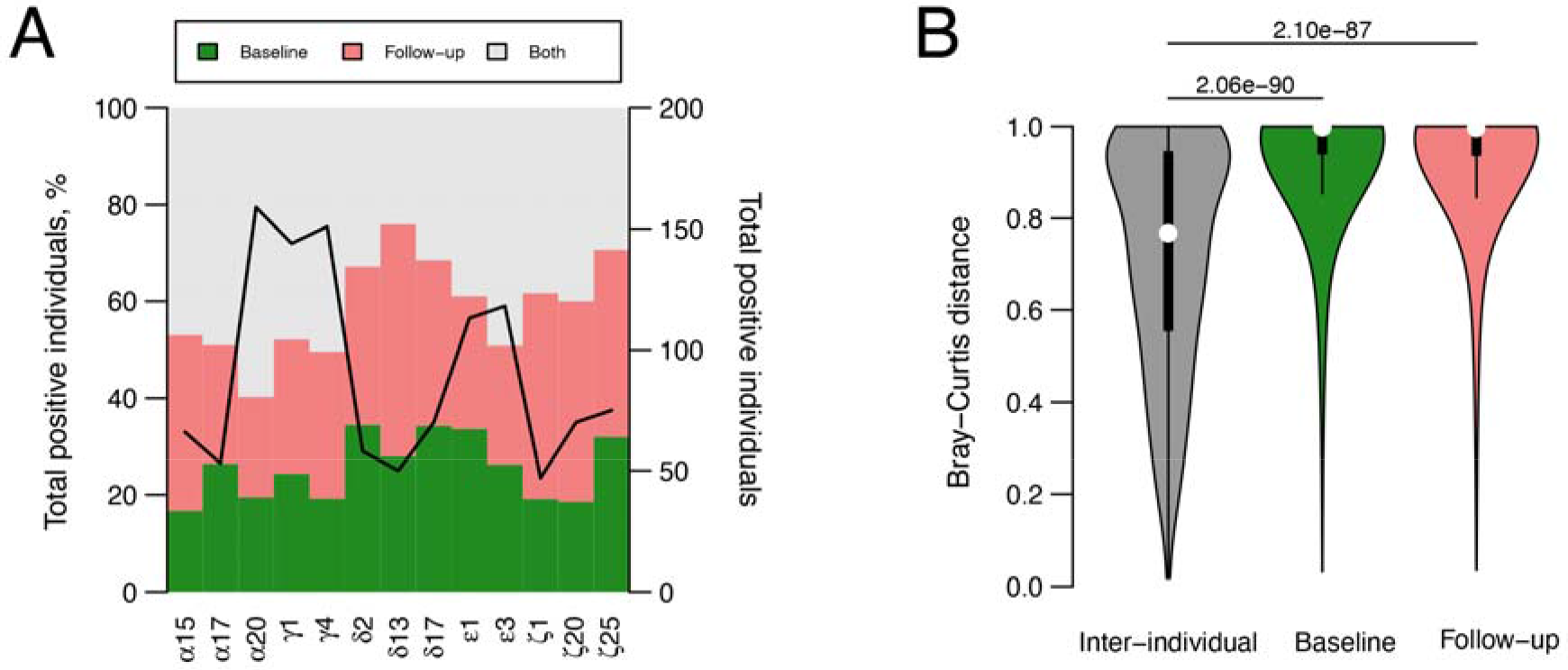
Temporal variation of the crAss-like phages composition in LLD and LLD follow-up longitudinal samples. (**A**) Variation at the level of individual genus-level clusters of crAss-like phages. The 13 clusters detected in > 10% of LLD or LLD follow-up samples were considered. The number of study subjects positive for a cluster in at least one timepoint is shown by a black line. The proportion of study subjects positive for a cluster only at the first, only at the second, or at both timepoints is indicated by green, red and grey colors, respectively. (**B**) Variation in composition of the crAss-like fraction of the virome. A distribution of the Bray-Curtis distances, quantifying the dissimilarity of the composition between samples, is shown for pairs of samples collected from the same individual at the baseline and follow-up timepoints (grey), and pairs of samples collected from different individuals at the same timepoint (green and red). P-values of the one-sided Wilcoxon signed-rank tests comparing the distributions are indicated above the violin plots.

### Potential hosts of crAss-like phages and relation to bacterial structural variants

Next, we predicted the potential hosts of the crAss-like phages using two approaches: we analyzed matches between microbial CRISPR spacers and crAss-like phage genomes and explored the co-abundance of crAss-like phages and gut microbes in the samples from the four cohorts.

Sequence similarity of CRISPR spacers in microbial genomes to regions of crAss-like phage genomes allowed us to link crAss-like phages from 32 genus-level clusters to 10 bacterial genera (Figure 5, Table S5). Many of these links were previously reported in (Yutin *et al.*, 2021). The majority (98%) of the links identified bacteria from the genera *Bacteroides*, *Prevotella*, *Porphyromonas* and *Parabacteroides*, all belonging to the order Bacteroidales, as potential hosts, consistent with the prediction made by the discoverers of the first crAssphage (Dutilh *et al.*, 2014). However, we also identified bacteria from phyla Firmicutes and Fusobacteria as potential hosts of a small number of crAss-like phages. There does not seem to be a one-to-one correspondence between groups of crAss-like phages and bacterial genera, as the alpha, delta and epsilon groups of crAss-like phages were each linked to three out of the four Bacteroidales genera mentioned above (Figure 5).

**Figure 5.**
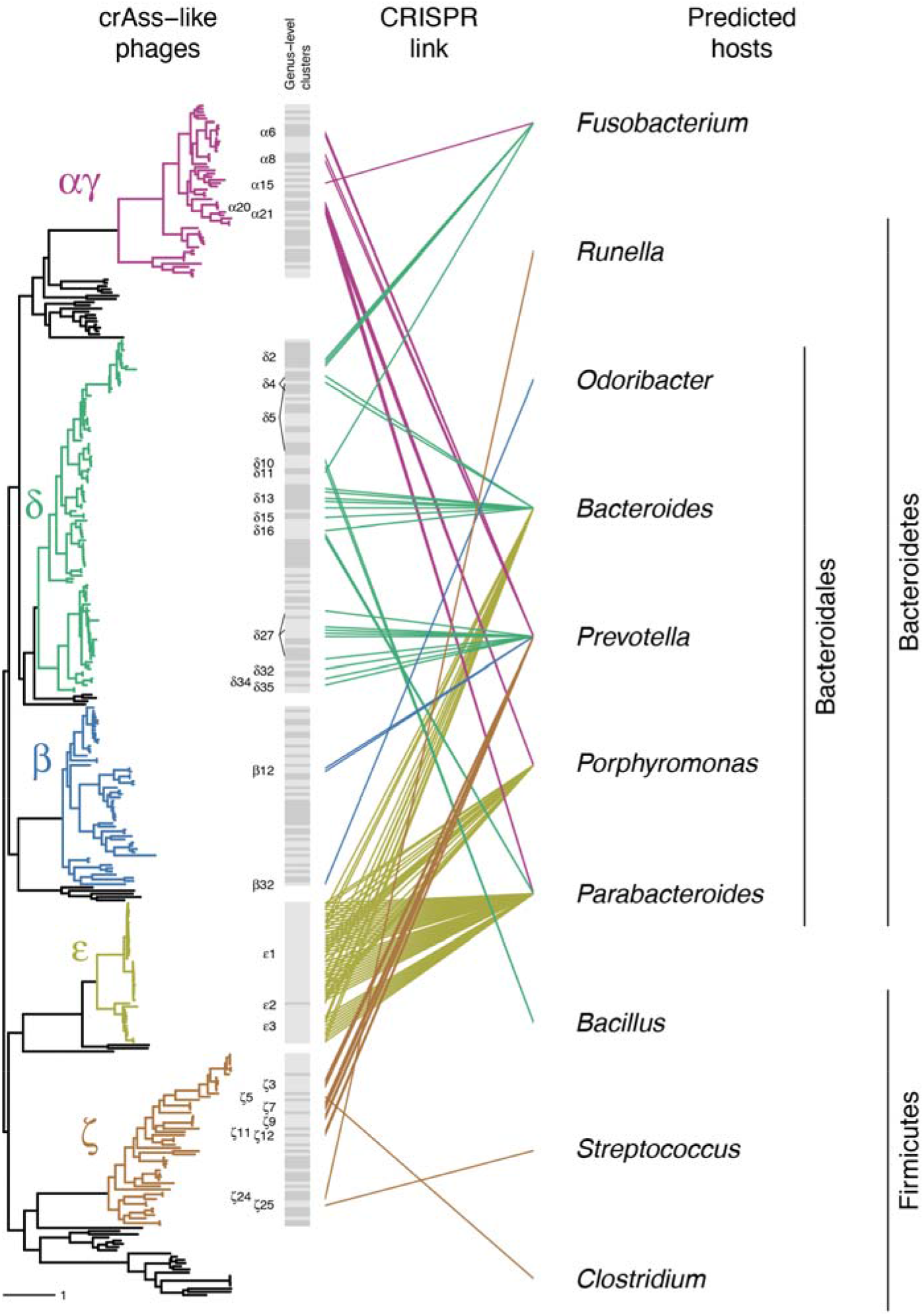
Links between crAss-like phages and bacterial genera based on CRISPR spacer matches. For details about the phylogenetic tree see the legend of Figure 1.

To detect correlation between the abundance of crAss-like phages and microbes, which may be a sign of a phage–host relationship, we calculated the Spearman correlation coefficient for pairs of phage and microbe taxa in each cohort, followed by meta-analysis of the results from three independent cohorts: LLD, 300OB and IBD cohort (Table S6). The meta-analysis identified 19 pairs of taxa characterized by an absolute value of the correlation coefficient > 0.2. All of these correlations were positive, and 16 of them involved bacteria from the order Bacteroidales, while the remaining three were with bacteria from the phyla Firmicutes and Actinobacteria. The phage–microbe pair with the highest absolute value of the correlation coefficient in each cohort and meta-analysis (r_meta_ = 0.447, P-value = 1.29e-78) was crAss-like phage genus-level cluster delta27 and bacterial species *Prevotella copri* (Figure S4, Table S6). This observation, together with the fact that a number of delta27 crAss-like phages were linked to *P. copri* based on CRISPR analysis (Figure 5, Table S5), identifies *P. copri* as a likely host of delta27 crAss-like phages. Overall, the results of the two methods of host prediction were consistent, both with each other and with previous studies, and identified members of the phylum Bacteroidetes as the major hosts of the crAss-like phages associated with the human gut.

Next, we explored associations between the abundance of crAss-like phages and structural variants (SVs) in bacterial genomes (Wang et al., 2021; Zeevi et al., 2019). Based on the collection of samples from the three independent cohorts LLD, 300OB and IBD, we detected 7,999 deletion structural variants (dSVs, genome segments absent in 25% to 75% samples) and 3,559 variable structural variants (vSVs, genome segments absent in < 25% samples). We analyzed associations between the detected SVs and the abundance of crAss-like phages in each of the three cohorts separately and integrated the results via meta-analysis and heterogeneity analysis. In total, we identified 22 replicable significant SV associations (19 vSV associations and 3 dSV associations) with the relative abundance of crAss-like phages at the genus and higher taxonomic levels (Table S7). The crAss-like phage taxonomic clusters involved in these associations included crAss-like phage assemblage, groups beta and delta, and genus-level clusters alpha15, gamma1, gamma4, delta27 and epsilon1. All the bacteria involved in these associations belonged to the phyla Actinobacteria and Firmicutes. The SVs in these associations ranged from 1 to 14 kb, and according to proGenomes database (Mende et al., 2017) annotation, encoded functionally diverse proteins including DNA primases and components of the ABC-type transport system (Table S7).

### CrAss-like phage associations with human phenotypes

To identify if crAss-like taxonomic clusters are associated with intrinsic, dietary and clinical phenotypes in a general population context, we analyzed 1135 metagenomic samples and 207 phenotypes from the population-based LLD cohort. The analysis was conducted for 30 taxonomic clusters of crAss-like phages detected in > 5% of LLD samples (crAss-like phage assemblage, 5 groups, 24 genus-level clusters) and utilized five different methods that demonstrated overall agreement (Table S8). Associations were considered significant at a false discovery rate (FDR) < 0.05.

When Spearman correlation analysis was applied to the crAss-like phage presence-absence data, the fecal level of the secretory protein chromogranin A (CgA), which is a precursor to peptides with regulatory, antimicrobial and antifungal activities (Bartolomucci et al., 2011), was negatively correlated with the presence of crAss-like phage assemblage, groups alpha/gamma and beta, and genus-level clusters gamma1 and gamma4, while the fecal level of the antimicrobial peptide human beta-defensin-2 (HBD-2) was positively correlated with the presence of genus-level cluster gamma1. This analysis also linked crAss-like phages to three dietary phenotypes: frequency of coffee intake was positively correlated with the presence of crAss-like phage assemblage and group beta, meat intake was positively correlated with the presence of group epsilon and genus-level cluster epsilon1, and the amount of carbohydrates in the diet was positively correlated with the presence of genus-level cluster alpha20. In addition, the presence of the alpha/gamma group was negatively associated with irritable bowel syndrome (IBS) diagnosis, stomach ulcer diagnosis and usage of laxatives (Table S8).

In contrast, when Spearman correlation analysis was applied to the crAss-like phage abundance data, only five associations were statistically significant (FDR < 0.05): the three associations with chromogranin A level described above and two of the dietary phenotype associations described above (Table S8). Unadjusted logistic regression and logistic regression adjusted for the age and sex of cohort participants yielded almost identical results: six and seven associations described above (with the level of chromogranin A and health conditions) were recognized as statistically significant, respectively (FDR < 0.05; Table S8). Interestingly, when the logistic regression was further adjusted for the log-transformed abundance of the bacterial genera *Bacteroides*, *Prevotella*, *Porphyromonas* and *Parabacteroides*, which were most frequently identified as potential hosts of crAss-like phages based on CRISPR spacer matches (Figure 5), there were no significant associations (FDR < 0.05), and all of the associations described above were only nominally significant.

Overall, crAss-like phage taxonomic clusters were associated with a limited number of human phenotypes, and the strongest, most consistent between analyses, were associations with the level of chromogranin A. This finding is in line with the massive effect of chromogranin A on the gut bacteriome reported earlier for this cohort (Zhernakova *et al.*, 2016). The associations were partially independent of the human age and sex but were influenced by the abundance of the potential host genera.

### CrAss-like phages in the context of IBD and obesity

Next, we aimed to explore the crAss-like phage composition in two different disease contexts: the metabolically altered environment represented by a cohort of overweight or obese individuals (300OB cohort; BMI > 27 kg/m^2^) (Ter Horst *et al.*, 2020) and the inflammatory environment represented by a cohort of patients with inflammatory bowel disease (IBD cohort) (Imhann *et al.*, 2019; Vich Vila *et al.*, 2018). We studied the potential effect of the two disease contexts on the crAss-like phage composition in the gut by comparing the prevalence of crAss-like phage assemblage, 5 groups and 24 genus-level clusters, which were detected in > 5% samples of the population-based LLD cohort, between the respective cohort and LLD. The analyses were performed using logistic regression, correcting for age and sex, as well as for carbohydrates and meat consumption, frequency of coffee consumption, usage of laxatives, and CgA and HBD-2 levels where applicable (see Methods). The prevalence of crAss-like phage assemblage, 5 groups and 19 genus-level clusters was significantly lower in IBD samples compared to LLD samples (FDR < 0.05; Table S9). All but two of these associations remained significant when the regression was further adjusted for the log-transformed abundance of the four potential host genera: *Bacteroides*, *Prevotella*, *Porphyromonas* and *Parabacteroides* (FDR < 0.05; Table S9). In contrast, we observed no significant difference in the prevalence of any of the crAss-like phage taxonomic clusters between LLD and 300OB cohorts (FDR > 0.05; Table S9).

Previous reports have shown that the fecal bacterial composition differs in IBD patients with Crohn’s disease (CD) versus ulcerative colitis (UC), as well as in IBD patients with varying disease locations (Franzosa et al., 2019; Vich Vila *et al.*, 2018). To investigate if these differences also extend to crAss-like phages, we performed differential prevalence analyses by comparing samples from IBD patients with (1) CD versus UC and (2) an exclusively colonic versus an ileal-inclusive disease location. In both tests, we did not observe any significant differences between the test groups at any of the taxonomic levels (FDR > 0.05; Table S9).

Finally, to investigate whether metabolic syndrome is associated with the crAss-like phage prevalence, we stratified samples from the 300OB cohort based on the presence or absence of metabolic syndrome according to the National Cholesterol Education Program ATP III criteria (Ter Horst *et al.*, 2020). We found no significant associations between the two groups at any of the taxonomic levels (FDR > 0.05; Table S9).

## Discussion

In this study, we relied on existing knowledge about the genomes and proteomes of crAss-like phages (Dutilh *et al.*, 2014; Guerin *et al.*, 2018; Yutin *et al.*, 2021; Yutin *et al.*, 2018) to systematically analyze the diversity and abundance of crAss-like phages in gut metagenomic samples collected from participants of four cohorts set up in the Netherlands: LLD, LLD follow-up, 300OB and IBD. Using the three conserved capsid and genome-packaging proteins of crAss-like phages (TerL, portal and MCP) as markers, we were able to detect 1556 crAss-like contigs. 76% of these contigs clustered into species-level vOTUs populated by previously discovered crAss-like phages, while the remaining 24% formed 125 novel vOTUs. At the genus level, 96% of the newly discovered contigs clustered with the known crAss-like phages, while the remaining 4% formed 32 novel genus-level clusters. Notably, all the crAss-like contigs we detected fell into the five known groups associated with the human gut (Guerin *et al.*, 2018; Yutin *et al.*, 2021). This may indicate that most groups of gut-associated crAss-like phages present in the Western European population have already been discovered, although, importantly, our detection method was based on information about the known groups and may be less sensitive for detection of other distantly related groups.

Interestingly, we observed that crAss-like phage genomes can be almost identical throughout a large portion of their length but possess strikingly different genome regions responsible for transcription. This phenomenon is likely to be widespread among crAss-like phages, as its manifestation – differential sequencing read coverage of the transcription gene module – was observed in multiple representatives of the alpha/gamma, beta, delta and zeta groups of crAss-like phages. Lower coverage of this region compared to the rest of the genome can be explained by the presence of a crAss-like phage that is closely related to the reference genome used for read mapping but possesses a highly divergent transcription gene module. Higher coverage of this region compared to the rest of the genome can be explained by the presence of a distant crAss-like phage that possesses a transcription gene module similar to the one in the reference genome. The likely mechanism driving the emergence of diverse genome regions responsible for transcription among closely related crAss-like phages is recombination. Acquisition of a divergent functional region through recombination may be beneficial for a virus, for instance allowing it to adapt to changing conditions. For example, non-segmented (+)ssRNA genomes of the *Enterovirus C* virus species include a type-specific capsid coding region and a non-structural protein coding region subject to frequent inter-typic homologous recombination, which may lead to the emergence of novel virus lineages (Smura et al., 2014). A similar mechanism of genetic diversification can be envisioned for the crAss-like phages.

Analysis of samples from 388 individuals taken at two timepoints (Chen *et al.*, 2021; Zhernakova *et al.*, 2016) allowed us to explore the temporal stability of crAss-like phages. Despite the 4-year period separating the two timepoints, we observed a degree of stability both at the level of individual genus-level clusters (on average, 41% of individuals positive for a frequently detected genus-level cluster were positive at both timepoints) and at the level of the overall composition of the crAss-like fraction of the virome (on average, dissimilarity between samples collected from the same individual at the two timepoints was lower than the dissimilarity between pairs of samples collected from different individuals at the same timepoint). These observations are in line with previous reports of temporal stability of crAss-like phages (Edwards *et al.*, 2019; Shkoporov *et al.*, 2019; Siranosian *et al.*, 2020), which may be explained by a relatively peaceful co-existence of phage and host (Guerin *et al.*, 2021; Shkoporov *et al.*, 2018; Shkoporov *et al.*, 2021).

Consistent with the results of previous studies (Dutilh *et al.*, 2014; Yutin *et al.*, 2021), our analysis of matches between CRISPR spacers from microbial genomes and crAss-like phage genomes, as well as our analysis of co-abundance of microbial and crAss-like phage taxa, pointed primarily to bacteria of the phylum Bacteroidetes as the hosts of the crAss-like phages. Predicting the hosts of phages is challenging because CRISPR spacer sequences are short and may match phage genomes just by chance, while only a fraction of microbes utilize the CRISPR defense system (Shmakov et al., 2017). Co-abundance analysis can detect a linear or monotonous relationship, but may not be able to capture a more complex relationship, e.g. a time-lagged phage–host relationship (Edwards et al., 2016), or distinguish between a phageChost relationship and other ecological interactions. Combining multiple techniques of host prediction can increase confidence in the results. In this study, both CRISPR-based and co-abundance analyses strongly pointed to *Prevotella copri* as a host of the crAss-like phages belonging to the genus-level cluster delta27. The fact that the correlation between the relative abundance of the delta27 phages and their potential host is positive suggests that the dynamics in this phage–host pair may be similar to the co-existence in equilibrium reported for the four crAss-like phages isolated in cultures of their hosts (Guerin *et al.*, 2021; Hryckowian *et al.*, 2020; Shkoporov *et al.*, 2018; Shkoporov *et al.*, 2021).

The interplay between phages and bacteria may be influenced by structural variation in bacterial genomes. For instance, several studies have linked phase variation in the bacterial genome loci responsible for cell surface coating biosynthesis to acquisition of resistance to phage infection (Porter et al., 2020; Shkoporov *et al.*, 2021). In our study, we identified 22 replicable significant associations between bacterial structural variants and crAss-like phage taxonomic clusters. All the bacteria involved in these associations were from the phyla Actinobacteria and Firmicutes, while the majority of the taxonomic clusters of crAss-like phages involved were linked to potential hosts from the order Bacteroidales by CRISPR spacer matches. Consequently, the detected associations are unlikely to stem from phage– host relationships and may instead reflect yet unknown interactions in the broader ecological community of the gut.

Our analysis of the correlations between human phenotype levels and crAss-like phage taxonomic clusters in the population-based LLD cohort revealed several associations with intrinsic (levels of chromogranin A and beta-defensin-2 secretory proteins in stool), dietary (coffee, meat and carbohydrate intake) and health (IBS and stomach ulcer diagnosis, usage of laxatives) phenotypes. An association between a phage and a human phenotype may reflect the association between the microbial host of the phage and the phenotype in question. Although the exact hosts of the majority of the crAss-like phages remain unknown, both previous reports (Dutilh *et al.*, 2014; Yutin *et al.*, 2021) and our data point to the bacteria of the phylum Bacteroidetes, and in particular the order Bacteroidales, as the major hosts of the crAss-like phages associated with the human gut. Accordingly, four of the phenotypes mentioned above were reported to be associated with bacteria belonging to the order Bacteroidales, and in three of these cases (levels of chromogranin A and beta-defensin-2, IBS) the directionality of the association matched that of the crAss-like phage associations with the phenotype in question, while in one case (laxative usage) the directionality was opposite (Zhernakova *et al.*, 2016). The notion that phage-phenotype associations may reflect the host–phenotype associations is further supported by our observation that, after the adjustment of the logistic regression model of the phage–phenotype relationship for the abundance of the four potential host genera (*Bacteroides*, *Prevotella*, *Porphyromonas* and *Parabacteroides*), the identified phage–phenotype associations became only nominally significant.

The prevalence of the crAss-like phages was strongly associated with an IBD diagnosis: the assemblage, all 5 groups and 19 genus-level clusters of crAss-like phages were less prevalent in the IBD patient cohort compared to the population-based LLD cohort. In contrast, there was no statistically significant difference in prevalence of crAss-like phages between the LLD and 300OB cohorts, at any taxonomic level. The depletion of crAss-like phages in the IBD cohort may be explained by the depletion of their hosts. A decrease in bacterial biomass and bacterial richness has consistently been described in patients with IBD, including the depletion of several members of the order Bacteroidales (Schirmer et al., 2019; Vieira-Silva et al., 2019). Although, interestingly, the majority of the associations between prevalence of crAss-like phages and IBD diagnosis remained significant after the adjustment of the logistic regression model for the abundance of the four potential host genera.

In conclusion, we have expanded the known diversity of crAss-like phages by a third at the species level, thus contributing to the ongoing discovery and cataloging of the human gut virome. Our study shows that identifying, classifying and analyzing novel viral genomes can both offer new clues about the biology of viruses and further our understanding of the interactions between viruses and their surrounding ecosystem and the role of viruses in human health and disease.

## Methods

### Datasets

Data used in the project included gut metagenome sequencing data and human phenotype data for 1135 LLD cohort participants (Tigchelaar *et al.*, 2015; Zhernakova *et al.*, 2016), 338 LLD follow-up cohort participants (Chen *et al.*, 2021) and 298 participants of the 300OB cohort (Ter Horst *et al.*, 2020). A total of 520 gut metagenome samples from the IBD cohort (Imhann *et al.*, 2019) containing ≥10 million sequencing reads were screened for the presence of crAss-like phages. To conduct association analyses, we excluded IBD samples taken from participants with stoma and ileoanal pouch, one duplicated sample and two samples for which no metadata was available, resulting in a subset of 455 IBD samples. For all cohorts, DNA isolation and metagenomic sequencing was done in the same laboratory and used the same protocols and pipelines.

### Sequencing reads quality control

Sequencing reads from each sample were independently processed using the KneadData 0.7.4 pipeline (https://huttenhower.sph.harvard.edu/kneaddata/). Specifically, reads mapping to the human reference genome GRCh38.p13 (Schneider et al., 2017) were filtered out, and adapter and quality trimming of the reads was performed using Trimmomatic 0.33 (Bolger et al., 2014) with the parameters “ILLUMINACLIP:NexteraPE-PE.fa:2:30:10:1:true LEADING:20 TRAILING:20 SLIDINGWINDOW:4:20 MINLEN:50”. Read quality was assessed using FastQC 0.11.7 (Wingett and Andrews, 2018) and MultiQC 1.9 (Ewels et al., 2016).

### Contigs assembly

Contigs were assembled for each sample independently using metaSPAdes 3.14.1 (Nurk et al., 2017). In some cases (14, 1, 6 and 24 samples from LLD, LLD follow-up, 300OB and IBD cohorts, respectively), metaSPAdes did not progress to finish its read error correction step. For these samples, read error correction was conducted using BBMap 37.93 program tadpole.sh with “mode=correct ecc=t prefilter=2” parameters (https://sourceforge.net/projects/bbmap/), followed by metaSPAdes assembly with the “-- only-assembler” parameter.

### crAss-like phage detection

The datasets screened for the presence of crAss-like phages included sets of contigs ≥ 10 kb from individual samples belonging to the LLD, LLD follow-up, 300OB and IBD cohorts, as well as genomes from four control datasets: Viral RefSeq 202 (Brister *et al.*, 2015), a database of 249 crAss-like phages described in (Guerin *et al.*, 2018), a database of 673 crAss-like phages described in (Yutin *et al.*, 2021) and a database of 146 nucleotide sequences ≥ 10 kb recognized as “crAss-like” by the NCBI Taxonomy resource on 24.12.2020 (Schoch *et al.*, 2020). Genomes and contigs were translated in six frames by the EMBOSS 6.6.0 program transeq with a “-clean” parameter (Rice et al., 2000), and their proteomes were predicted by Prodigal 2.6.3 in the “meta” mode (Hyatt et al., 2010). The HMMER 3.3 program hmmsearch with an E-value threshold of 0.001 (http://hmmer.org/) was used to compare proteomes, and, separately, translations in six frames to the following three highly conserved structural and genome-packaging proteins of crAss-like phages: terminase large subunit (three profiles from (Yutin *et al.*, 2021)), portal protein (three profiles from (Yutin *et al.*, 2021)) and major capsid protein (nine profiles from (Yutin *et al.*, 2021; Yutin *et al.*, 2018)). The output was processed with the help of R package *rhmmer* 0.1.0 (https://CRAN.R-project.org/package=rhmmer). A genome or a contig was considered as crAss-like if either its predicted proteome or its translations in six frames were hit by profiles representing each of the three proteins.

### Nucleotide sequence characterization

To identify potentially complete genomes, direct overlaps of ≥10 nt at the ends of the genome sequences were detected using the approach from (Roux et al., 2015). The nucleotide content was estimated as the percent of each of the four nucleotides in a window sliding along the genome. GC and AT skew were calculated as (G − C)/(G + C) and (A − T)/(A + T) in a window sliding along the genome, respectively (Lobry, 1996). A 1001 nt window with a 200 nt step was used in all calculations; the most 3’-terminal window was extended to include up to 200 3’-terminal nucleotides. Cumulative GC (AT) skew corresponding to a window was calculated as a sum of GC (AT) skew values in this and all preceding windows (Grigoriev, 1998).

### Species- and genus-level clustering of crAss-like phage genomes

Species-level virus operational taxonomic units (vOTUs) were defined under the 95% ANI over the 85% AF threshold using the Cluster_genomes_5.1.pl script (https://github.com/simroux/ClusterGenomes) relying on MUMmer 4.0.0beta2 (Kurtz et al., 2004; Roux *et al.*, 2019; Roux et al., 2017). If available, a tentatively complete genome belonging to a vOTU was selected to represent it in all subsequent analyses. Otherwise, the longest genome fragment belonging to the vOTU was used. Three potentially chimeric contigs (artifacts of sequencing and assembly or genuine biological entities such as proviruses integrated into bacterial genomes or recombinant viruses) representing vOTUs were recognized based on abrupt changes in nucleotide content and coverage by reads along the length of the contig and CheckV contamination assessment (Nayfach et al., 2020). The non-crAss-like regions of these contigs were excluded from consideration (Table S2), followed by re-calculation of the species-level clusters. Representatives of the vOTUs were further clustered into genus-level clusters under the 50% AF threshold by the Cluster_genomes_5.1.pl script (Adriaenssens and Brister, 2017).

### Open reading frame prediction

Open reading frames (ORFs) of every crAss-like phage representing a vOTU were independently predicted under the standard genetic code (translation table 11, c11) and under the two alternative genetic codes employed by crAss-like phages: c11 genetic code with stop codon TAG reassignment for glutamine and c11 genetic code with stop codon TGA reassignment for tryptophan (Yutin *et al.*, 2021). The prediction under the standard genetic code was made by Prodigal 2.6.3 (Hyatt *et al.*, 2010). Predictions under the alternative genetic codes were made by a specifically designed modification of Prodigal published in (Yutin *et al.*, 2021). The predictions were made for the individual genomes in the “meta” mode, unless the genome sequence was longer than 100 kb and the “single” mode was used.

### Functional annotation of crAss-like phage proteomes

Predicted proteins of crAss-like phages were functionally annotated by the hits to profiles representing the highly conserved proteins of crAss-like phages (Yutin *et al.*, 2021). The HMMER 3.3 program hmmsearch with an E-value threshold of 0.001 and the “--domtblout” output parameter was used (http://hmmer.org/). Hit envelope coordinates were used to define domain position. When envelopes of hits identifying the same domain overlapped, their coordinates were combined with the help of R package *IRanges* 2.22.2 (Lawrence et al., 2013).

### Determining the genetic code of crAss-like phage genomes

For each crAss-like phage representing a vOTU (Table S3), we considered three genome maps, each with ORFs predicted using a different genetic code as described above. The proteomes corresponding to each genome map were functionally annotated as described above. The genome maps (Material S2) were inspected to determine if a crAss-like phage may employ an alternative genetic code. If conserved genes of a crAss-like phage were split into multiple small ORFs under a standard genetic code, an alternative genetic code allowing for expression of the conserved genes with less interruption was assigned to the genome in question.

### Phylogeny reconstruction

A multiple sequence alignment (MSA) of the TerL protein sequences was built based on the MSA from (Yutin *et al.*, 2021). A subset of the TerL MSA from (Yutin *et al.*, 2021) corresponding to the crAss-like phages representing the vOTUs in this study was supplemented with TerL sequences of crAss-like phages that represent vOTUs and are specific to this study using MAFFT 7.453 with the “--add” parameter (Katoh and Standley, 2013). Protein fragments (potentially split by introns (Yutin *et al.*, 2021)) were joined, when necessary, with the help of R package *seqinr* 3.6-1 (Charif and Lobry), and the alignment was inspected with the help of Jalview 2.11.1.3 (Waterhouse et al., 2009). The MSA was used to reconstruct a phylogenetic tree using IQ-TREE 2.0.3 with 1000 replicates of ultrafast bootstrap (Minh et al., 2013; Nguyen et al., 2015). The tree was midpoint-rooted using R package *phangorn* 2.5.5 and visualized using R package *ape* 5.4-1 (Paradis and Schliep, 2019; Schliep, 2011). An identical procedure was used to reconstruct a phylogenetic tree based on the portal protein sequences.

### Assignment of crAss-like phage genomes to groups

Assignments of crAss-like phages to the five groups made in (Yutin *et al.*, 2021) were extended to the vOTUs of this study. First, the most recent common ancestor (MRCA) of all crAss-like phages that received an assignment to a group in (Yutin *et al.*, 2021) was identified on the TerL protein phylogenetic tree. Next, we verified that all descendants of the MRCA were either assigned to the group in question in (Yutin *et al.*, 2021) or were not part of that study. Finally, all descendants of the MRCA were assigned to the group in question. The procedure was conducted using R packages *phangorn* 2.5.5 and *ape* 5.4-1 (Paradis and Schliep, 2019; Schliep, 2011).

### Sequencing reads mapping to reference genomes

Sequencing reads of each individual sample, filtered and quality-trimmed as described above, were mapped to the crAss-like phage genomes representing vOTUs using Bowtie2 2.4.2 with a “--very-sensitive” parameter (Langmead and Salzberg, 2012). Breadth of genome coverage by reads was calculated using the BEDTools 2.29.2 command coverage (Quinlan and Hall, 2010). Depth of genome coverage by reads was calculated using the SAMtools 1.10 command depth (Li et al., 2009). Abundance of a vOTU in a sample was estimated as (*N* · 10^6^)/(*L* · *S*), where *N* is the number of reads mapped to a genome, *L* is the length of a genome and *S* is the number of sample reads after filtering and quality trimming.

### Analysis of nucleotide identity along the genomes

All genomes belonging to a genus-level cluster that were tentatively complete or nearly complete and represented a vOTU were collected with the help of Seqtk 1.3 (https://github.com/lh3/seqtk), reverse-transcribed and permuted where necessary to make their termini match those of a selected genome, and aligned using MAFFT 7.453 (Katoh and Standley, 2013). To estimate nucleotide identity between the selected genome and any other genome, a pairwise alignment was derived from the MSA using R package *Bio3D* 2.4-1 (Grant et al., 2006), its columns containing gaps in the selected genome row were removed, and the identity was recorded using a 1001 nt window sliding along the selected genome with a 200 nt step. The most 3’-terminal window was extended to include up to 200 3’-terminal nucleotides. A fragment of the delta27 genome MSA corresponding to the 50656-72821 nt of the OLNE01000081 genome (RNAP and VP02740 ORFs predicted under the TAG stop codon reassignment for glutamine) was extracted and used to reconstruct a phylogenetic tree using IQ-TREE 2.0.3 with the “-B 1000” parameter (Minh et al., 2020). The tree was subsequently midpoint-rooted using *phangorn* 2.5.5 (Schliep, 2011). The RNAP and VP02740 ORFs of OLNE01000081 were also used to query the set of 2616 crAss-like phage genomes and GenBank (14.02.2021, parameters “-db’nt’-remote”) using BLASTN 2.10.1+ (Altschul et al., 1990).

### Ecological measurements

The Bray-Curtis distances between samples were calculated based on the genus-level data using the function vegdist() from the R package *vegan* 2.5-6. Genus-level clusters that were not detected in any sample and samples without crAss-like phages were excluded from consideration. Distributions of the Bray-Curtis distances were visualized using R package *vioplot* 0.3.6. Wilcoxon signed-rank tests comparing intra-individual and inter-individual distances were performed using the R function wilcox.test() with the “alternative = ‘less’, paired = FALSE” parameters (R Core Team, 2020).

### Taxonomic profiling of microbial communities

The relative abundance of microbial taxa was estimated using MetaPhlAn 3.0.7 (Beghini et al., 2020).

### Correlation between abundances of crAss-like phages and microbes

Correlation between the relative abundance of microbial taxa (from kingdoms to species) and the abundance of crAss-like phages (assemblage, groups, genus-level clusters) was assessed using the R function cor.test() with the “method = ‘spearman’” parameter (R Core Team, 2020) for each cohort independently. The IBD cohort was represented by the selected 455 samples (see above). Only the taxa present in > 10 samples in a given cohort were considered. Meta-analysis of the results obtained for the LLD, 300OB and IBD cohorts was conducted using the R package *meta* 4.18-0 (Balduzzi et al., 2019), function metacor() employing Fisher’s Z-transformation of correlations and the Sidik-Jonkman between-study variance estimation method (parameters “sm =’ZCOR’, method.tau =’SJ’”). Multiple testing correction was conducted by the R function p.adjust() using the Benjamini-Hochberg procedure (Benjamini and Hochberg, 1995).

### Host prediction based on CRISPR spacers

The database of CRISPR spacers published in (Shmakov *et al.*, 2017) and the CRISPR-Cas++ spacers database (21.01.2021) (Pourcel et al., 2020) were independently compared to the set of 2616 crAss-like phage genomes using BLASTN 2.10.1+ (Altschul *et al.*, 1990) with the parameters “-task blastn-short -dust no -evalue 1-max_target_seqs 1000000”. A crAss-like genome was linked to a host if there was a spacer–protospacer match characterized by ≥ 95% identity over the length of the spacer, or multiple spacer–protospacer matches characterized by ≥ 80% identity over the length of each spacer (Roux et al., 2021). When multiple spacers of a host organism matched exactly the same region of a crAss-like phage genome, or if multiple regions of a crAss-like phage genome matched the same spacer of a host organism, a single spacer–protospacer match characterized by the highest bit-score was considered. Host taxonomy was retrieved from Genbank with the help of BioPerl 1.6.924 (Stajich et al., 2002).

### Metagenomic sequencing data cleaning prior to structural variant calling

We filtered out human genome reads and low-quality reads from raw sequencing data using KneadData 0.7.4, Bowtie 2.3.4.3 and Trimmomatic 0.39. Two main steps were included in this procedure: (1) removal of human genome reads by aligning metagenomic reads to the human reference genome (GRCh37/hg19) and (2) removal of low quality reads and adaptor sequences using Trimmomatic (parameters: SLIDINGWINDOW:4:20 MINLEN:50).

### Detection of bacterial structural variants

We detected bacterial structural variants (SVs) from cleaned metagenomic data using a previously published tool (Zeevi *et al.*, 2019), which is composed of two main steps. The first step aligns the cleaned metagenomic reads to reference bacterial genomes and solves ambiguously aligned reads using the iterative coverage-based read assignment algorithm. (2) The second step splits the reference genomes into genomic bins and examines the coverage of these bins to detect highly variable genomic segments for species with enough coverage and generate variable SV and deletion SV profiles. We used the reference database provided by SGVFinder, which is based on the proGenomes database (http://progenomes1.embl.de/) (Mende *et al.*, 2017). In total, we detected 7,999 deletion SVs and 3,559 variable SVs from 55 bacteria using default parameters. All bacterial species with SV calling were present in at least 5% of total samples.

### Association analysis between bacterial SVs and crAss-like phages

Before association analysis, all continuous variables were standardized to follow a standard normal distribution using empirical normal quantile transformation. Associations between SVs and relative abundances of crAss-like phages at genus and higher taxonomic levels (30 taxonomic clusters present in > 5% LLD samples) were assessed in the LLD, 300OB and IBD cohorts separately using linear models with the following formula:

crAss-like phage relative abundance ~ SV + Age + Sex + BMI + Reads number

The association results from different cohorts were then integrated via meta-analysis (random-effect model) and heterogeneity analysis. To control the FDR, Benjamini-Hochberg P-value corrections were performed using the p.adjust() function in R. The association analysis and P-value correction were conducted for vSVs and dSVs separately. Replicable significant associations were defined using the following criteria: FDR_meta_ < 0.05, P-value_heterogeneity_ > 0.05, P-value_LLD_ < 0.05, P-value_300OB_ < 0.05 and P-value_IBD_ < 0.05.

### Associations between crAss-like phages and human phenotypes in the LLD cohort

The analysis was applied to 1135 LLD cohort samples, 207 phenotypes (missing values imputed) and 30 taxonomic clusters of crAss-like phages present in > 5% LLD samples. The associations were analyzed using five methods: (1) Spearman correlation with abundance of crAss-like taxonomic clusters, (2) Spearman correlation with presence-absence of crAss-like taxonomic clusters, (3) unadjusted logistic regression, (4) logistic regression adjusted for the age and sex of the cohort participants and (5) logistic regression adjusted for the age and sex of the cohort participants and for the log-transformed abundance of the bacterial genera *Bacteroides*, *Prevotella*, *Porphyromonas* and *Parabacteroides*. Spearman correlation was assessed using R function cor.test() with the “method = ‘spearman’” parameter (R Core Team, 2020). Logistic regression was fitted using the R function glm() with the “family =’binomial’” parameter. Multiple testing correction was conducted using the R function p.adjust() using the Benjamini-Hochberg procedure (Benjamini and Hochberg, 1995).

### Association between crAss-like phages and disease phenotypes

The analysis involved 455 IBD cohort samples from patients without a stoma (colostomy or ileostomy) or ileoanal pouch, 298 300OB cohort samples and 1135 LLD cohort samples; 30 taxonomic clusters of crAss-like phages present in > 5% LLD samples were considered. Logistic regression was performed to compare the prevalence of the crAss-like phage taxonomic clusters between the following test groups: (1) LLD vs. IBD cohort, (2) LLD vs. 300OB cohort, (3) within IBD cohort: CD vs. UC, (4) within IBD cohort: exclusively colonic vs. ileal-inclusive disease location, (5) within 300OB cohort: absence vs. presence of metabolic syndrome. CrAss-like phage abundances were coded as 0 for absence (zero values) and 1 for presence (non-zero values). The logistic regression tests were performed using the R function glm() with the parameters “family = binomial(link =’logit’)”. Age and sex were included as covariates in the regression model for all comparisons. For comparisons involving IBD cohort samples, we also included carbohydrates and meat consumption, frequency of coffee consumption, usage of laxatives, and CgA and HBD-2 levels as covariates; these variables were not available for the 300OB cohort. In addition, we conducted a version of the LLD vs. IBD cohort and the LLD vs. 300OB cohort comparisons with the logistic regression further adjusted for the log-transformed abundance of the bacterial genera *Bacteroides*, *Prevotella*, *Porphyromonas* and *Parabacteroides*. Benjamini-Hochberg correction for multiple testing was applied using the R function p.adjust() with the parameter “method = ‘fdr’”. A significance threshold of FDR < 0.05 was used.

## Acknowledgments

We thank Natalya Yutin and Eugene V. Koonin for helpful discussion and advice. We would like to thank Lianmin Chen and Sergio Andreu-Sánchez for technical support, Kate McIntyre for editing the manuscript, the Center for Information Technology of the University of Groningen for their support and for providing access to the Peregrine high performance computing cluster and the volunteers of the LifeLines-DEEP, 300OB and IBD cohorts for their participation. S.G. and R.A.A.A.R. hold scholarships from the Graduate School of Medical Sciences, University of Groningen and the Junior Scientific Masterclass, University of Groningen, respectively. A.Z. is supported by the European Research Council (ERC) Starting Grant 715772, the Netherlands Organization for Scientific Research (NWO) VIDI grant 016.178.056 and NWO Gravitation grant ExposomeNL 024.004.017. J.F. is supported by NWO Gravitation grant Netherlands Organ-on-Chip Initiative 024.003.001, ERC Consolidator grant 101001678 and NWO VICI grant VI.C.202.022. N.P.R., M.G.N., J.F. and A.Z. are supported by the Netherlands Heart Foundation CVON grant 2018-27. R.K.W. is supported by the Seerave Foundation and the Dutch Digestive Foundation (16-14). C.W. is supported by NWO Gravitation grant 024.003.001 and NWO Spinoza Prize SPI 92-266.

## Author contributions

A.G. and A.Z. designed the study. N.P.R., M.G.N., C. W., R.K.W., J.F. and A.Z. provided the data. A.G., S.G., R.A.A.A.R., D.W., A.V.V. and A.K. performed data analysis. A.G., S.G., R.A.A.A.R., D.W., A.V.V., A.K. and A.Z. wrote the manuscript. All authors reviewed and edited the manuscript.

---

**Figure S1.**
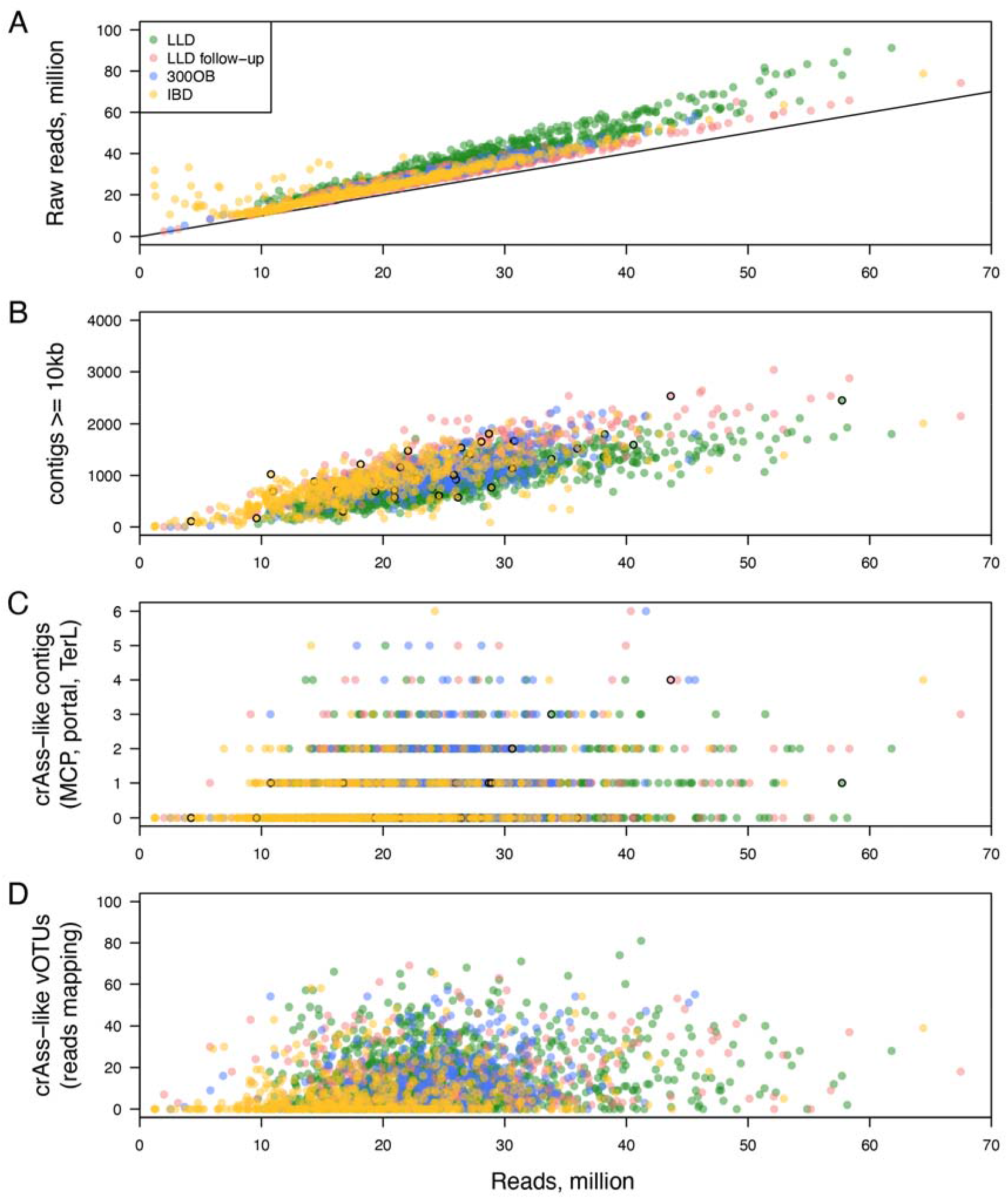
Quantitative characteristics of the crAss-like phage genome assembly and identification. For each sample, the number of sequencing reads after filtering and quality trimming is compared to the number of (**A**) raw sequencing reads, (**B**) assembled contigs of 10 kb or longer, (**C**) crAss-like contigs, and (**D**) crAss-like phage vOTUs detected based on reads mapping. Samples from different cohorts are distinguished by the color of the points. Black borders of points on panels B and C are used to indicate samples where BBMap was used for read error correction during contig assembly. Black line on panel A is the diagonal. Note that the two ends of each read were counted separately.

**Figure S2.**
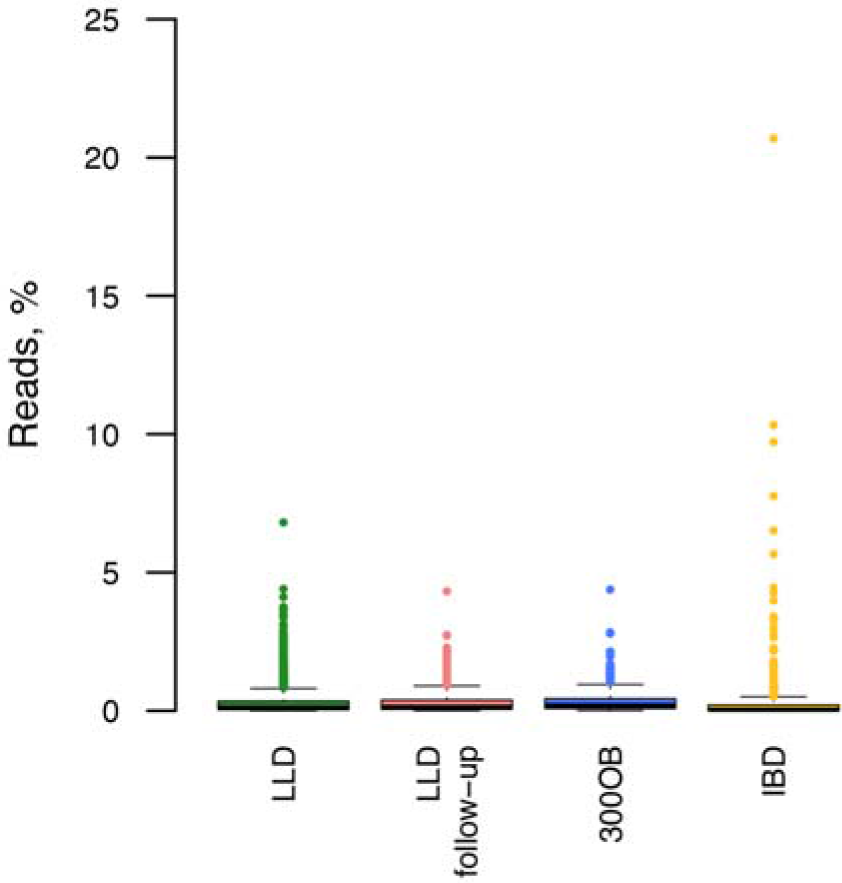
Percent of sequencing reads mapping to crAss-like phage genomes per sample in different cohorts. The reads were considered after filtering and trimming.

**Figure S3.**
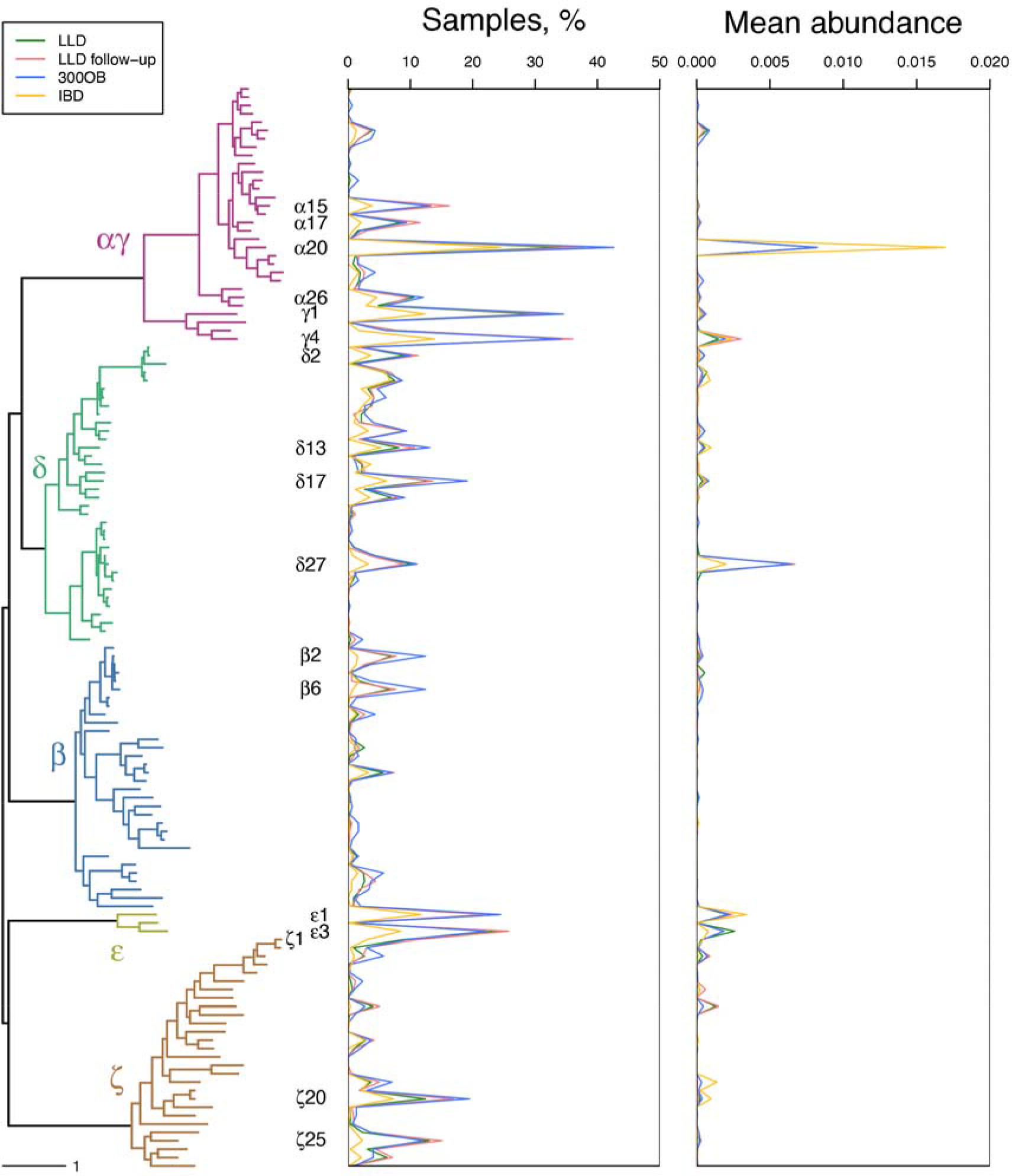
Abundance of crAss-like phage genus-level clusters in the four cohorts. For genus-level clusters belonging to the five groups associated with the human gut, the number of samples where a cluster was detected and its mean abundance in samples are presented per cohort. The genus-level clusters are ordered according to their position on a TerL-based phylogenetic tree, derived from the tree presented on Figure 1 by preserving a single tree tip per genus-level cluster from groups associated with the human gut. Names are indicated for the genus-level clusters present in > 10% of samples in any cohort.

**Figure S4.**
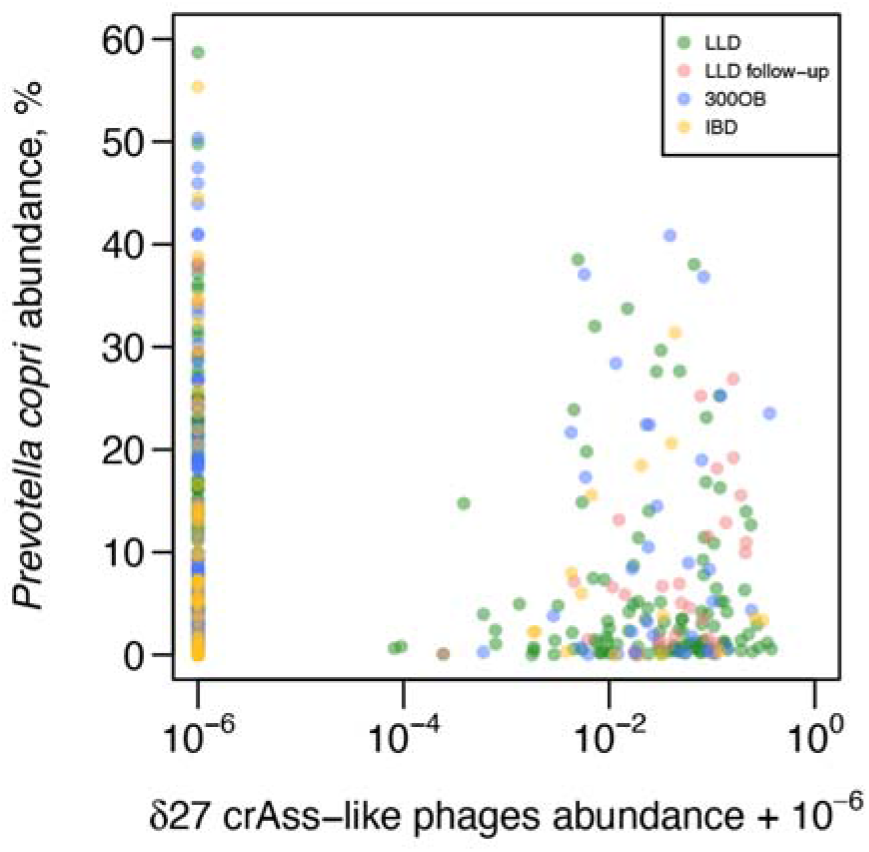
Relationship between the relative abundance of *P. copri* and crAss-like phages belonging to the delta27 genus-level cluster.

**Table S1 | Results of crAss-like phages detection in the control datasets.**

**Table S2 | CrAss-like phage genome fragments detected in this study**.

**Table S3 | Composition of the crAss-like phage vOTUs**.

**Table S4 | Characteristics of the crAss-like phage vOTUs**.

**Table S5 | Pairs of crAss-like phages and bacteria linked by CRISPR spacer matches**.

**Table S6 | Correlation between abundances of crAss-like phage and microbial taxa**. Results of the analysis of individual cohorts and the meta-analysis are presented. Rows are ordered based on the Spearman correlation coefficient (r) estimate obtained in meta-analysis. Assemblage, groups and genus-level clusters of crAss-like phages are designated as “crAss”, “crAss|group_name” and “crAss|group_name|cluster_name”, respectively. Microbial taxa are designated using the Metaphlan software format, with “k__”, “p__”, “c__”, “o__”, “f__”, “g__” and “s__” standing for kingdom, phylum, class, order, family, genus and species, respectively.

**Table S7 | Associations between bacterial structural variants and abundance of crAss-like phage taxonomic clusters**. Associations with variable and deletion SVs are presented on separate pages. All associations that were nominally significant in at least one cohort are shown, with replicable significant associations identified in meta-analysis highlighted by a green background. Assemblage, groups and genus-level clusters of crAss-like phages are designated as “crAss”, “crAss|group_name” and “crAss|group_name|cluster_name”, respectively. Column “N” specifies the number of samples where coverage of an SV by sequencing reads was sufficient to conduct the analysis and values of all phenotypes included in the linear model were available. Column “Unique N” specifies the number of unique abundance values observed for a crAss-like phage taxonomic cluster in samples satisfying the column “N” inclusion criteria. SV identifiers are displayed in the following format: <NCBI Taxonomy ID of the bacteria>.<NCBI BioProject Accession of the bacteria>:<genomic coordinates of the SV segments>, where the genomic coordinates of the SV segments are specified in kilobases and separated by semicolons, and start and end coordinates of each segment are separated by an underscore. Genes and gene products corresponding to each SV are specified according to the proGenomes database annotation.

**Table S8 | Associations between human phenotypes and abundance or presence-absence of crAss-like phages**. Results of the analysis conducted by five different methods are presented. Significant associations (FDR < 0.05) are highlighted by a green background. Assemblage, groups and genus-level clusters of crAss-like phages are designated as “crAss”, “crAss|group_name” and “crAss|group_name|cluster_name”, respectively.

**Table S9 | Prevalence of crAss-like phage taxonomic clusters in the context of IBD and obesity**. Results of the analyses comparing the prevalence of crAss-like phage taxonomic clusters between the following groups are each presented on a separate page: (1) LLD vs. IBD cohort, (2) LLD vs. 300OB cohort, (3) within IBD cohort: CD vs. UC diagnosis, (4) within IBD cohort: exclusively colonic vs. ileum-inclusive disease location, (5) within 300OB cohort: absence vs. presence of metabolic syndrome. Significant associations (FDR < 0.05) are highlighted by a green background. Assemblage, groups and genus-level clusters of crAss-like phages are designated as “crAss”, “crAss|group_name” and “crAss|group_name|cluster_name”, respectively.

**Material S1 | FASTA file with 1556 crAss-like genome fragments detected in the LLD, LLD follow-up, 300OB and IBD cohorts**.

**Material S2 | Genome maps of crAss-like phages representing vOTUs**. For each genome, three genome maps are presented on a separate page: top, genome map with ORFs predicted under the standard genetic code; middle, ORFs predicted under the code with stop codon TAG reassignment for glutamine; bottom, ORFs predicted under the code with stop codon TGA reassignment for tryptophan. The name of the genetic code assigned to the genome is highlighted by bold font. A genome is presented as a black rectangular contour, three forward and three reverse frames of the genome are indicated, and ORFs are presented as grey bars. ORF regions displaying similarity to conserved domains of the crAss-like phages are indicated by colored bars (see legend on the first page: Tstab, tail stabilization protein; PolA and PolB, DNA polymerase family A and B, respectively; RNApA, RNApB and RNApB’, RNA polymerase subunits A, B and B’, respectively; BACON, Bacteroides-associated carbohydrate-binding often N-terminal domain; RT_G2, reverse transcriptase with group II intron origin (Yutin *et al.*, 2021)). Regular and cumulative GC and AT skew are presented below the genome maps. For the crAss-like phages belonging to one of the five groups associated with the human gut, the name of the genus-level cluster is specified at the top right corner of the page.

**Material S3 | TerL-based phylogenetic tree of crAss-like phages**. The tree is midpoint pseudo-rooted. Tree tips correspond to crAss-like phages representing vOTUs. Seven vOTUs represented by partial genomes lacking a TerL gene are not shown. The five clades of crAss-like phages associated with the human gut are designated by the color of the branches. Genus-level clusters of the crAss-like phages belonging to the five groups associated with the human gut are depicted next to the tree. Bootstrap support (BP) values are indicated by black, grey and white circles.

**Material S4 | Portal-based phylogenetic tree of crAss-like phages**. See the legend of Material S3 for details.

**Material S5 | Depth of crAss-like phage genomes coverage by sequencing reads**. Information about each crAss-like phage genome representing a vOTU and detected at least once is presented on a separate page. Each of the four top plots represents a cohort. Each transparent grey line on these plots represents mean depth of the genome coverage by reads from a sample. On the bottom plot, the content of the four nucleotides is shown along the genome. All plots were produced with a 1001 nt sliding window. For the crAss-like phages belonging to one of the five groups associated with the human gut, the name of the genus-level cluster is specified at the top right corner of the page.

**Material S6 | Coverage anomaly and nucleotide sequence divergence in the transcription gene module of alpha6, beta8 and zeta9 genomes**. Information about each genus-level cluster is presented on a separate page and is based on a selected genome: OLOZ01000098 for alpha6, IAS_virus_KJ003983 for beta8 and NL_crAss000848 for zeta9. Each page contains the name of a genus-level cluster in the top-right corner and three plots. Top plot, depth of the selected genome coverage by reads from the LLD cohort (for alpha6 and zeta9) or 300OB cohort (for beta8, as 300OB is the only cohort where this genus-level cluster was detected). Each transparent grey line corresponds to a sample and represents mean coverage depth in a 1001 nt sliding window. Middle plot, genome map of the selected genome (see legend of the Material S2 for details). Bottom plot, nucleotide identity between the selected genome and other (nearly) complete genomes belonging to the same genus-level cluster. Nucleotide identity was calculated using a 1001 nt sliding window. Data for genomes belonging to different vOTUs are distinguished by color.

