## Supplementary material for "Discovery, diversity and functional associations of crAss-like phages in human gut metagenomes from four Dutch cohorts": Material S2

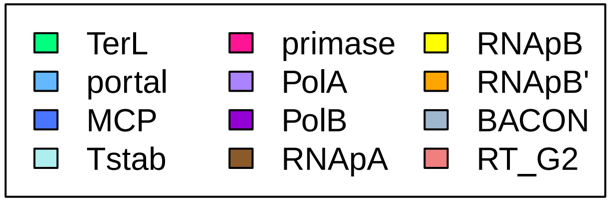
TerL
portal
MCP
Tstab
primase
PolA
PolB
RNApA
RNApB
RNApB'
BACON
RT_G2

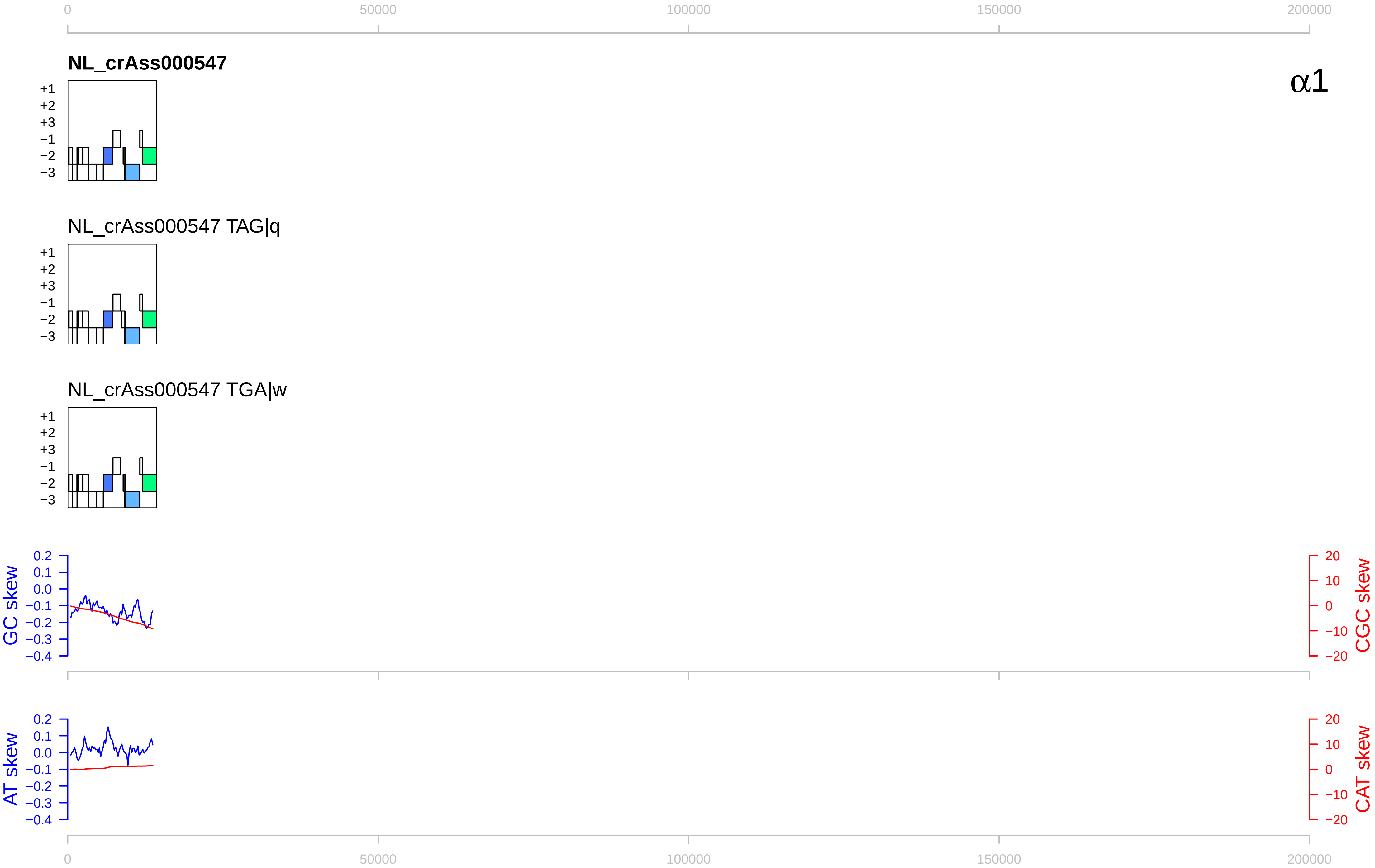

0
50000
100000
150000
200000
NL_crAss000547
α1
+1
+2
+3
−1
−2
−3
NL_crAss000547 TAG|q
NL_crAss000547 TGA|w
GC skew
CGC skew
0.2
0.1
0.0
−0.1
−0.2
−0.3
−0.4
20
10
0
−10
−20
AT skew
CAT skew

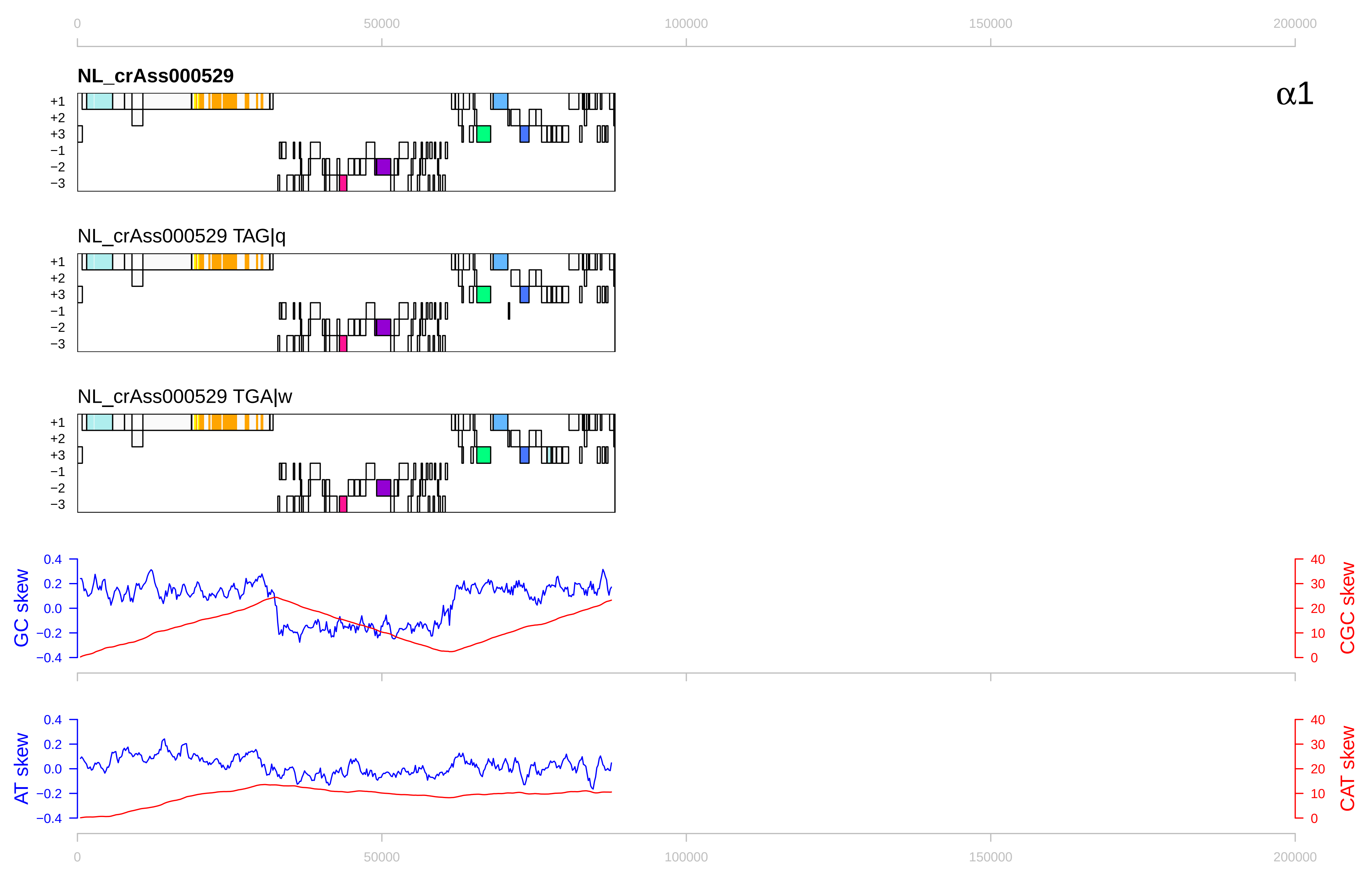

0
50000
100000
150000
200000
NL_crAss000529
α1
+1
+2
+3
−1
−2
−3
NL_crAss000529 TAG|q
NL_crAss000529 TGA|w
GC skew
CGC skew
AT skew
CAT skew
0.4
0.2
0.0
−0.2
−0.4
40
30
20
10
0

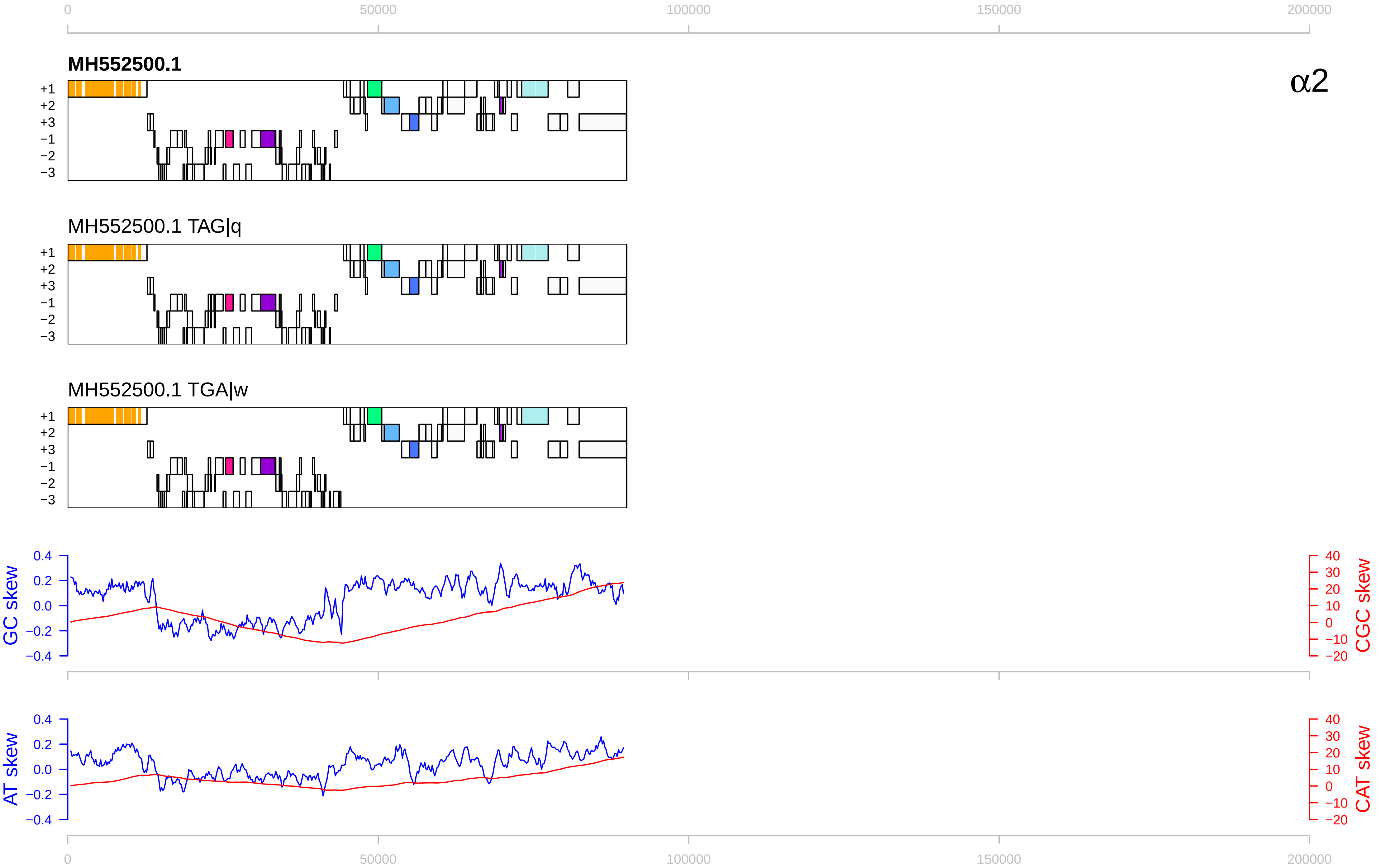
0
50000
100000
150000
200000
MH552500.1
α2
+1
+2
+3
−1
−2
−3
MH552500.1 TAG|q
MH552500.1 TGA|w
GC skew
CGC skew
0.4
0.2
0.0
−0.2
−0.4
40
30
20
10
0
−10
−20
AT skew
CAT skew

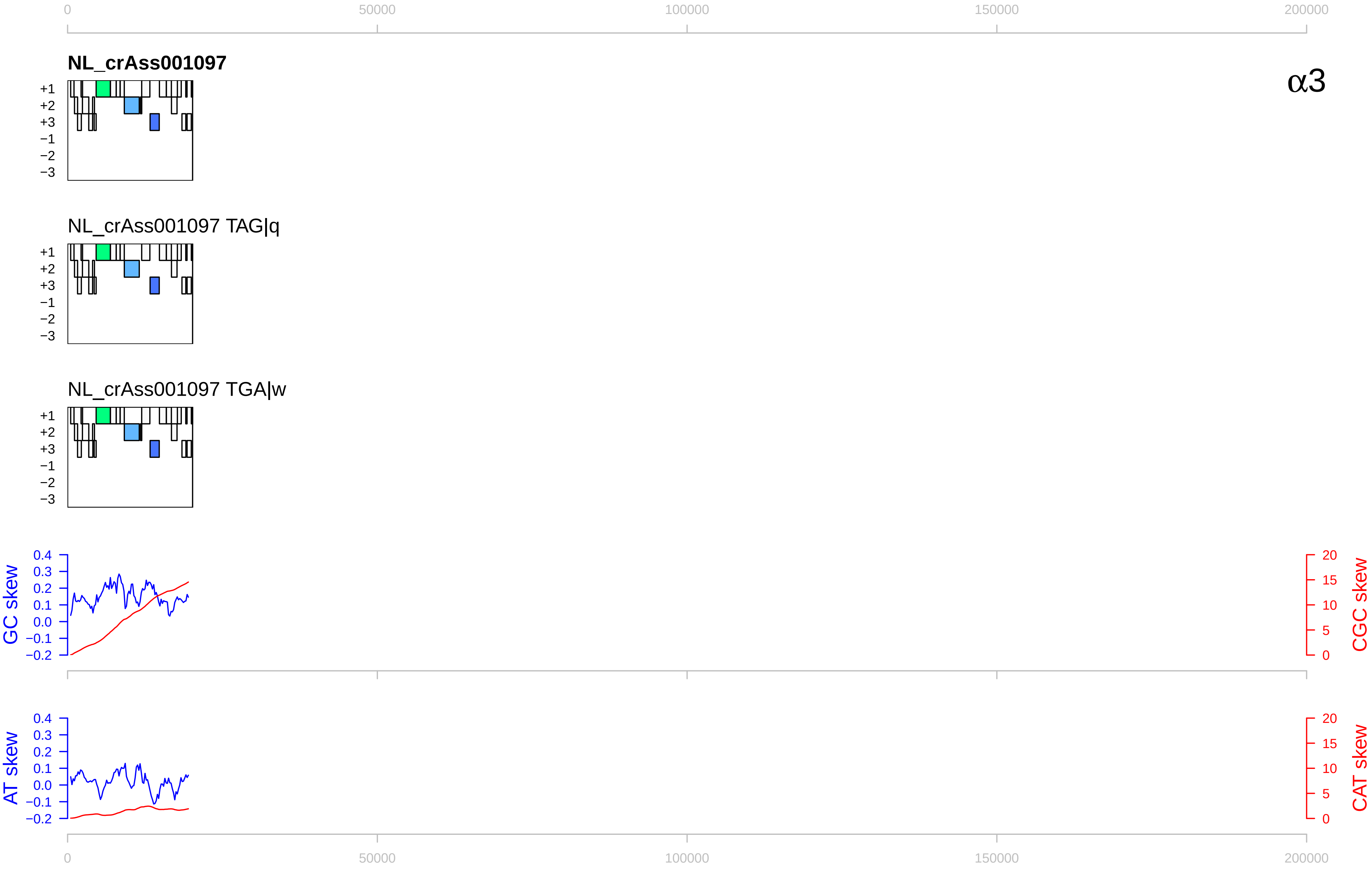

0
50000
100000
150000
200000
NL_crAss001097
α3
+1
+2
+3
−1
−2
−3
NL_crAss001097 TAG|q
NL_crAss001097 TGA|w
GC skew
CGC skew
AT skew
CAT skew
0.4
0.3
0.2
0.1
0.0
−0.1
−0.2
20
15
10
5
0

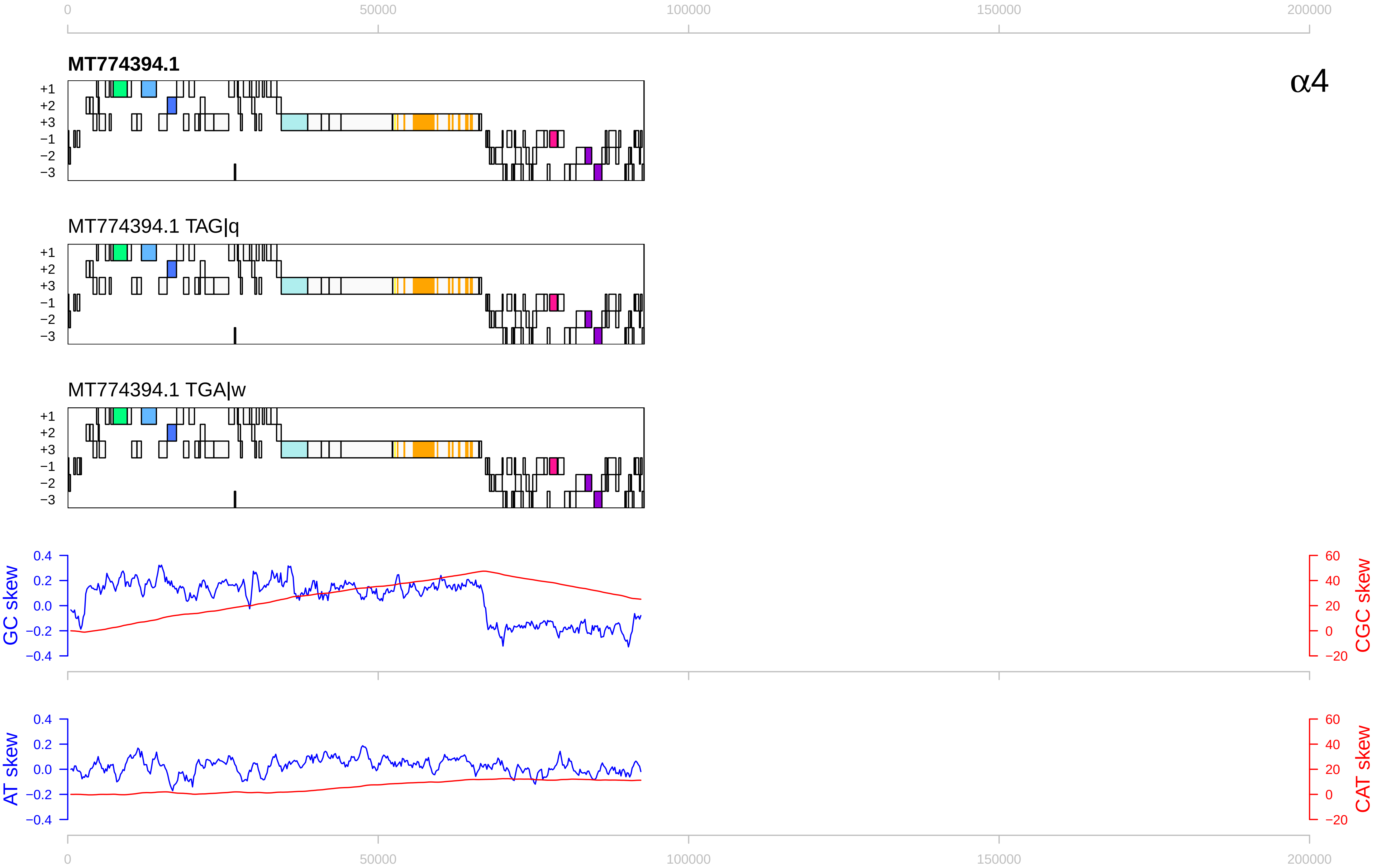

0
50000
100000
150000
200000
MT774394.1
α4
+1
+2
+3
−1
−2
−3
MT774394.1 TAG|q
MT774394.1 TGA|w
GC skew
0.4
0.2
0.0
−0.2
−0.4
CGC skew
60
40
20
0
−20
AT skew
CAT skew

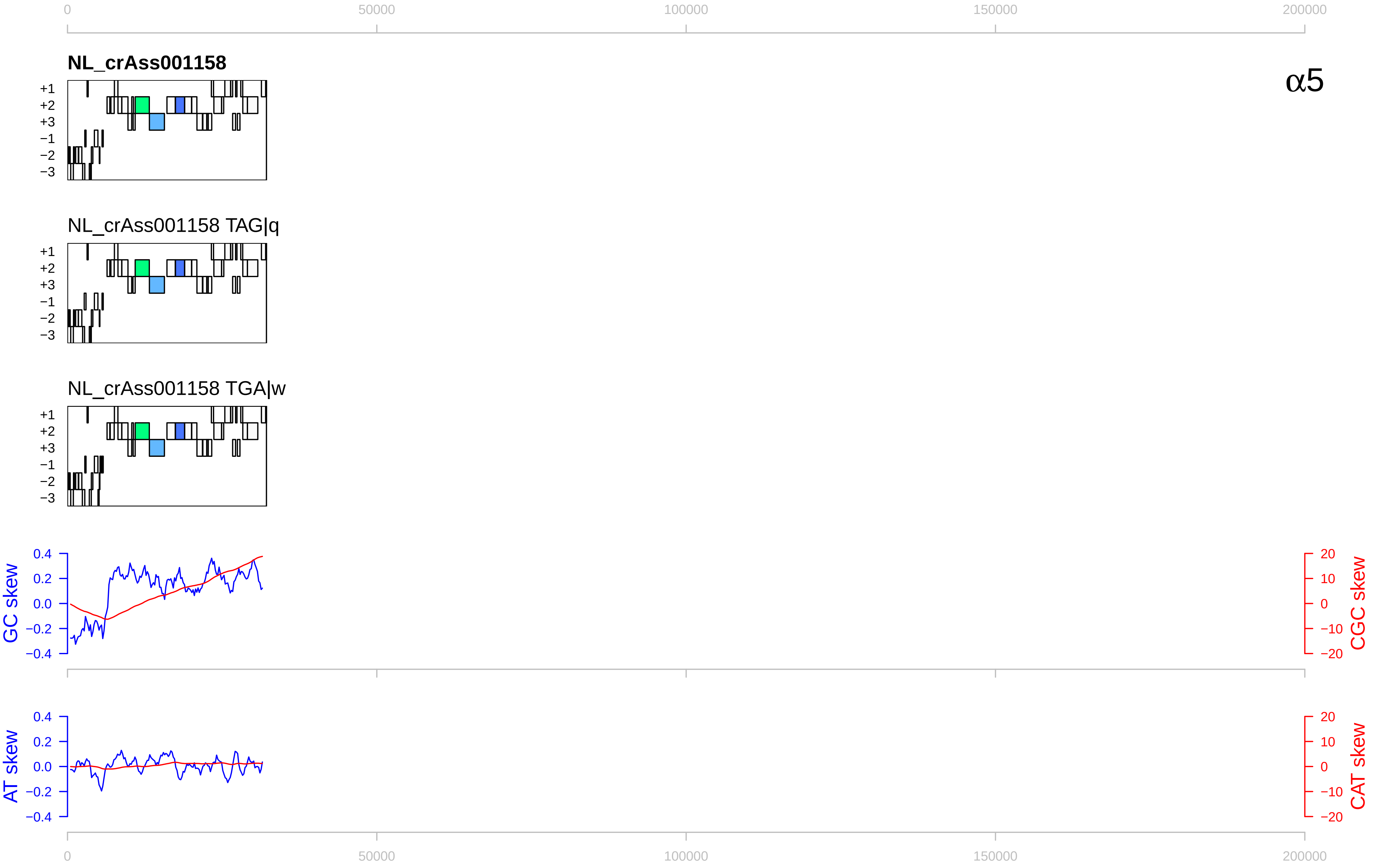

0
50000
100000
150000
200000
NL_crAss001158
α5
+1
+2
+3
−1
−2
−3
NL_crAss001158 TAG|q
NL_crAss001158 TGA|w
GC skew
0.4
0.2
0.0
−0.2
−0.4
CGC skew
20
10
0
−10
−20
AT skew
CAT skew

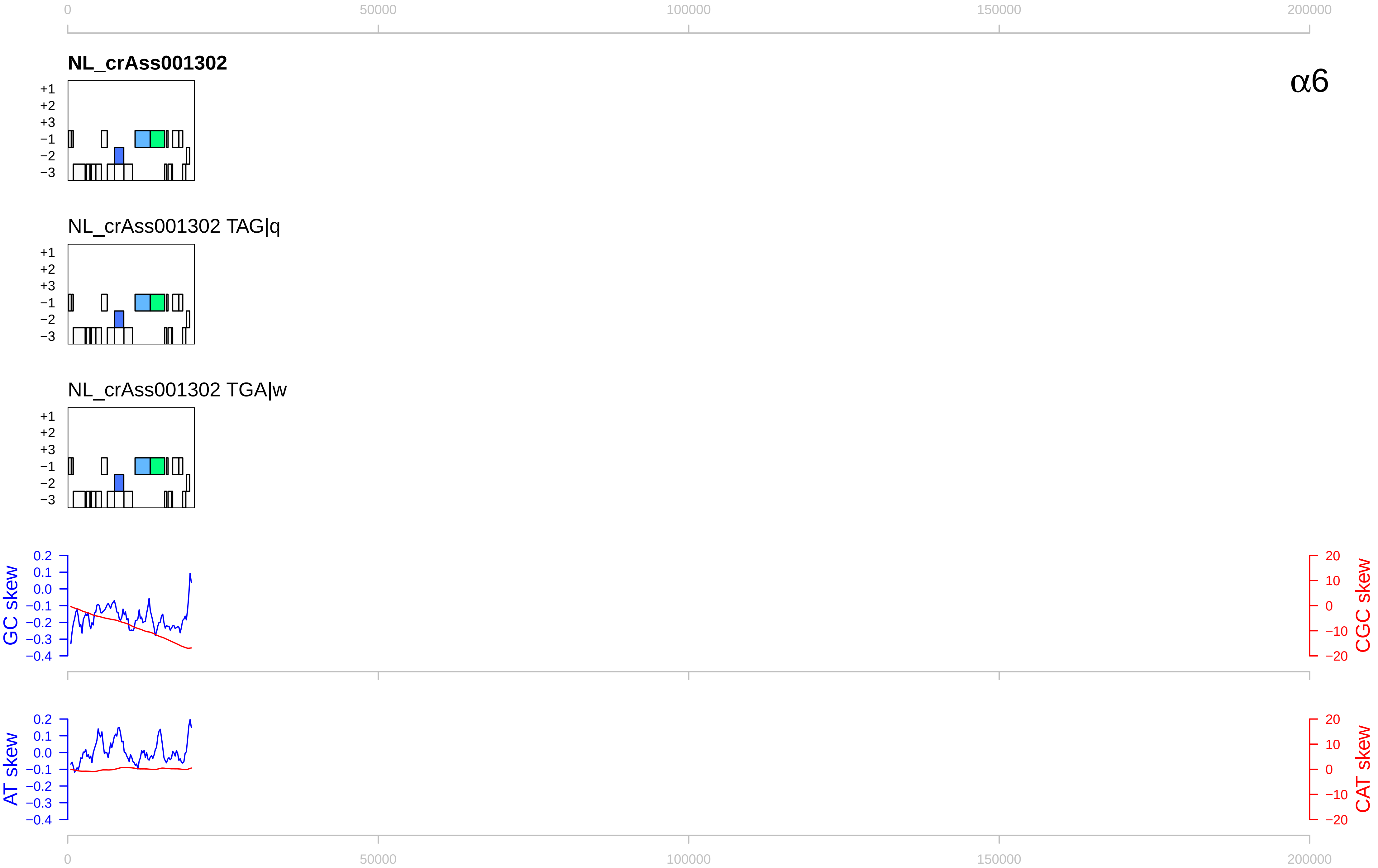

0
50000
100000
150000
200000
NL_crAss001302
α6
+1
+2
+3
−1
−2
−3
NL_crAss001302 TAG|q
NL_crAss001302 TGA|w
GC skew
CGC skew
AT skew
CAT skew
0.2
0.1
0.0
−0.1
−0.2
−0.3
−0.4
20
10
0
−10
−20

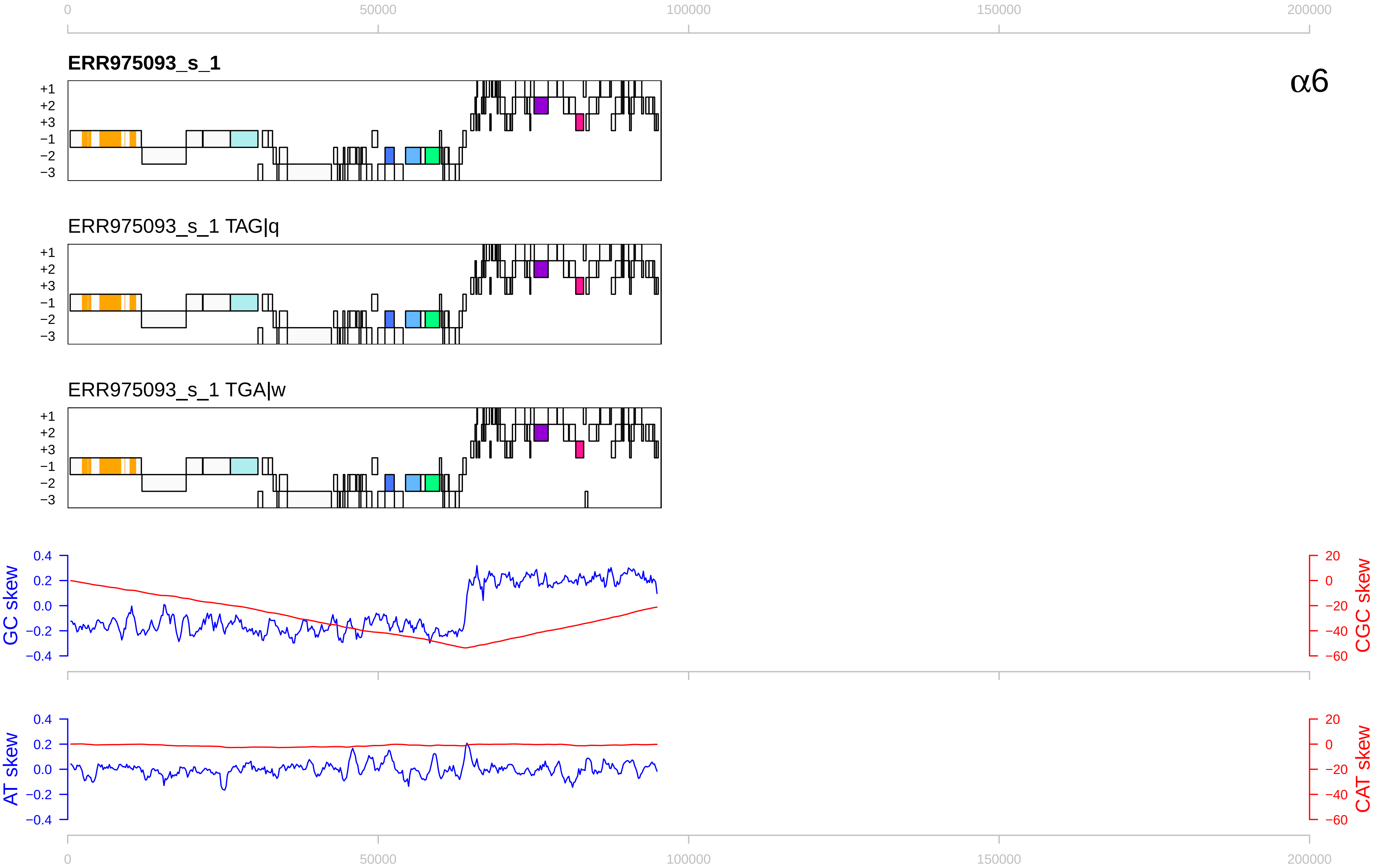

α6
ERR975093_s_1
ERR975093_s_1 TAG|q
ERR975093_s_1 TGA|w
+1
+2
+3
−1
−2
−3
GC skew
CGC skew
AT skew
CAT skew
0
50000
100000
150000
200000

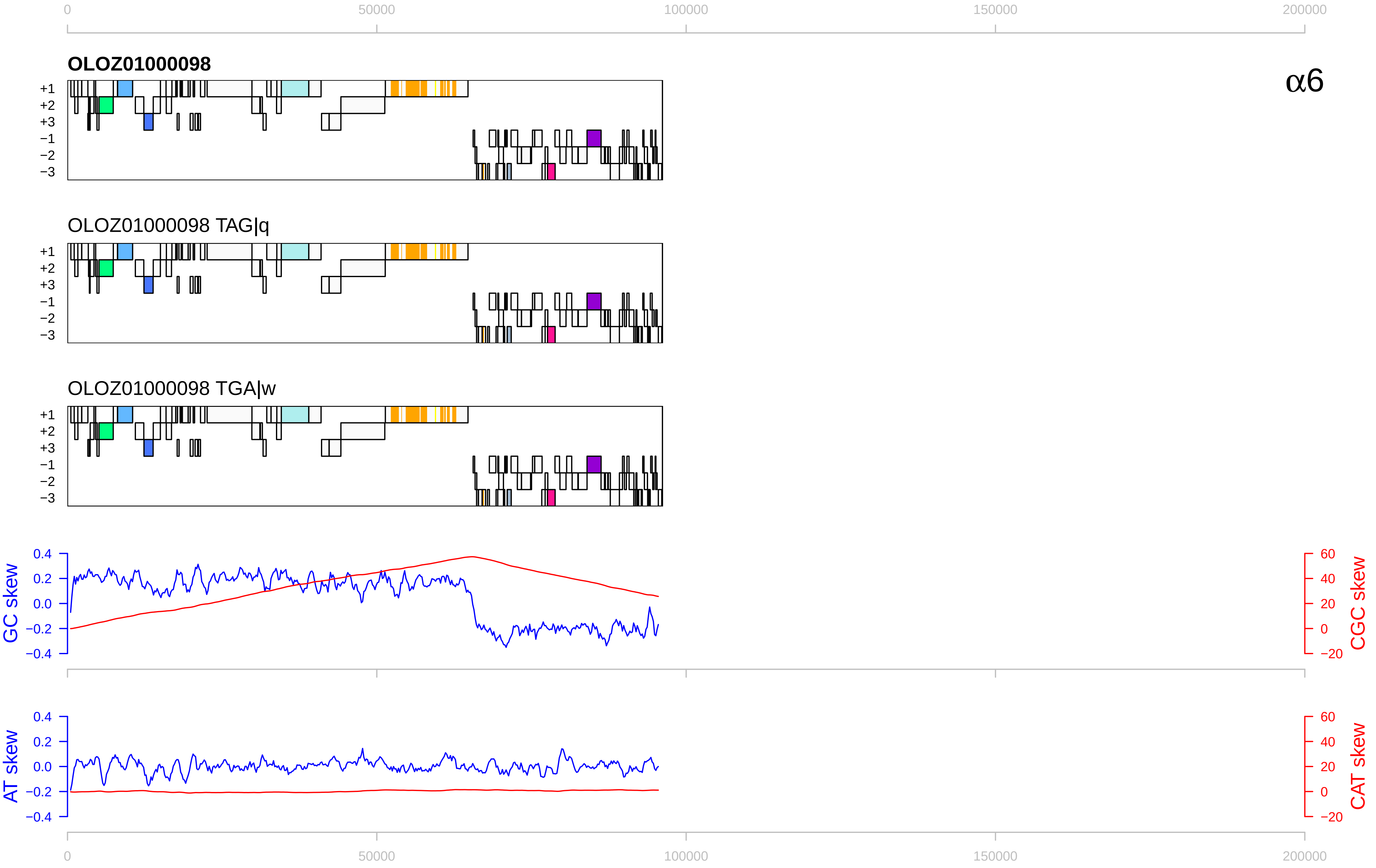

0
50000
100000
150000
200000
OLOZ01000098
α6
+1
+2
+3
−1
−2
−3
OLOZ01000098 TAG|q
OLOZ01000098 TGA|w
GC skew
CGC skew
AT skew
CAT skew
0.4
0.2
0.0
−0.2
−0.4
60
40
20
0
−20

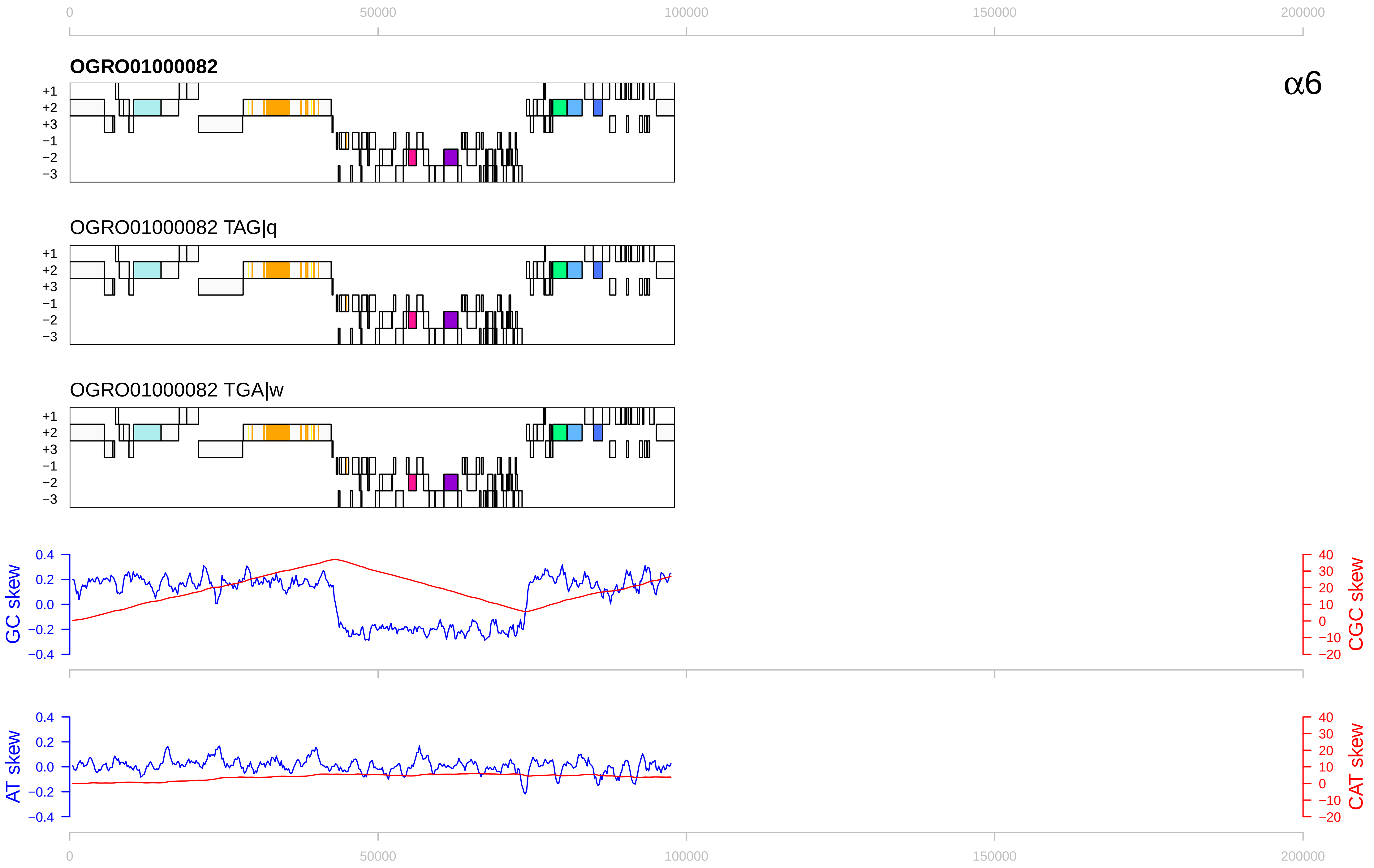
0
50000
100000
150000
200000
OGRO01000082
α6
+1
+2
+3
−1
−2
−3
OGRO01000082 TAG|q
OGRO01000082 TGA|w
GC skew
CGC skew
0.4
0.2
0.0
−0.2
−0.4
40
30
20
10
0
−10
−20
AT skew
CAT skew

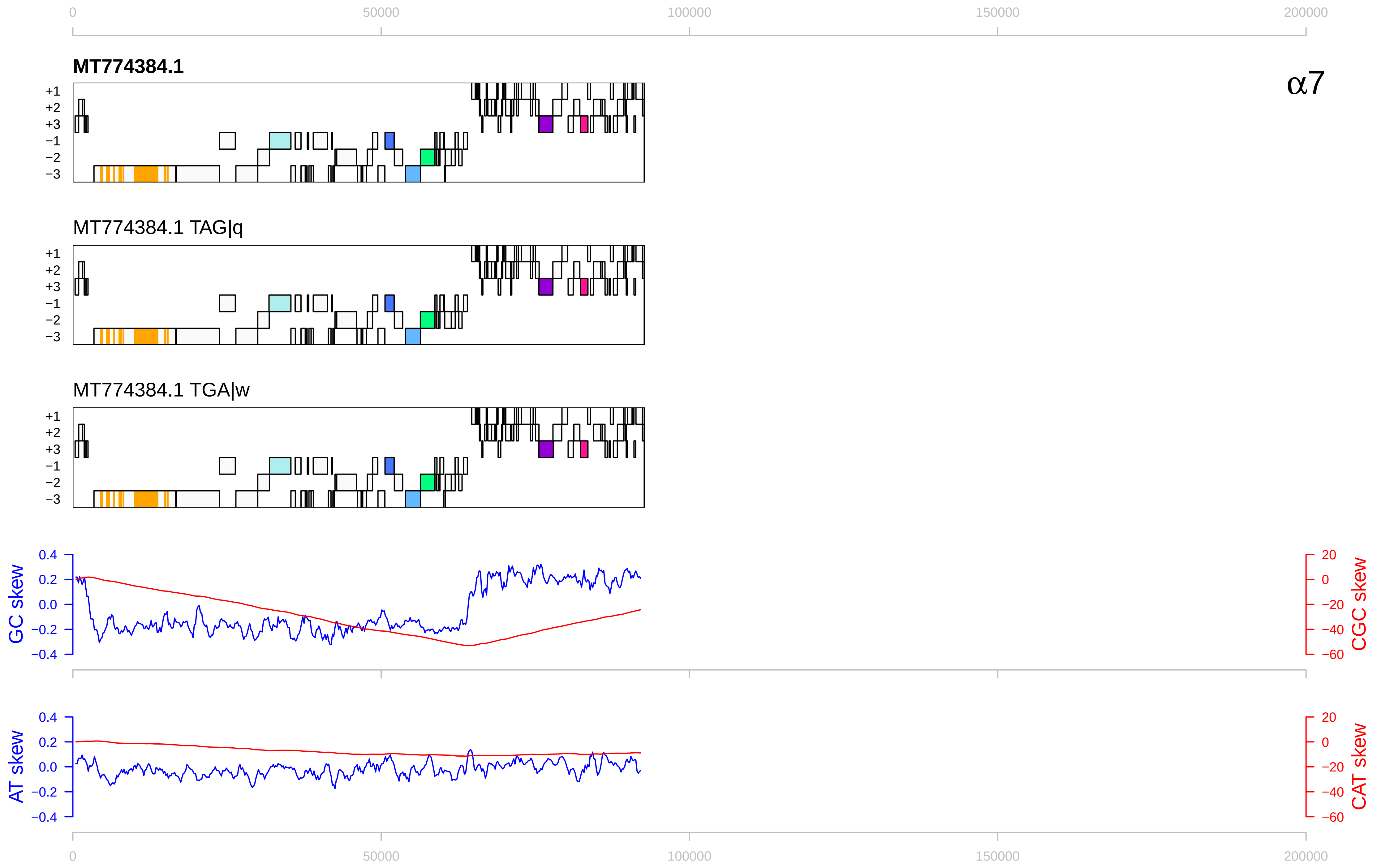

0
50000
100000
150000
200000
MT774384.1
α7
+1
+2
+3
−1
−2
−3
MT774384.1 TAG|q
MT774384.1 TGA|w
GC skew
CGC skew
0.4
0.2
0.0
−0.2
−0.4
20
0
−20
−40
−60
AT skew
CAT skew

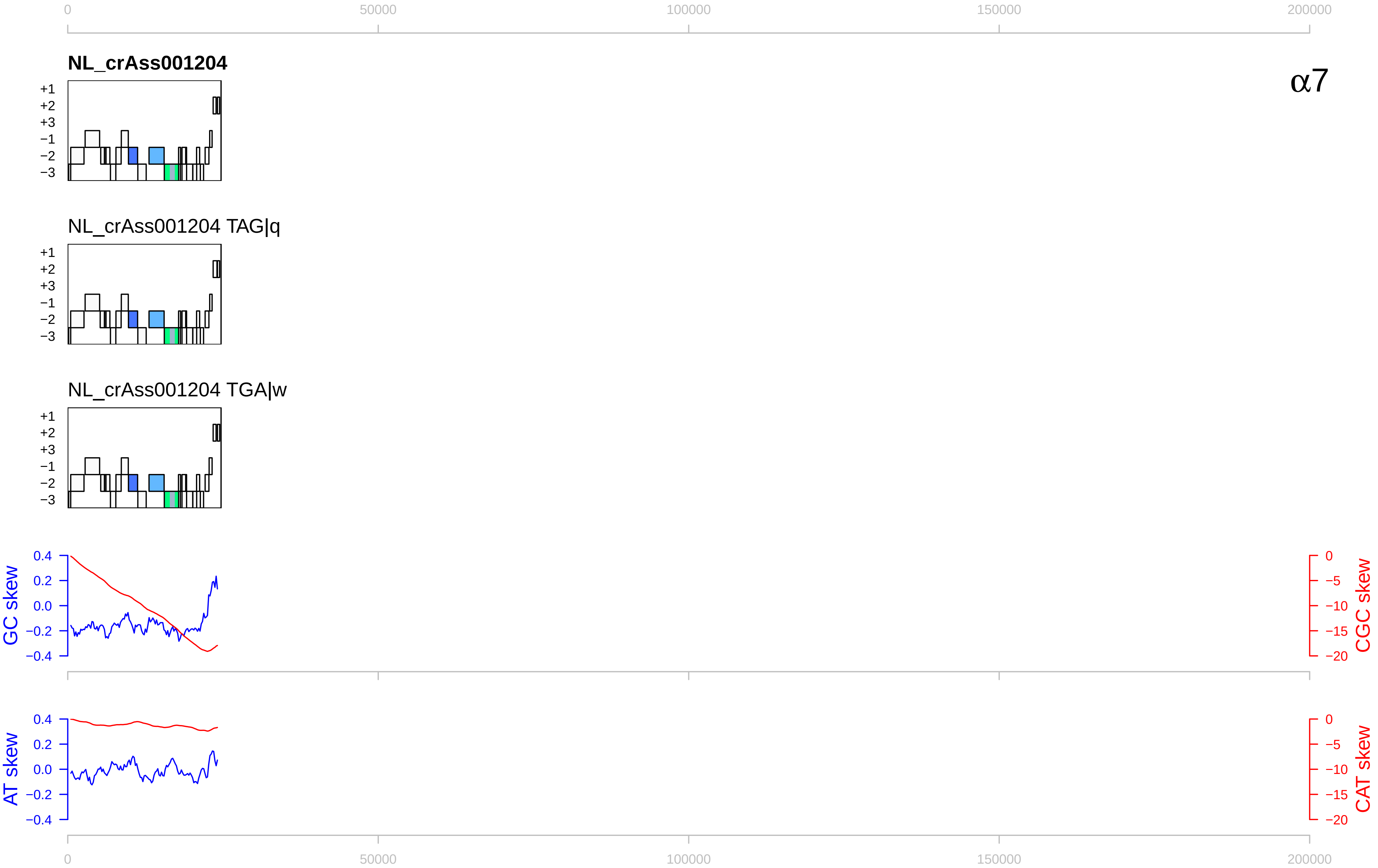
0
50000
100000
150000
200000
NL_crAss001204
α7
+1
+2
+3
−1
−2
−3
NL_crAss001204 TAG|q
NL_crAss001204 TGA|w
GC skew
0.4
0.2
0.0
−0.2
−0.4
CGC skew
0
−5
−10
−15
−20
AT skew
CAT skew

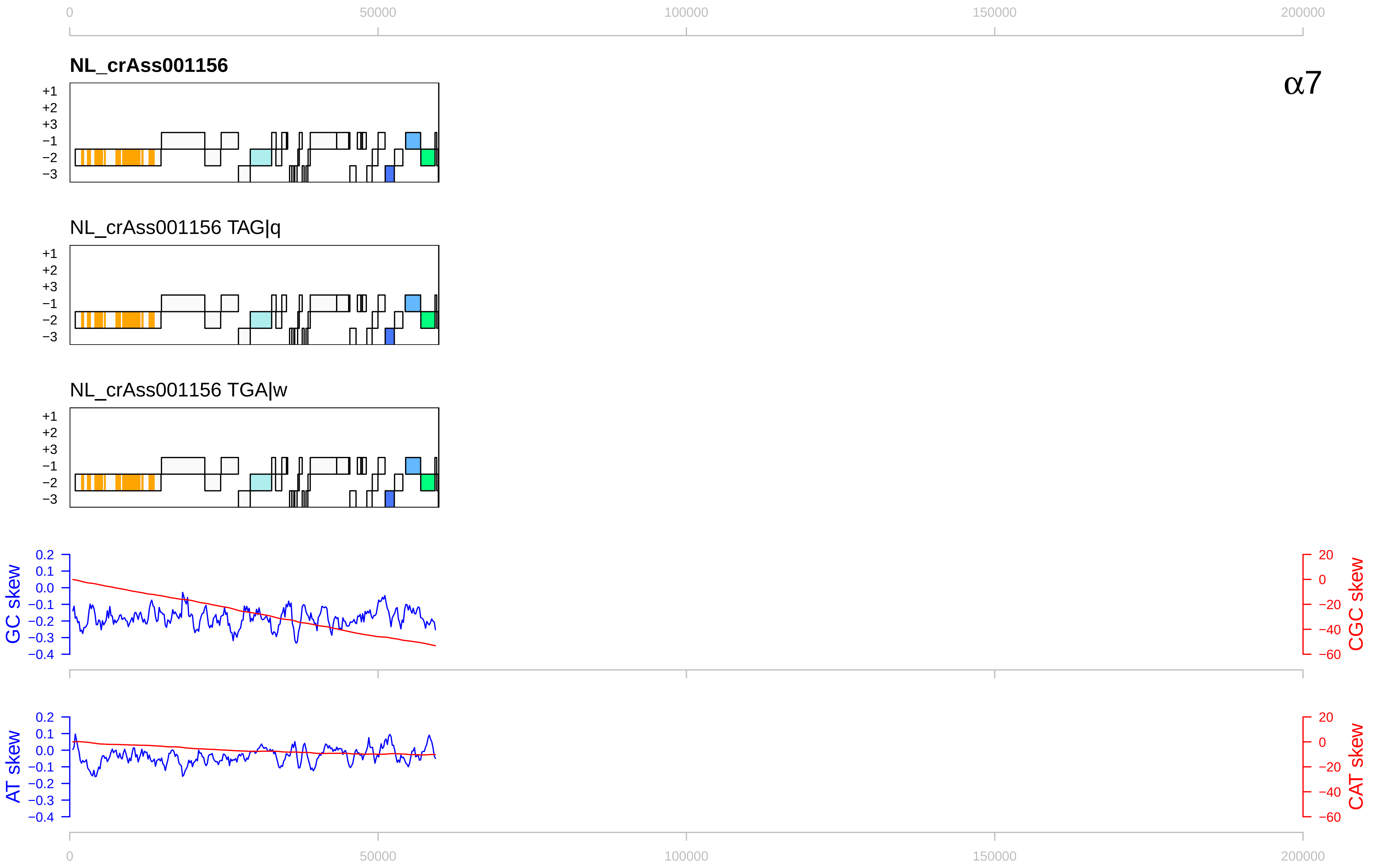

0
50000
100000
150000
200000
NL_crAss001156
α7
+1
+2
+3
−1
−2
−3
NL_crAss001156 TAG|q
NL_crAss001156 TGA|w
GC skew
0.2
0.1
0.0
−0.1
−0.2
−0.3
−0.4
CGC skew
20
0
−20
−40
−60
AT skew
CAT skew

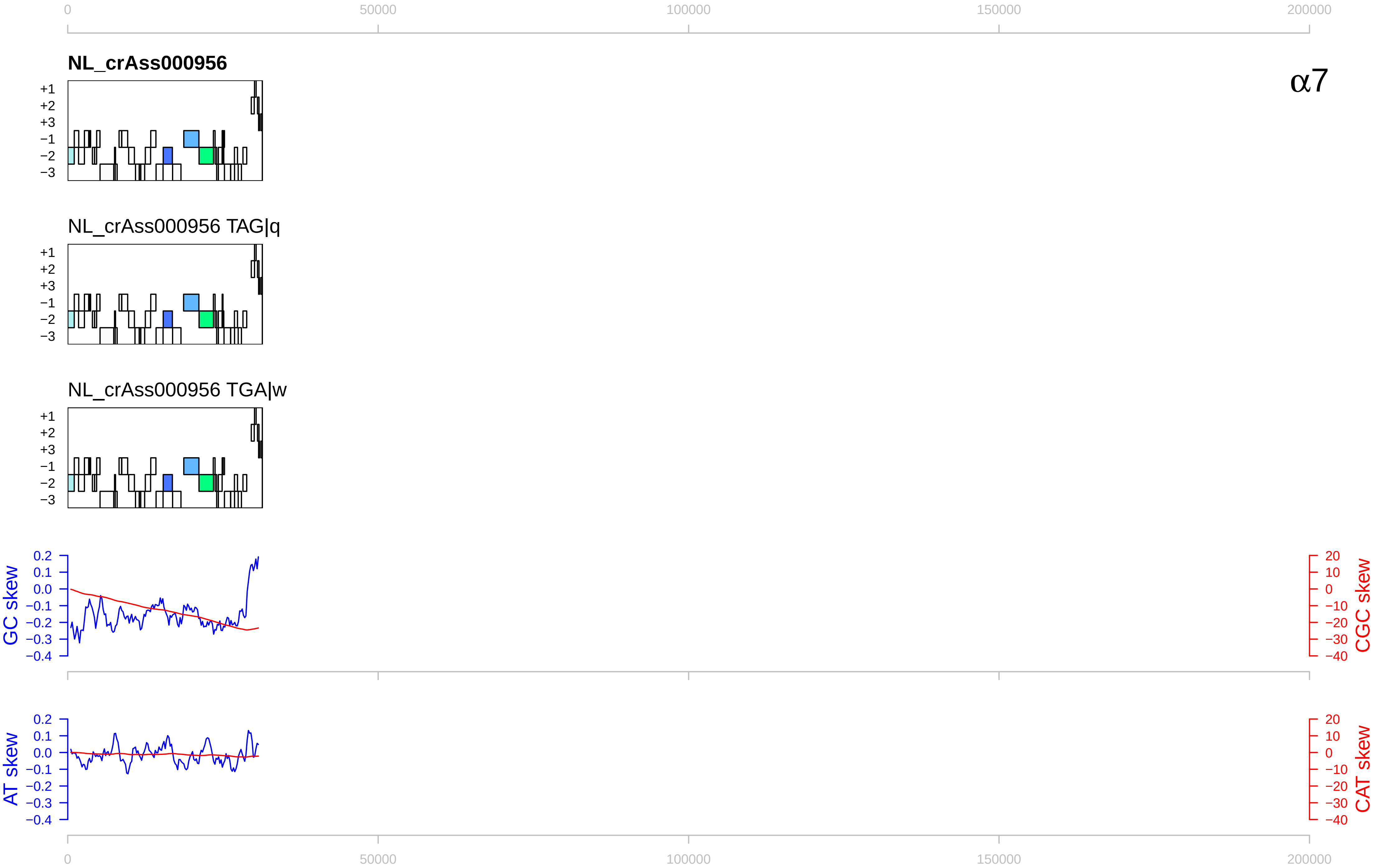

0
50000
100000
150000
200000
NL_crAss000956
α7
+1
+2
+3
−1
−2
−3
NL_crAss000956 TAG|q
NL_crAss000956 TGA|w
GC skew
CGC skew
AT skew
CAT skew
0.2
0.1
0.0
−0.1
−0.2
−0.3
−0.4
20
10
0
−10
−20
−30
−40

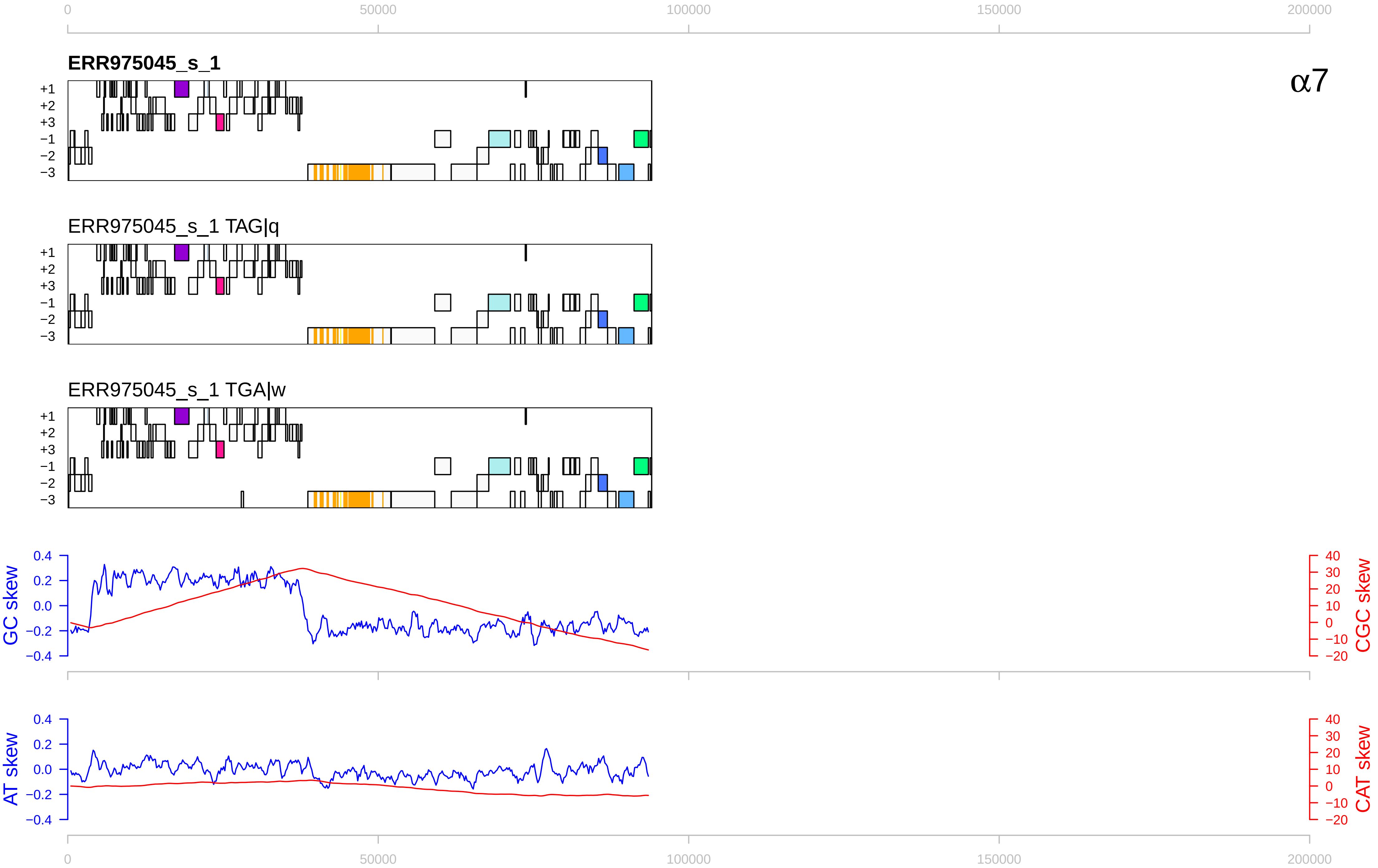
ERR975045_s_1
α7
ERR975045_s_1 TAG|q
ERR975045_s_1 TGA|w
+1
+2
+3
−1
−2
−3
GC skew
CGC skew
AT skew
CAT skew
0
50000
100000
150000
200000

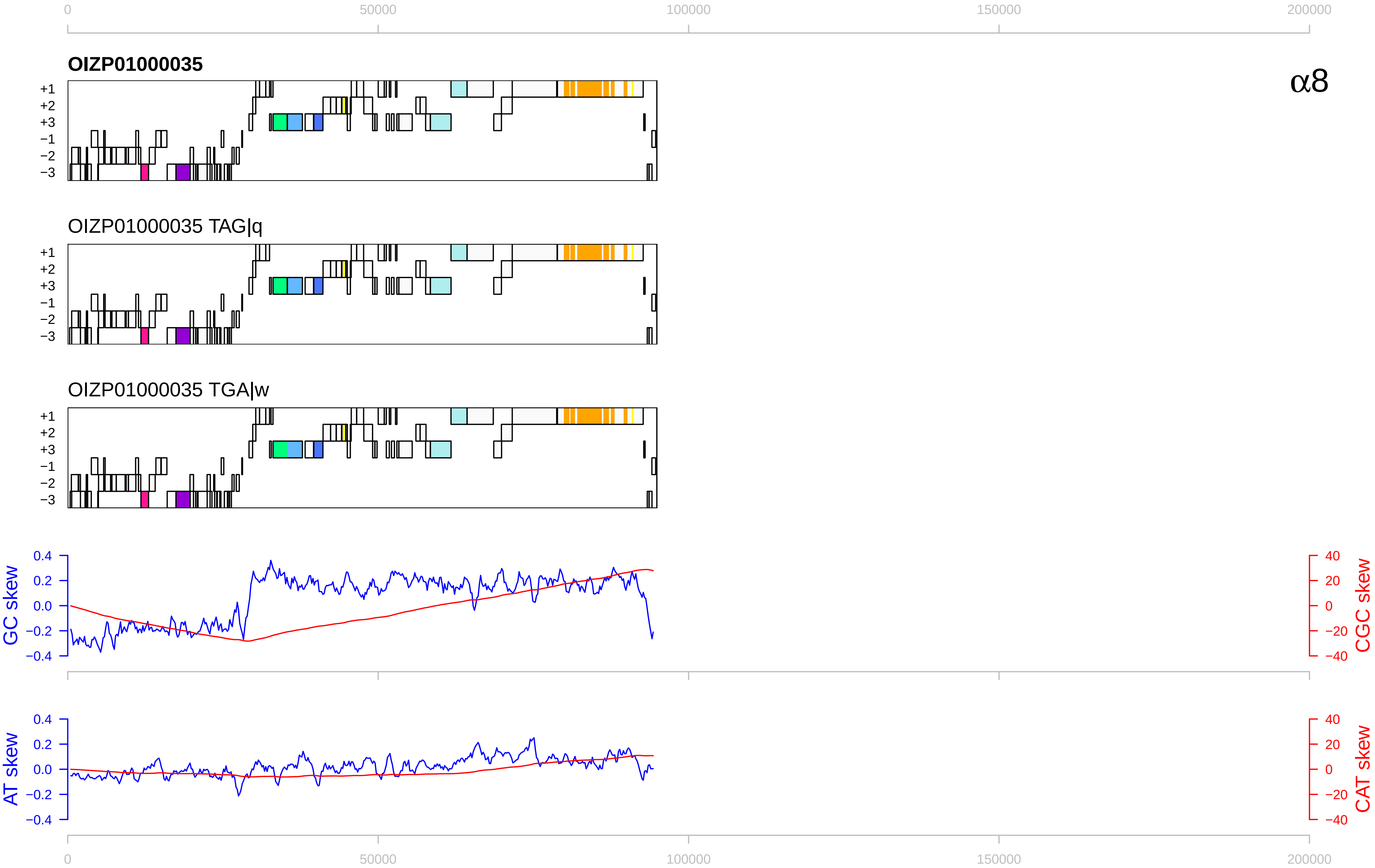

0
50000
100000
150000
200000
OIZP01000035
α8
+1
+2
+3
−1
−2
−3
OIZP01000035 TAG|q
OIZP01000035 TGA|w
GC skew
CGC skew
AT skew
CAT skew
0.4
0.2
0.0
−0.2
−0.4
40
20
0
−20
−40

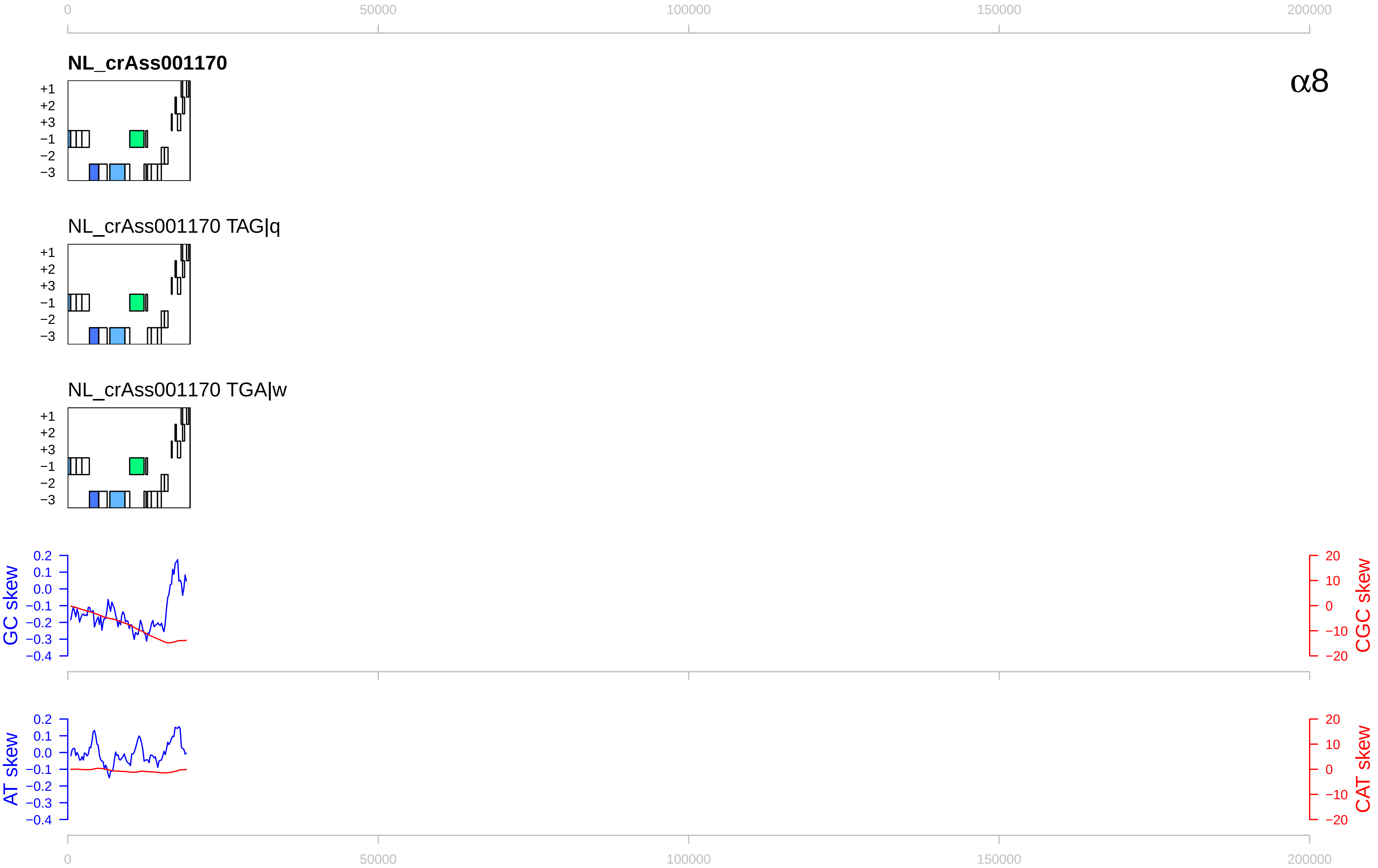

0
50000
100000
150000
200000
NL_crAss001170
α8
+1
+2
+3
−1
−2
−3
NL_crAss001170 TAG|q
NL_crAss001170 TGA|w
GC skew
CGC skew
AT skew
CAT skew
0.2
0.1
0.0
−0.1
−0.2
−0.3
−0.4
20
10
0
−10
−20

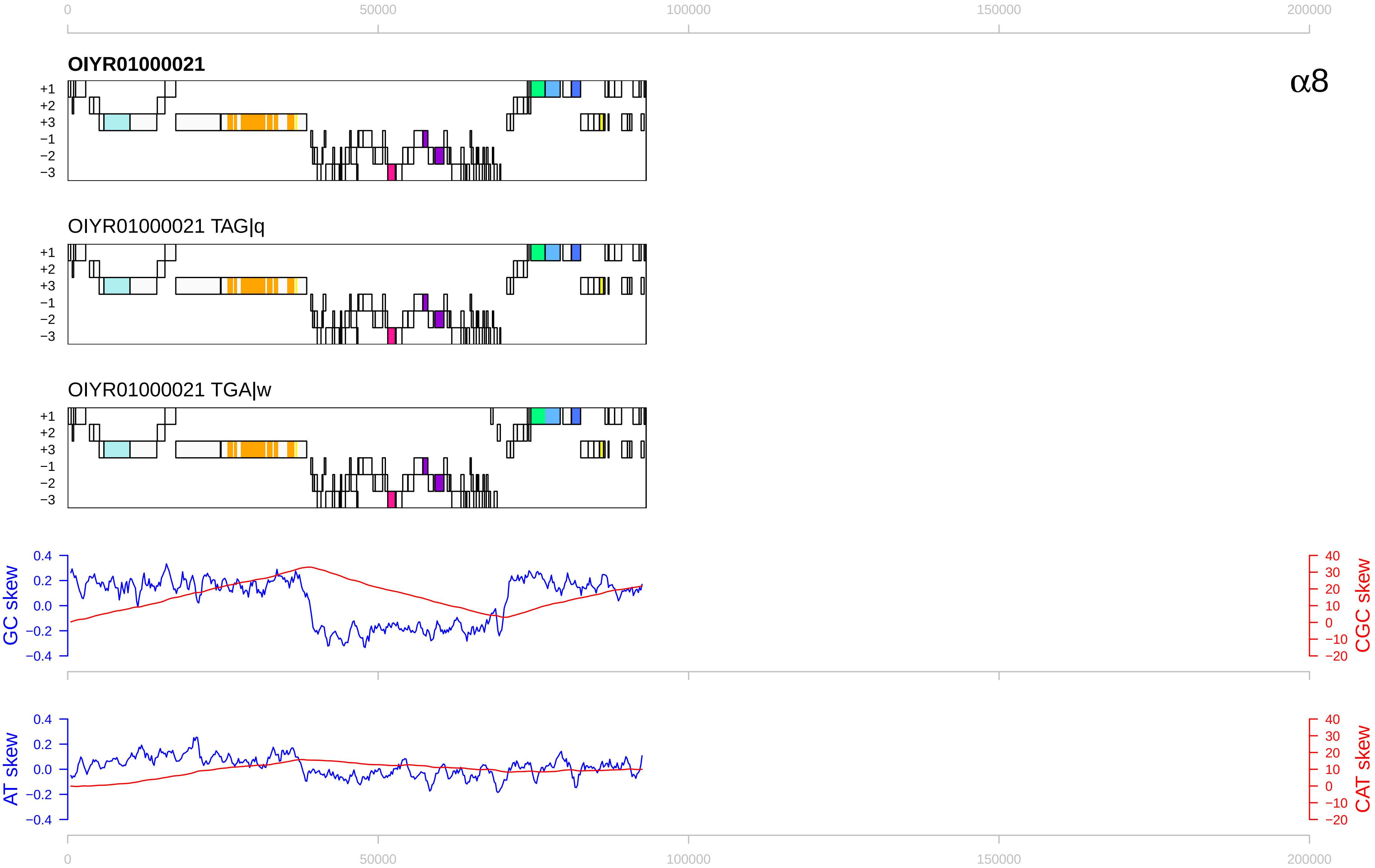

0
50000
100000
150000
200000
OIYR01000021
α8
+1
+2
+3
−1
−2
−3
OIYR01000021 TAG|q
OIYR01000021 TGA|w
GC skew
CGC skew
AT skew
CAT skew
0.4
0.2
0.0
−0.2
−0.4
40
30
20
10
0
−10
−20

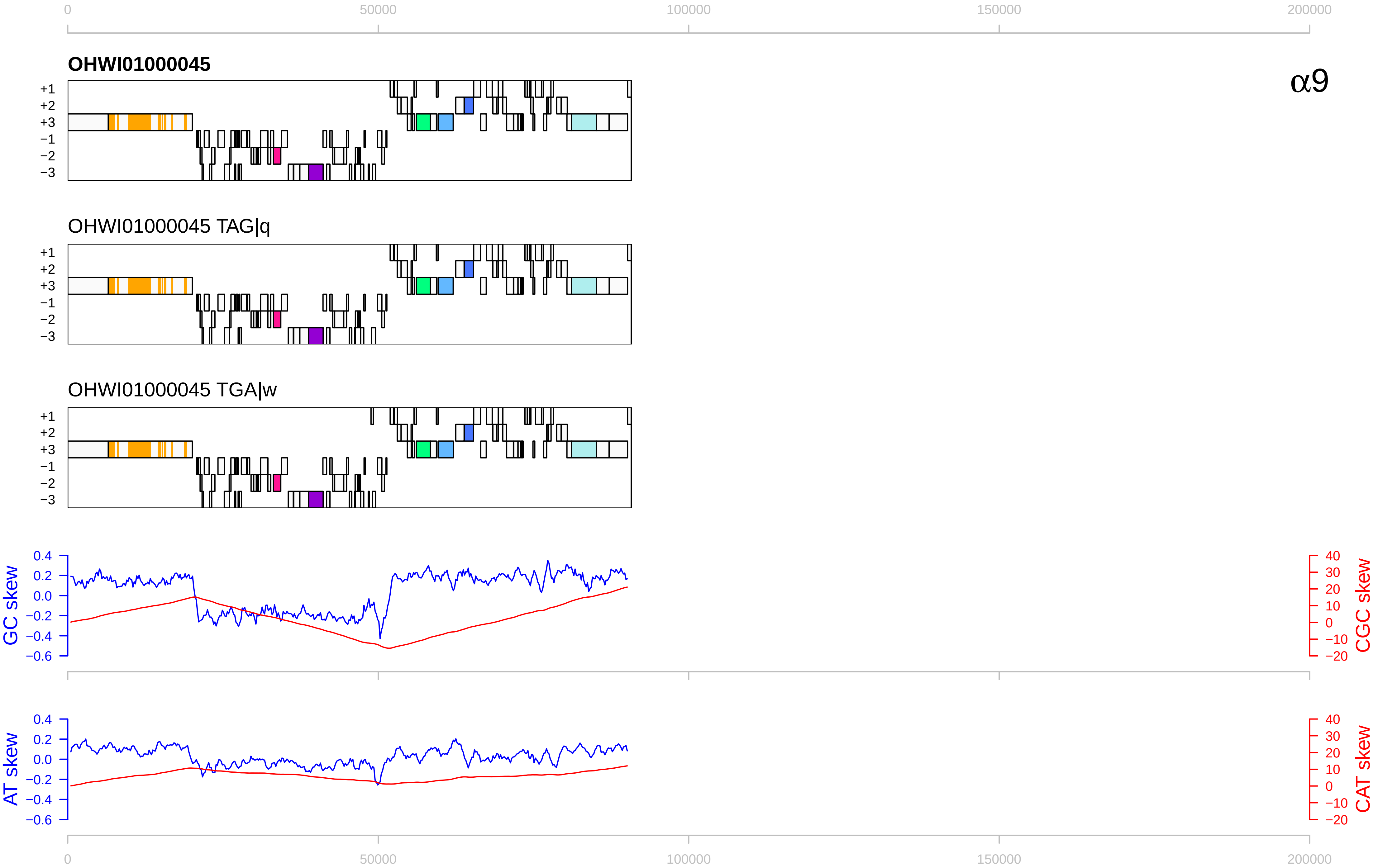

0
50000
100000
150000
200000
OHWI01000045
α9
+1
+2
+3
−1
−2
−3
OHWI01000045 TAG|q
OHWI01000045 TGA|w
GC skew
CGC skew
AT skew
CAT skew
0.4
0.2
0.0
−0.2
−0.4
−0.6
40
30
20
10
0
−10
−20

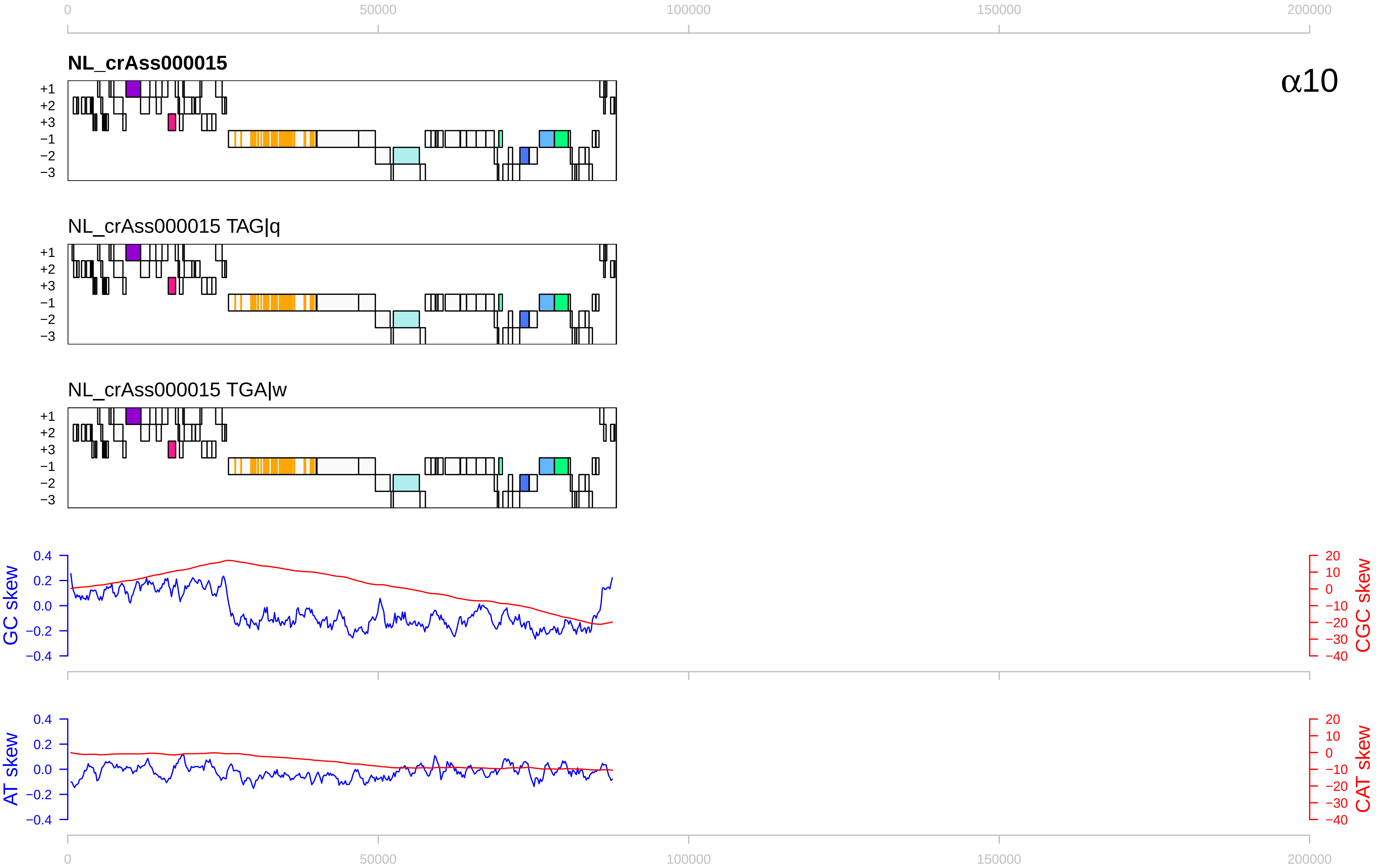

0
50000
100000
150000
200000
NL_crAss000015
α10
+1
+2
+3
−1
−2
−3
NL_crAss000015 TAG|q
NL_crAss000015 TGA|w
GC skew
CGC skew
AT skew
CAT skew
0.4
0.2
0.0
−0.2
−0.4
20
10
0
−10
−20
−30
−40

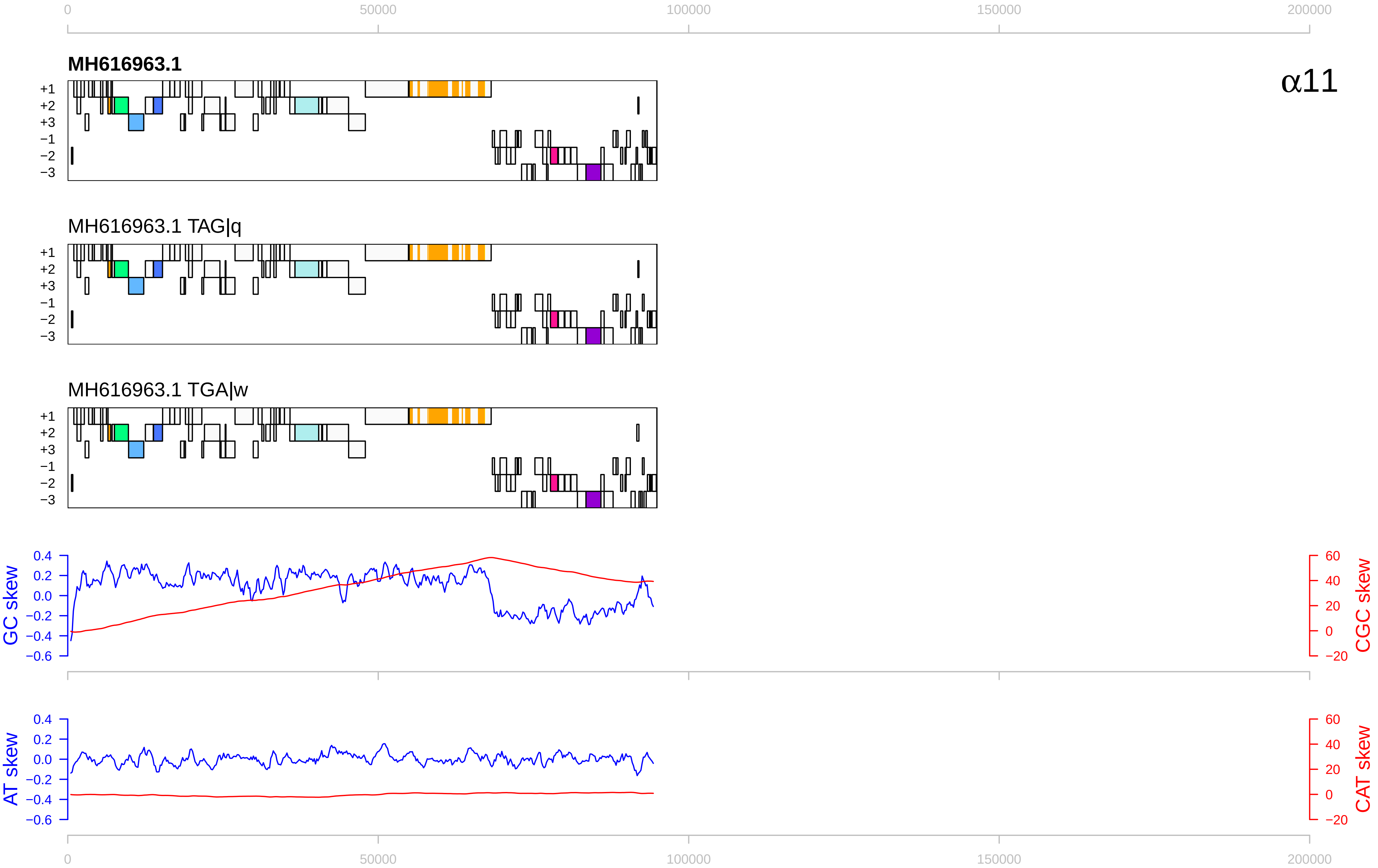

0
50000
100000
150000
200000
MH616963.1
α11
+1
+2
+3
−1
−2
−3
MH616963.1 TAG|q
MH616963.1 TGA|w
GC skew
CGC skew
0.4
0.2
0.0
−0.2
−0.4
−0.6
60
40
20
0
−20
AT skew
CAT skew

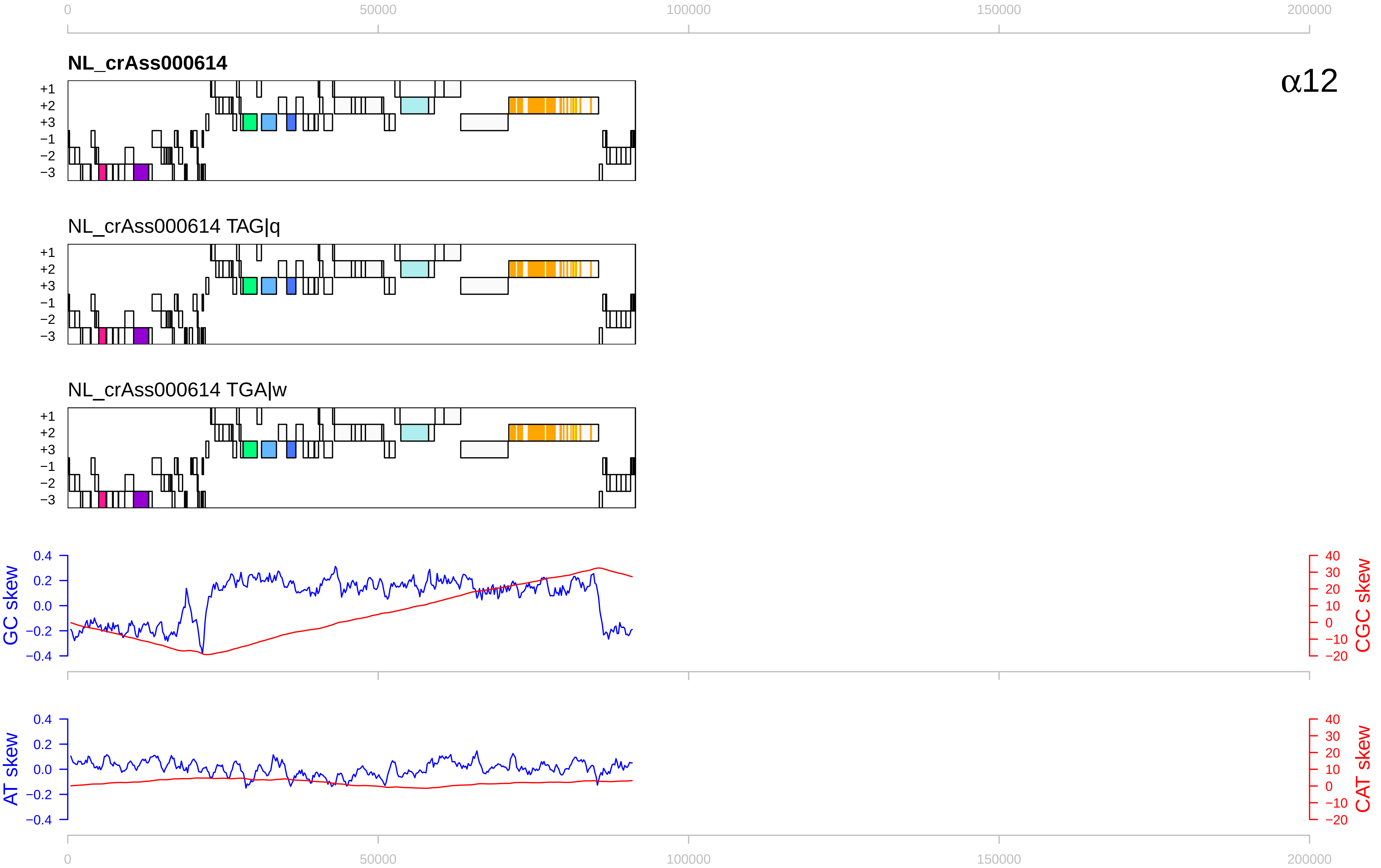

0
50000
100000
150000
200000
NL_crAss000614
α12
+1
+2
+3
−1
−2
−3
NL_crAss000614 TAG|q
NL_crAss000614 TGA|w
GC skew
CGC skew
AT skew
CAT skew
0.4
0.2
0.0
−0.2
−0.4
40
30
20
10
0
−10
−20

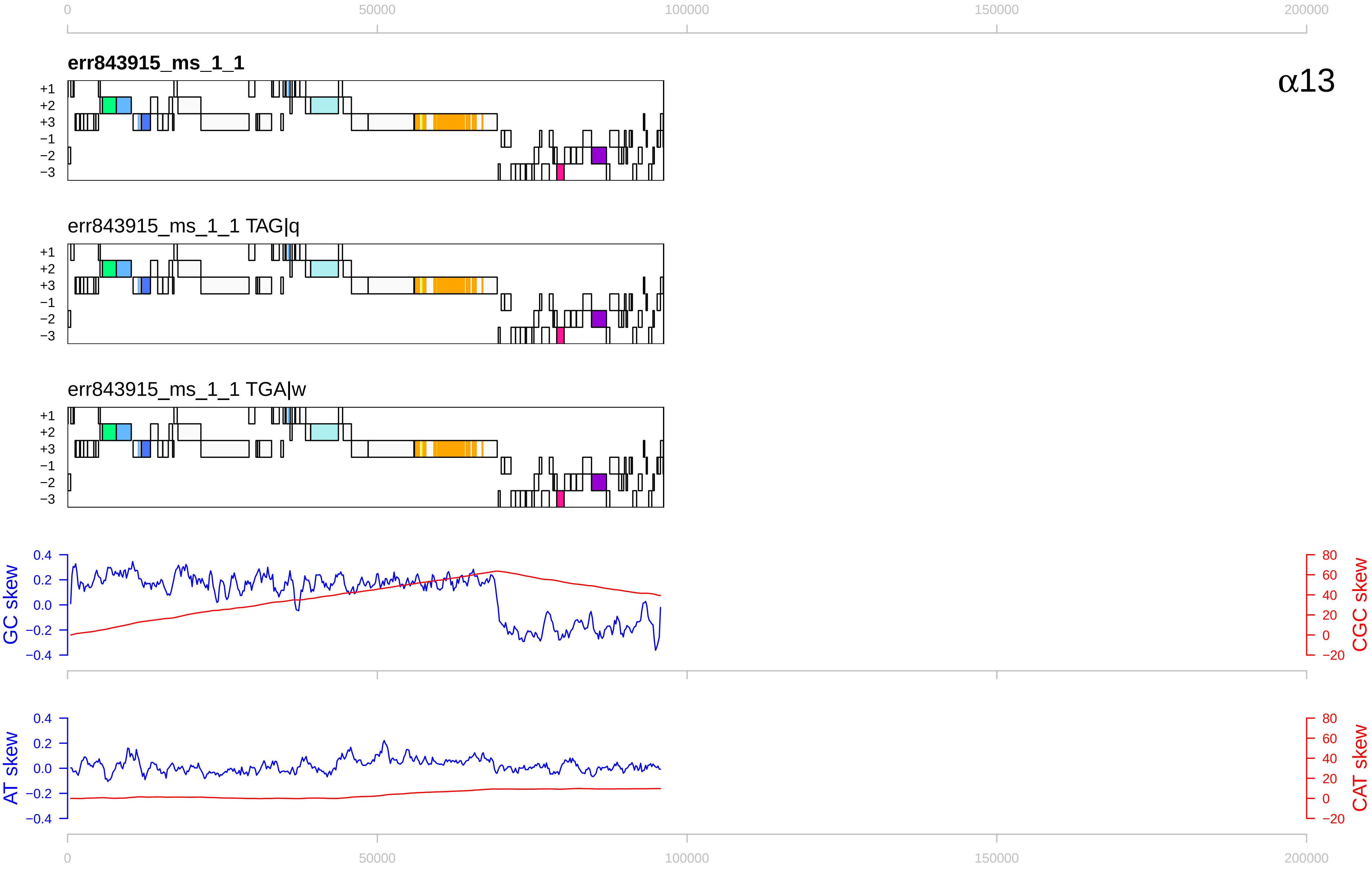

err843915_ms_1_1
err843915_ms_1_1 TAG|q
err843915_ms_1_1 TGA|w
α13
GC skew
CGC skew
AT skew
CAT skew
0
50000
100000
150000
200000
+1
+2
+3
−1
−2
−3

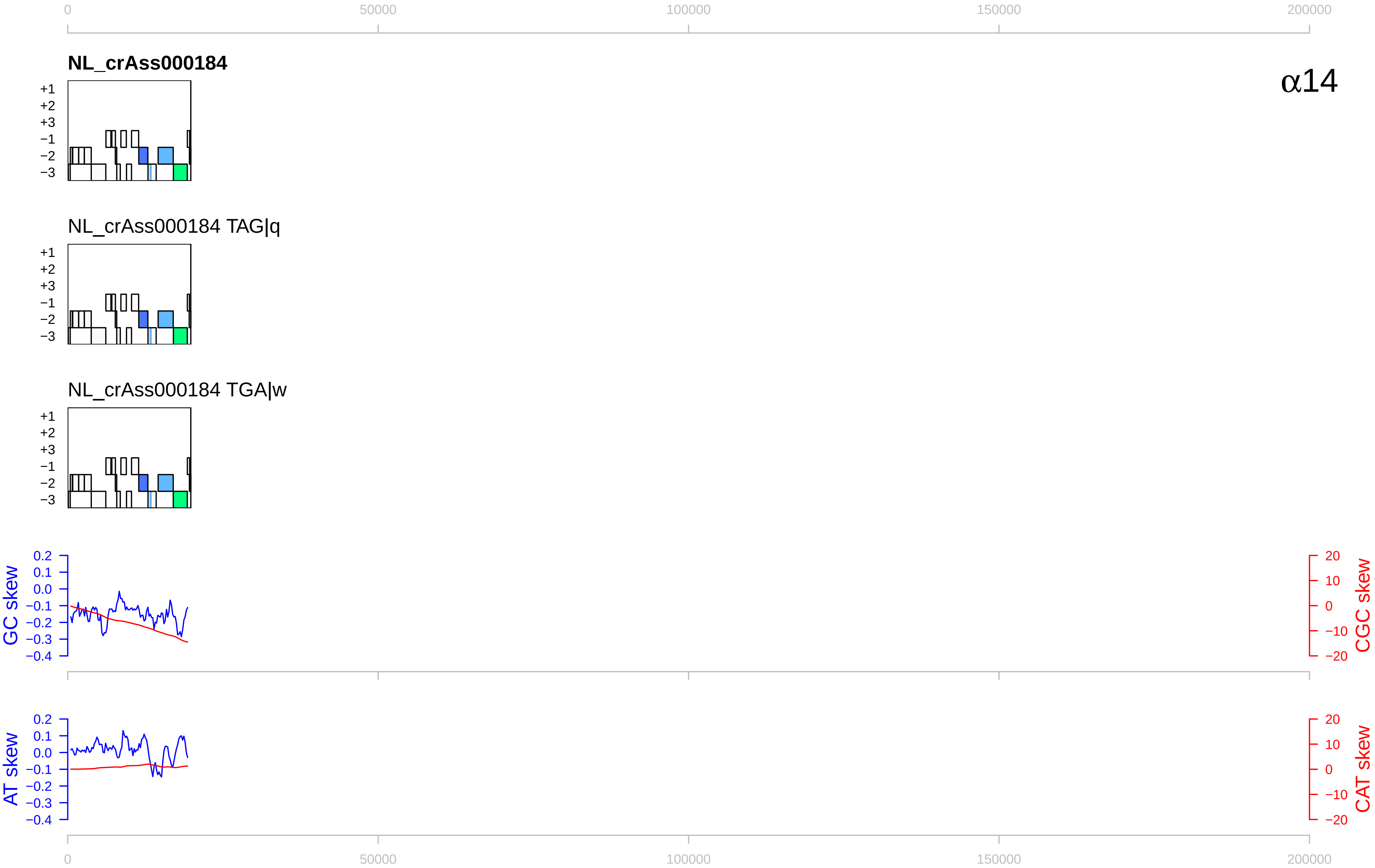

0
50000
100000
150000
200000
NL_crAss000184
α14
+1
+2
+3
−1
−2
−3
NL_crAss000184 TAG|q
NL_crAss000184 TGA|w
GC skew
0.2
0.1
0.0
−0.1
−0.2
−0.3
−0.4
CGC skew
20
10
0
−10
−20
AT skew
CAT skew

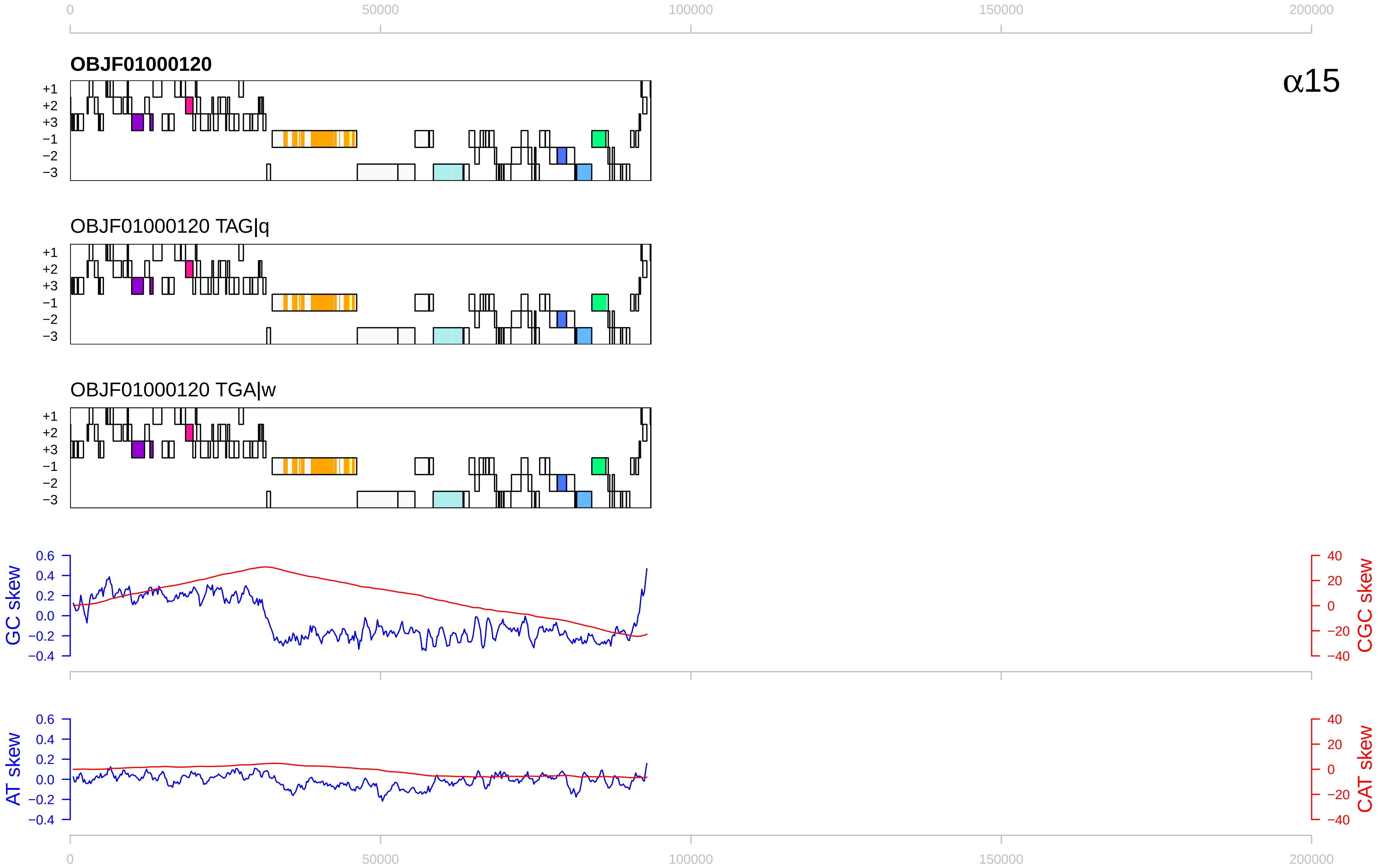

0
50000
100000
150000
200000
OBJF01000120
α15
+1
+2
+3
−1
−2
−3
OBJF01000120 TAG|q
OBJF01000120 TGA|w
GC skew
CGC skew
AT skew
CAT skew
0.6
0.4
0.2
0.0
−0.2
−0.4
40
20
0
−20
−40

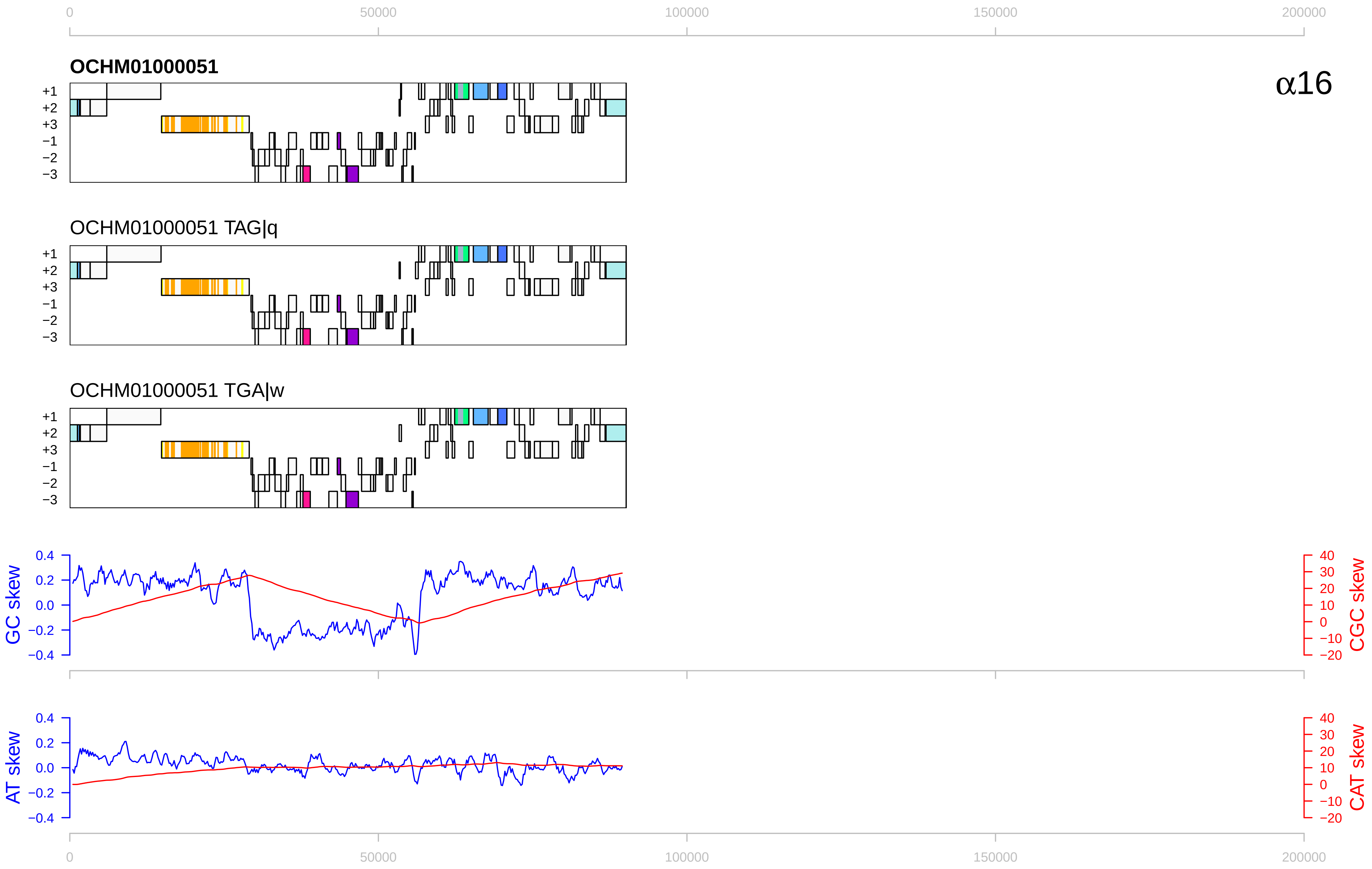

0
50000
100000
150000
200000
OCHM01000051
α16
+1
+2
+3
−1
−2
−3
OCHM01000051 TAG|q
OCHM01000051 TGA|w
GC skew
CGC skew
AT skew
CAT skew
0.4
0.2
0.0
−0.2
−0.4
40
30
20
10
0
−10
−20

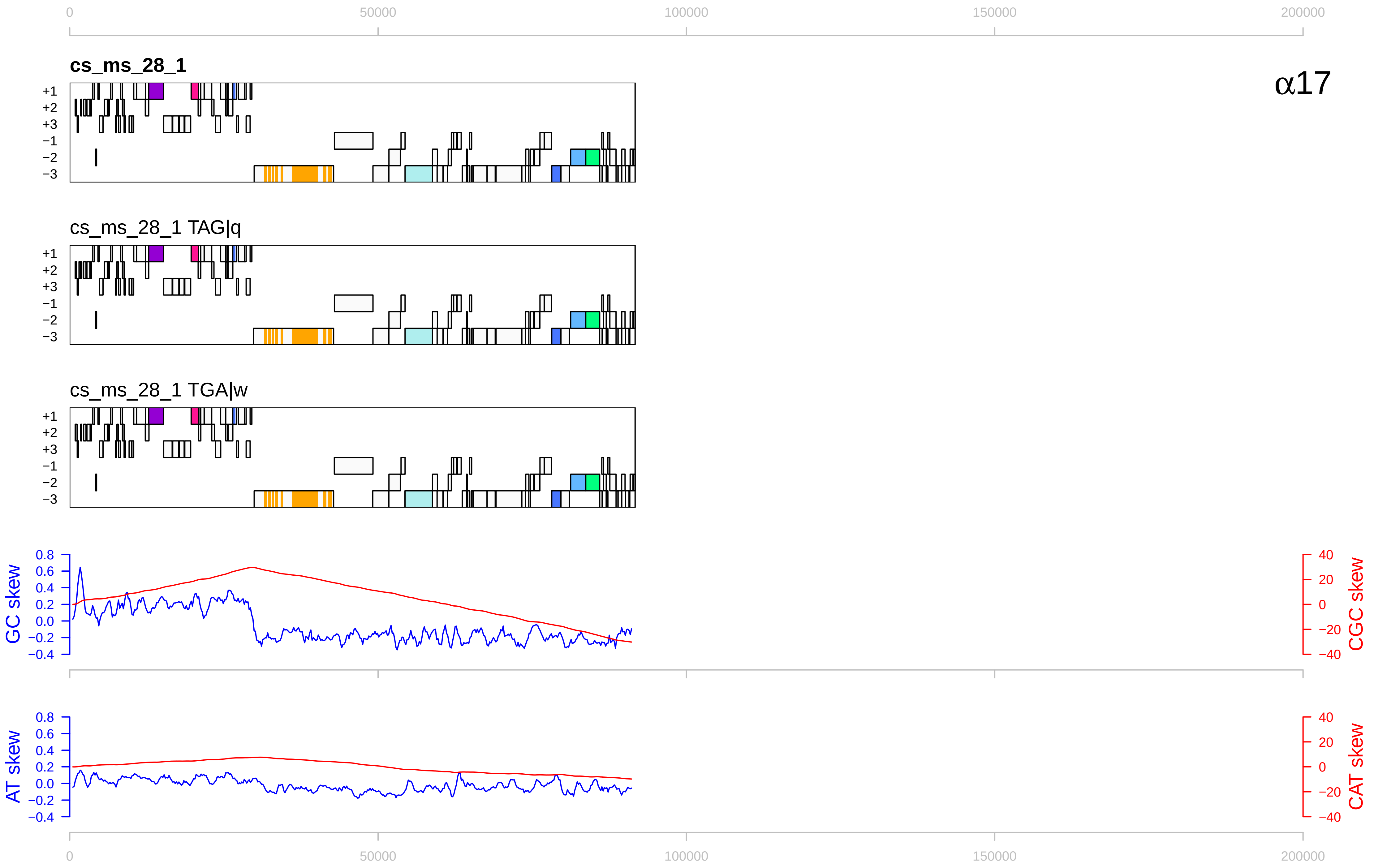

0
50000
100000
150000
200000
cs_ms_28_1
α17
+1
+2
+3
−1
−2
−3
cs_ms_28_1 TAG|q
cs_ms_28_1 TGA|w
GC skew
0.8
0.6
0.4
0.2
0.0
−0.2
−0.4
40
20
0
−20
−40
CGC skew
AT skew
CAT skew

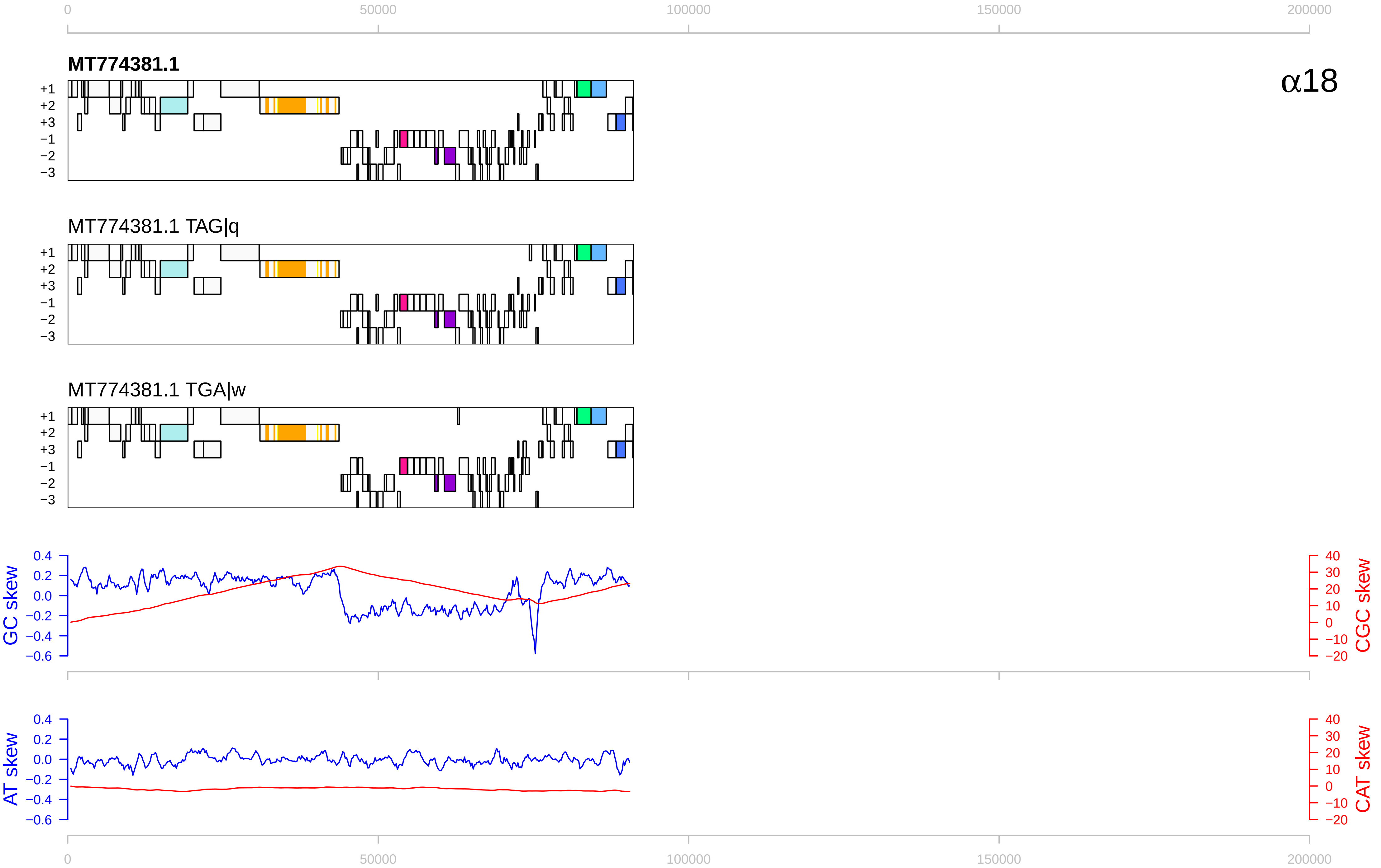

0
50000
100000
150000
200000
MT774381.1
α18
+1
+2
+3
−1
−2
−3
MT774381.1 TAG|q
MT774381.1 TGA|w
GC skew
0.4
0.2
0.0
−0.2
−0.4
−0.6
CGC skew
40
30
20
10
0
−10
−20
AT skew
CAT skew

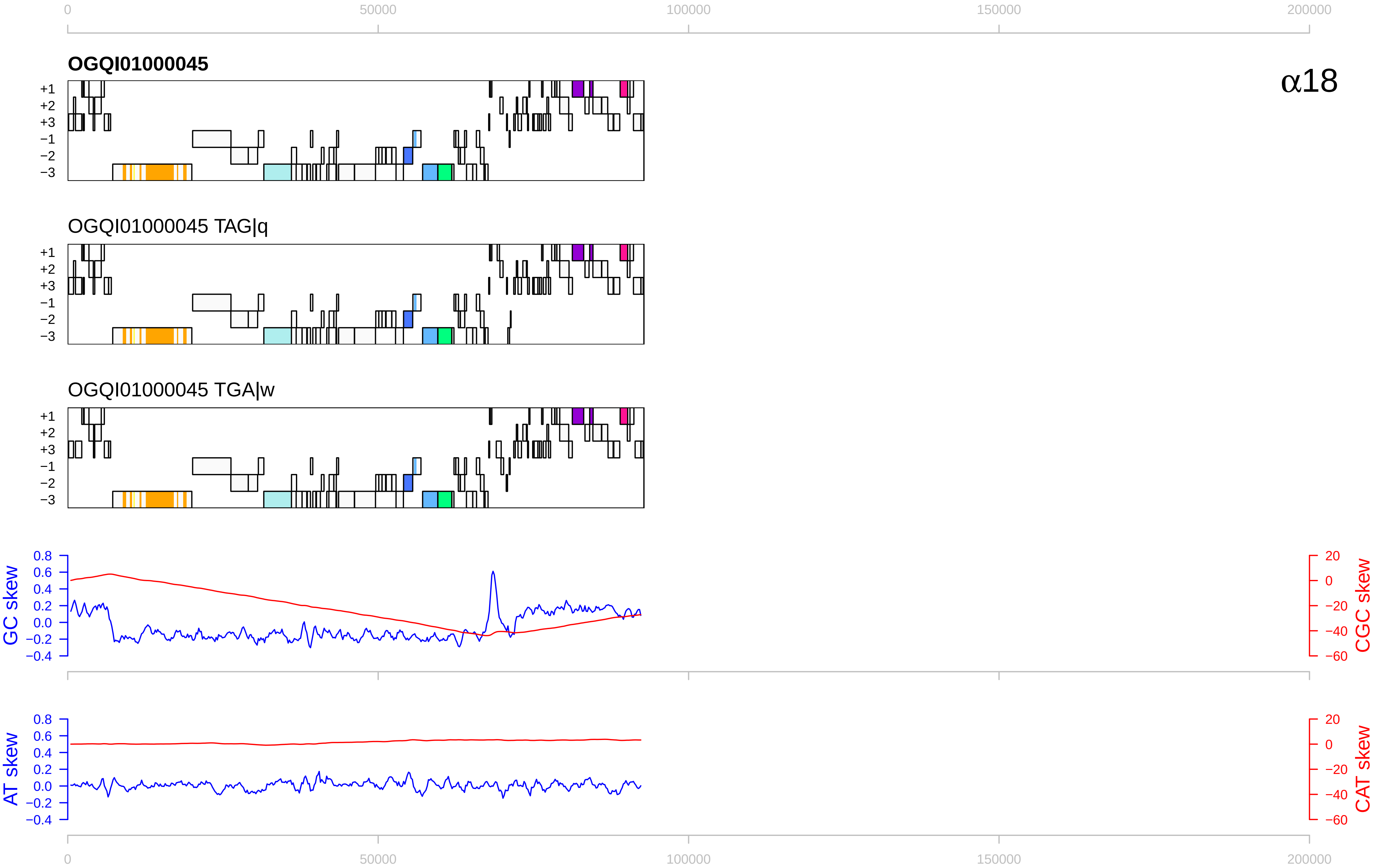

0
50000
100000
150000
200000
OGQI01000045
α18
+1
+2
+3
−1
−2
−3
OGQI01000045 TAG|q
OGQI01000045 TGA|w
GC skew
CGC skew
AT skew
CAT skew
0.8
0.6
0.4
0.2
0.0
−0.2
−0.4
20
0
−20
−40
−60

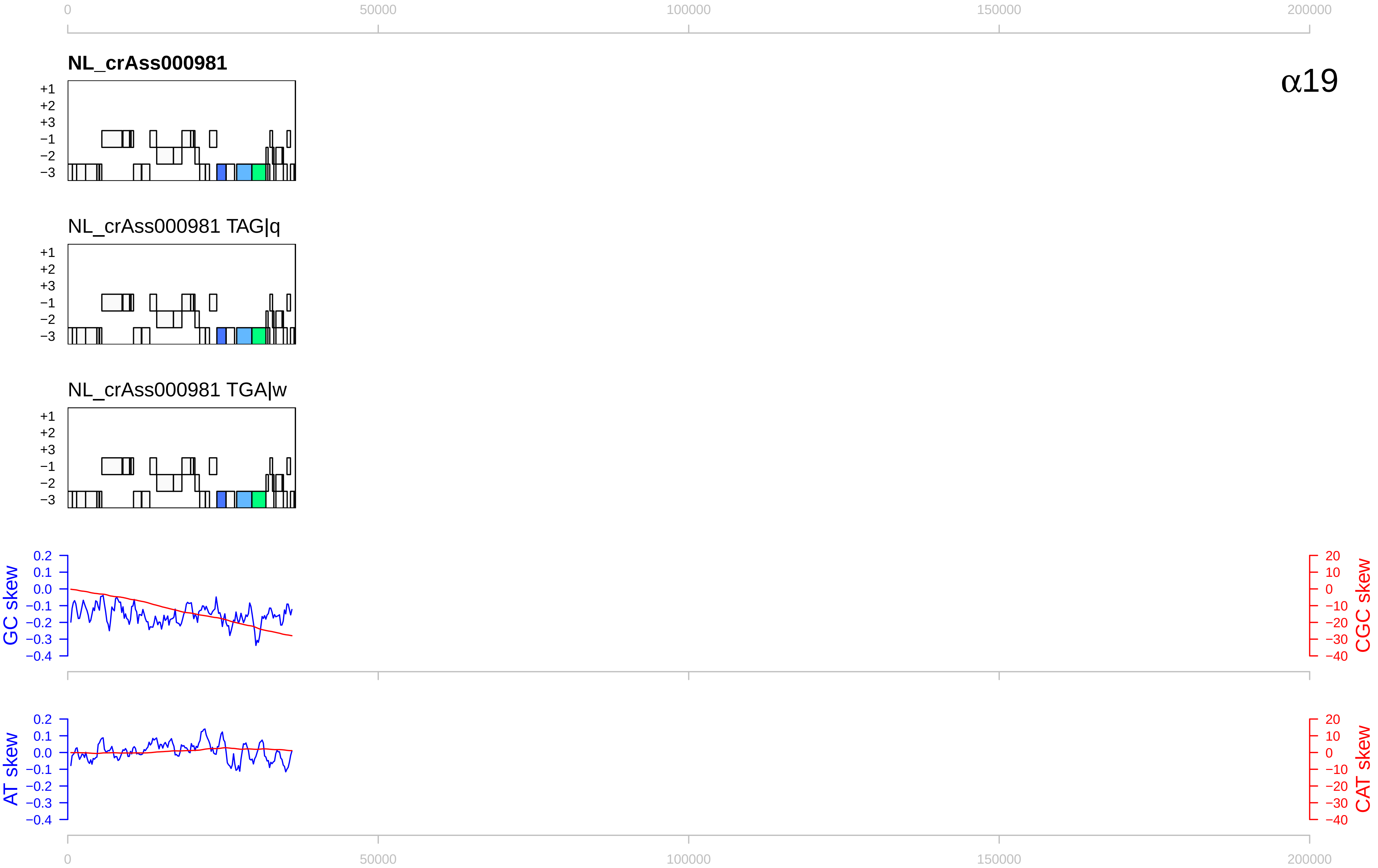

0
50000
100000
150000
200000
NL_crAss000981
α19
+1
+2
+3
−1
−2
−3
NL_crAss000981 TAG|q
NL_crAss000981 TGA|w
GC skew
0.2
0.1
0.0
−0.1
−0.2
−0.3
−0.4
CGC skew
20
10
0
−10
−20
−30
−40
AT skew
CAT skew

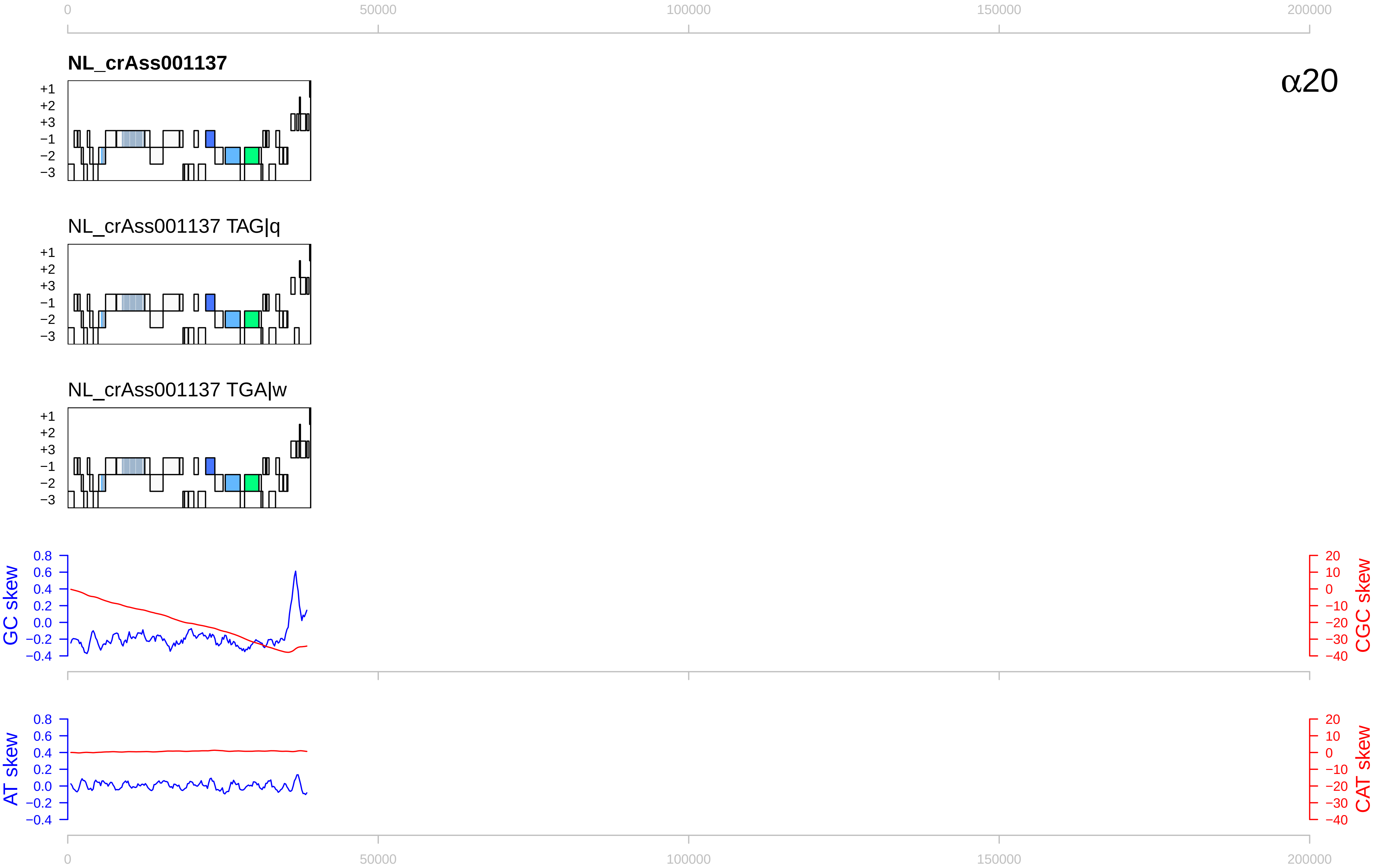

0
50000
100000
150000
200000
NL_crAss001137
α20
+1
+2
+3
−1
−2
−3
NL_crAss001137 TAG|q
NL_crAss001137 TGA|w
GC skew
CGC skew
AT skew
CAT skew
0.8
0.6
0.4
0.2
0.0
−0.2
−0.4
20
10
0
−10
−20
−30
−40

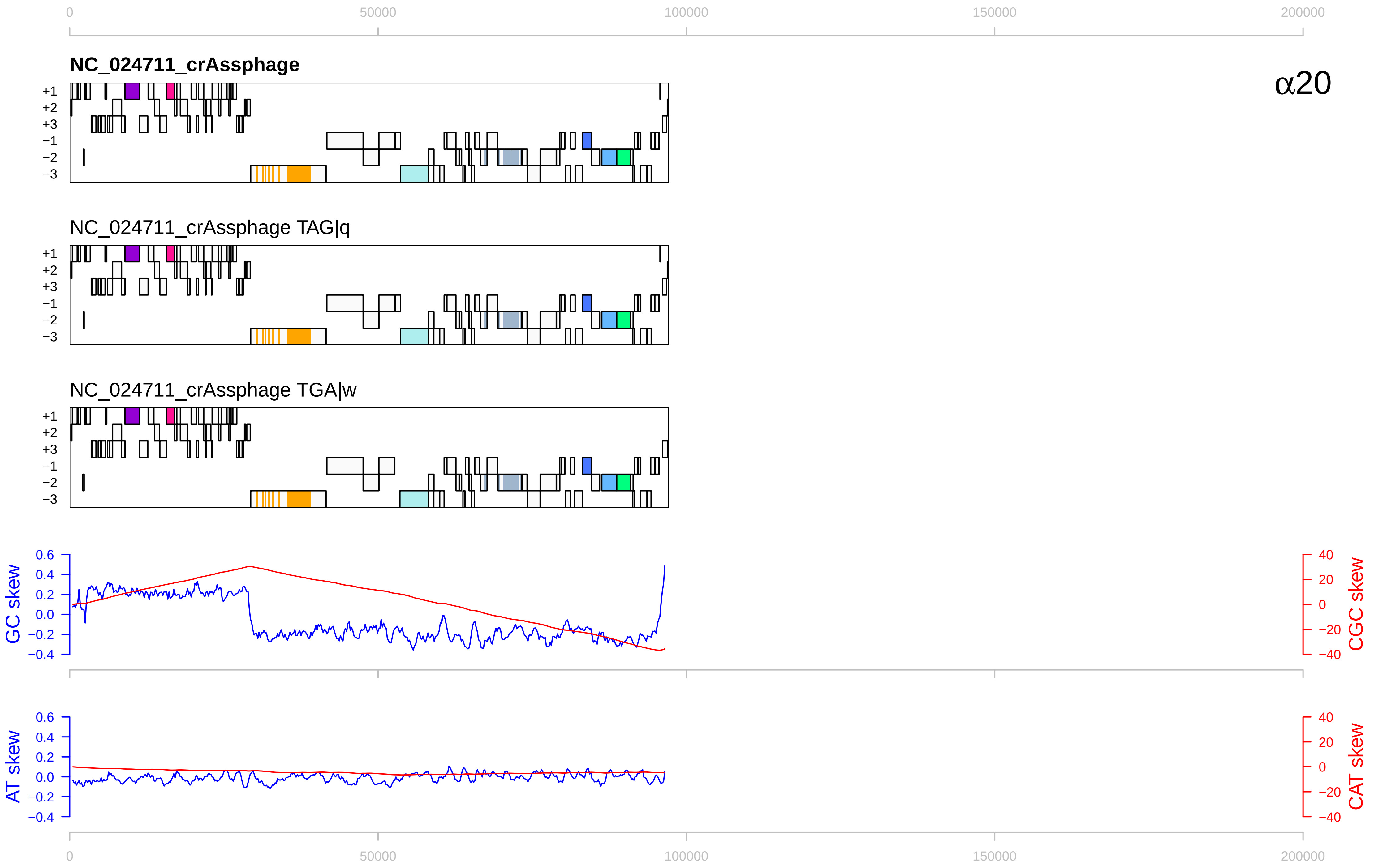

0
50000
100000
150000
200000
NC_024711_crAssphage
α20
+1
+2
+3
−1
−2
−3
NC_024711_crAssphage TAG|q
NC_024711_crAssphage TGA|w
GC skew
CGC skew
AT skew
CAT skew
0.6
0.4
0.2
0.0
−0.2
−0.4
40
20
0
−20
−40

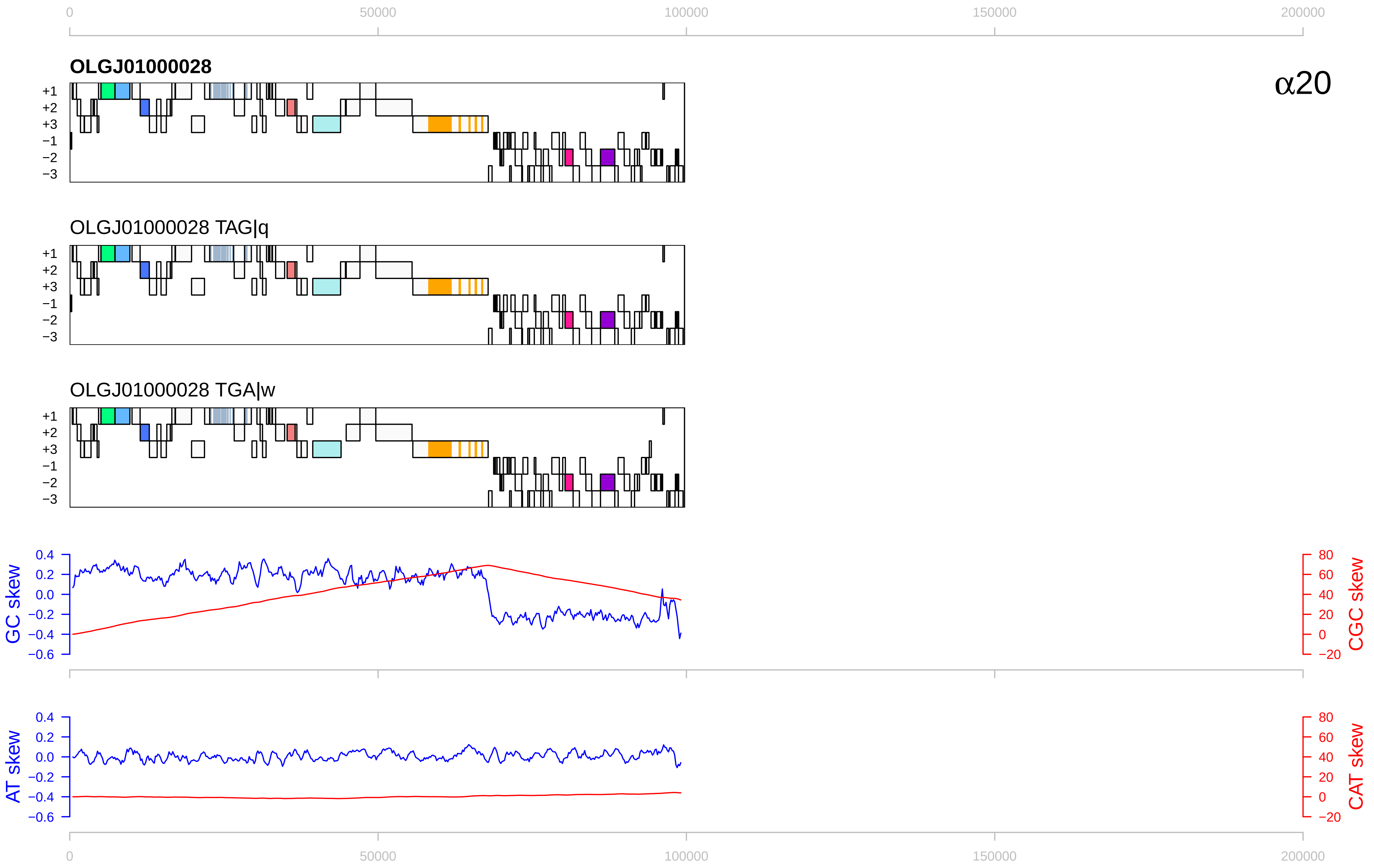

0
50000
100000
150000
200000
OLGJ01000028
α20
+1
+2
+3
−1
−2
−3
OLGJ01000028 TAG|q
OLGJ01000028 TGA|w
GC skew
CGC skew
AT skew
CAT skew
0.4
0.2
0.0
−0.2
−0.4
−0.6
80
60
40
20
0
−20

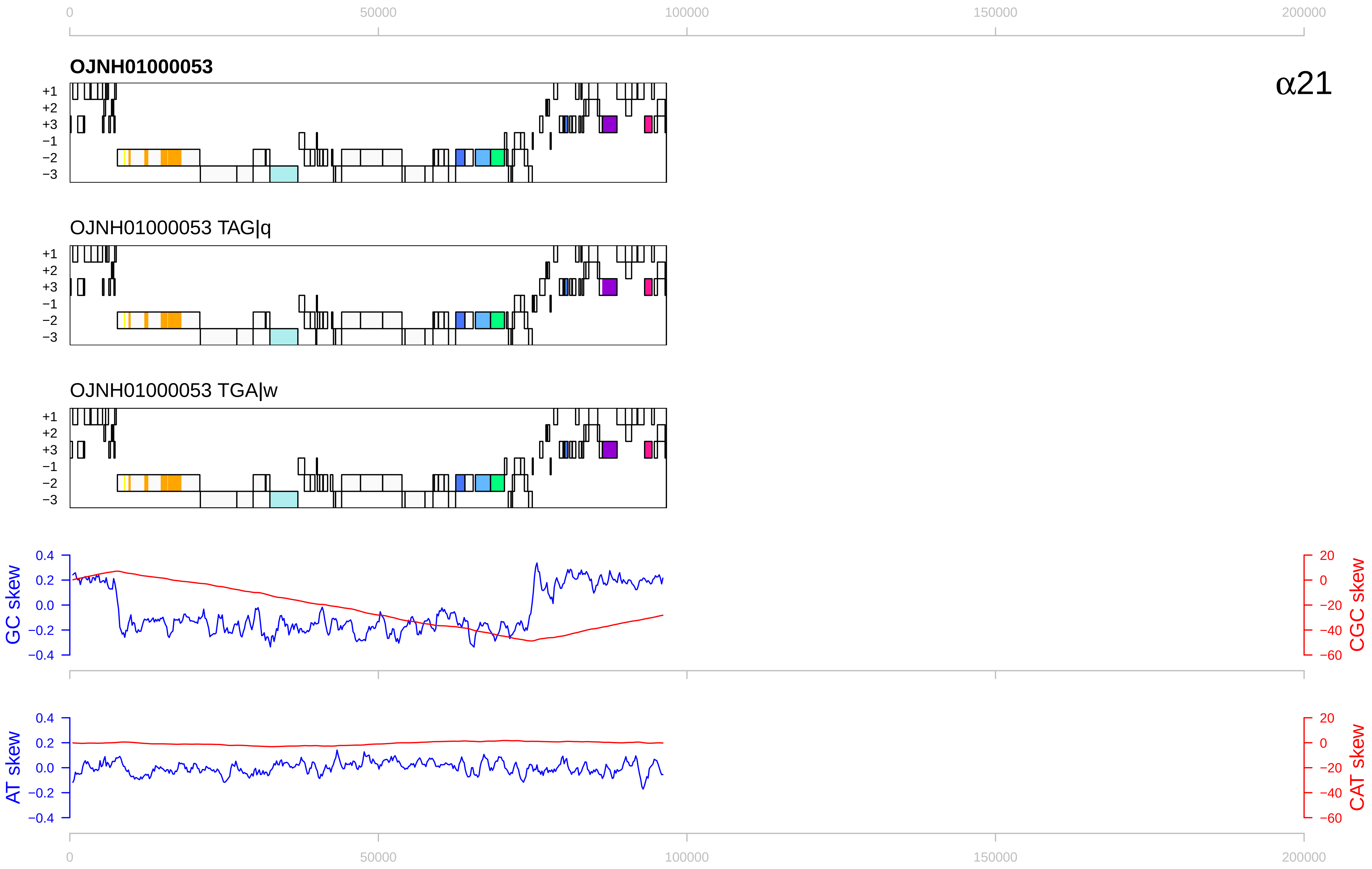

0
50000
100000
150000
200000
OJNH01000053
α21
+1
+2
+3
−1
−2
−3
OJNH01000053 TAG|q
OJNH01000053 TGA|w
GC skew
CGC skew
AT skew
CAT skew
0.4
0.2
0.0
−0.2
−0.4
20
0
−20
−40
−60

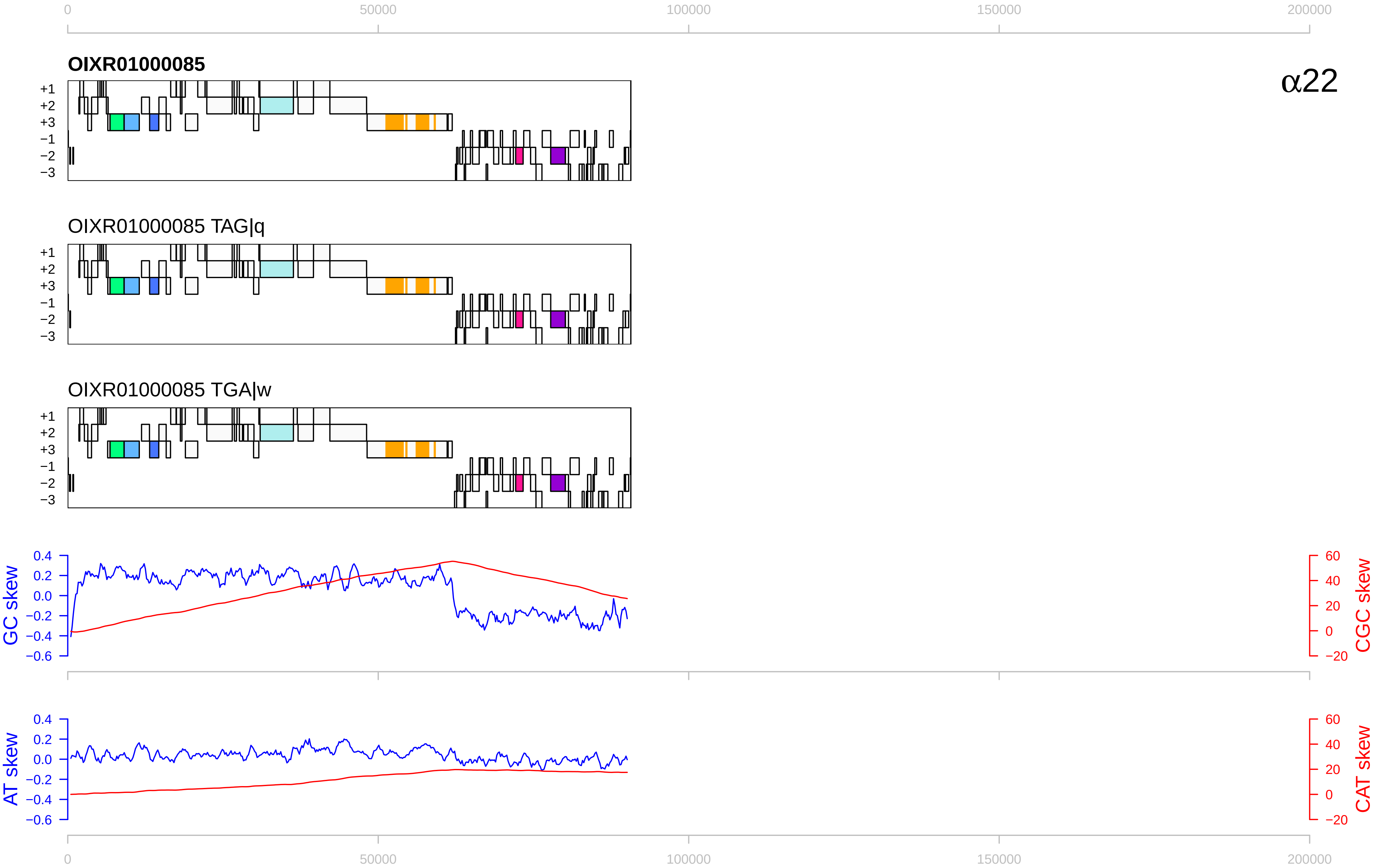

0
50000
100000
150000
200000
OIXR01000085
α22
+1
+2
+3
−1
−2
−3
OIXR01000085 TAG|q
OIXR01000085 TGA|w
GC skew
CGC skew
AT skew
CAT skew
0.4
0.2
0.0
−0.2
−0.4
−0.6
60
40
20
0
−20

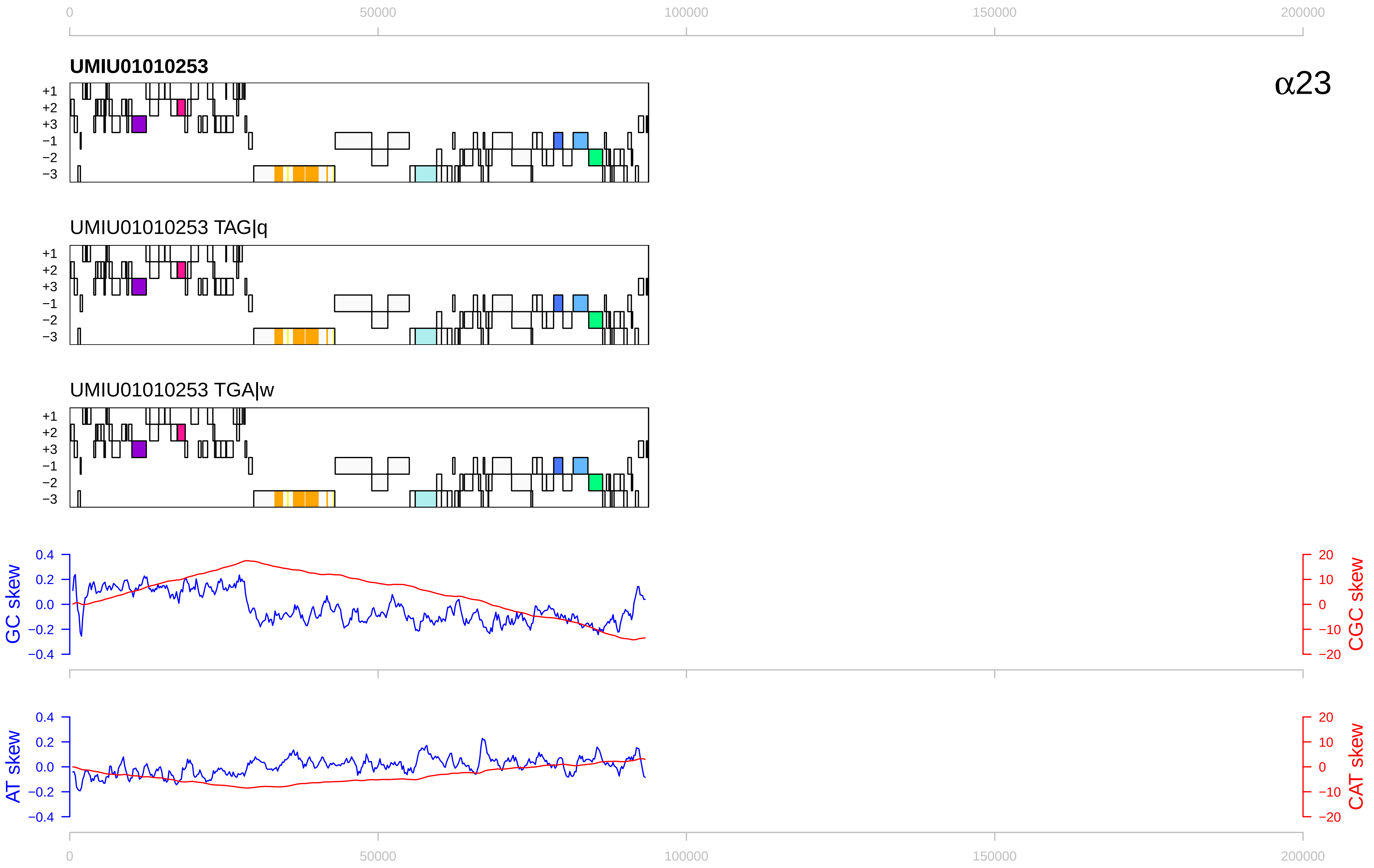

0
50000
100000
150000
200000
UMIU01010253
α23
+1
+2
+3
−1
−2
−3
UMIU01010253 TAG|q
UMIU01010253 TGA|w
GC skew
CGC skew
0.4
0.2
0.0
−0.2
−0.4
20
10
0
−10
−20
AT skew
CAT skew

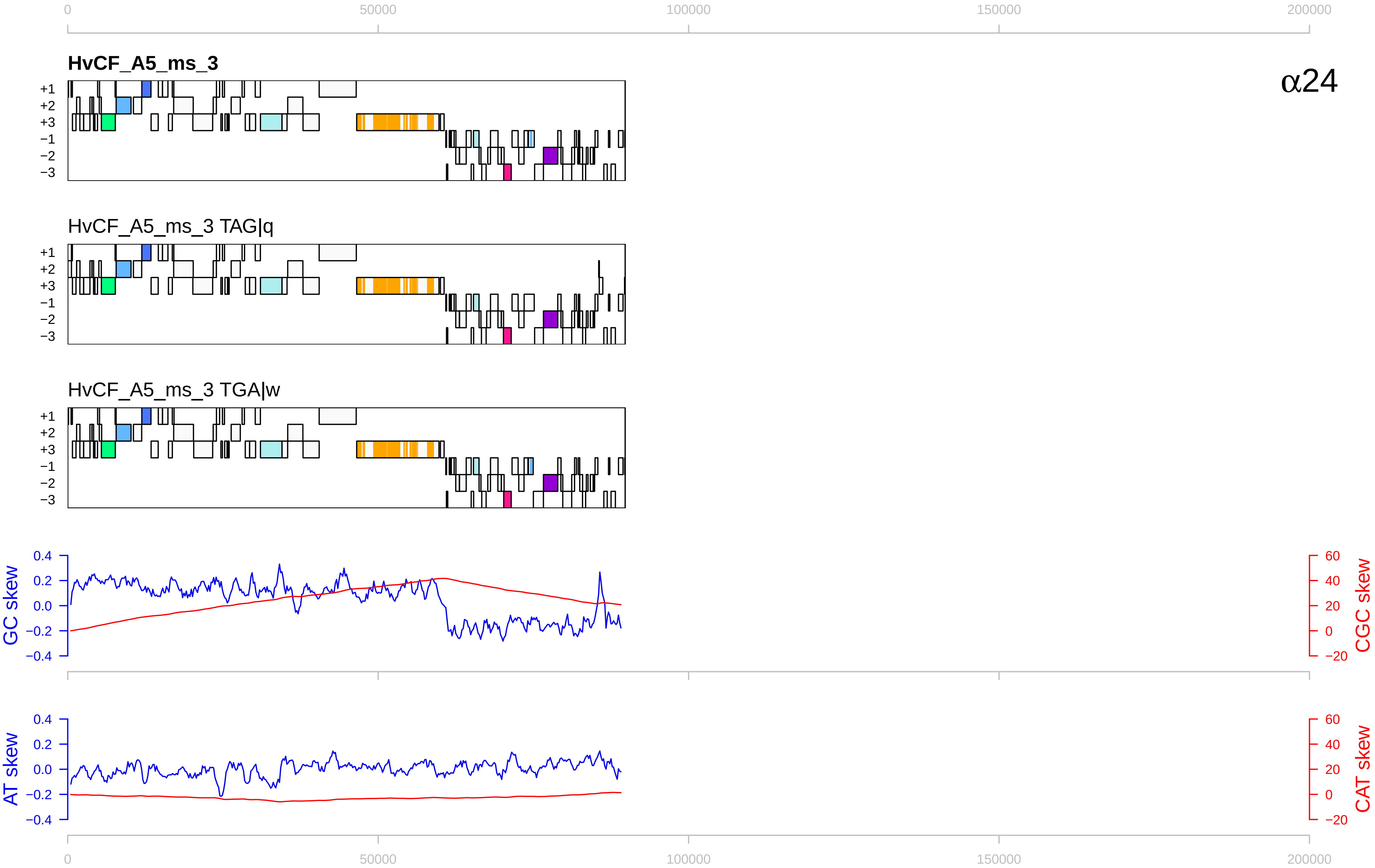

0
50000
100000
150000
200000
HvCF_A5_ms_3
α24
+1
+2
+3
−1
−2
−3
HvCF_A5_ms_3 TAG|q
HvCF_A5_ms_3 TGA|w
GC skew
0.4
0.2
0.0
−0.2
−0.4
CGC skew
60
40
20
0
−20
AT skew
CAT skew

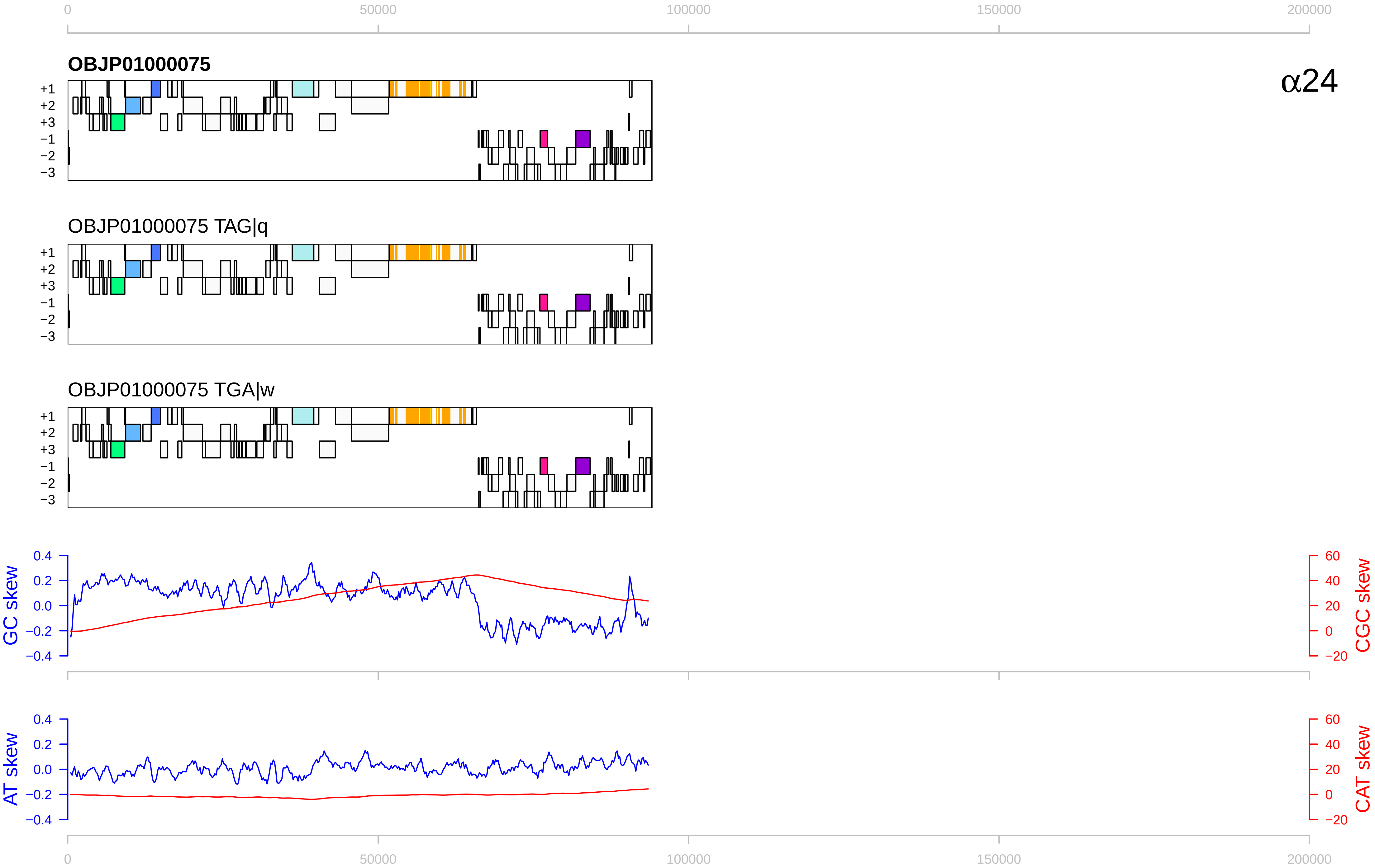
0
50000
100000
150000
200000
OBJP01000075
α24
+1
+2
+3
−1
−2
−3
OBJP01000075 TAG|q
OBJP01000075 TGA|w
GC skew
CGC skew
0.4
0.2
0.0
−0.2
−0.4
60
40
20
0
−20
AT skew
CAT skew

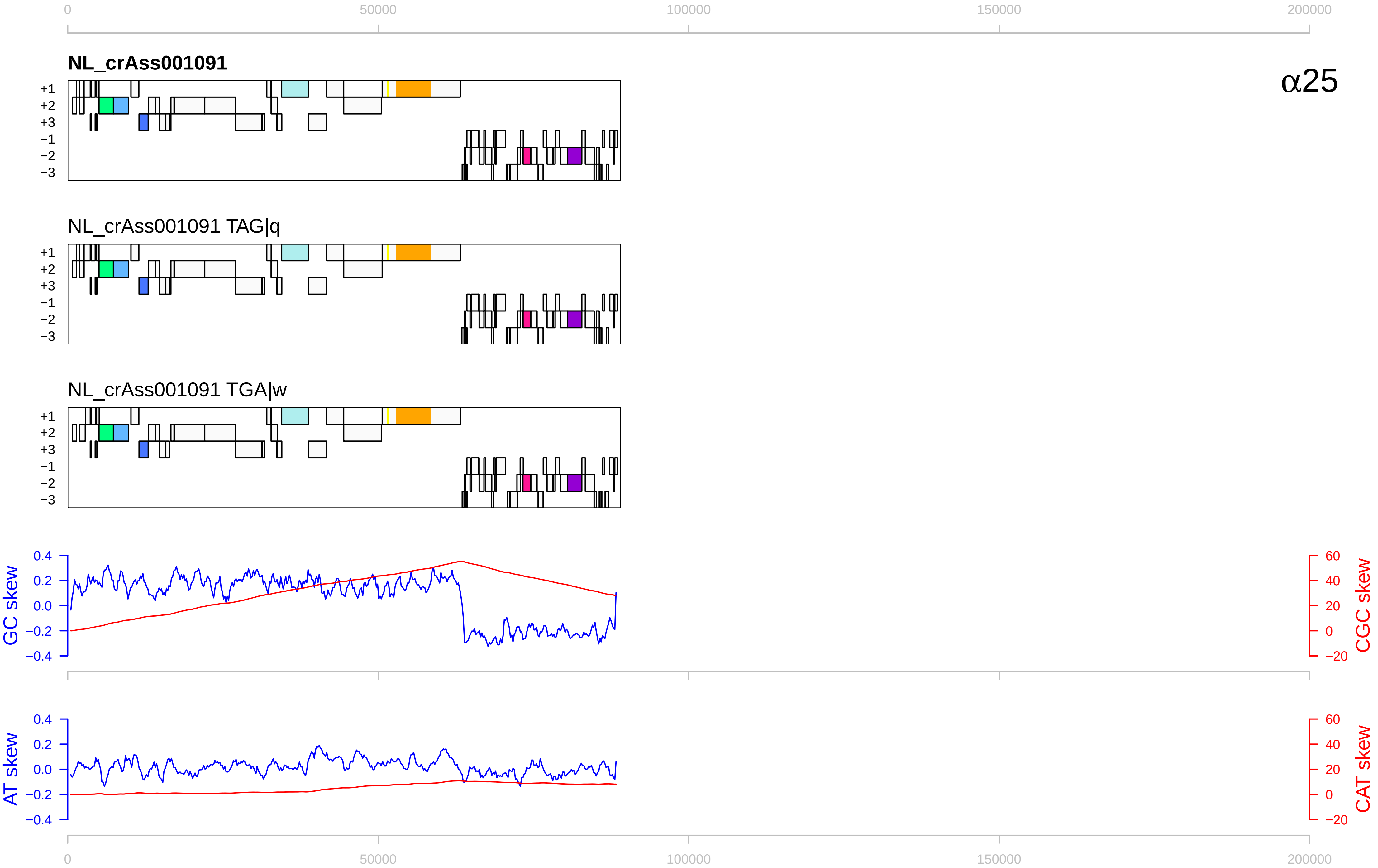

0
50000
100000
150000
200000
NL_crAss001091
α25
+1
+2
+3
−1
−2
−3
NL_crAss001091 TAG|q
NL_crAss001091 TGA|w
GC skew
CGC skew
0.4
0.2
0.0
−0.2
−0.4
60
40
20
0
−20
AT skew
CAT skew

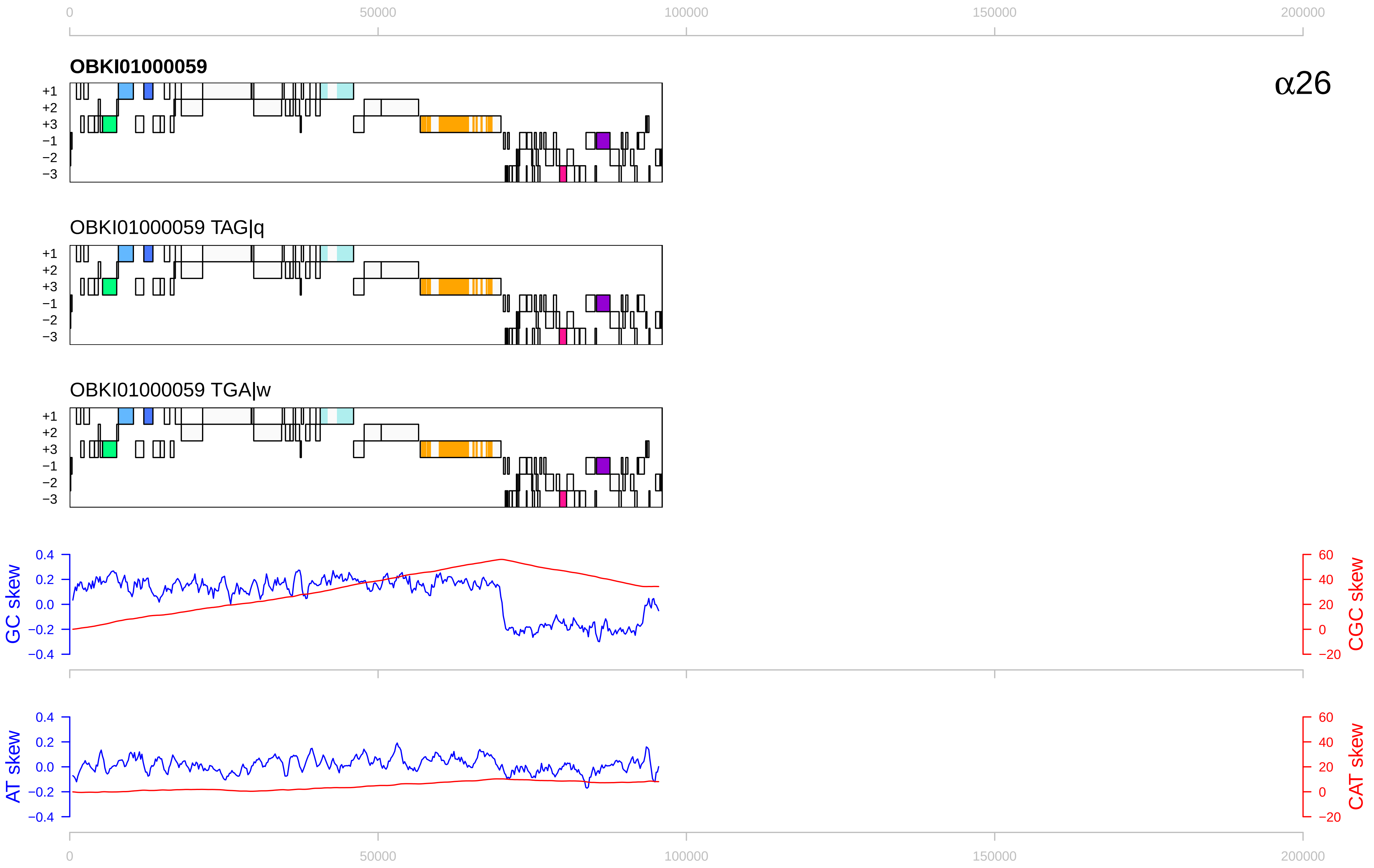

0
50000
100000
150000
200000
OBKI01000059
α26
+1
+2
+3
−1
−2
−3
OBKI01000059 TAG|q
OBKI01000059 TGA|w
GC skew
CGC skew
0.4
0.2
0.0
−0.2
−0.4
60
40
20
0
−20
AT skew
CAT skew

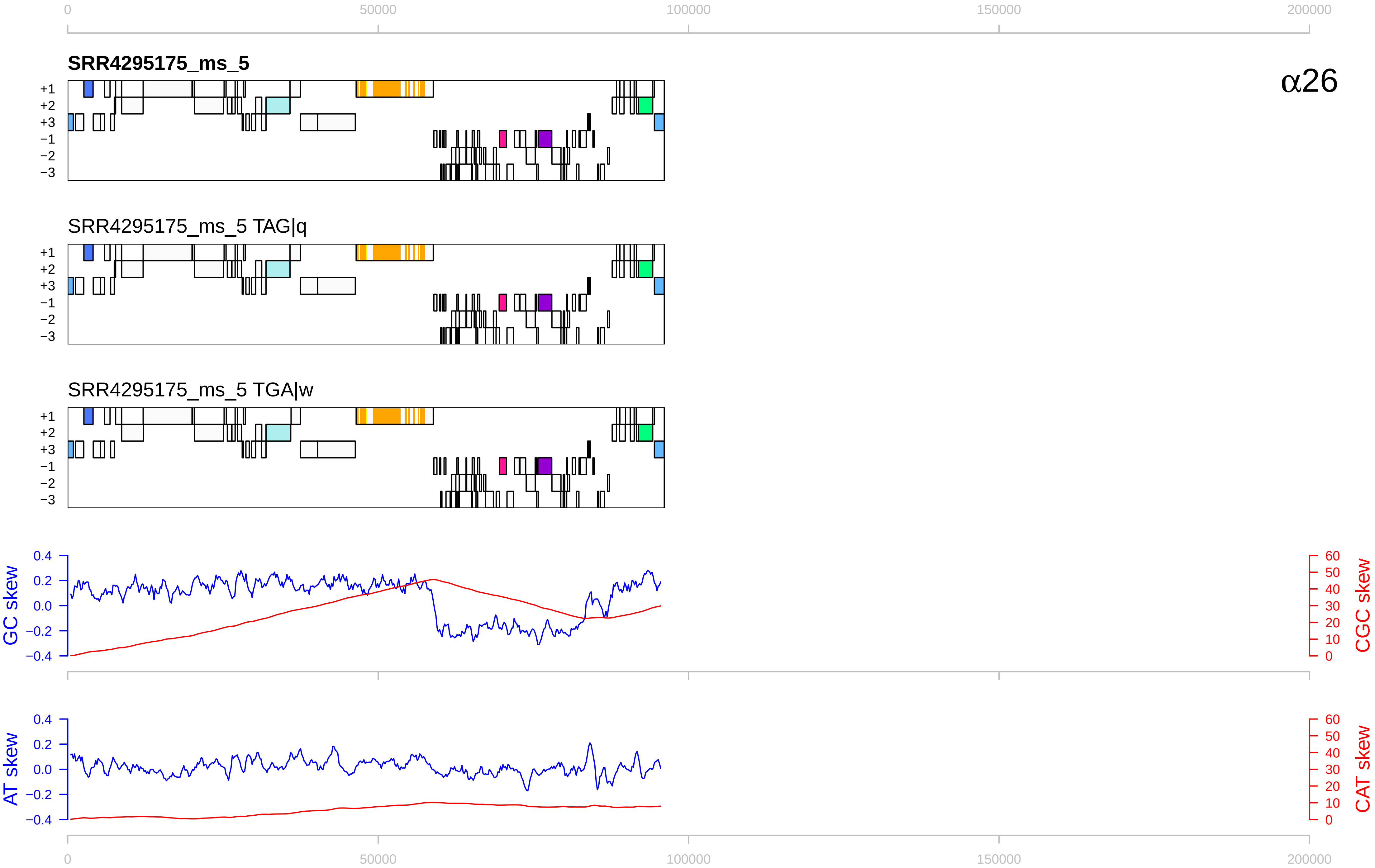

0
50000
100000
150000
200000
SRR4295175_ms_5
α26
+1
+2
+3
−1
−2
−3
SRR4295175_ms_5 TAG|q
SRR4295175_ms_5 TGA|w
GC skew
CGC skew
AT skew
CAT skew
0.4
0.2
0.0
−0.2
−0.4
60
50
40
30
20
10
0

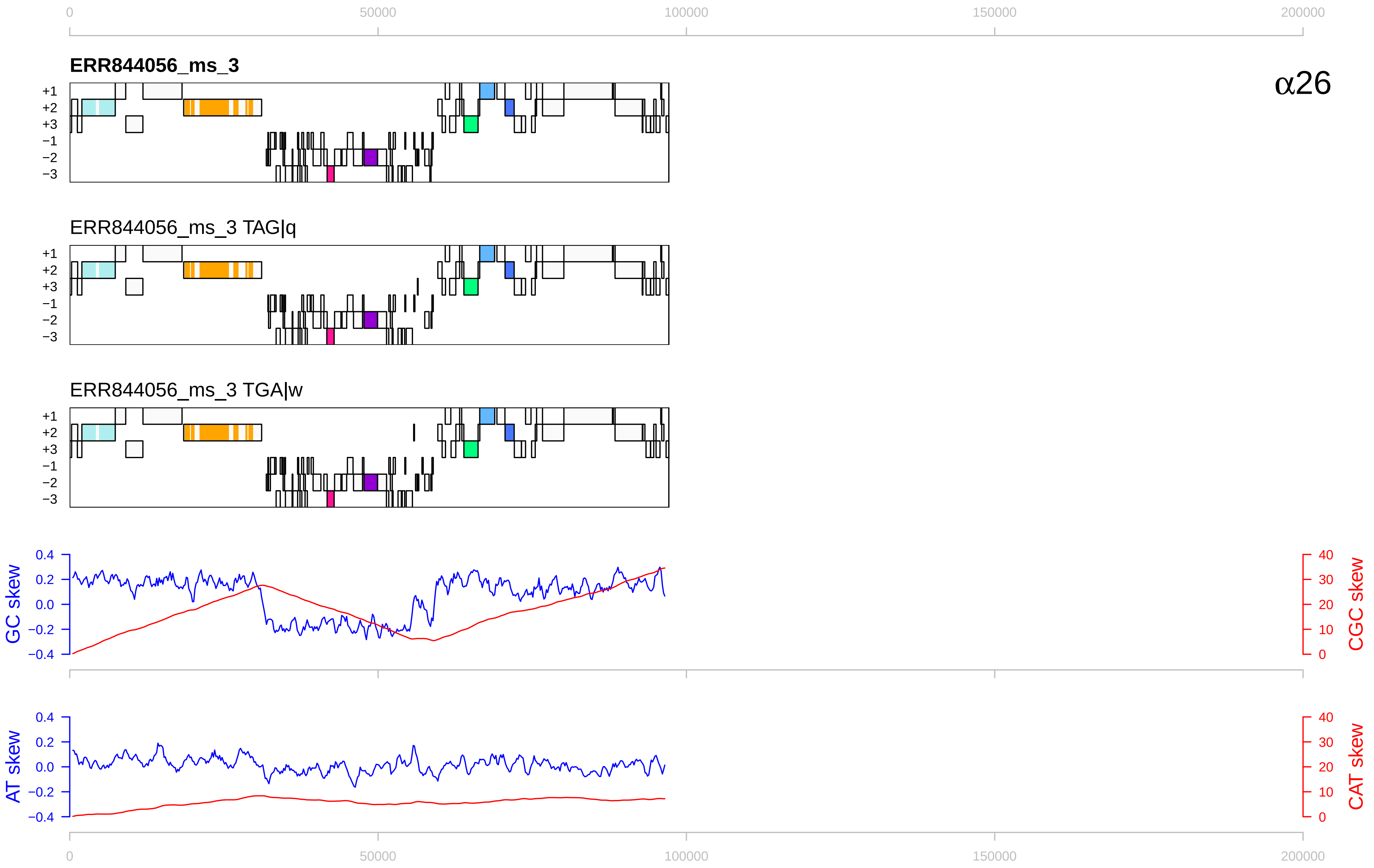

0
50000
100000
150000
200000
ERR844056_ms_3
α26
+1
+2
+3
−1
−2
−3
ERR844056_ms_3 TAG|q
ERR844056_ms_3 TGA|w
GC skew
CGC skew
0.4
0.2
0.0
−0.2
−0.4
40
30
20
10
0
AT skew
CAT skew

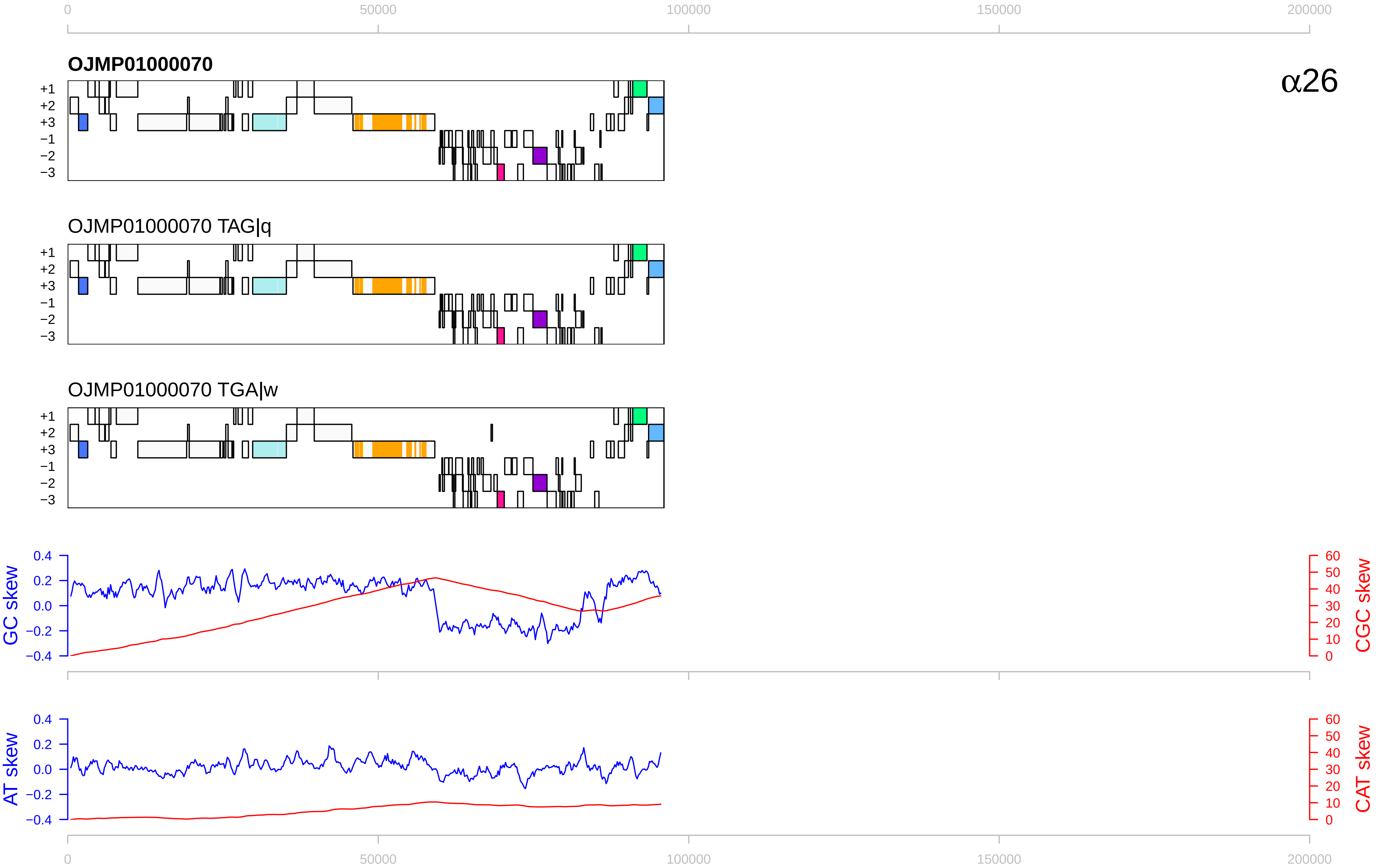

0
50000
100000
150000
200000
OJMP01000070
α26
+1
+2
+3
−1
−2
−3
OJMP01000070 TAG|q
OJMP01000070 TGA|w
GC skew
CGC skew
AT skew
CAT skew
0.4
0.2
0.0
−0.2
−0.4
60
50
40
30
20
10
0

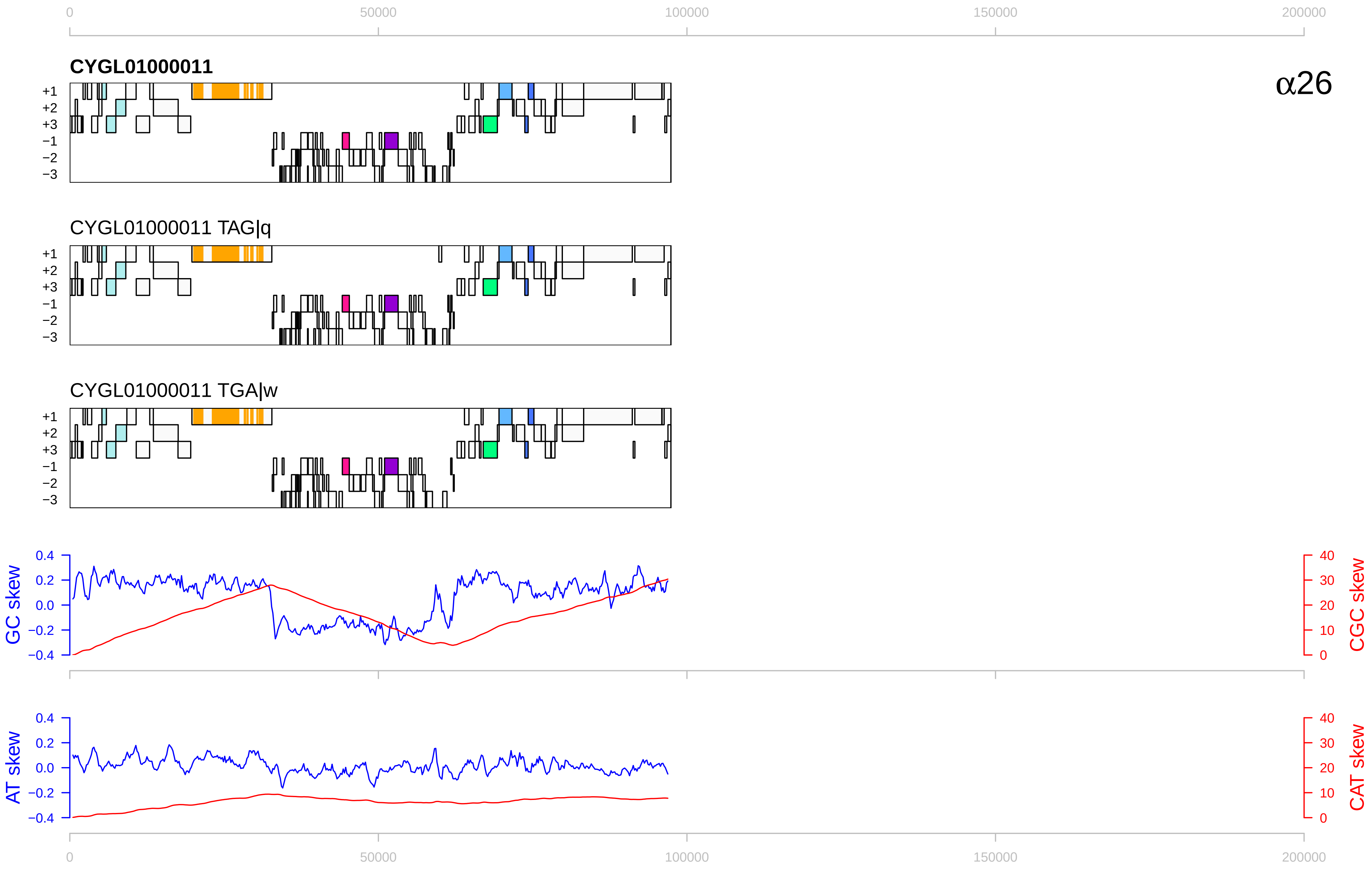

0
50000
100000
150000
200000
CYGL01000011
α26
+1
+2
+3
−1
−2
−3
CYGL01000011 TAG|q
CYGL01000011 TGA|w
GC skew
CGC skew
AT skew
CAT skew
0.4
0.2
0.0
−0.2
−0.4
40
30
20
10
0

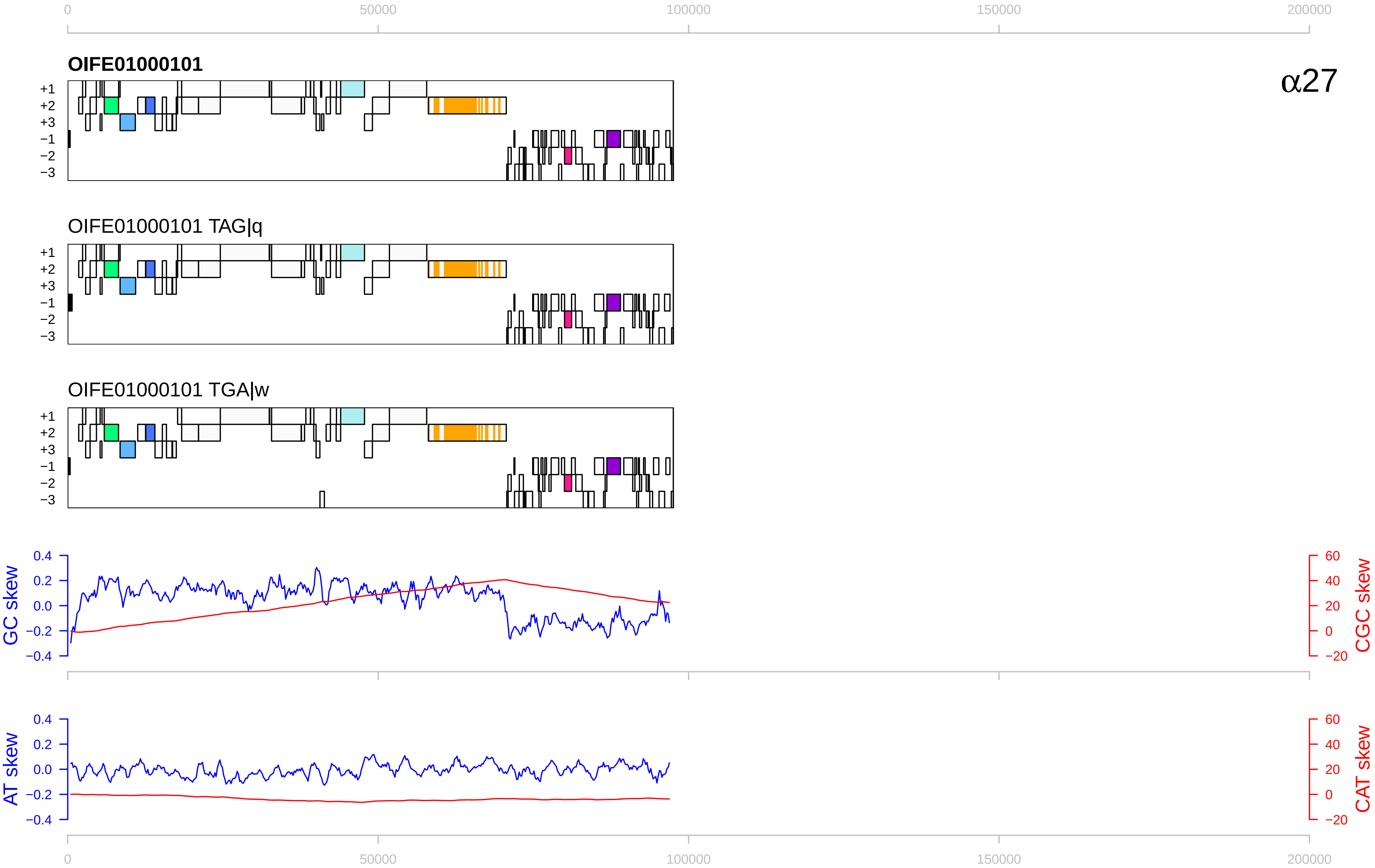

0
50000
100000
150000
200000
OIFE01000101
α27
+1
+2
+3
−1
−2
−3
OIFE01000101 TAG|q
OIFE01000101 TGA|w
GC skew
CGC skew
0.4
0.2
0.0
−0.2
−0.4
60
40
20
0
−20
AT skew
CAT skew

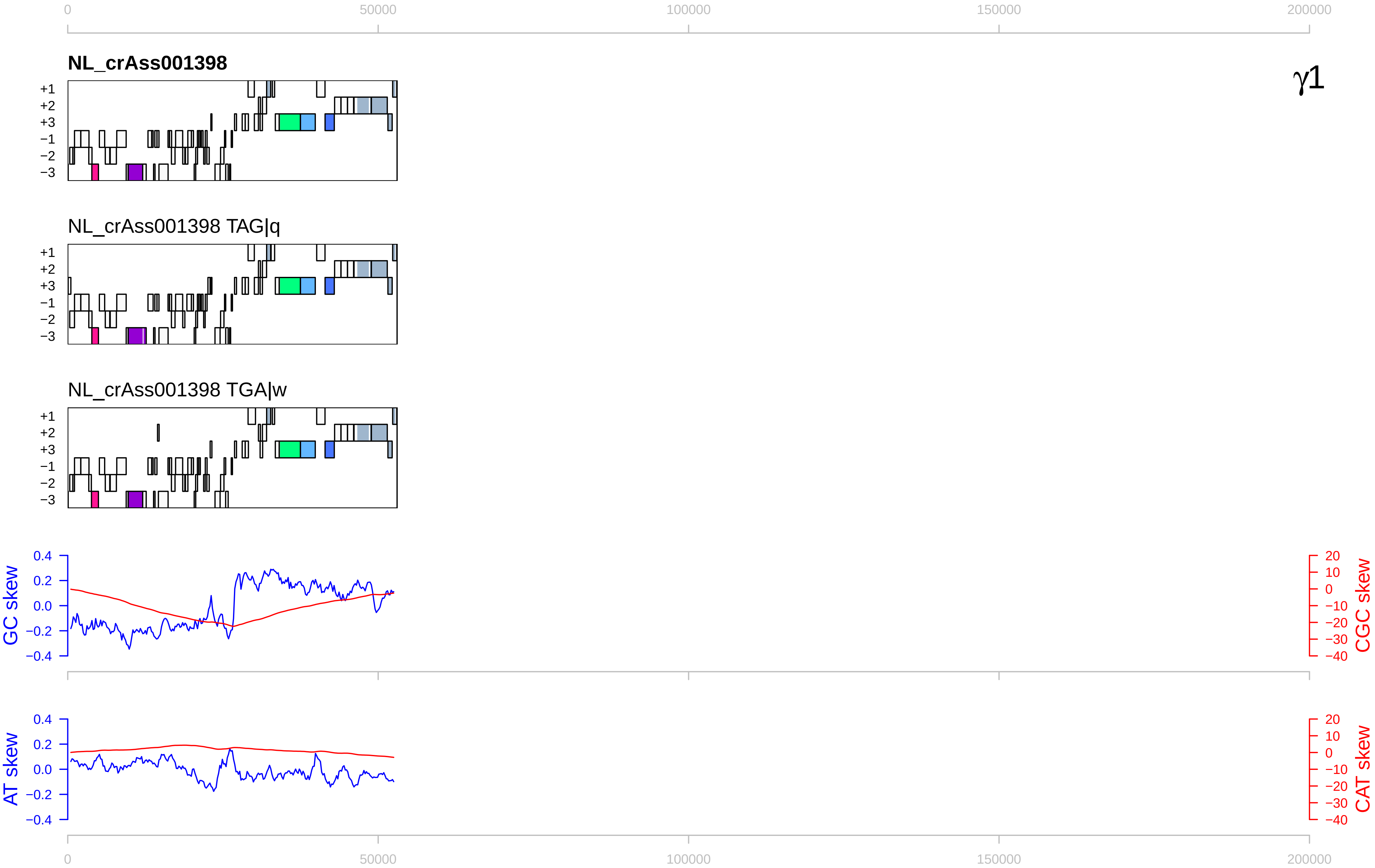

0
50000
100000
150000
200000
NL_crAss001398
γ1
+1
+2
+3
−1
−2
−3
NL_crAss001398 TAG|q
NL_crAss001398 TGA|w
GC skew
CGC skew
AT skew
CAT skew
0.4
0.2
0.0
−0.2
−0.4
20
10
0
−10
−20
−30
−40

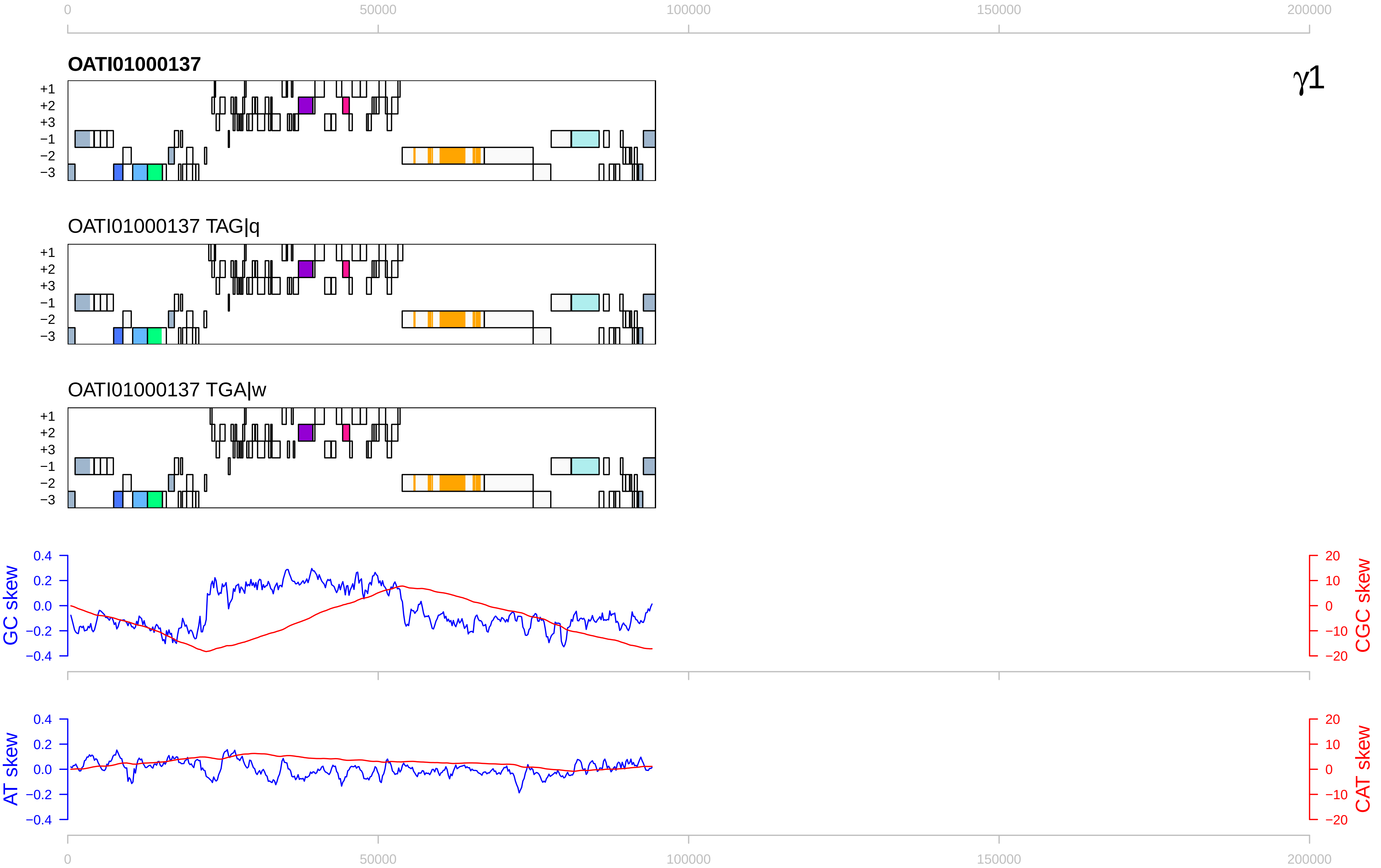

0
50000
100000
150000
200000
OATI01000137
γ1
+1
+2
+3
−1
−2
−3
OATI01000137 TAG|q
OATI01000137 TGA|w
GC skew
CGC skew
AT skew
CAT skew
0.4
0.2
0.0
−0.2
−0.4
20
10
−10
−20

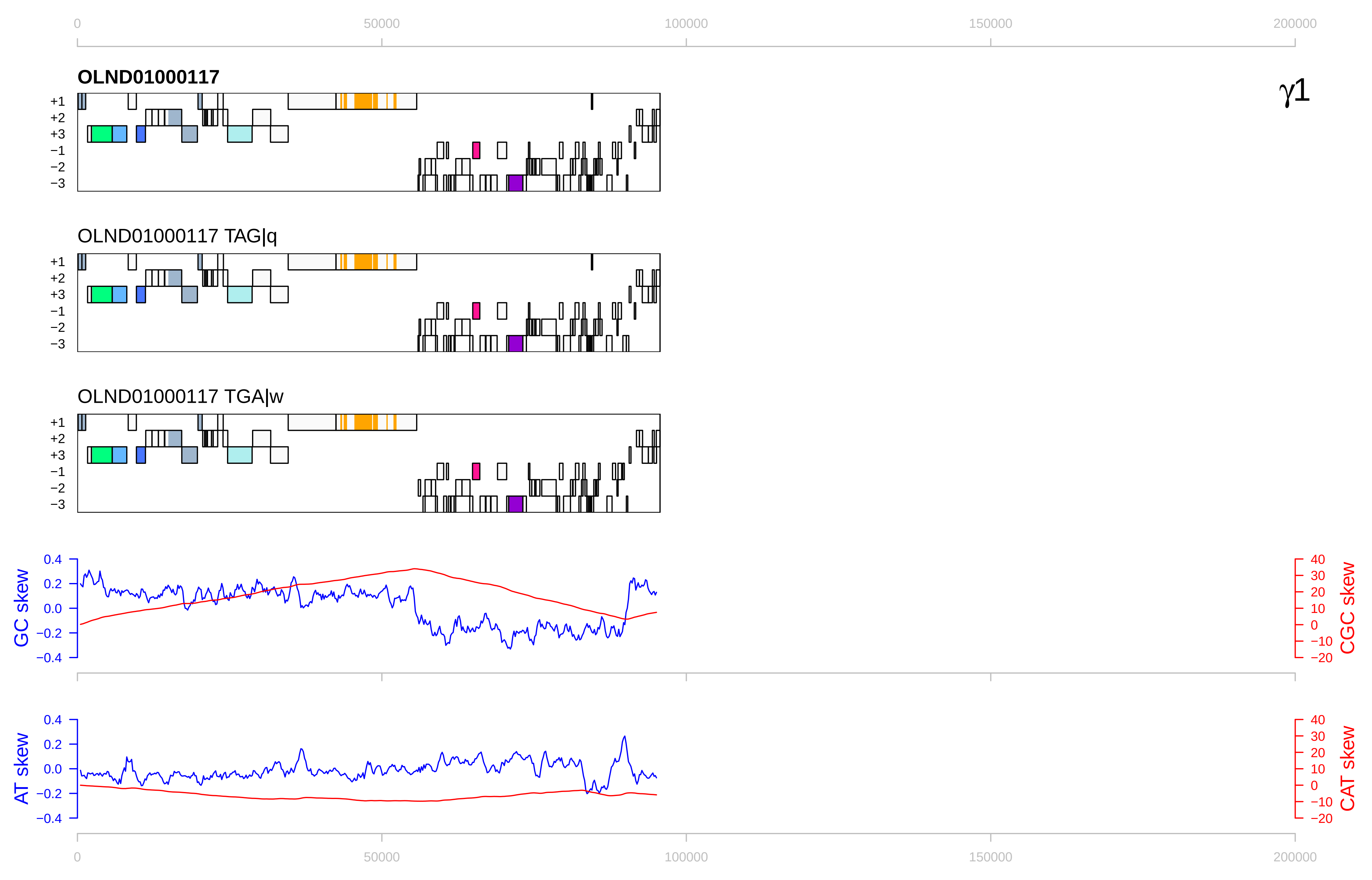

0
50000
100000
150000
200000
OLND01000117
γ1
+1
+2
+3
−1
−2
−3
OLND01000117 TAG|q
OLND01000117 TGA|w
GC skew
CGC skew
AT skew
CAT skew
0.4
0.2
0.0
−0.2
−0.4
40
30
20
10
0
−10
−20

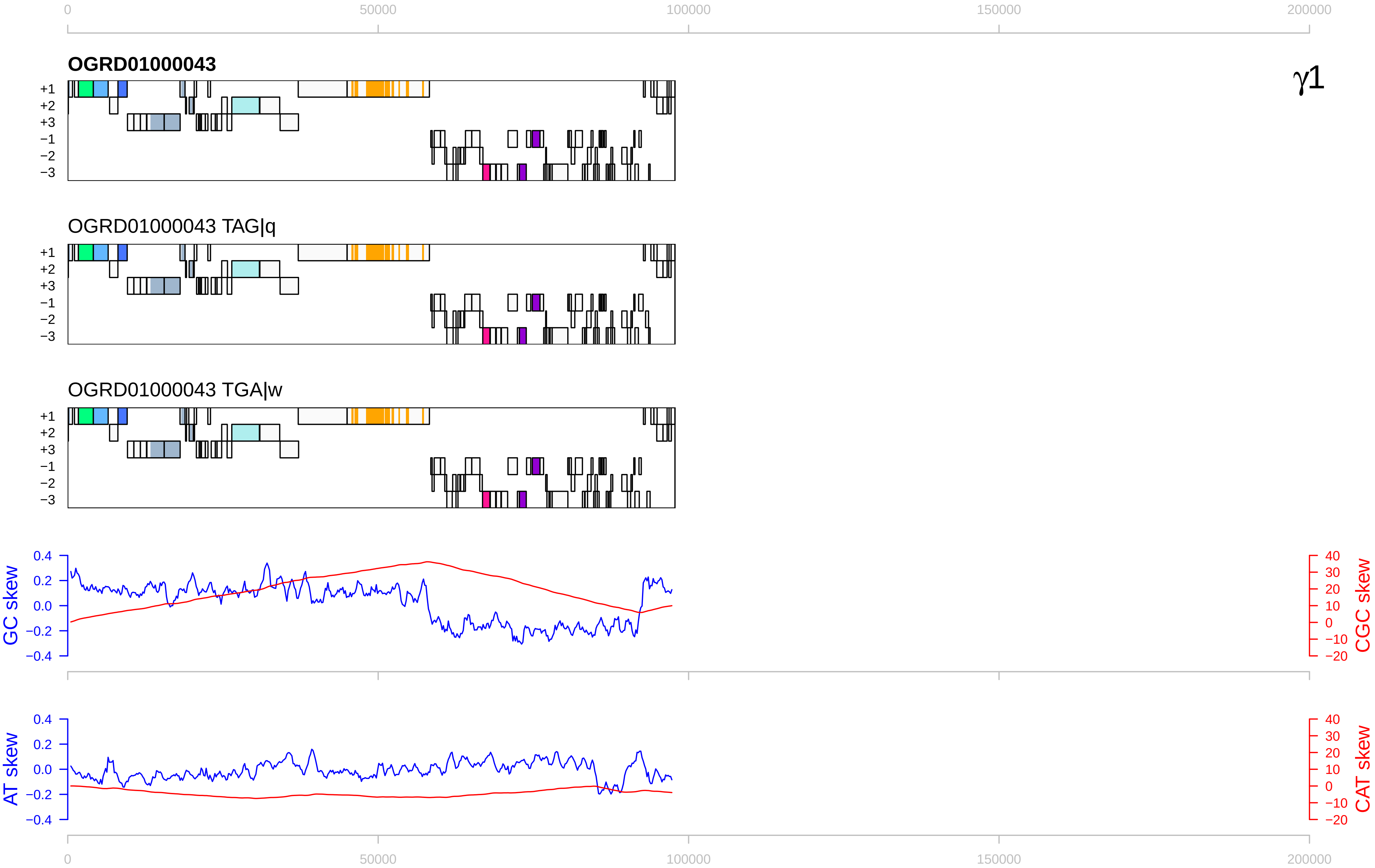

0
50000
100000
150000
200000
OGRD01000043
γ1
+1
+2
+3
−1
−2
−3
OGRD01000043 TAG|q
OGRD01000043 TGA|w
GC skew
CGC skew
AT skew
CAT skew
0.4
0.2
0.0
−0.2
−0.4
40
30
20
10
0
−10
−20

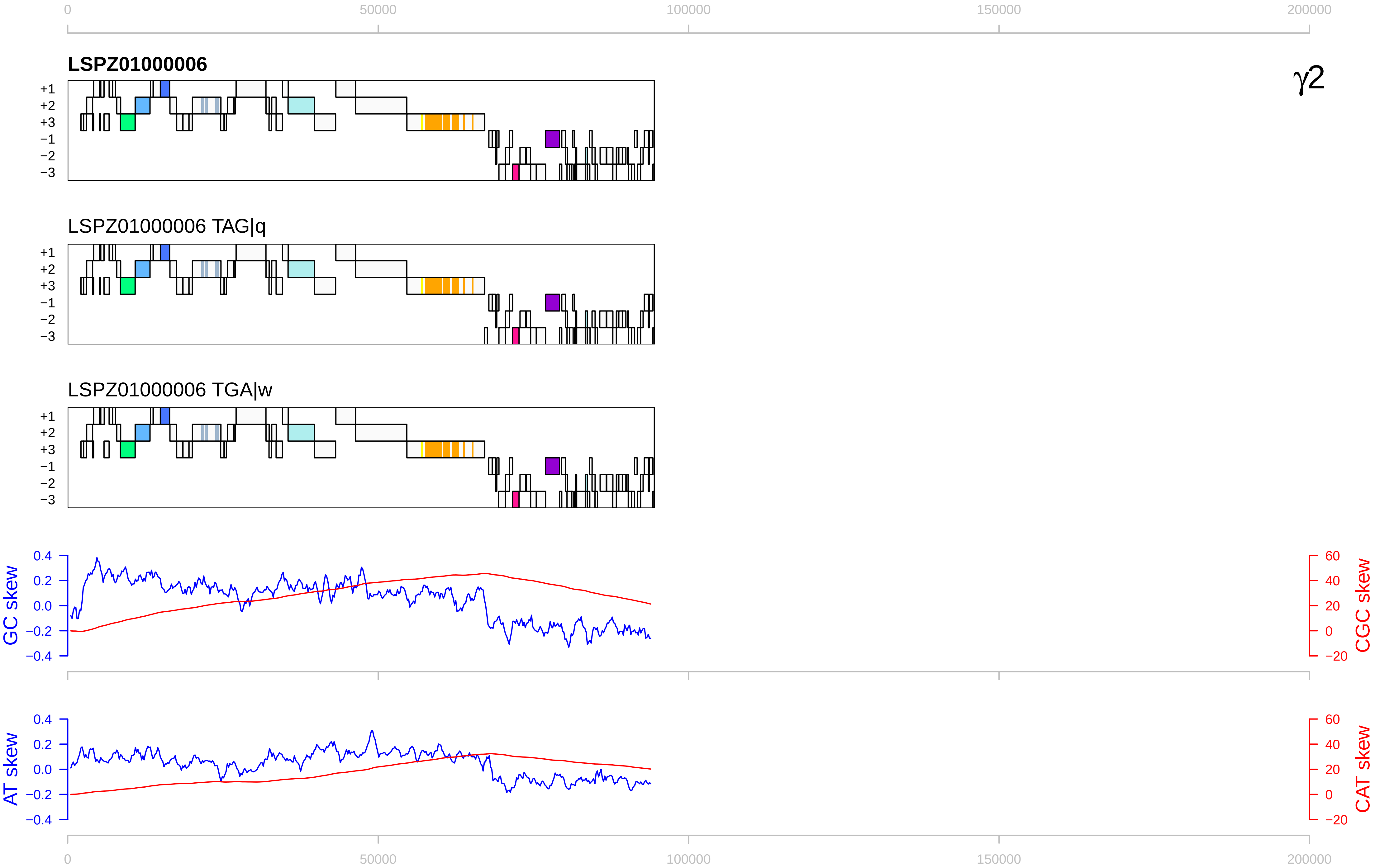

0
50000
100000
150000
200000
LSPZ01000006
γ2
+1
+2
+3
−1
−2
−3
LSPZ01000006 TAG|q
LSPZ01000006 TGA|w
GC skew
CGC skew
0.4
0.2
0.0
−0.2
−0.4
60
40
20
0
−20
AT skew
CAT skew

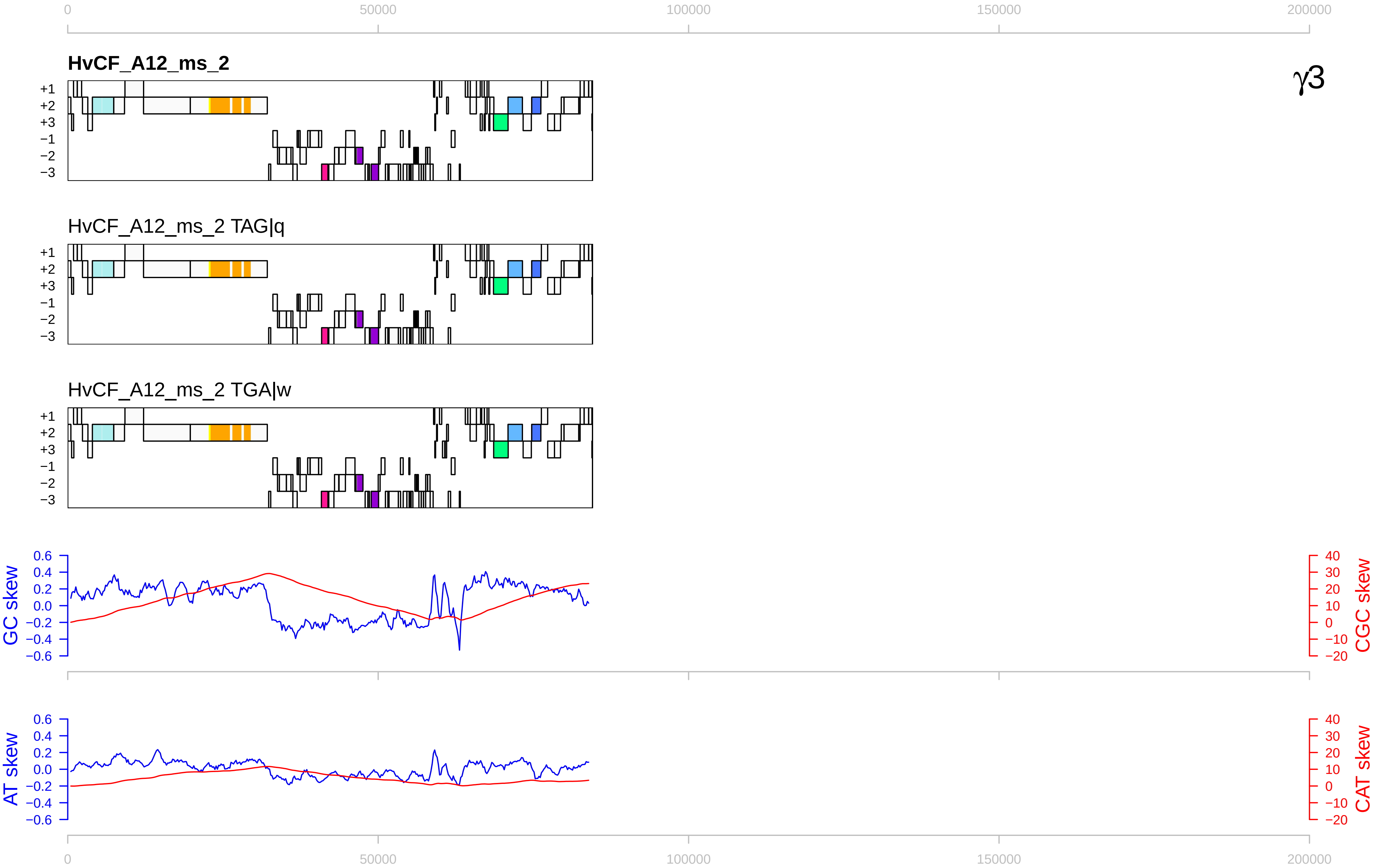
0
50000
100000
150000
200000
HvCF_A12_ms_2
γ3
+1
+2
+3
−1
−2
−3
HvCF_A12_ms_2 TAG|q
HvCF_A12_ms_2 TGA|w
GC skew
CGC skew
AT skew
CAT skew
0.6
0.4
0.2
0.0
−0.2
−0.4
−0.6
40
30
20
10
0
−10
−20

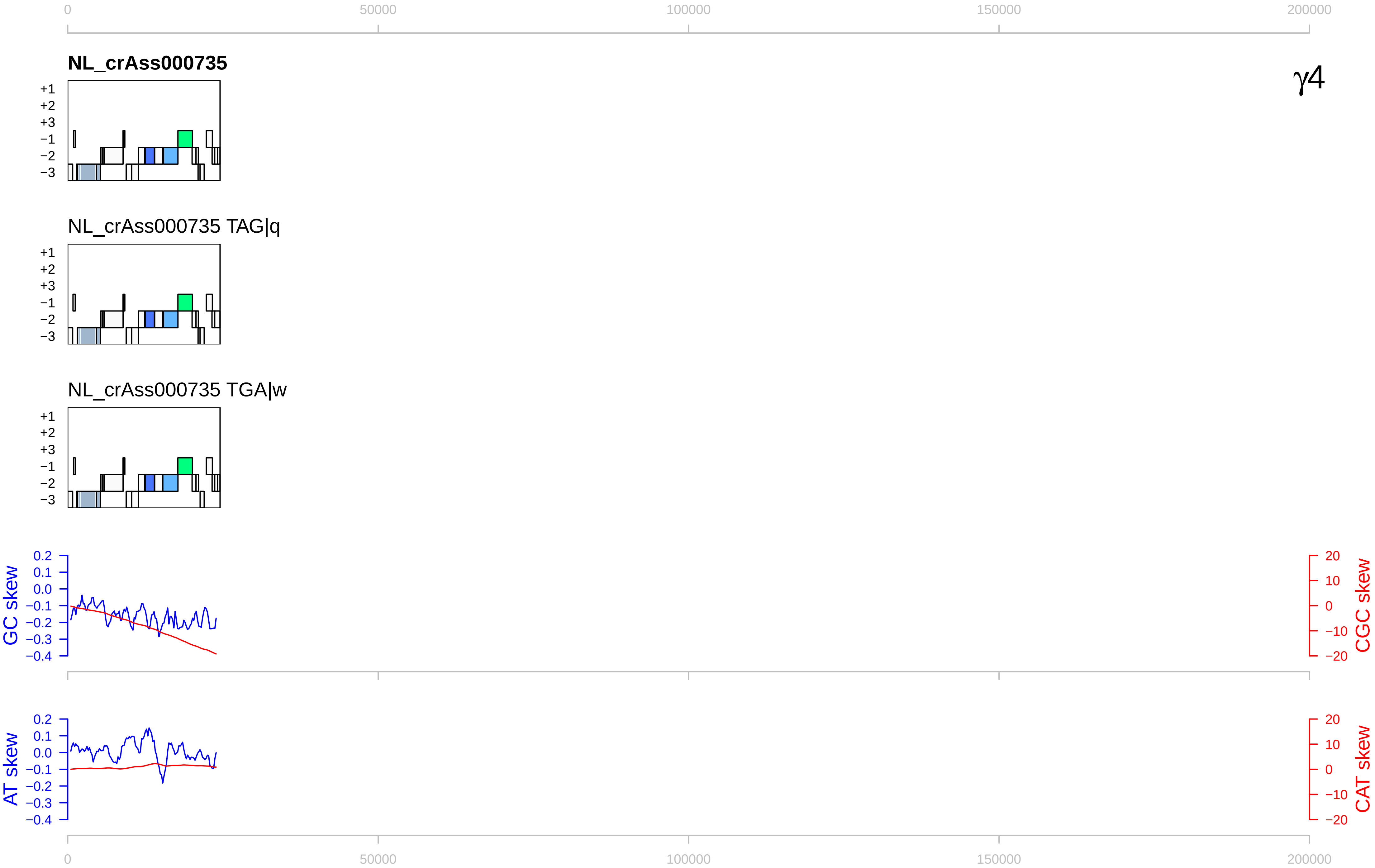

0
50000
100000
150000
200000
NL_crAss000735
γ4
+1
+2
+3
−1
−2
−3
NL_crAss000735 TAG|q
NL_crAss000735 TGA|w
GC skew
0.2
0.1
0.0
−0.1
−0.2
−0.3
−0.4
CGC skew
20
10
0
−10
−20
AT skew
CAT skew

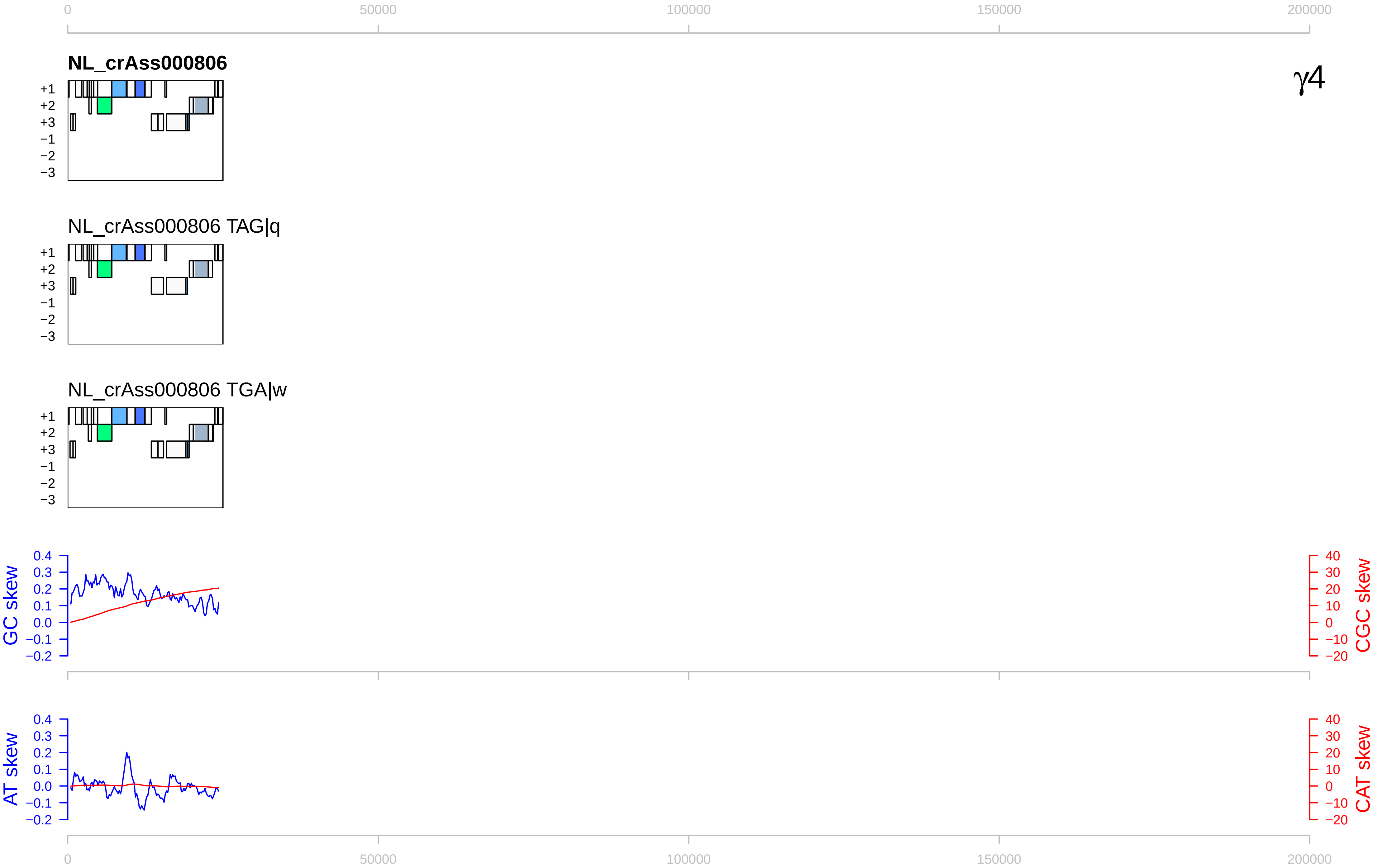
0
50000
100000
150000
200000
NL_crAss000806
γ4
+1
+2
+3
−1
−2
−3
NL_crAss000806 TAG|q
NL_crAss000806 TGA|w
GC skew
CGC skew
AT skew
CAT skew
0.4
0.3
0.2
0.1
0.0
−0.1
−0.2
40
30
20
10
0
−10
−20

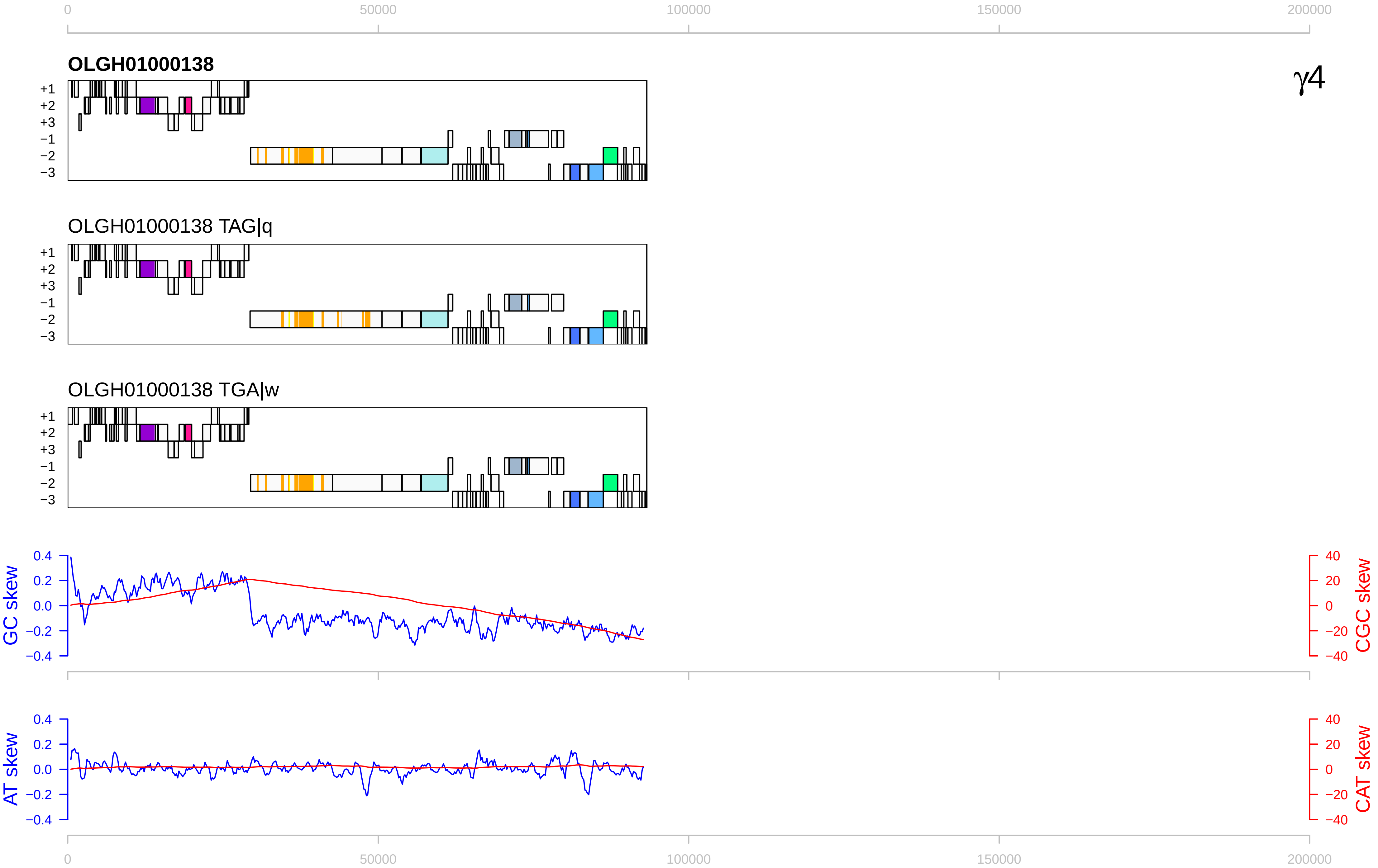

0
50000
100000
150000
200000
OLGH01000138
γ4
+1
+2
+3
−1
−2
−3
OLGH01000138 TAG|q
OLGH01000138 TGA|w
GC skew
CGC skew
AT skew
CAT skew
0.4
0.2
0.0
−0.2
−0.4
40
20
0
−20
−40

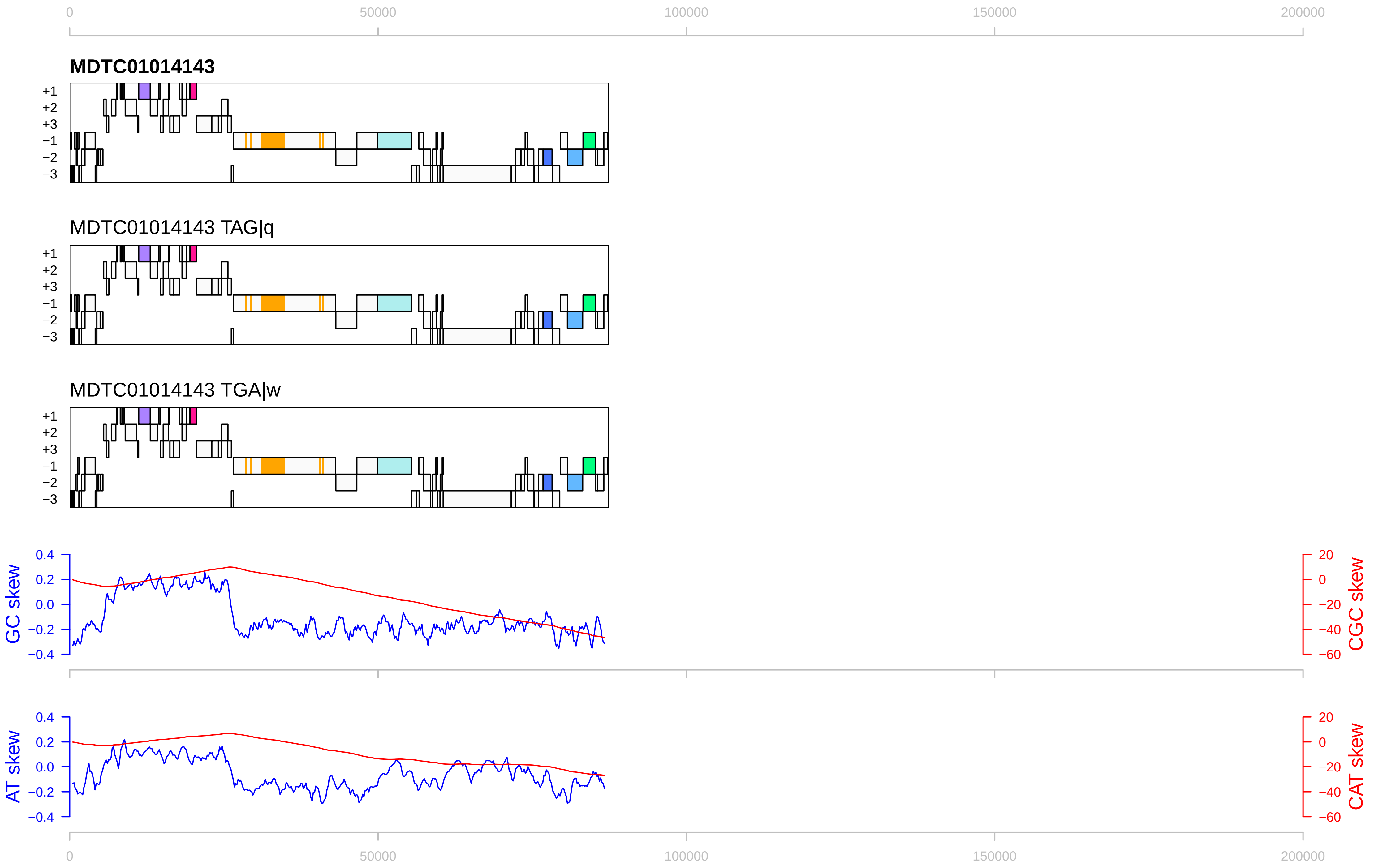

0
50000
100000
150000
200000
MDTC01014143
+1
+2
+3
−1
−2
−3
MDTC01014143 TAG|q
MDTC01014143 TGA|w
GC skew
CGC skew
AT skew
CAT skew
0.4
0.2
0.0
−0.2
−0.4
20
0
−20
−40
−60

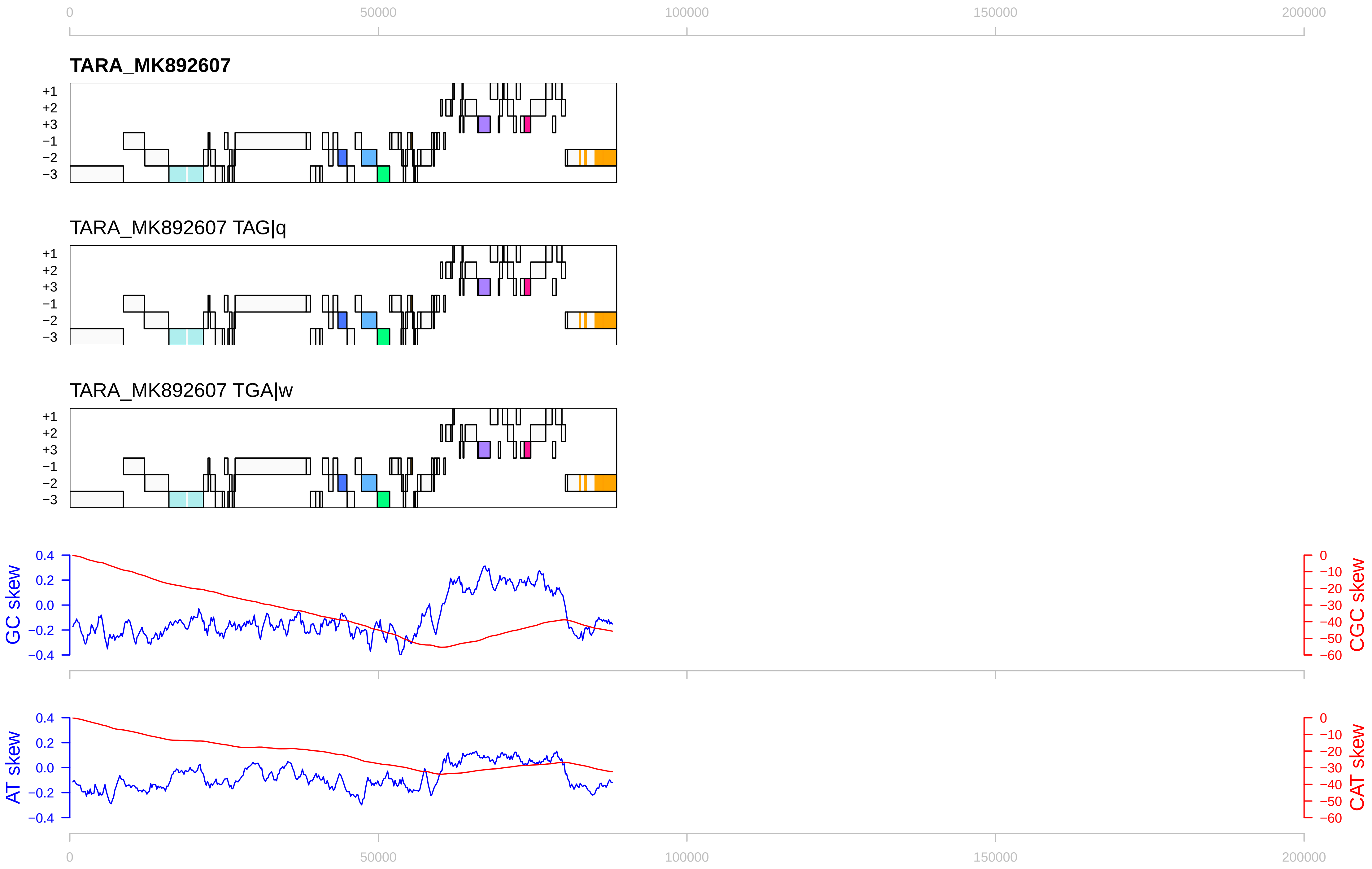

0
50000
100000
150000
200000
TARA_MK892607
+1
+2
+3
−1
−2
−3
TARA_MK892607 TAG|q
TARA_MK892607 TGA|w
GC skew
CGC skew
AT skew
CAT skew
0.4
0.2
0.0
−0.2
−0.4
0
−10
−20
−30
−40
−50
−60

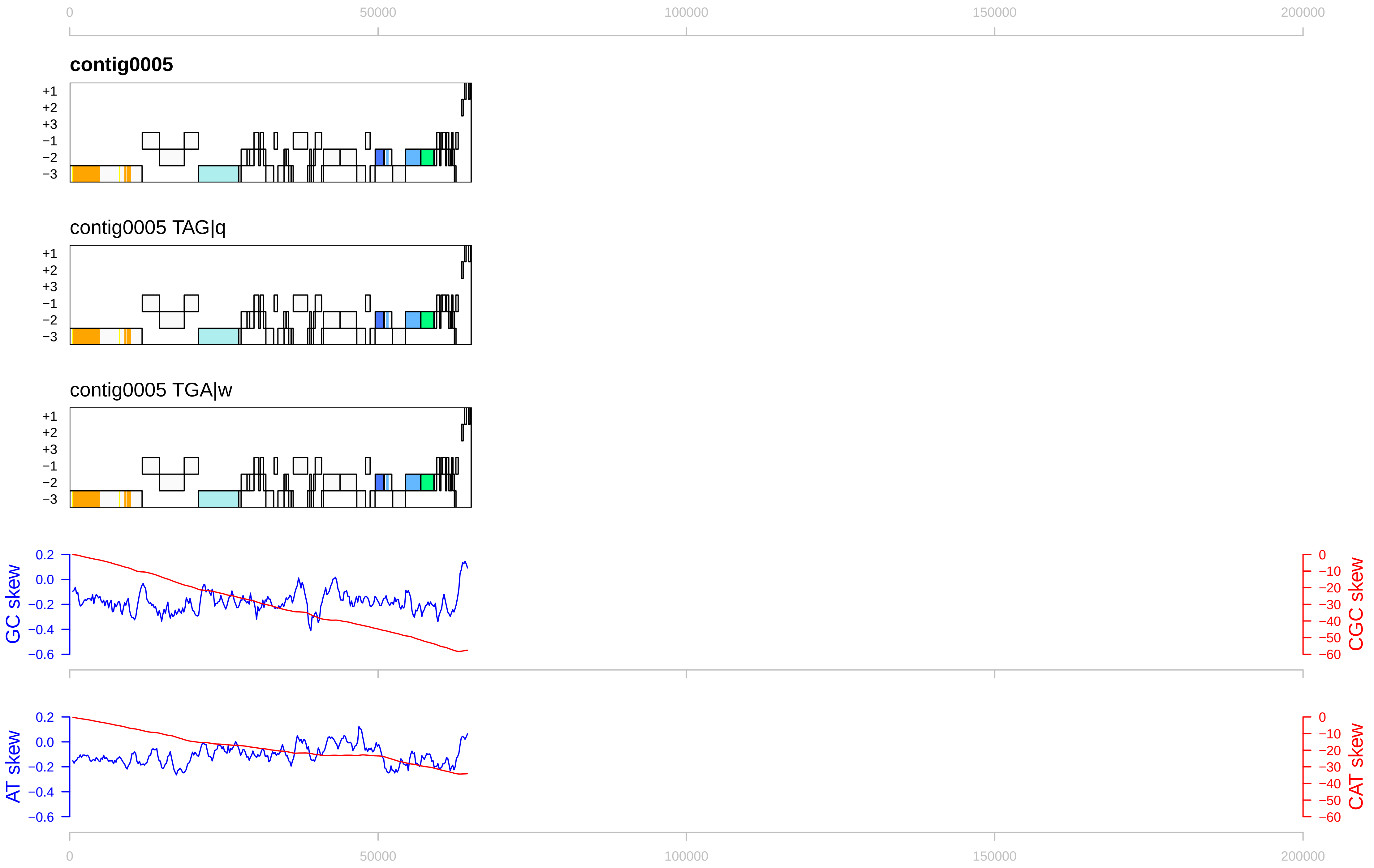

0
50000
100000
150000
200000
contig0005
+1
+2
+3
−1
−2
−3
contig0005 TAG|q
contig0005 TGA|w
GC skew
CGC skew
AT skew
CAT skew
0.2
0.0
−0.2
−0.4
−0.6
0
−10
−20
−30
−40
−50
−60

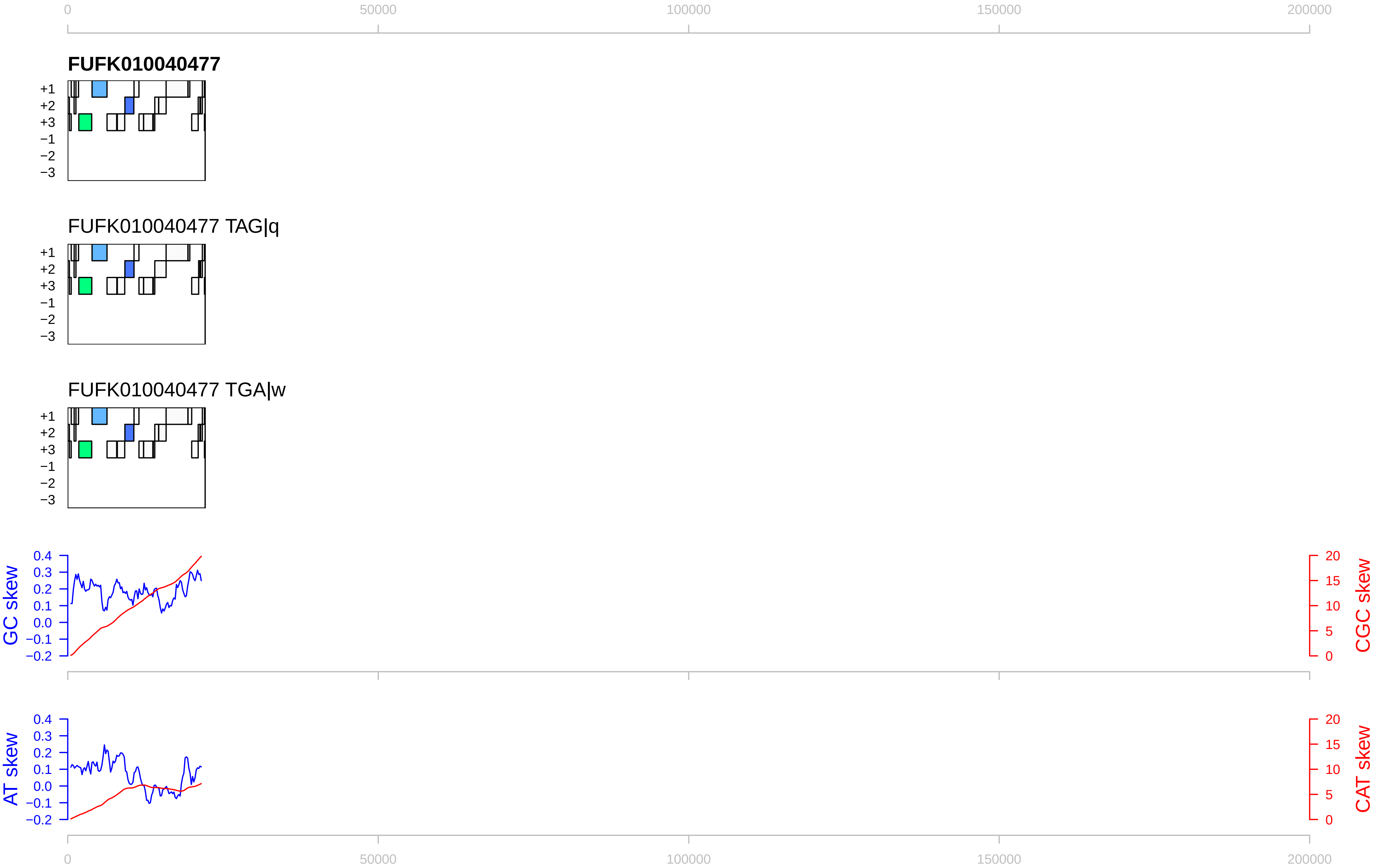
0
50000
100000
150000
200000
FUFK010040477
+1
+2
+3
−1
−2
−3
FUFK010040477 TAG|q
FUFK010040477 TGA|w
GC skew
CGC skew
AT skew
CAT skew
0.4
0.3
0.2
0.1
0.0
−0.1
−0.2
20
15
10
5
0

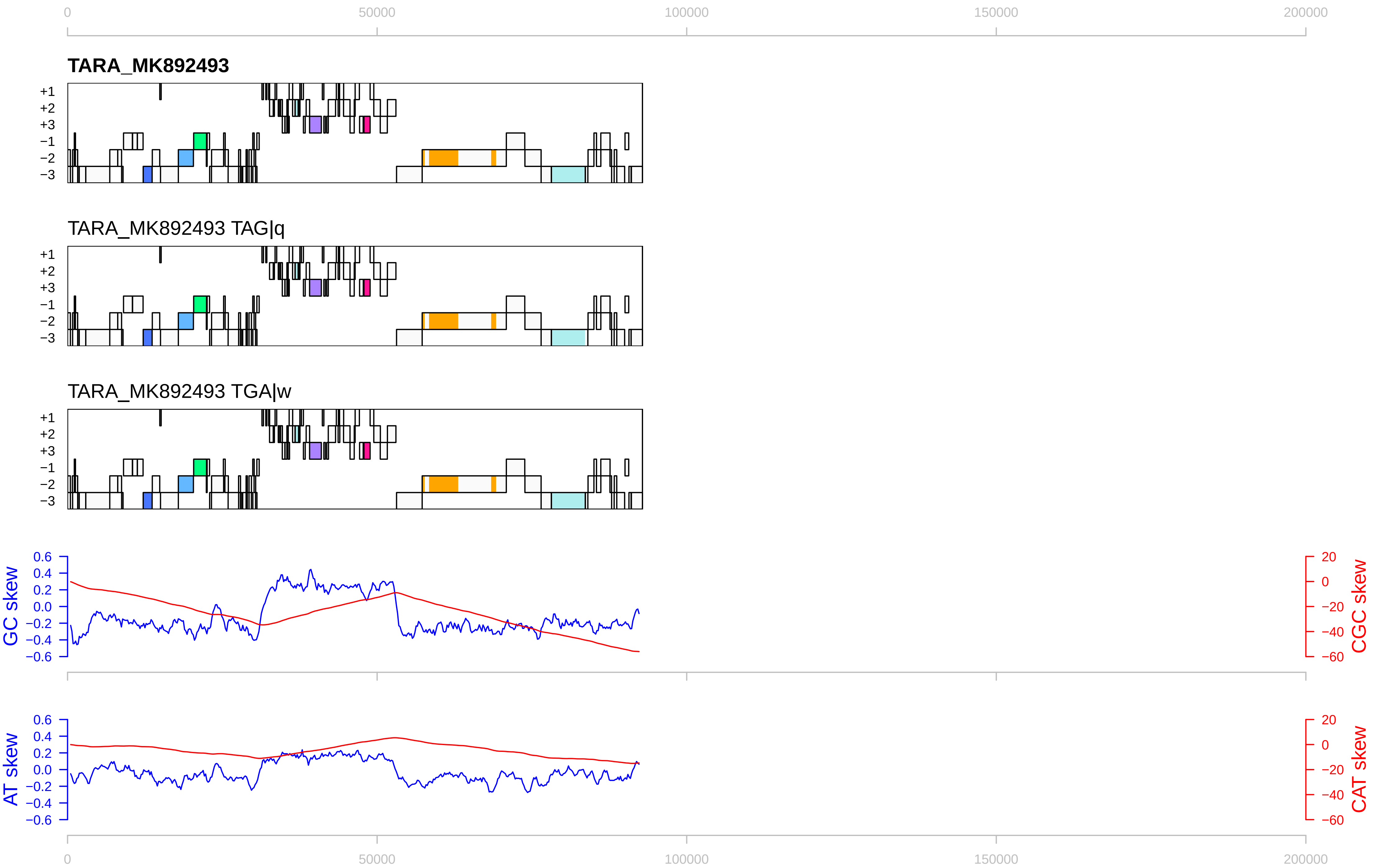

0
50000
100000
150000
200000
TARA_MK892493
+1
+2
+3
−1
−2
−3
TARA_MK892493 TAG|q
TARA_MK892493 TGA|w
GC skew
CGC skew
AT skew
CAT skew
0.6
0.4
0.2
0.0
−0.2
−0.4
−0.6
20
0
−20
−40
−60

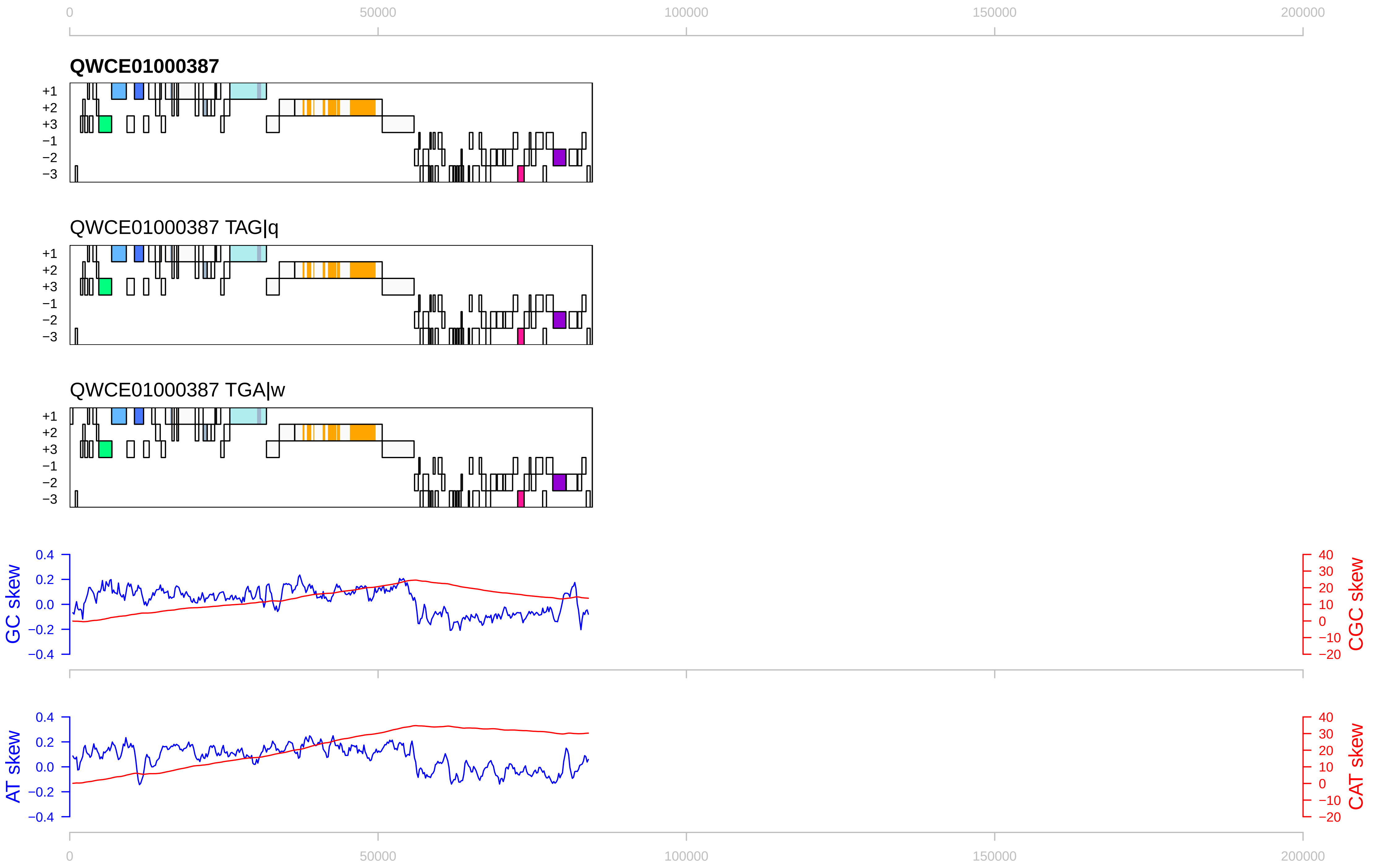

0
50000
100000
150000
200000
QWCE01000387
+1
+2
+3
−1
−2
−3
QWCE01000387 TAG|q
QWCE01000387 TGA|w
GC skew
CGC skew
AT skew
CAT skew
0.4
0.2
0.0
−0.2
−0.4
40
30
20
10
0
−10
−20

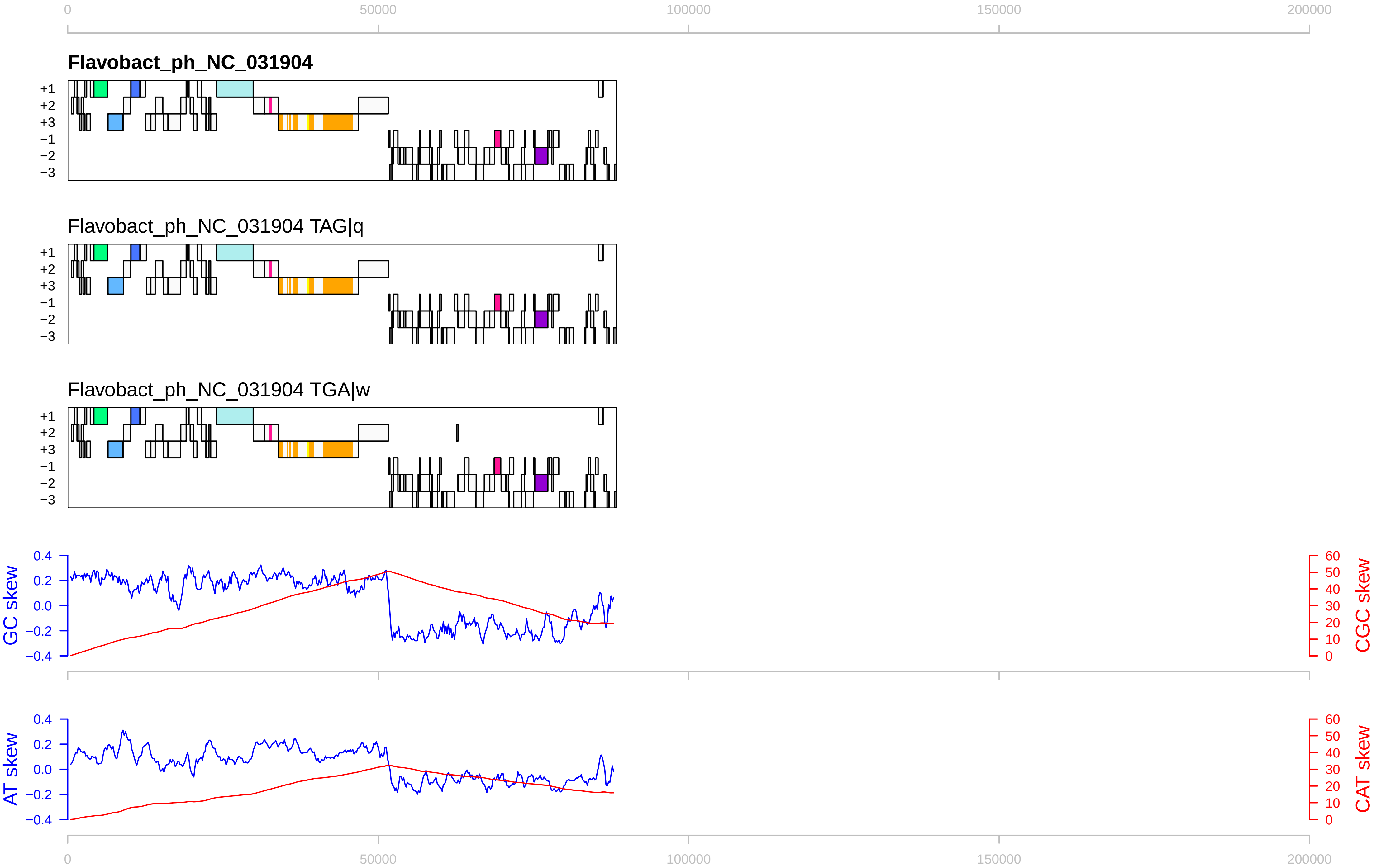

0
50000
100000
150000
200000
Flavobact_ph_NC_031904
+1
+2
+3
−1
−2
−3
Flavobact_ph_NC_031904 TAG|q
Flavobact_ph_NC_031904 TGA|w
GC skew
CGC skew
AT skew
CAT skew
0.4
0.2
0.0
−0.2
−0.4
60
50
40
30
20
10

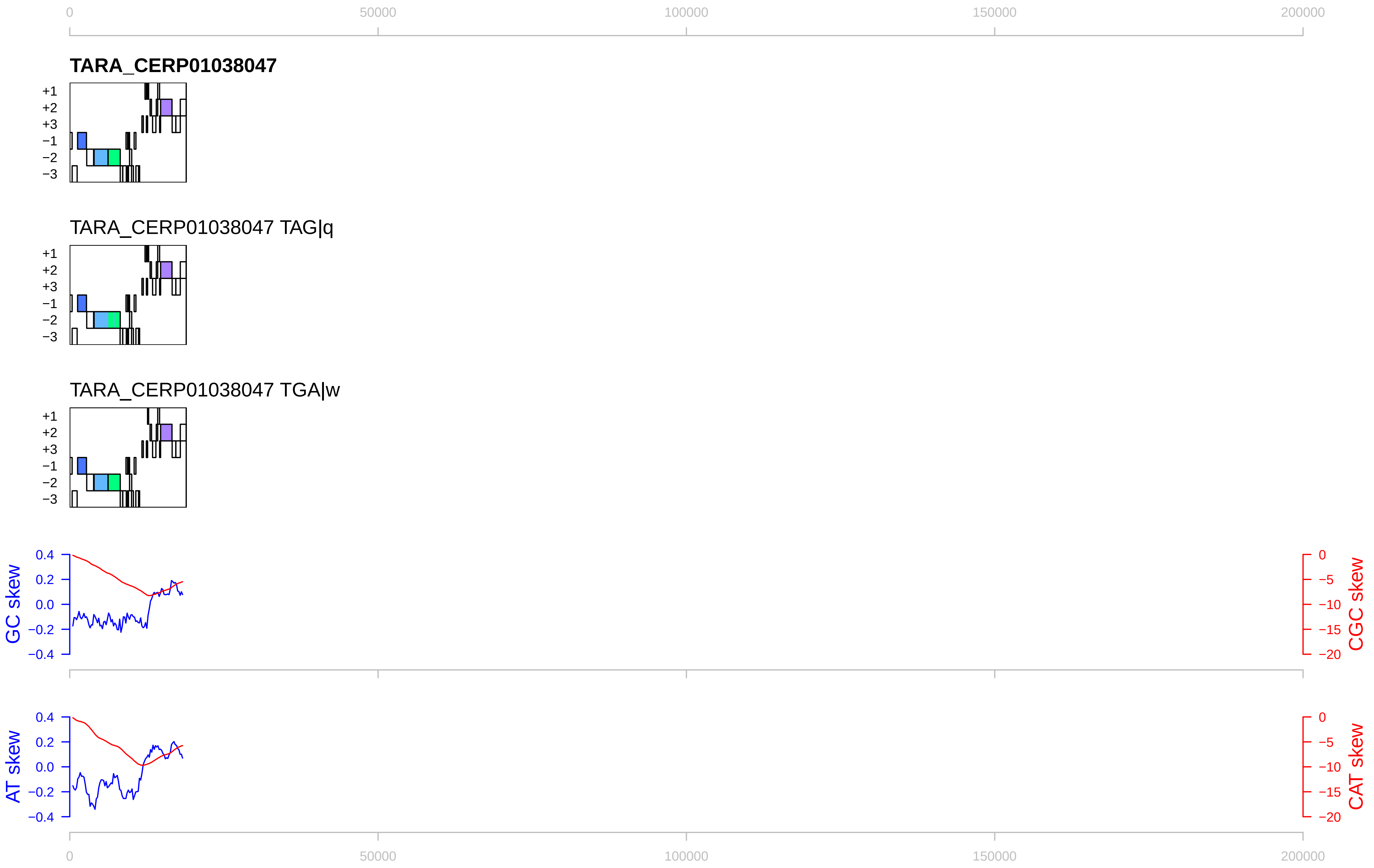

0
50000
100000
150000
200000
TARA_CERP01038047
+1
+2
+3
−1
−2
−3
TARA_CERP01038047 TAG|q
TARA_CERP01038047 TGA|w
GC skew
CGC skew
AT skew
CAT skew
0.4
0.2
0.0
−0.2
−0.4
0
−5
−10
−15
−20

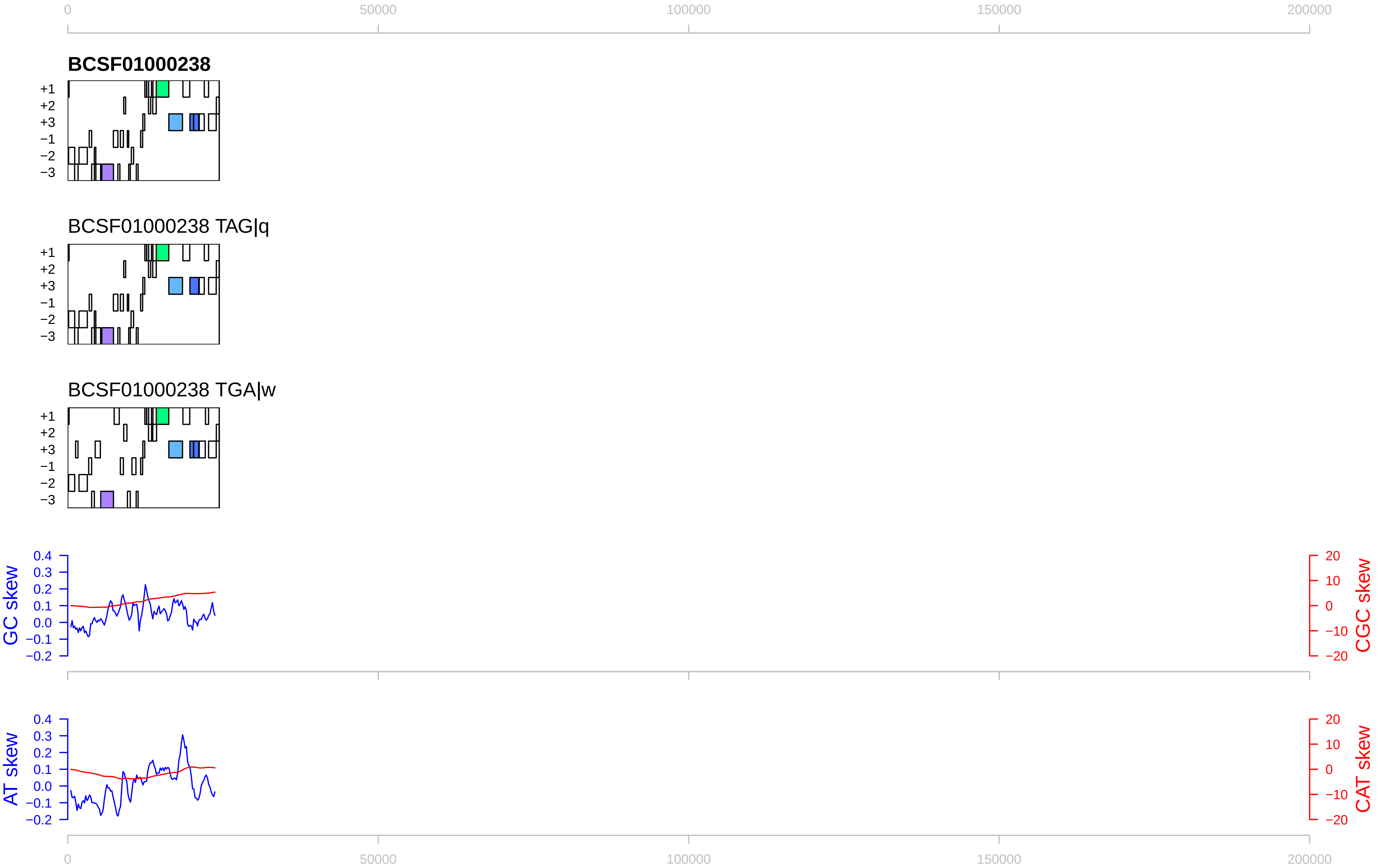
0
50000
100000
150000
200000
BCSF01000238
+1
+2
+3
−1
−2
−3
BCSF01000238 TAG|q
BCSF01000238 TGA|w
GC skew
CGC skew
AT skew
CAT skew
0.4
0.3
0.2
0.1
0.0
−0.1
−0.2
20
10
0
−10
−20

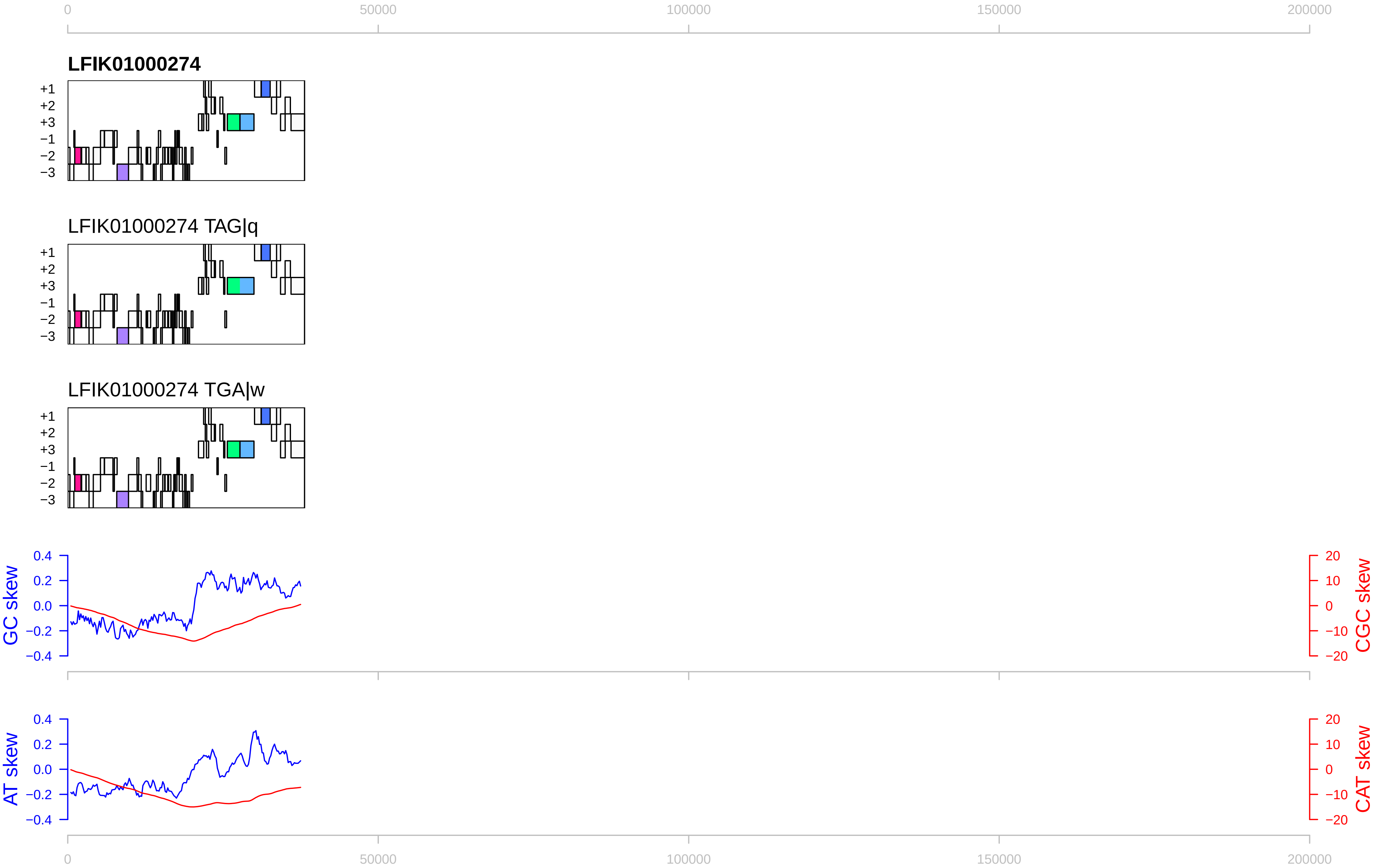

0
50000
100000
150000
200000
LFIK01000274
+1
+2
+3
−1
−2
−3
LFIK01000274 TAG|q
LFIK01000274 TGA|w
GC skew
CGC skew
0.4
0.2
0.0
−0.2
−0.4
20
10
0
−10
−20
AT skew
CAT skew

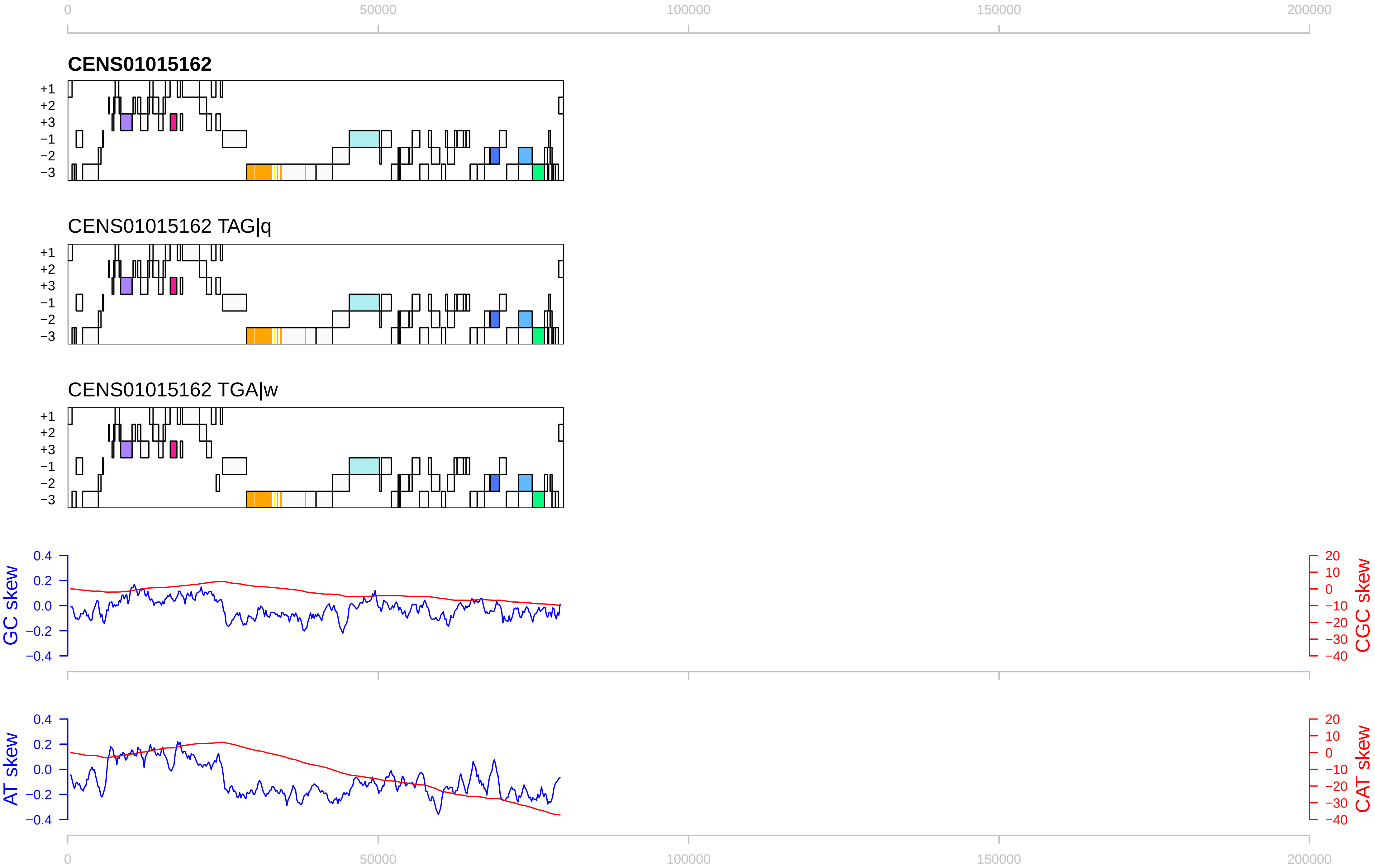

0
50000
100000
150000
200000
CENS01015162
+1
+2
+3
−1
−2
−3
CENS01015162 TAG|q
CENS01015162 TGA|w
GC skew
CGC skew
AT skew
CAT skew
0.4
0.2
0.0
−0.2
−0.4
20
10
0
−10
−20
−30
−40

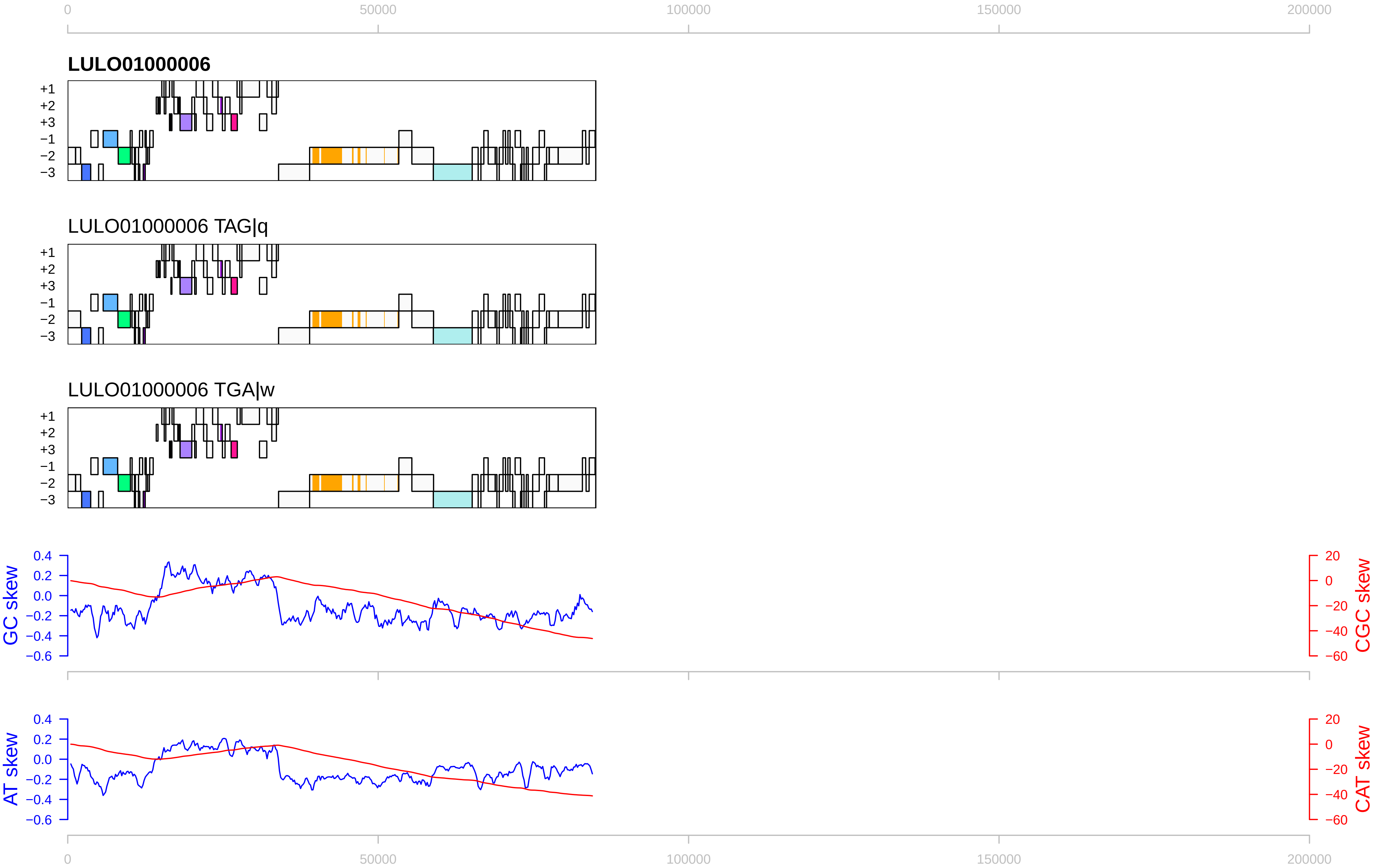

0
50000
100000
150000
200000
LULO01000006
+1
+2
+3
−1
−2
−3
LULO01000006 TAG|q
LULO01000006 TGA|w
GC skew
CGC skew
AT skew
CAT skew
0.4
0.2
0.0
−0.2
−0.4
−0.6
20
0
−20
−40
−60

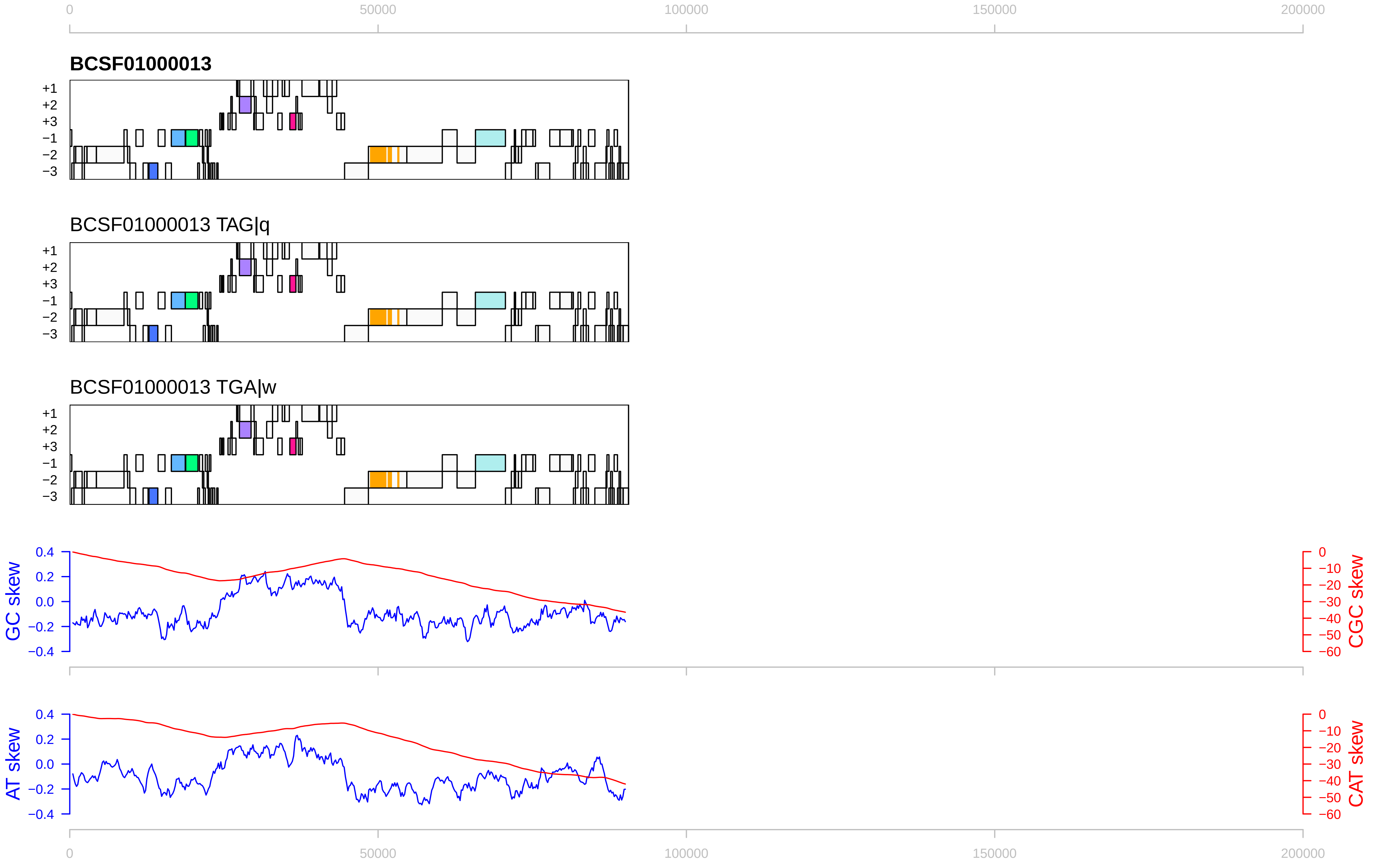

0
50000
100000
150000
200000
BCSF01000013
+1
+2
+3
−1
−2
−3
BCSF01000013 TAG|q
BCSF01000013 TGA|w
GC skew
CGC skew
AT skew
CAT skew
0.4
0.2
0.0
−0.2
−0.4
0
−10
−20
−30
−40
−50
−60

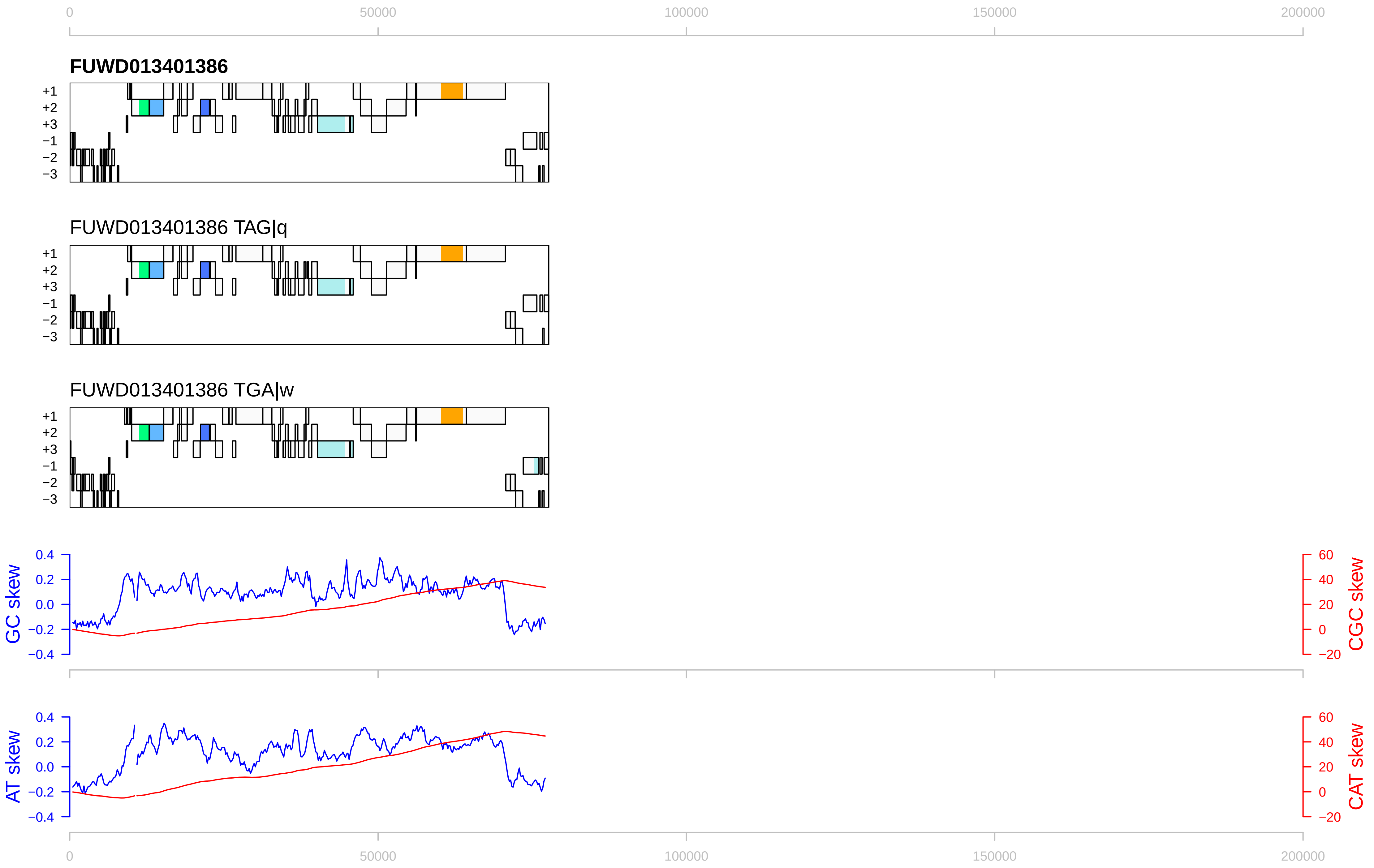

0
50000
100000
150000
200000
FUWD013401386
+1
+2
+3
−1
−2
−3
FUWD013401386 TAG|q
FUWD013401386 TGA|w
GC skew
CGC skew
AT skew
CAT skew
0.4
0.2
0.0
−0.2
−0.4
60
40
20
0
−20

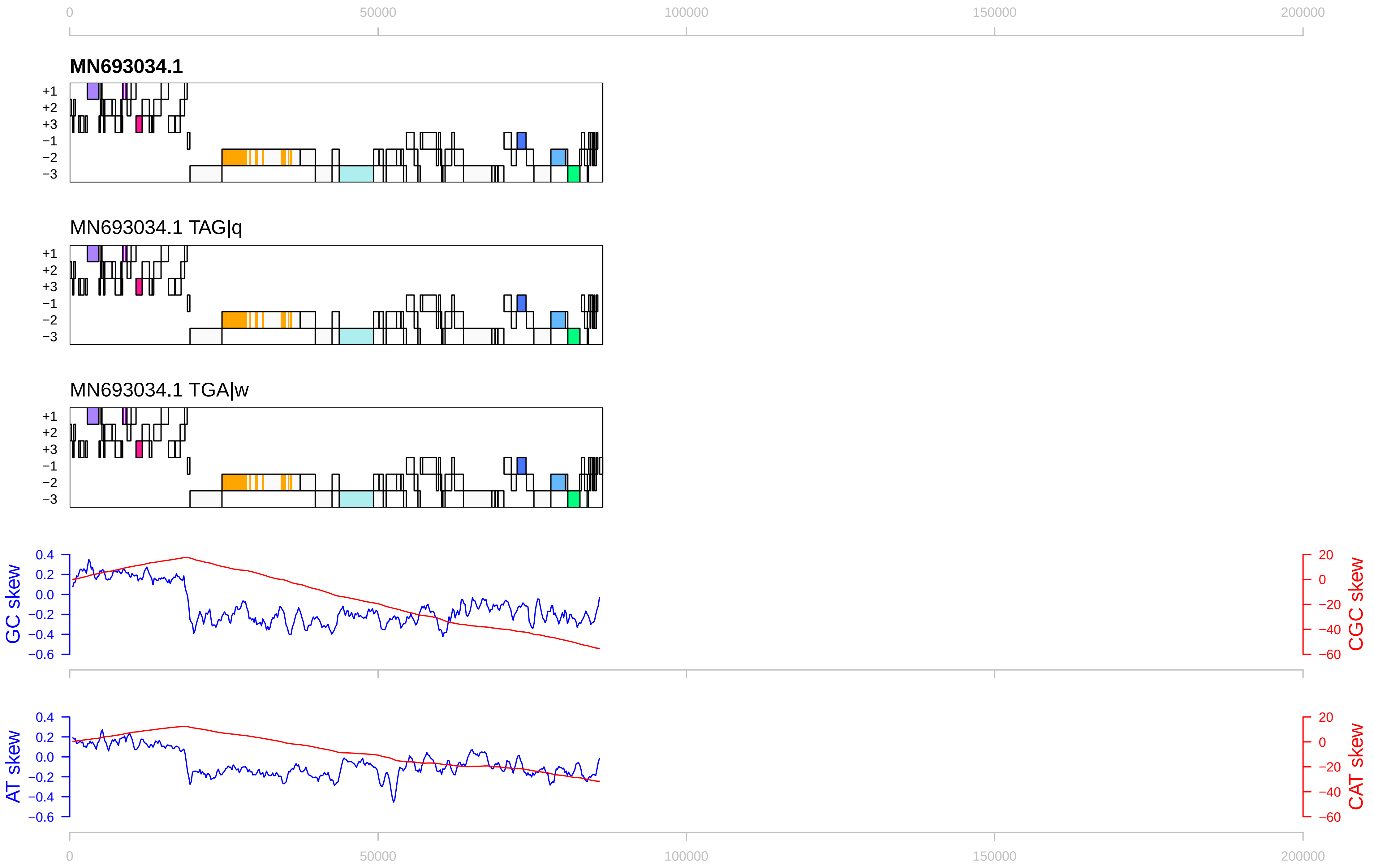

0
50000
100000
150000
200000
MN693034.1
+1
+2
+3
−1
−2
−3
MN693034.1 TAG|q
MN693034.1 TGA|w
GC skew
CGC skew
AT skew
CAT skew
0.4
0.2
0.0
−0.2
−0.4
−0.6
20
0
−20
−40
−60

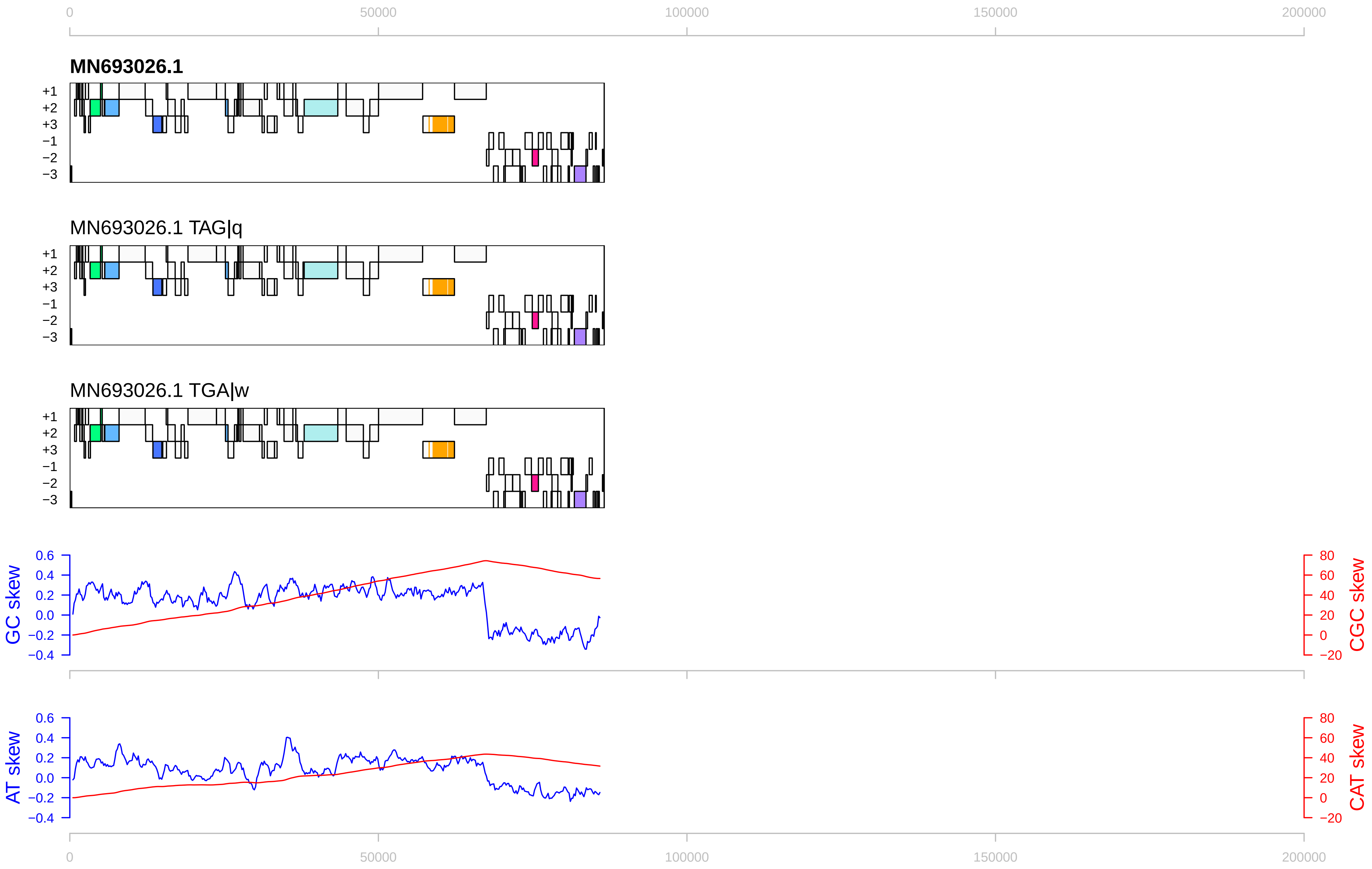

0
50000
100000
150000
200000
MN693026.1
+1
+2
+3
−1
−2
−3
MN693026.1 TAG|q
MN693026.1 TGA|w
GC skew
CGC skew
0.6
0.4
0.2
0.0
−0.2
−0.4
80
60
40
20
0
−20
AT skew
CAT skew

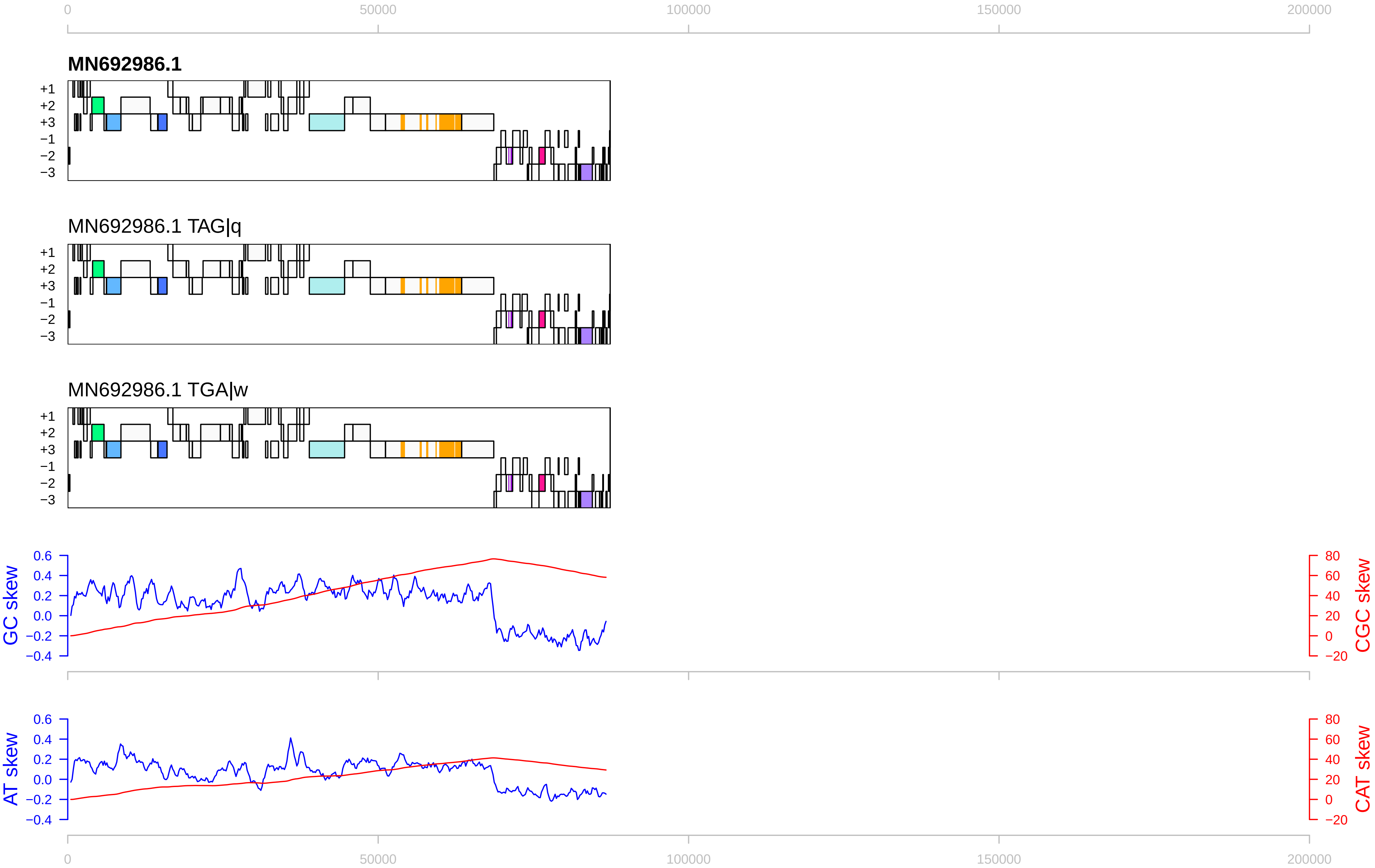

0
50000
100000
150000
200000
MN692986.1
+1
+2
+3
−1
−2
−3
MN692986.1 TAG|q
MN692986.1 TGA|w
GC skew
CGC skew
AT skew
CAT skew
0.6
0.4
0.2
0.0
−0.2
−0.4
80
60
40
20
0
−20

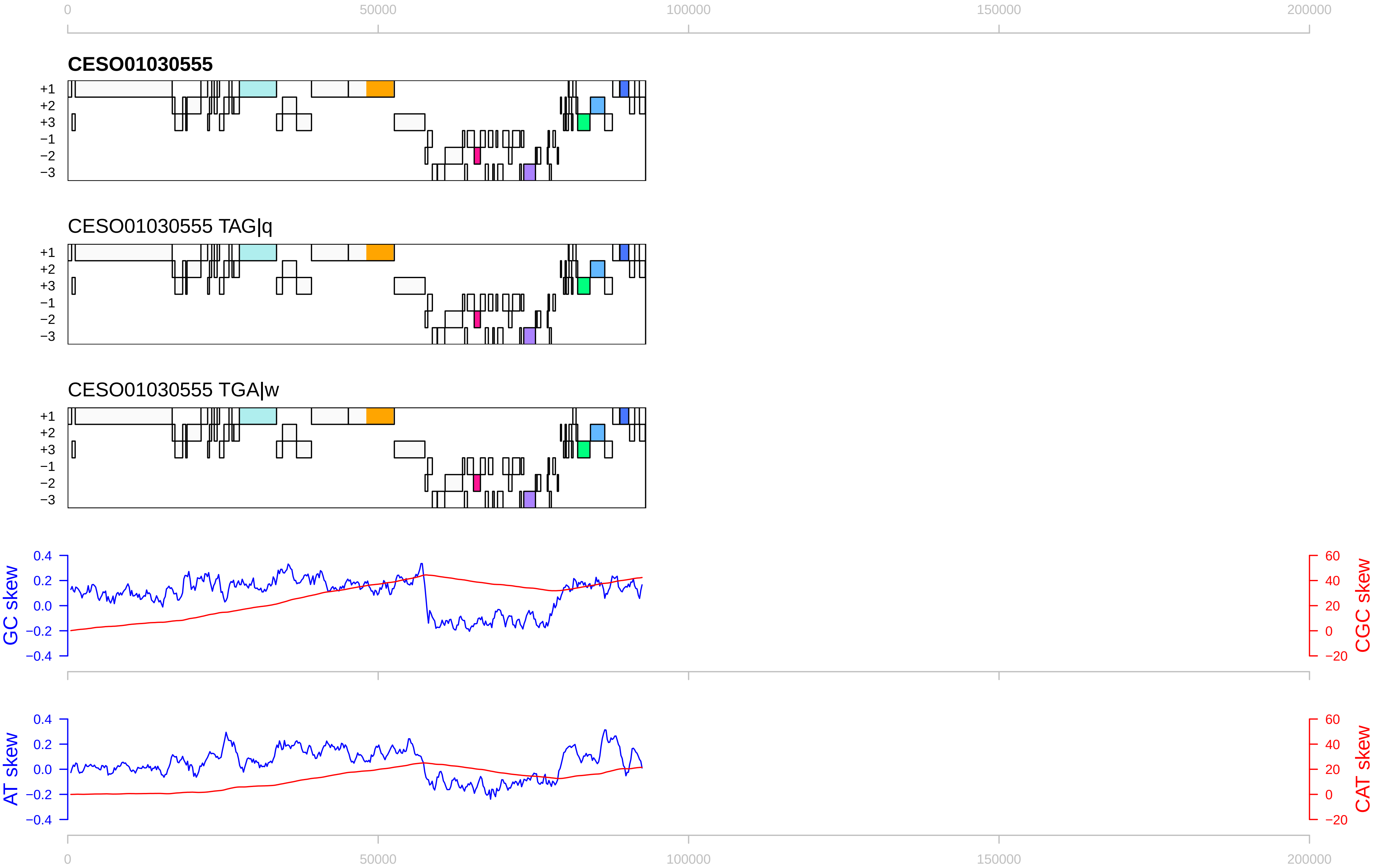

0
50000
100000
150000
200000
CESO01030555
+1
+2
+3
−1
−2
−3
CESO01030555 TAG|q
CESO01030555 TGA|w
GC skew
0.4
0.2
0.0
−0.2
−0.4
CGC skew
60
40
20
0
−20
AT skew
CAT skew

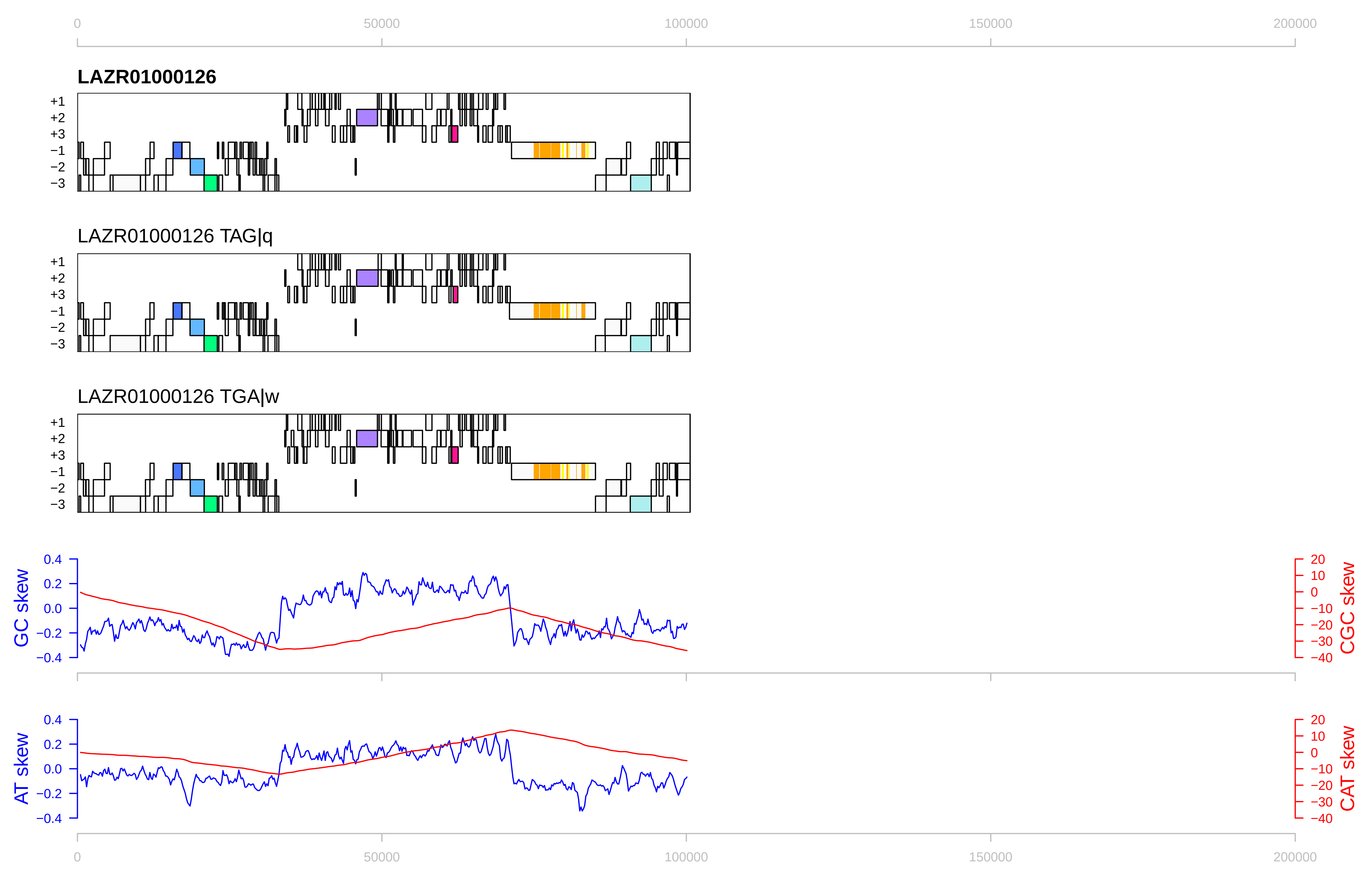

0
50000
100000
150000
200000
LAZR01000126
LAZR01000126 TAG|q
LAZR01000126 TGA|w
+1
+2
+3
−1
−2
−3
GC skew
CGC skew
AT skew
CAT skew
0.4
0.2
0.0
−0.2
−0.4
20
10
0
−10
−20
−30
−40

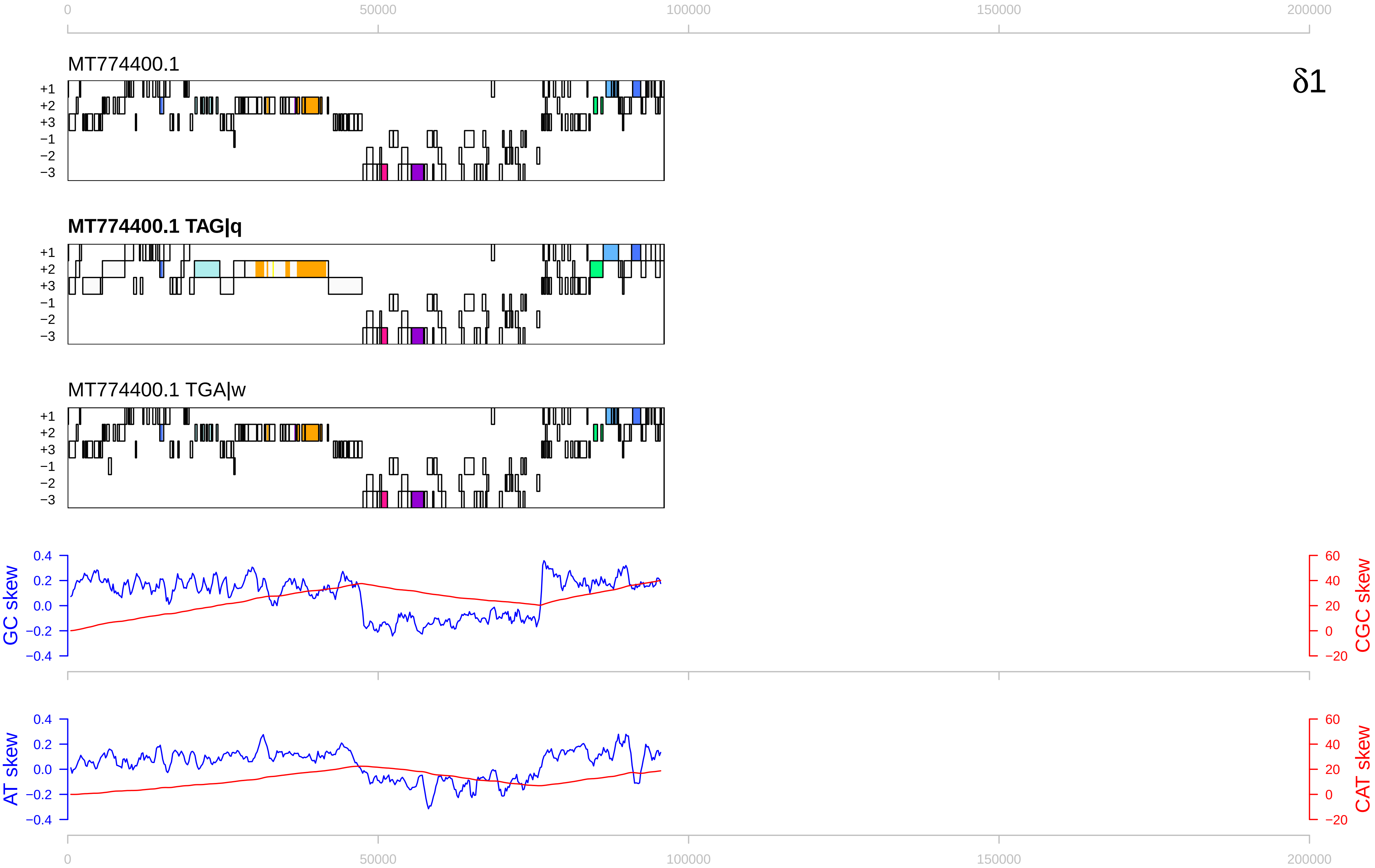

0
50000
100000
150000
200000
MT774400.1
δ1
+1
+2
+3
−1
−2
−3
MT774400.1 TAG|q
MT774400.1 TGA|w
GC skew
CGC skew
AT skew
CAT skew
0.4
0.2
0.0
−0.2
−0.4
60
40
20
0
−20

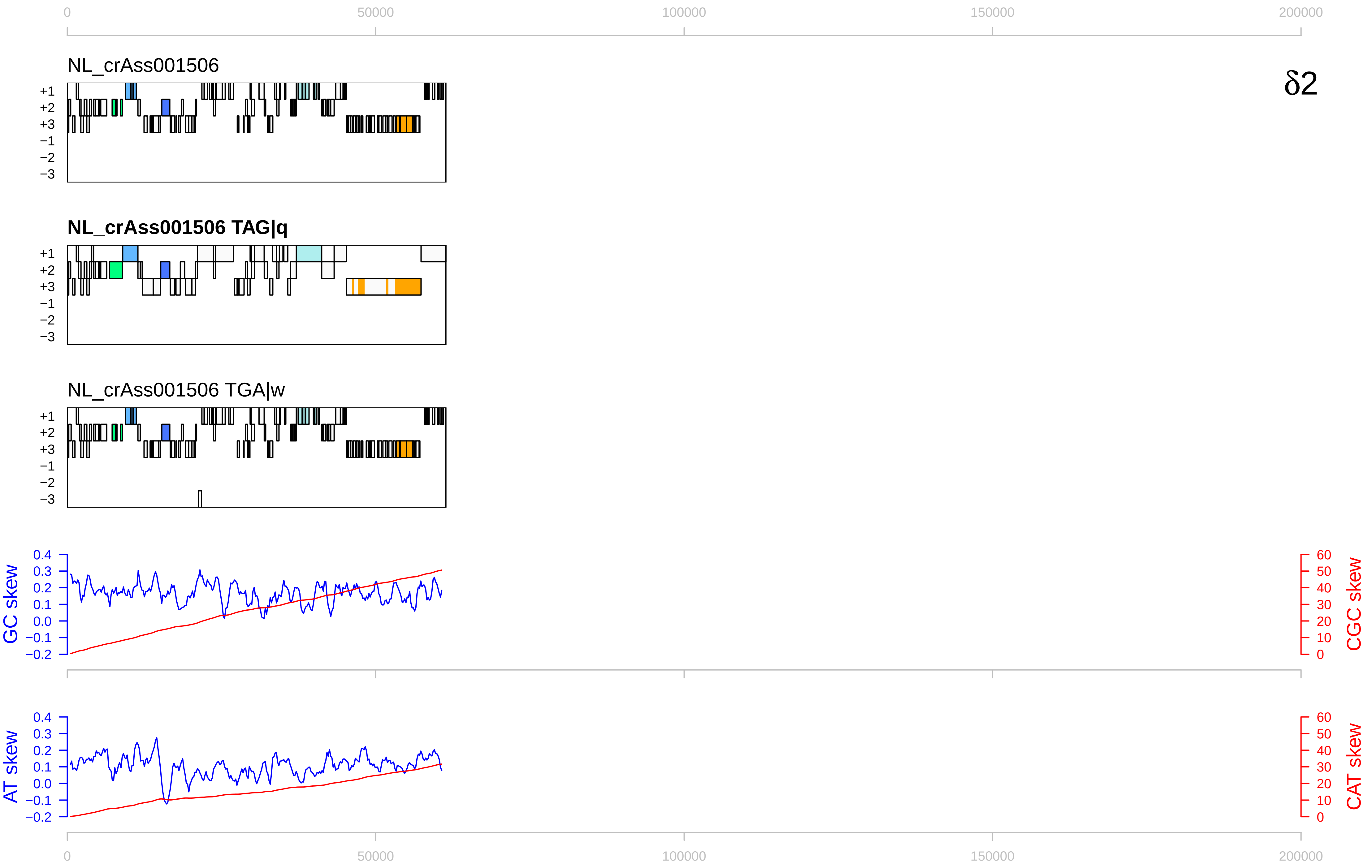

δ2
NL_crAss001506
NL_crAss001506 TAG|q
NL_crAss001506 TGA|w
+1
+2
+3
−1
−2
−3
GC skew
CGC skew
AT skew
CAT skew
0
50000
100000
150000
200000

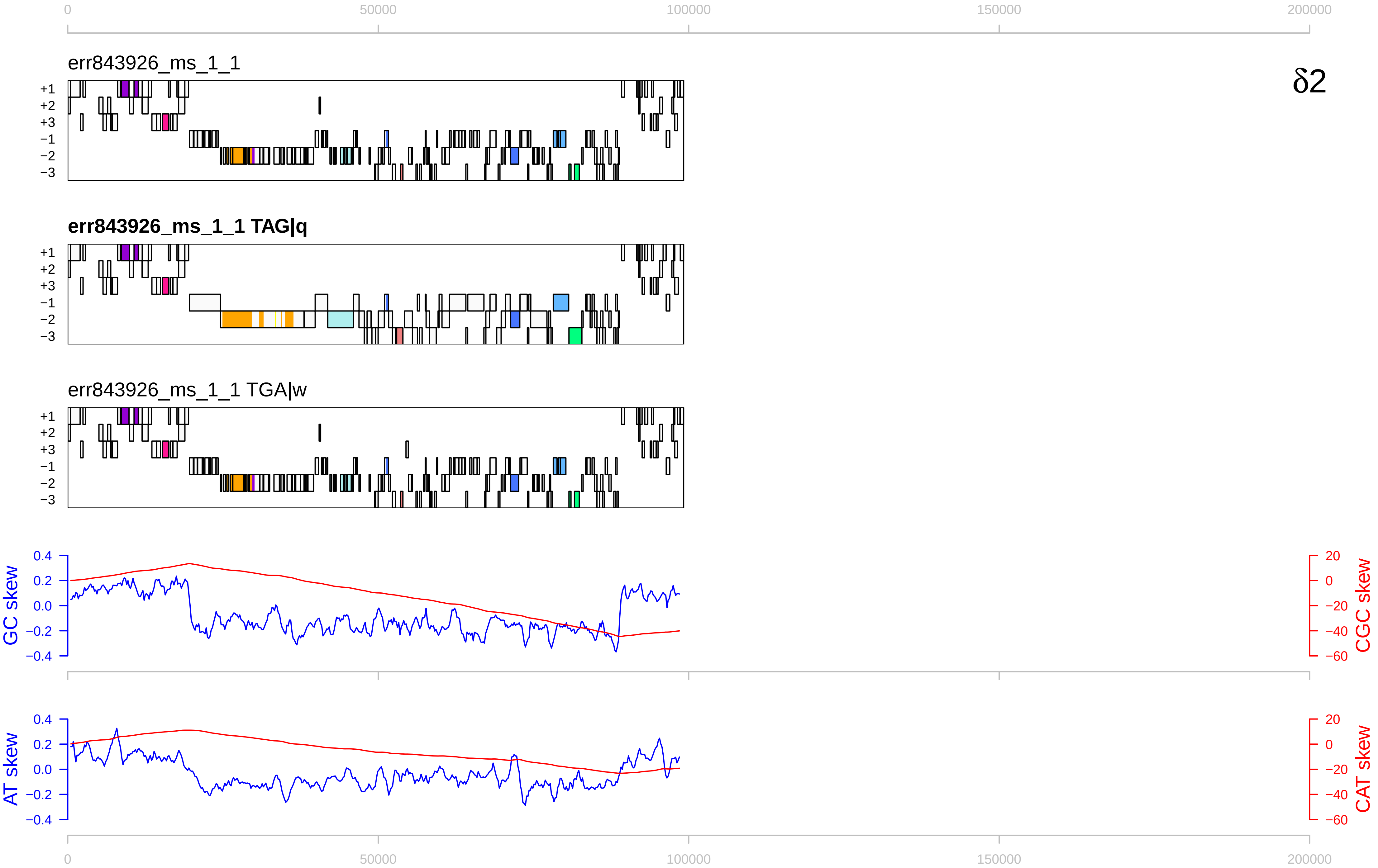

0
50000
100000
150000
200000
err843926_ms_1_1
δ2
+1
+2
+3
−1
−2
−3
err843926_ms_1_1 TAG|q
err843926_ms_1_1 TGA|w
GC skew
CGC skew
AT skew
CAT skew
0.4
0.2
0.0
−0.2
−0.4
20
0
−20
−40
−60

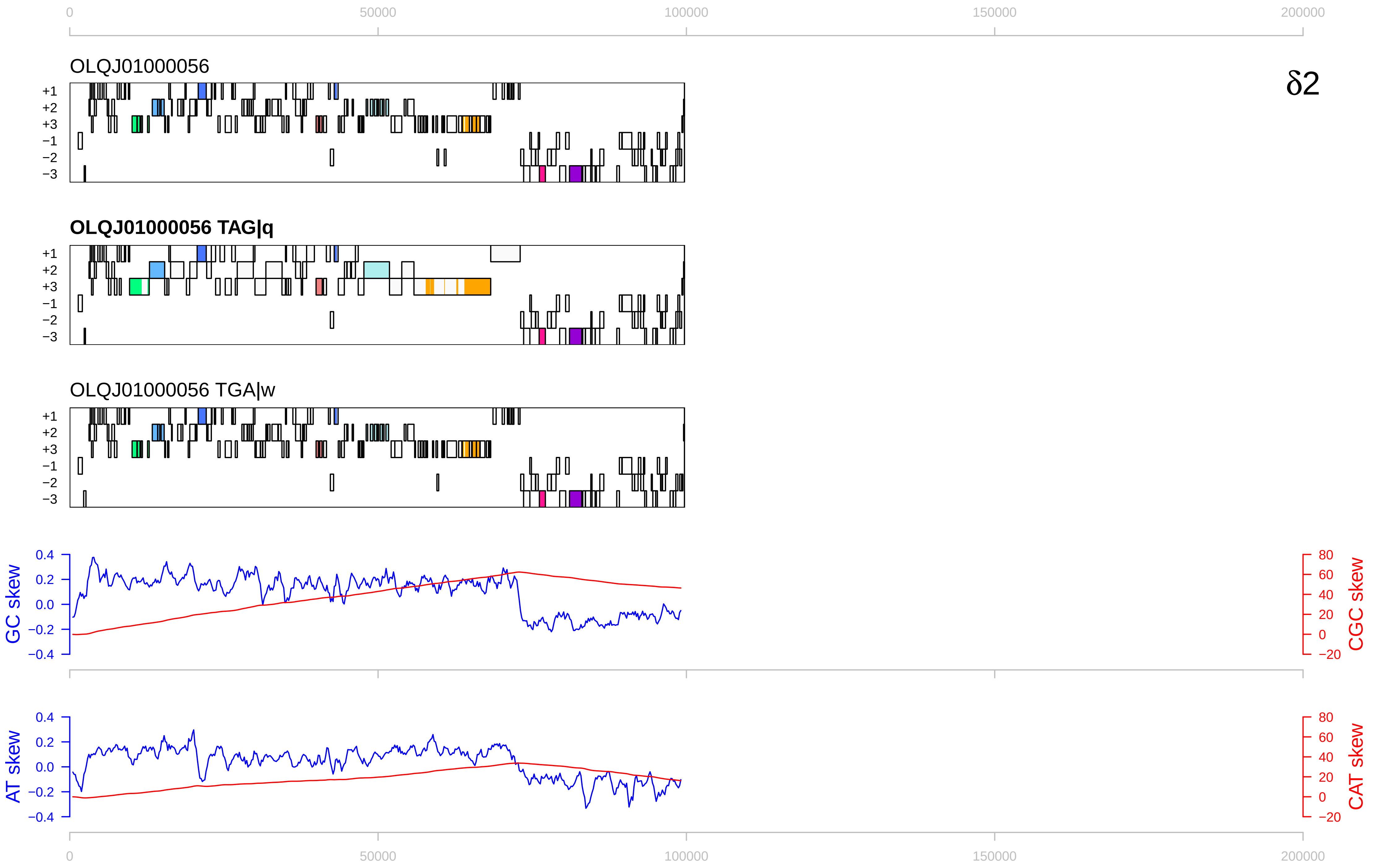

δ2
OLQJ01000056
OLQJ01000056 TAG|q
OLQJ01000056 TGA|w
+1
+2
+3
−1
−2
−3
GC skew
CGC skew
AT skew
CAT skew
0
50000
100000
150000
200000
0.4
0.2
0.0
−0.2
−0.4
80
60
40
20
0
−20

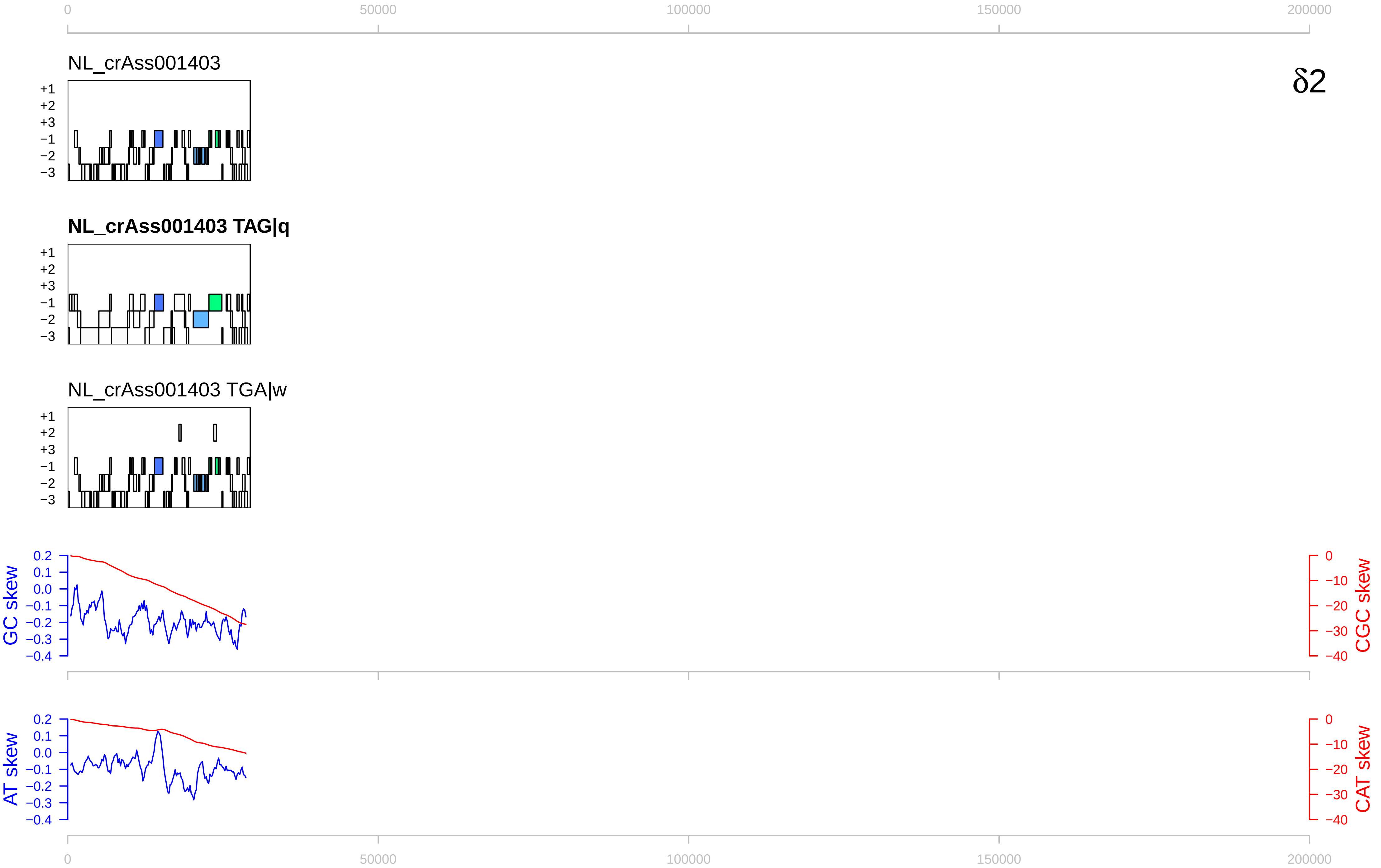

0
50000
100000
150000
200000
NL_crAss001403
δ2
+1
+2
+3
−1
−2
−3
NL_crAss001403 TAG|q
NL_crAss001403 TGA|w
GC skew
CGC skew
AT skew
CAT skew
0.2
0.1
0.0
−0.1
−0.2
−0.3
−0.4
0
−10
−20
−30
−40

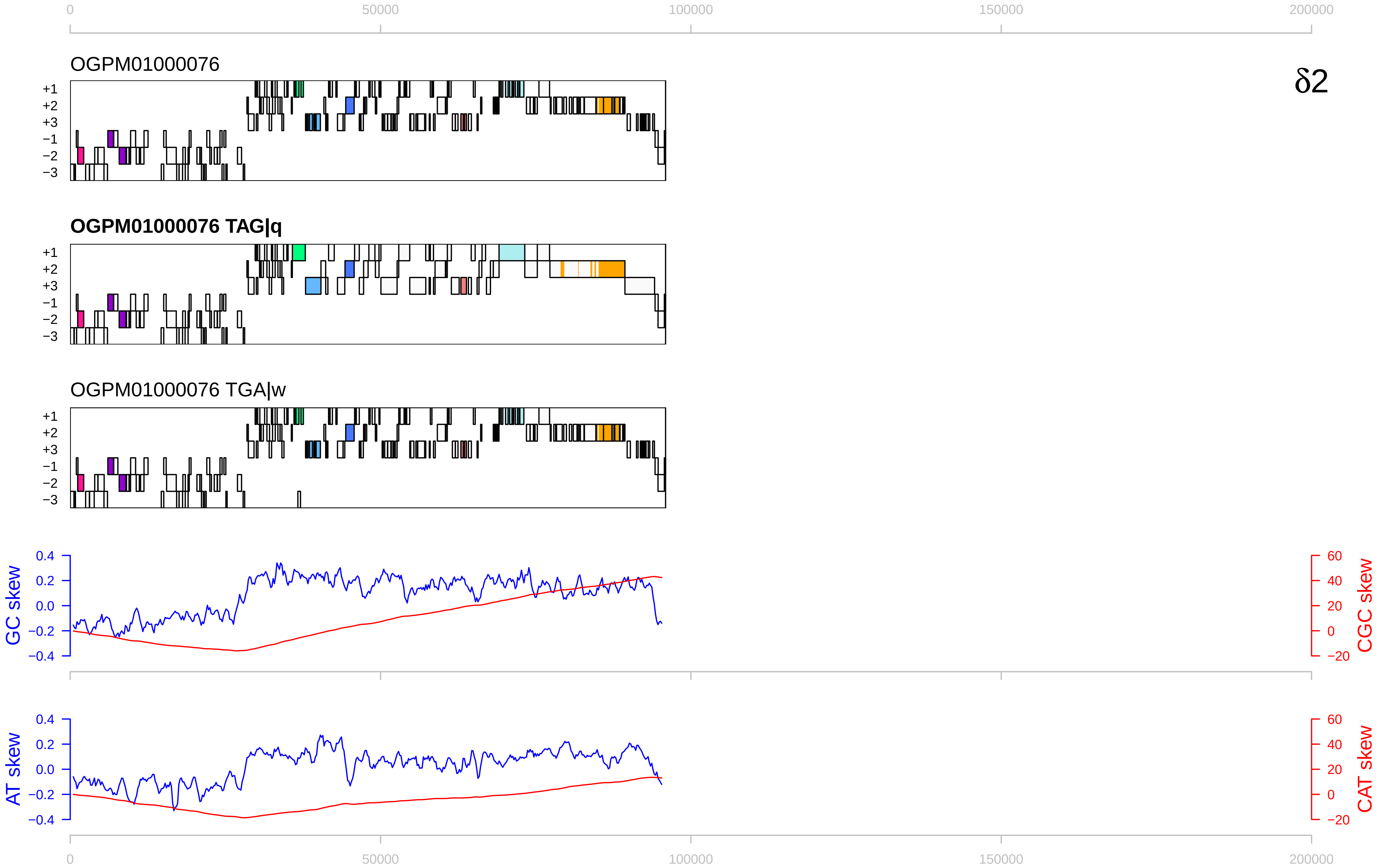

0
50000
100000
150000
200000
OGPM01000076
δ2
+1
+2
+3
−1
−2
−3
OGPM01000076 TAG|q
OGPM01000076 TGA|w
GC skew
CGC skew
AT skew
CAT skew
0.4
0.2
0.0
−0.2
−0.4
60
40
20
0
−20

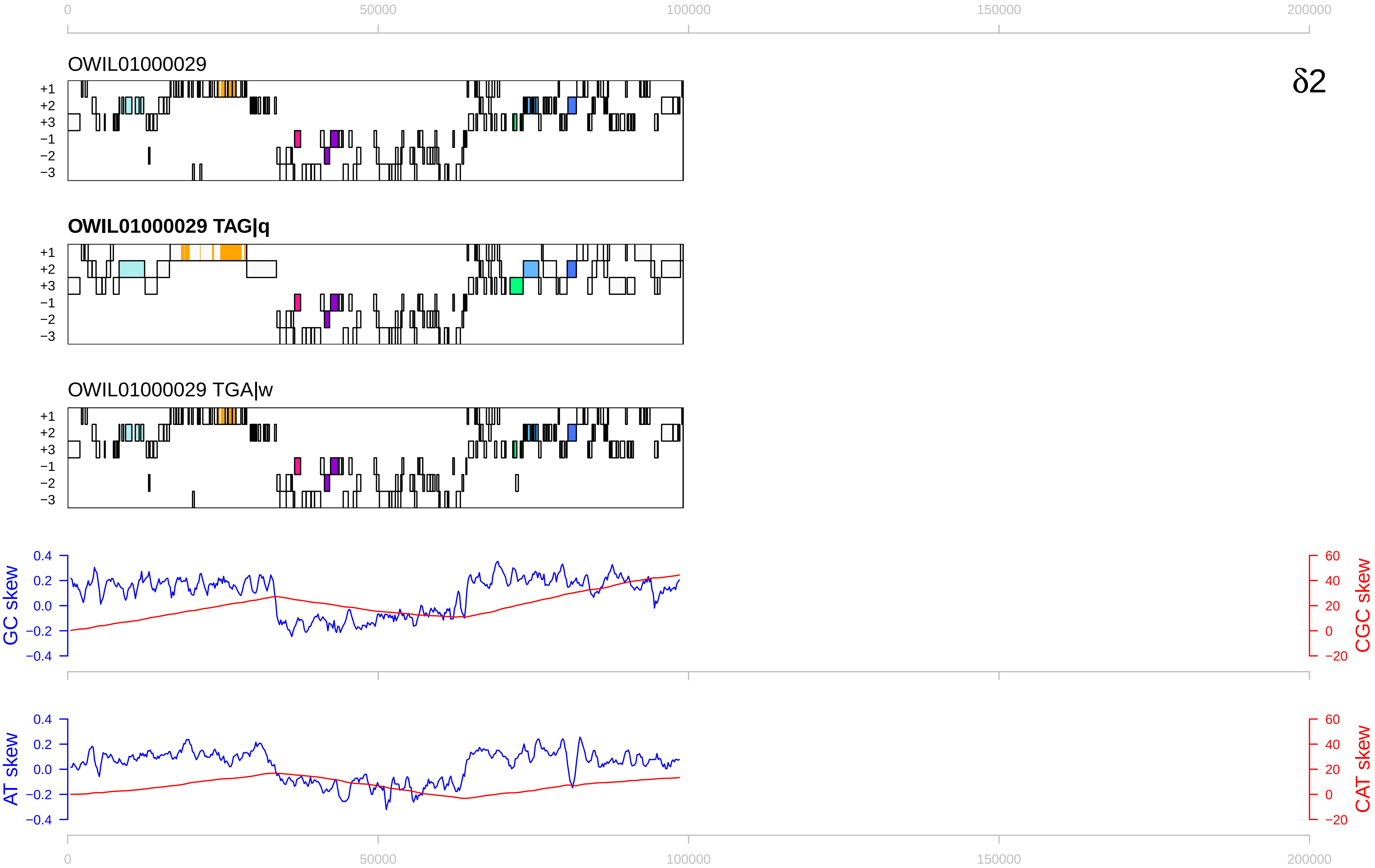

0
50000
100000
150000
200000
OWIL01000029
δ2
+1
+2
+3
−1
−2
−3
OWIL01000029 TAG|q
OWIL01000029 TGA|w
GC skew
CGC skew
0.4
0.2
0.0
−0.2
−0.4
60
40
20
0
−20
AT skew
CAT skew

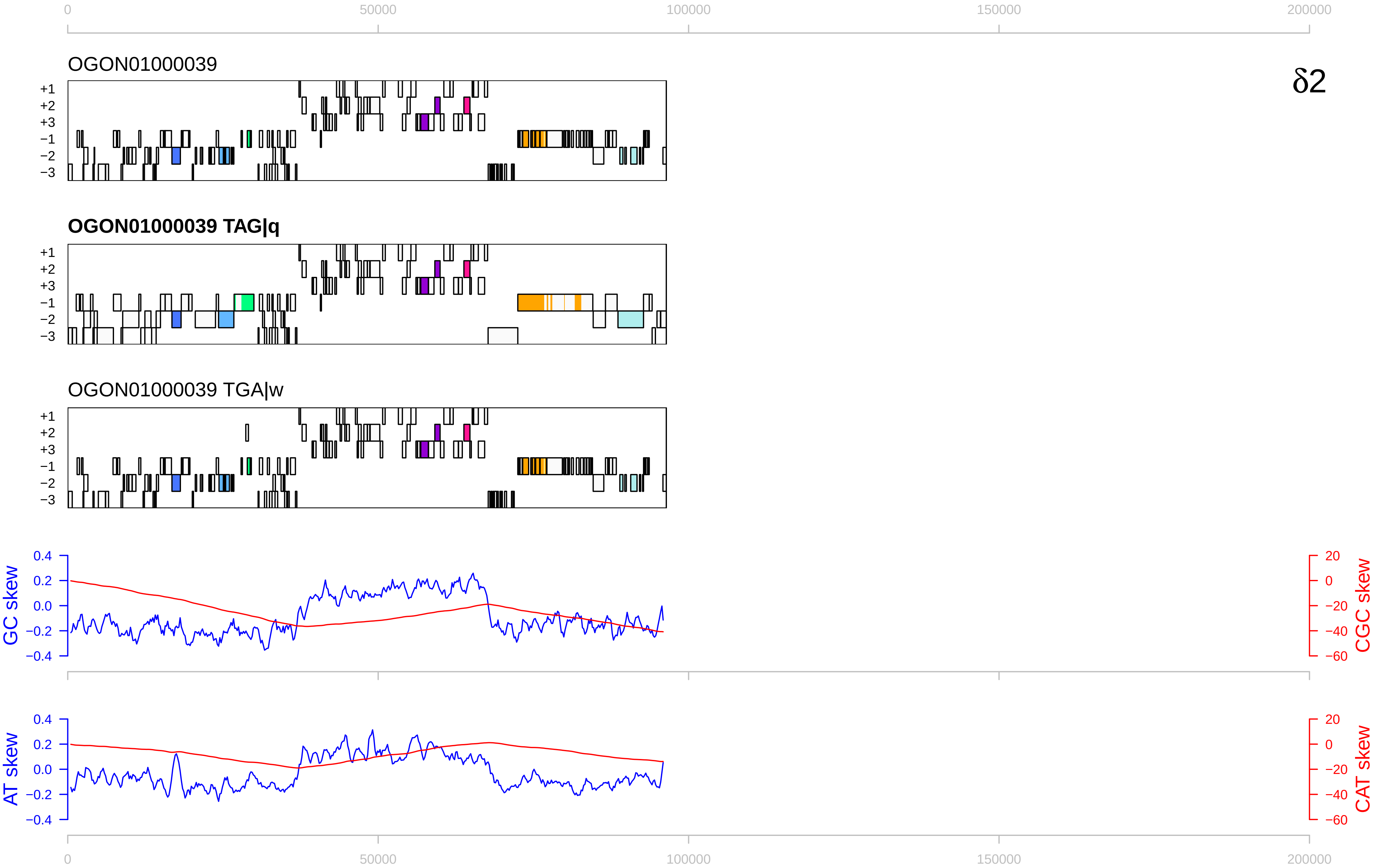

0
50000
100000
150000
200000
OGON01000039
δ2
+1
+2
+3
−1
−2
−3
OGON01000039 TAG|q
OGON01000039 TGA|w
GC skew
CGC skew
0.4
0.2
0.0
−0.2
−0.4
20
0
−20
−40
−60
AT skew
CAT skew

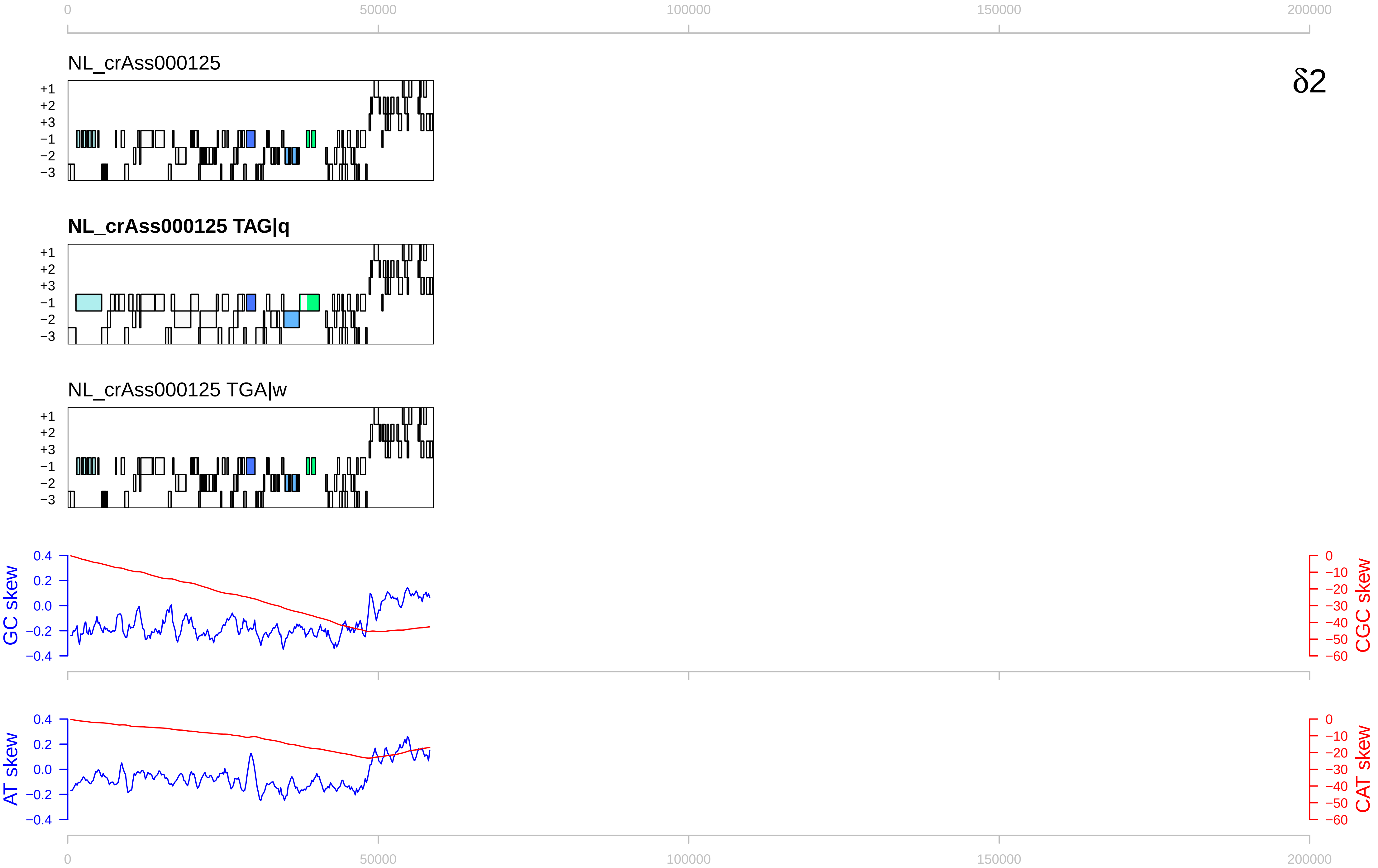

0
50000
100000
150000
200000
NL_crAss000125
δ2
+1
+2
+3
−1
−2
−3
NL_crAss000125 TAG|q
NL_crAss000125 TGA|w
GC skew
CGC skew
AT skew
CAT skew
0.4
0.2
0.0
−0.2
−0.4
0
−10
−20
−30
−40
−50
−60

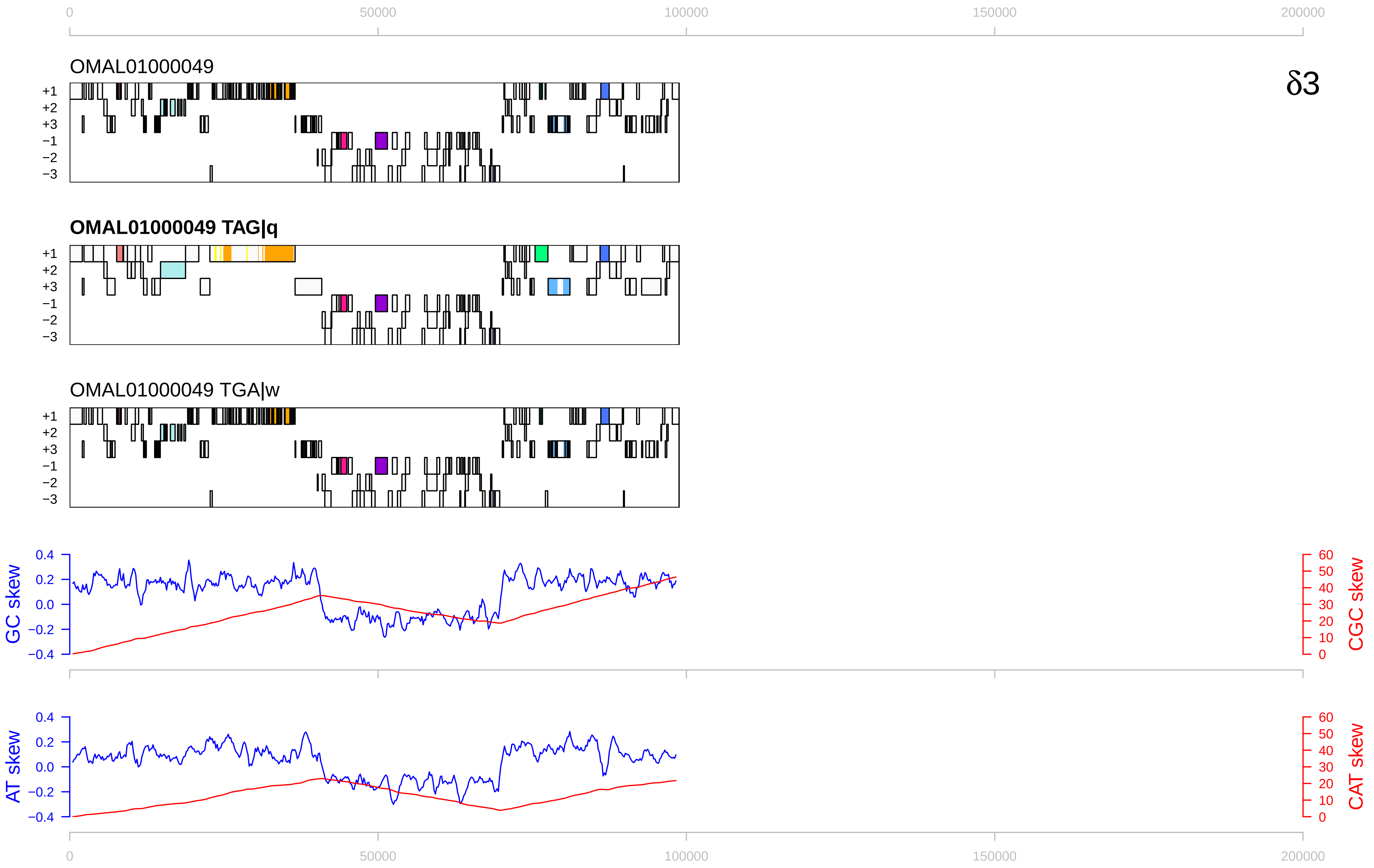

0
50000
100000
150000
200000
OMAL01000049
δ3
+1
+2
+3
−1
−2
−3
OMAL01000049 TAG|q
OMAL01000049 TGA|w
GC skew
CGC skew
AT skew
CAT skew
0.4
0.2
0.0
−0.2
−0.4
60
50
40
30
20
10
0

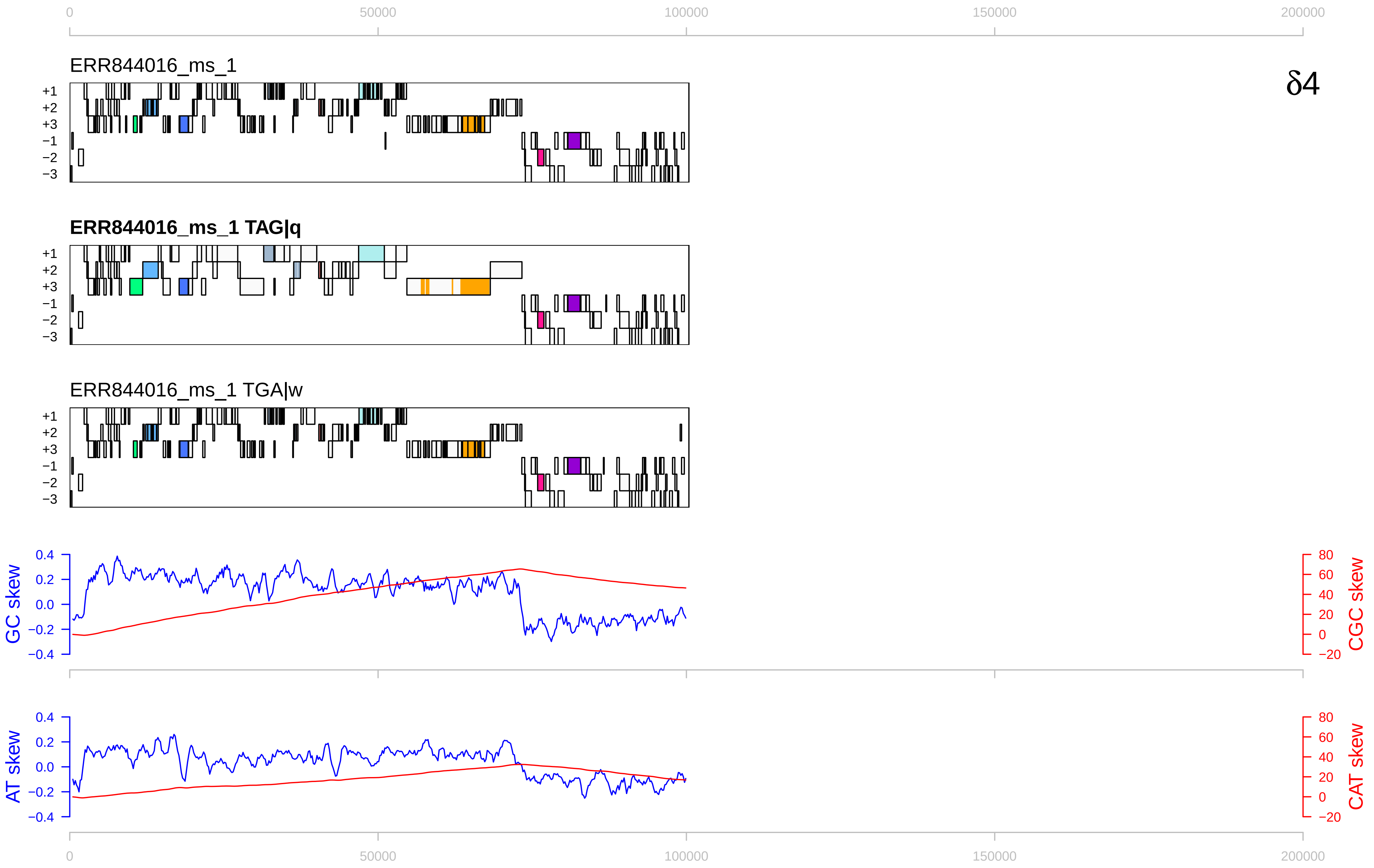

0
50000
100000
150000
200000
δ4
ERR844016_ms_1
+1
+2
+3
−1
−2
−3
ERR844016_ms_1 TAG|q
ERR844016_ms_1 TGA|w
GC skew
CGC skew
AT skew
CAT skew
0.4
0.2
0.0
−0.2
−0.4
80
60
40
20
0
−20

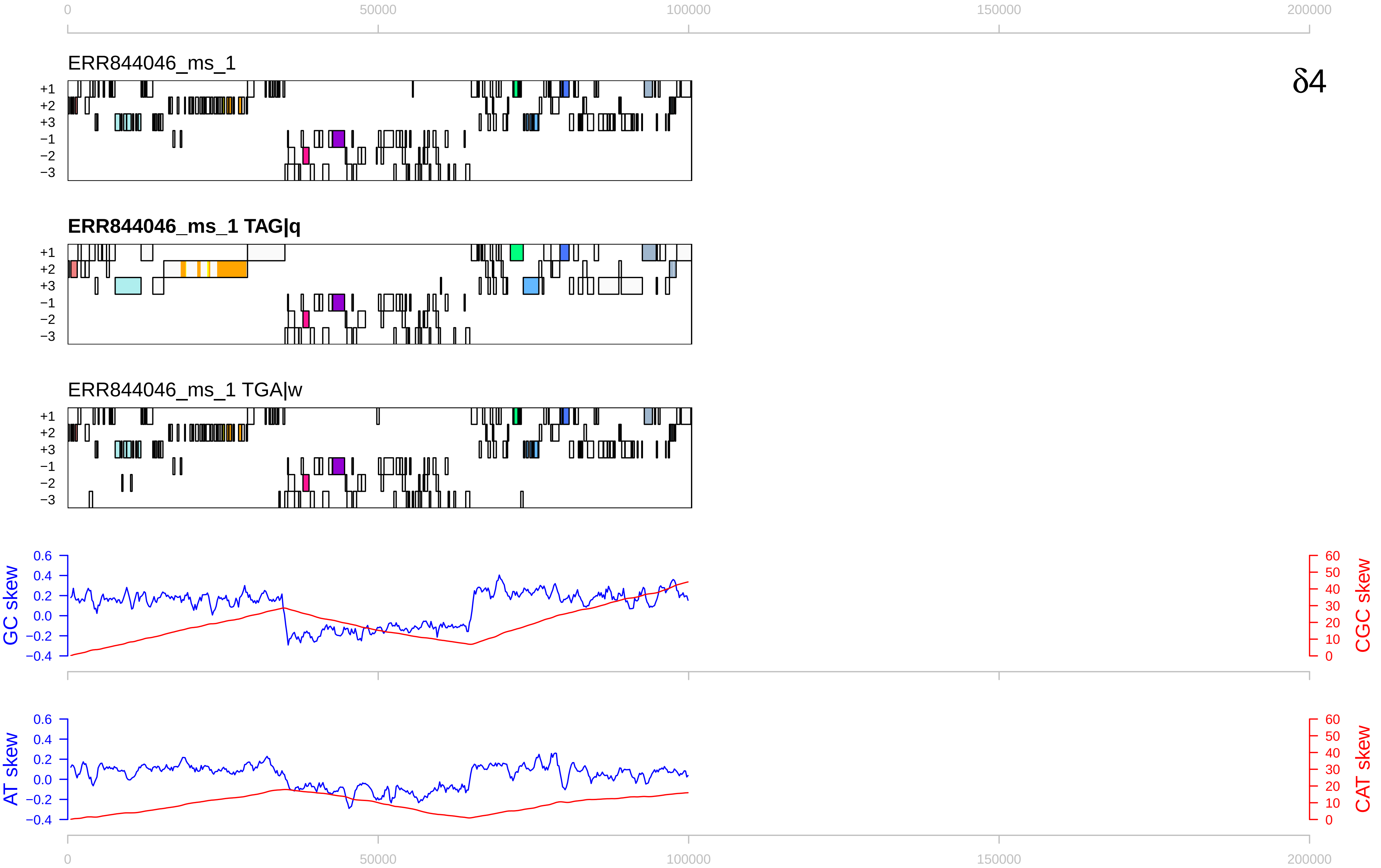
δ4
ERR844046_ms_1
ERR844046_ms_1 TAG|q
ERR844046_ms_1 TGA|w
+1
+2
+3
−1
−2
−3
GC skew
CGC skew
AT skew
CAT skew
0
50000
100000
150000
200000
0.6
0.4
0.2
0.0
−0.2
−0.4
60
50
40
30
20
10
0

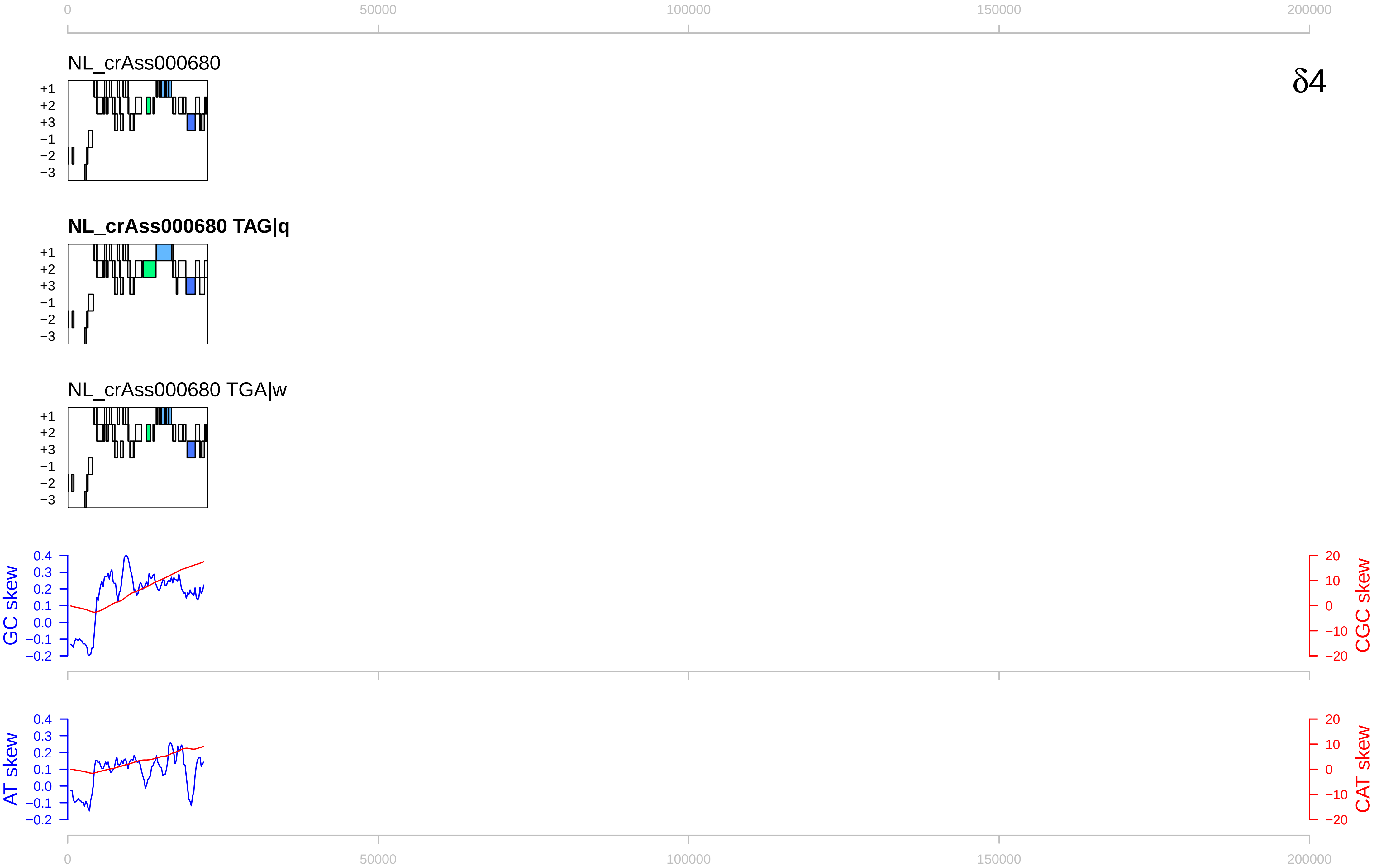

0
50000
100000
150000
200000
NL_crAss000680
δ4
+1
+2
+3
−1
−2
−3
NL_crAss000680 TAG|q
NL_crAss000680 TGA|w
GC skew
CGC skew
AT skew
CAT skew
0.4
0.3
0.2
0.1
0.0
−0.1
−0.2
20
10
0
−10
−20

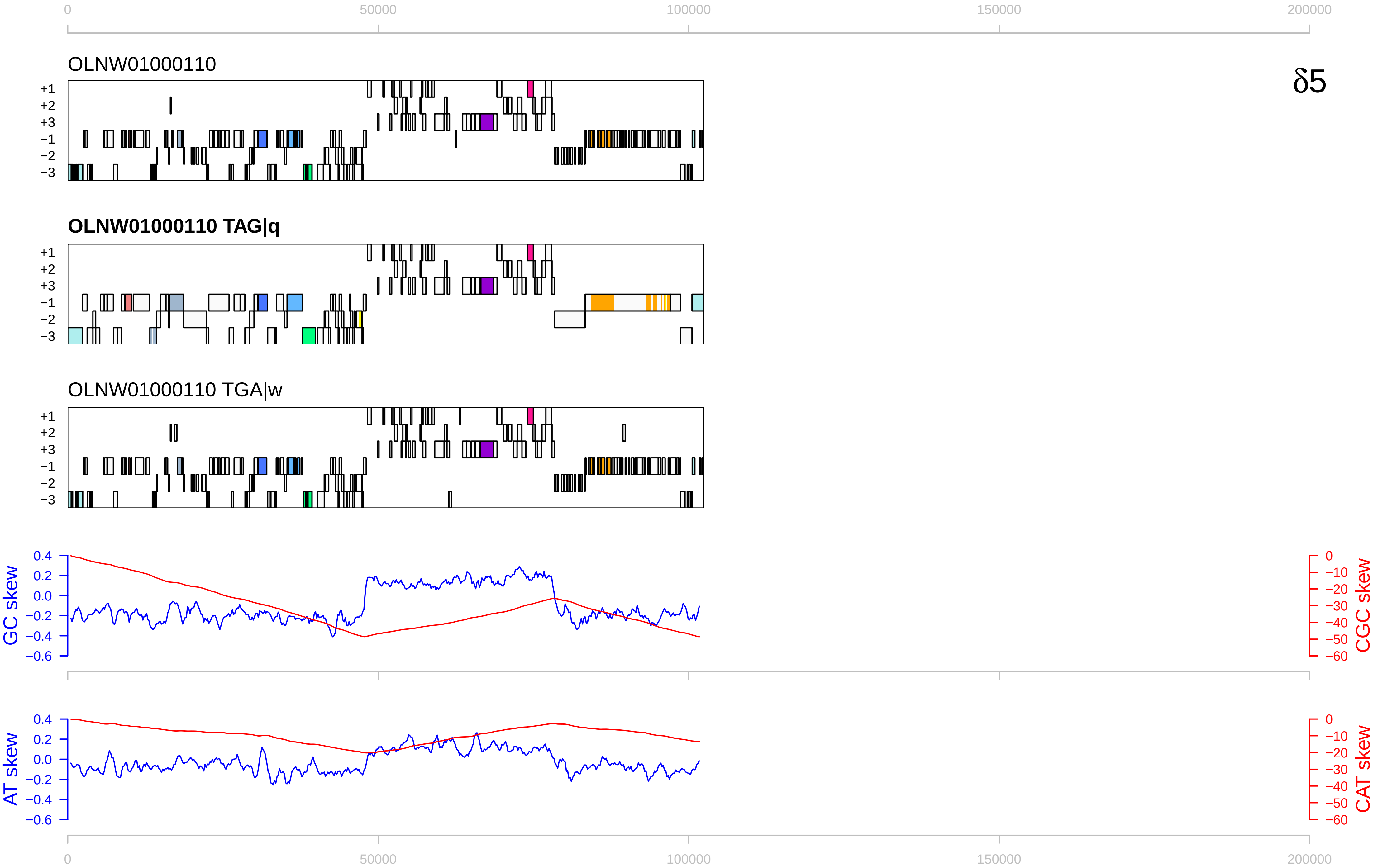

0
50000
100000
150000
200000
OLNW01000110
δ5
+1
+2
+3
−1
−2
−3
OLNW01000110 TAG|q
OLNW01000110 TGA|w
GC skew
CGC skew
0.4
0.2
0.0
−0.2
−0.4
−0.6
0
−10
−20
−30
−40
−50
−60
AT skew
CAT skew

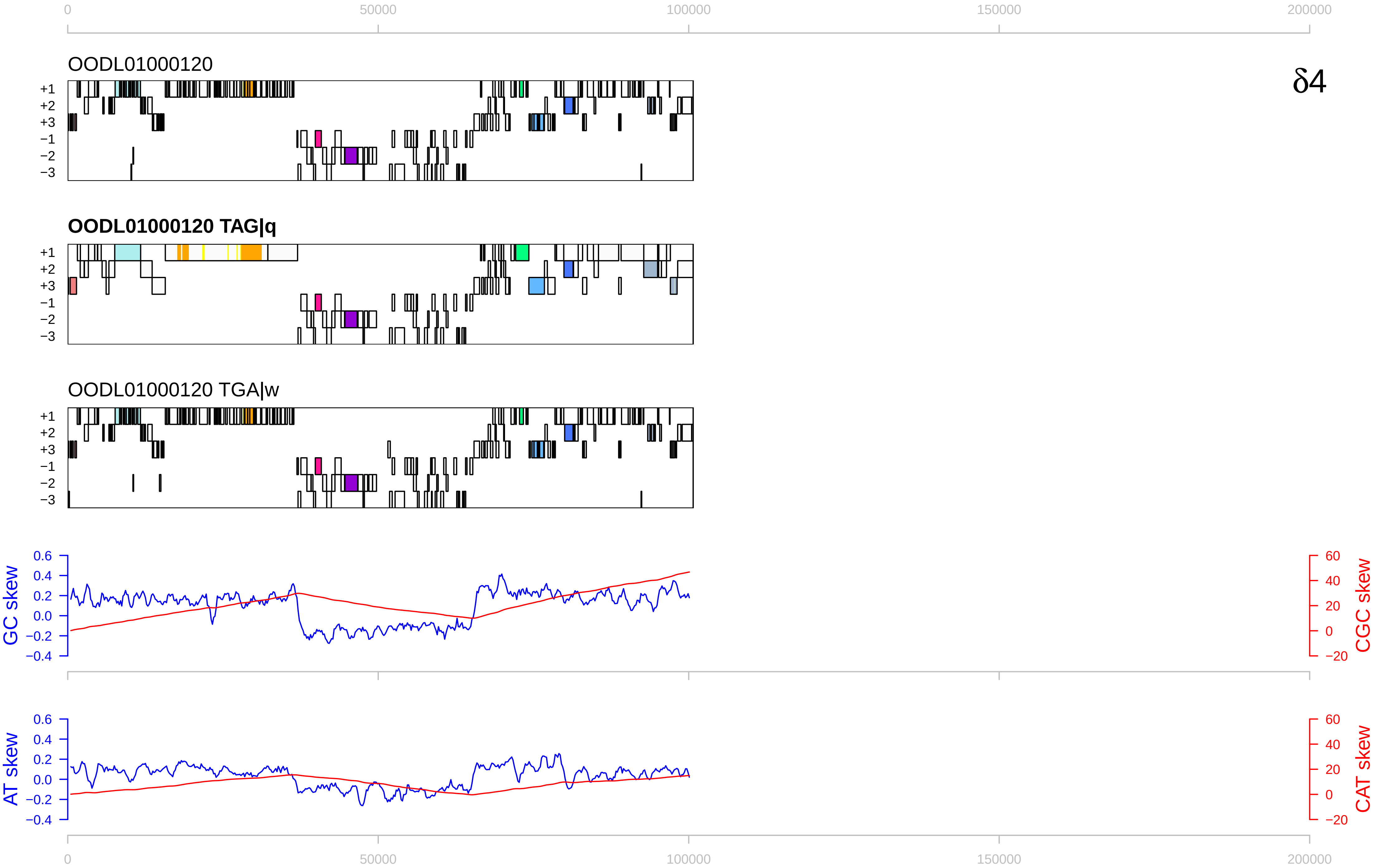

0
50000
100000
150000
200000
OODL01000120
δ4
+1
+2
+3
−1
−2
−3
OODL01000120 TAG|q
OODL01000120 TGA|w
GC skew
CGC skew
0.6
0.4
0.2
0.0
−0.2
−0.4
60
40
20
0
−20
AT skew
CAT skew

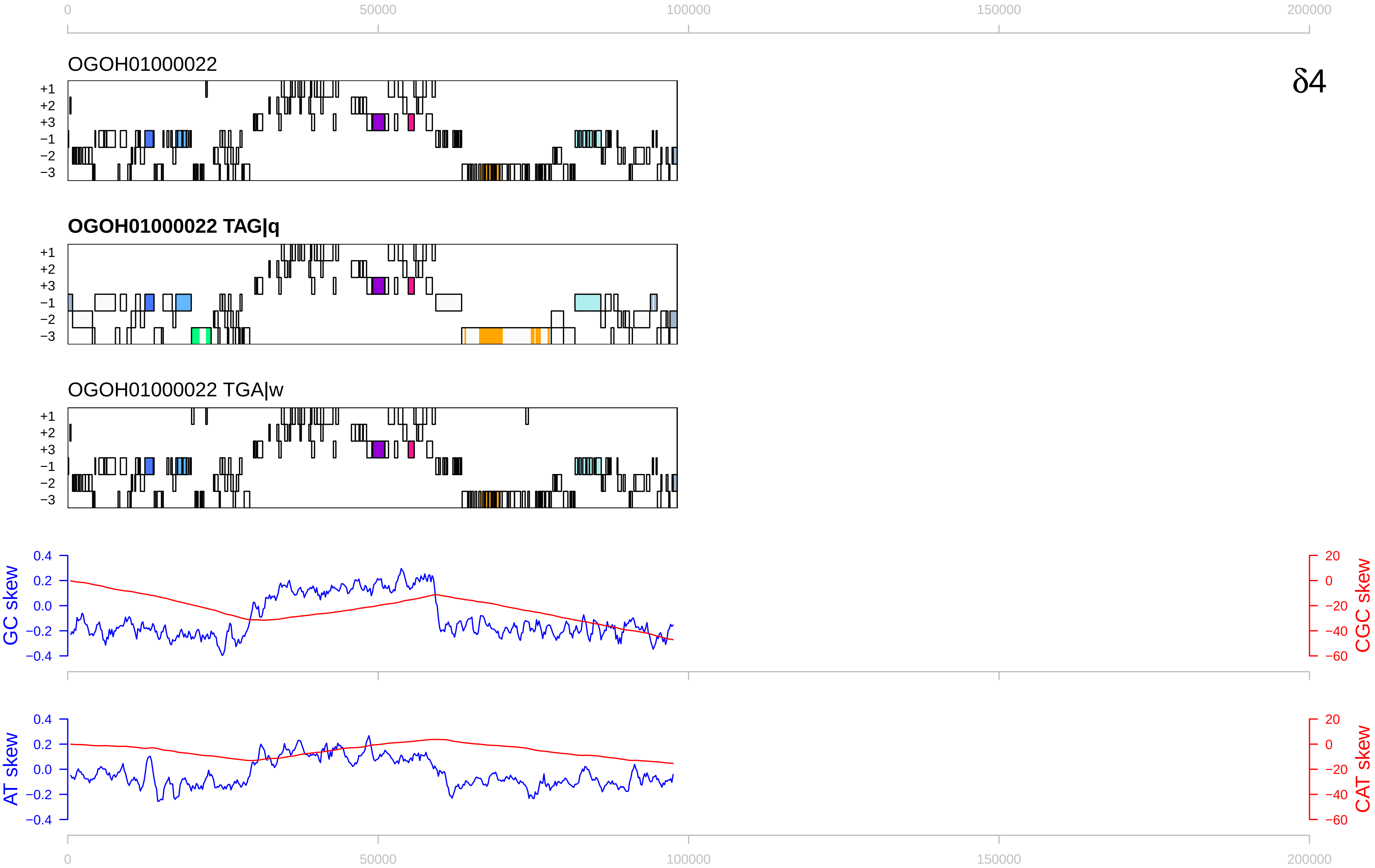

0
50000
100000
150000
200000
OGOH01000022
δ4
+1
+2
+3
−1
−2
−3
OGOH01000022 TAG|q
OGOH01000022 TGA|w
GC skew
CGC skew
AT skew
CAT skew
0.4
0.2
0.0
−0.2
−0.4
20
0
−20
−40
−60

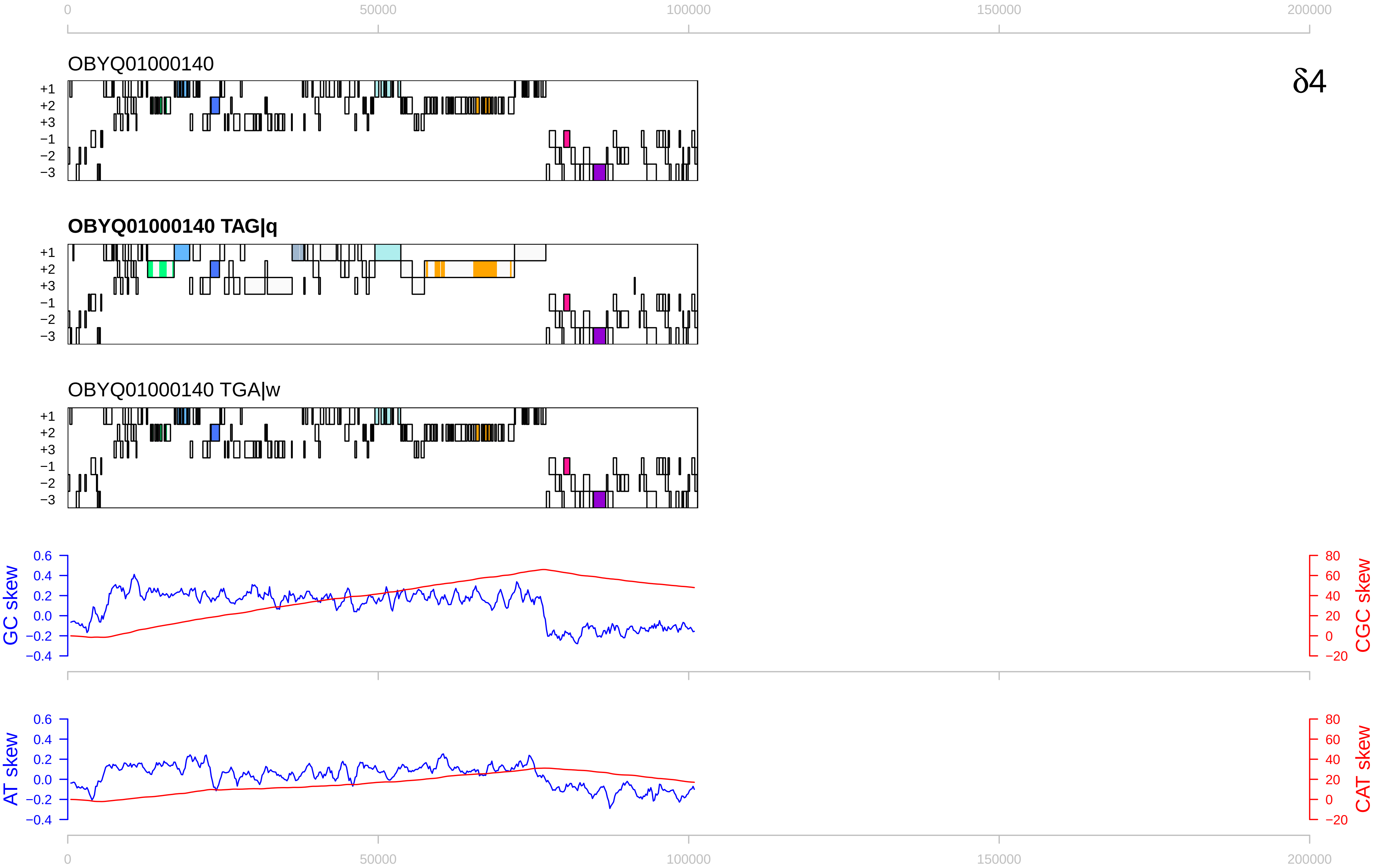

0
50000
100000
150000
200000
OBYQ01000140
δ4
+1
+2
+3
−1
−2
−3
OBYQ01000140 TAG|q
OBYQ01000140 TGA|w
GC skew
CGC skew
AT skew
CAT skew
0.6
0.4
0.2
0.0
−0.2
−0.4
80
60
40
20
0
−20

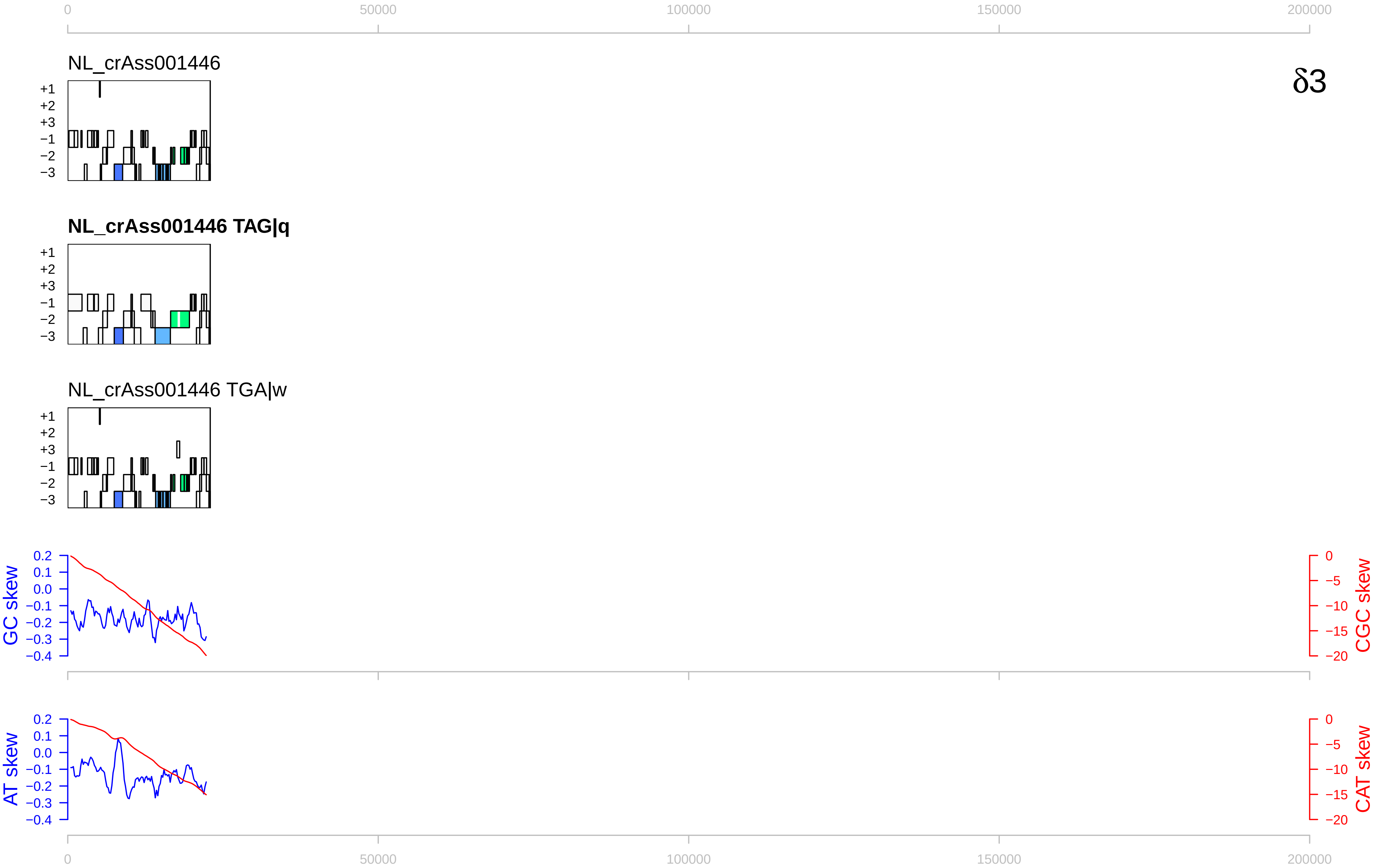
δ3
NL_crAss001446
NL_crAss001446 TAG|q
NL_crAss001446 TGA|w
+1
+2
+3
−1
−2
−3
GC skew
CGC skew
AT skew
CAT skew
0
50000
100000
150000
200000
0.2
0.1
0.0
−0.1
−0.2
−0.3
−0.4
0
−5
−10
−15
−20

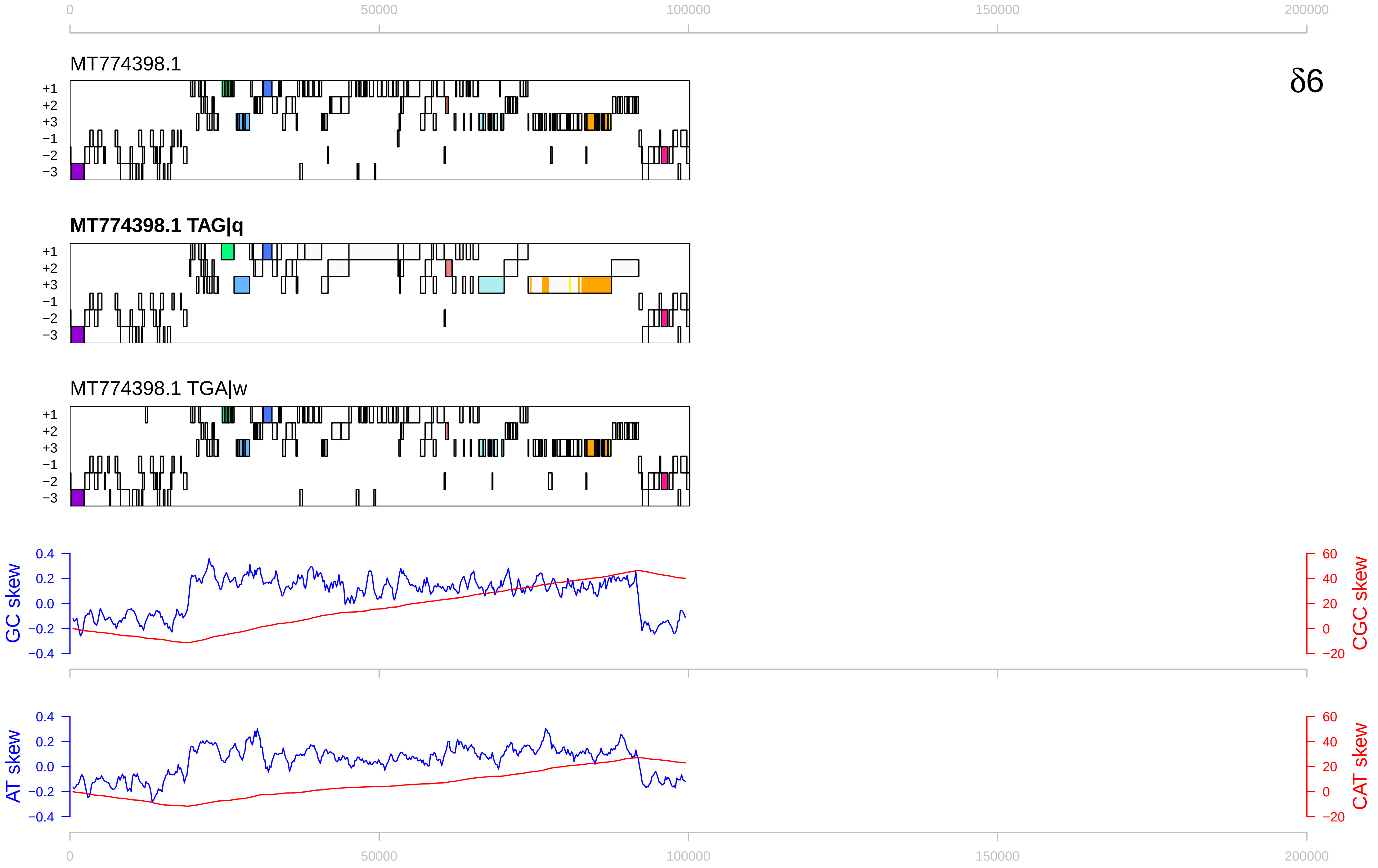

δ6
MT774398.1
MT774398.1 TAG|q
MT774398.1 TGA|w
+1
+2
+3
−1
−2
−3
GC skew
CGC skew
AT skew
CAT skew
0.4
0.2
0.0
−0.2
−0.4
60
40
20
0
−20
0
50000
100000
150000
200000

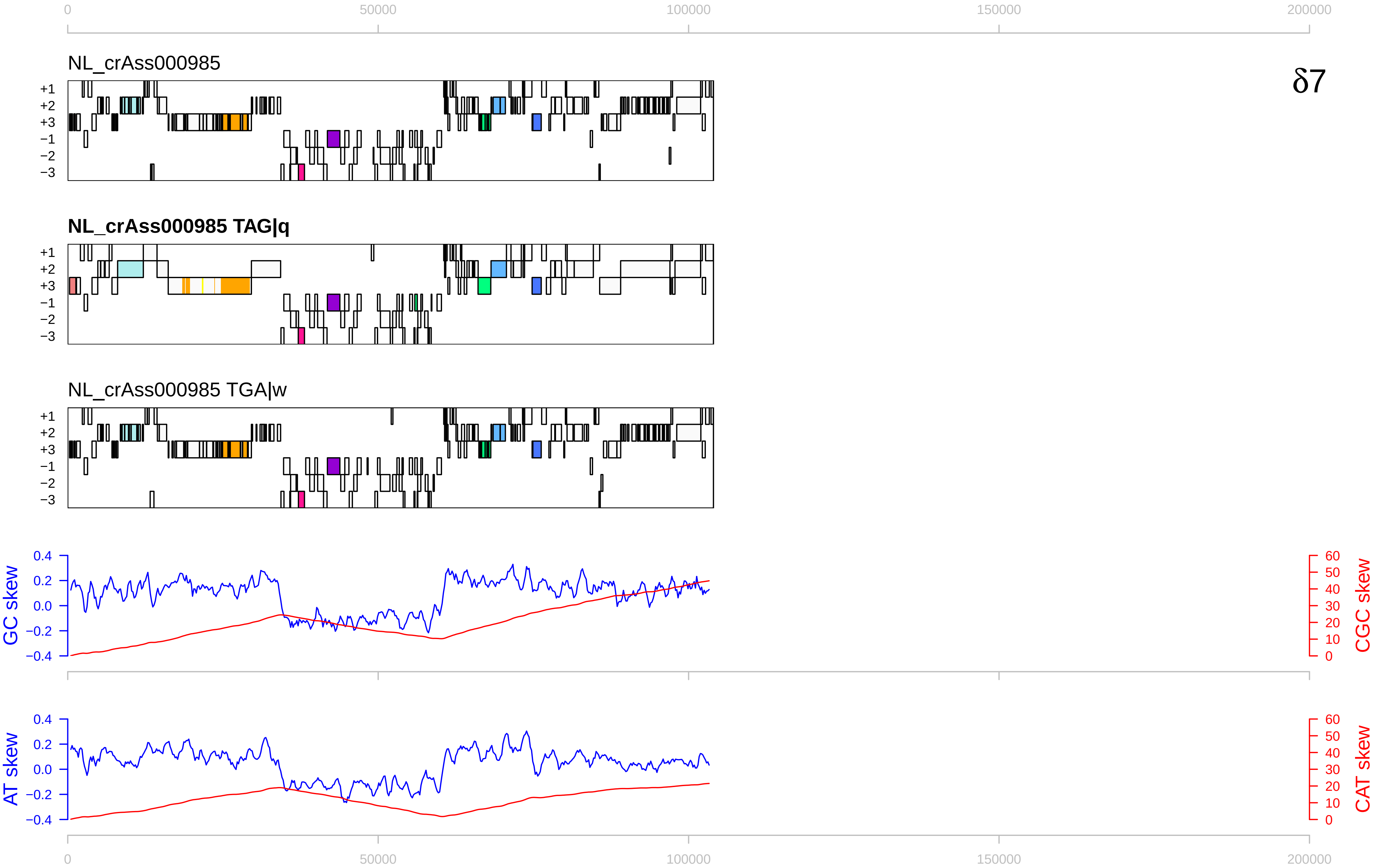
δ7
NL_crAss000985
NL_crAss000985 TAG|q
NL_crAss000985 TGA|w
+1
+2
+3
−1
−2
−3
GC skew
CGC skew
AT skew
CAT skew
0
50000
100000
150000
200000
0.4
0.2
0.0
−0.2
−0.4
60
50
40
30
20
10
0

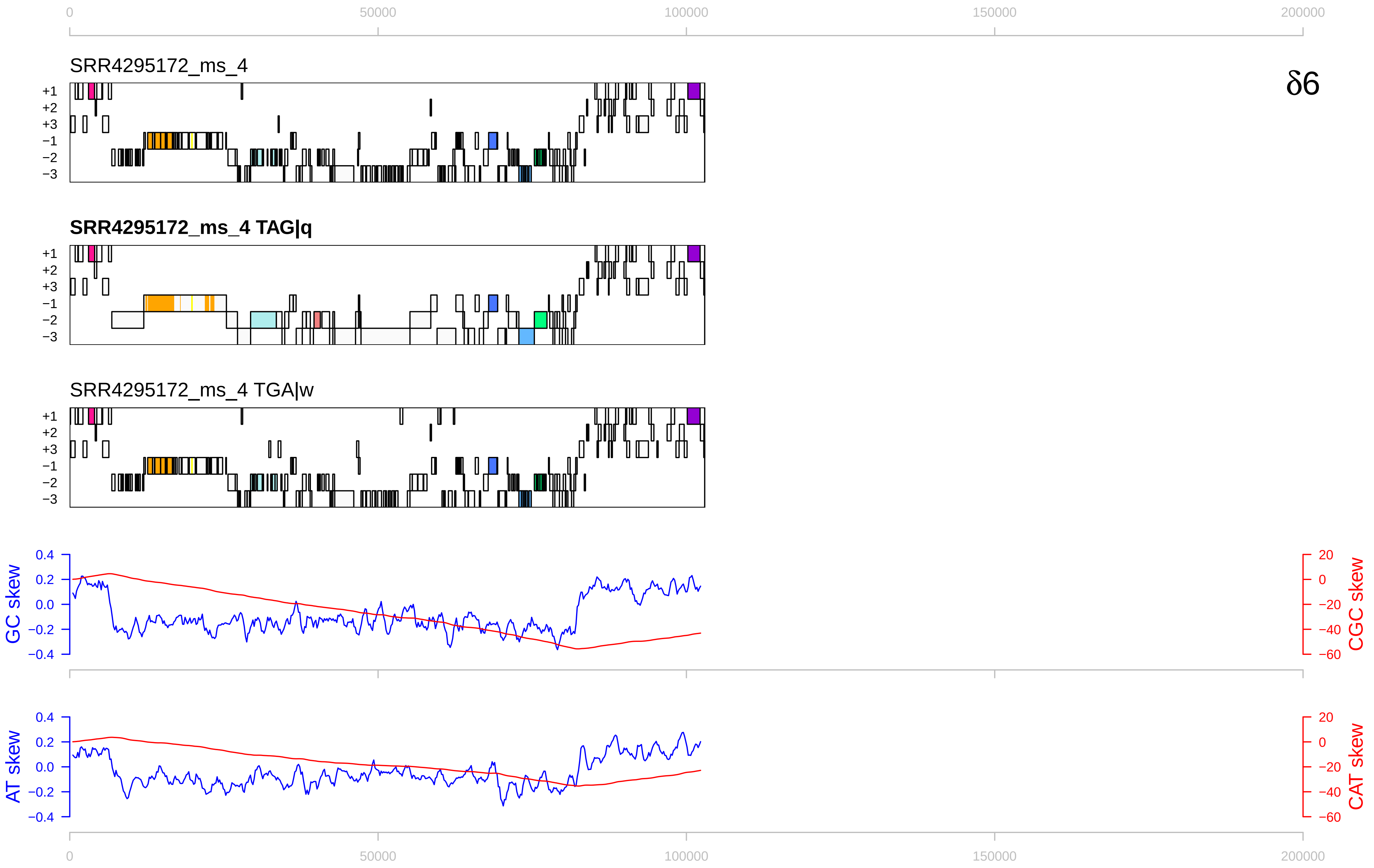

0
50000
100000
150000
200000
SRR4295172_ms_4
δ6
SRR4295172_ms_4 TAG|q
SRR4295172_ms_4 TGA|w
+1
+2
+3
−1
−2
−3
GC skew
CGC skew
AT skew
CAT skew
0.4
0.2
0.0
−0.2
−0.4
20
0
−20
−40
−60

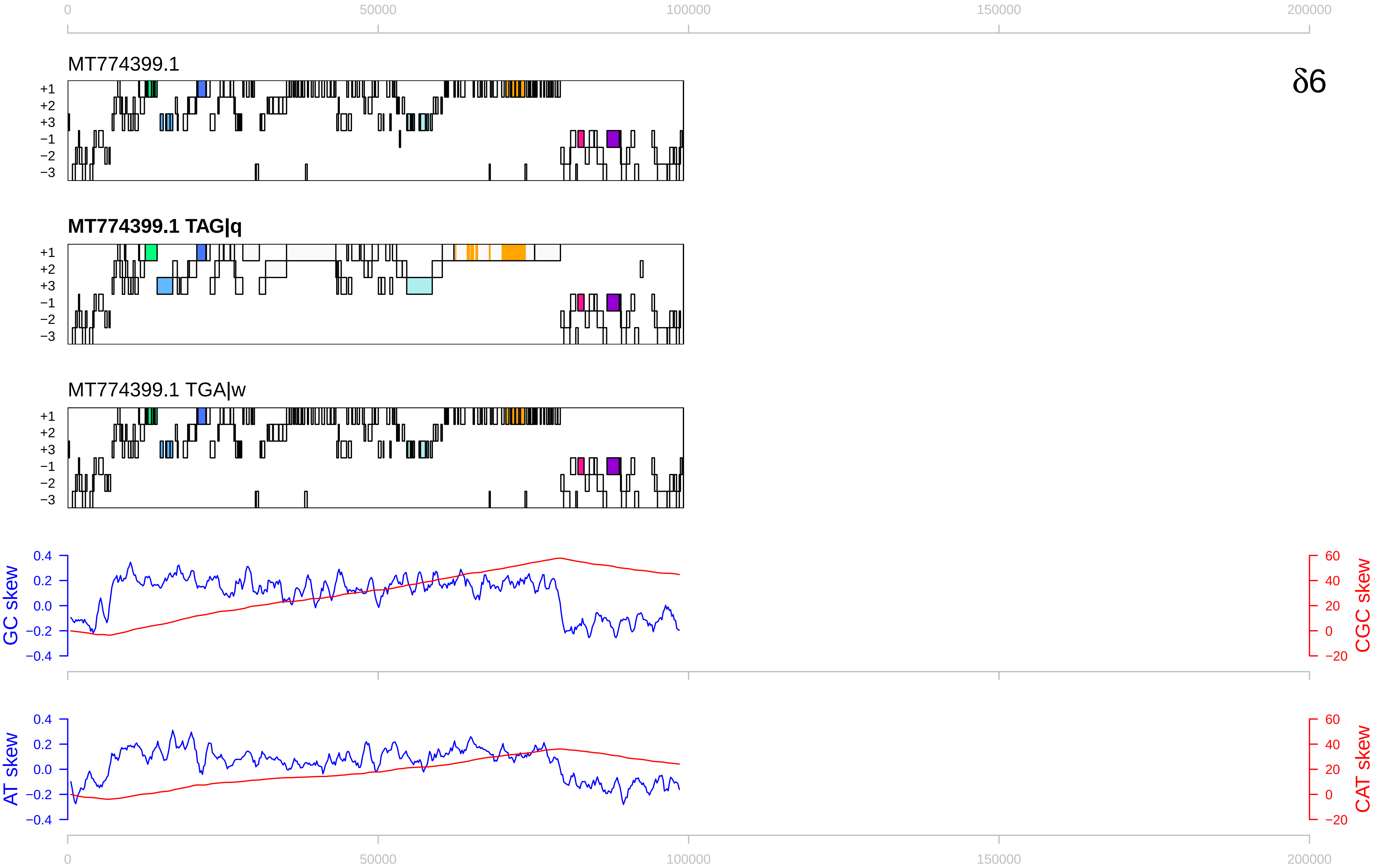

0
50000
100000
150000
200000
MT774399.1
δ6
+1
+2
+3
−1
−2
−3
MT774399.1 TAG|q
MT774399.1 TGA|w
GC skew
CGC skew
AT skew
CAT skew
0.4
0.2
0.0
−0.2
−0.4
60
40
20
0
−20

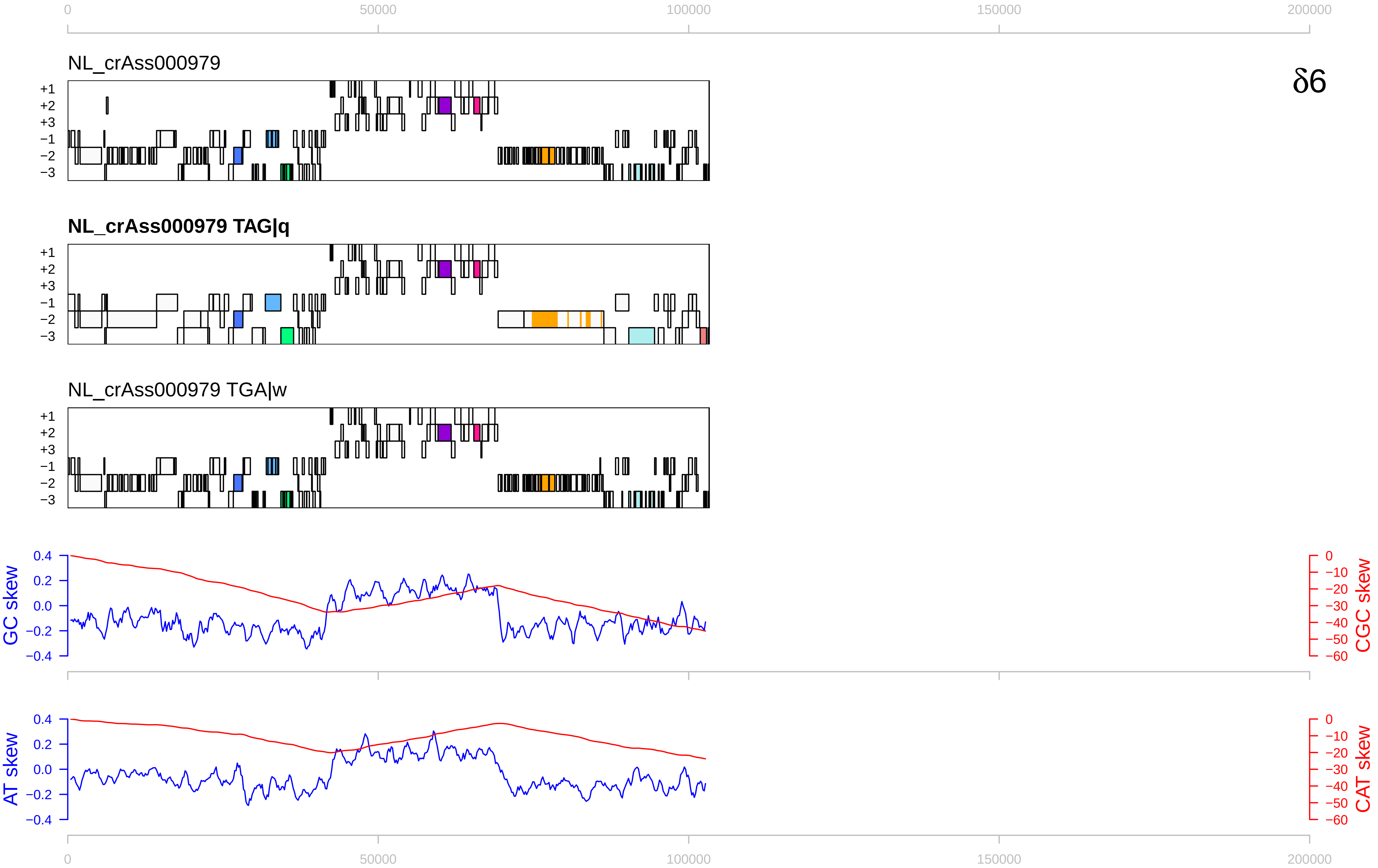

0
50000
100000
150000
200000
NL_crAss000979
δ6
+1
+2
+3
−1
−2
−3
NL_crAss000979 TAG|q
NL_crAss000979 TGA|w
GC skew
0.4
0.2
0.0
−0.2
−0.4
CGC skew
0
−10
−20
−30
−40
−50
−60
AT skew
CAT skew

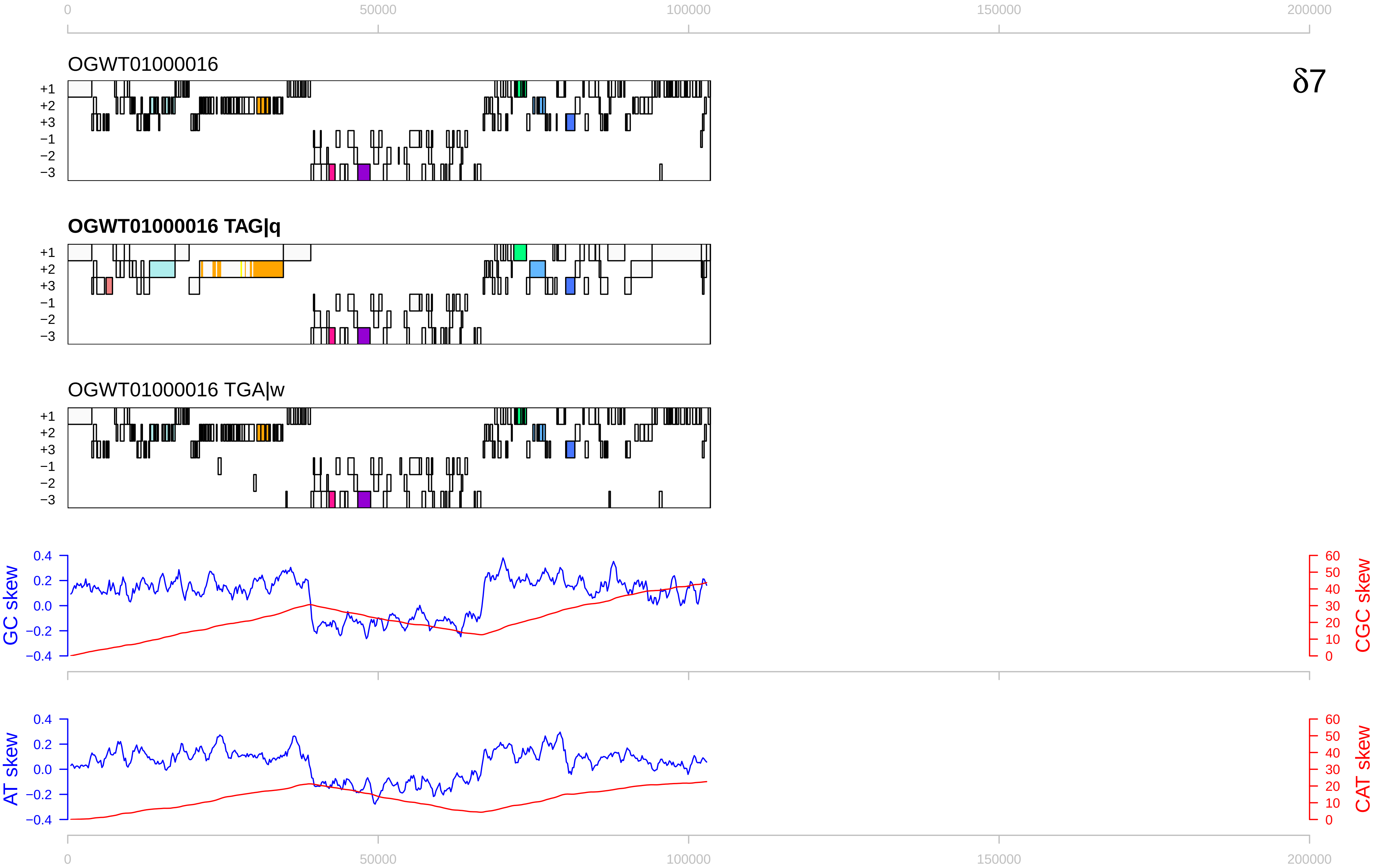

0
50000
100000
150000
200000
OGWT01000016
δ7
+1
+2
+3
−1
−2
−3
OGWT01000016 TAG|q
OGWT01000016 TGA|w
GC skew
CGC skew
AT skew
CAT skew
0.4
0.2
0.0
−0.2
−0.4
60
50
40
30
20
10
0

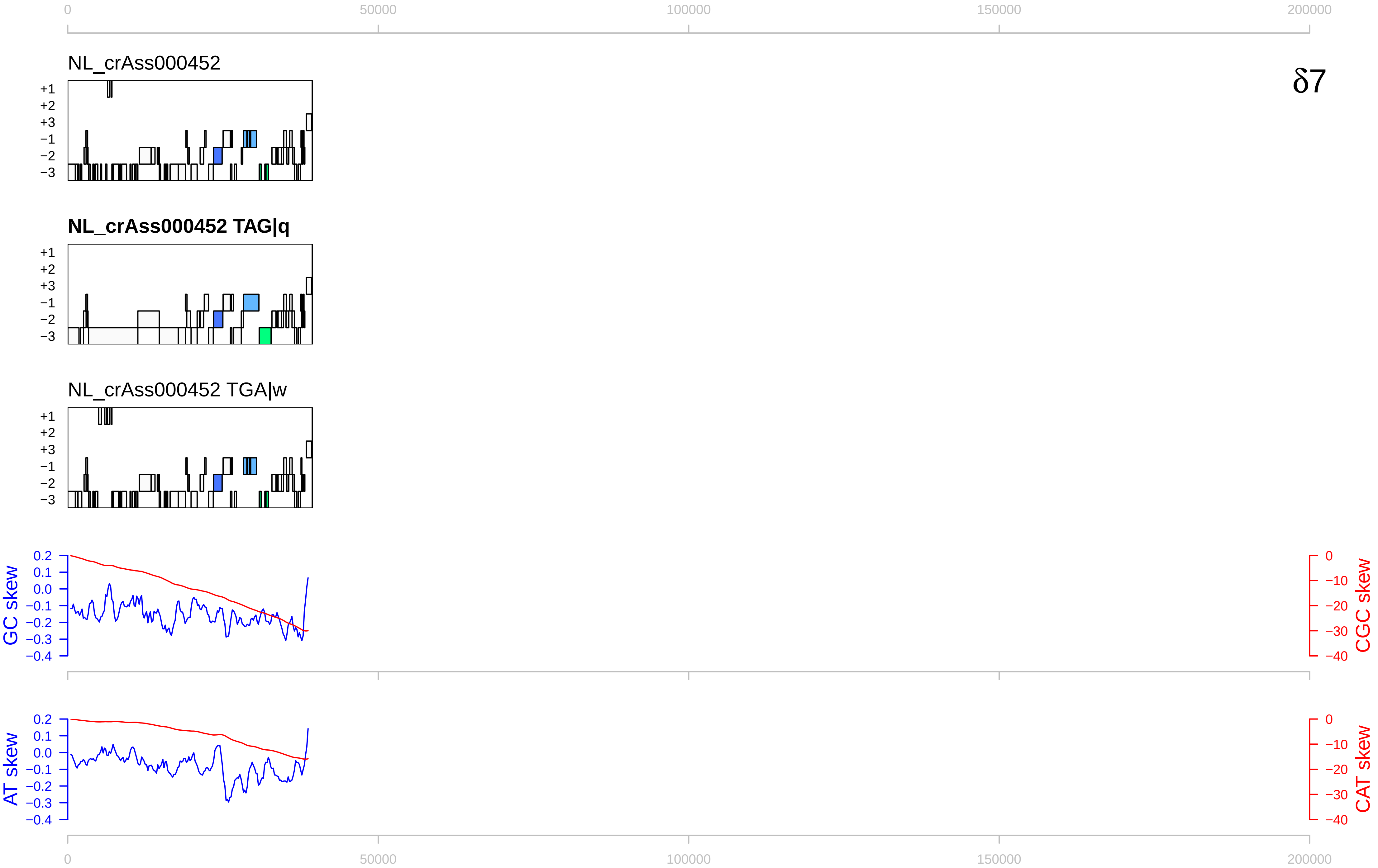
δ7
NL_crAss000452
NL_crAss000452 TAG|q
NL_crAss000452 TGA|w
+1
+2
+3
−1
−2
−3
GC skew
CGC skew
AT skew
CAT skew
0
50000
100000
150000
200000

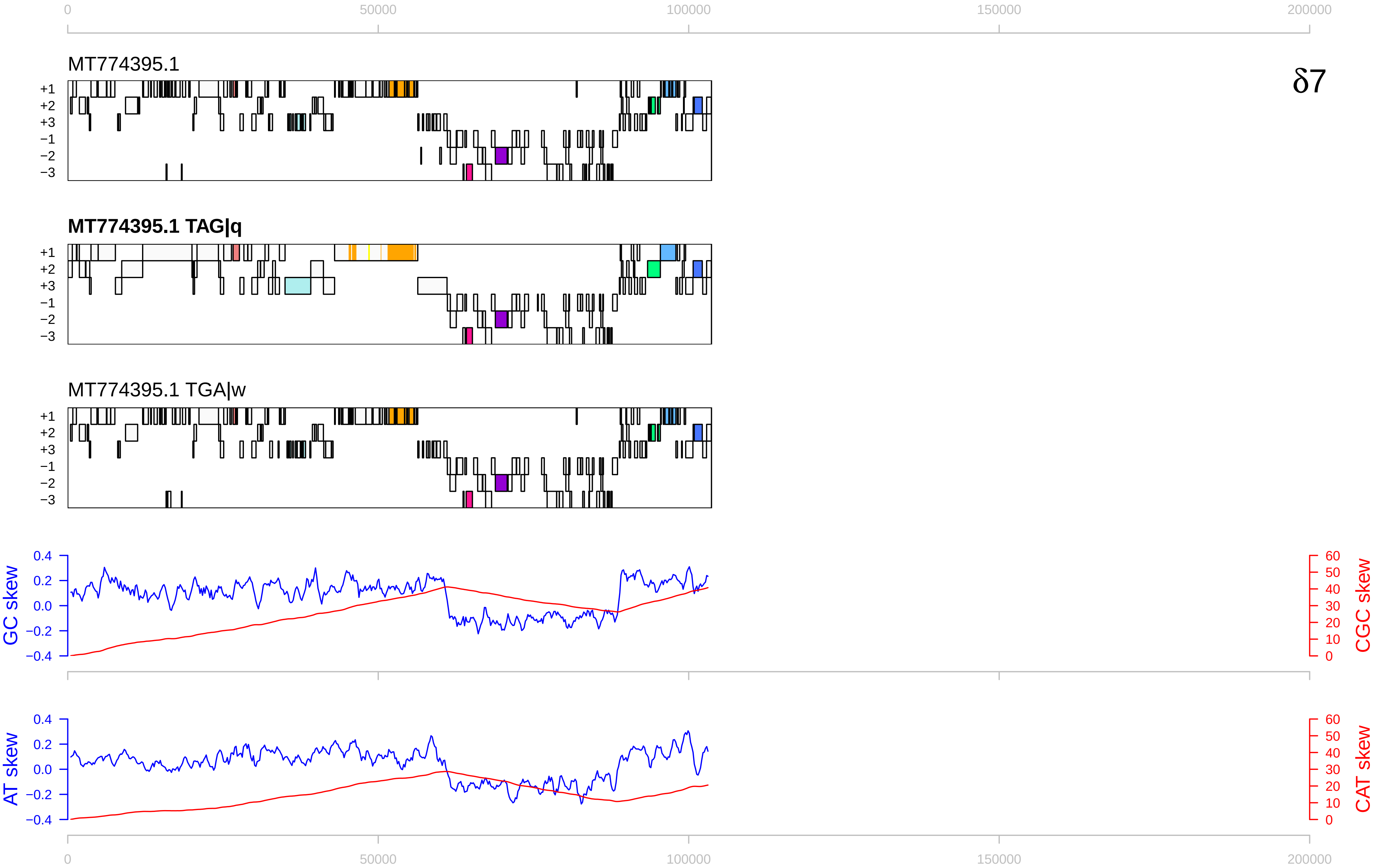

0
50000
100000
150000
200000
MT774395.1
δ7
+1
+2
+3
−1
−2
−3
MT774395.1 TAG|q
MT774395.1 TGA|w
GC skew
CGC skew
AT skew
CAT skew
0.4
0.2
0.0
−0.2
−0.4
60
50
40
30
20
10
0

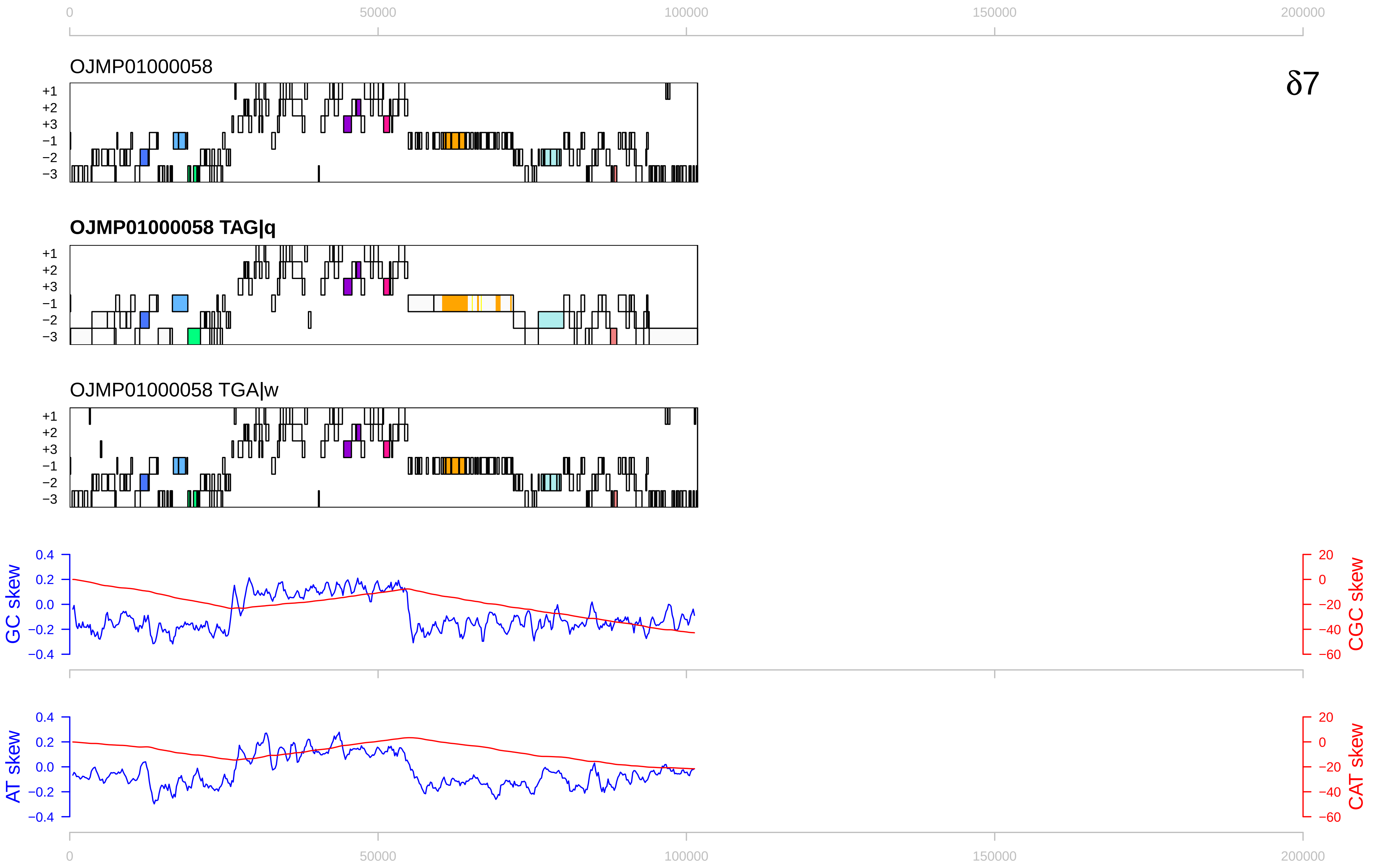

0
50000
100000
150000
200000
OJMP01000058
δ7
+1
+2
+3
−1
−2
−3
OJMP01000058 TAG|q
OJMP01000058 TGA|w
GC skew
CGC skew
AT skew
CAT skew
0.4
0.2
0.0
−0.2
−0.4
20
0
−20
−40
−60

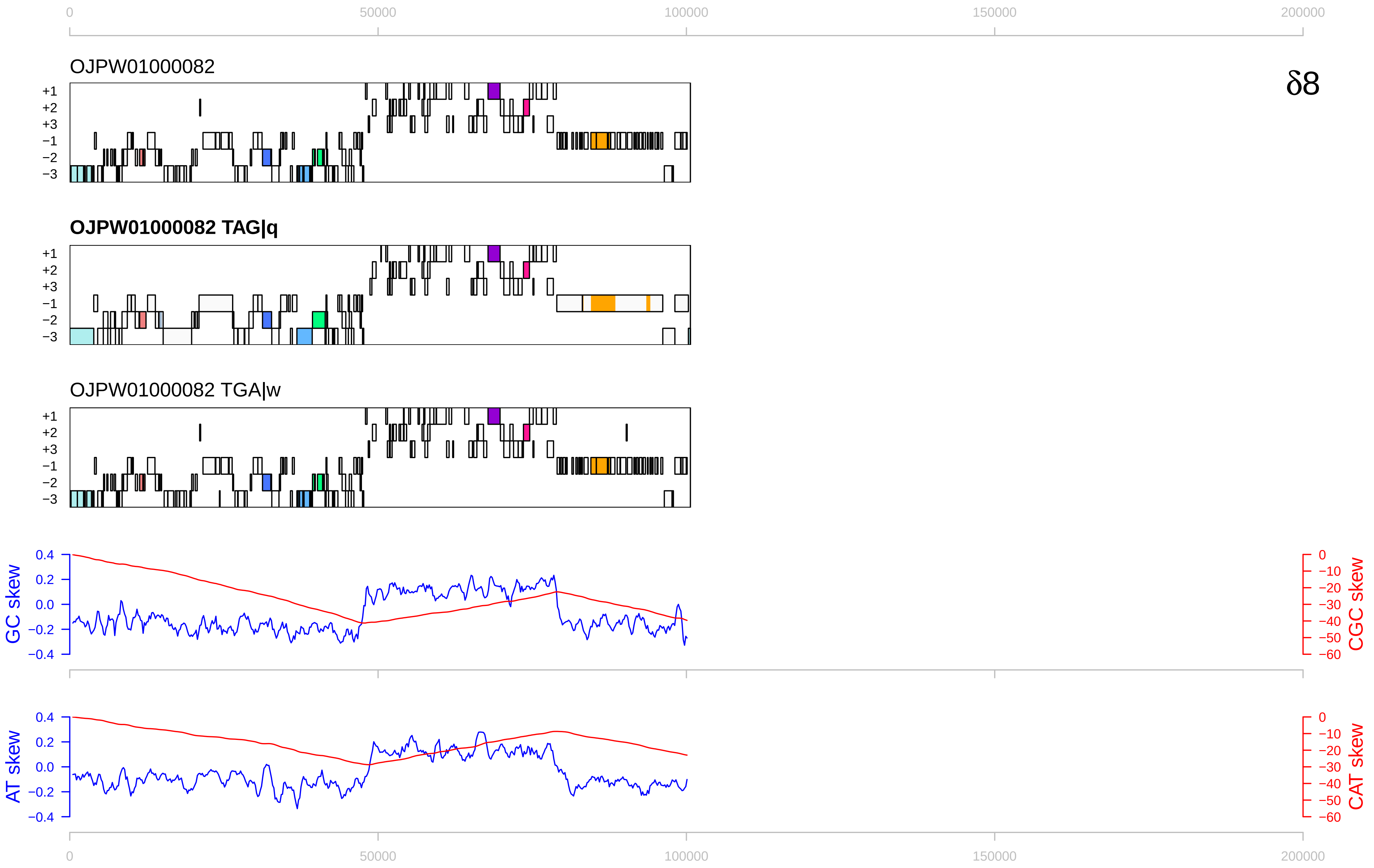

0
50000
100000
150000
200000
OJPW01000082
δ8
+1
+2
+3
−1
−2
−3
OJPW01000082 TAG|q
OJPW01000082 TGA|w
GC skew
CGC skew
AT skew
CAT skew
0.4
0.2
0.0
−0.2
−0.4
0
−10
−20
−30
−40
−50
−60

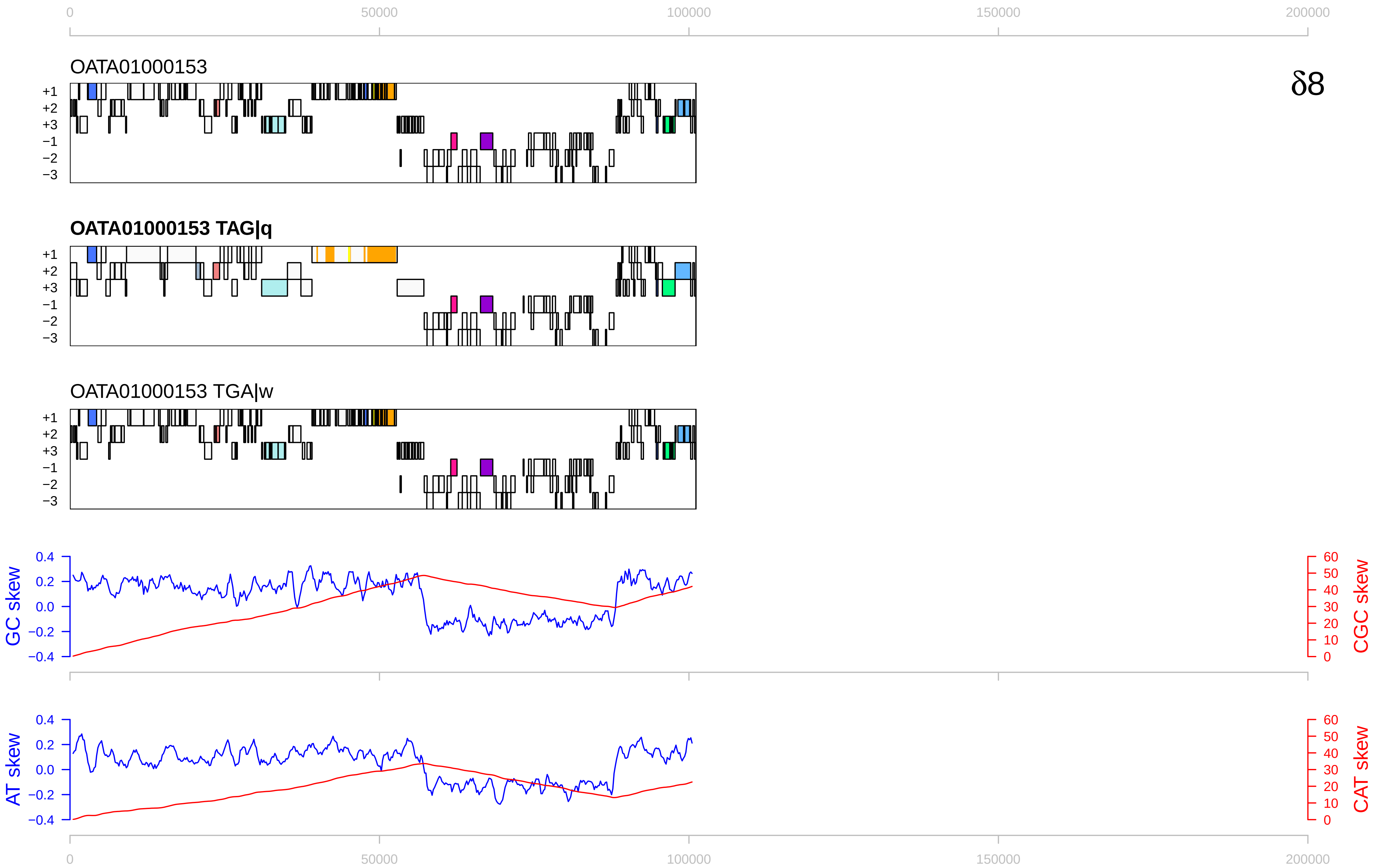

0
50000
100000
150000
200000
OATA01000153
δ8
+1
+2
+3
−1
−2
−3
OATA01000153 TAG|q
OATA01000153 TGA|w
GC skew
CGC skew
AT skew
CAT skew
0.4
0.2
0.0
−0.2
−0.4
60
50
40
30
20
10

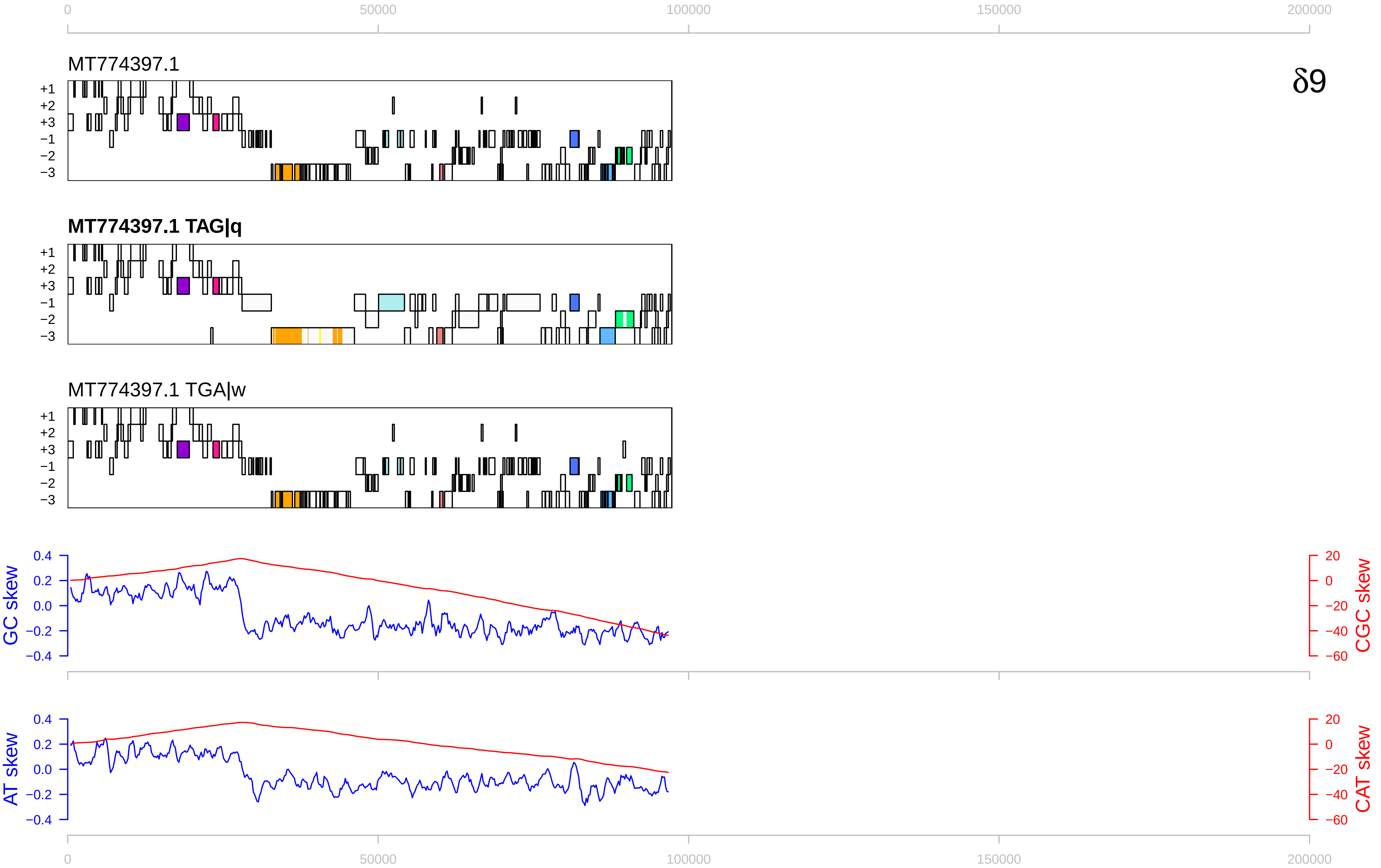

δ9
MT774397.1
MT774397.1 TAG|q
MT774397.1 TGA|w
+1
+2
+3
−1
−2
−3
GC skew
CGC skew
AT skew
CAT skew
0
50000
100000
150000
200000
0.4
0.2
0.0
−0.2
−0.4
20
0
−20
−40
−60

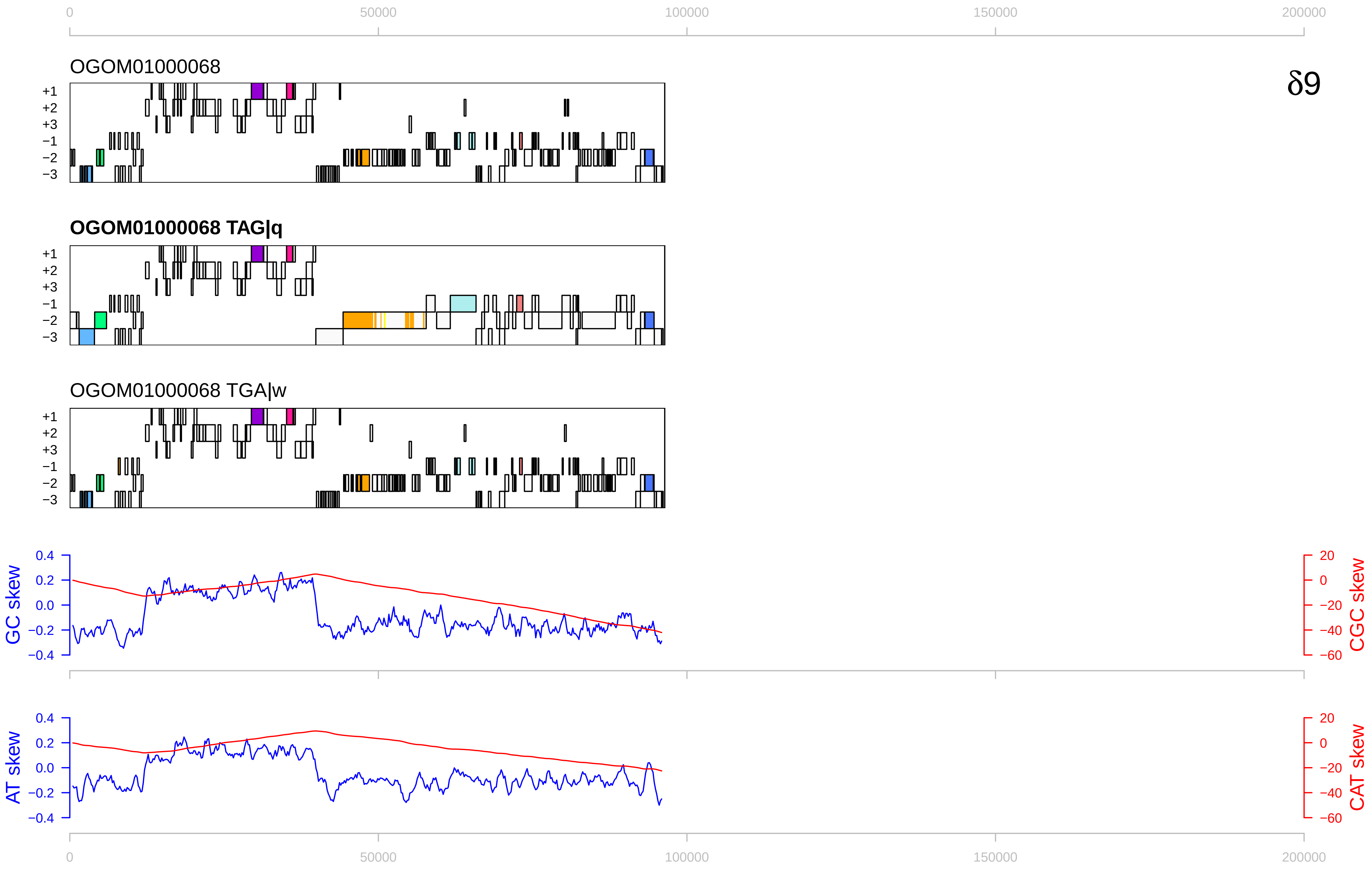

0
50000
100000
150000
200000
OGOM01000068
δ9
+1
+2
+3
−1
−2
−3
OGOM01000068 TAG|q
OGOM01000068 TGA|w
GC skew
0.4
0.2
0.0
−0.2
−0.4
CGC skew
20
0
−20
−40
−60
AT skew
CAT skew

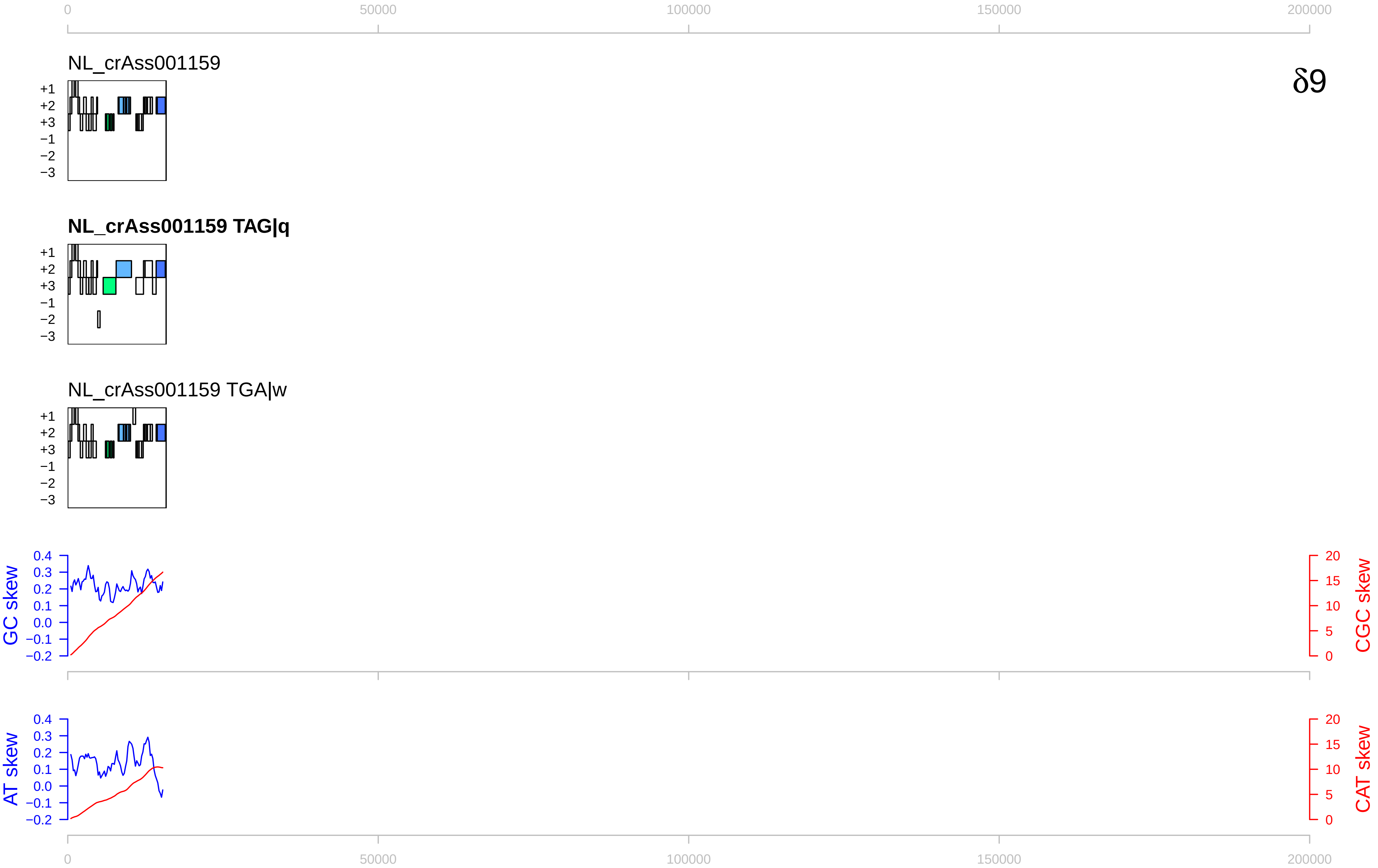

0
50000
100000
150000
200000
NL_crAss001159
δ9
+1
+2
+3
−1
−2
−3
NL_crAss001159 TAG|q
NL_crAss001159 TGA|w
GC skew
CGC skew
AT skew
CAT skew
0.4
0.3
0.2
0.1
0.0
−0.1
−0.2
20
15
10
5
0

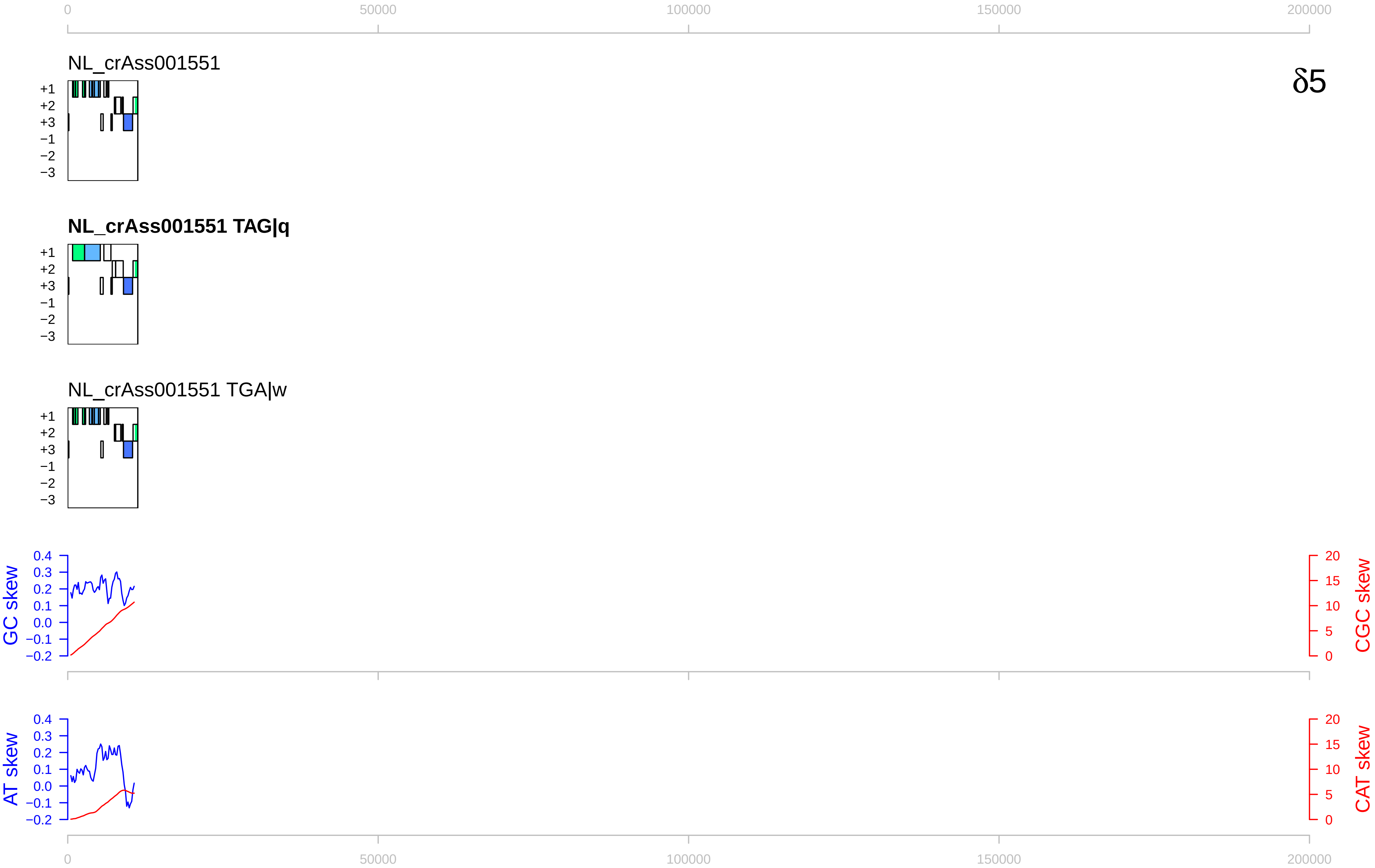

0
50000
100000
150000
200000
NL_crAss001551
δ5
+1
+2
+3
−1
−2
−3
NL_crAss001551 TAG|q
NL_crAss001551 TGA|w
GC skew
CGC skew
0.4
0.3
0.2
0.1
0.0
−0.1
−0.2
20
15
10
5
0
AT skew
CAT skew

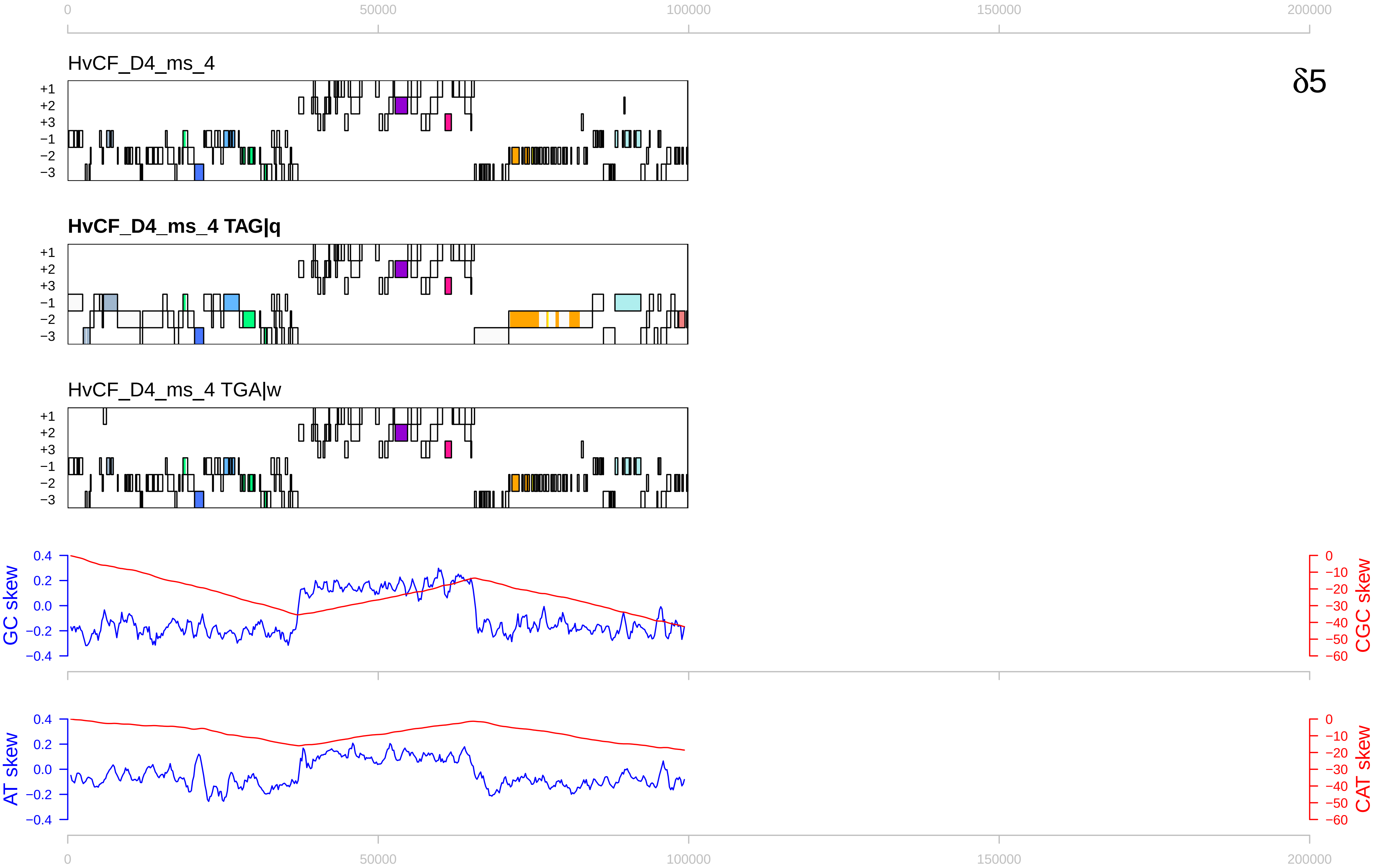
0
50000
100000
150000
200000
HvCF_D4_ms_4
δ5
+1
+2
+3
−1
−2
−3
HvCF_D4_ms_4 TAG|q
HvCF_D4_ms_4 TGA|w
GC skew
CGC skew
AT skew
CAT skew
0.4
0.2
0.0
−0.2
−0.4
0
−10
−20
−30
−40
−50
−60

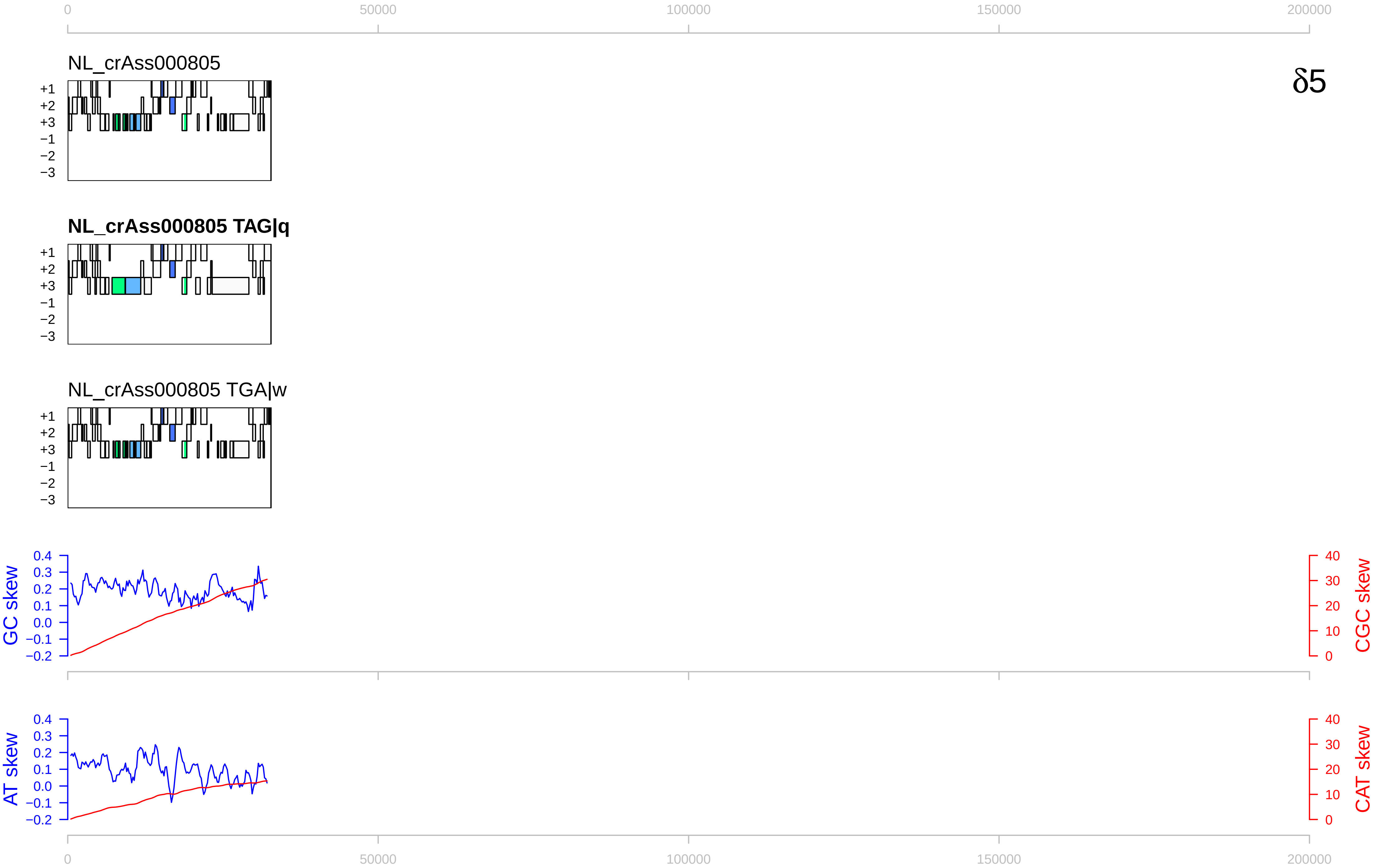

0
50000
100000
150000
200000
NL_crAss000805
δ5
+1
+2
+3
−1
−2
−3
NL_crAss000805 TAG|q
NL_crAss000805 TGA|w
GC skew
CGC skew
AT skew
CAT skew
0.4
0.3
0.2
0.1
0.0
−0.1
−0.2
40
30
20
10

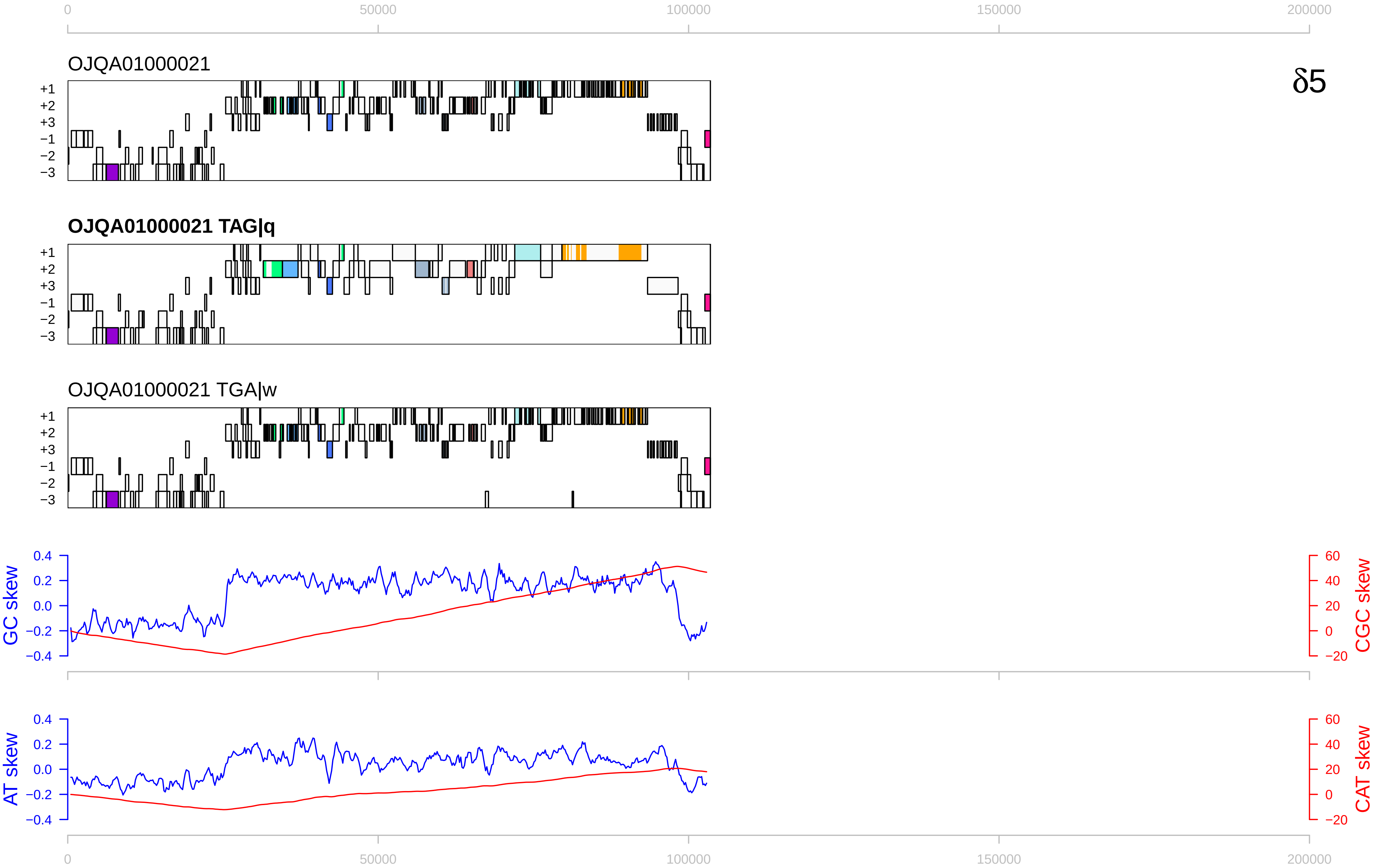

0
50000
100000
150000
200000
OJQA01000021
δ5
+1
+2
+3
−1
−2
−3
OJQA01000021 TAG|q
OJQA01000021 TGA|w
GC skew
CGC skew
AT skew
CAT skew
0.4
0.2
0.0
−0.2
−0.4
60
40
20
0
−20

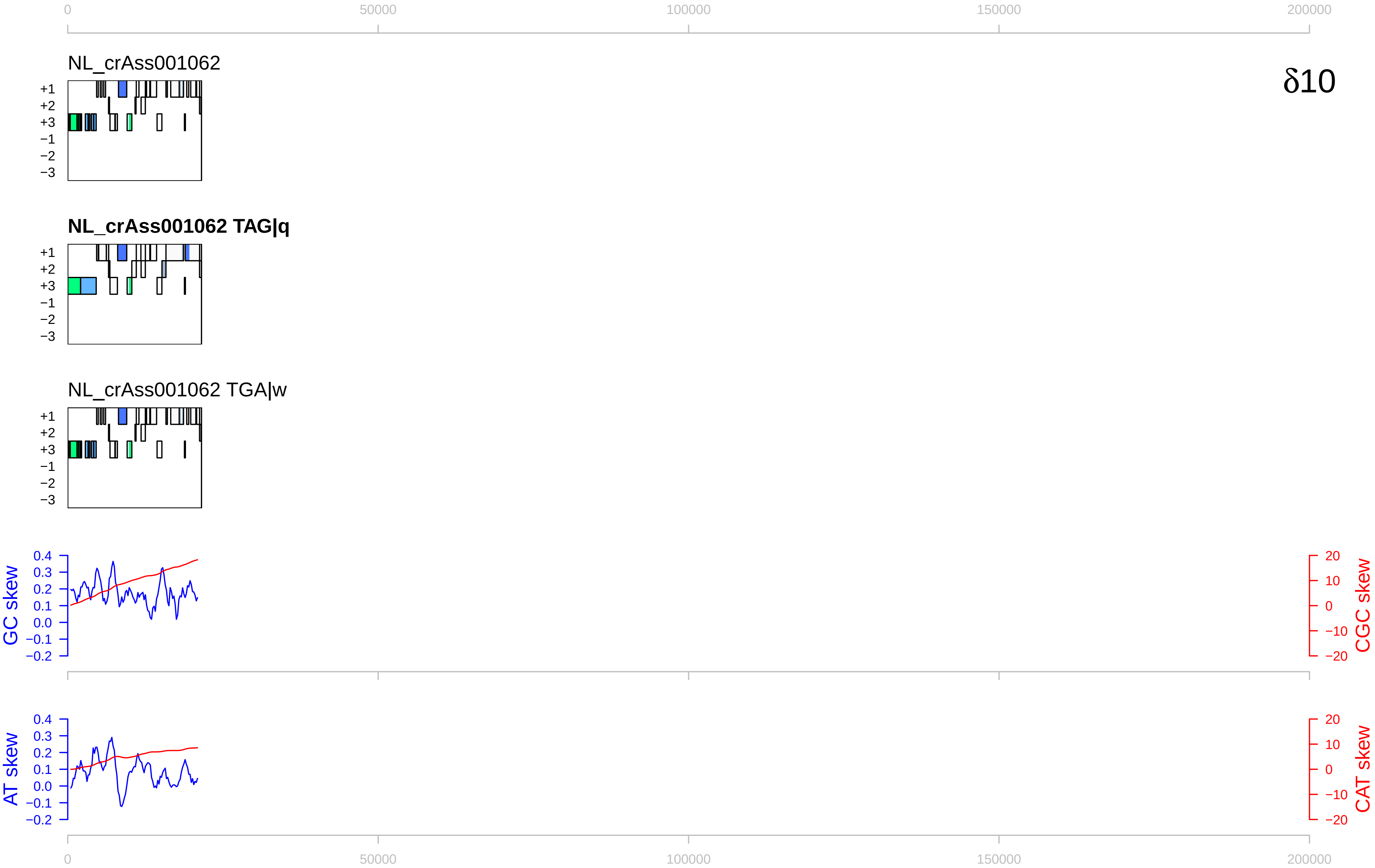

0
50000
100000
150000
200000
NL_crAss001062
δ10
+1
+2
+3
−1
−2
−3
NL_crAss001062 TAG|q
NL_crAss001062 TGA|w
GC skew
CGC skew
AT skew
CAT skew
0.4
0.3
0.2
0.1
0.0
−0.1
−0.2
20
10
0
−10
−20

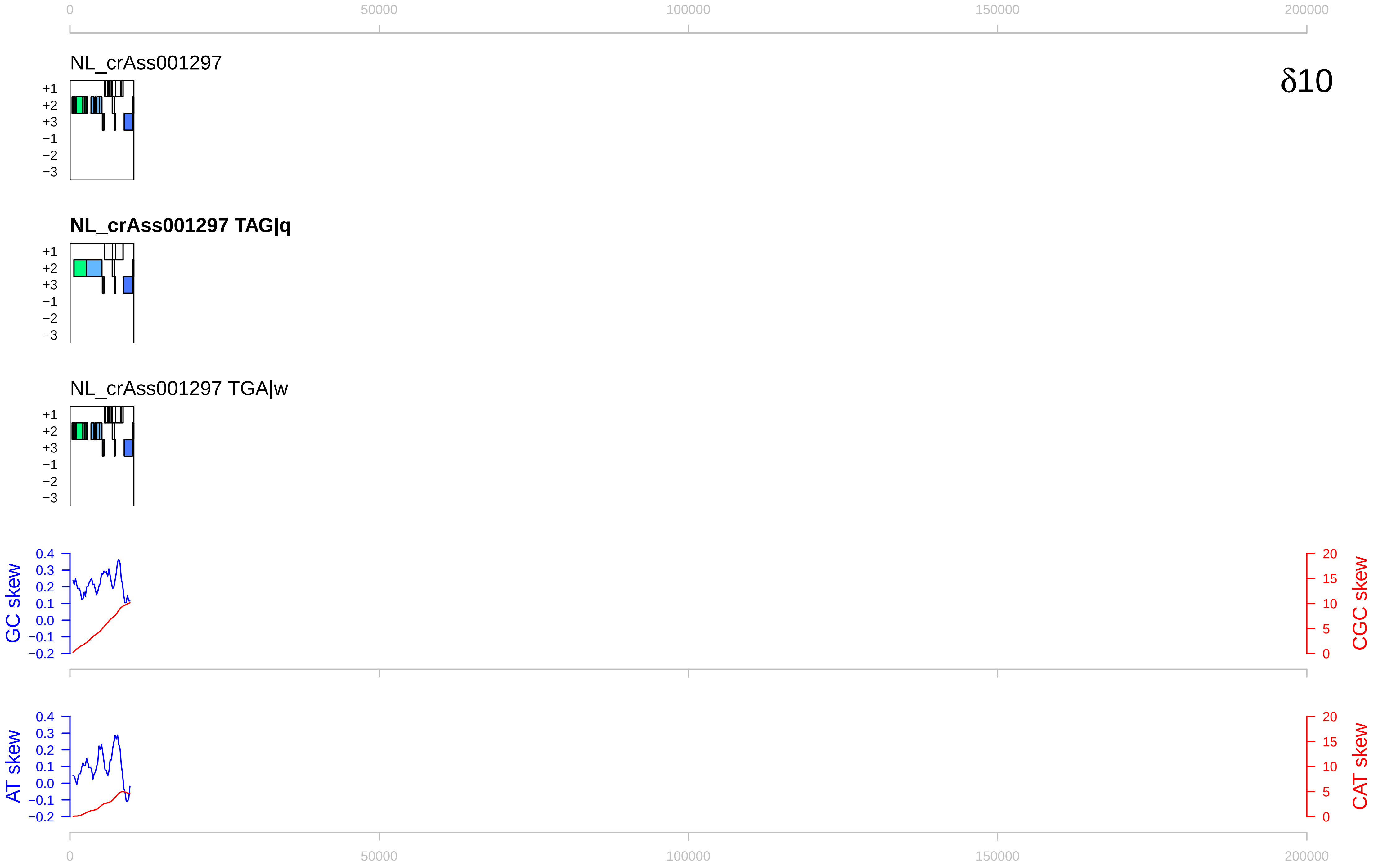

0
50000
100000
150000
200000
NL_crAss001297
δ10
+1
+2
+3
−1
−2
−3
NL_crAss001297 TAG|q
NL_crAss001297 TGA|w
GC skew
CGC skew
AT skew
CAT skew
0.4
0.3
0.2
0.1
0.0
−0.1
−0.2
20
15
10
5
0

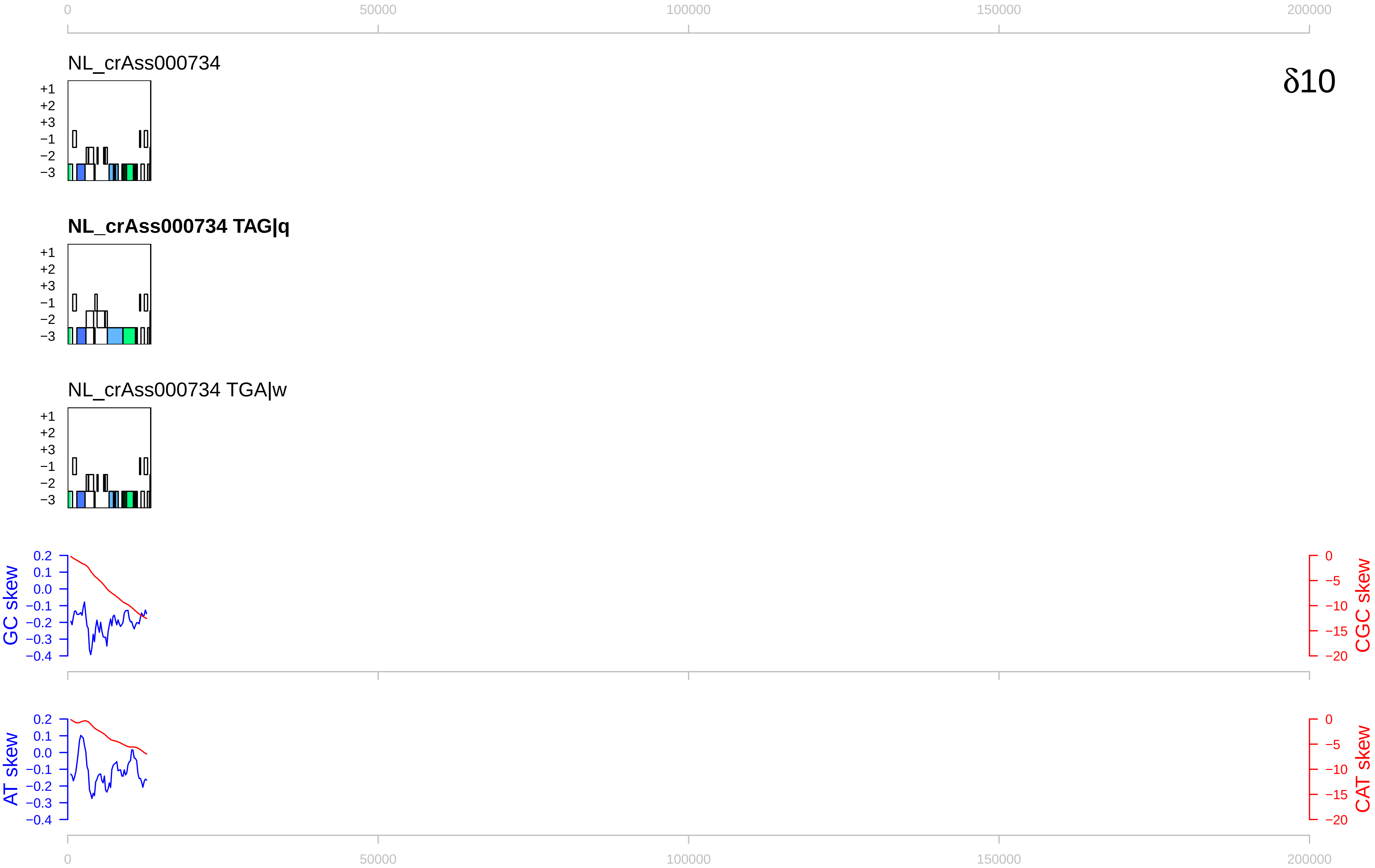

0
50000
100000
150000
200000
NL_crAss000734
δ10
+1
+2
+3
−1
−2
−3
NL_crAss000734 TAG|q
NL_crAss000734 TGA|w
GC skew
CGC skew
AT skew
CAT skew
0.2
0.1
0.0
−0.1
−0.2
−0.3
−0.4
−5
−10
−15
−20

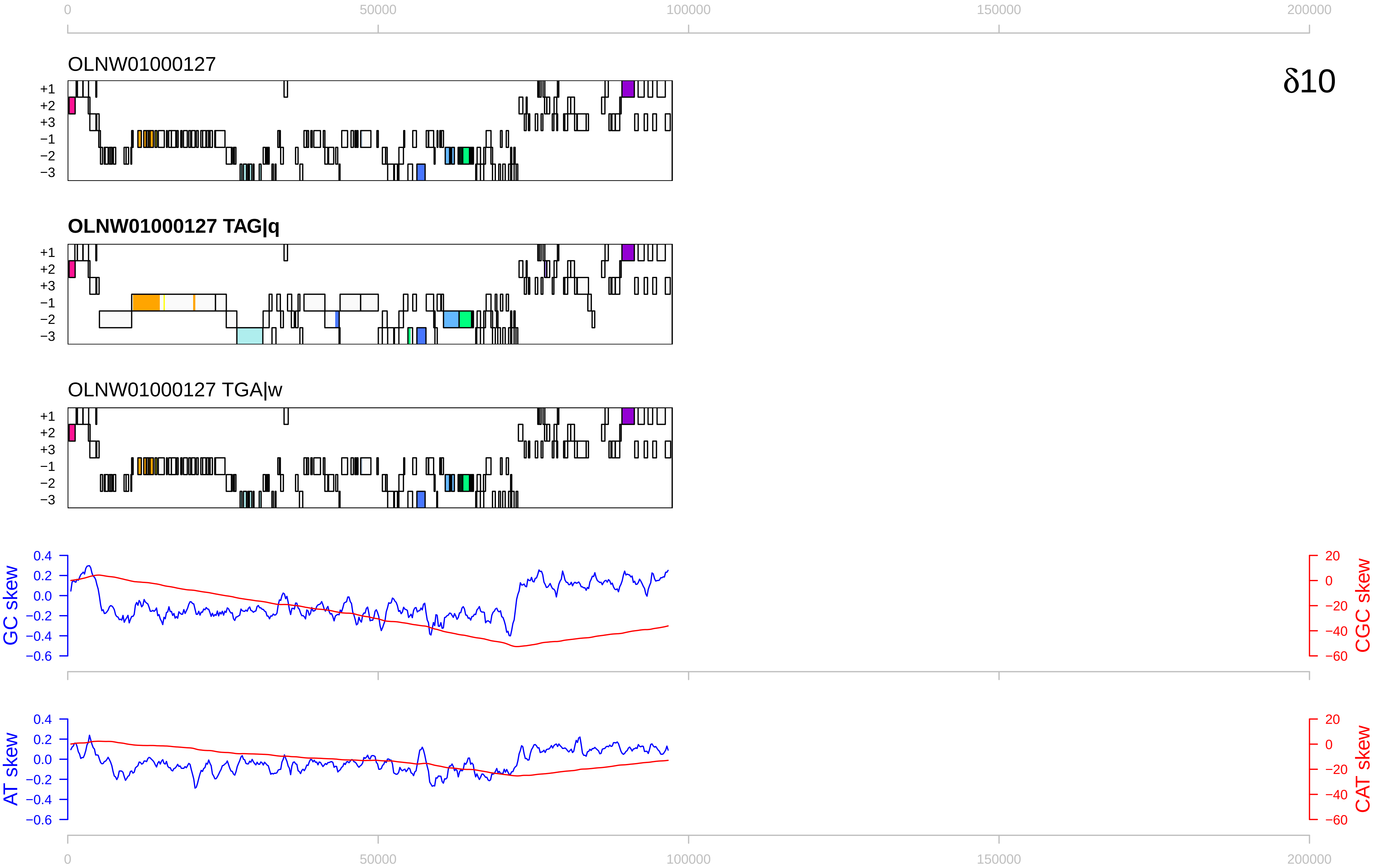

0
50000
100000
150000
200000
OLNW01000127
δ10
+1
+2
+3
−1
−2
−3
OLNW01000127 TAG|q
OLNW01000127 TGA|w
GC skew
CGC skew
0.4
0.2
0.0
−0.2
−0.4
−0.6
20
0
−20
−40
−60
AT skew
CAT skew

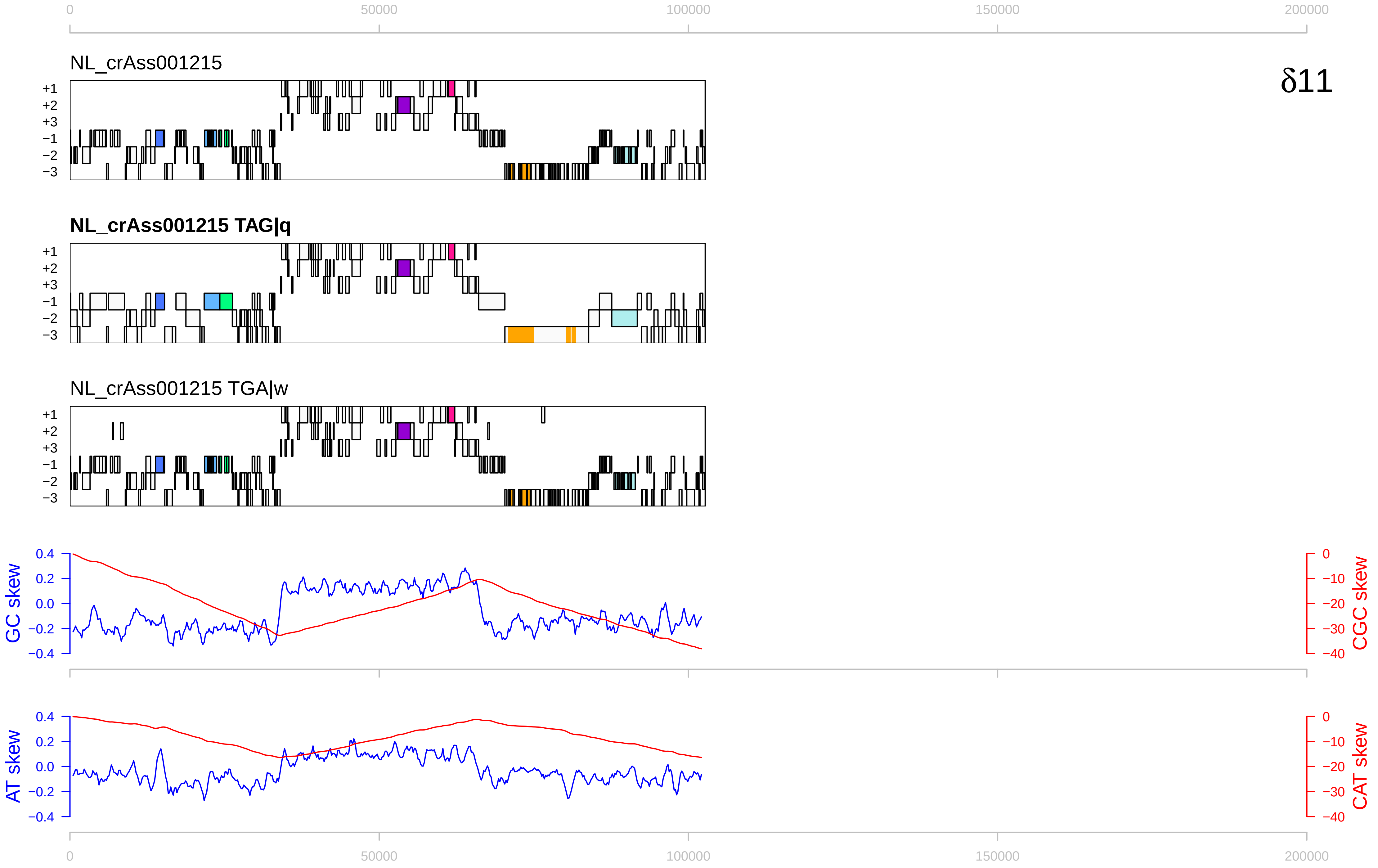

0
50000
100000
150000
200000
NL_crAss001215
δ11
+1
+2
+3
−1
−2
−3
NL_crAss001215 TAG|q
NL_crAss001215 TGA|w
GC skew
CGC skew
0.4
0.2
0.0
−0.2
−0.4
0
−10
−20
−30
−40
AT skew
CAT skew

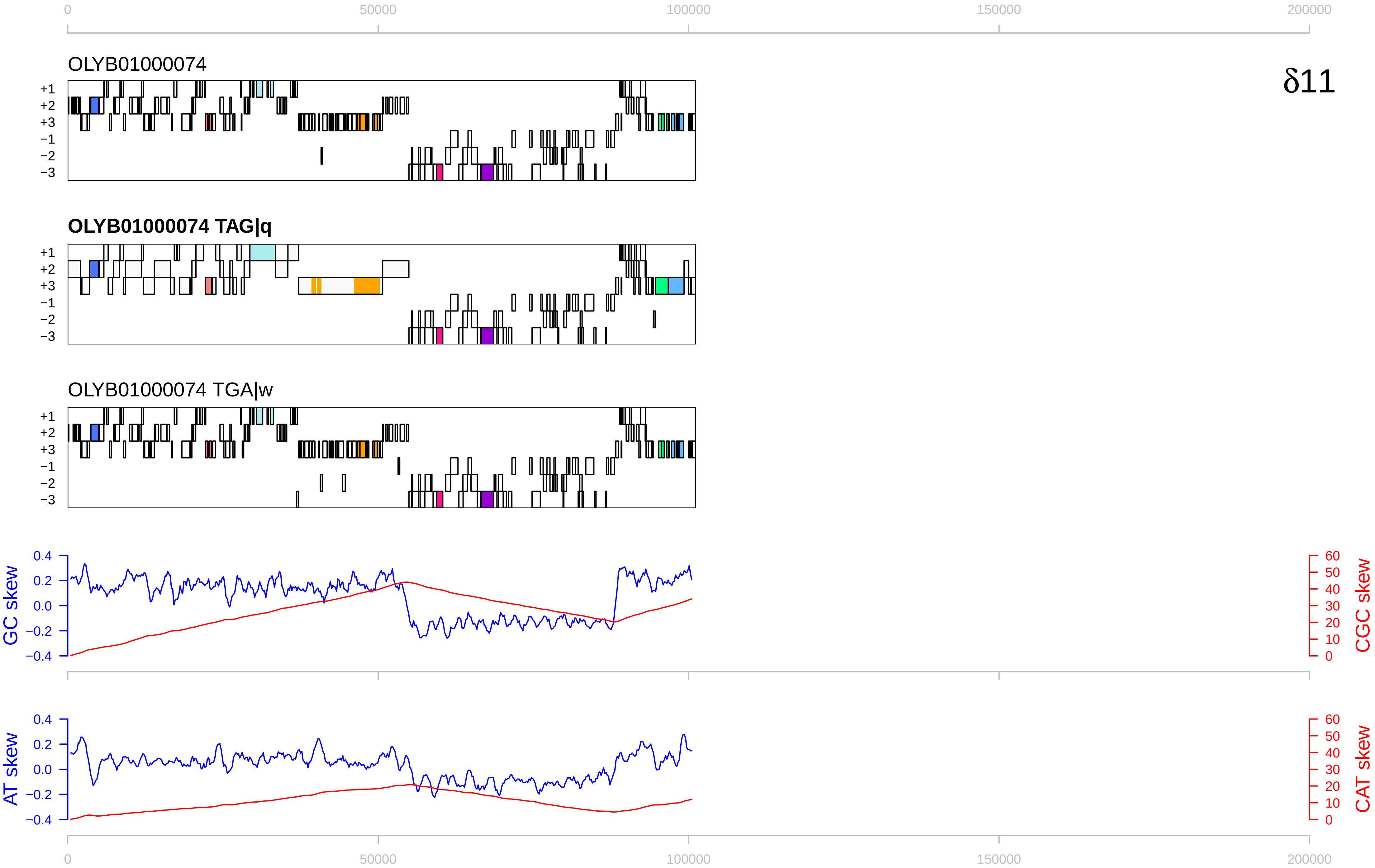
δ11
OLYB01000074
OLYB01000074 TAG|q
OLYB01000074 TGA|w
+1
+2
+3
−1
−2
−3
GC skew
CGC skew
AT skew
CAT skew
0
50000
100000
150000
200000

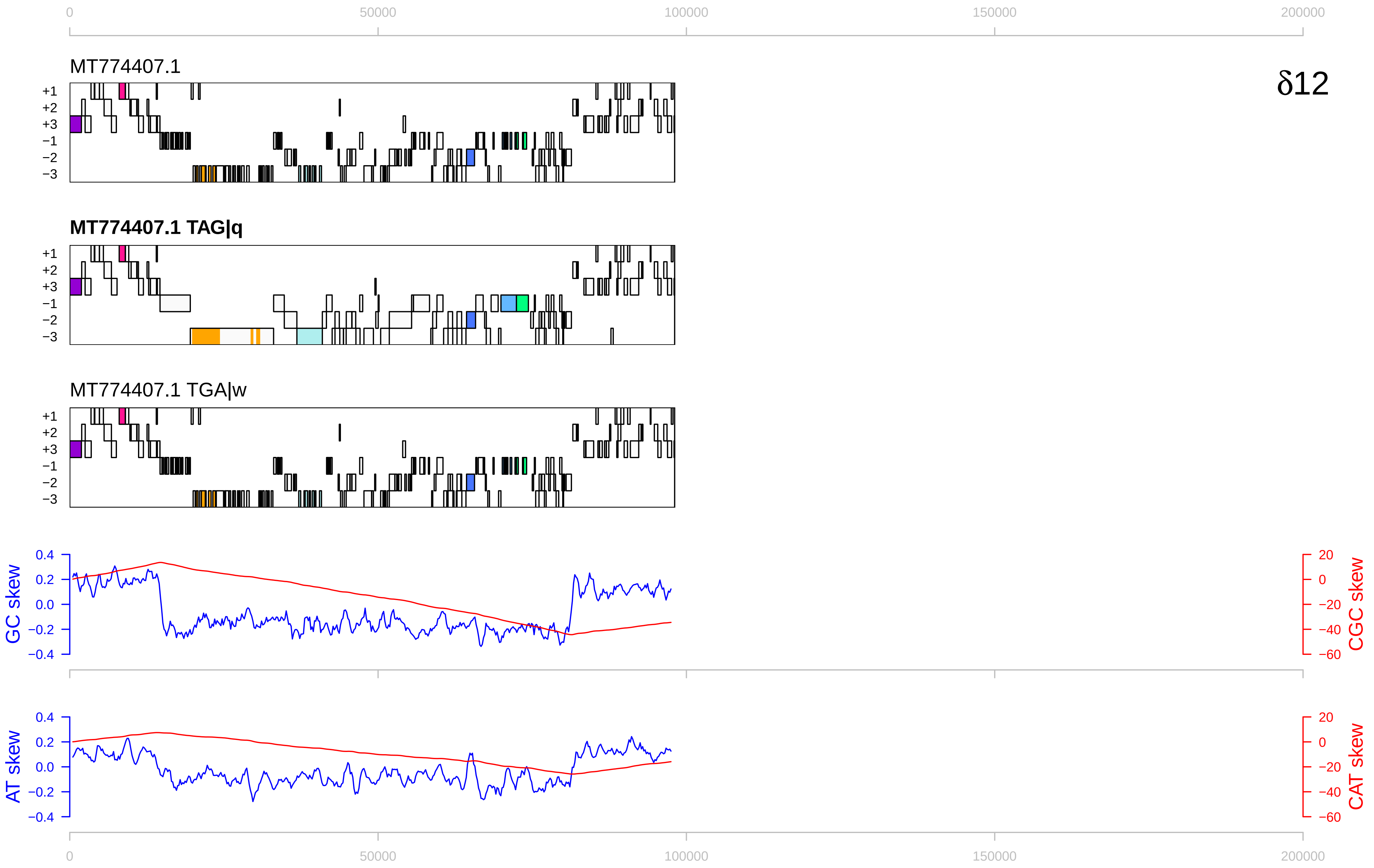

0
50000
100000
150000
200000
MT774407.1
δ12
+1
+2
+3
−1
−2
−3
MT774407.1 TAG|q
MT774407.1 TGA|w
GC skew
CGC skew
AT skew
CAT skew
0.4
0.2
0.0
−0.2
−0.4
20
0
−20
−40
−60

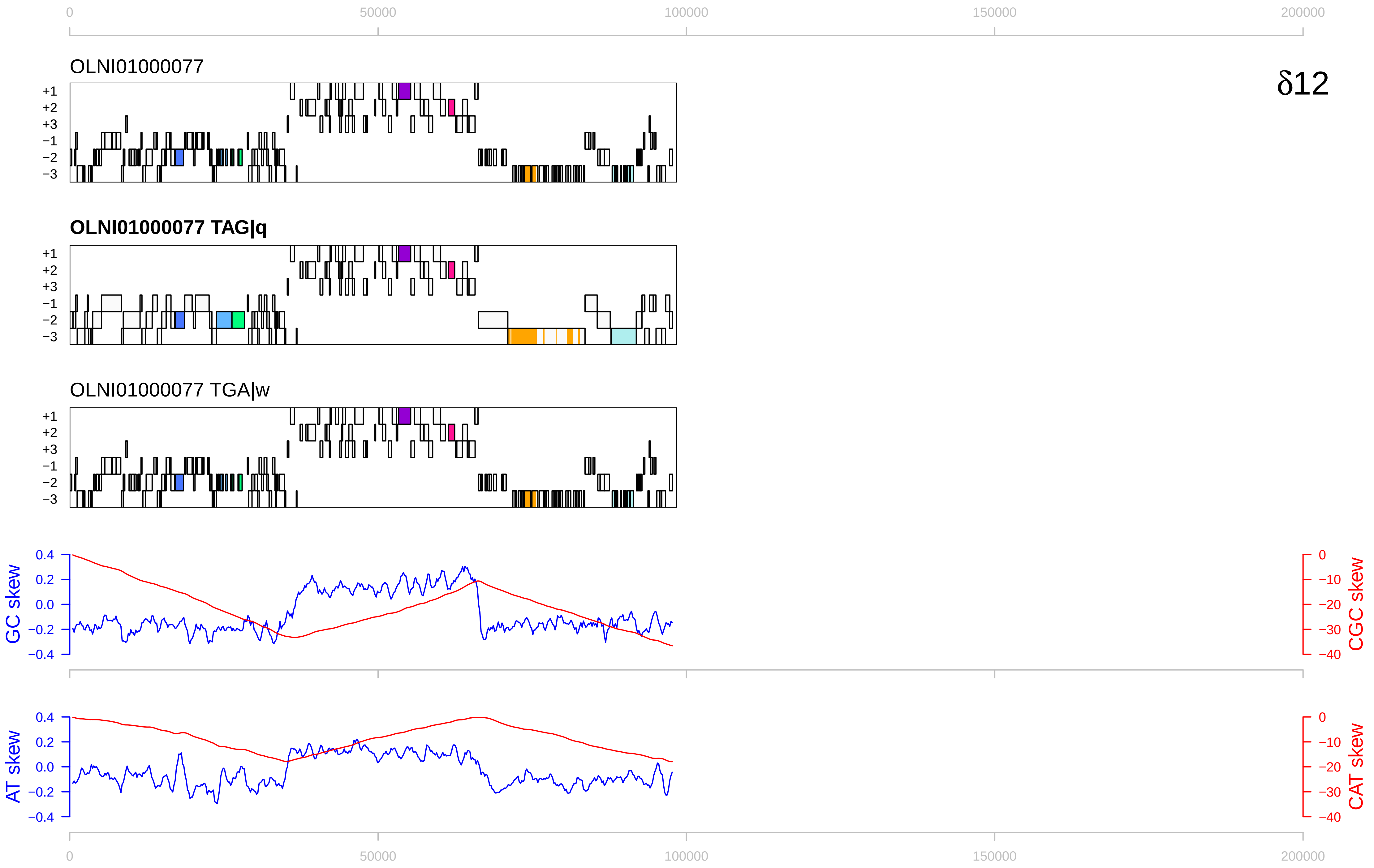

0
50000
100000
150000
200000
OLNI01000077
δ12
+1
+2
+3
−1
−2
−3
OLNI01000077 TAG|q
OLNI01000077 TGA|w
GC skew
CGC skew
AT skew
CAT skew
0.4
0.2
0.0
−0.2
−0.4
0
−10
−20
−30
−40

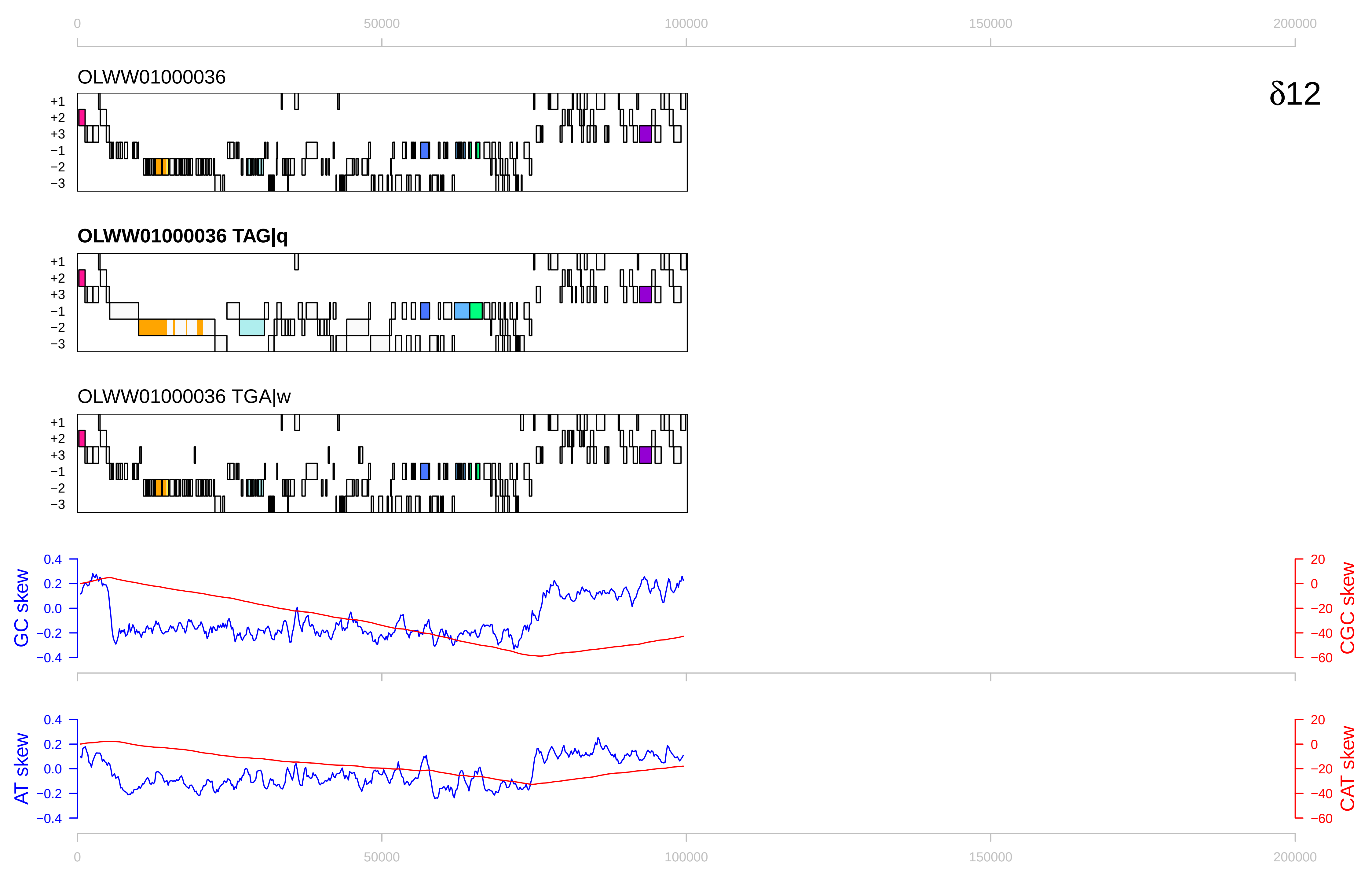

δ12
OLWW01000036
OLWW01000036 TAG|q
OLWW01000036 TGA|w
+1
+2
+3
−1
−2
−3
GC skew
CGC skew
AT skew
CAT skew
0.4
0.2
0.0
−0.2
−0.4
20
0
−20
−40
−60
0
50000
100000
150000
200000

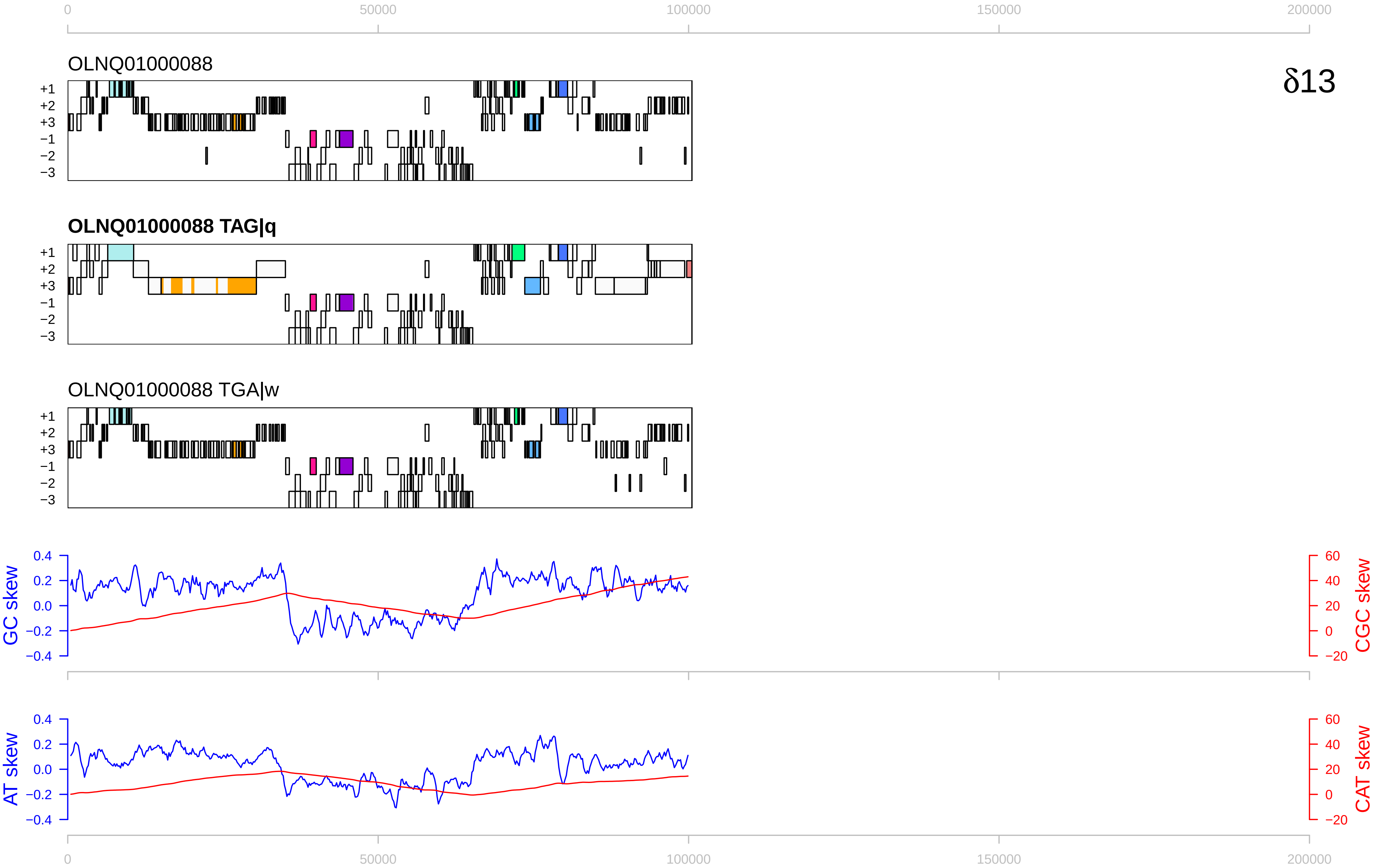

δ13
OLNQ01000088
OLNQ01000088 TAG|q
OLNQ01000088 TGA|w
+1
+2
+3
−1
−2
−3
GC skew
CGC skew
AT skew
CAT skew
0
50000
100000
150000
200000
0.4
0.2
0.0
−0.2
−0.4
60
40
20
0
−20

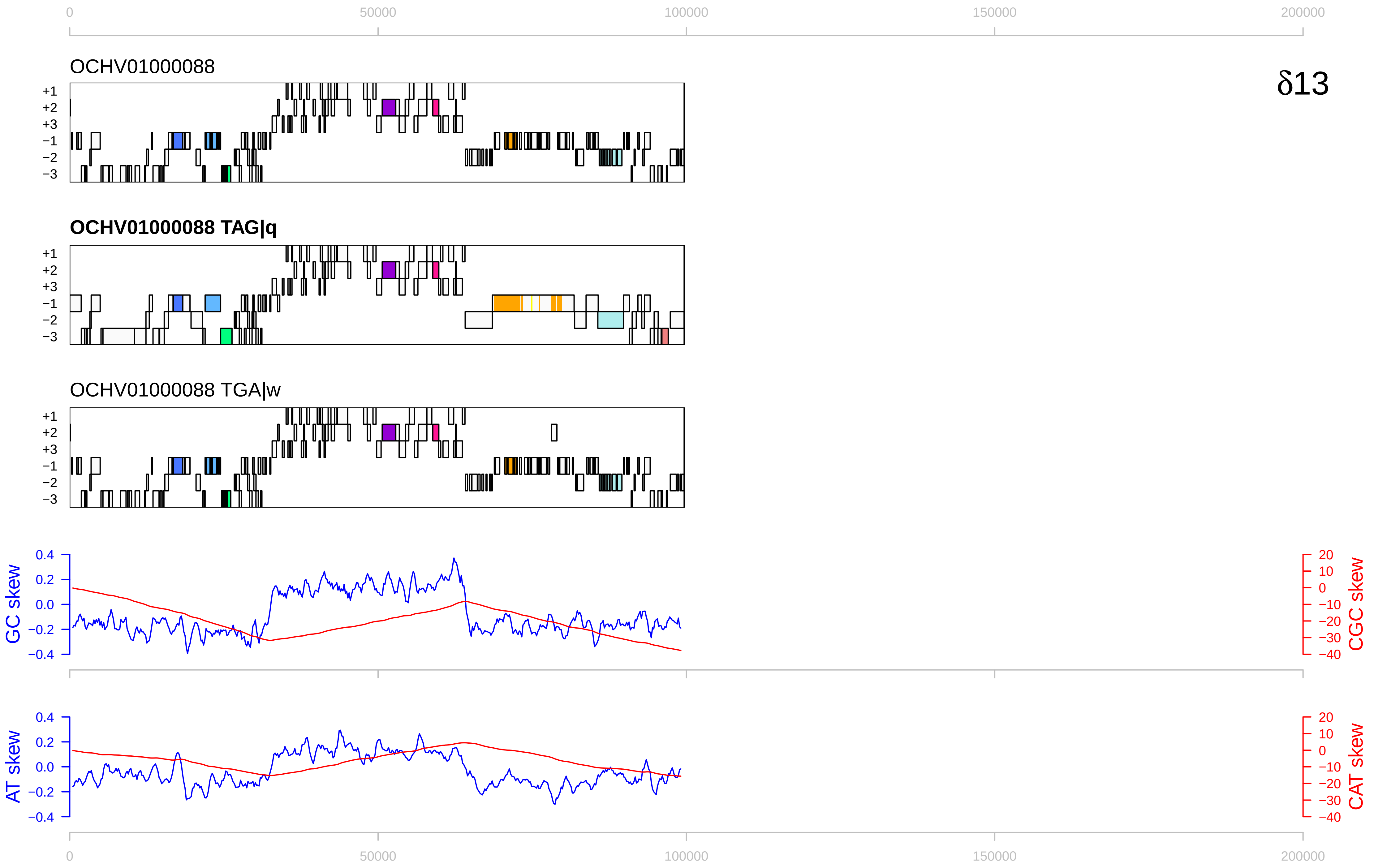

δ13
OCHV01000088
OCHV01000088 TAG|q
OCHV01000088 TGA|w
+1
+2
+3
−1
−2
−3
GC skew
CGC skew
AT skew
CAT skew
0
50000
100000
150000
200000
0.4
0.2
0.0
−0.2
−0.4
20
10
0
−10
−20
−30
−40

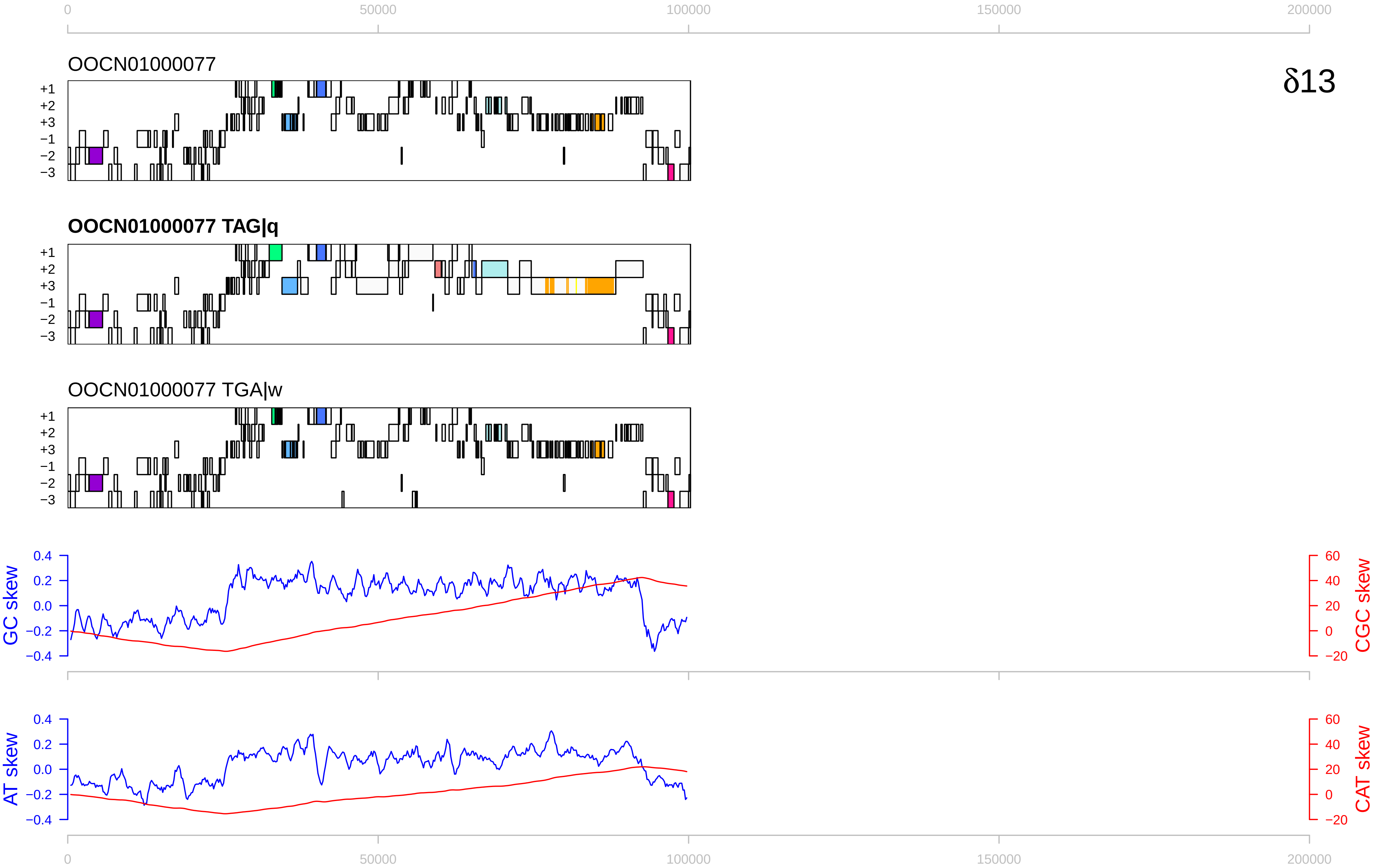

δ13
OOCN01000077
OOCN01000077 TAG|q
OOCN01000077 TGA|w
+1
+2
+3
−1
−2
−3
0
50000
100000
150000
200000
GC skew
CGC skew
AT skew
CAT skew
0.4
0.2
0.0
−0.2
−0.4
60
40
20
0
−20

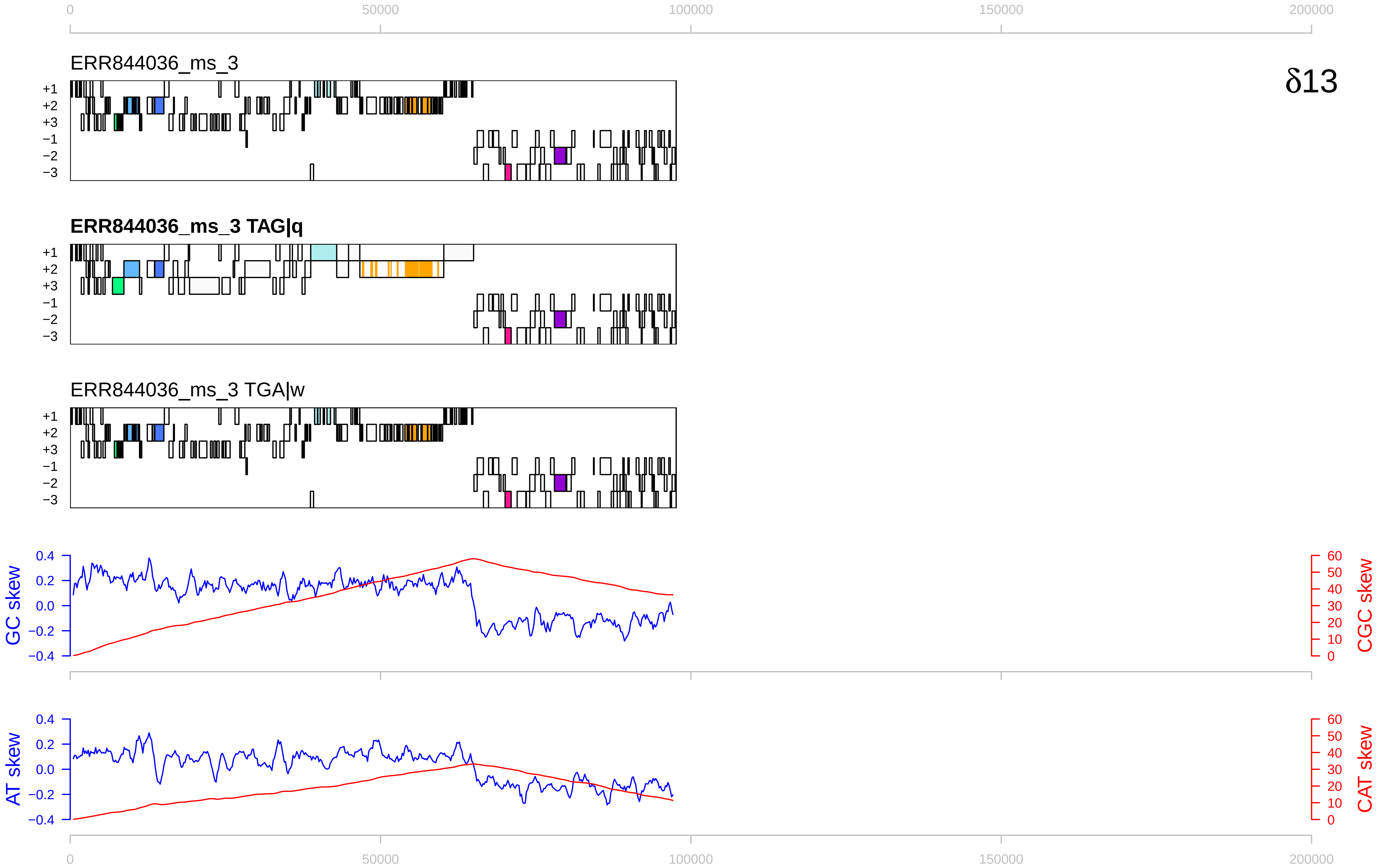

0
50000
100000
150000
200000
ERR844036_ms_3
δ13
+1
+2
+3
−1
−2
−3
ERR844036_ms_3 TAG|q
ERR844036_ms_3 TGA|w
GC skew
CGC skew
AT skew
CAT skew
0.4
0.2
0.0
−0.2
−0.4
60
50
40
30
20
10
0

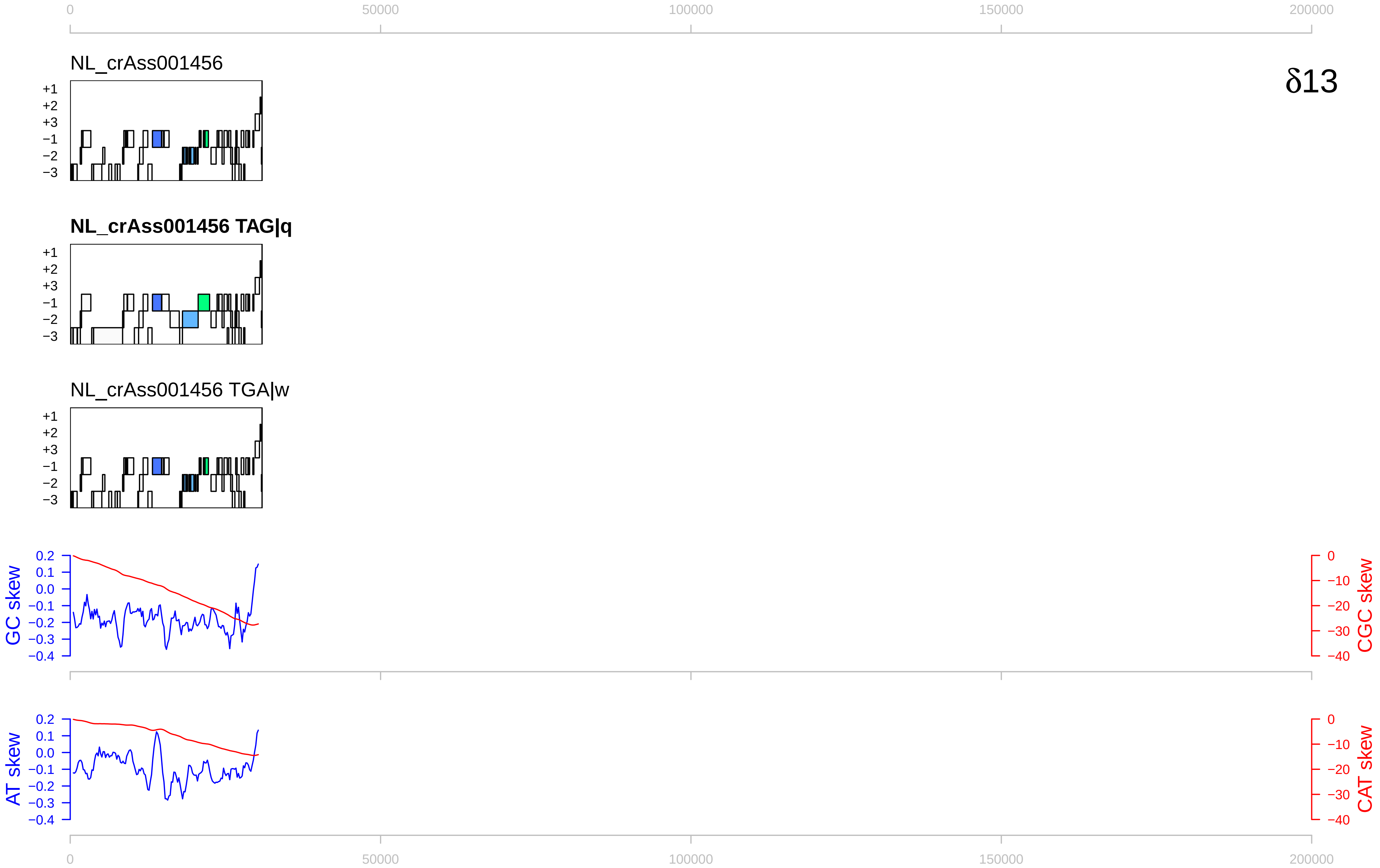

0
50000
100000
150000
200000
NL_crAss001456
δ13
+1
+2
+3
−1
−2
−3
NL_crAss001456 TAG|q
NL_crAss001456 TGA|w
GC skew
CGC skew
0.2
0.1
0.0
−0.1
−0.2
−0.3
−0.4
0
−10
−20
−30
−40
AT skew
CAT skew

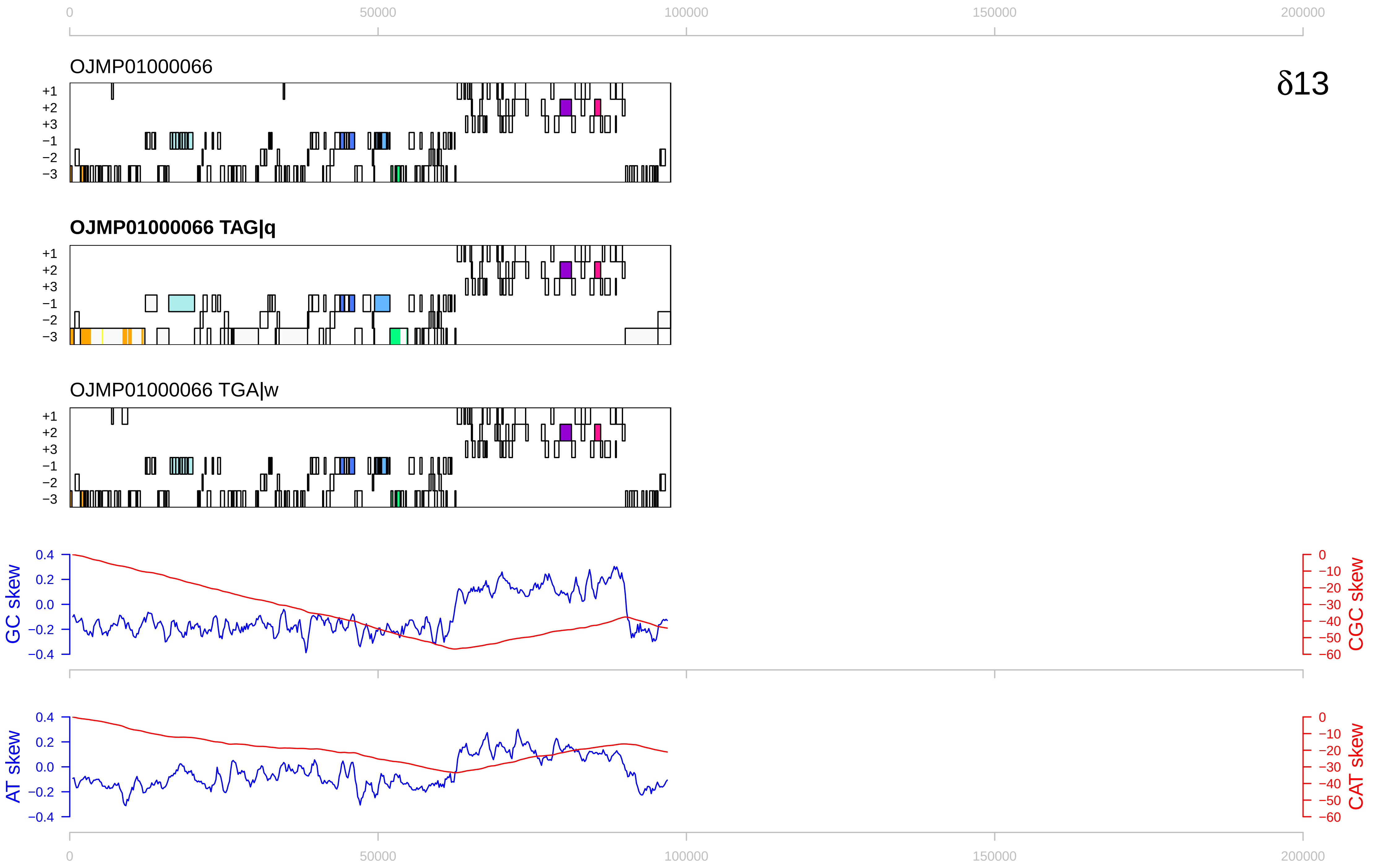

δ13
OJMP01000066
OJMP01000066 TAG|q
OJMP01000066 TGA|w
+1
+2
+3
−1
−2
−3
GC skew
CGC skew
AT skew
CAT skew
0
50000
100000
150000
200000

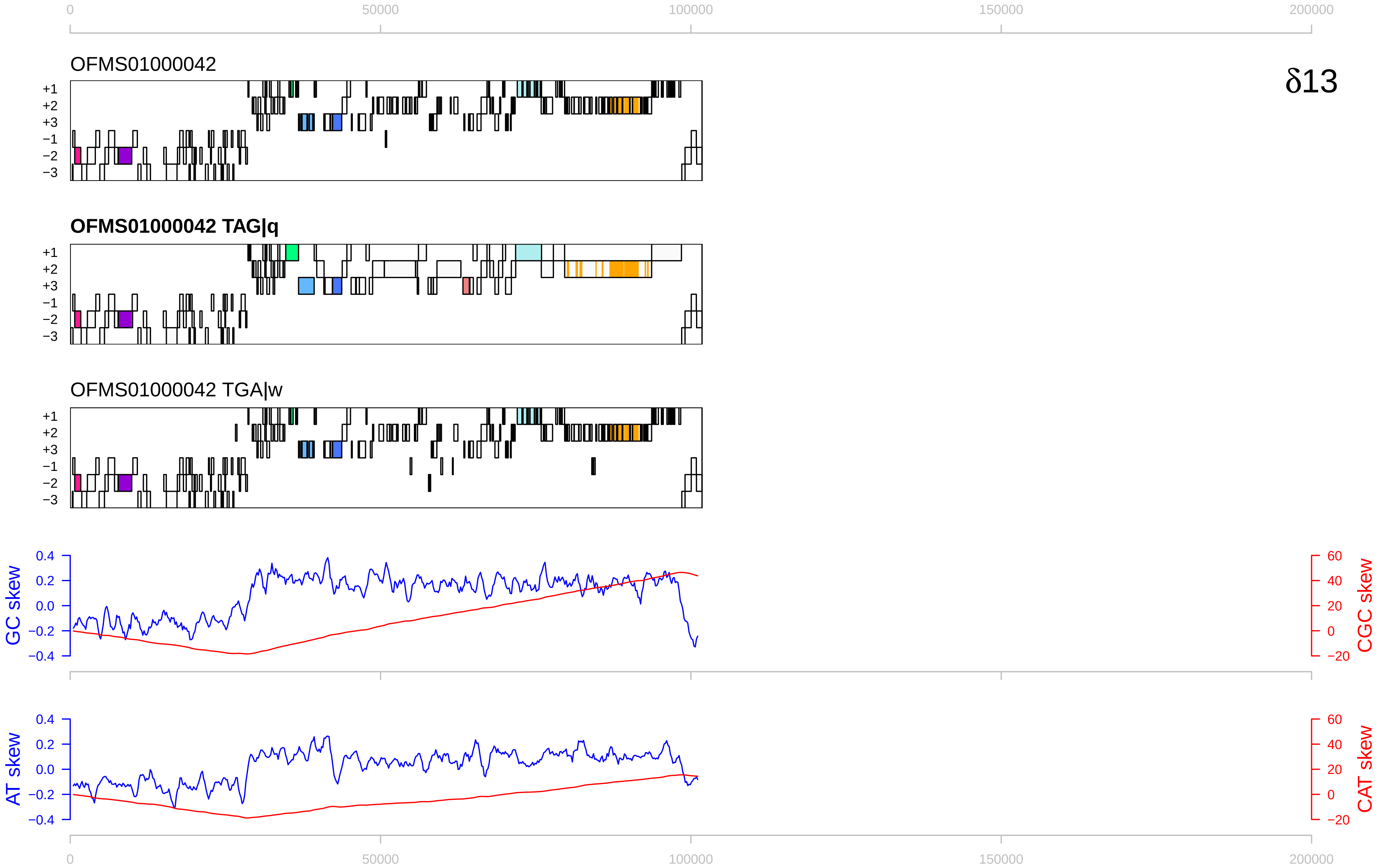
0
50000
100000
150000
200000
OFMS01000042
δ13
+1
+2
+3
−1
−2
−3
OFMS01000042 TAG|q
OFMS01000042 TGA|w
GC skew
CGC skew
0.4
0.2
0.0
−0.2
−0.4
60
40
20
0
−20
AT skew
CAT skew

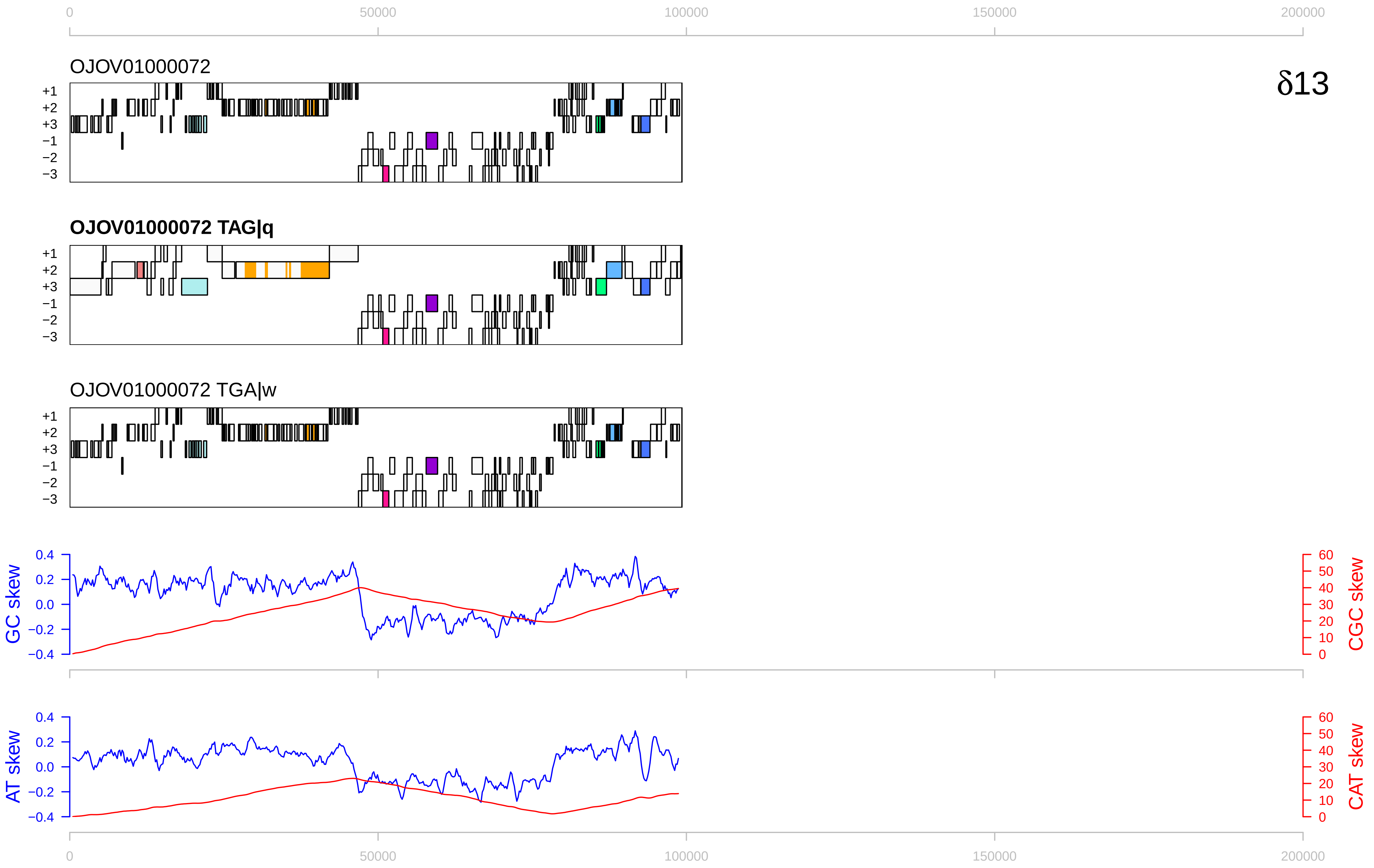
0
50000
100000
150000
200000
OJOV01000072
δ13
+1
+2
+3
−1
−2
−3
OJOV01000072 TAG|q
OJOV01000072 TGA|w
GC skew
CGC skew
AT skew
CAT skew
0.4
0.2
0.0
−0.2
−0.4
60
50
40
30
20
10
0

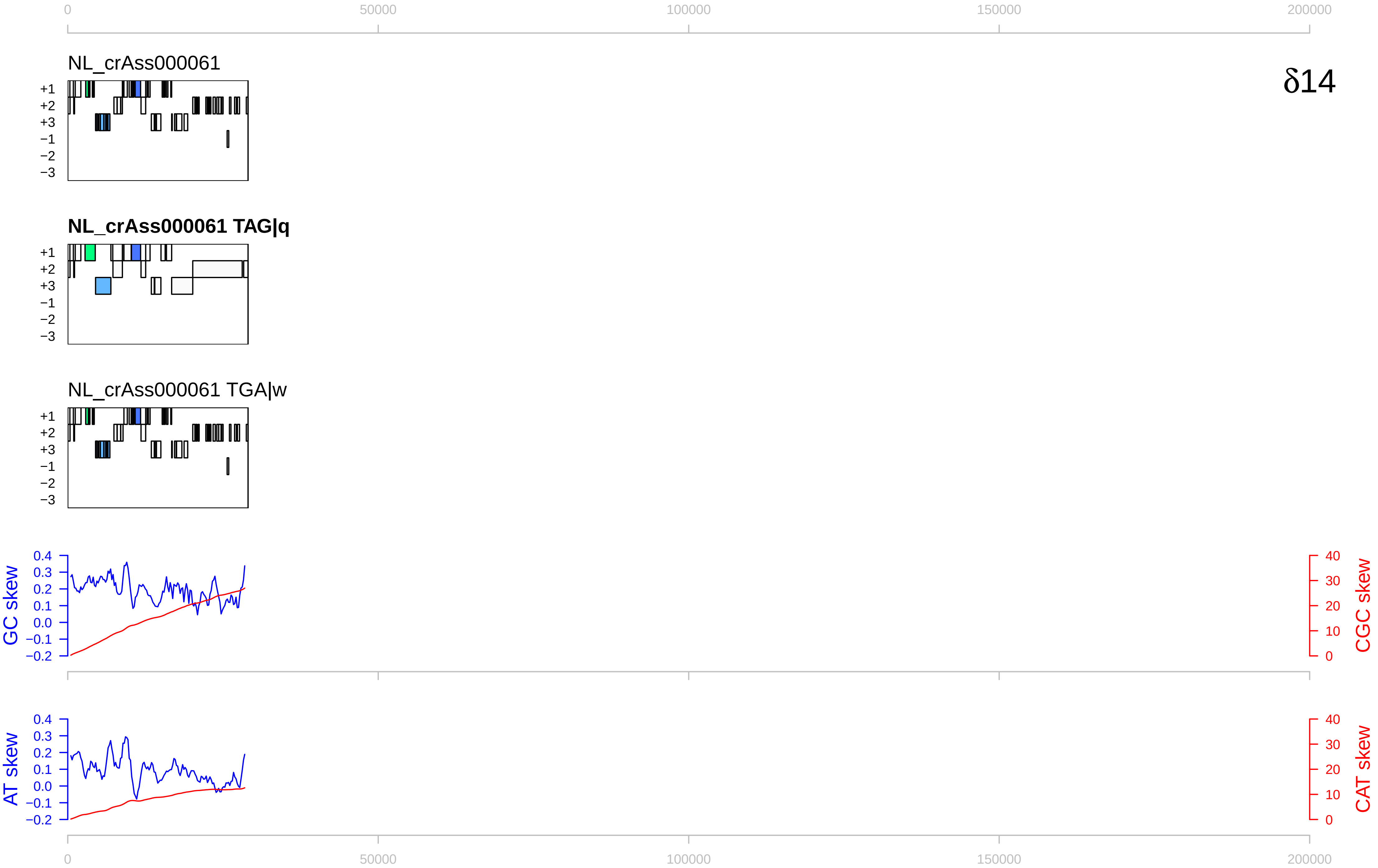

0
50000
100000
150000
200000
NL_crAss000061
δ14
+1
+2
+3
−1
−2
−3
NL_crAss000061 TAG|q
NL_crAss000061 TGA|w
GC skew
CGC skew
AT skew
CAT skew
0.4
0.3
0.2
0.1
0.0
−0.1
−0.2
40
30
20
10
0

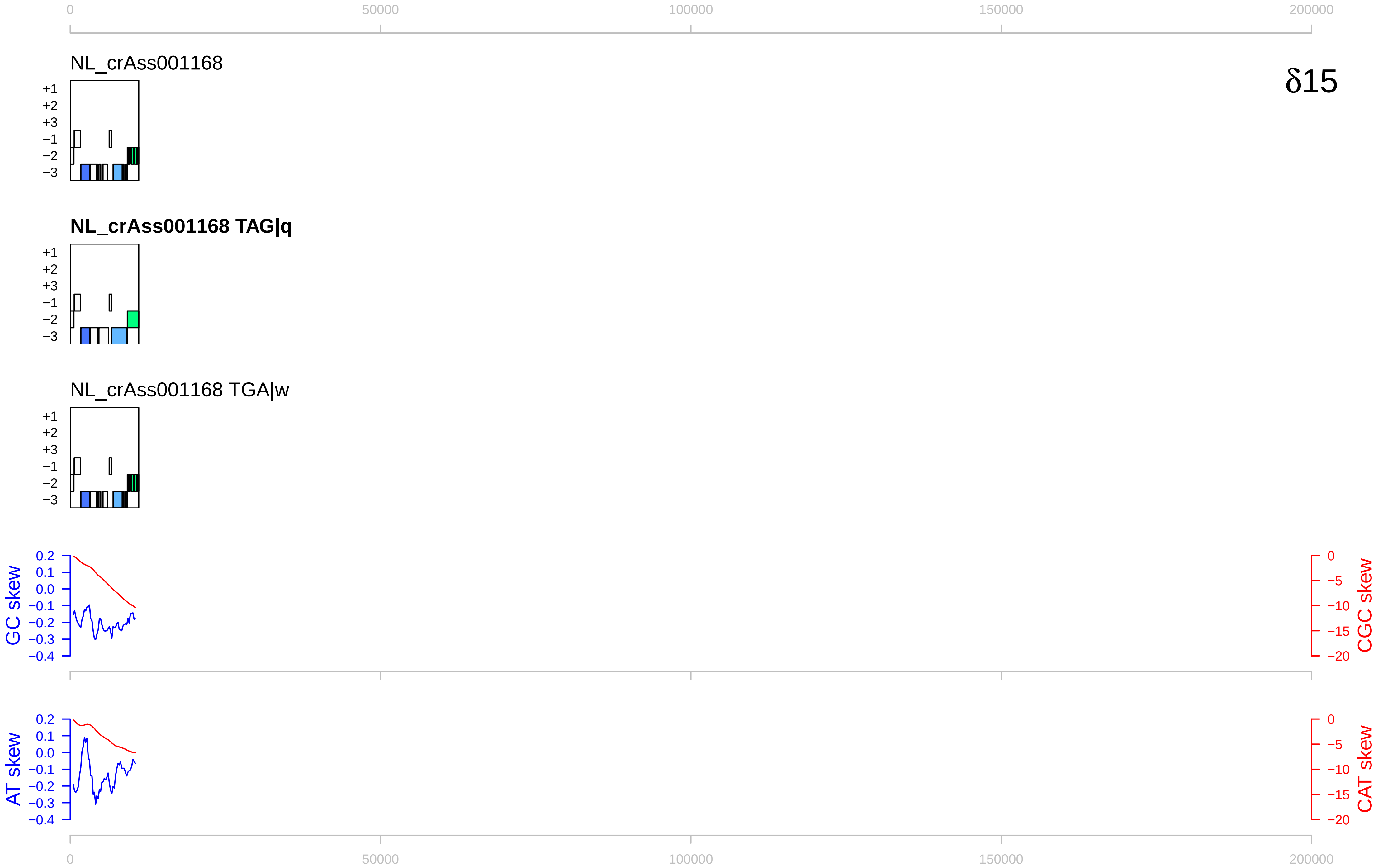

δ15
NL_crAss001168
NL_crAss001168 TAG|q
NL_crAss001168 TGA|w
+1
+2
+3
−1
−2
−3
GC skew
CGC skew
AT skew
CAT skew
0
50000
100000
150000
200000

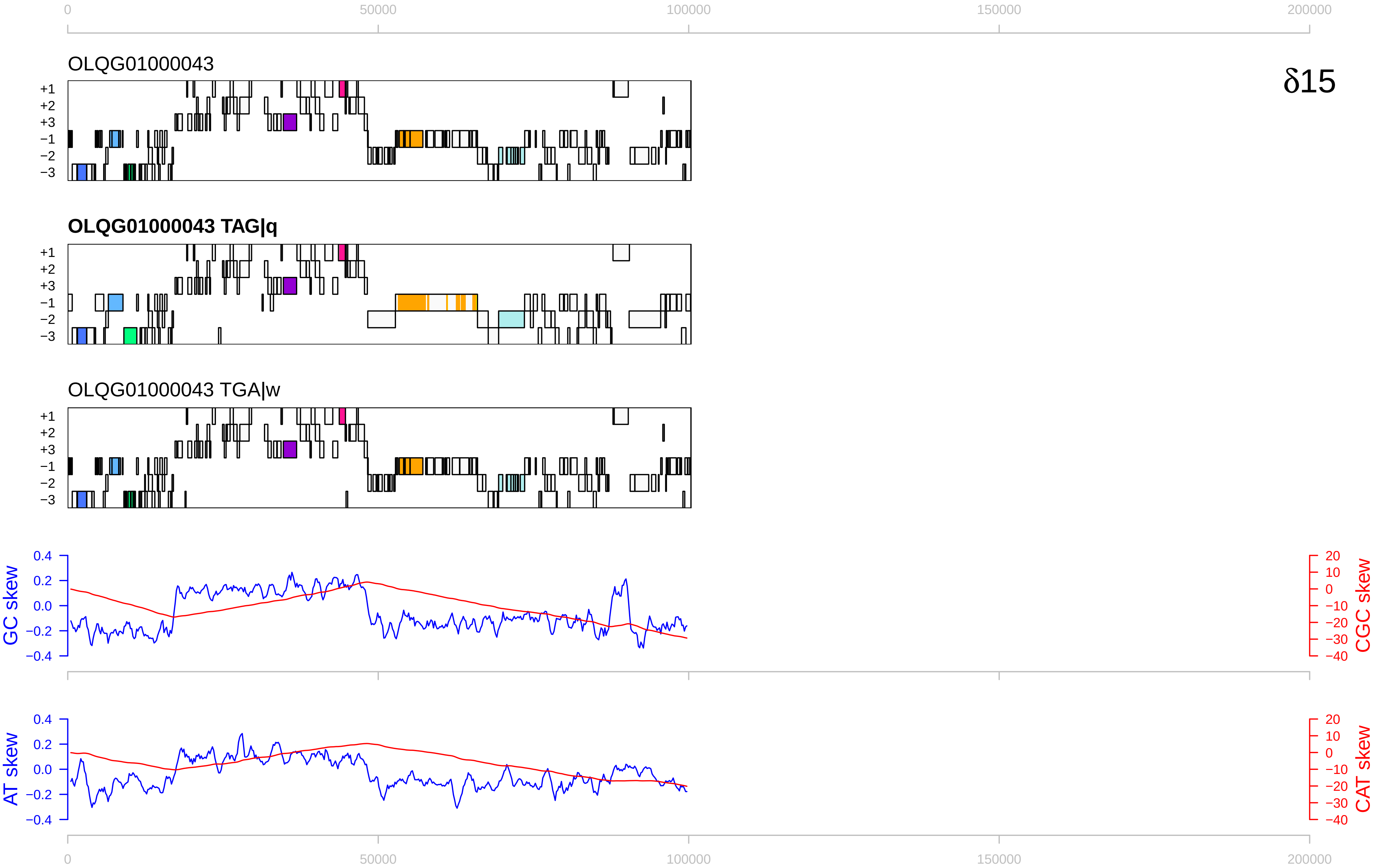

0
50000
100000
150000
200000
OLQG01000043
δ15
+1
+2
+3
−1
−2
−3
OLQG01000043 TAG|q
OLQG01000043 TGA|w
GC skew
CGC skew
AT skew
CAT skew
0.4
0.2
0.0
−0.2
−0.4
20
10
0
−10
−20
−30
−40

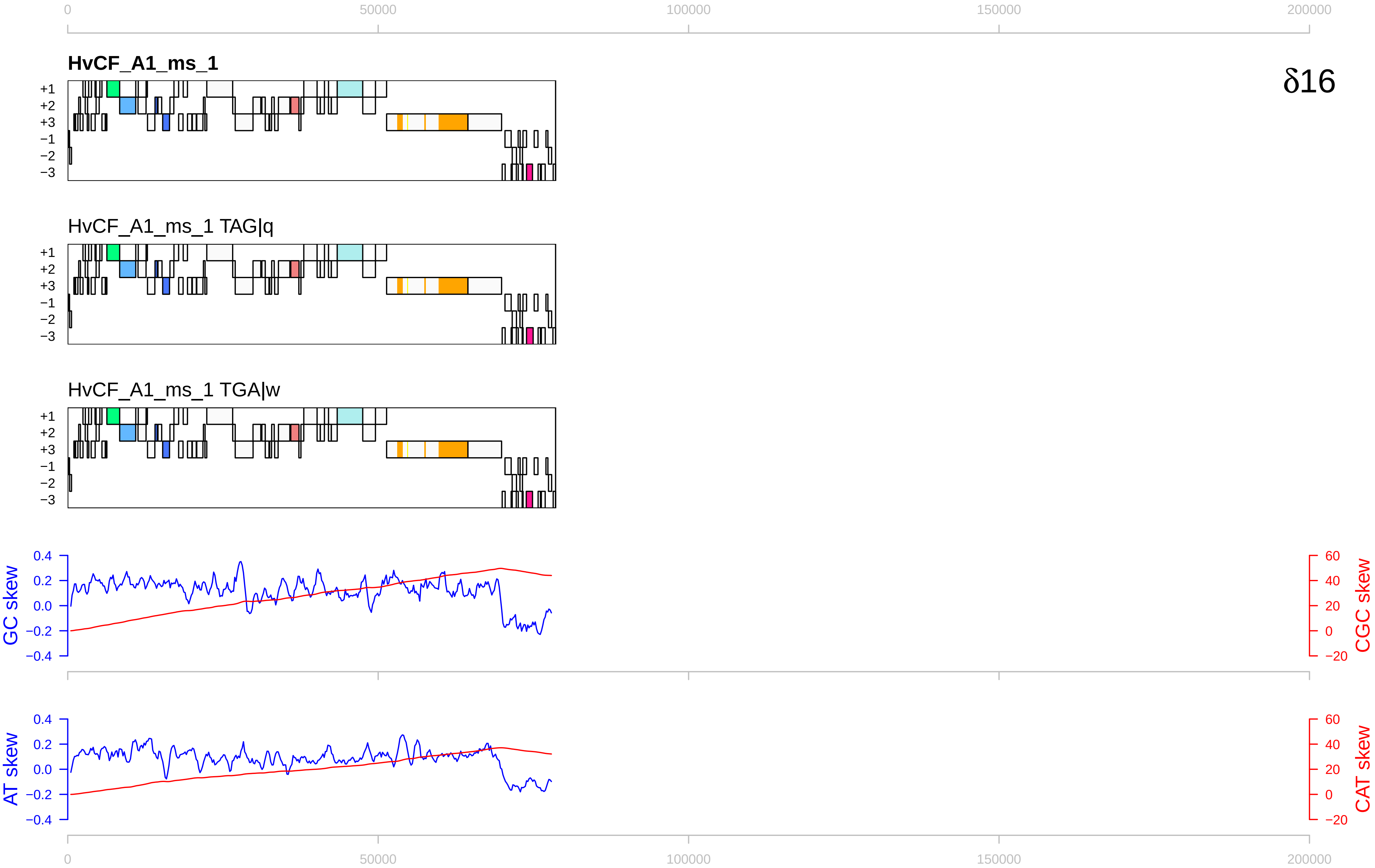

0
50000
100000
150000
200000
HvCF_A1_ms_1
δ16
+1
+2
+3
−1
−2
−3
HvCF_A1_ms_1 TAG|q
HvCF_A1_ms_1 TGA|w
GC skew
CGC skew
AT skew
CAT skew
0.4
0.2
0.0
−0.2
−0.4
60
40
20
0
−20

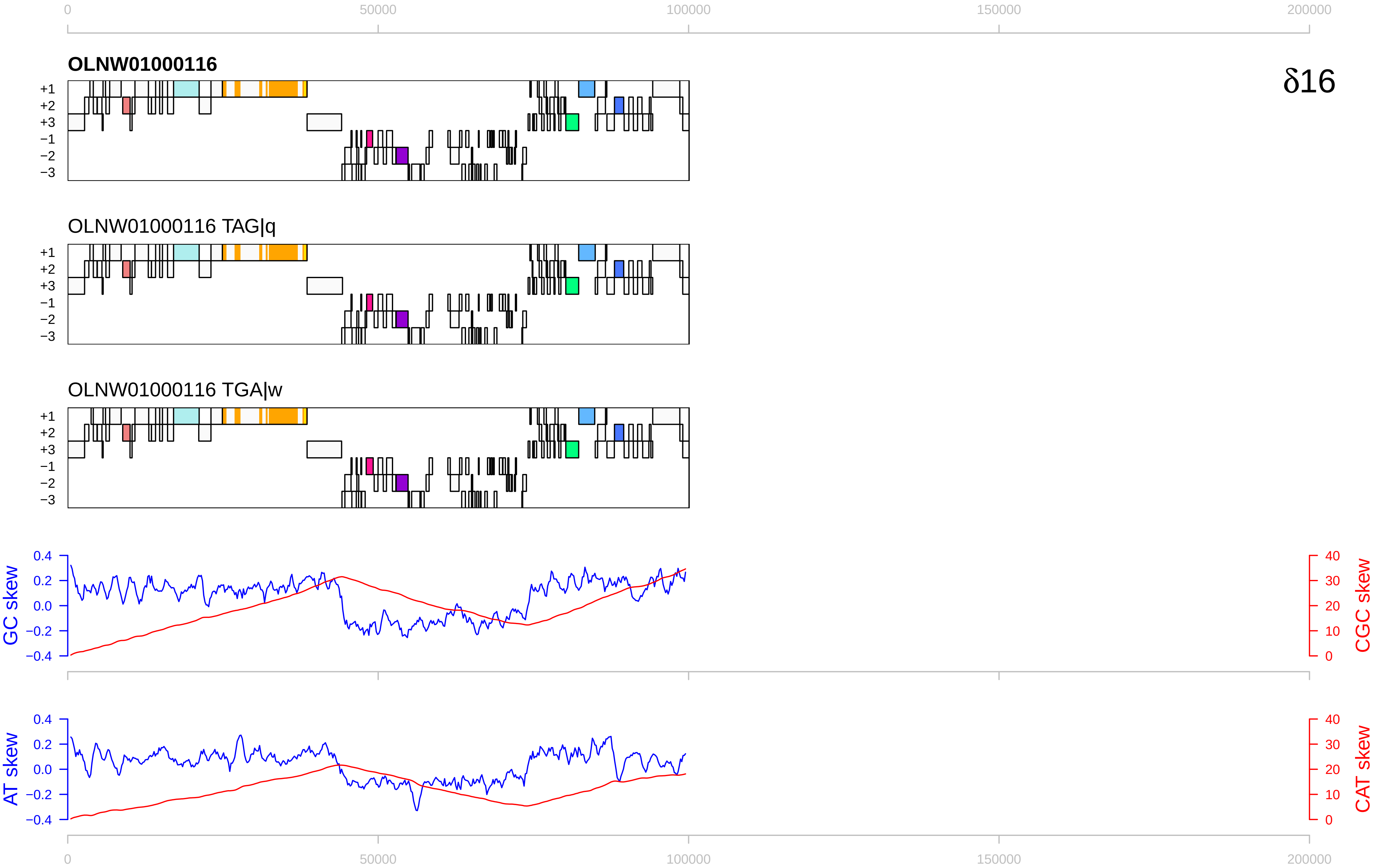

0
50000
100000
150000
200000
OLNW01000116
δ16
+1
+2
+3
−1
−2
−3
OLNW01000116 TAG|q
OLNW01000116 TGA|w
GC skew
CGC skew
AT skew
CAT skew
0.4
0.2
0.0
−0.2
−0.4
40
30
20
10

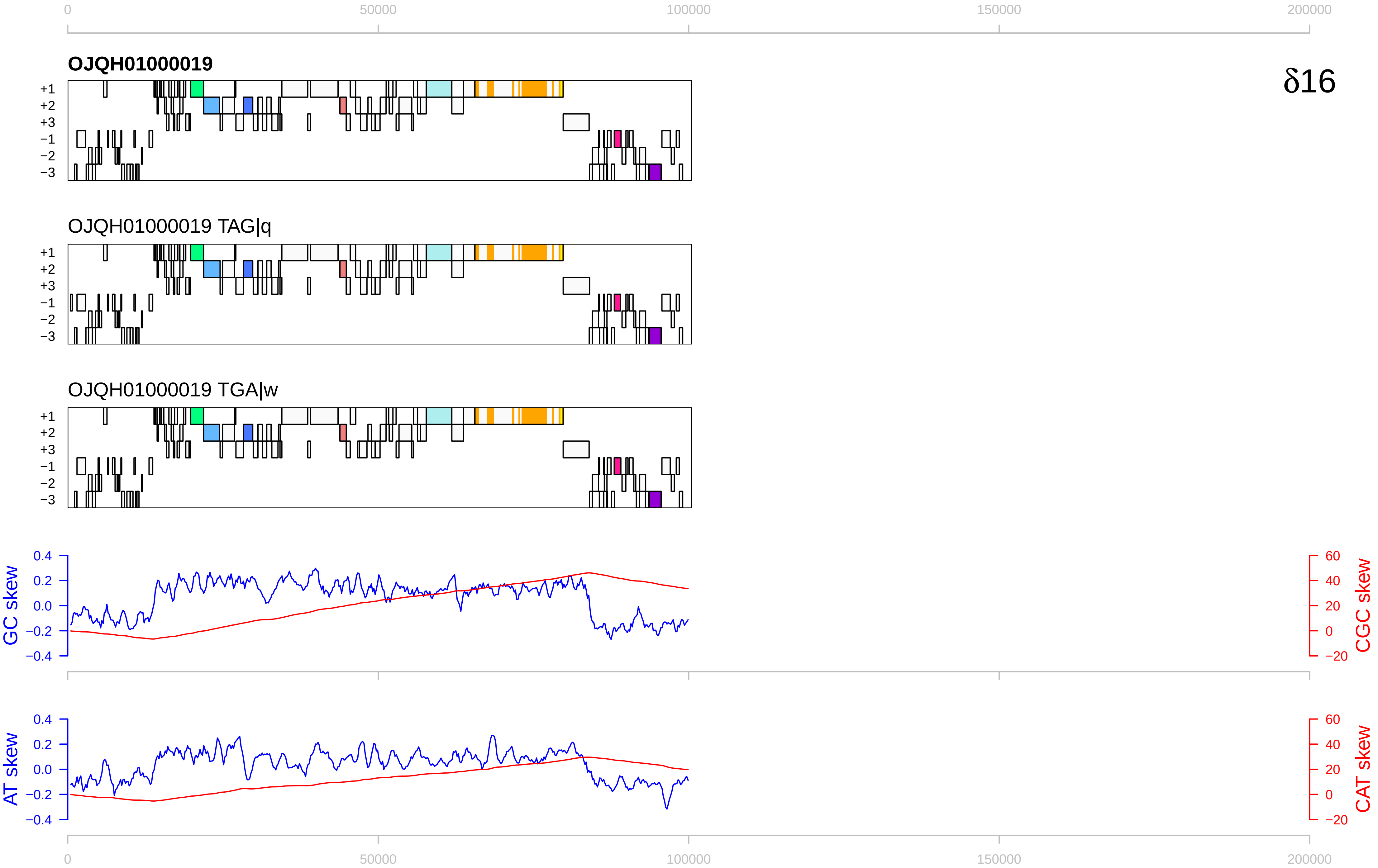

0
50000
100000
150000
200000
OJQH01000019
δ16
+1
+2
+3
−1
−2
−3
OJQH01000019 TAG|q
OJQH01000019 TGA|w
GC skew
CGC skew
0.4
0.2
0.0
−0.2
−0.4
60
40
20
0
−20
AT skew
CAT skew

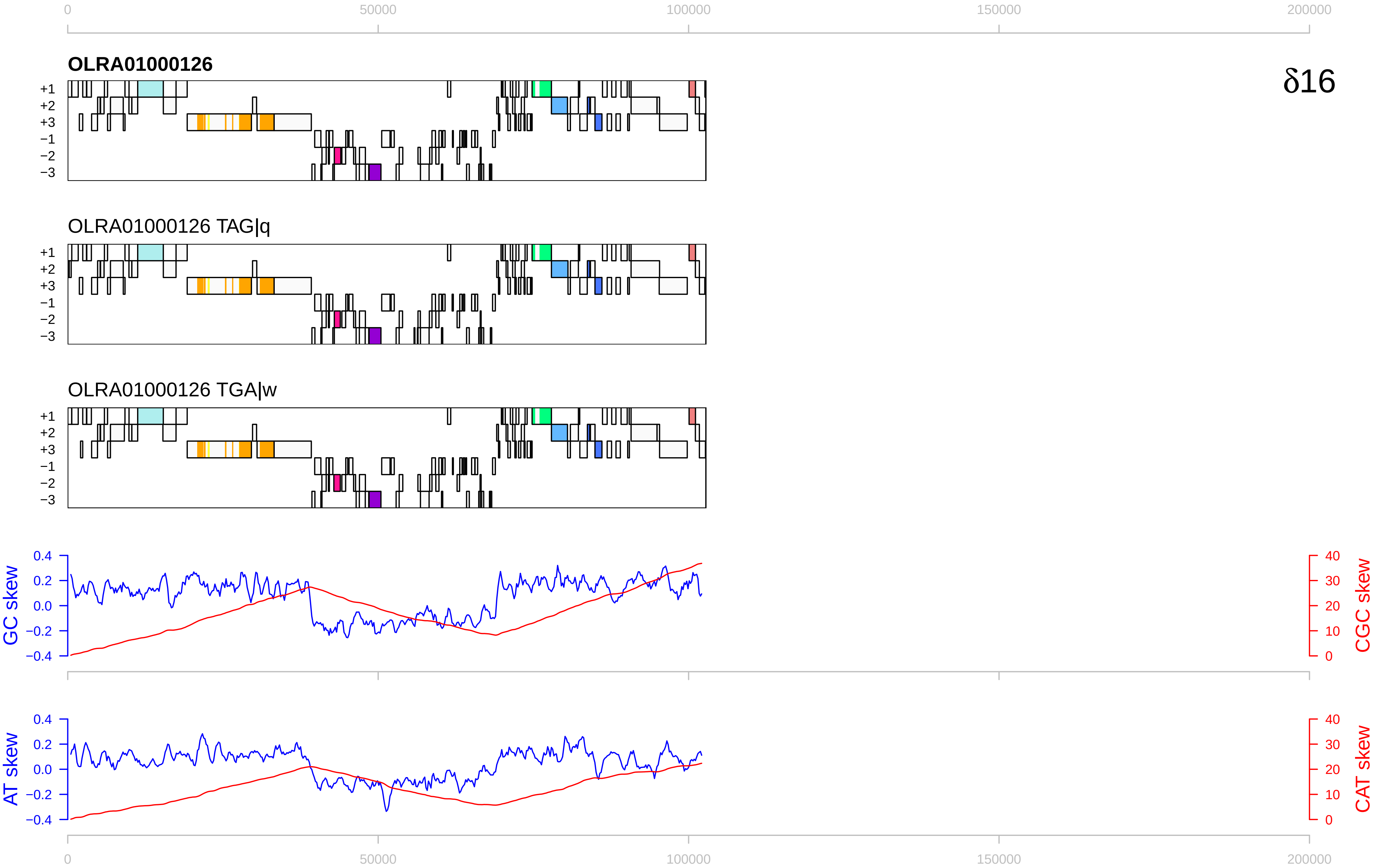

0
50000
100000
150000
200000
OLRA01000126
δ16
+1
+2
+3
−1
−2
−3
OLRA01000126 TAG|q
OLRA01000126 TGA|w
GC skew
CGC skew
0.4
0.2
0.0
−0.2
−0.4
40
30
20
10
0
AT skew
CAT skew

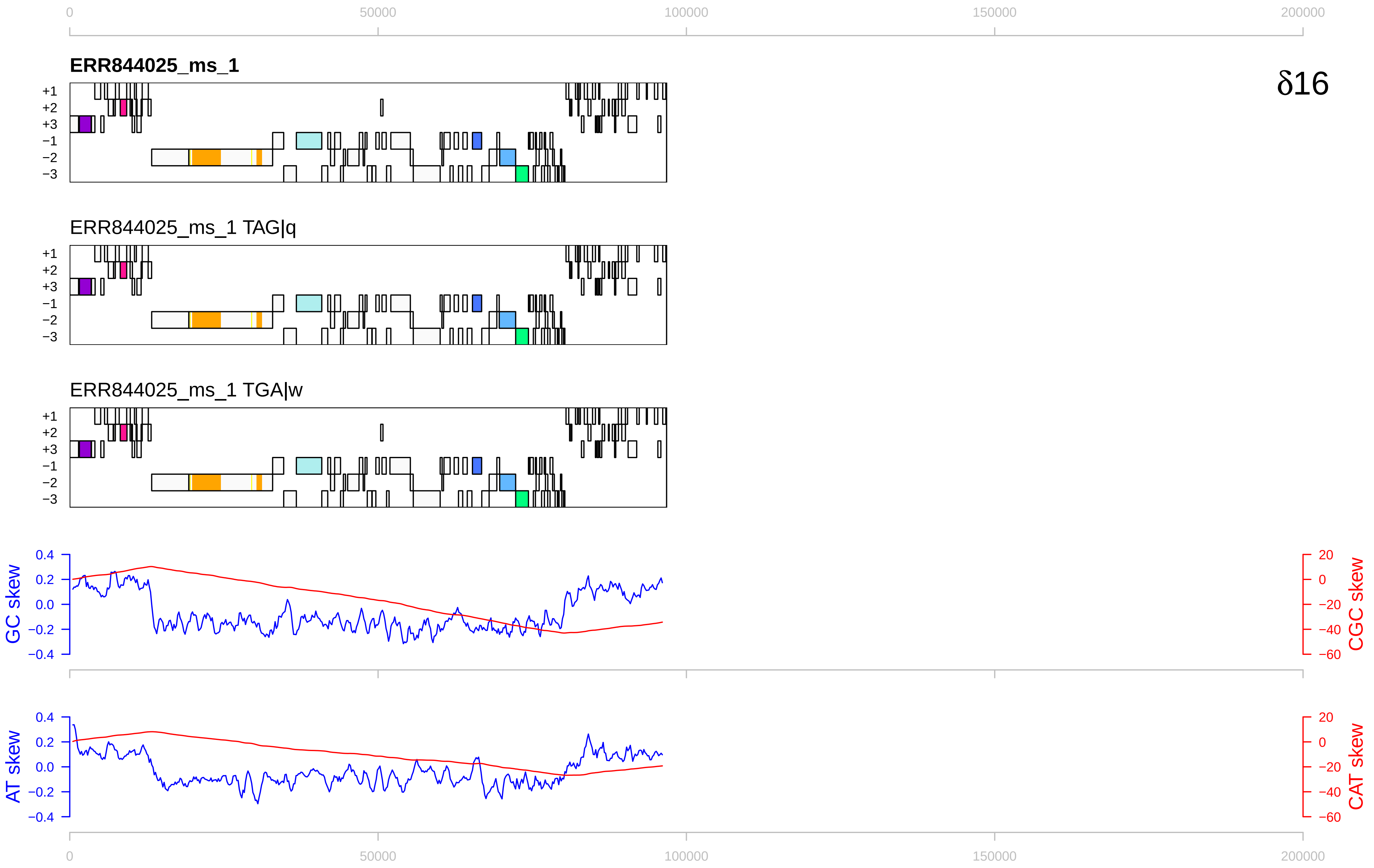

δ16
ERR844025_ms_1
ERR844025_ms_1 TAG|q
ERR844025_ms_1 TGA|w
+1
+2
+3
−1
−2
−3
GC skew
CGC skew
AT skew
CAT skew
0
50000
100000
150000
200000
0.4
0.2
0.0
−0.2
−0.4
20
0
−20
−40
−60

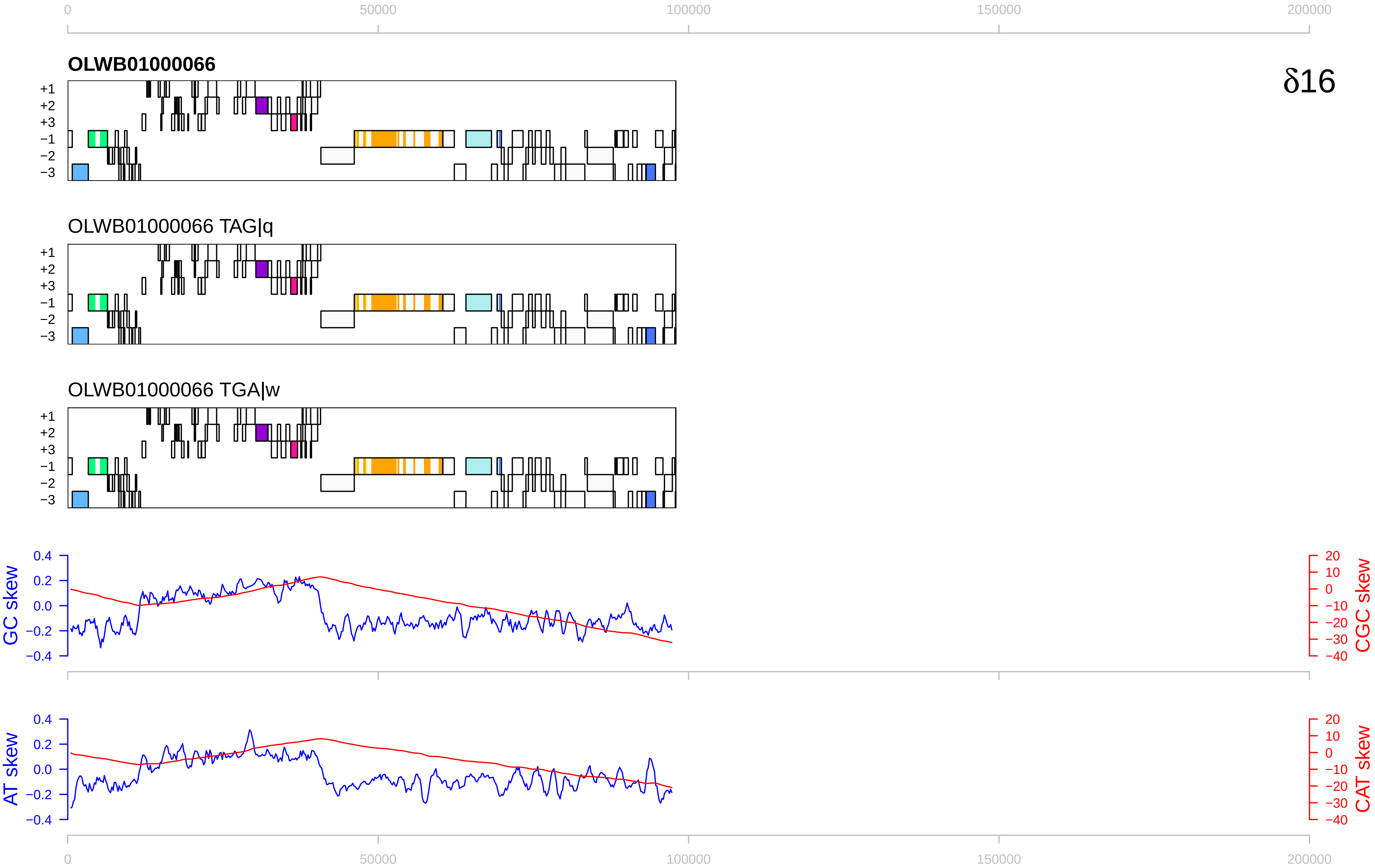

0
50000
100000
150000
200000
OLWB01000066
δ16
+1
+2
+3
−1
−2
−3
OLWB01000066 TAG|q
OLWB01000066 TGA|w
GC skew
CGC skew
AT skew
CAT skew
0.4
0.2
0.0
−0.2
−0.4
20
10
0
−10
−20
−30
−40

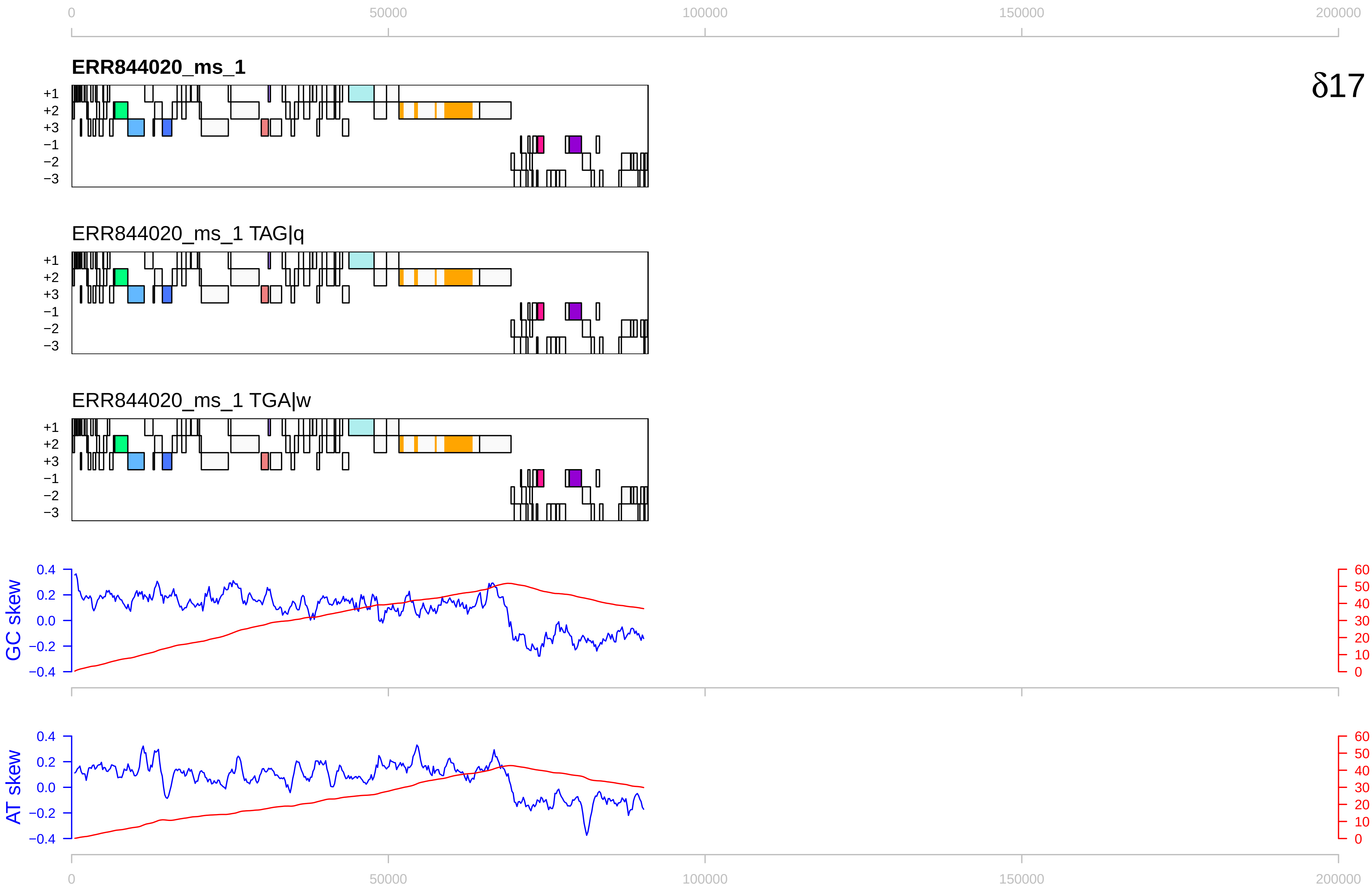

δ17
ERR844020_ms_1
ERR844020_ms_1 TAG|q
ERR844020_ms_1 TGA|w
+1 +2 +3 −1 −2 −3
GC skew
CGC skew
AT skew
CAT skew
0 50000 100000 150000 200000
0.4 0.2 0.0 −0.2 −0.4
60 50 40 30 20 10 0

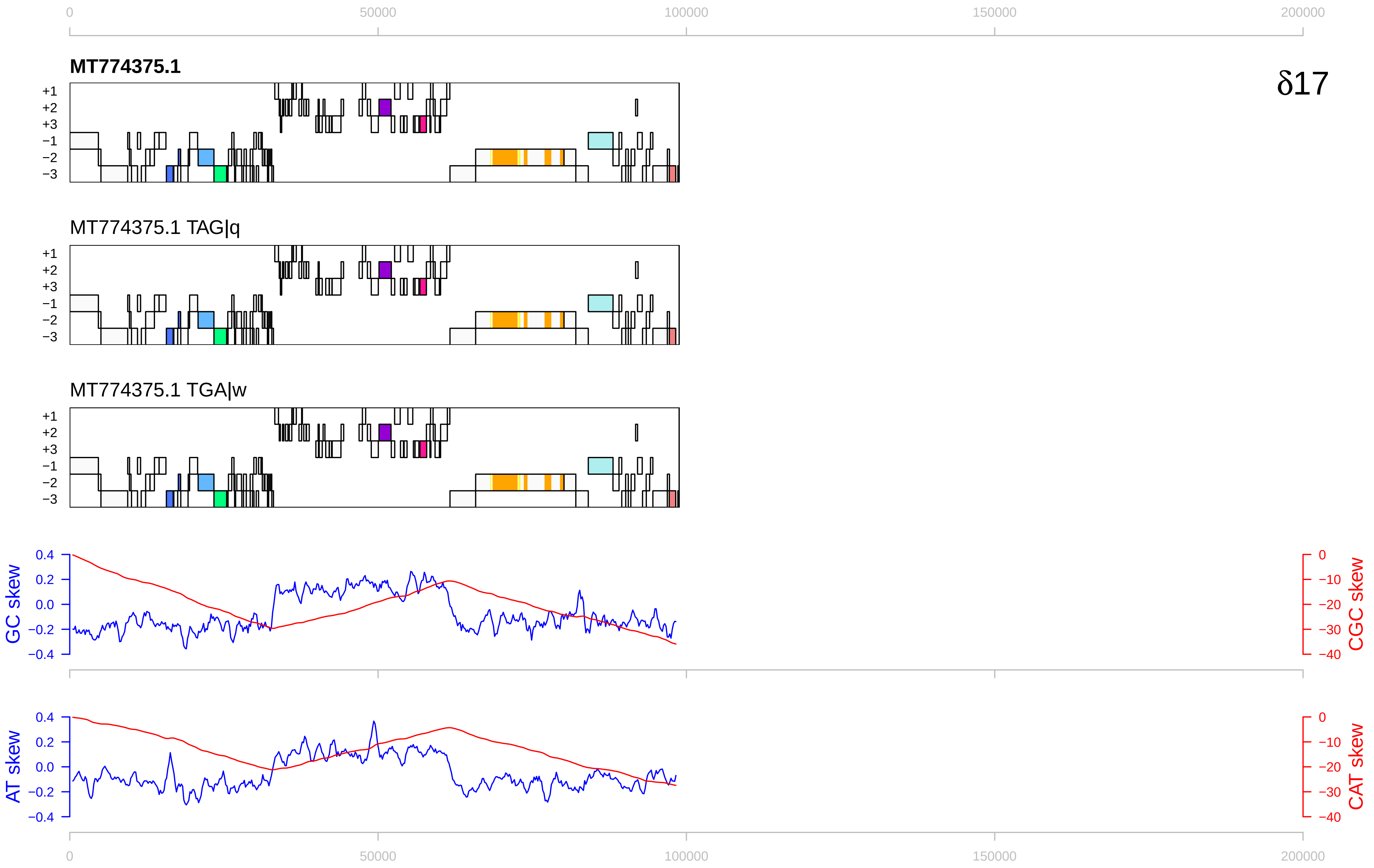

0
50000
100000
150000
200000
MT774375.1
δ17
+1
+2
+3
−1
−2
−3
MT774375.1 TAG|q
MT774375.1 TGA|w
GC skew
CGC skew
AT skew
CAT skew
0.4
0.2
0.0
−0.2
−0.4
0
−10
−20
−30
−40

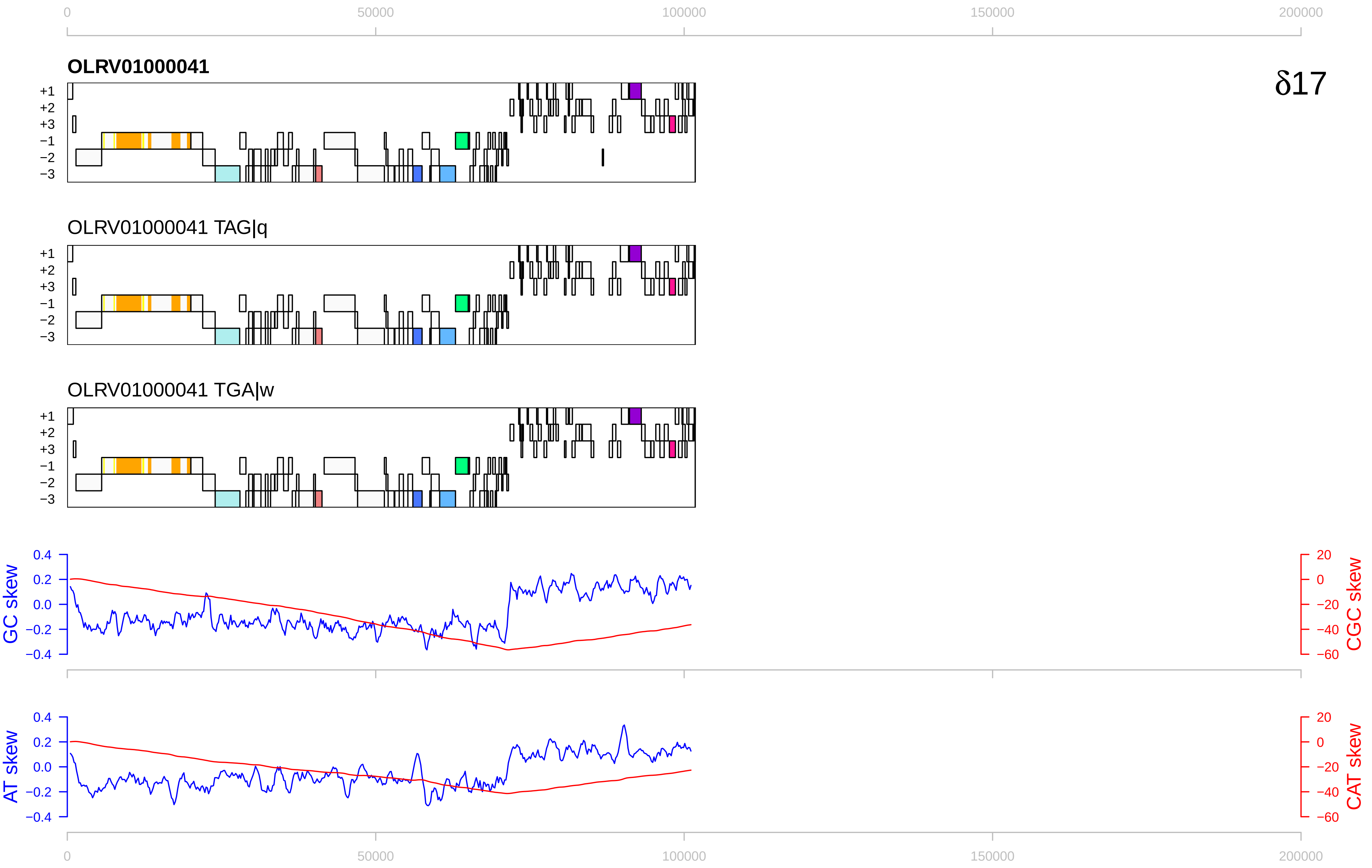

0
50000
100000
150000
200000
OLRV01000041
δ17
+1
+2
+3
−1
−2
−3
OLRV01000041 TAG|q
OLRV01000041 TGA|w
GC skew
CGC skew
AT skew
CAT skew
0.4
0.2
0.0
−0.2
−0.4
20
0
−20
−40
−60

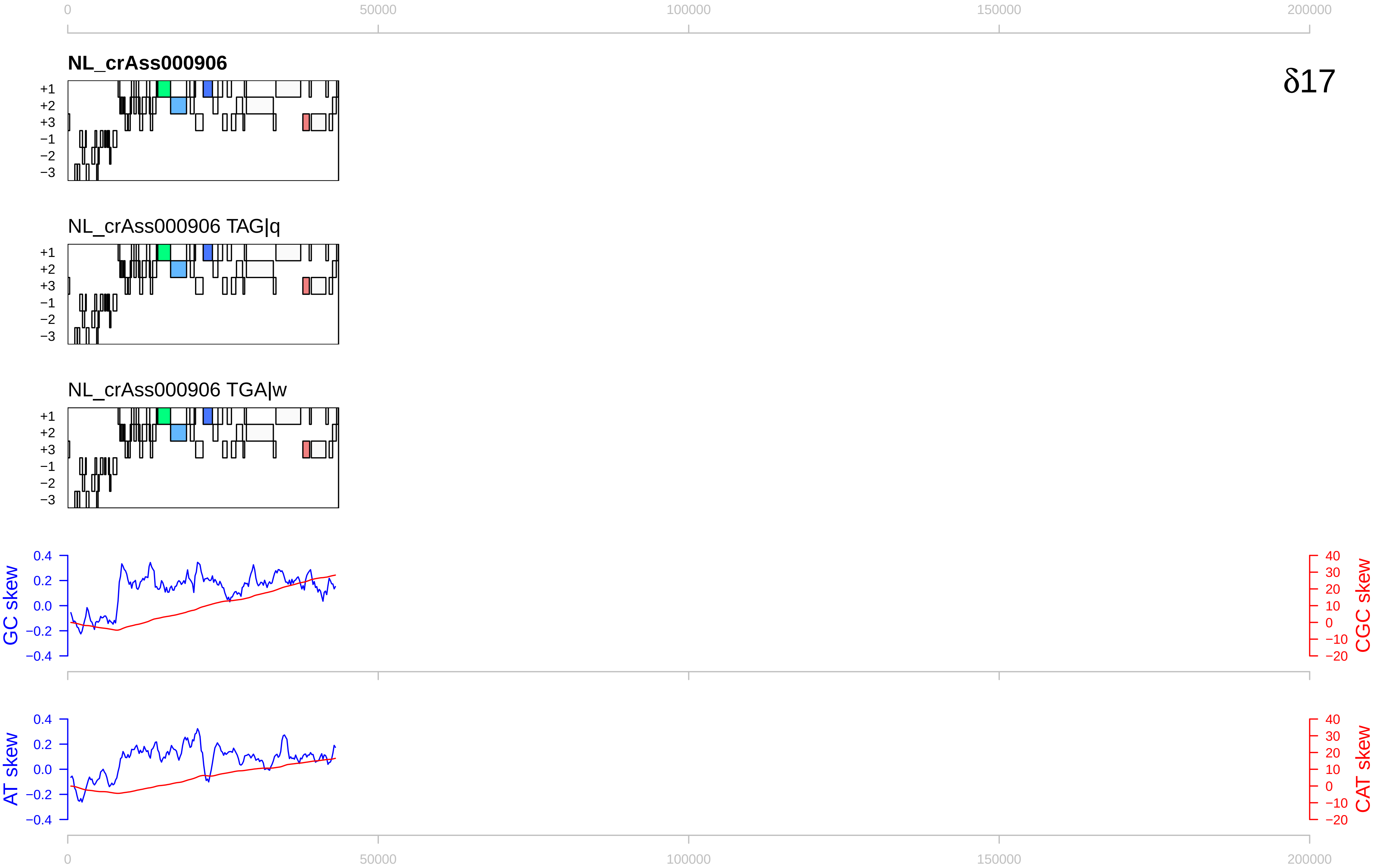

0
50000
100000
150000
200000
NL_crAss000906
δ17
+1
+2
+3
−1
−2
−3
NL_crAss000906 TAG|q
NL_crAss000906 TGA|w
GC skew
CGC skew
AT skew
CAT skew
0.4
0.2
0.0
−0.2
−0.4
40
30
20
10
0
−10
−20

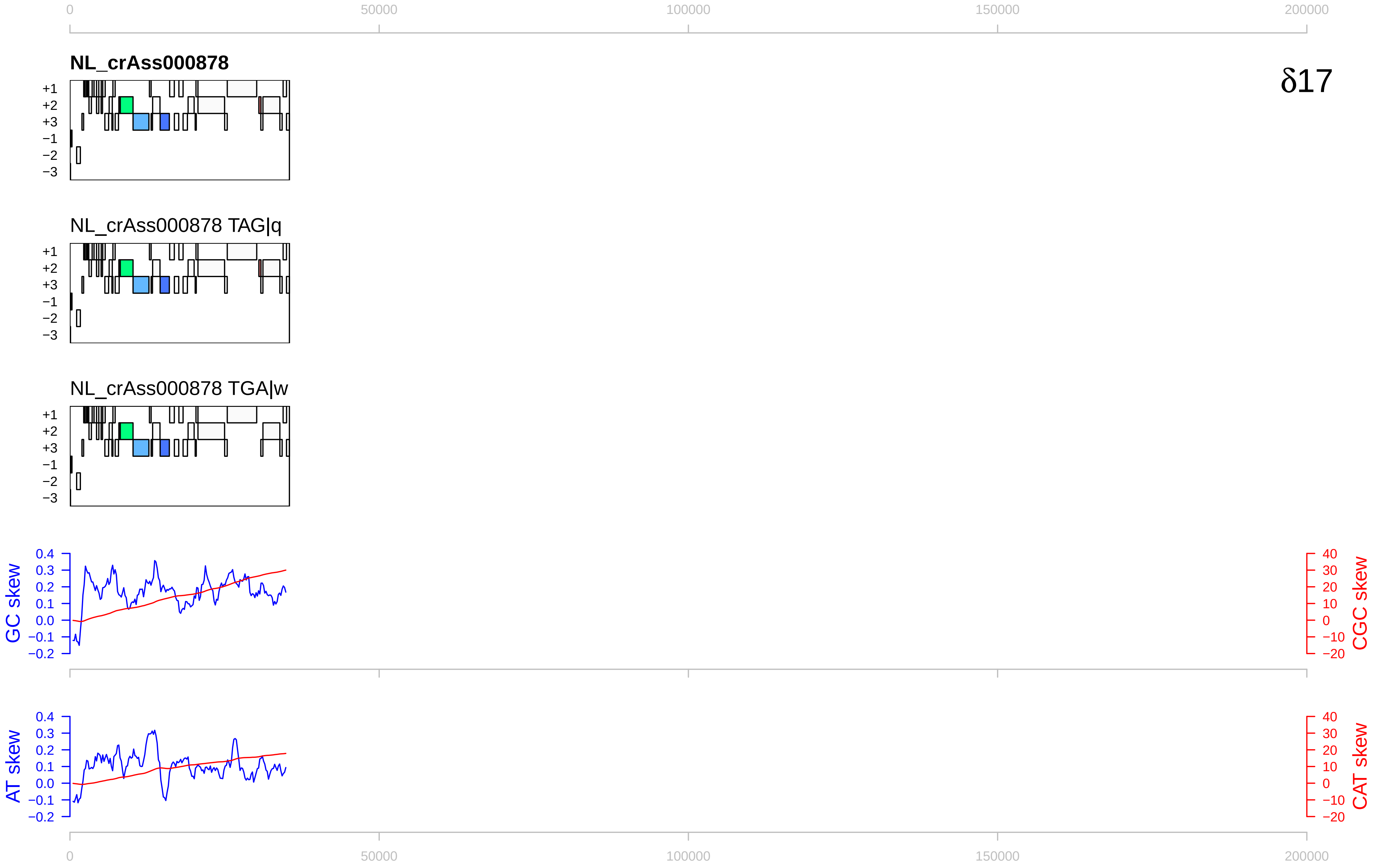

0
50000
100000
150000
200000
NL_crAss000878
δ17
+1
+2
+3
−1
−2
−3
NL_crAss000878 TAG|q
NL_crAss000878 TGA|w
GC skew
CGC skew
AT skew
CAT skew
0.4
0.3
0.2
0.1
0.0
−0.1
−0.2
40
30
20
10
0
−10
−20

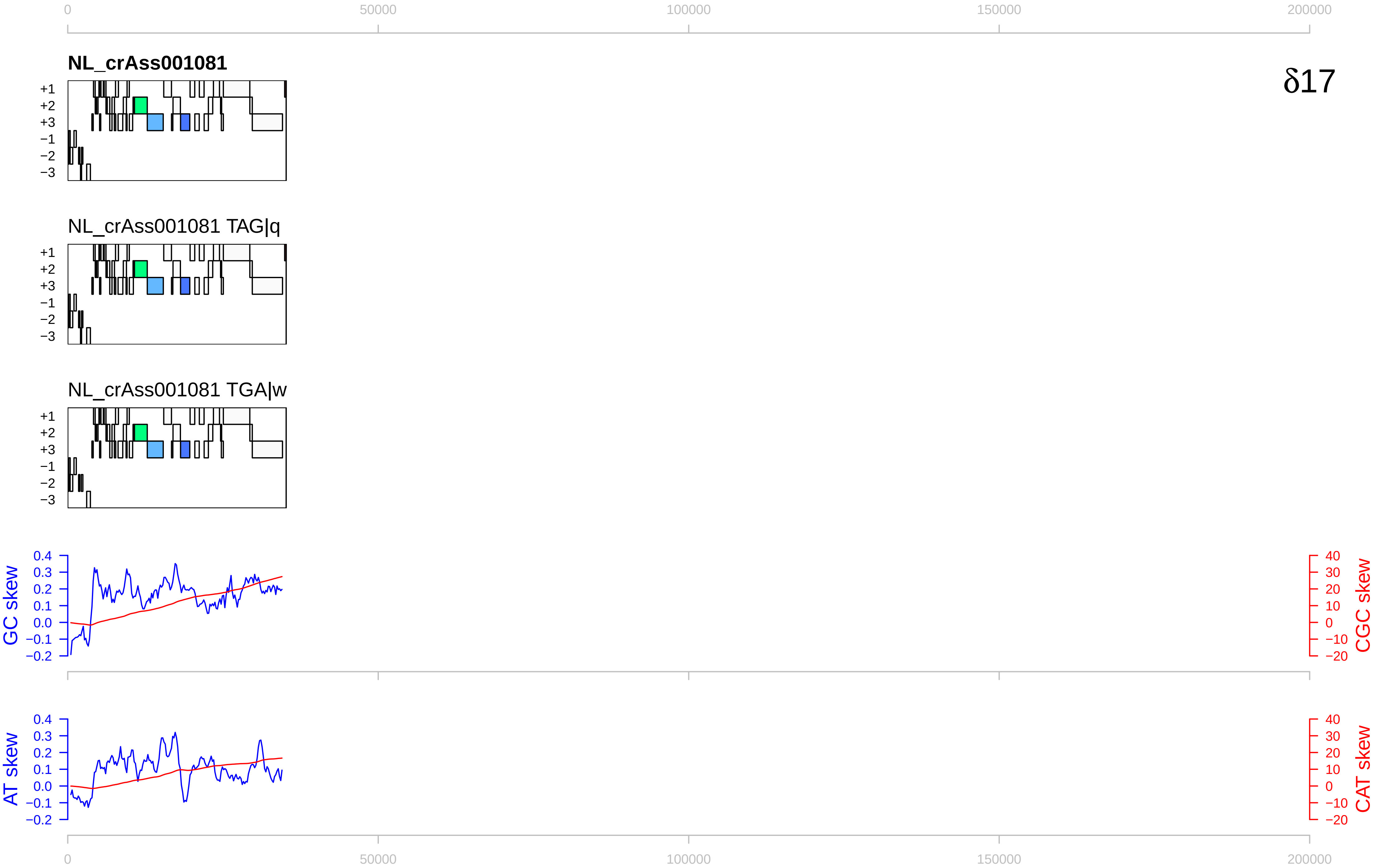

0
50000
100000
150000
200000
NL_crAss001081
δ17
+1
+2
+3
−1
−2
−3
NL_crAss001081 TAG|q
NL_crAss001081 TGA|w
GC skew
CGC skew
0.4
0.3
0.2
0.1
0.0
−0.1
−0.2
40
30
20
10
0
−10
−20
AT skew
CAT skew

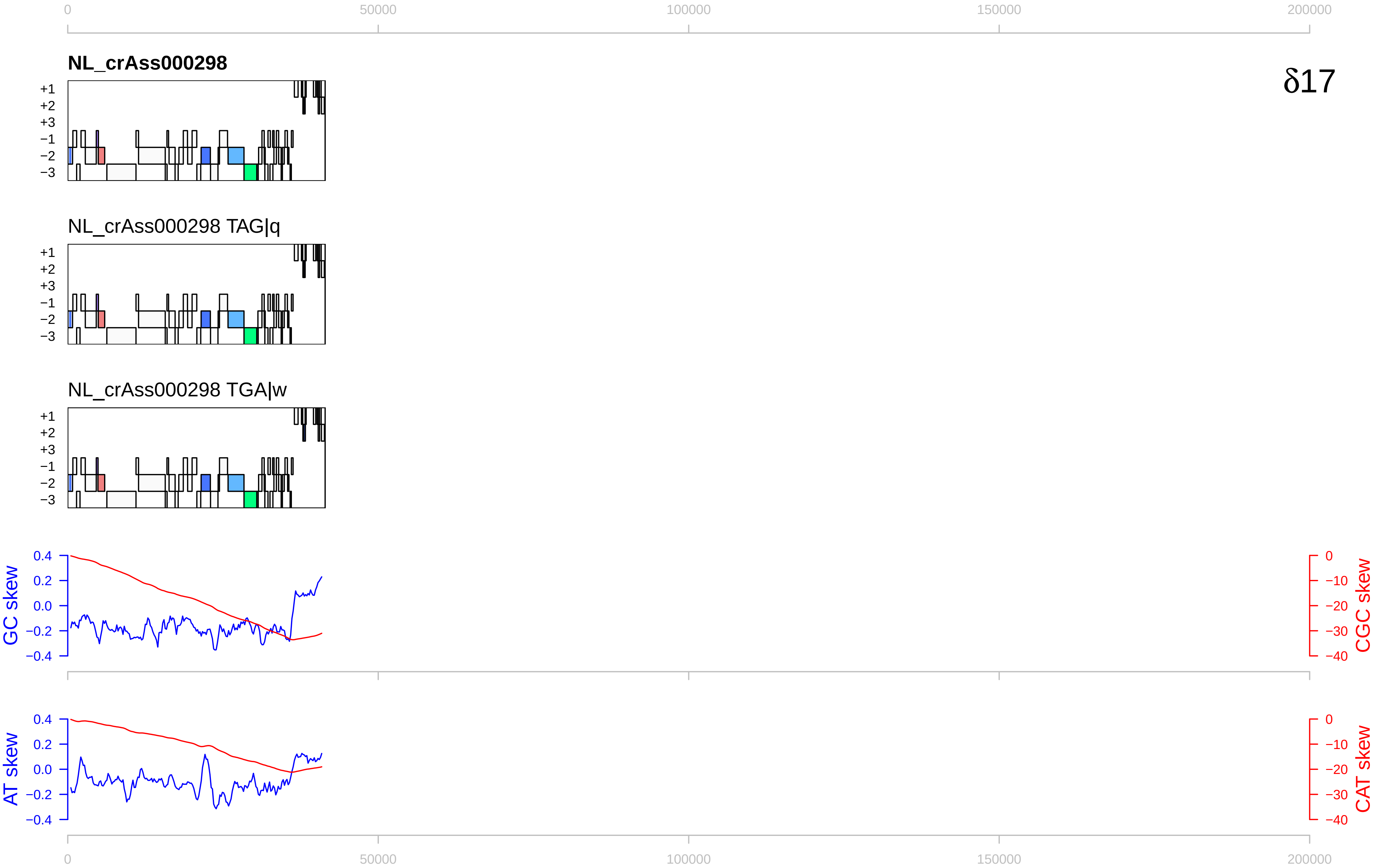

δ17
NL_crAss000298
NL_crAss000298 TAG|q
NL_crAss000298 TGA|w
+1
+2
+3
−1
−2
−3
GC skew
CGC skew
AT skew
CAT skew
0
50000
100000
150000
200000

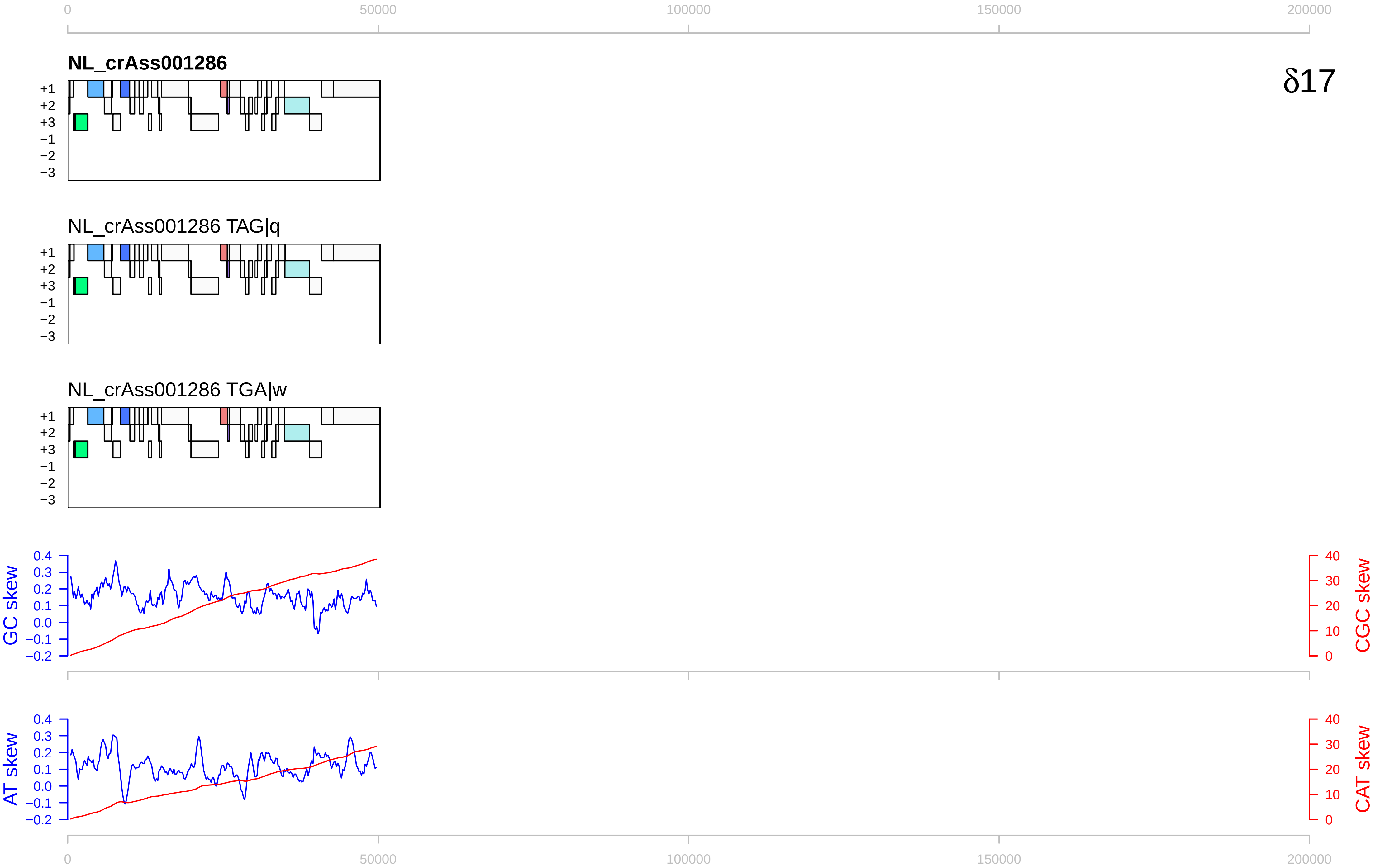

0
50000
100000
150000
200000
NL_crAss001286
δ17
+1
+2
+3
−1
−2
−3
NL_crAss001286 TAG|q
NL_crAss001286 TGA|w
GC skew
CGC skew
0.4
0.3
0.2
0.1
0.0
−0.1
−0.2
40
30
20
10
0
AT skew
CAT skew

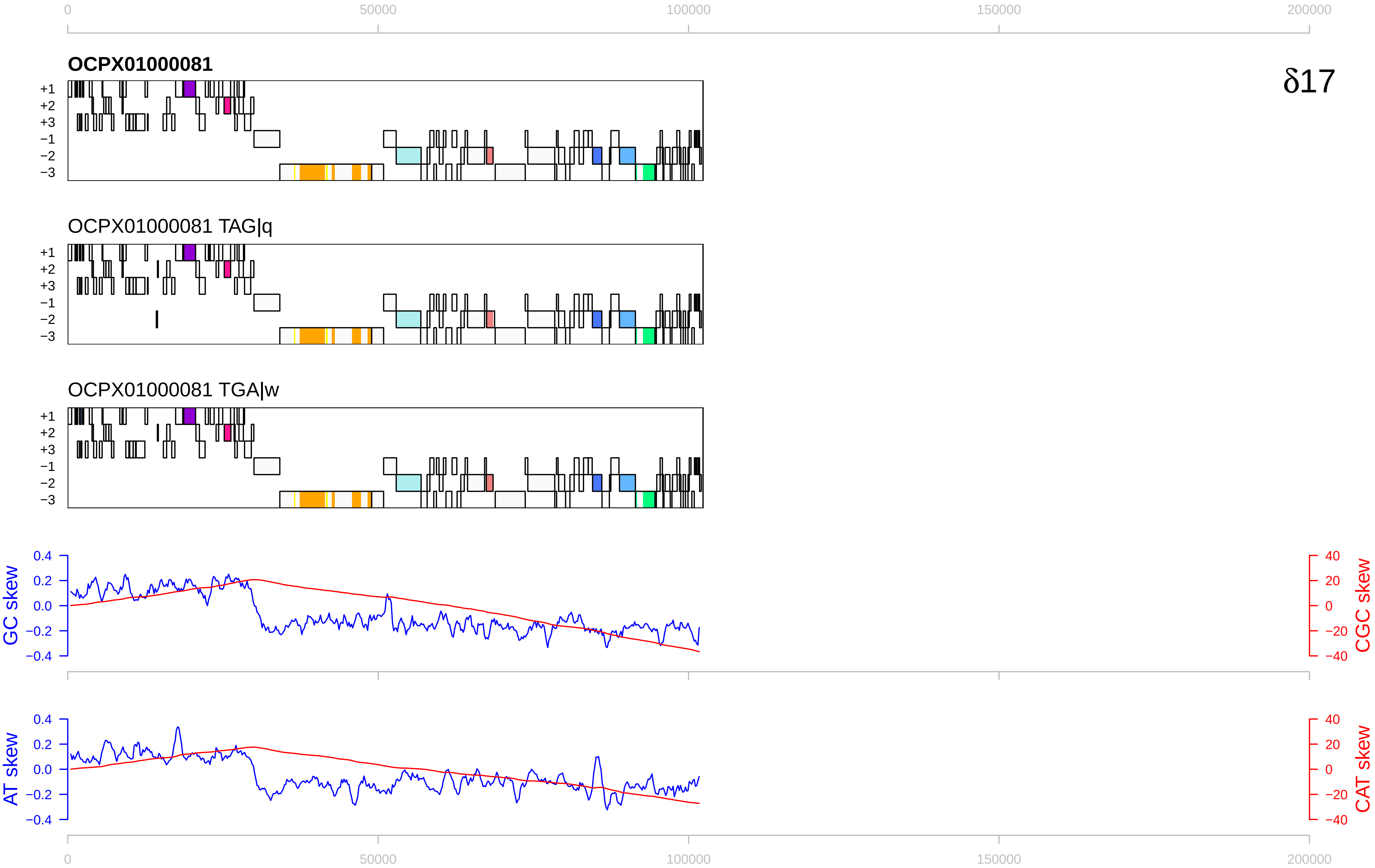

0
50000
100000
150000
200000
OCPX01000081
δ17
+1
+2
+3
−1
−2
−3
OCPX01000081 TAG|q
OCPX01000081 TGA|w
GC skew
CGC skew
AT skew
CAT skew
0.4
0.2
0.0
−0.2
−0.4
40
20
0
−20
−40

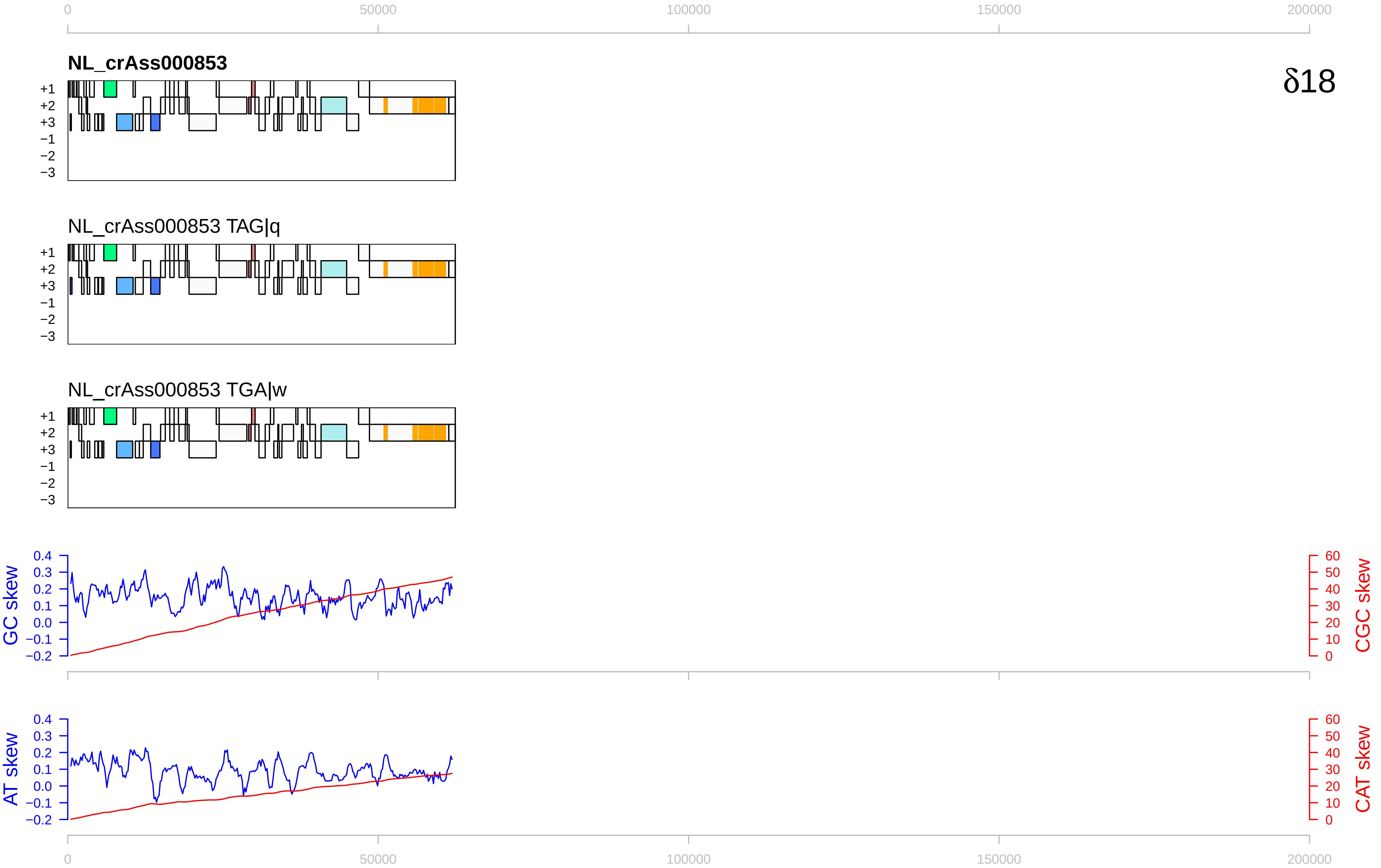

0
50000
100000
150000
200000
NL_crAss000853
δ18
+1
+2
+3
−1
−2
−3
NL_crAss000853 TAG|q
NL_crAss000853 TGA|w
GC skew
CGC skew
AT skew
CAT skew
0.4
0.3
0.2
0.1
0.0
−0.1
−0.2
60
50
40
30
20
10

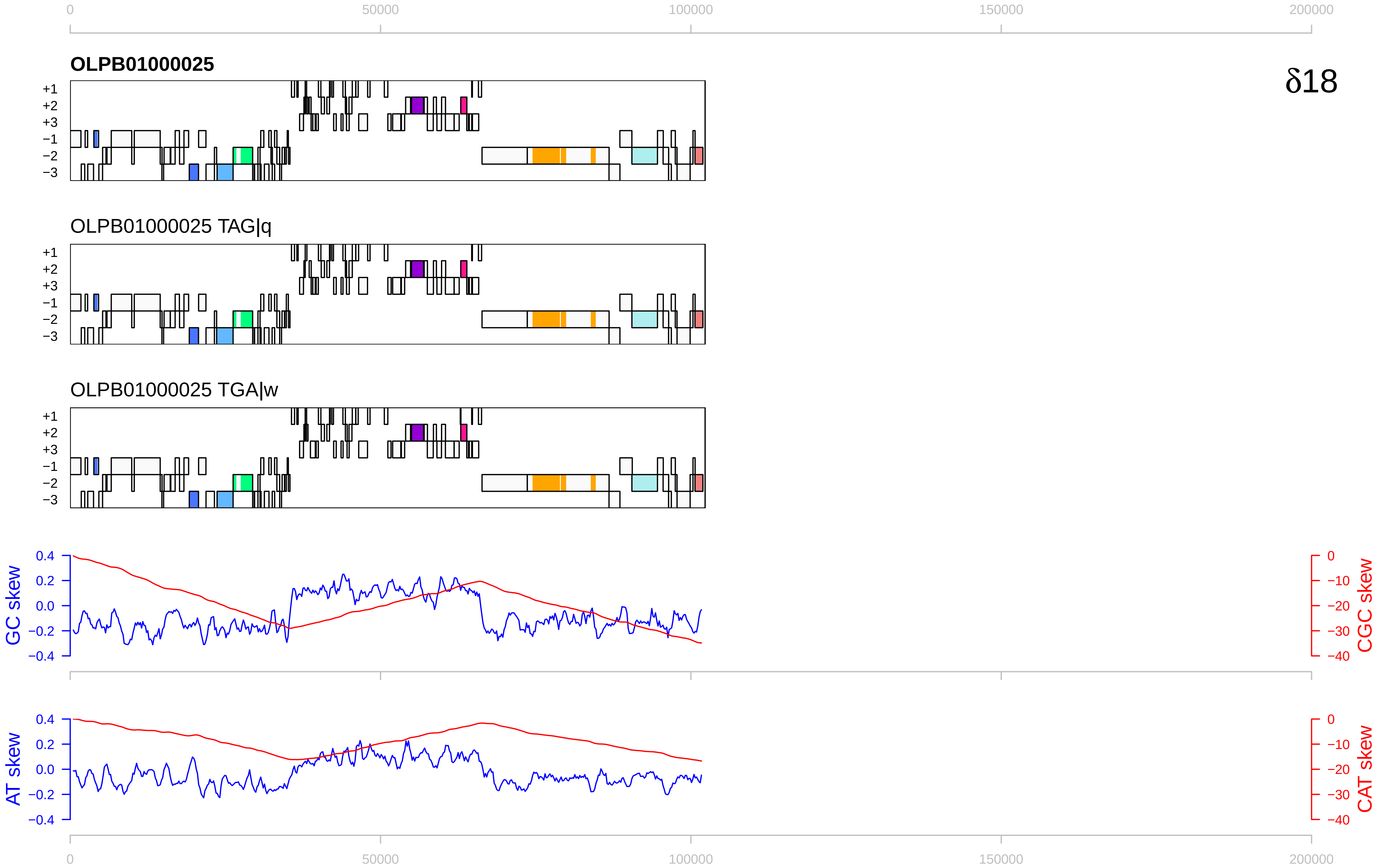

δ18
OLPB01000025
OLPB01000025 TAG|q
OLPB01000025 TGA|w
+1
+2
+3
−1
−2
−3
GC skew
CGC skew
AT skew
CAT skew
0
50000
100000
150000
200000

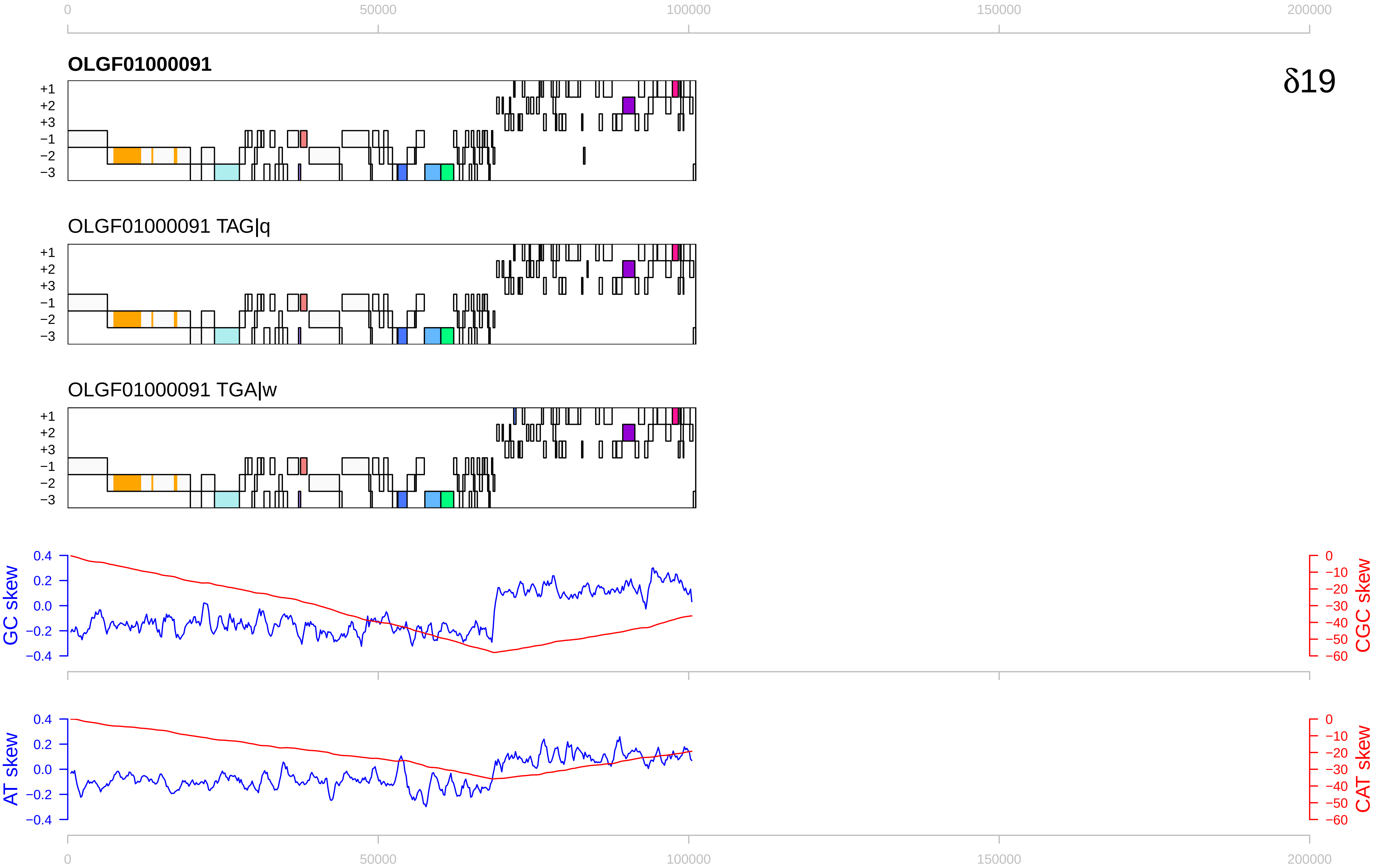

0
50000
100000
150000
200000
OLGF01000091
δ19
+1
+2
+3
−1
−2
−3
OLGF01000091 TAG|q
OLGF01000091 TGA|w
GC skew
CGC skew
0.4
0.2
0.0
−0.2
−0.4
0
−10
−20
−30
−40
−50
−60
AT skew
CAT skew

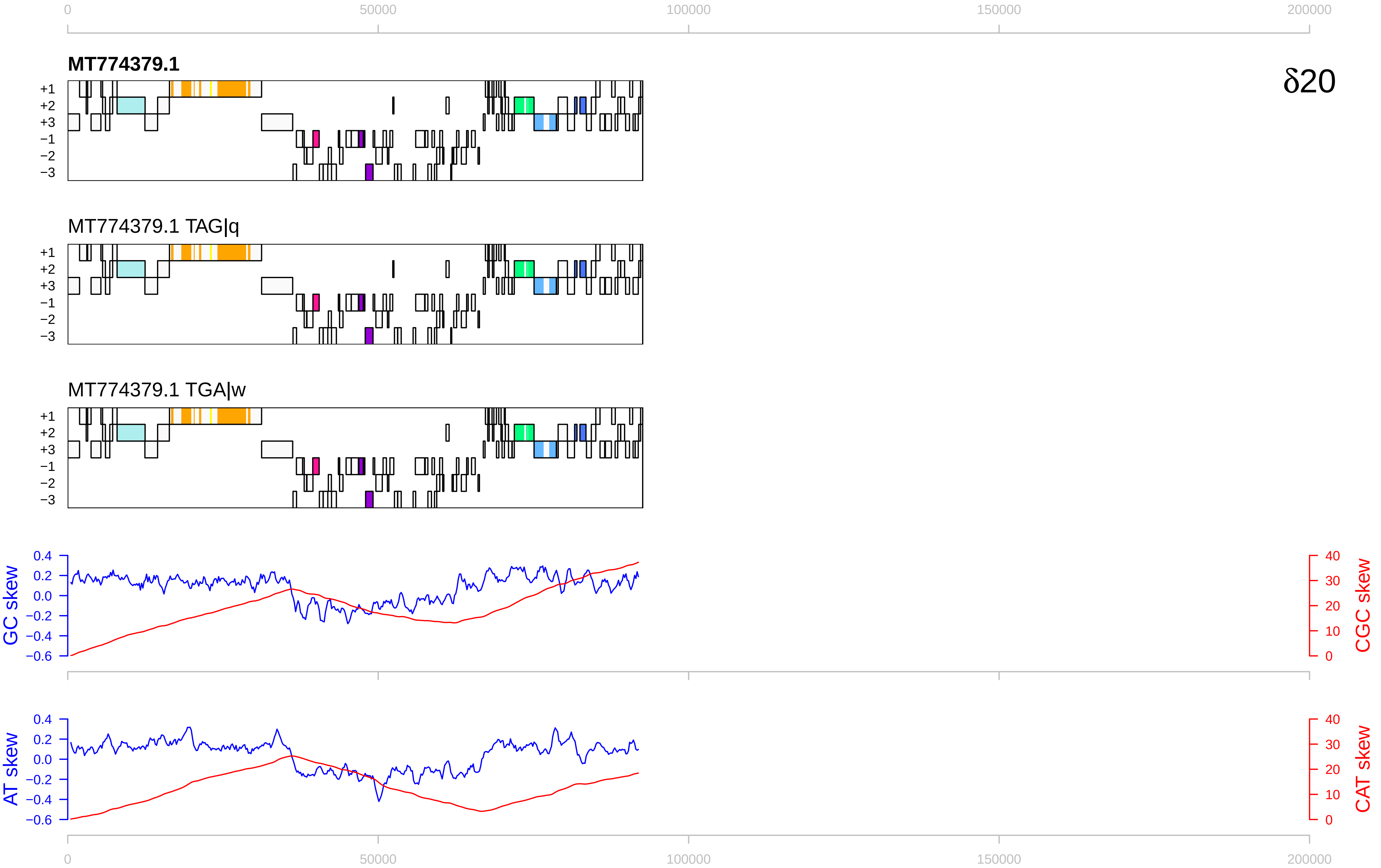

0
50000
100000
150000
200000
MT774379.1
δ20
+1
+2
+3
−1
−2
−3
MT774379.1 TAG|q
MT774379.1 TGA|w
GC skew
CGC skew
AT skew
CAT skew
0.4
0.2
0.0
−0.2
−0.4
−0.6
40
30
20
10
0

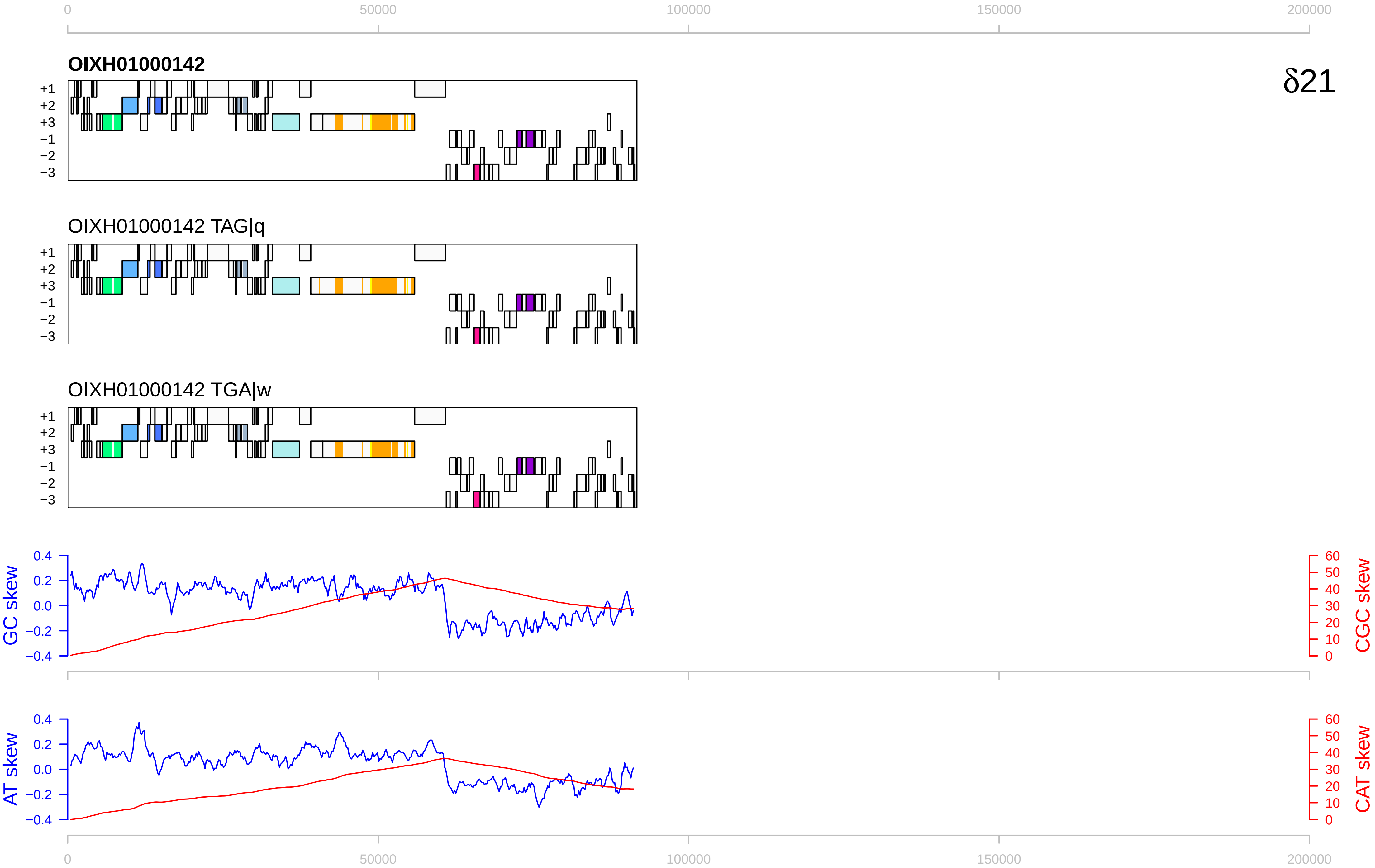

0
50000
100000
150000
200000
OIXH01000142
δ21
+1
+2
+3
−1
−2
−3
OIXH01000142 TAG|q
OIXH01000142 TGA|w
GC skew
CGC skew
0.4
0.2
0.0
−0.2
−0.4
60
50
40
30
20
10
0
AT skew
CAT skew

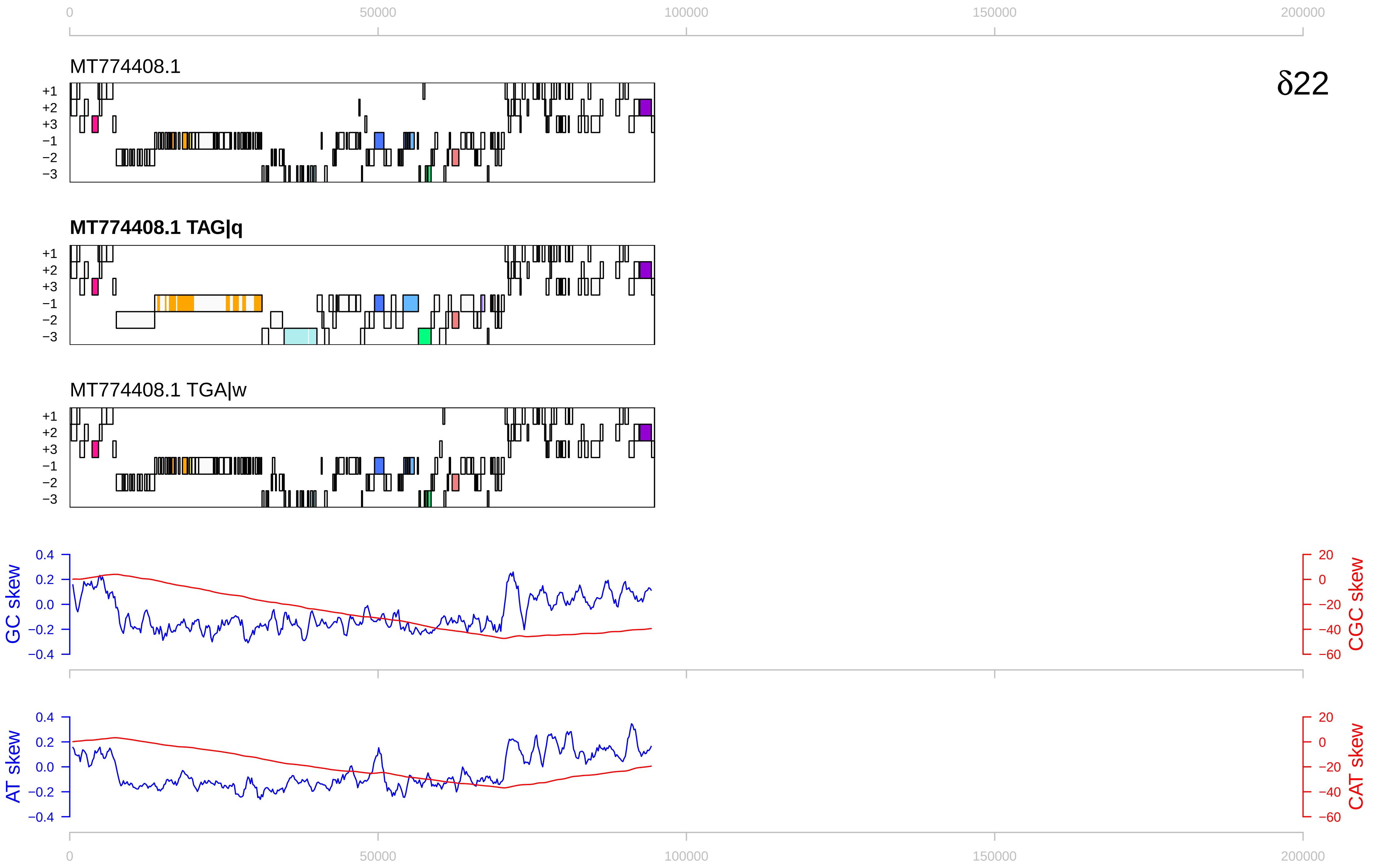

0
50000
100000
150000
200000
MT774408.1
δ22
+1
+2
+3
−1
−2
−3
MT774408.1 TAG|q
MT774408.1 TGA|w
GC skew
CGC skew
AT skew
CAT skew
0.4
0.2
0.0
−0.2
−0.4
20
0
−20
−40
−60

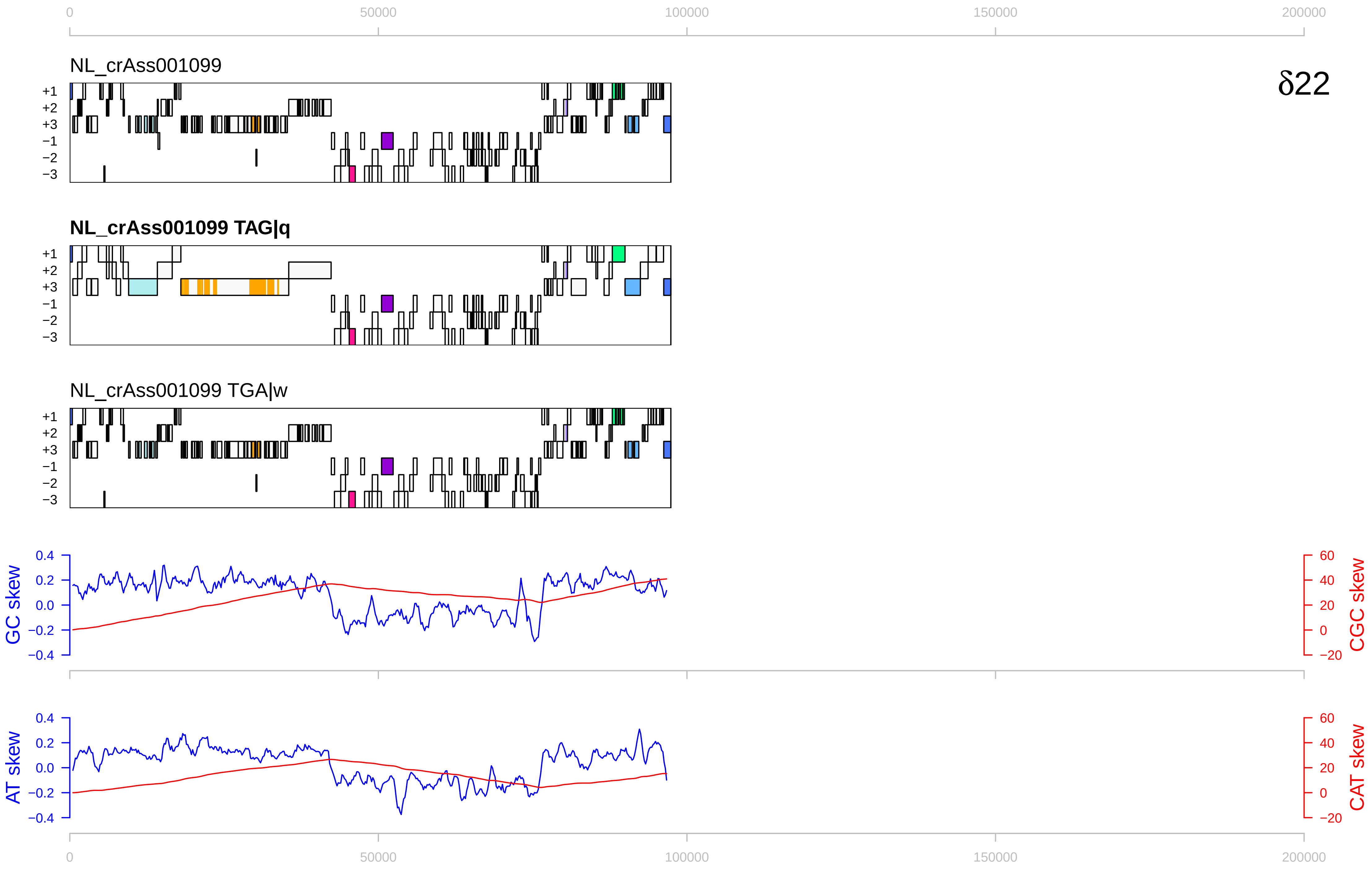

0
50000
100000
150000
200000
NL_crAss001099
δ22
+1
+2
+3
−1
−2
−3
NL_crAss001099 TAG|q
NL_crAss001099 TGA|w
GC skew
CGC skew
AT skew
CAT skew
0.4
0.2
0.0
−0.2
−0.4
60
40
20
0
−20

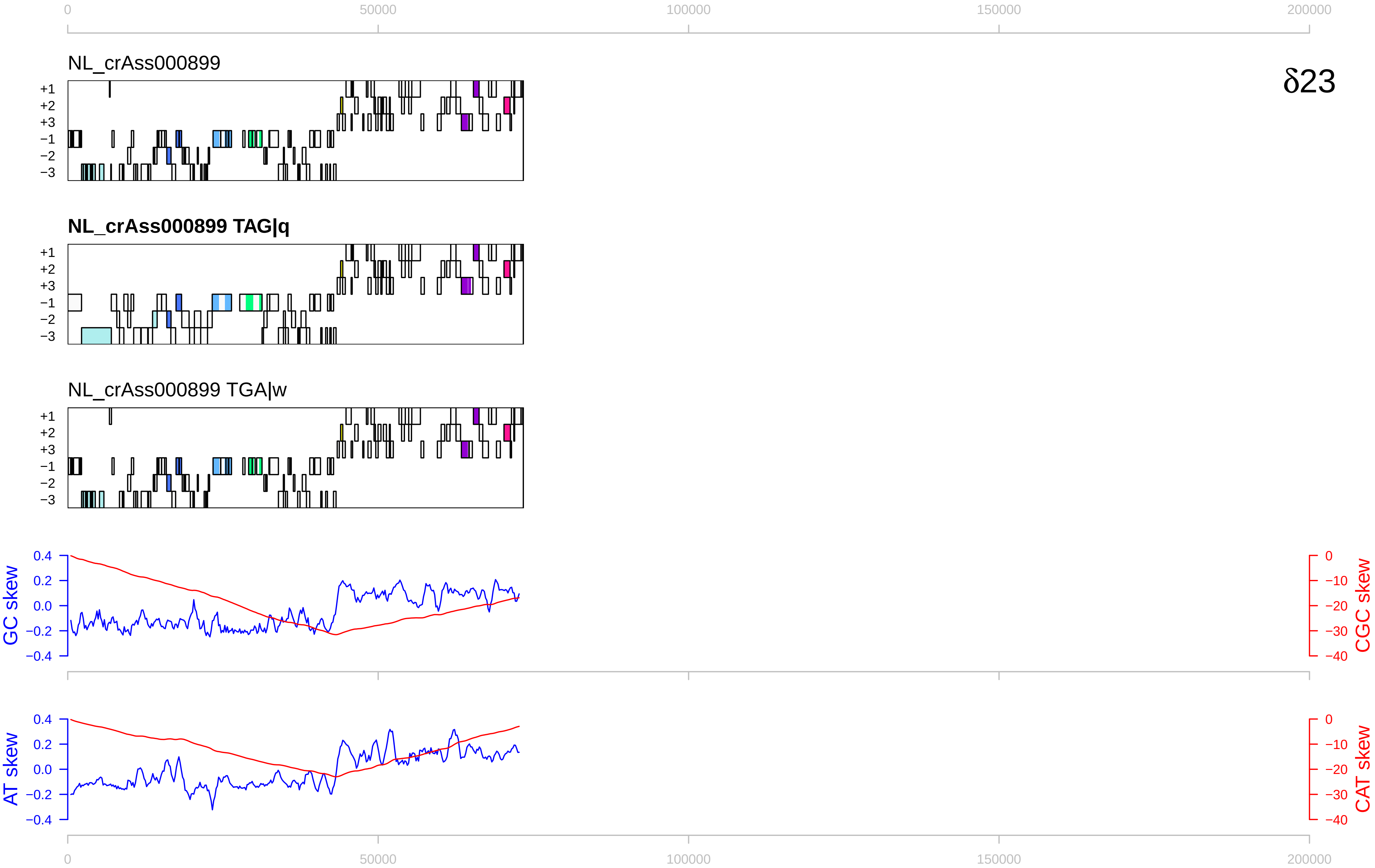

0
50000
100000
150000
200000
NL_crAss000899
δ23
+1
+2
+3
−1
−2
−3
NL_crAss000899 TAG|q
NL_crAss000899 TGA|w
GC skew
0.4
0.2
0.0
−0.2
−0.4
CGC skew
0
−10
−20
−30
−40
AT skew
CAT skew

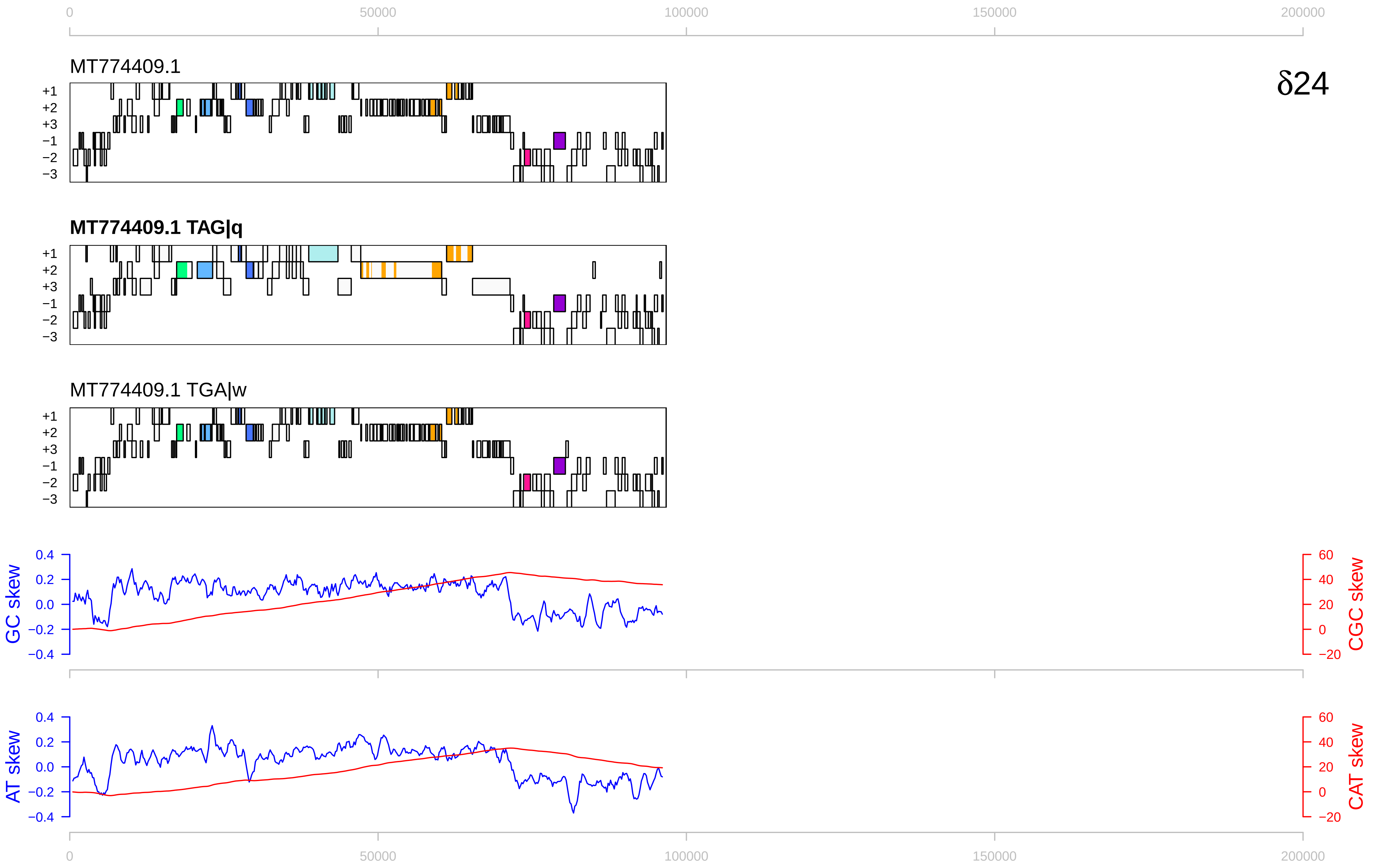

0
50000
100000
150000
200000
δ24
MT774409.1
MT774409.1 TAG|q
MT774409.1 TGA|w
+1
+2
+3
−1
−2
−3
GC skew
CGC skew
AT skew
CAT skew
0.4
0.2
0.0
−0.2
−0.4
60
40
20
0
−20

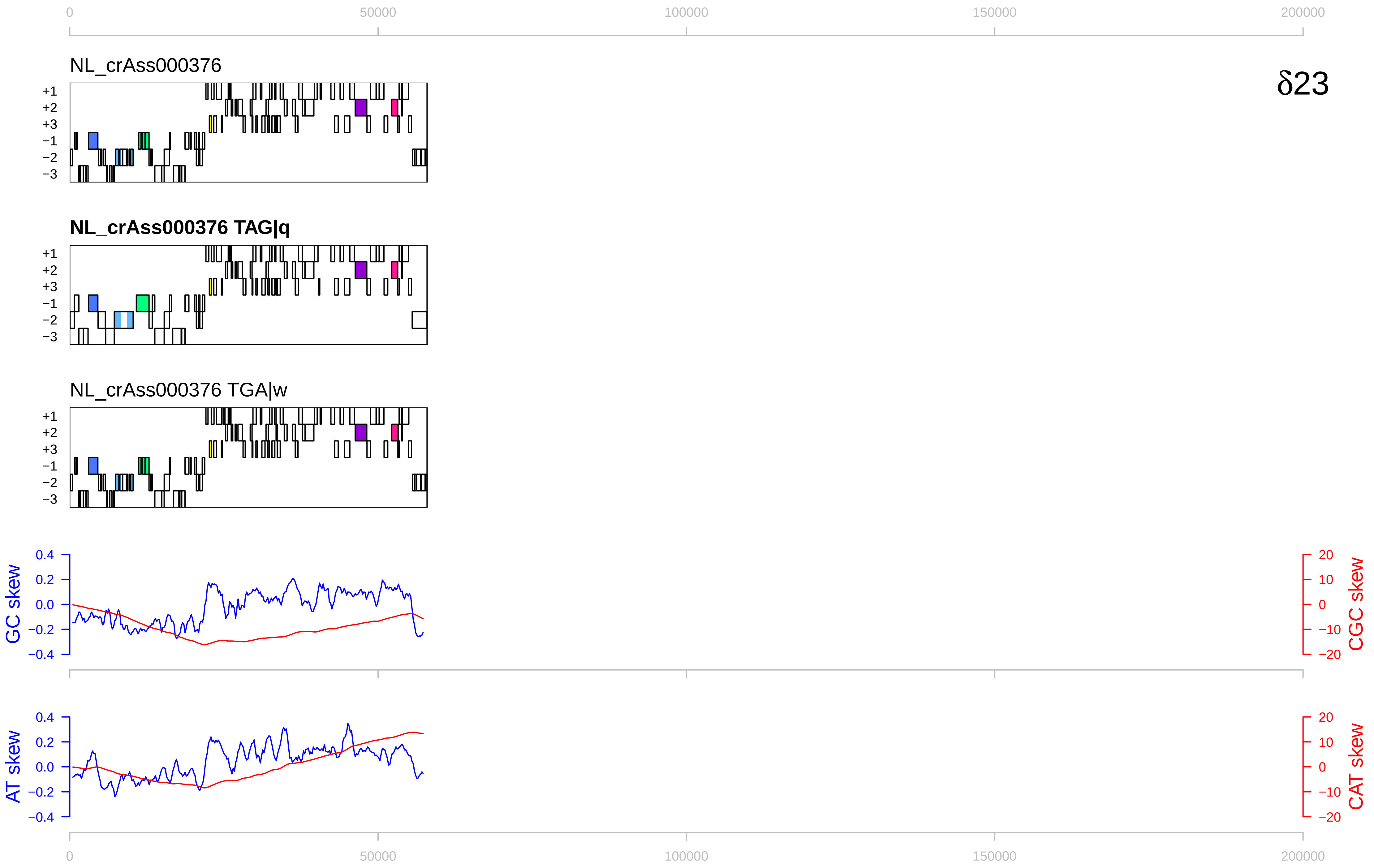

0
50000
100000
150000
200000
NL_crAss000376
δ23
+1
+2
+3
−1
−2
−3
NL_crAss000376 TAG|q
NL_crAss000376 TGA|w
GC skew
CGC skew
AT skew
CAT skew
0.4
0.2
0.0
−0.2
−0.4
20
10
0
−10
−20

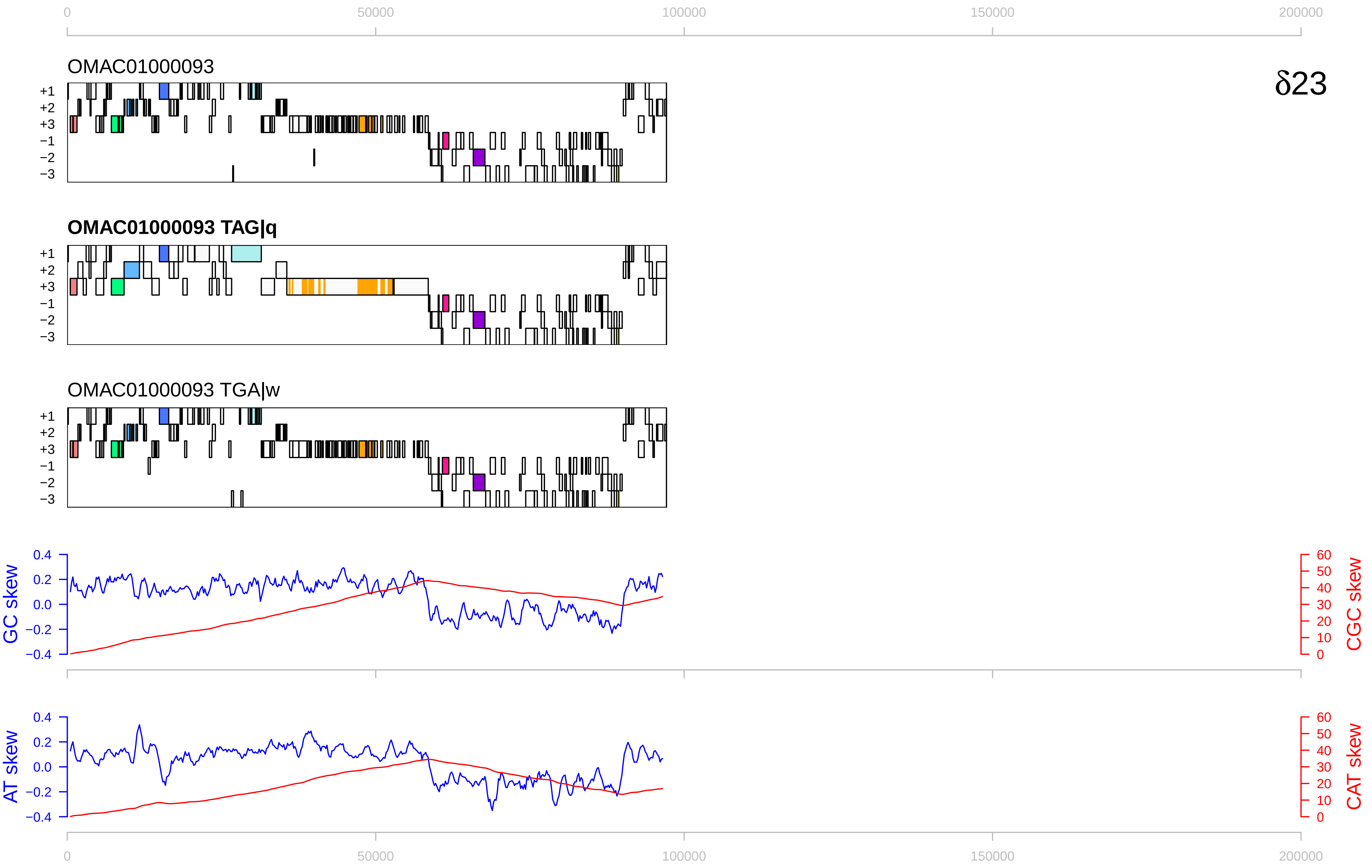

δ23
OMAC01000093
OMAC01000093 TAG|q
OMAC01000093 TGA|w
+1
+2
+3
−1
−2
−3
GC skew
CGC skew
AT skew
CAT skew
0
50000
100000
150000
200000
0.4
0.2
0.0
−0.2
−0.4
60
50
40
30
20
10
0

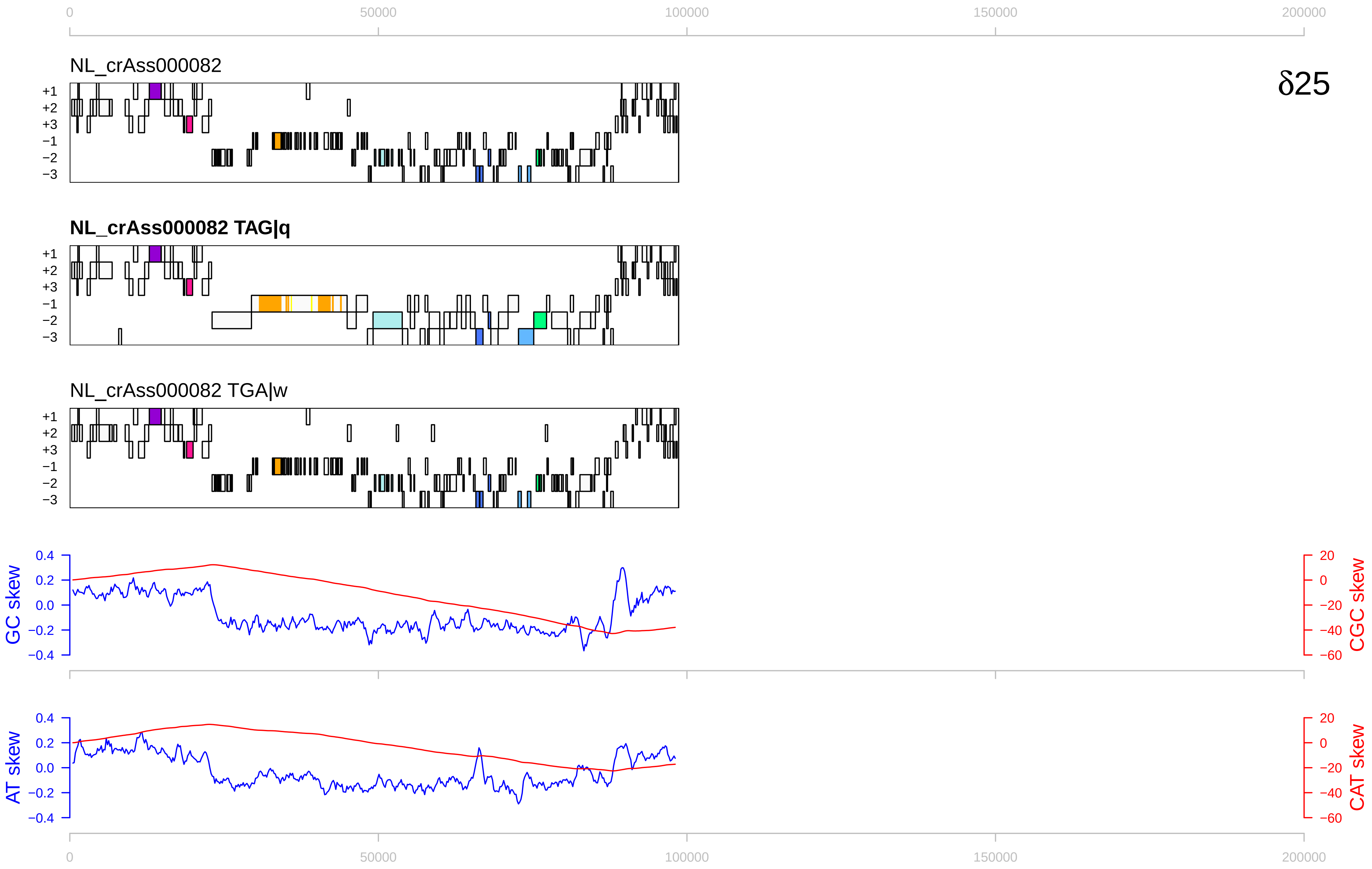

0
50000
100000
150000
200000
NL_crAss000082
δ25
+1
+2
+3
−1
−2
−3
NL_crAss000082 TAG|q
NL_crAss000082 TGA|w
GC skew
CGC skew
AT skew
CAT skew
0.4
0.2
0.0
−0.2
−0.4
20
0
−20
−40
−60

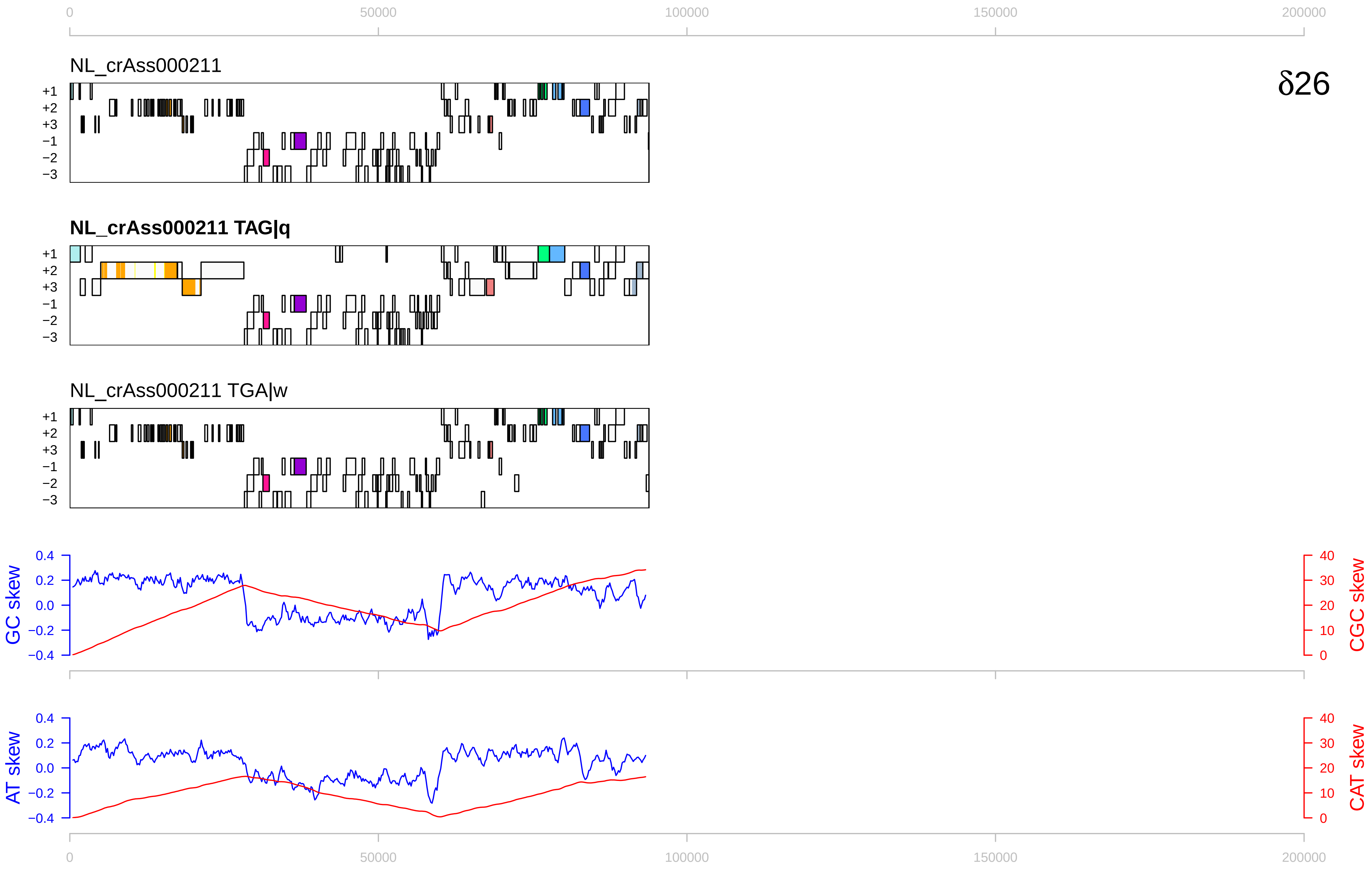

0
50000
100000
150000
200000
NL_crAss000211
δ26
+1
+2
+3
−1
−2
−3
NL_crAss000211 TAG|q
NL_crAss000211 TGA|w
GC skew
CGC skew
AT skew
CAT skew
0.4
0.2
0.0
−0.2
−0.4
40
30
20
10
0

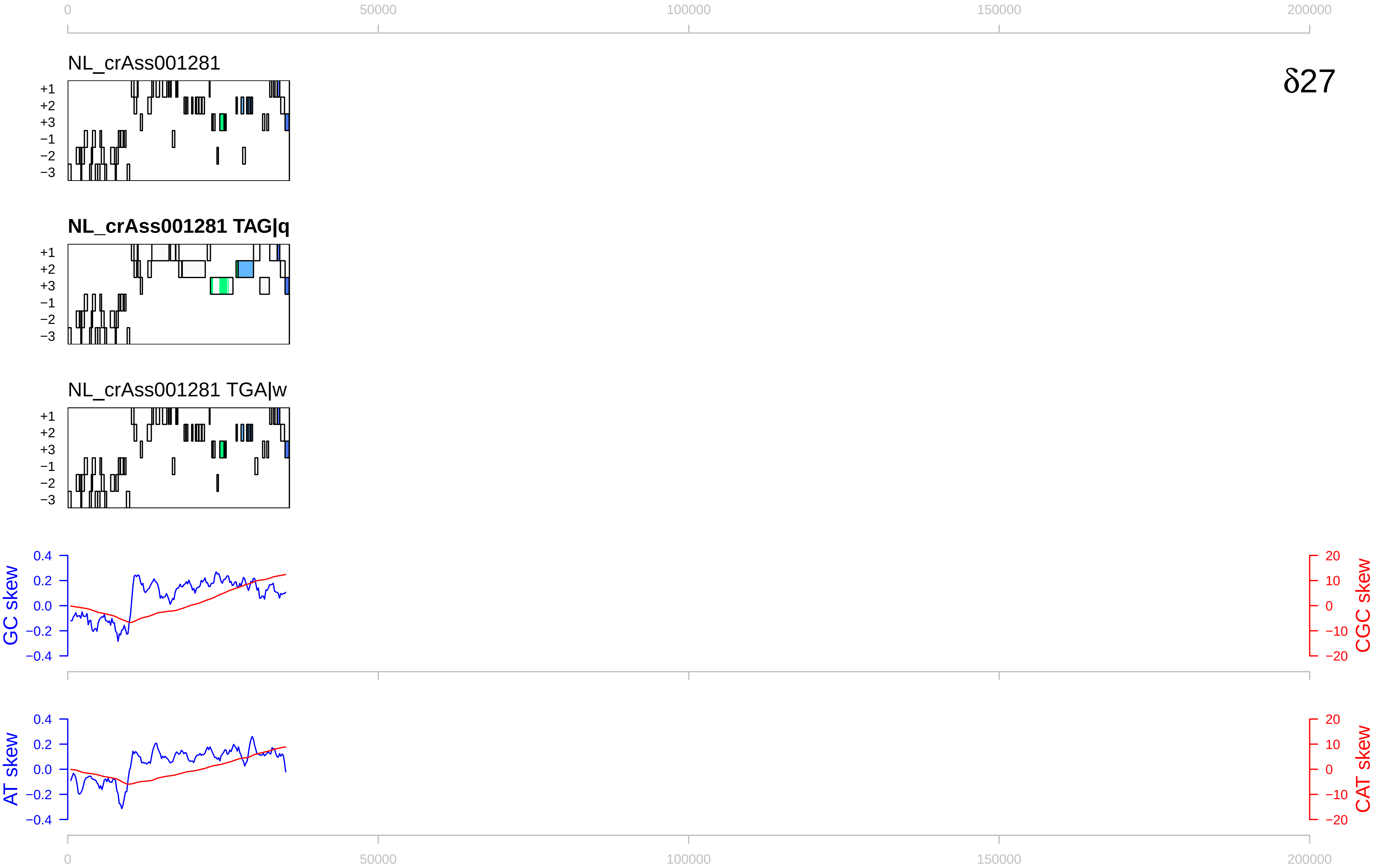

δ27
NL_crAss001281
NL_crAss001281 TAG|q
NL_crAss001281 TGA|w
+1
+2
+3
−1
−2
−3
GC skew
CGC skew
AT skew
CAT skew
0
50000
100000
150000
200000

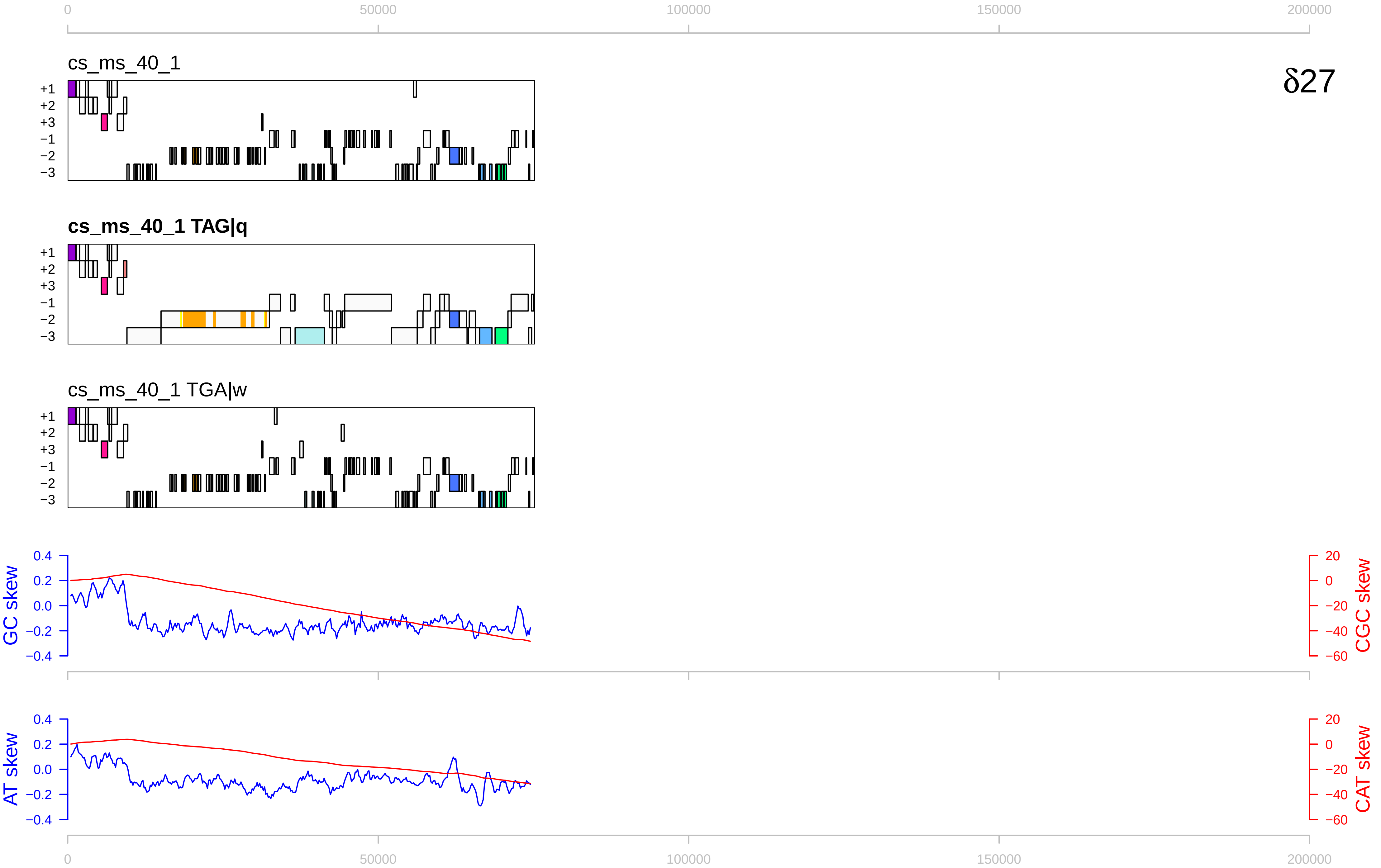

0
50000
100000
150000
200000
cs_ms_40_1
δ27
+1
+2
+3
−1
−2
−3
cs_ms_40_1 TAG|q
cs_ms_40_1 TGA|w
GC skew
CGC skew
0.4
0.2
0.0
−0.2
−0.4
20
0
−20
−40
−60
AT skew
CAT skew

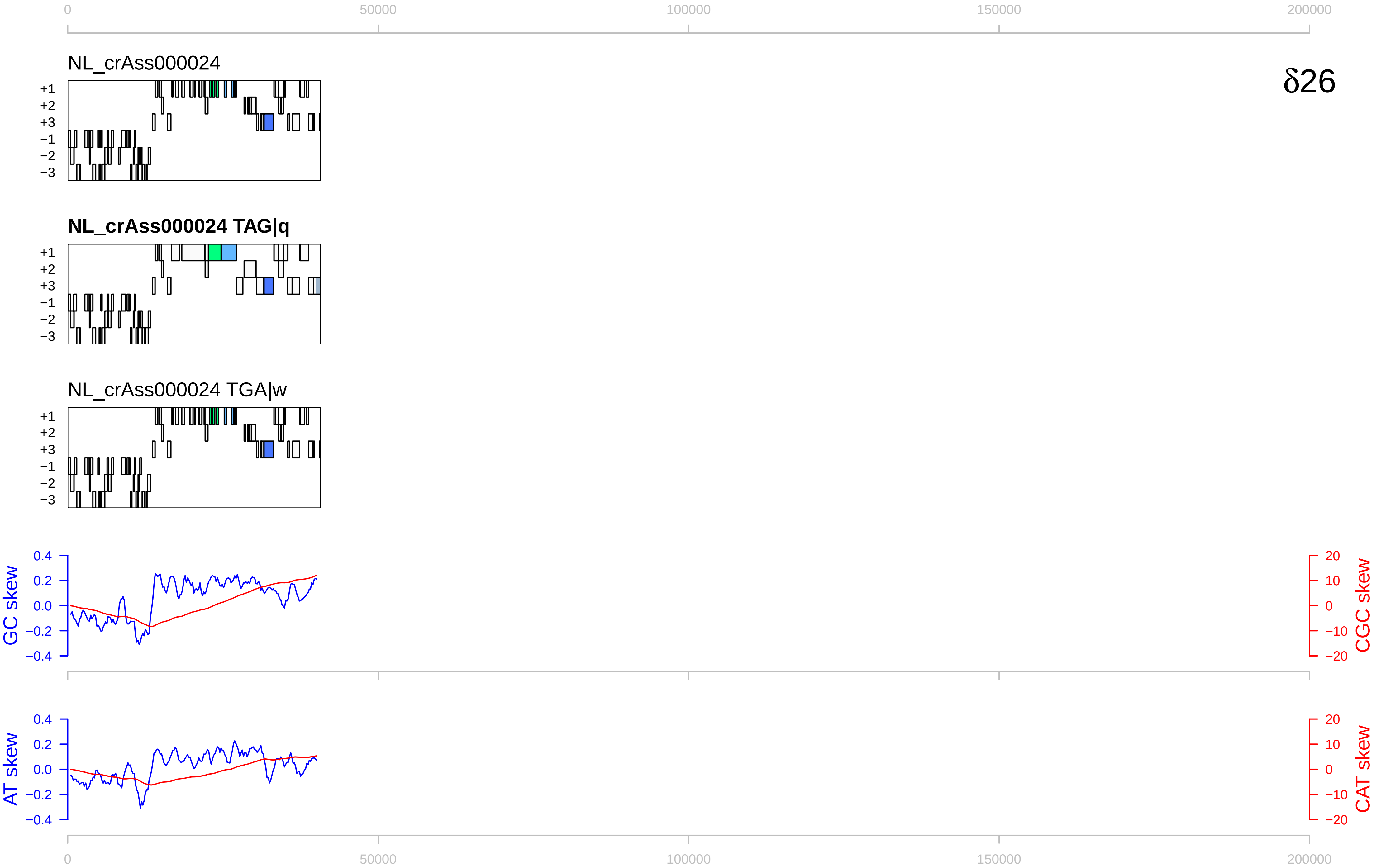

0
50000
100000
150000
200000
NL_crAss000024
δ26
+1
+2
+3
−1
−2
−3
NL_crAss000024 TAG|q
NL_crAss000024 TGA|w
GC skew
CGC skew
AT skew
CAT skew
0.4
0.2
0.0
−0.2
−0.4
20
10
0
−10
−20

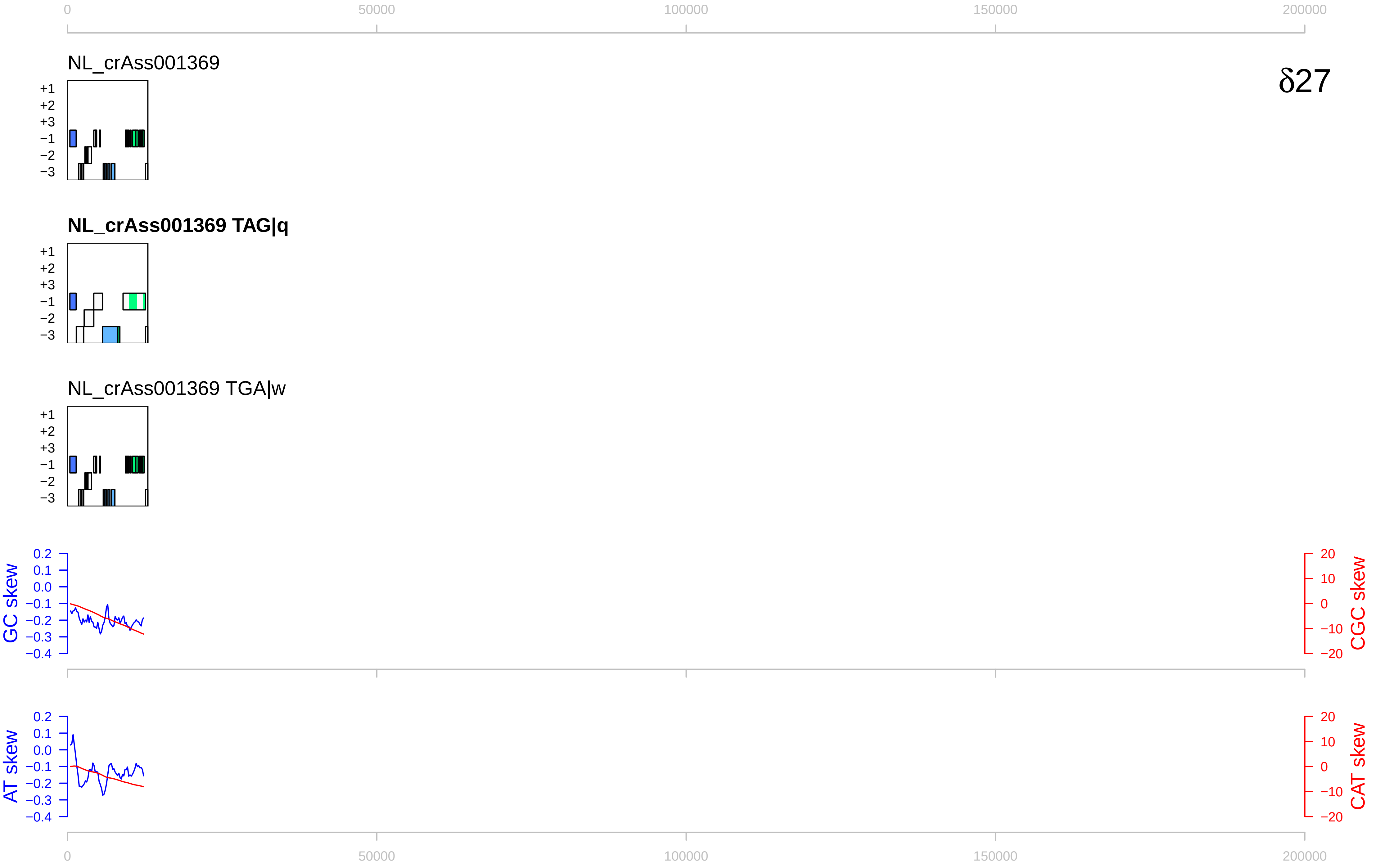

0
50000
100000
150000
200000
NL_crAss001369
δ27
+1
+2
+3
−1
−2
−3
NL_crAss001369 TAG|q
NL_crAss001369 TGA|w
GC skew
CGC skew
AT skew
CAT skew
0.2
0.1
0.0
−0.1
−0.2
−0.3
−0.4
20
10
0
−10
−20

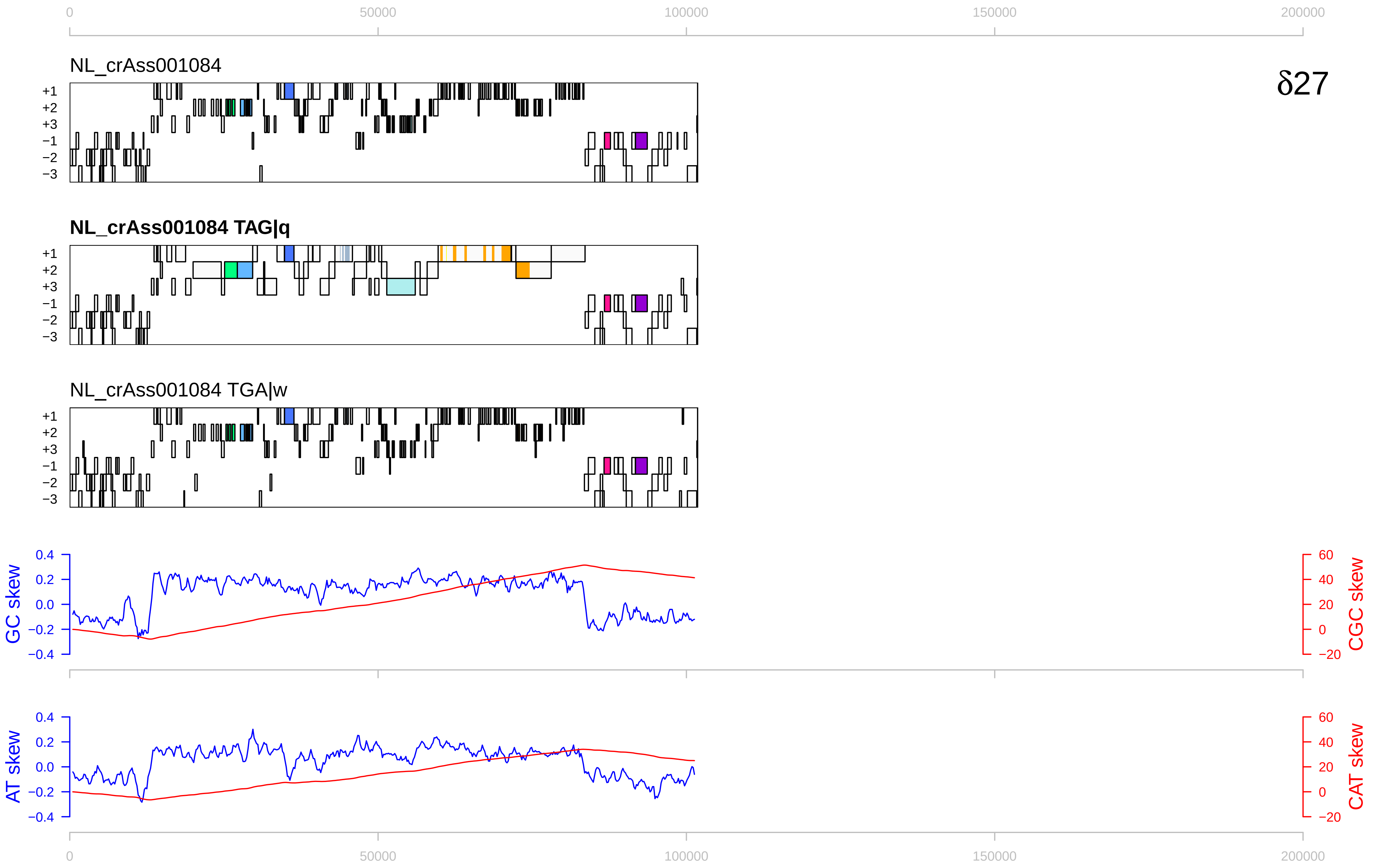

0
50000
100000
150000
200000
NL_crAss001084
δ27
+1
+2
+3
−1
−2
−3
NL_crAss001084 TAG|q
NL_crAss001084 TGA|w
GC skew
CGC skew
AT skew
CAT skew
0.4
0.2
0.0
−0.2
−0.4
60
40
20
0
−20

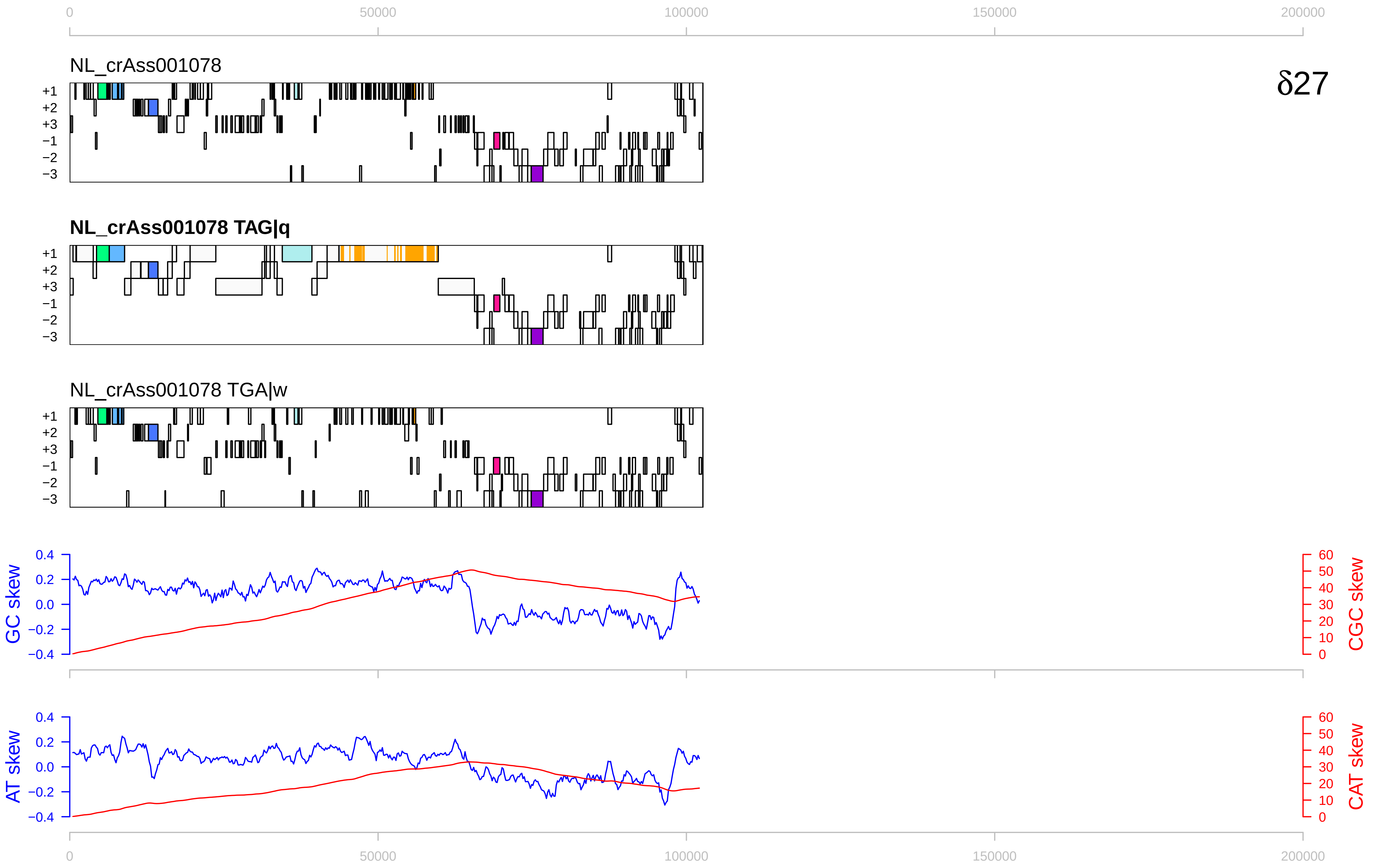

0
50000
100000
150000
200000
NL_crAss001078
δ27
+1
+2
+3
−1
−2
−3
NL_crAss001078 TAG|q
NL_crAss001078 TGA|w
GC skew
CGC skew
AT skew
CAT skew
0.4
0.2
0.0
−0.2
−0.4
60
50
40
30
20
10
0

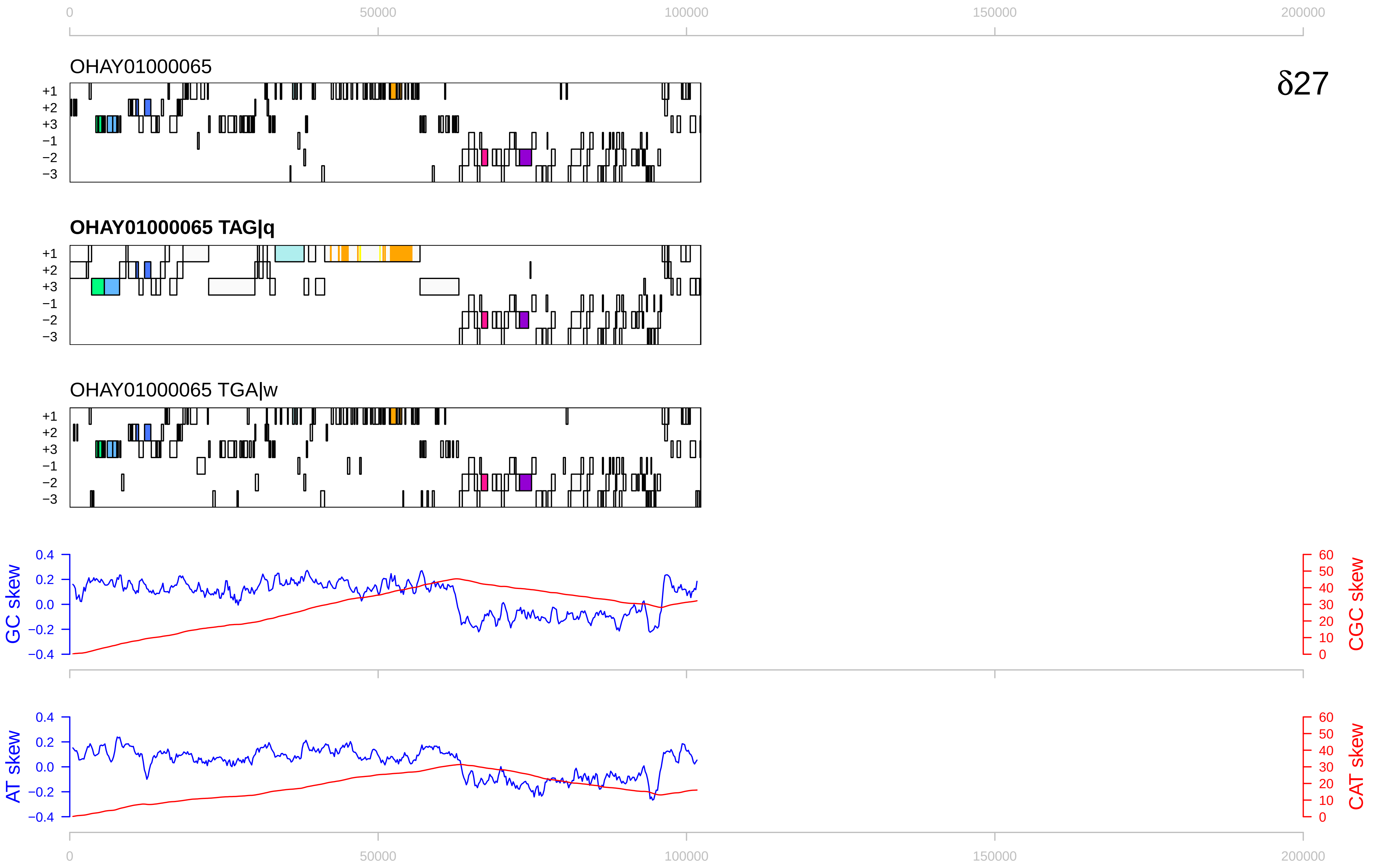

δ27
OHAY01000065
OHAY01000065 TAG|q
OHAY01000065 TGA|w
+1
+2
+3
−1
−2
−3
GC skew
CGC skew
AT skew
CAT skew
0
50000
100000
150000
200000

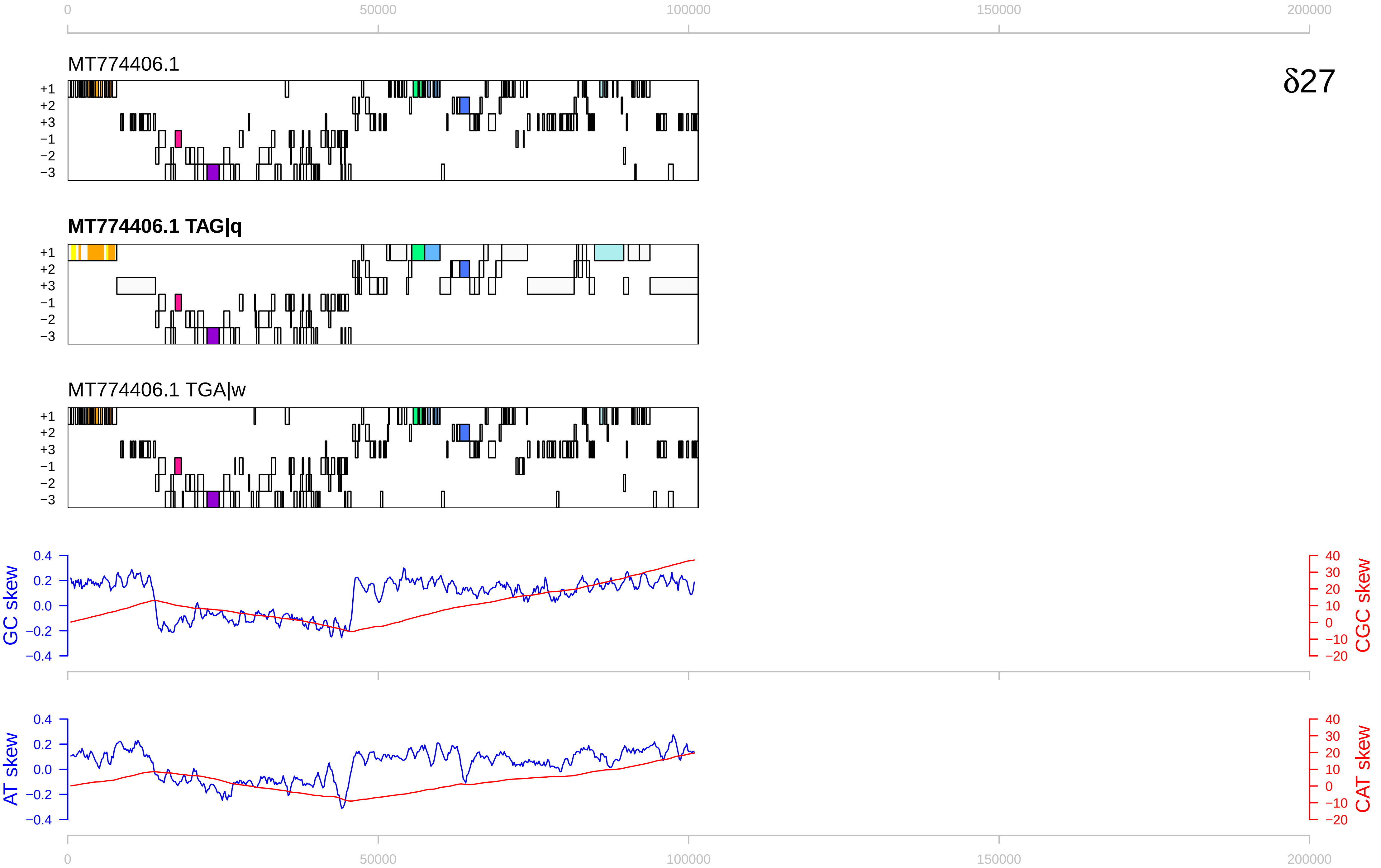

0
50000
100000
150000
200000
MT774406.1
δ27
+1
+2
+3
−1
−2
−3
MT774406.1 TAG|q
MT774406.1 TGA|w
GC skew
CGC skew
0.4
0.2
0.0
−0.2
−0.4
40
30
20
10
0
−10
−20
AT skew
CAT skew

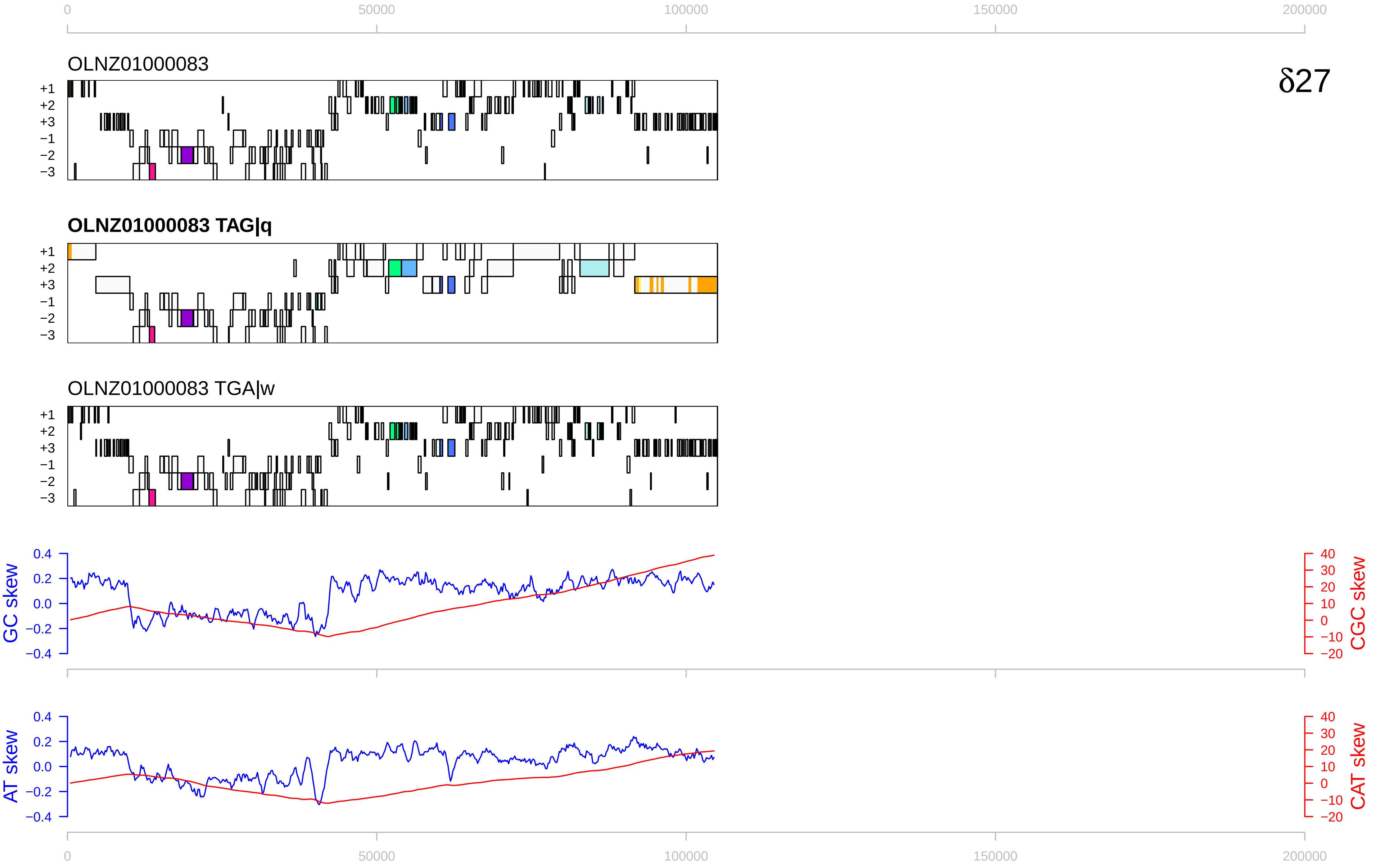

0
50000
100000
150000
200000
OLNZ01000083
δ27
+1
+2
+3
−1
−2
−3
OLNZ01000083 TAG|q
OLNZ01000083 TGA|w
GC skew
CGC skew
AT skew
CAT skew
0.4
0.2
0.0
−0.2
−0.4
40
30
20
10
0
−10
−20

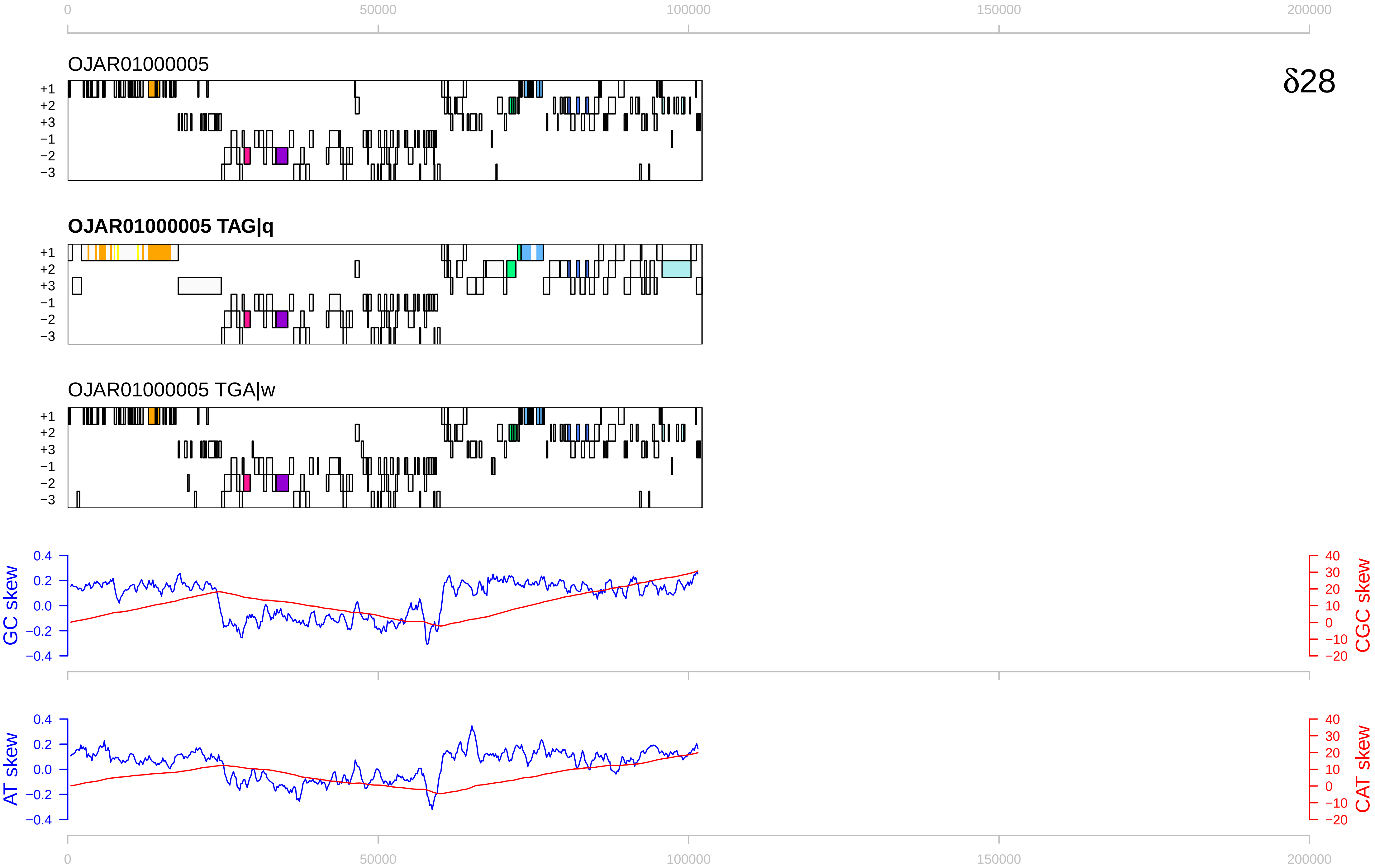

δ28
OJAR01000005
OJAR01000005 TAG|q
OJAR01000005 TGA|w
+1
+2
+3
−1
−2
−3
GC skew
CGC skew
AT skew
CAT skew
0
50000
100000
150000
200000

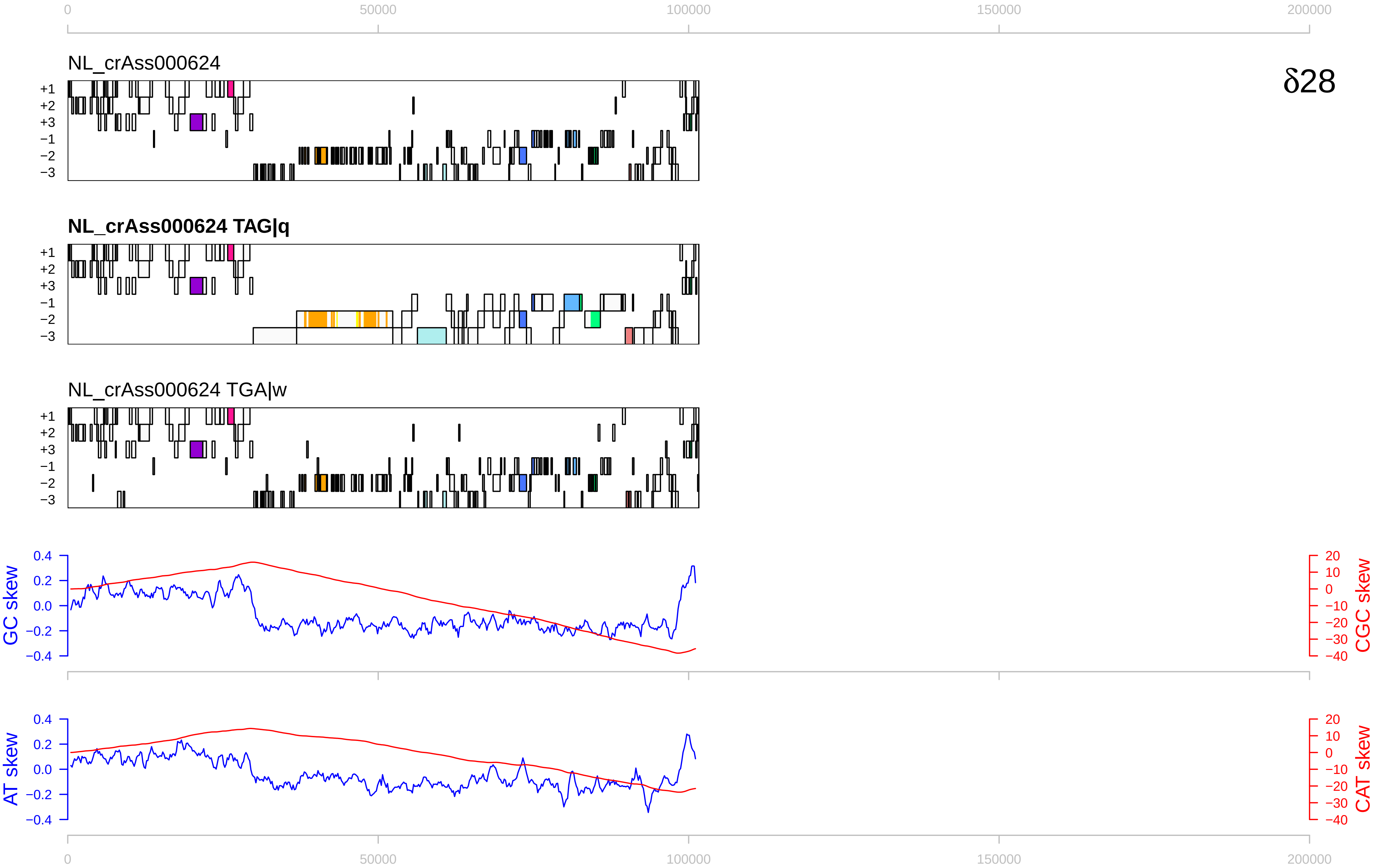

0
50000
100000
150000
200000
NL_crAss000624
δ28
+1
+2
+3
−1
−2
−3
NL_crAss000624 TAG|q
NL_crAss000624 TGA|w
GC skew
CGC skew
AT skew
CAT skew
0.4
0.2
0.0
−0.2
−0.4
20
10
0
−10
−20
−30
−40

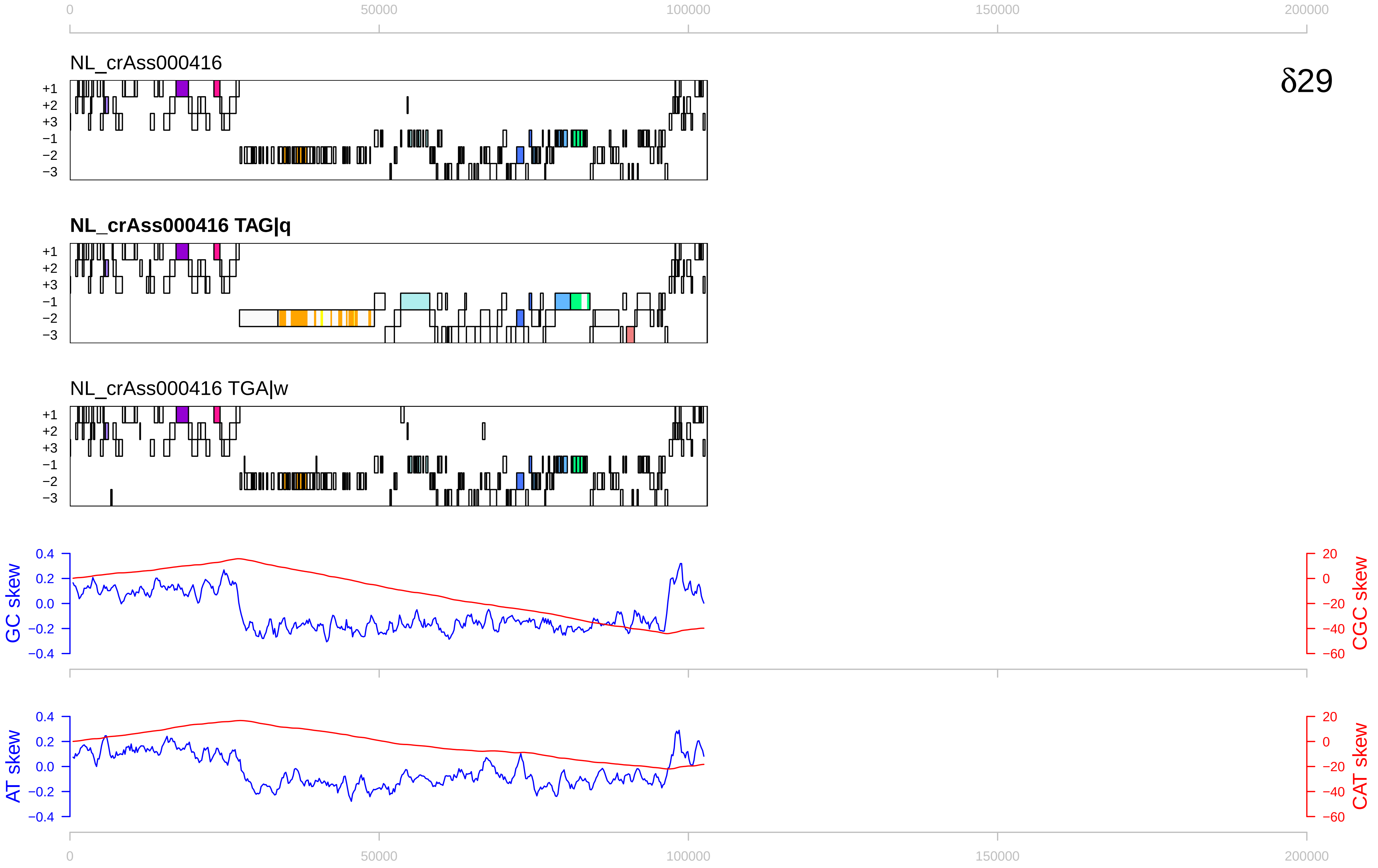

0
50000
100000
150000
200000
NL_crAss000416
δ29
+1
+2
+3
−1
−2
−3
NL_crAss000416 TAG|q
NL_crAss000416 TGA|w
GC skew
CGC skew
AT skew
CAT skew
0.4
0.2
0.0
−0.2
−0.4
20
0
−20
−40
−60

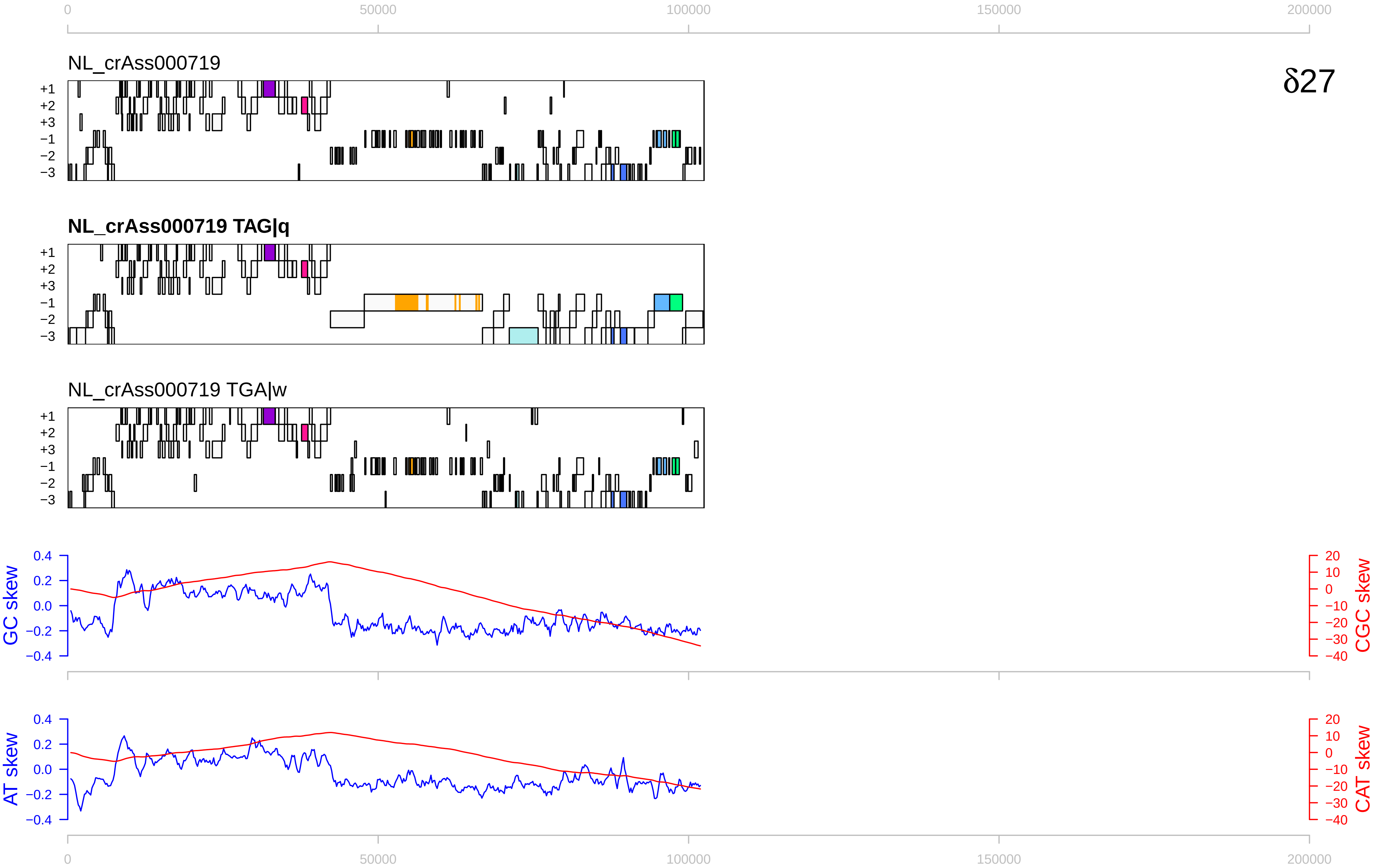
0
50000
100000
150000
200000
NL_crAss000719
δ27
+1
+2
+3
−1
−2
−3
NL_crAss000719 TAG|q
NL_crAss000719 TGA|w
GC skew
CGC skew
AT skew
CAT skew
0.4
0.2
0.0
−0.2
−0.4
20
10
0
−10
−20
−30
−40

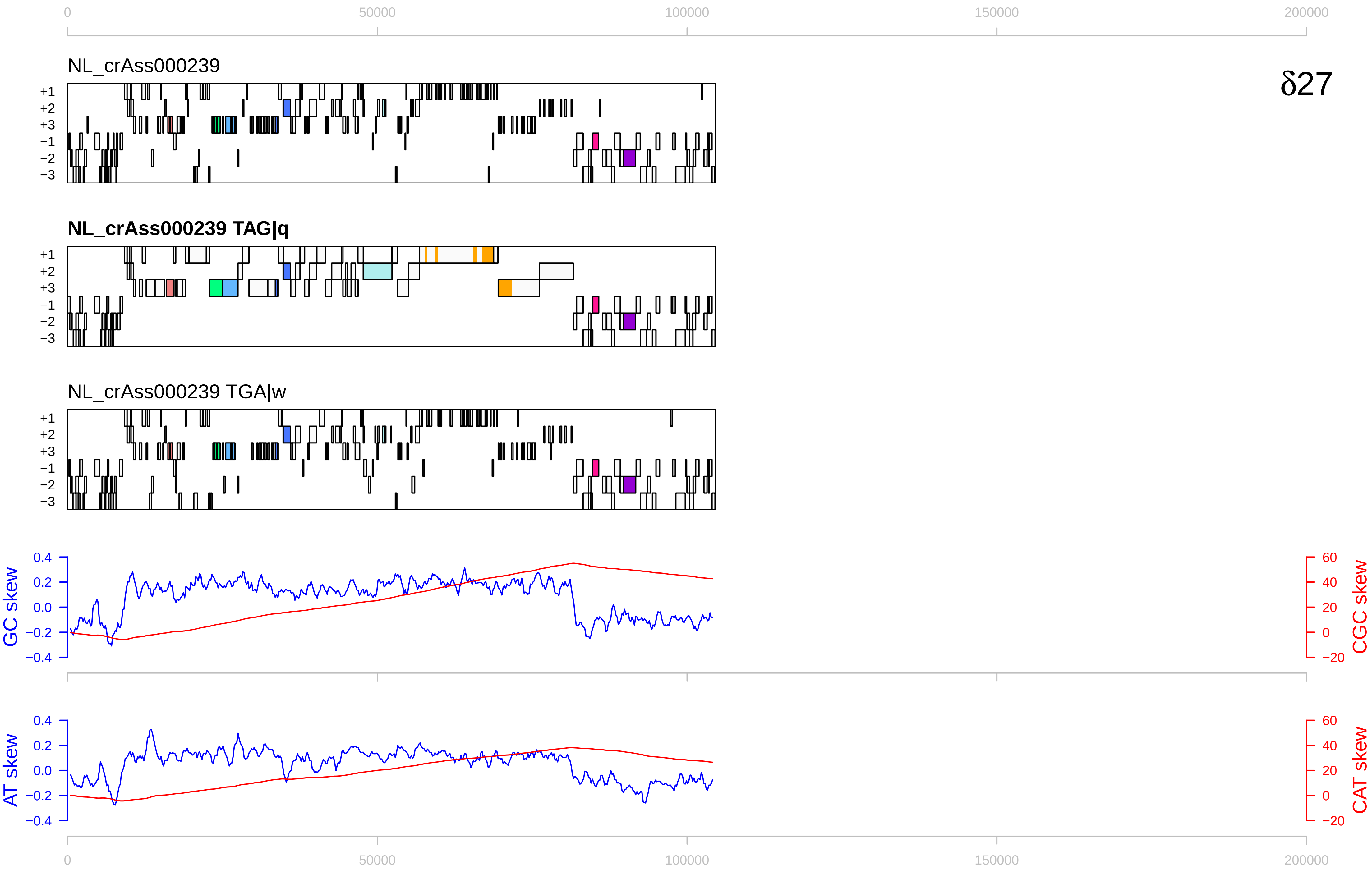

δ27
NL_crAss000239
NL_crAss000239 TAG|q
NL_crAss000239 TGA|w
+1
+2
+3
−1
−2
−3
GC skew
CGC skew
AT skew
CAT skew
0
50000
100000
150000
200000
0.4
0.2
0.0
−0.2
−0.4
60
40
20
0
−20

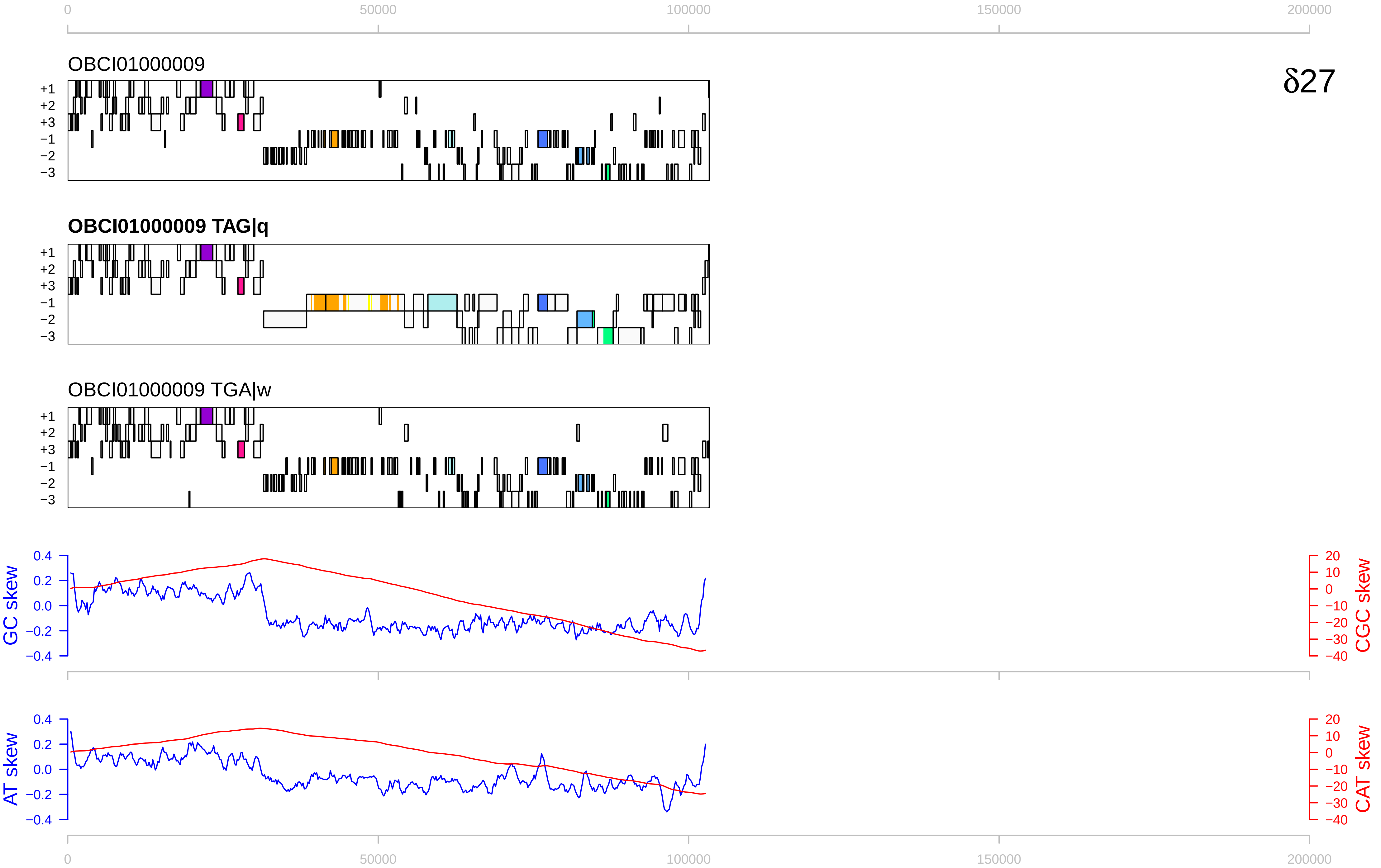

δ27
OBCI01000009
OBCI01000009 TAG|q
OBCI01000009 TGA|w
+1
+2
+3
−1
−2
−3
GC skew
CGC skew
AT skew
CAT skew
0
50000
100000
150000
200000

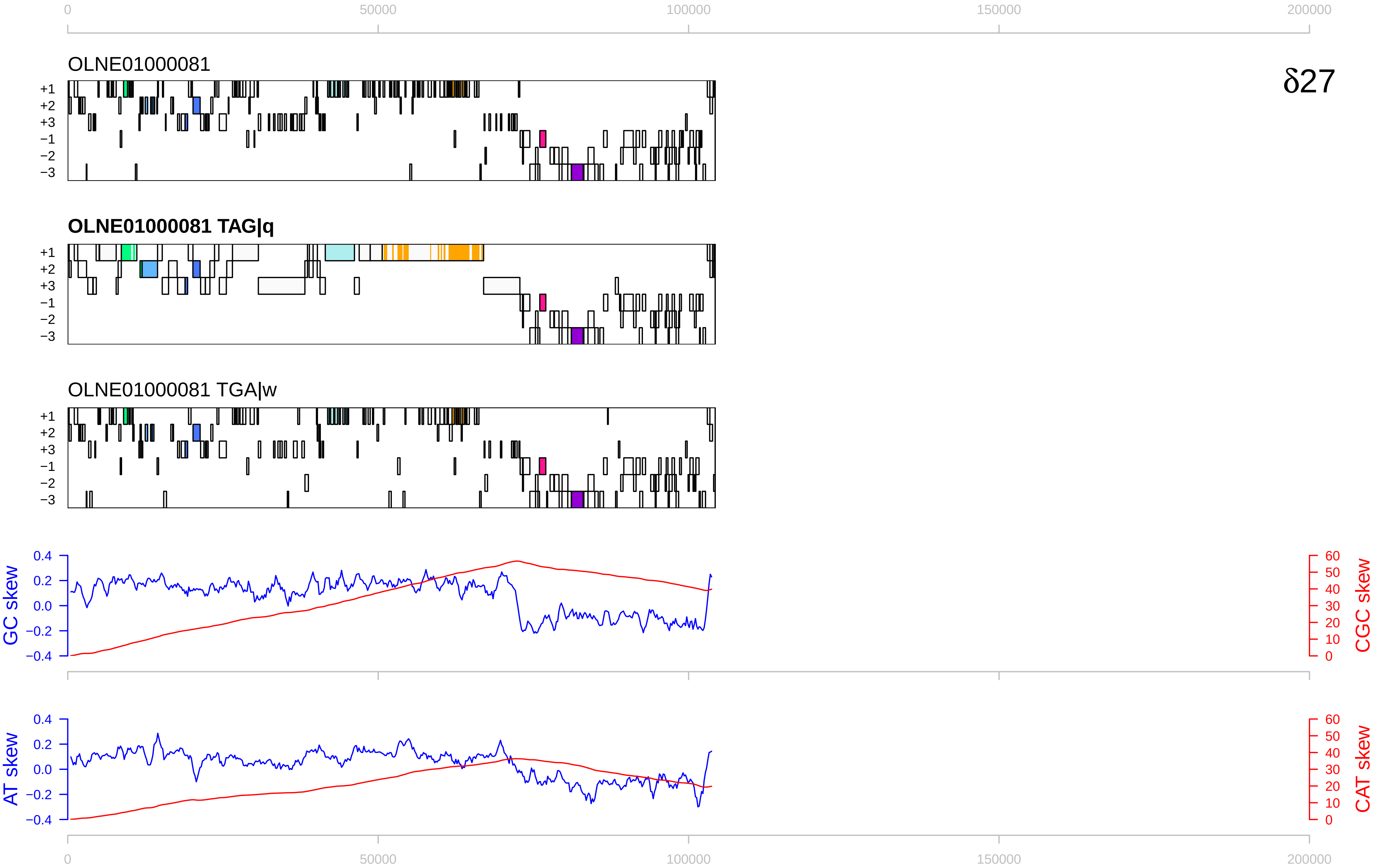

0
50000
100000
150000
200000
OLNE01000081
δ27
+1
+2
+3
−1
−2
−3
OLNE01000081 TAG|q
OLNE01000081 TGA|w
GC skew
CGC skew
AT skew
CAT skew
0.4
0.2
0.0
−0.2
−0.4
60
50
40
30
20
10
0

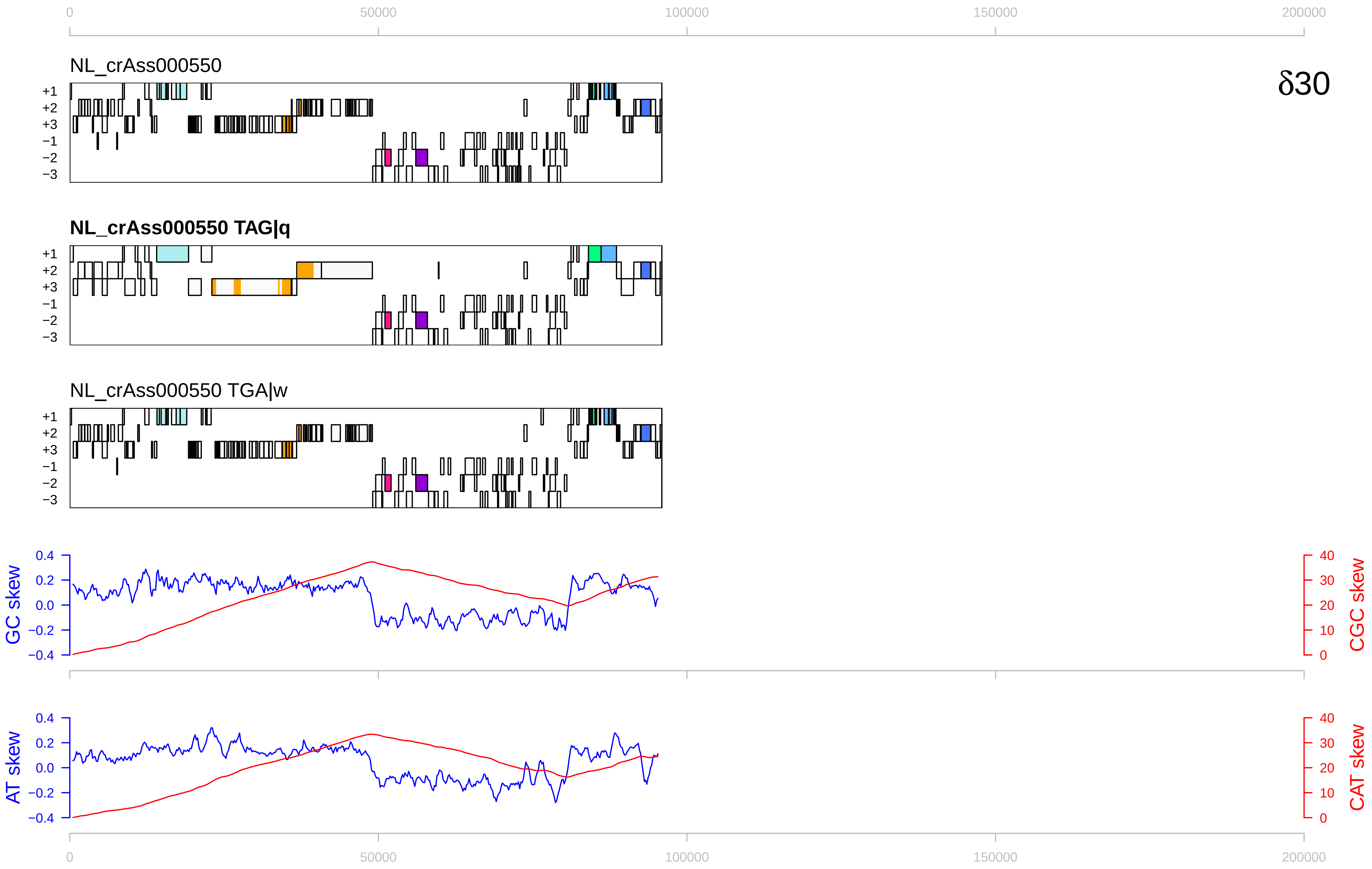

0
50000
100000
150000
200000
NL_crAss000550
δ30
+1
+2
+3
−1
−2
−3
NL_crAss000550 TAG|q
NL_crAss000550 TGA|w
GC skew
CGC skew
AT skew
CAT skew
0.4
0.2
0.0
−0.2
−0.4
40
30
20
10
0

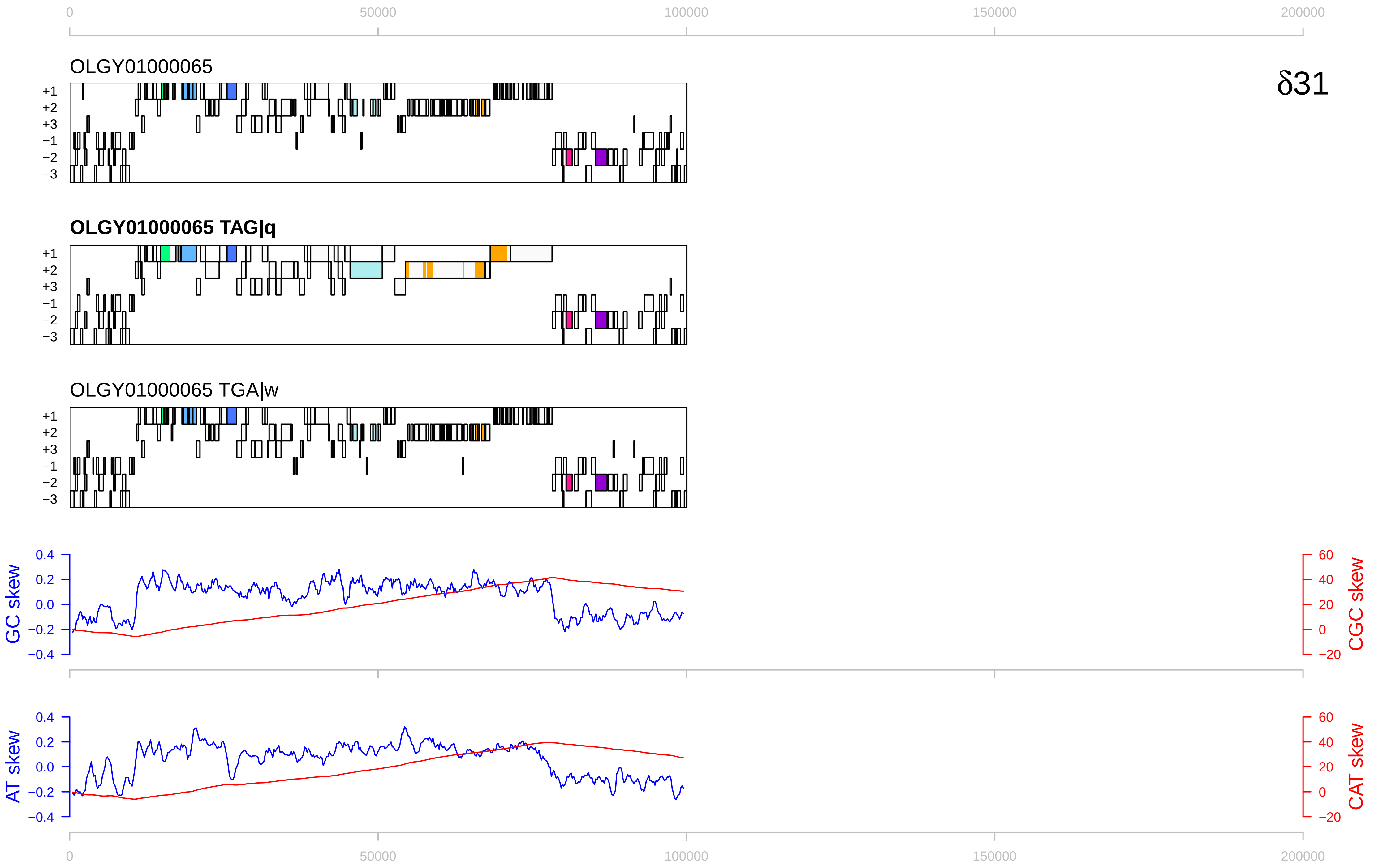

δ31
OLGY01000065
OLGY01000065 TAG|q
OLGY01000065 TGA|w
+1
+2
+3
−1
−2
−3
GC skew
CGC skew
AT skew
CAT skew
0
50000
100000
150000
200000
0.4
0.2
0.0
−0.2
−0.4
60
40
20
0
−20

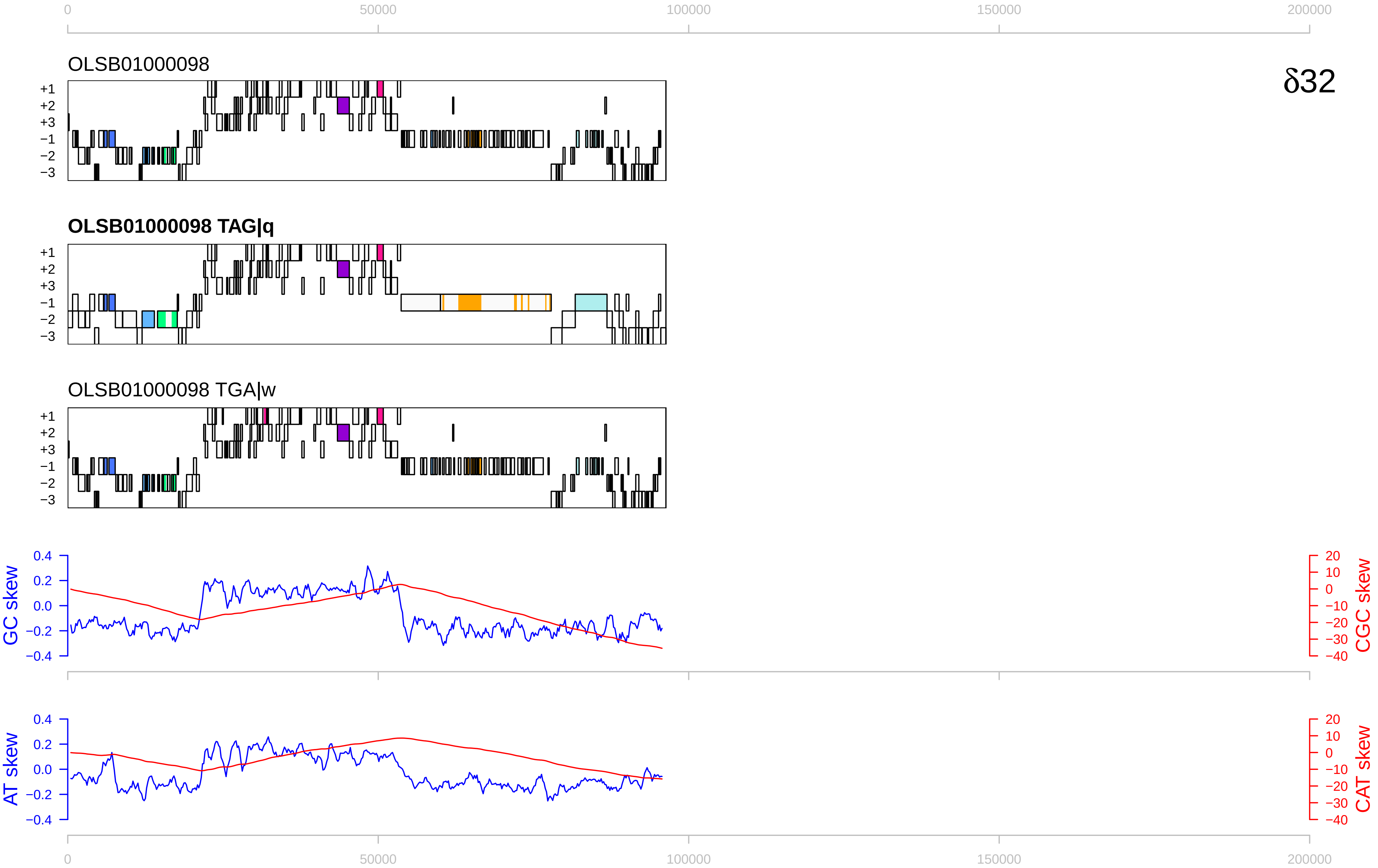

δ32
OLSB01000098
OLSB01000098 TAG|q
OLSB01000098 TGA|w
+1
+2
+3
−1
−2
−3
GC skew
CGC skew
AT skew
CAT skew
0
50000
100000
150000
200000
0.4
0.2
0.0
−0.2
−0.4
20
10
0
−10
−20
−30
−40

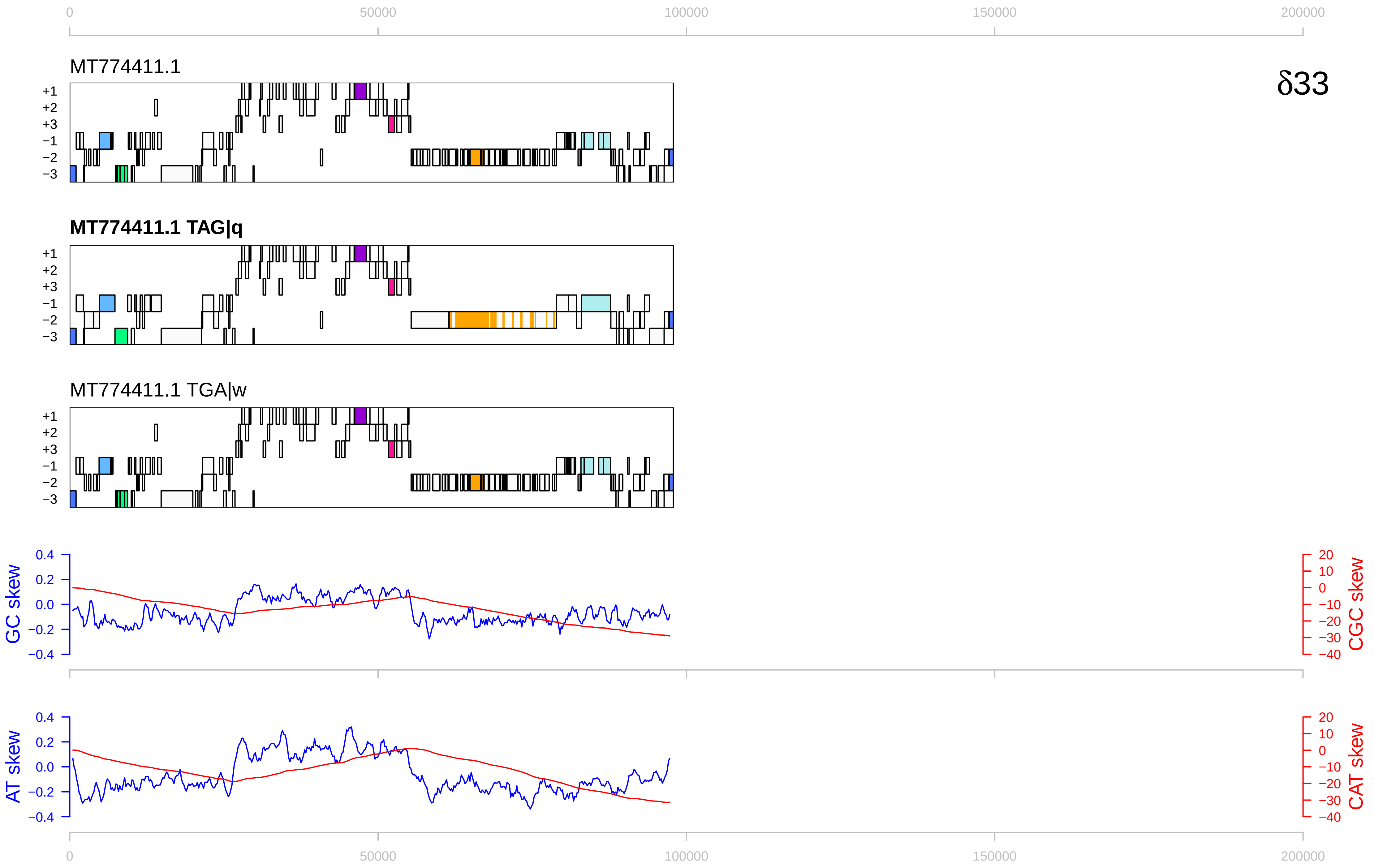

0
50000
100000
150000
200000
MT774411.1
δ33
+1
+2
+3
−1
−2
−3
MT774411.1 TAG|q
MT774411.1 TGA|w
GC skew
CGC skew
AT skew
CAT skew
0.4
0.2
0.0
−0.2
−0.4
20
10
0
−10
−20
−30
−40

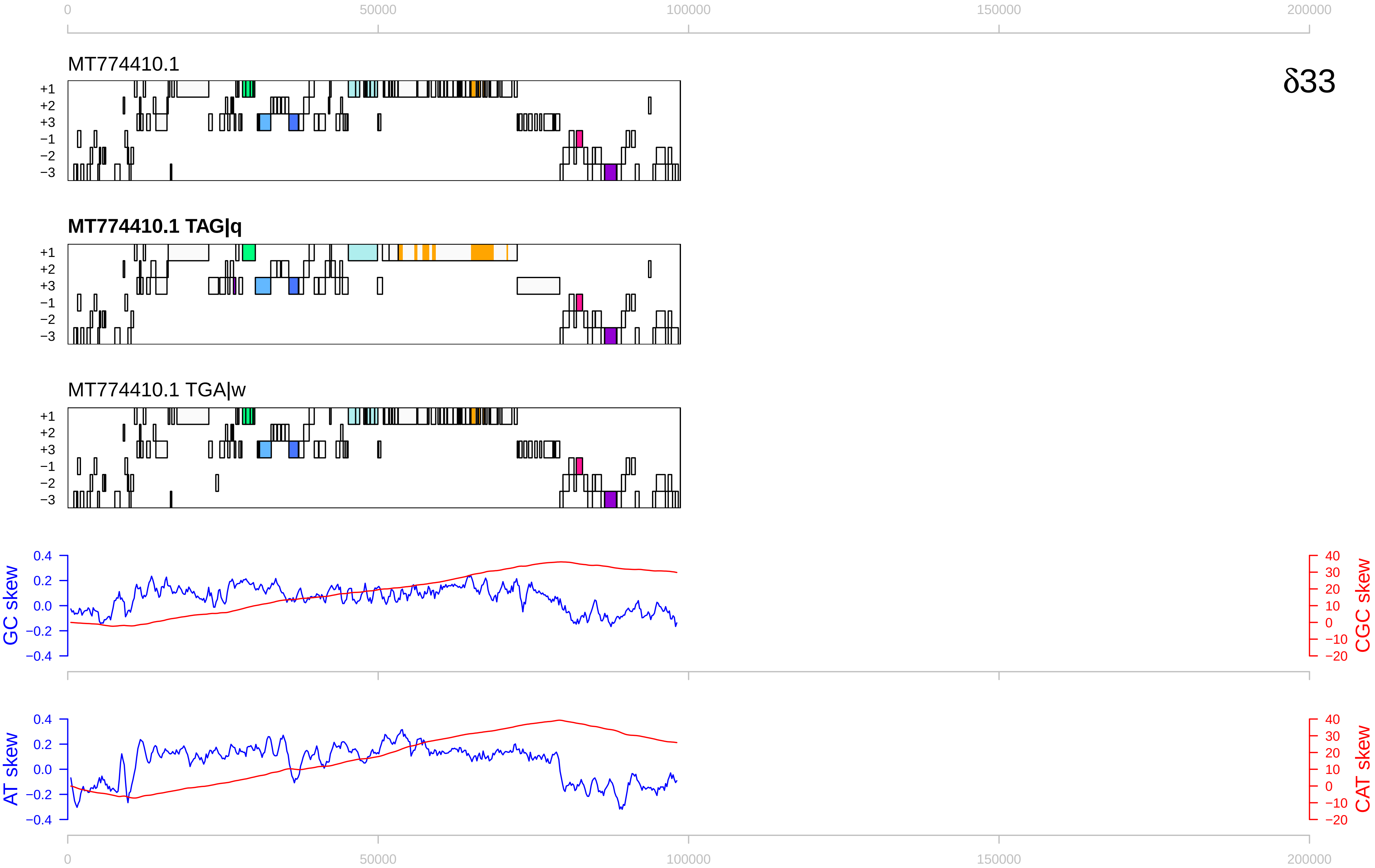

0
50000
100000
150000
200000
MT774410.1
δ33
+1
+2
+3
−1
−2
−3
MT774410.1 TAG|q
MT774410.1 TGA|w
GC skew
CGC skew
AT skew
CAT skew
0.4
0.2
0.0
−0.2
−0.4
40
30
20
10
0
−10
−20

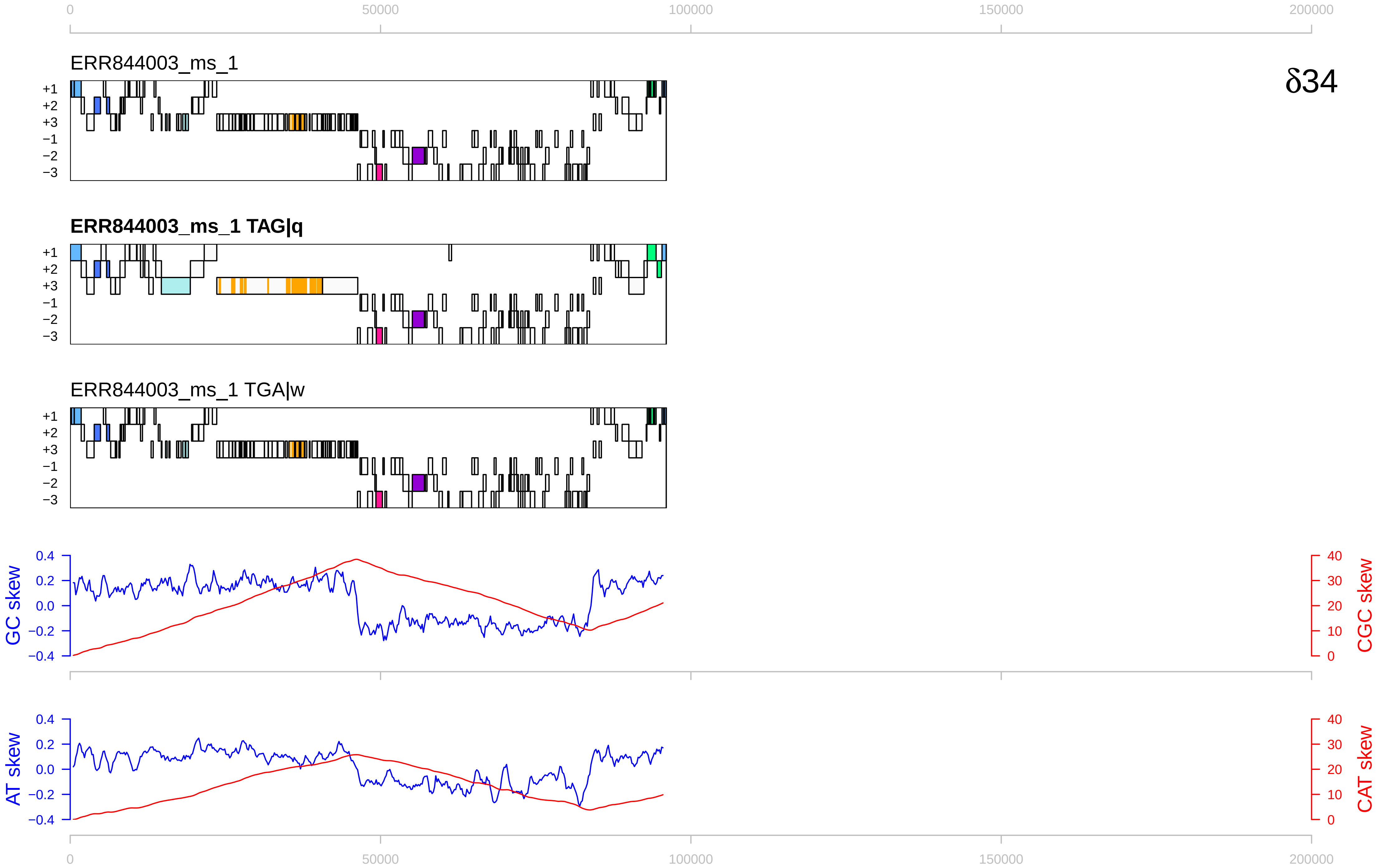

0
50000
100000
150000
200000
ERR844003_ms_1
δ34
+1
+2
+3
−1
−2
−3
ERR844003_ms_1 TAG|q
ERR844003_ms_1 TGA|w
GC skew
CGC skew
AT skew
CAT skew
0.4
0.2
0.0
−0.2
−0.4
40
30
20
10
0

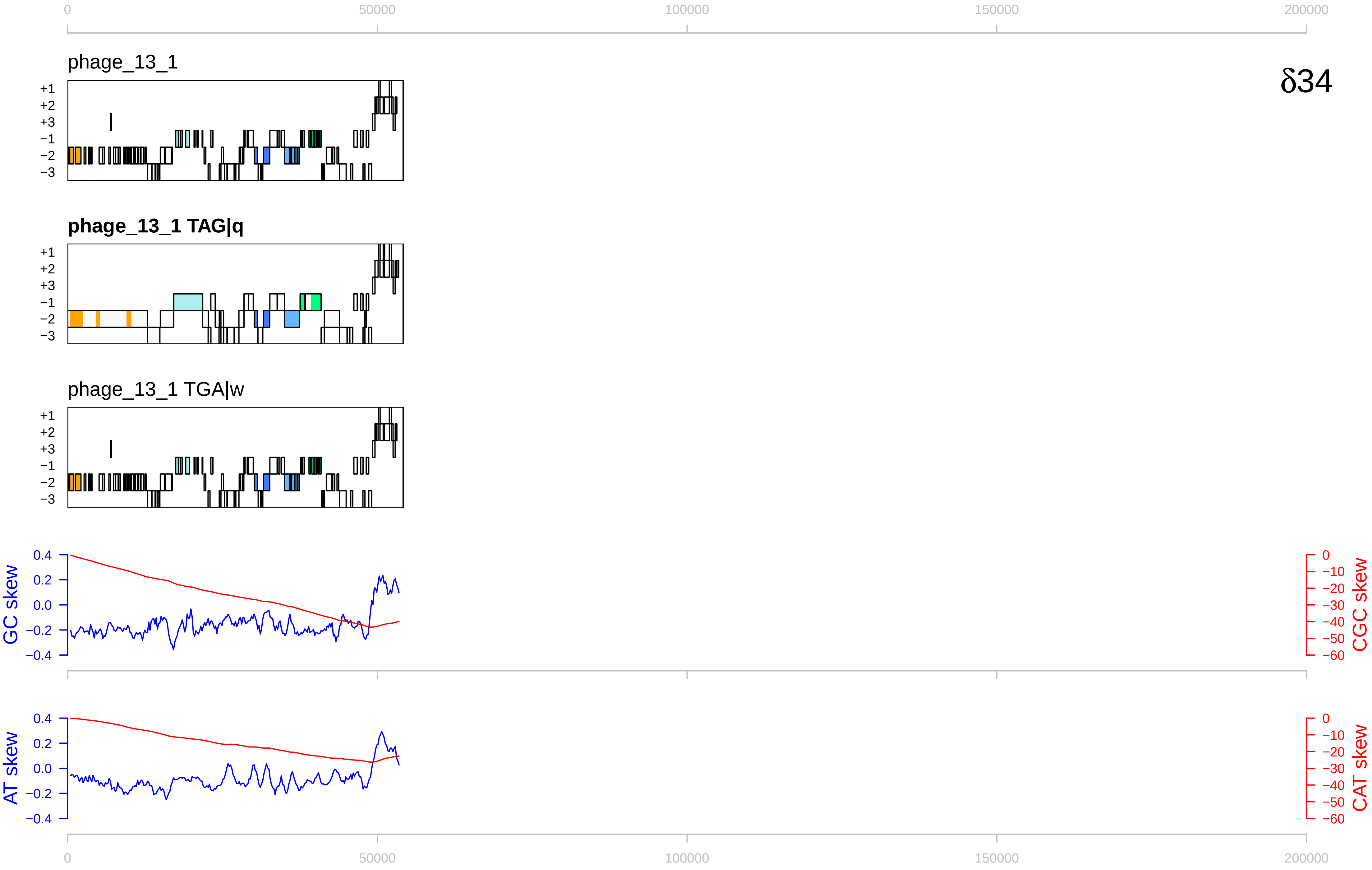

0
50000
100000
150000
200000
phage_13_1
δ34
+1
+2
+3
−1
−2
−3
phage_13_1 TAG|q
phage_13_1 TGA|w
GC skew
CGC skew
AT skew
CAT skew
0.4
0.2
0.0
−0.2
−0.4
0
−10
−20
−30
−40
−50
−60

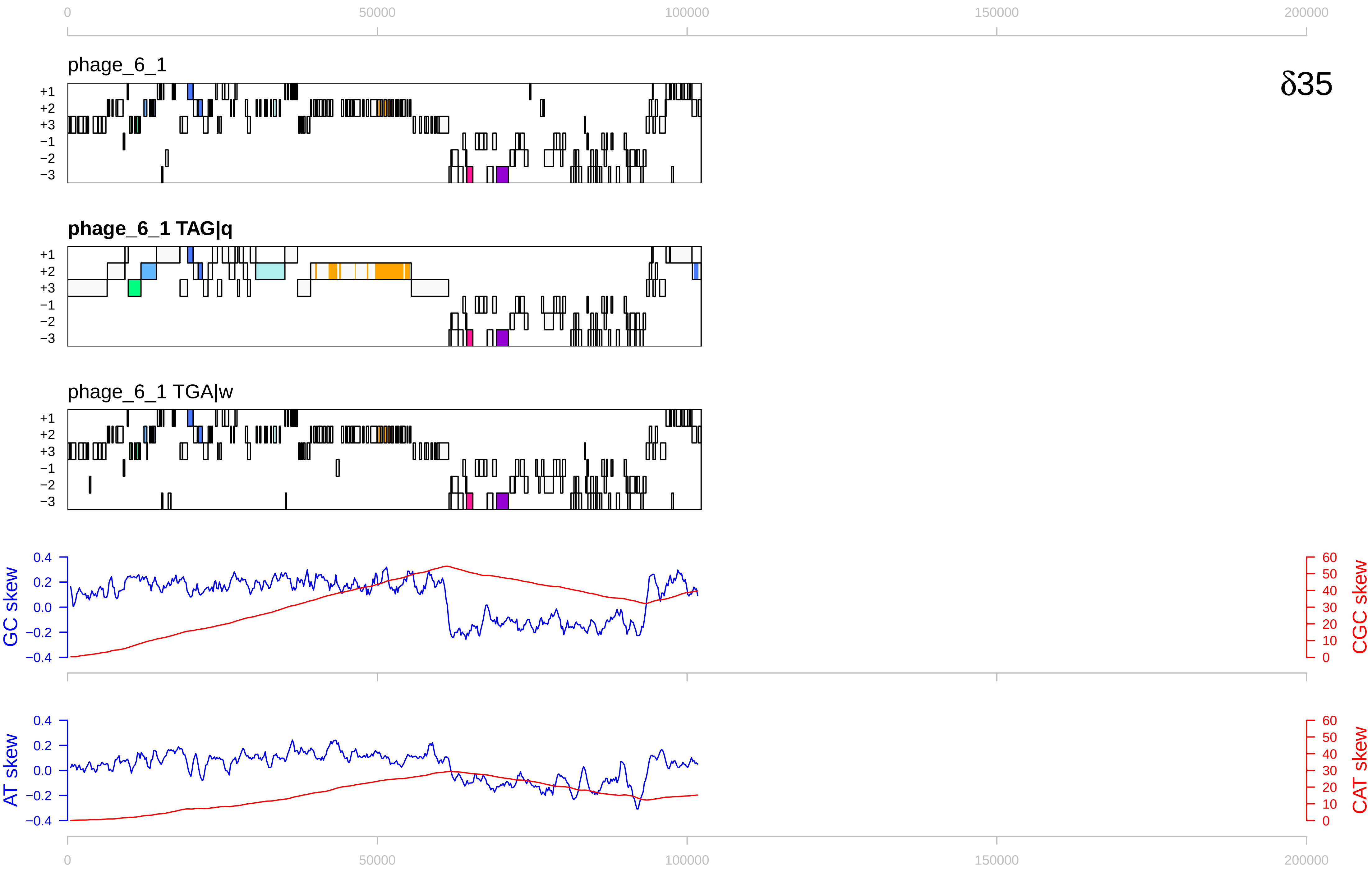
δ35
phage_6_1
phage_6_1 TAG|q
phage_6_1 TGA|w
+1
+2
+3
−1
−2
−3
GC skew
CGC skew
AT skew
CAT skew
0
50000
100000
150000
200000
0.4
0.2
0.0
−0.2
−0.4
60
50
40
30
20
10
0

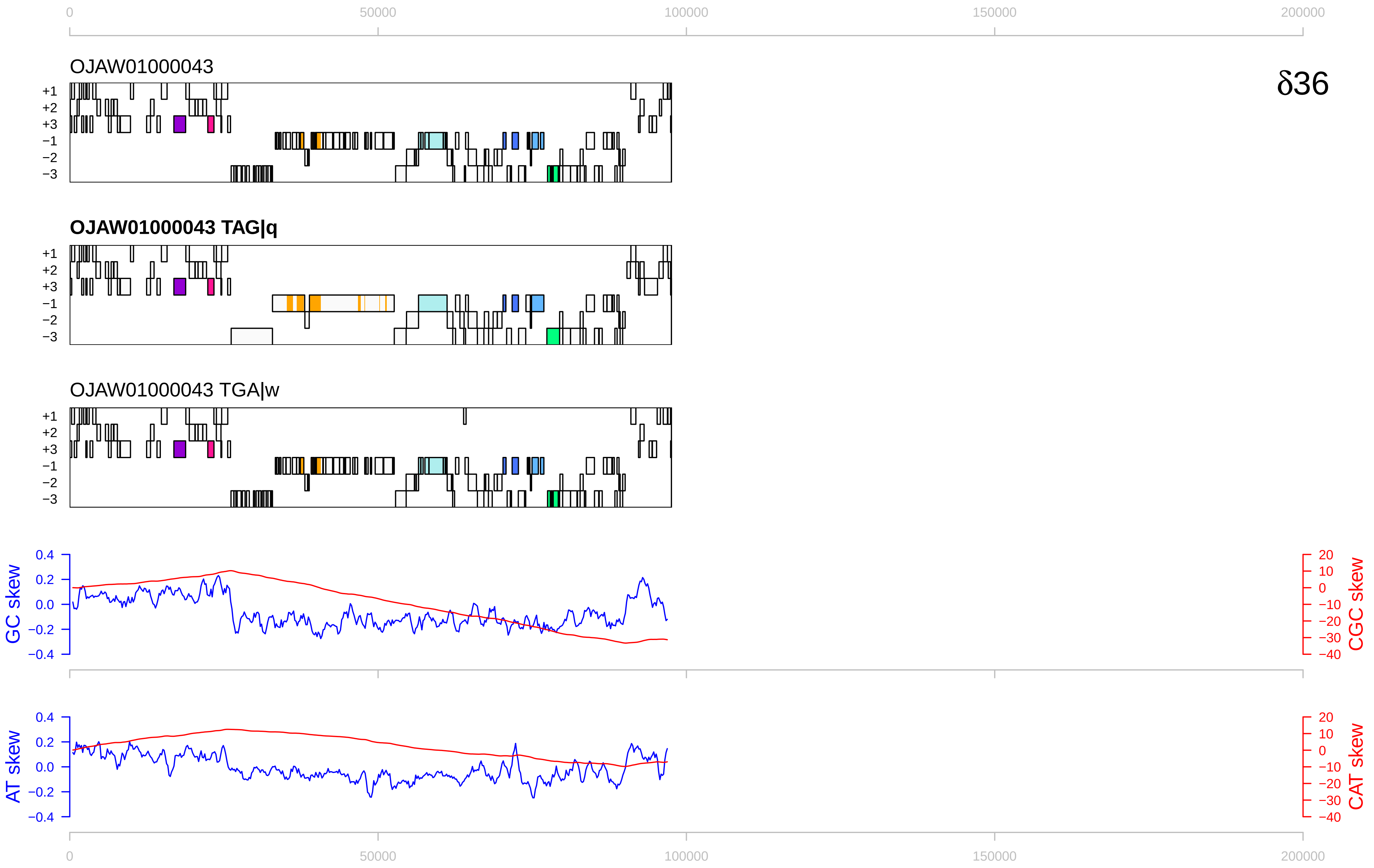

0
50000
100000
150000
200000
OJAW01000043
δ36
+1
+2
+3
−1
−2
−3
OJAW01000043 TAG|q
OJAW01000043 TGA|w
GC skew
CGC skew
0.4
0.2
0.0
−0.2
−0.4
20
10
0
−10
−20
−30
−40
AT skew
CAT skew

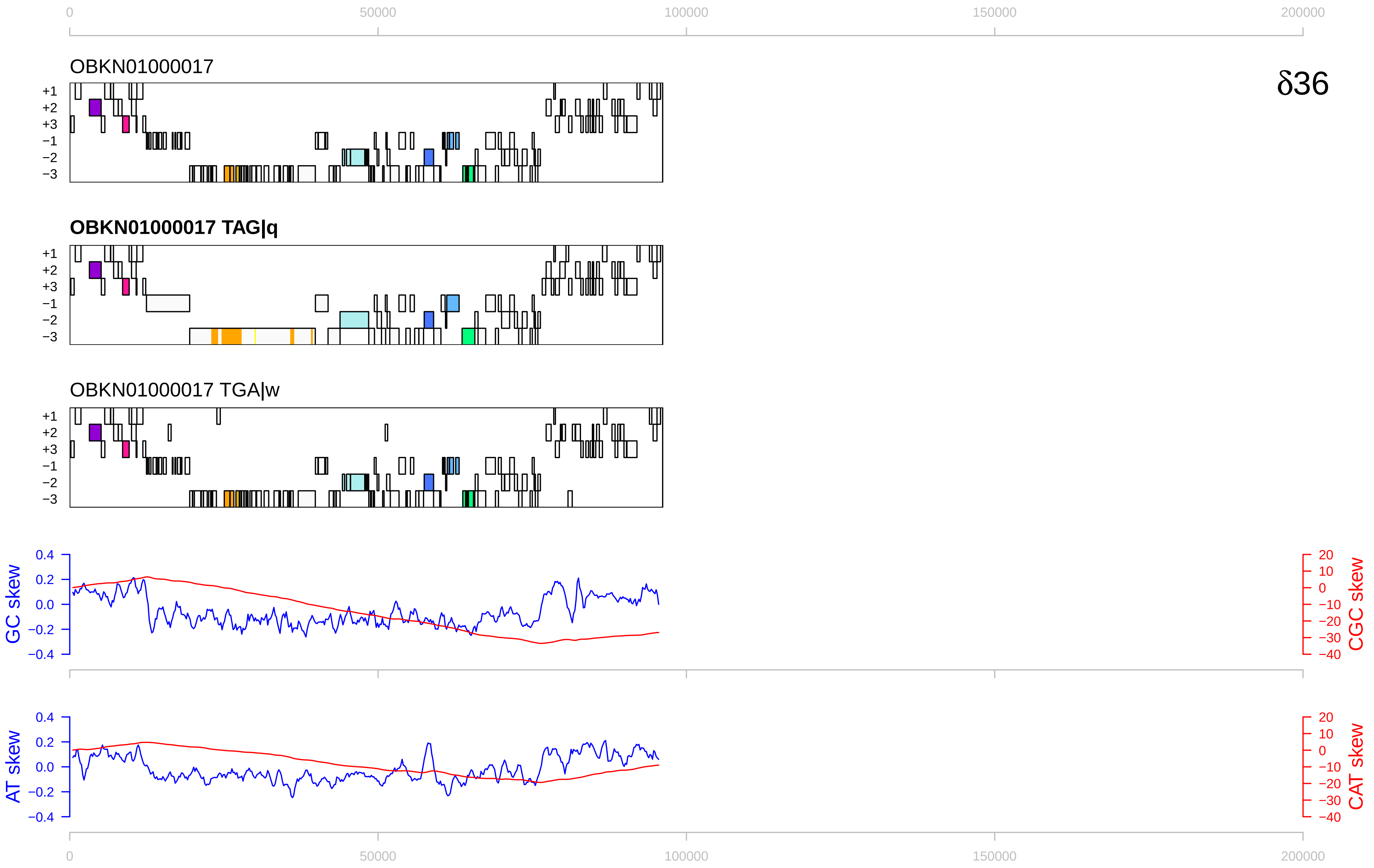
0
50000
100000
150000
200000
OBKN01000017
δ36
+1
+2
+3
−1
−2
−3
OBKN01000017 TAG|q
OBKN01000017 TGA|w
GC skew
CGC skew
AT skew
CAT skew
0.4
0.2
0.0
−0.2
−0.4
20
10
0
−10
−20
−30
−40

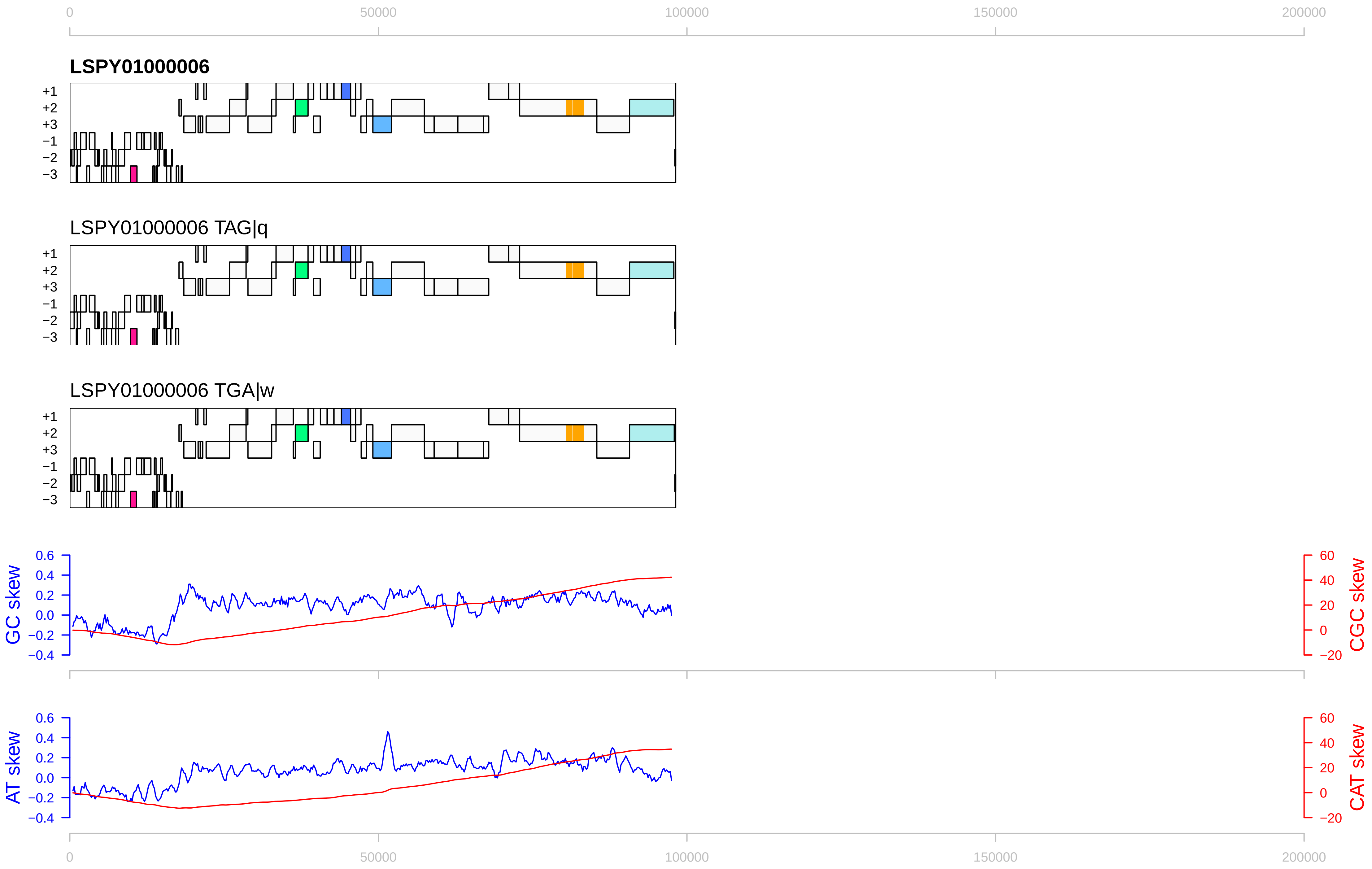

0
50000
100000
150000
200000
LSPY01000006
+1
+2
+3
−1
−2
−3
LSPY01000006 TAG|q
LSPY01000006 TGA|w
GC skew
CGC skew
AT skew
CAT skew
0.6
0.4
0.2
0.0
−0.2
−0.4
60
40
20
0
−20

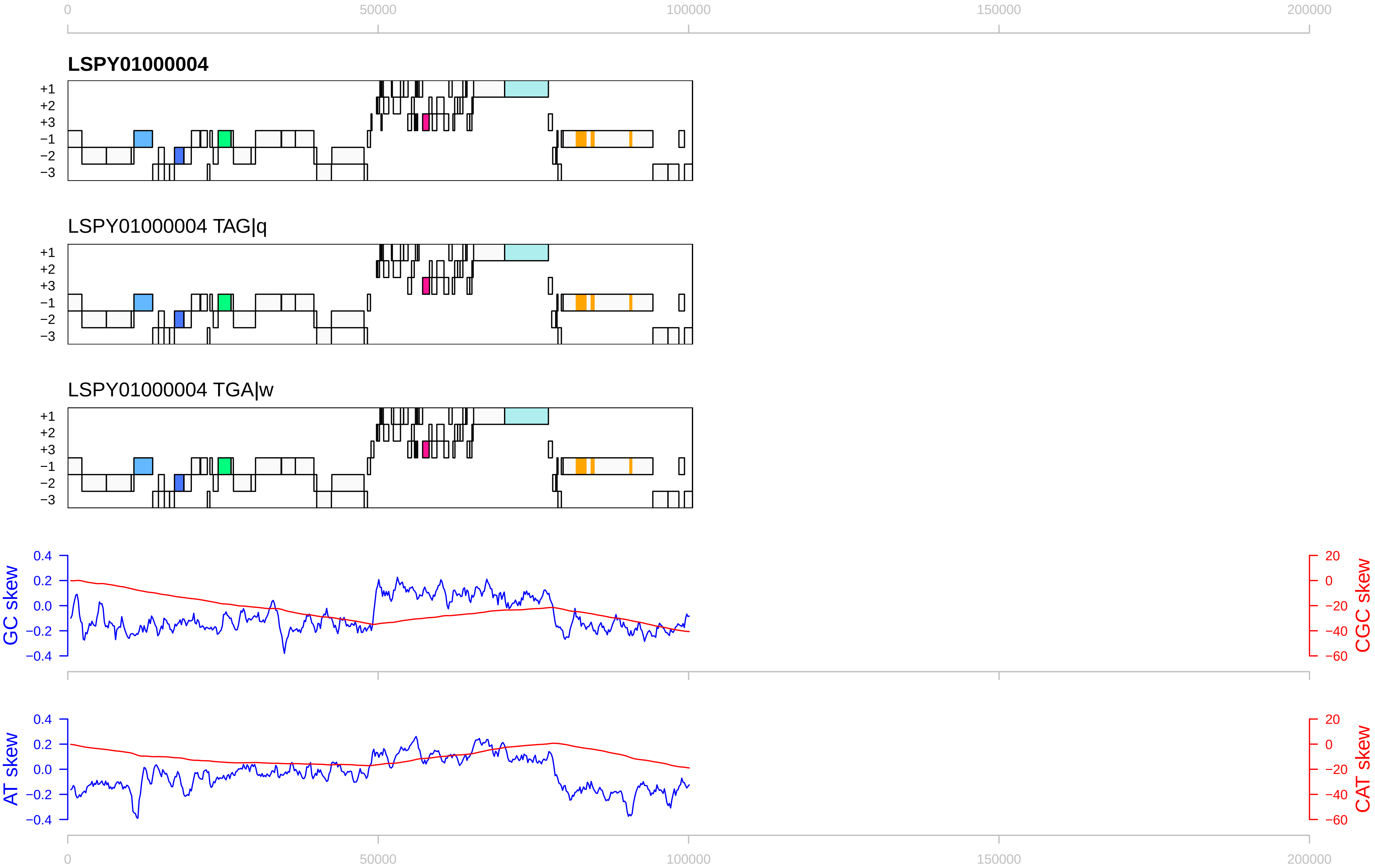

0
50000
100000
150000
200000
LSPY01000004
LSPY01000004 TAG|q
LSPY01000004 TGA|w
+1
+2
+3
−1
−2
−3
GC skew
CGC skew
AT skew
CAT skew
0.4
0.2
0.0
−0.2
−0.4
20
0
−20
−40
−60

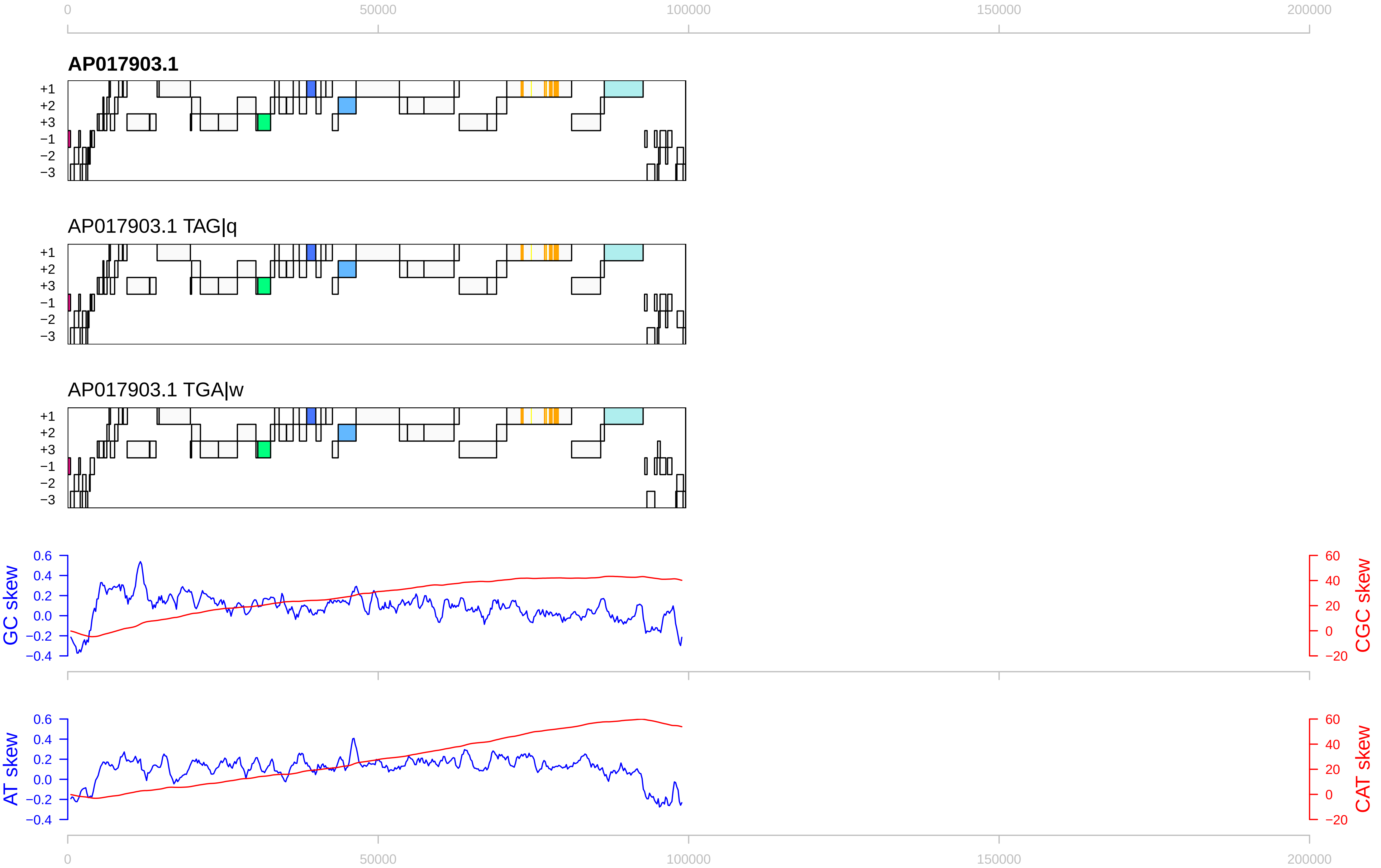

0
50000
100000
150000
200000
AP017903.1
+1
+2
+3
−1
−2
−3
AP017903.1 TAG|q
AP017903.1 TGA|w
GC skew
CGC skew
AT skew
CAT skew
0.6
0.4
0.2
0.0
−0.2
−0.4
60
40
20
0
−20

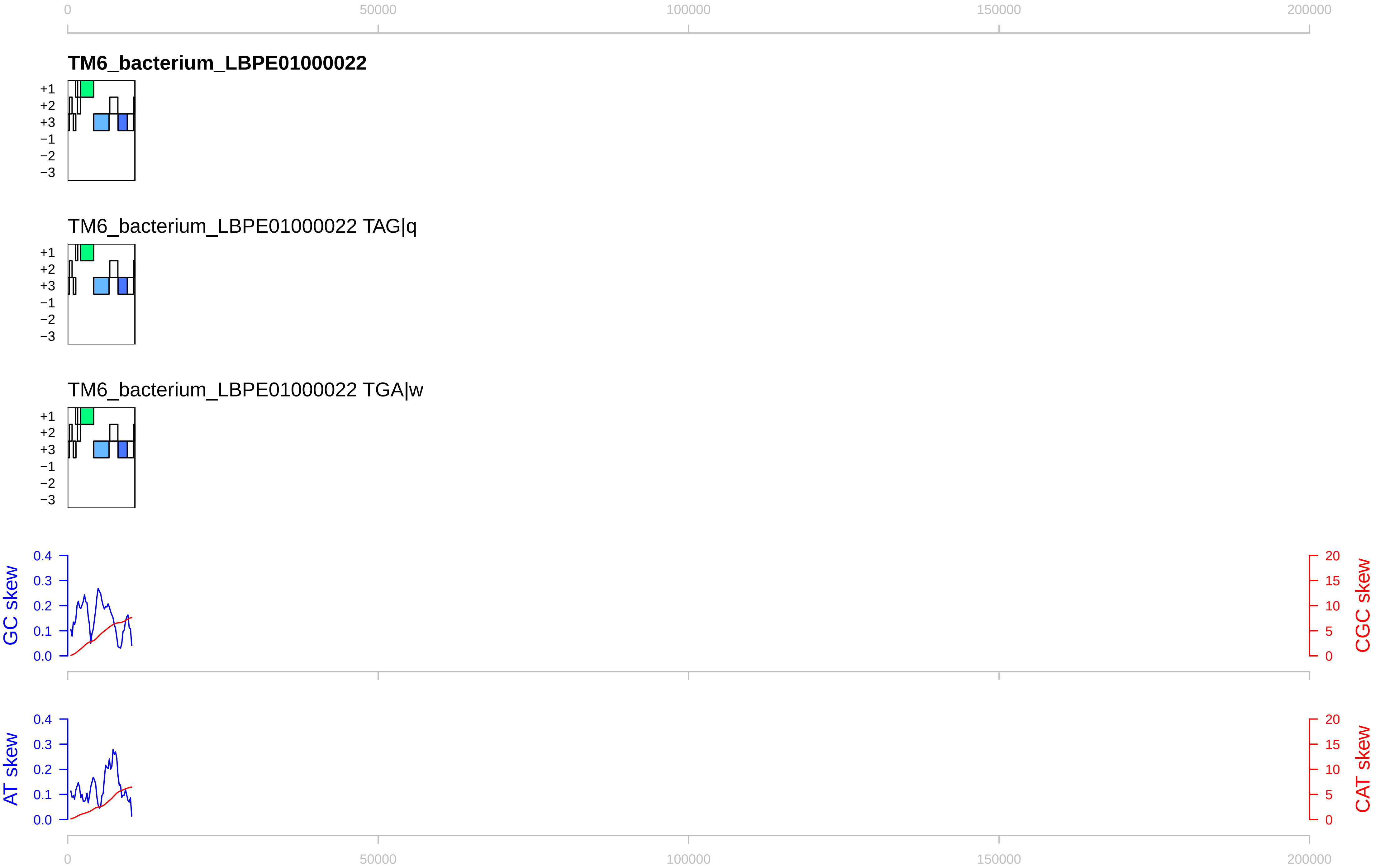

0
50000
100000
150000
200000
TM6_bacterium_LBPE01000022
+1
+2
+3
−1
−2
−3
TM6_bacterium_LBPE01000022 TAG|q
+1
+2
+3
−1
−2
−3
TM6_bacterium_LBPE01000022 TGA|w
+1
+2
+3
−1
−2
−3
GC skew
0.4
0.3
0.2
0.1
0.0
CGC skew
20
15
10
5
0
AT skew
0.4
0.3
0.2
0.1
0.0
CAT skew
20
15
10
5
0
0
50000
100000
150000
200000

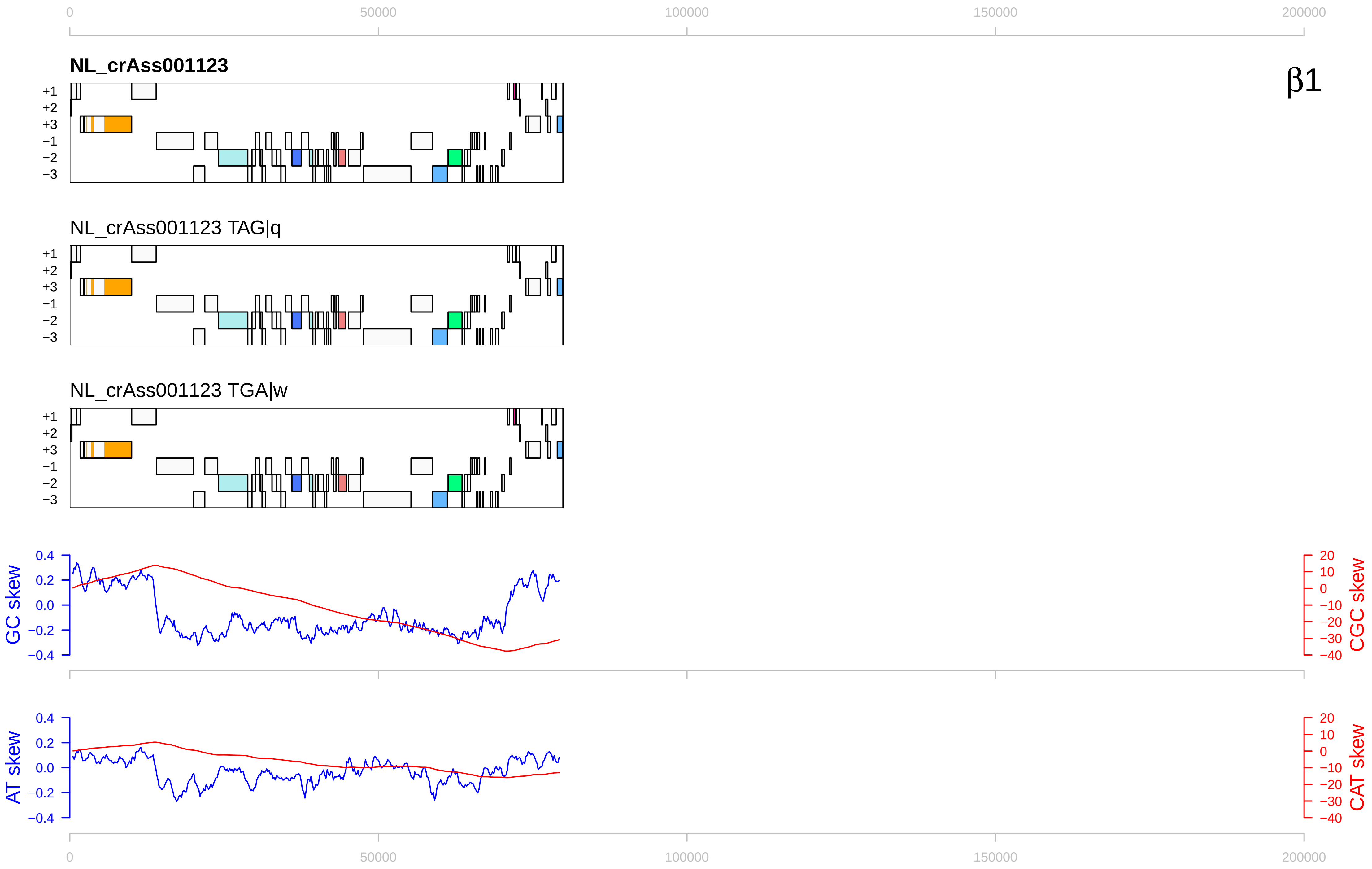

0
50000
100000
150000
200000
NL_crAss001123
β1
+1
+2
+3
−1
−2
−3
NL_crAss001123 TAG|q
NL_crAss001123 TGA|w
GC skew
0.4
0.2
0.0
−0.2
−0.4
CGC skew
20
10
0
−10
−20
−30
−40
AT skew
CAT skew

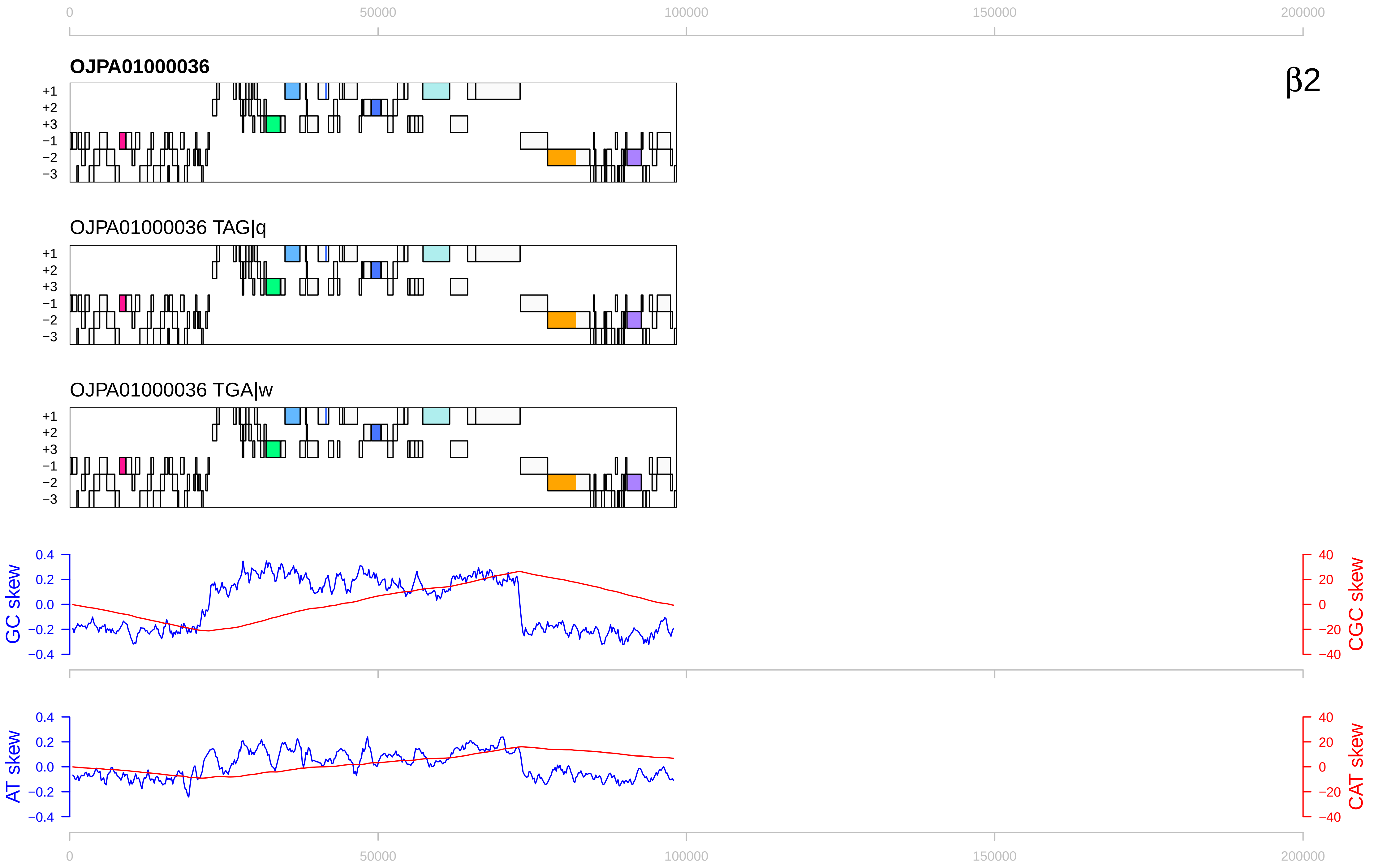

0
50000
100000
150000
200000
OJPA01000036
β2
+1
+2
+3
−1
−2
−3
OJPA01000036 TAG|q
OJPA01000036 TGA|w
GC skew
CGC skew
0.4
0.2
0.0
−0.2
−0.4
40
20
0
−20
−40
AT skew
CAT skew

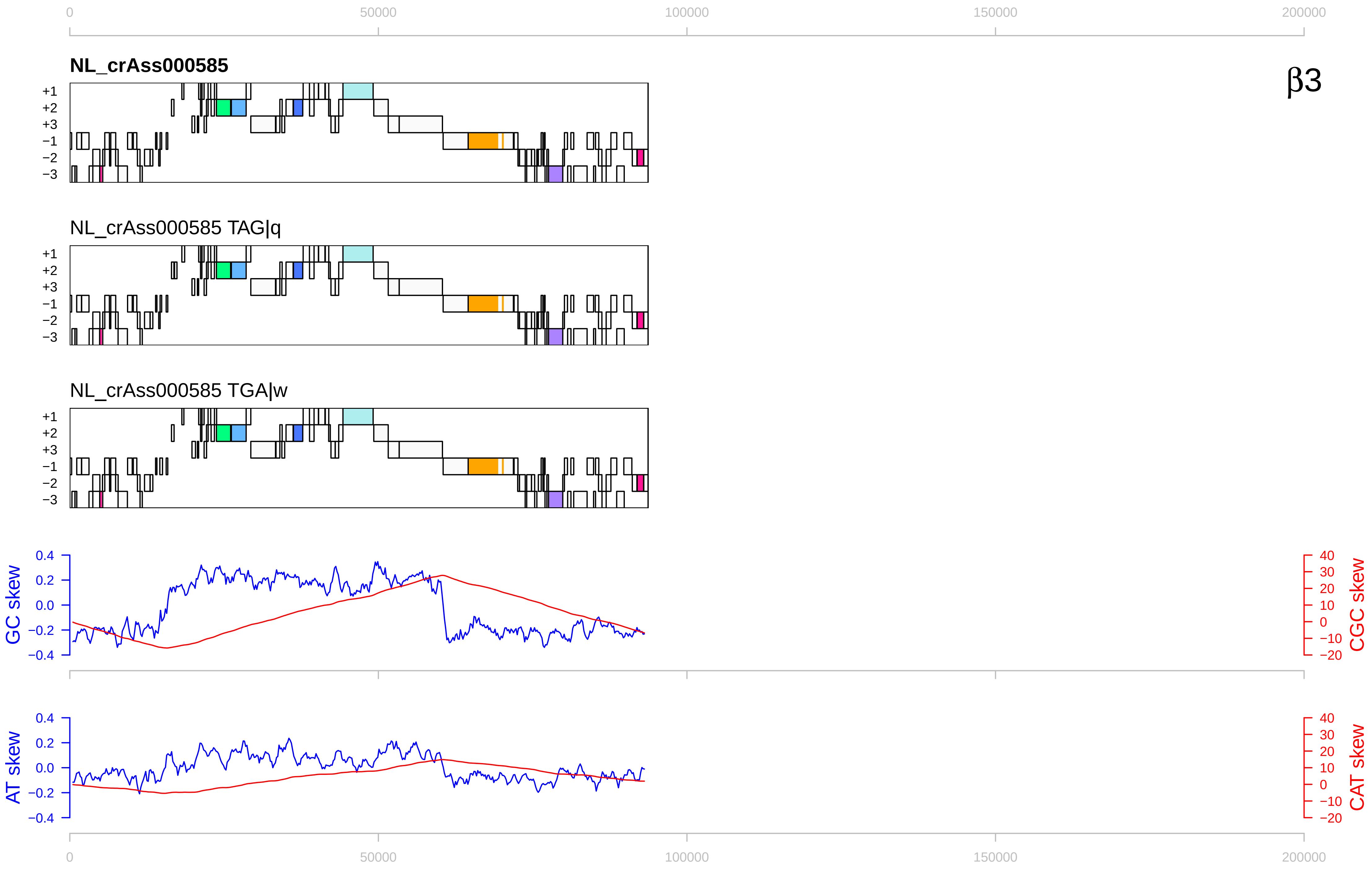

0
50000
100000
150000
200000
NL_crAss000585
β3
+1
+2
+3
−1
−2
−3
NL_crAss000585 TAG|q
NL_crAss000585 TGA|w
GC skew
CGC skew
AT skew
CAT skew
0.4
0.2
0.0
−0.2
−0.4
40
30
20
10
0
−10
−20

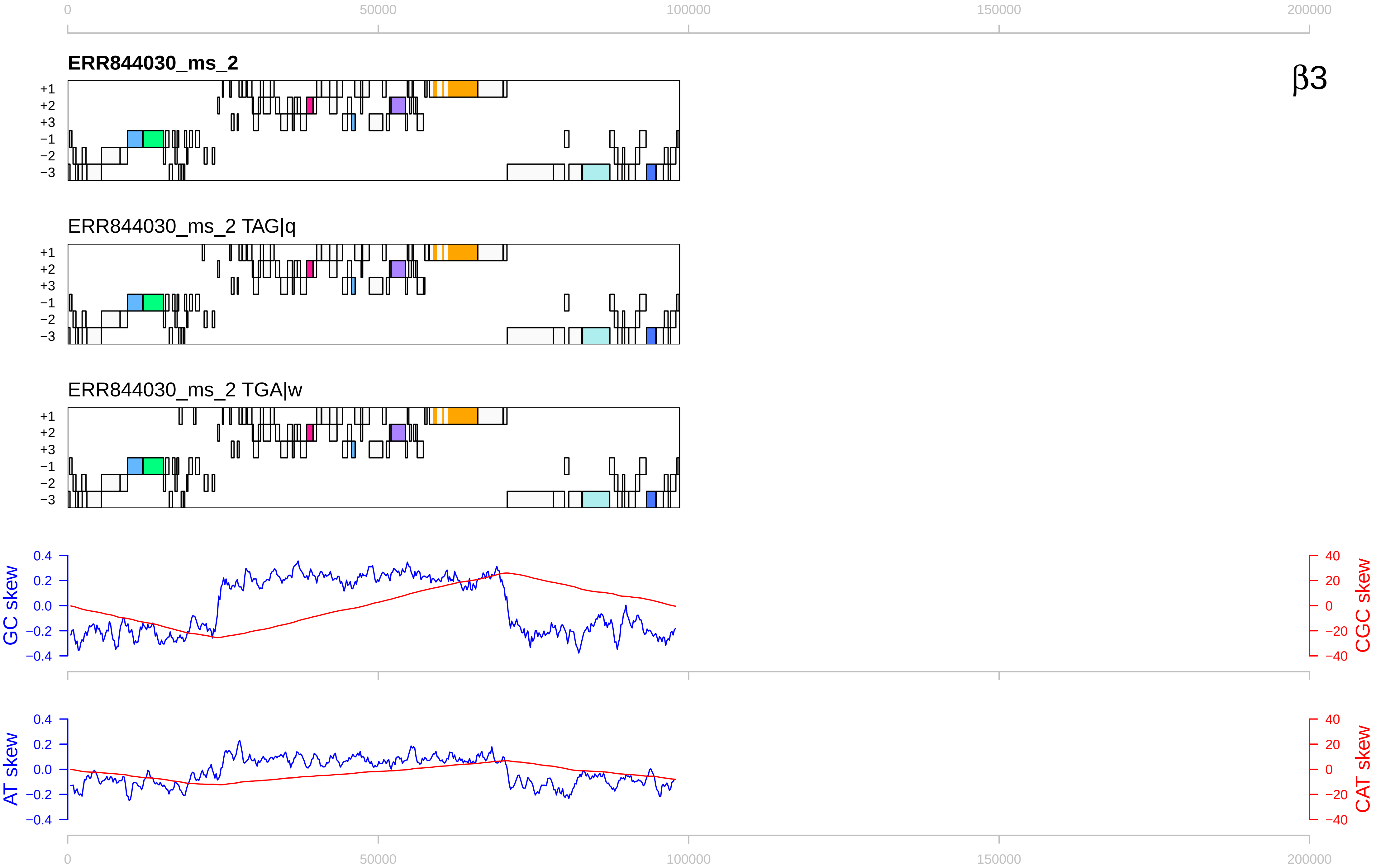
0
50000
100000
150000
200000
ERR844030_ms_2
β3
+1
+2
+3
−1
−2
−3
ERR844030_ms_2 TAG|q
ERR844030_ms_2 TGA|w
GC skew
CGC skew
AT skew
CAT skew
0.4
0.2
0.0
−0.2
−0.4
40
20
0
−20
−40

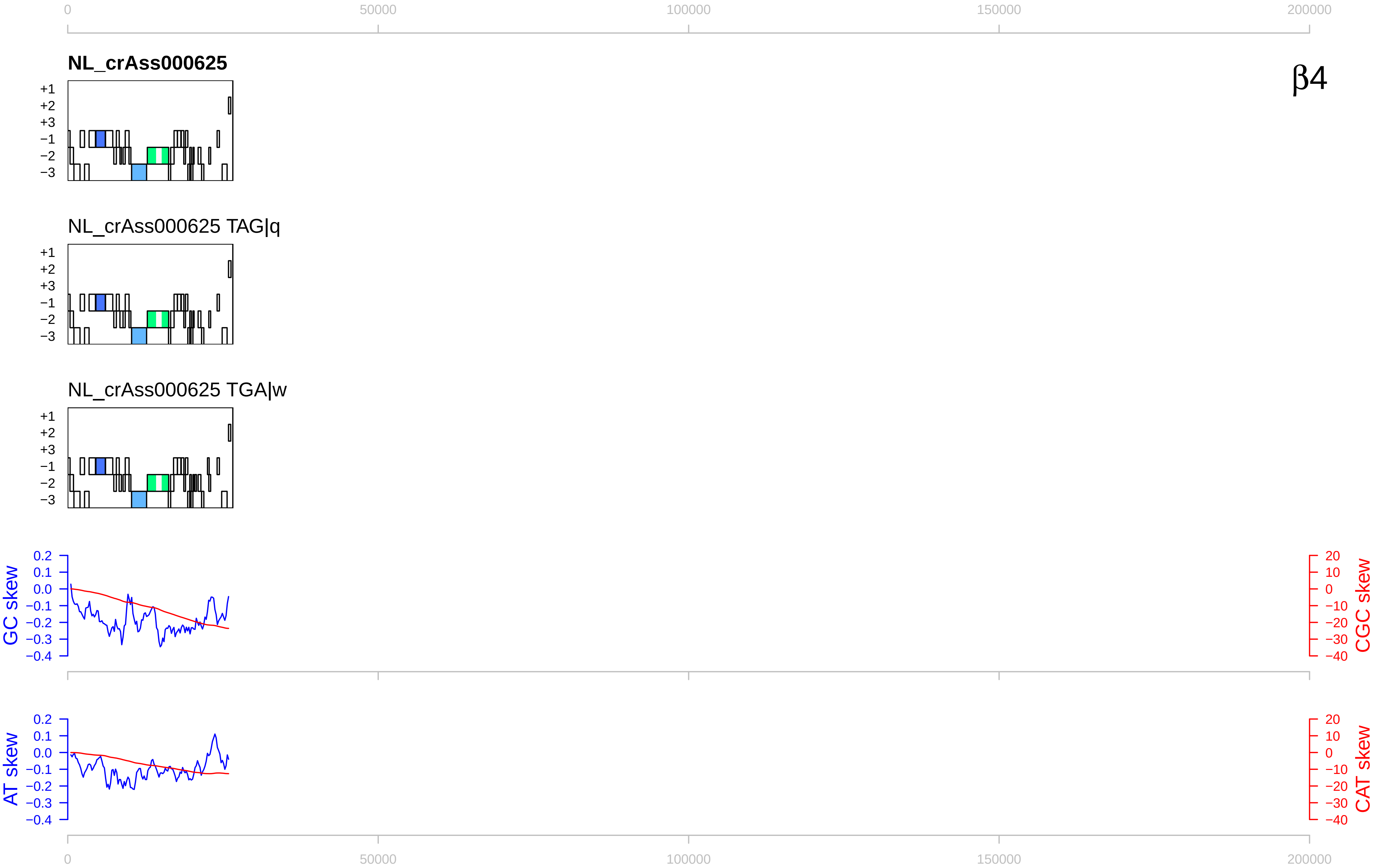

0
50000
100000
150000
200000
NL_crAss000625
β4
+1
+2
+3
−1
−2
−3
NL_crAss000625 TAG|q
NL_crAss000625 TGA|w
GC skew
CGC skew
AT skew
CAT skew
0.2
0.1
0.0
−0.1
−0.2
−0.3
−0.4
20
10
0
−10
−20
−30
−40

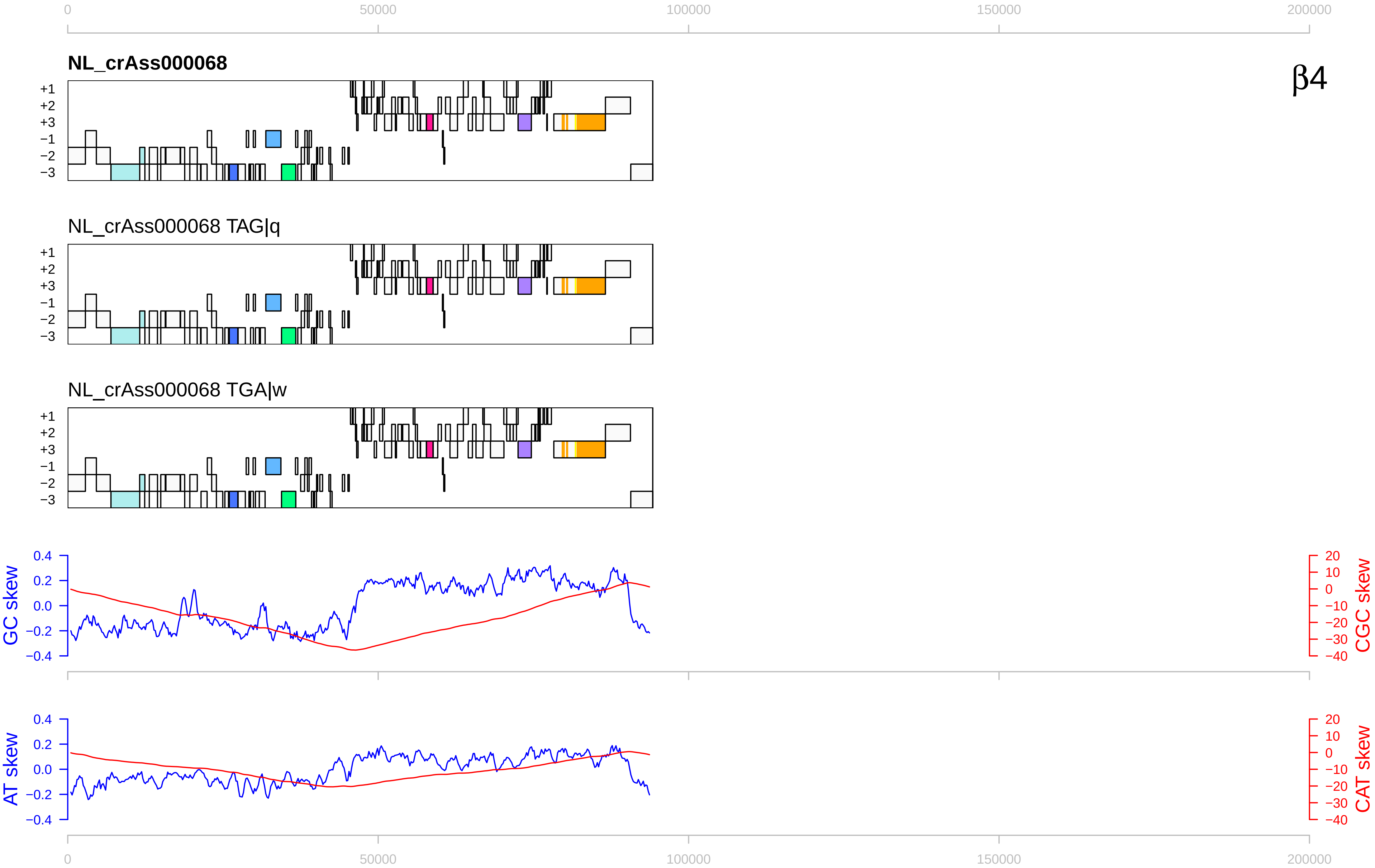

0
50000
100000
150000
200000
NL_crAss000068
β4
+1
+2
+3
−1
−2
−3
NL_crAss000068 TAG|q
NL_crAss000068 TGA|w
GC skew
CGC skew
AT skew
CAT skew
0.4
0.2
0.0
−0.2
−0.4
20
10
0
−10
−20
−30
−40

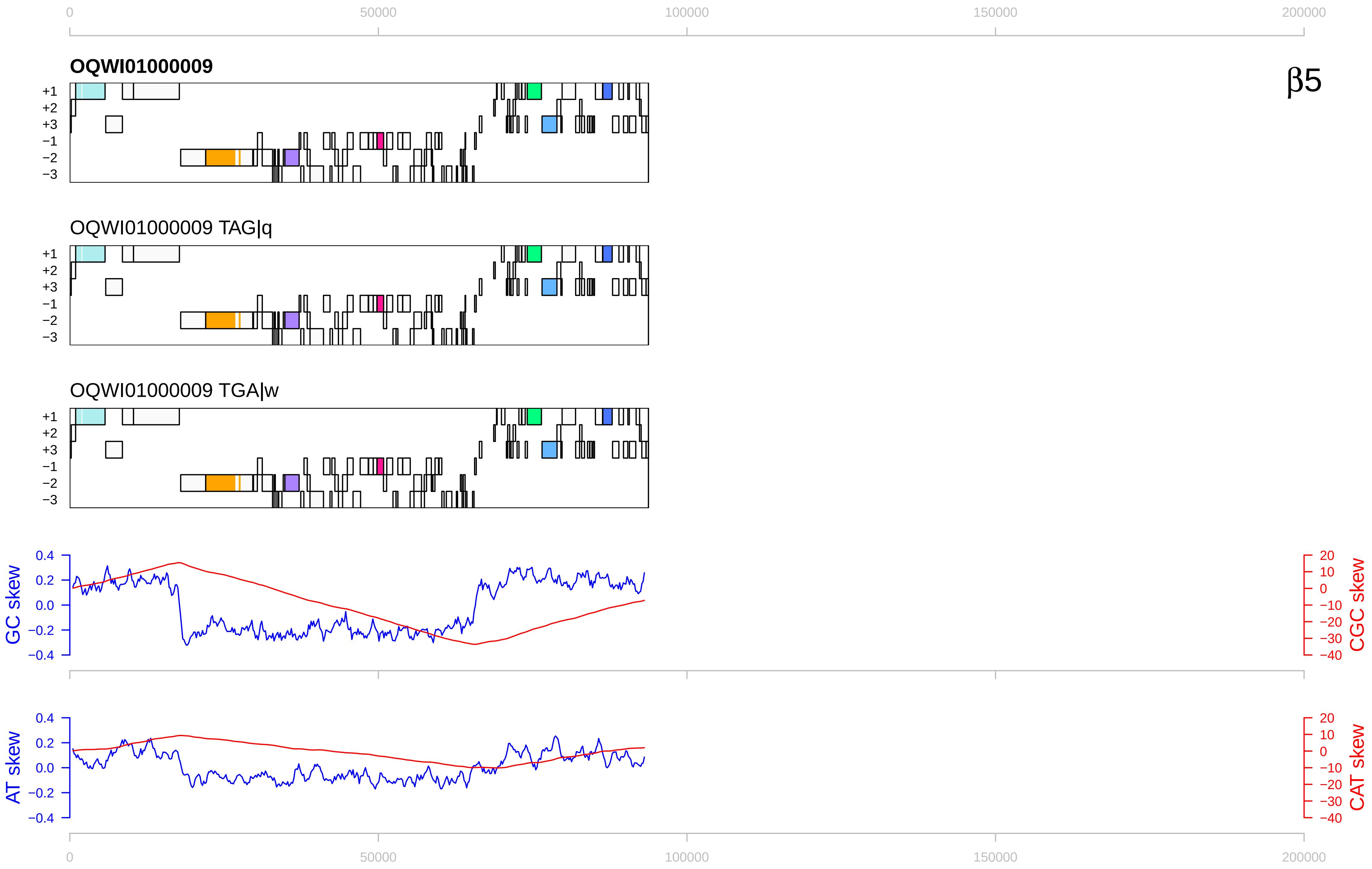

0
50000
100000
150000
200000
OQWI01000009
β5
+1
+2
+3
−1
−2
−3
OQWI01000009 TAG|q
OQWI01000009 TGA|w
GC skew
CGC skew
AT skew
CAT skew
0.4
0.2
0.0
−0.2
−0.4
20
10
0
−10
−20
−30
−40

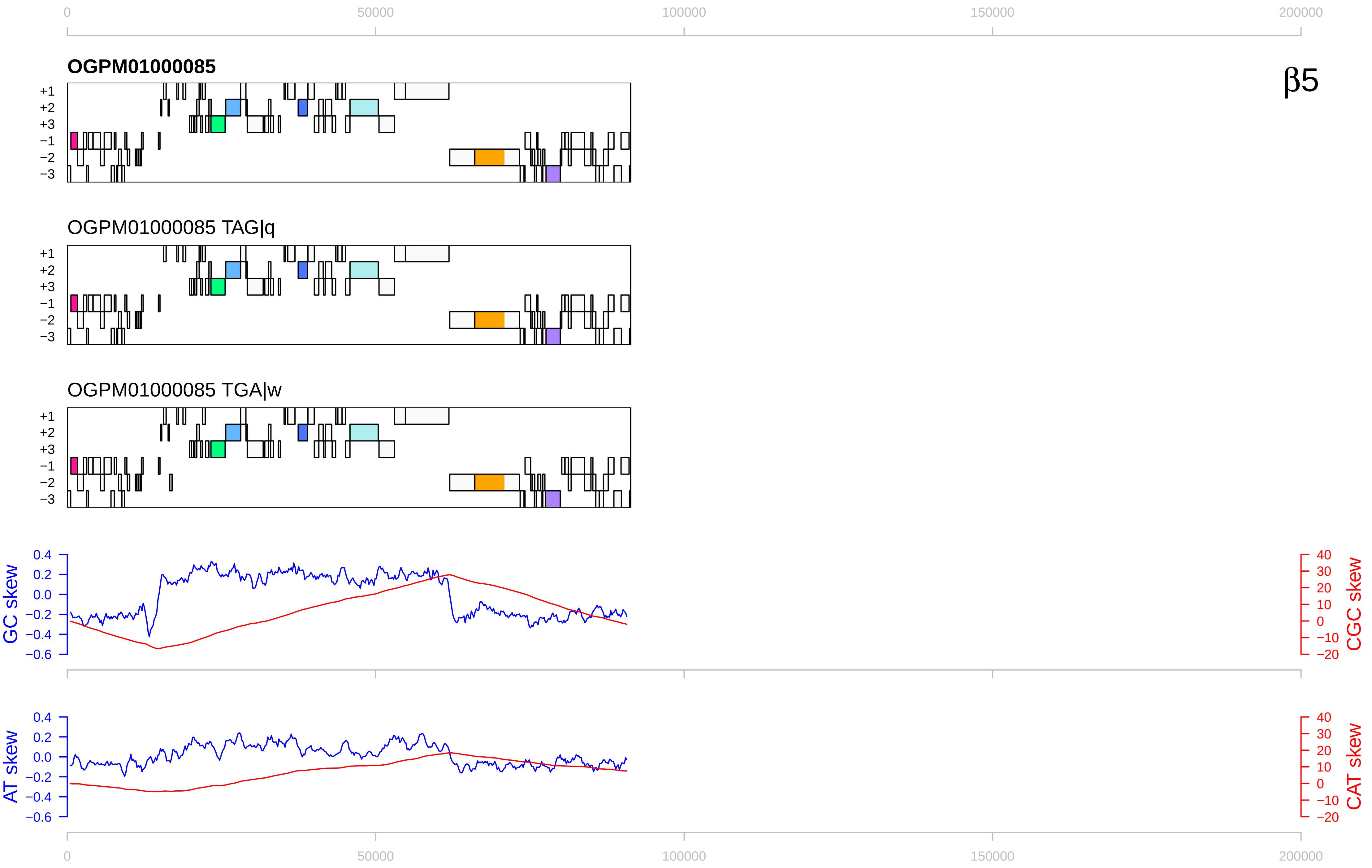

0
50000
100000
150000
200000
OGPM01000085
β5
+1
+2
+3
−1
−2
−3
OGPM01000085 TAG|q
OGPM01000085 TGA|w
GC skew
CGC skew
0.4
0.2
0.0
−0.2
−0.4
−0.6
40
30
20
10
0
−10
−20
AT skew
CAT skew

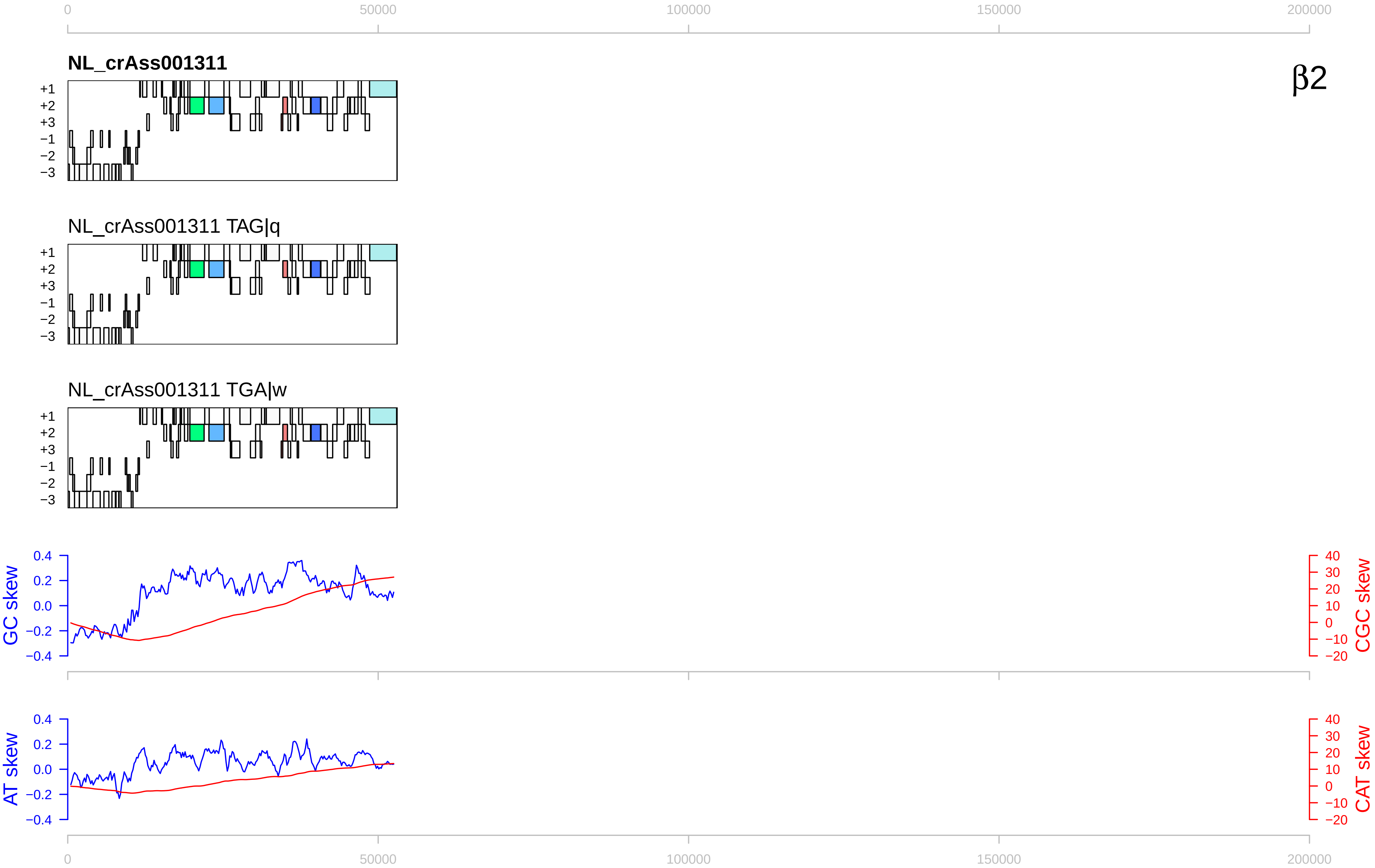

NL_crAss001311
β2
NL_crAss001311 TAG|q
NL_crAss001311 TGA|w
+1
+2
+3
−1
−2
−3
GC skew
CGC skew
AT skew
CAT skew
0
50000
100000
150000
200000

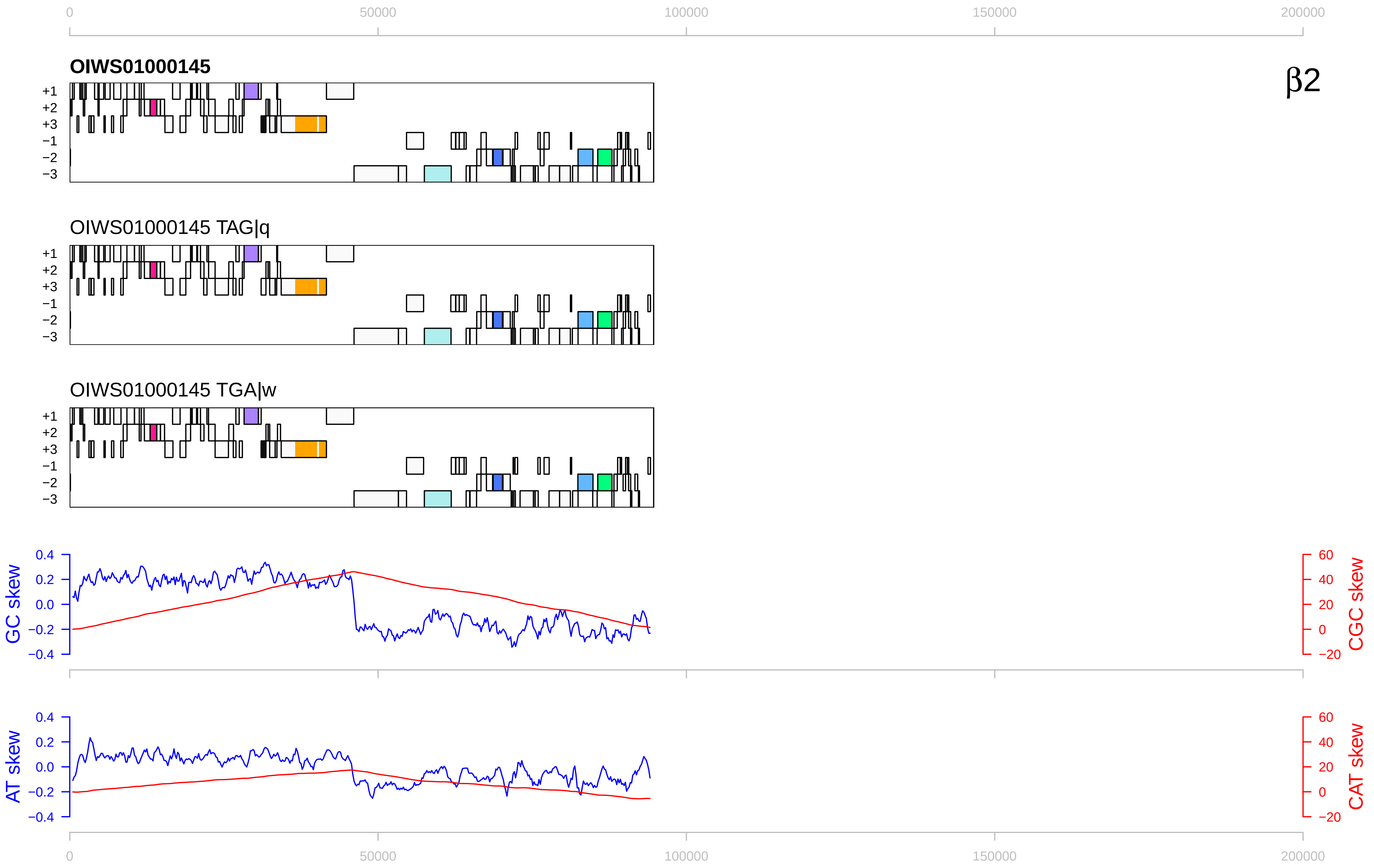

0
50000
100000
150000
200000
β2
OIWS01000145
OIWS01000145 TAG|q
OIWS01000145 TGA|w
+1
+2
+3
−1
−2
−3
GC skew
CGC skew
AT skew
CAT skew
0.4
0.2
0.0
−0.2
−0.4
60
40
20
0
−20

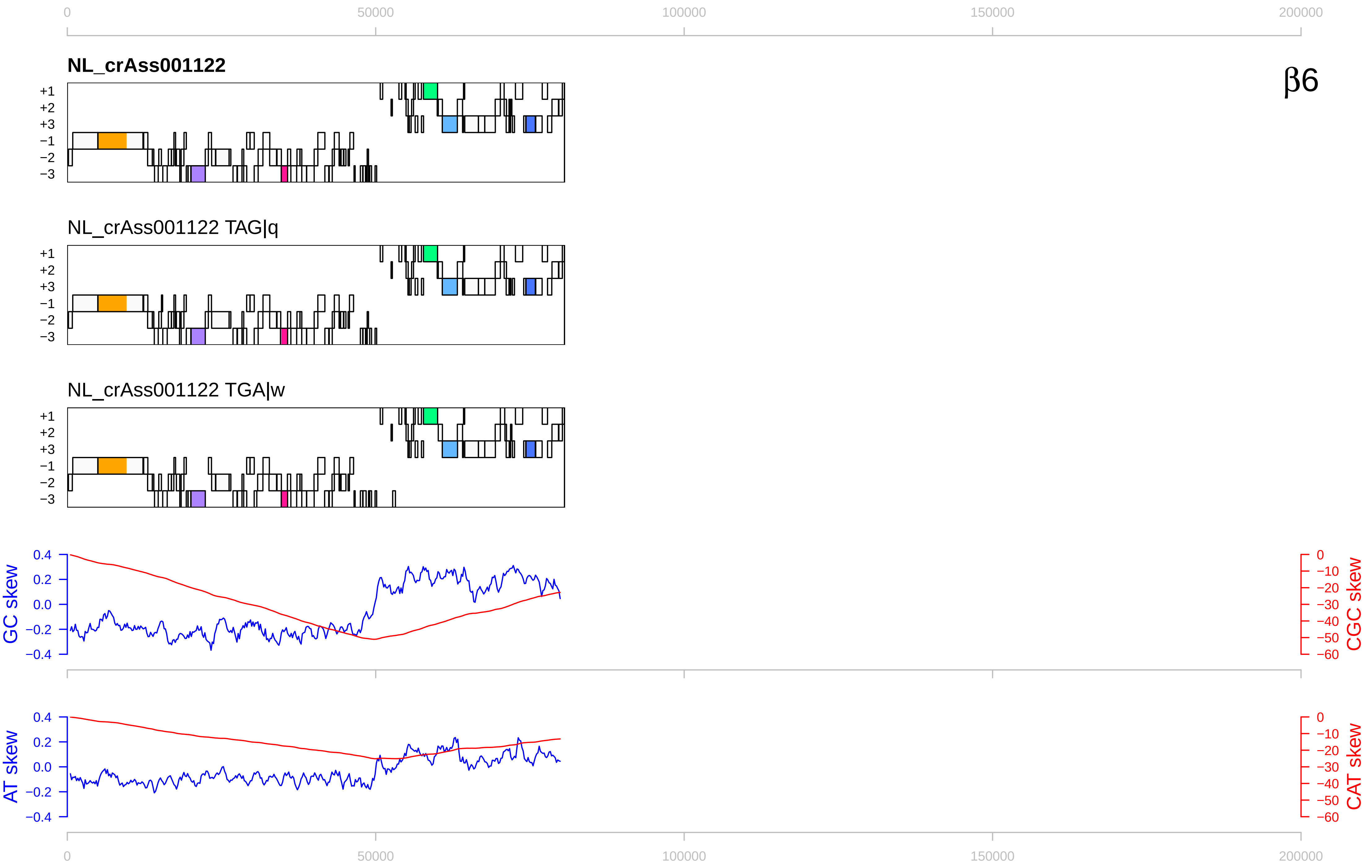

0
50000
100000
150000
200000
NL_crAss001122
β6
+1
+2
+3
−1
−2
−3
NL_crAss001122 TAG|q
NL_crAss001122 TGA|w
GC skew
CGC skew
AT skew
CAT skew
0.4
0.2
0.0
−0.2
−0.4
0
−10
−20
−30
−40
−50
−60

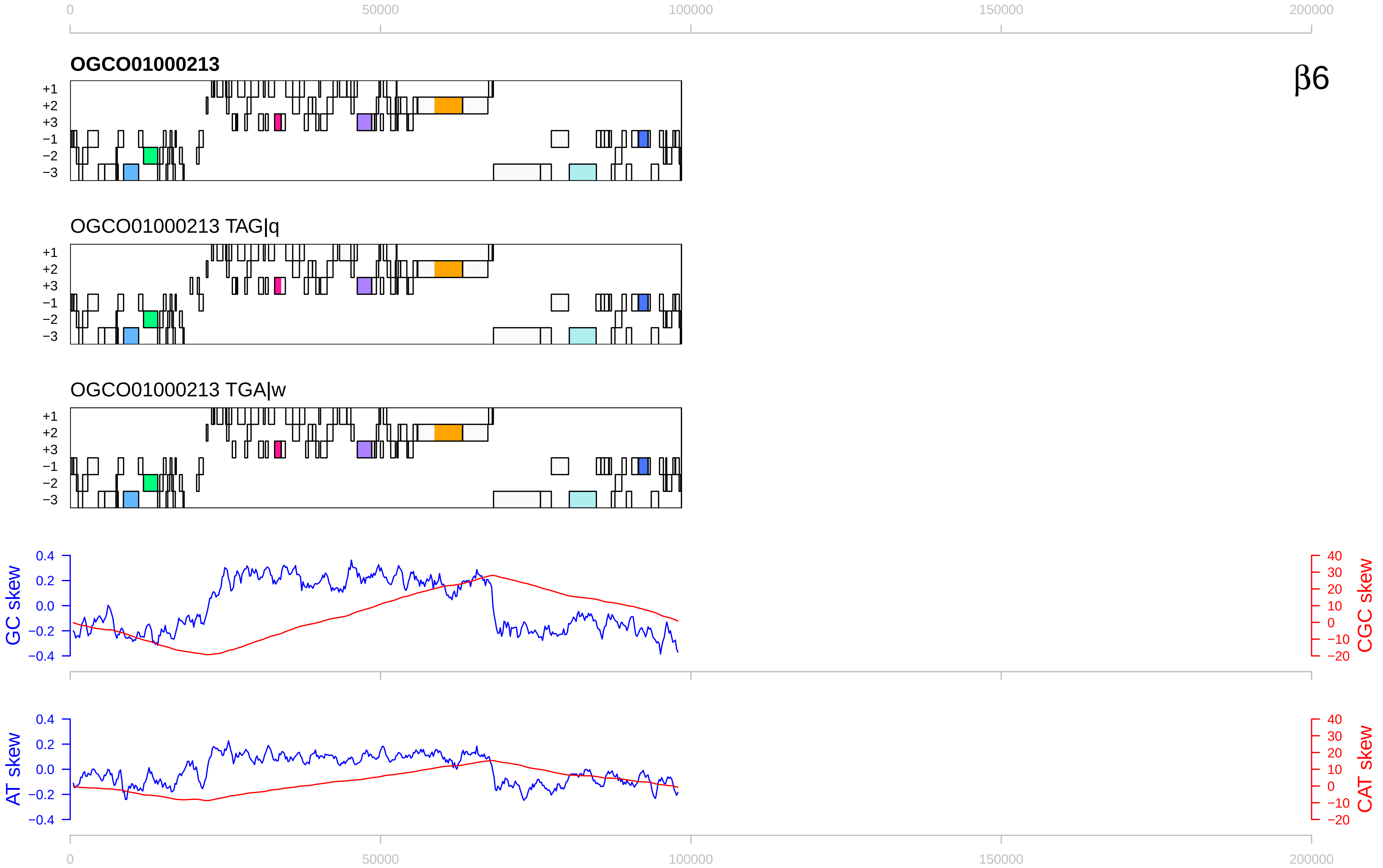

0
50000
100000
150000
200000
OGCO01000213
β6
+1
+2
+3
−1
−2
−3
OGCO01000213 TAG|q
OGCO01000213 TGA|w
GC skew
CGC skew
AT skew
CAT skew
0.4
0.2
0.0
−0.2
−0.4
40
30
20
10
0
−10
−20

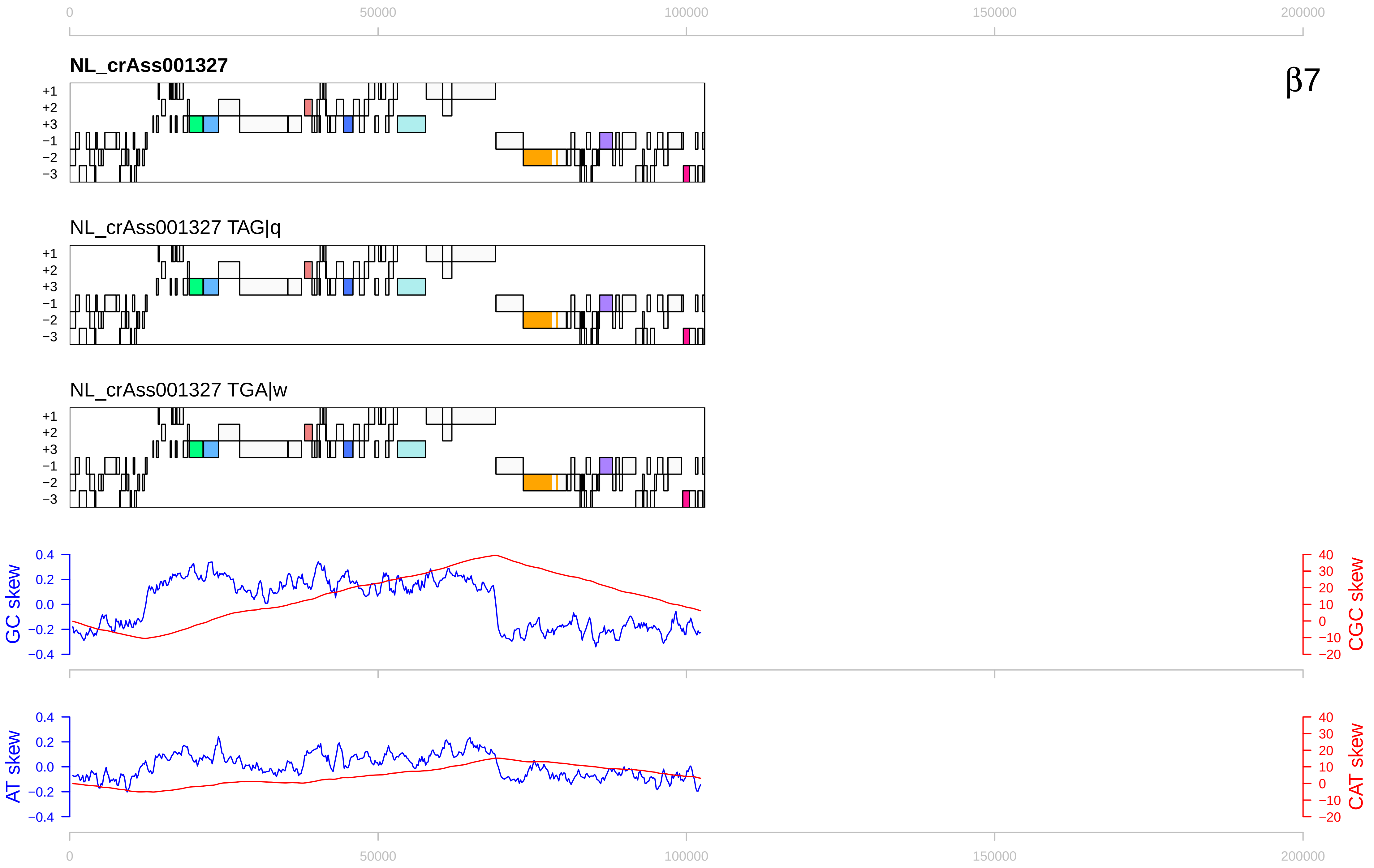

0
50000
100000
150000
200000
NL_crAss001327
β7
+1
+2
+3
−1
−2
−3
NL_crAss001327 TAG|q
NL_crAss001327 TGA|w
GC skew
CGC skew
AT skew
CAT skew
0.4
0.2
0.0
−0.2
−0.4
40
30
20
10
0
−10
−20

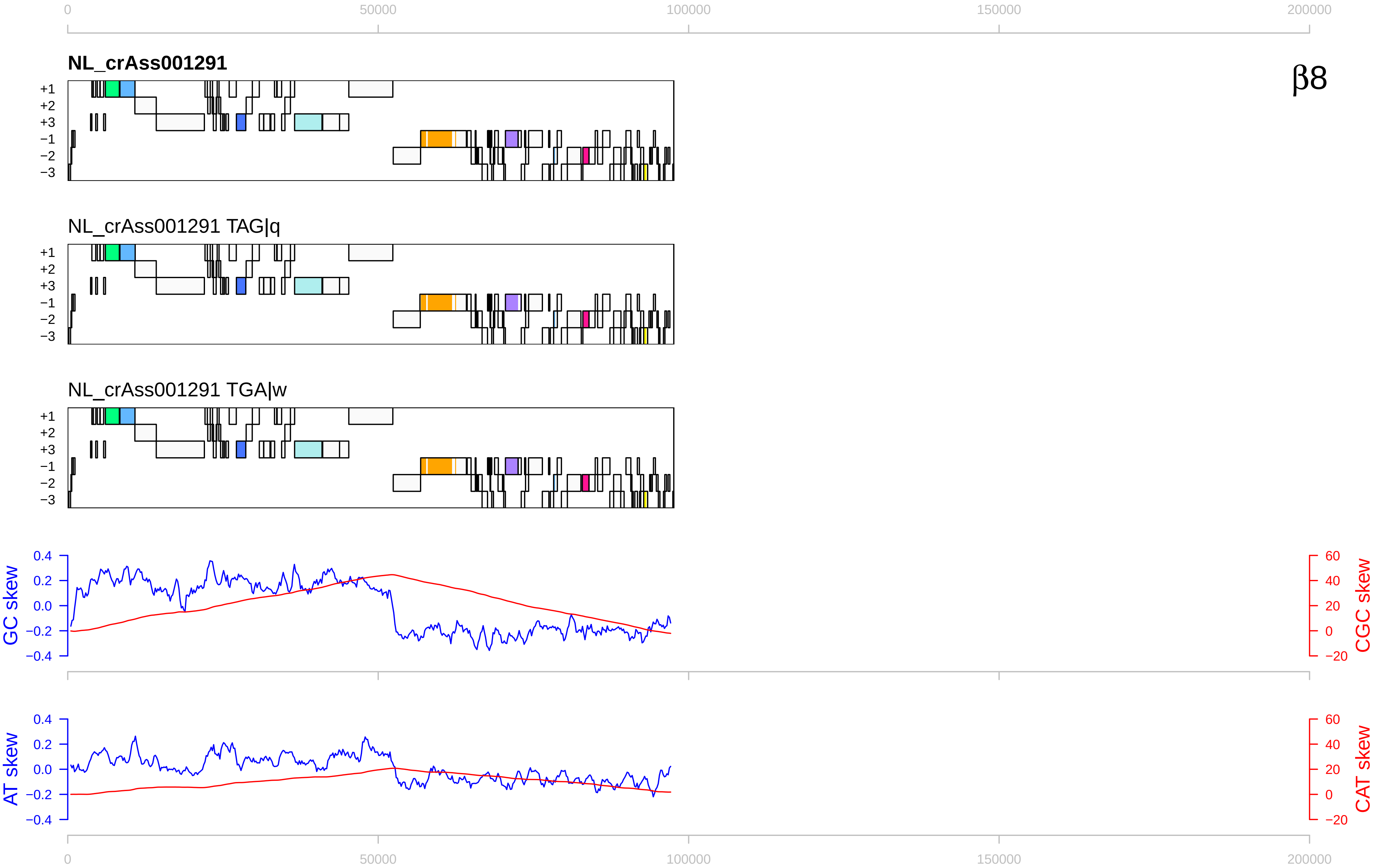
β8
NL_crAss001291
NL_crAss001291 TAG|q
NL_crAss001291 TGA|w
+1
+2
+3
−1
−2
−3
GC skew
CGC skew
AT skew
CAT skew
0
50000
100000
150000
200000
0.4
0.2
0.0
−0.2
−0.4
60
40
20
0
−20

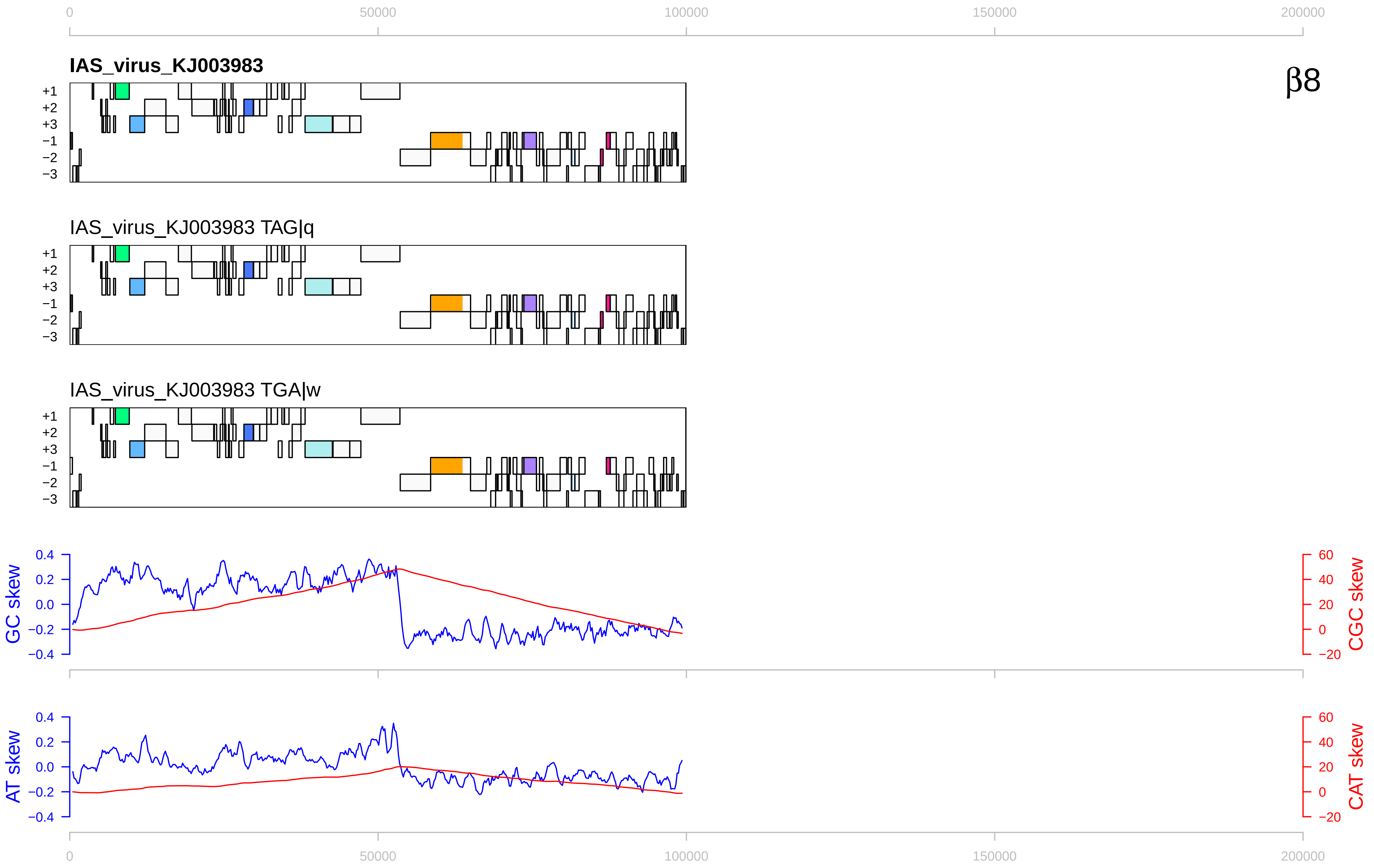

β8
IAS_virus_KJ003983
IAS_virus_KJ003983 TAG|q
IAS_virus_KJ003983 TGA|w
+1
+2
+3
−1
−2
−3
GC skew
CGC skew
AT skew
CAT skew
0
50000
100000
150000
200000
0.4
0.2
0.0
−0.2
−0.4
60
40
20
0
−20

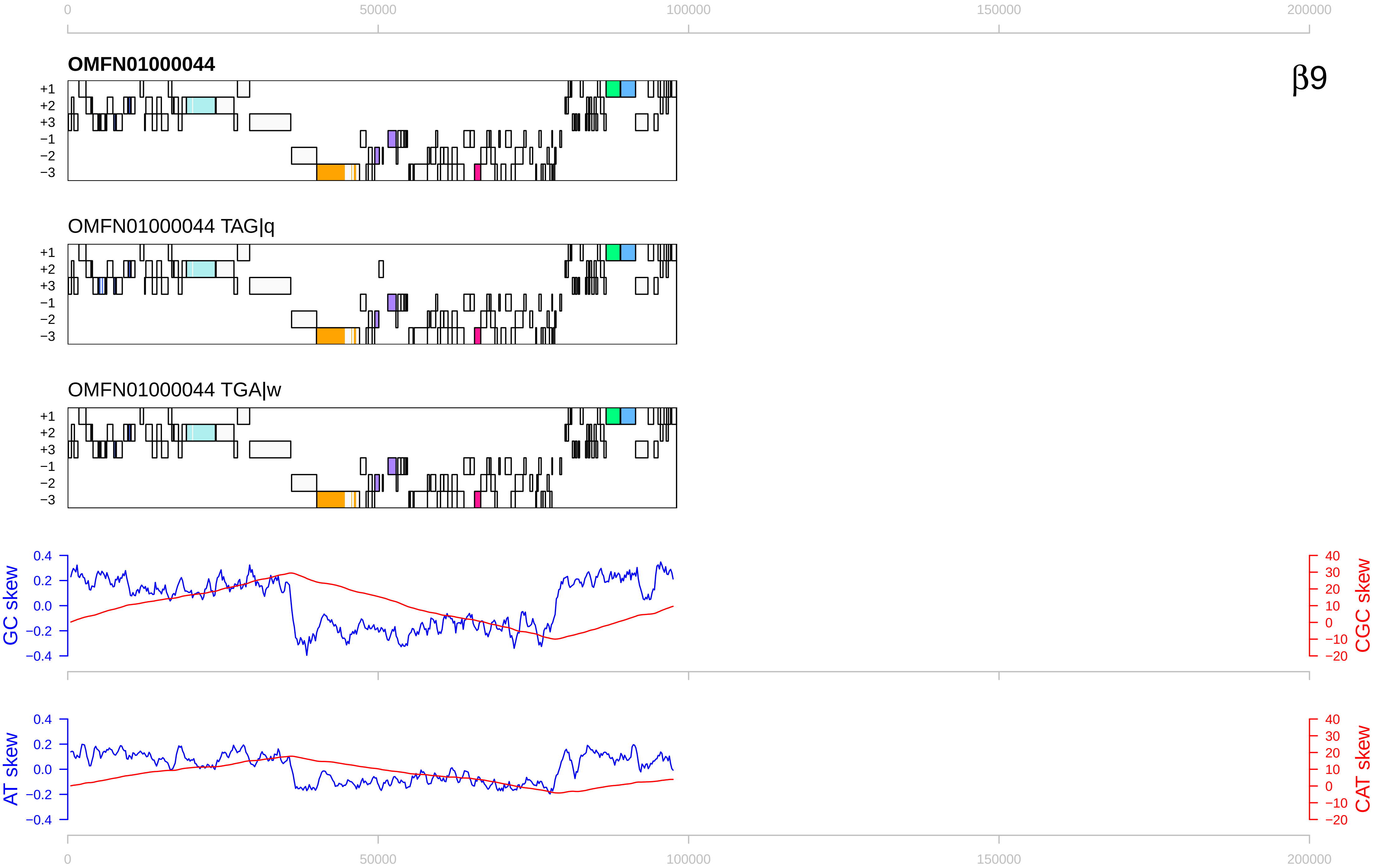

0
50000
100000
150000
200000
OMFN01000044
β9
+1
+2
+3
−1
−2
−3
OMFN01000044 TAG|q
OMFN01000044 TGA|w
GC skew
CGC skew
AT skew
CAT skew
0.4
0.2
0.0
−0.2
−0.4
40
30
20
10
0
−10
−20

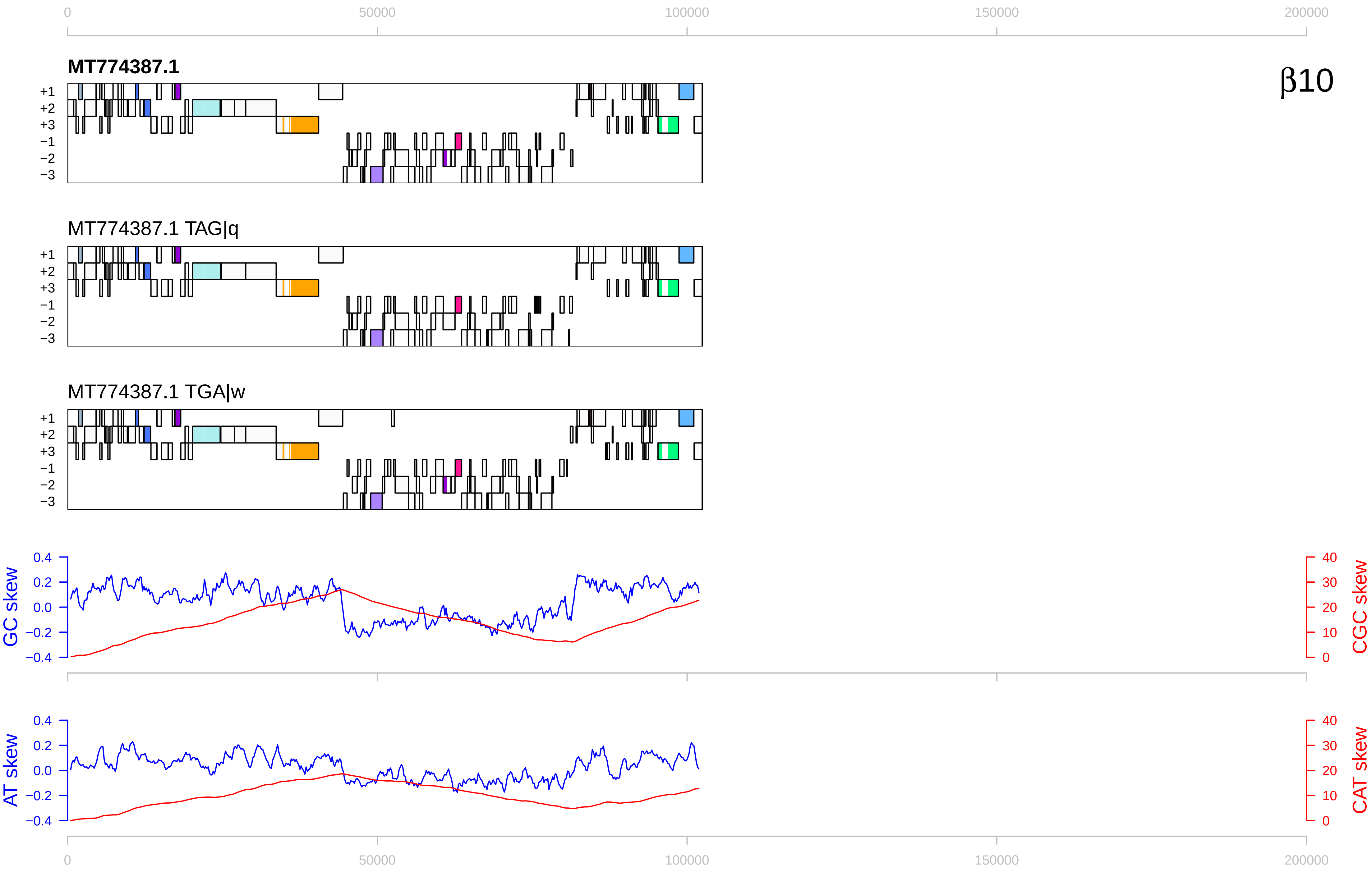

β10
MT774387.1
MT774387.1 TAG|q
MT774387.1 TGA|w
+1
+2
+3
−1
−2
−3
GC skew
CGC skew
AT skew
CAT skew
0
50000
100000
150000
200000
0.4
0.2
0.0
−0.2
−0.4
40
30
20
10

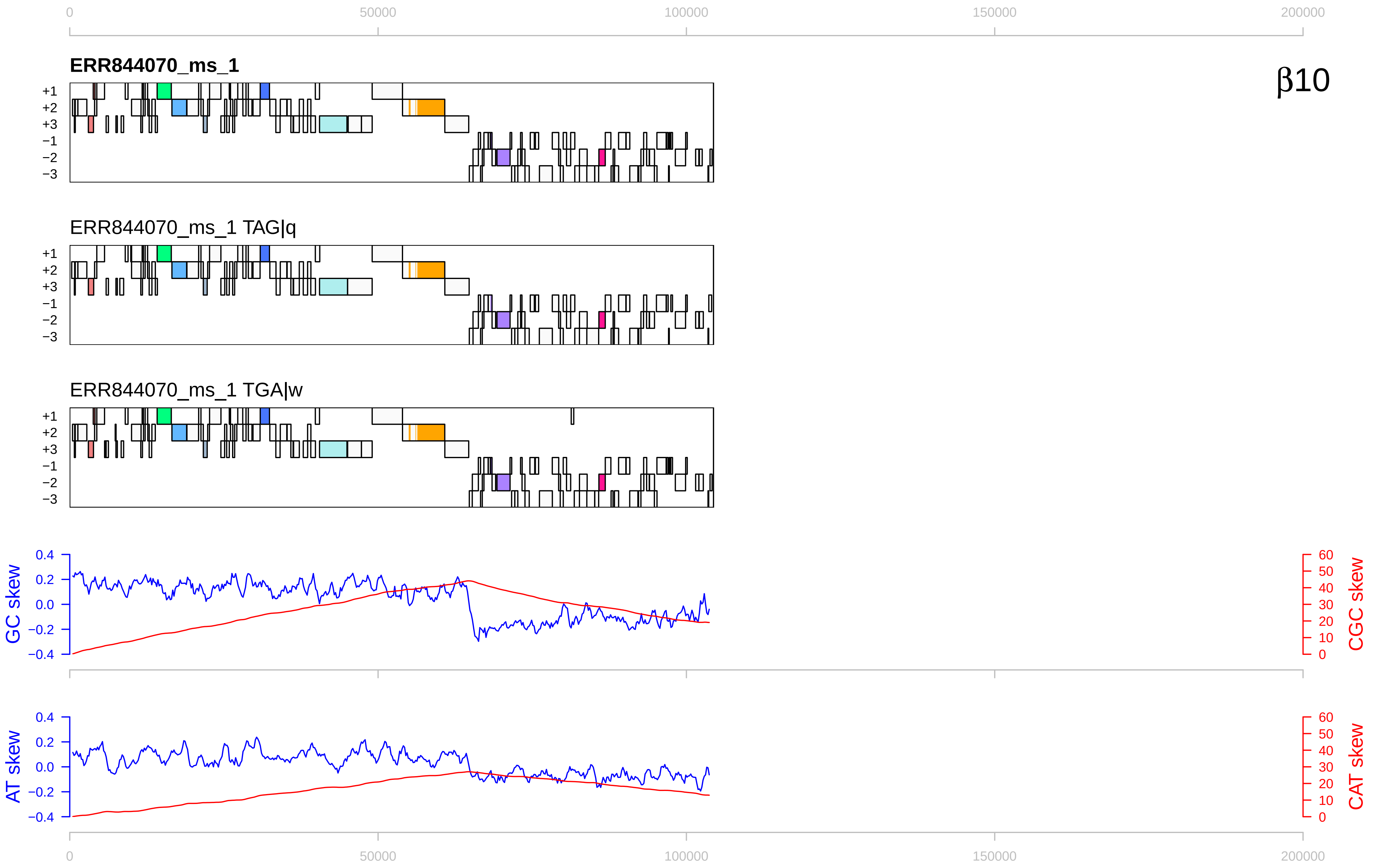

0
50000
100000
150000
200000
ERR844070_ms_1
β10
+1
+2
+3
−1
−2
−3
ERR844070_ms_1 TAG|q
ERR844070_ms_1 TGA|w
GC skew
CGC skew
AT skew
CAT skew
0.4
0.2
0.0
−0.2
−0.4
60
50
40
30
20
10

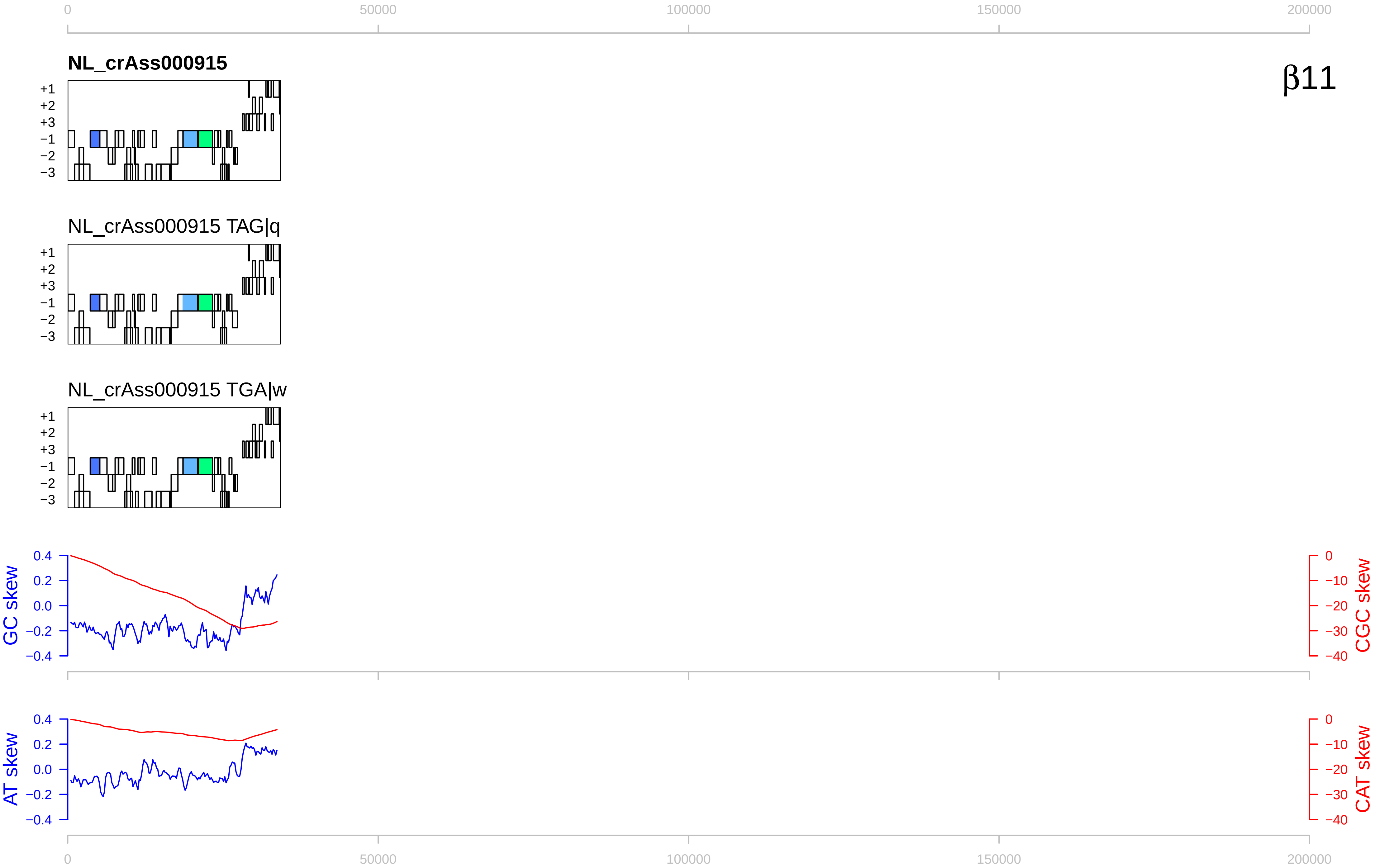

β11
NL_crAss000915
NL_crAss000915 TAG|q
NL_crAss000915 TGA|w
+1
+2
+3
−1
−2
−3
GC skew
CGC skew
AT skew
CAT skew
0
50000
100000
150000
200000

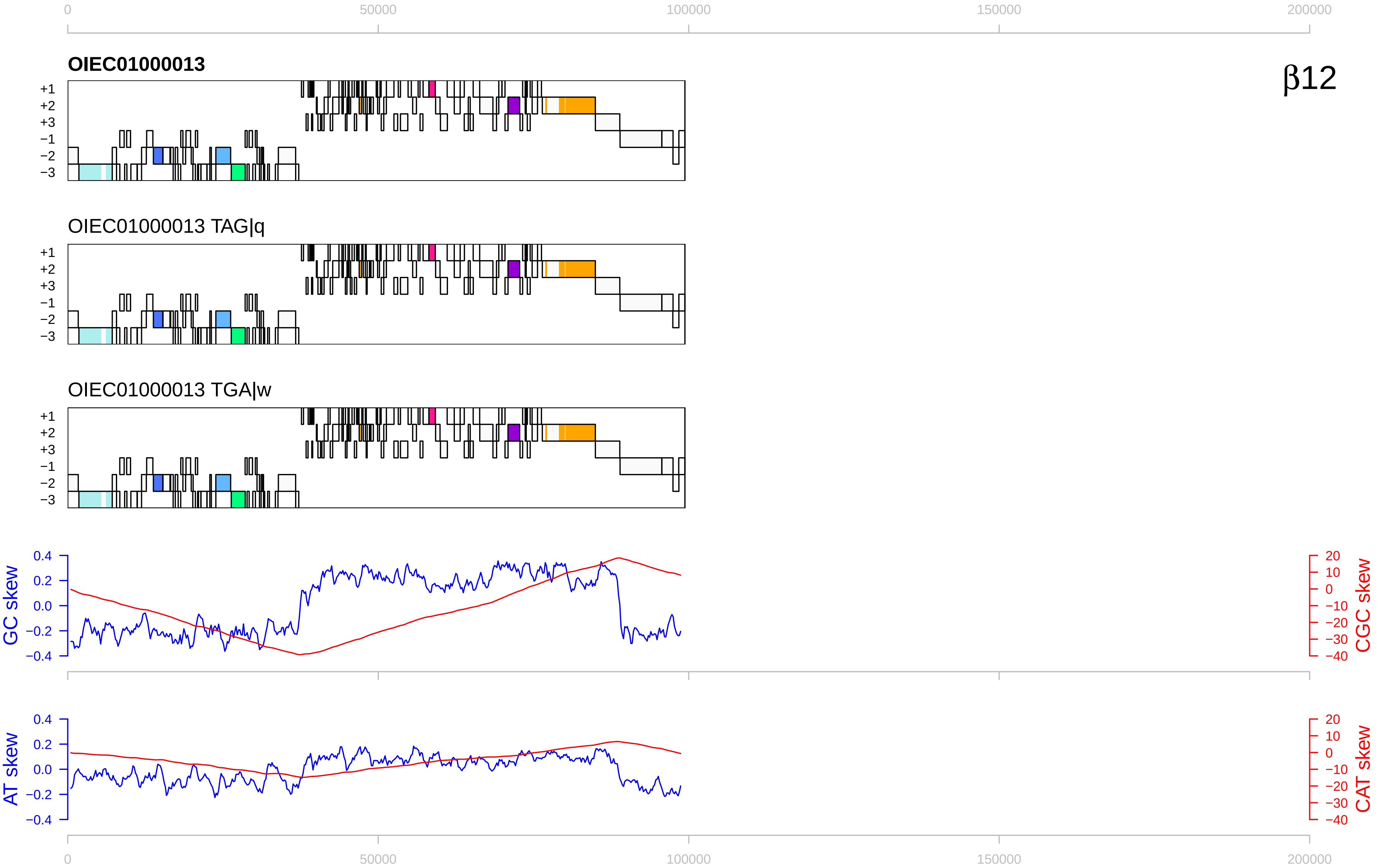

β12
OIEC01000013
OIEC01000013 TAG|q
OIEC01000013 TGA|w
+1
+2
+3
−1
−2
−3
GC skew
CGC skew
AT skew
CAT skew
0
50000
100000
150000
200000

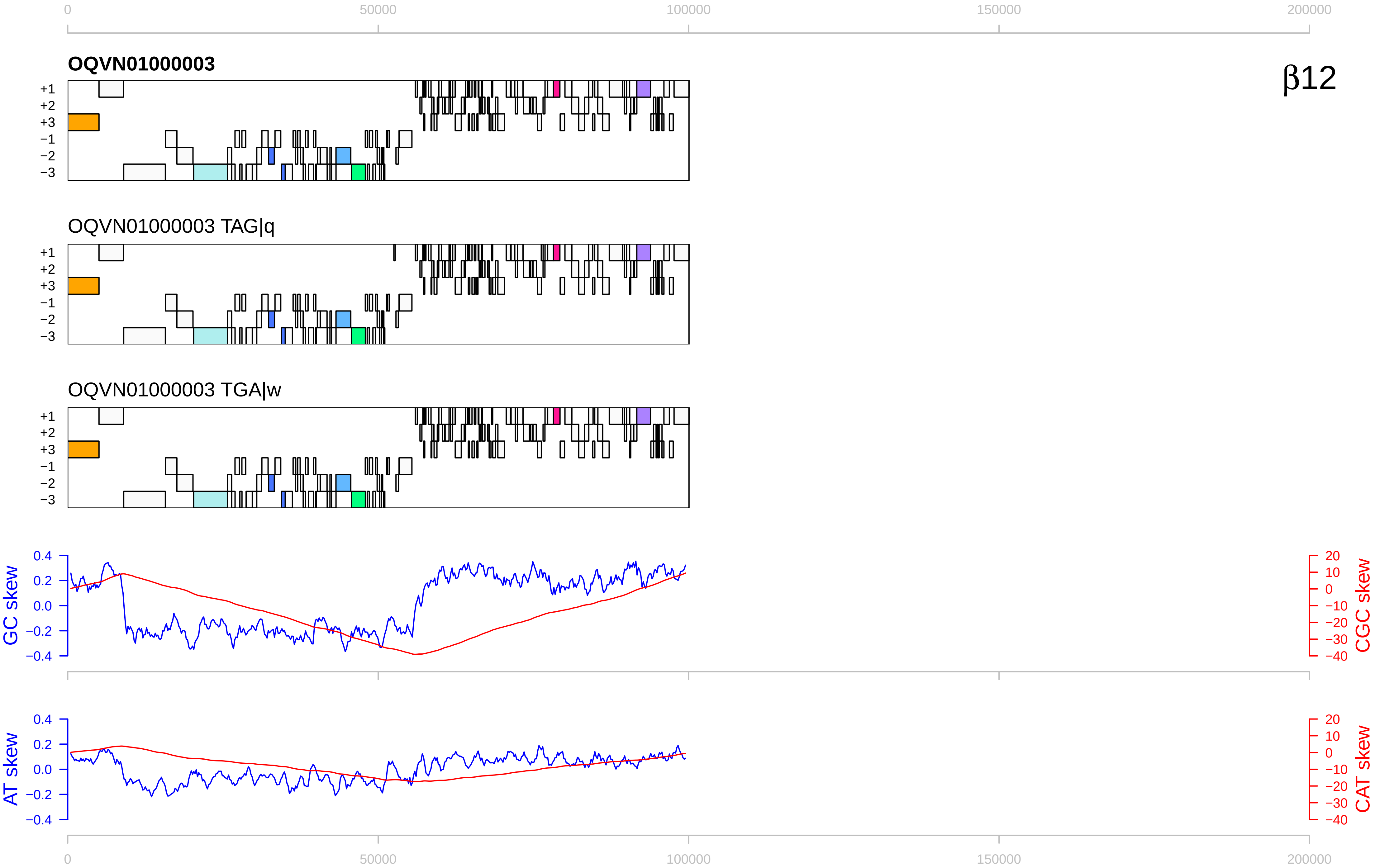

0
50000
100000
150000
200000
OQVN01000003
β12
+1
+2
+3
−1
−2
−3
OQVN01000003 TAG|q
OQVN01000003 TGA|w
GC skew
CGC skew
AT skew
CAT skew
0.4
0.2
0.0
−0.2
−0.4
20
10
0
−10
−20
−30
−40

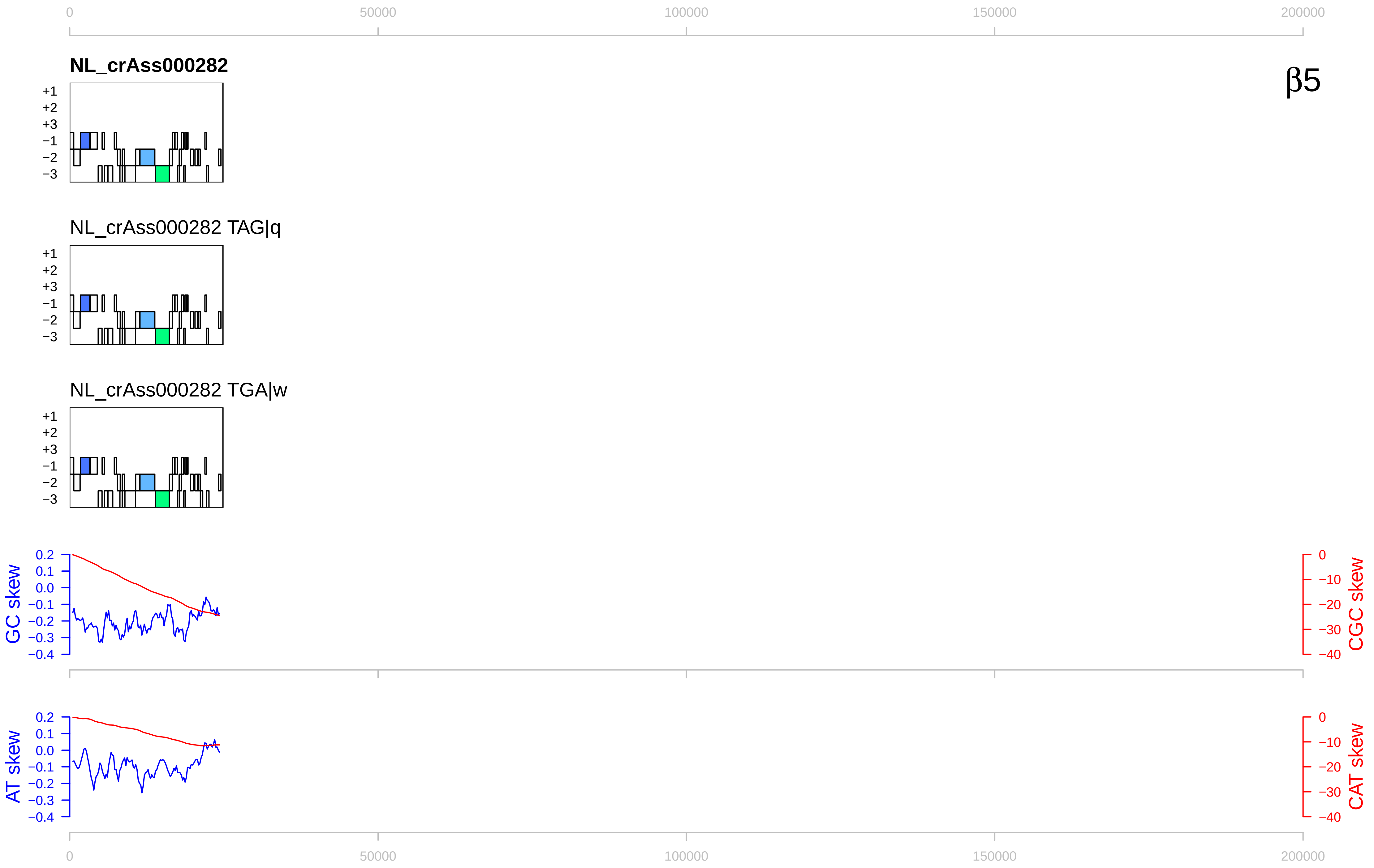

0
50000
100000
150000
200000
NL_crAss000282
β5
+1
+2
+3
−1
−2
−3
NL_crAss000282 TAG|q
NL_crAss000282 TGA|w
GC skew
CGC skew
AT skew
CAT skew
0.2
0.1
0.0
−0.1
−0.2
−0.3
−0.4
0
−10
−20
−30
−40

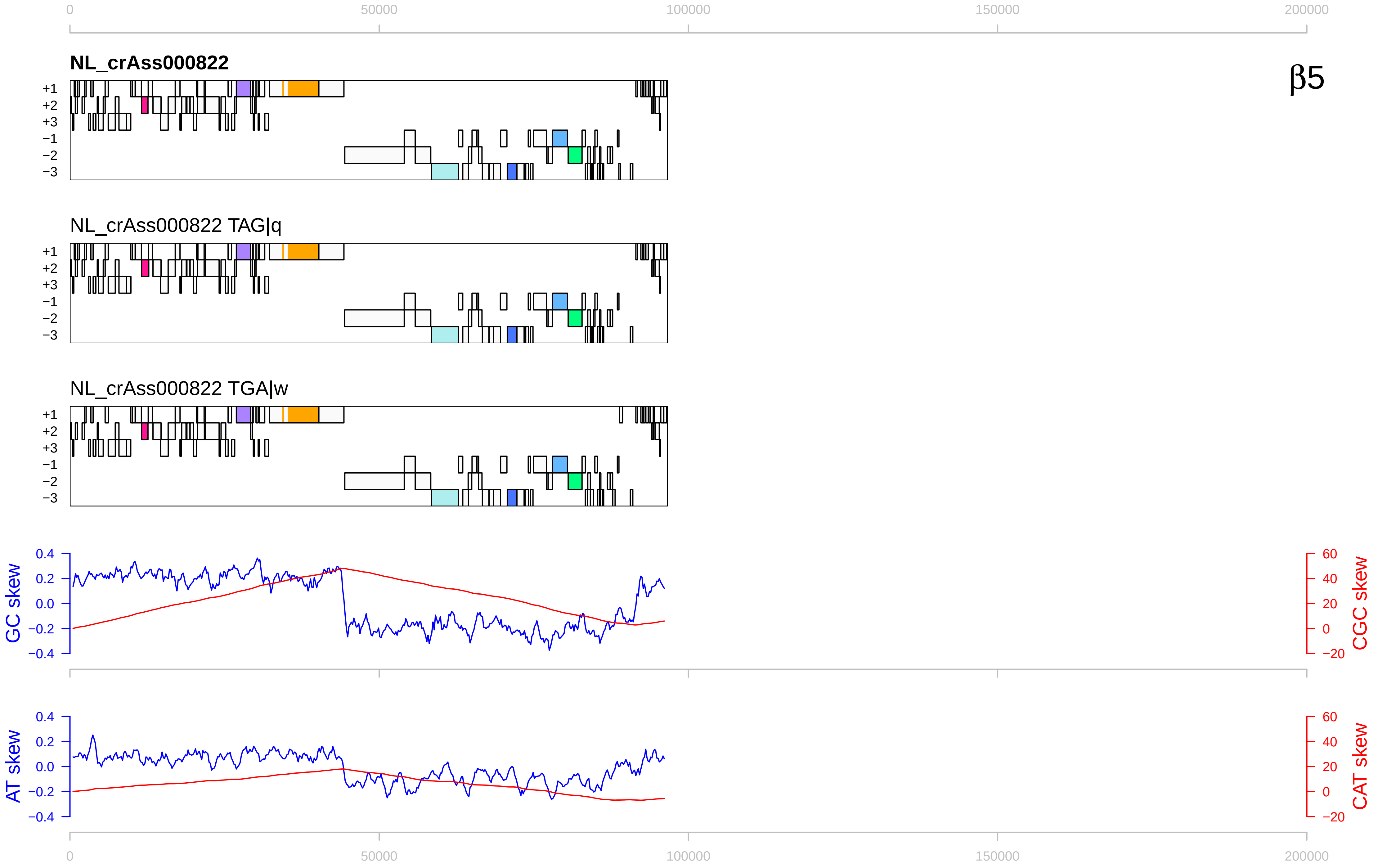

β5
NL_crAss000822
NL_crAss000822 TAG|q
NL_crAss000822 TGA|w
+1
+2
+3
−1
−2
−3
GC skew
CGC skew
AT skew
CAT skew
0
50000
100000
150000
200000
0.4
0.2
0.0
−0.2
−0.4
60
40
20
0
−20

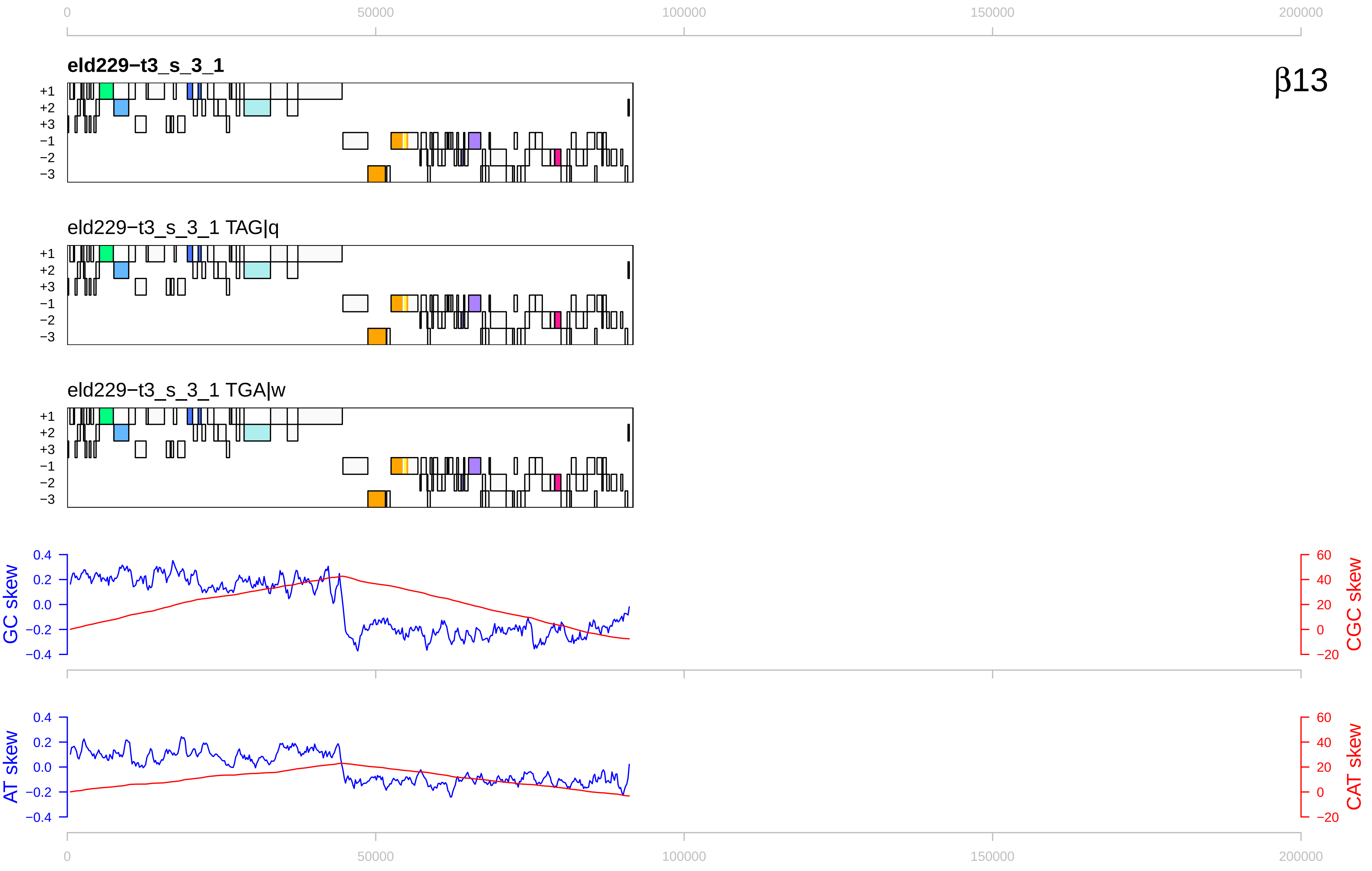

0
50000
100000
150000
200000
β13
eld229−t3_s_3_1
eld229−t3_s_3_1 TAG|q
eld229−t3_s_3_1 TGA|w
+1
+2
+3
−1
−2
−3
GC skew
CGC skew
AT skew
CAT skew
0.4
0.2
0.0
−0.2
−0.4
60
40
20
0
−20

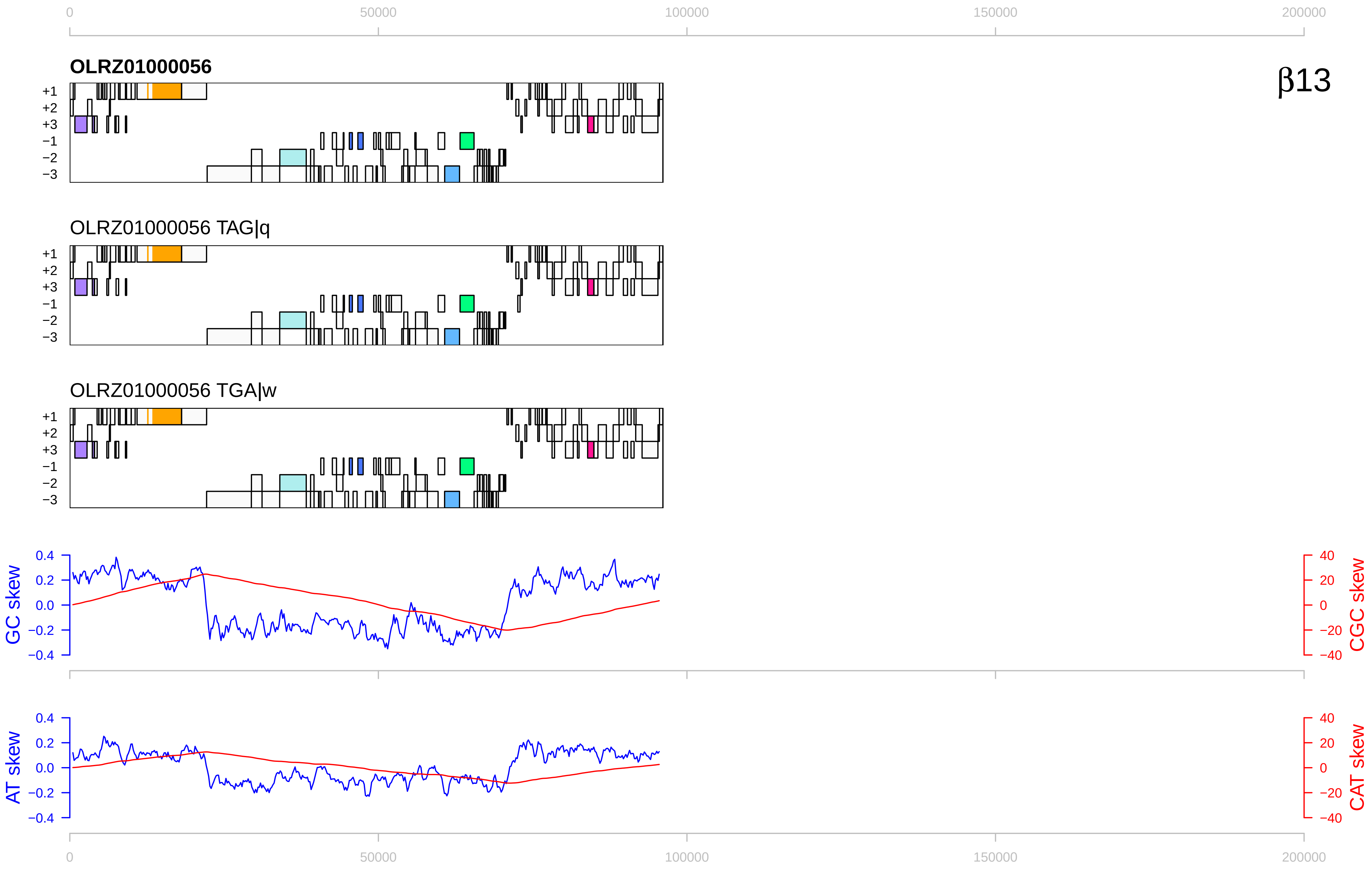

0
50000
100000
150000
200000
OLRZ01000056
β13
+1
+2
+3
−1
−2
−3
OLRZ01000056 TAG|q
OLRZ01000056 TGA|w
GC skew
CGC skew
AT skew
CAT skew
0.4
0.2
0.0
−0.2
−0.4
40
20
0
−20
−40

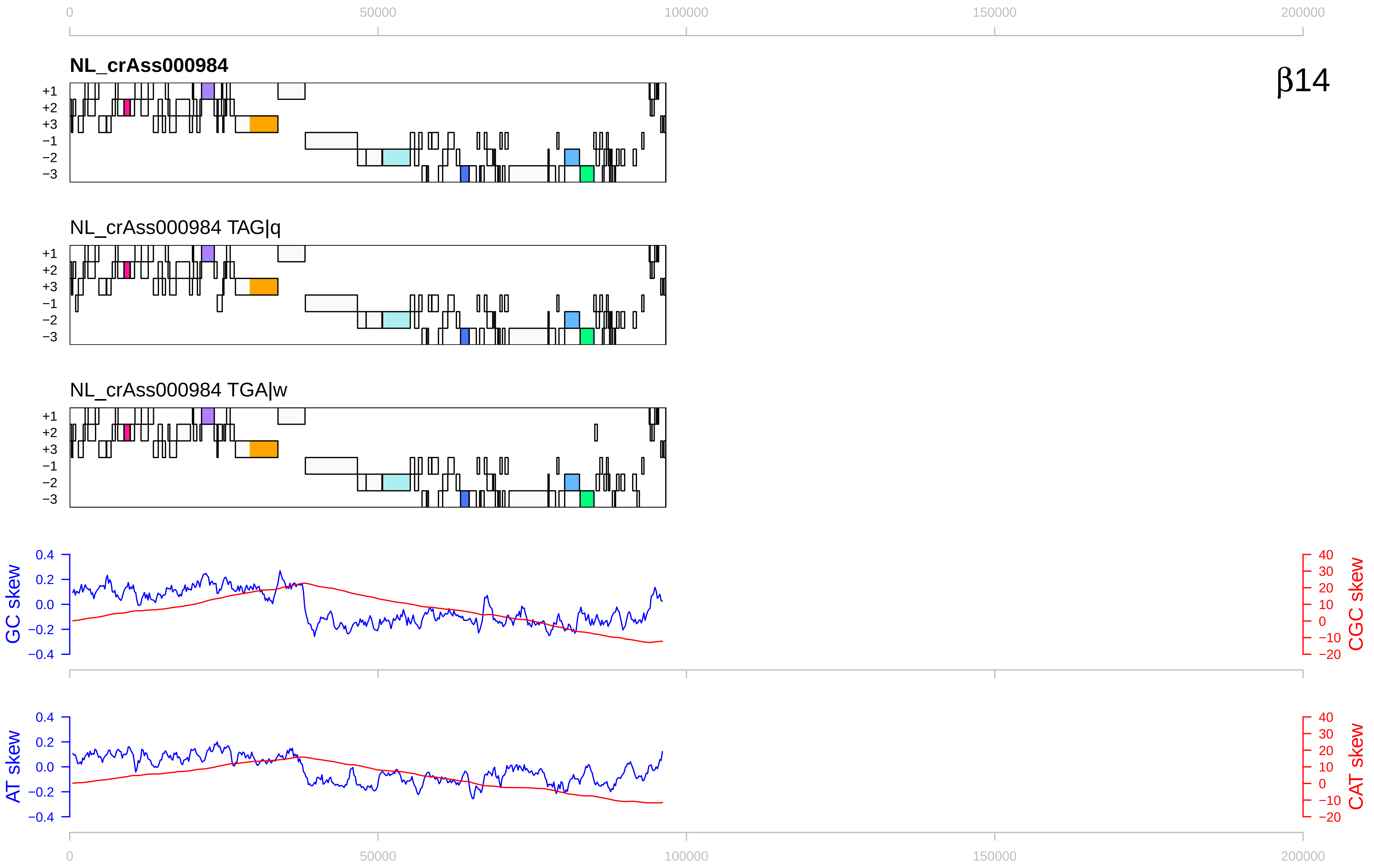

0
50000
100000
150000
200000
NL_crAss000984
β14
+1
+2
+3
−1
−2
−3
NL_crAss000984 TAG|q
NL_crAss000984 TGA|w
GC skew
CGC skew
AT skew
CAT skew
0.4
0.2
0.0
−0.2
−0.4
40
30
20
10
0
−10
−20

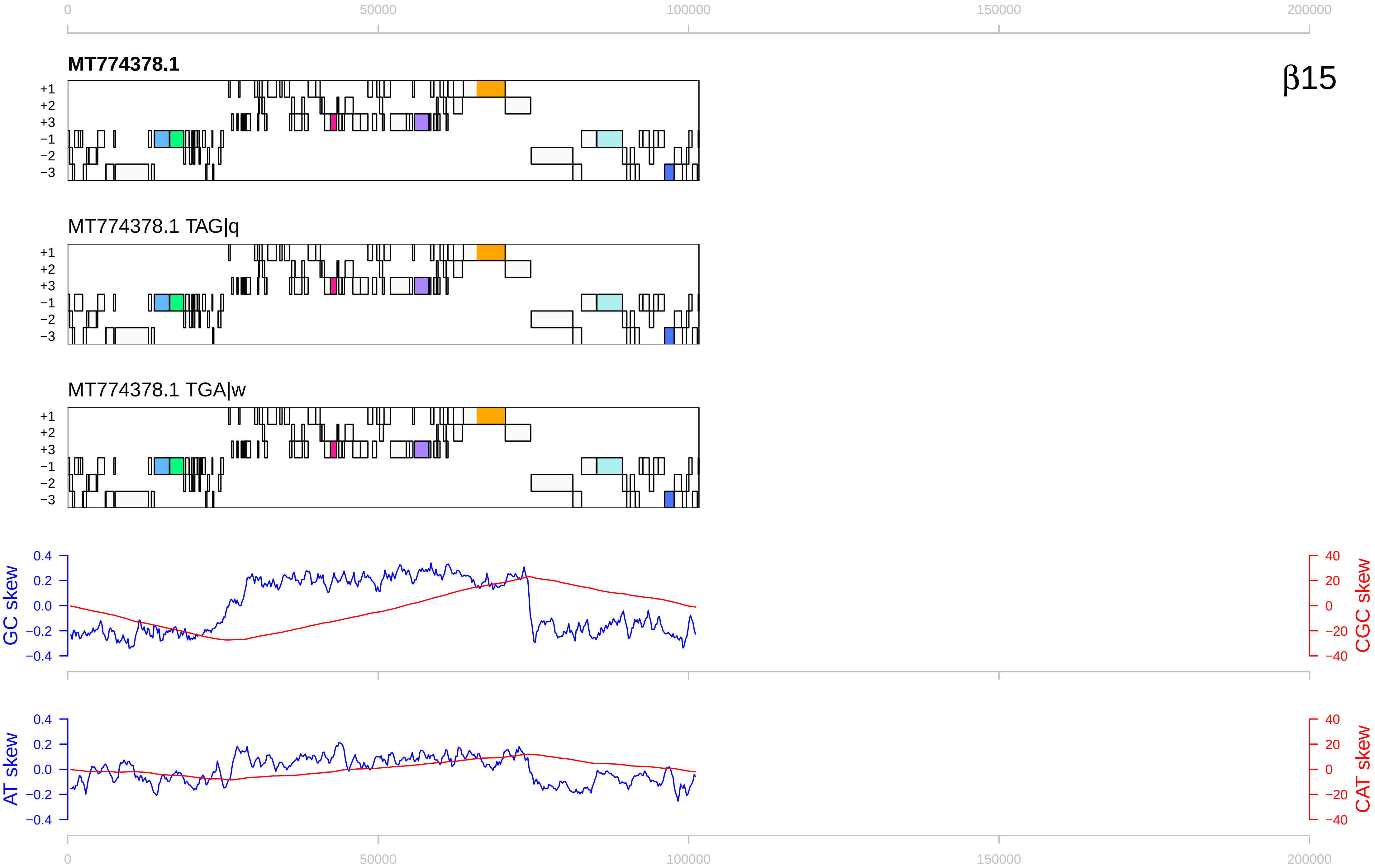

MT774378.1
β15
MT774378.1 TAG|q
MT774378.1 TGA|w
+1
+2
+3
−1
−2
−3
GC skew
CGC skew
AT skew
CAT skew
0
50000
100000
150000
200000

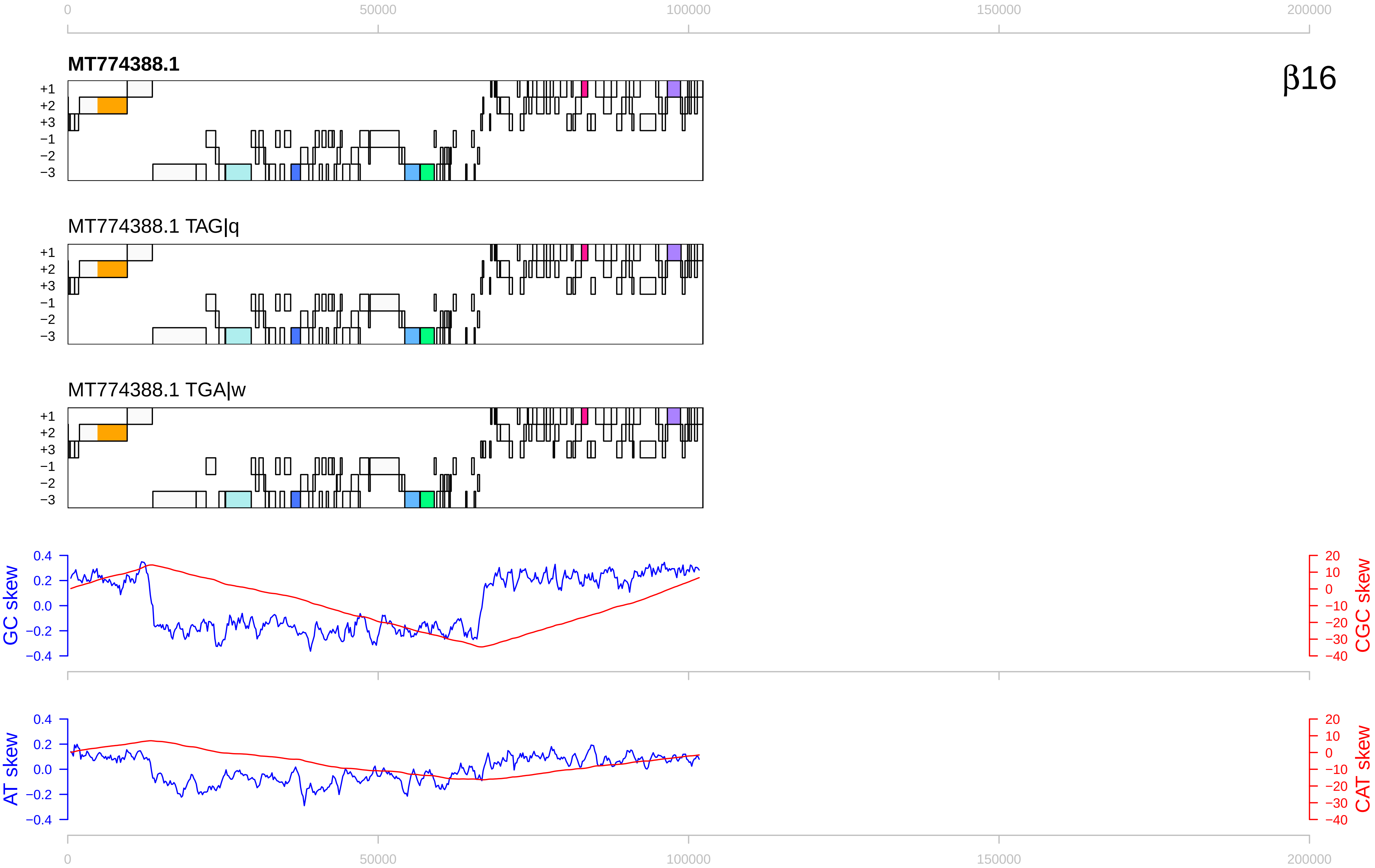

0
50000
100000
150000
200000
MT774388.1
β16
+1
+2
+3
−1
−2
−3
MT774388.1 TAG|q
MT774388.1 TGA|w
GC skew
CGC skew
AT skew
CAT skew
0.4
0.2
0.0
−0.2
−0.4
20
10
0
−10
−20
−30
−40

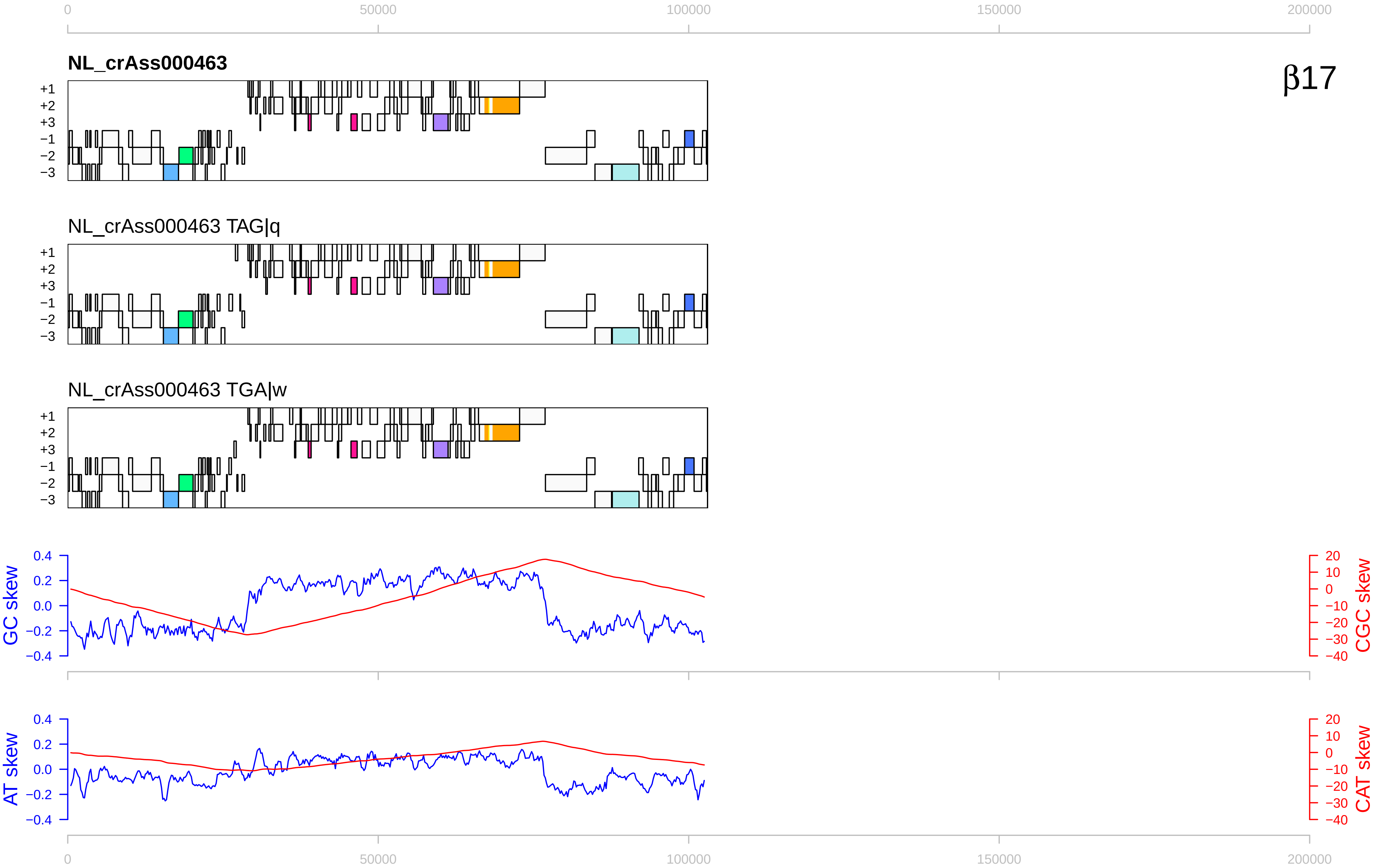

0
50000
100000
150000
200000
NL_crAss000463
β17
+1
+2
+3
−1
−2
−3
NL_crAss000463 TAG|q
NL_crAss000463 TGA|w
GC skew
CGC skew
AT skew
CAT skew
0.4
0.2
0.0
−0.2
−0.4
20
10
0
−10
−20
−30
−40

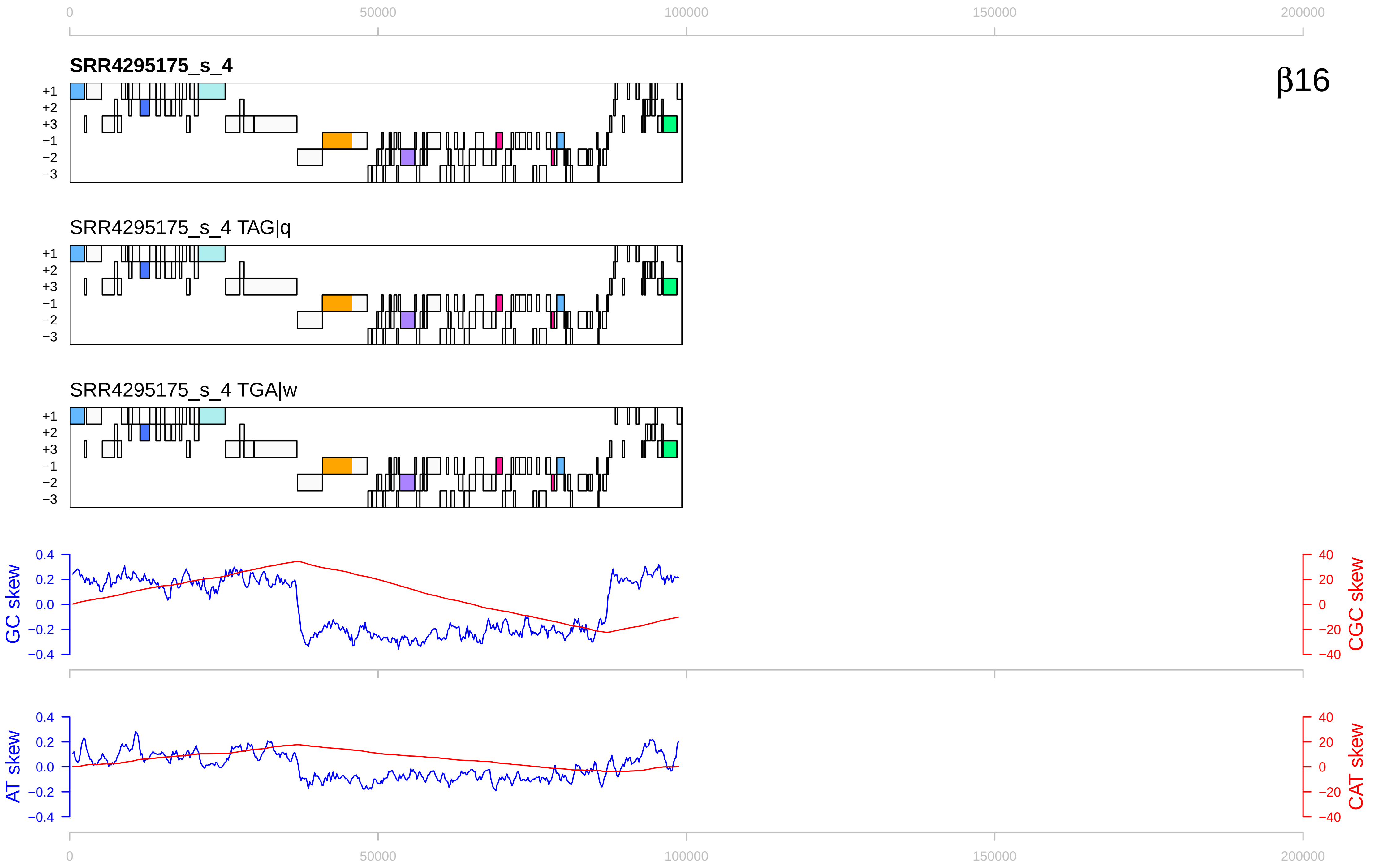

0
50000
100000
150000
200000
SRR4295175_s_4
β16
+1
+2
+3
−1
−2
−3
SRR4295175_s_4 TAG|q
SRR4295175_s_4 TGA|w
GC skew
CGC skew
0.4
0.2
0.0
−0.2
−0.4
40
20
0
−20
−40
AT skew
CAT skew

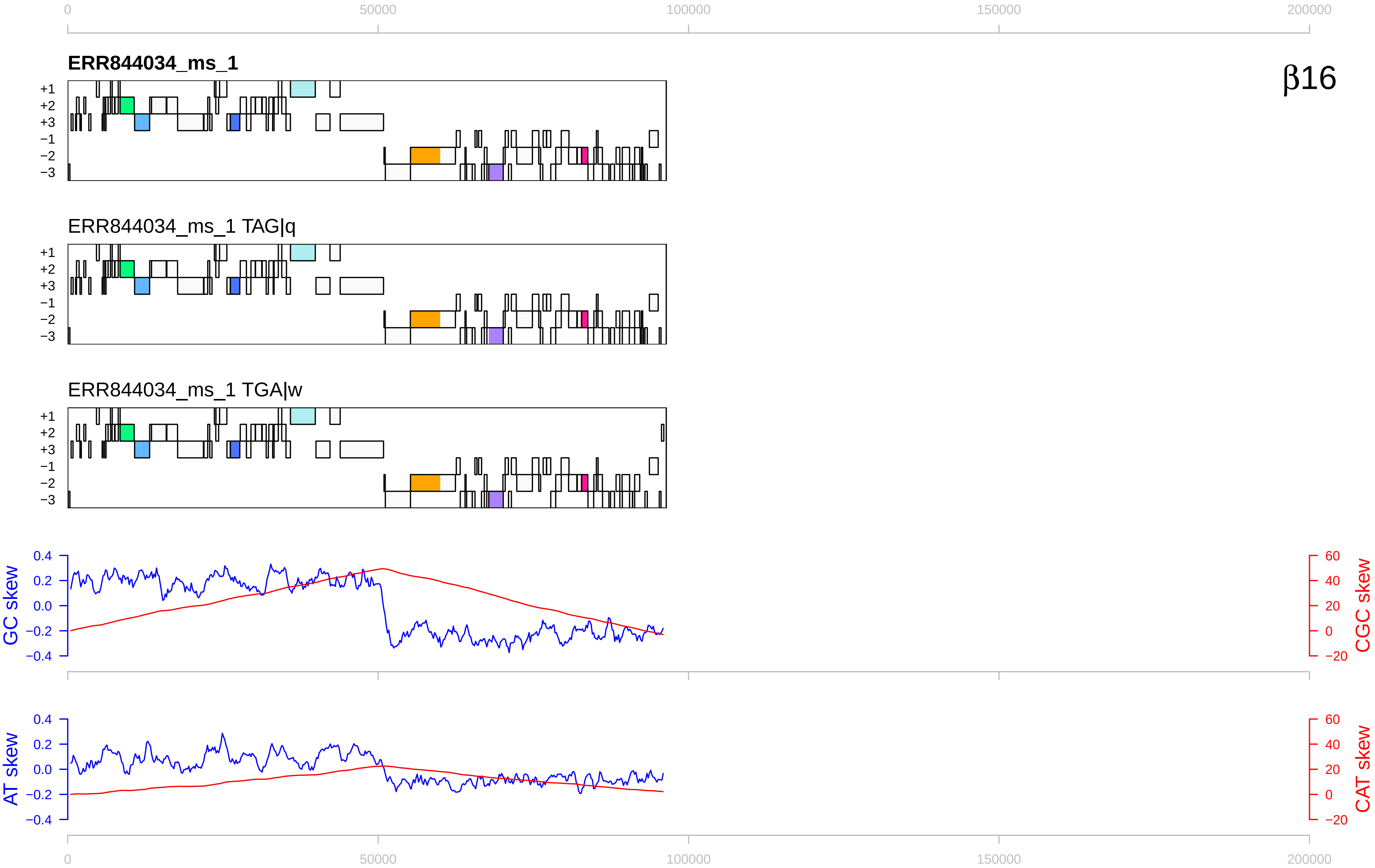

β16
ERR844034_ms_1
ERR844034_ms_1 TAG|q
ERR844034_ms_1 TGA|w
+1
+2
+3
−1
−2
−3
GC skew
CGC skew
AT skew
CAT skew
0
50000
100000
150000
200000

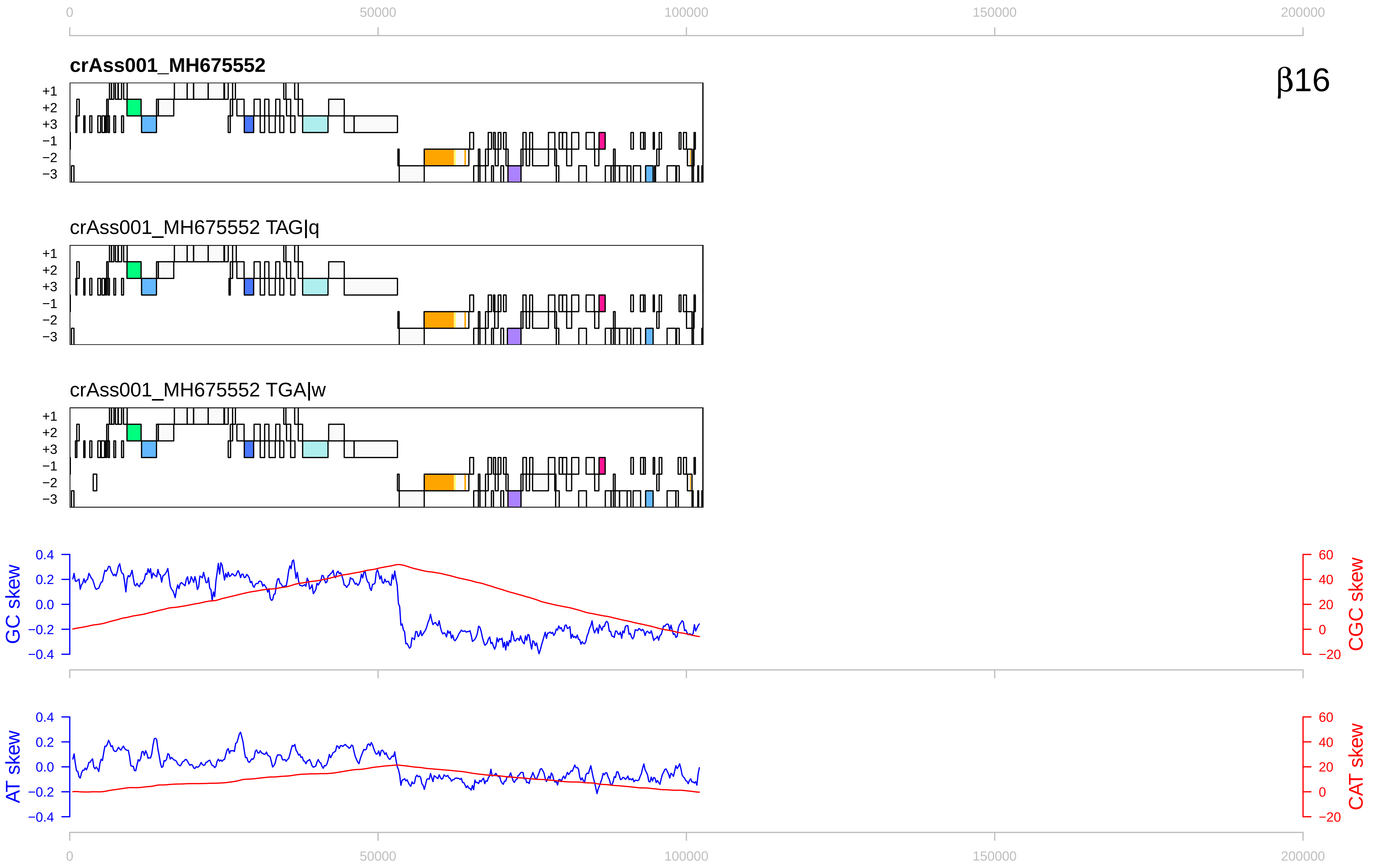

β16
crAss001_MH675552
crAss001_MH675552 TAG|q
crAss001_MH675552 TGA|w
+1
+2
+3
−1
−2
−3
GC skew
CGC skew
AT skew
CAT skew
0
50000
100000
150000
200000
0.4
0.2
0.0
−0.2
−0.4
60
40
20
0
−20

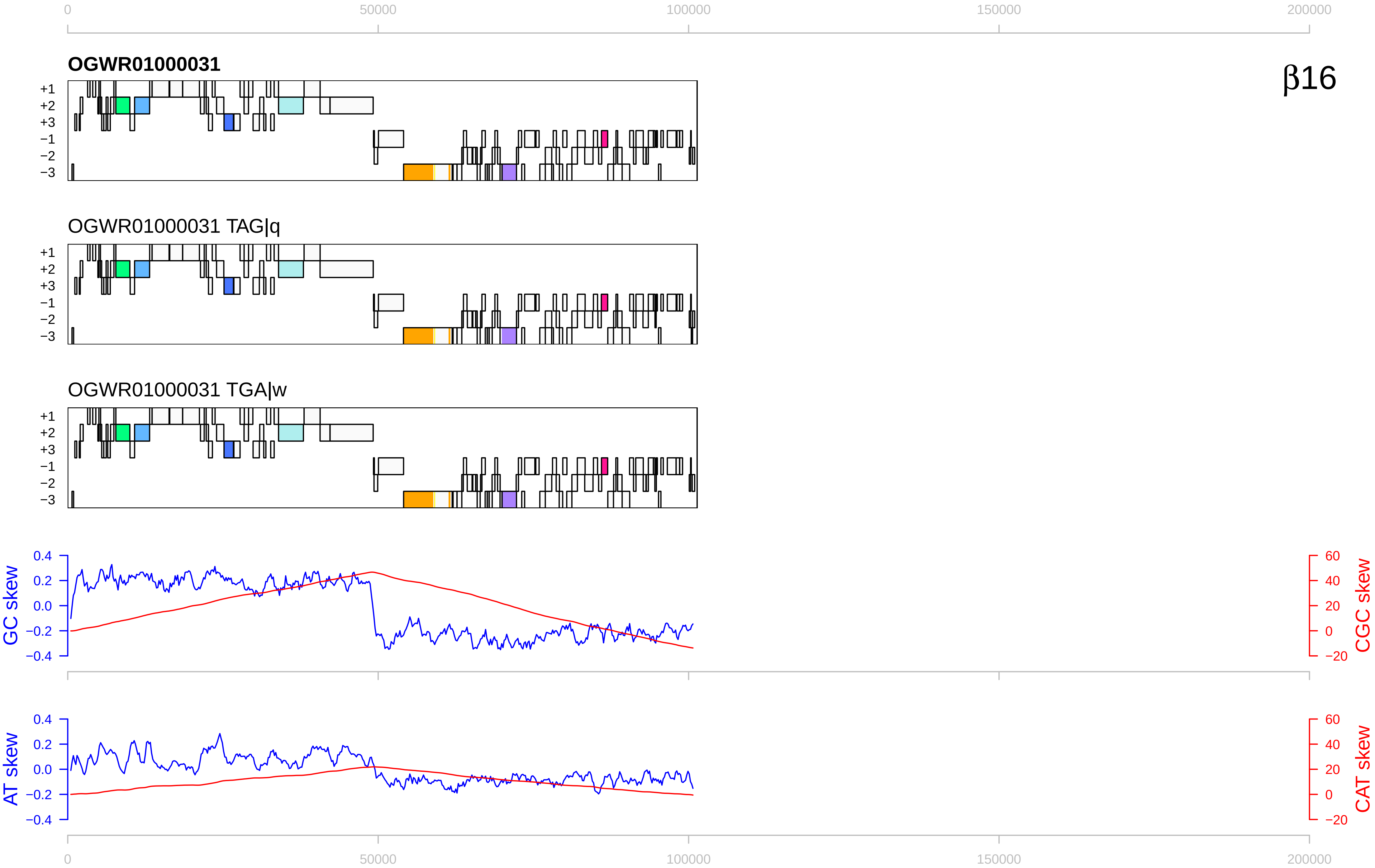
0
50000
100000
150000
200000
OGWR01000031
β16
+1
+2
+3
−1
−2
−3
OGWR01000031 TAG|q
OGWR01000031 TGA|w
GC skew
CGC skew
AT skew
CAT skew
0.4
0.2
0.0
−0.2
−0.4
60
40
20
0
−20

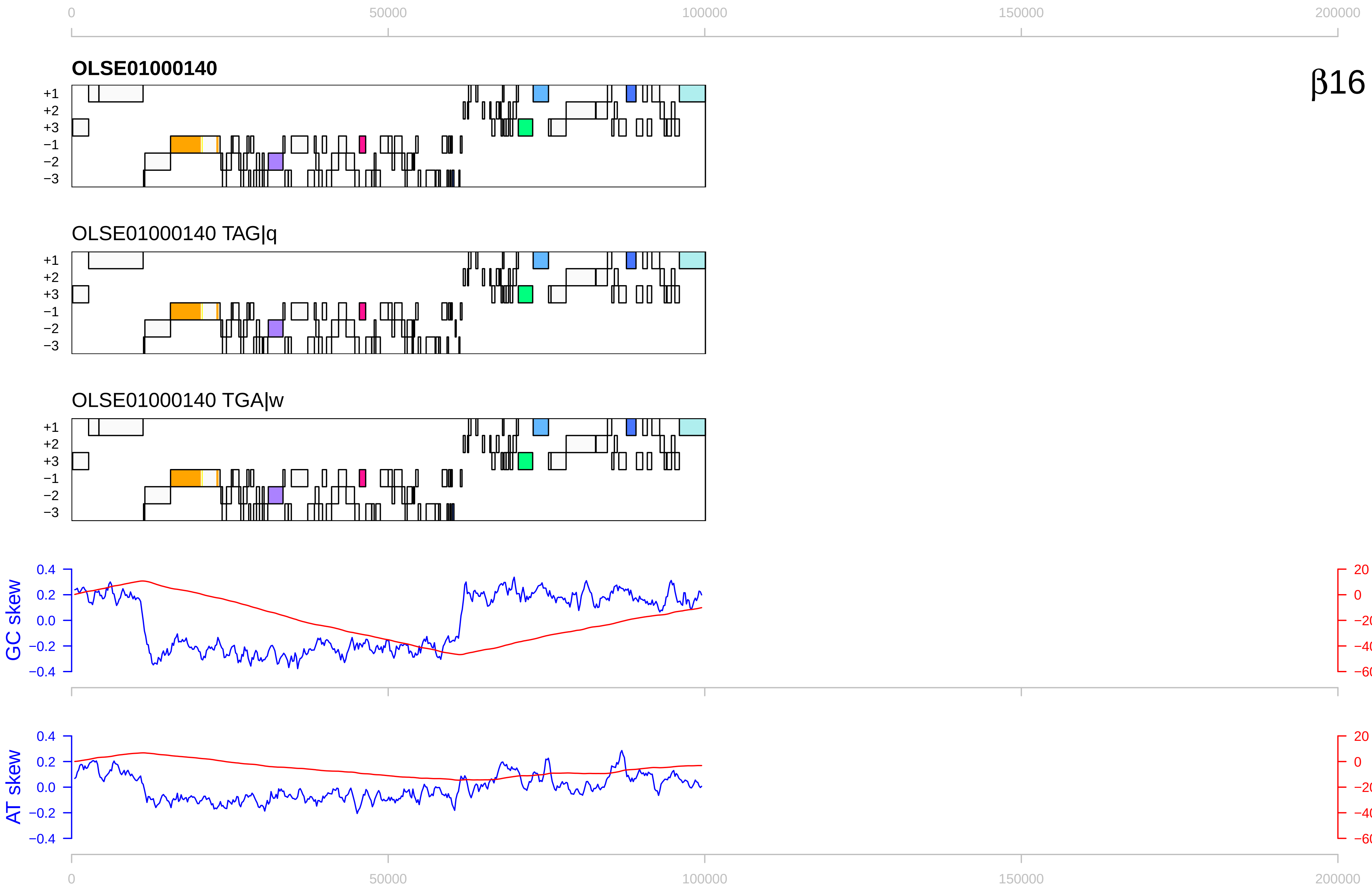

0
50000
100000
150000
200000
OLSE01000140
β16
+1
+2
+3
−1
−2
−3
OLSE01000140 TAG|q
OLSE01000140 TGA|w
GC skew
CGC skew
0.4
0.2
0.0
−0.2
−0.4
20
0
−20
−40
−60
AT skew
CAT skew

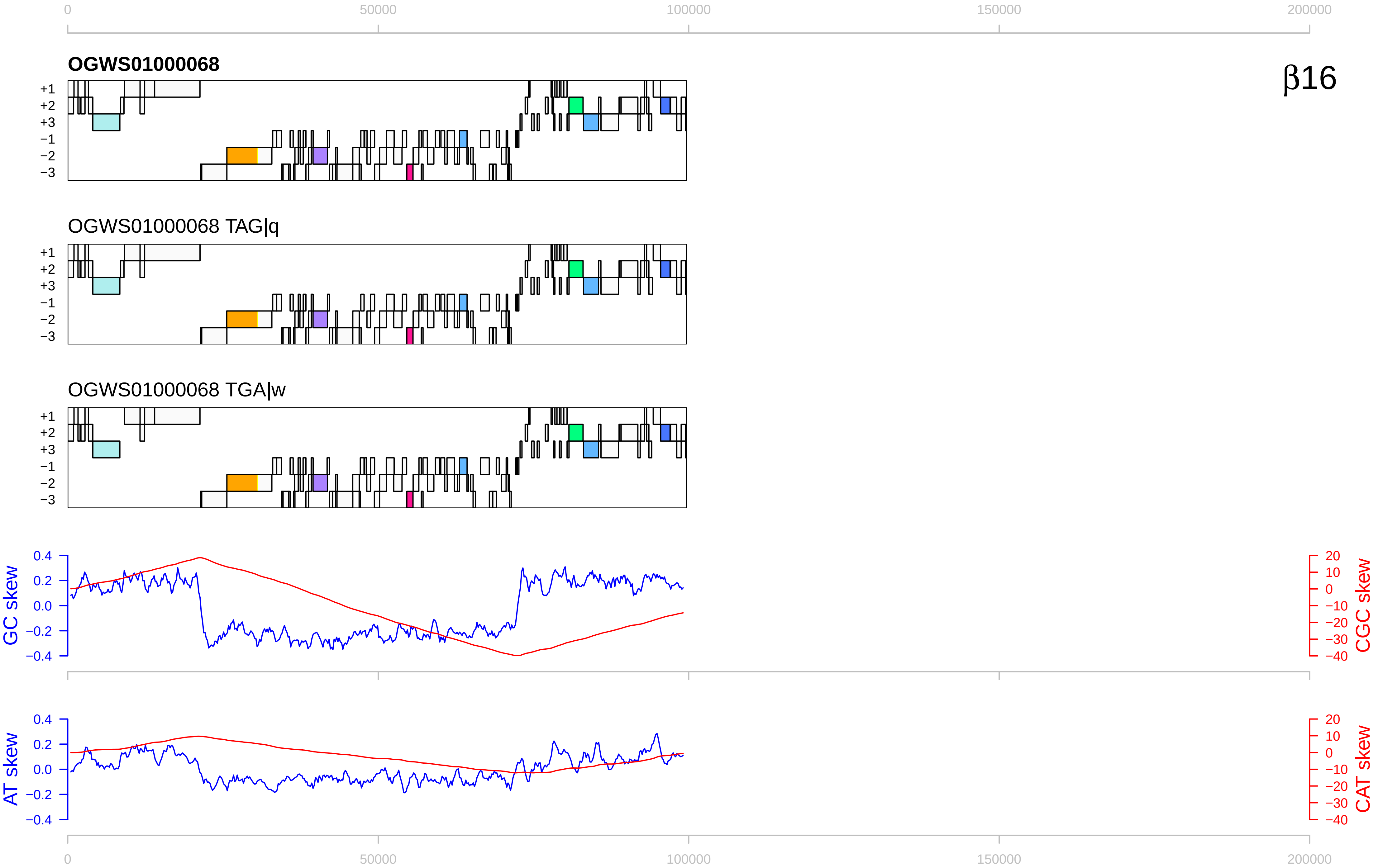

0
50000
100000
150000
200000
OGWS01000068
β16
+1
+2
+3
−1
−2
−3
OGWS01000068 TAG|q
OGWS01000068 TGA|w
GC skew
CGC skew
AT skew
CAT skew
0.4
0.2
0.0
−0.2
−0.4
20
10
0
−10
−20
−30
−40

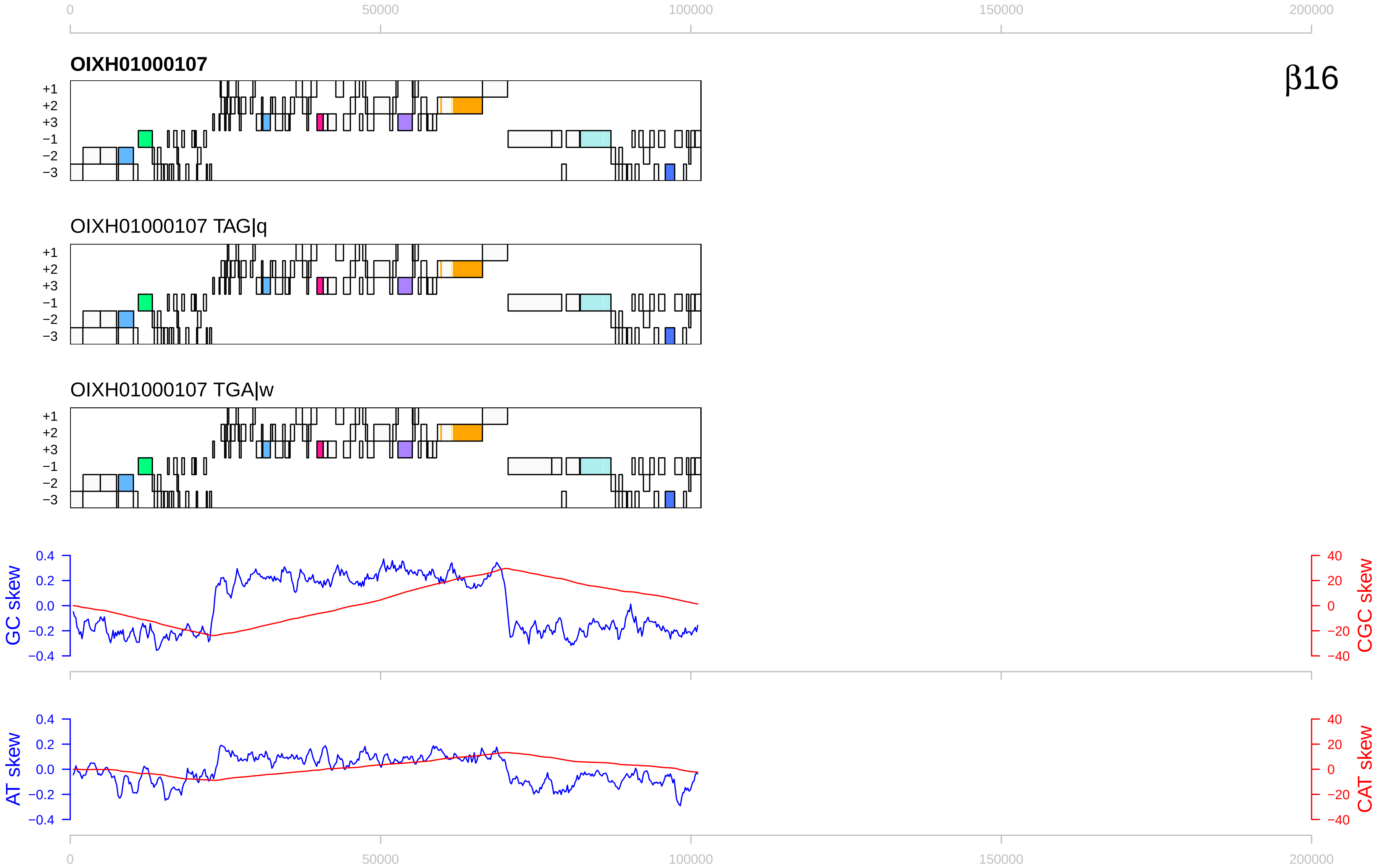

β16
OIXH01000107
OIXH01000107 TAG|q
OIXH01000107 TGA|w
+1
+2
+3
−1
−2
−3
GC skew
CGC skew
AT skew
CAT skew
0
50000
100000
150000
200000
0.4
0.2
0.0
−0.2
−0.4
40
20
0
−20
−40

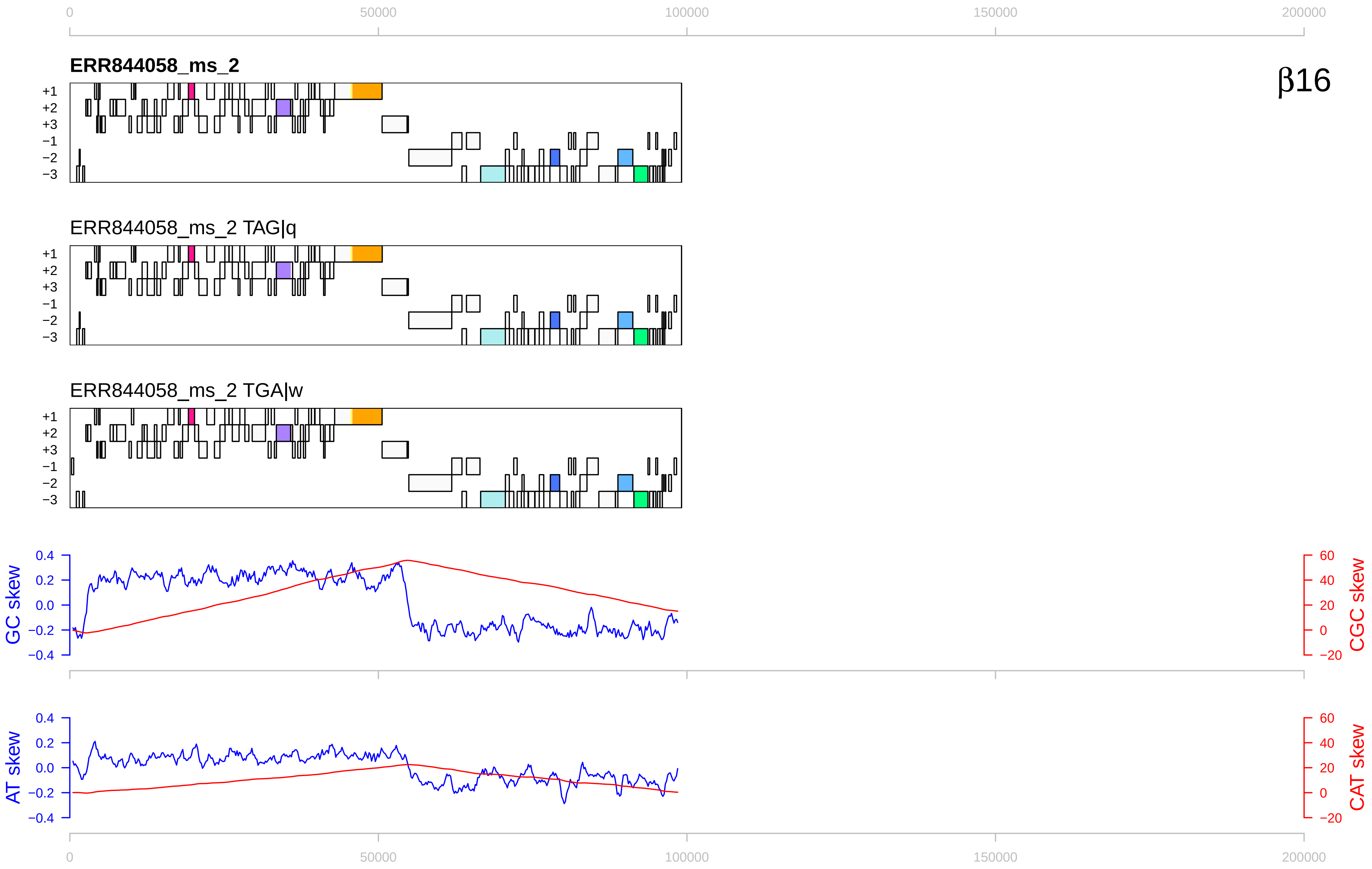

0
50000
100000
150000
200000
ERR844058_ms_2
β16
+1
+2
+3
−1
−2
−3
ERR844058_ms_2 TAG|q
ERR844058_ms_2 TGA|w
GC skew
0.4
0.2
0.0
−0.2
−0.4
CGC skew
60
40
20
0
−20
AT skew
CAT skew

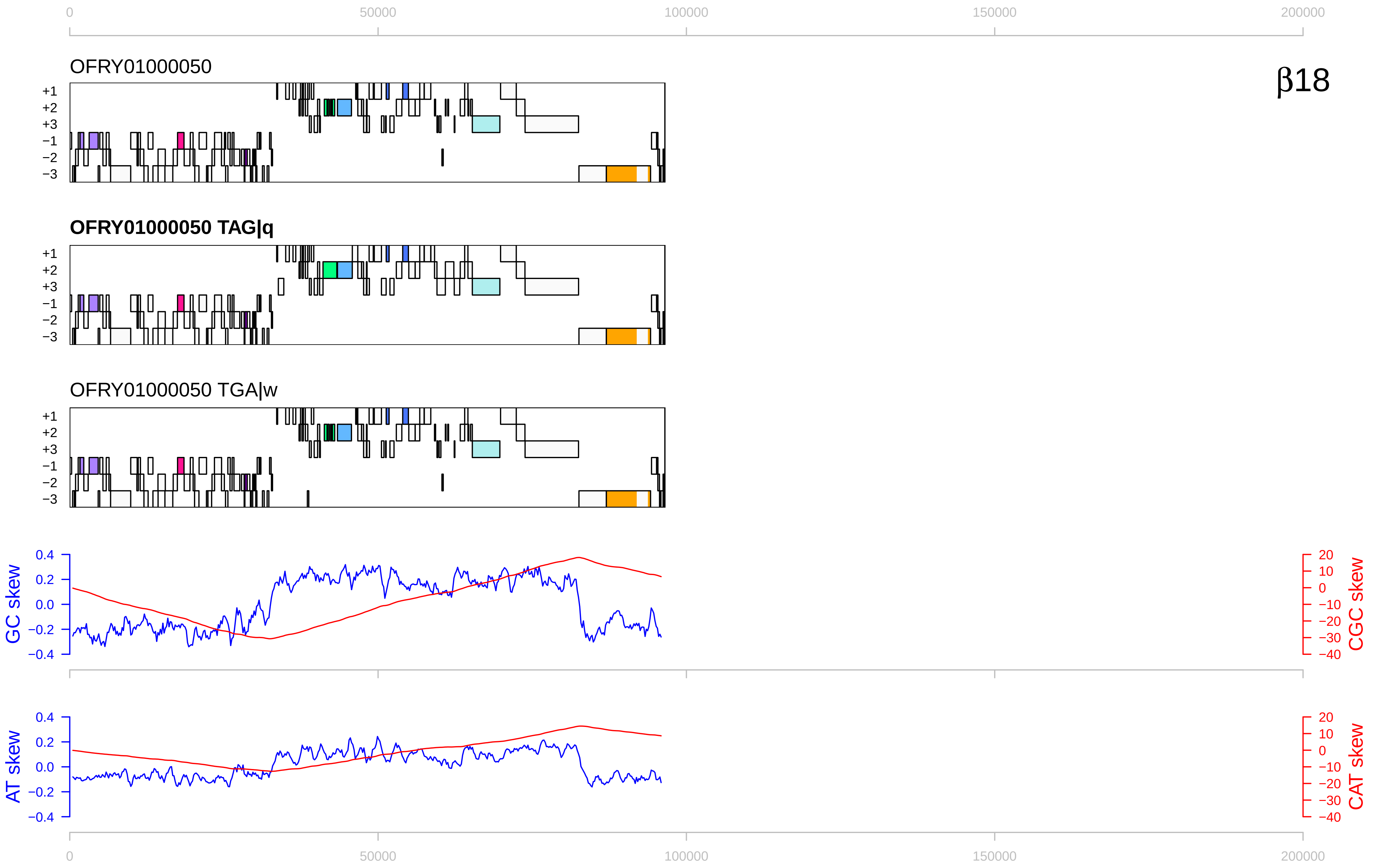

β18
OFRY01000050
OFRY01000050 TAG|q
OFRY01000050 TGA|w
+1
+2
+3
−1
−2
−3
GC skew
CGC skew
AT skew
CAT skew
0
50000
100000
150000
200000

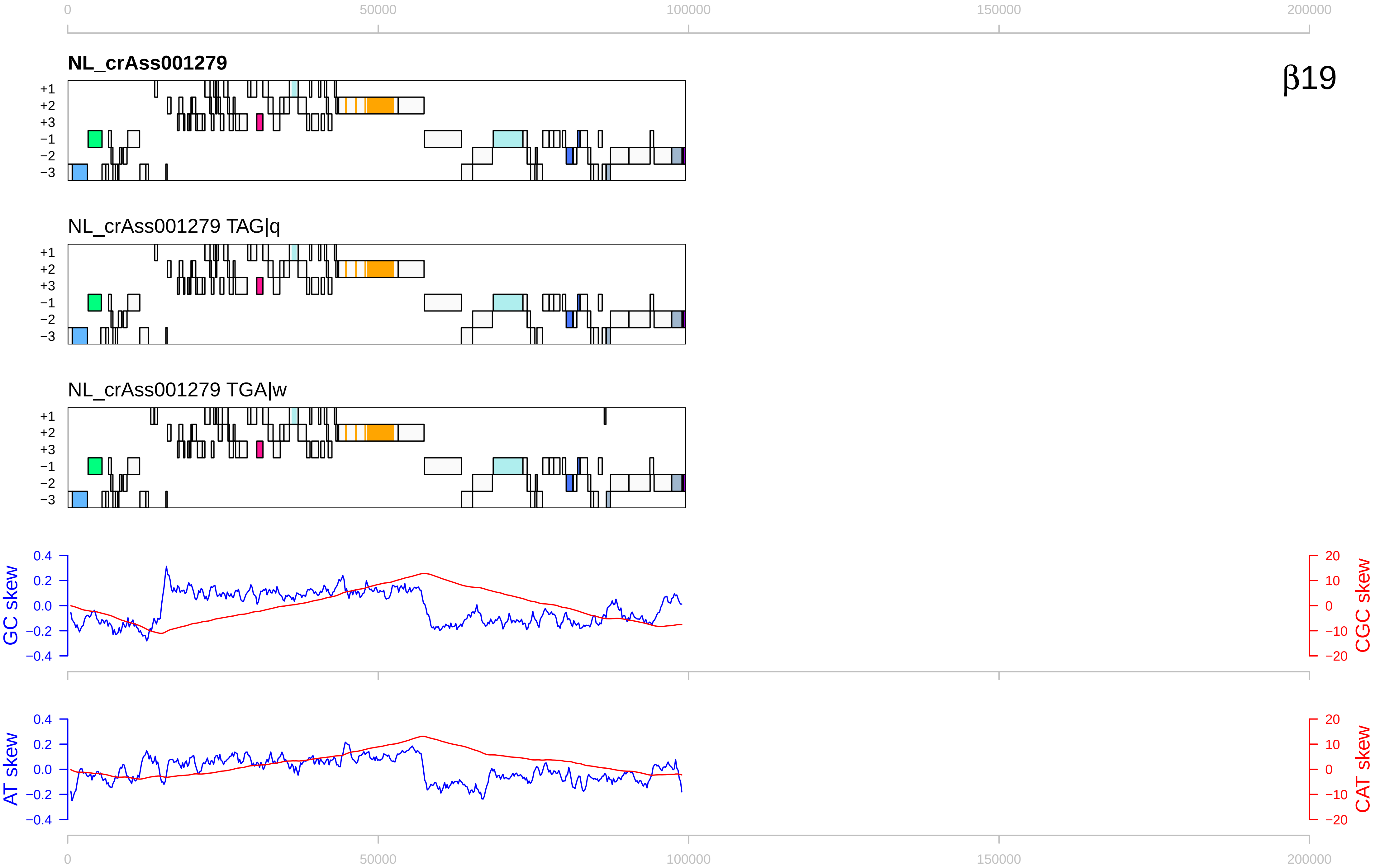
β19
NL_crAss001279
NL_crAss001279 TAG|q
NL_crAss001279 TGA|w
+1
+2
+3
−1
−2
−3
GC skew
CGC skew
AT skew
CAT skew
0
50000
100000
150000
200000
0.4
0.2
0.0
−0.2
−0.4
20
10
0
−10
−20

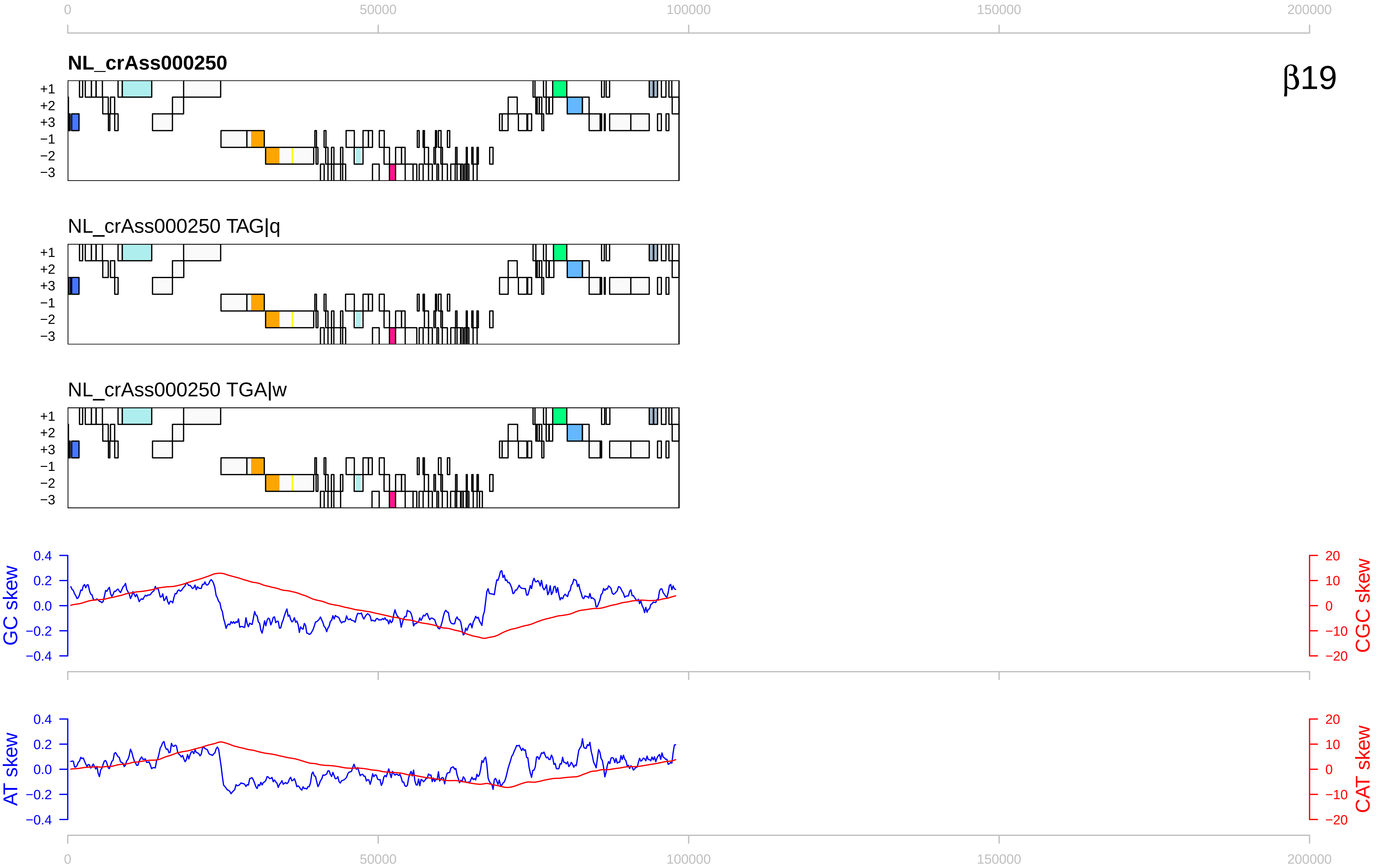

0
50000
100000
150000
200000
NL_crAss000250
β19
+1
+2
+3
−1
−2
−3
NL_crAss000250 TAG|q
NL_crAss000250 TGA|w
GC skew
CGC skew
AT skew
CAT skew
0.4
0.2
0.0
−0.2
−0.4
20
10
0
−10
−20

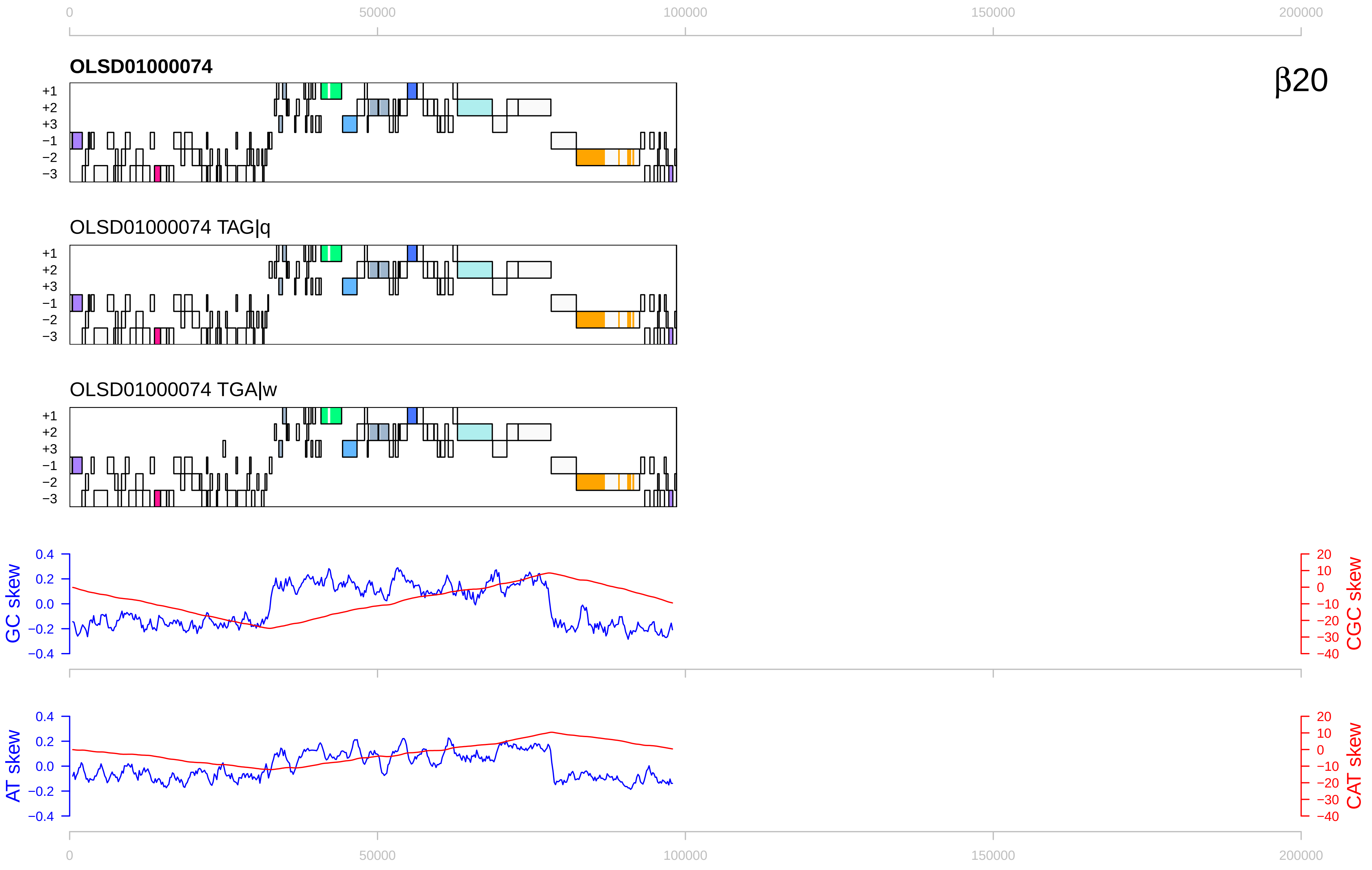

0
50000
100000
150000
200000
OLSD01000074
β20
+1
+2
+3
−1
−2
−3
OLSD01000074 TAG|q
OLSD01000074 TGA|w
GC skew
CGC skew
AT skew
CAT skew
0.4
0.2
0.0
−0.2
−0.4
20
10
0
−10
−20
−30
−40

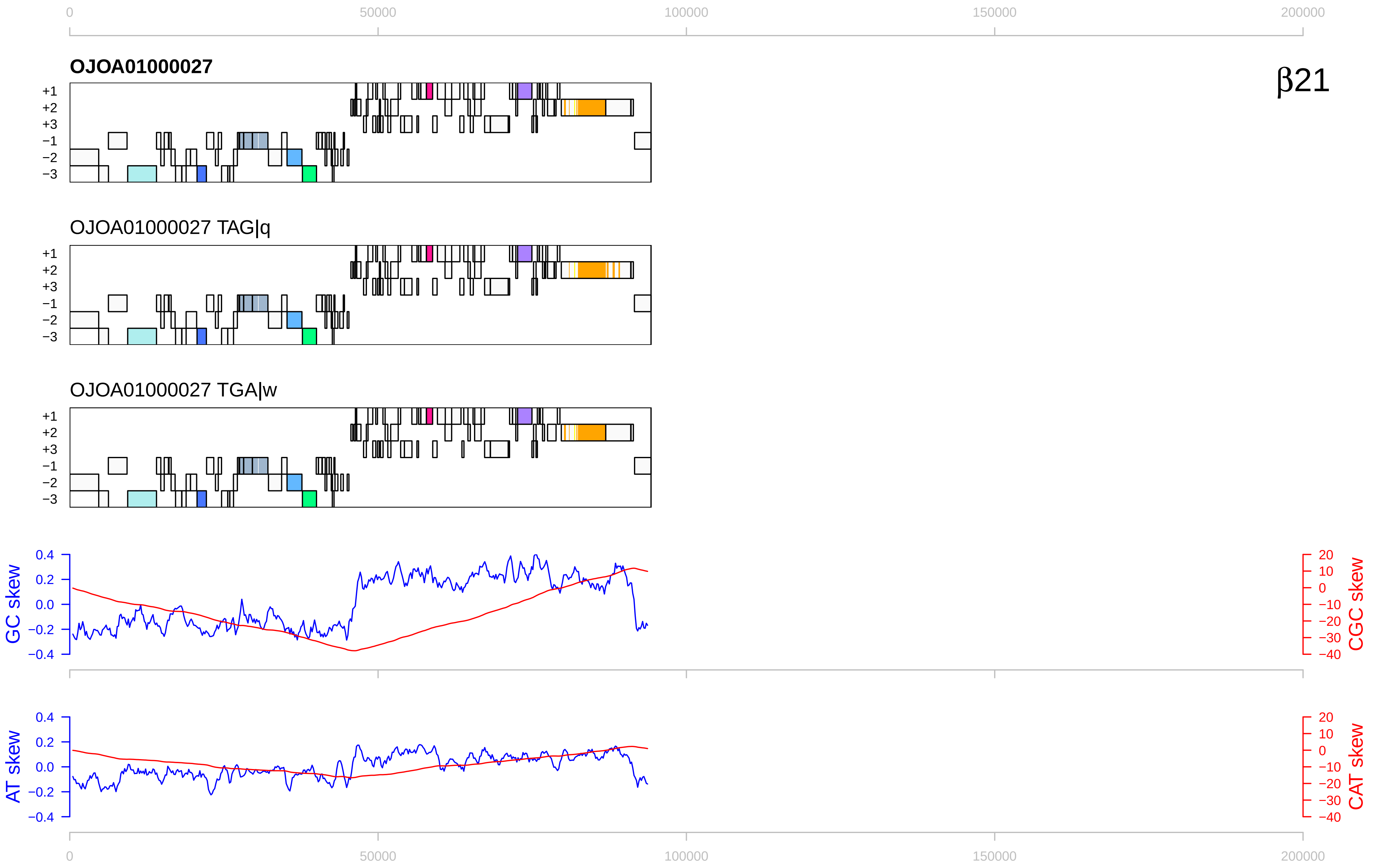

0
50000
100000
150000
200000
OJOA01000027
β21
+1
+2
+3
−1
−2
−3
OJOA01000027 TAG|q
OJOA01000027 TGA|w
GC skew
CGC skew
AT skew
CAT skew
0.4
0.2
0.0
−0.2
−0.4
20
10
0
−10
−20
−30
−40

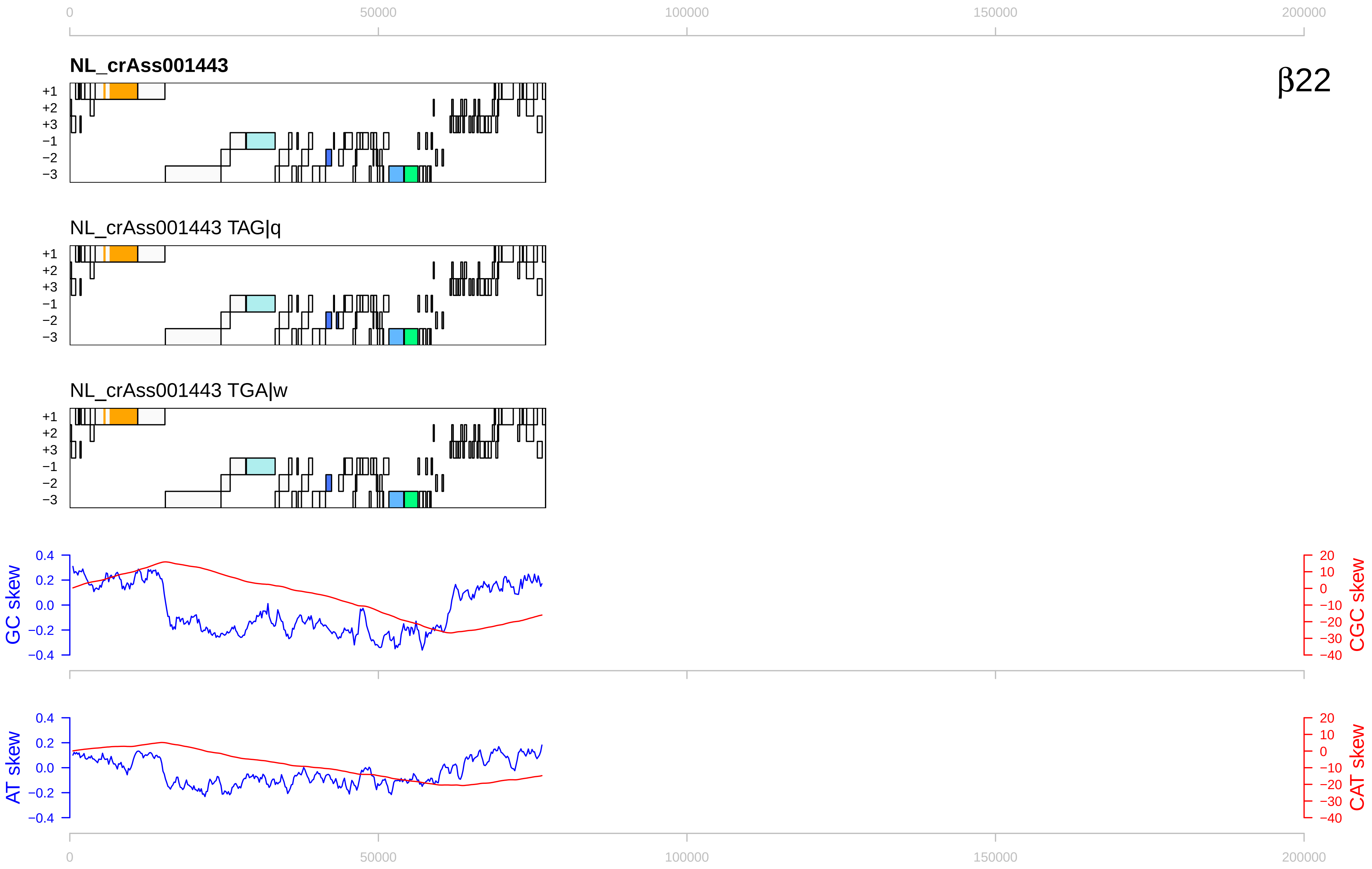

0
50000
100000
150000
200000
NL_crAss001443
β22
+1
+2
+3
−1
−2
−3
NL_crAss001443 TAG|q
NL_crAss001443 TGA|w
GC skew
CGC skew
AT skew
CAT skew
0.4
0.2
0.0
−0.2
−0.4
20
10
0
−10
−20
−30
−40

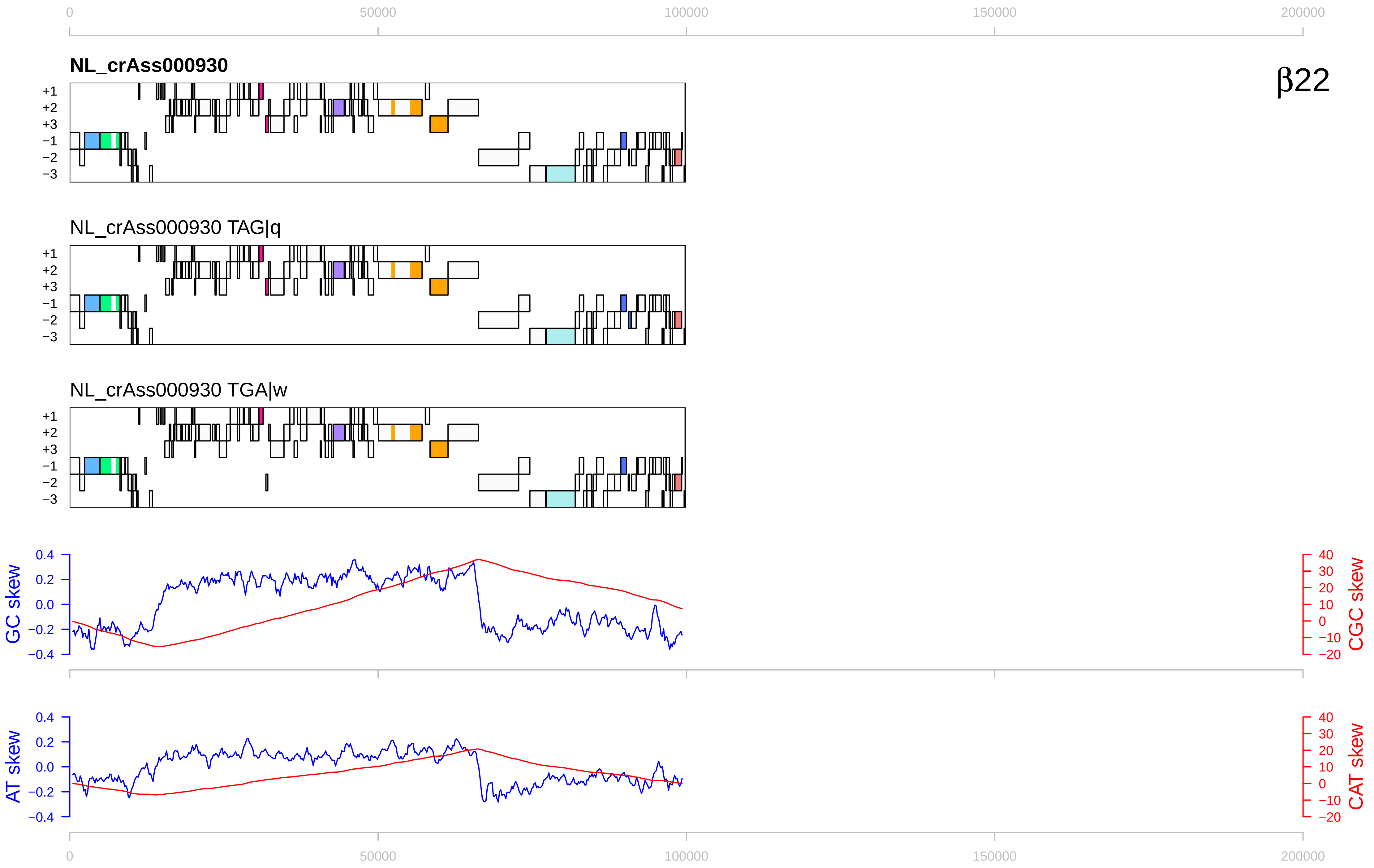

β22
NL_crAss000930
NL_crAss000930 TAG|q
NL_crAss000930 TGA|w
+1
+2
+3
−1
−2
−3
GC skew
CGC skew
AT skew
CAT skew
0
50000
100000
150000
200000

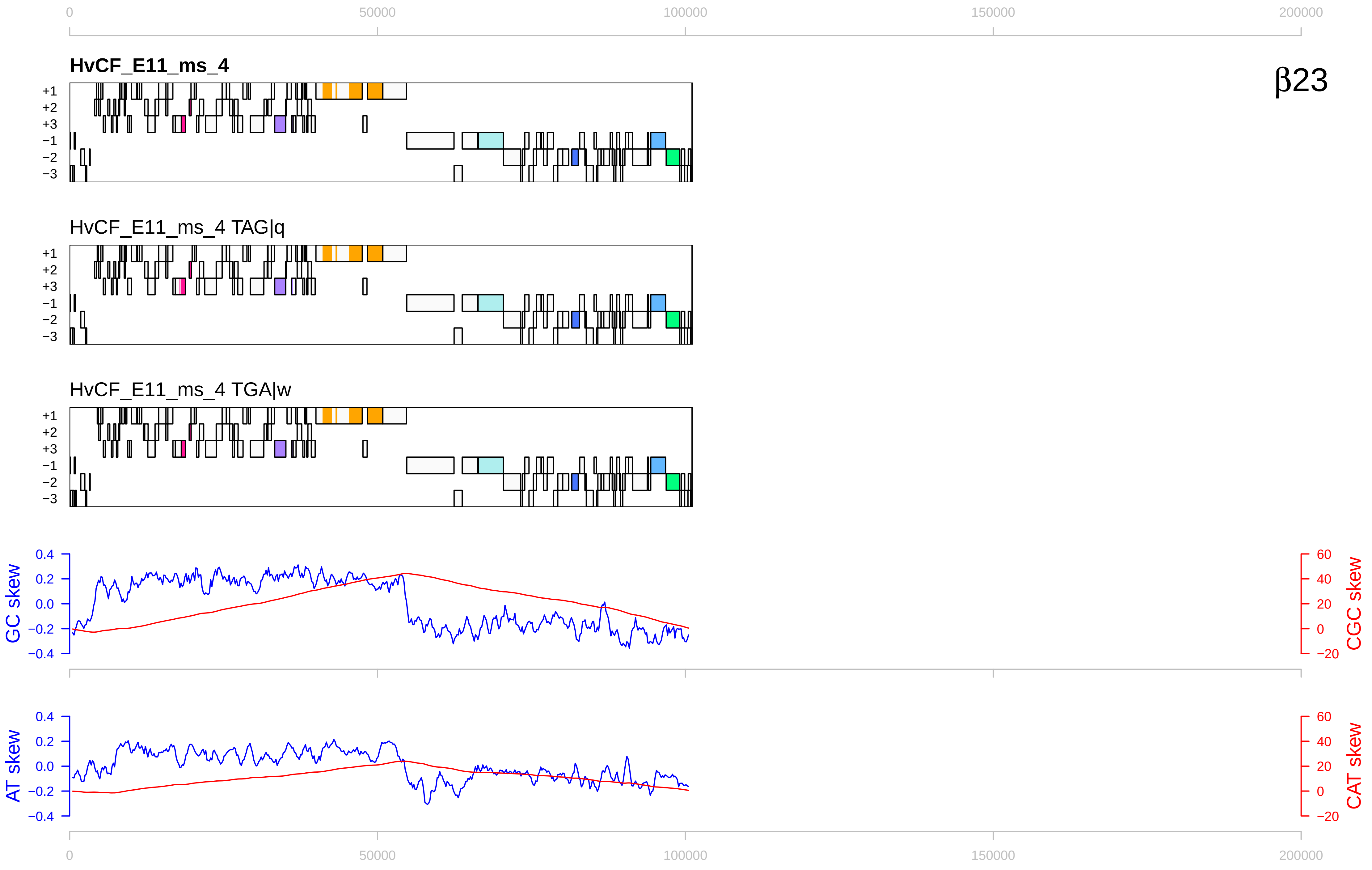

0
50000
100000
150000
200000
HvCF_E11_ms_4
β23
+1
+2
+3
−1
−2
−3
HvCF_E11_ms_4 TAG|q
HvCF_E11_ms_4 TGA|w
GC skew
CGC skew
AT skew
CAT skew
0.4
0.2
0.0
−0.2
−0.4
60
40
20
0
−20

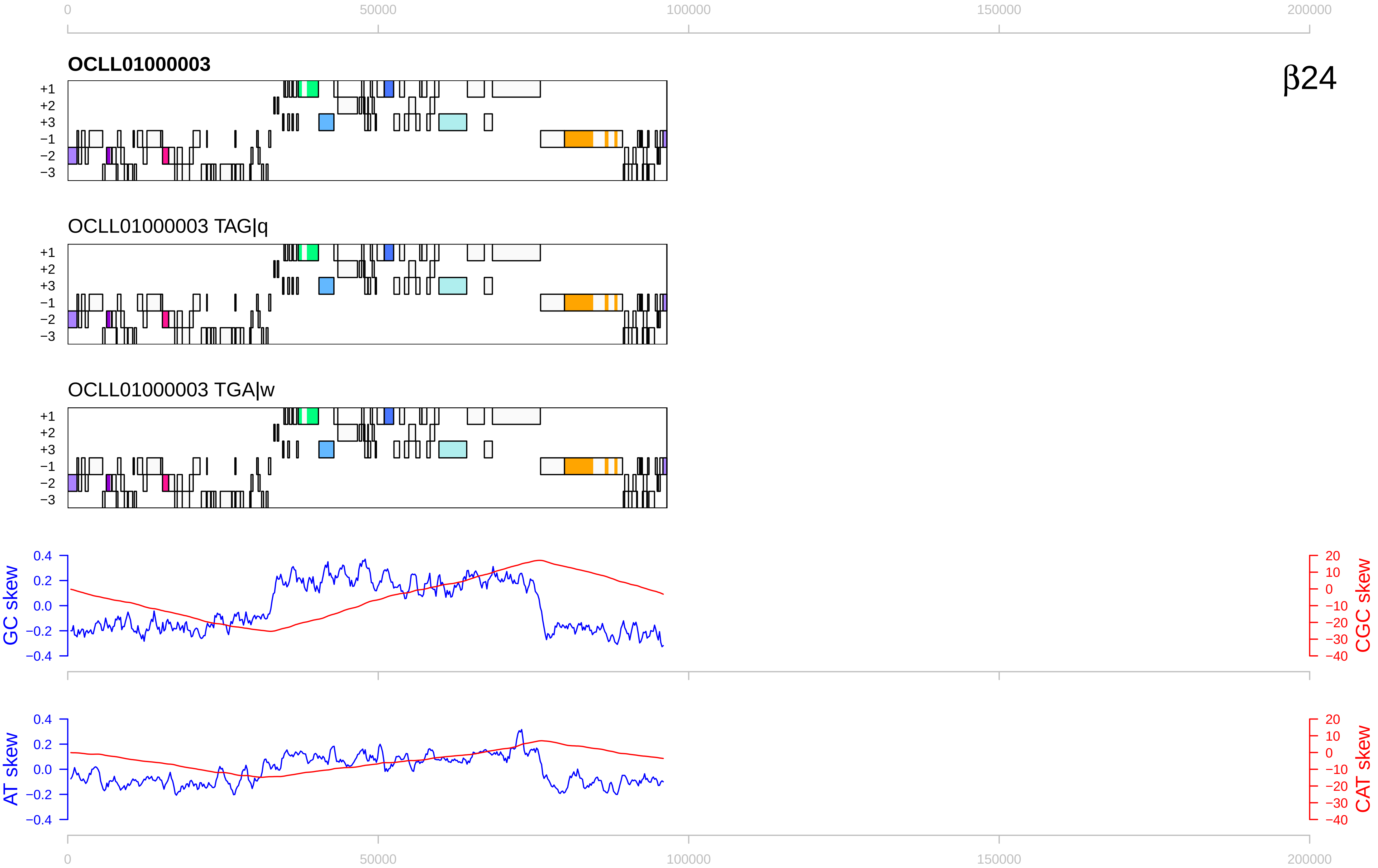

β24
OCLL01000003
OCLL01000003 TAG|q
OCLL01000003 TGA|w
+1
+2
+3
−1
−2
−3
GC skew
CGC skew
AT skew
CAT skew
0.4
0.2
0.0
−0.2
−0.4
20
10
0
−10
−20
−30
−40
0
50000
100000
150000
200000

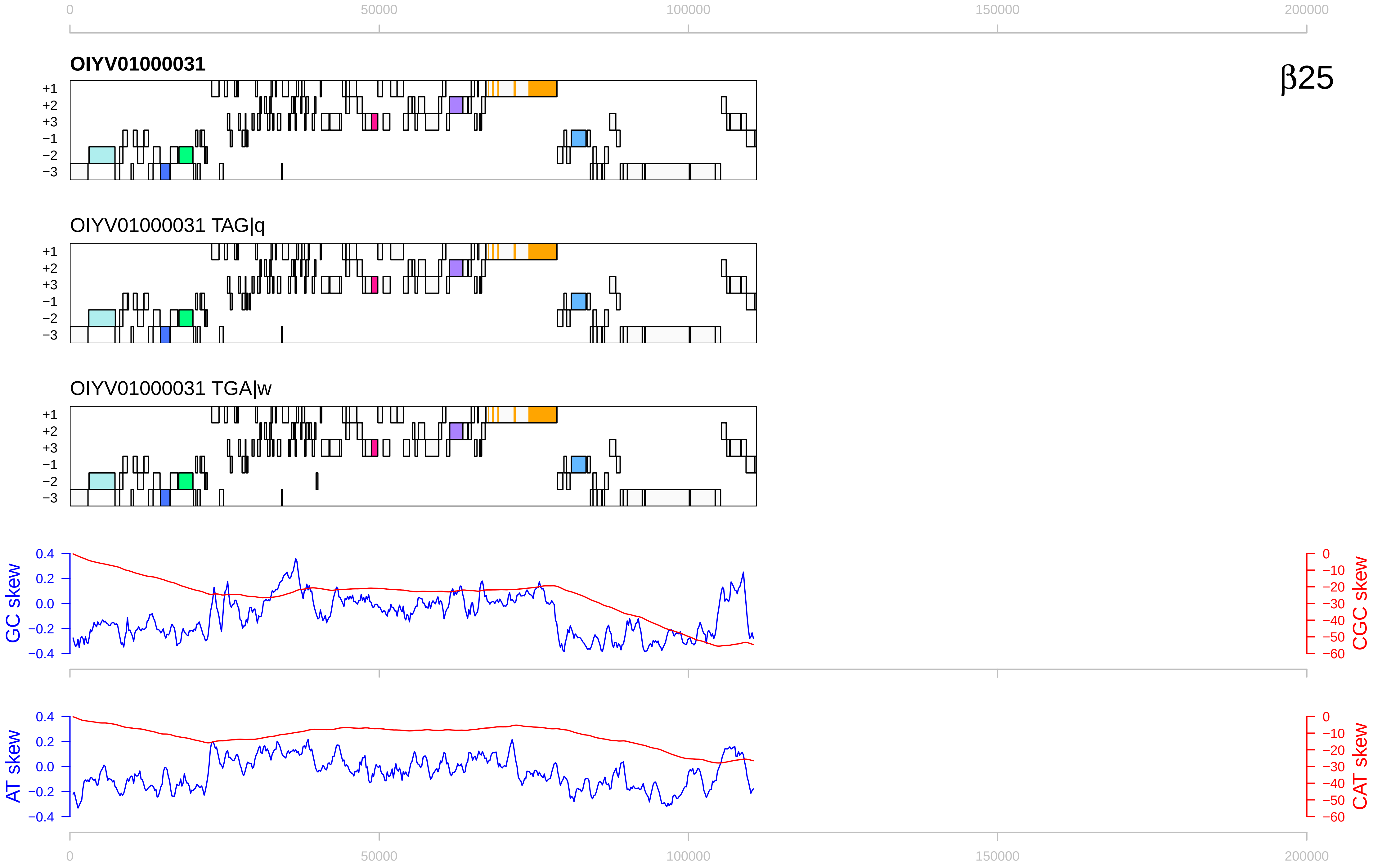

β25
OIYV01000031
OIYV01000031 TAG|q
OIYV01000031 TGA|w
+1
+2
+3
−1
−2
−3
GC skew
CGC skew
AT skew
CAT skew
0
50000
100000
150000
200000
0.4
0.2
0.0
−0.2
−0.4
0
−10
−20
−30
−40
−50
−60

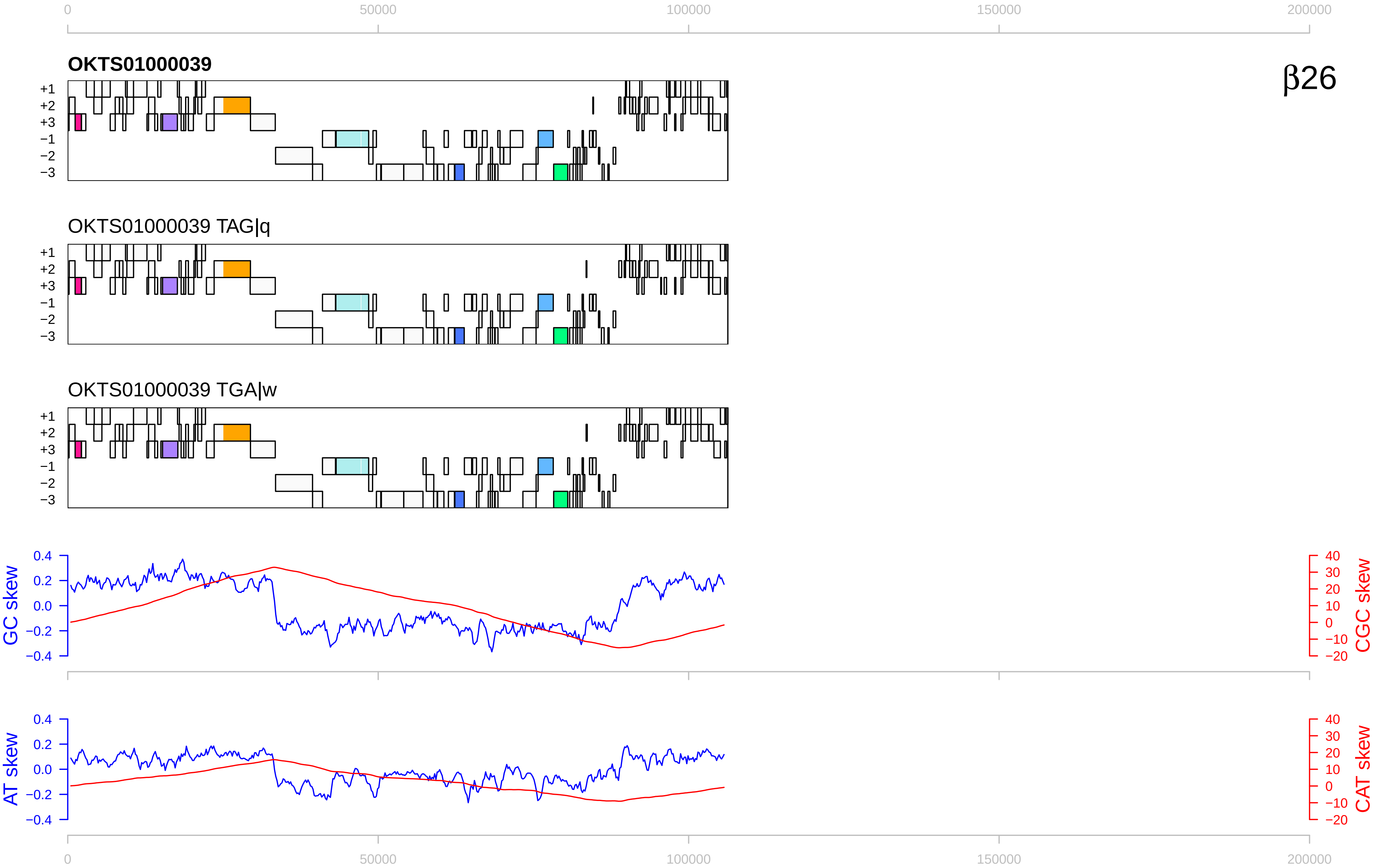

β26
OKTS01000039
OKTS01000039 TAG|q
OKTS01000039 TGA|w
+1
+2
+3
−1
−2
−3
GC skew
CGC skew
AT skew
CAT skew
0
50000
100000
150000
200000
0.4
0.2
0.0
−0.2
−0.4
40
30
20
10
0
−10
−20

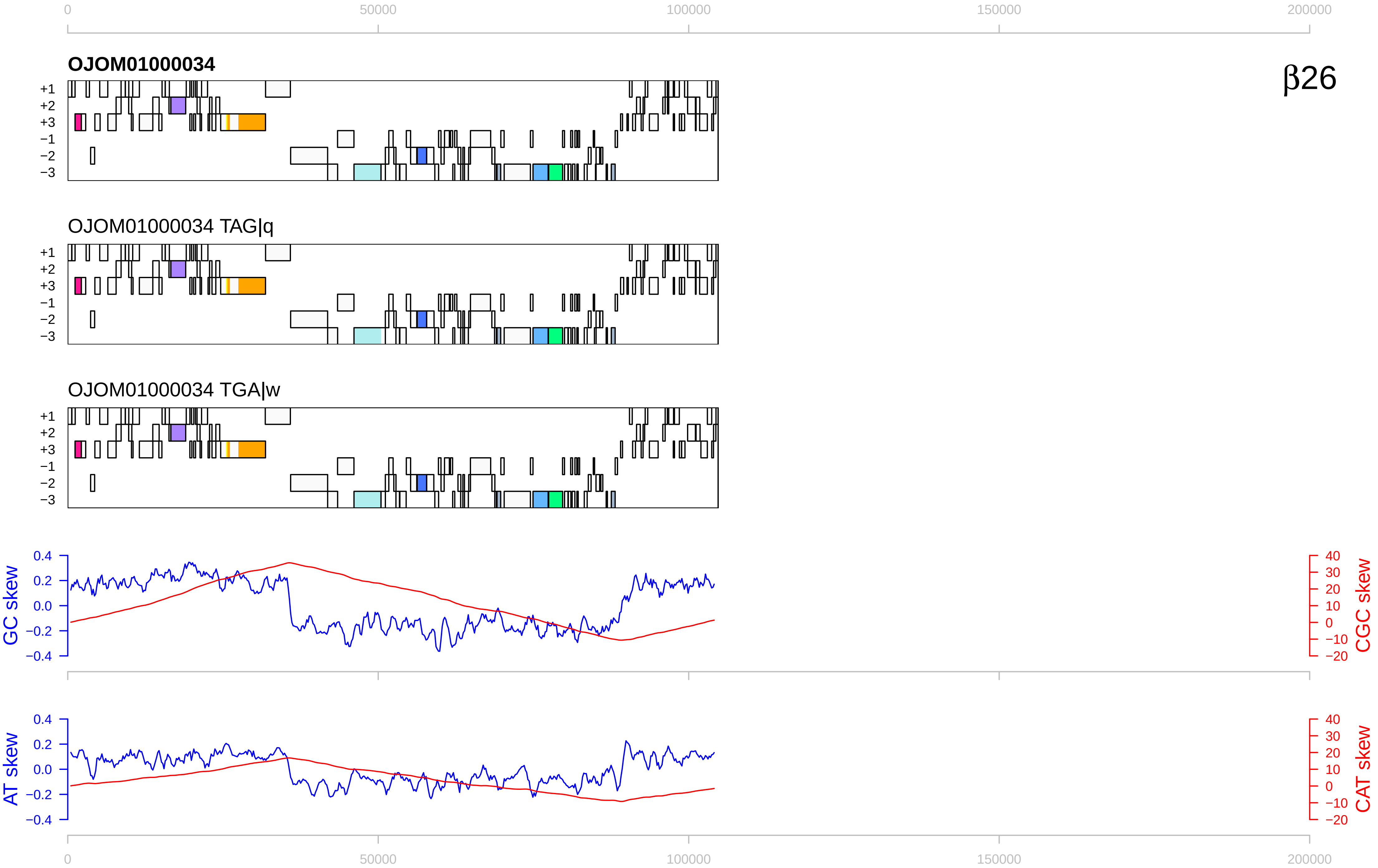

0
50000
100000
150000
200000
OJOM01000034
β26
+1
+2
+3
−1
−2
−3
OJOM01000034 TAG|q
OJOM01000034 TGA|w
GC skew
CGC skew
AT skew
CAT skew
0.4
0.2
0.0
−0.2
−0.4
40
30
20
10
0
−10
−20

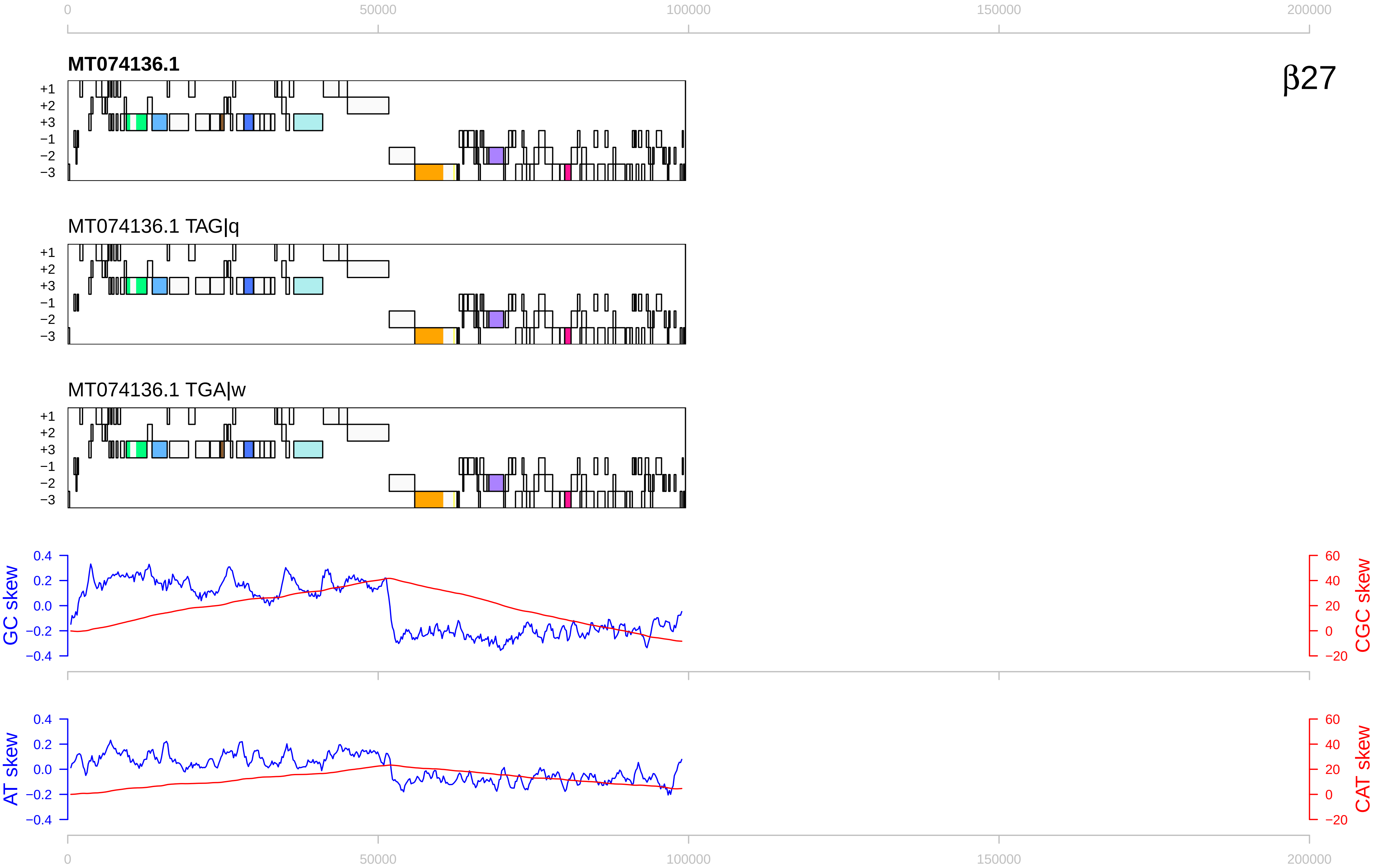

β27
MT074136.1
MT074136.1 TAG|q
MT074136.1 TGA|w
+1
+2
+3
−1
−2
−3
GC skew
CGC skew
AT skew
CAT skew
0
50000
100000
150000
200000
0.4
0.2
0.0
−0.2
−0.4
60
40
20
0
−20

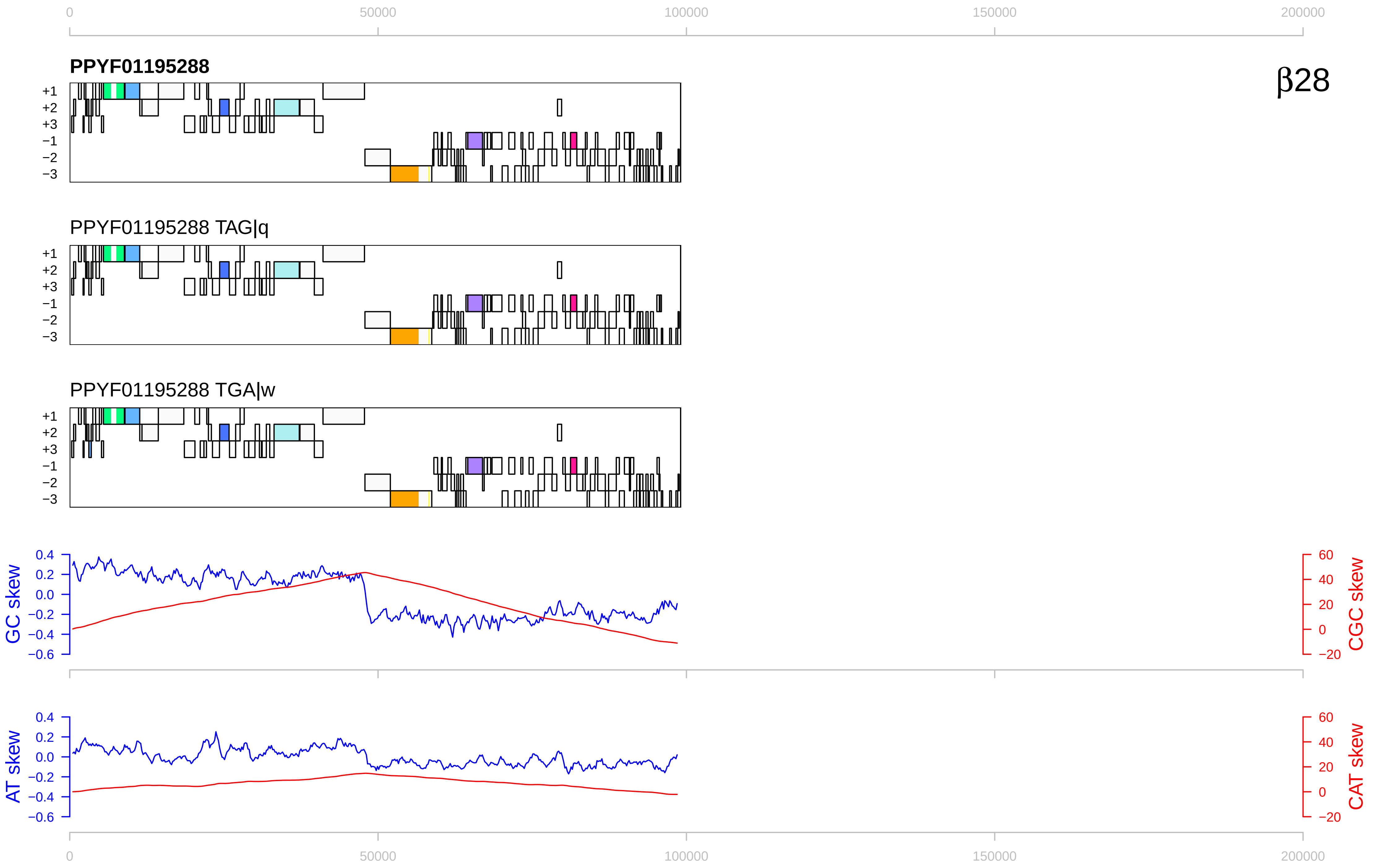
β28
PPYF01195288
PPYF01195288 TAG|q
PPYF01195288 TGA|w
+1
+2
+3
−1
−2
−3
GC skew
CGC skew
AT skew
CAT skew
0
50000
100000
150000
200000

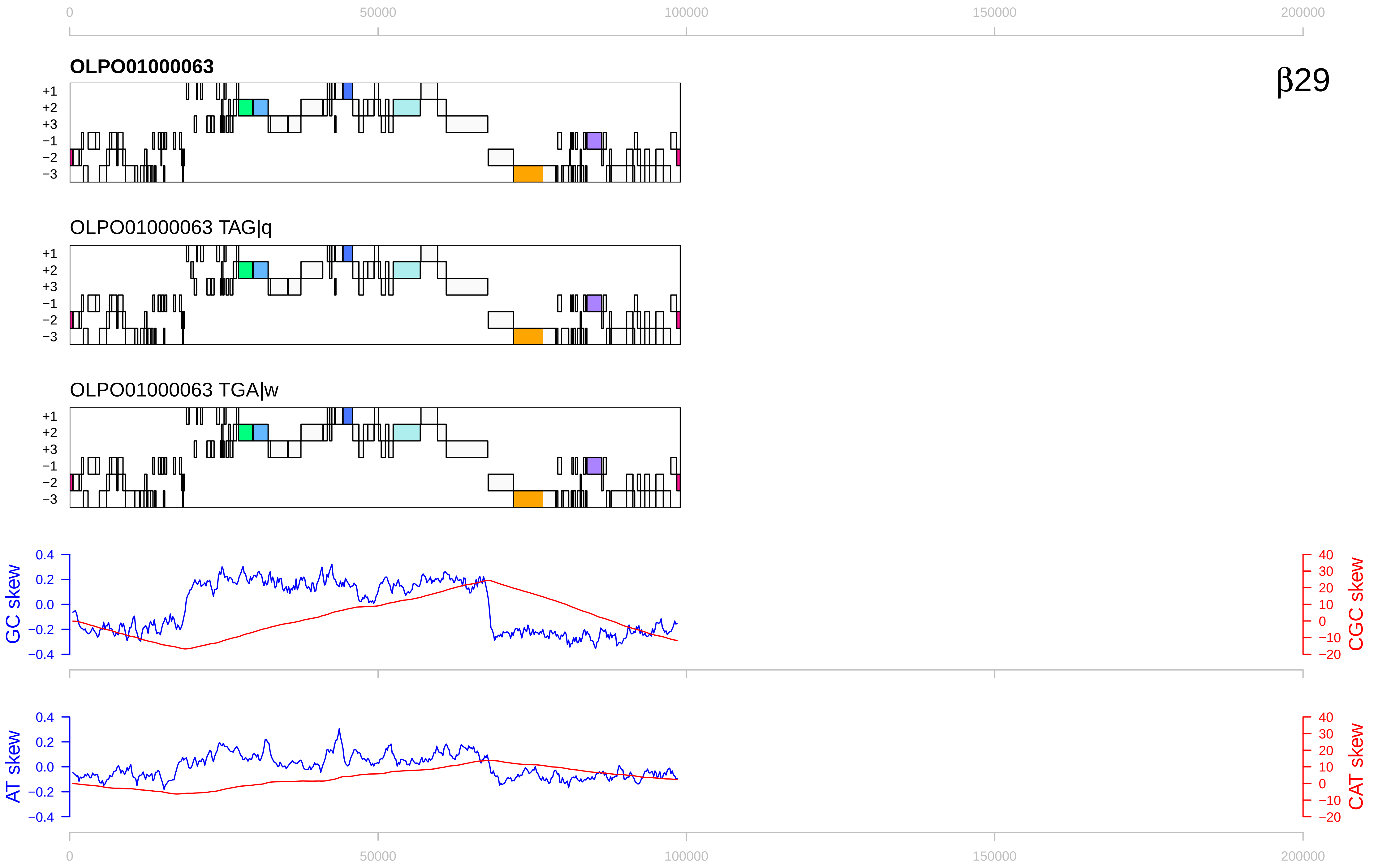

0
50000
100000
150000
200000
OLPO01000063
β29
+1
+2
+3
−1
−2
−3
OLPO01000063 TAG|q
OLPO01000063 TGA|w
GC skew
CGC skew
0.4
0.2
0.0
−0.2
−0.4
40
30
20
10
0
−10
−20
AT skew
CAT skew

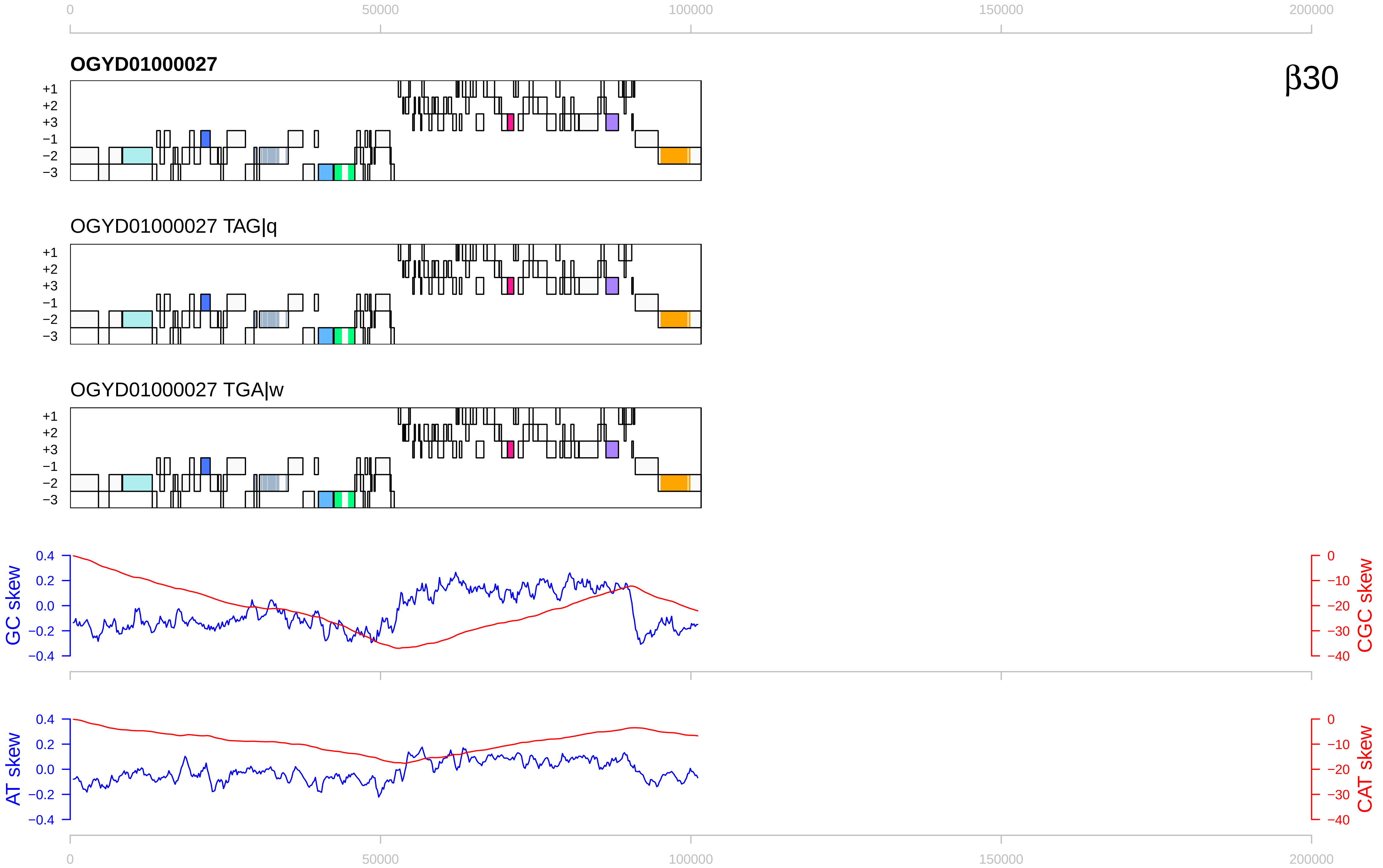

0
50000
100000
150000
200000
OGYD01000027
β30
+1
+2
+3
−1
−2
−3
OGYD01000027 TAG|q
OGYD01000027 TGA|w
GC skew
CGC skew
0.4
0.2
0.0
−0.2
−0.4
0
−10
−20
−30
−40
AT skew
CAT skew

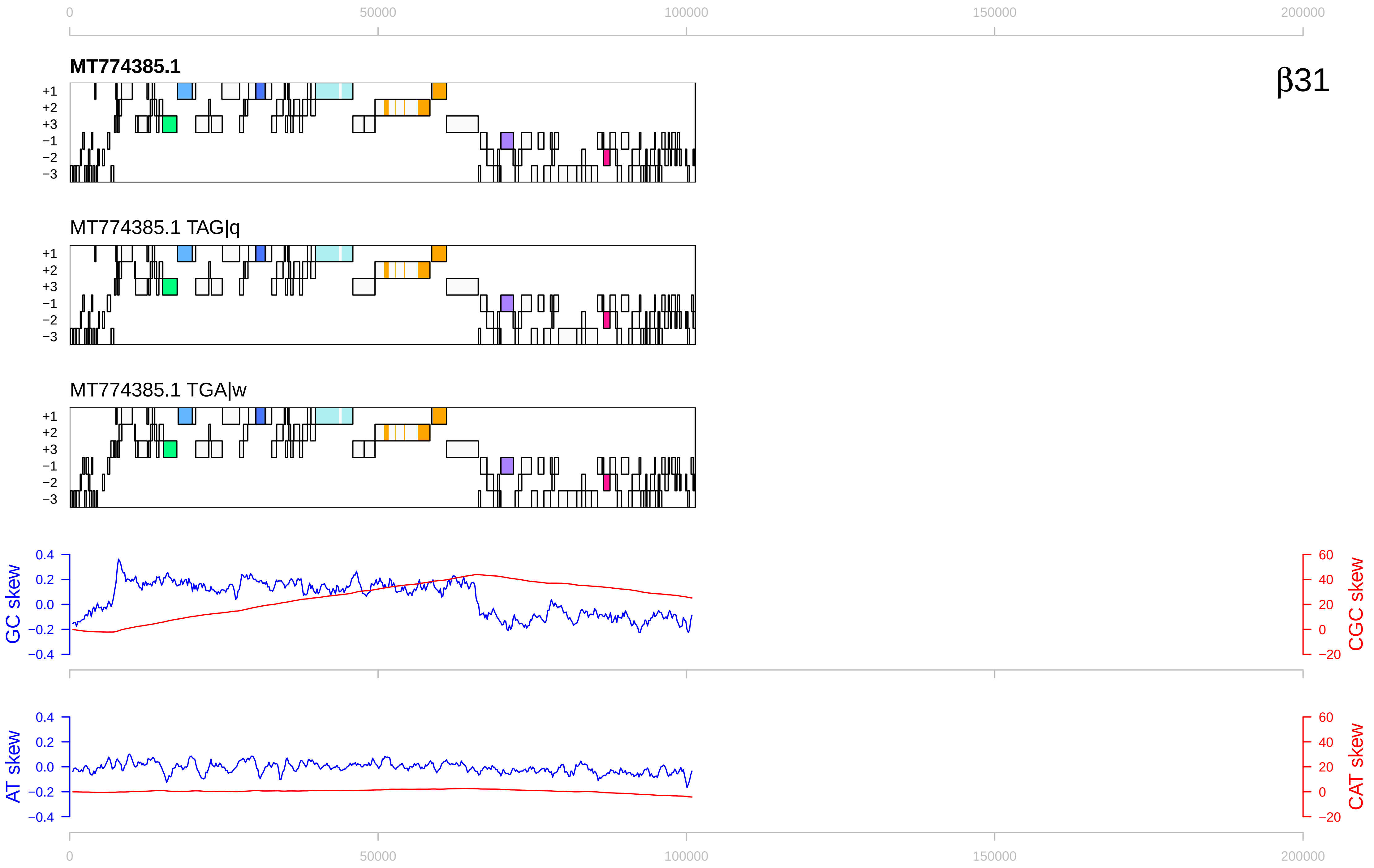

0
50000
100000
150000
200000
MT774385.1
β31
+1
+2
+3
−1
−2
−3
MT774385.1 TAG|q
MT774385.1 TGA|w
GC skew
CGC skew
AT skew
CAT skew
0.4
0.2
0.0
−0.2
−0.4
60
40
20
0
−20

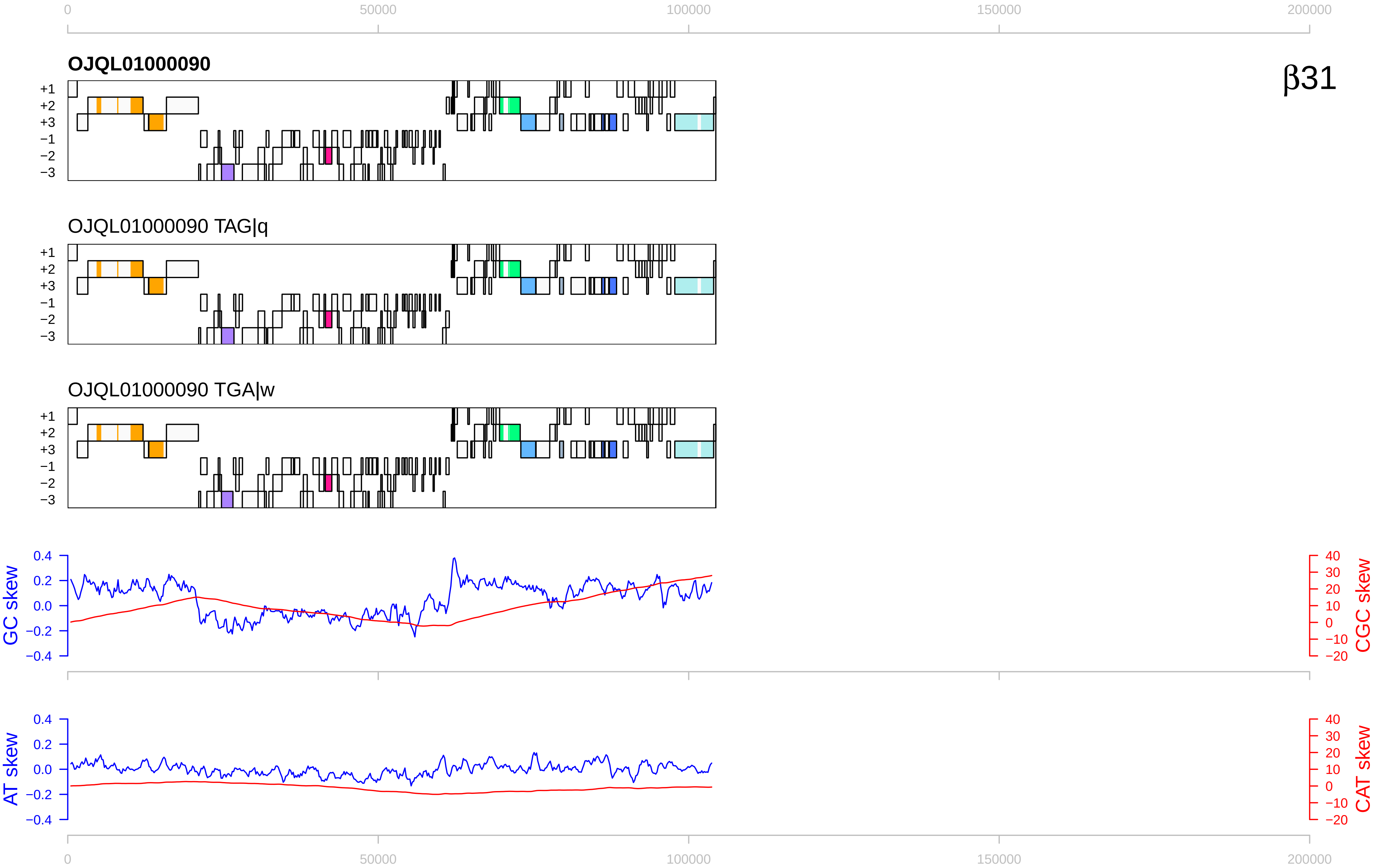
β31
OJQL01000090
OJQL01000090 TAG|q
OJQL01000090 TGA|w
+1
+2
+3
−1
−2
−3
GC skew
CGC skew
AT skew
CAT skew
0
50000
100000
150000
200000

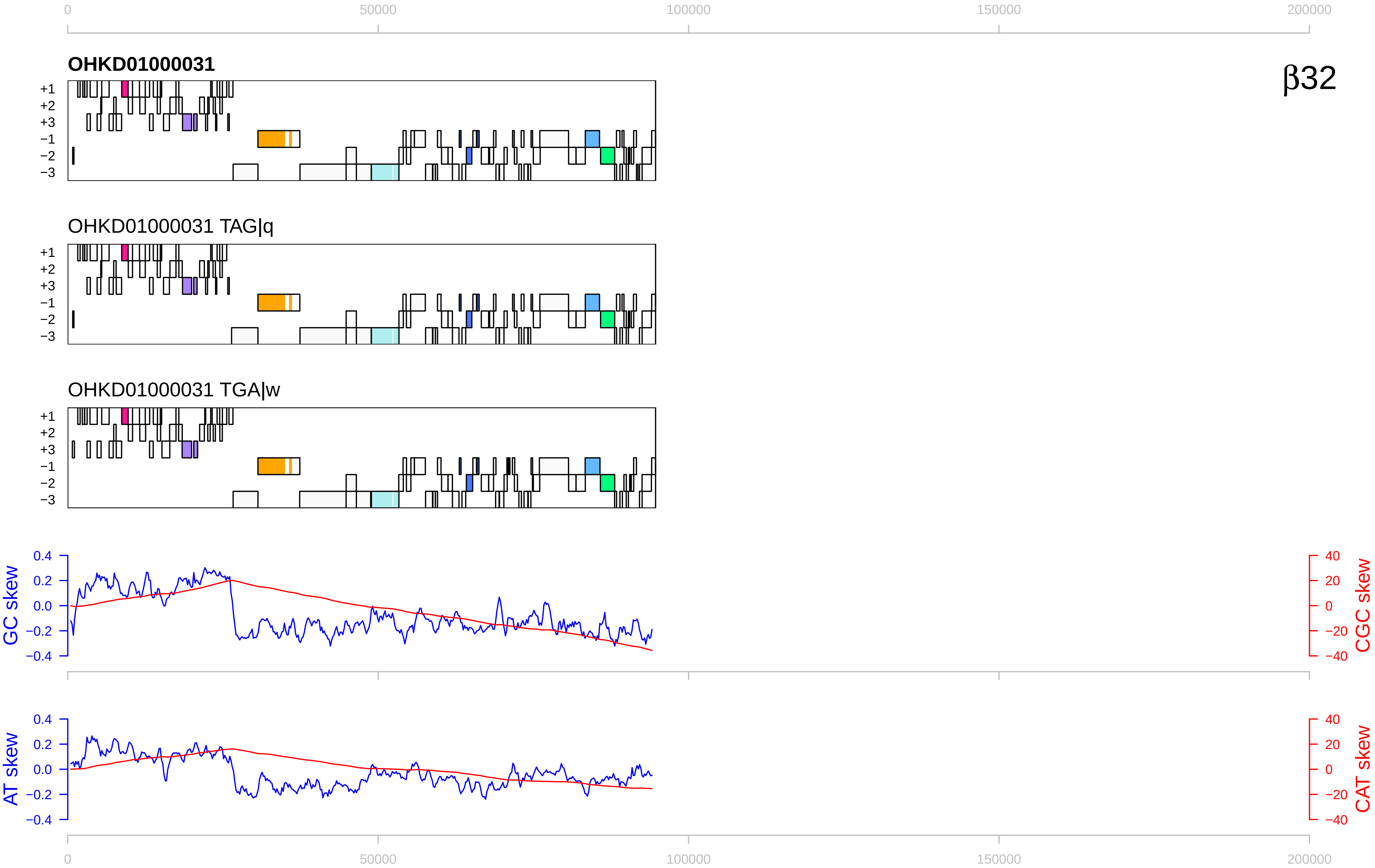

0
50000
100000
150000
200000
OHKD01000031
β32
+1
+2
+3
−1
−2
−3
OHKD01000031 TAG|q
OHKD01000031 TGA|w
GC skew
CGC skew
AT skew
CAT skew
0.4
0.2
0.0
−0.2
−0.4
40
20
0
−20
−40

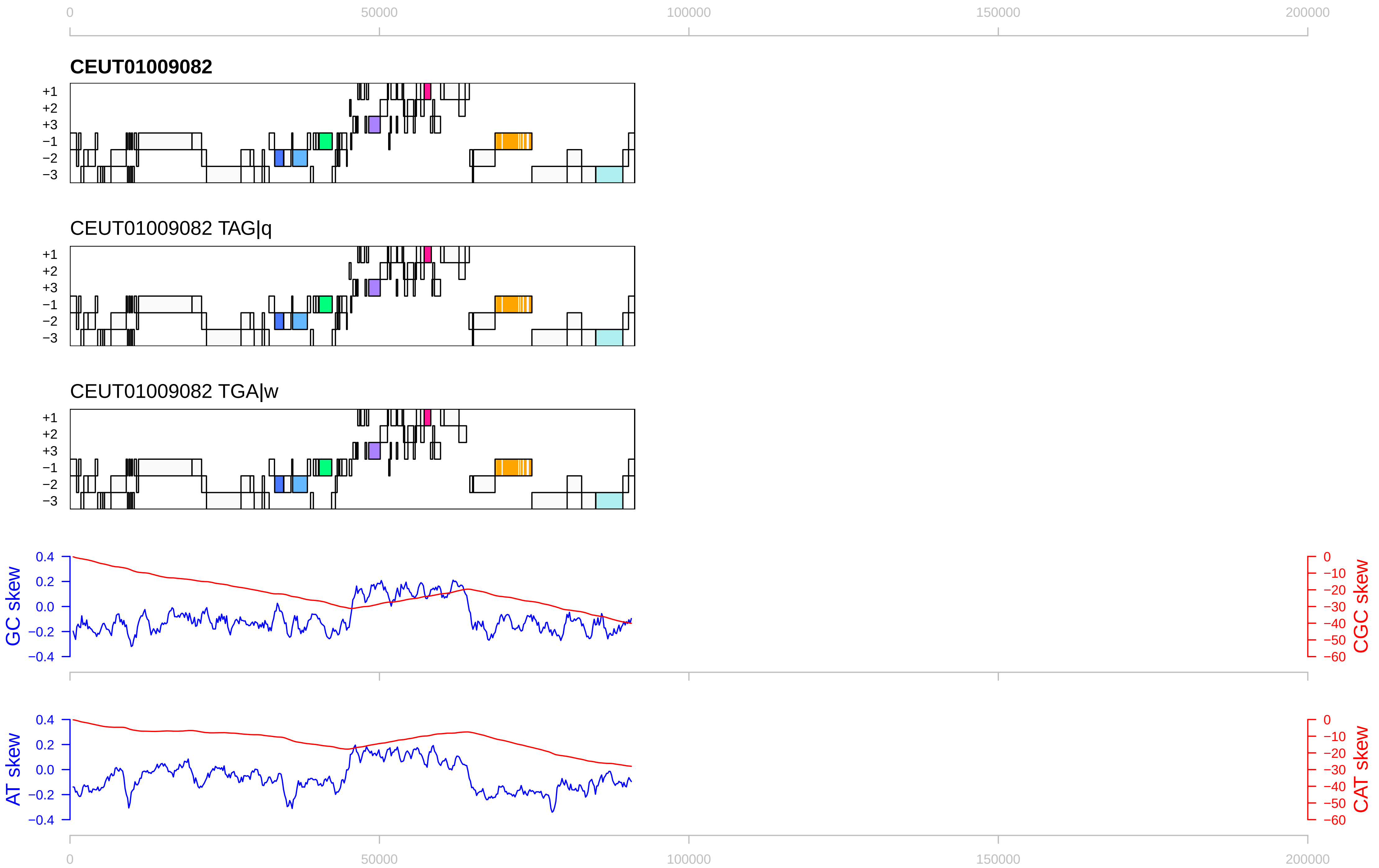

0
50000
100000
150000
200000
CEUT01009082
+1
+2
+3
−1
−2
−3
CEUT01009082 TAG|q
CEUT01009082 TGA|w
GC skew
CGC skew
AT skew
CAT skew
0.4
0.2
0.0
−0.2
−0.4
0
−10
−20
−30
−40
−50
−60

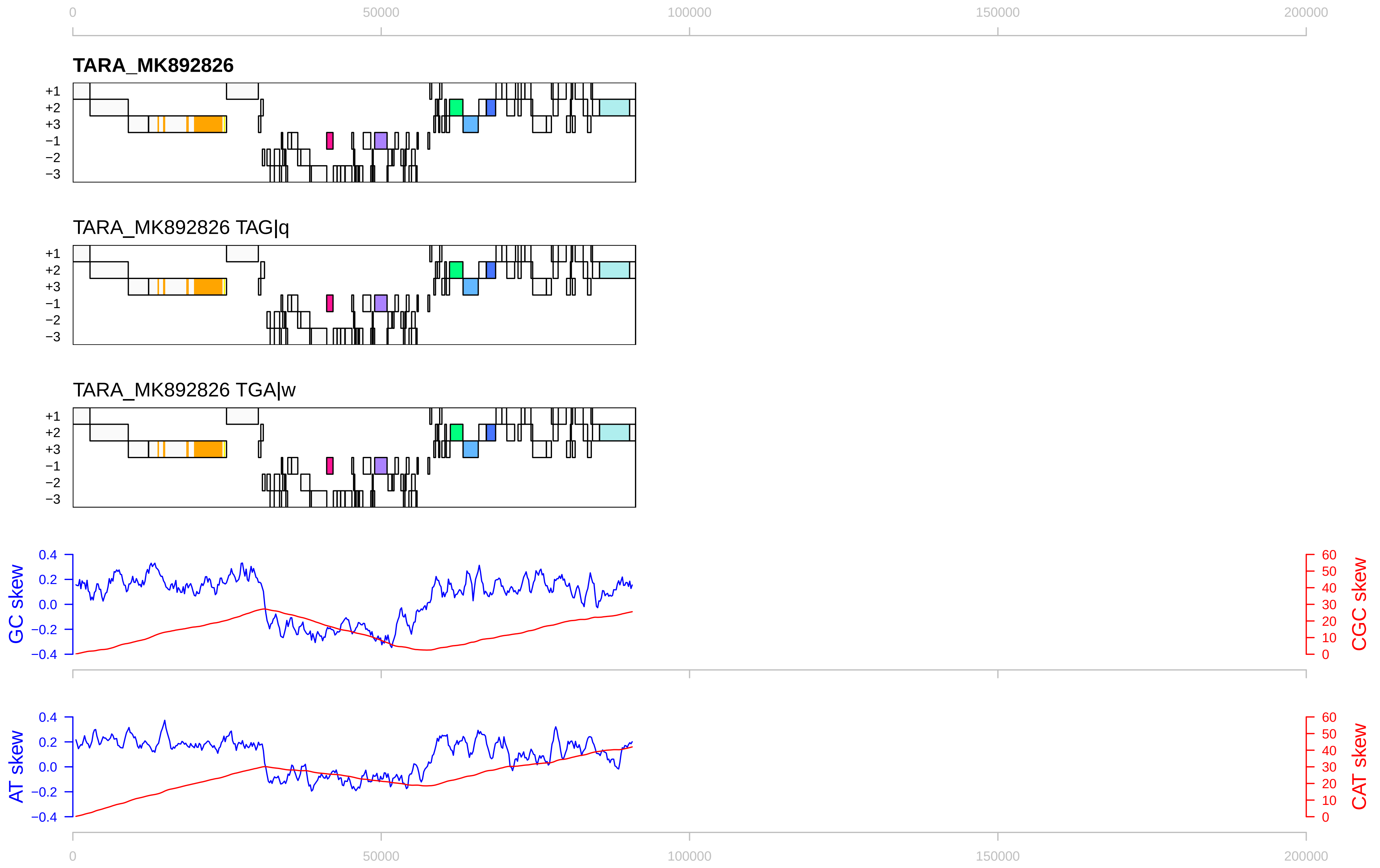

0
50000
100000
150000
200000
TARA_MK892826
+1
+2
+3
−1
−2
−3
TARA_MK892826 TAG|q
TARA_MK892826 TGA|w
GC skew
CGC skew
AT skew
CAT skew
0.4
0.2
0.0
−0.2
−0.4
60
50
40
30
20
10
0

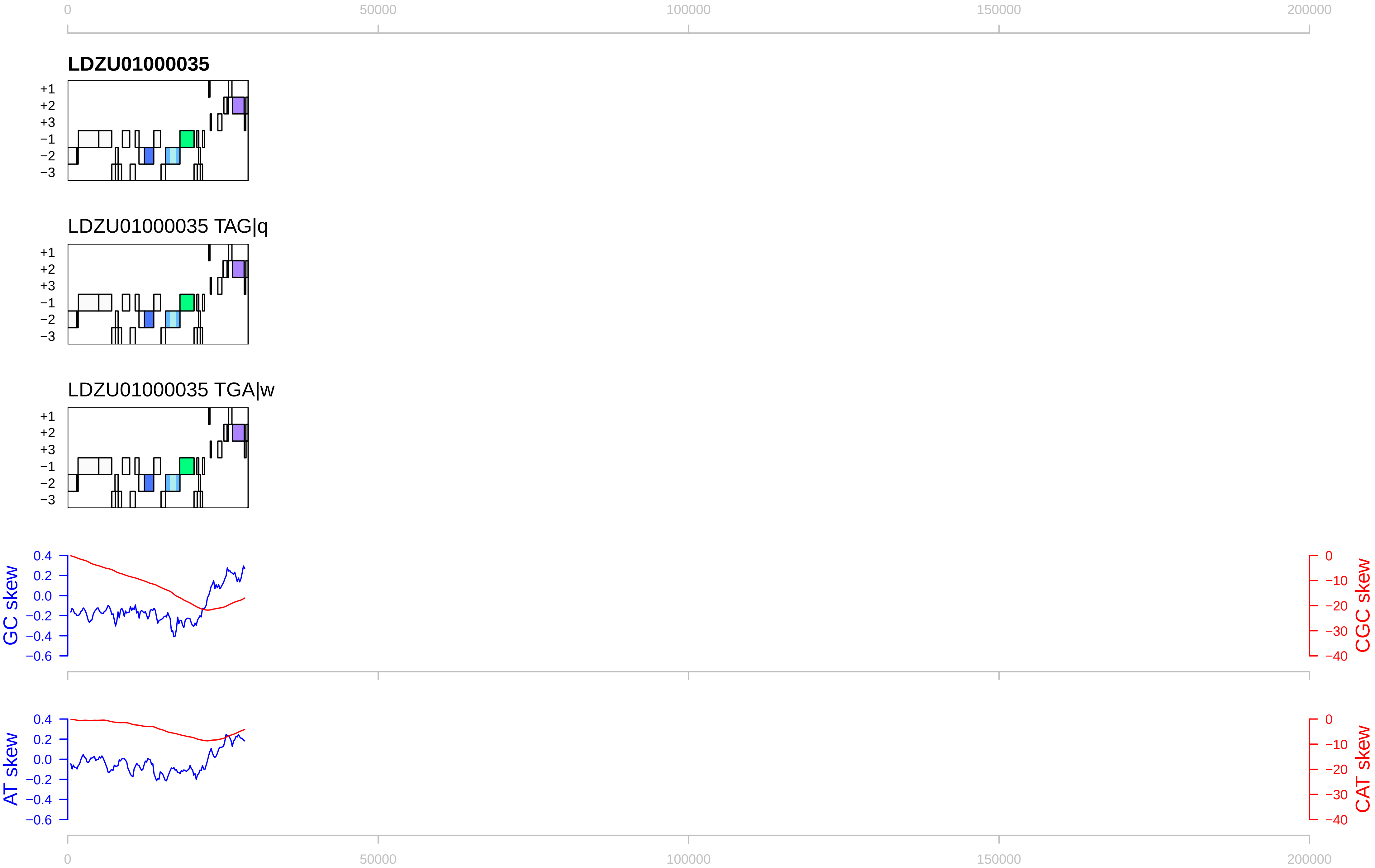

0
50000
100000
150000
200000
LDZU01000035
+1
+2
+3
−1
−2
−3
LDZU01000035 TAG|q
LDZU01000035 TGA|w
GC skew
CGC skew
AT skew
CAT skew
0.4
0.2
0.0
−0.2
−0.4
−0.6
0
−10
−20
−30
−40

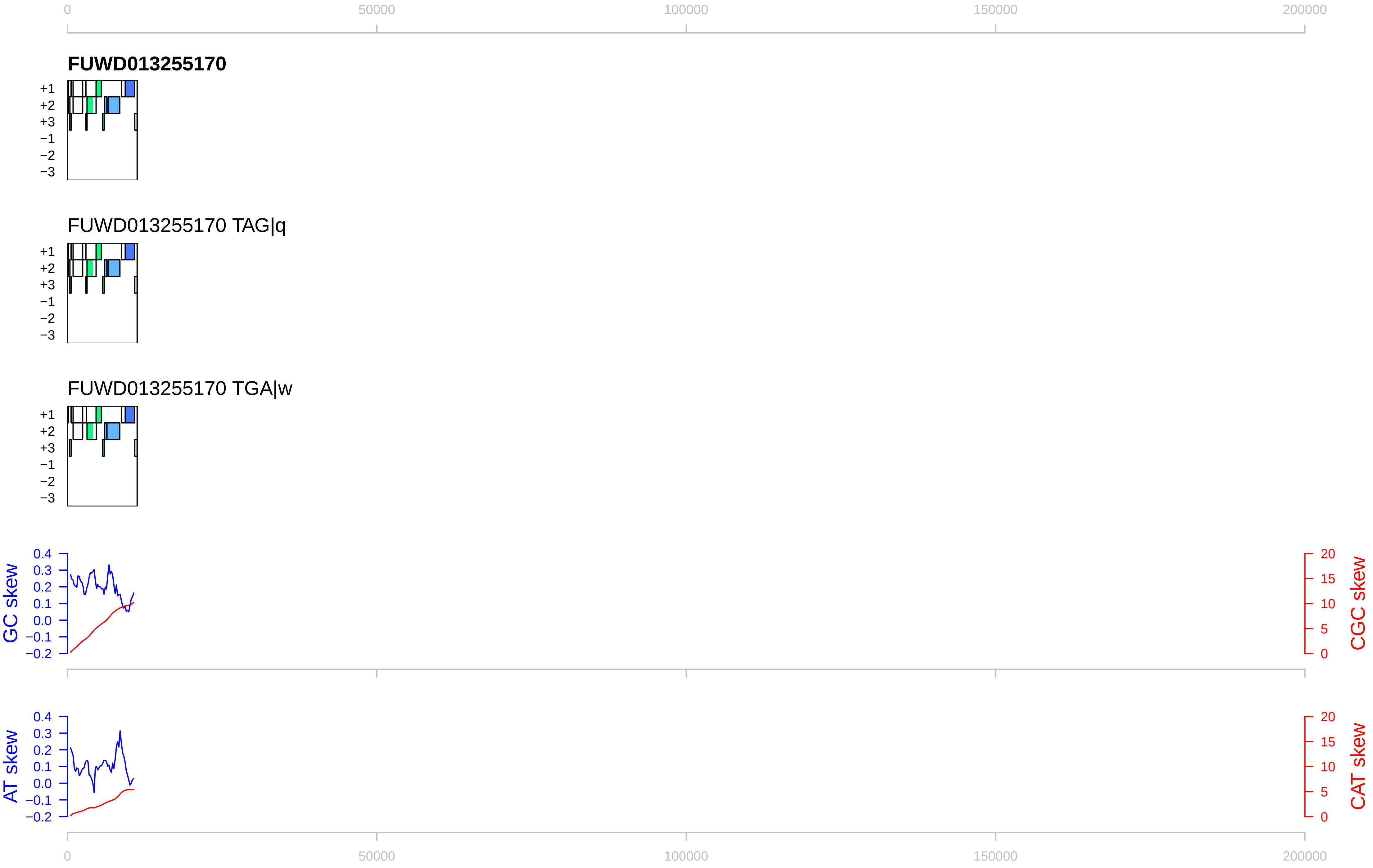

0
50000
100000
150000
200000
FUWD013255170
+1
+2
+3
−1
−2
−3
FUWD013255170 TAG|q
FUWD013255170 TGA|w
GC skew
0.4
0.3
0.2
0.1
0.0
−0.1
−0.2
CGC skew
20
15
10
5
0
AT skew
CAT skew

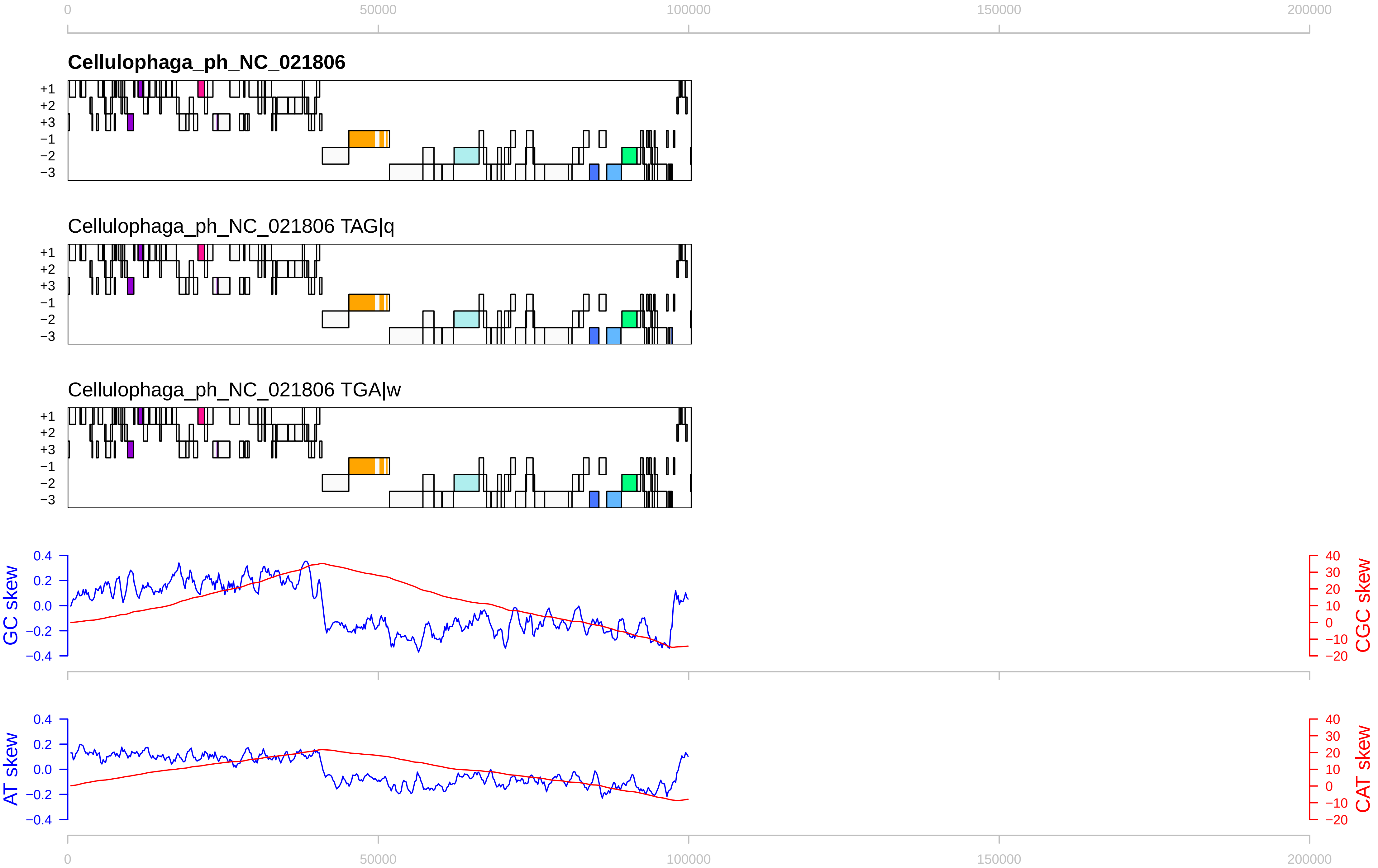

0
50000
100000
150000
200000
Cellulophaga_ph_NC_021806
+1
+2
+3
−1
−2
−3
Cellulophaga_ph_NC_021806 TAG|q
Cellulophaga_ph_NC_021806 TGA|w
GC skew
CGC skew
AT skew
CAT skew
0.4
0.2
0.0
−0.2
−0.4
40
30
20
10
0
−10
−20

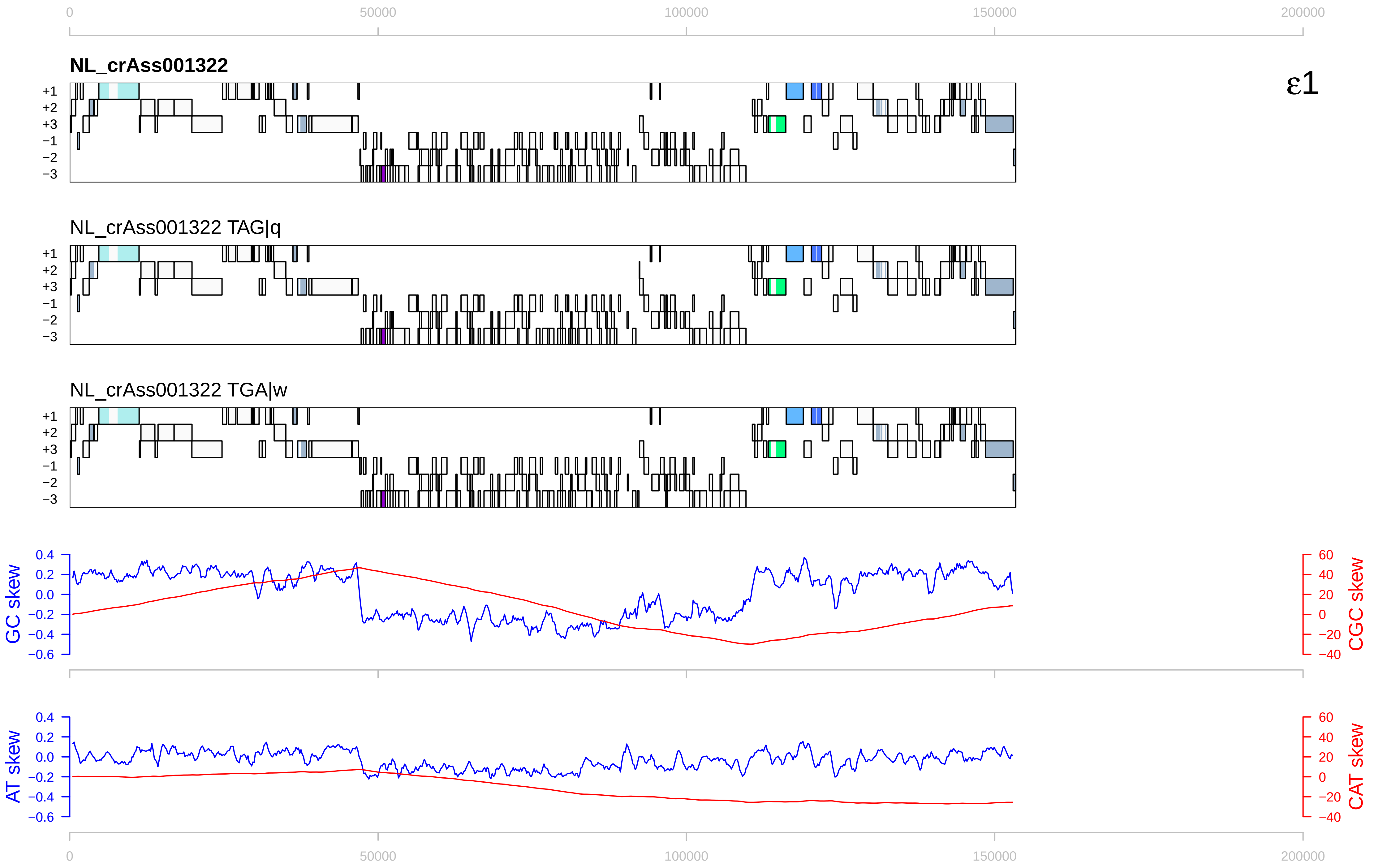

0
50000
100000
150000
200000
NL_crAss001322
ε1
+1
+2
+3
−1
−2
−3
NL_crAss001322 TAG|q
NL_crAss001322 TGA|w
GC skew
CGC skew
AT skew
CAT skew
0.4
0.2
0.0
−0.2
−0.4
−0.6
60
40
20
0
−20
−40

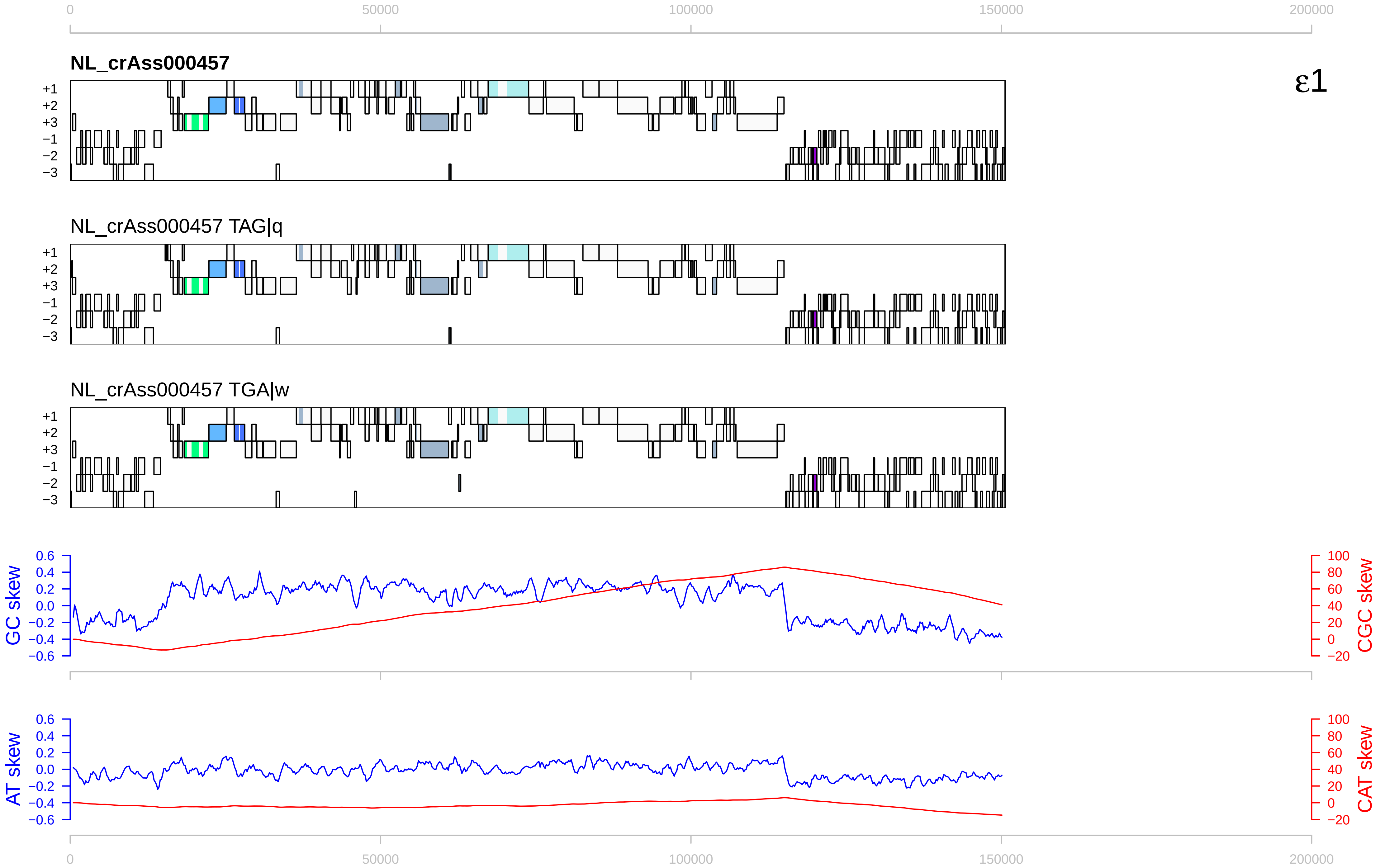

0
50000
100000
150000
200000
NL_crAss000457
ε1
+1
+2
+3
−1
−2
−3
NL_crAss000457 TAG|q
NL_crAss000457 TGA|w
GC skew
CGC skew
AT skew
CAT skew
0.6
0.4
0.2
0.0
−0.2
−0.4
−0.6
100
80
60
40
20
0
−20

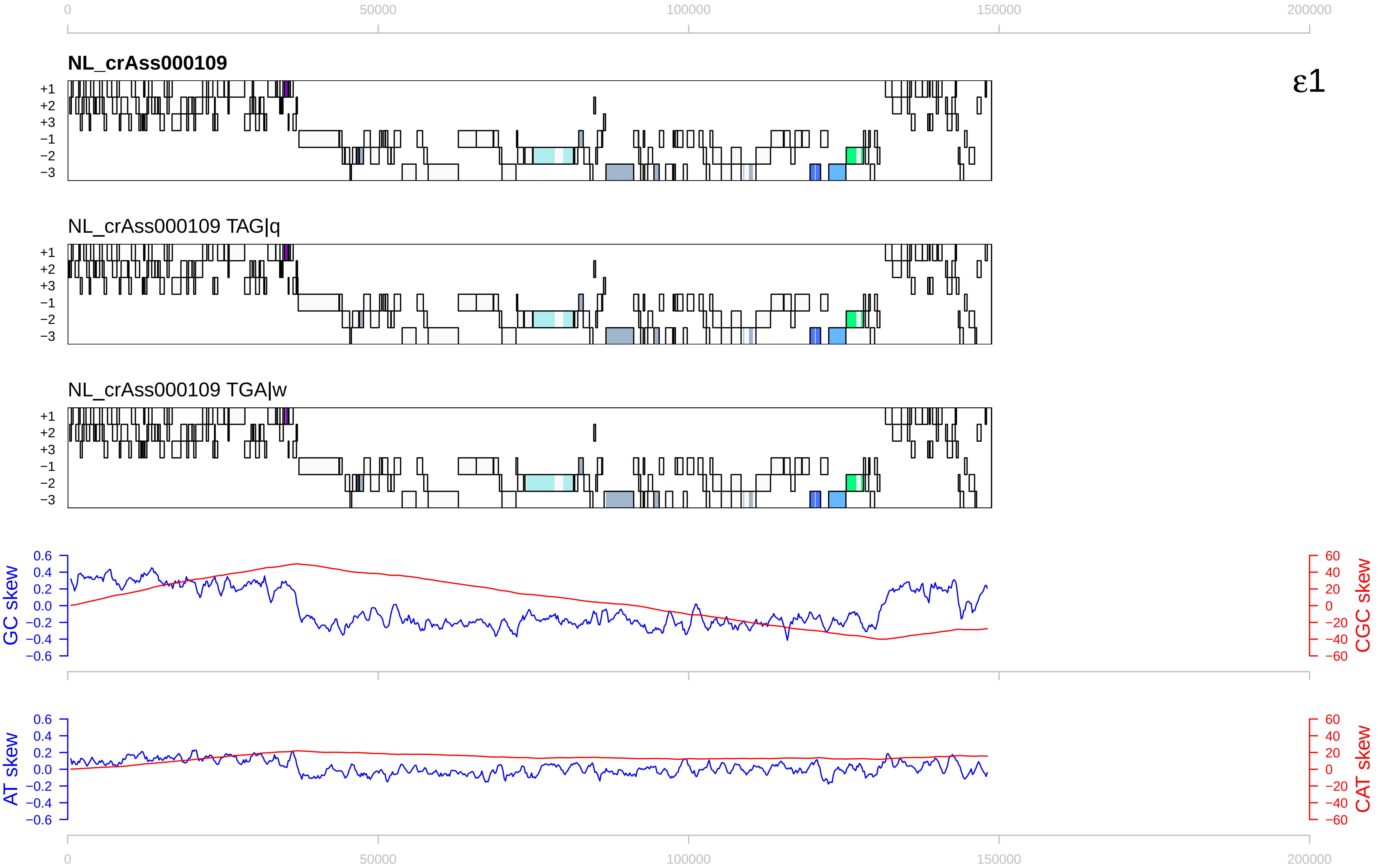

0
50000
100000
150000
200000
NL_crAss000109
ε1
+1
+2
+3
−1
−2
−3
NL_crAss000109 TAG|q
NL_crAss000109 TGA|w
GC skew
CGC skew
AT skew
CAT skew
0.6
0.4
0.2
0.0
−0.2
−0.4
−0.6
60
40
20
0
−20
−40
−60

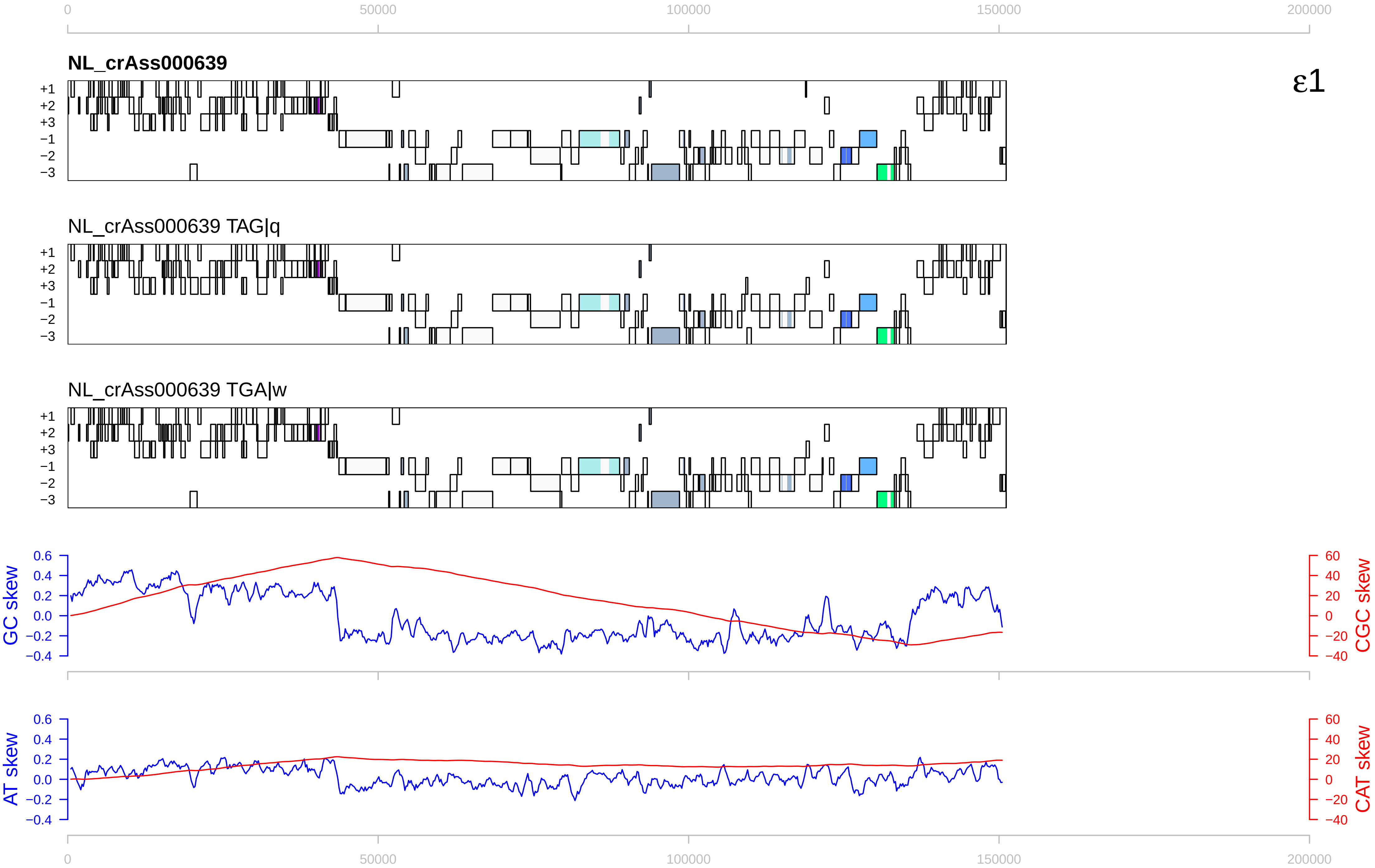

0
50000
100000
150000
200000
NL_crAss000639
ε1
+1
+2
+3
−1
−2
−3
NL_crAss000639 TAG|q
NL_crAss000639 TGA|w
GC skew
CGC skew
AT skew
CAT skew
0.6
0.4
0.2
0.0
−0.2
−0.4
60
40
20
0
−20
−40

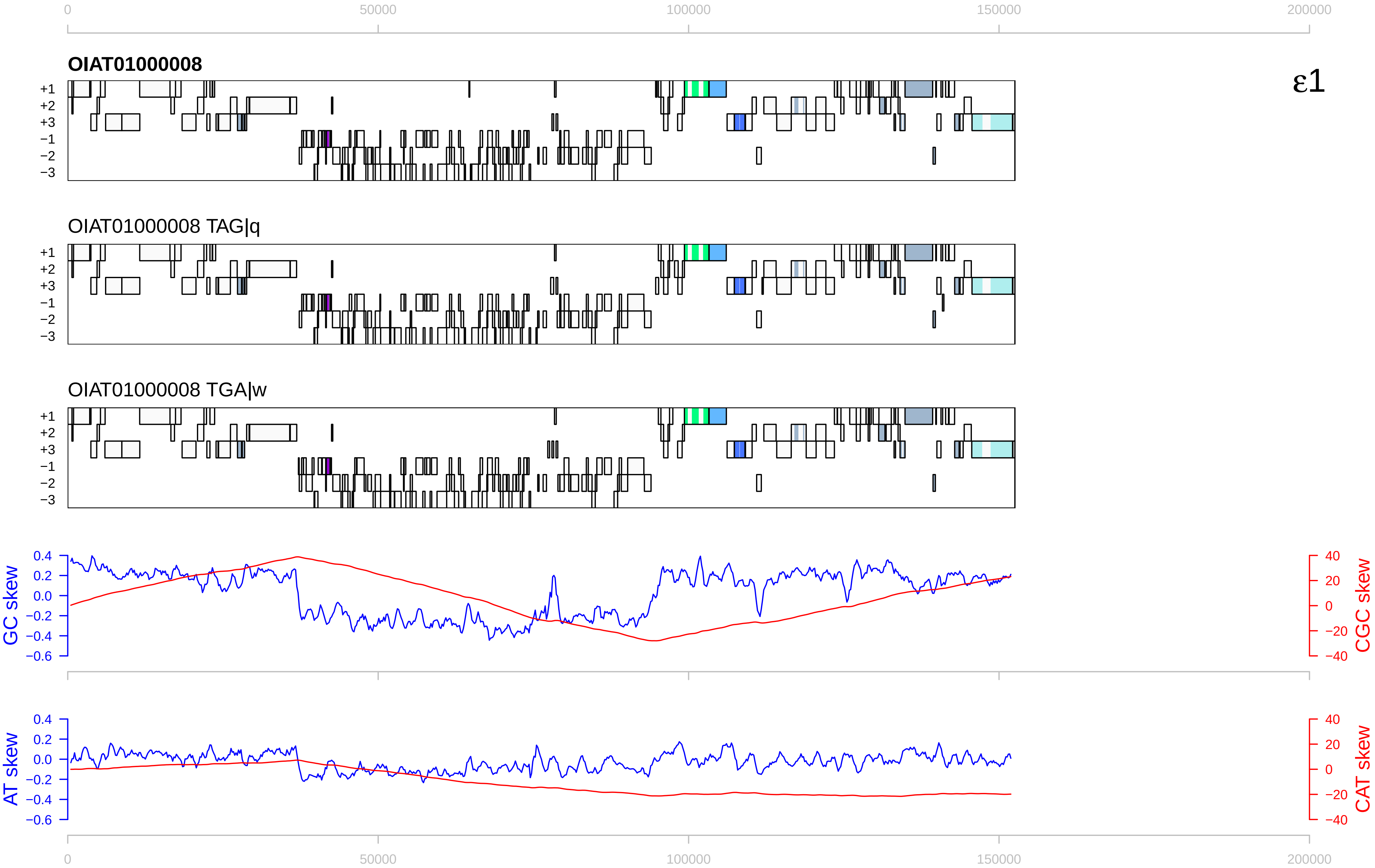

0
50000
100000
150000
200000
OIAT01000008
ε1
+1
+2
+3
−1
−2
−3
OIAT01000008 TAG|q
OIAT01000008 TGA|w
GC skew
CGC skew
AT skew
CAT skew
0.4
0.2
0.0
−0.2
−0.4
−0.6
40
20
0
−20
−40

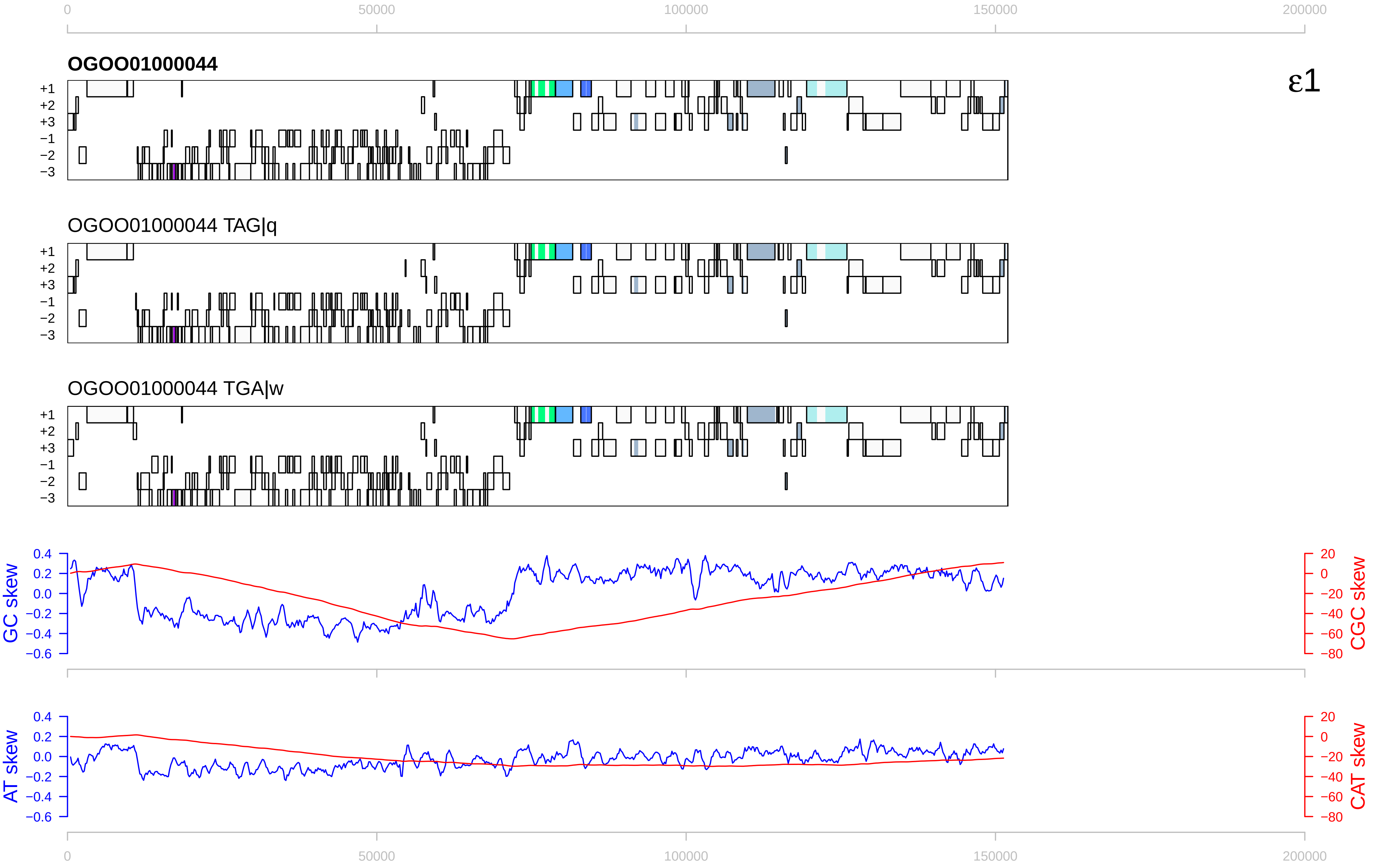

0
50000
100000
150000
200000
OGOO01000044
ε1
+1
+2
+3
−1
−2
−3
OGOO01000044 TAG|q
OGOO01000044 TGA|w
GC skew
CGC skew
AT skew
CAT skew
0.4
0.2
0.0
−0.2
−0.4
−0.6
20
0
−20
−40
−60
−80

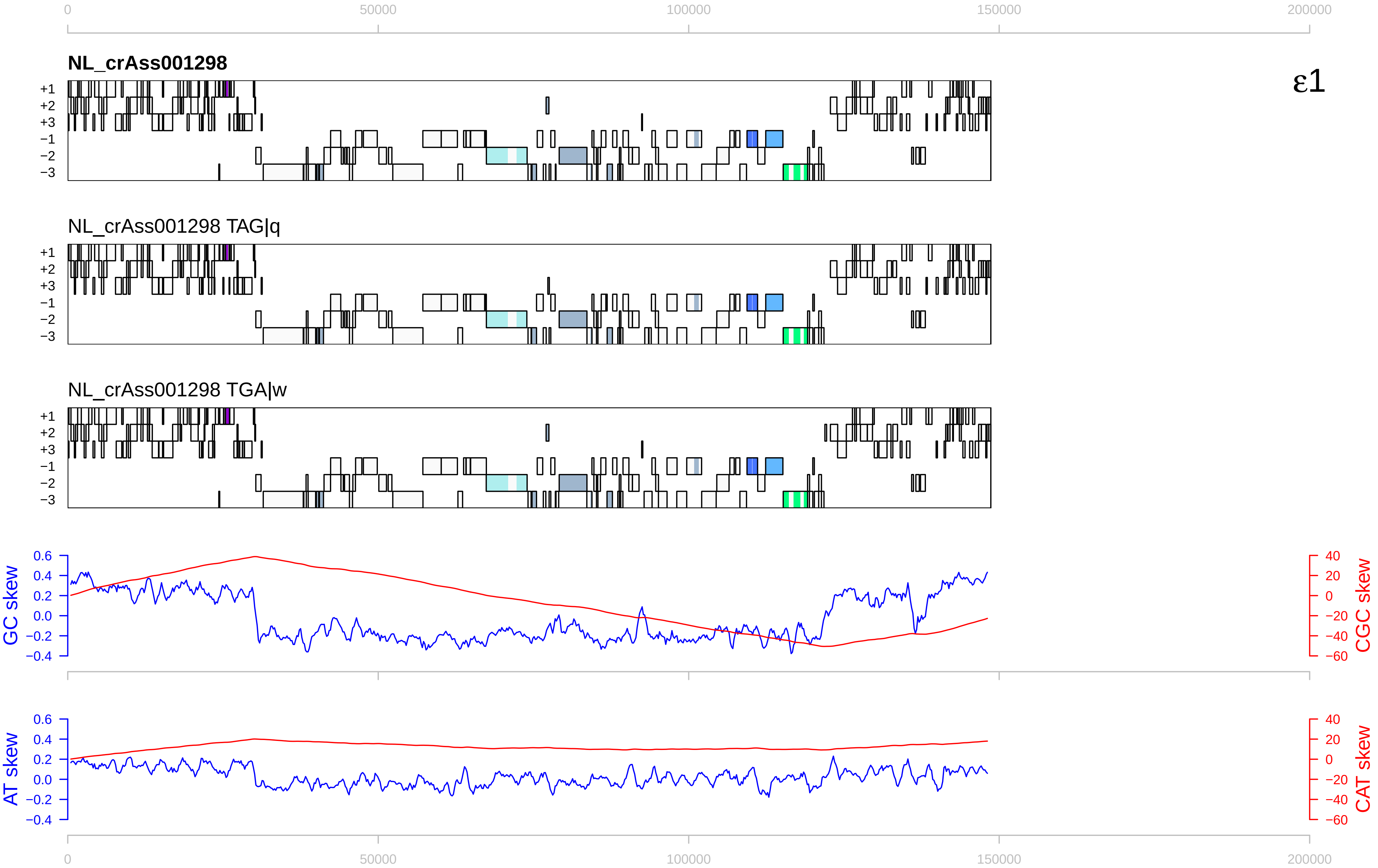

0
50000
100000
150000
200000
NL_crAss001298
ε1
+1
+2
+3
−1
−2
−3
NL_crAss001298 TAG|q
NL_crAss001298 TGA|w
GC skew
CGC skew
AT skew
CAT skew
0.6
0.4
0.2
0.0
−0.2
−0.4
40
20
0
−20
−40
−60

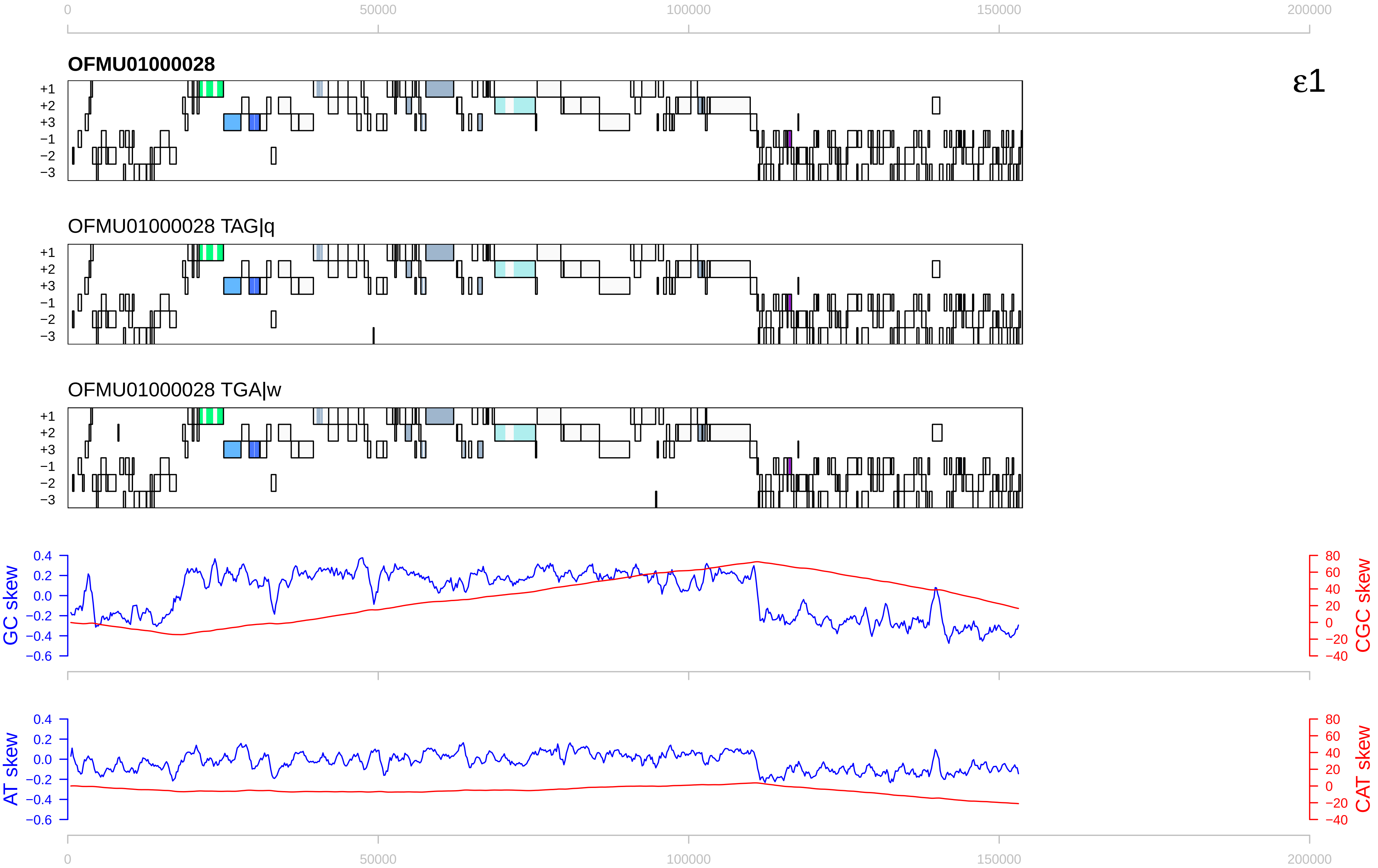

OFMU01000028
ε1
OFMU01000028 TAG|q
OFMU01000028 TGA|w
+1
+2
+3
−1
−2
−3
0
50000
100000
150000
200000
GC skew
CGC skew
AT skew
CAT skew
0.4
0.2
0.0
−0.2
−0.4
−0.6
80
60
40
20
0
−20
−40

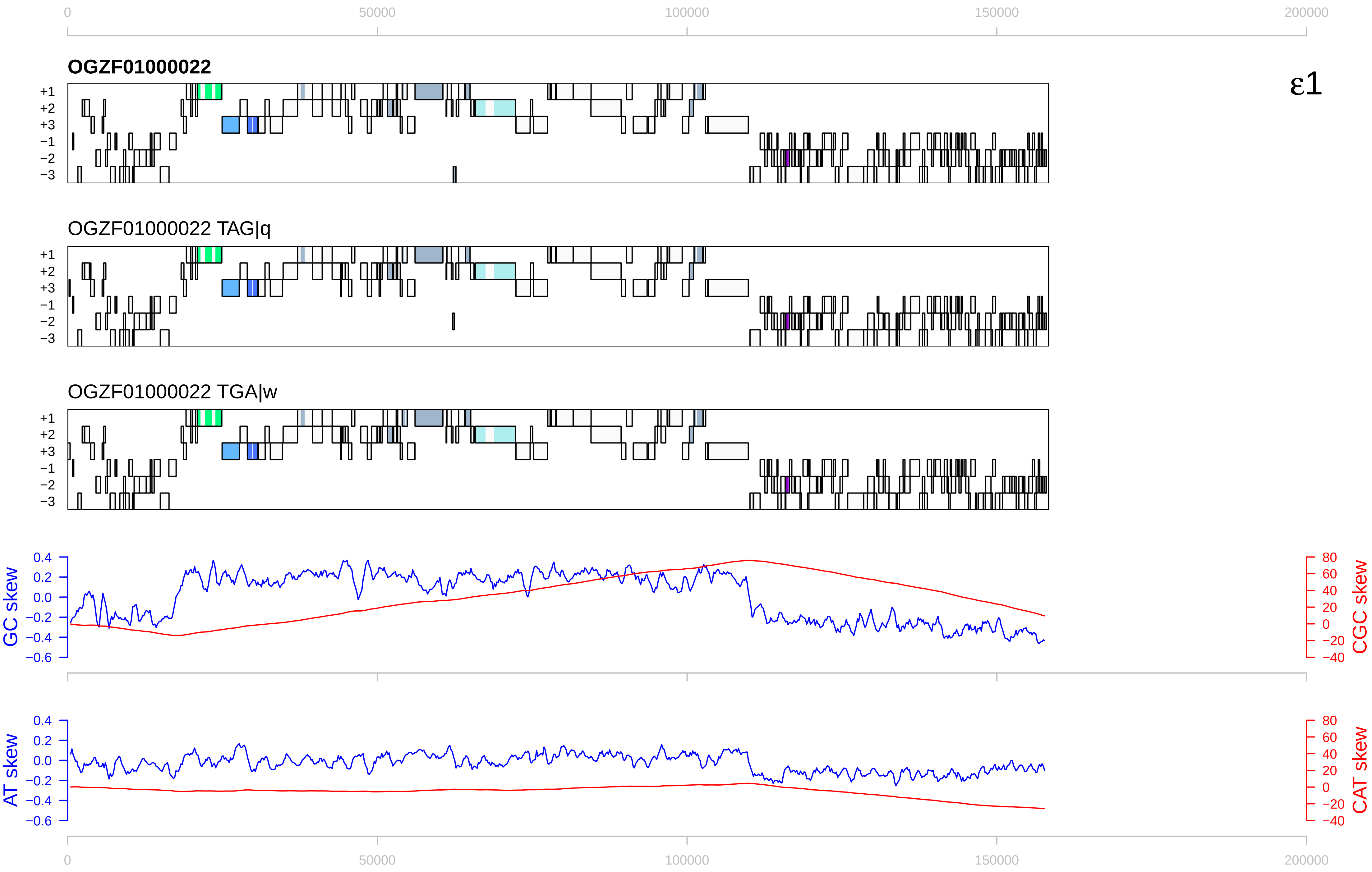

0
50000
100000
150000
200000
OGZF01000022
ε1
+1
+2
+3
−1
−2
−3
OGZF01000022 TAG|q
OGZF01000022 TGA|w
GC skew
CGC skew
AT skew
CAT skew
0.4
0.2
0.0
−0.2
−0.4
−0.6
80
60
40
20
0
−20
−40

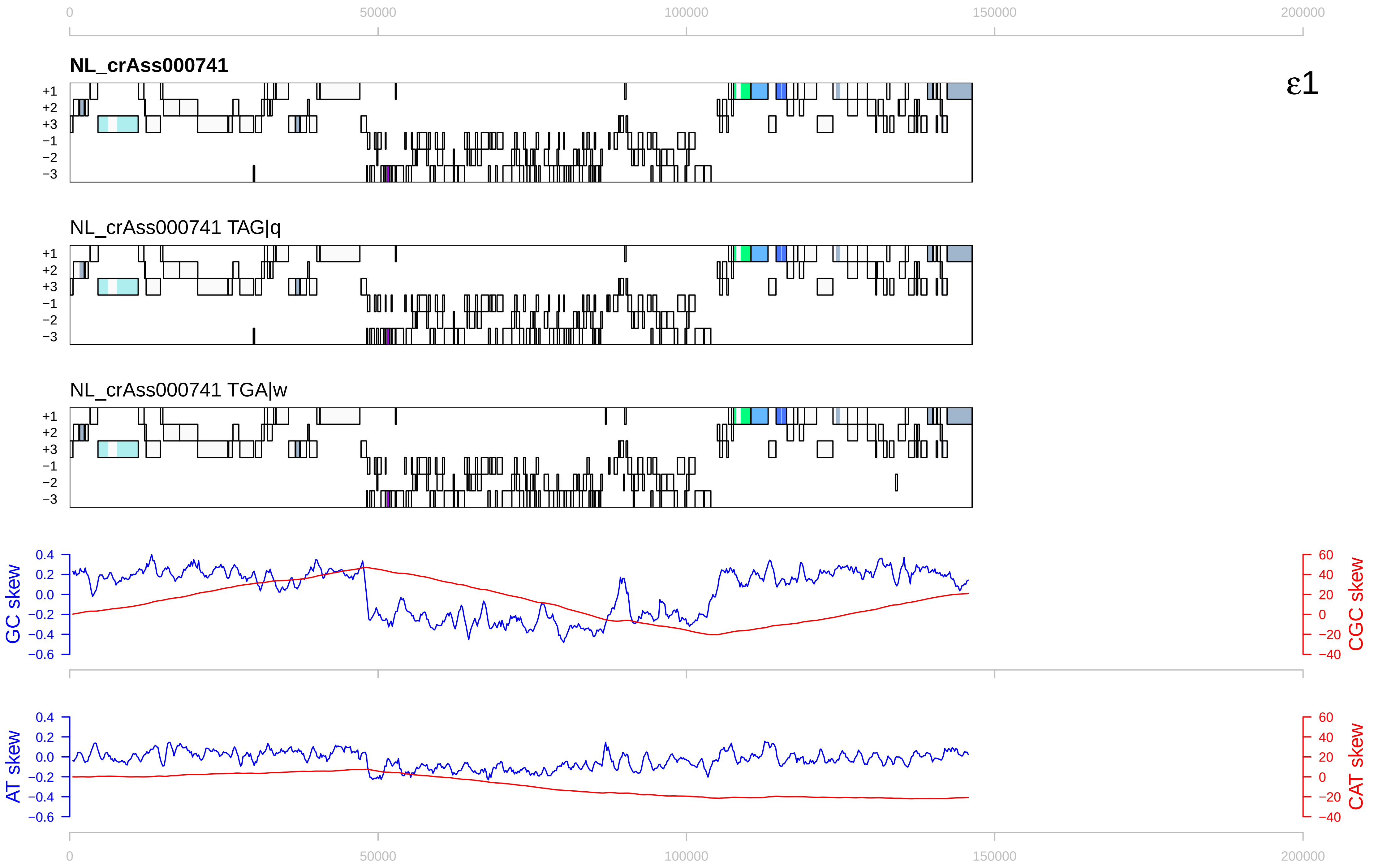

0
50000
100000
150000
200000
NL_crAss000741
ε1
+1
+2
+3
−1
−2
−3
NL_crAss000741 TAG|q
NL_crAss000741 TGA|w
GC skew
CGC skew
AT skew
CAT skew
0.4
0.2
0.0
−0.2
−0.4
−0.6
60
40
20
0
−20
−40

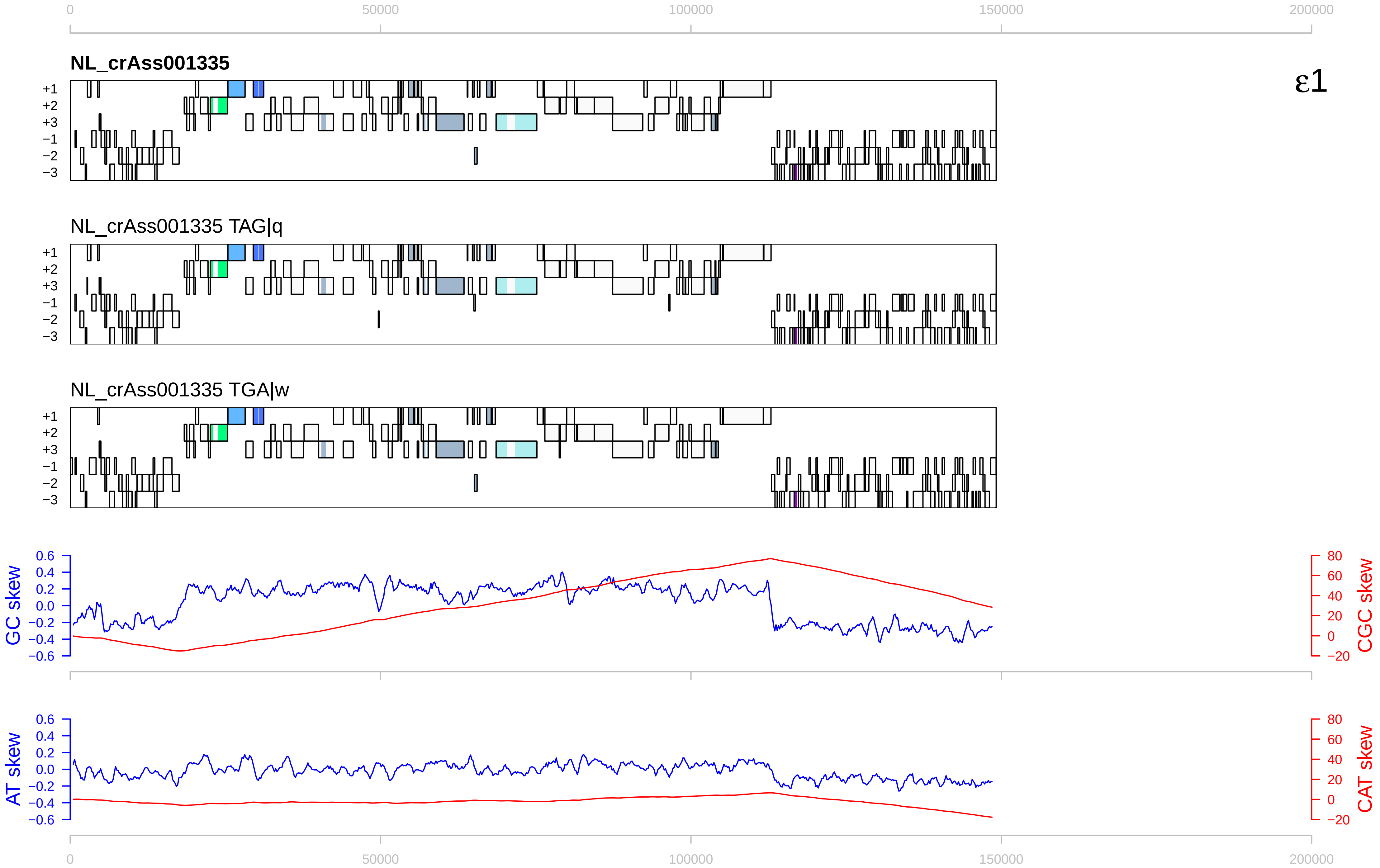

0
50000
100000
150000
200000
NL_crAss001335
ε1
+1
+2
+3
−1
−2
−3
NL_crAss001335 TAG|q
NL_crAss001335 TGA|w
GC skew
CGC skew
AT skew
CAT skew
0.6
0.4
0.2
0.0
−0.2
−0.4
−0.6
80
60
40
20
0
−20

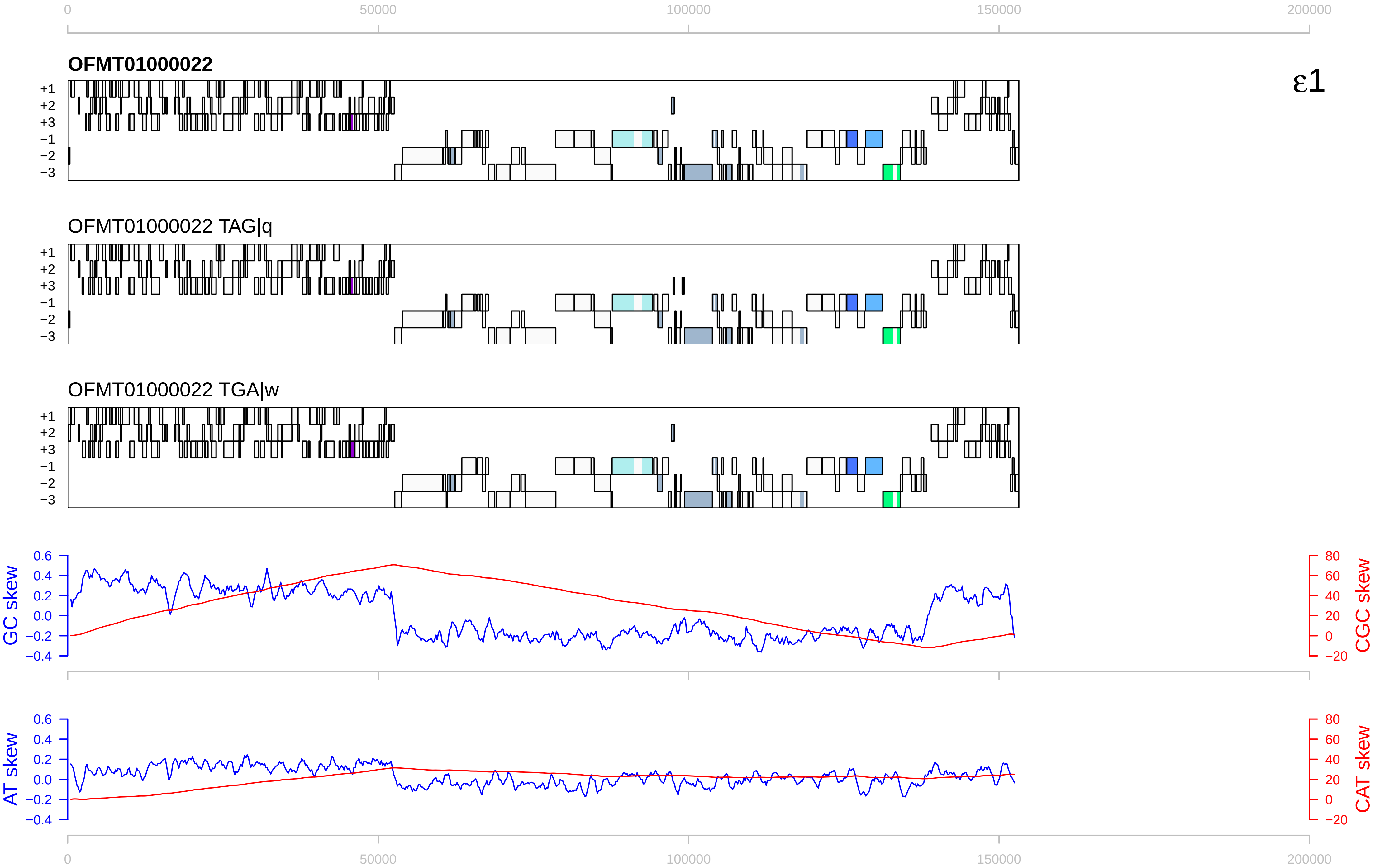

0
50000
100000
150000
200000
OFMT01000022
ε1
+1
+2
+3
−1
−2
−3
OFMT01000022 TAG|q
OFMT01000022 TGA|w
GC skew
CGC skew
AT skew
CAT skew
0.6
0.4
0.2
0.0
−0.2
−0.4
80
60
40
20
0
−20

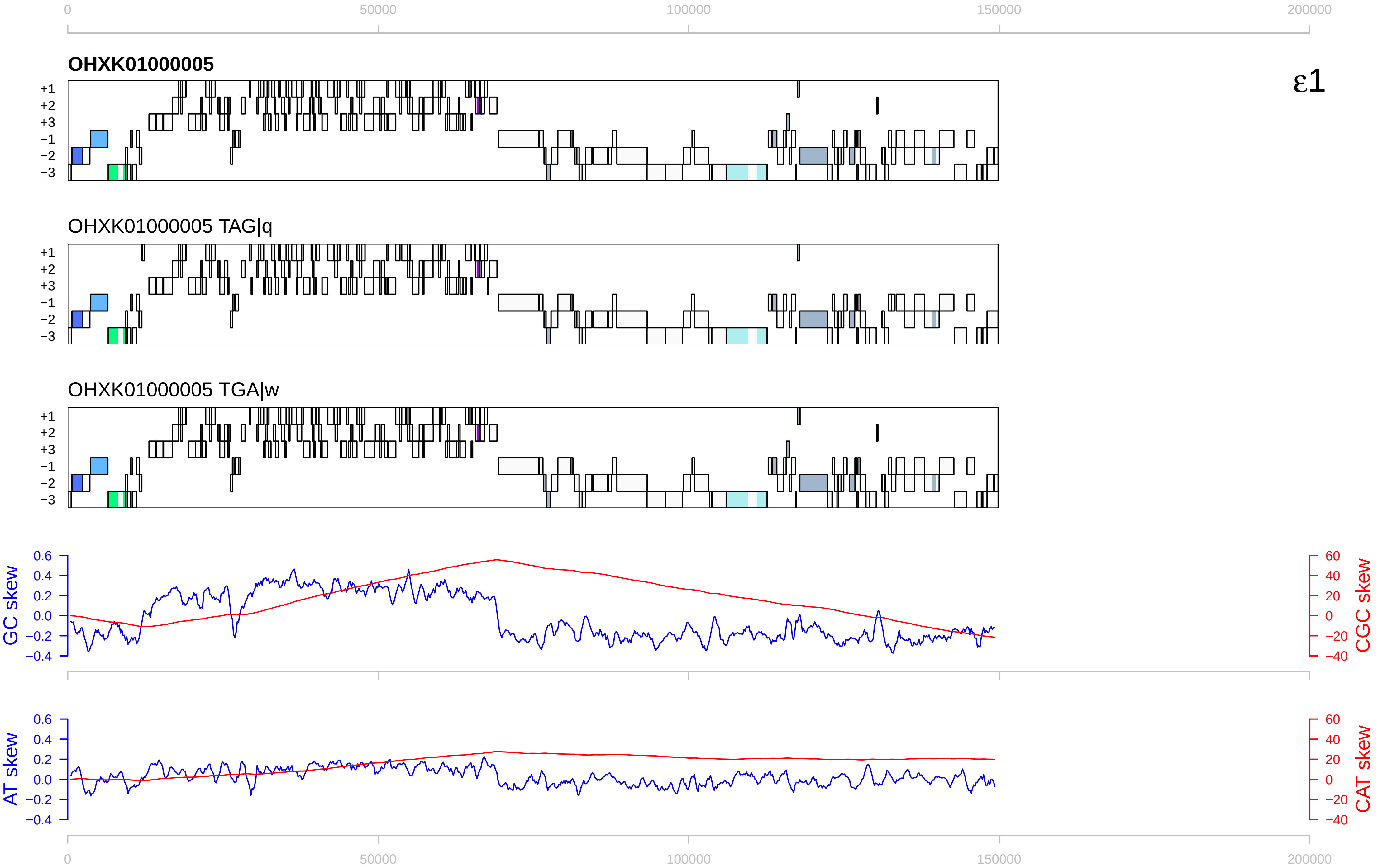

0
50000
100000
150000
200000
OHXK01000005
ε1
+1
+2
+3
−1
−2
−3
OHXK01000005 TAG|q
OHXK01000005 TGA|w
GC skew
CGC skew
AT skew
CAT skew
0.6
0.4
0.2
0.0
−0.2
−0.4
60
40
20
0
−20
−40

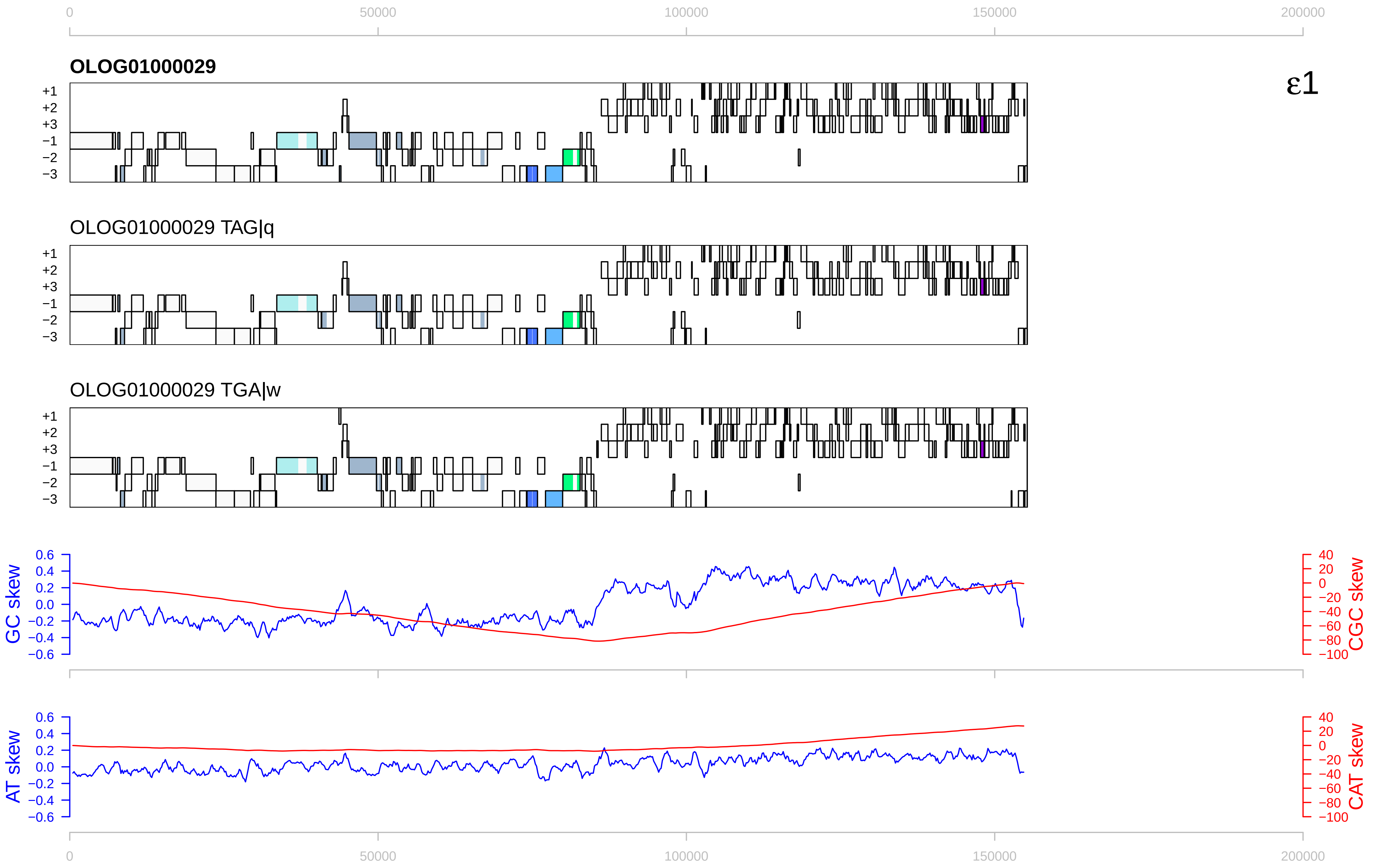

0
50000
100000
150000
200000
OLOG01000029
ε1
+1
+2
+3
−1
−2
−3
OLOG01000029 TAG|q
OLOG01000029 TGA|w
GC skew
CGC skew
AT skew
CAT skew
0.6
0.4
0.2
0.0
−0.2
−0.4
−0.6
40
20
0
−20
−40
−60
−80
−100

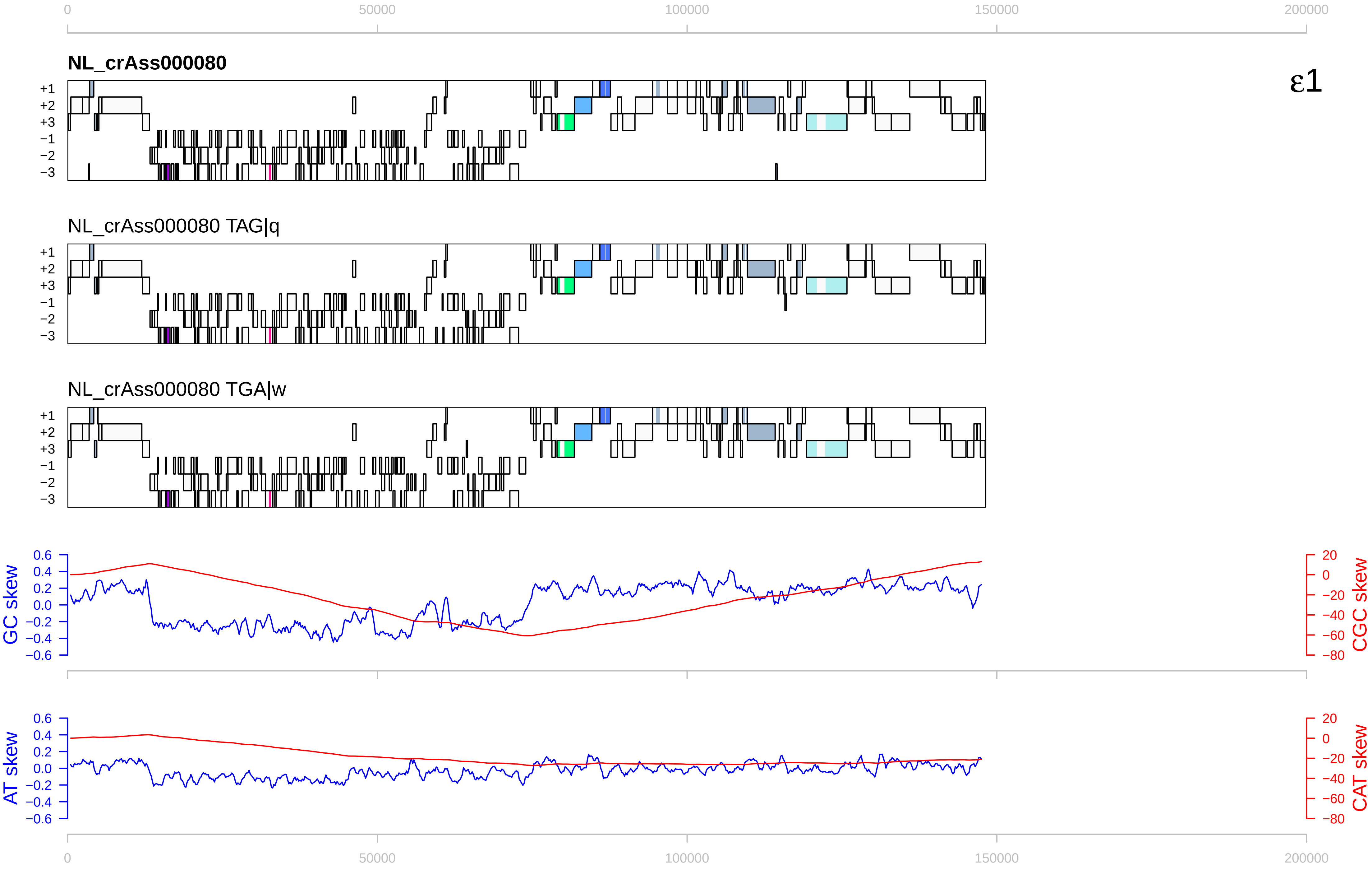

0
50000
100000
150000
200000
NL_crAss000080
ε1
+1
+2
+3
−1
−2
−3
NL_crAss000080 TAG|q
NL_crAss000080 TGA|w
GC skew
CGC skew
AT skew
CAT skew
0.6
0.4
0.2
0.0
−0.2
−0.4
−0.6
20
0
−20
−40
−60
−80

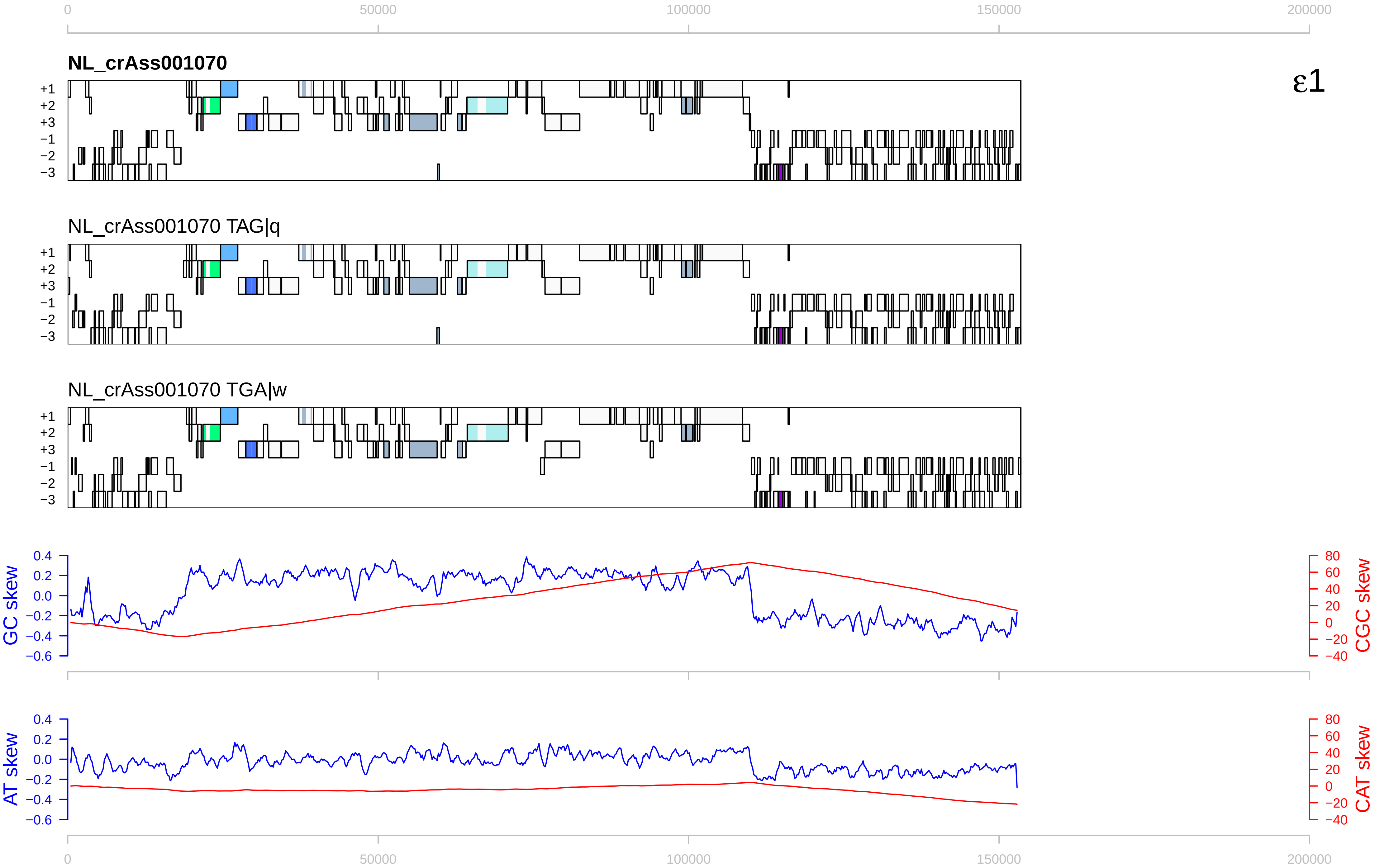

NL_crAss001070
ε1
NL_crAss001070 TAG|q
NL_crAss001070 TGA|w
+1
+2
+3
−1
−2
−3
GC skew
CGC skew
AT skew
CAT skew
0
50000
100000
150000
200000

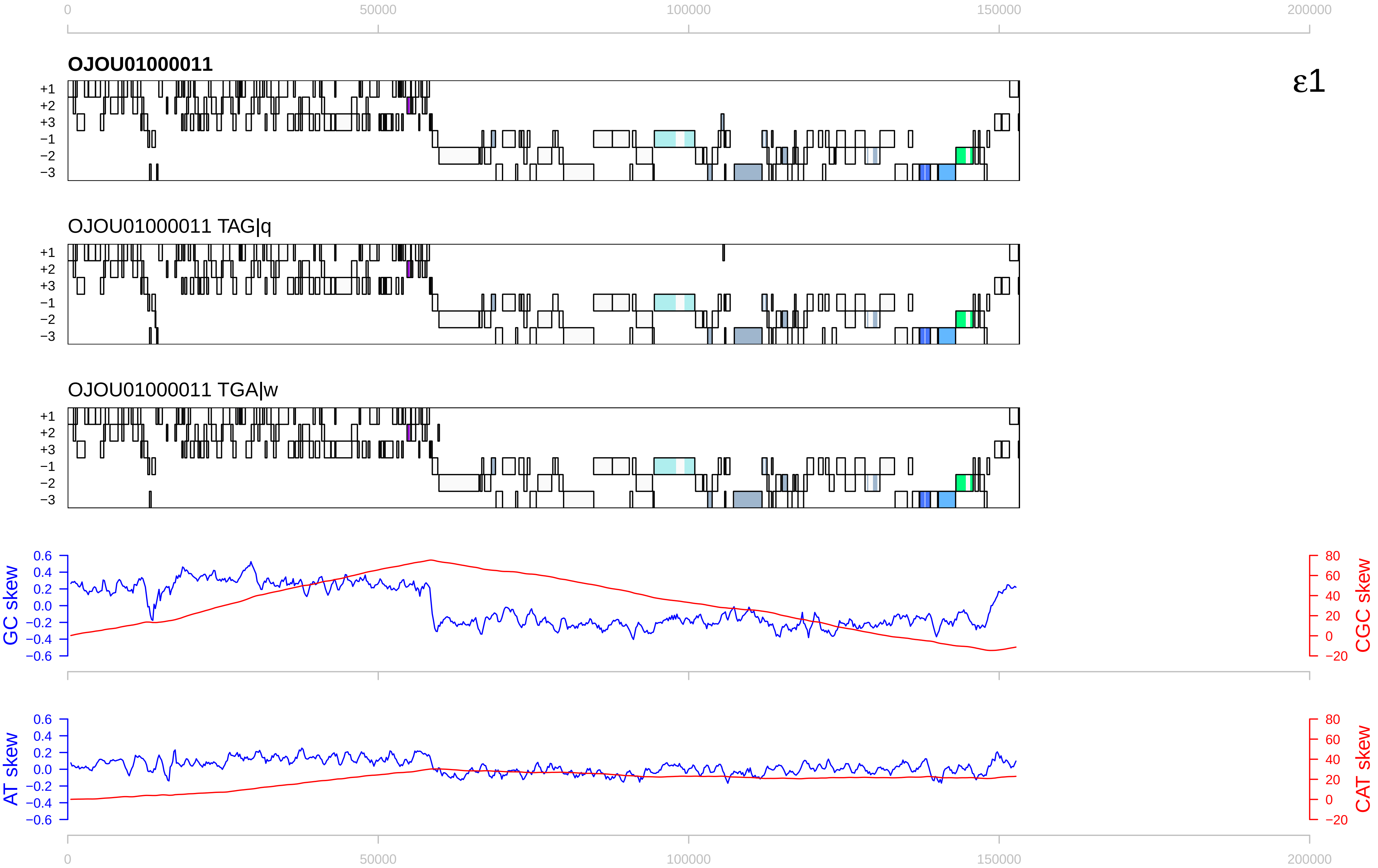

0
50000
100000
150000
200000
OJOU01000011
ε1
+1
+2
+3
−1
−2
−3
OJOU01000011 TAG|q
OJOU01000011 TGA|w
GC skew
CGC skew
AT skew
CAT skew
0.6
0.4
0.2
0.0
−0.2
−0.4
−0.6
80
60
40
20
0
−20

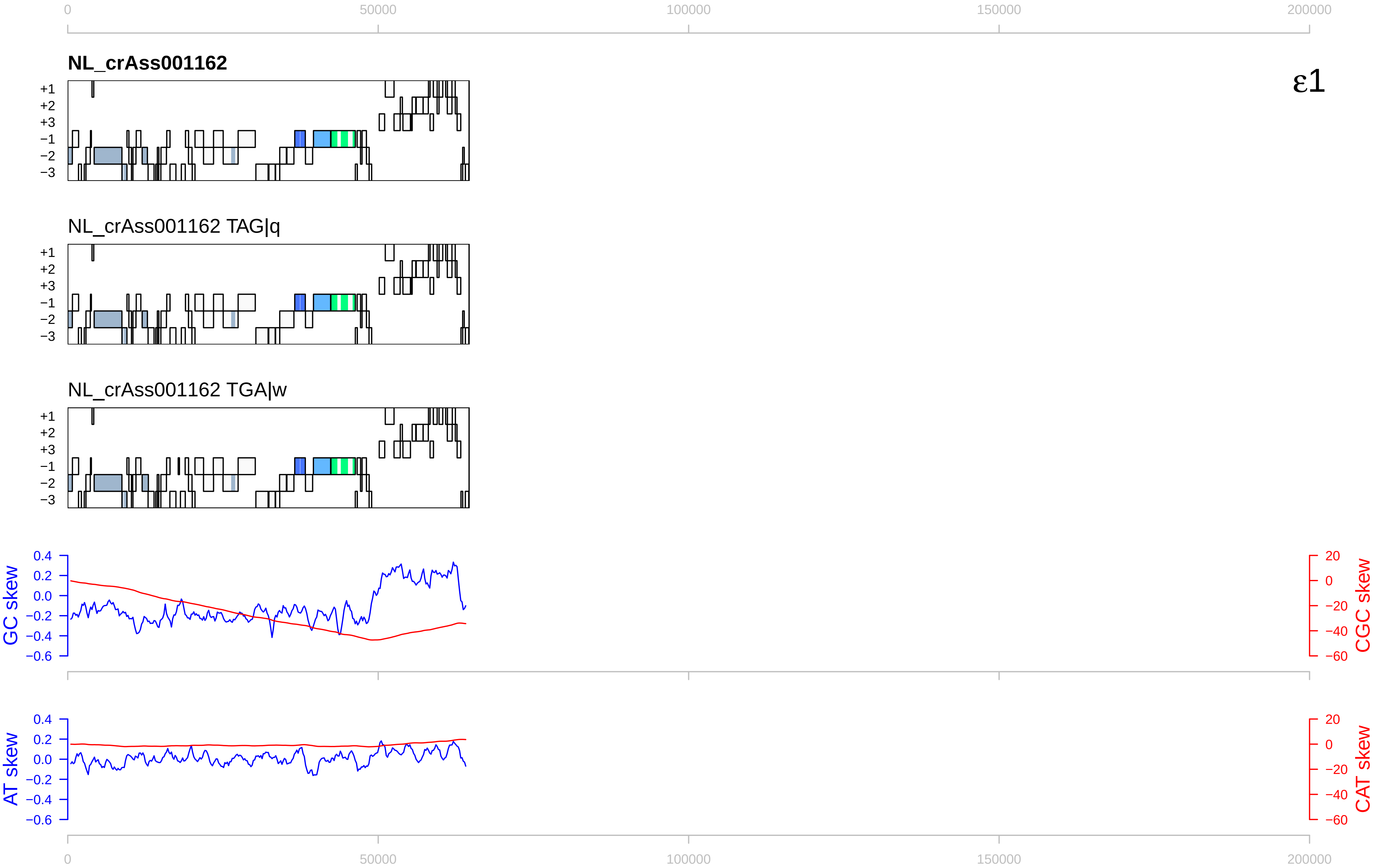

0
50000
100000
150000
200000
NL_crAss001162
ε1
+1
+2
+3
−1
−2
−3
NL_crAss001162 TAG|q
NL_crAss001162 TGA|w
GC skew
CGC skew
0.4
0.2
0.0
−0.2
−0.4
−0.6
20
0
−20
−40
−60
AT skew
CAT skew

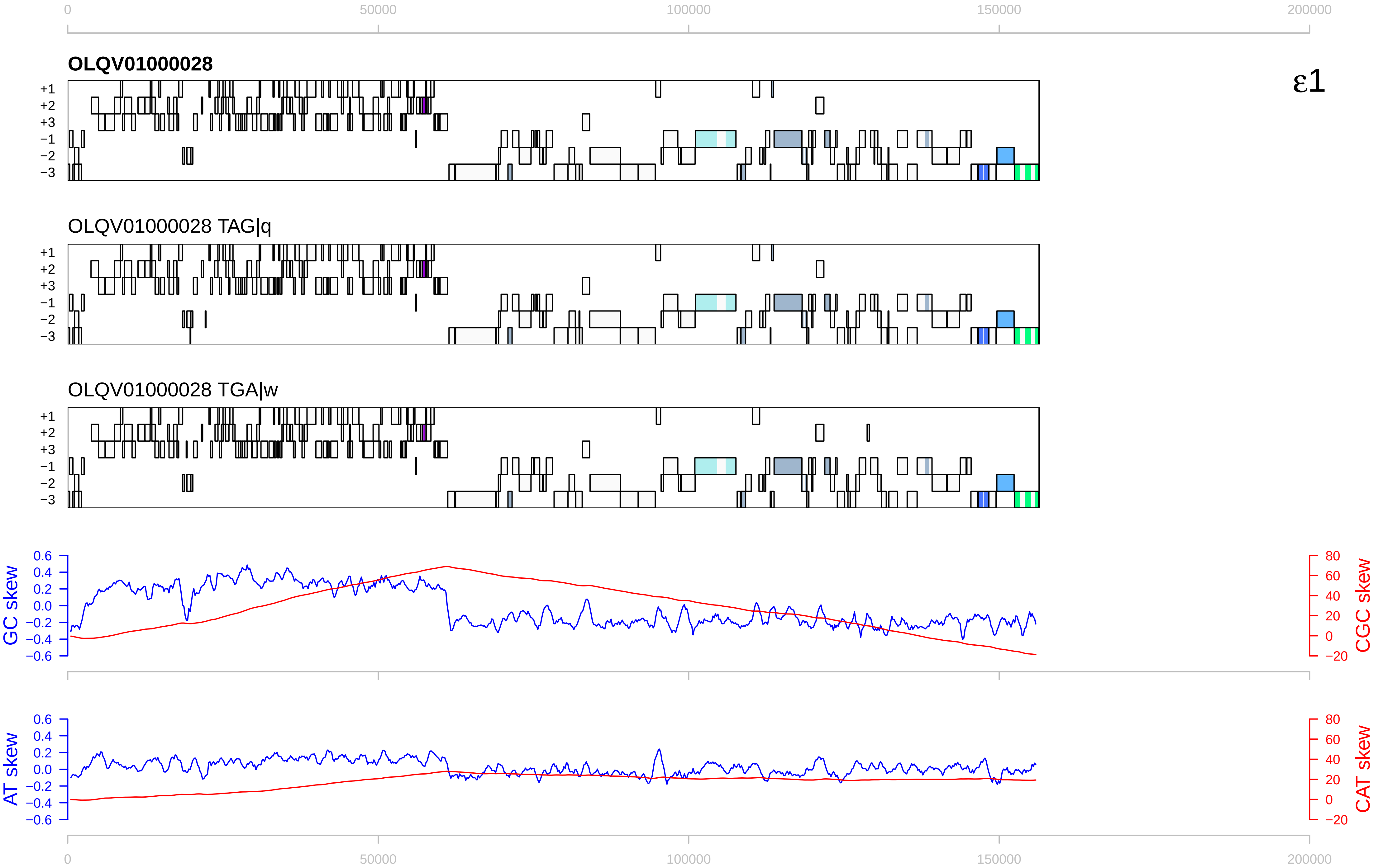
ε1
OLQV01000028
OLQV01000028 TAG|q
OLQV01000028 TGA|w
+1
+2
+3
−1
−2
−3
GC skew
CGC skew
AT skew
CAT skew
0
50000
100000
150000
200000

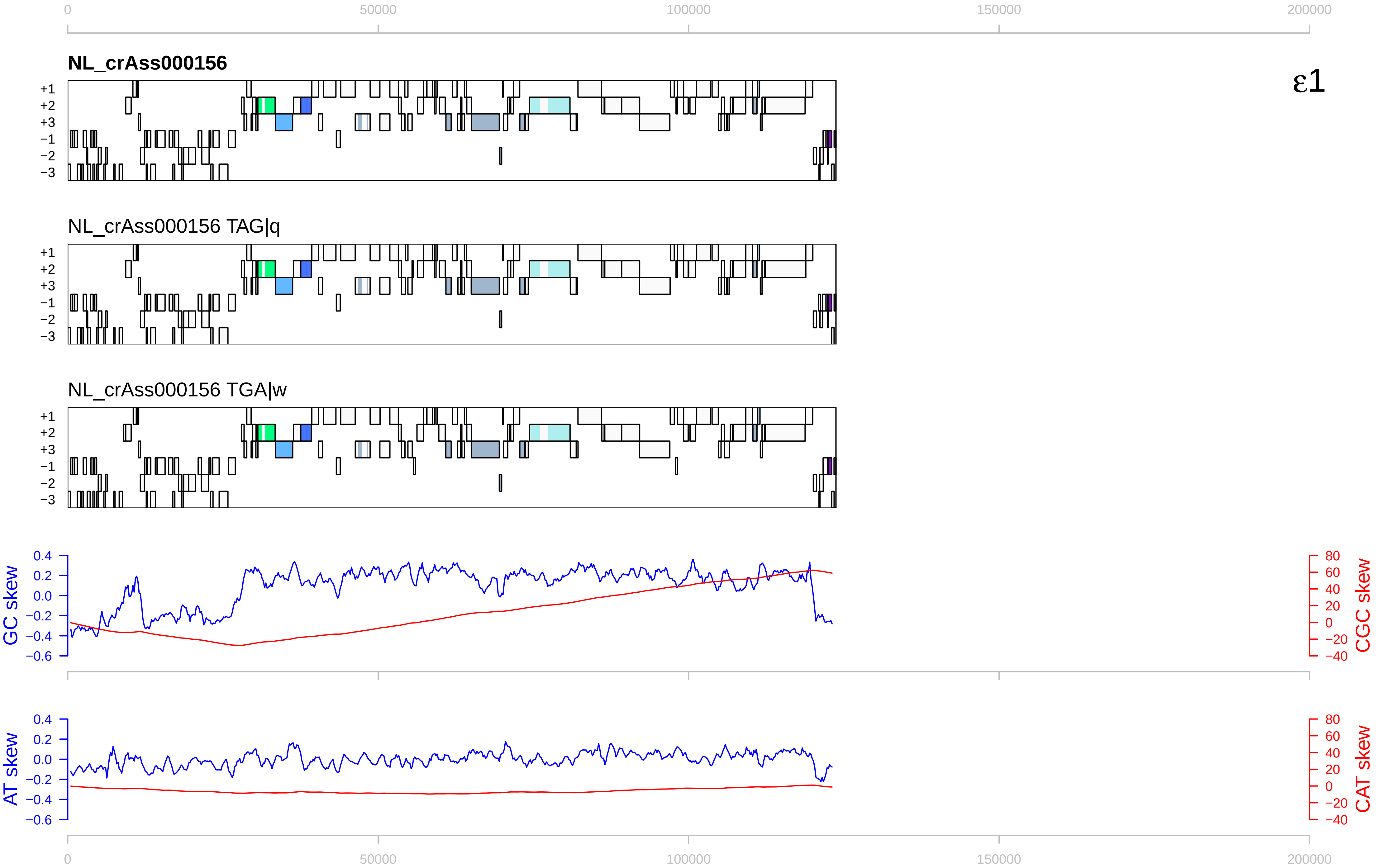

ε1
NL_crAss000156
NL_crAss000156 TAG|q
NL_crAss000156 TGA|w
+1
+2
+3
−1
−2
−3
GC skew
CGC skew
AT skew
CAT skew
0
50000
100000
150000
200000

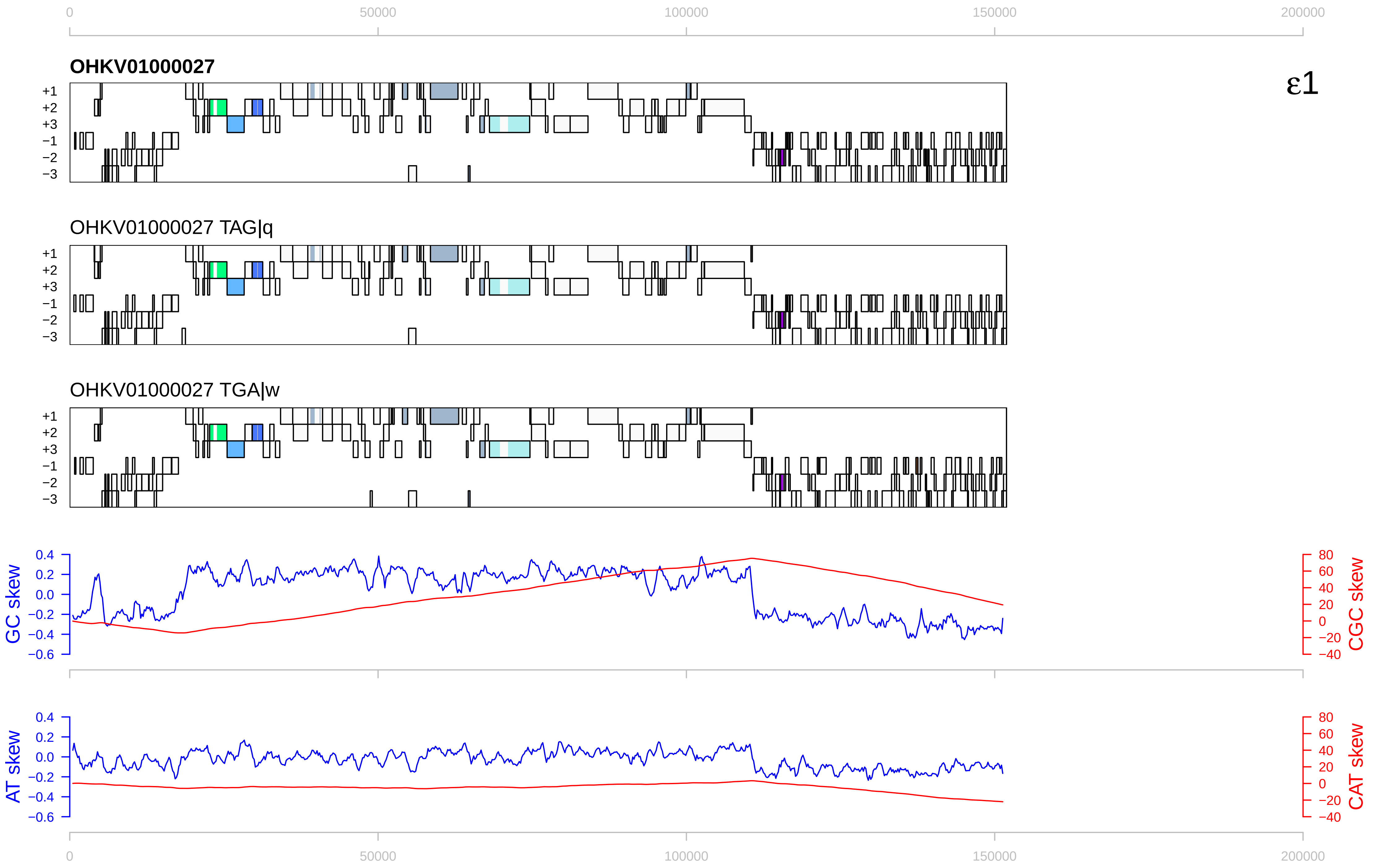

ε1
OHKV01000027
OHKV01000027 TAG|q
OHKV01000027 TGA|w
+1
+2
+3
−1
−2
−3
0
50000
100000
150000
200000
GC skew
CGC skew
AT skew
CAT skew
0.4
0.2
0.0
−0.2
−0.4
−0.6
80
60
40
20
0
−20
−40

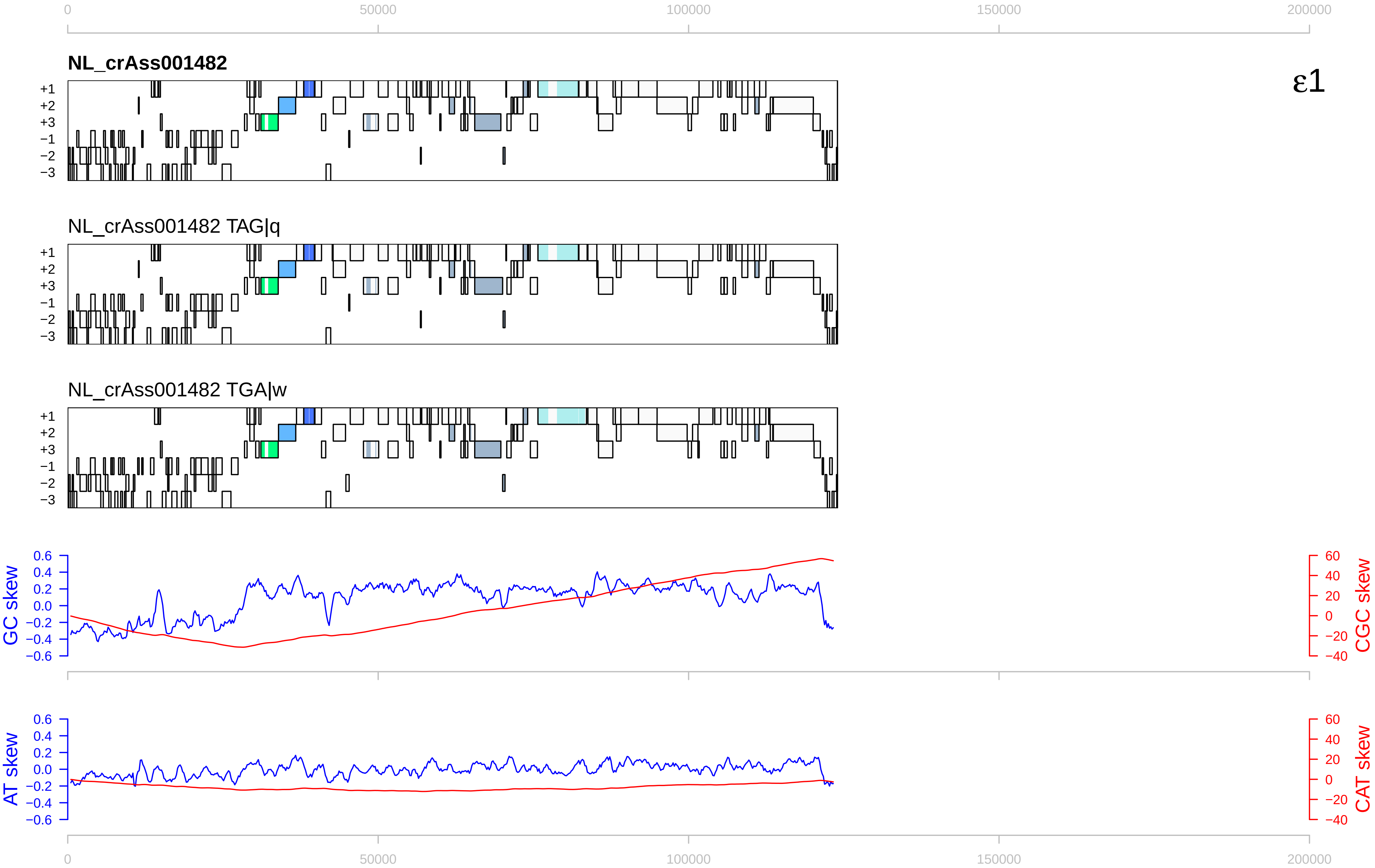

0
50000
100000
150000
200000
NL_crAss001482
ε1
+1
+2
+3
−1
−2
−3
NL_crAss001482 TAG|q
NL_crAss001482 TGA|w
GC skew
CGC skew
AT skew
CAT skew
0.6
0.4
0.2
0.0
−0.2
−0.4
−0.6
60
40
20
0
−20
−40

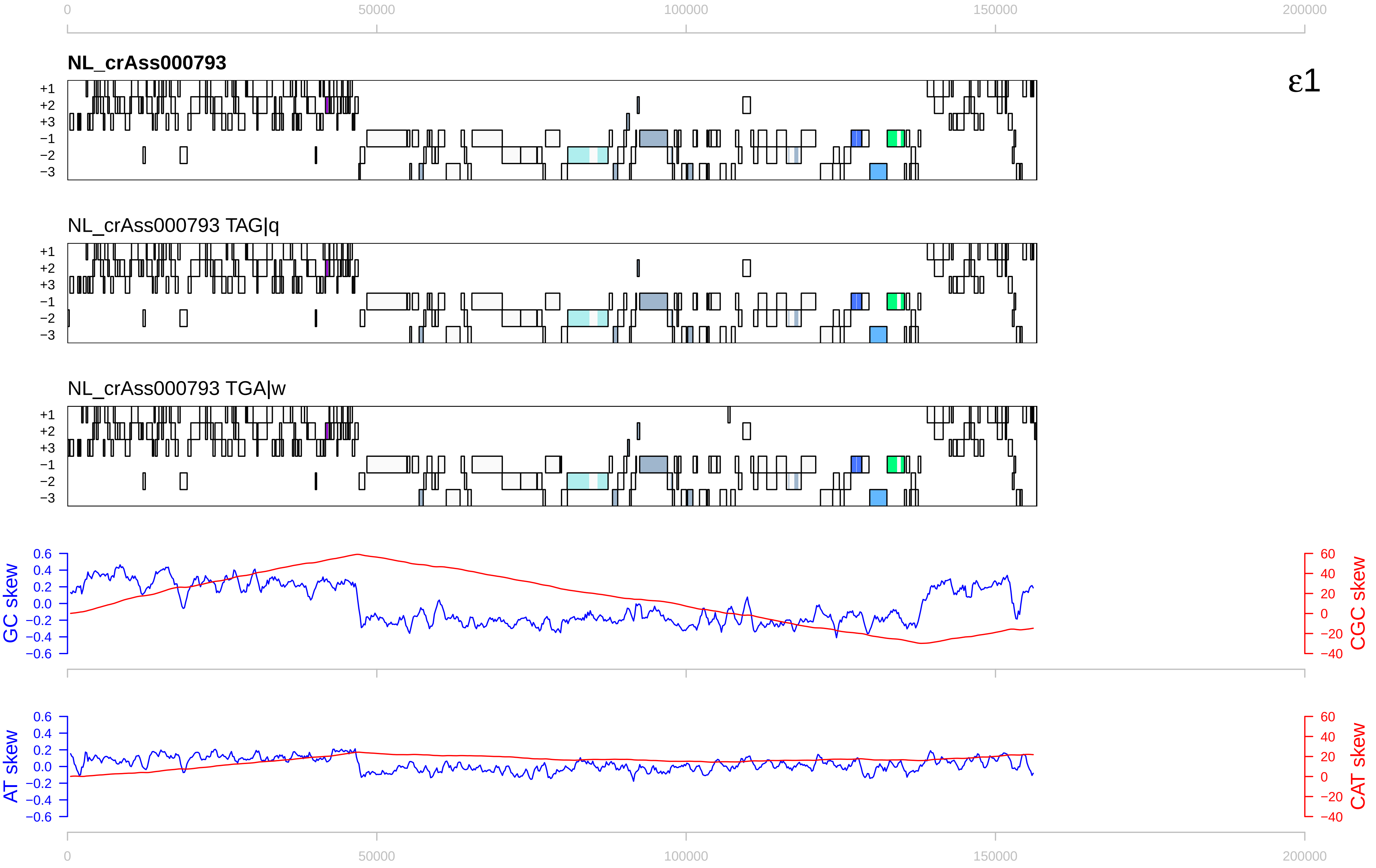

ε1
NL_crAss000793
NL_crAss000793 TAG|q
NL_crAss000793 TGA|w
+1
+2
+3
−1
−2
−3
0
50000
100000
150000
200000
GC skew
CGC skew
AT skew
CAT skew

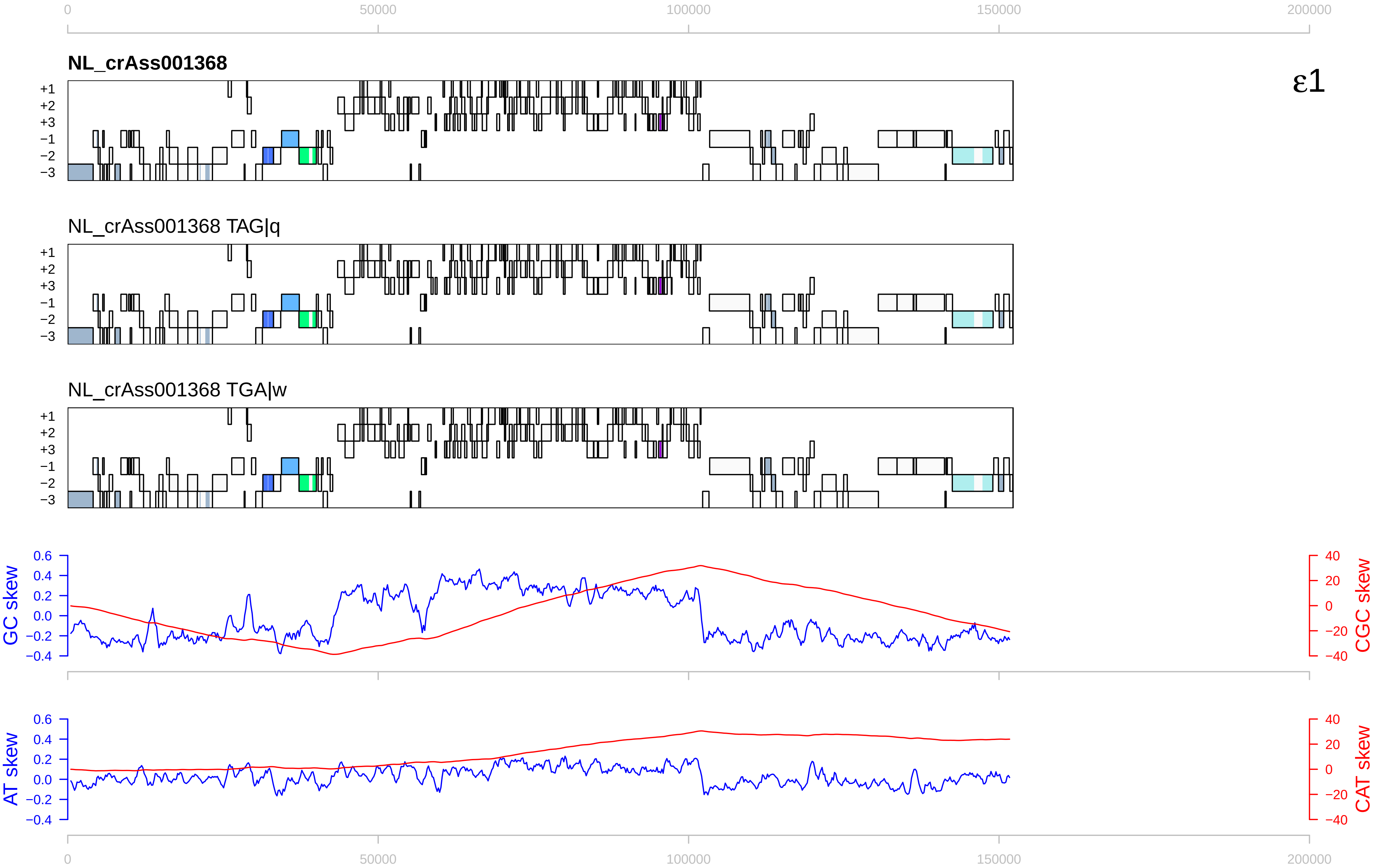

NL_crAss001368
ε1
NL_crAss001368 TAG|q
NL_crAss001368 TGA|w
+1
+2
+3
−1
−2
−3
GC skew
CGC skew
AT skew
CAT skew
0
50000
100000
150000
200000

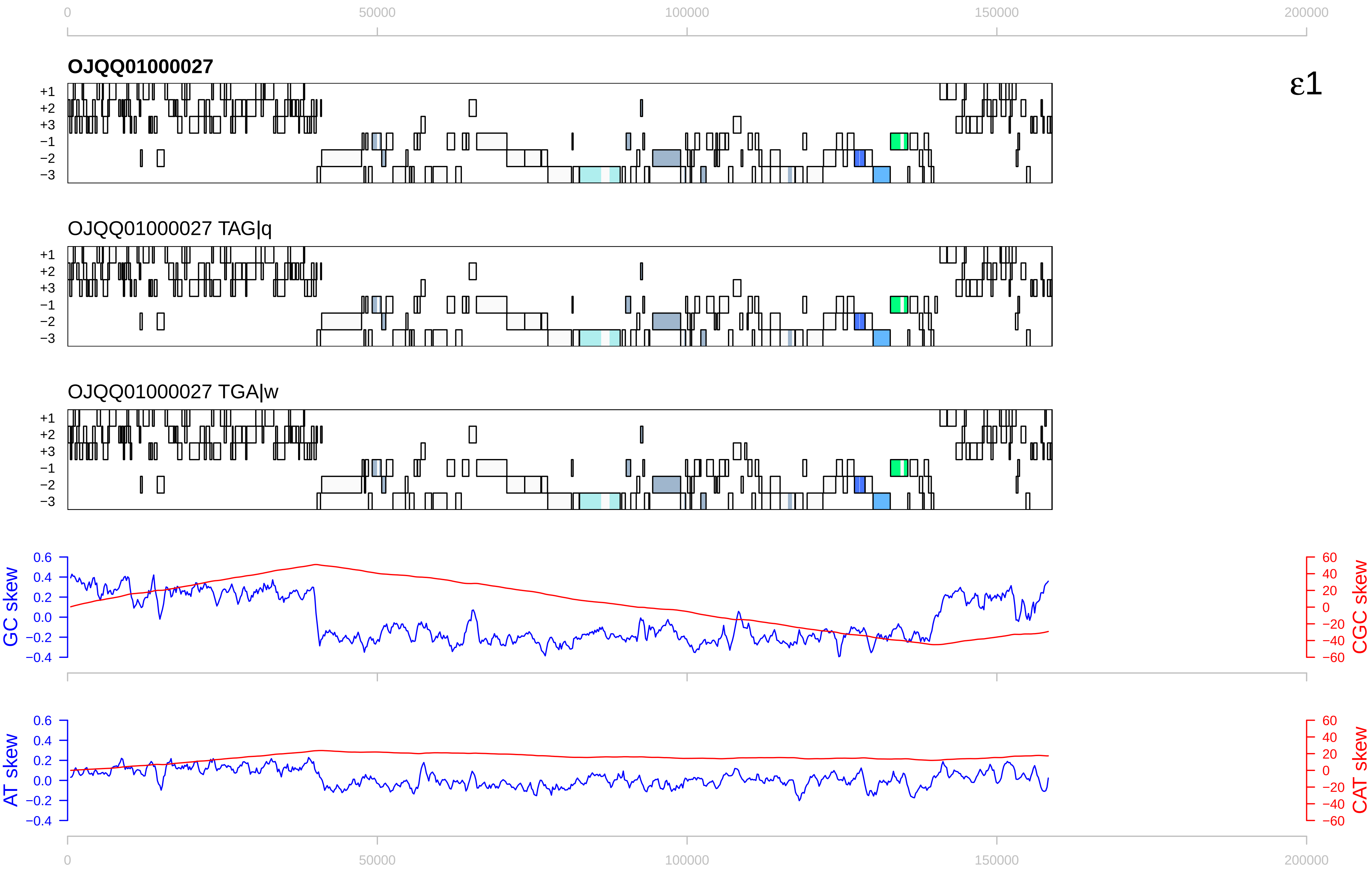

0
50000
100000
150000
200000
OJQQ01000027
ε1
+1
+2
+3
−1
−2
−3
OJQQ01000027 TAG|q
OJQQ01000027 TGA|w
GC skew
CGC skew
AT skew
CAT skew
0.6
0.4
0.2
0.0
−0.2
−0.4
60
40
20
0
−20
−40
−60

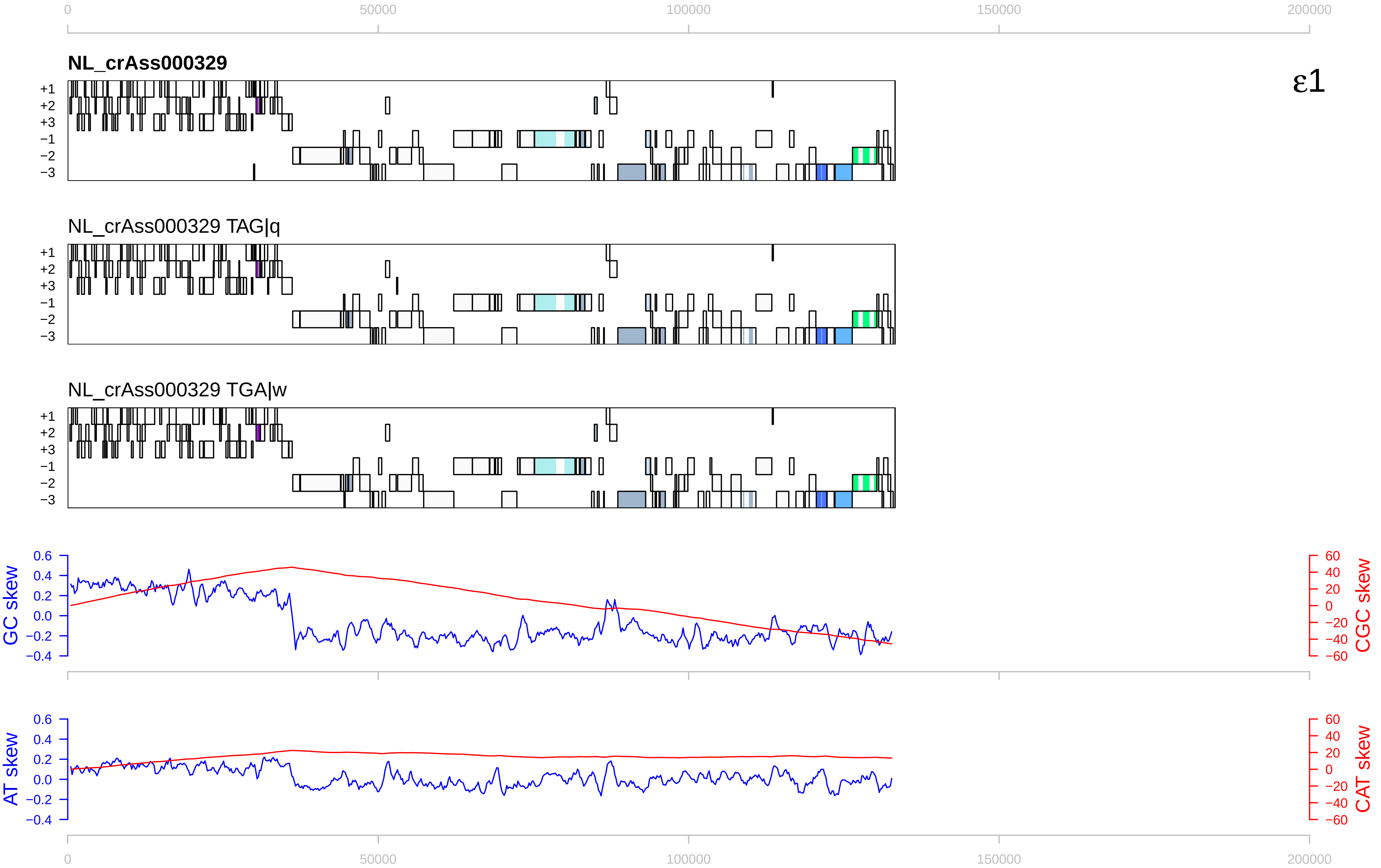

0
50000
100000
150000
200000
NL_crAss000329
ε1
+1
+2
+3
−1
−2
−3
NL_crAss000329 TAG|q
NL_crAss000329 TGA|w
GC skew
CGC skew
AT skew
CAT skew
0.6
0.4
0.2
0.0
−0.2
−0.4
60
40
20
0
−20
−40
−60

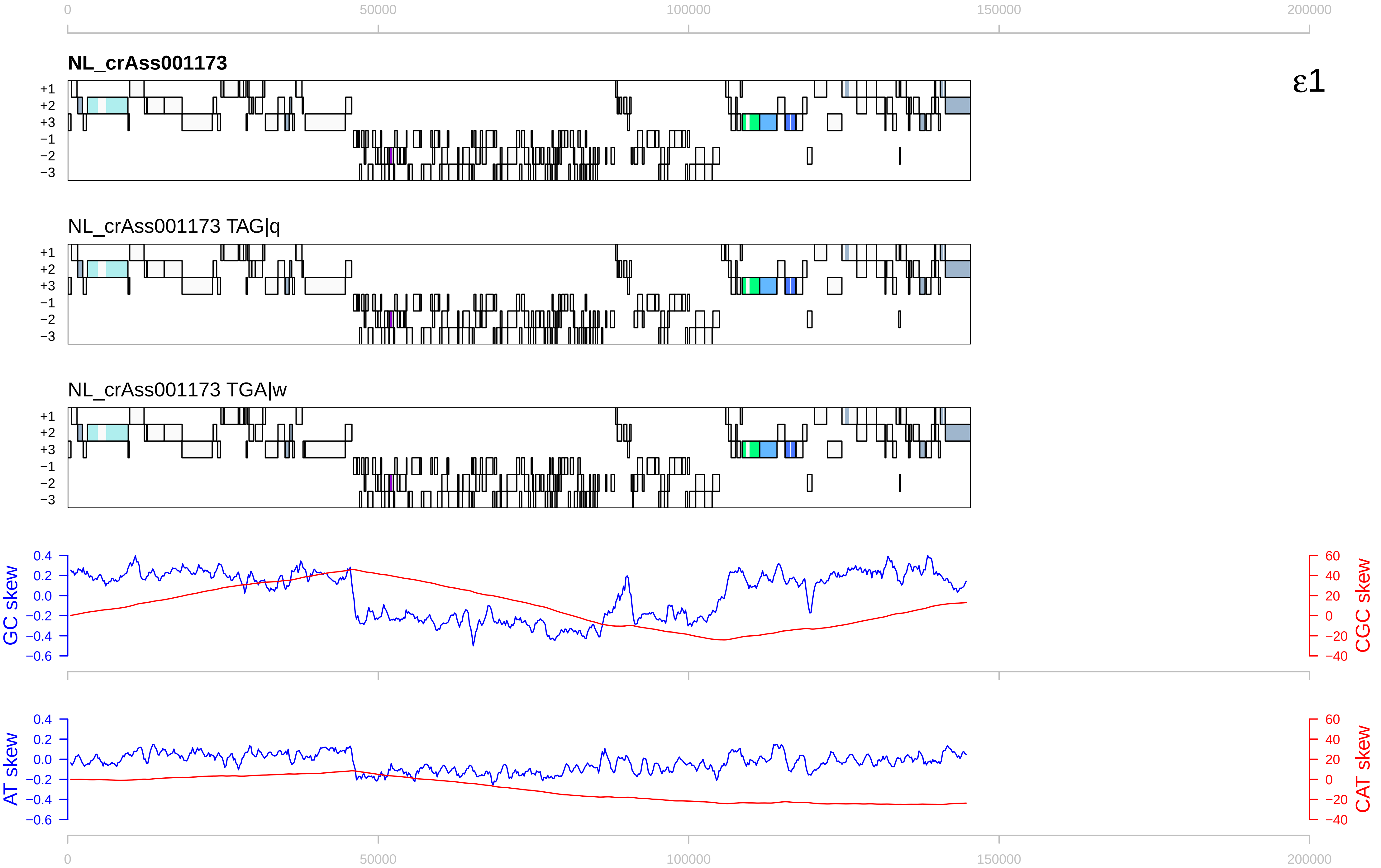

ε1
NL_crAss001173
NL_crAss001173 TAG|q
NL_crAss001173 TGA|w
+1
+2
+3
−1
−2
−3
0
50000
100000
150000
200000
GC skew
CGC skew
AT skew
CAT skew
0.4
0.2
0.0
−0.2
−0.4
−0.6
60
40
20
0
−20
−40

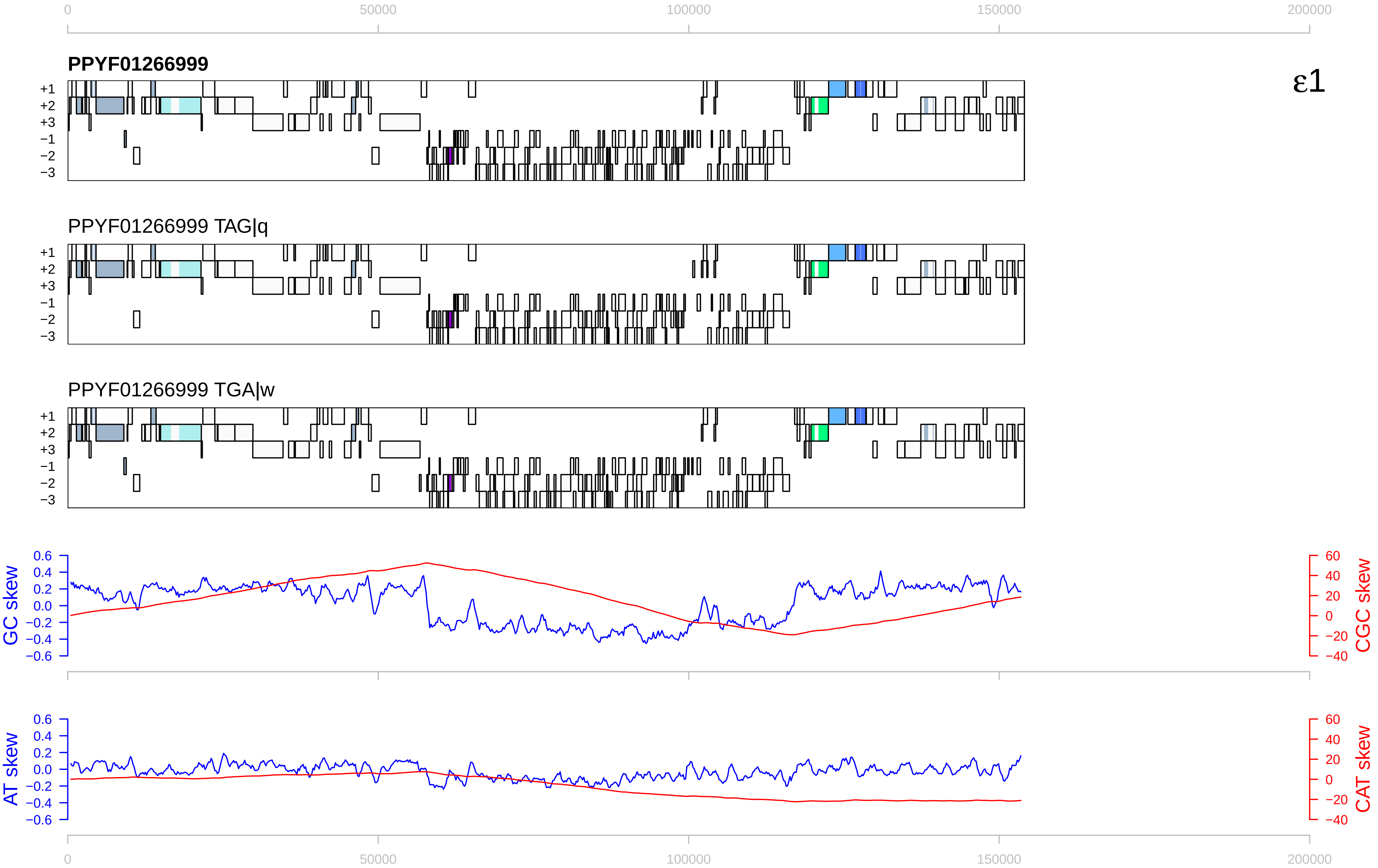

0
50000
100000
150000
200000
PPYF01266999
ε1
+1
+2
+3
−1
−2
−3
PPYF01266999 TAG|q
PPYF01266999 TGA|w
GC skew
CGC skew
AT skew
CAT skew
0.6
0.4
0.2
0.0
−0.2
−0.4
−0.6
60
40
20
0
−20
−40

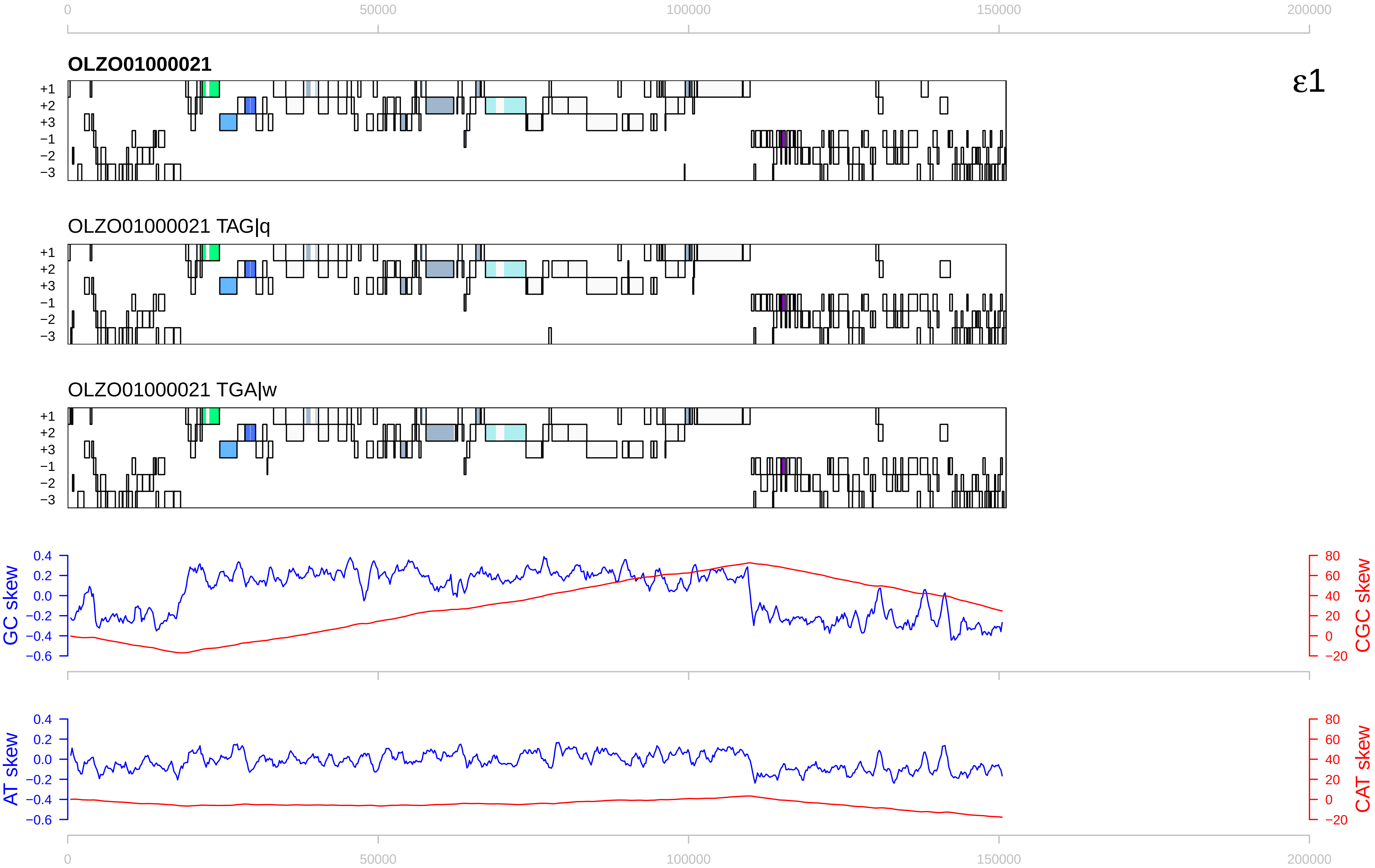

ε1
OLZO01000021
OLZO01000021 TAG|q
OLZO01000021 TGA|w
+1
+2
+3
−1
−2
−3
GC skew
CGC skew
AT skew
CAT skew
0
50000
100000
150000
200000

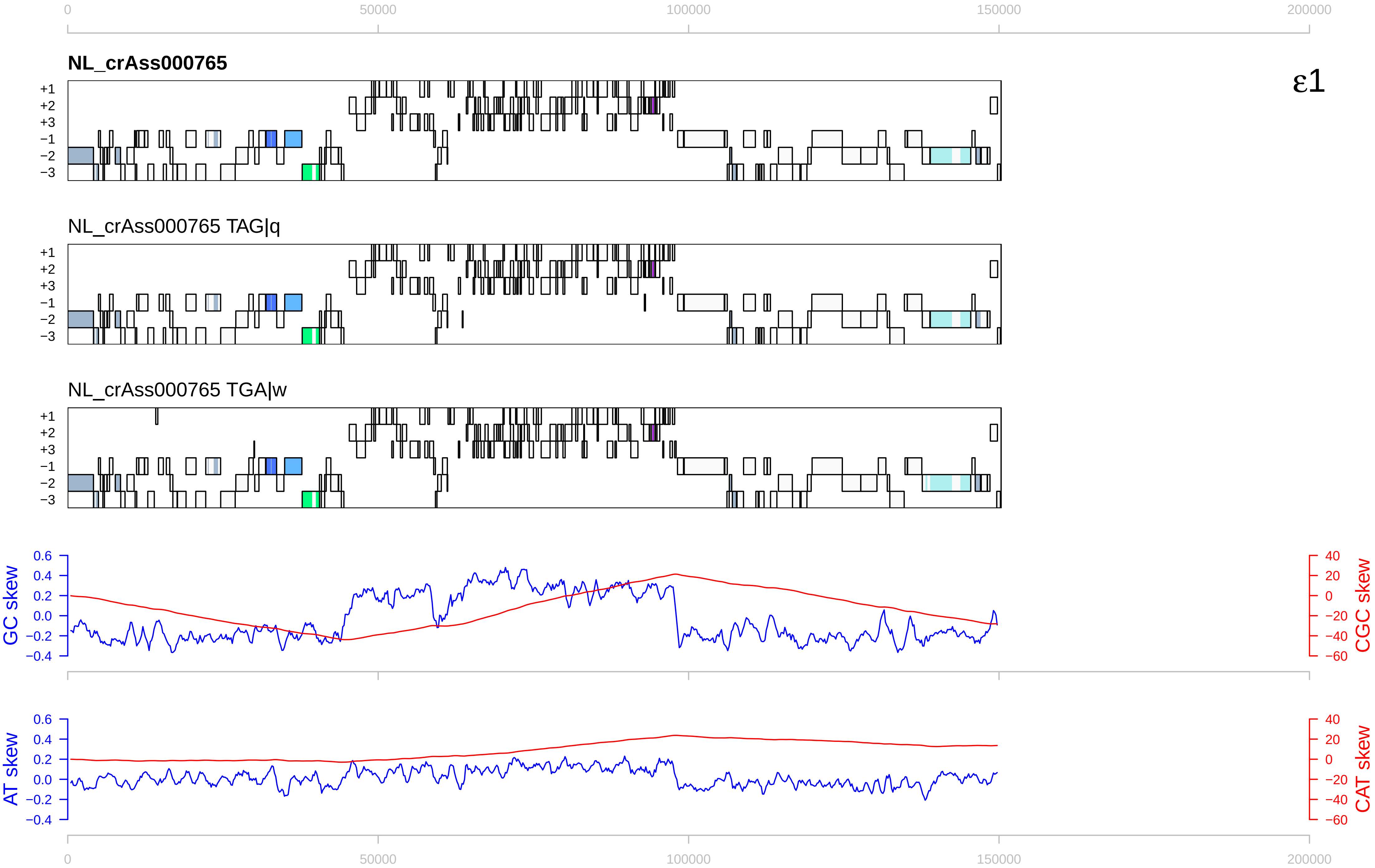

0
50000
100000
150000
200000
NL_crAss000765
ε1
+1
+2
+3
−1
−2
−3
NL_crAss000765 TAG|q
NL_crAss000765 TGA|w
GC skew
CGC skew
AT skew
CAT skew
0.6
0.4
0.2
0.0
−0.2
−0.4
40
20
0
−20
−40
−60

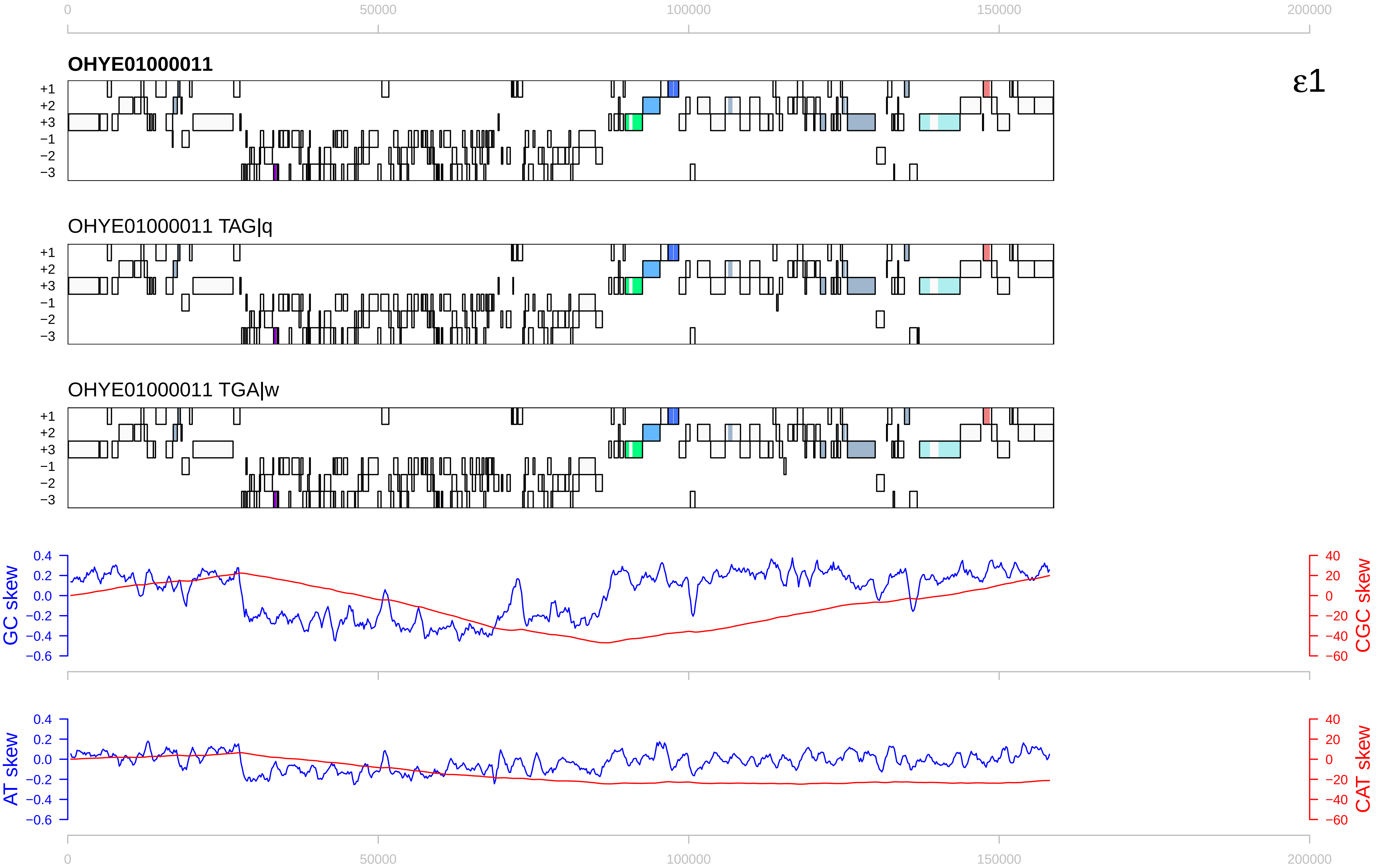

0
50000
100000
150000
200000
OHYE01000011
ε1
+1
+2
+3
−1
−2
−3
OHYE01000011 TAG|q
OHYE01000011 TGA|w
GC skew
CGC skew
AT skew
CAT skew
0.4
0.2
0.0
−0.2
−0.4
−0.6
40
20
0
−20
−40
−60

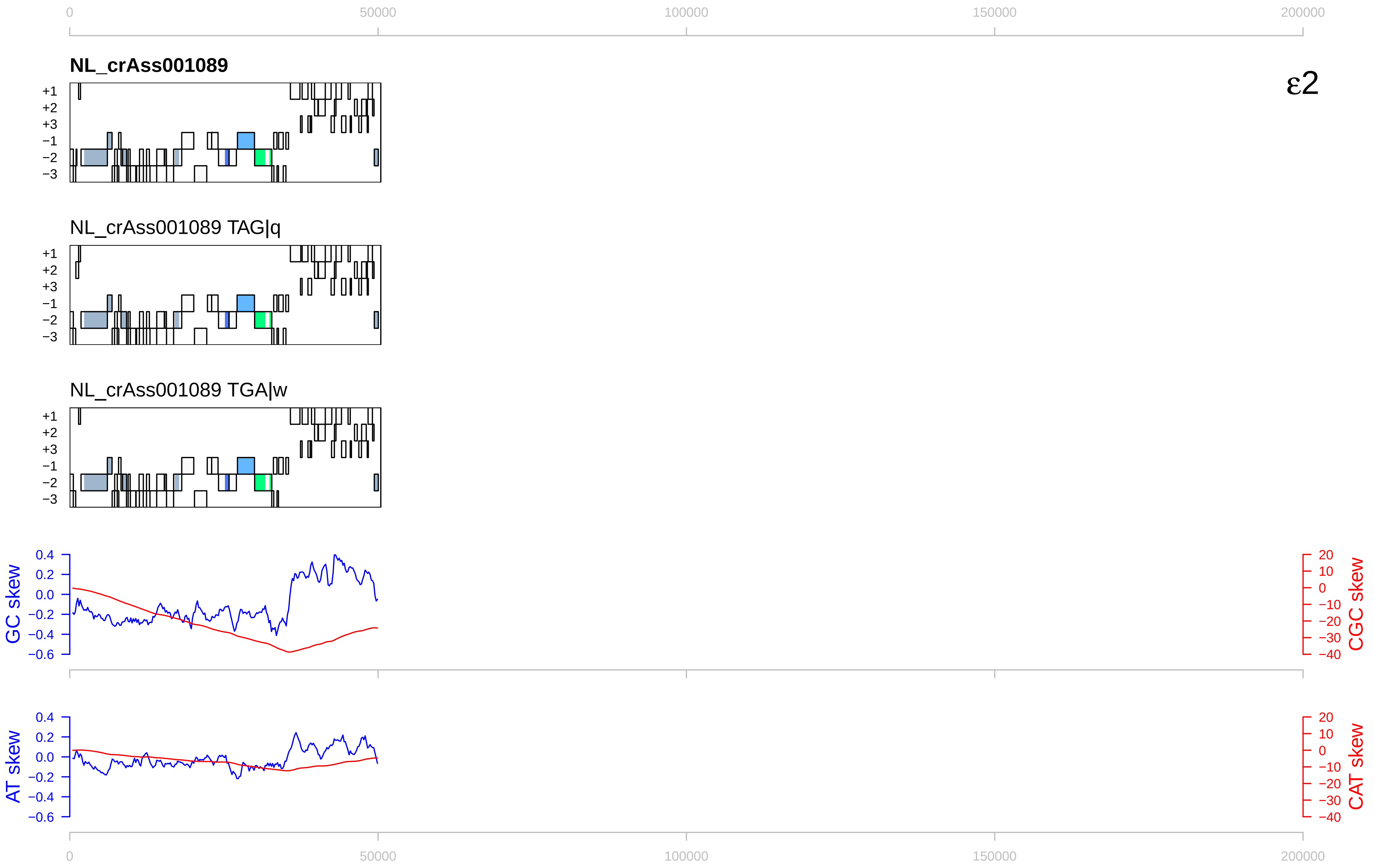

0
50000
100000
150000
200000
NL_crAss001089
ε2
+1
+2
+3
−1
−2
−3
NL_crAss001089 TAG|q
NL_crAss001089 TGA|w
GC skew
CGC skew
AT skew
CAT skew
0.4
0.2
0.0
−0.2
−0.4
−0.6
20
10
−10
−20
−30
−40

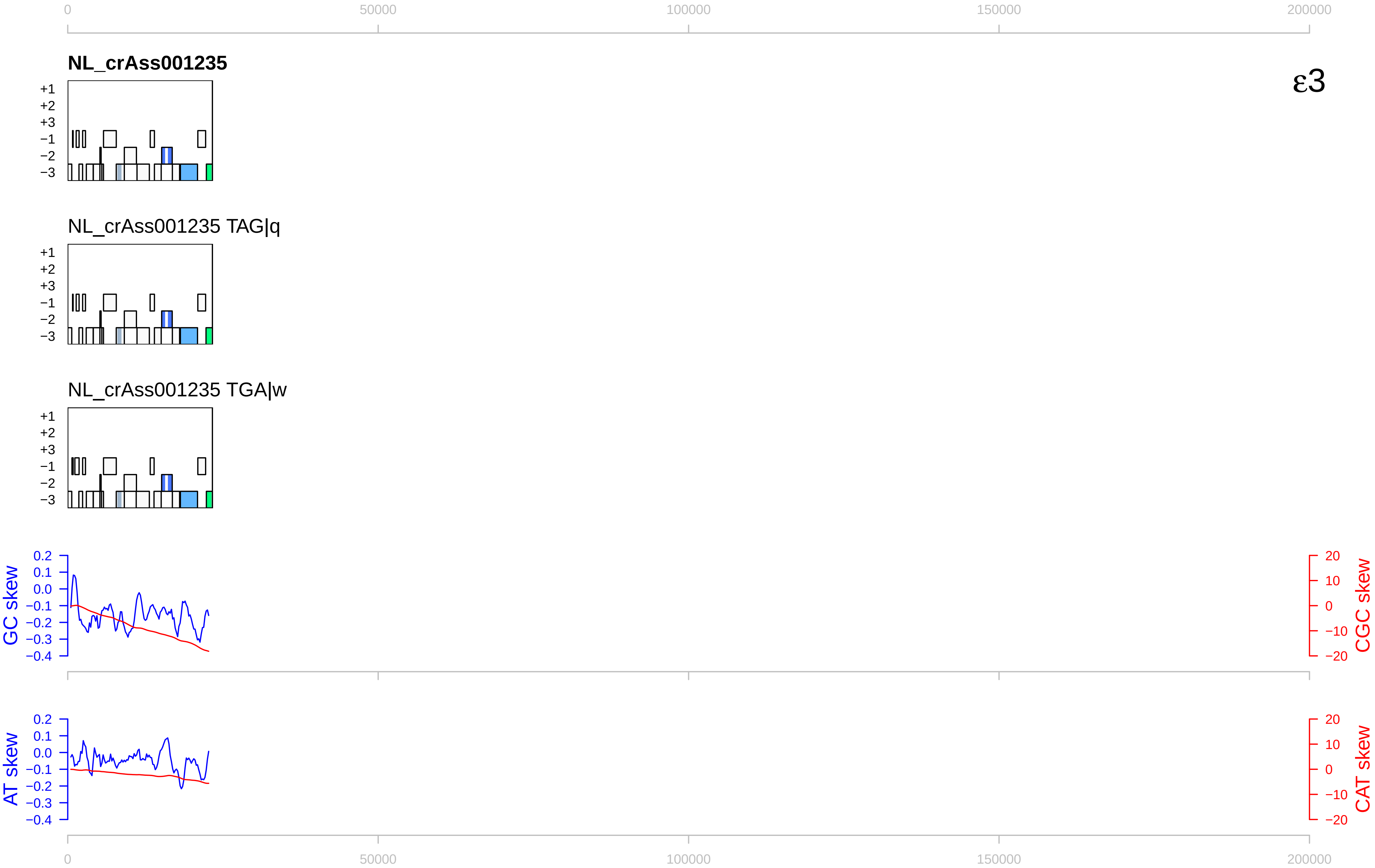

NL_crAss001235
ε3
NL_crAss001235 TAG|q
NL_crAss001235 TGA|w
+1
+2
+3
−1
−2
−3
GC skew
CGC skew
AT skew
CAT skew
0
50000
100000
150000
200000

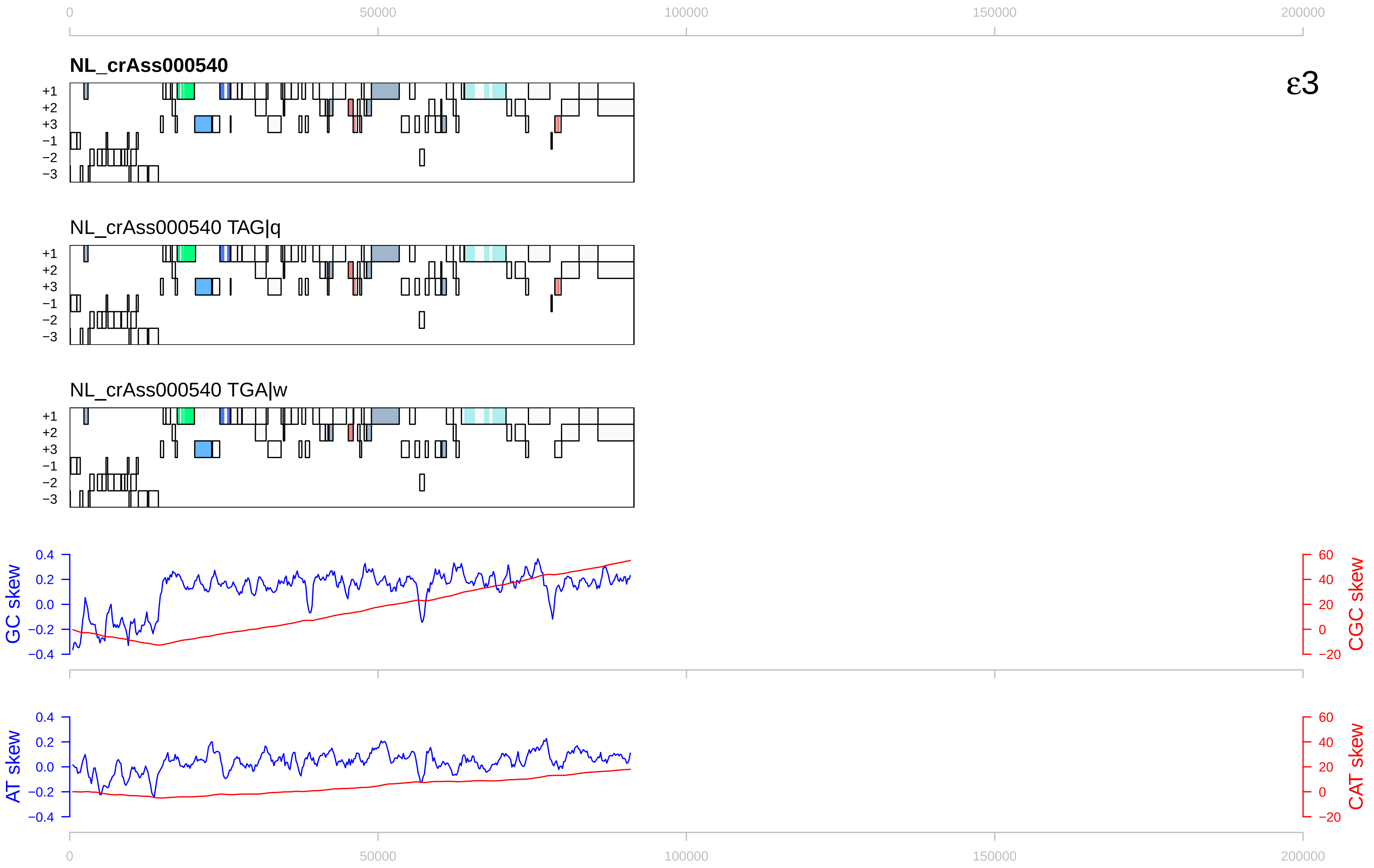

NL_crAss000540
ε3
NL_crAss000540 TAG|q
NL_crAss000540 TGA|w
+1
+2
+3
−1
−2
−3
GC skew
CGC skew
AT skew
CAT skew
0
50000
100000
150000
200000

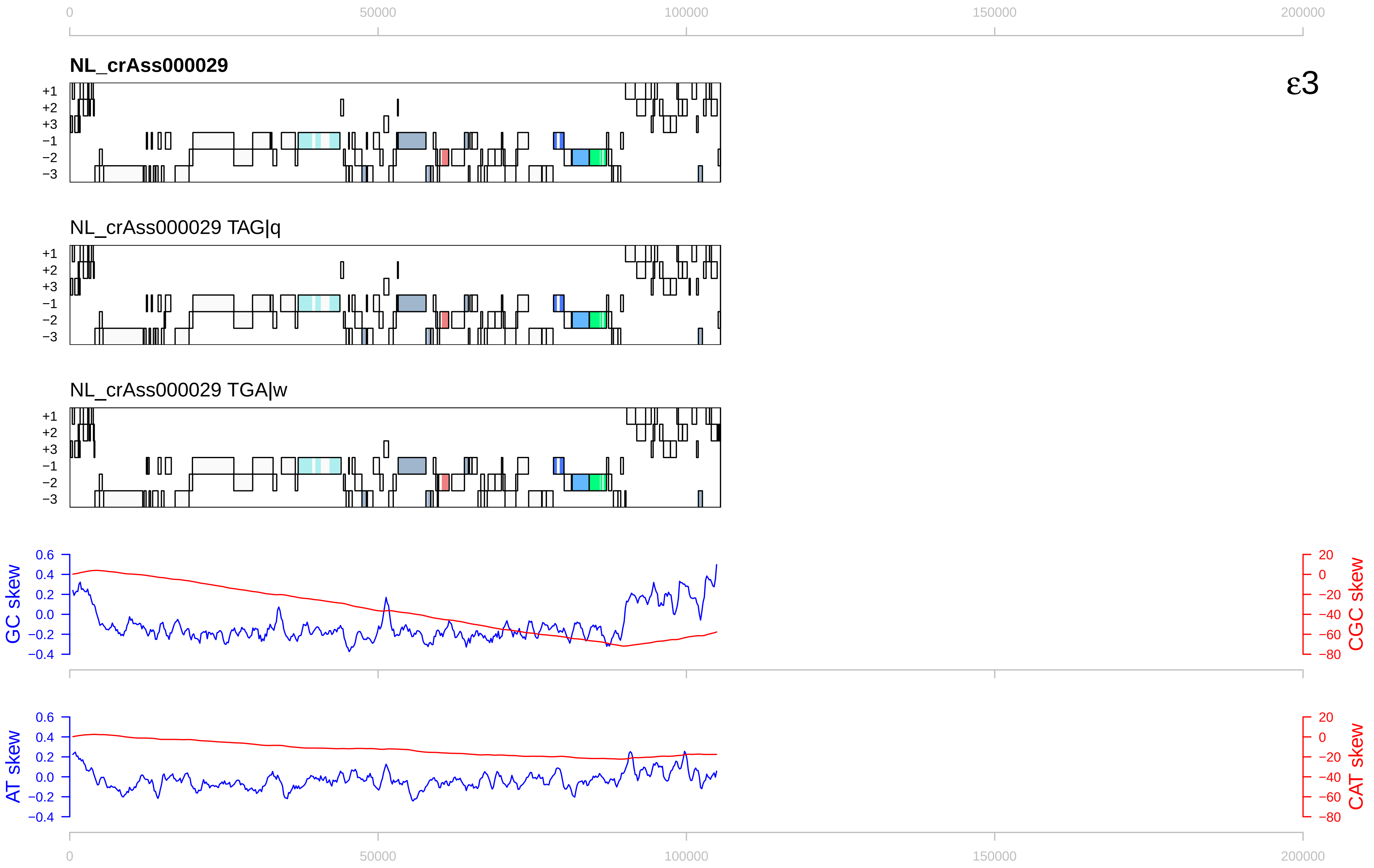
ε3
NL_crAss000029
NL_crAss000029 TAG|q
NL_crAss000029 TGA|w
+1
+2
+3
−1
−2
−3
GC skew
CGC skew
AT skew
CAT skew
0
50000
100000
150000
200000

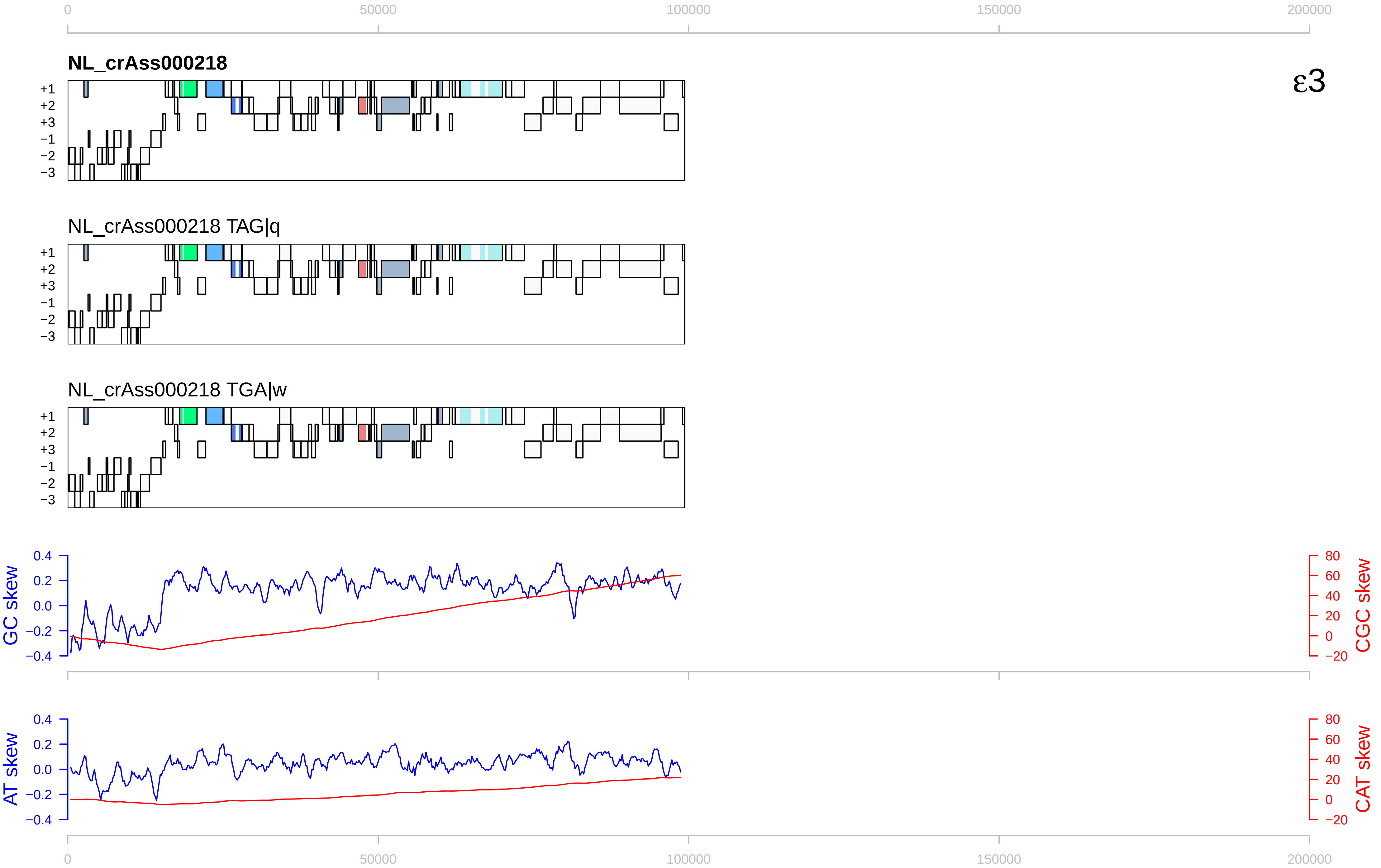

0
50000
100000
150000
200000
NL_crAss000218
ε3
+1
+2
+3
−1
−2
−3
NL_crAss000218 TAG|q
NL_crAss000218 TGA|w
GC skew
0.4
0.2
0.0
−0.2
−0.4
CGC skew
80
60
40
20
0
−20
AT skew
CAT skew

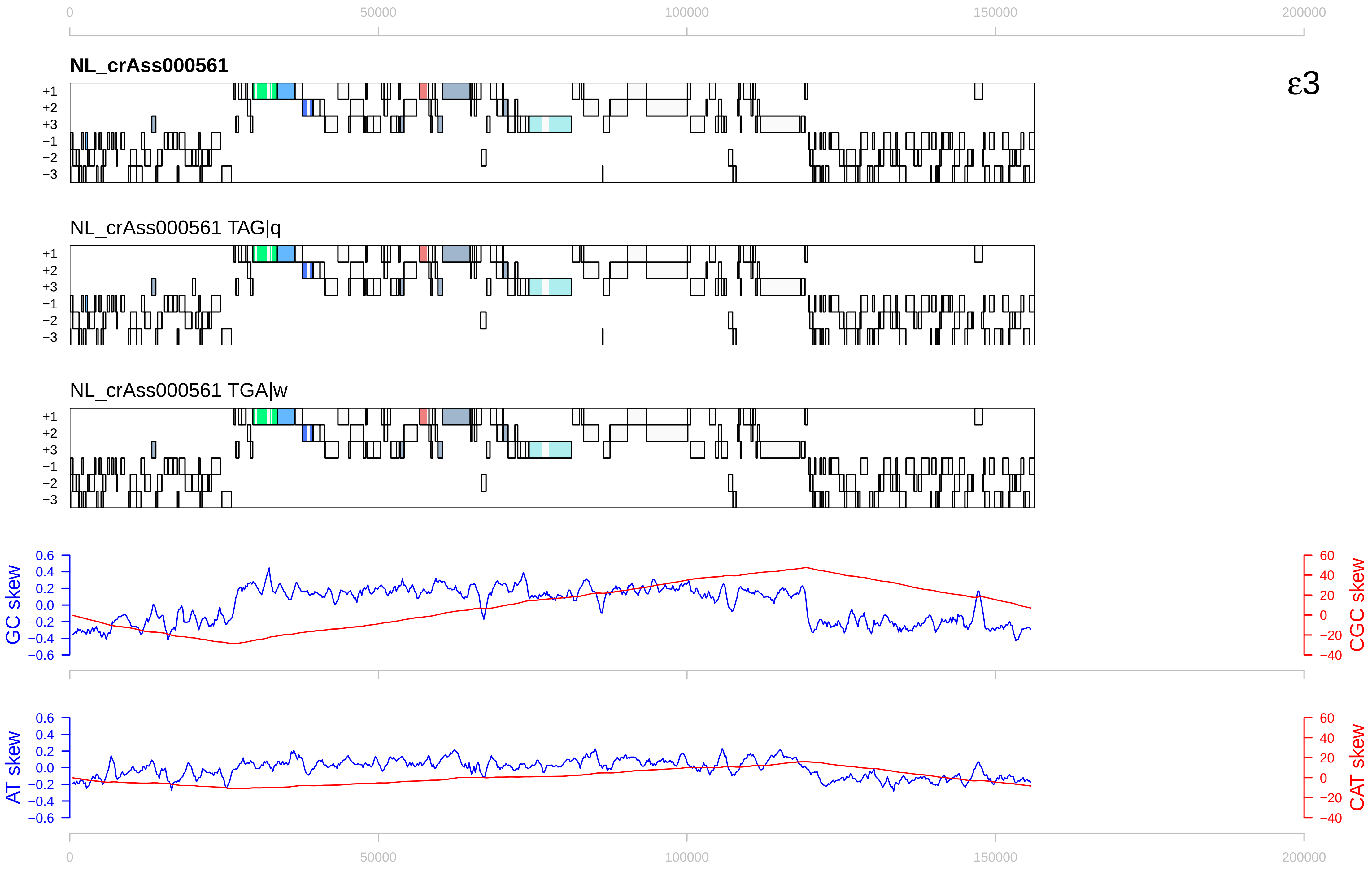
NL_crAss000561
NL_crAss000561 TAG|q
NL_crAss000561 TGA|w
ε3
+1
+2
+3
−1
−2
−3
0
50000
100000
150000
200000
GC skew
CGC skew
AT skew
CAT skew

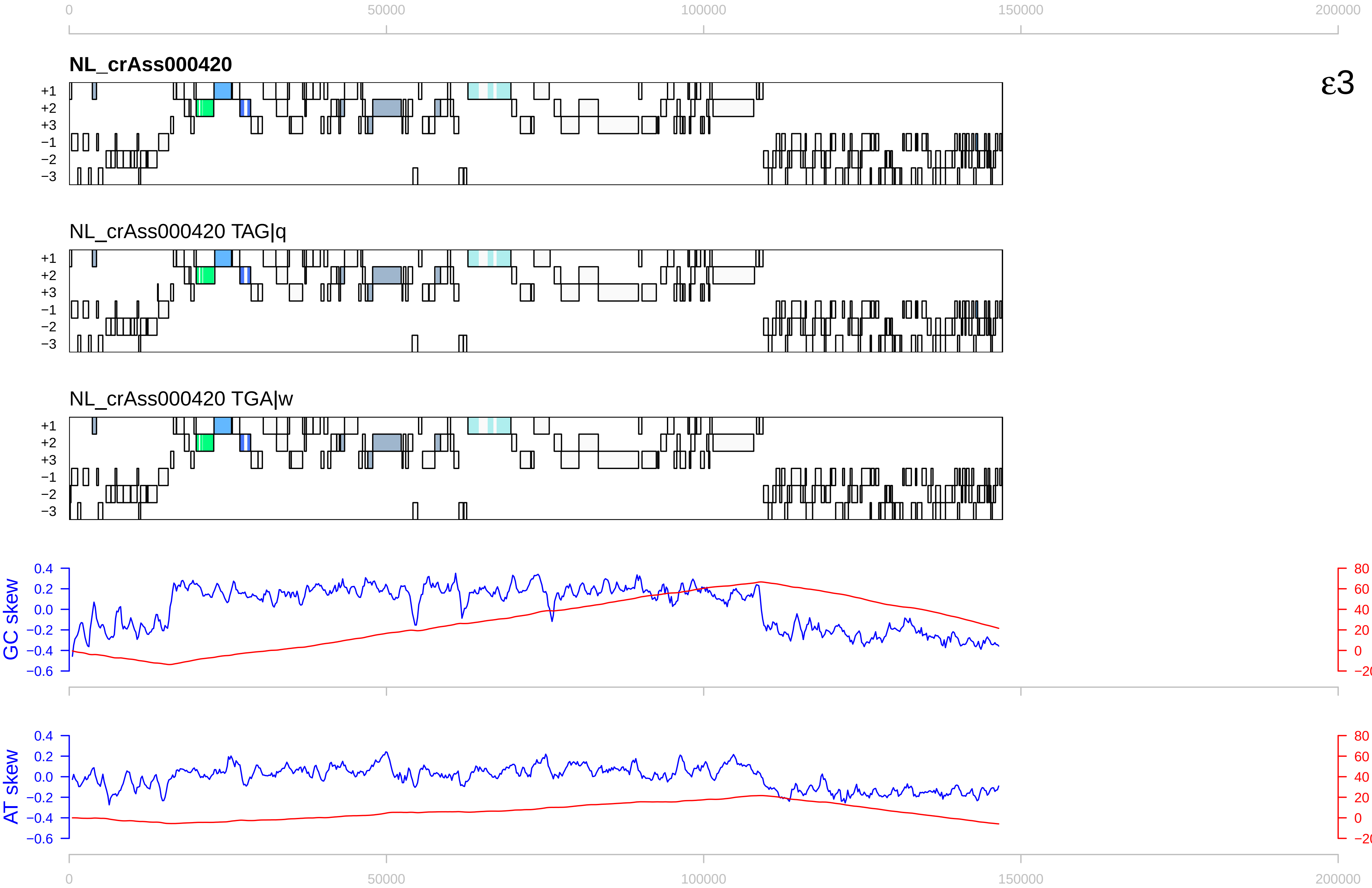
ε3
NL_crAss000420
NL_crAss000420 TAG|q
NL_crAss000420 TGA|w
+1
+2
+3
−1
−2
−3
GC skew
CGC skew
AT skew
CAT skew
0
50000
100000
150000
200000

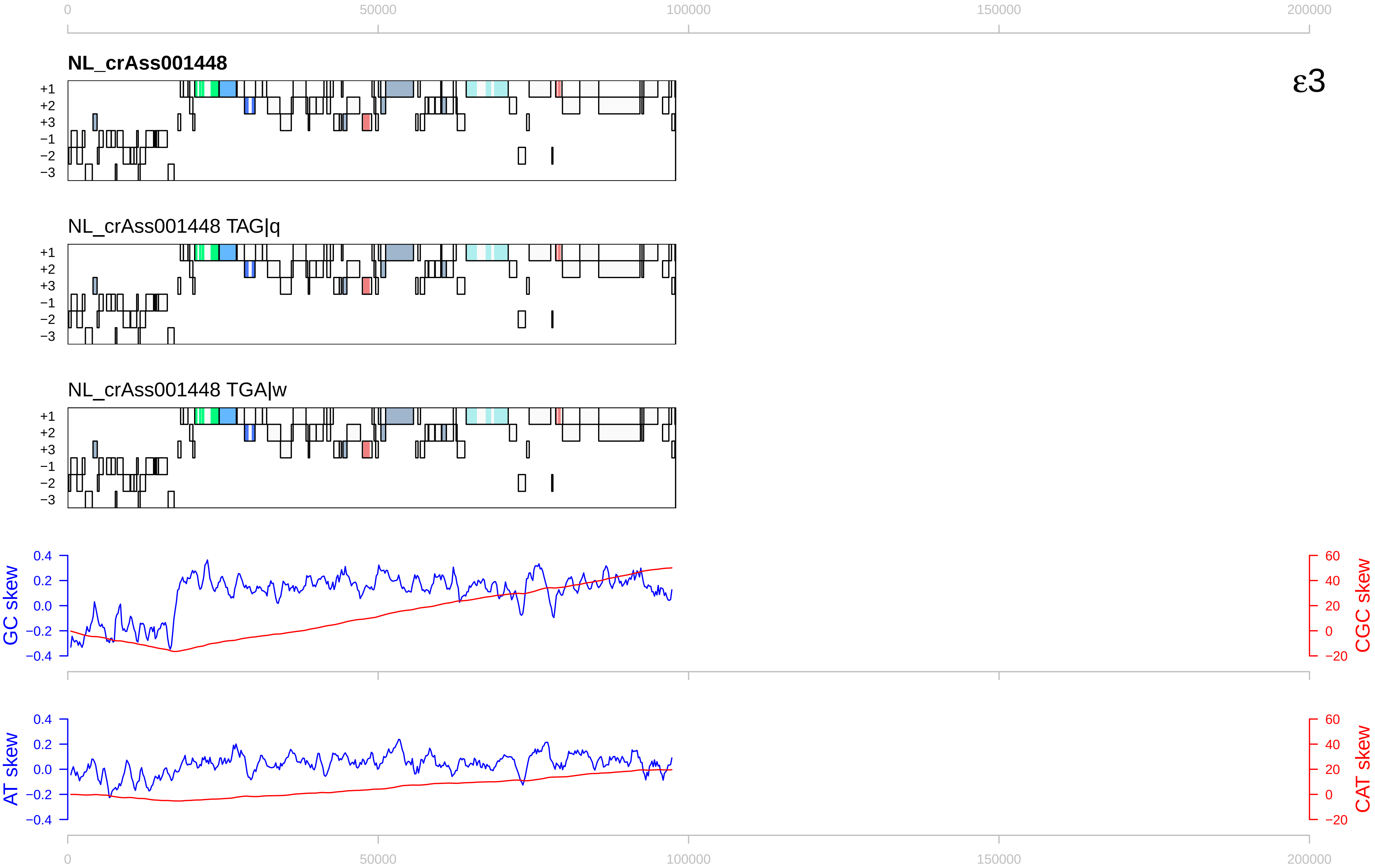

ε3
NL_crAss001448
NL_crAss001448 TAG|q
NL_crAss001448 TGA|w
+1
+2
+3
−1
−2
−3
GC skew
CGC skew
AT skew
CAT skew
0
50000
100000
150000
200000
0.4
0.2
0.0
−0.2
−0.4
60
40
20
0
−20

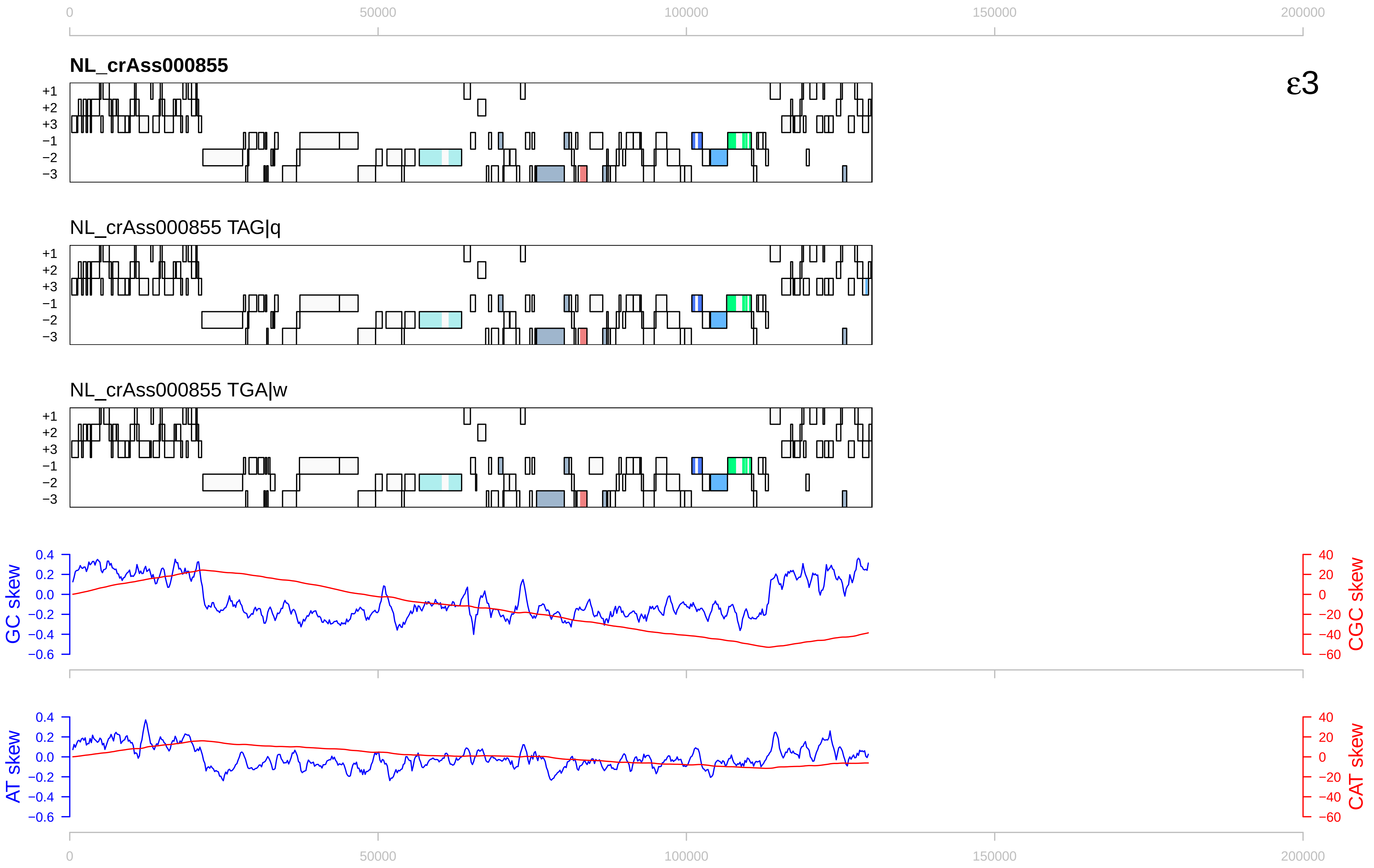

0
50000
100000
150000
200000
NL_crAss000855
ε3
+1
+2
+3
−1
−2
−3
NL_crAss000855 TAG|q
NL_crAss000855 TGA|w
GC skew
CGC skew
AT skew
CAT skew
0.4
0.2
0.0
−0.2
−0.4
−0.6
40
20
0
−20
−40
−60

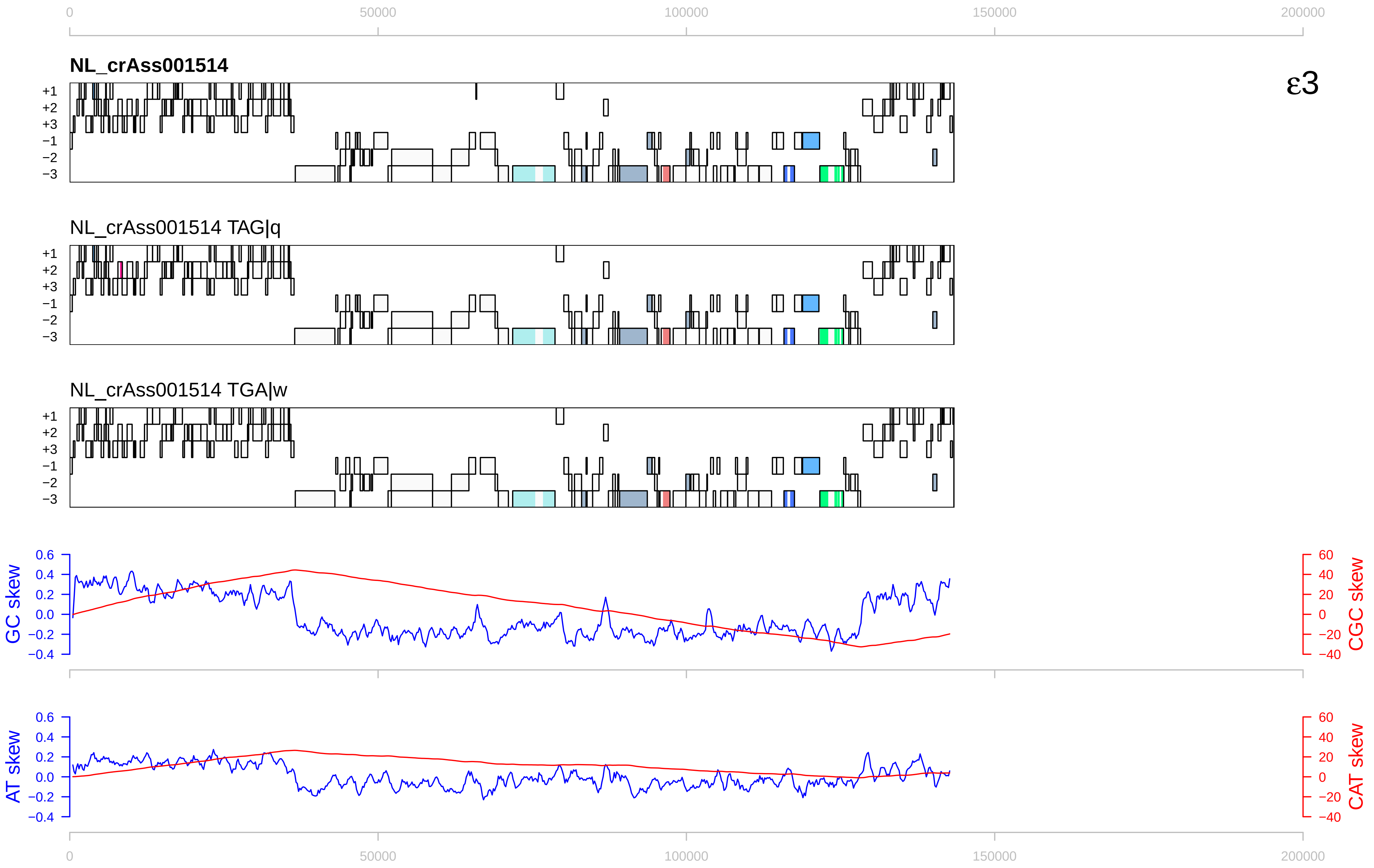
0
50000
100000
150000
200000
NL_crAss001514
ε3
+1
+2
+3
−1
−2
−3
NL_crAss001514 TAG|q
NL_crAss001514 TGA|w
GC skew
CGC skew
AT skew
CAT skew
0.6
0.4
0.2
0.0
−0.2
−0.4
60
40
20
0
−20
−40

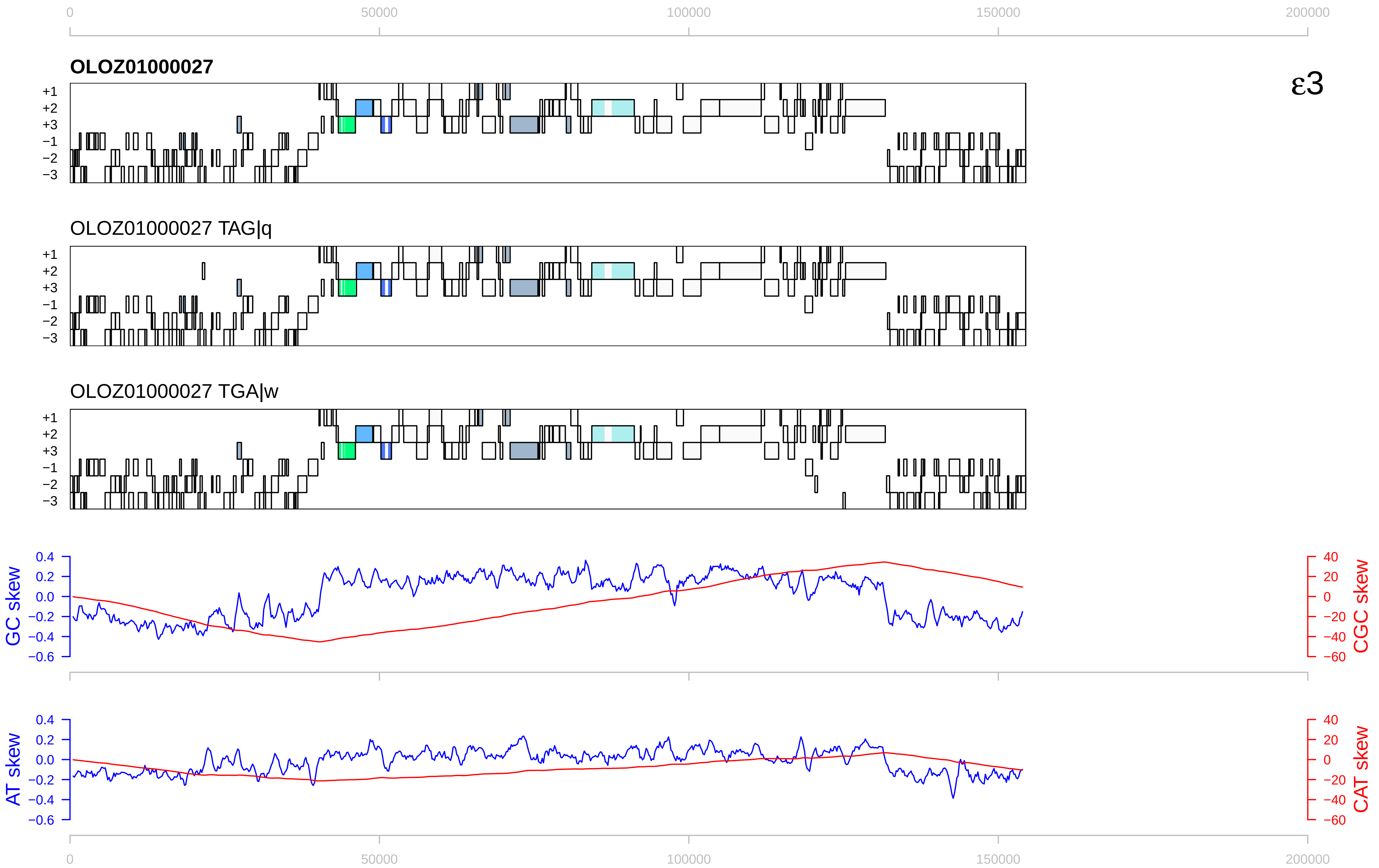

ε3
OLOZ01000027
OLOZ01000027 TAG|q
OLOZ01000027 TGA|w
+1
+2
+3
−1
−2
−3
GC skew
CGC skew
AT skew
CAT skew
0
50000
100000
150000
200000

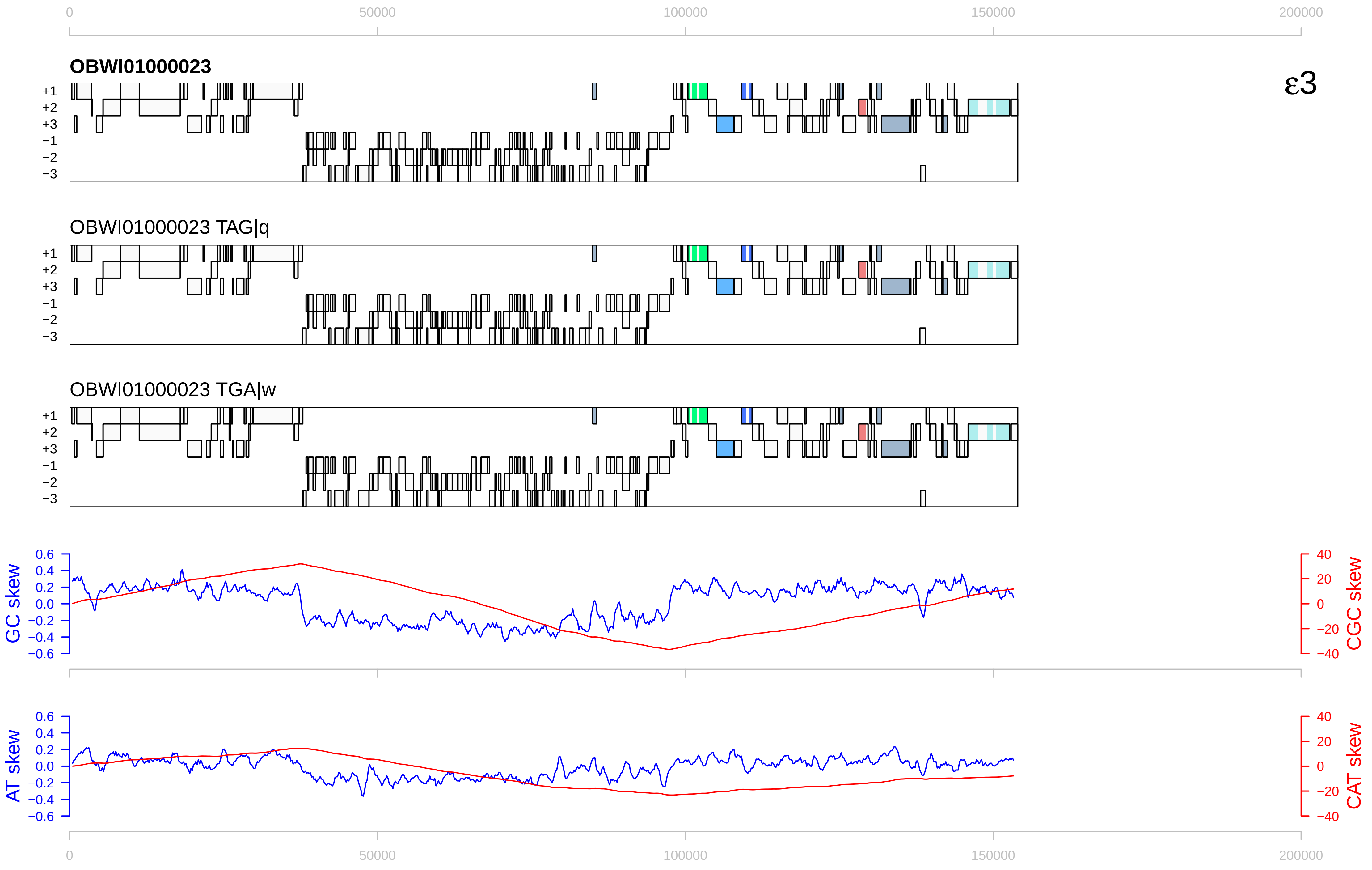

0
50000
100000
150000
200000
OBWI01000023
ε3
+1
+2
+3
−1
−2
−3
OBWI01000023 TAG|q
OBWI01000023 TGA|w
GC skew
CGC skew
AT skew
CAT skew
0.6
0.4
0.2
0.0
−0.2
−0.4
−0.6
40
20
0
−20
−40

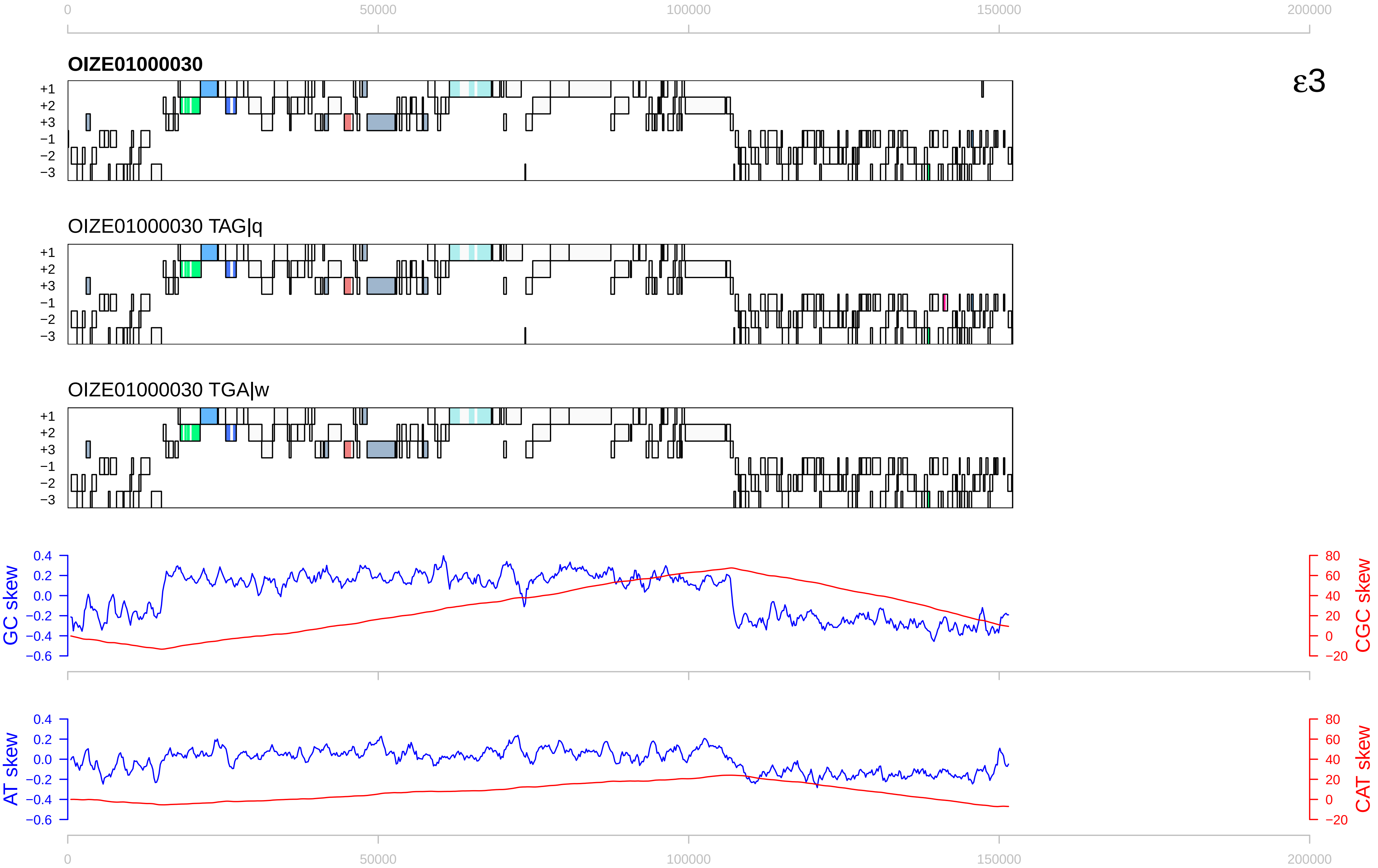

0
50000
100000
150000
200000
OIZE01000030
ε3
+1
+2
+3
−1
−2
−3
OIZE01000030 TAG|q
OIZE01000030 TGA|w
GC skew
CGC skew
AT skew
CAT skew
0.4
0.2
0.0
−0.2
−0.4
−0.6
80
60
40
20
0
−20

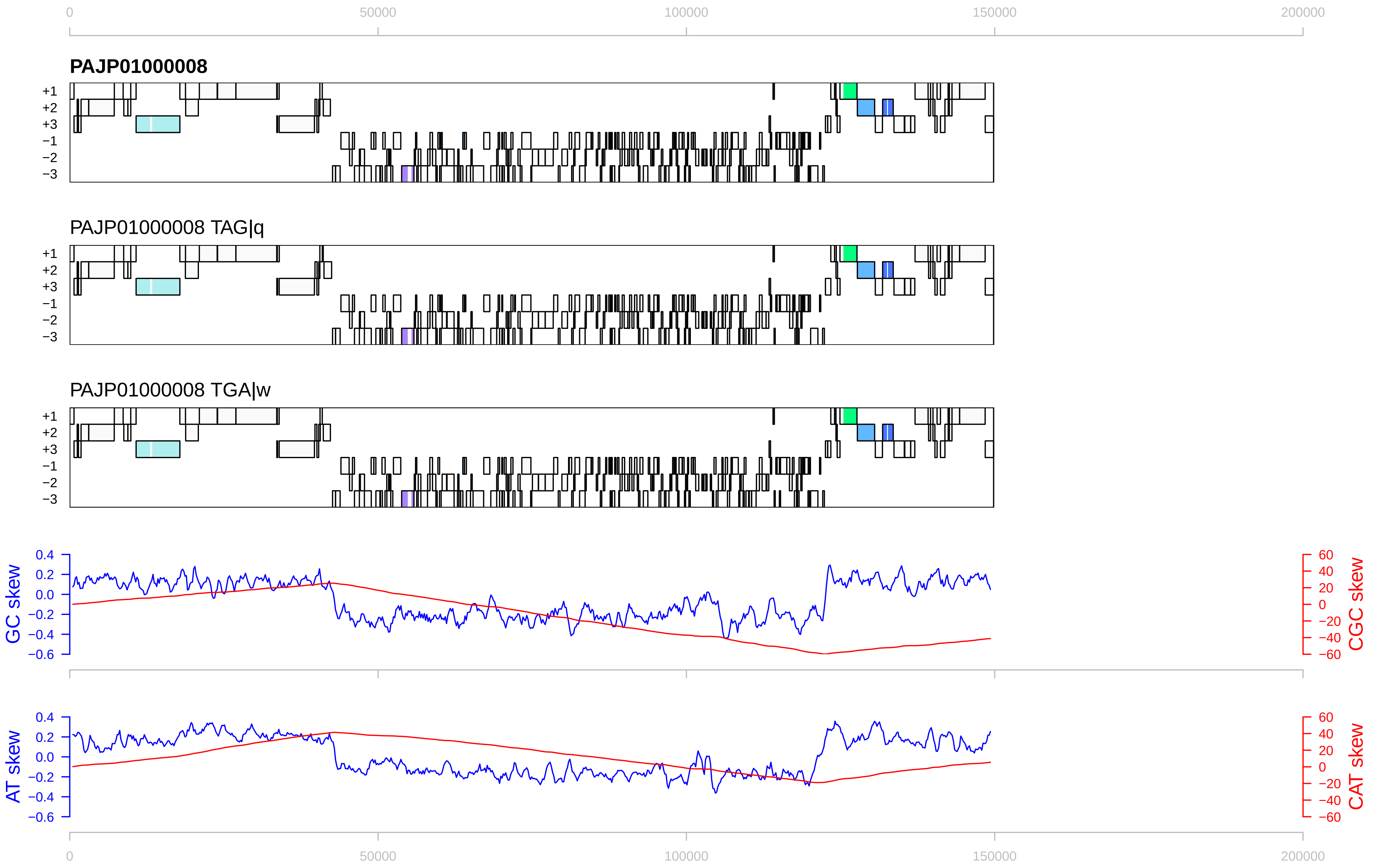

0
50000
100000
150000
200000
PAJP01000008
+1
+2
+3
−1
−2
−3
PAJP01000008 TAG|q
PAJP01000008 TGA|w
GC skew
CGC skew
AT skew
CAT skew
0.4
0.2
0.0
−0.2
−0.4
−0.6
60
40
20
0
−20
−40
−60

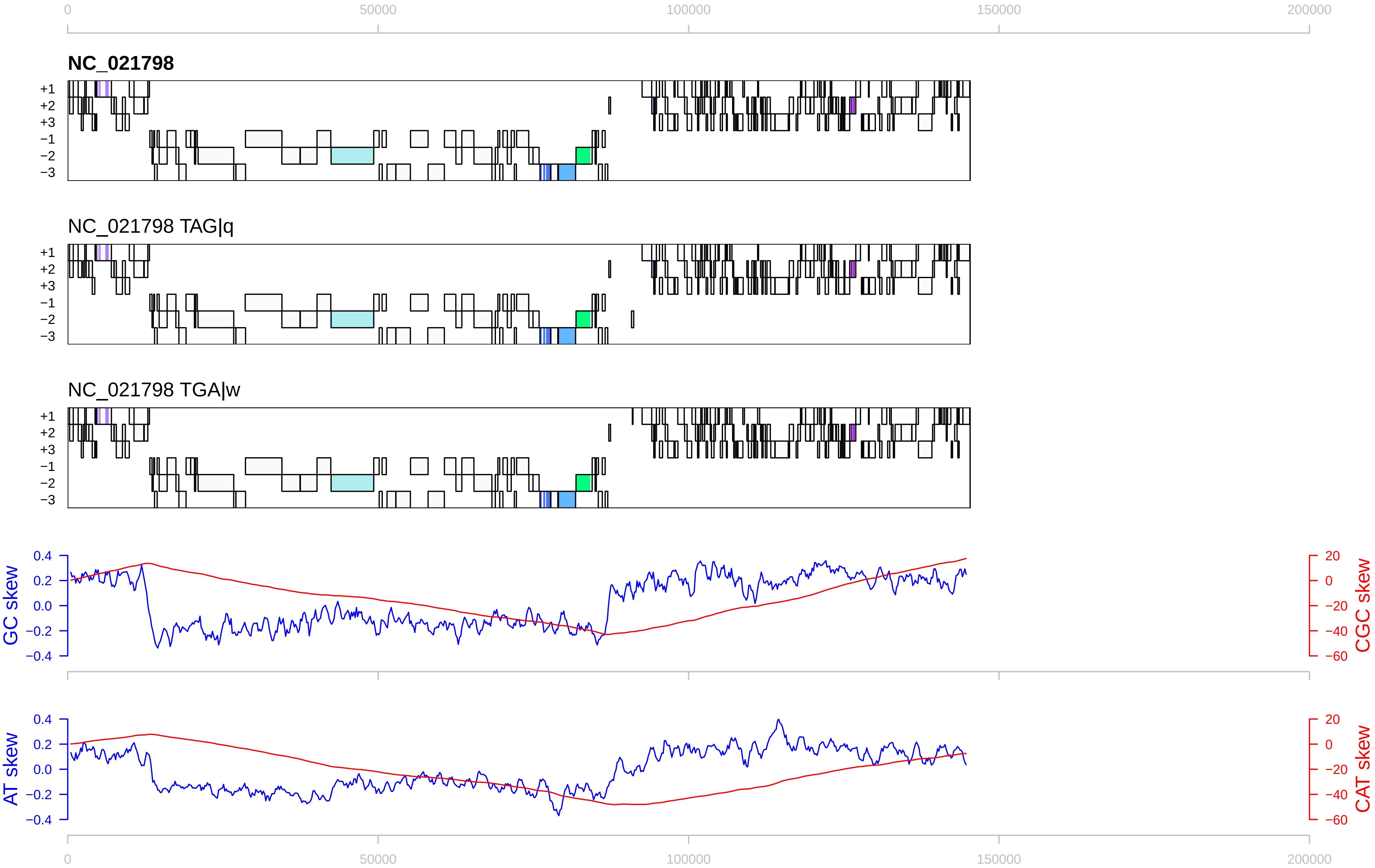

0
50000
100000
150000
200000
NC_021798
NC_021798 TAG|q
NC_021798 TGA|w
+1
+2
+3
−1
−2
−3
GC skew
CGC skew
AT skew
CAT skew
0.4
0.2
0.0
−0.2
−0.4
20
0
−20
−40
−60

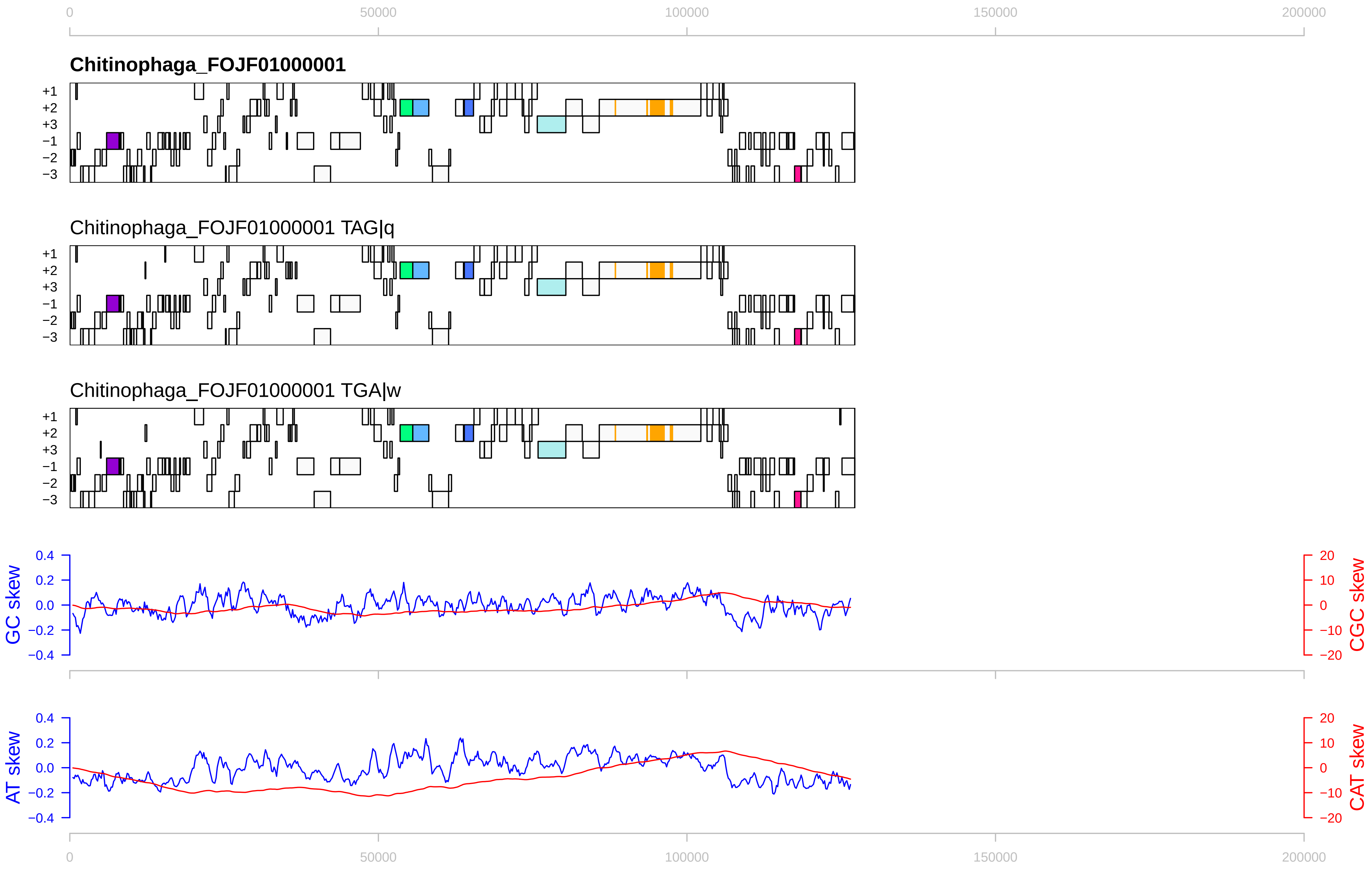

0
50000
100000
150000
200000
Chitinophaga_FOJF01000001
+1
+2
+3
−1
−2
−3
Chitinophaga_FOJF01000001 TAG|q
Chitinophaga_FOJF01000001 TGA|w
GC skew
CGC skew
0.4
0.2
0.0
−0.2
−0.4
20
10
0
−10
−20
AT skew
CAT skew

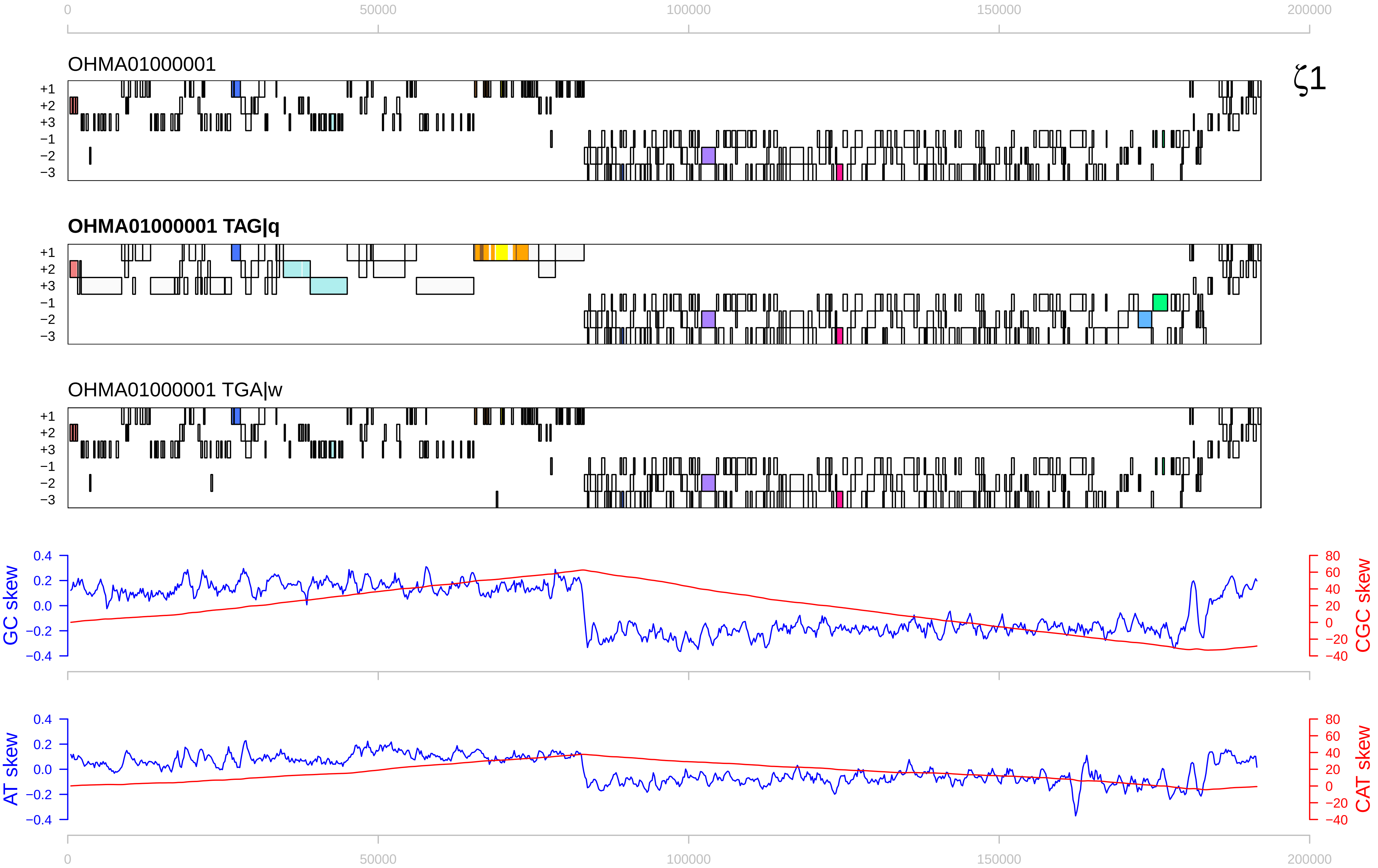

0
50000
100000
150000
200000
OHMA01000001
ζ1
+1
+2
+3
−1
−2
−3
OHMA01000001 TAG|q
OHMA01000001 TGA|w
GC skew
CGC skew
AT skew
CAT skew
0.4
0.2
0.0
−0.2
−0.4
80
60
40
20
0
−20
−40

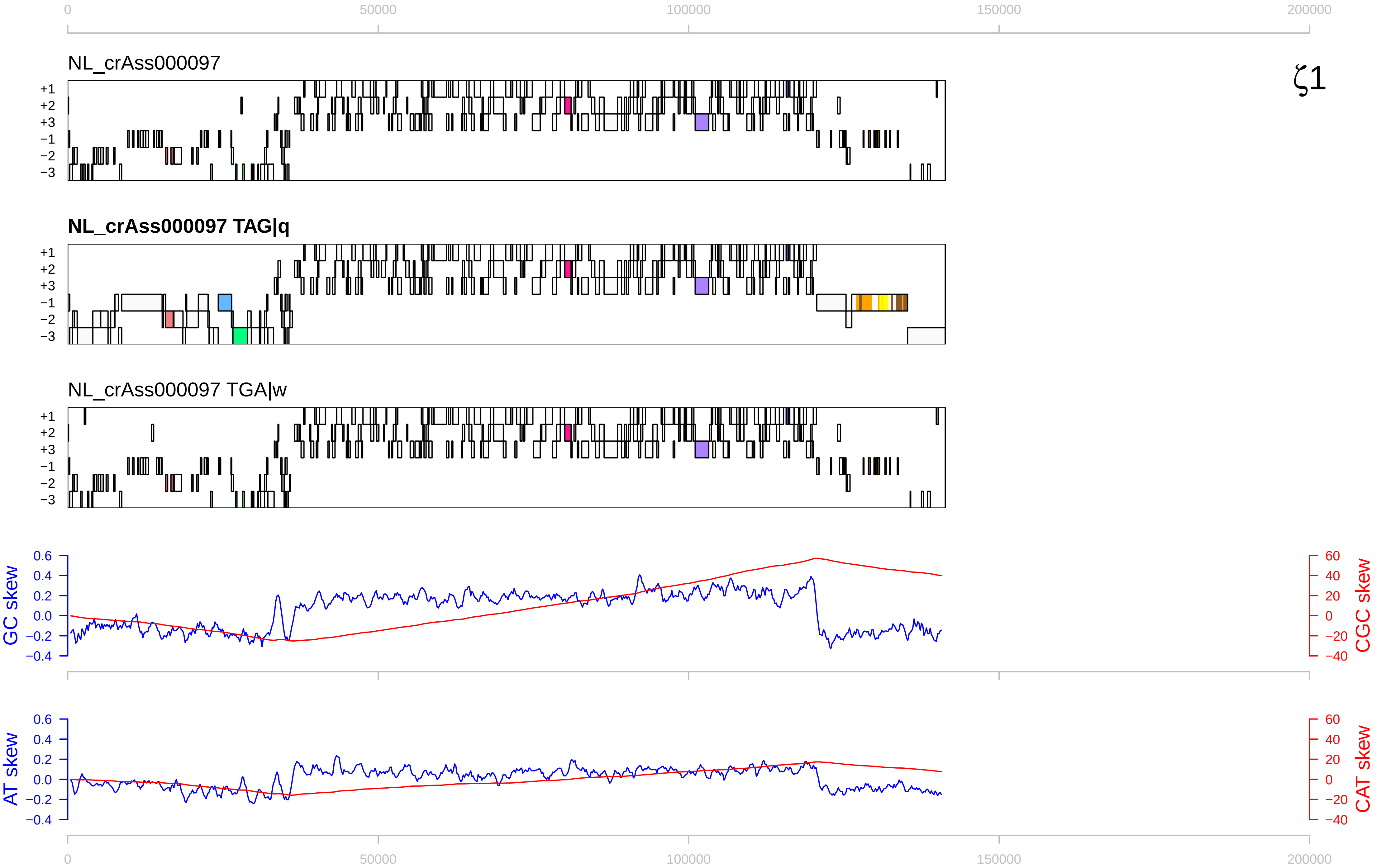

0
50000
100000
150000
200000
NL_crAss000097
ζ1
+1
+2
+3
−1
−2
−3
NL_crAss000097 TAG|q
NL_crAss000097 TGA|w
GC skew
CGC skew
AT skew
CAT skew
0.6
0.4
0.2
0.0
−0.2
−0.4
60
40
20
0
−20
−40

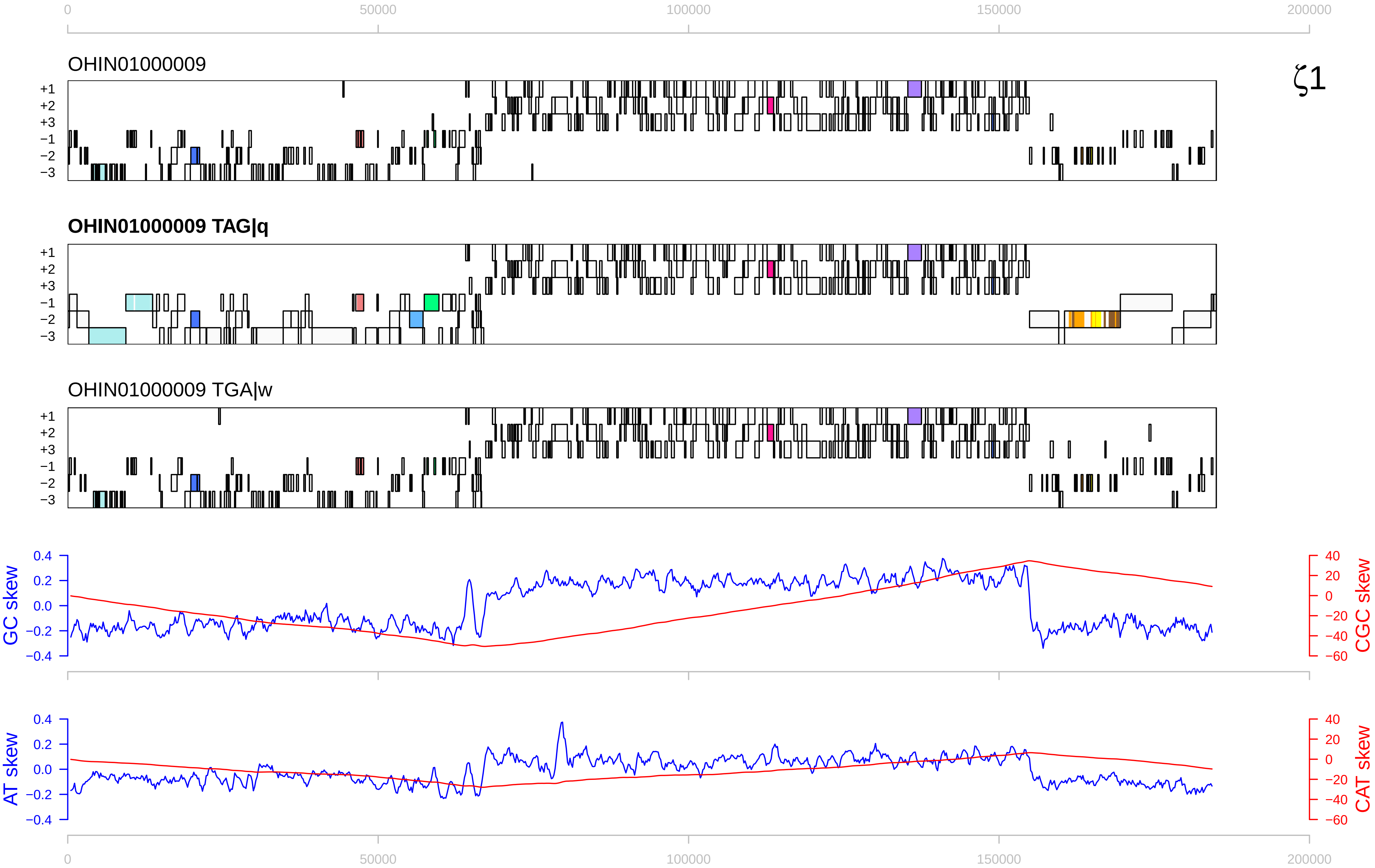

0
50000
100000
150000
200000
OHIN01000009
ζ1
+1
+2
+3
−1
−2
−3
OHIN01000009 TAG|q
OHIN01000009 TGA|w
GC skew
CGC skew
AT skew
CAT skew
0.4
0.2
0.0
−0.2
−0.4
40
20
0
−20
−40
−60

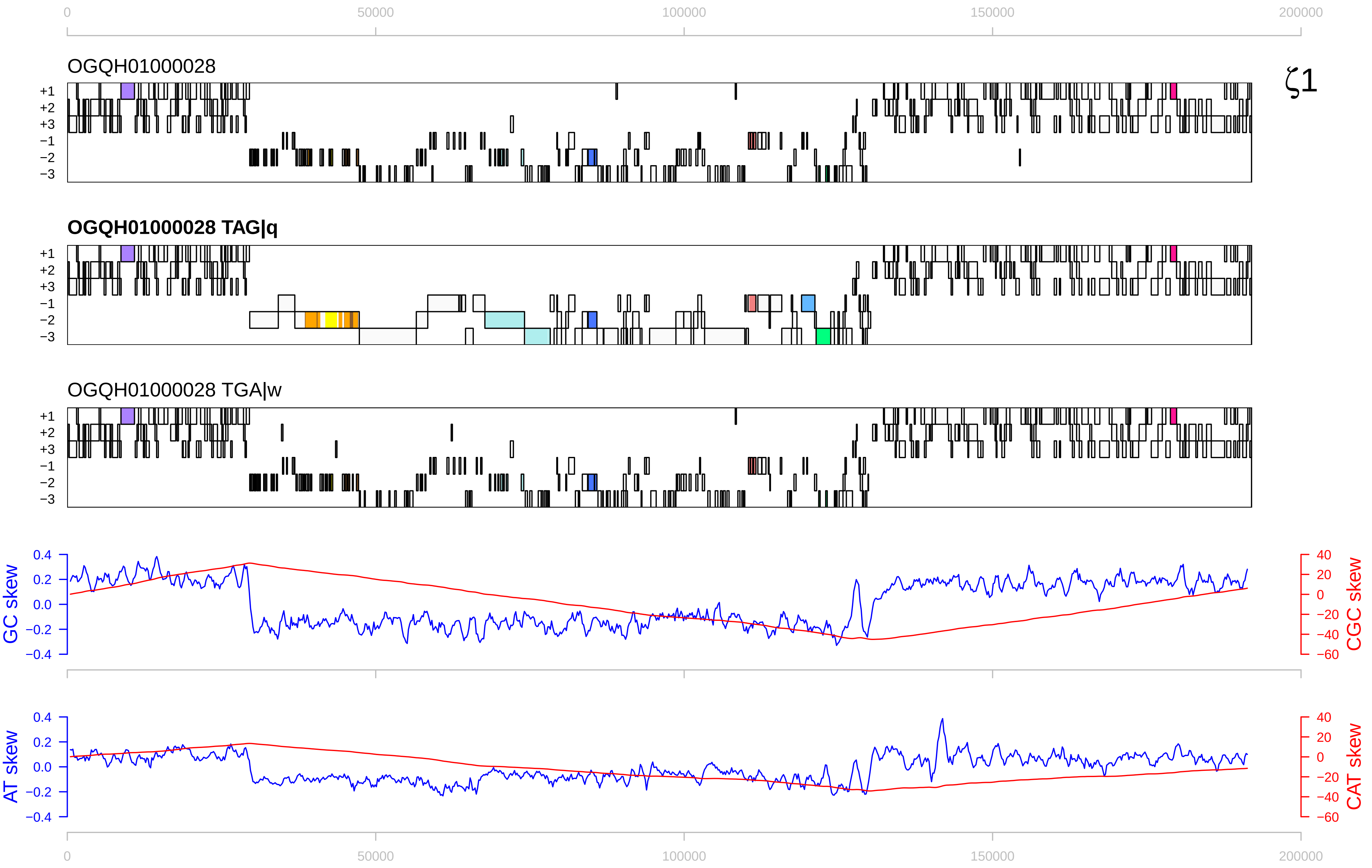

0
50000
100000
150000
200000
OGQH01000028
ζ1
+1
+2
+3
−1
−2
−3
OGQH01000028 TAG|q
OGQH01000028 TGA|w
GC skew
CGC skew
0.4
0.2
0.0
−0.2
−0.4
40
20
0
−20
−40
−60
AT skew
CAT skew

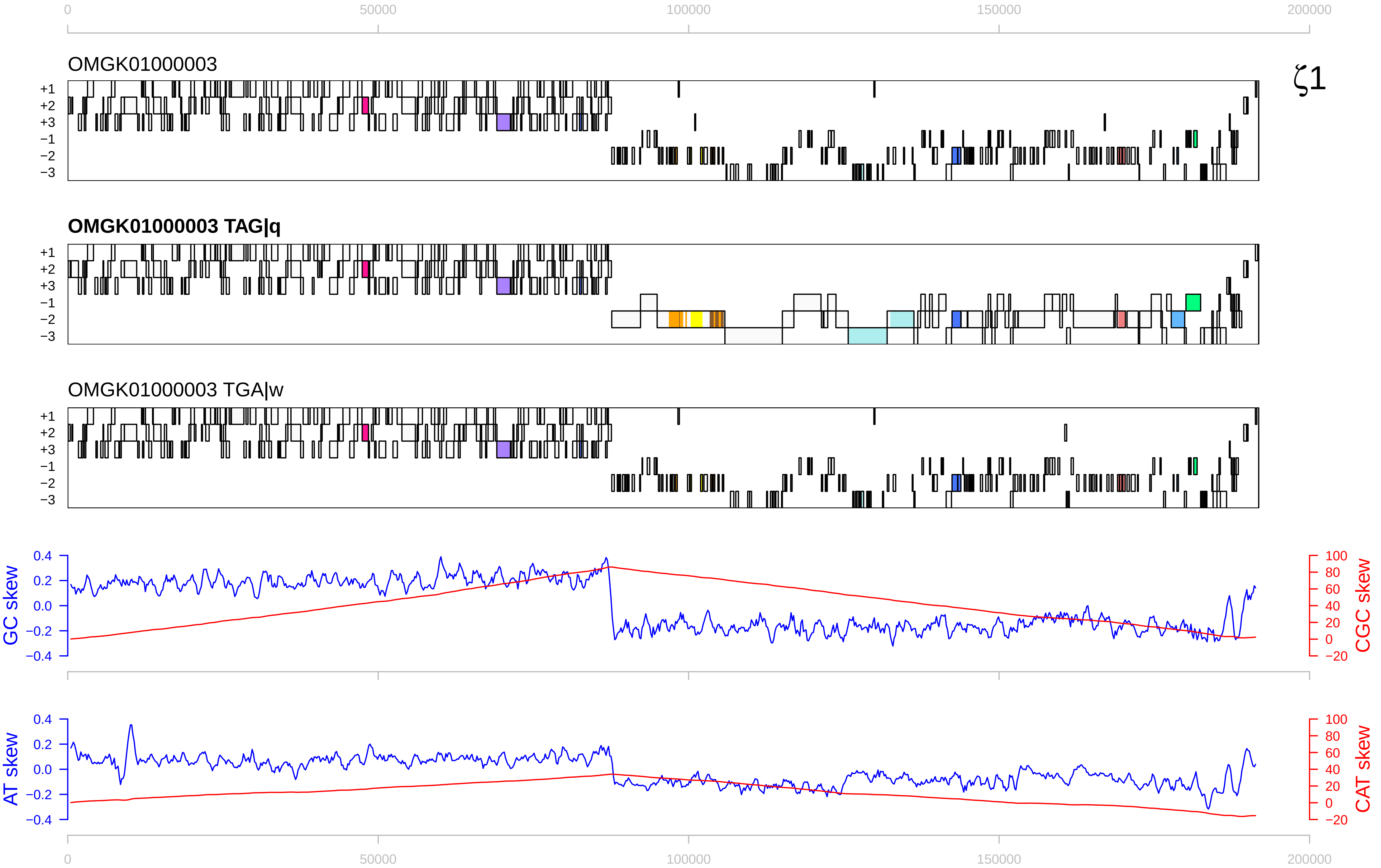

0
50000
100000
150000
200000
OMGK01000003
ζ1
+1
+2
+3
−1
−2
−3
OMGK01000003 TAG|q
OMGK01000003 TGA|w
GC skew
CGC skew
0.4
0.2
0.0
−0.2
−0.4
100
80
60
40
20
0
−20
AT skew
CAT skew

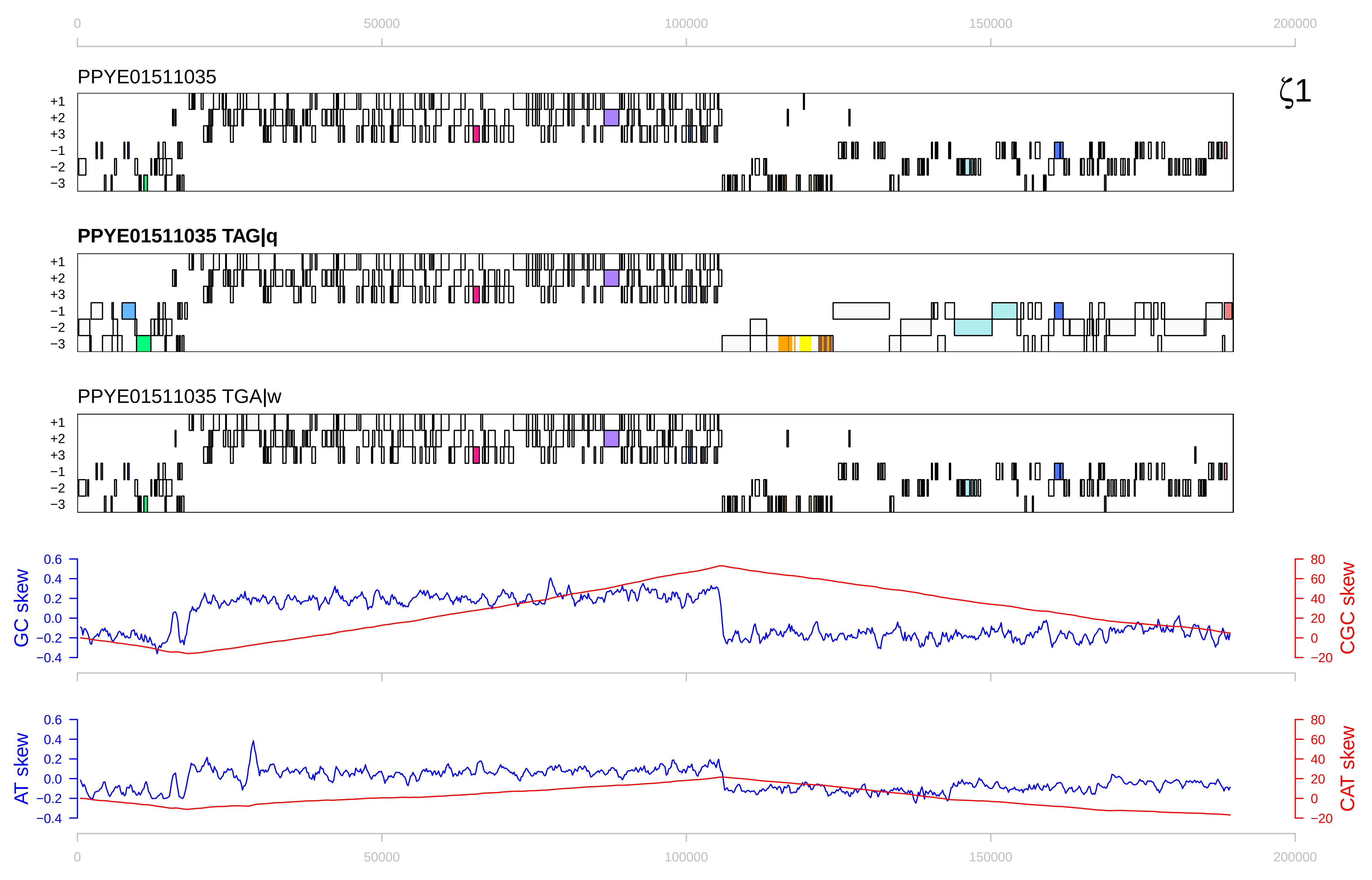

PPYE01511035
ζ1
PPYE01511035 TAG|q
PPYE01511035 TGA|w
+1
+2
+3
−1
−2
−3
GC skew
CGC skew
AT skew
CAT skew
0
50000
100000
150000
200000

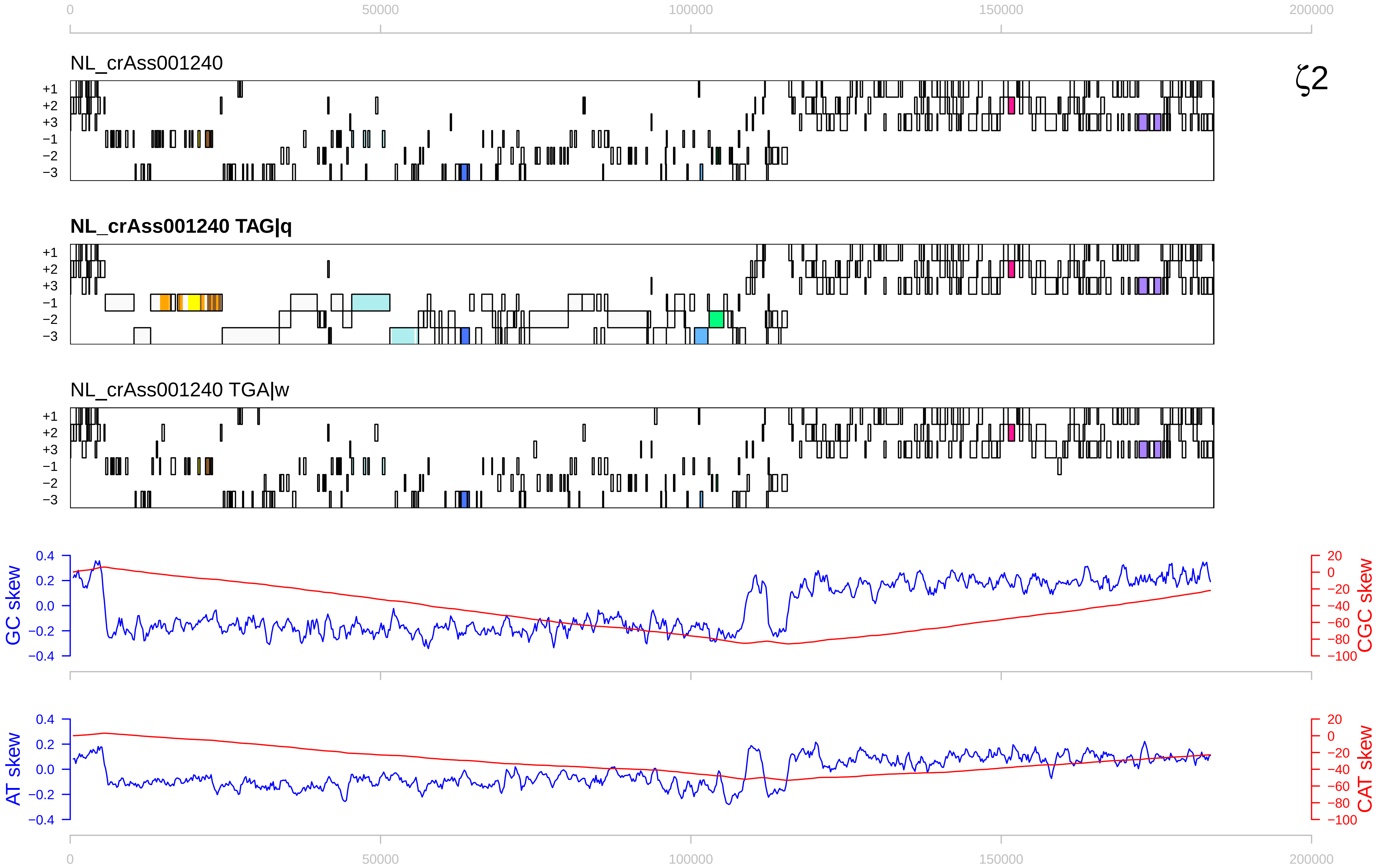

0
50000
100000
150000
200000
NL_crAss001240
ζ2
+1
+2
+3
−1
−2
−3
NL_crAss001240 TAG|q
NL_crAss001240 TGA|w
GC skew
CGC skew
AT skew
CAT skew
0.4
0.2
0.0
−0.2
−0.4
20
0
−20
−40
−60
−80
−100

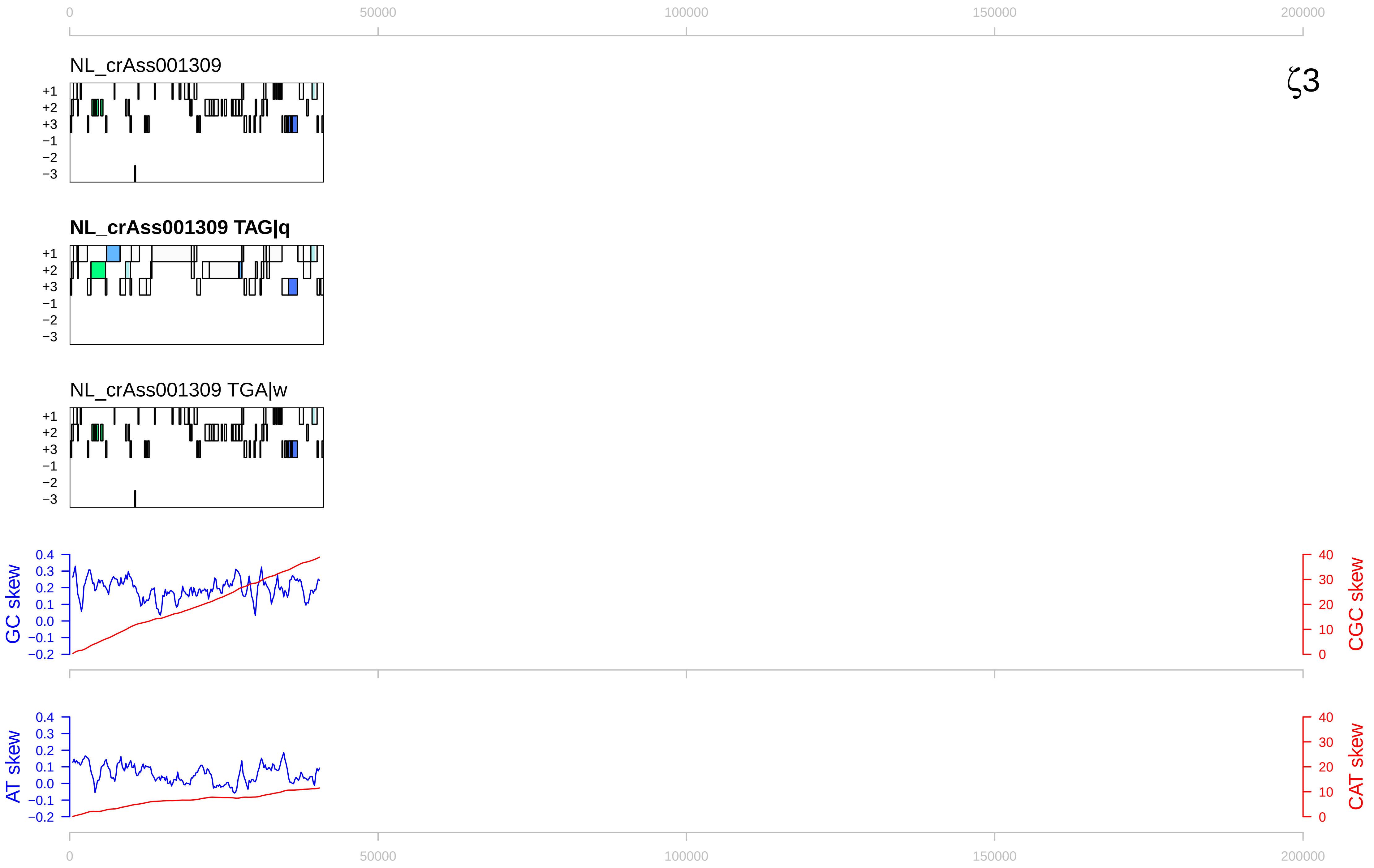

ζ3
NL_crAss001309
NL_crAss001309 TAG|q
NL_crAss001309 TGA|w
+1
+2
+3
−1
−2
−3
GC skew
CGC skew
AT skew
CAT skew
0
50000
100000
150000
200000

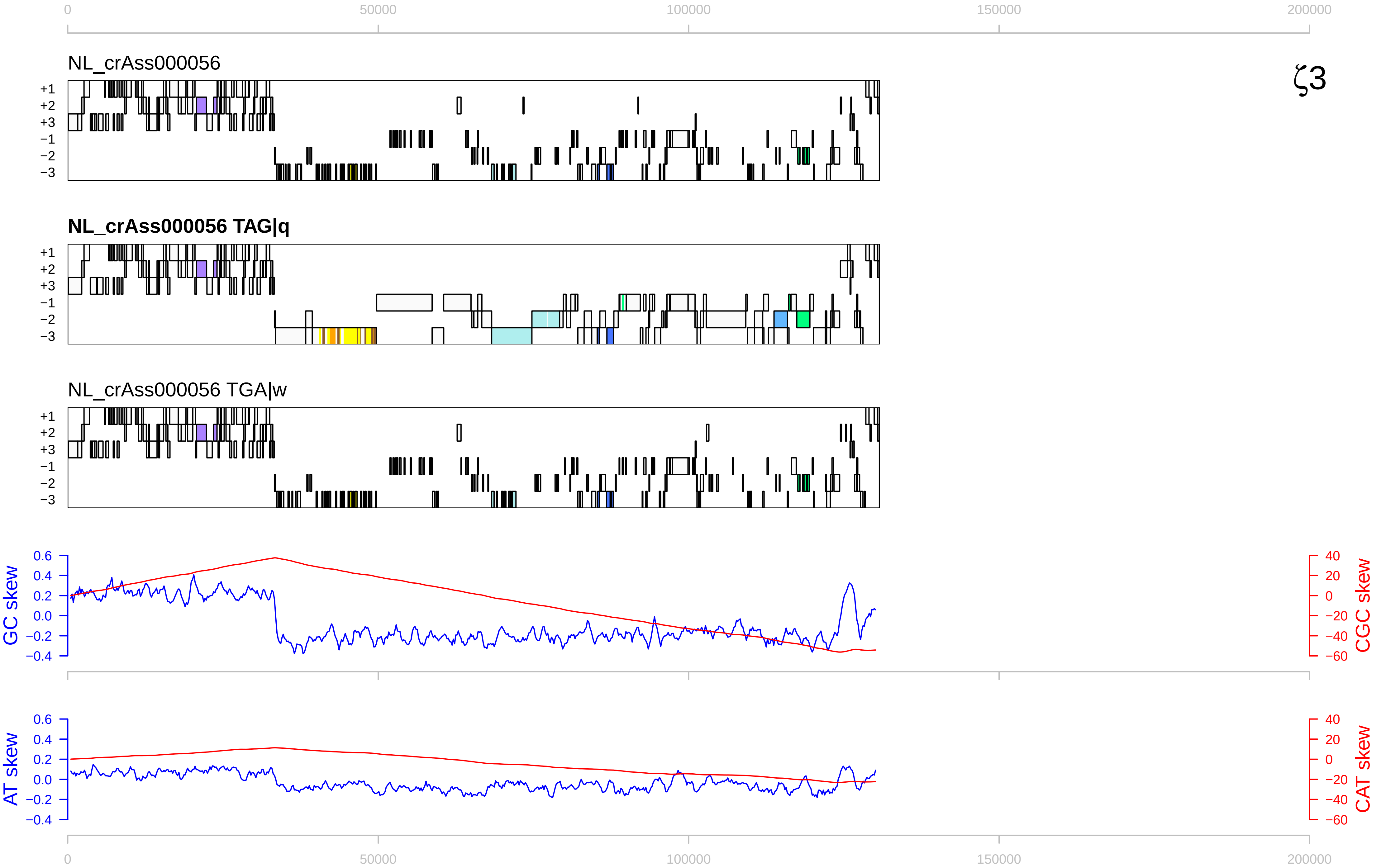

0
50000
100000
150000
200000
NL_crAss000056
ζ3
+1
+2
+3
−1
−2
−3
NL_crAss000056 TAG|q
NL_crAss000056 TGA|w
GC skew
CGC skew
AT skew
CAT skew
0.6
0.4
0.2
0.0
−0.2
−0.4
40
20
0
−20
−40
−60

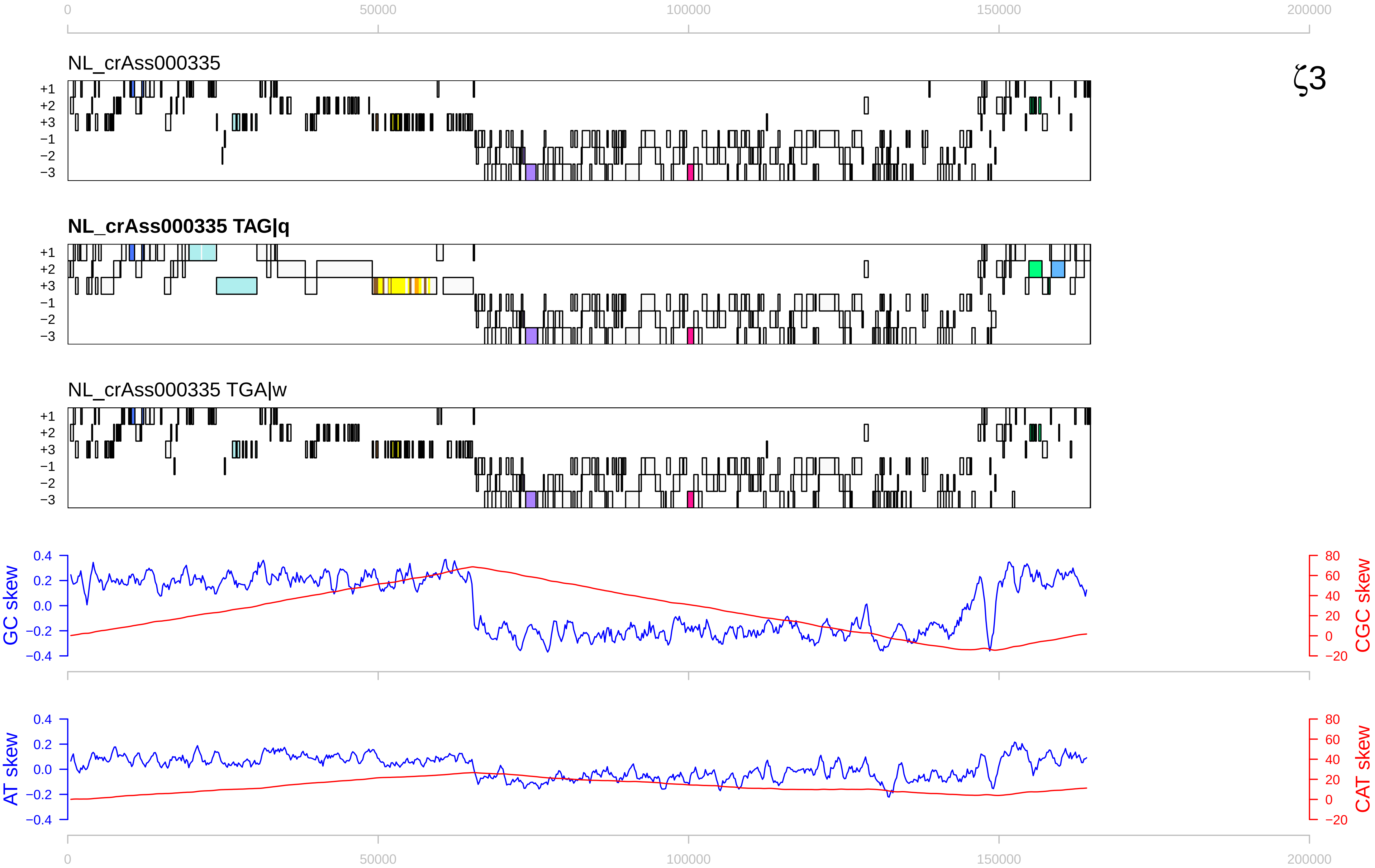

0
50000
100000
150000
200000
NL_crAss000335
ζ3
+1
+2
+3
−1
−2
−3
NL_crAss000335 TAG|q
NL_crAss000335 TGA|w
GC skew
CGC skew
AT skew
CAT skew
0.4
0.2
0.0
−0.2
−0.4
80
60
40
20
0
−20

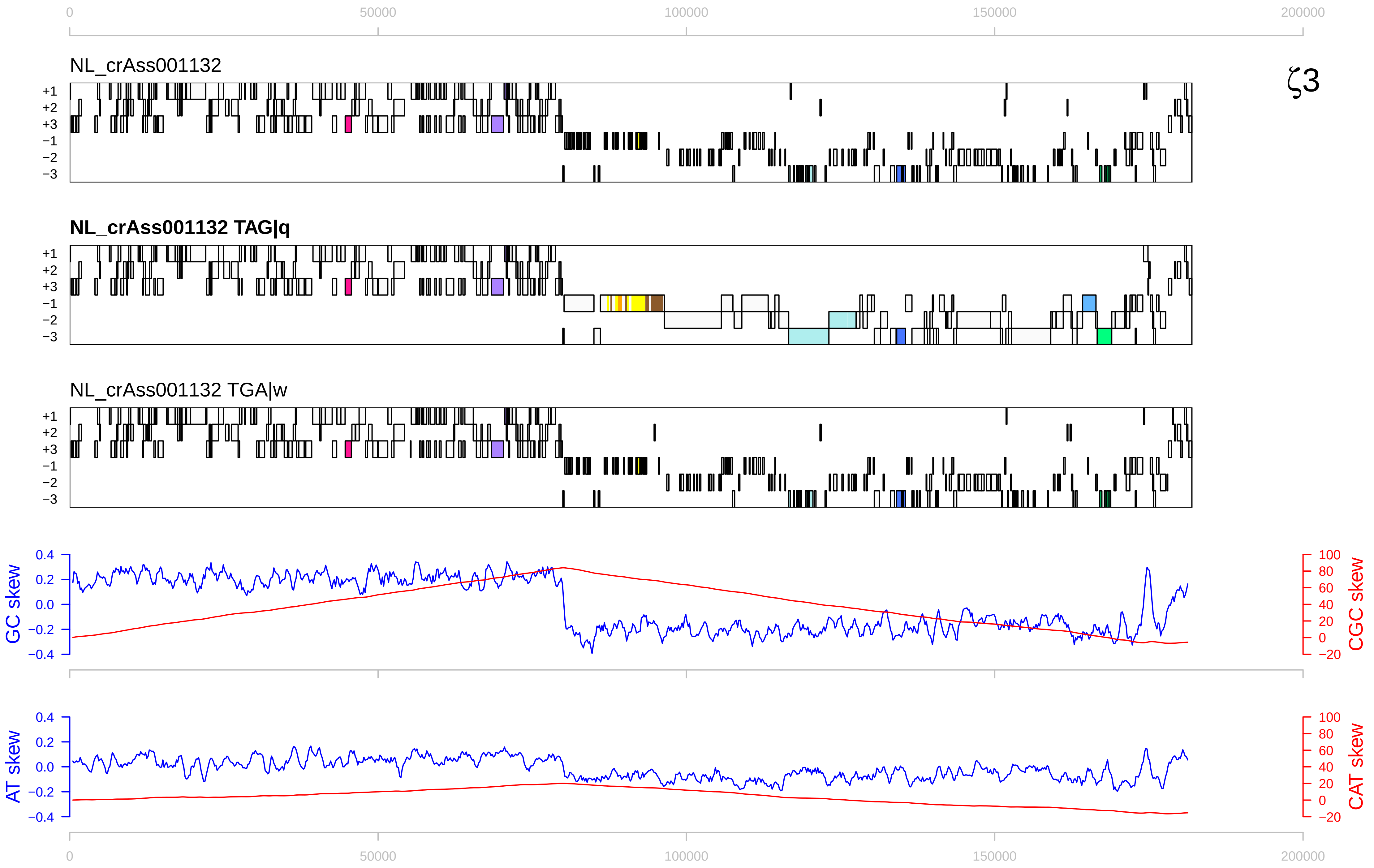

0
50000
100000
150000
200000
NL_crAss001132
ζ3
+1
+2
+3
−1
−2
−3
NL_crAss001132 TAG|q
NL_crAss001132 TGA|w
GC skew
CGC skew
AT skew
CAT skew
0.4
0.2
0.0
−0.2
−0.4
100
80
60
40
20
0
−20

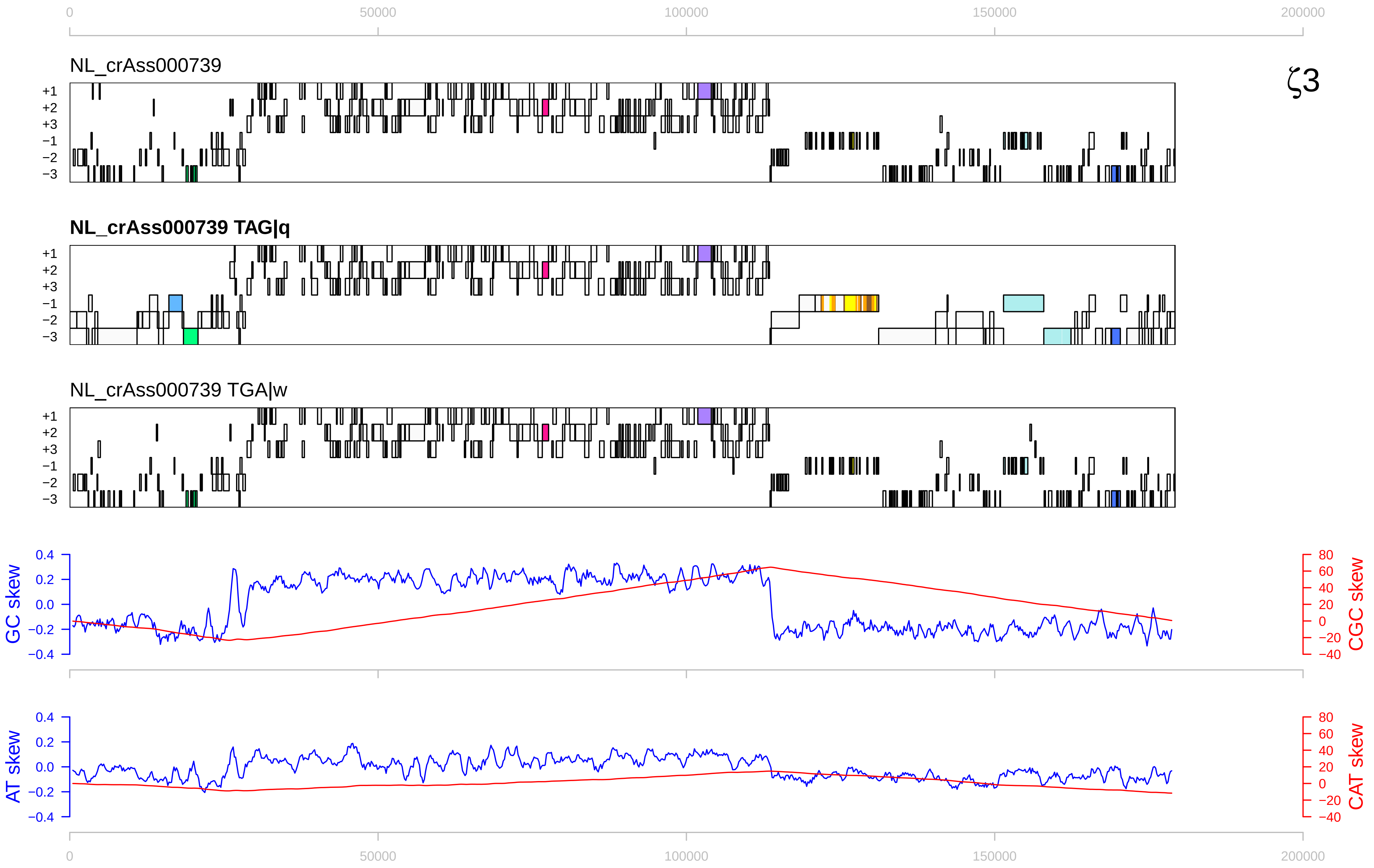
ζ3
NL_crAss000739
NL_crAss000739 TAG|q
NL_crAss000739 TGA|w
GC skew
CGC skew
AT skew
CAT skew
0
50000
100000
150000
200000

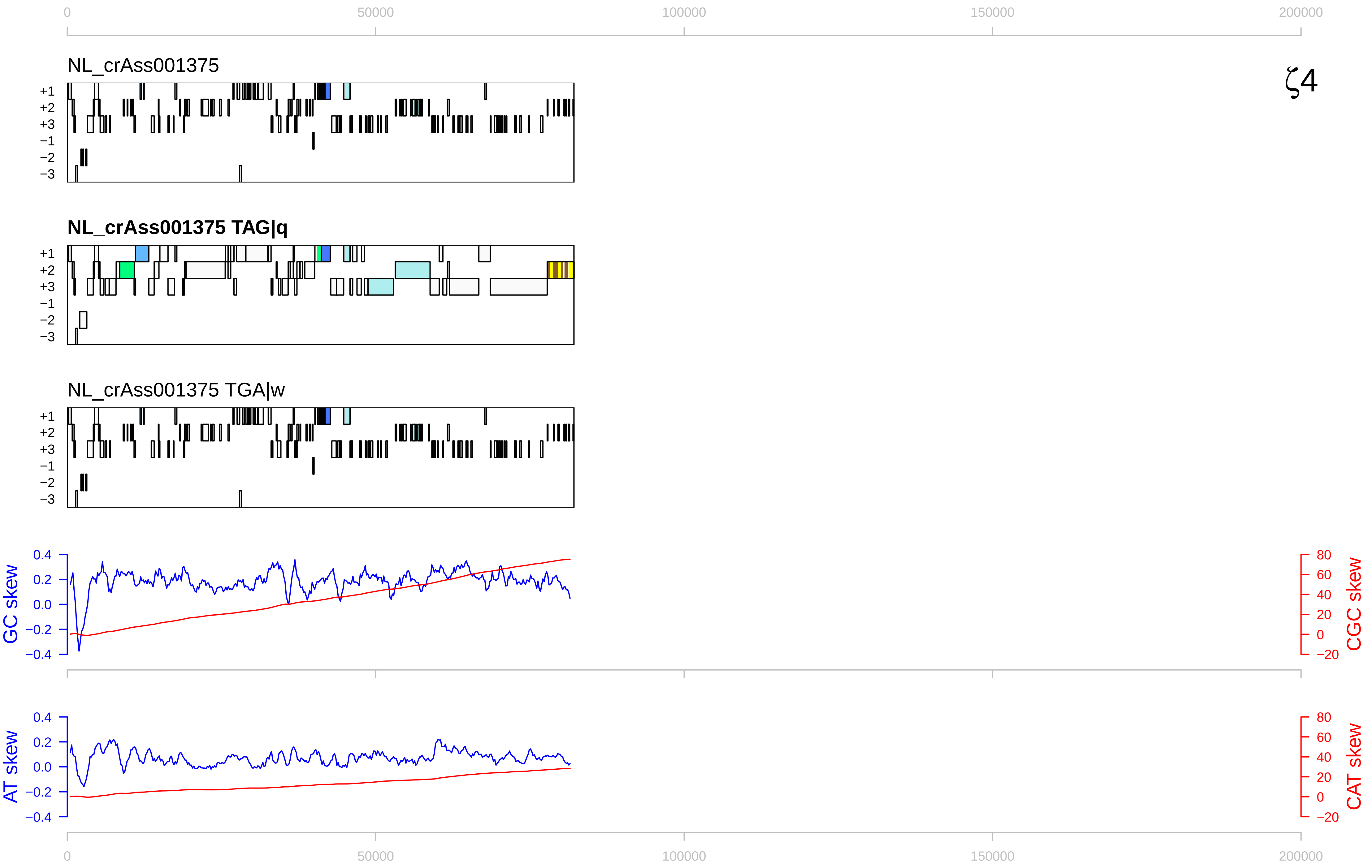

ζ4
NL_crAss001375
NL_crAss001375 TAG|q
NL_crAss001375 TGA|w
+1
+2
+3
−1
−2
−3
0
50000
100000
150000
200000
GC skew
CGC skew
AT skew
CAT skew
0.4
0.2
0.0
−0.2
−0.4
80
60
40
20
0
−20

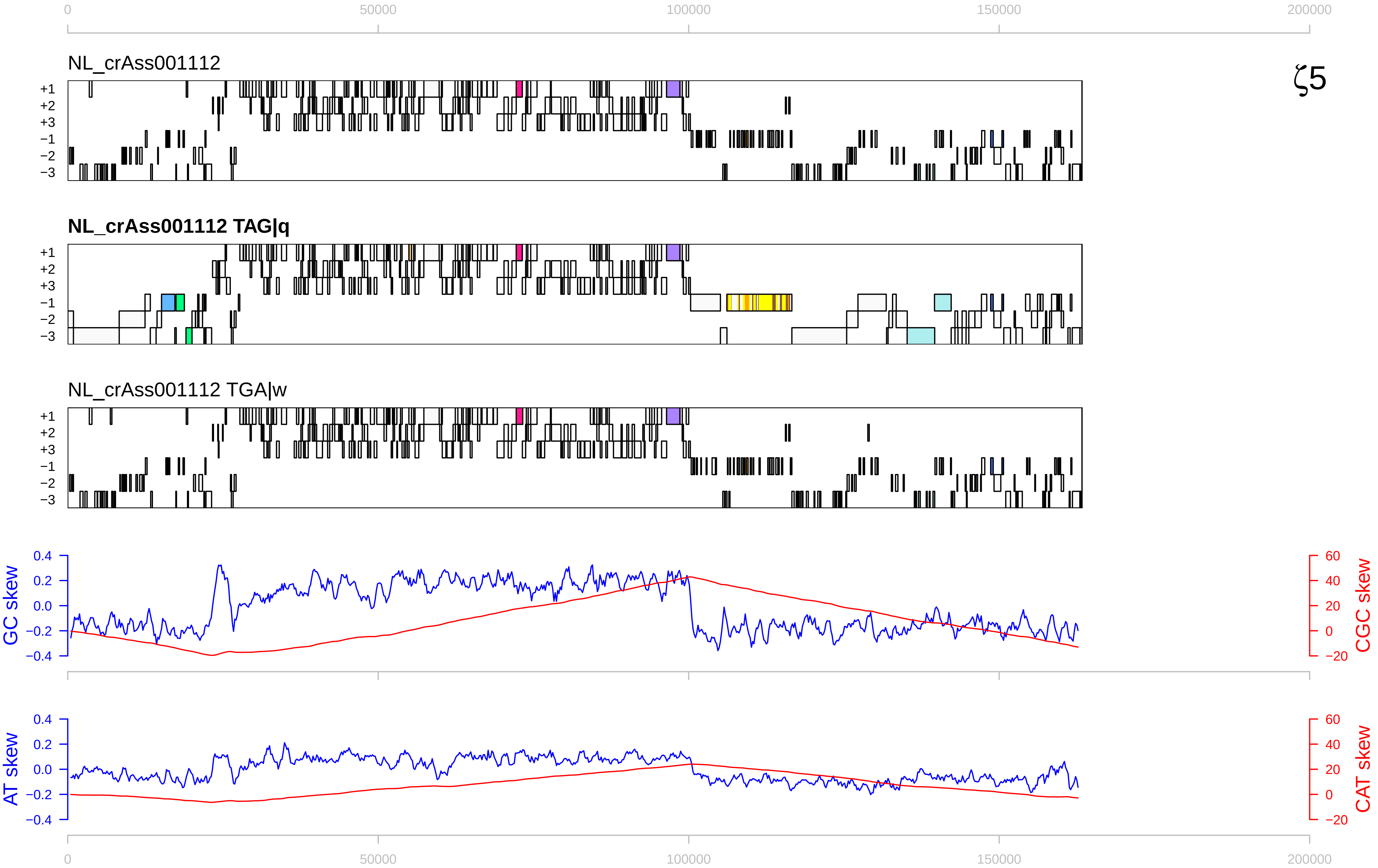

0
50000
100000
150000
200000
NL_crAss001112
ζ5
+1
+2
+3
−1
−2
−3
NL_crAss001112 TAG|q
NL_crAss001112 TGA|w
GC skew
CGC skew
AT skew
CAT skew
0.4
0.2
0.0
−0.2
−0.4
60
40
20
0
−20

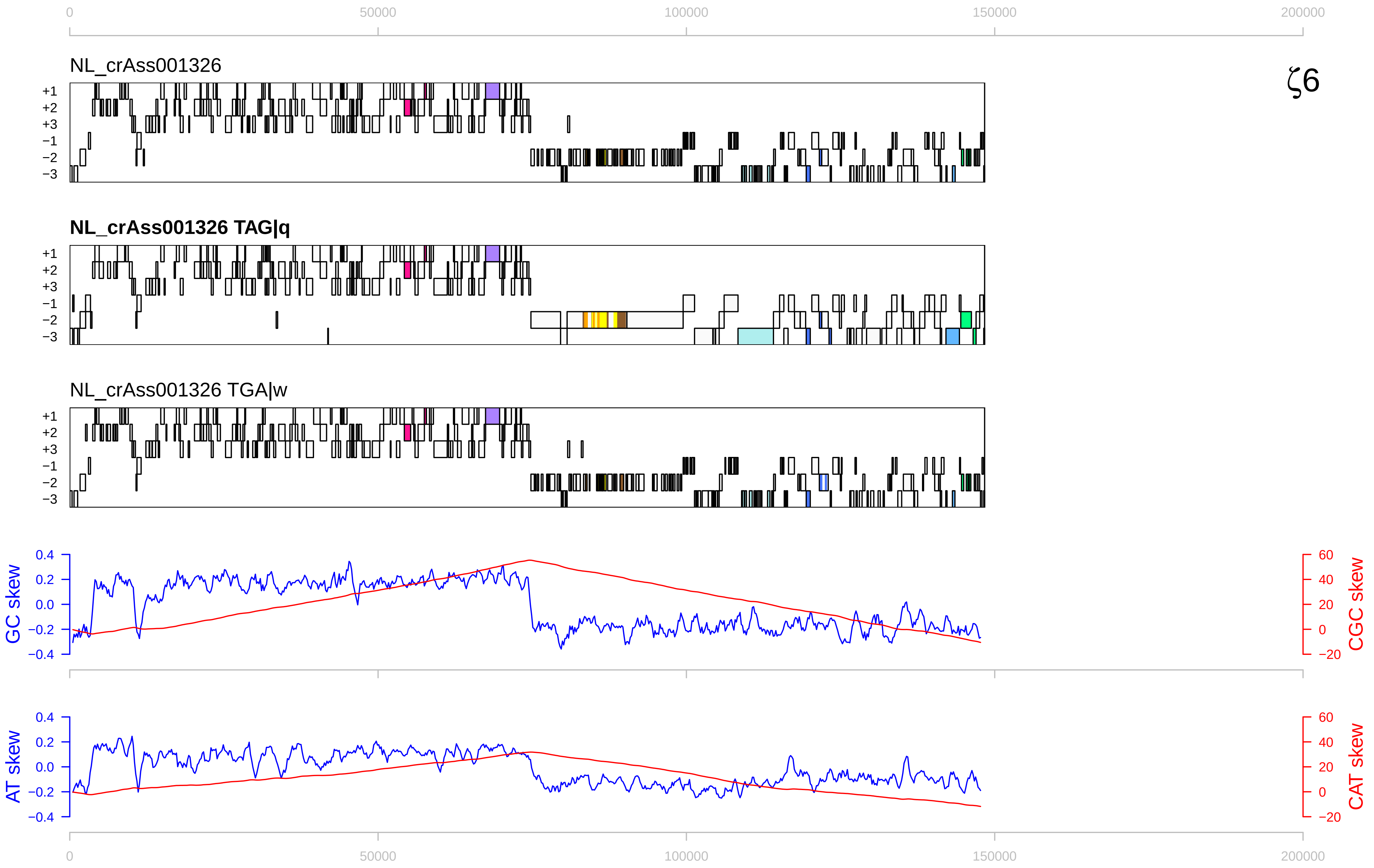

0
50000
100000
150000
200000
NL_crAss001326
ζ6
+1
+2
+3
−1
−2
−3
NL_crAss001326 TAG|q
NL_crAss001326 TGA|w
GC skew
CGC skew
0.4
0.2
0.0
−0.2
−0.4
60
40
20
0
−20
AT skew
CAT skew

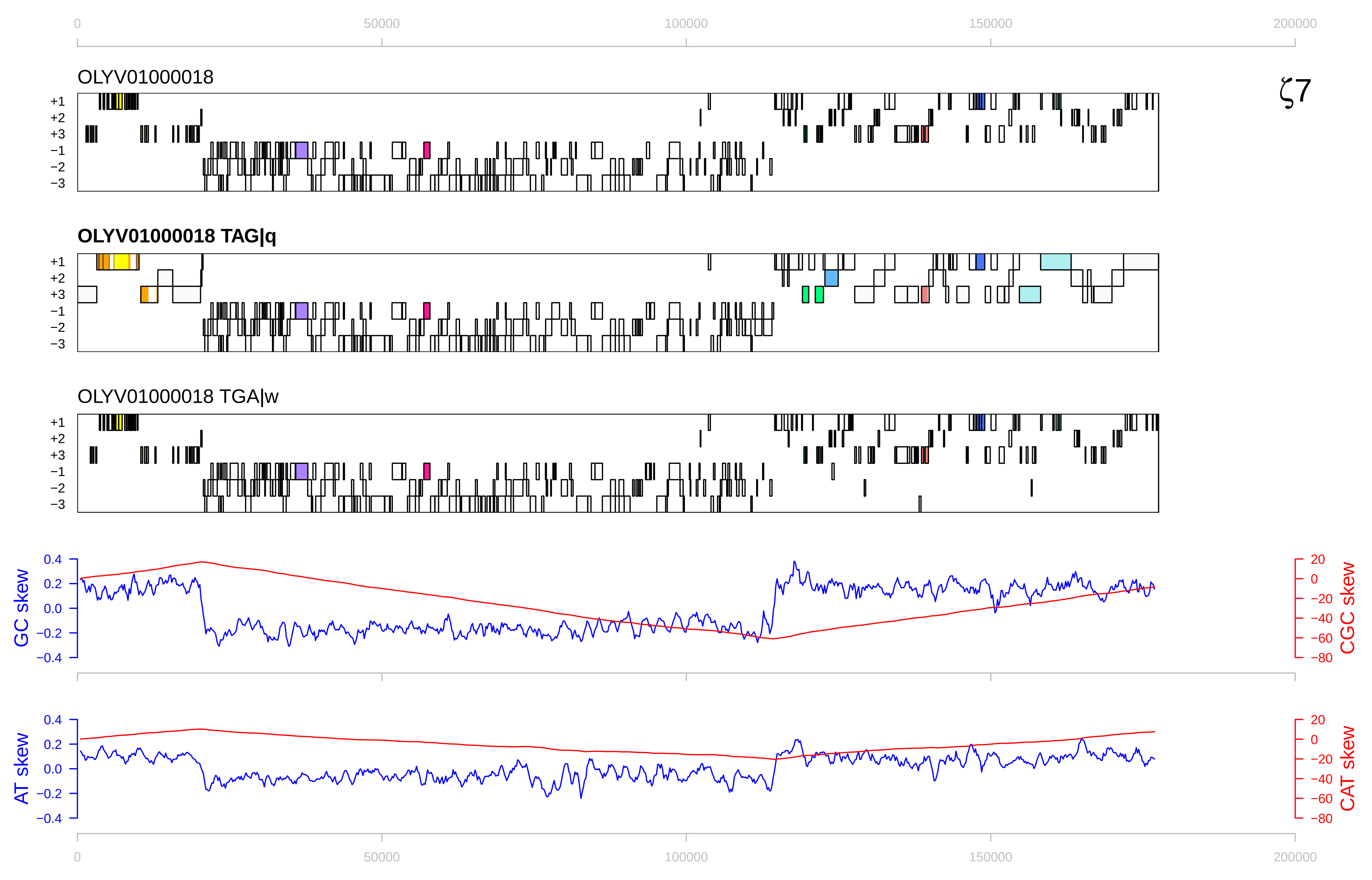

0
50000
100000
150000
200000
OLYV01000018
ζ7
+1
+2
+3
−1
−2
−3
OLYV01000018 TAG|q
OLYV01000018 TGA|w
GC skew
CGC skew
0.4
0.2
0.0
−0.2
−0.4
20
0
−20
−40
−60
−80
AT skew
CAT skew

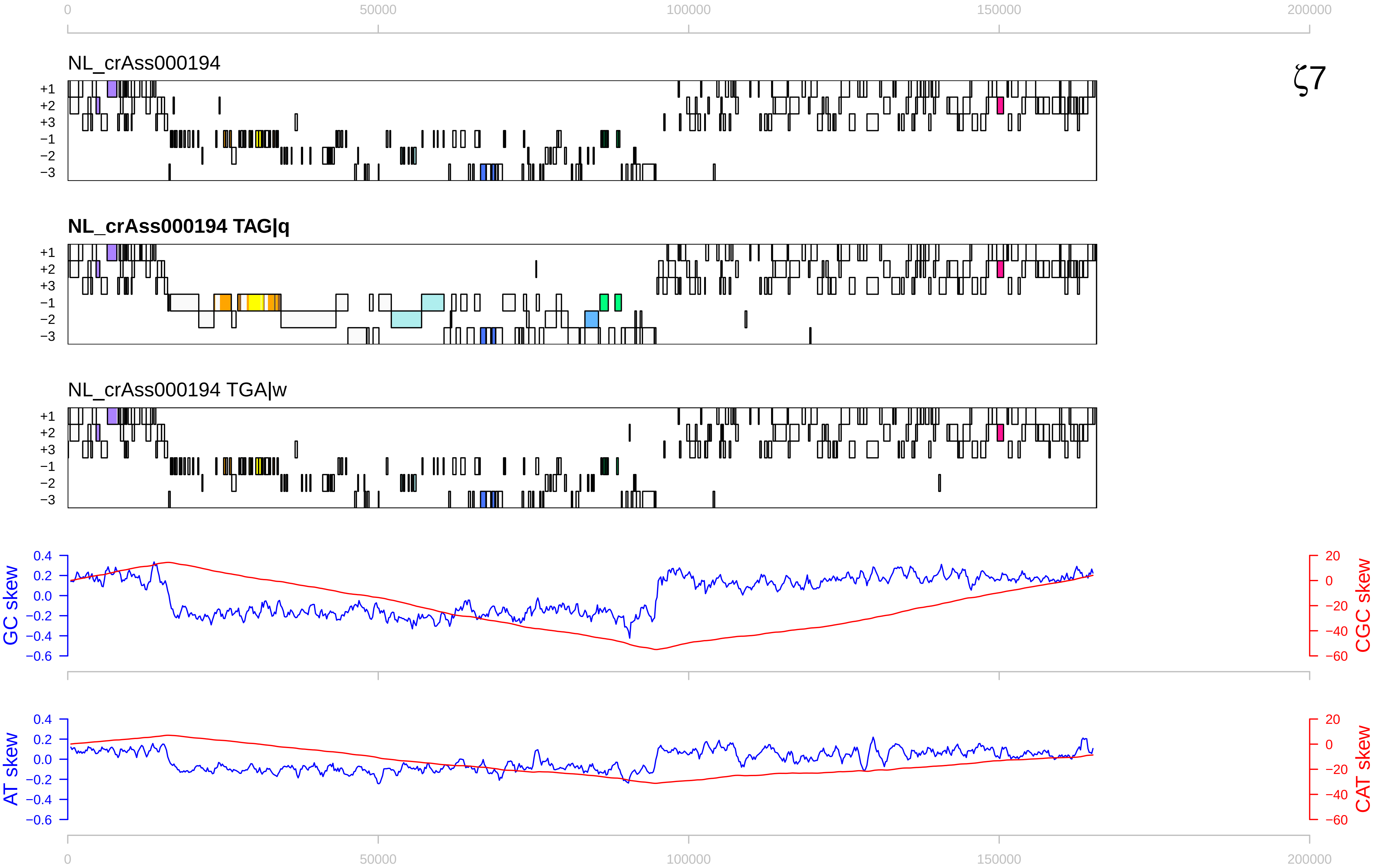

0
50000
100000
150000
200000
NL_crAss000194
ζ7
+1
+2
+3
−1
−2
−3
NL_crAss000194 TAG|q
NL_crAss000194 TGA|w
GC skew
CGC skew
0.4
0.2
0.0
−0.2
−0.4
−0.6
20
0
−20
−40
−60
AT skew
CAT skew

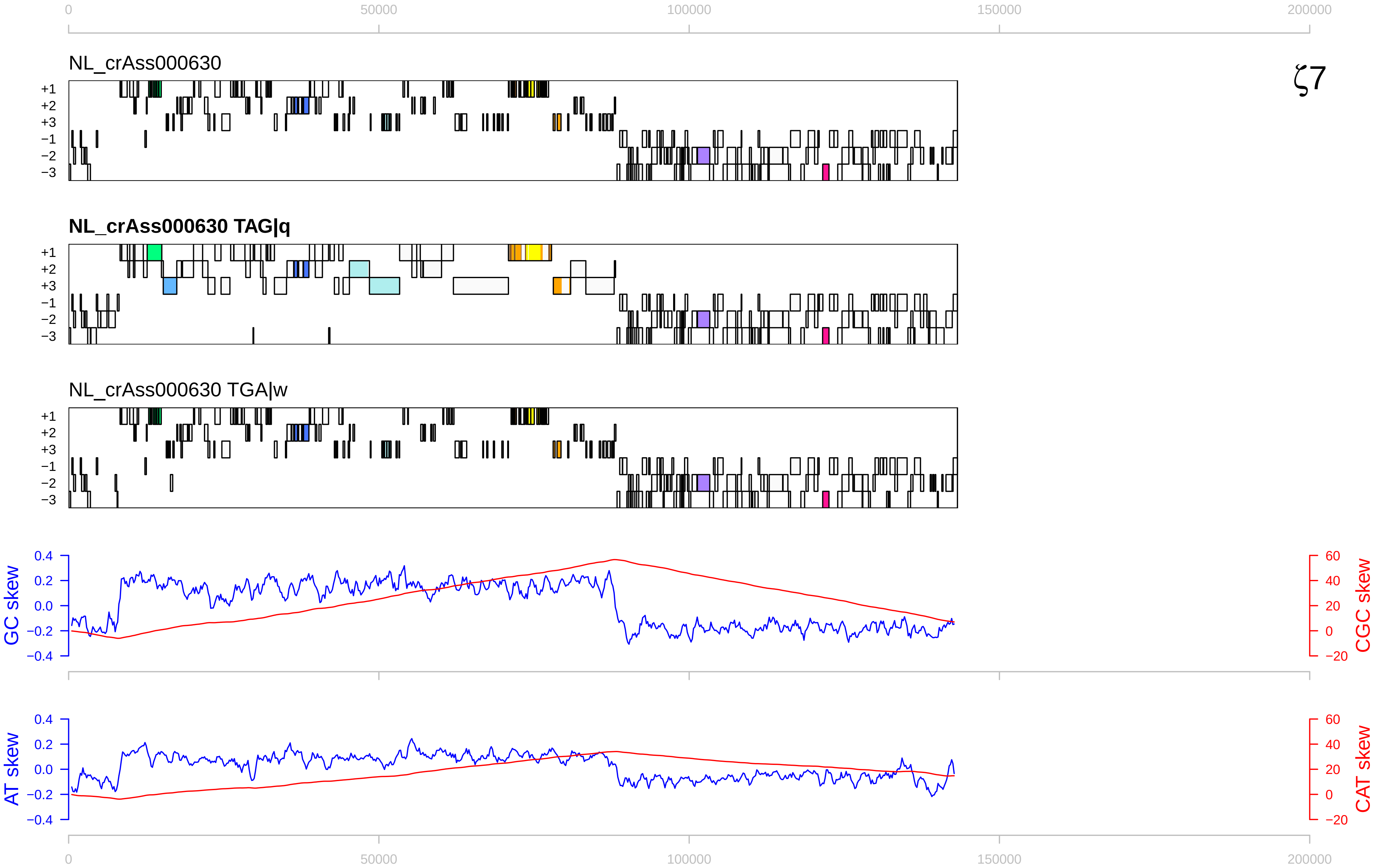
ζ7
0
50000
100000
150000
200000
NL_crAss000630
NL_crAss000630 TAG|q
NL_crAss000630 TGA|w
+1
+2
+3
−1
−2
−3
GC skew
CGC skew
AT skew
CAT skew
0.4
0.2
0.0
−0.2
−0.4
60
40
20
0
−20

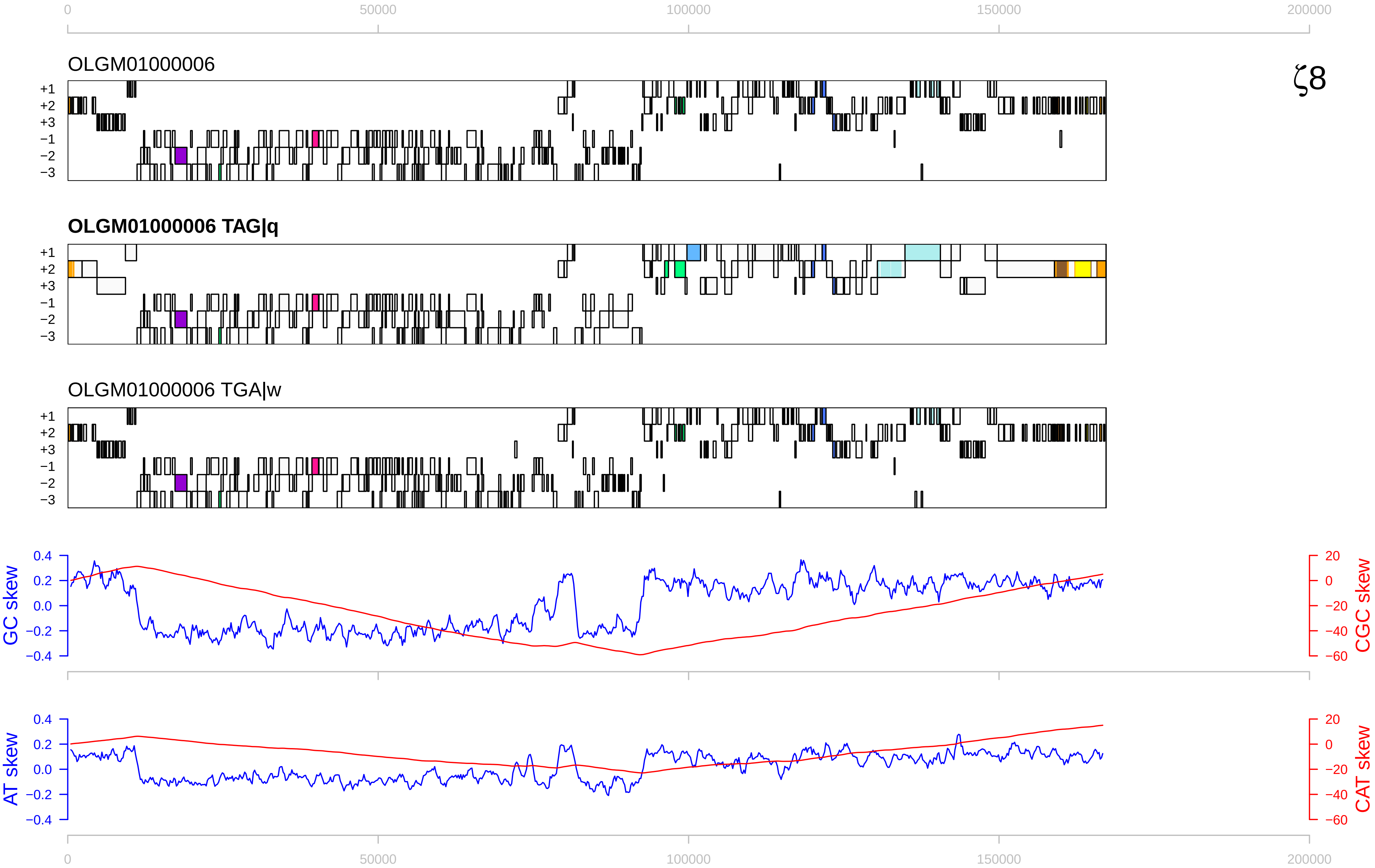

0
50000
100000
150000
200000
OLGM01000006
ζ8
+1
+2
+3
−1
−2
−3
OLGM01000006 TAG|q
OLGM01000006 TGA|w
GC skew
CGC skew
AT skew
CAT skew
0.4
0.2
0.0
−0.2
−0.4
20
0
−20
−40
−60

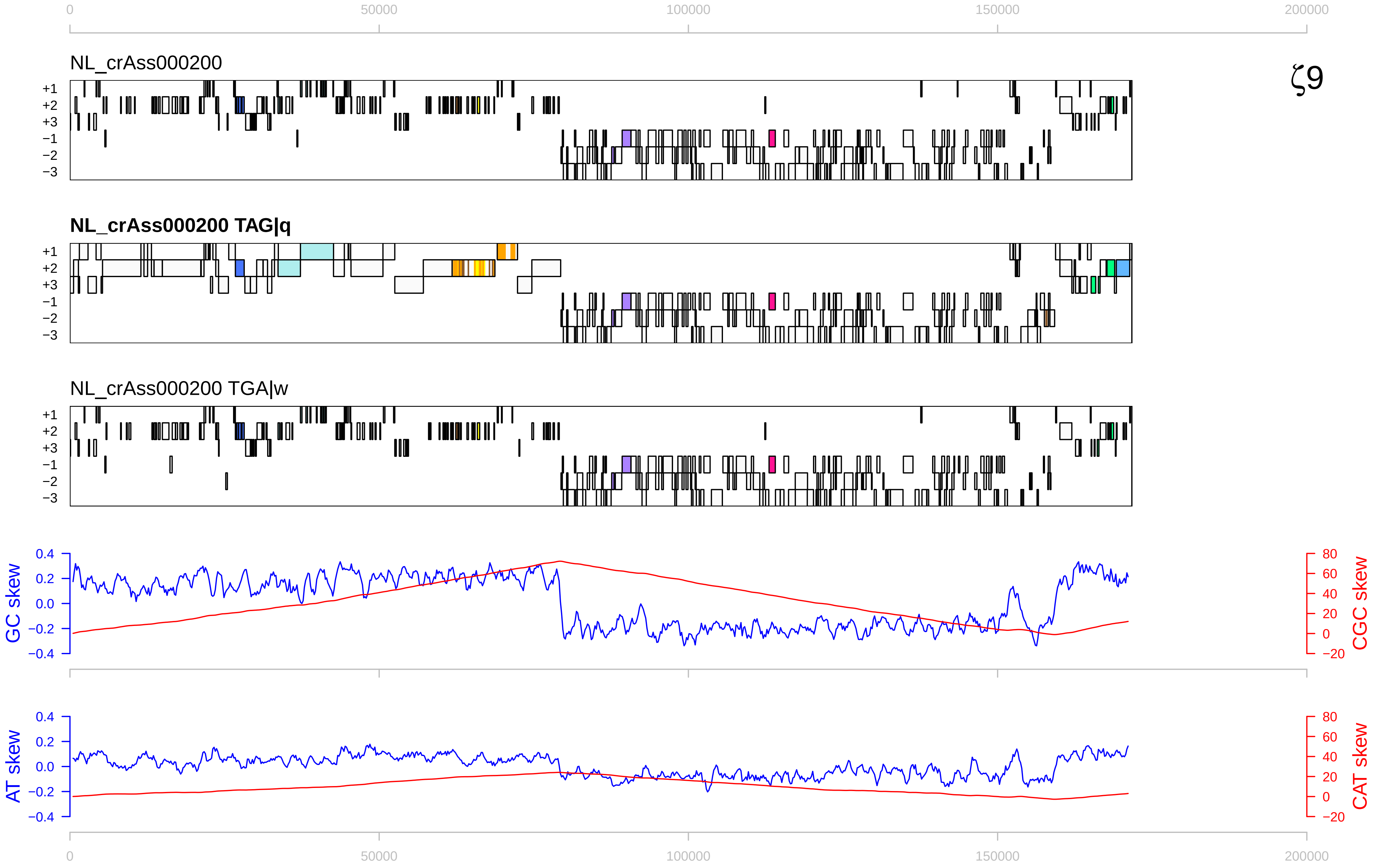

0
50000
100000
150000
200000
NL_crAss000200
ζ9
+1
+2
+3
−1
−2
−3
NL_crAss000200 TAG|q
NL_crAss000200 TGA|w
GC skew
CGC skew
0.4
0.2
0.0
−0.2
−0.4
80
60
40
20
0
−20
AT skew
CAT skew

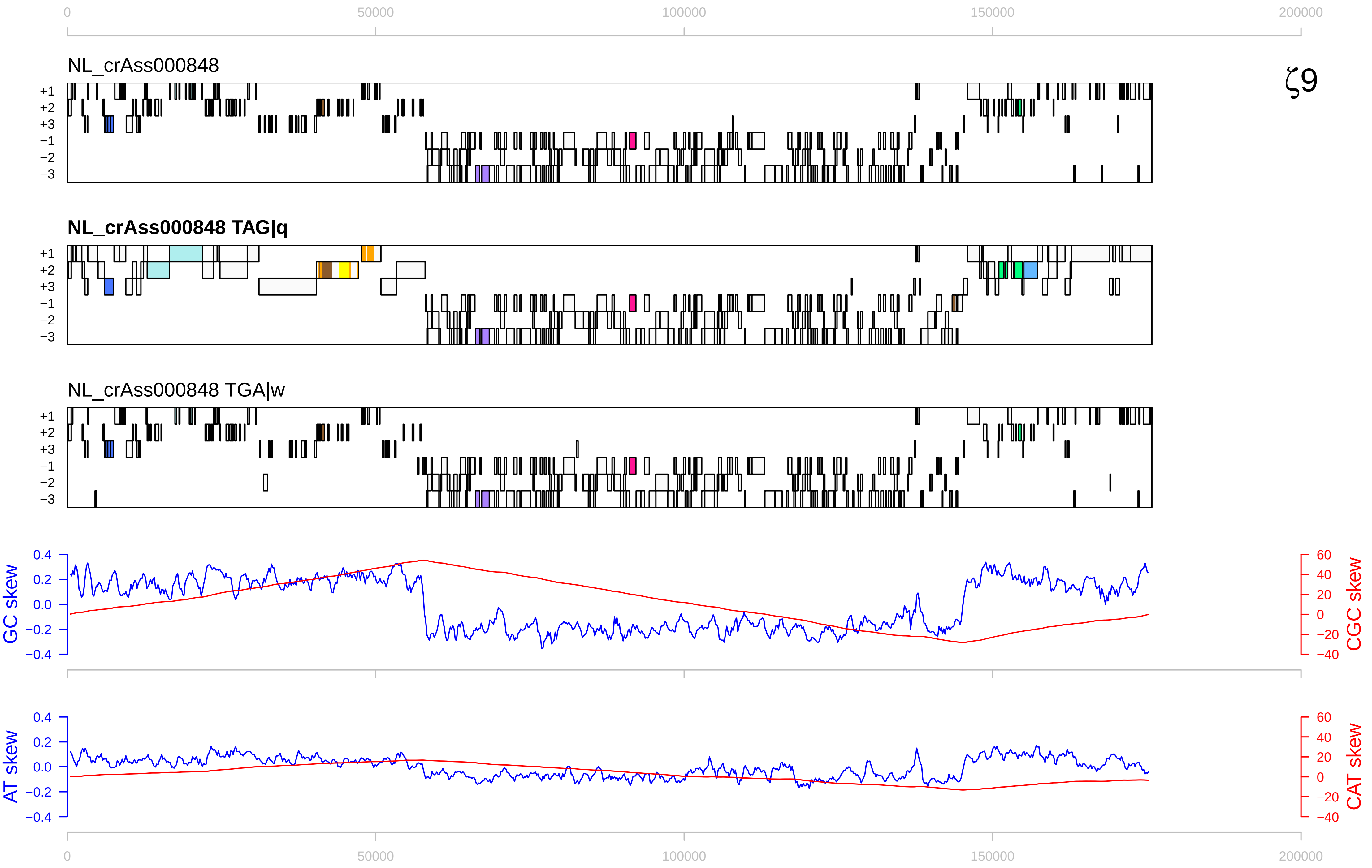

0
50000
100000
150000
200000
NL_crAss000848
ζ9
+1
+2
+3
−1
−2
−3
NL_crAss000848 TAG|q
NL_crAss000848 TGA|w
GC skew
CGC skew
AT skew
CAT skew
0.4
0.2
0.0
−0.2
−0.4
60
40
20
0
−20
−40

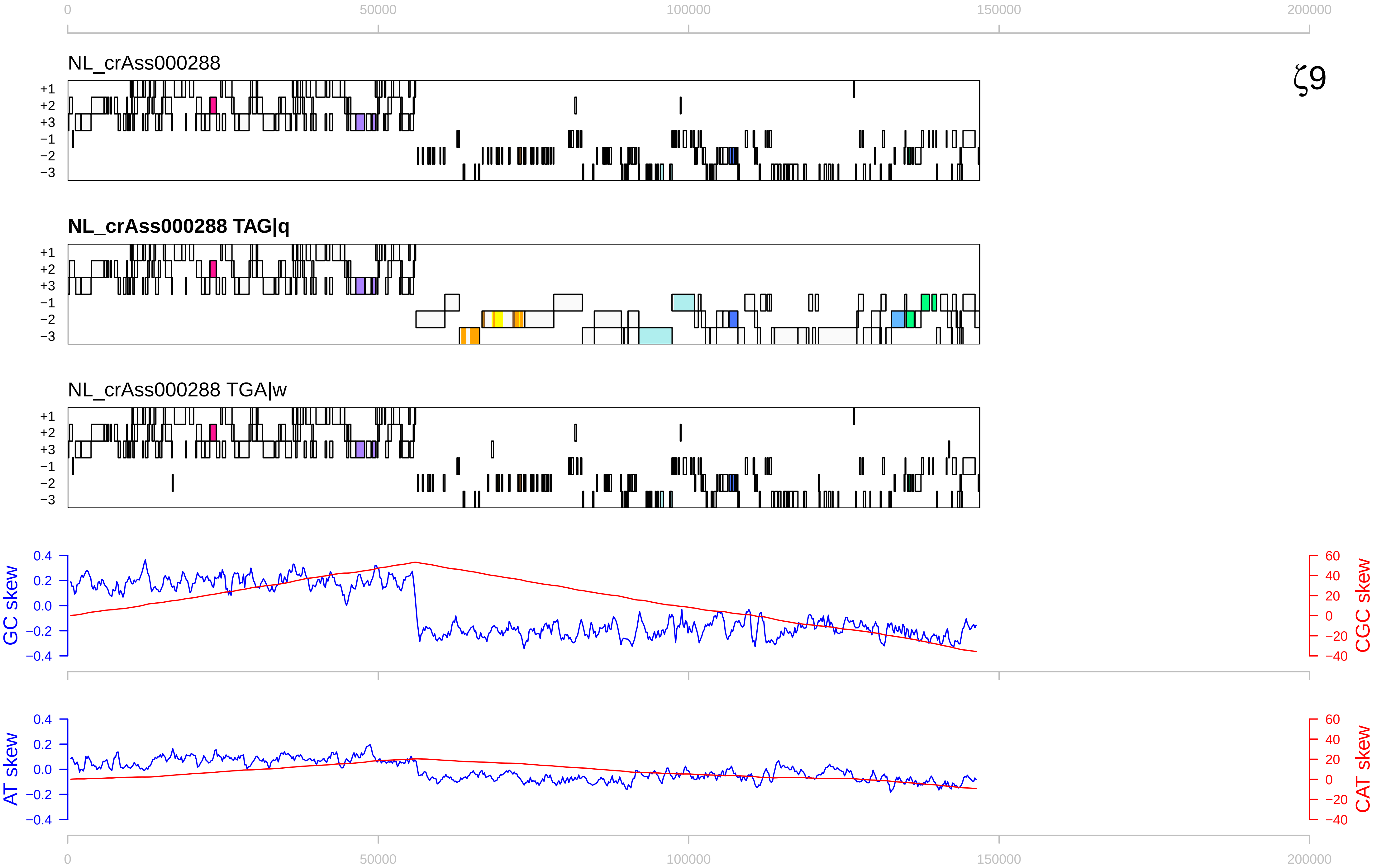

ζ9
NL_crAss000288
NL_crAss000288 TAG|q
NL_crAss000288 TGA|w
+1
+2
+3
−1
−2
−3
0
50000
100000
150000
200000
GC skew
CGC skew
AT skew
CAT skew
0.4
0.2
0.0
−0.2
−0.4
60
40
20
0
−20
−40

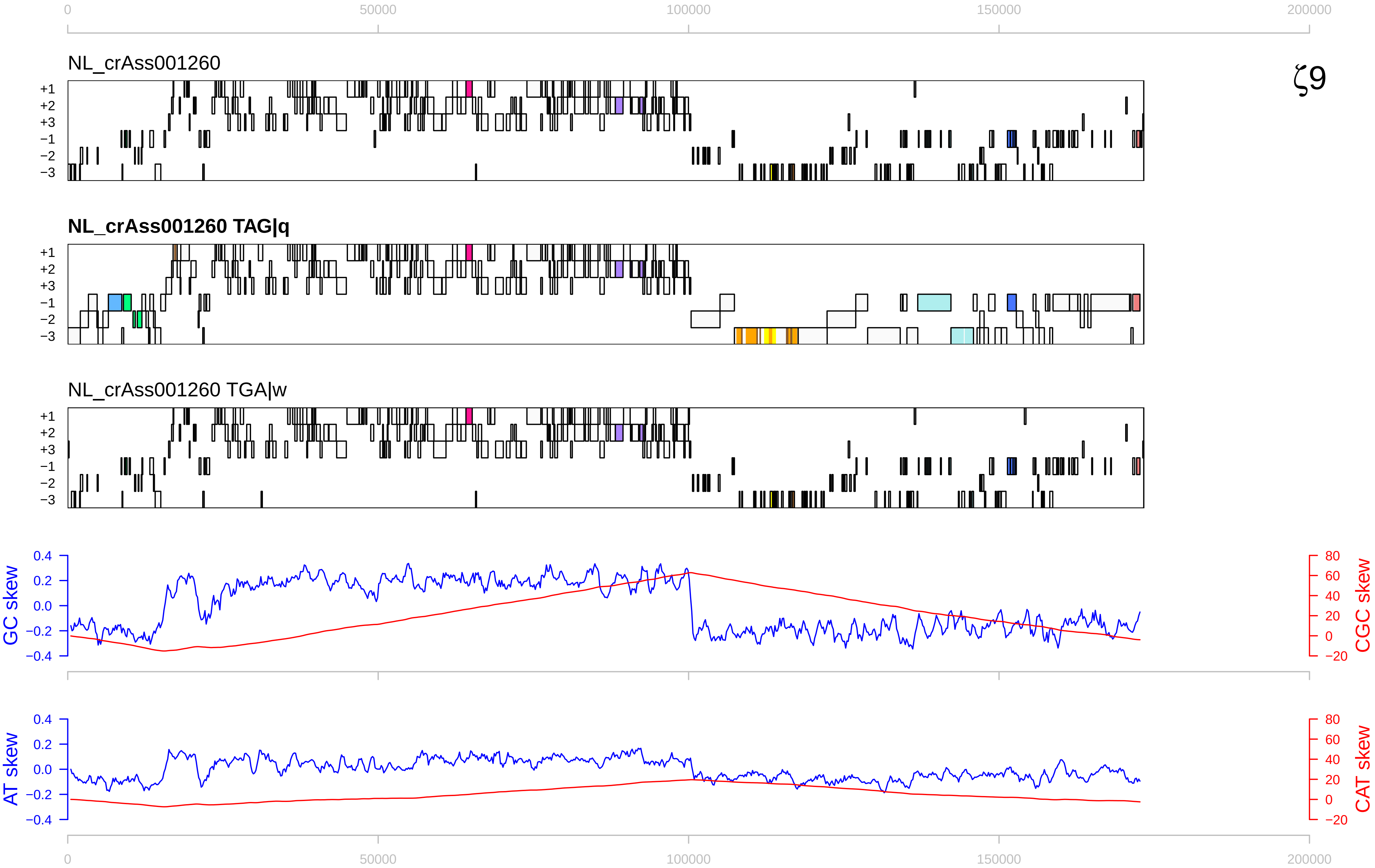

ζ9
NL_crAss001260
NL_crAss001260 TAG|q
NL_crAss001260 TGA|w
+1
+2
+3
−1
−2
−3
0
50000
100000
150000
200000
GC skew
CGC skew
AT skew
CAT skew
0.4
0.2
0.0
−0.2
−0.4
80
60
40
20
0
−20

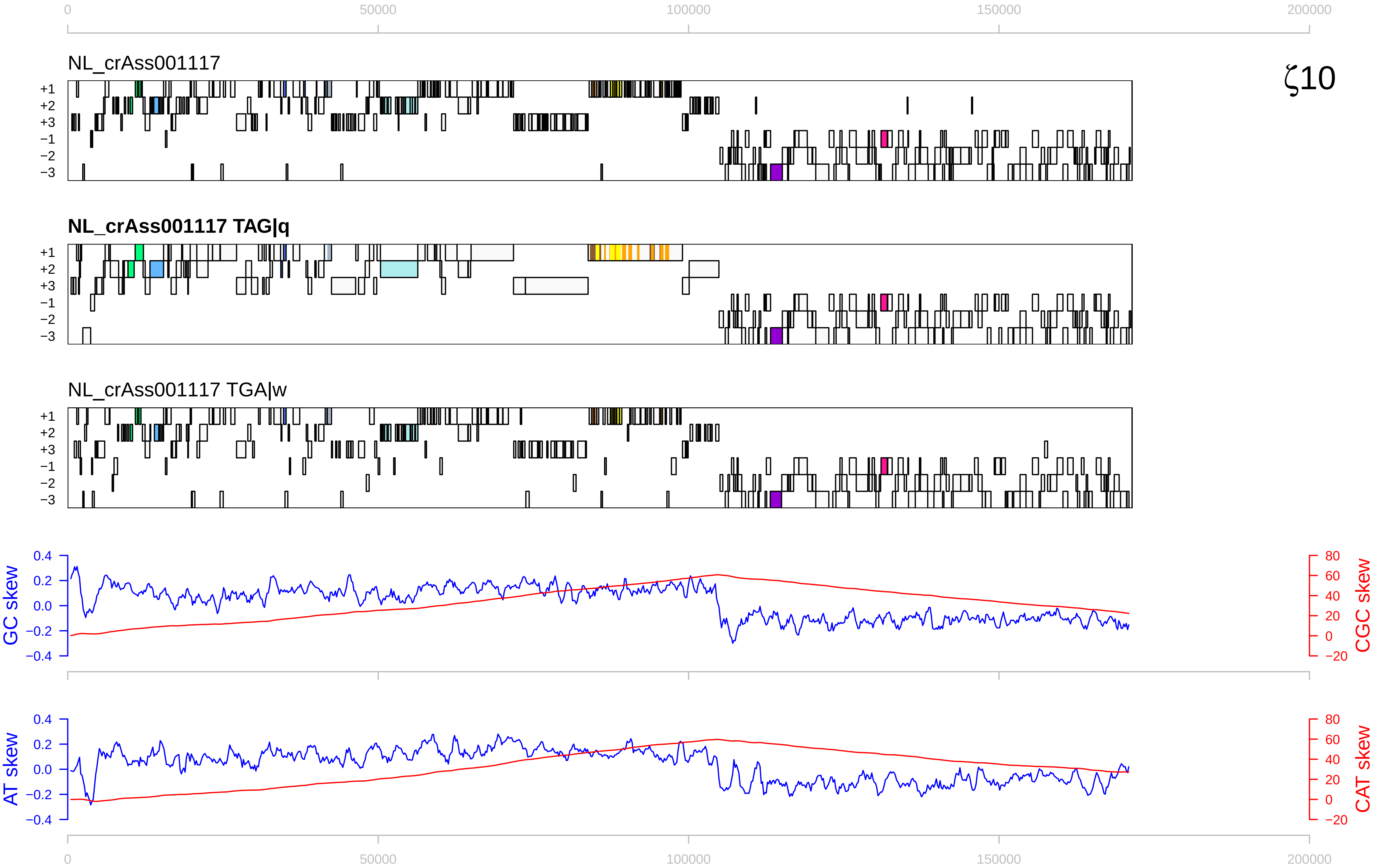

0
50000
100000
150000
200000
NL_crAss001117
ζ10
NL_crAss001117 TAG|q
NL_crAss001117 TGA|w
+1
+2
+3
−1
−2
−3
GC skew
CGC skew
AT skew
CAT skew
0.4
0.2
0.0
−0.2
−0.4
80
60
40
20
0
−20

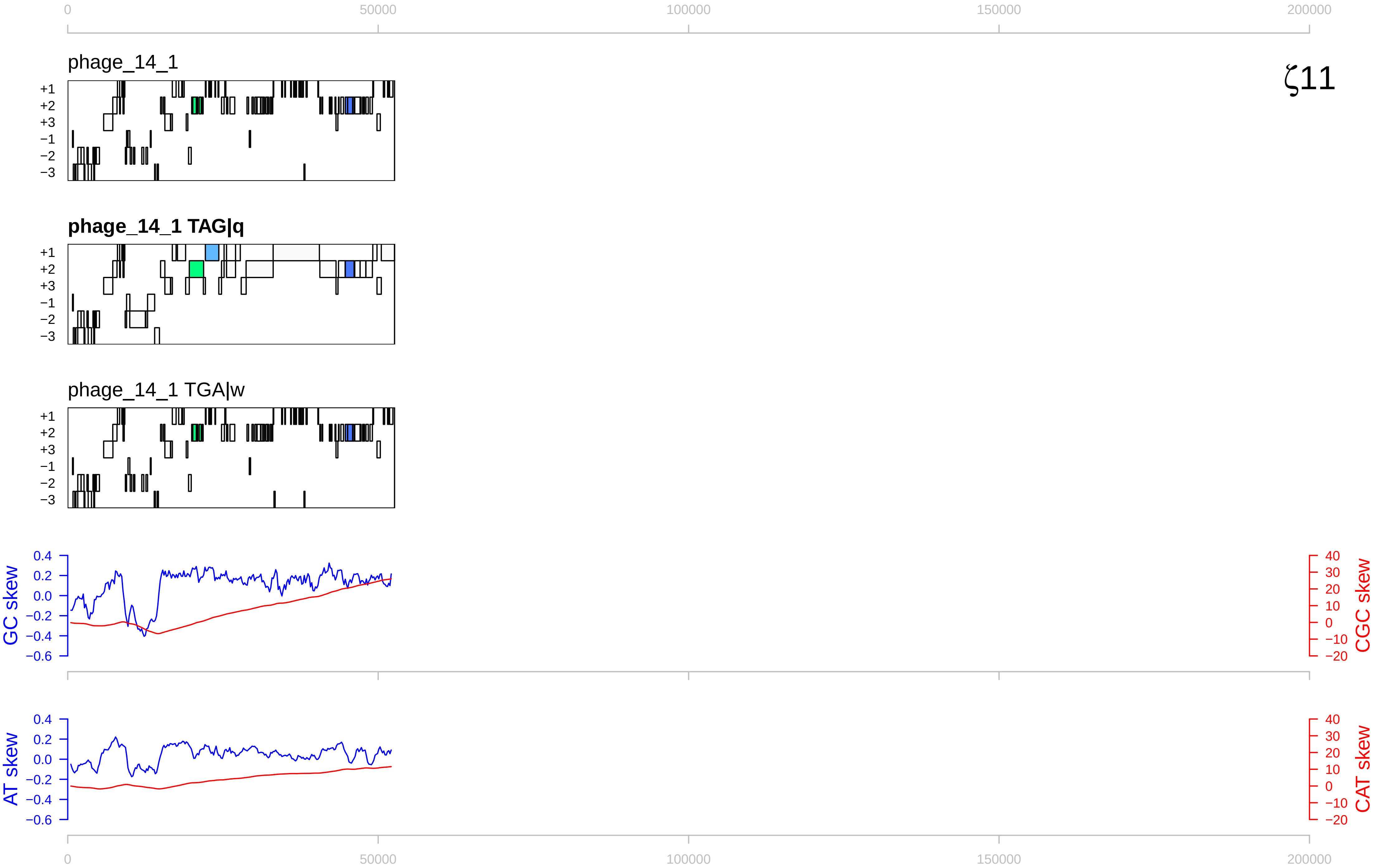

ζ11
phage_14_1
phage_14_1 TAG|q
phage_14_1 TGA|w
GC skew
CGC skew
AT skew
CAT skew
0
50000
100000
150000
200000

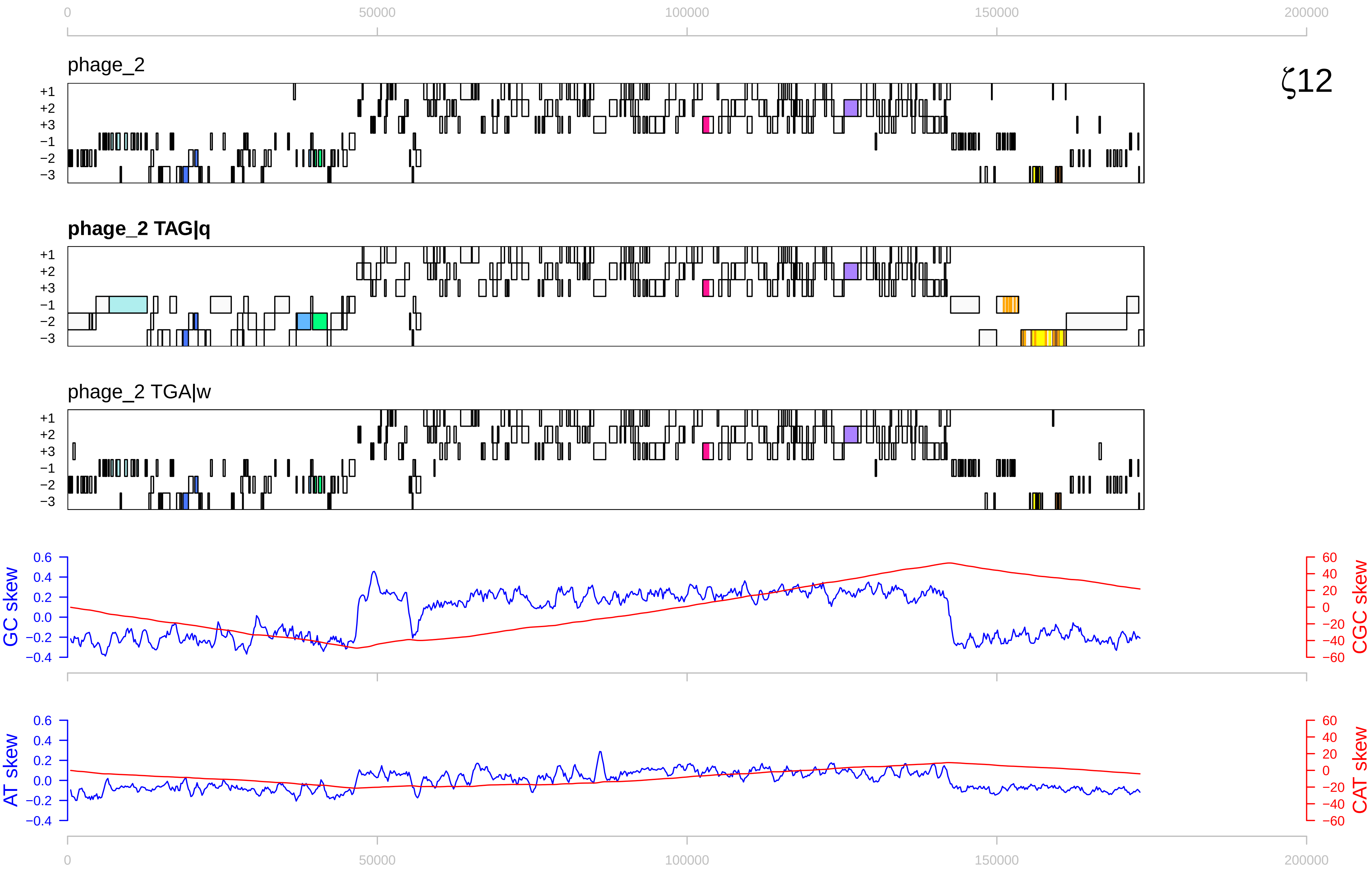

ζ12
0
50000
100000
150000
200000
phage_2
phage_2 TAG|q
phage_2 TGA|w
+1
+2
+3
−1
−2
−3
GC skew
CGC skew
AT skew
CAT skew
0.6
0.4
0.2
0.0
−0.2
−0.4
60
40
20
0
−20
−40
−60

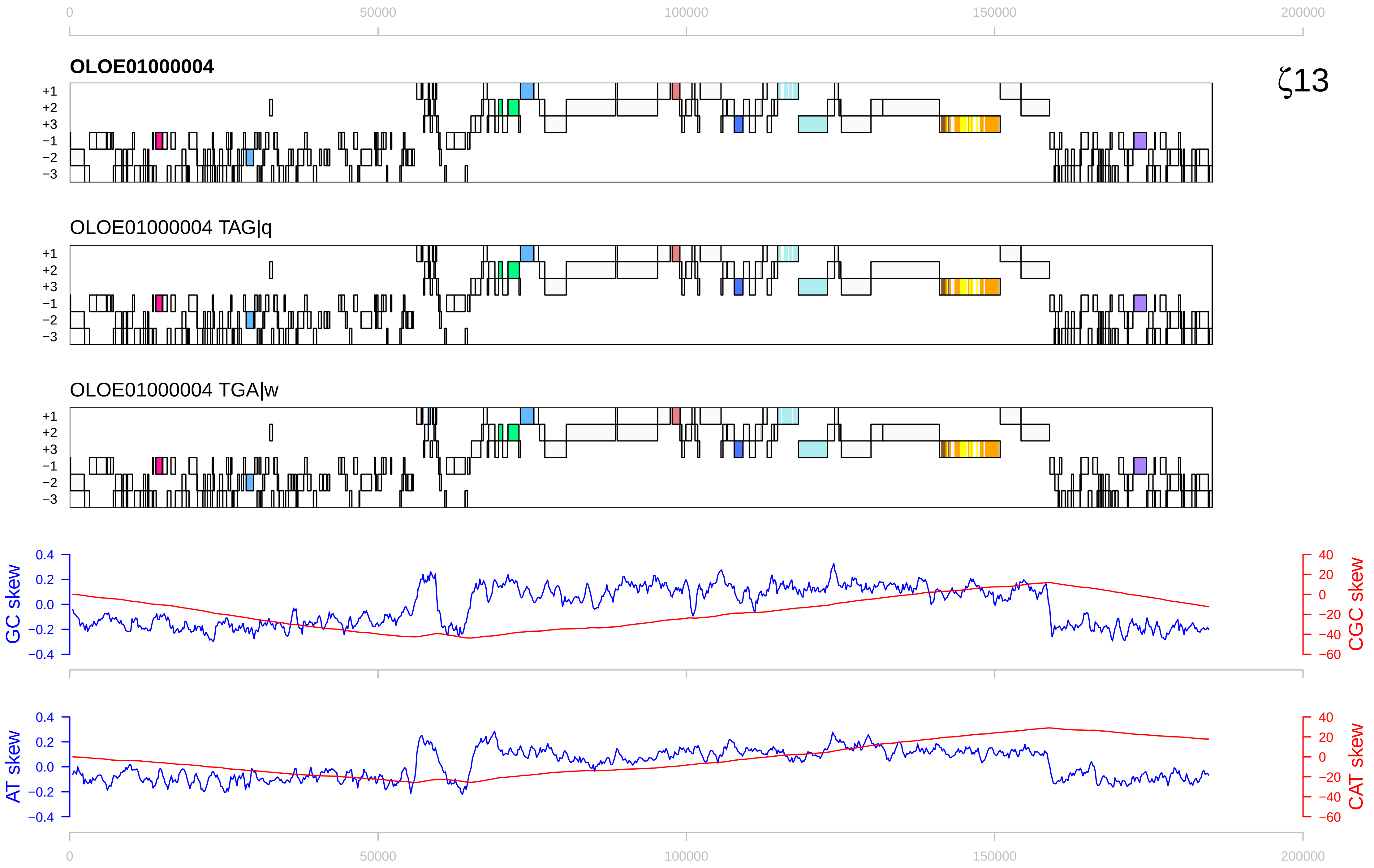

ζ13
OLOE01000004
OLOE01000004 TAG|q
OLOE01000004 TGA|w
+1
+2
+3
−1
−2
−3
GC skew
CGC skew
AT skew
CAT skew
0
50000
100000
150000
200000

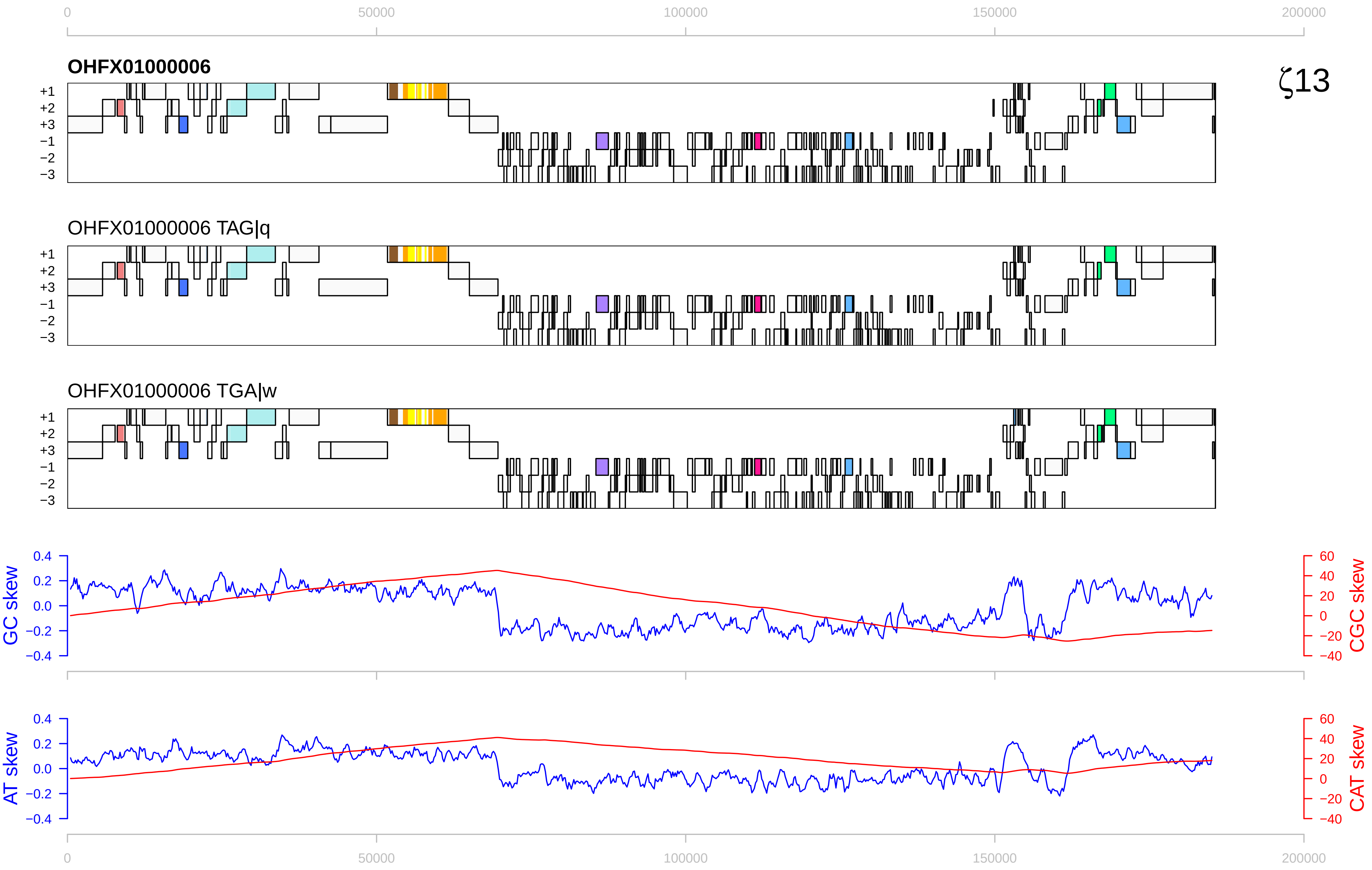

0
50000
100000
150000
200000
OHFX01000006
ζ13
+1
+2
+3
−1
−2
−3
OHFX01000006 TAG|q
OHFX01000006 TGA|w
GC skew
CGC skew
AT skew
CAT skew
0.4
0.2
0.0
−0.2
−0.4
60
40
20
0
−20
−40

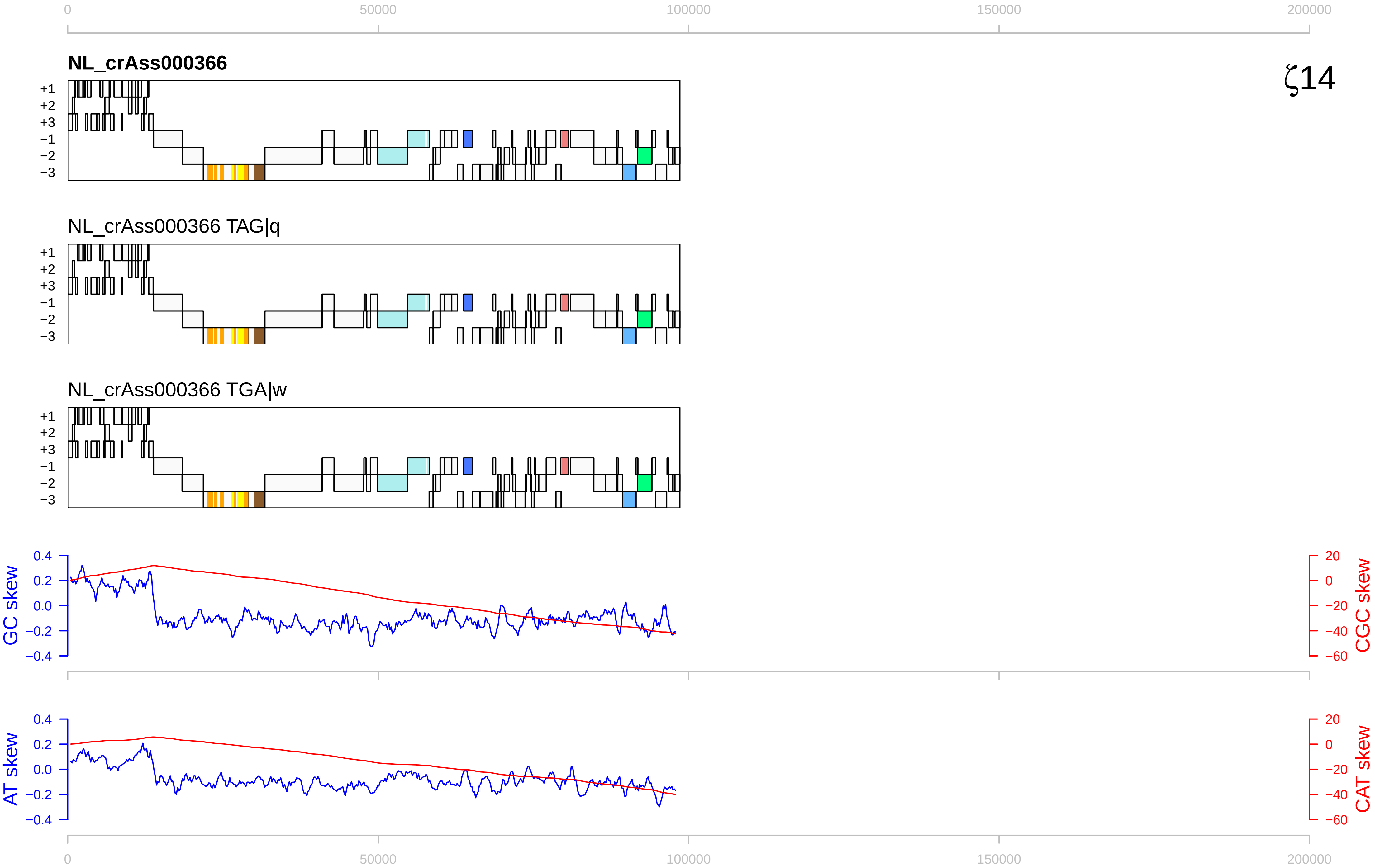

ζ14
NL_crAss000366
NL_crAss000366 TAG|q
NL_crAss000366 TGA|w
+1
+2
+3
−1
−2
−3
GC skew
CGC skew
AT skew
CAT skew
0
50000
100000
150000
200000
0.4
0.2
0.0
−0.2
−0.4
20
0
−20
−40
−60

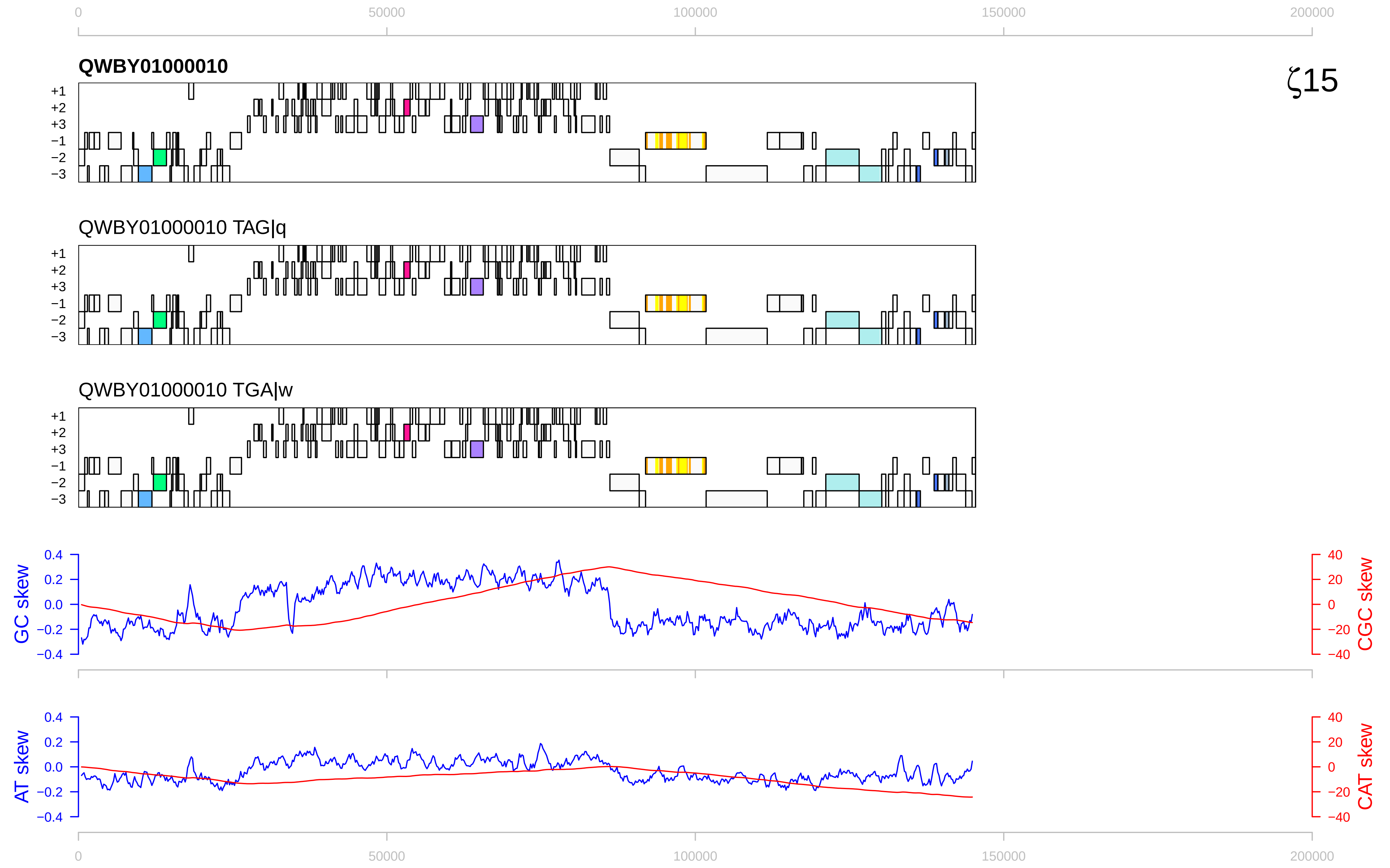
QWBY01000010
QWBY01000010 TAG|q
QWBY01000010 TGA|w
ζ15
GC skew
CGC skew
AT skew
CAT skew
0
50000
100000
150000
200000

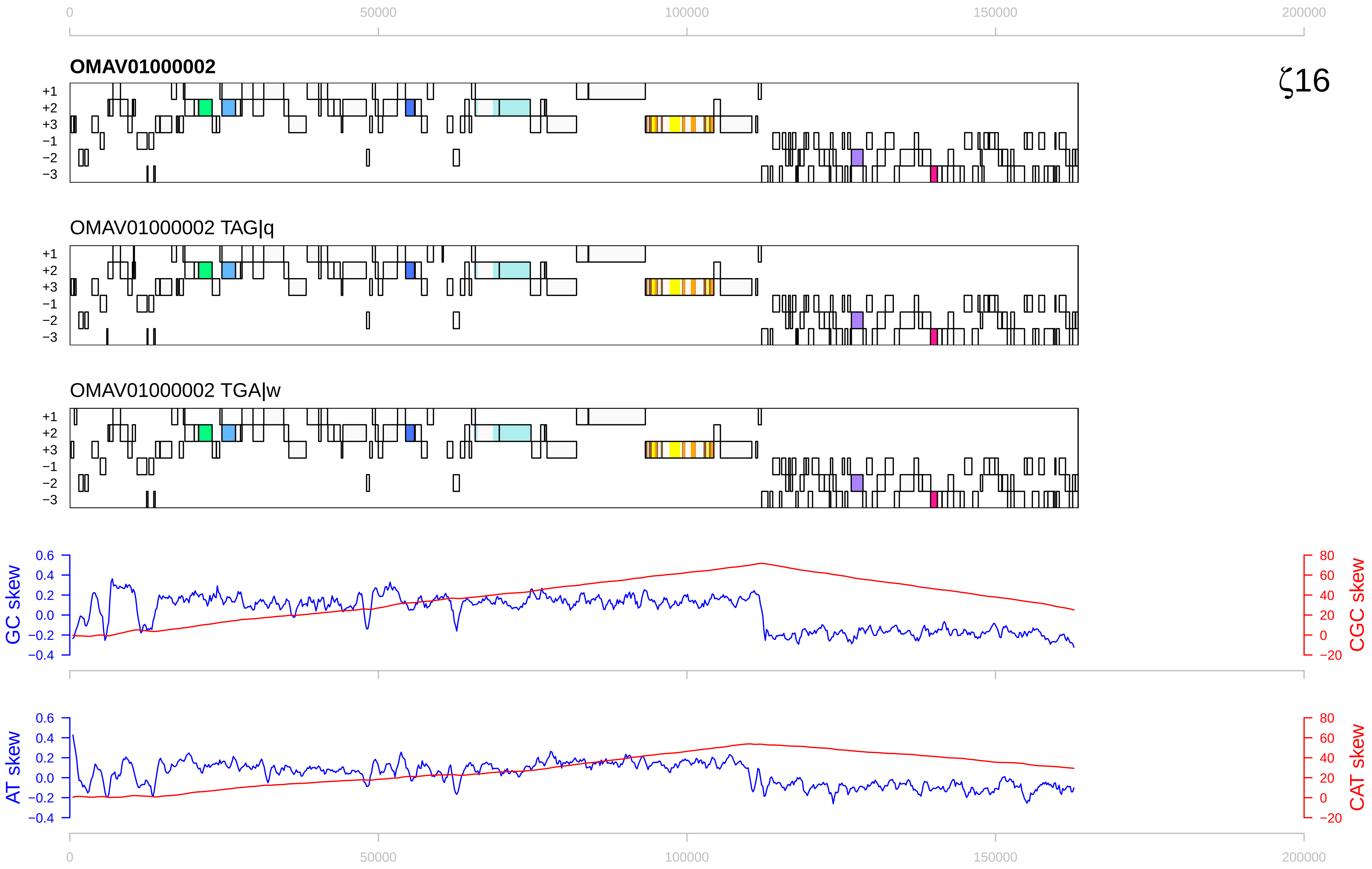

0
50000
100000
150000
200000
OMAV01000002
ζ16
+1
+2
+3
−1
−2
−3
OMAV01000002 TAG|q
OMAV01000002 TGA|w
GC skew
CGC skew
AT skew
CAT skew
0.6
0.4
0.2
0.0
−0.2
−0.4
80
60
40
20
0
−20

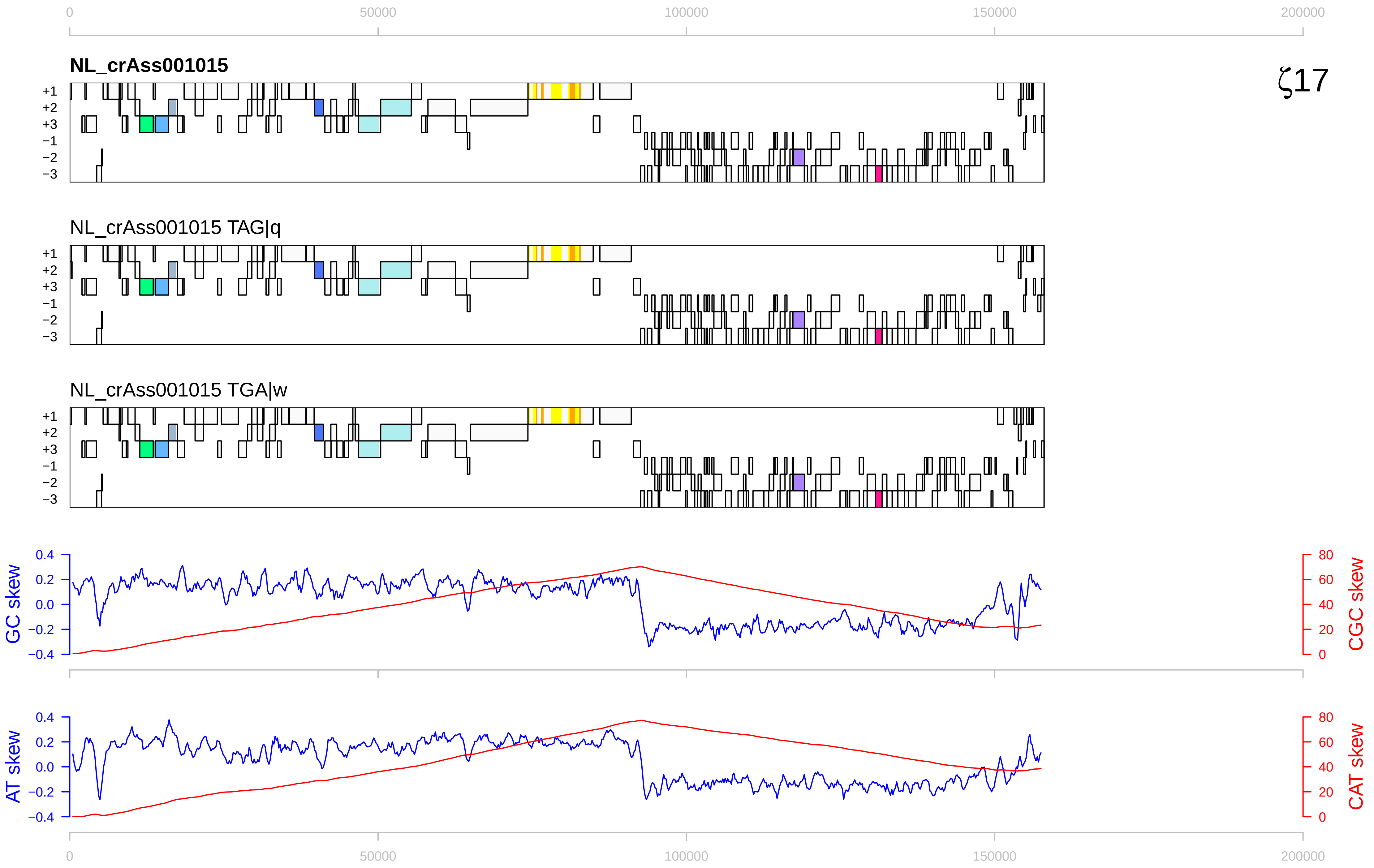

0
50000
100000
150000
200000
NL_crAss001015
ζ17
+1
+2
+3
−1
−2
−3
NL_crAss001015 TAG|q
NL_crAss001015 TGA|w
GC skew
CGC skew
AT skew
CAT skew
0.4
0.2
0.0
−0.2
−0.4
80
60
40
20
0

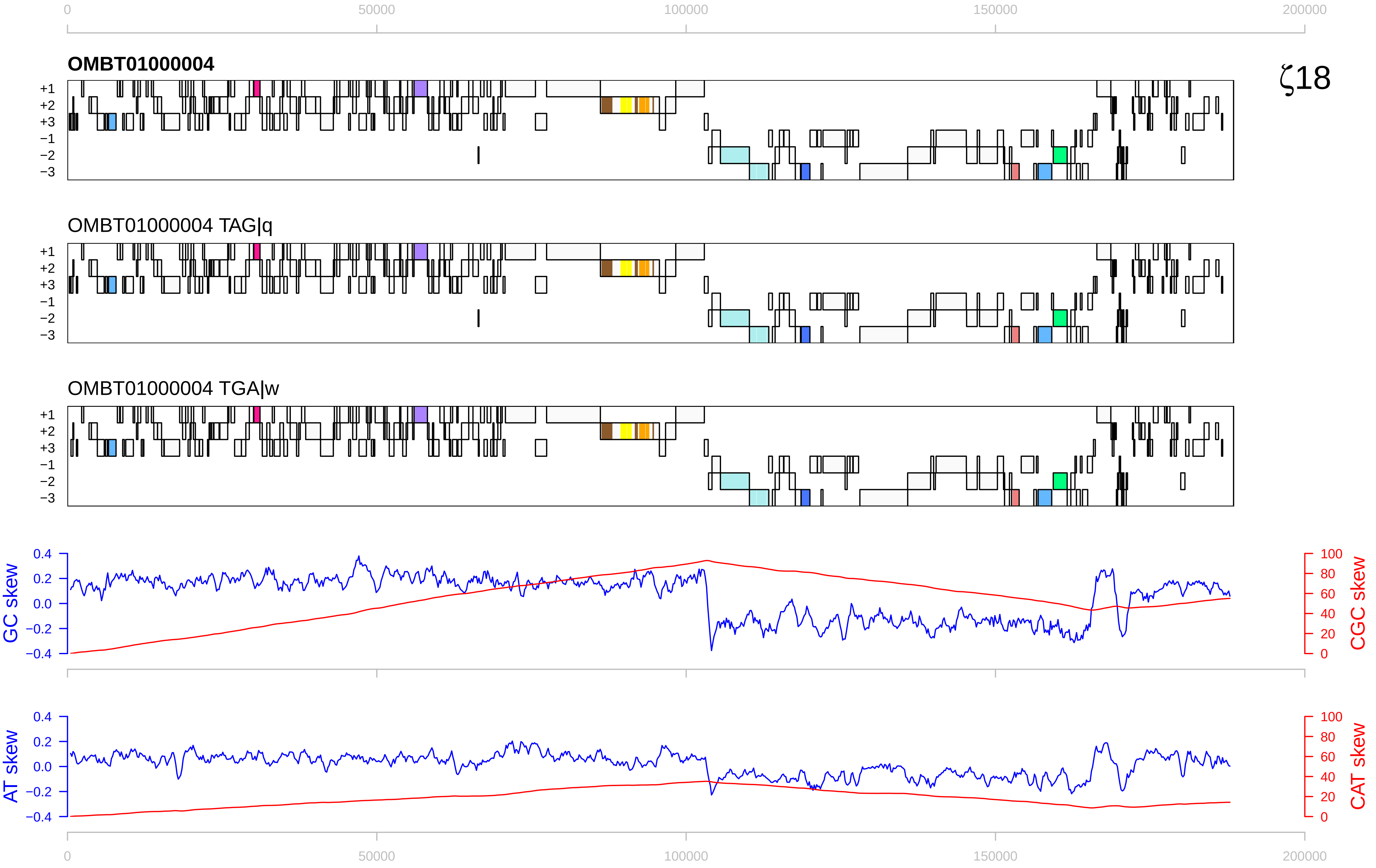

0
50000
100000
150000
200000
OMBT01000004
ζ18
+1
+2
+3
−1
−2
−3
OMBT01000004 TAG|q
OMBT01000004 TGA|w
GC skew
CGC skew
AT skew
CAT skew
0.4
0.2
0.0
−0.2
−0.4
100
80
60
40
20
0

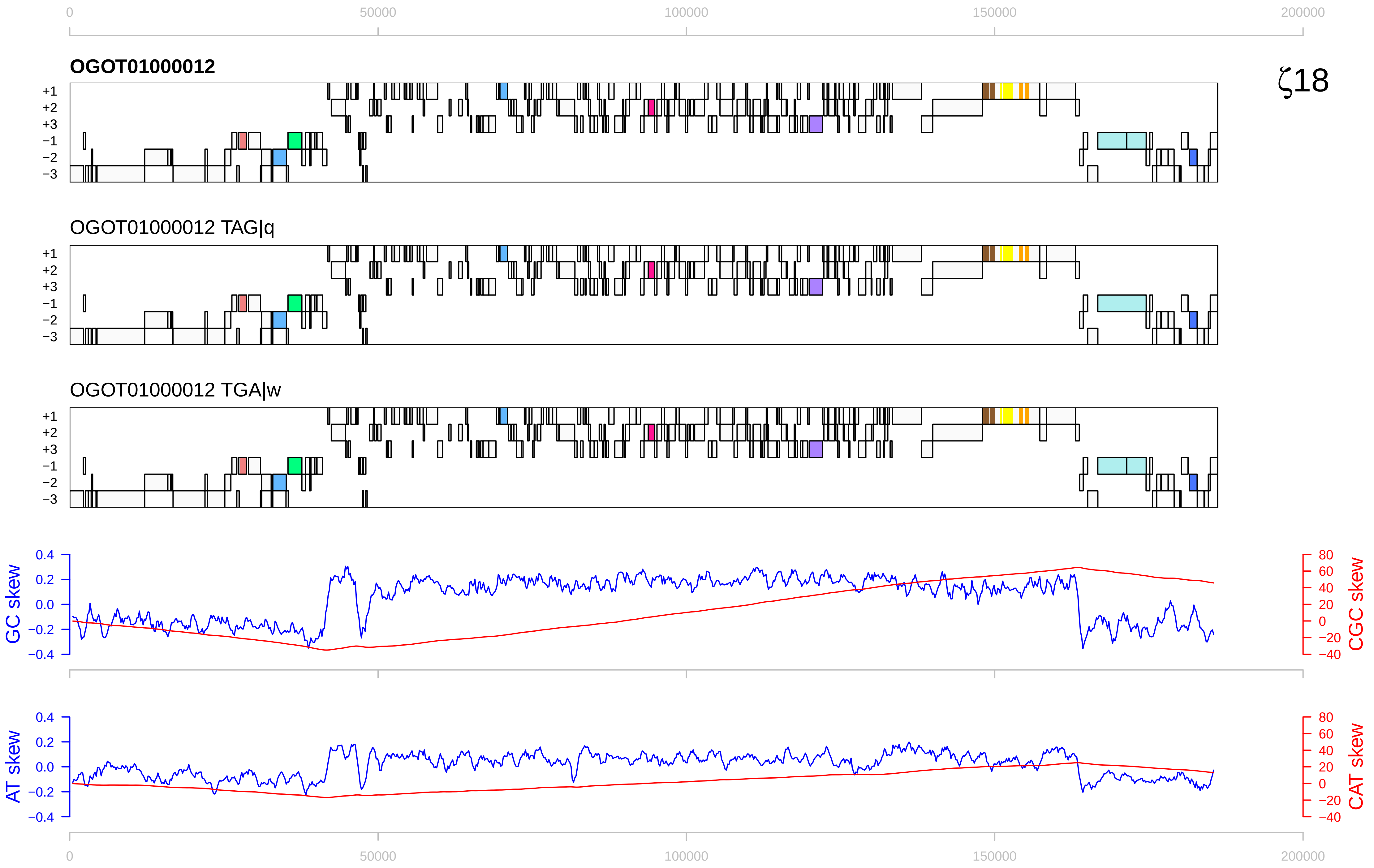

0
50000
100000
150000
200000
OGOT01000012
ζ18
+1
+2
+3
−1
−2
−3
OGOT01000012 TAG|q
OGOT01000012 TGA|w
GC skew
CGC skew
AT skew
CAT skew
0.4
0.2
0.0
−0.2
−0.4
80
60
40
20
0
−20
−40

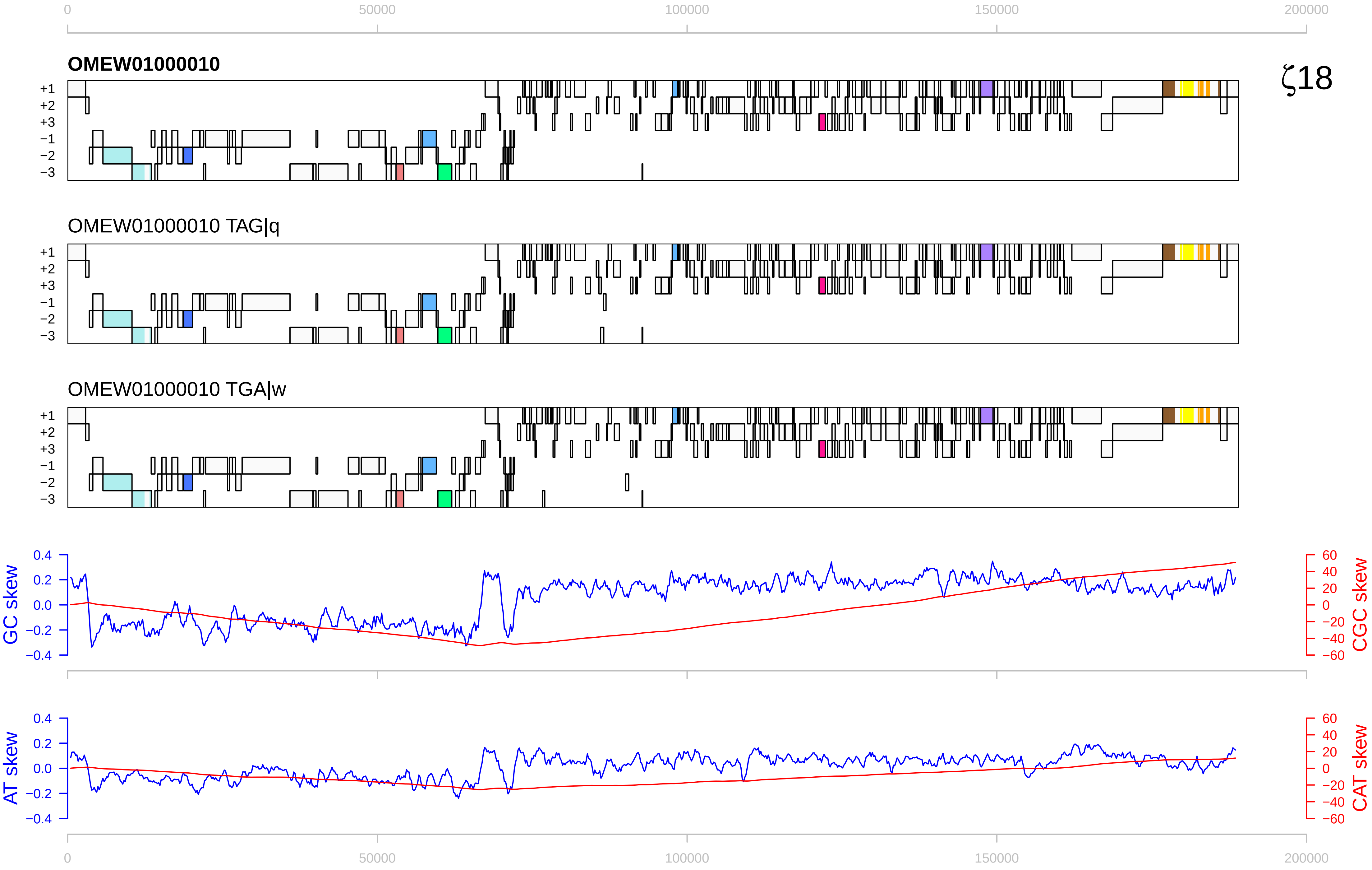

0
50000
100000
150000
200000
OMEW01000010
ζ18
+1
+2
+3
−1
−2
−3
OMEW01000010 TAG|q
OMEW01000010 TGA|w
GC skew
CGC skew
AT skew
CAT skew
0.4
0.2
0.0
−0.2
−0.4
60
40
20
0
−20
−40
−60

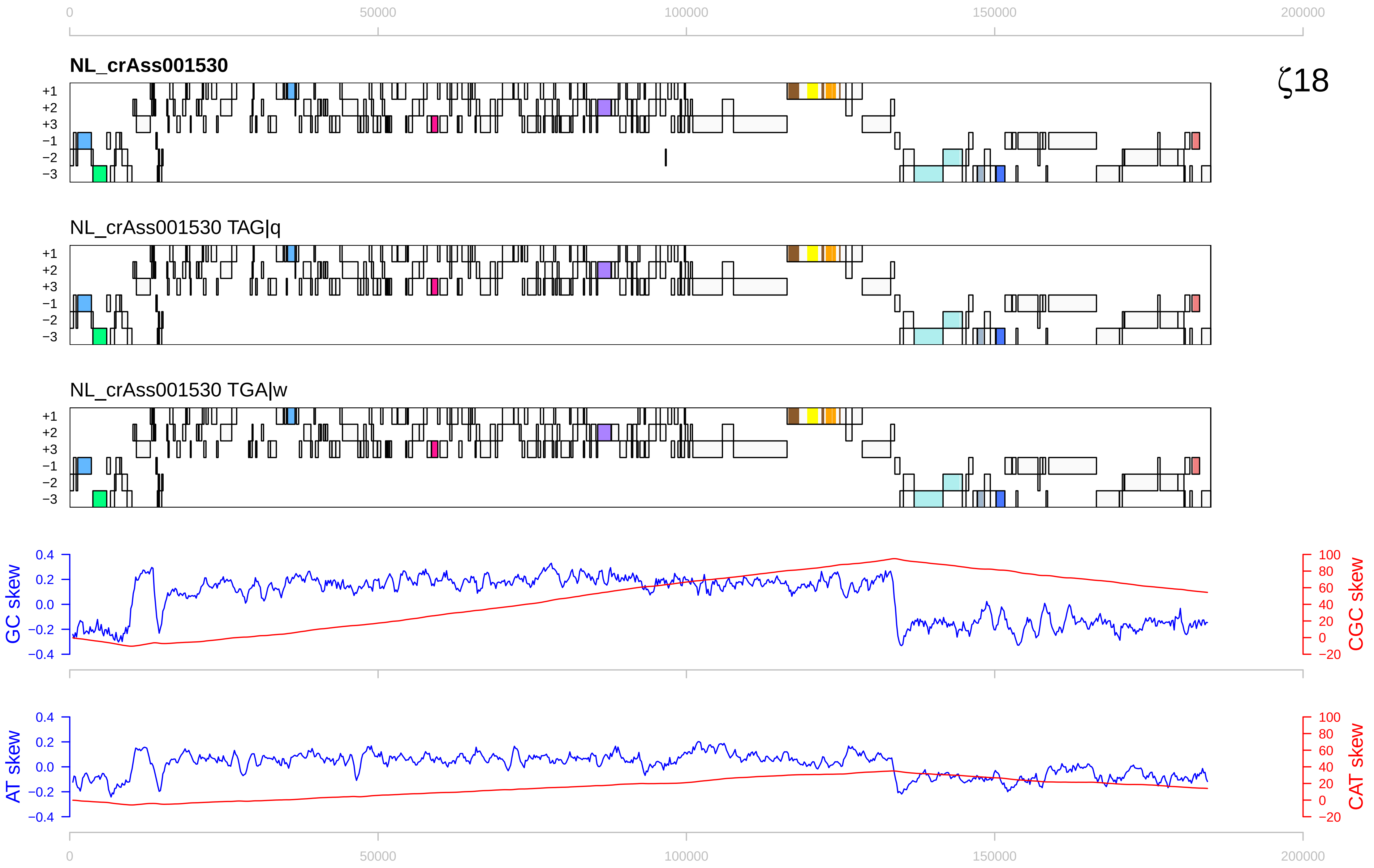
ζ18
NL_crAss001530
NL_crAss001530 TAG|q
NL_crAss001530 TGA|w
+1
+2
+3
−1
−2
−3
GC skew
CGC skew
AT skew
CAT skew
0
50000
100000
150000
200000

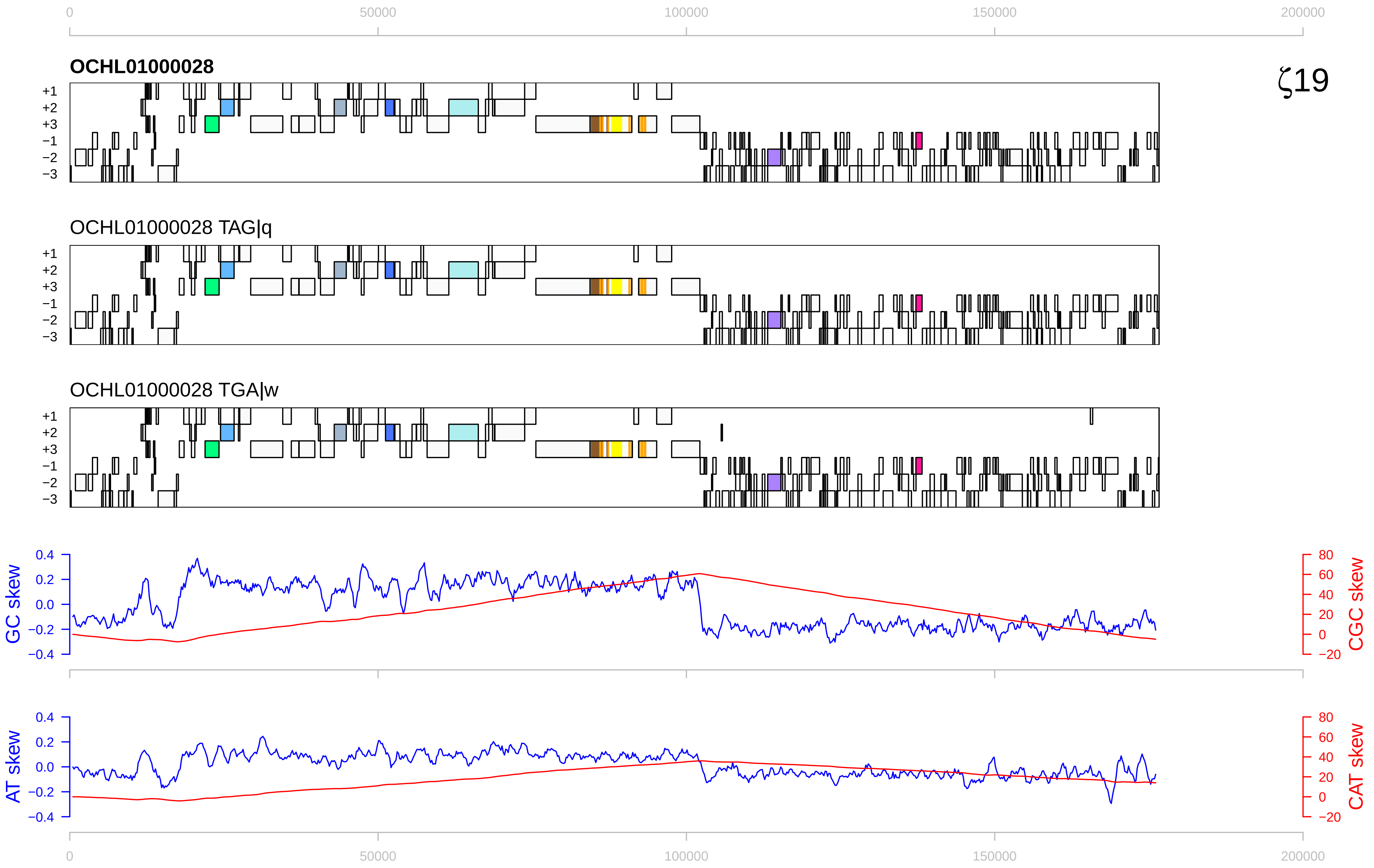

ζ19
OCHL01000028
OCHL01000028 TAG|q
OCHL01000028 TGA|w
+1
+2
+3
−1
−2
−3
GC skew
CGC skew
AT skew
CAT skew
0
50000
100000
150000
200000

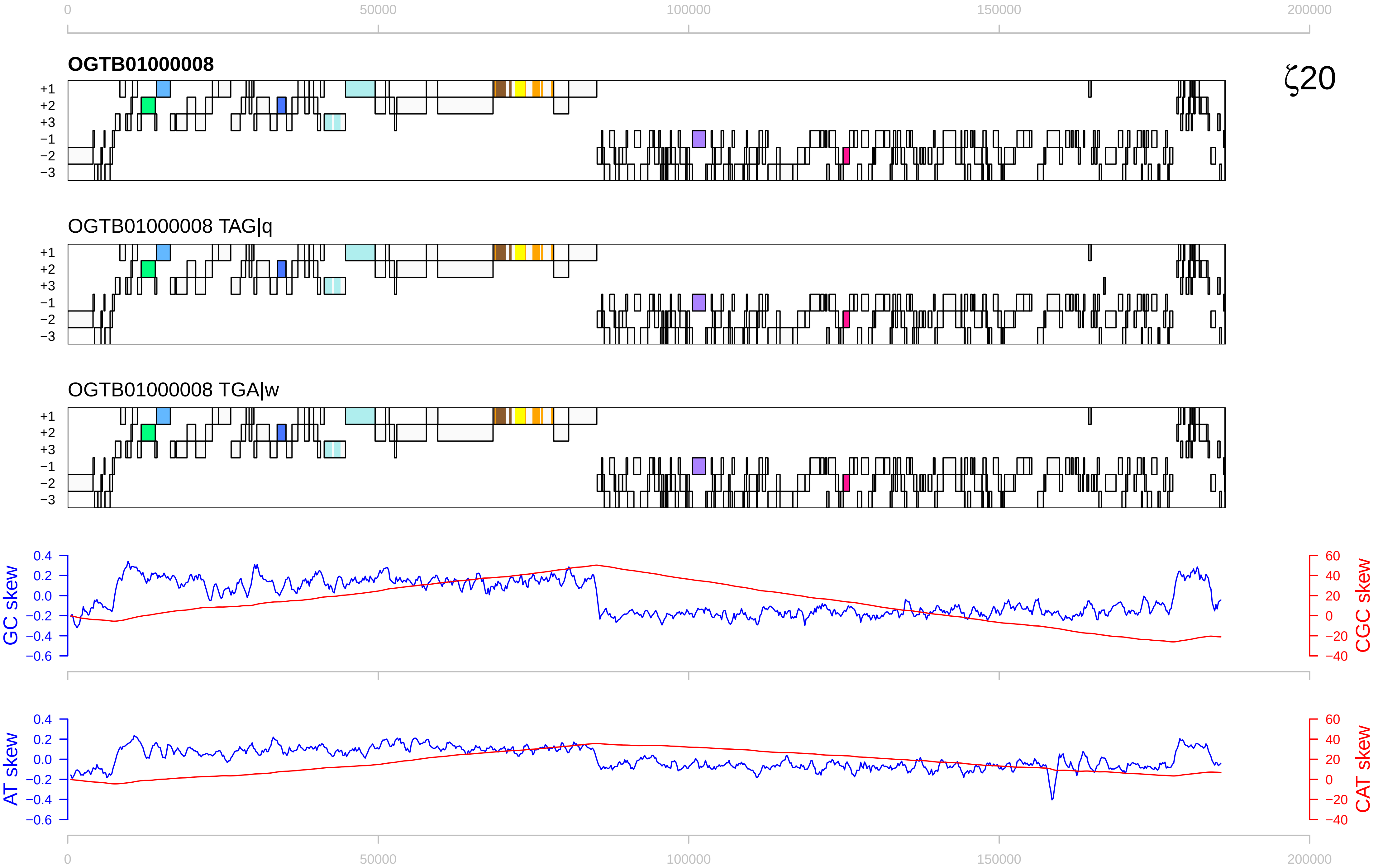

0
50000
100000
150000
200000
OGTB01000008
ζ20
+1
+2
+3
−1
−2
−3
OGTB01000008 TAG|q
OGTB01000008 TGA|w
GC skew
CGC skew
AT skew
CAT skew
0.4
0.2
0.0
−0.2
−0.4
−0.6
60
40
20
0
−20
−40

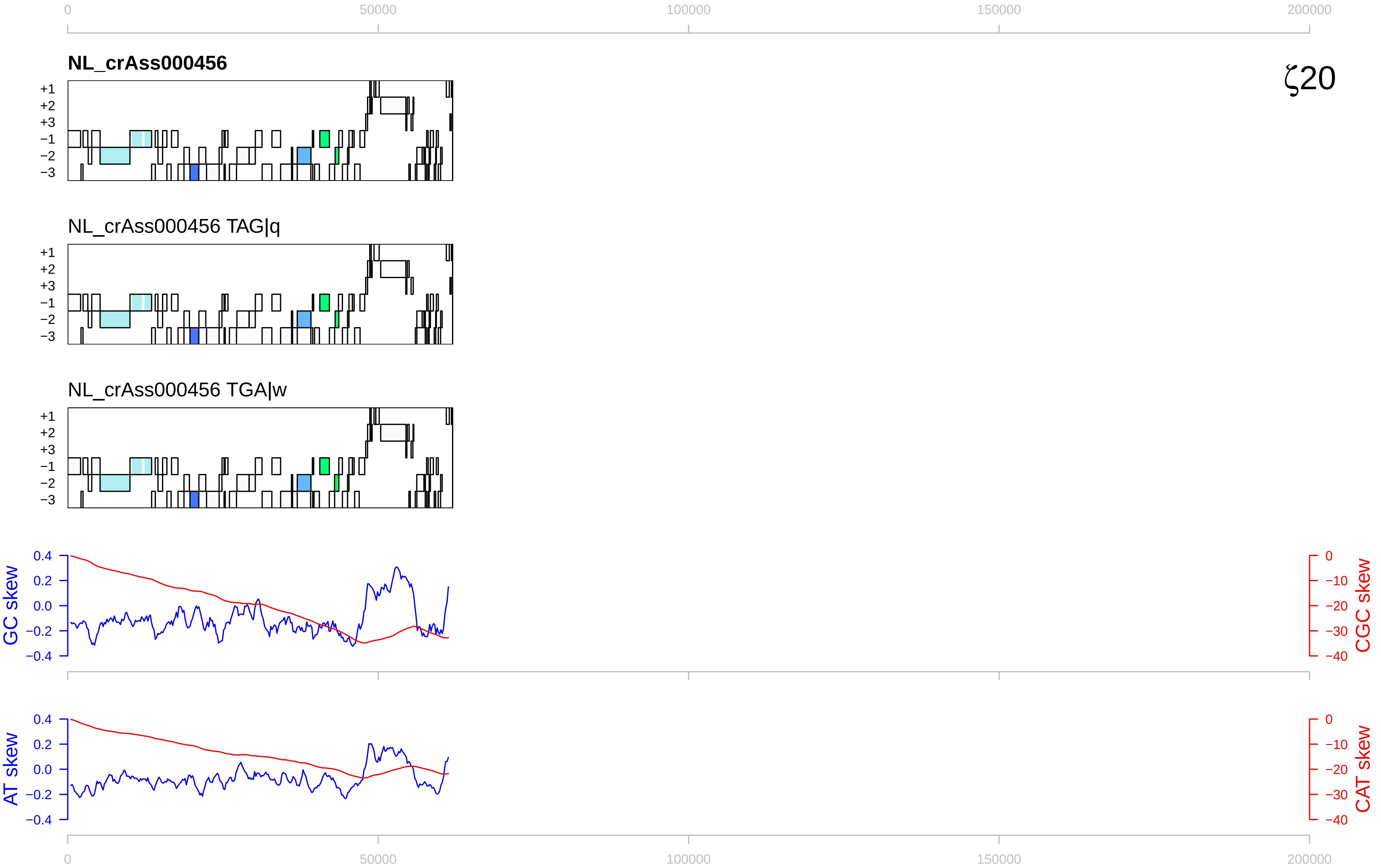

0
50000
100000
150000
200000
NL_crAss000456
ζ20
+1
+2
+3
−1
−2
−3
NL_crAss000456 TAG|q
NL_crAss000456 TGA|w
GC skew
CGC skew
AT skew
CAT skew
0.4
0.2
0.0
−0.2
−0.4
0
−10
−20
−30
−40

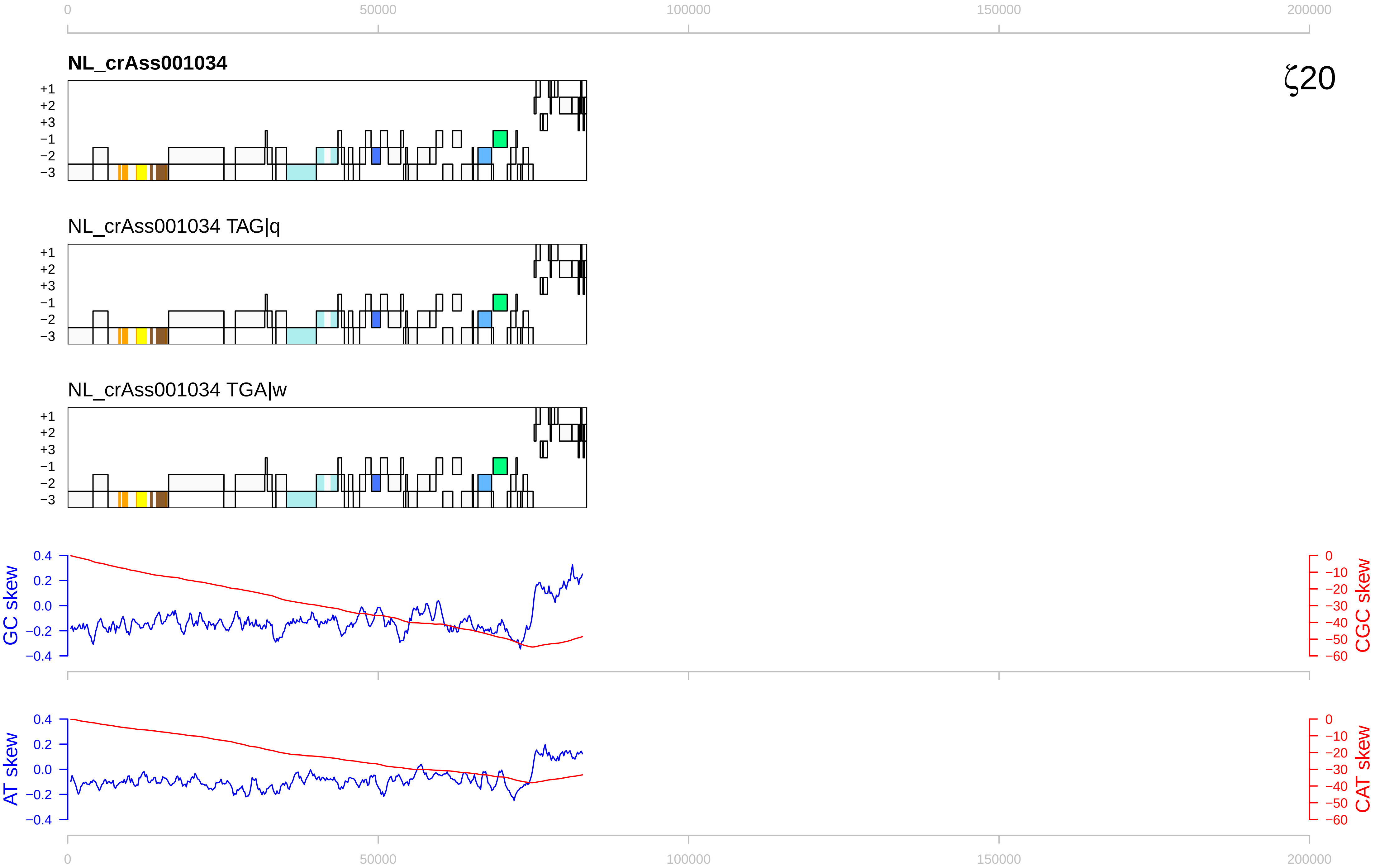
ζ20
NL_crAss001034
NL_crAss001034 TAG|q
NL_crAss001034 TGA|w
+1
+2
+3
−1
−2
−3
GC skew
CGC skew
AT skew
CAT skew
0
50000
100000
150000
200000

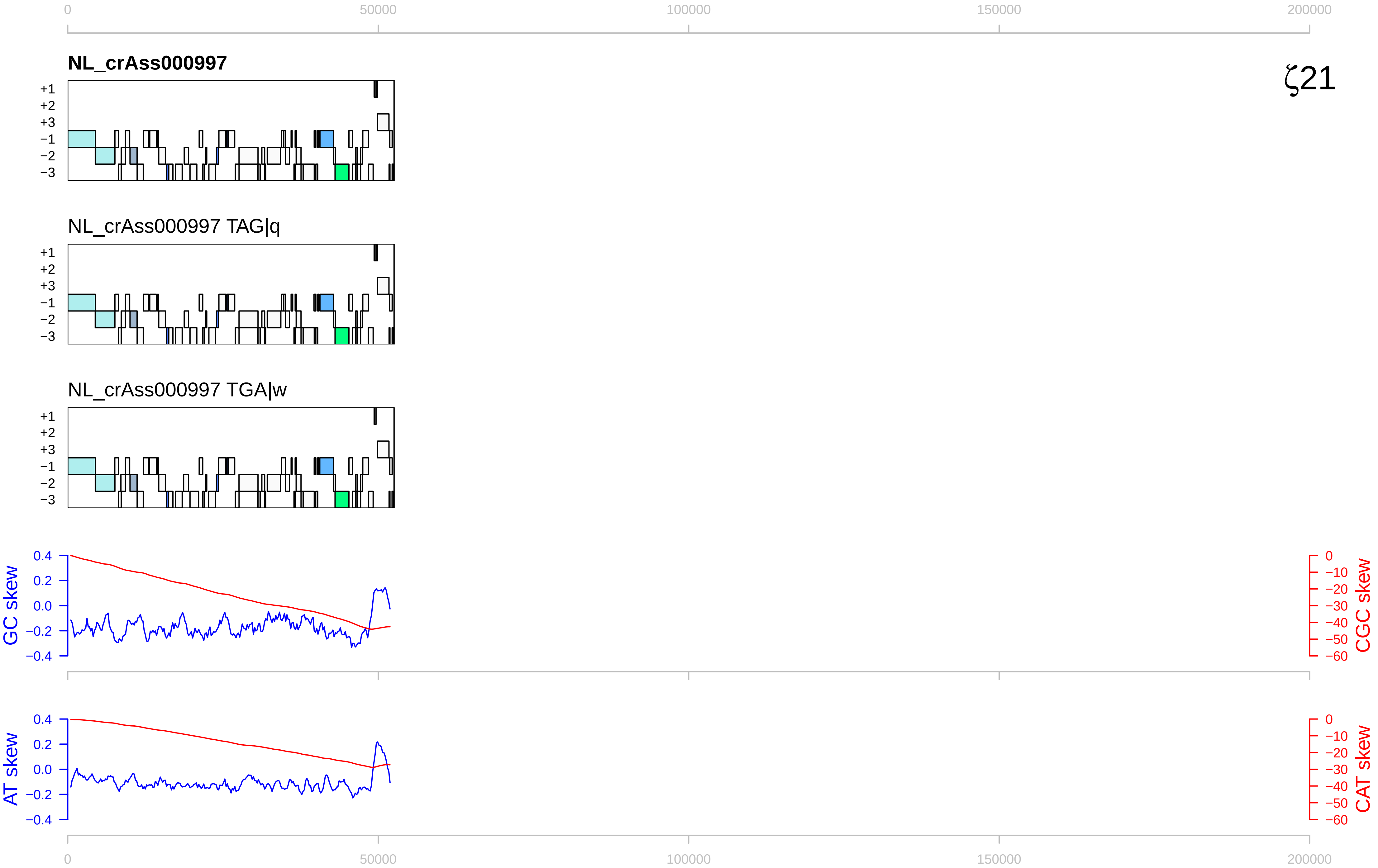
0
50000
100000
150000
200000
NL_crAss000997
ζ21
+1
+2
+3
−1
−2
−3
NL_crAss000997 TAG|q
NL_crAss000997 TGA|w
GC skew
0.4
0.2
0.0
−0.2
−0.4
CGC skew
0
−10
−20
−30
−40
−50
−60
AT skew
CAT skew

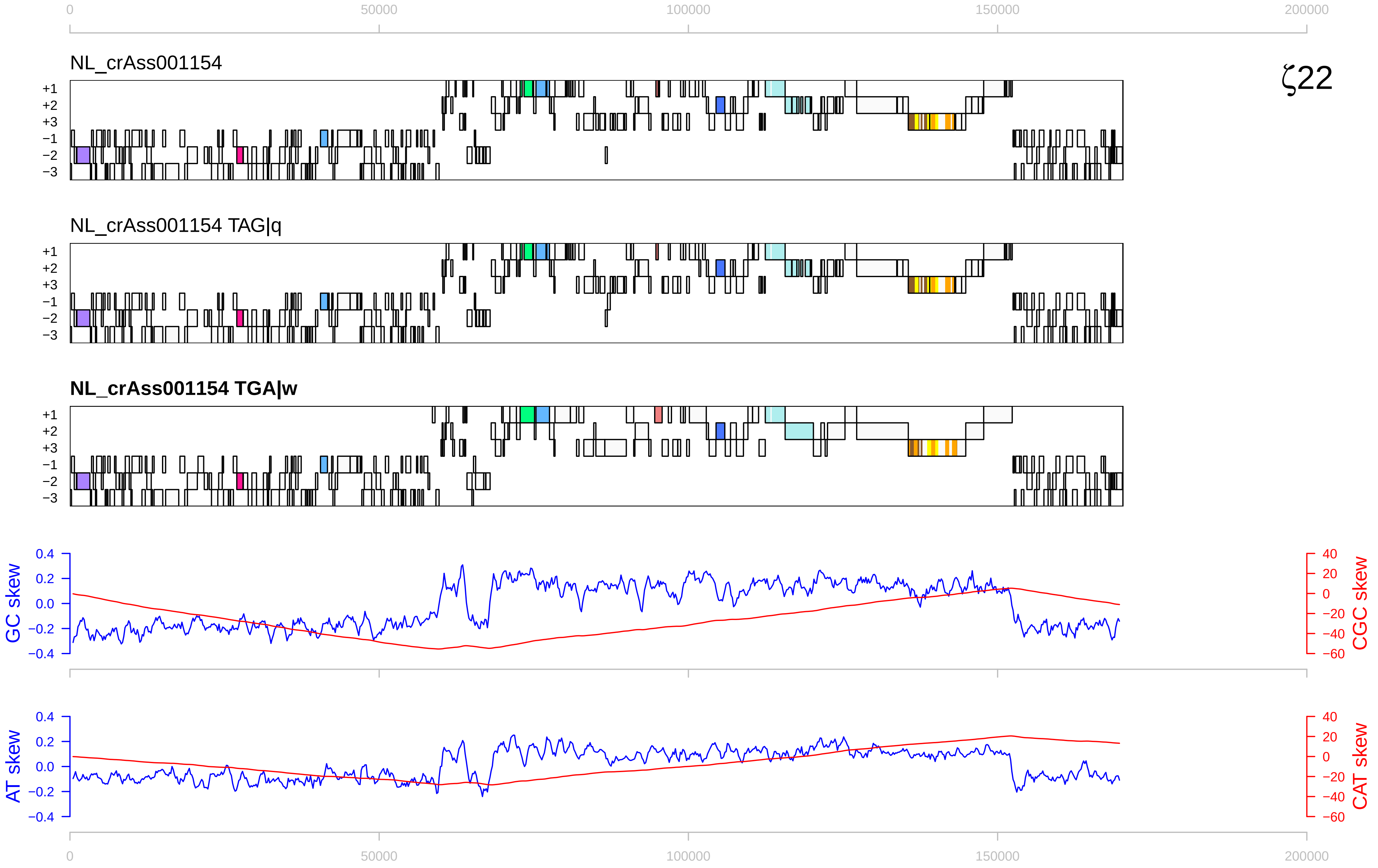

0
50000
100000
150000
200000
NL_crAss001154
ζ22
+1
+2
+3
−1
−2
−3
NL_crAss001154 TAG|q
NL_crAss001154 TGA|w
GC skew
CGC skew
AT skew
CAT skew
0.4
0.2
0.0
−0.2
−0.4
40
20
0
−20
−40
−60

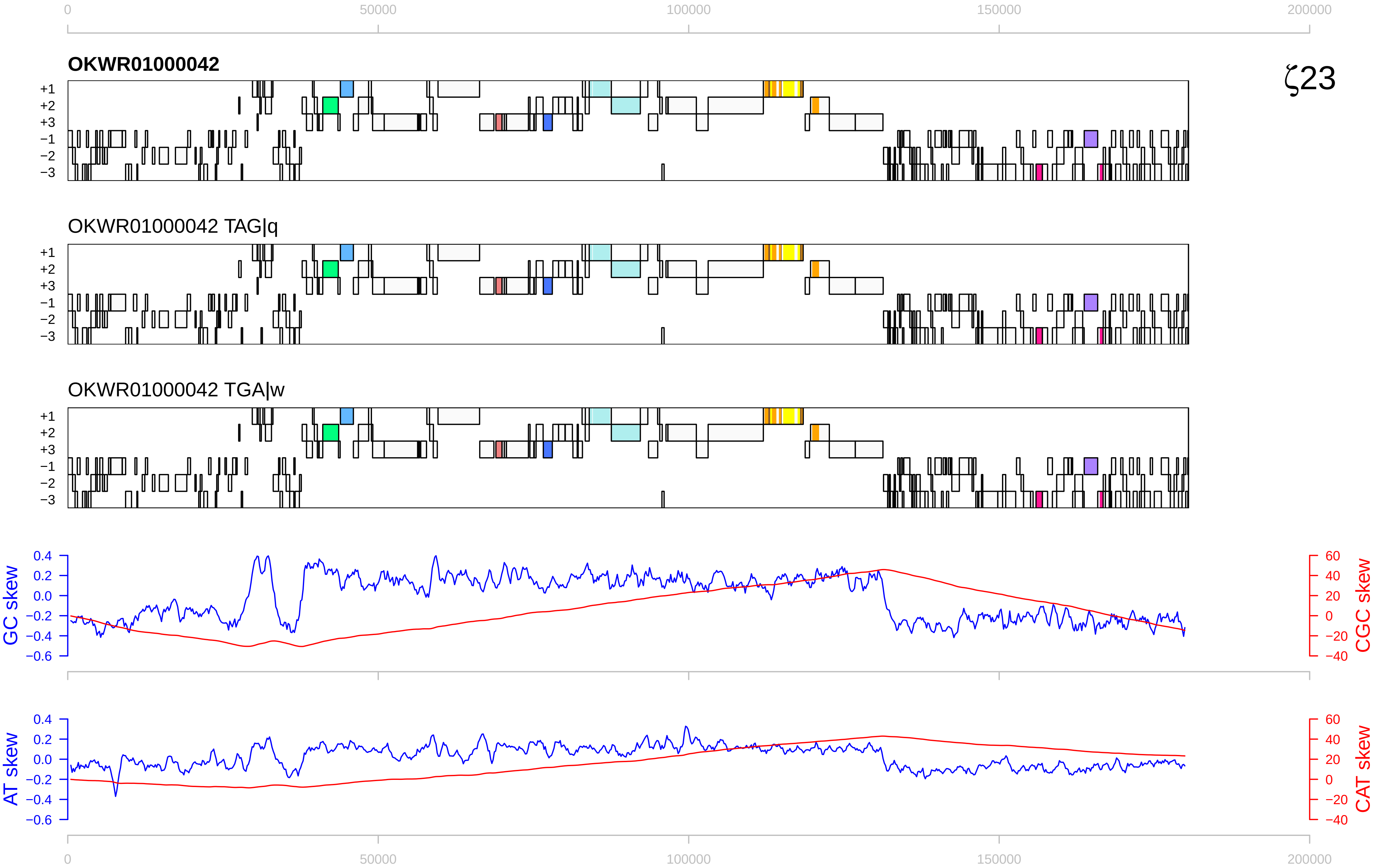

0
50000
100000
150000
200000
OKWR01000042
ζ23
+1
+2
+3
−1
−2
−3
OKWR01000042 TAG|q
OKWR01000042 TGA|w
GC skew
0.4
0.2
0.0
−0.2
−0.4
−0.6
CGC skew
60
40
20
0
−20
−40
AT skew
CAT skew

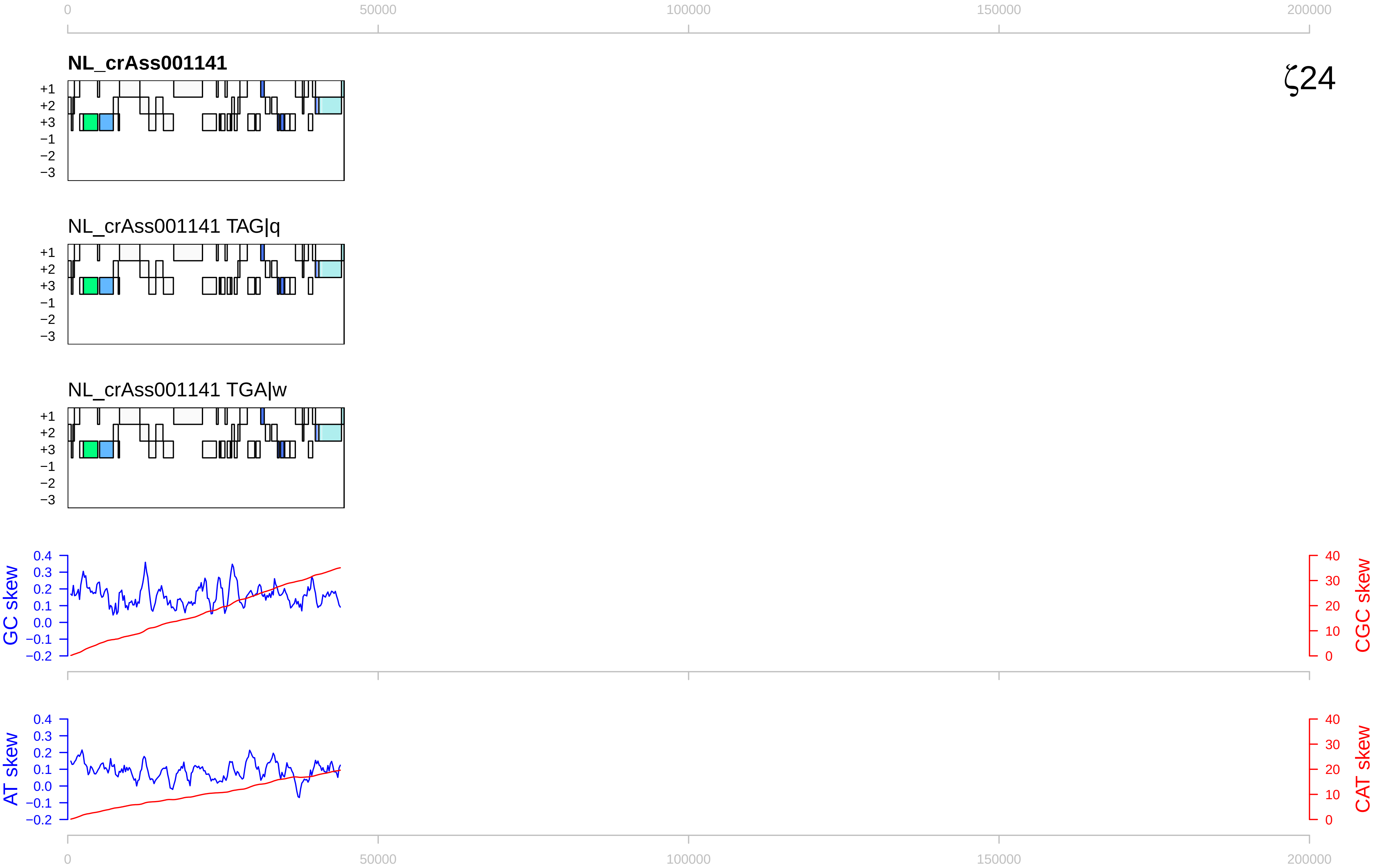

0
50000
100000
150000
200000
NL_crAss001141
ζ24
+1
+2
+3
−1
−2
−3
NL_crAss001141 TAG|q
NL_crAss001141 TGA|w
GC skew
CGC skew
AT skew
CAT skew
0.4
0.3
0.2
0.1
0.0
−0.1
−0.2
40
30
20
10
0

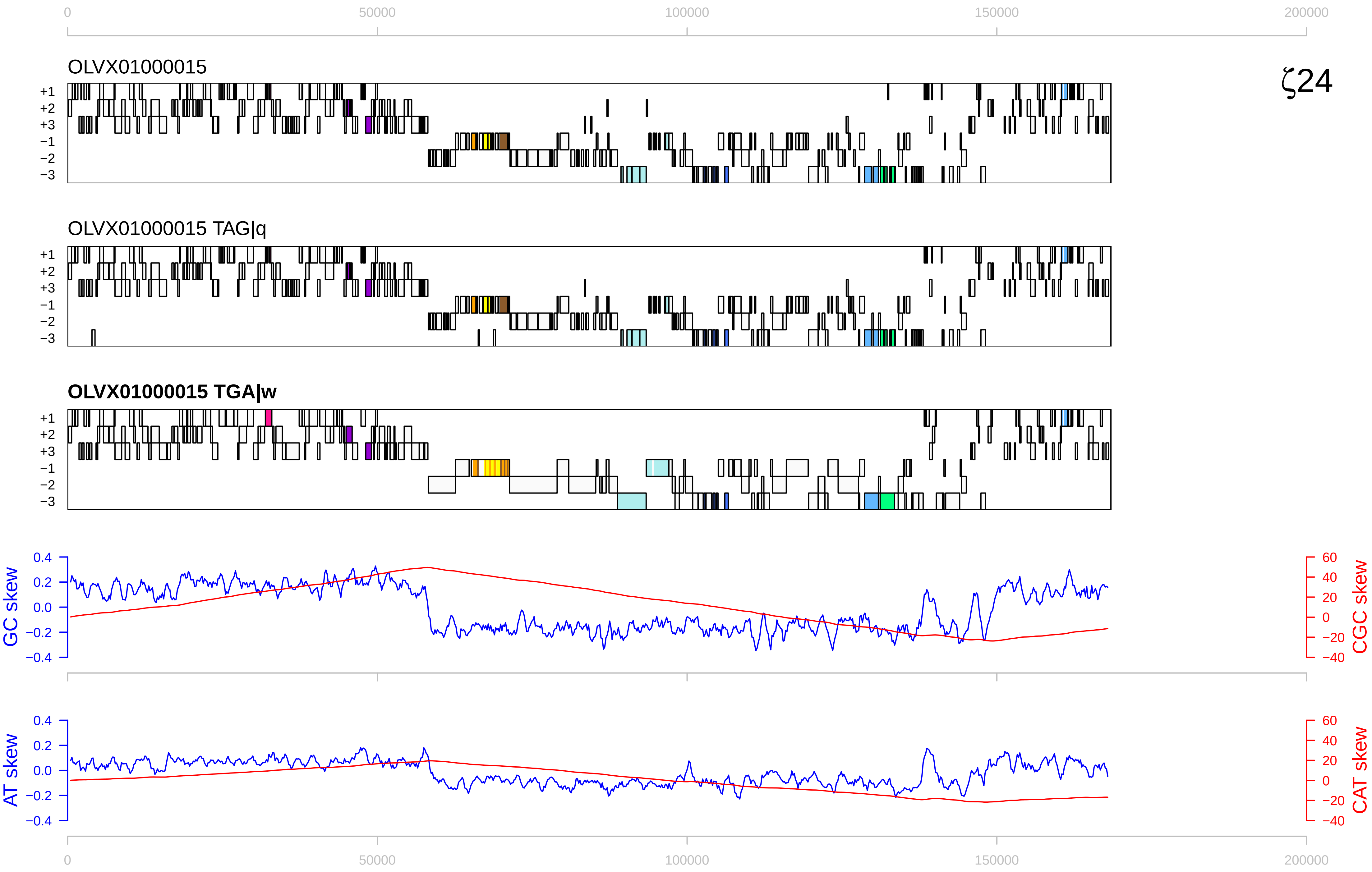

ζ24
OLVX01000015
OLVX01000015 TAG|q
OLVX01000015 TGA|w
+1
+2
+3
−1
−2
−3
GC skew
CGC skew
AT skew
CAT skew
0
50000
100000
150000
200000

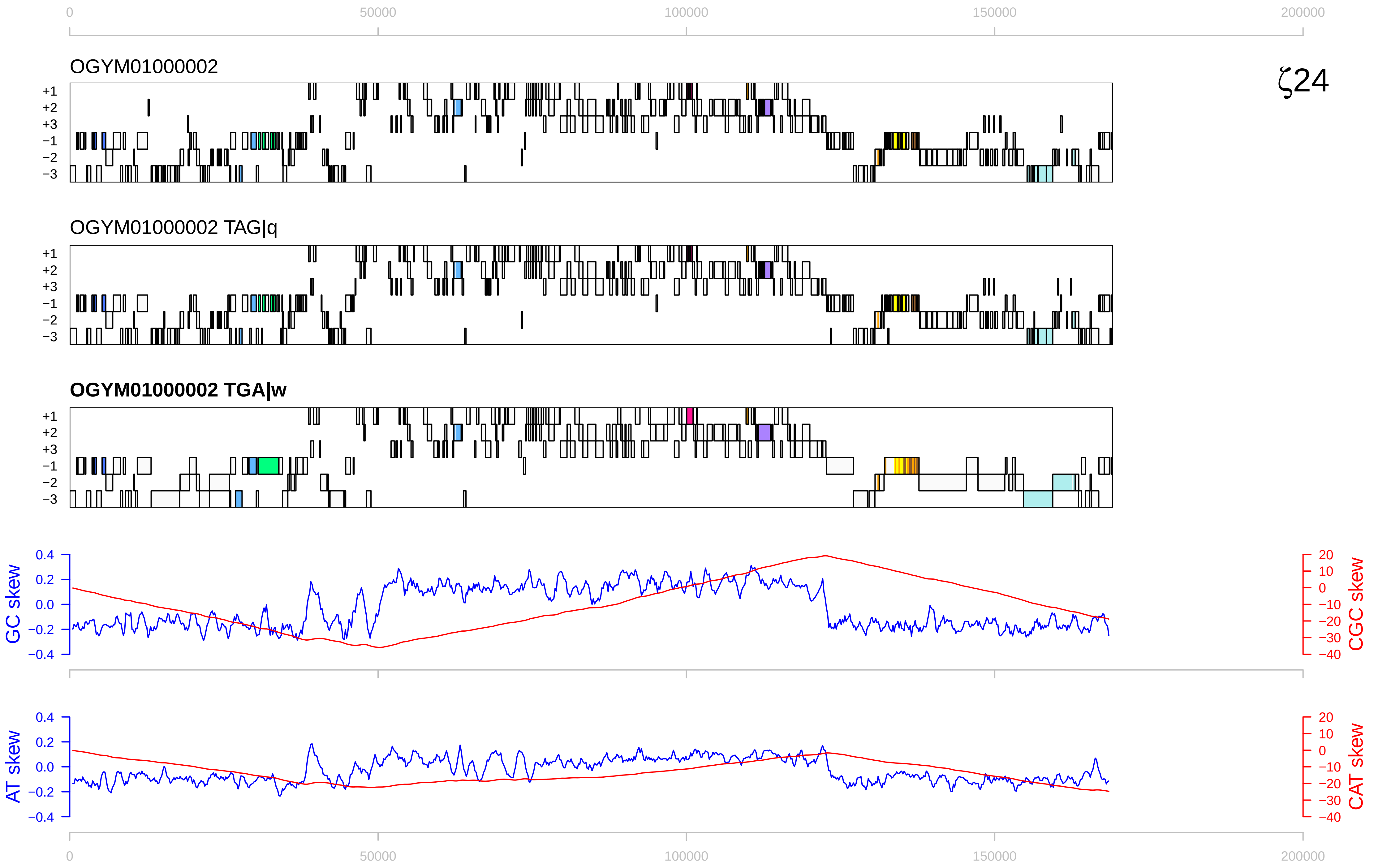

0
50000
100000
150000
200000
OGYM01000002
ζ24
+1
+2
+3
−1
−2
−3
OGYM01000002 TAG|q
OGYM01000002 TGA|w
GC skew
CGC skew
0.4
0.2
0.0
−0.2
−0.4
20
10
0
−10
−20
−30
−40
AT skew
CAT skew

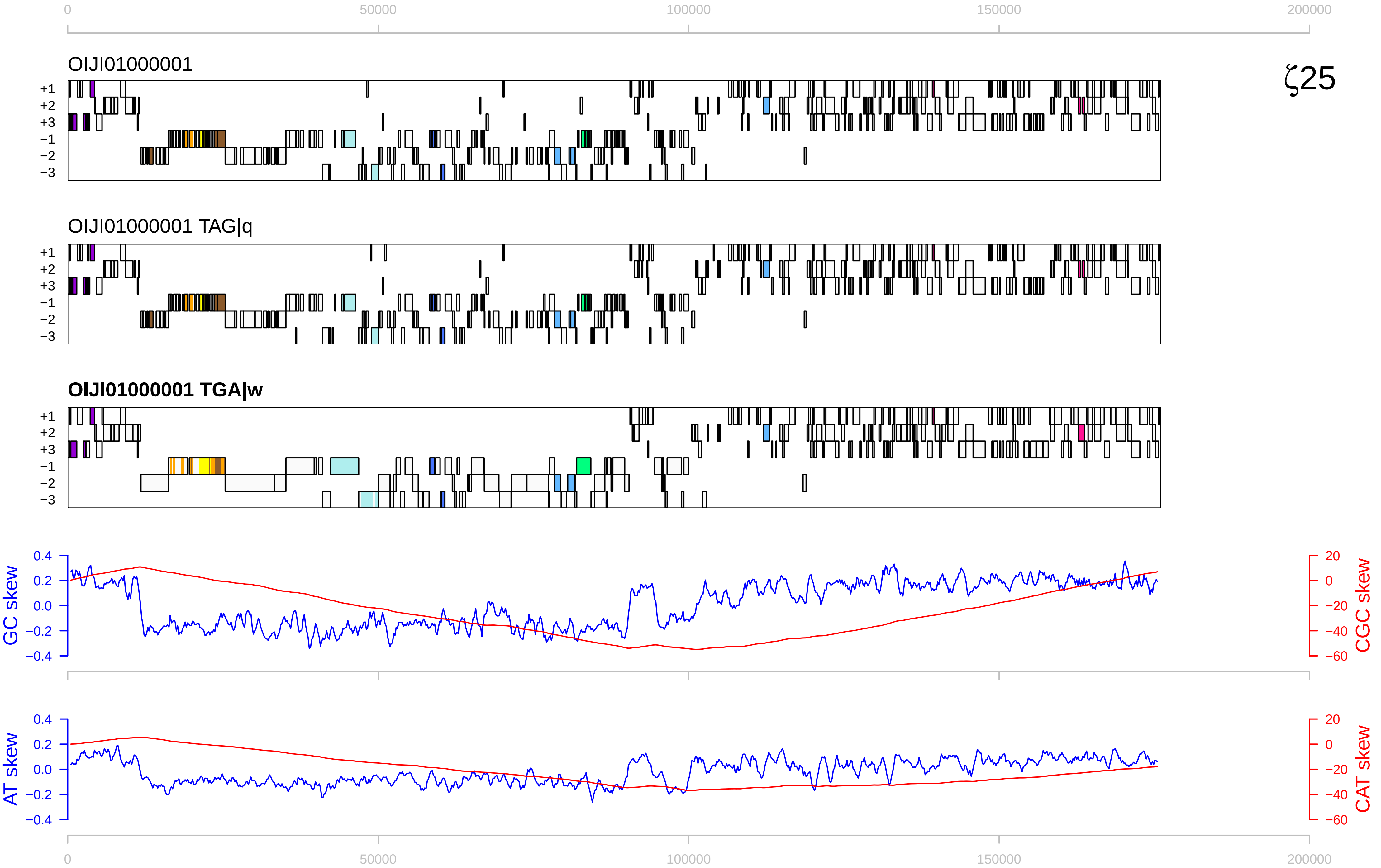

ζ25
OIJI01000001
OIJI01000001 TAG|q
OIJI01000001 TGA|w
+1
+2
+3
−1
−2
−3
0
50000
100000
150000
200000
GC skew
CGC skew
AT skew
CAT skew
0.4
0.2
0.0
−0.2
−0.4
20
0
−20
−40
−60

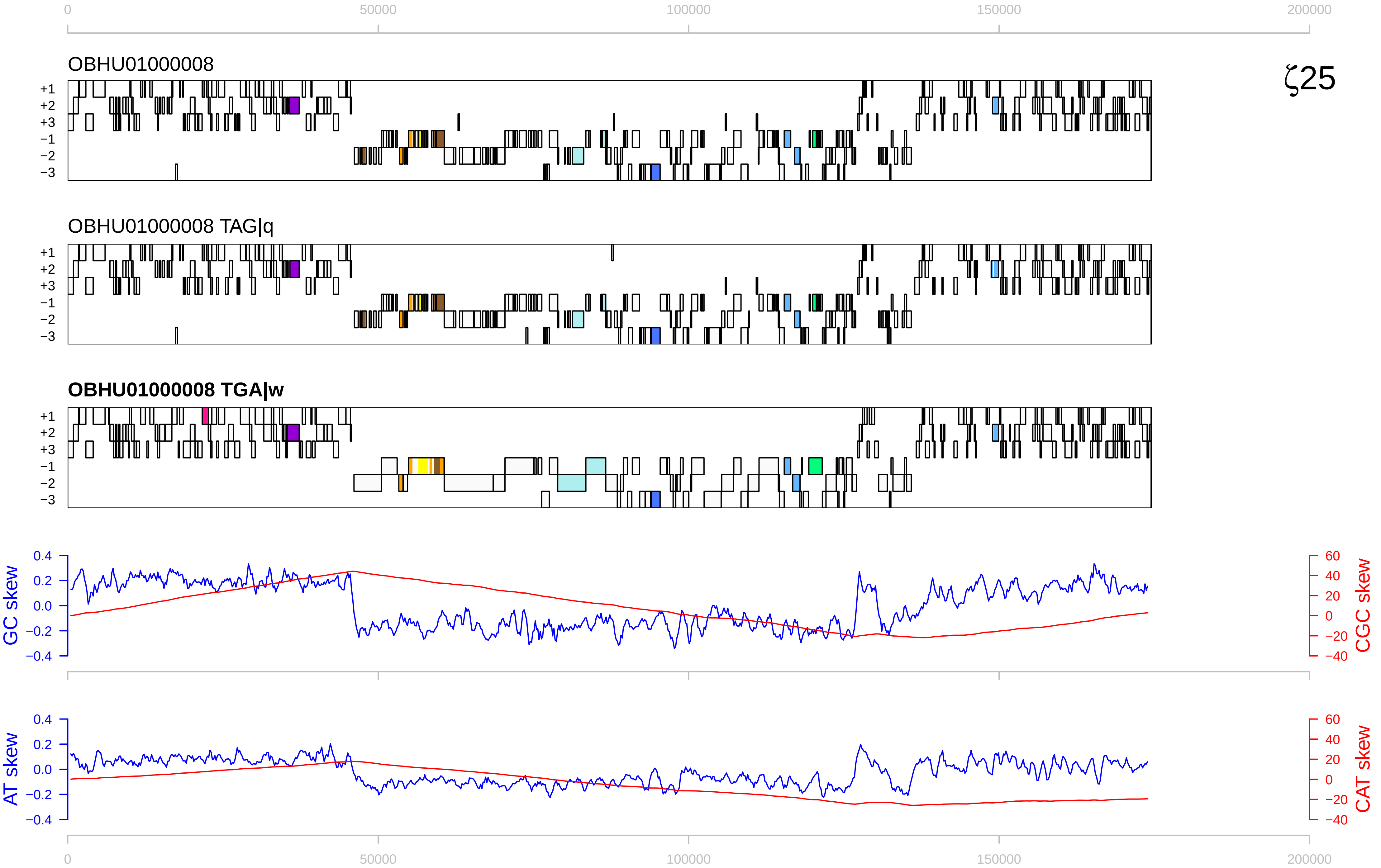

0
50000
100000
150000
200000
OBHU01000008
ζ25
+1
+2
+3
−1
−2
−3
OBHU01000008 TAG|q
OBHU01000008 TGA|w
GC skew
CGC skew
AT skew
CAT skew
0.4
0.2
0.0
−0.2
−0.4
60
40
20
0
−20
−40

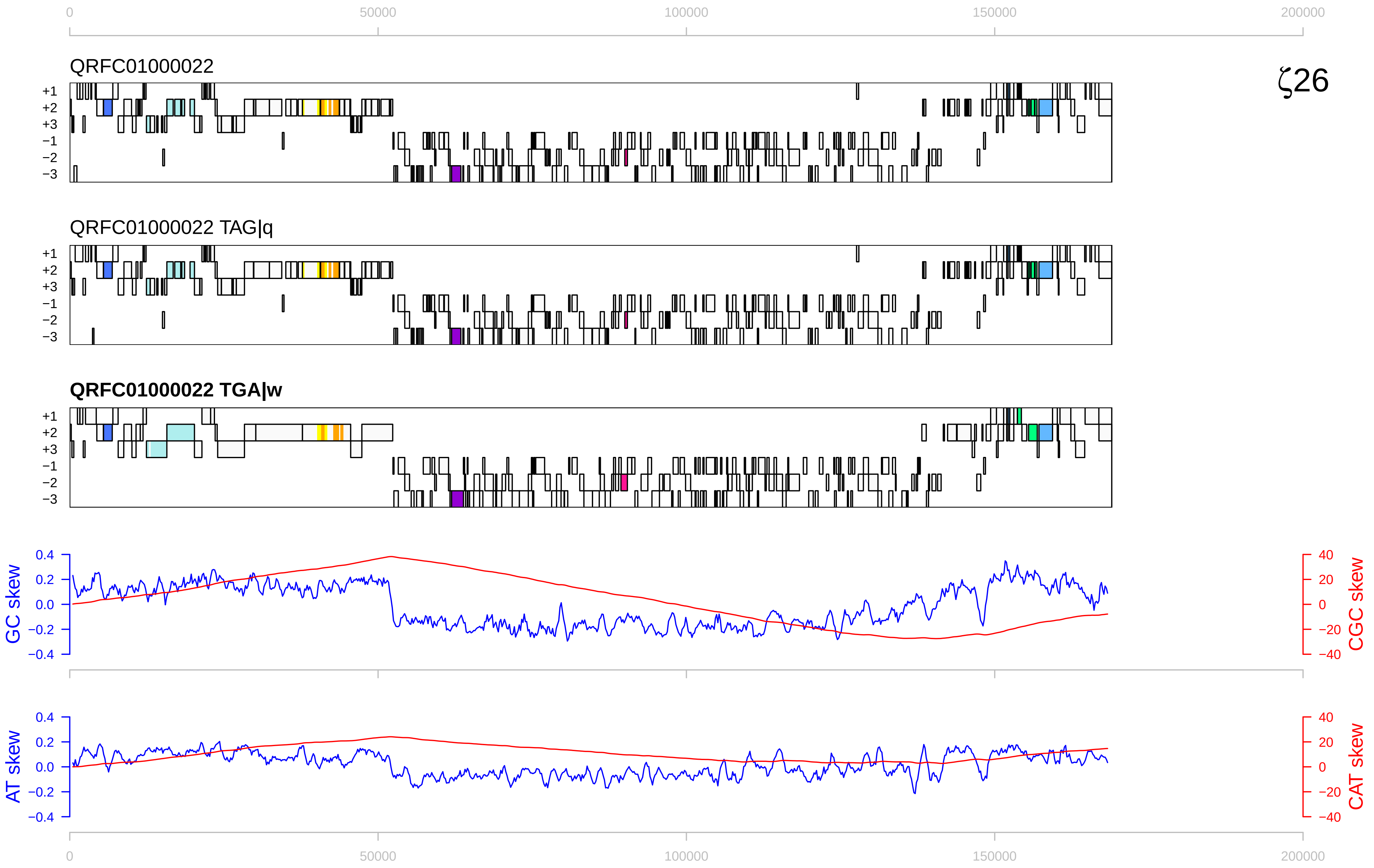

0
50000
100000
150000
200000
QRFC01000022
ζ26
+1
+2
+3
−1
−2
−3
QRFC01000022 TAG|q
QRFC01000022 TGA|w
GC skew
CGC skew
AT skew
CAT skew
0.4
0.2
0.0
−0.2
−0.4
40
20
0
−20
−40

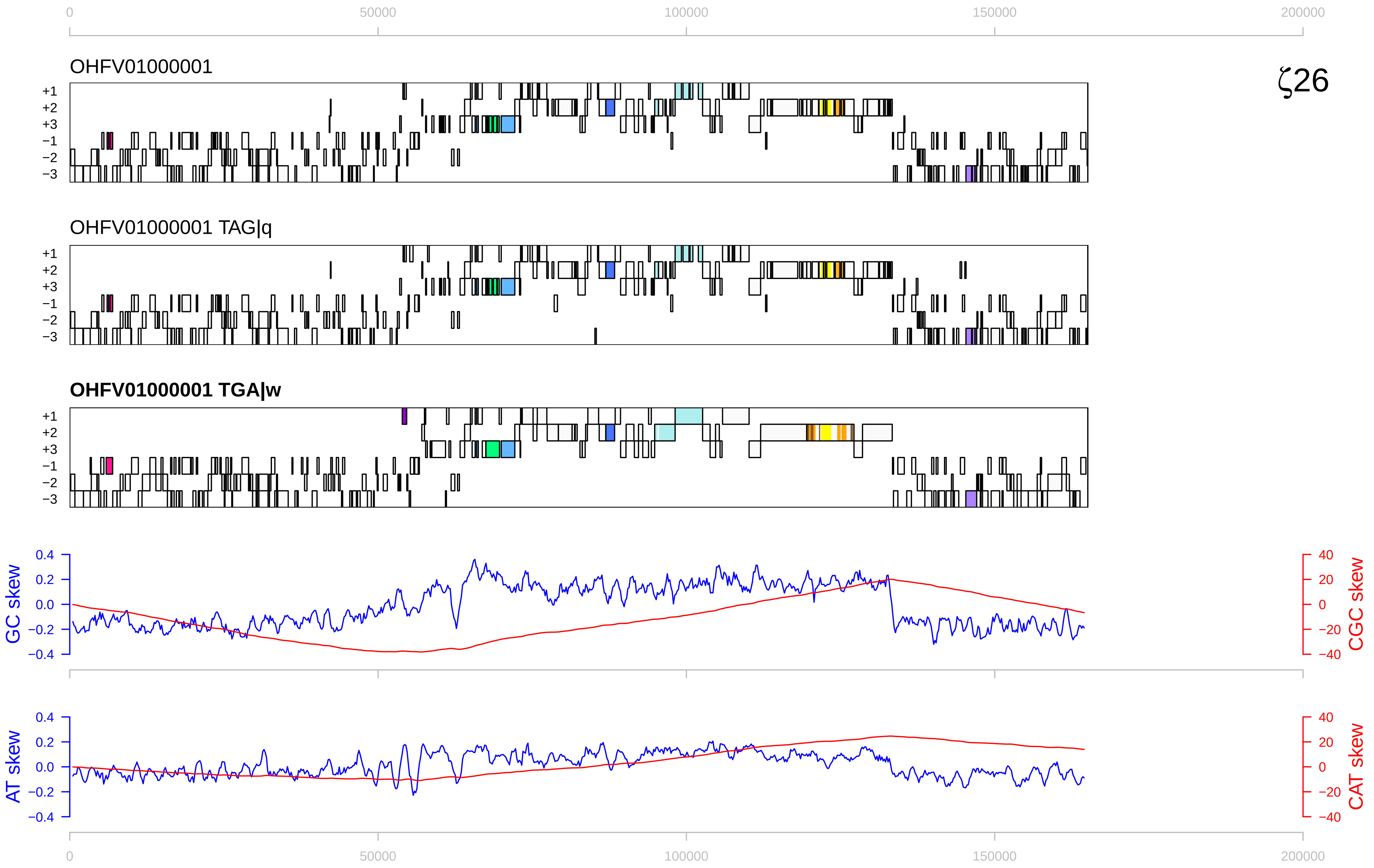

0
50000
100000
150000
200000
OHFV01000001
ζ26
+1
+2
+3
−1
−2
−3
OHFV01000001 TAG|q
OHFV01000001 TGA|w
GC skew
CGC skew
0.4
0.2
0.0
−0.2
−0.4
40
20
0
−20
−40
AT skew
CAT skew

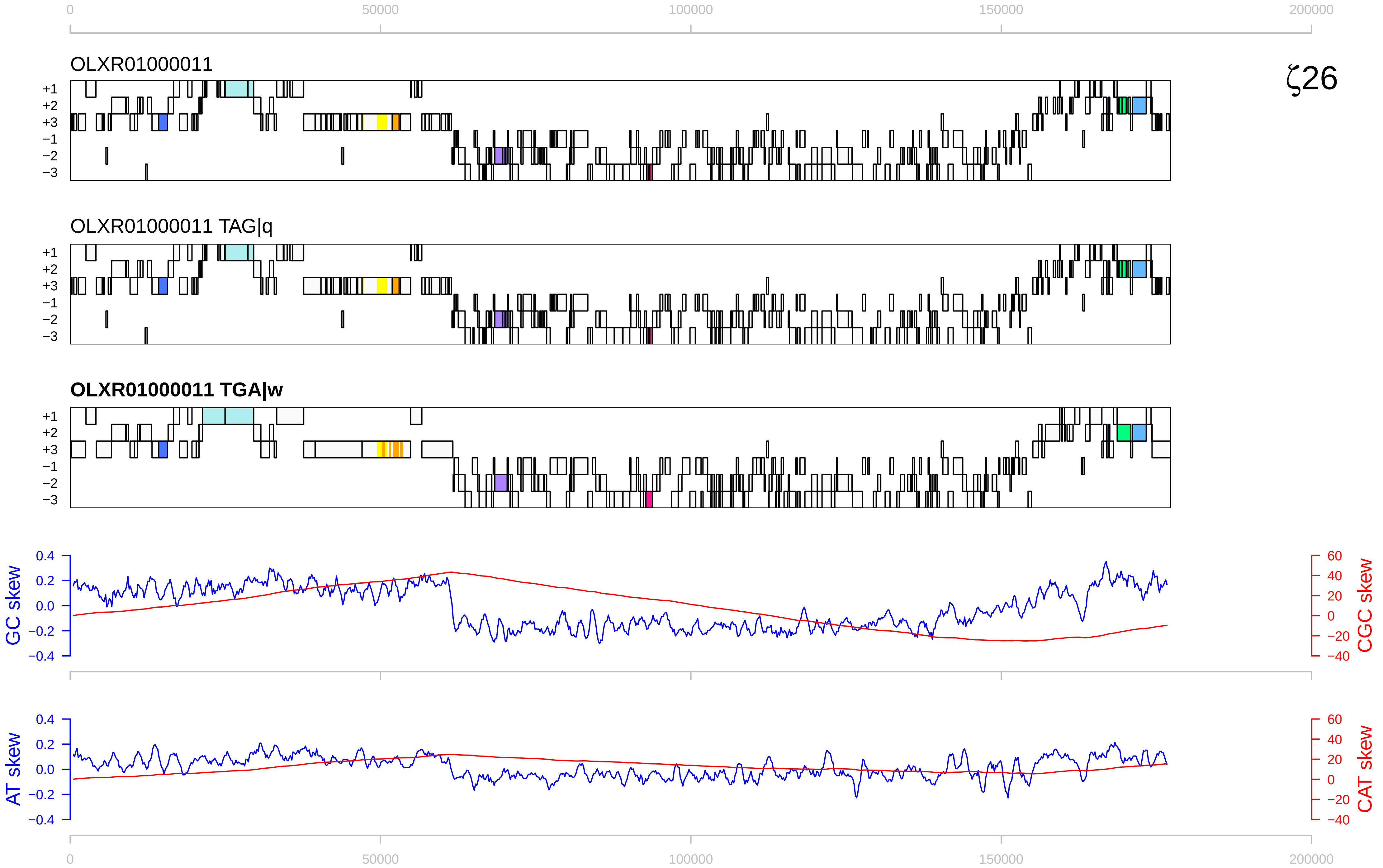

0
50000
100000
150000
200000
OLXR01000011
ζ26
+1
+2
+3
−1
−2
−3
OLXR01000011 TAG|q
OLXR01000011 TGA|w
GC skew
CGC skew
AT skew
CAT skew
0.4
0.2
0.0
−0.2
−0.4
60
40
20
0
−20
−40

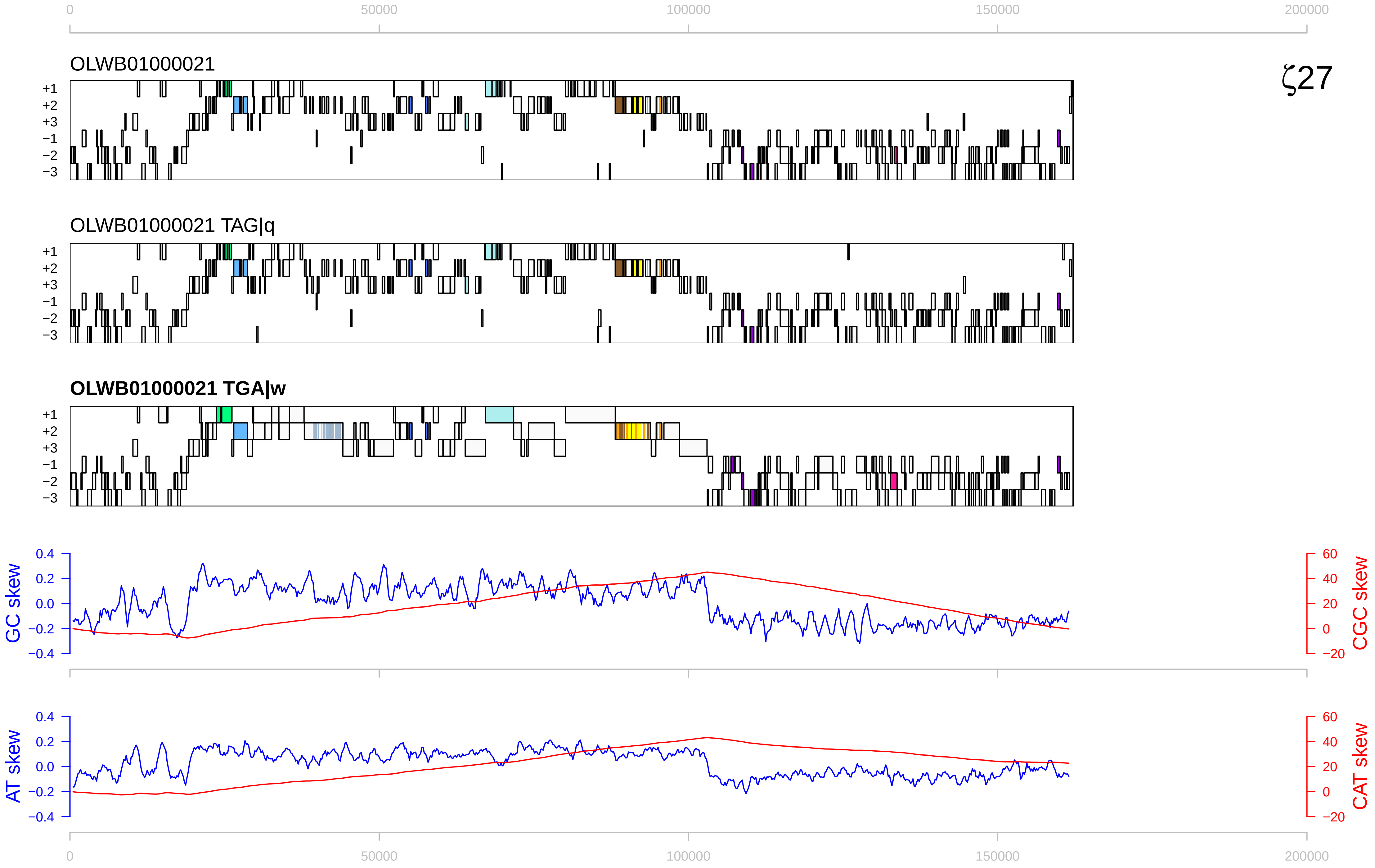

0
50000
100000
150000
200000
OLWB01000021
ζ27
+1
+2
+3
−1
−2
−3
OLWB01000021 TAG|q
OLWB01000021 TGA|w
GC skew
CGC skew
0.4
0.2
0.0
−0.2
−0.4
60
40
20
0
−20
AT skew
CAT skew

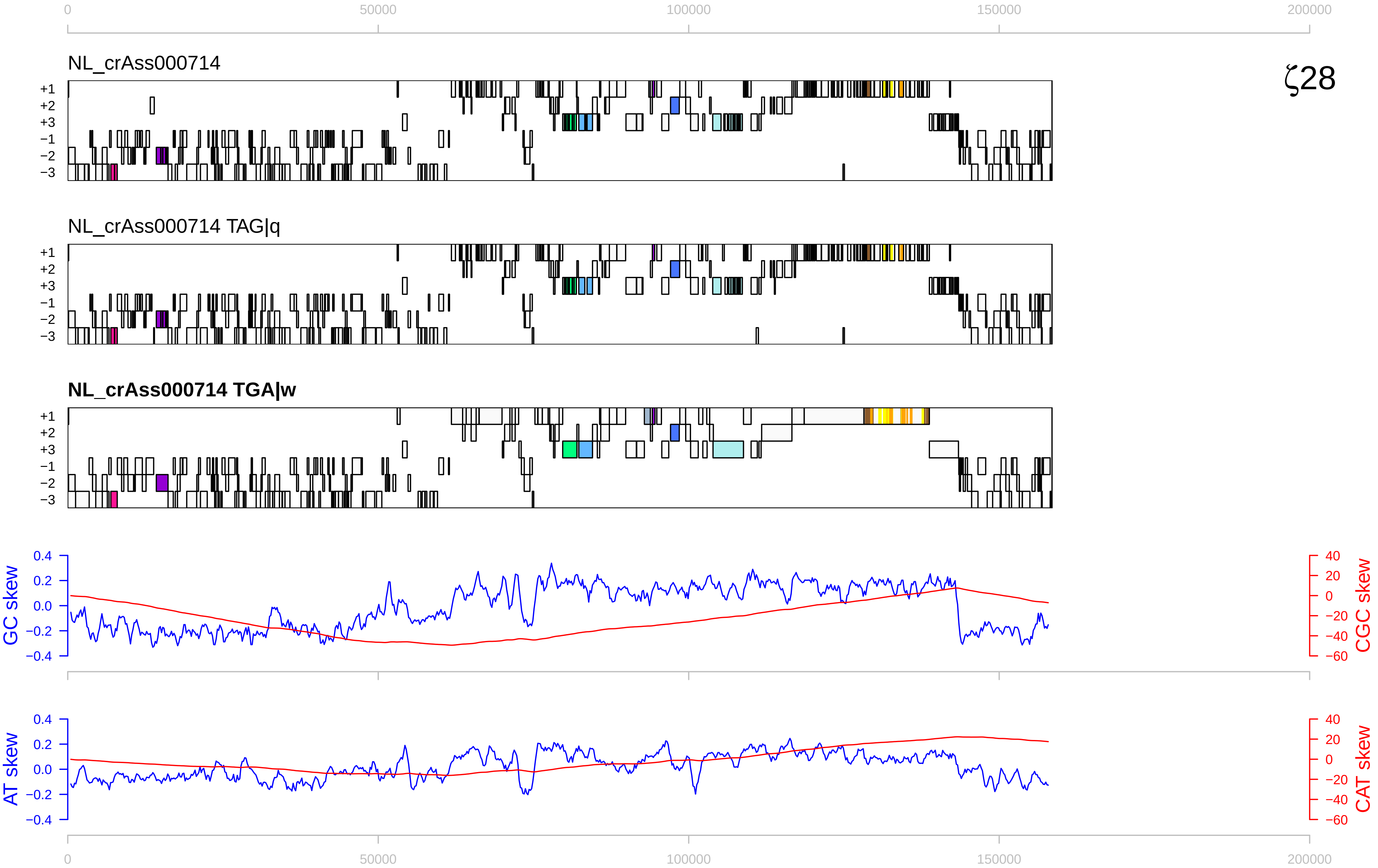

ζ28
NL_crAss000714
NL_crAss000714 TAG|q
NL_crAss000714 TGA|w
+1
+2
+3
−1
−2
−3
GC skew
CGC skew
AT skew
CAT skew
0
50000
100000
150000
200000

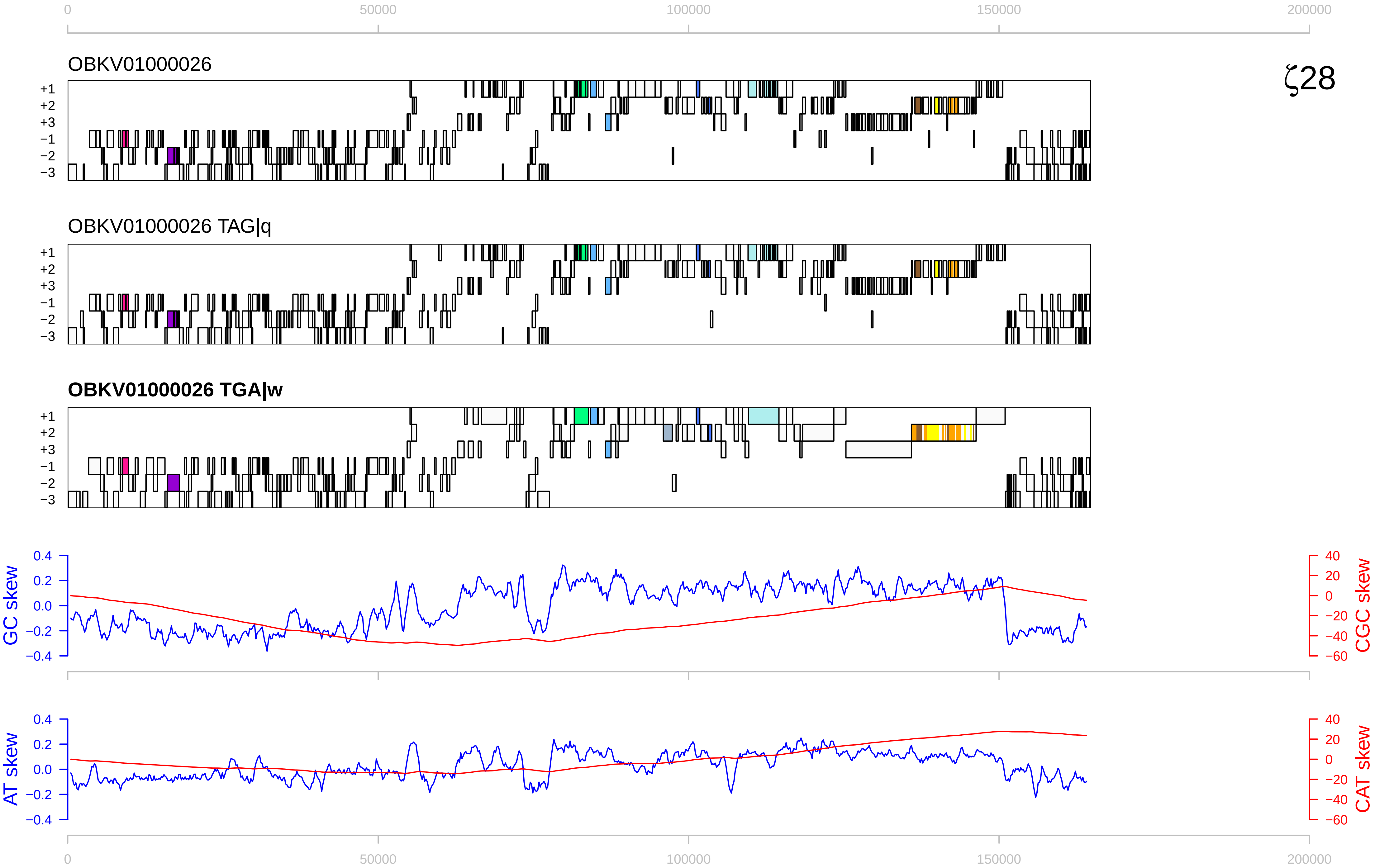

0
50000
100000
150000
200000
OBKV01000026
ζ28
+1
+2
+3
−1
−2
−3
OBKV01000026 TAG|q
OBKV01000026 TGA|w
GC skew
CGC skew
AT skew
CAT skew
0.4
0.2
0.0
−0.2
−0.4
40
20
0
−20
−40
−60

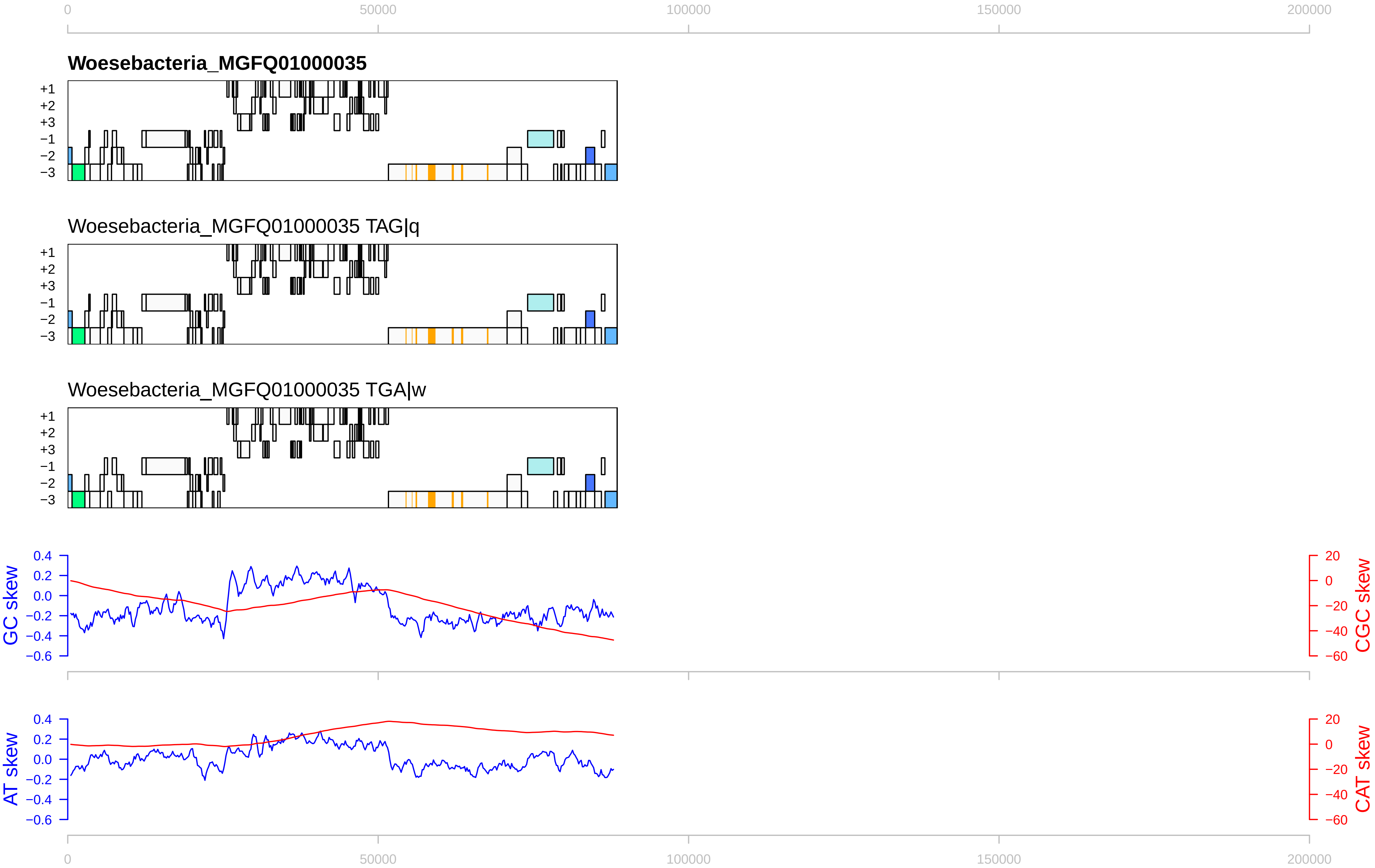

0
50000
100000
150000
200000
Woesebacteria_MGFQ01000035
+1
+2
+3
−1
−2
−3
Woesebacteria_MGFQ01000035 TAG|q
Woesebacteria_MGFQ01000035 TGA|w
GC skew
CGC skew
AT skew
CAT skew
0.4
0.2
0.0
−0.2
−0.4
−0.6
20
0
−20
−40
−60

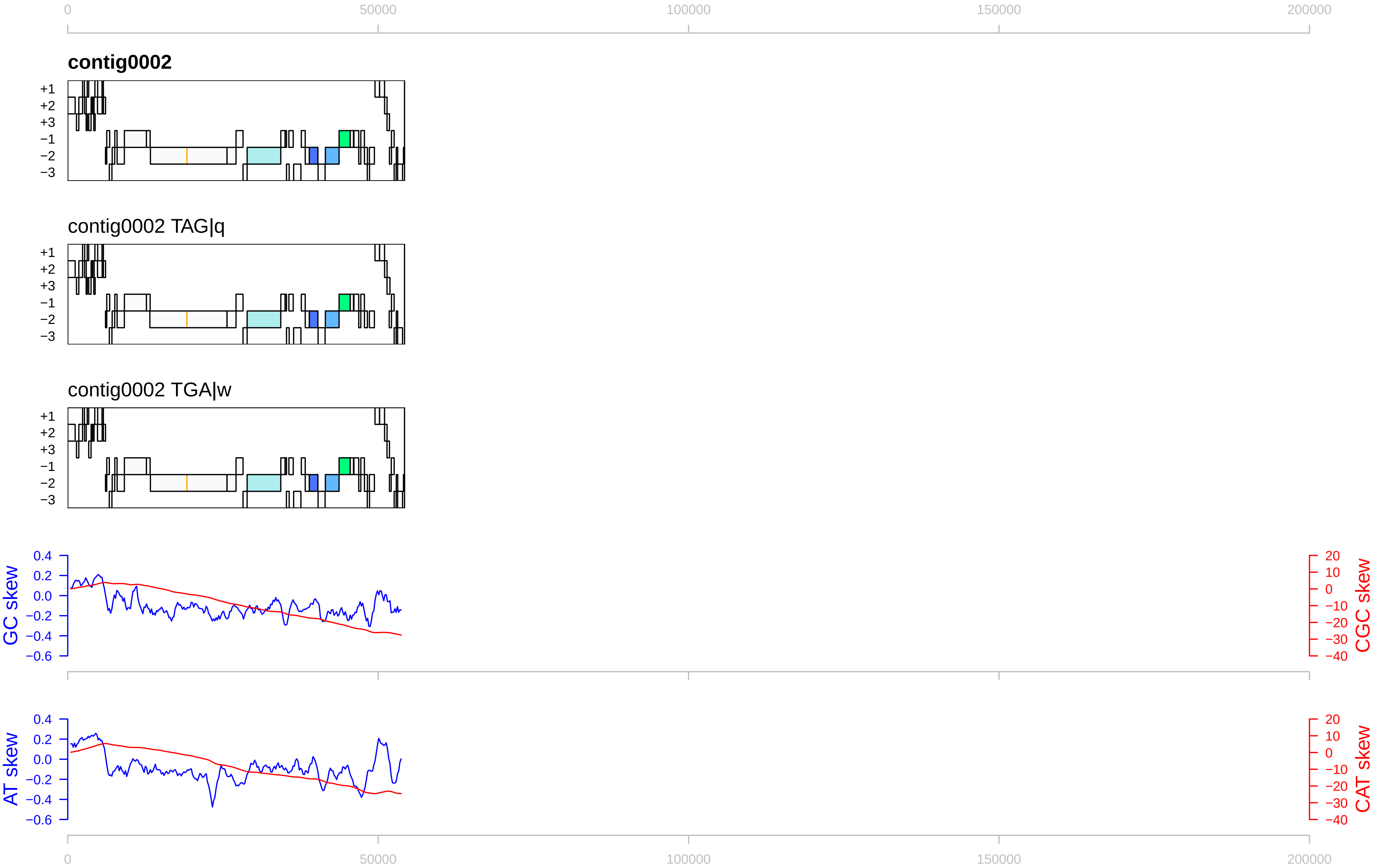

0
50000
100000
150000
200000
contig0002
+1
+2
+3
−1
−2
−3
contig0002 TAG|q
contig0002 TGA|w
GC skew
CGC skew
AT skew
CAT skew
0.4
0.2
0.0
−0.2
−0.4
−0.6
20
10
0
−10
−20
−30
−40

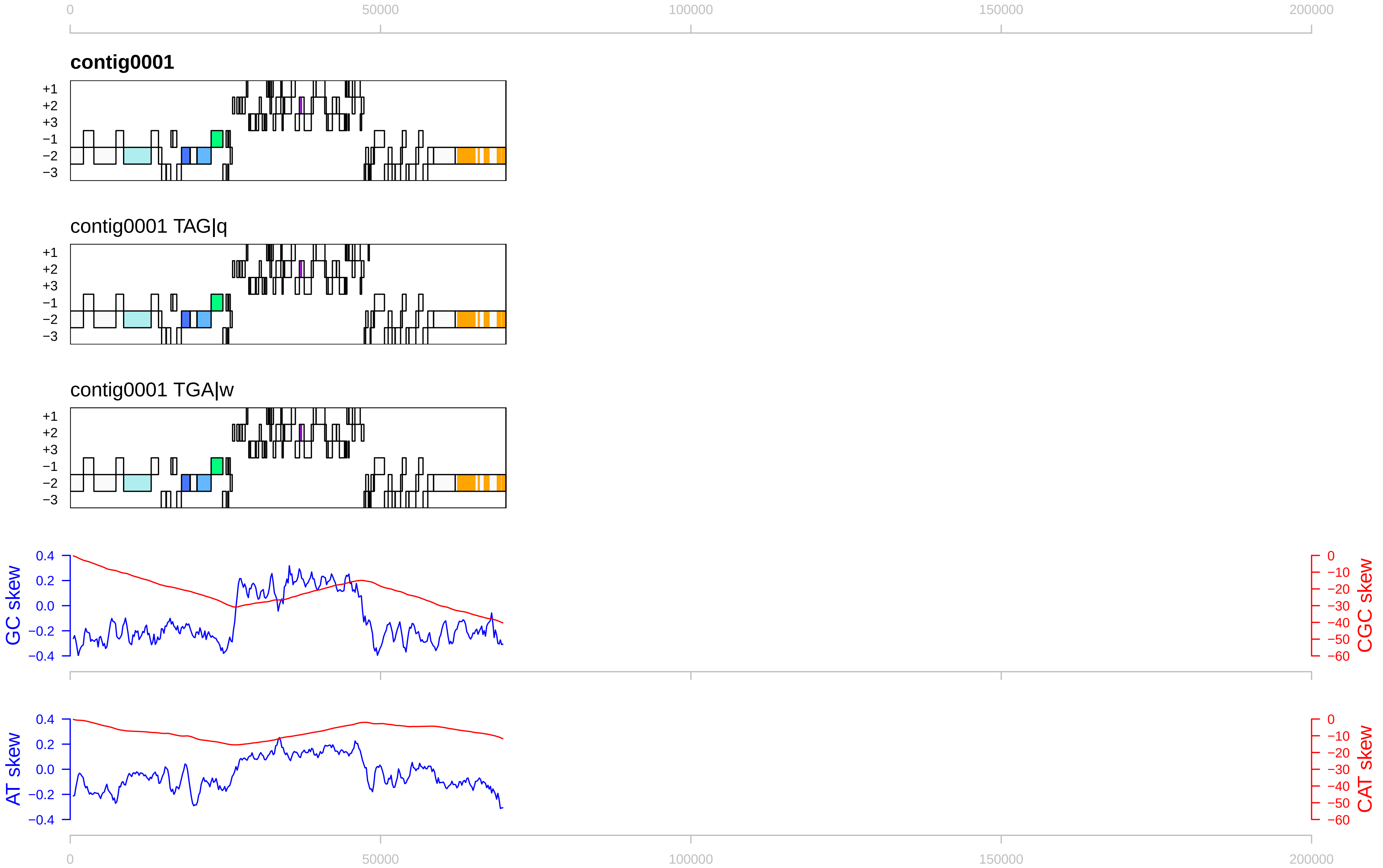

0
50000
100000
150000
200000
contig0001
contig0001 TAG|q
contig0001 TGA|w
+1
+2
+3
−1
−2
−3
GC skew
CGC skew
AT skew
CAT skew
0.4
0.2
0.0
−0.2
−0.4
0
−10
−20
−30
−40
−50
−60

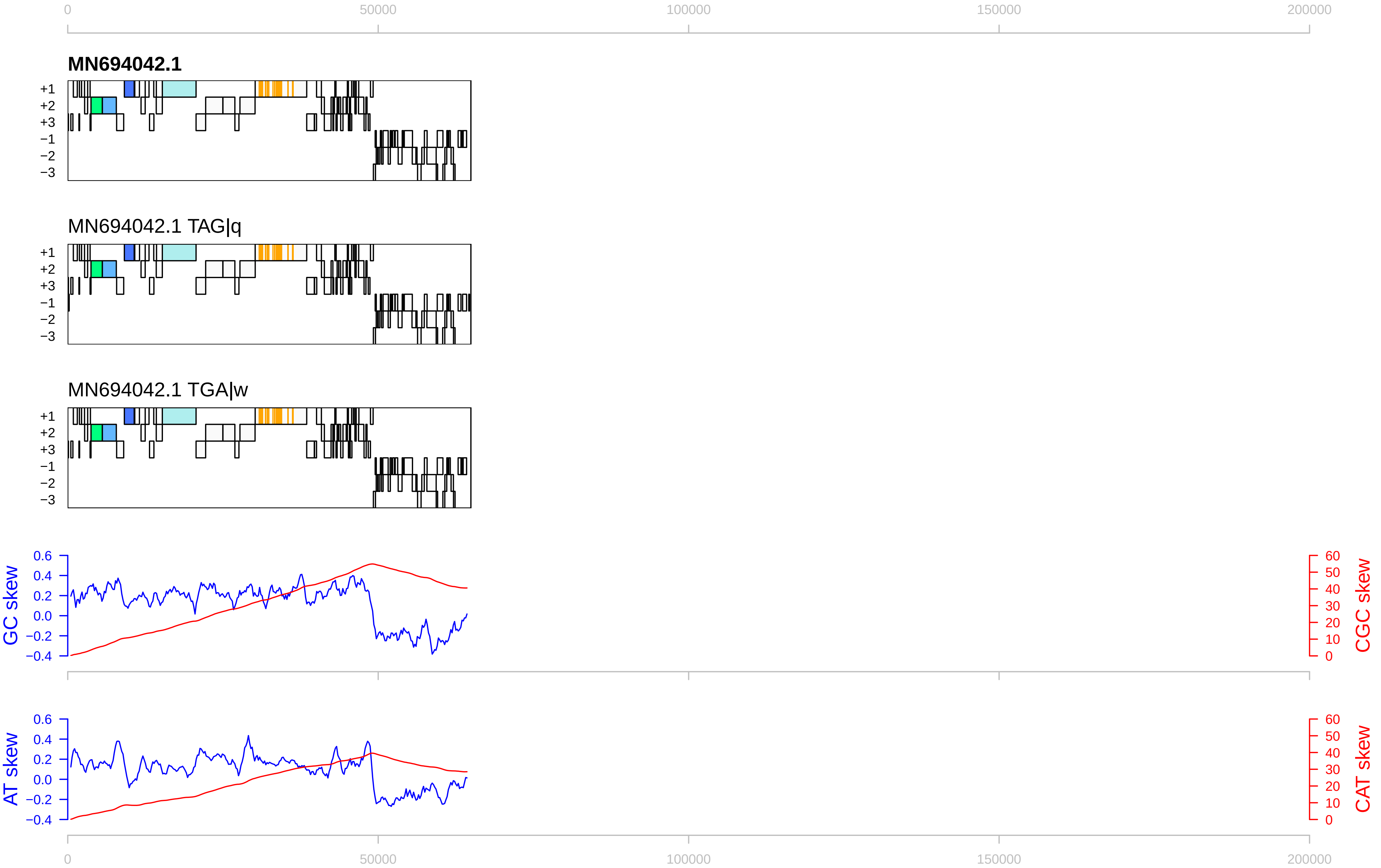

0
50000
100000
150000
200000
MN694042.1
+1
+2
+3
−1
−2
−3
MN694042.1 TAG|q
MN694042.1 TGA|w
GC skew
CGC skew
AT skew
CAT skew
0.6
0.4
0.2
0.0
−0.2
−0.4
60
50
40
30
20
10
0

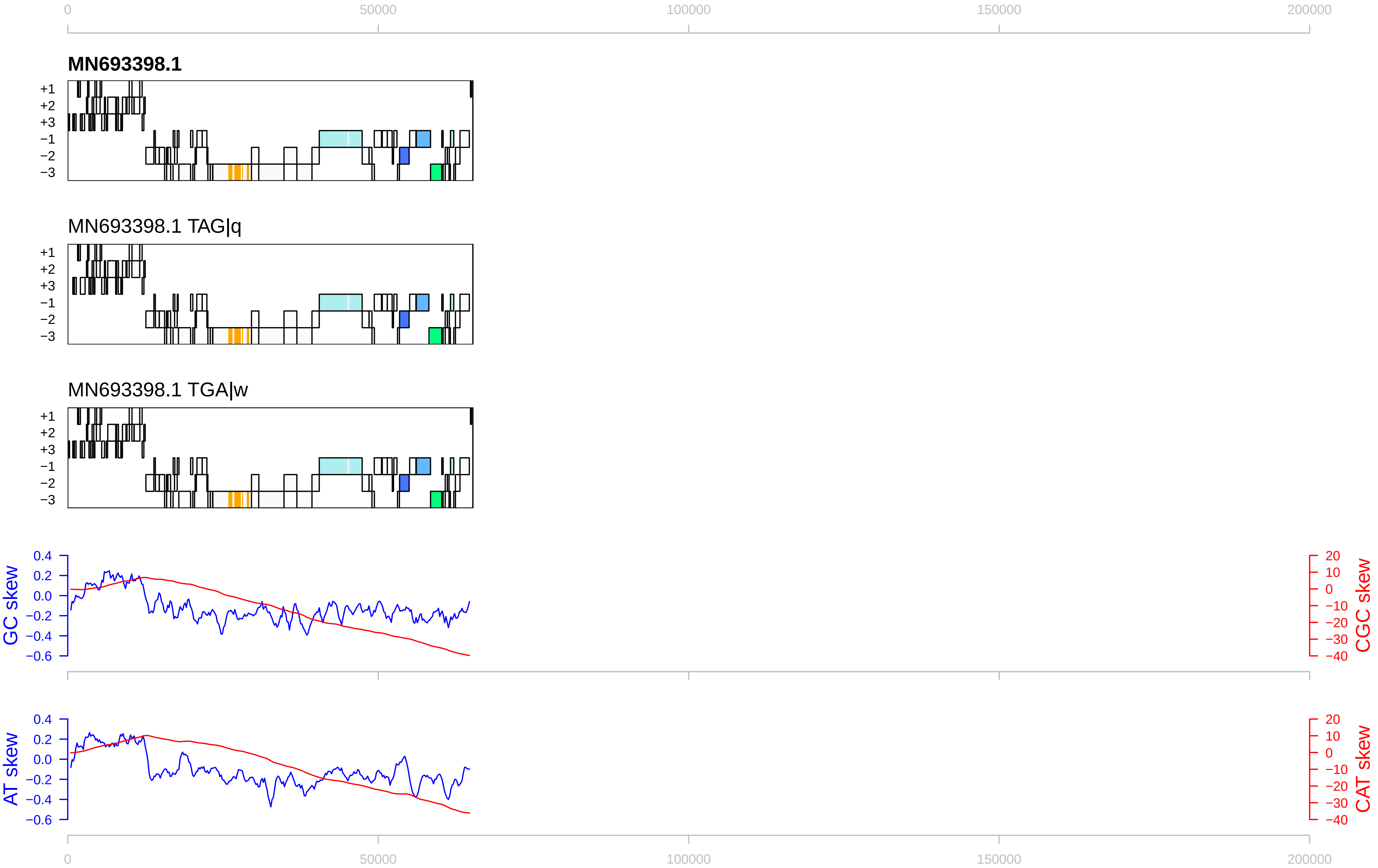
0
50000
100000
150000
200000
MN693398.1
+1
+2
+3
−1
−2
−3
MN693398.1 TAG|q
MN693398.1 TGA|w
GC skew
CGC skew
AT skew
CAT skew
0.4
0.2
0.0
−0.2
−0.4
−0.6
20
10
0
−10
−20
−30
−40

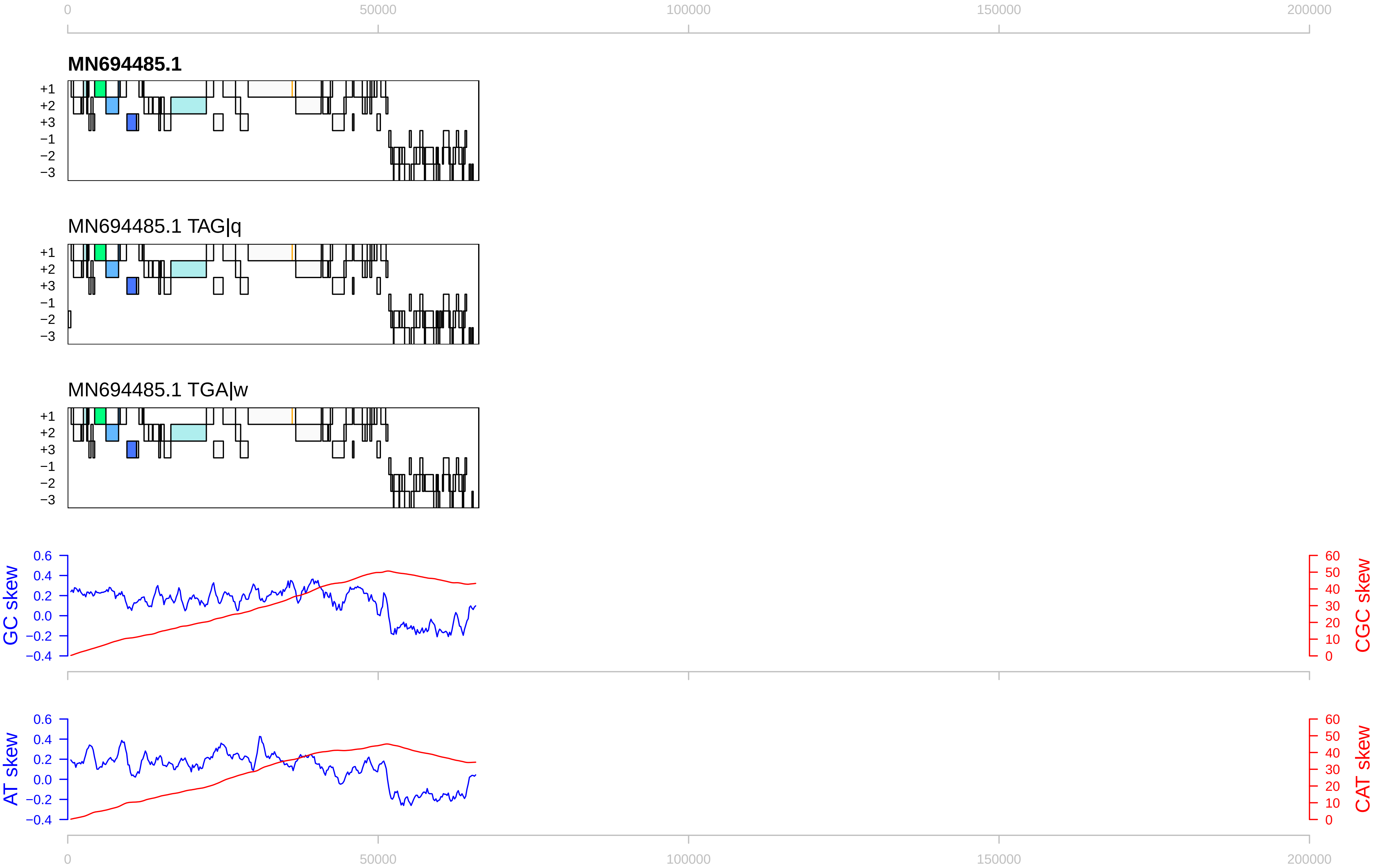

0
50000
100000
150000
200000
MN694485.1
+1
+2
+3
−1
−2
−3
MN694485.1 TAG|q
MN694485.1 TGA|w
GC skew
CGC skew
AT skew
CAT skew
0.6
0.4
0.2
0.0
−0.2
−0.4
60
50
40
30
20
10
0

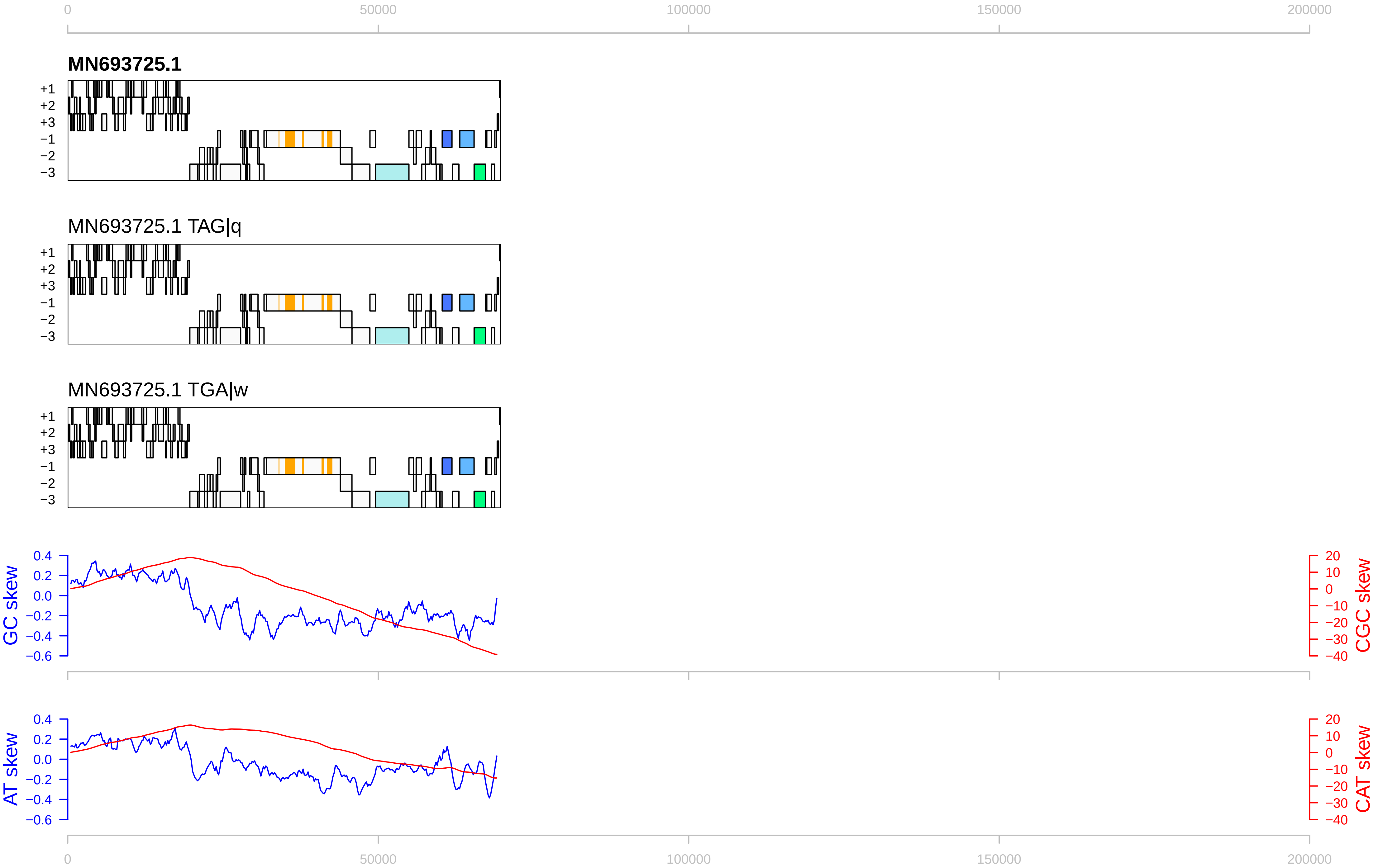

0
50000
100000
150000
200000
MN693725.1
+1
+2
+3
−1
−2
−3
MN693725.1 TAG|q
MN693725.1 TGA|w
GC skew
CGC skew
AT skew
CAT skew
0.4
0.2
0.0
−0.2
−0.4
−0.6
20
10
0
−10
−20
−30
−40

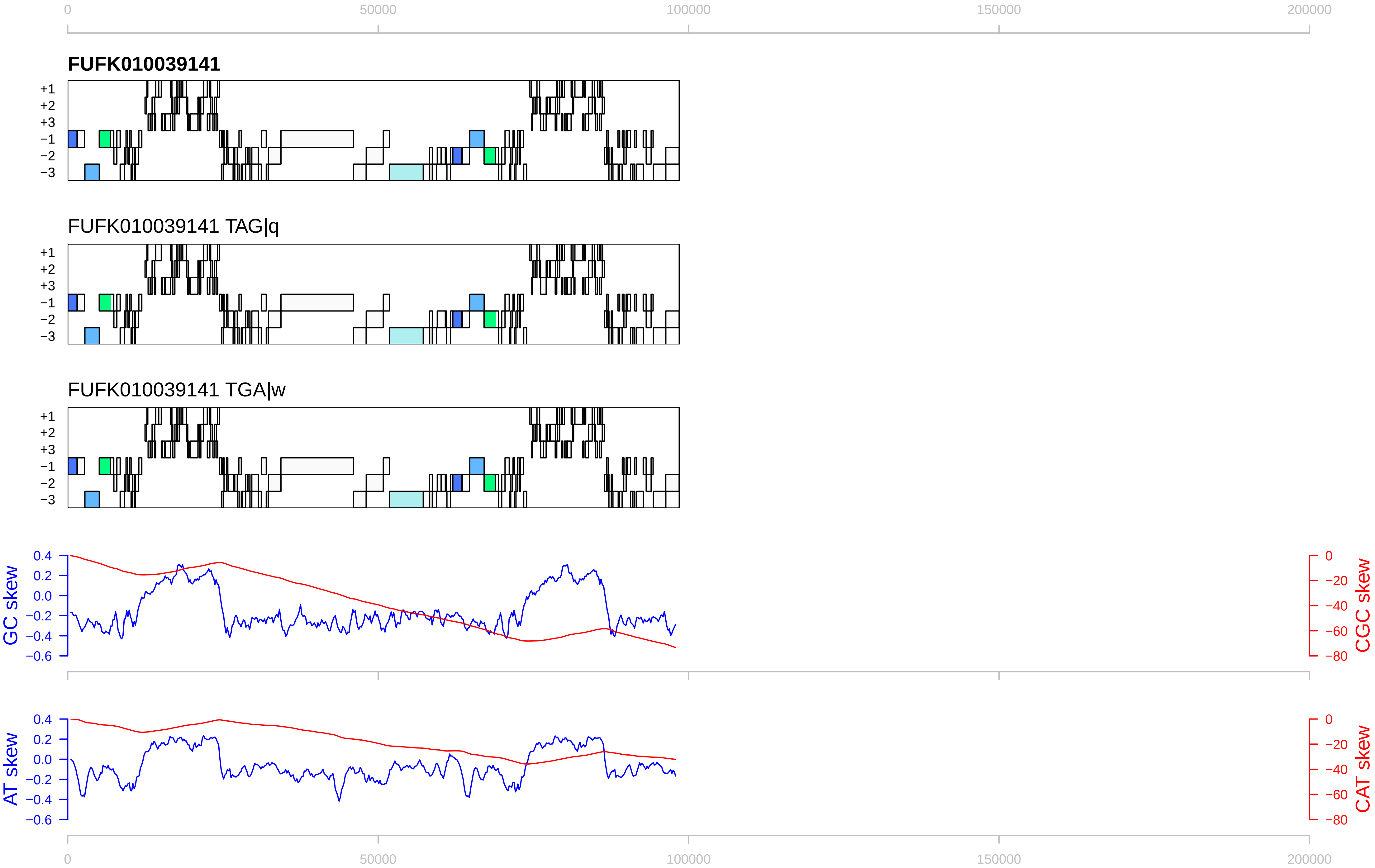

0
50000
100000
150000
200000
FUFK010039141
+1
+2
+3
−1
−2
−3
FUFK010039141 TAG|q
FUFK010039141 TGA|w
GC skew
CGC skew
AT skew
CAT skew
0.4
0.2
0.0
−0.2
−0.4
−0.6
0
−20
−40
−60
−80

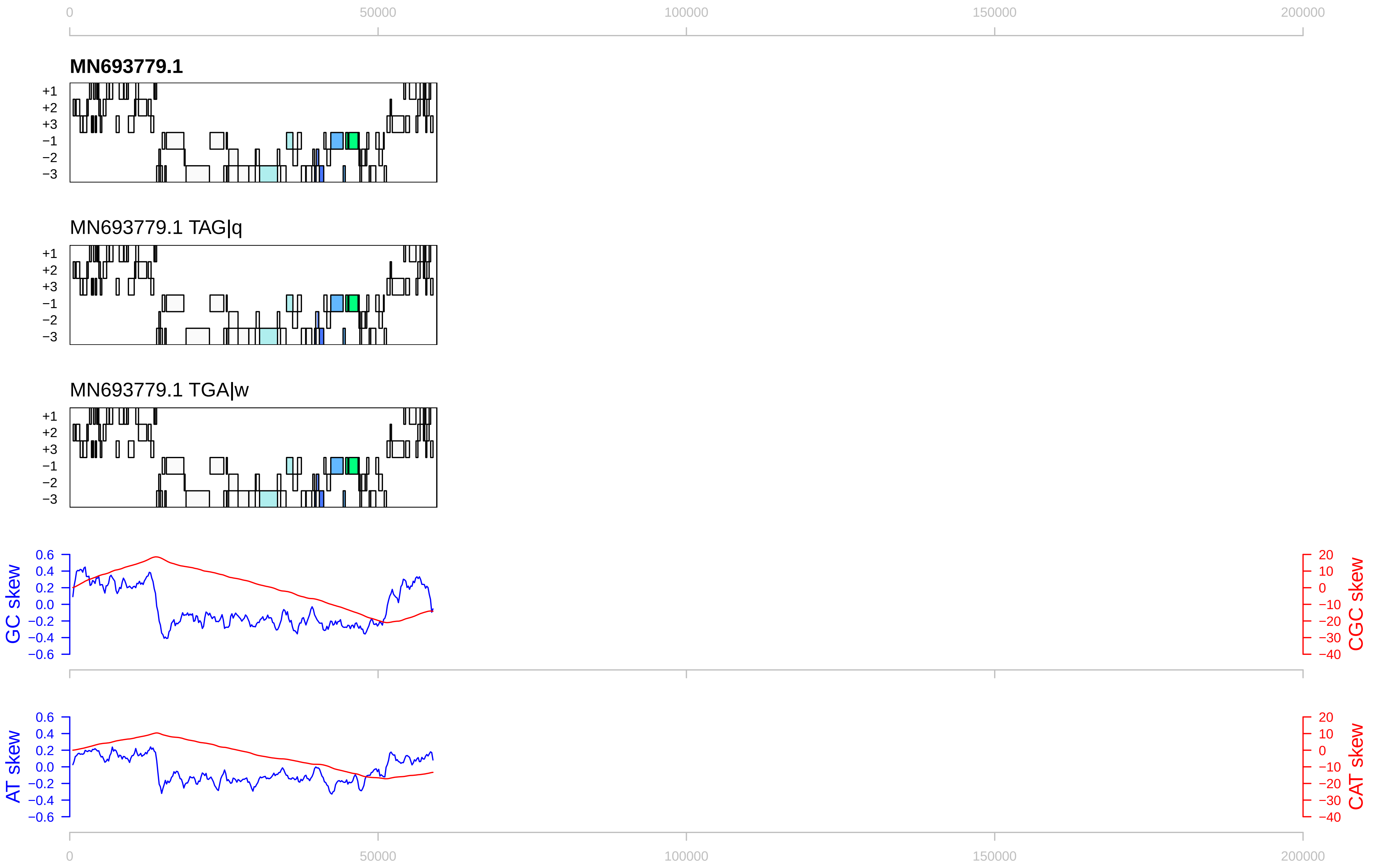

0
50000
100000
150000
200000
MN693779.1
+1
+2
+3
−1
−2
−3
MN693779.1 TAG|q
MN693779.1 TGA|w
GC skew
CGC skew
AT skew
CAT skew
0.6
0.4
0.2
0.0
−0.2
−0.4
−0.6
20
10
0
−10
−20
−30
−40

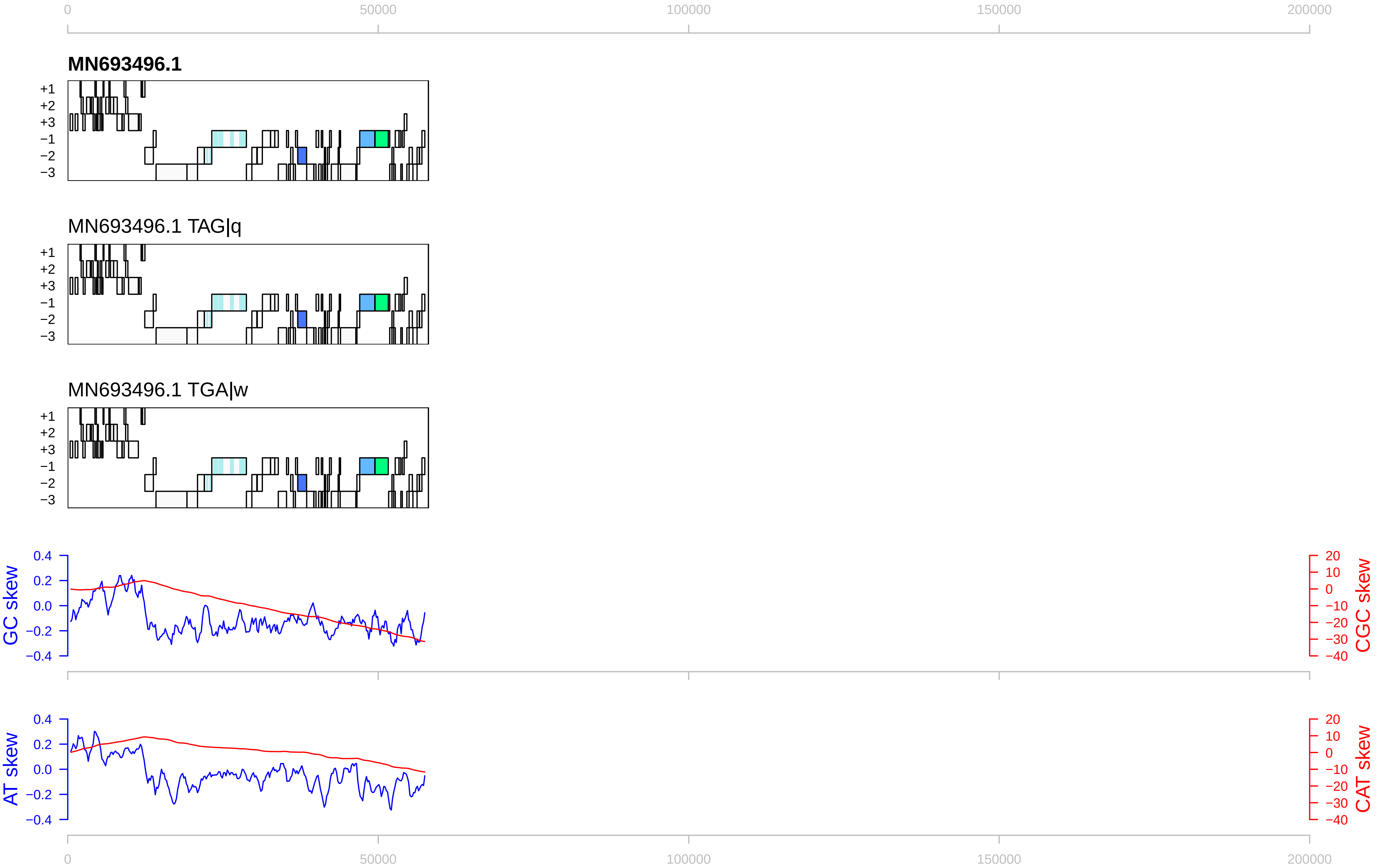
0
50000
100000
150000
200000
MN693496.1
+1
+2
+3
−1
−2
−3
MN693496.1 TAG|q
MN693496.1 TGA|w
GC skew
CGC skew
AT skew
CAT skew
0.4
0.2
0.0
−0.2
−0.4
20
10
0
−10
−20
−30
−40

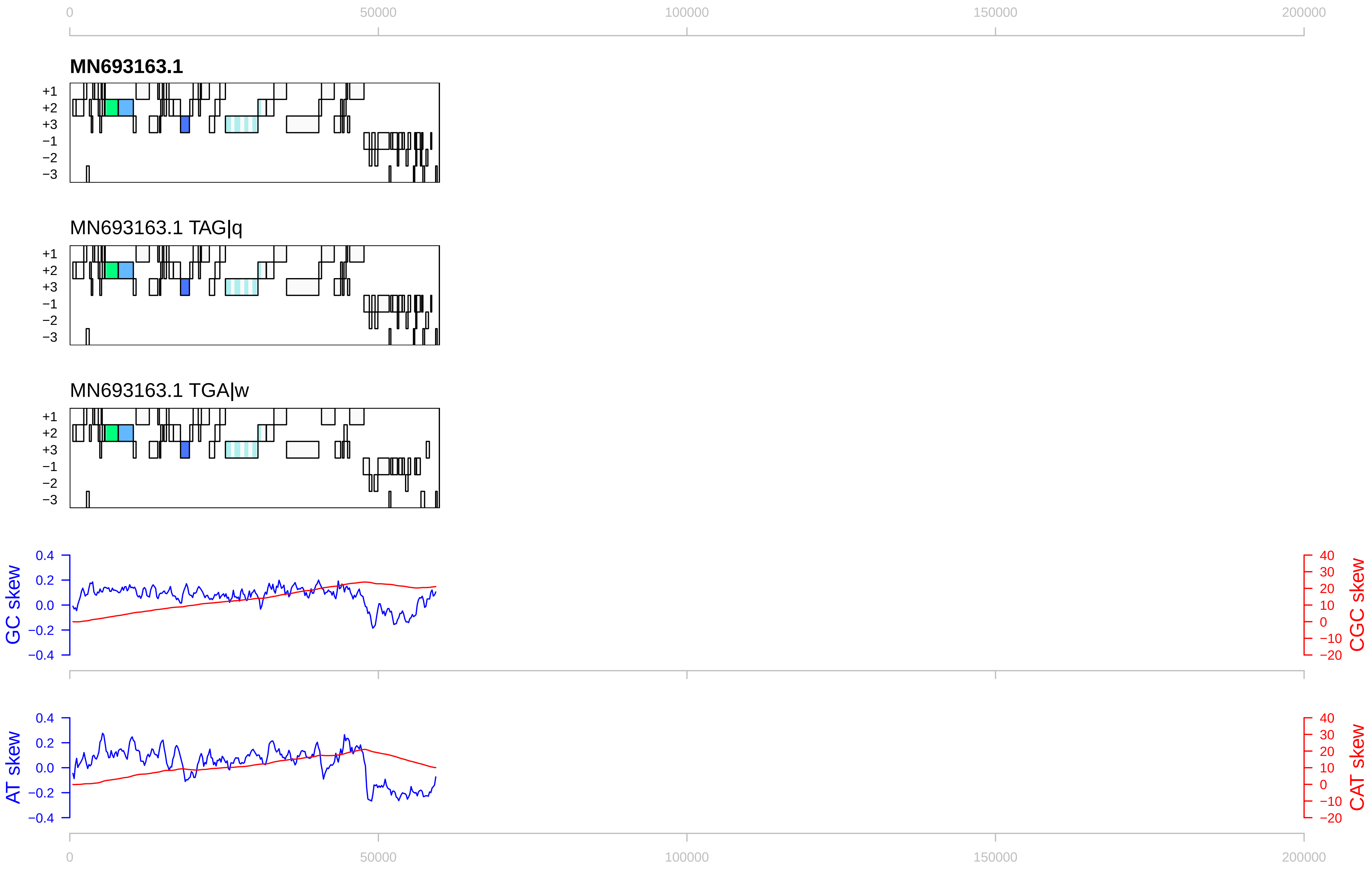

0
50000
100000
150000
200000
MN693163.1
+1
+2
+3
−1
−2
−3
MN693163.1 TAG|q
MN693163.1 TGA|w
GC skew
CGC skew
AT skew
CAT skew
0.4
0.2
0.0
−0.2
−0.4
40
30
20
10
0
−10
−20

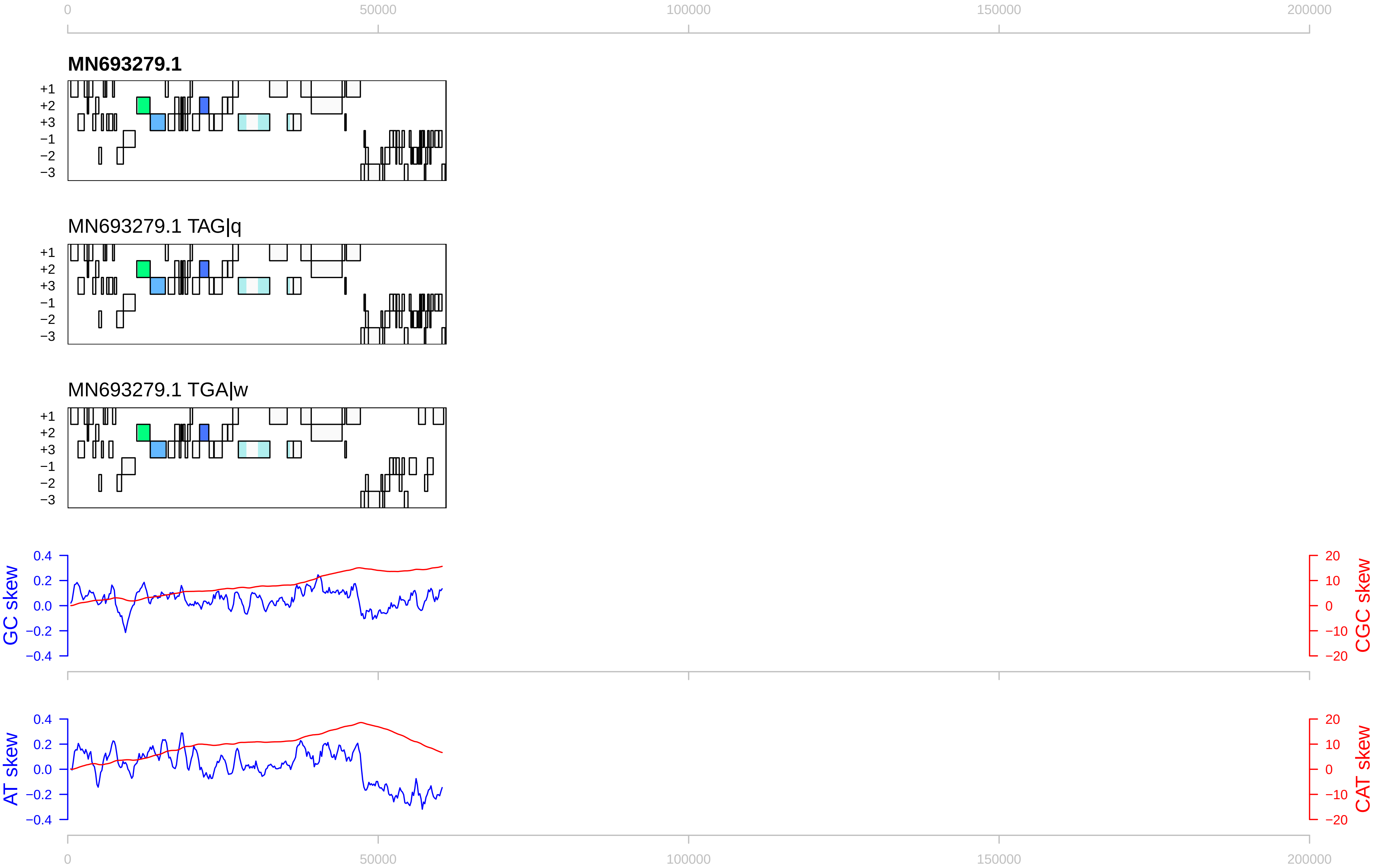
0
50000
100000
150000
200000
MN693279.1
+1
+2
+3
−1
−2
−3
MN693279.1 TAG|q
MN693279.1 TGA|w
GC skew
CGC skew
0.4
0.2
0.0
−0.2
−0.4
20
10
0
−10
−20
AT skew
CAT skew

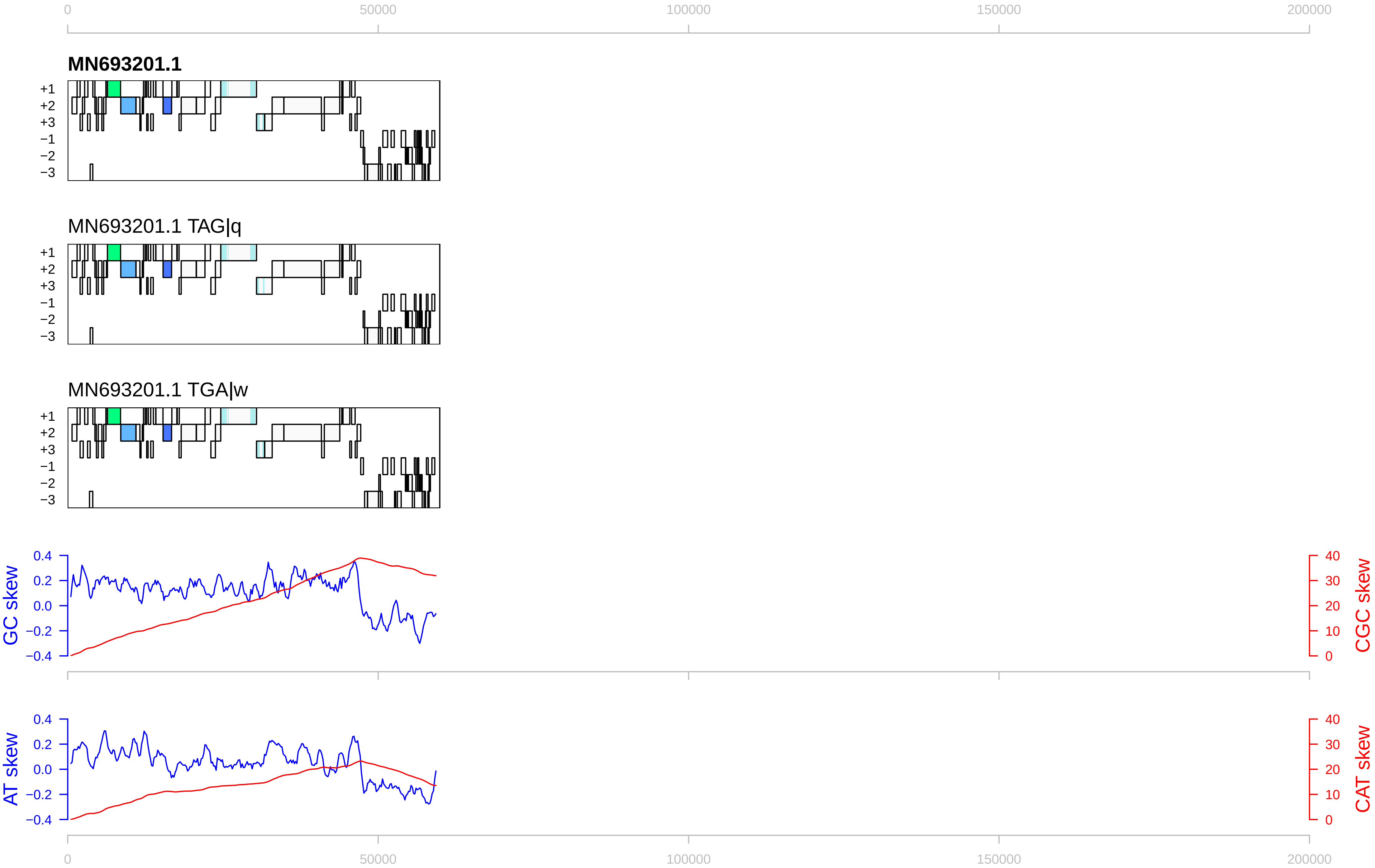

0
50000
100000
150000
200000
MN693201.1
+1
+2
+3
−1
−2
−3
MN693201.1 TAG|q
MN693201.1 TGA|w
GC skew
CGC skew
AT skew
CAT skew
0.4
0.2
0.0
−0.2
−0.4
40
30
20
10
0

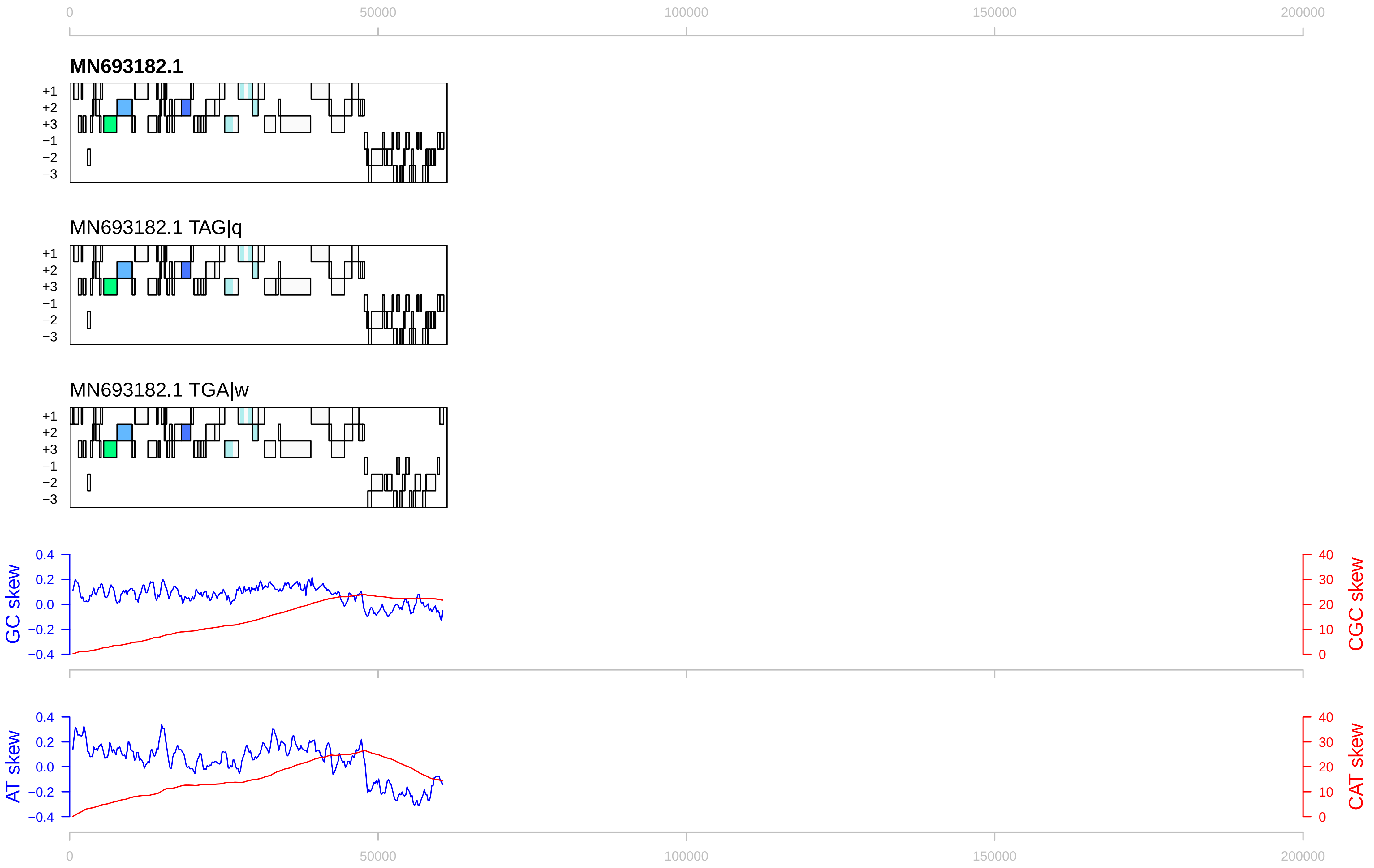

0
50000
100000
150000
200000
MN693182.1
+1
+2
+3
−1
−2
−3
MN693182.1 TAG|q
MN693182.1 TGA|w
GC skew
CGC skew
AT skew
CAT skew
0.4
0.2
0.0
−0.2
−0.4
40
30
20
10
0

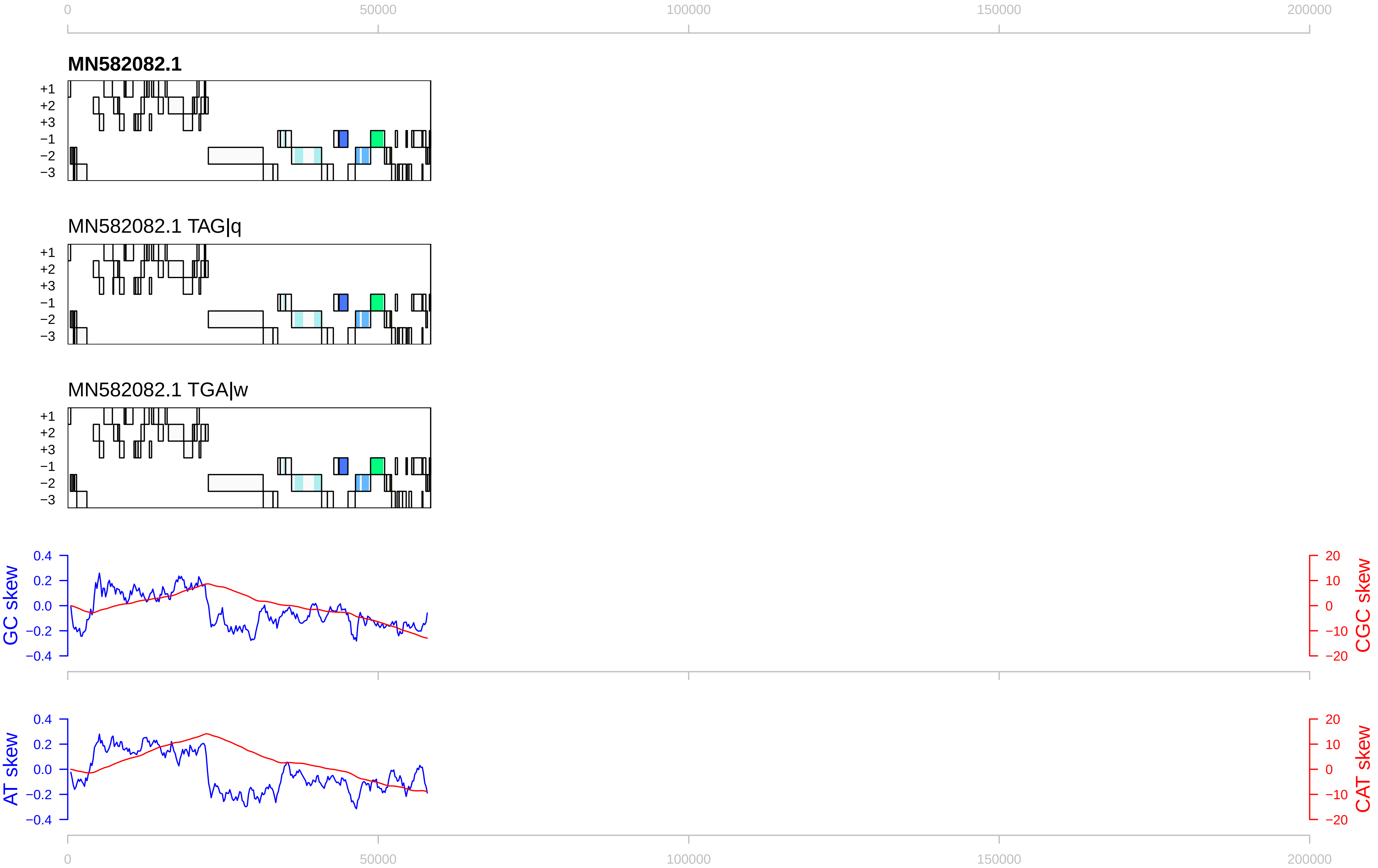

0
50000
100000
150000
200000
MN582082.1
+1
+2
+3
−1
−2
−3
MN582082.1 TAG|q
MN582082.1 TGA|w
GC skew
CGC skew
AT skew
CAT skew
0.4
0.2
0.0
−0.2
−0.4
20
10
0
−10
−20

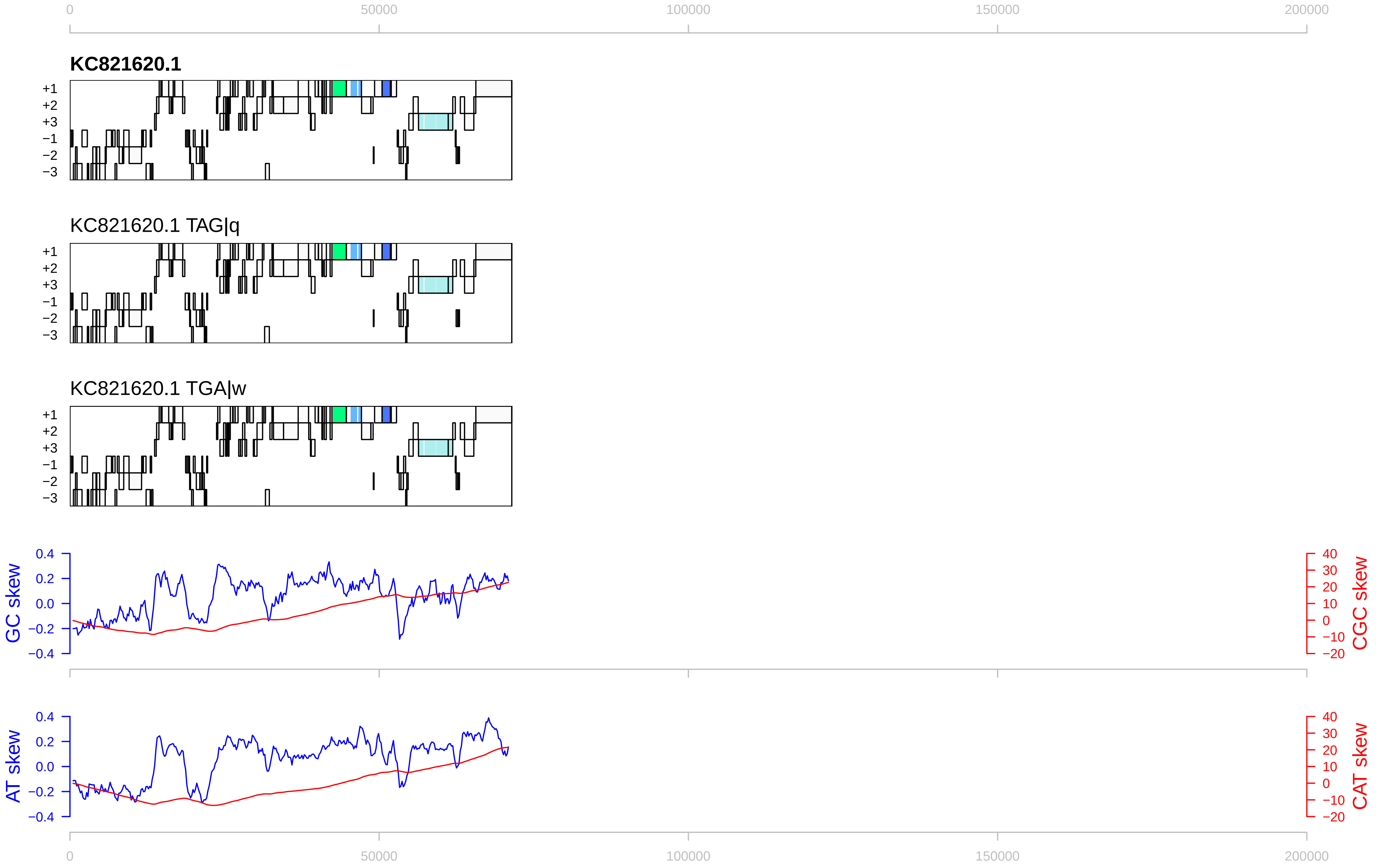

0
50000
100000
150000
200000
KC821620.1
KC821620.1 TAG|q
KC821620.1 TGA|w
+1
+2
+3
−1
−2
−3
GC skew
CGC skew
AT skew
CAT skew
0.4
0.2
0.0
−0.2
−0.4
40
30
20
10
0
−10
−20

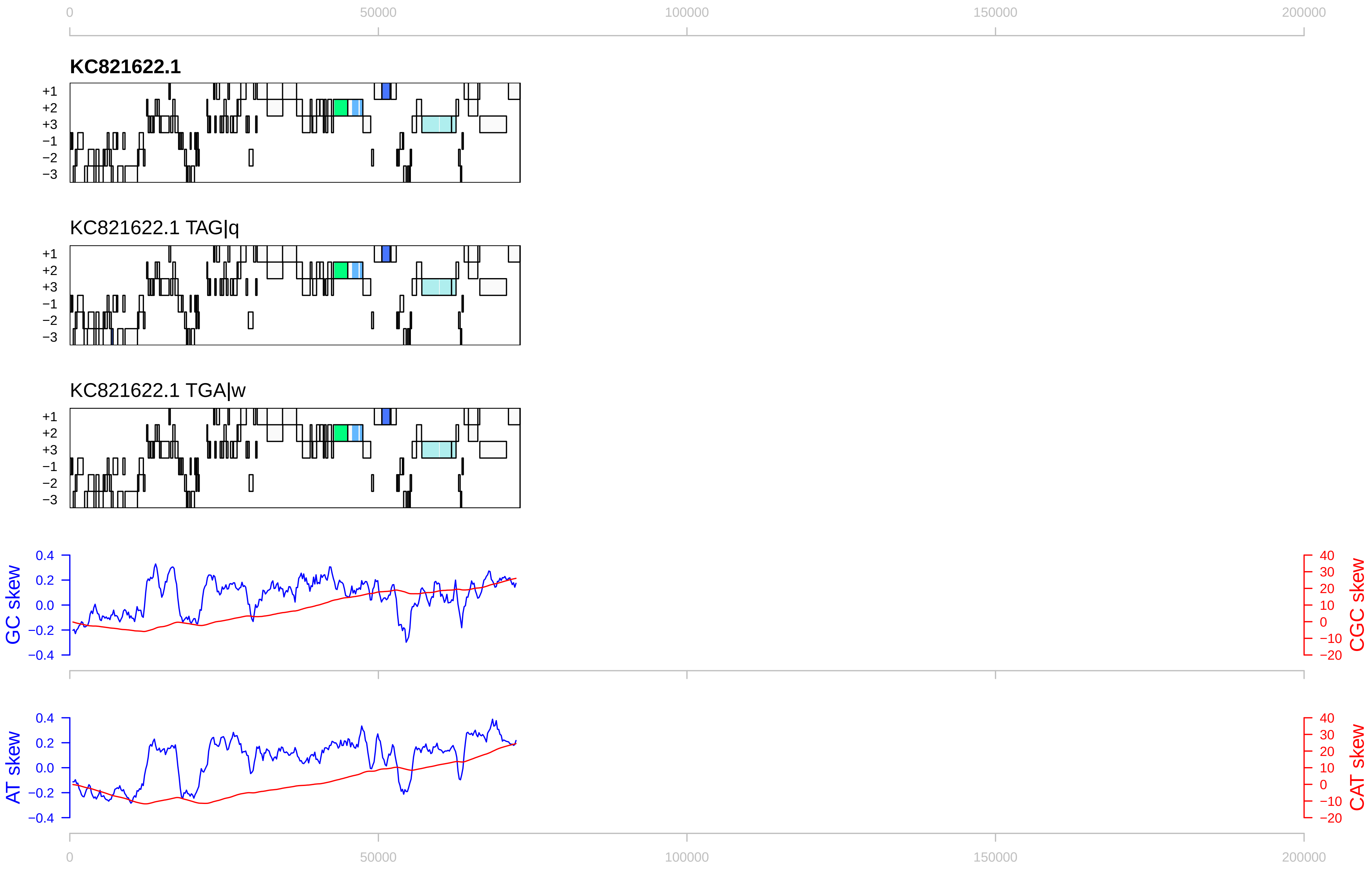

0
50000
100000
150000
200000
KC821622.1
+1
+2
+3
−1
−2
−3
KC821622.1 TAG|q
KC821622.1 TGA|w
GC skew
CGC skew
AT skew
CAT skew
0.4
0.2
0.0
−0.2
−0.4
40
30
20
10
0
−10
−20

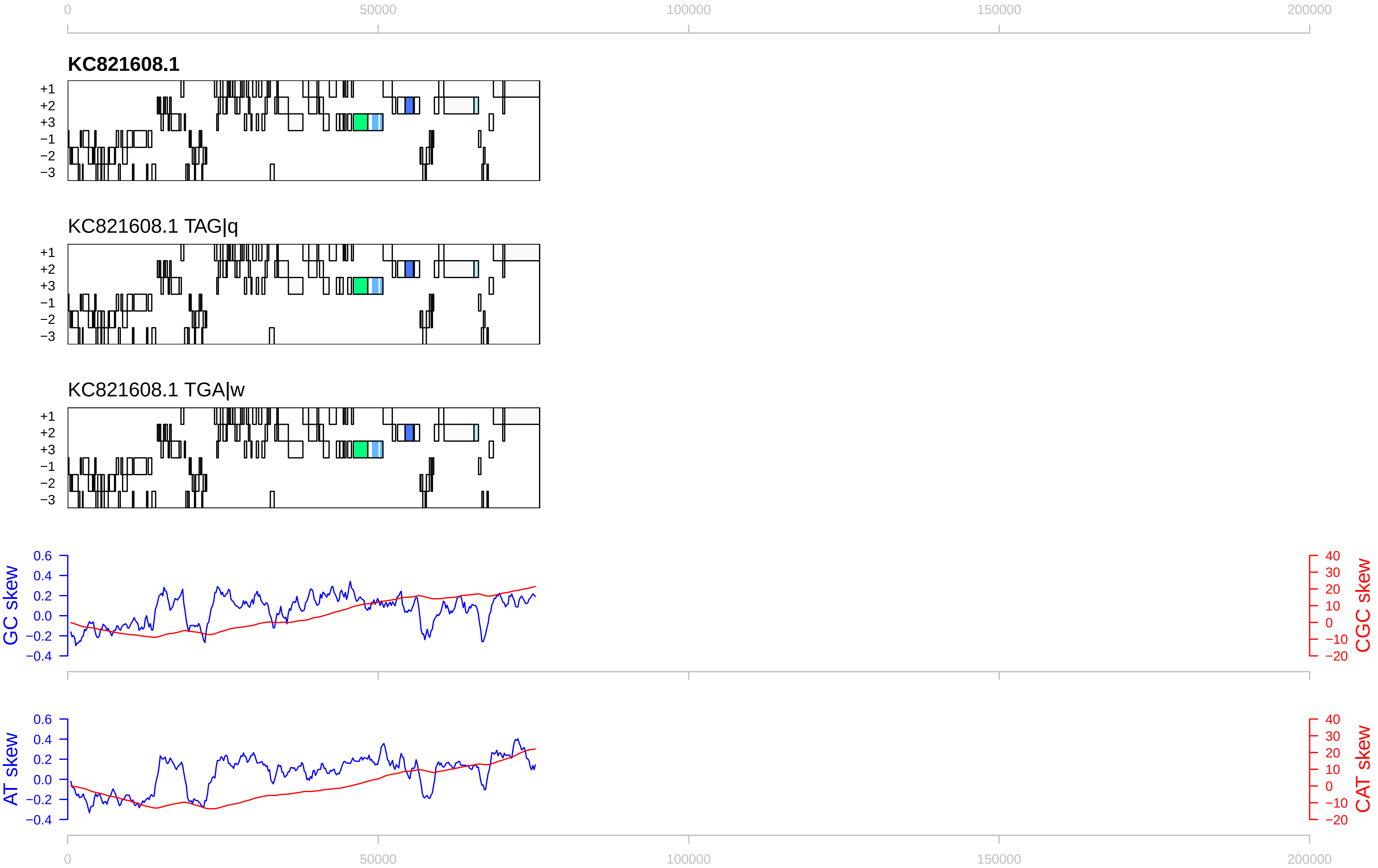

0
50000
100000
150000
200000
KC821608.1
+1
+2
+3
−1
−2
−3
KC821608.1 TAG|q
KC821608.1 TGA|w
GC skew
0.6
0.4
0.2
0.0
−0.2
−0.4
CGC skew
40
30
20
10
0
−10
−20
AT skew
CAT skew

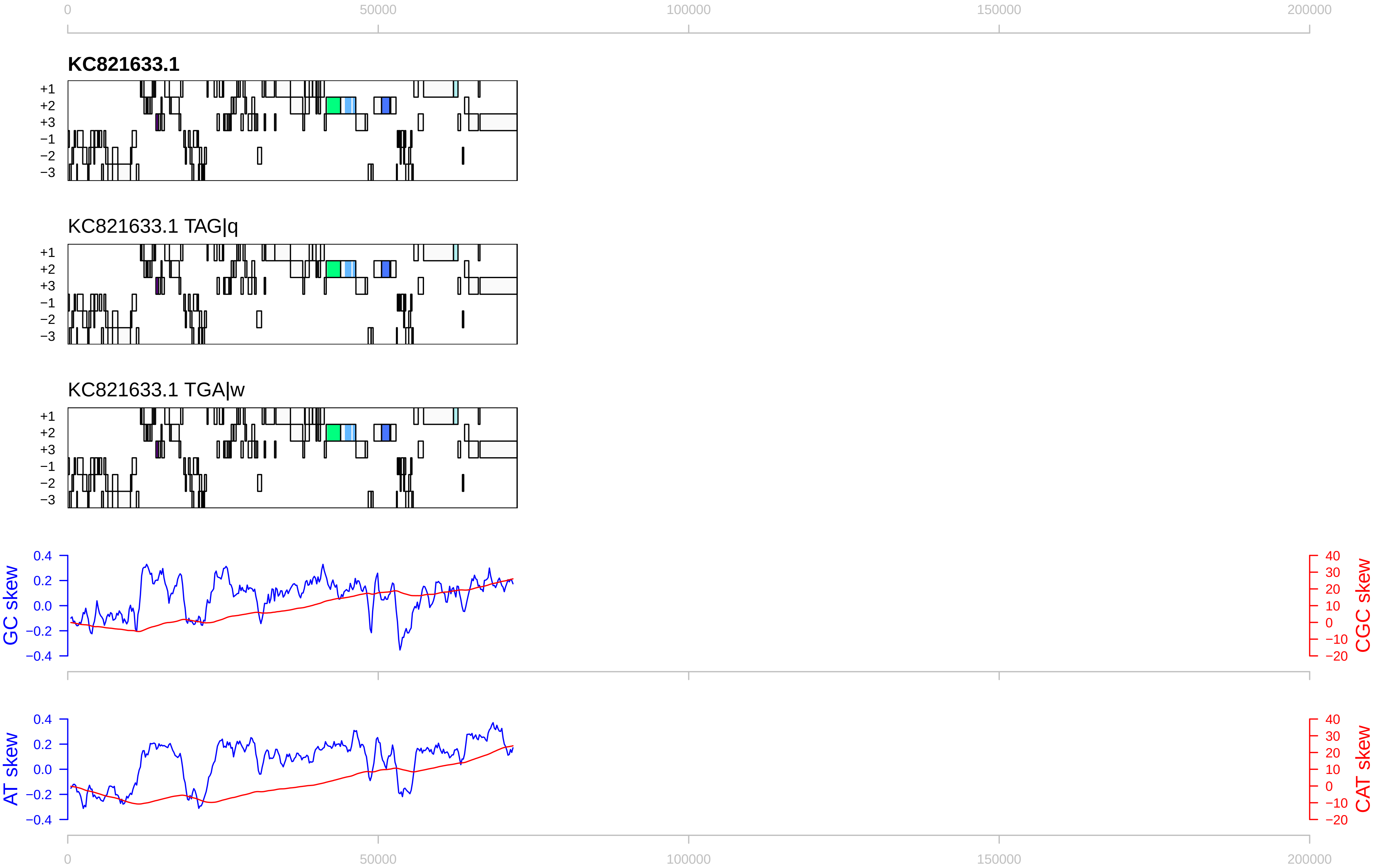

0
50000
100000
150000
200000
KC821633.1
KC821633.1 TAG|q
KC821633.1 TGA|w
+1
+2
+3
−1
−2
−3
GC skew
CGC skew
AT skew
CAT skew
0.4
0.2
0.0
−0.2
−0.4
40
30
20
10
0
−10
−20

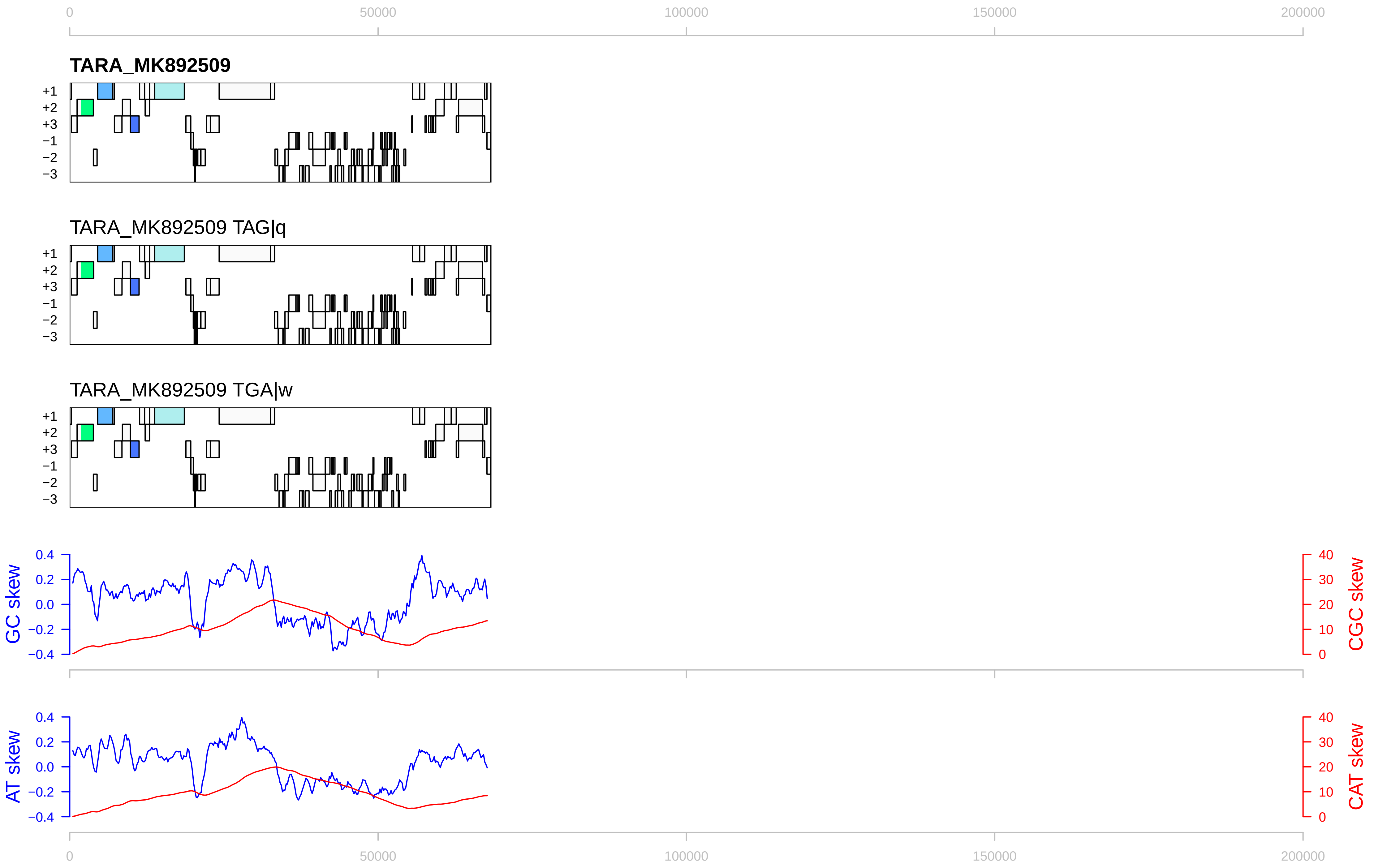

0
50000
100000
150000
200000
TARA_MK892509
+1
+2
+3
−1
−2
−3
TARA_MK892509 TAG|q
TARA_MK892509 TGA|w
GC skew
CGC skew
0.4
0.2
0.0
−0.2
−0.4
40
30
20
10
0
AT skew
CAT skew

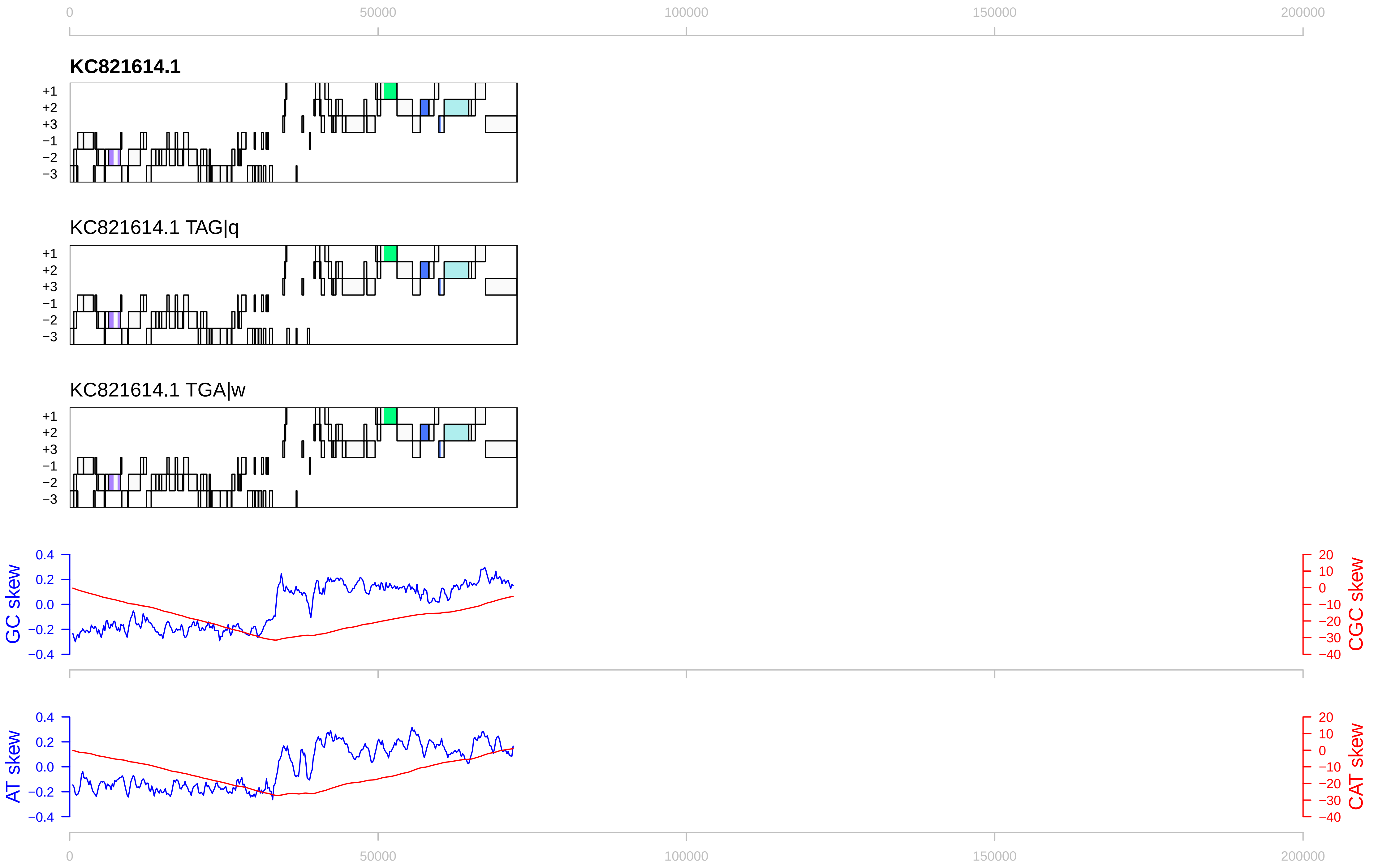
0
50000
100000
150000
200000
KC821614.1
+1
+2
+3
−1
−2
−3
KC821614.1 TAG|q
KC821614.1 TGA|w
GC skew
0.4
0.2
0.0
−0.2
−0.4
CGC skew
20
10
0
−10
−20
−30
−40
AT skew
CAT skew

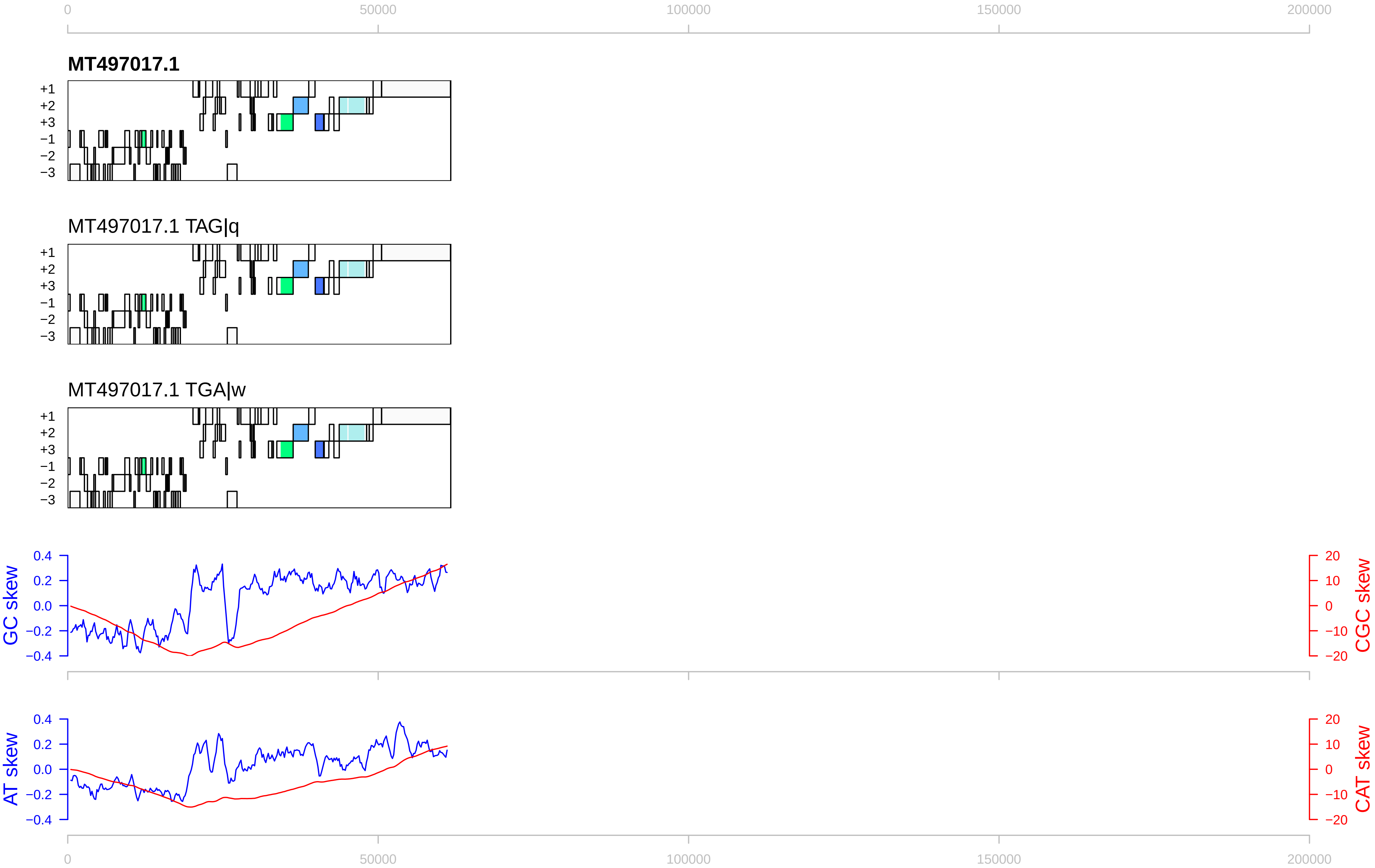
0
50000
100000
150000
200000
MT497017.1
+1
+2
+3
−1
−2
−3
MT497017.1 TAG|q
MT497017.1 TGA|w
GC skew
CGC skew
0.4
0.2
0.0
−0.2
−0.4
20
10
0
−10
−20
AT skew
CAT skew

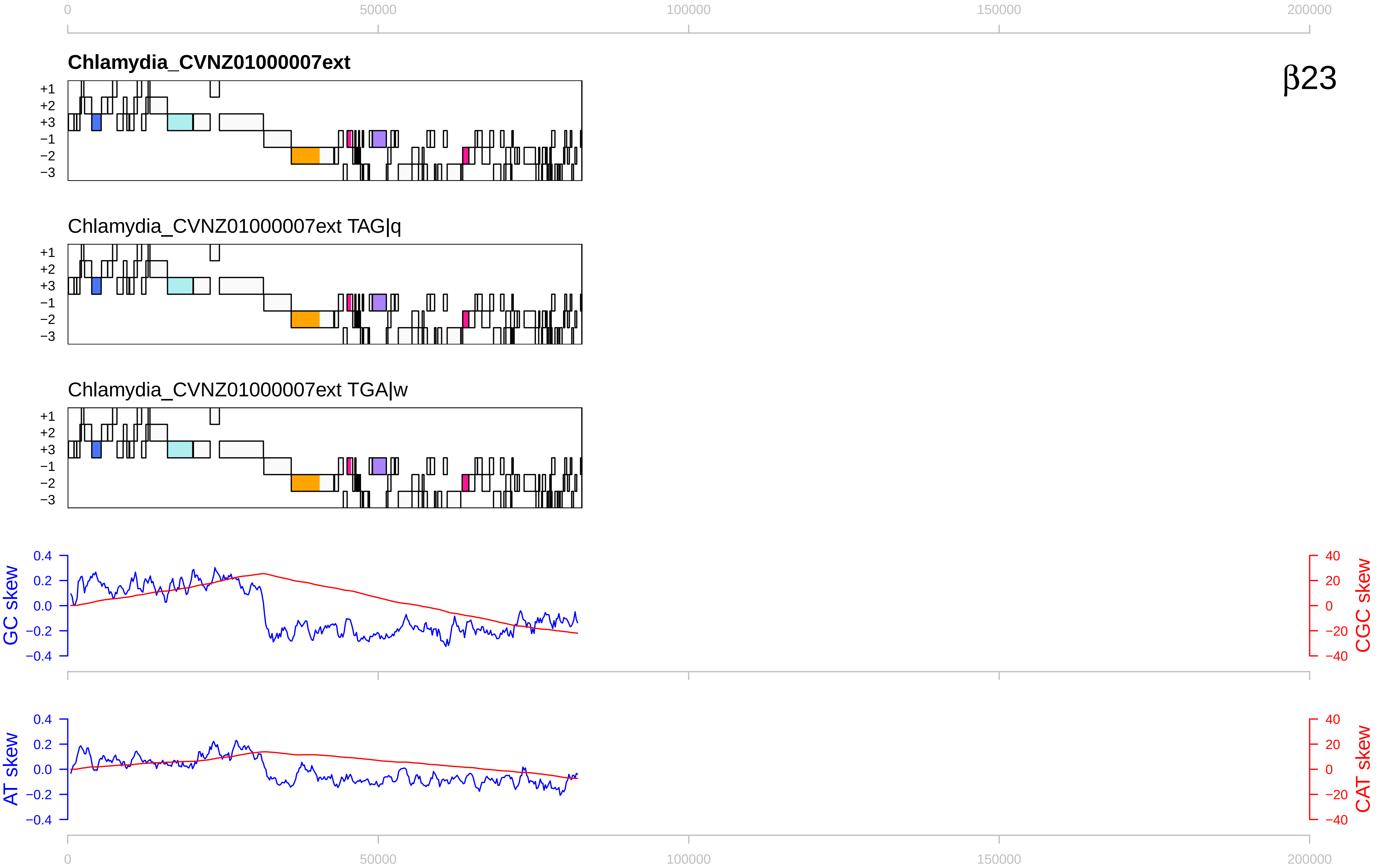

0
50000
100000
150000
200000
Chlamydia_CVNZ01000007ext
β23
+1
+2
+3
−1
−2
−3
Chlamydia_CVNZ01000007ext TAG|q
Chlamydia_CVNZ01000007ext TGA|w
GC skew
CGC skew
AT skew
CAT skew
0.4
0.2
0.0
−0.2
−0.4
40
20
0
−20
−40

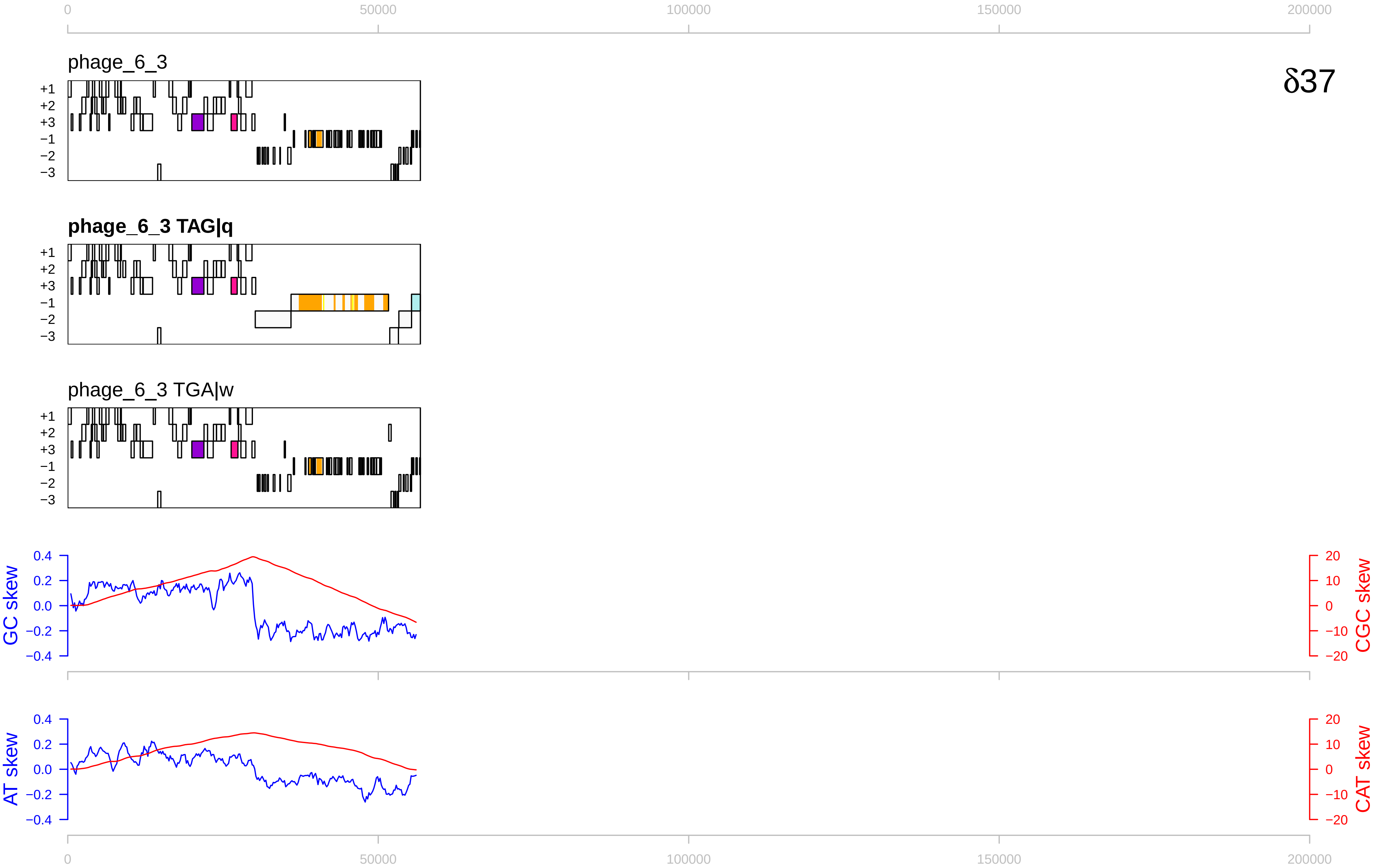

0
50000
100000
150000
200000
phage_6_3
δ37
+1
+2
+3
−1
−2
−3
phage_6_3 TAG|q
phage_6_3 TGA|w
GC skew
CGC skew
0.4
0.2
0.0
−0.2
−0.4
20
10
0
−10
−20
AT skew
CAT skew

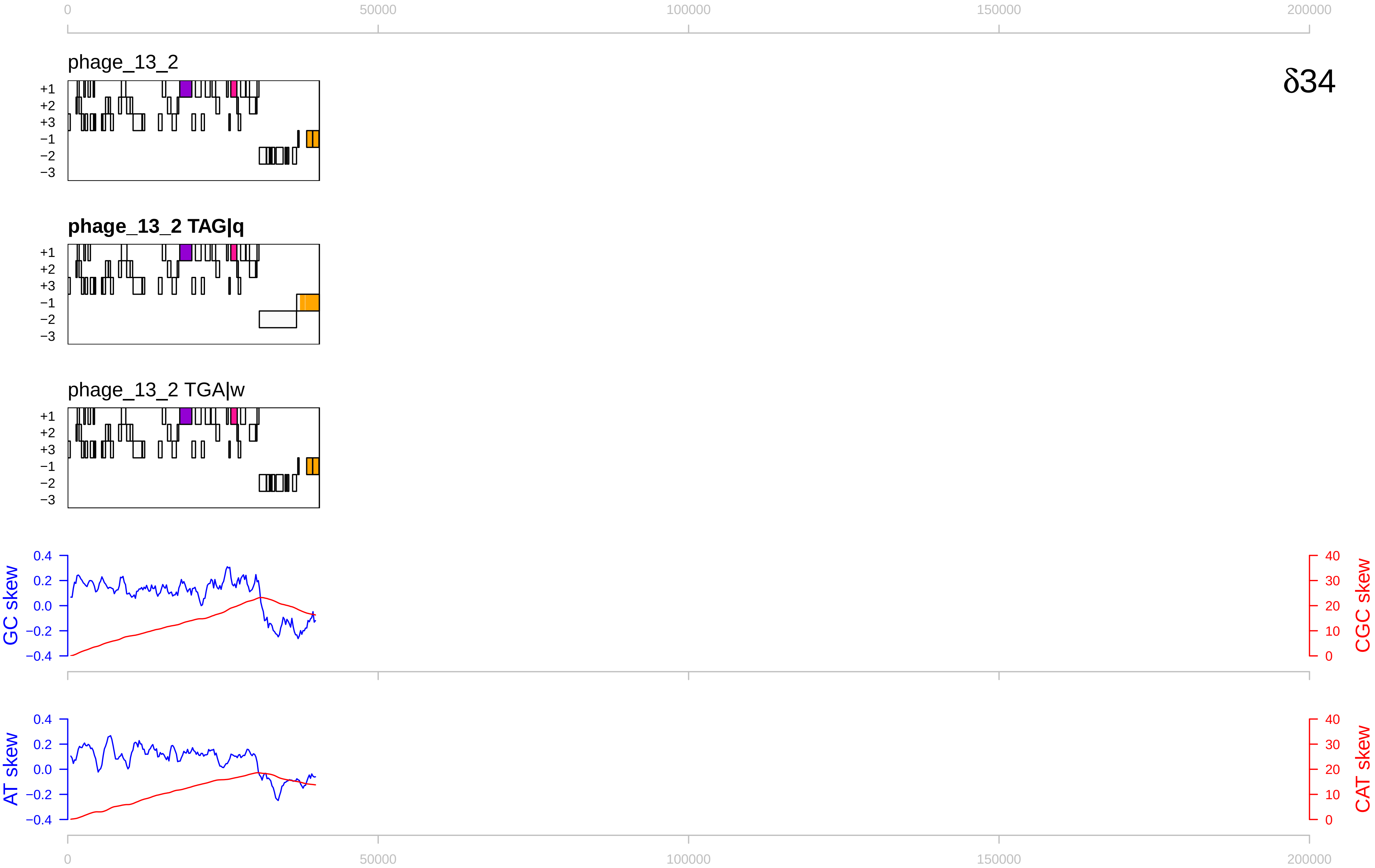

0
50000
100000
150000
200000
phage_13_2
δ34
+1
+2
+3
−1
−2
−3
phage_13_2 TAG|q
phage_13_2 TGA|w
GC skew
CGC skew
0.4
0.2
0.0
−0.2
−0.4
40
30
20
10
AT skew
CAT skew

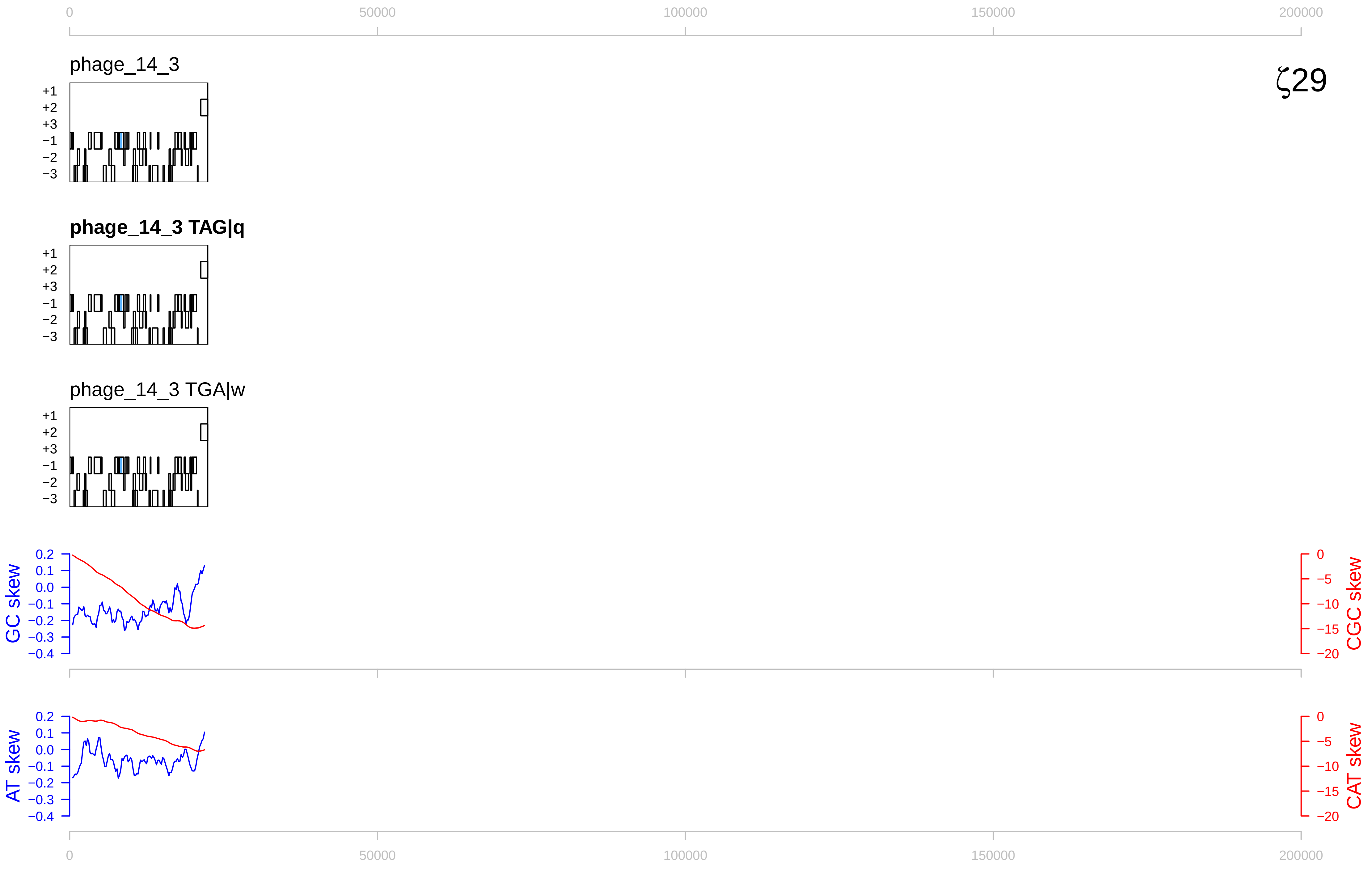

0
50000
100000
150000
200000
phage_14_3
ζ29
+1
+2
+3
−1
−2
−3
phage_14_3 TAG|q
phage_14_3 TGA|w
GC skew
CGC skew
AT skew
CAT skew
0.2
0.1
0.0
−0.1
−0.2
−0.3
−0.4
−5
−10
−15
−20

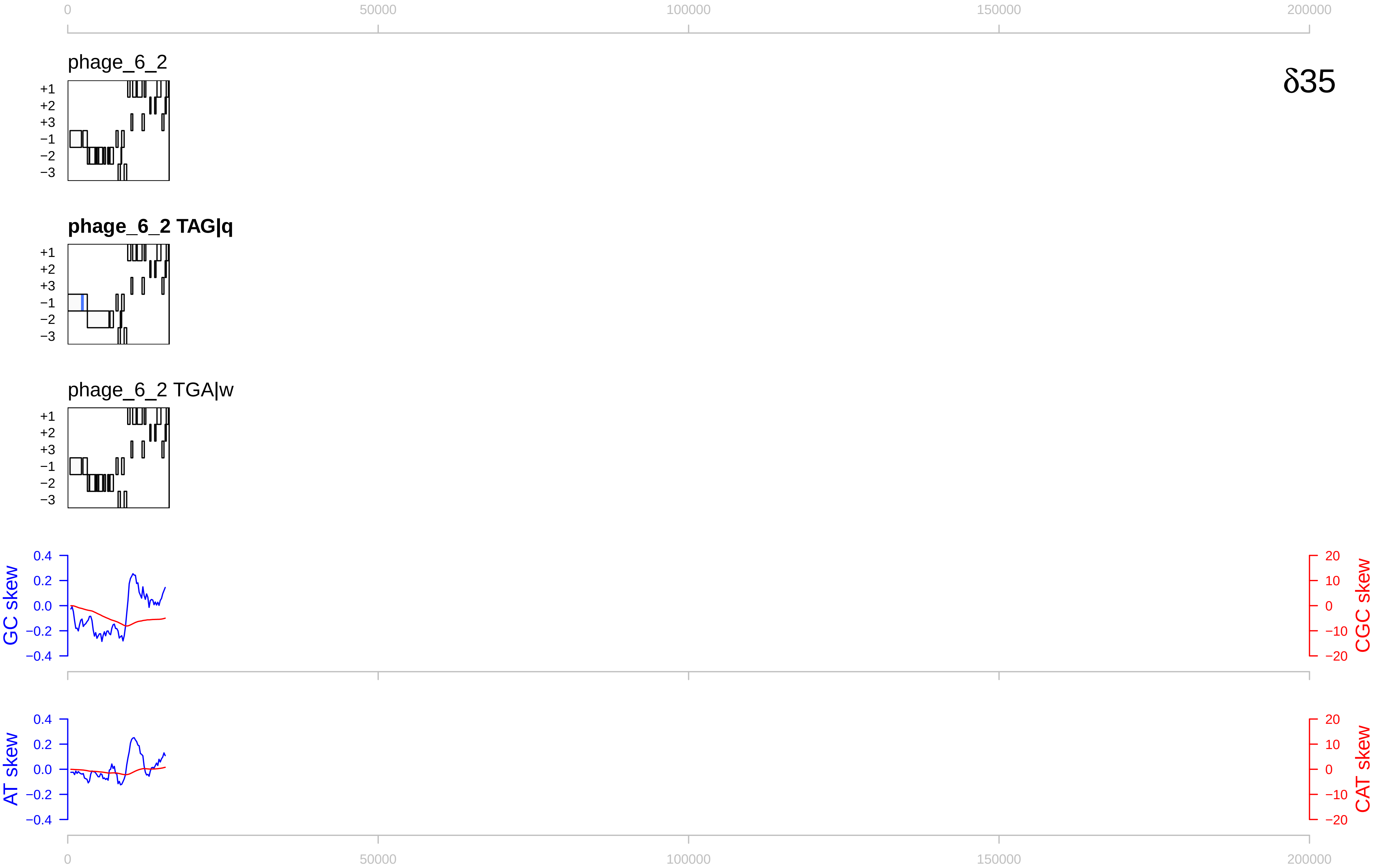
0
50000
100000
150000
200000
phage_6_2
δ35
+1
+2
+3
−1
−2
−3
phage_6_2 TAG|q
phage_6_2 TGA|w
GC skew
CGC skew
AT skew
CAT skew
0.4
0.2
0.0
−0.2
−0.4
20
10
0
−10
−20

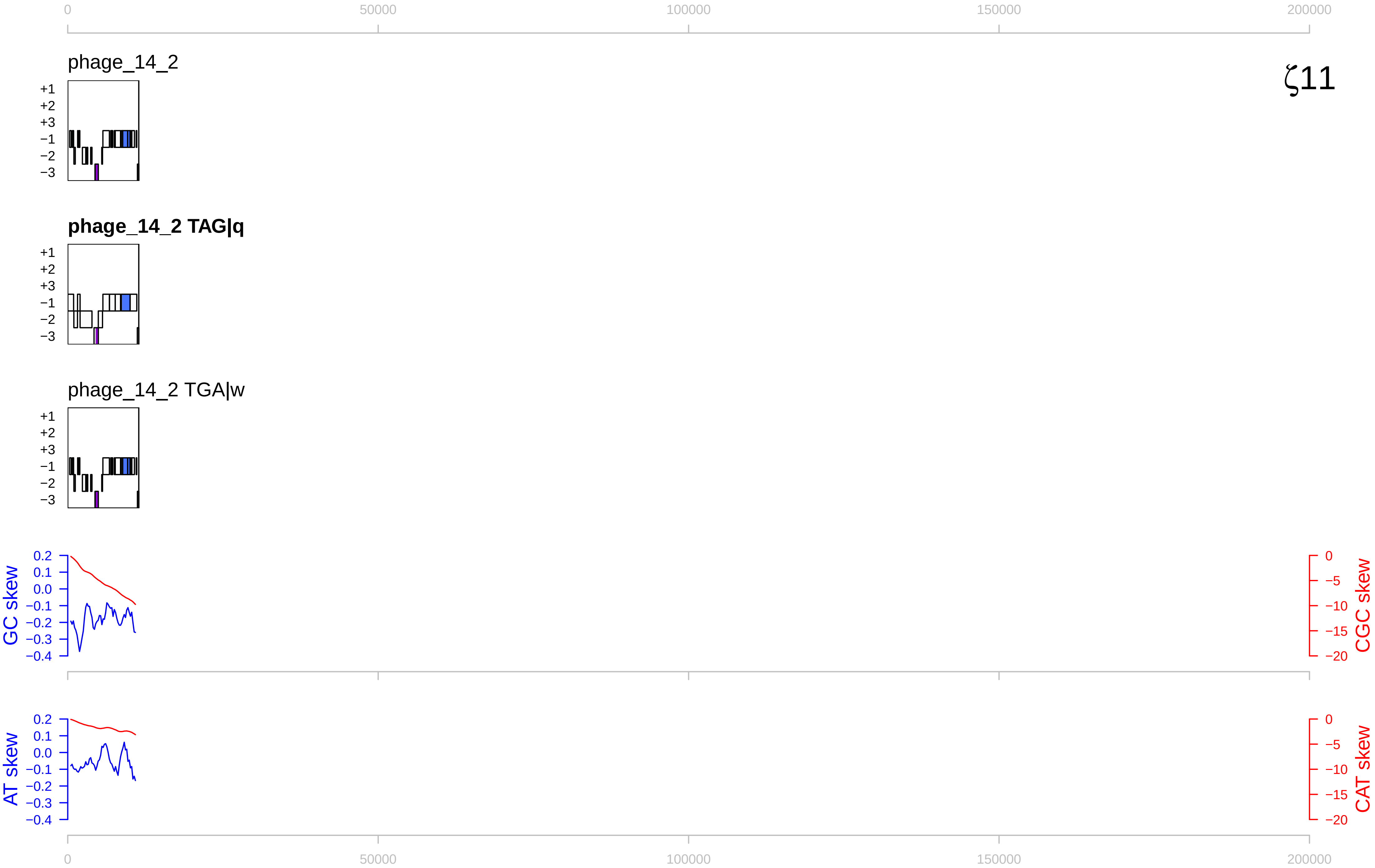

0
50000
100000
150000
200000
phage_14_2
ζ11
+1
+2
+3
−1
−2
−3
phage_14_2 TAG|q
phage_14_2 TGA|w
GC skew
CGC skew
0.2
0.1
0.0
−0.1
−0.2
−0.3
−0.4
0
−5
−10
−15
−20
AT skew
CAT skew

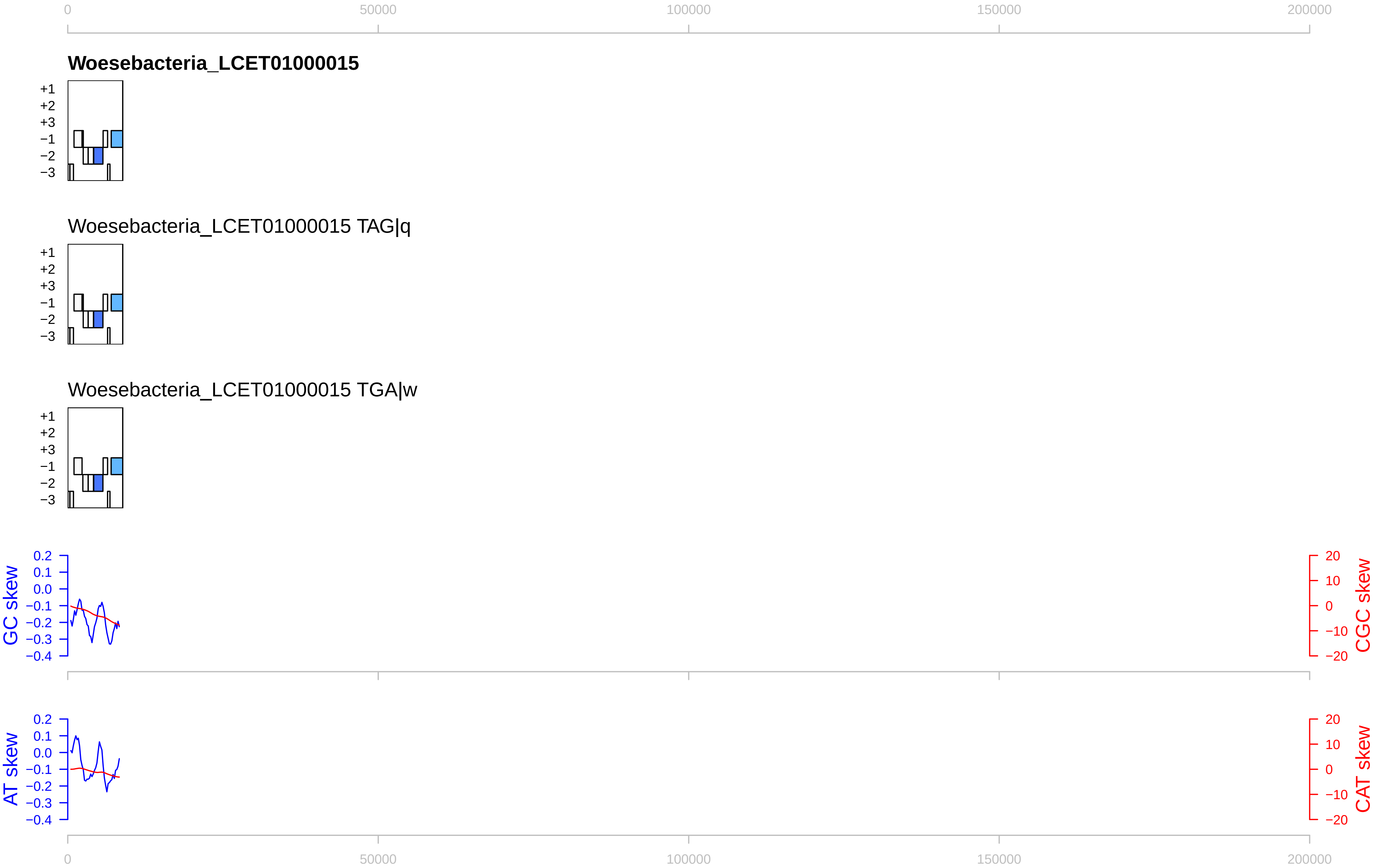

0
50000
100000
150000
200000
Woesebacteria_LCET01000015
+1
+2
+3
−1
−2
−3
Woesebacteria_LCET01000015 TAG|q
Woesebacteria_LCET01000015 TGA|w
GC skew
CGC skew
AT skew
CAT skew
0.2
0.1
0.0
−0.1
−0.2
−0.3
−0.4
20
10
0
−10
−20
