## Supplementary figures and images for "Discovery, diversity and functional associations of crAss-like phages in human gut metagenomes from four Dutch cohorts"

### Material S3

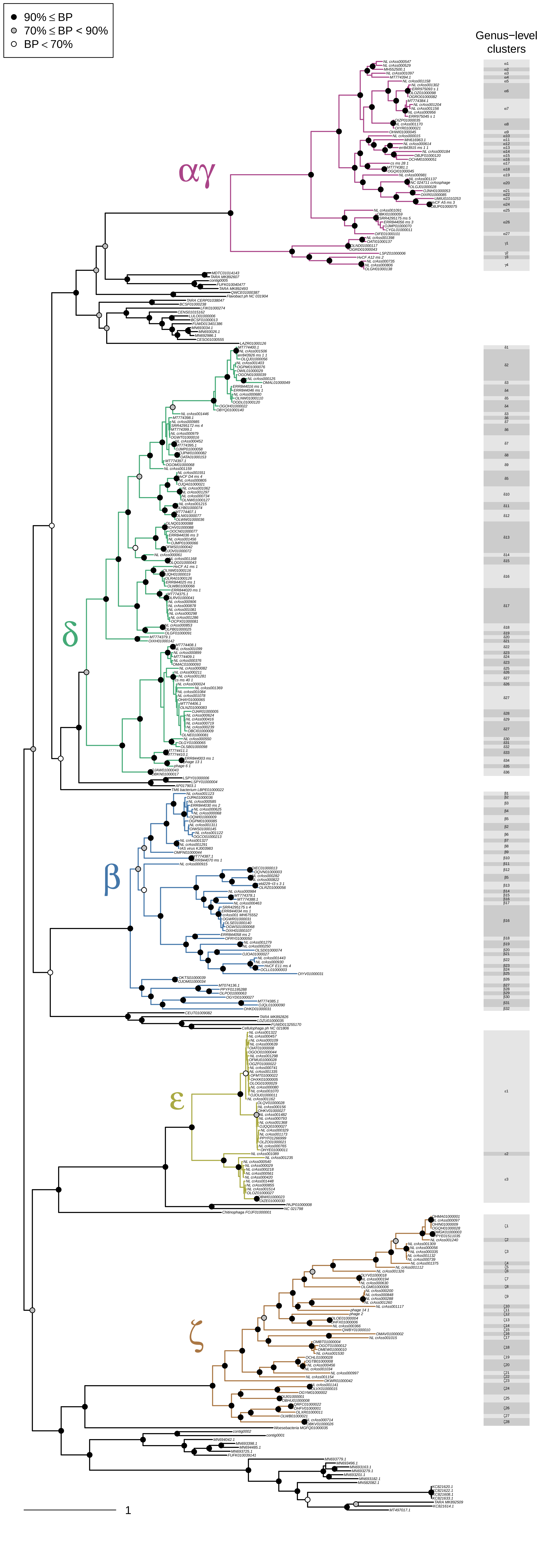

### Material S4

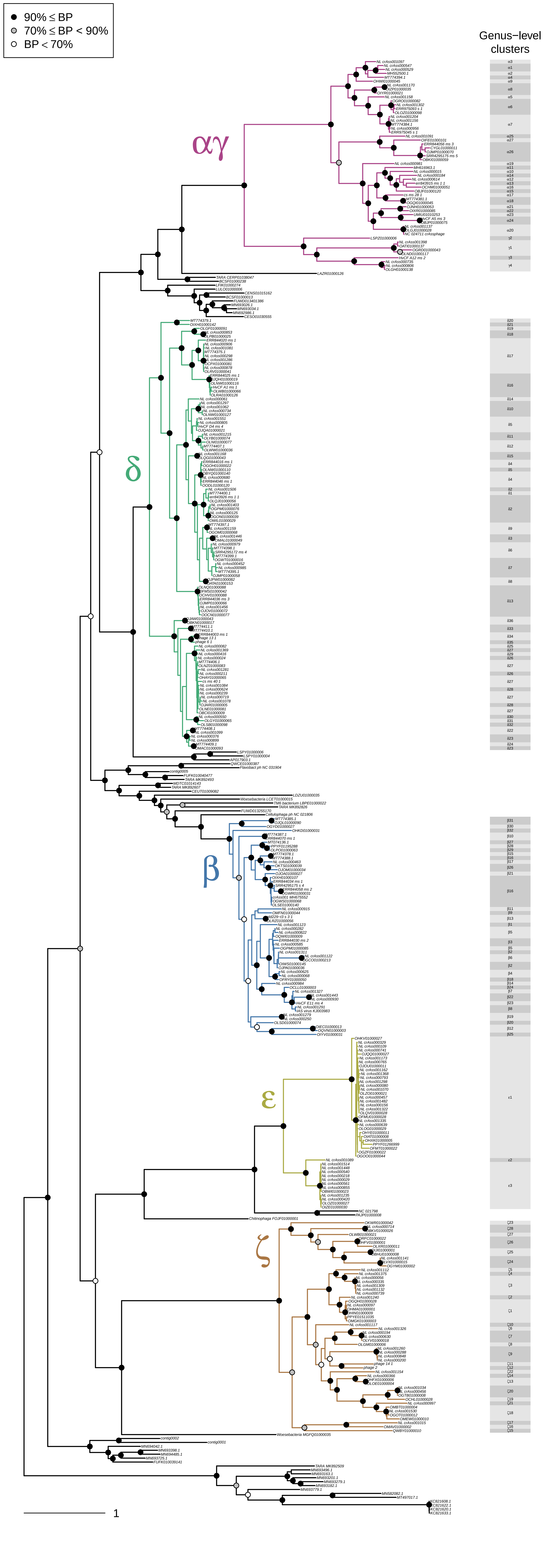

### Material S6

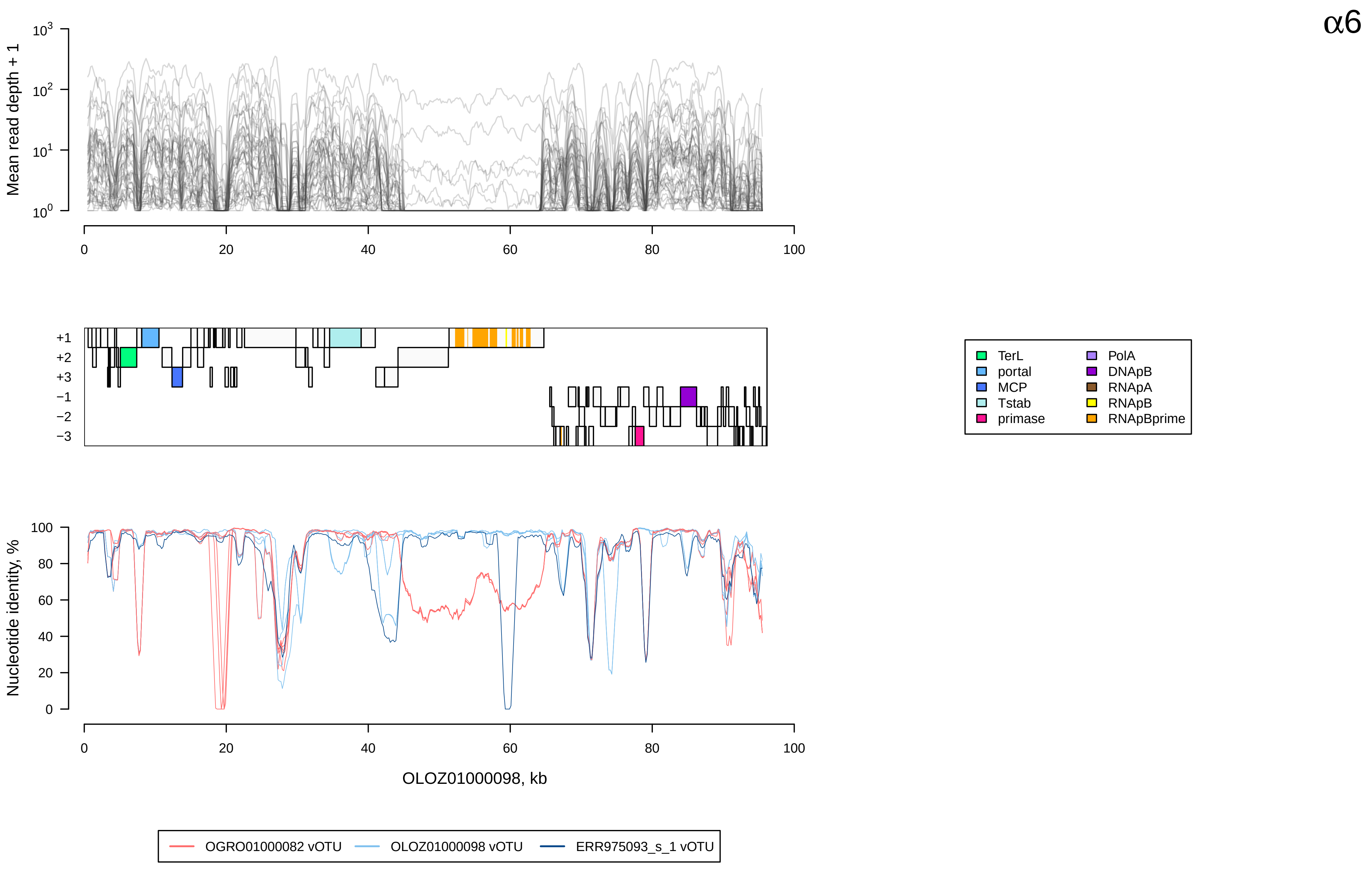

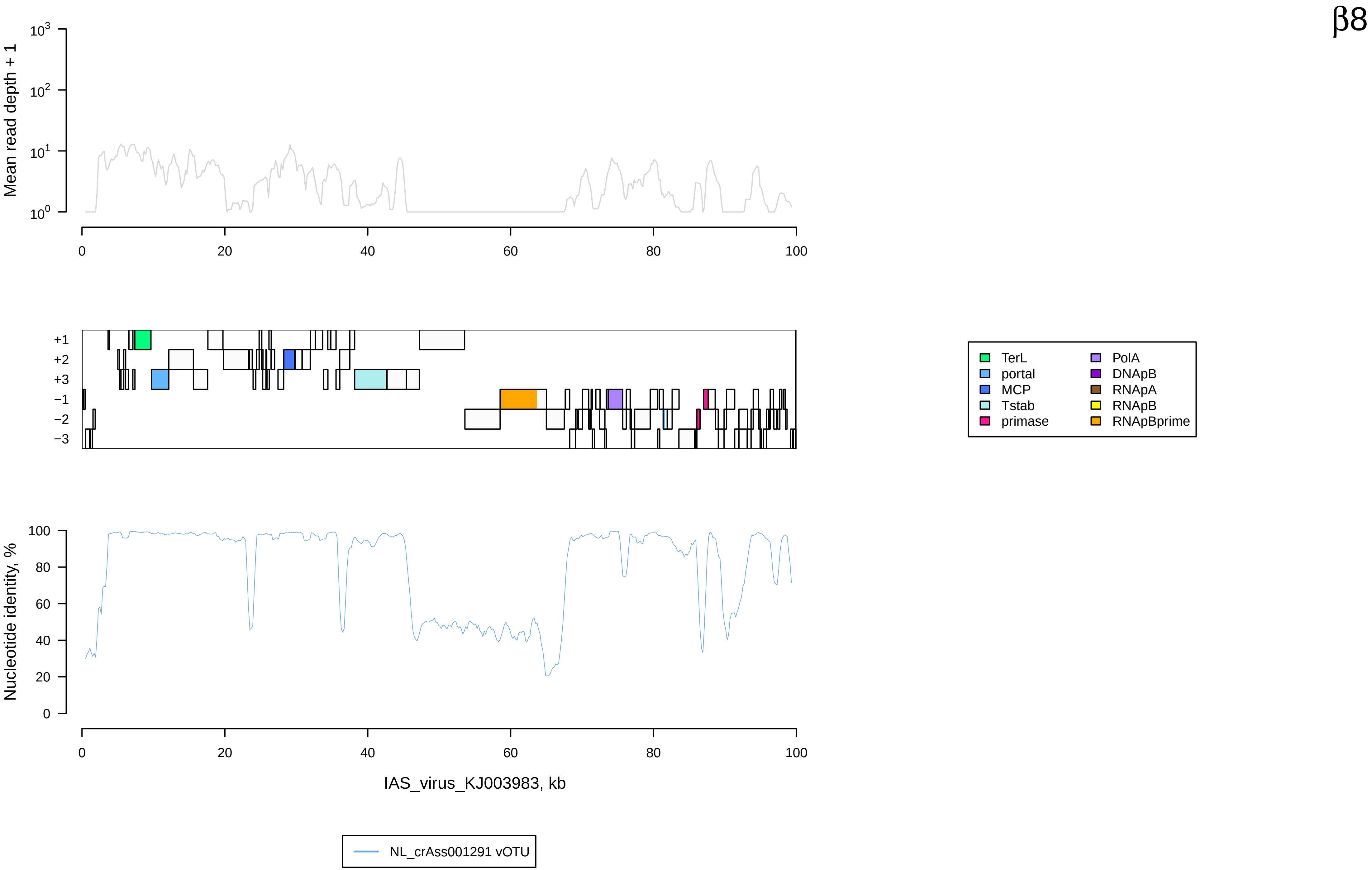

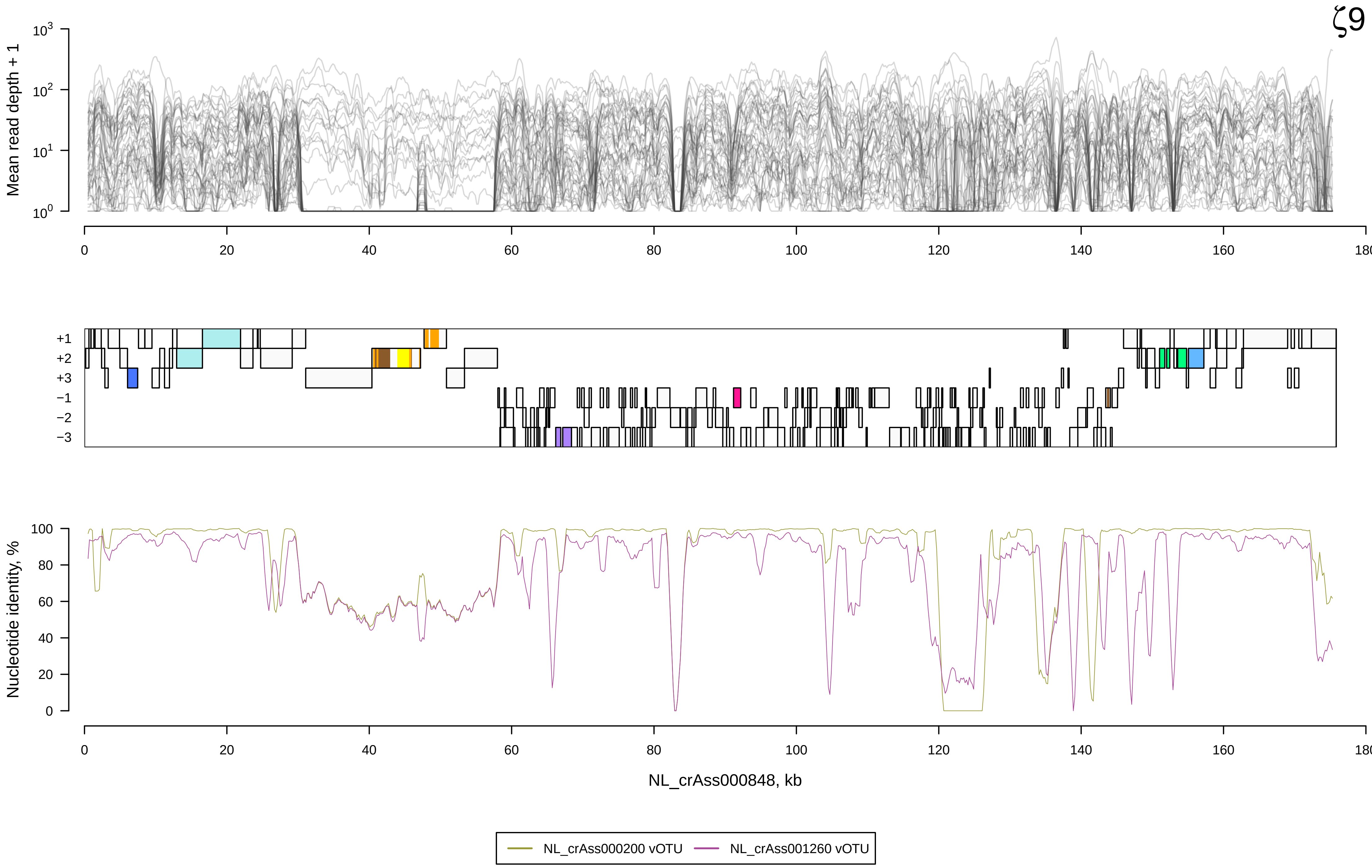
