## Supplementary material for "Discovery, diversity and functional associations of crAss-like phages in human gut metagenomes from four Dutch cohorts": Material S5

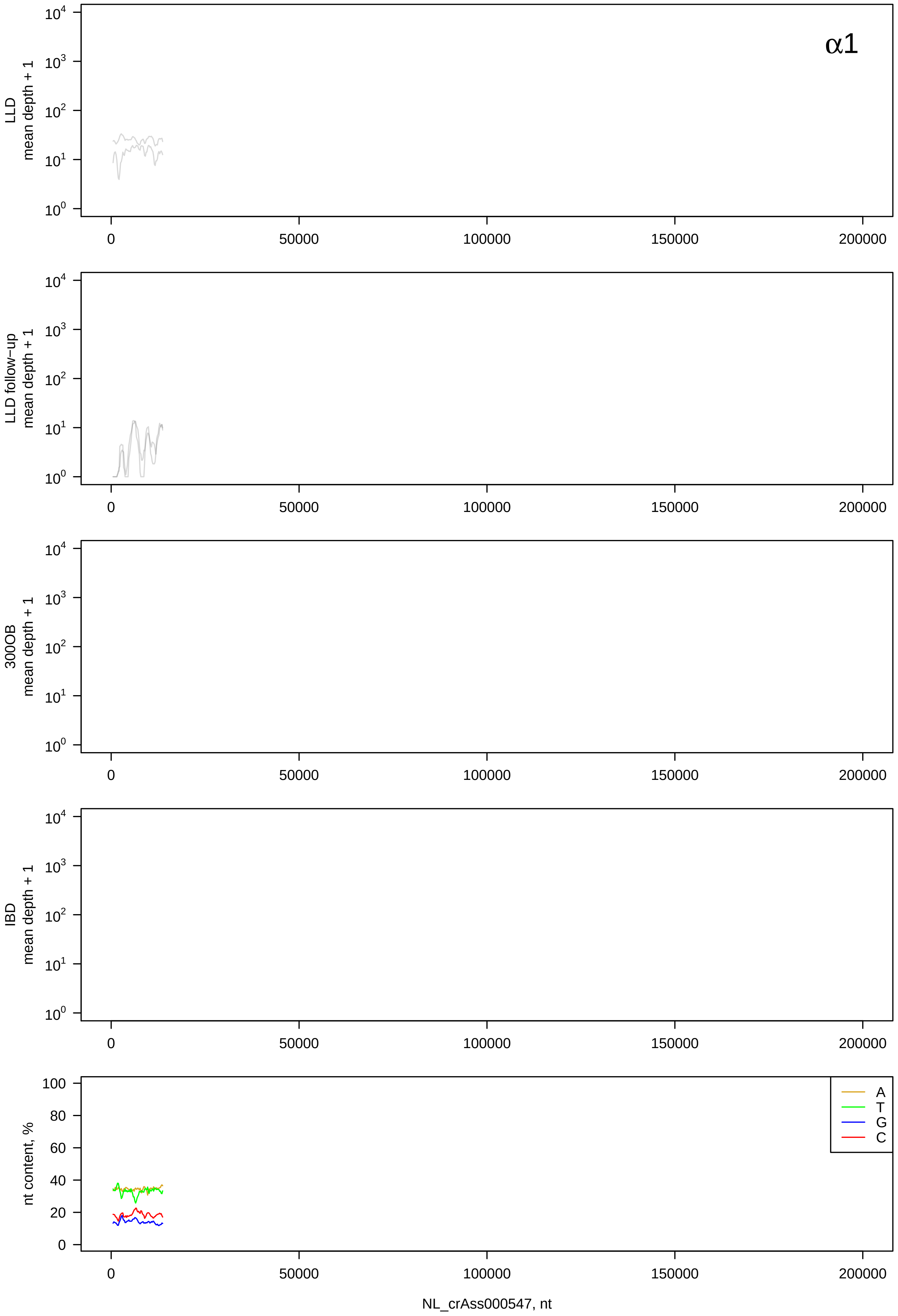
α1
LLD mean depth + 1
LLD follow-up mean depth + 1
300OB mean depth + 1
IBD mean depth + 1
nt content, %
A
T
G
C
0
50000
100000
150000
200000
NL_crAss000547, nt

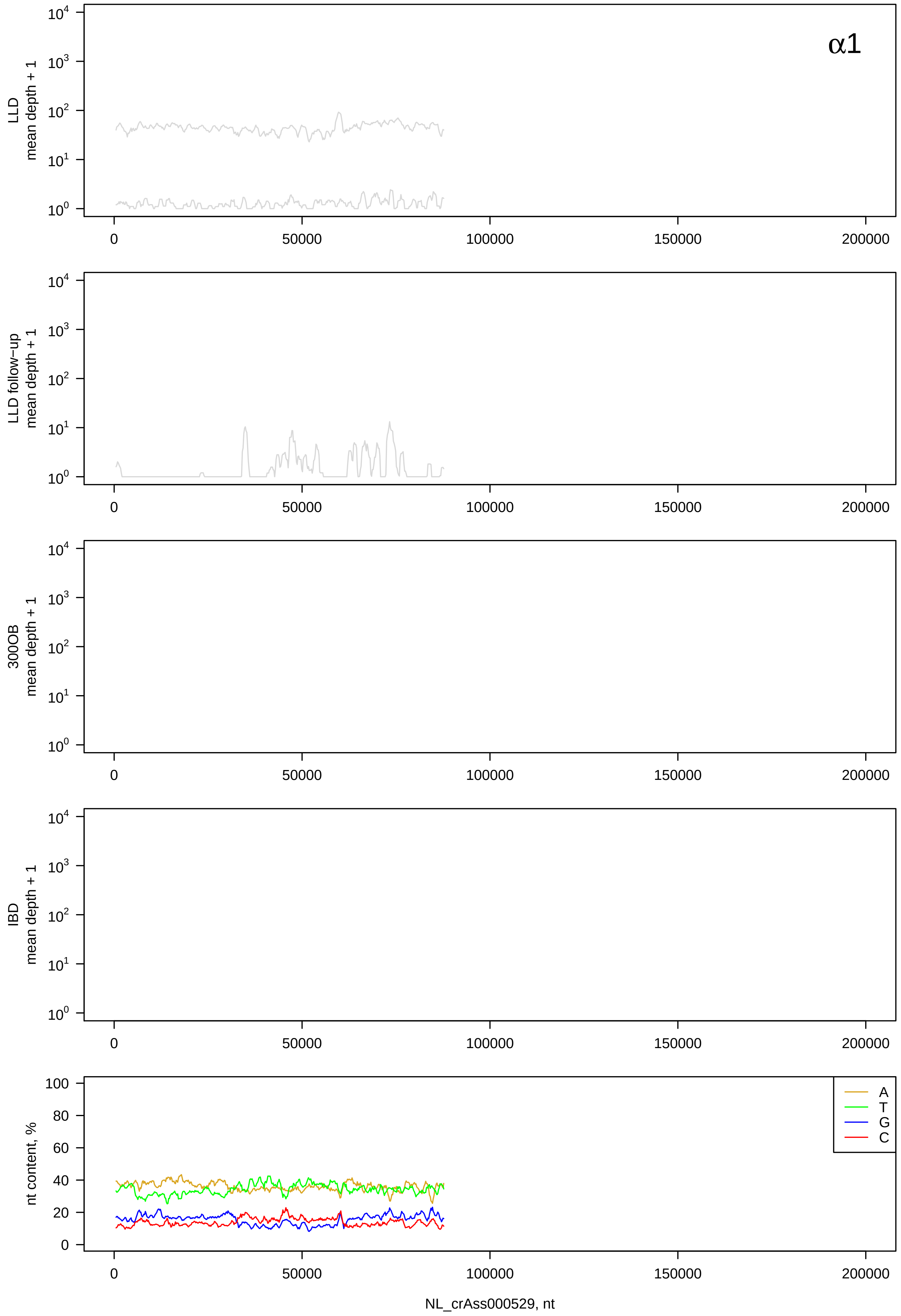
α1
LLD mean depth + 1
LLD follow−up mean depth + 1
300OB mean depth + 1
IBD mean depth + 1
nt content, %
A
T
G
C
NL_crAss000529, nt

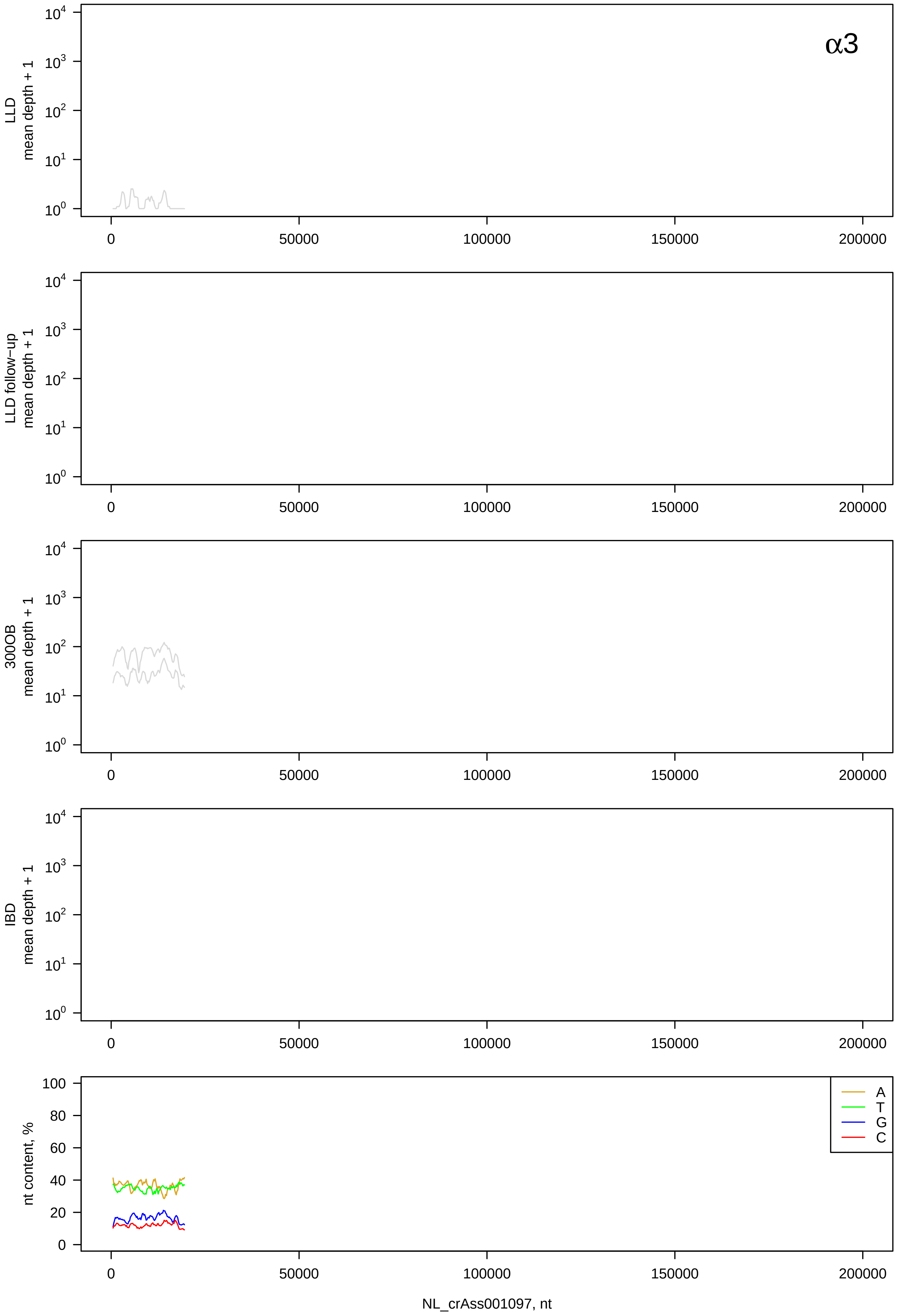
α3
LLD mean depth + 1
LLD follow−up mean depth + 1
300OB mean depth + 1
IBD mean depth + 1
nt content, %
A
T
G
C
0
50000
100000
150000
200000
NL_crAss001097, nt

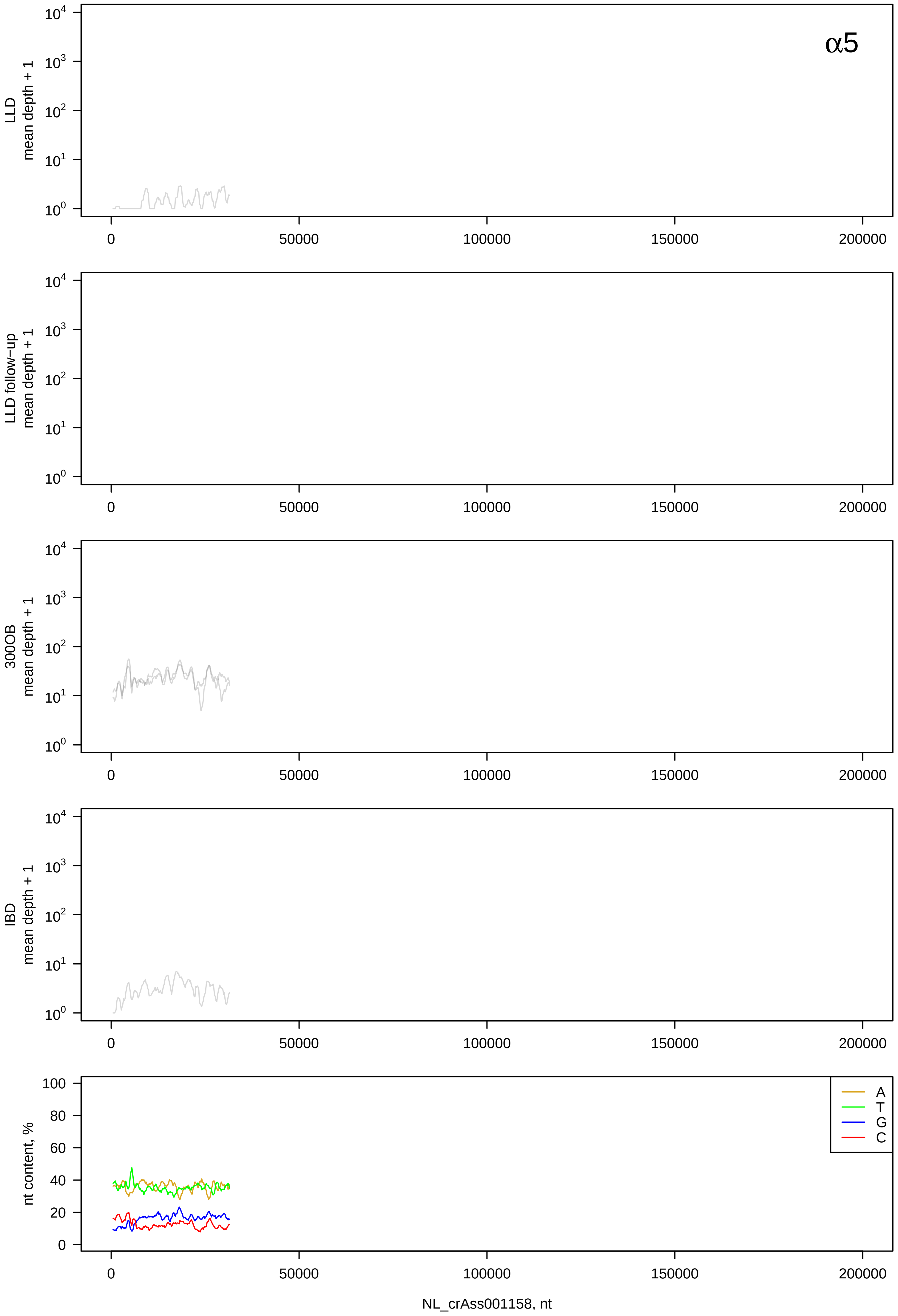

α5
LLD mean depth + 1
LLD follow−up mean depth + 1
300OB mean depth + 1
IBD mean depth + 1
nt content, %
A
T
G
C
0
50000
100000
150000
200000
NL_crAss001158, nt

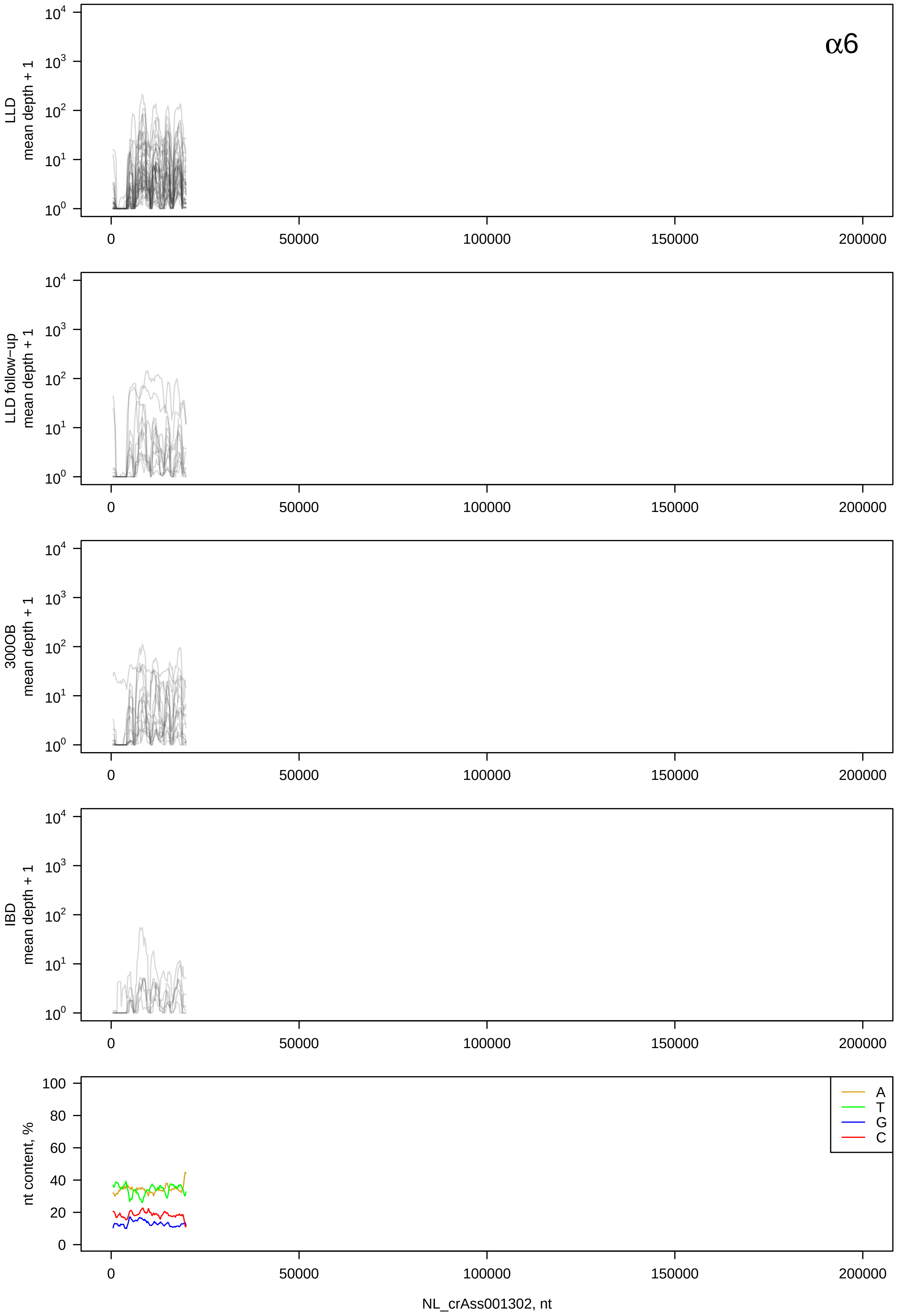

α6
LLD
mean depth + 1
LLD follow−up
mean depth + 1
300OB
mean depth + 1
IBD
mean depth + 1
$10^0$
$10^1$
$10^2$
$10^3$
$10^4$
0
50000
100000
150000
200000
nt content, %
0
20
40
60
80
100
A
T
G
C
NL_crAss001302, nt

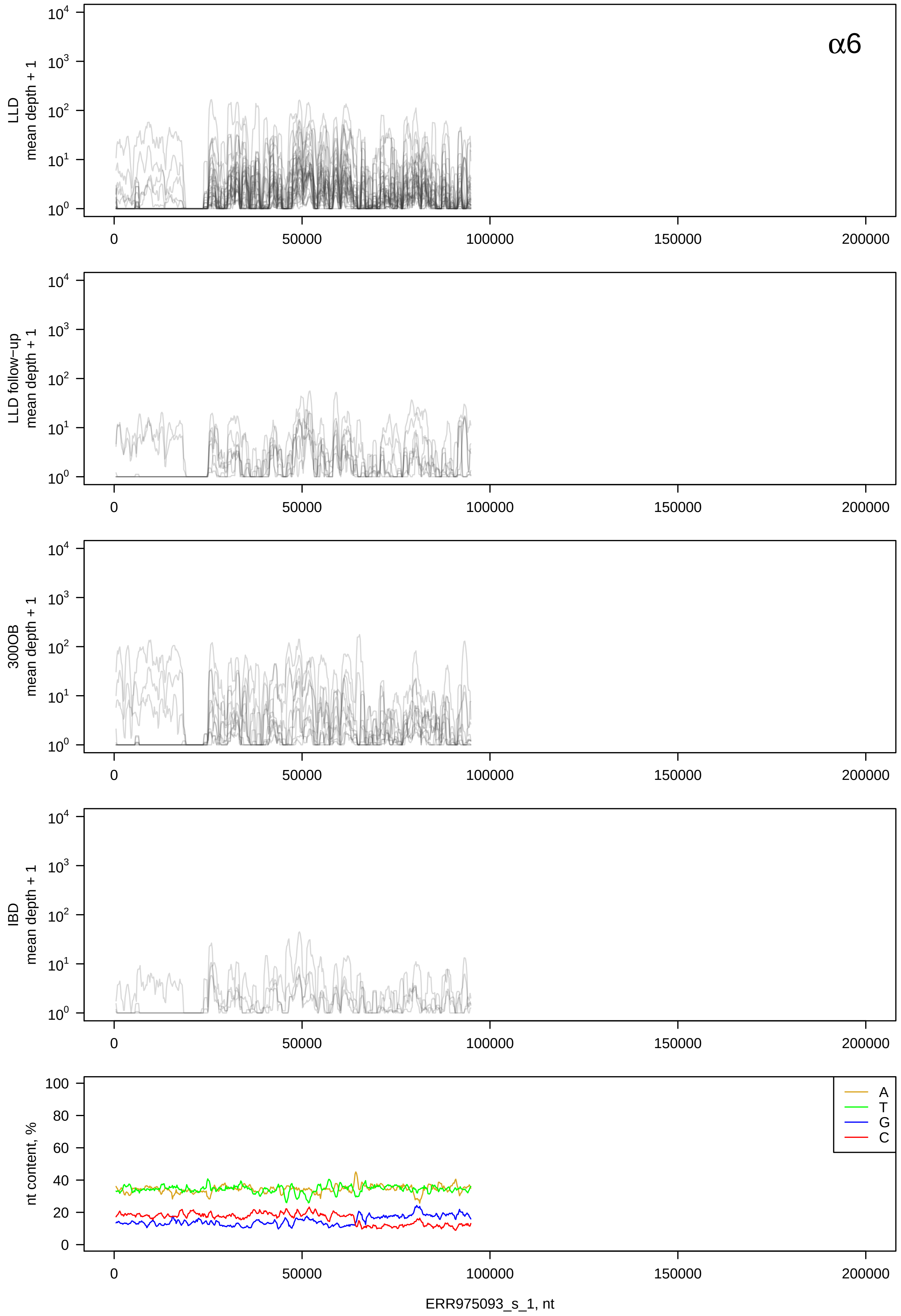

α6
LLD
mean depth + 1
LLD follow−up
mean depth + 1
300OB
mean depth + 1
IBD
mean depth + 1
$10^4$
$10^3$
$10^2$
$10^1$
$10^0$
0
50000
100000
150000
200000
nt content, %
100
80
60
40
20
0
A
T
G
C
ERR975093_s_1, nt

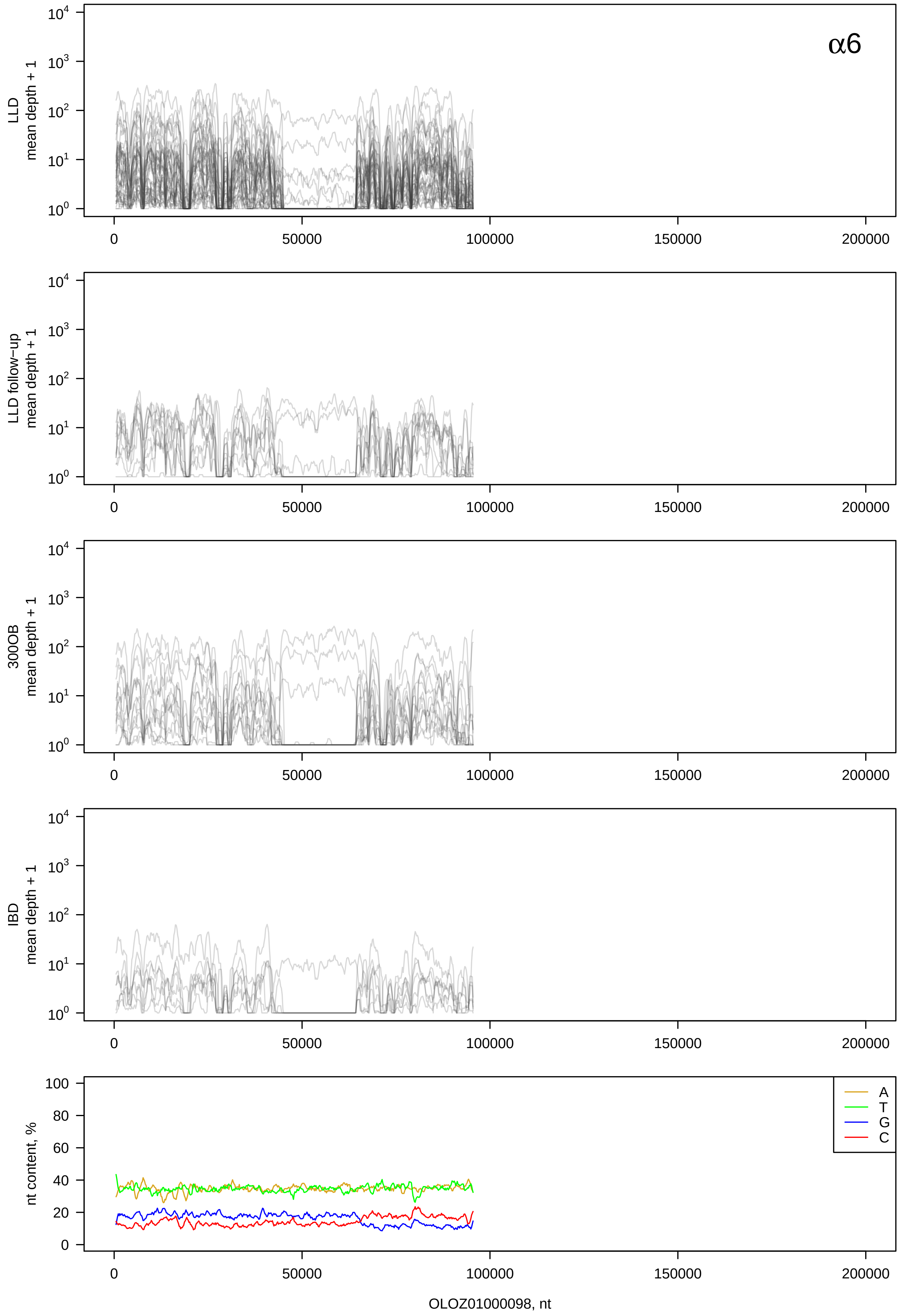

α6
LLD
mean depth + 1
LLD follow−up
mean depth + 1
300OB
mean depth + 1
IBD
mean depth + 1
nt content, %
A
T
G
C
0
50000
100000
150000
200000
OLOZ01000098, nt

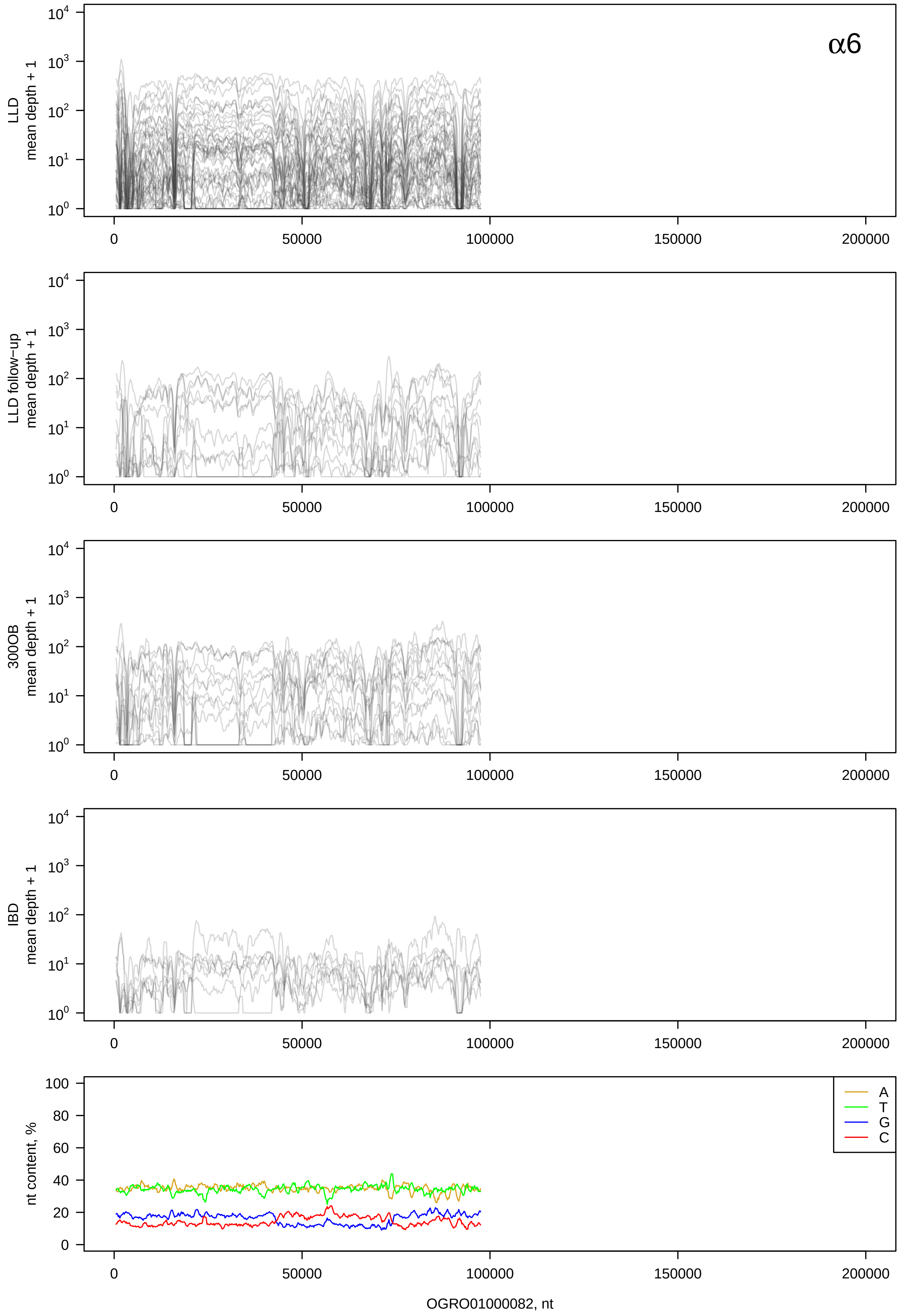

α6
LLD mean depth + 1
LLD follow-up mean depth + 1
300OB mean depth + 1
IBD mean depth + 1
nt content, %
A
T
G
C
0
50000
100000
150000
200000
OGRO01000082, nt

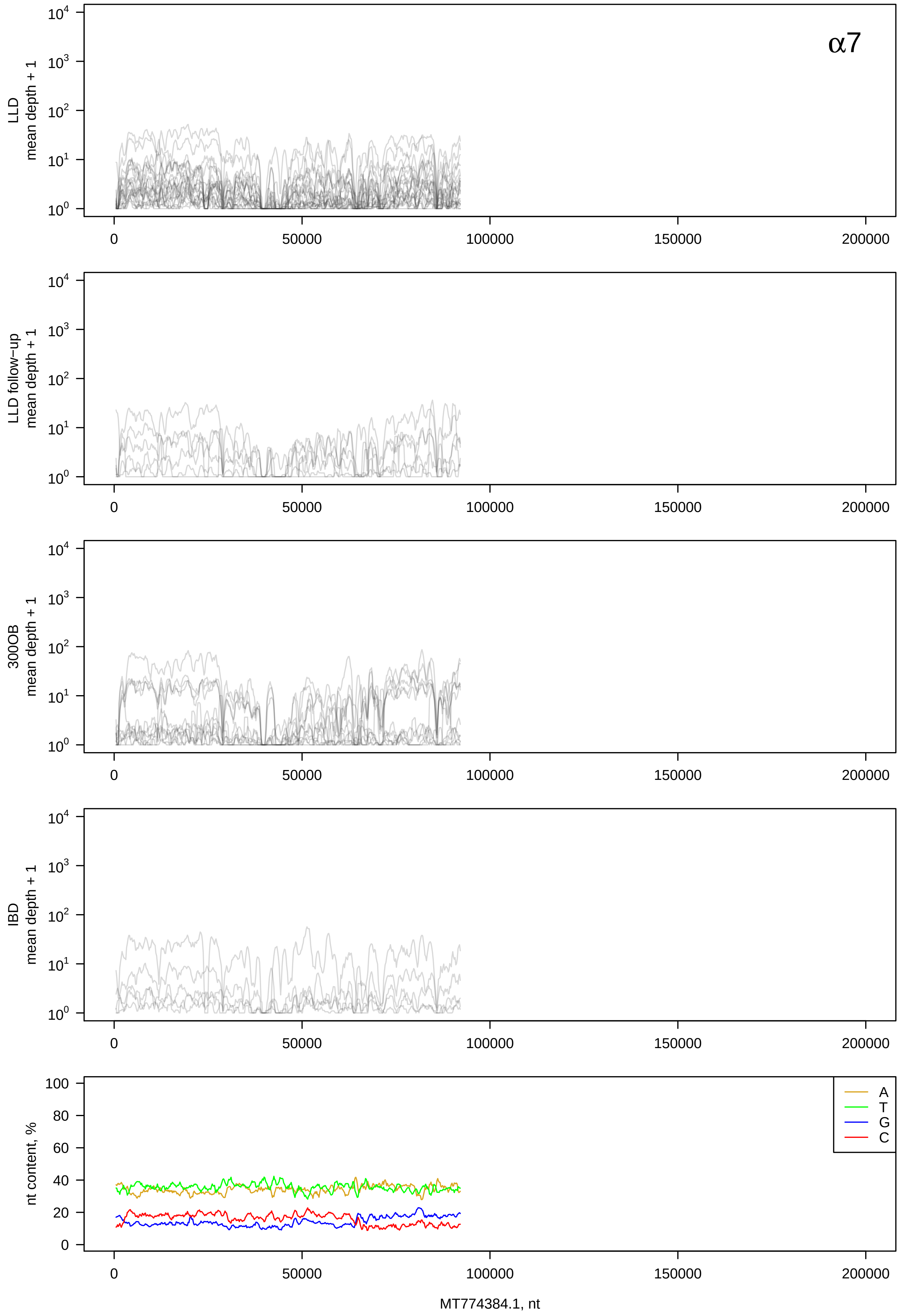

α7
LLD mean depth + 1
LLD follow-up mean depth + 1
300OB mean depth + 1
IBD mean depth + 1
nt content, %
A
T
G
C
MT774384.1, nt

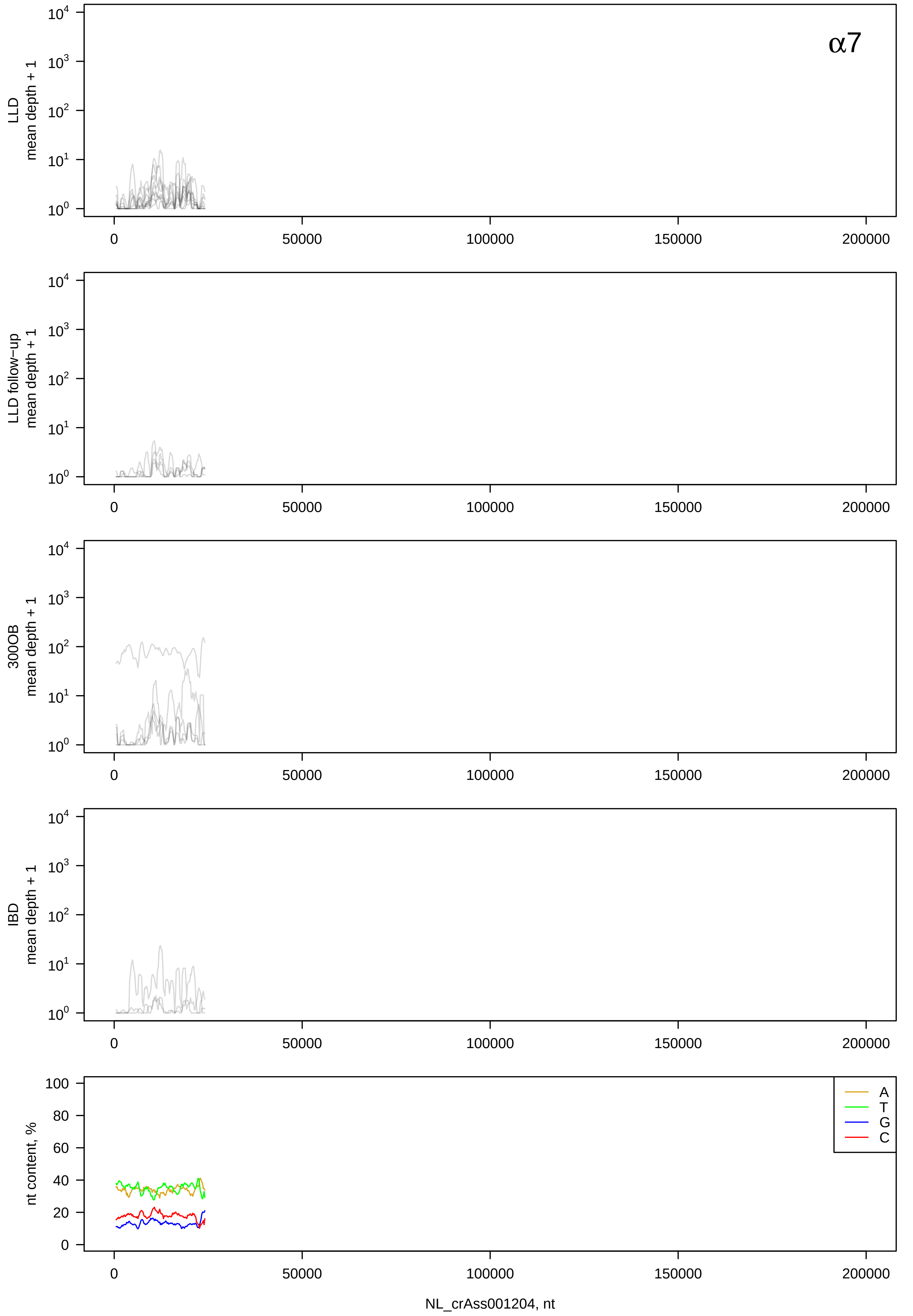
α7
LLD
mean depth + 1
LLD follow-up
mean depth + 1
300OB
mean depth + 1
IBD
mean depth + 1
nt content, %
A
T
G
C
NL_crAss001204, nt

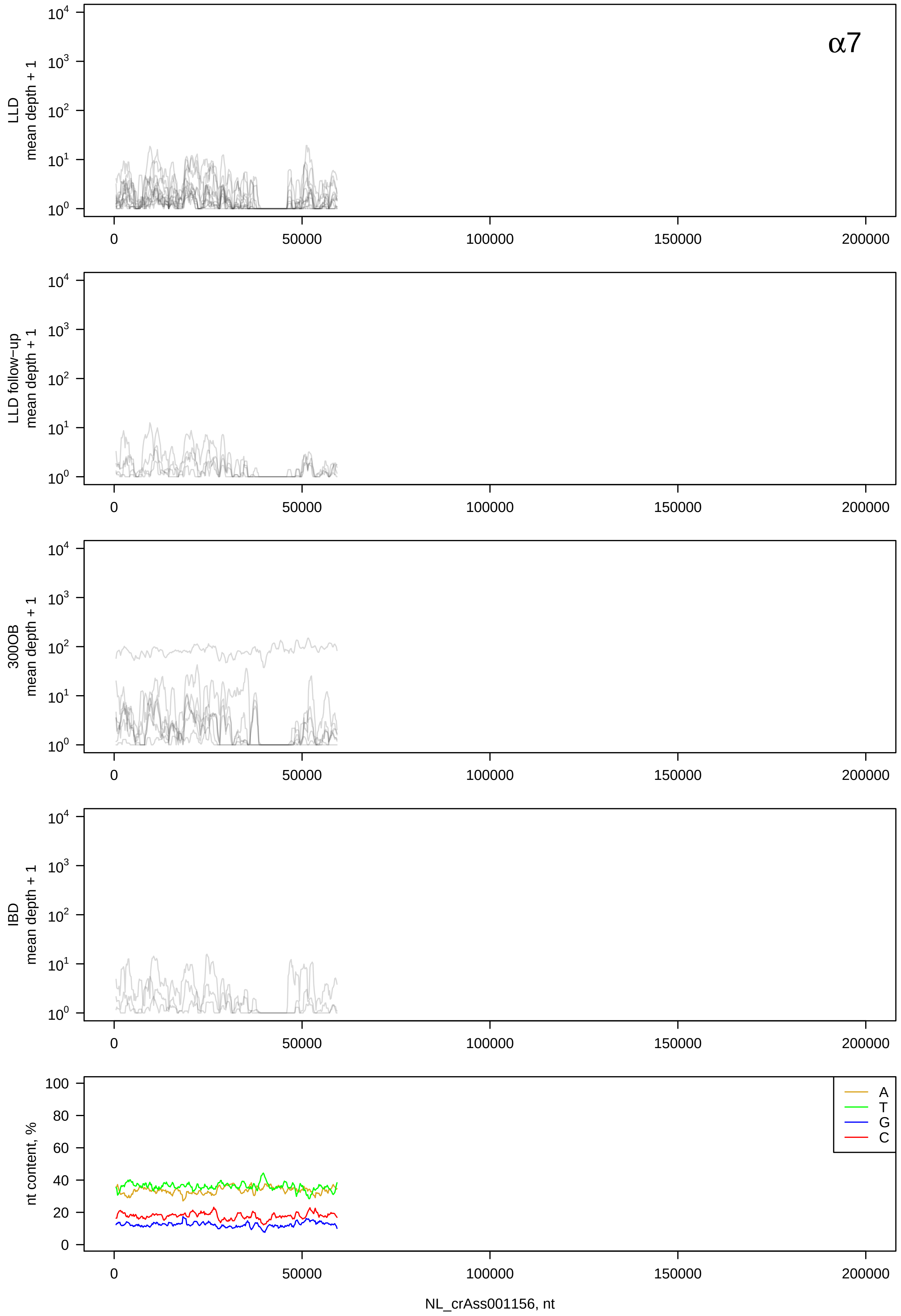

α7
LLD mean depth + 1
LLD follow−up mean depth + 1
300OB mean depth + 1
IBD mean depth + 1
nt content, %
A
T
G
C
NL_crAss001156, nt

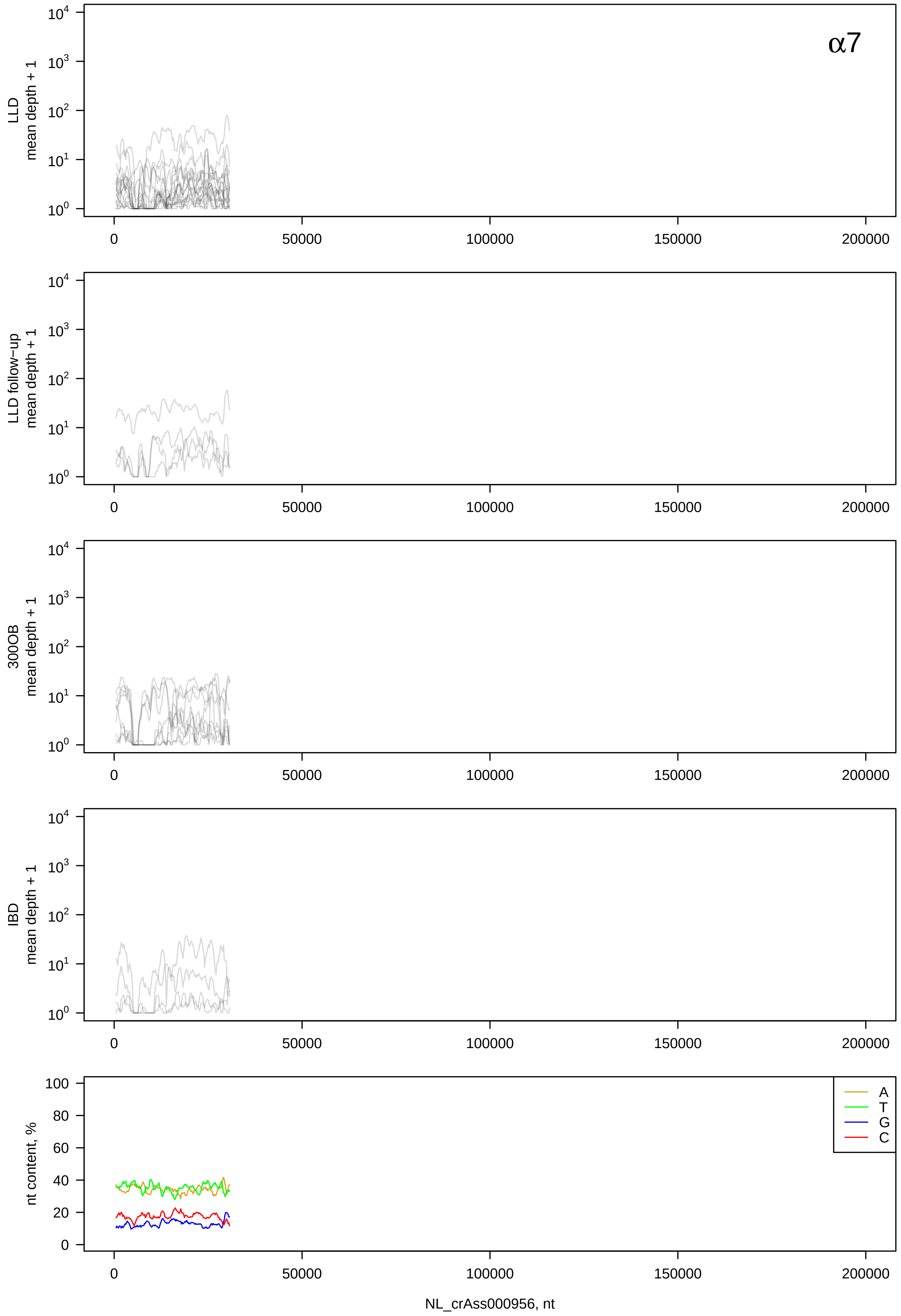

α7
LLD mean depth + 1
LLD follow−up mean depth + 1
300OB mean depth + 1
IBD mean depth + 1
nt content, %
A
T
G
C
0
50000
100000
150000
200000
NL_crAss000956, nt

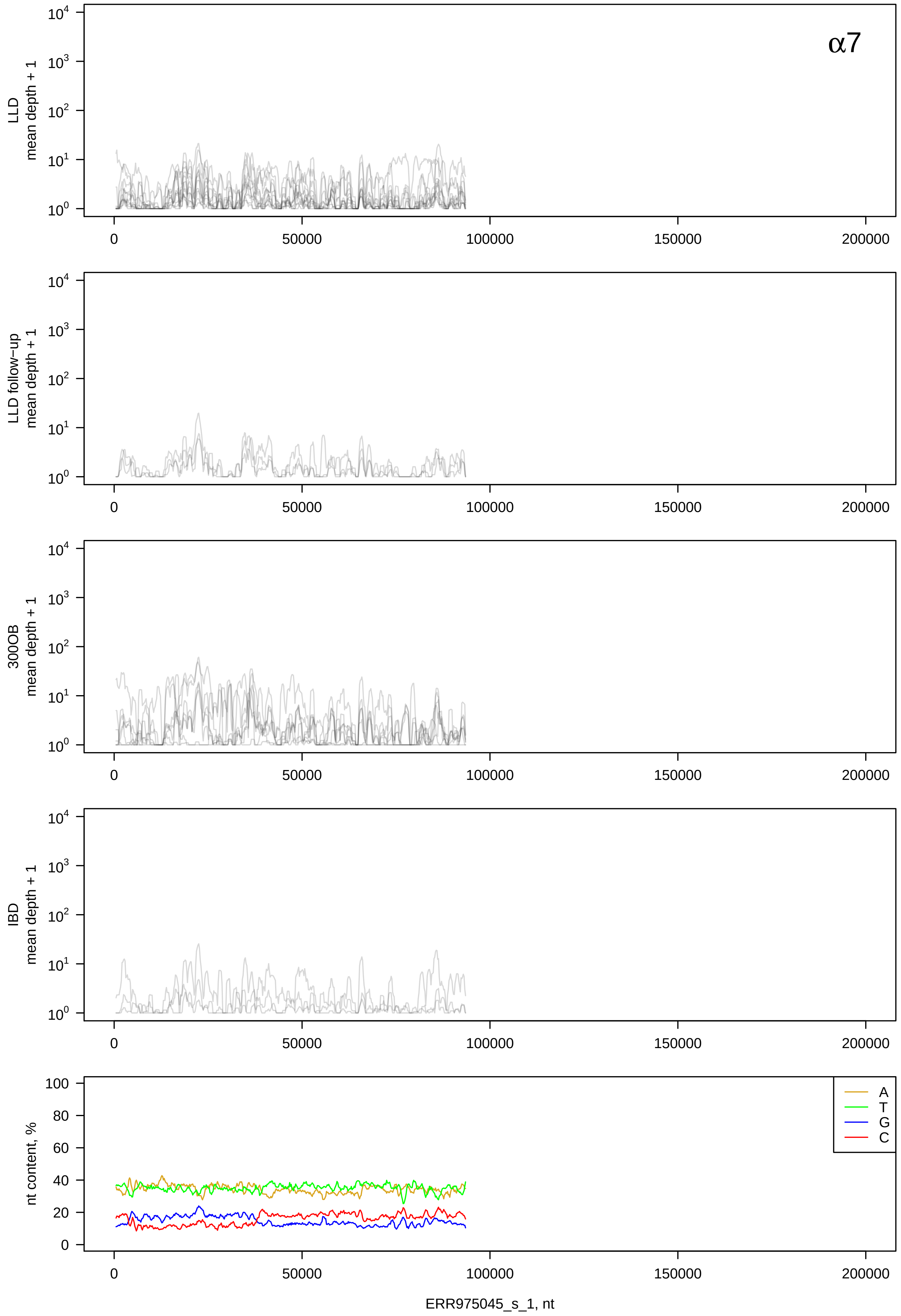

α7
LLD mean depth + 1
LLD follow-up mean depth + 1
300OB mean depth + 1
IBD mean depth + 1
nt content, %
A
T
G
C
0
50000
100000
150000
200000
ERR975045_s_1, nt

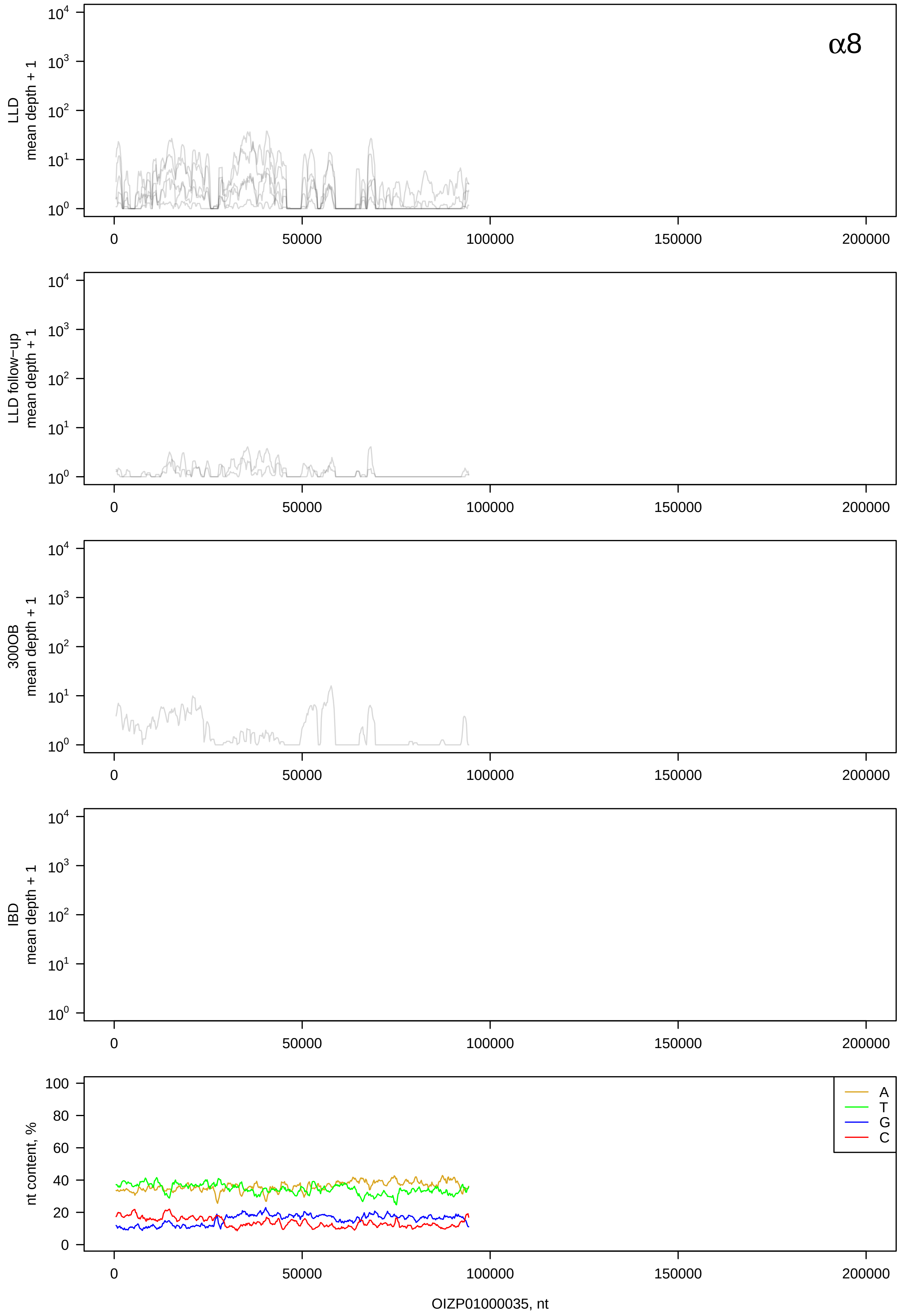
α8
LLD
mean depth + 1
LLD follow−up
mean depth + 1
300OB
mean depth + 1
IBD
mean depth + 1
nt content, %
A
T
G
C
OIZP01000035, nt

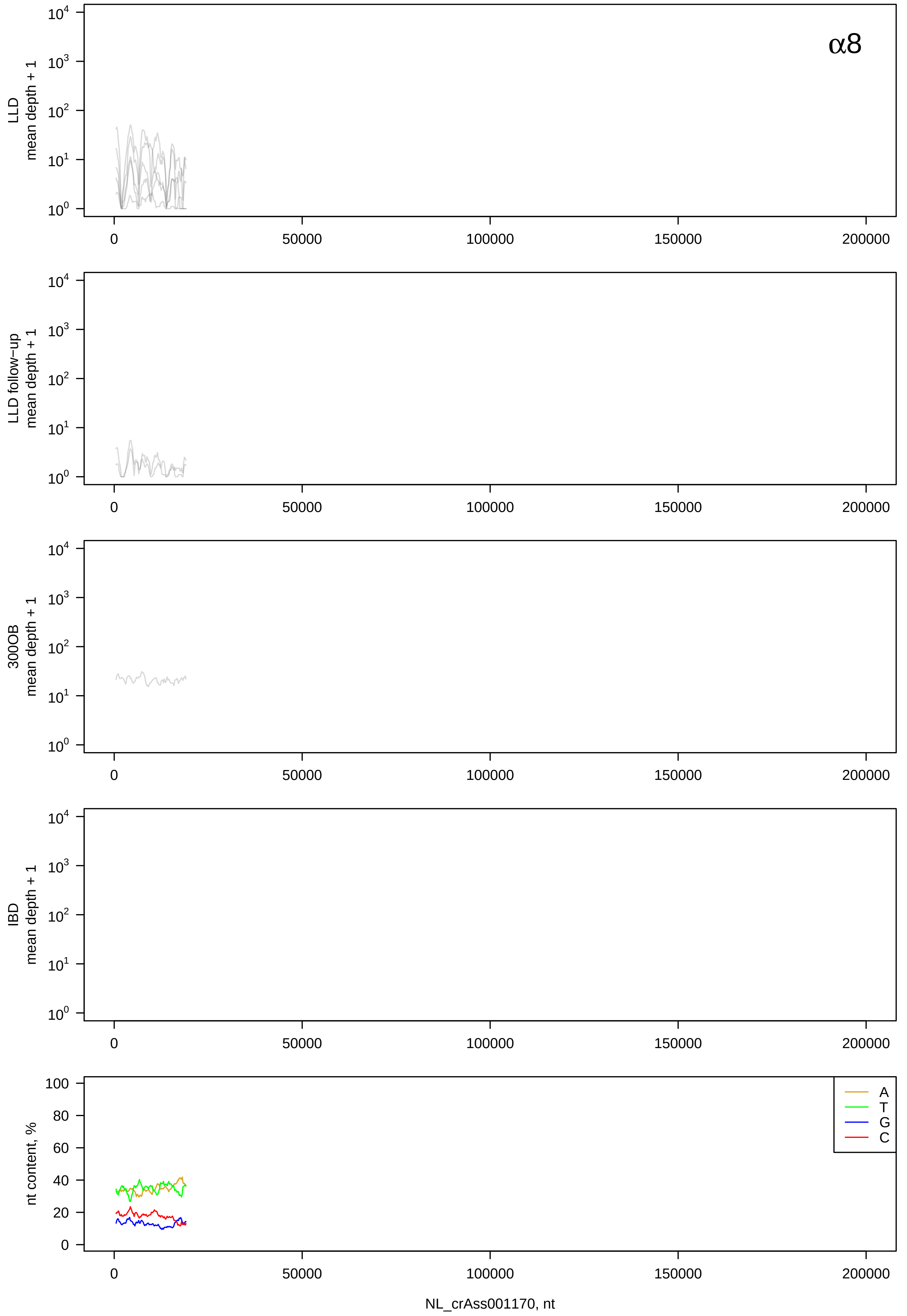
α8
LLD mean depth + 1
LLD follow-up mean depth + 1
300OB mean depth + 1
IBD mean depth + 1
nt content, %
A
T
G
C
0
50000
100000
150000
200000
NL_crAss001170, nt

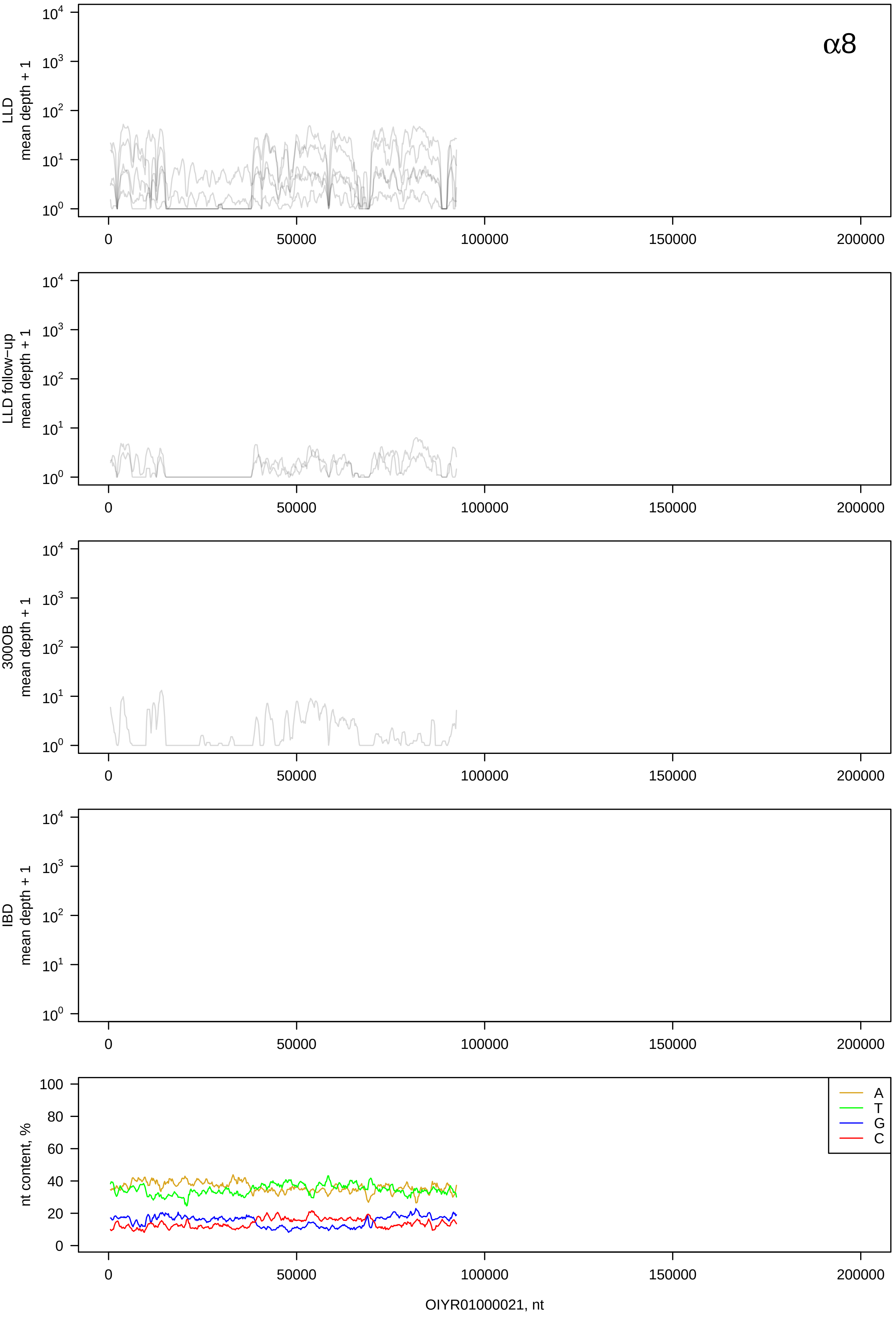
α8
LLD
mean depth + 1
LLD follow−up
mean depth + 1
300OB
mean depth + 1
IBD
mean depth + 1
nt content, %
A
T
G
C
OIYR01000021, nt

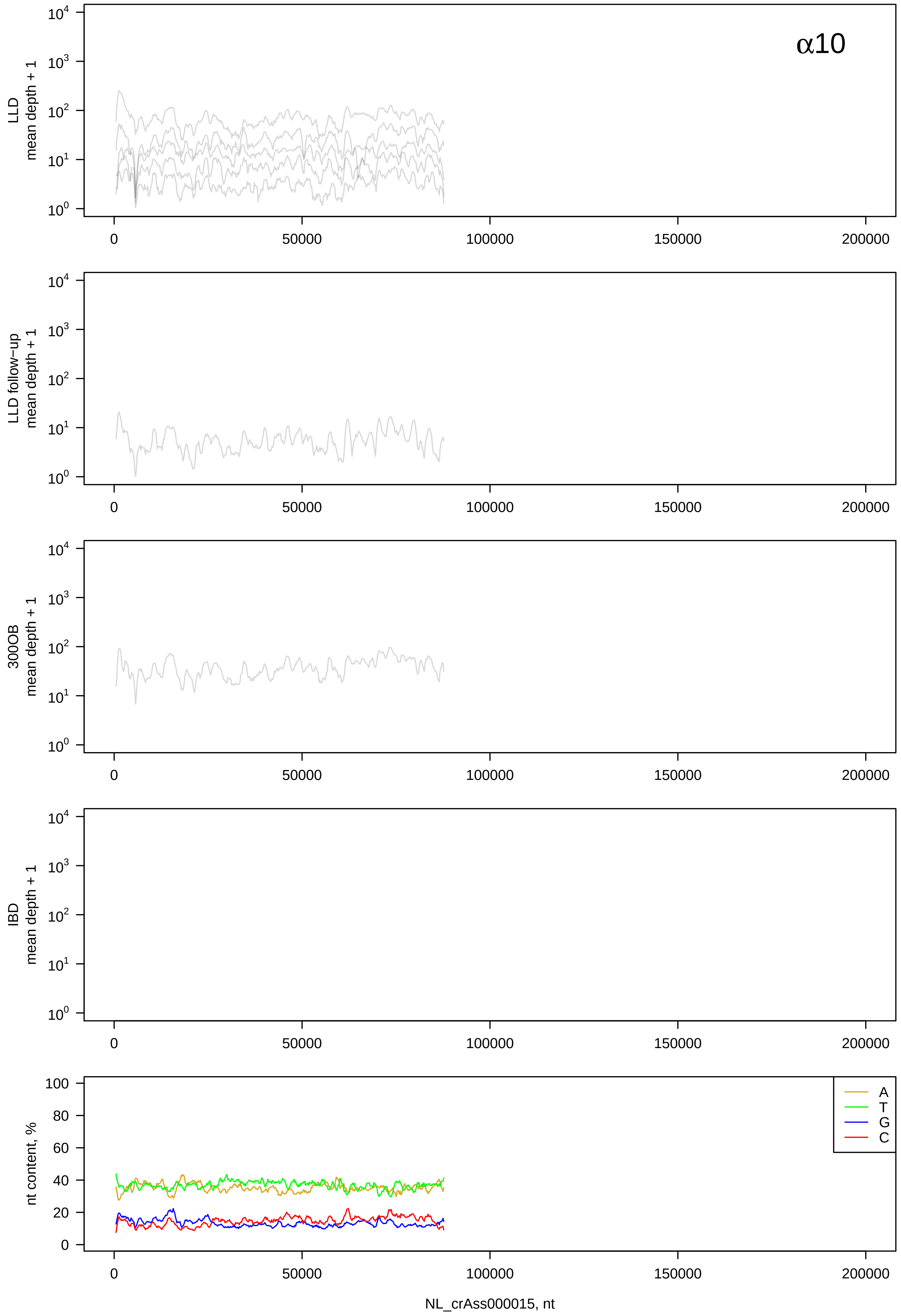
α10
LLD
mean depth + 1
LLD follow-up
mean depth + 1
300OB
mean depth + 1
IBD
mean depth + 1
nt content, %
A
T
G
C
NL_crAss000015, nt

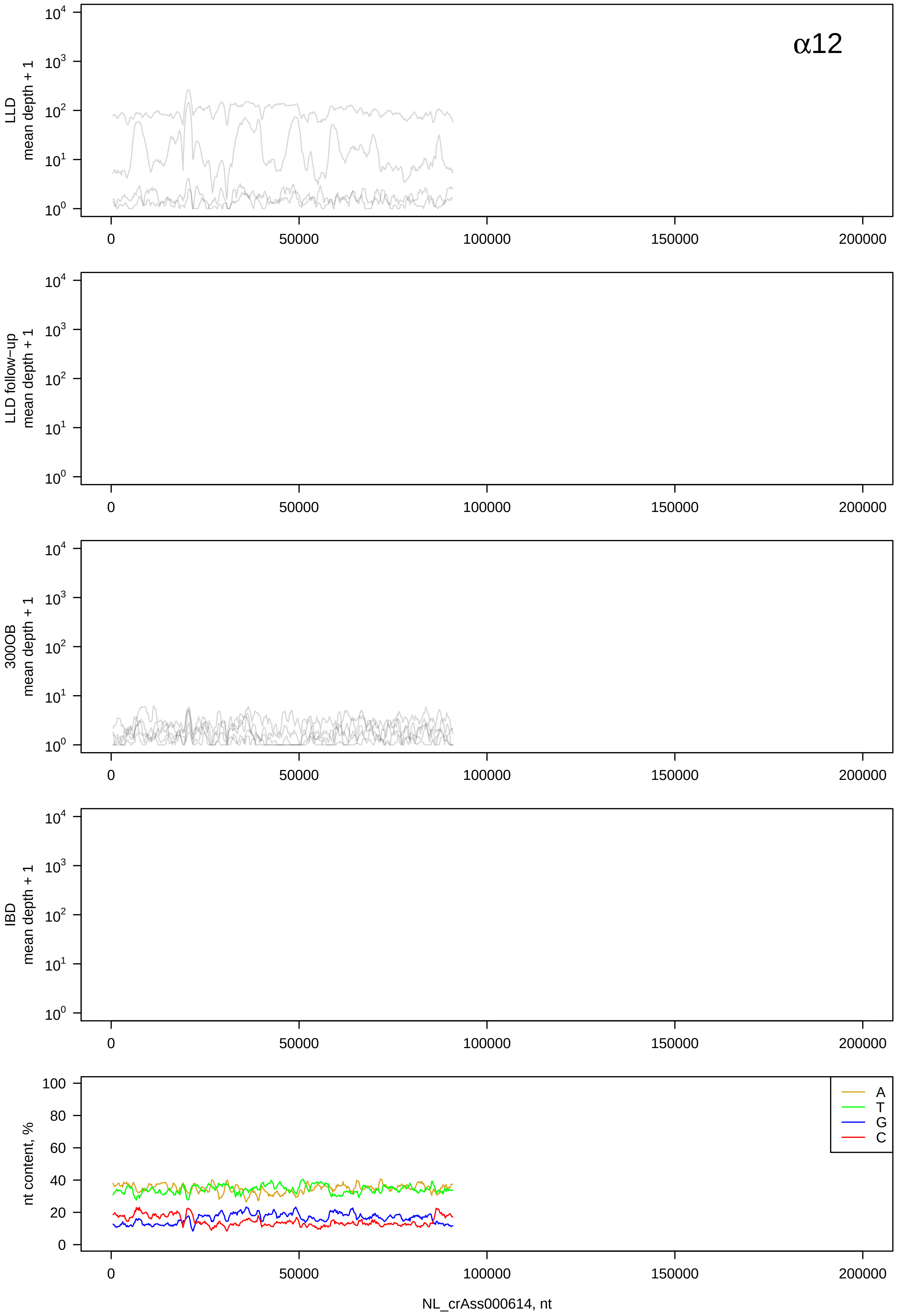
α12
LLD
mean depth + 1
LLD follow−up
mean depth + 1
300OB
mean depth + 1
IBD
mean depth + 1
nt content, %
A
T
G
C
NL_crAss000614, nt

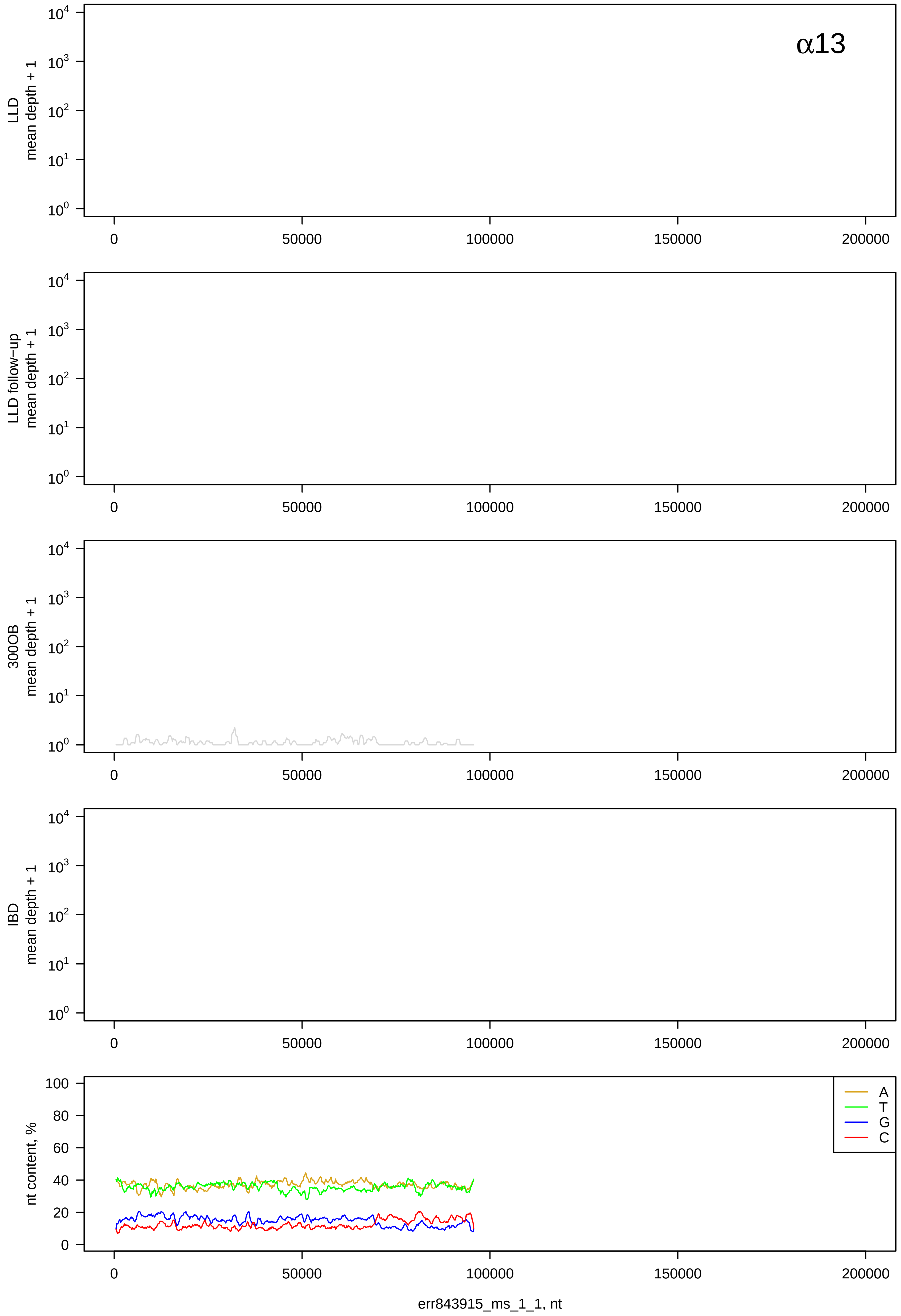
α13
LLD mean depth + 1
LLD follow−up mean depth + 1
300OB mean depth + 1
IBD mean depth + 1
nt content, %
A
T
G
C
err843915_ms_1_1, nt

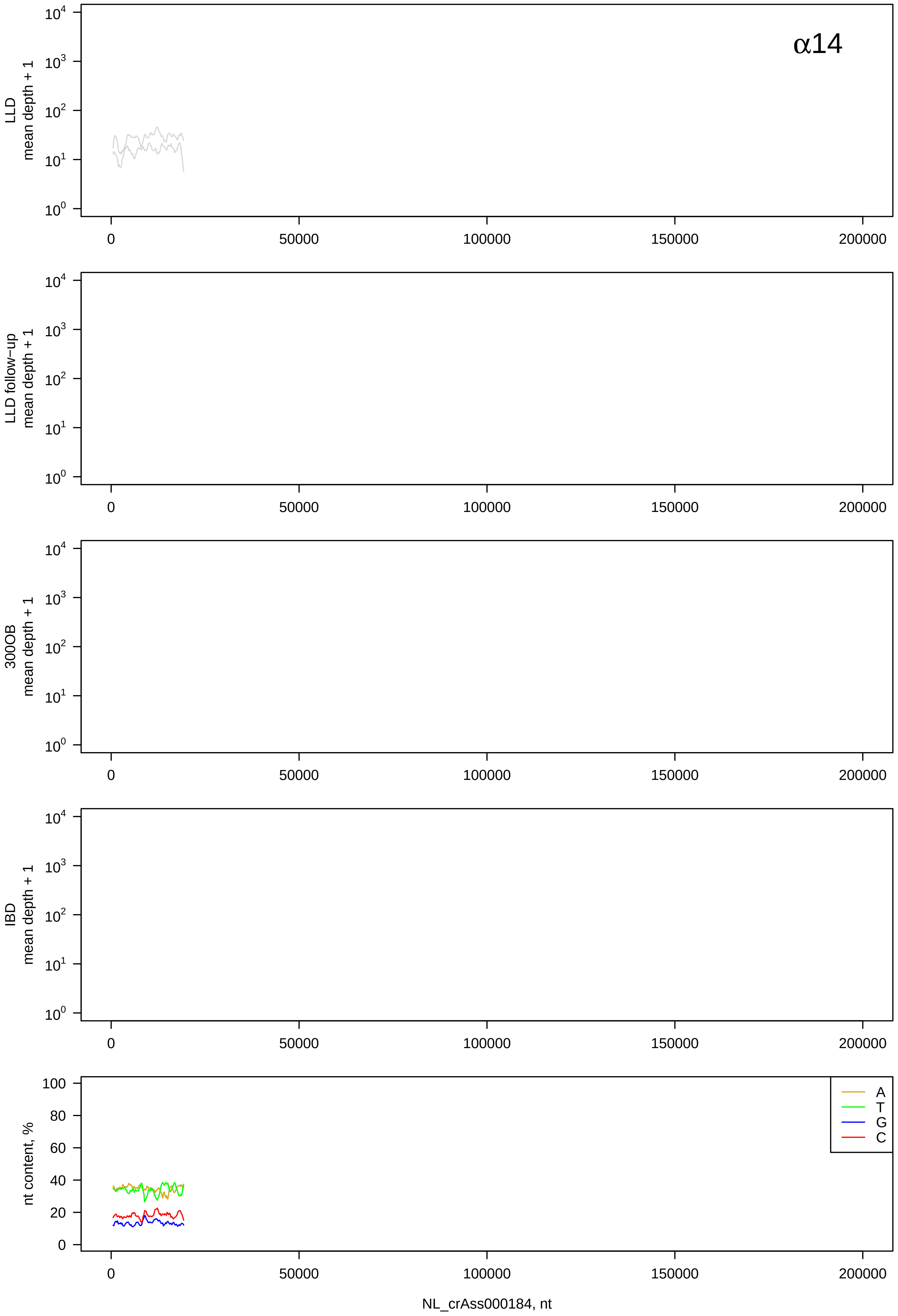
α14
LLD
mean depth + 1
LLD follow−up
mean depth + 1
300OB
mean depth + 1
IBD
mean depth + 1
nt content, %
A
T
G
C
0
50000
100000
150000
200000
NL_crAss000184, nt

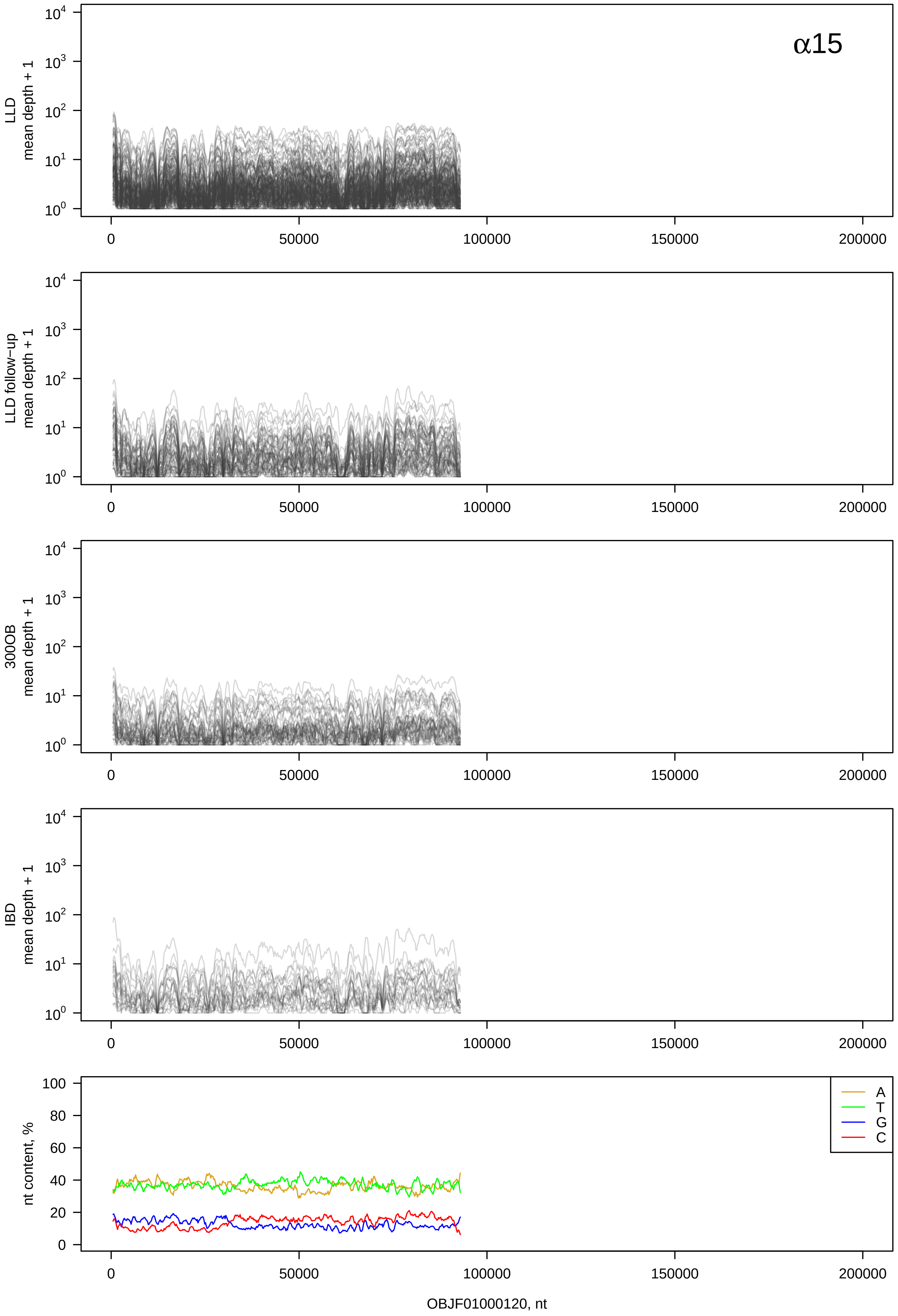

α15
LLD mean depth + 1
LLD follow−up mean depth + 1
300OB mean depth + 1
IBD mean depth + 1
nt content, %
A
T
G
C
0
50000
100000
150000
200000
OBJF01000120, nt

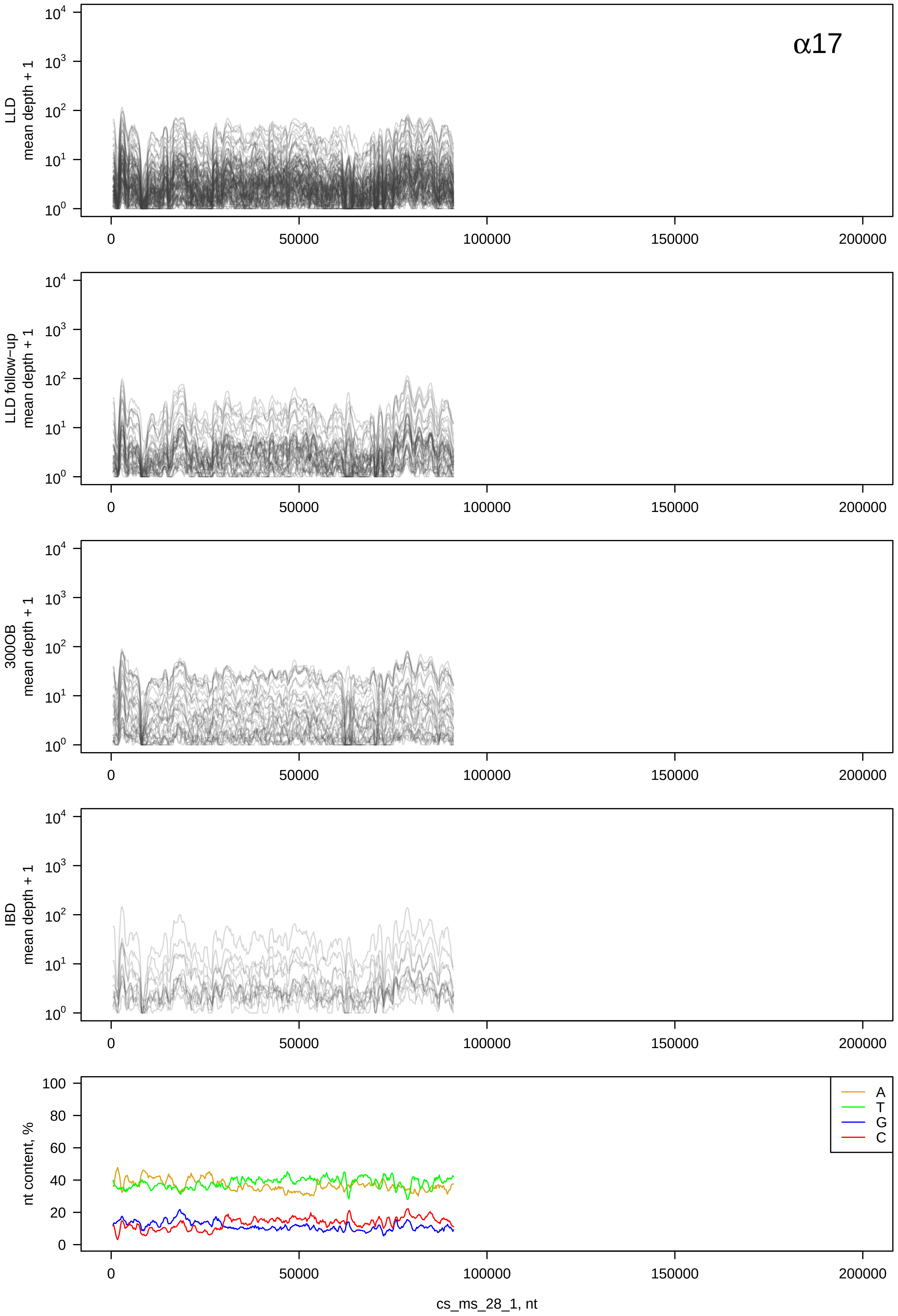

α17
LLD mean depth + 1
LLD follow−up mean depth + 1
300OB mean depth + 1
IBD mean depth + 1
nt content, %
A
T
G
C
cs_ms_28_1, nt

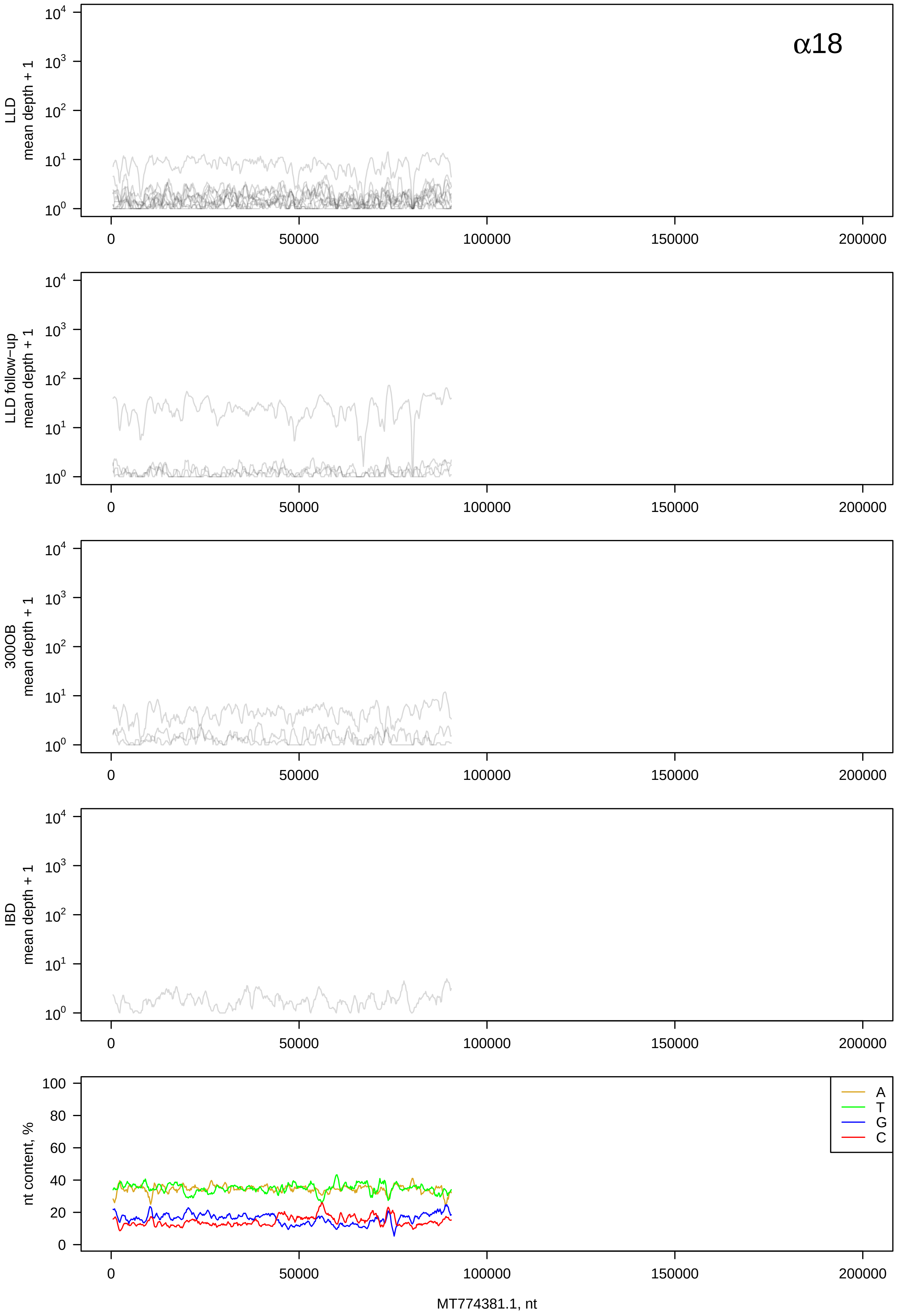

α18
LLD
mean depth + 1
LLD follow−up
mean depth + 1
300OB
mean depth + 1
IBD
mean depth + 1
nt content, %
A
T
G
C
0
50000
100000
150000
200000
MT774381.1, nt

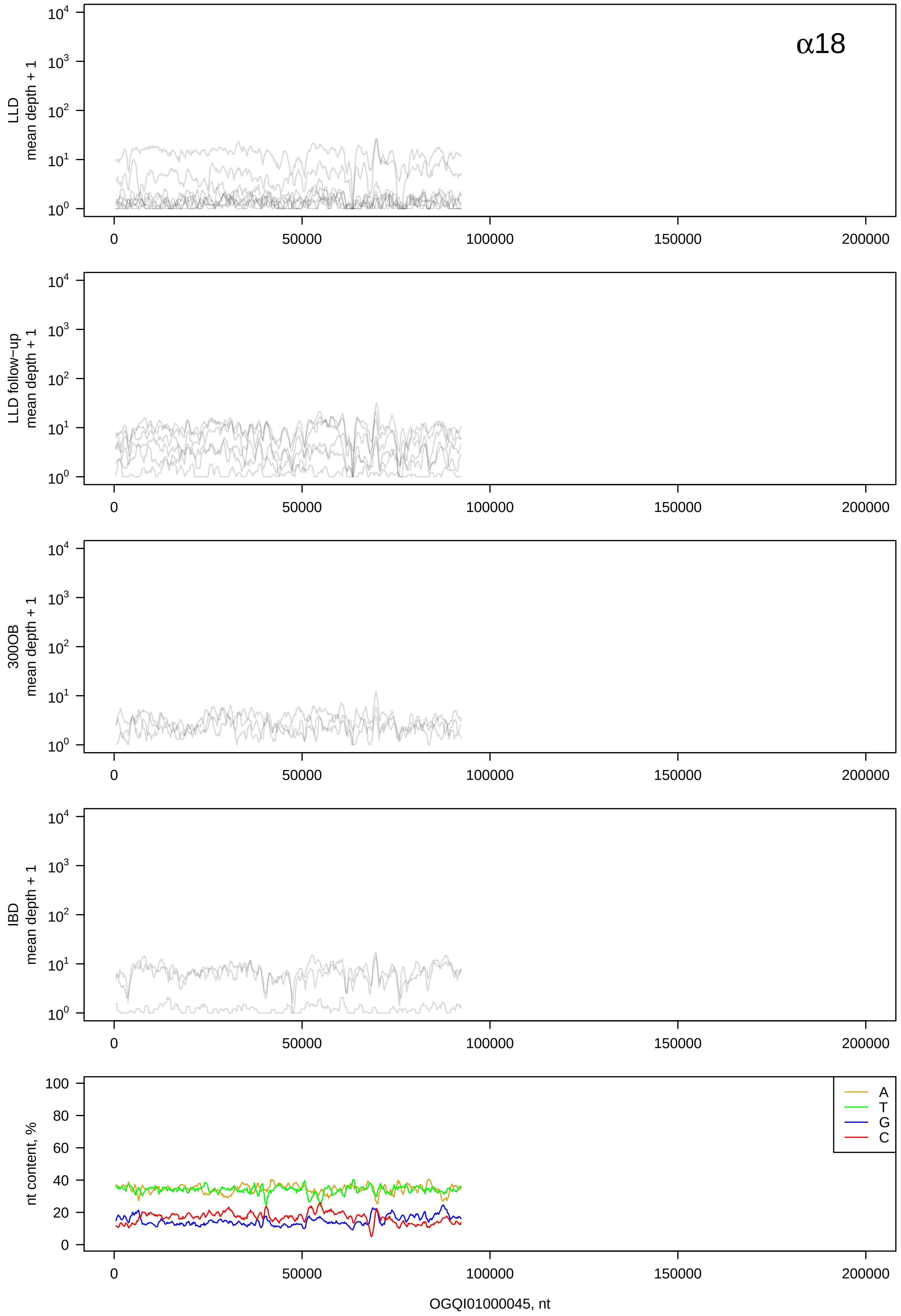

α18
LLD
mean depth + 1
LLD follow−up
mean depth + 1
300OB
mean depth + 1
IBD
mean depth + 1
nt content, %
A
T
G
C
OGQI01000045, nt

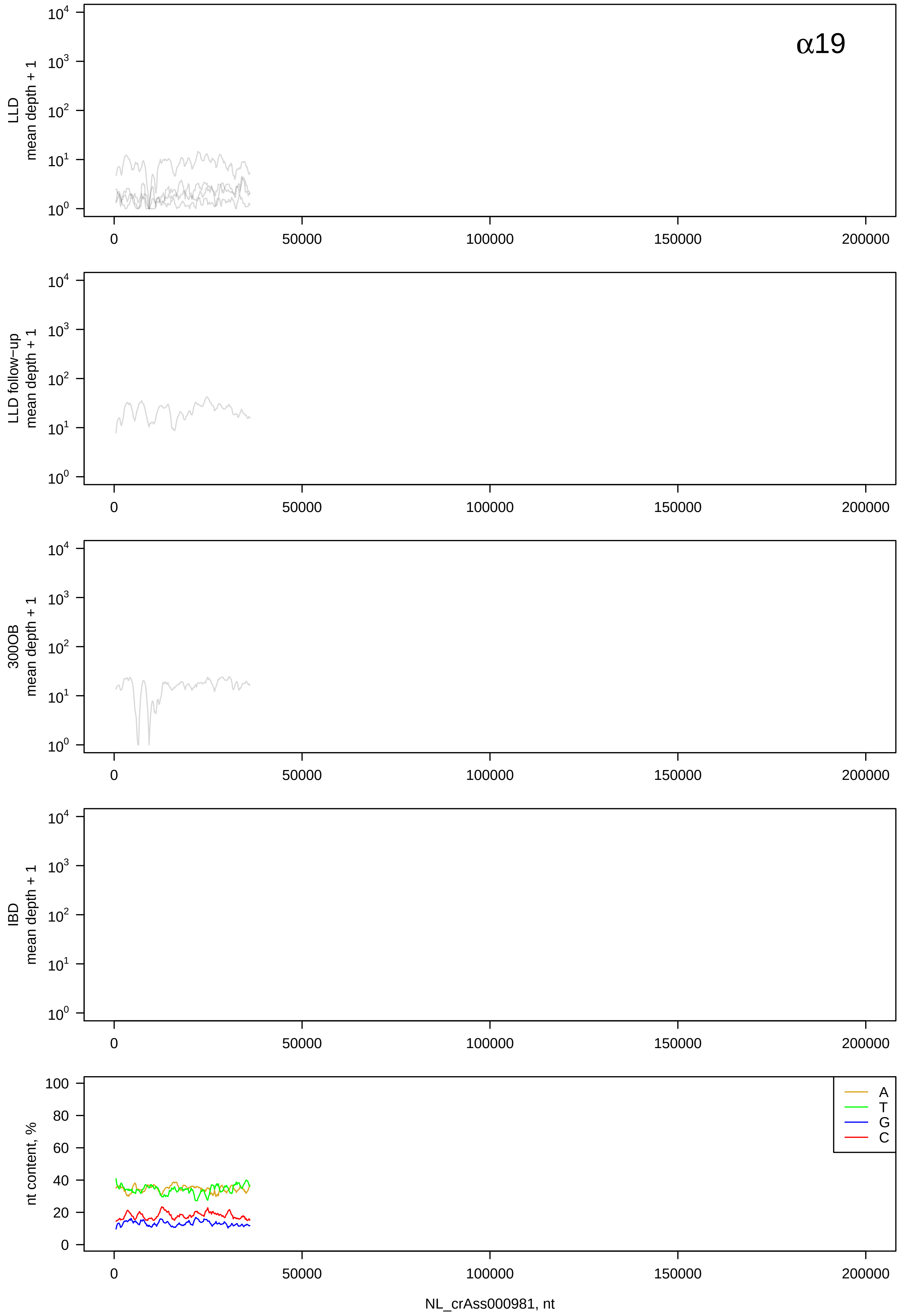

α19
LLD
mean depth + 1
LLD follow−up
mean depth + 1
300OB
mean depth + 1
IBD
mean depth + 1
nt content, %
A
T
G
C
NL_crAss000981, nt

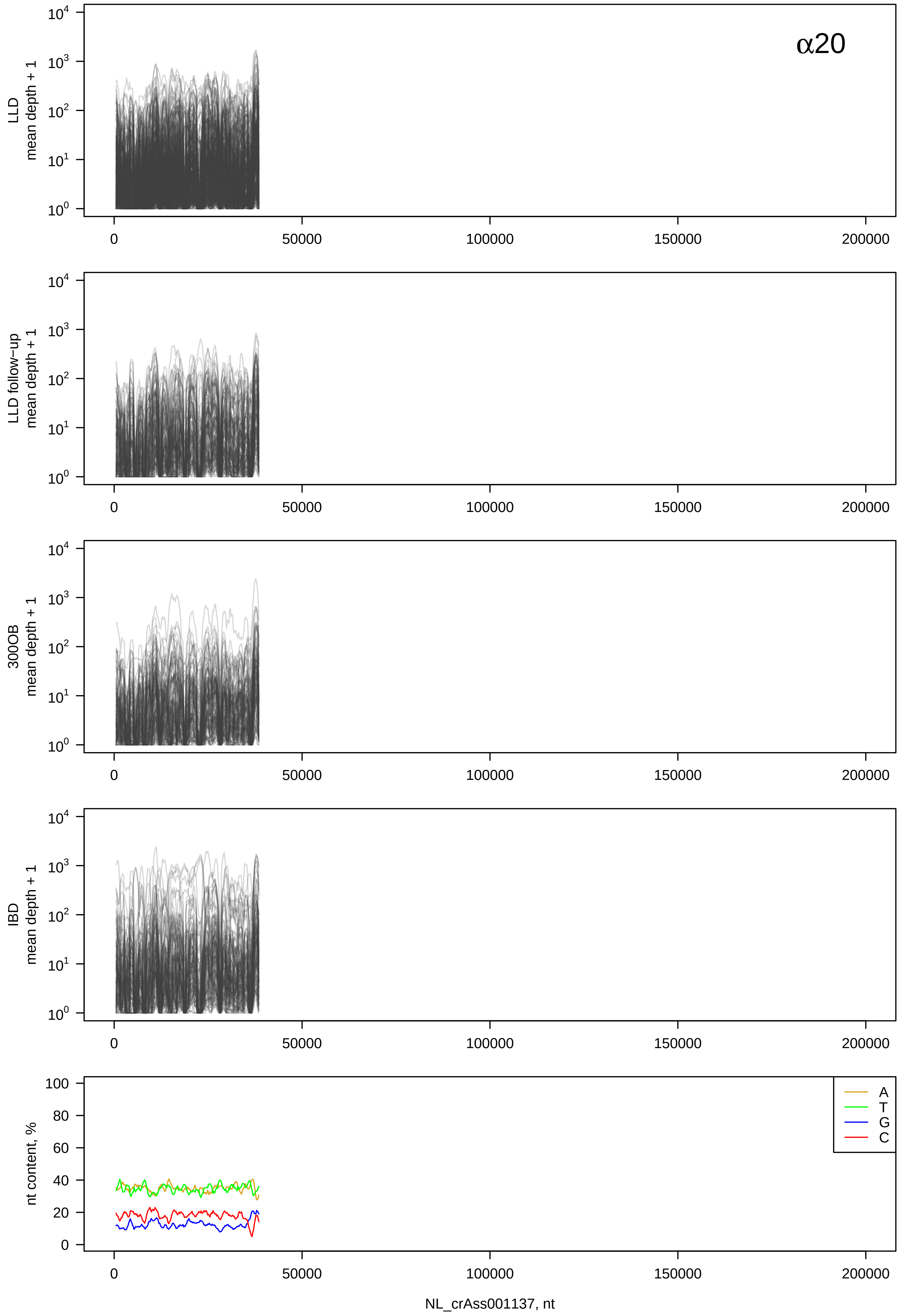

α20
LLD mean depth + 1
LLD follow-up mean depth + 1
300OB mean depth + 1
IBD mean depth + 1
nt content, %
A
T
G
C
0
50000
100000
150000
200000
NL_crAss001137, nt

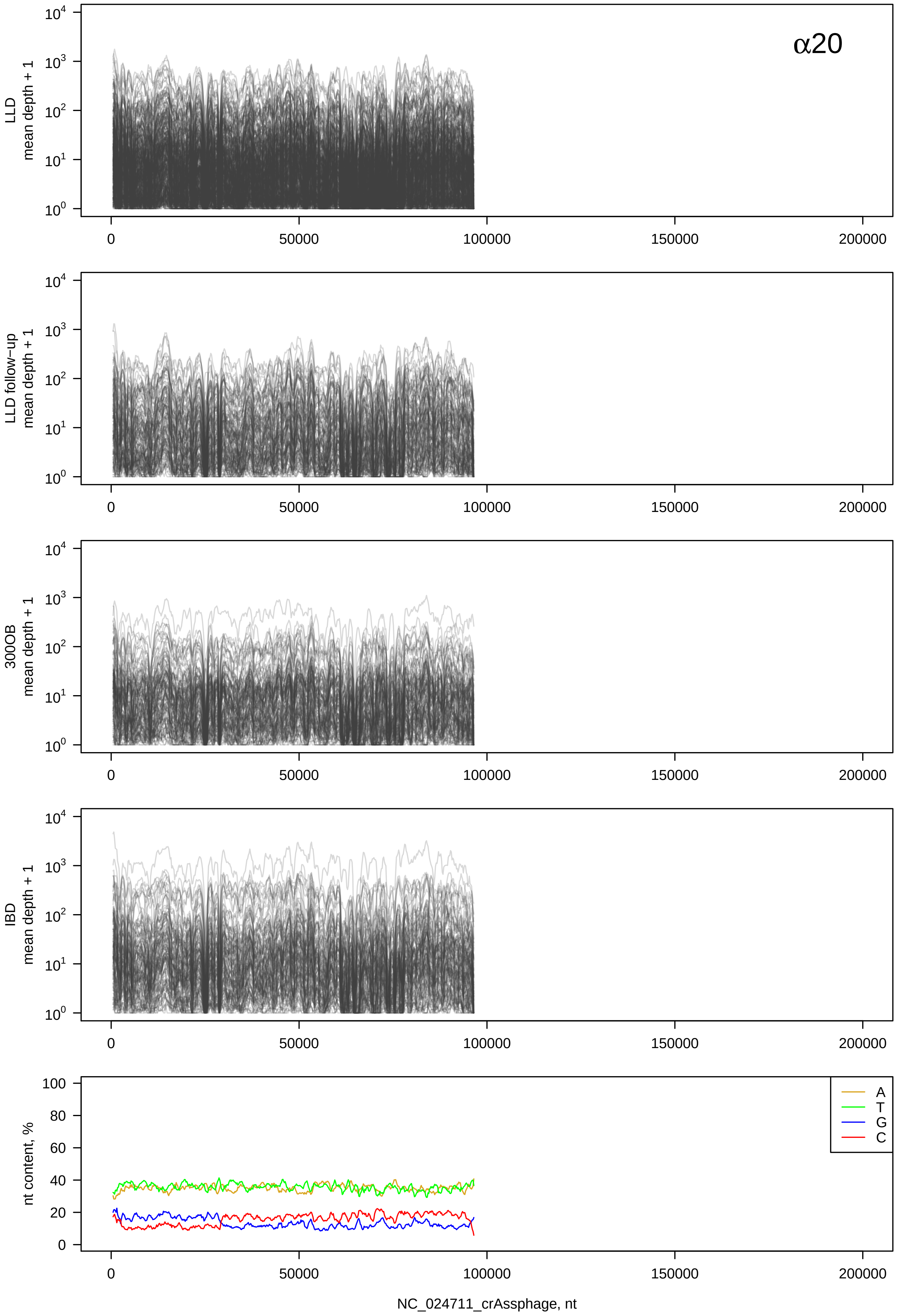

α20
LLD mean depth + 1
LLD follow-up mean depth + 1
300OB mean depth + 1
IBD mean depth + 1
nt content, %
A
T
G
C
0
50000
100000
150000
200000
NC_024711_crAssphage, nt

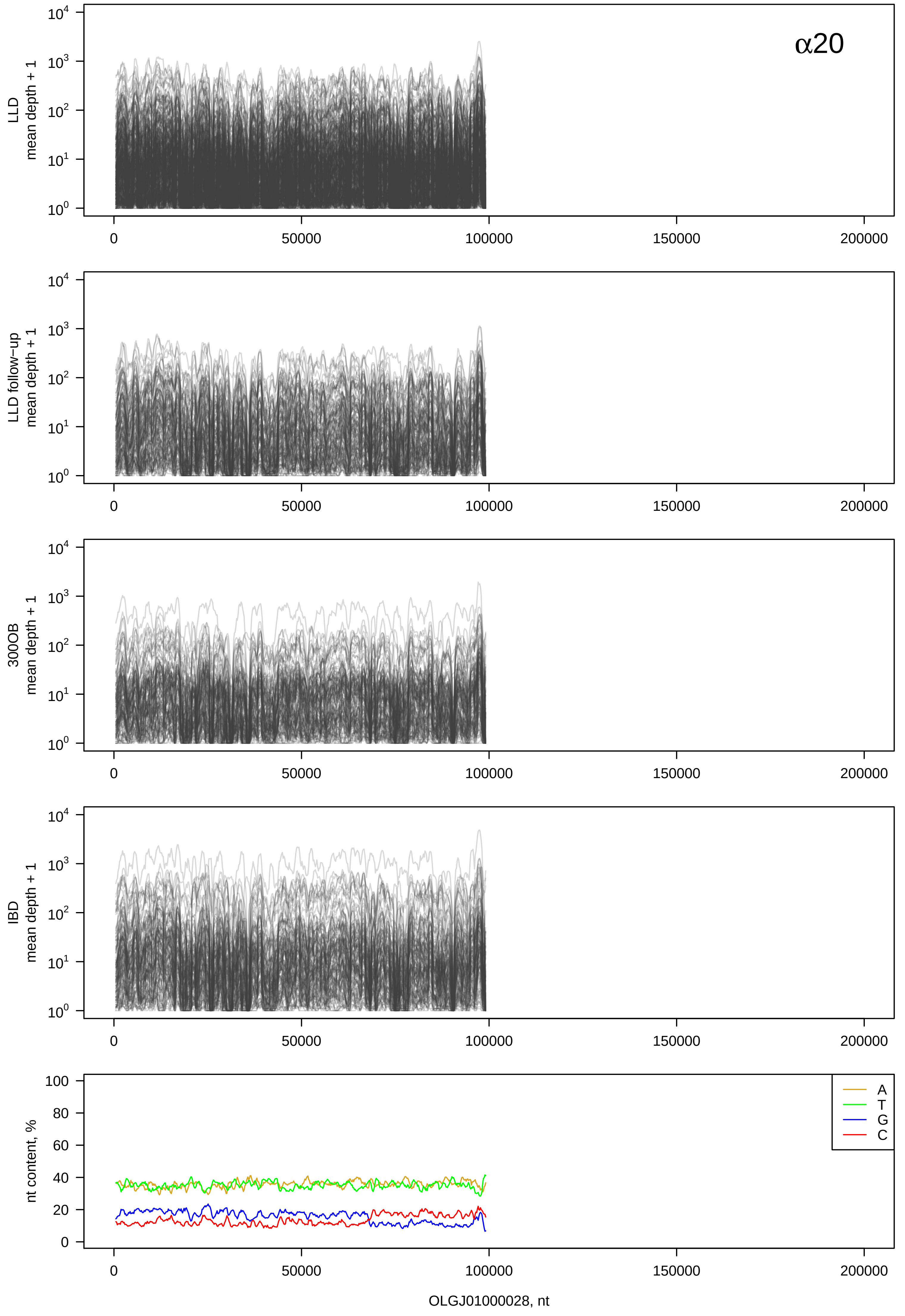

α20
LLD mean depth + 1
LLD follow−up mean depth + 1
300OB mean depth + 1
IBD mean depth + 1
nt content, %
A
T
G
C
OLGJ01000028, nt

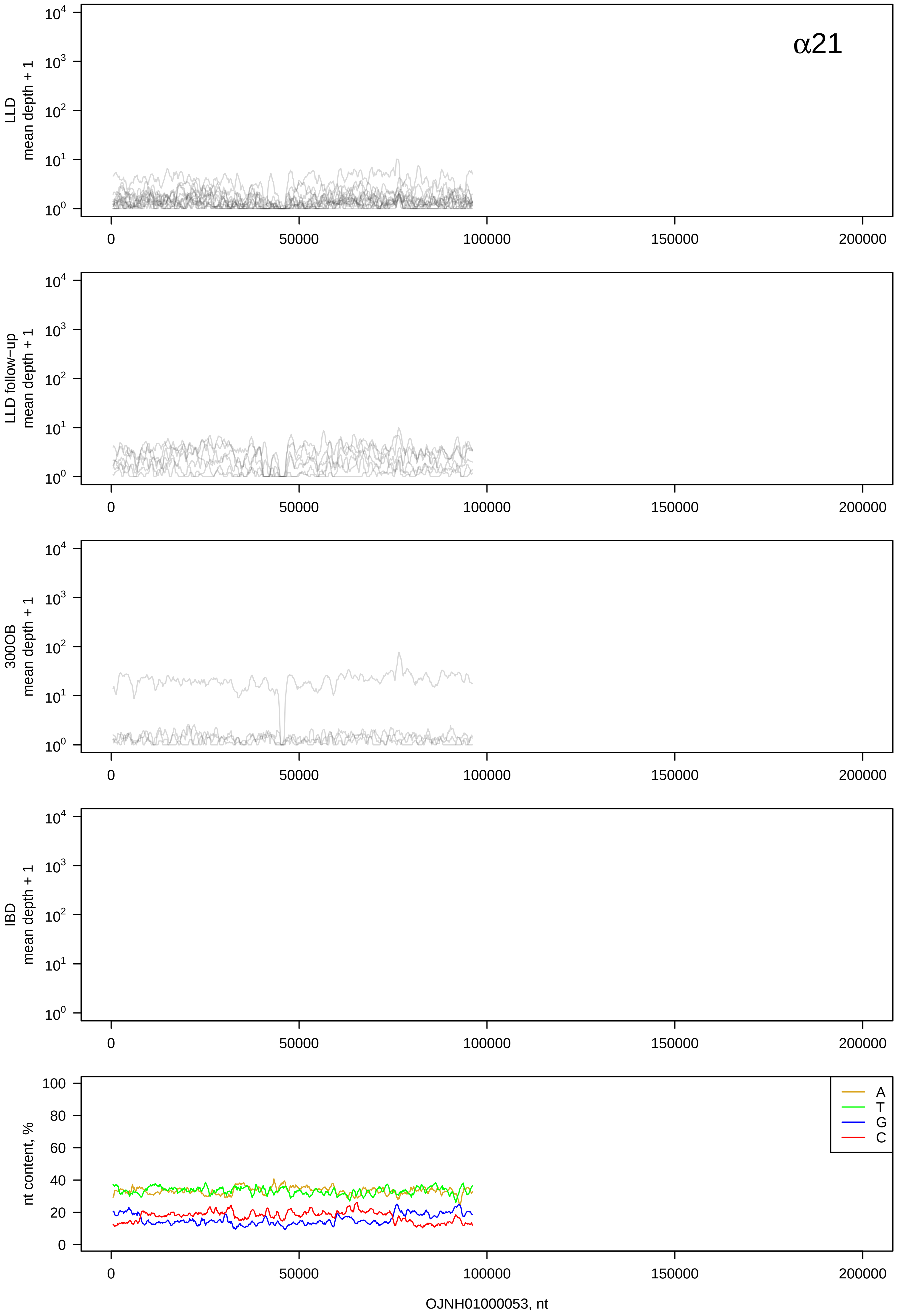

α21
LLD mean depth + 1
LLD follow−up mean depth + 1
300OB mean depth + 1
IBD mean depth + 1
nt content, %
A
T
G
C
0
50000
100000
150000
200000
OJNH01000053, nt

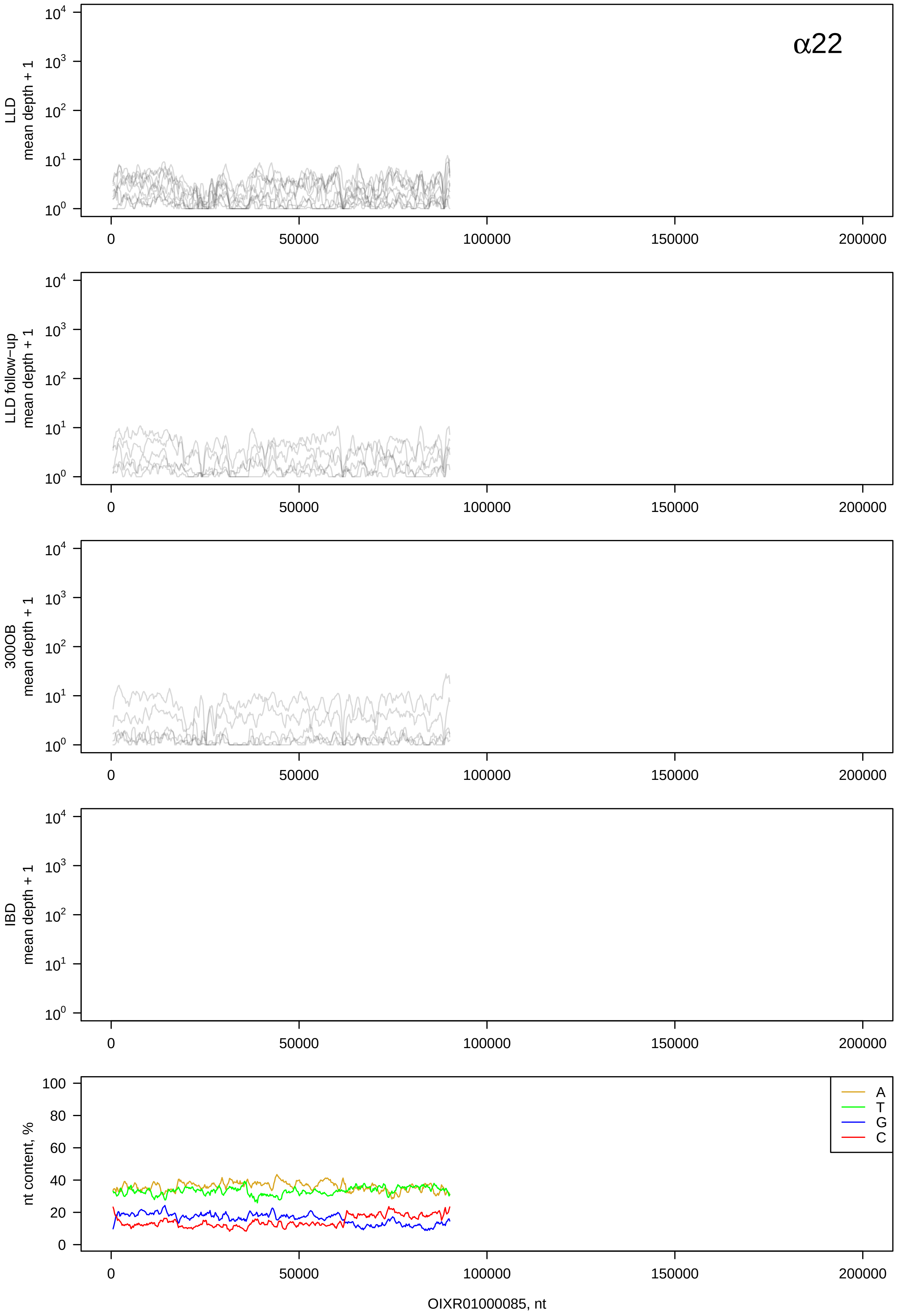

α22
LLD mean depth + 1
LLD follow−up mean depth + 1
300OB mean depth + 1
IBD mean depth + 1
nt content, %
A
T
G
C
OIXR01000085, nt

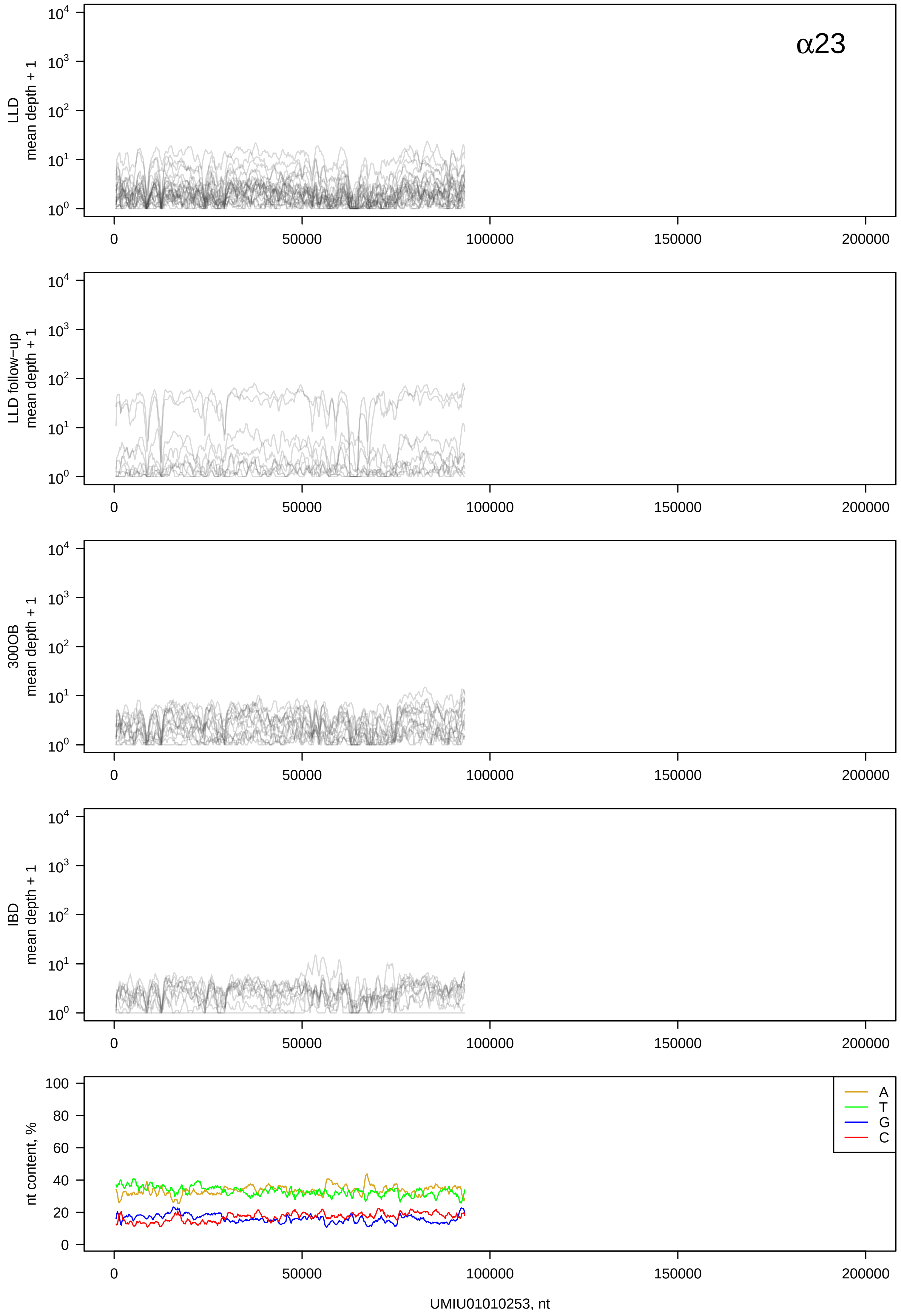

α23
LLD mean depth + 1
LLD follow-up mean depth + 1
300OB mean depth + 1
IBD mean depth + 1
nt content, %
A
T
G
C
UMIU01010253, nt

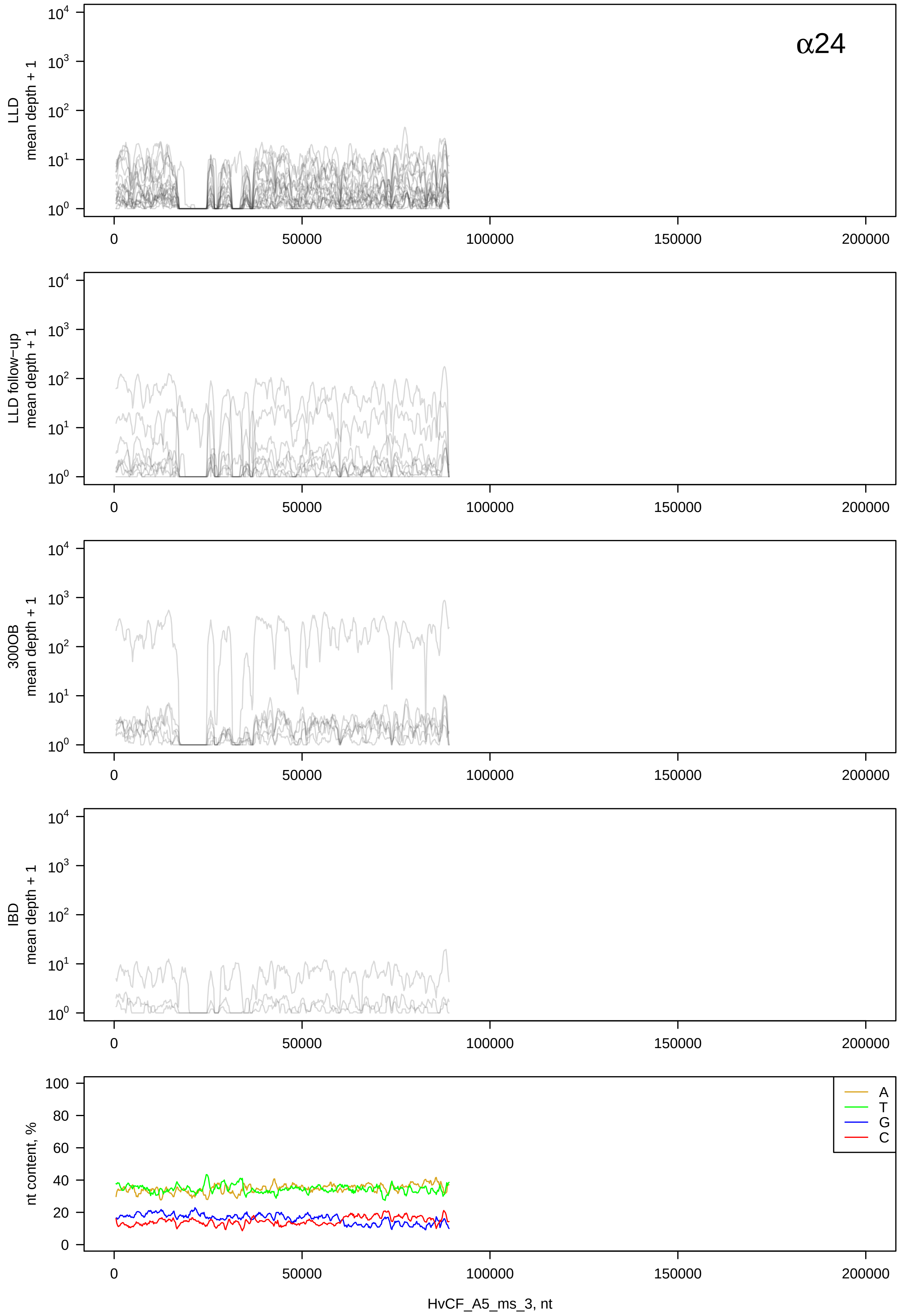

α24
LLD mean depth + 1
LLD follow−up mean depth + 1
300OB mean depth + 1
IBD mean depth + 1
nt content, %
A
T
G
C
0
50000
100000
150000
200000
HvCF_A5_ms_3, nt

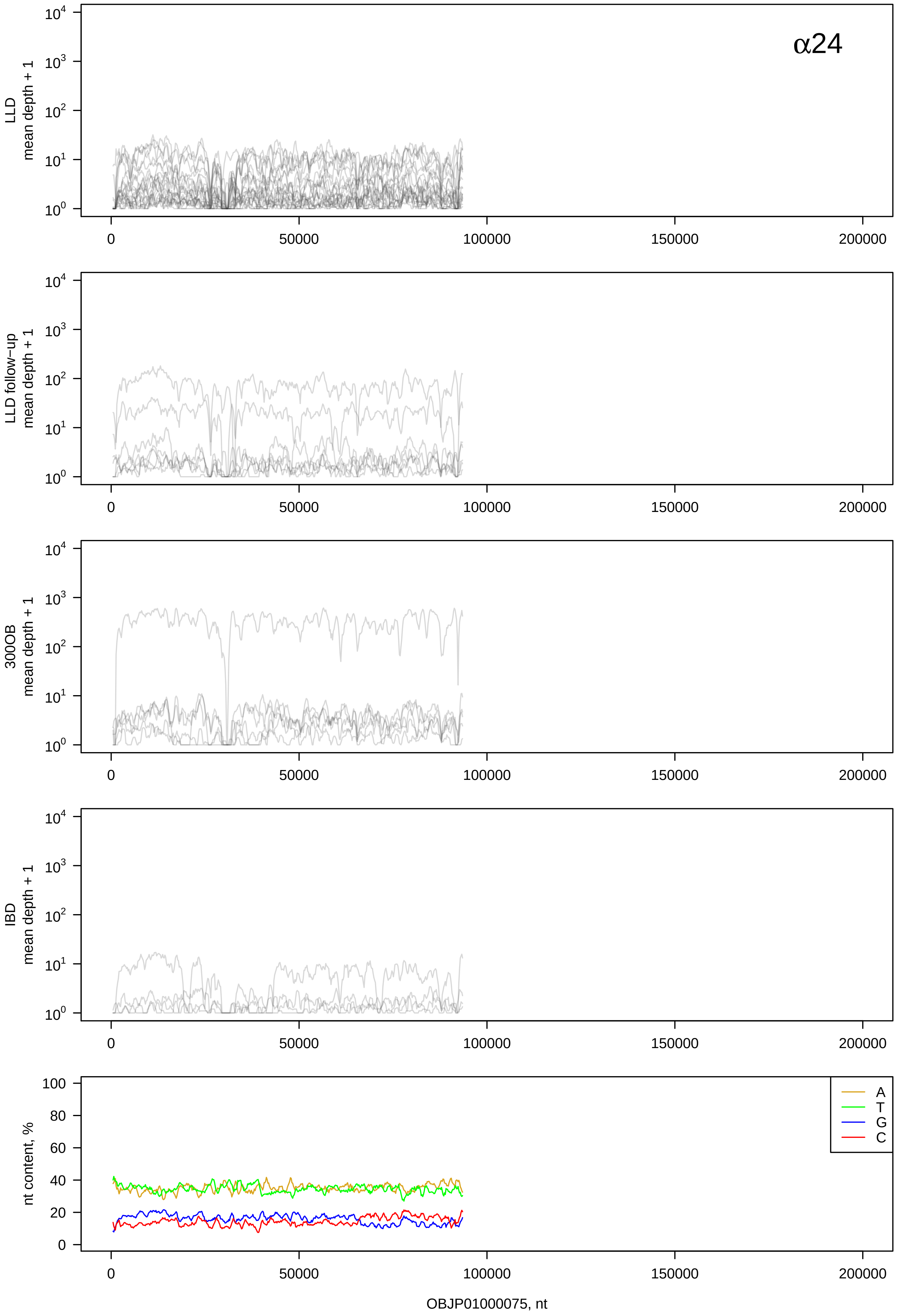
α24
LLD
mean depth + 1
LLD follow-up
mean depth + 1
300OB
mean depth + 1
IBD
mean depth + 1
nt content, %
A
T
G
C
OBJP01000075, nt

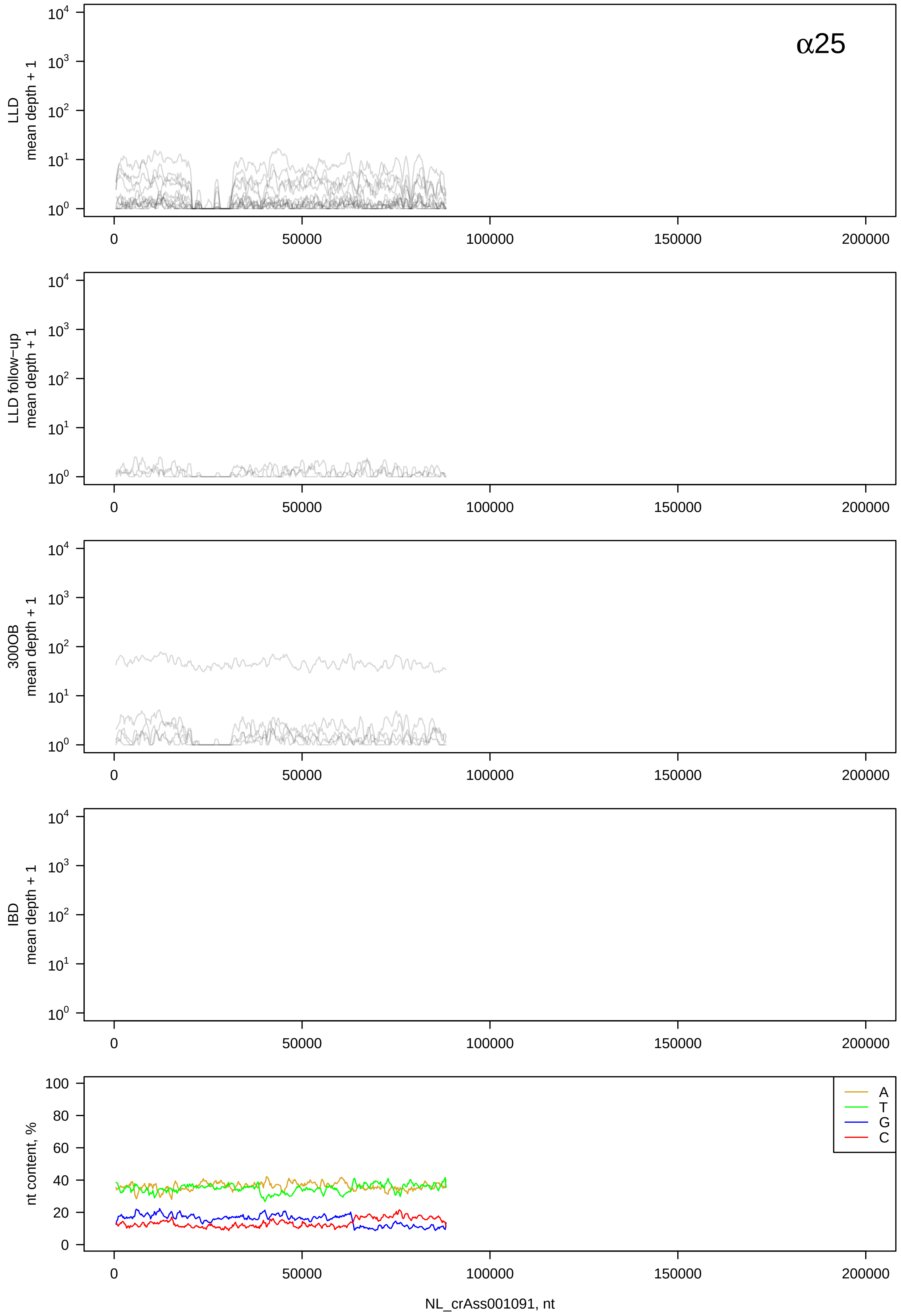
α25
LLD
mean depth + 1
LLD follow-up
mean depth + 1
300OB
mean depth + 1
IBD
mean depth + 1
nt content, %
A
T
G
C
NL_crAss001091, nt

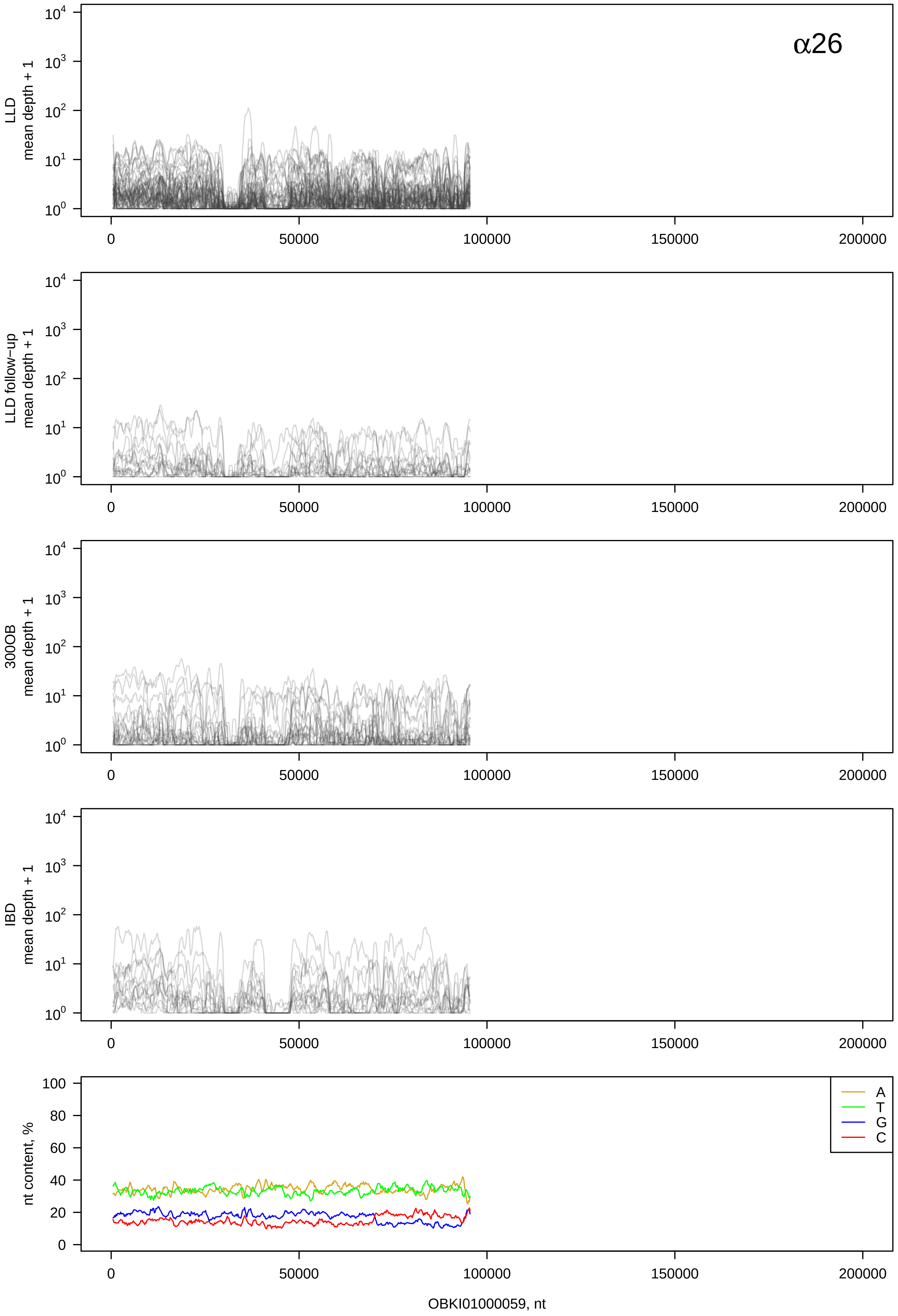

α26
LLD mean depth + 1
LLD follow−up mean depth + 1
300OB mean depth + 1
IBD mean depth + 1
nt content, %
A
T
G
C
0
50000
100000
150000
200000
OBKI01000059, nt

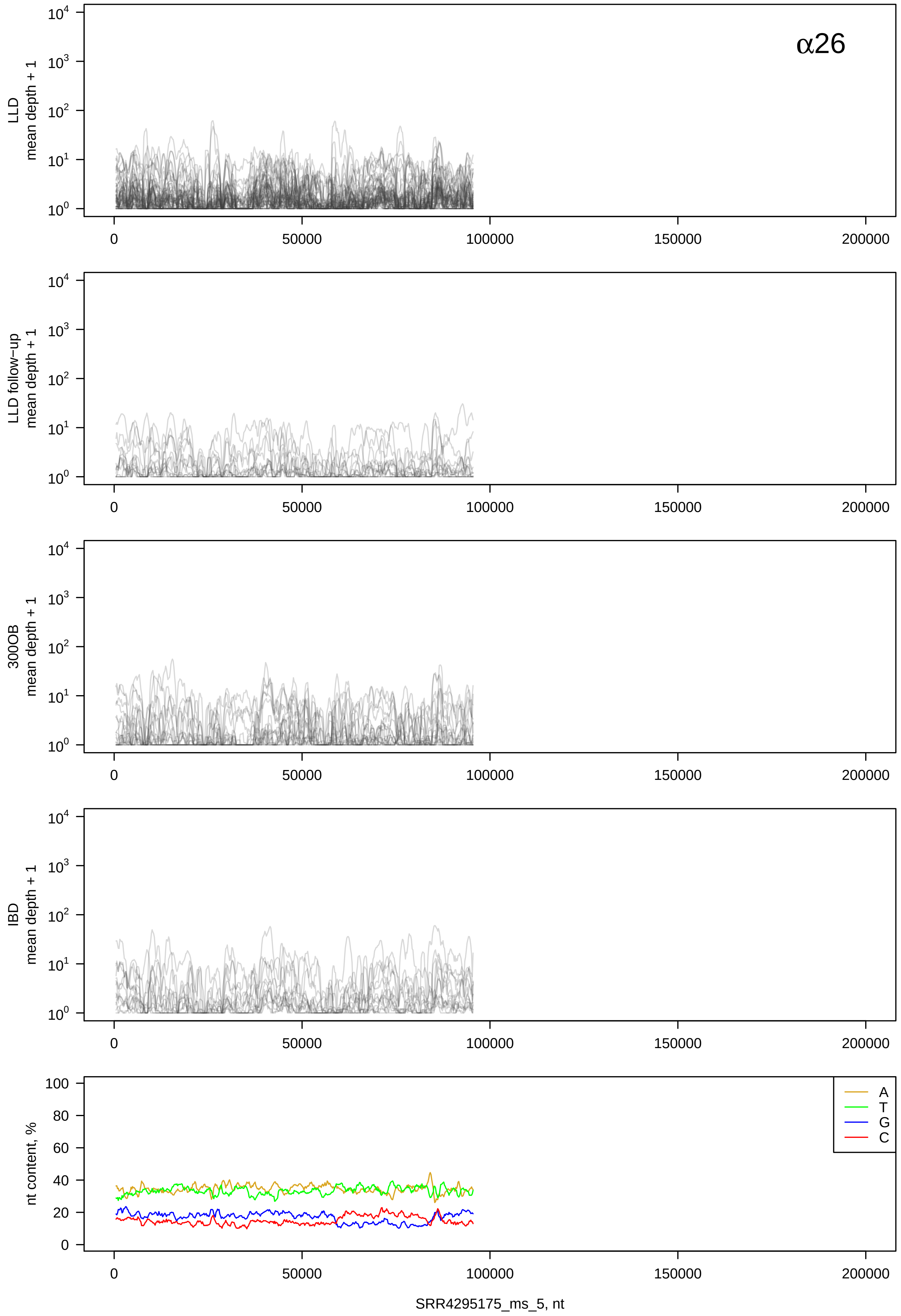
α26
LLD mean depth + 1
LLD follow-up mean depth + 1
300OB mean depth + 1
IBD mean depth + 1
nt content, %
A
T
G
C
SRR4295175_ms_5, nt

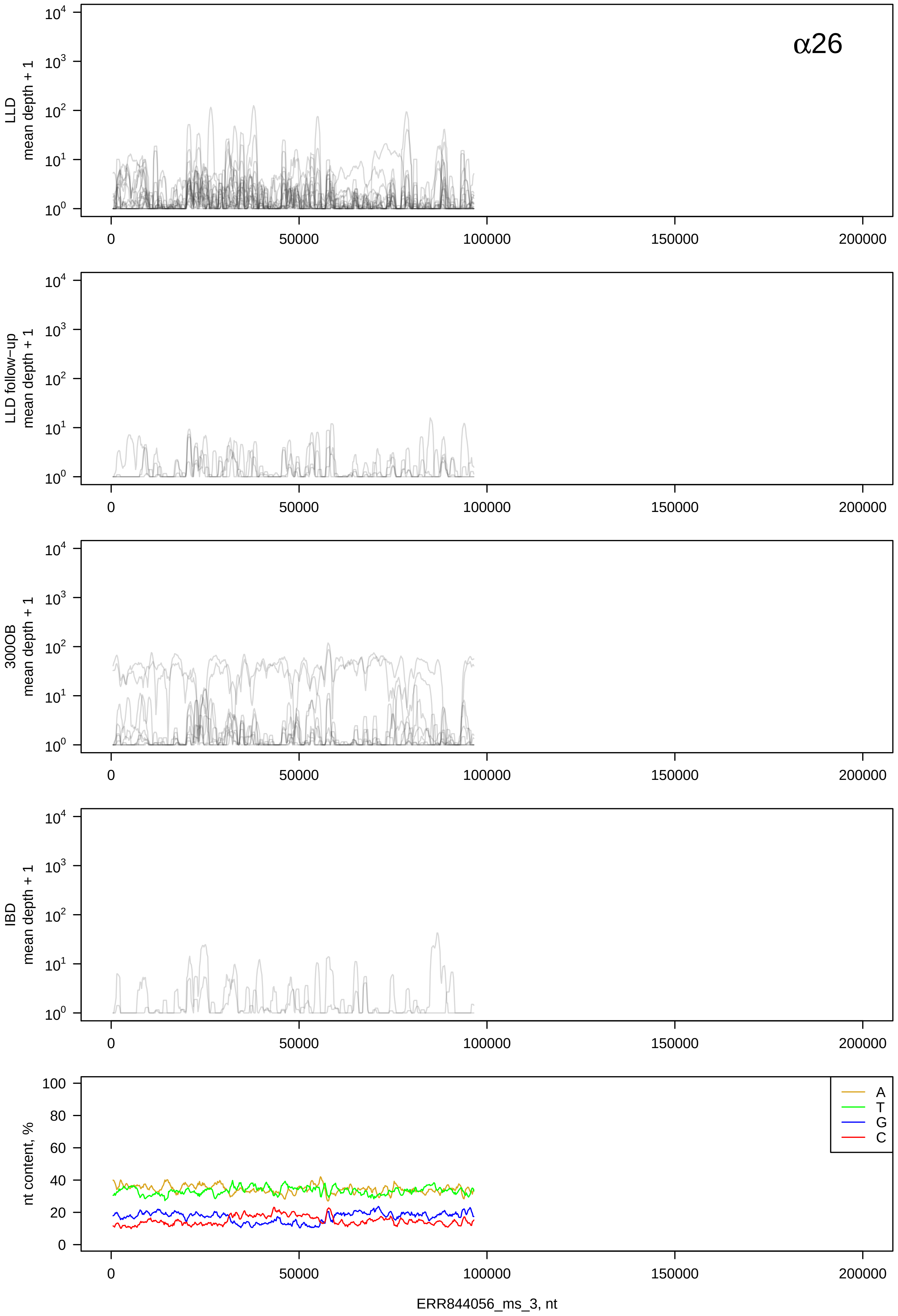

α26
LLD
mean depth + 1
LLD follow−up
mean depth + 1
300OB
mean depth + 1
IBD
mean depth + 1
nt content, %
A
T
G
C
0
50000
100000
150000
200000
ERR844056_ms_3, nt

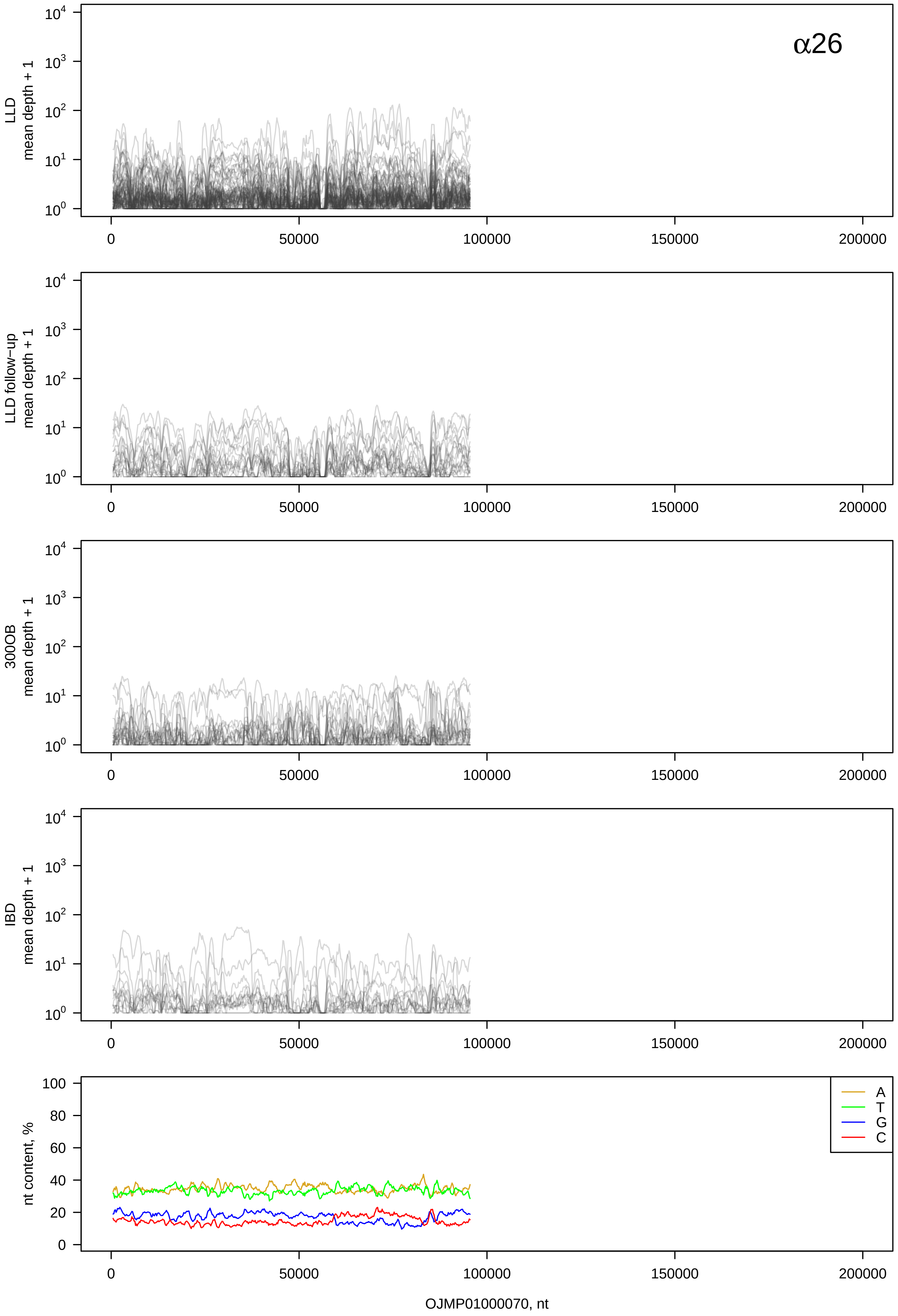
α26
LLD mean depth + 1
LLD follow-up mean depth + 1
300OB mean depth + 1
IBD mean depth + 1
nt content, %
$10^0$
$10^1$
$10^2$
$10^3$
$10^4$
0
20
40
60
80
100
0
50000
100000
150000
200000
A
T
G
C
OJMP01000070, nt

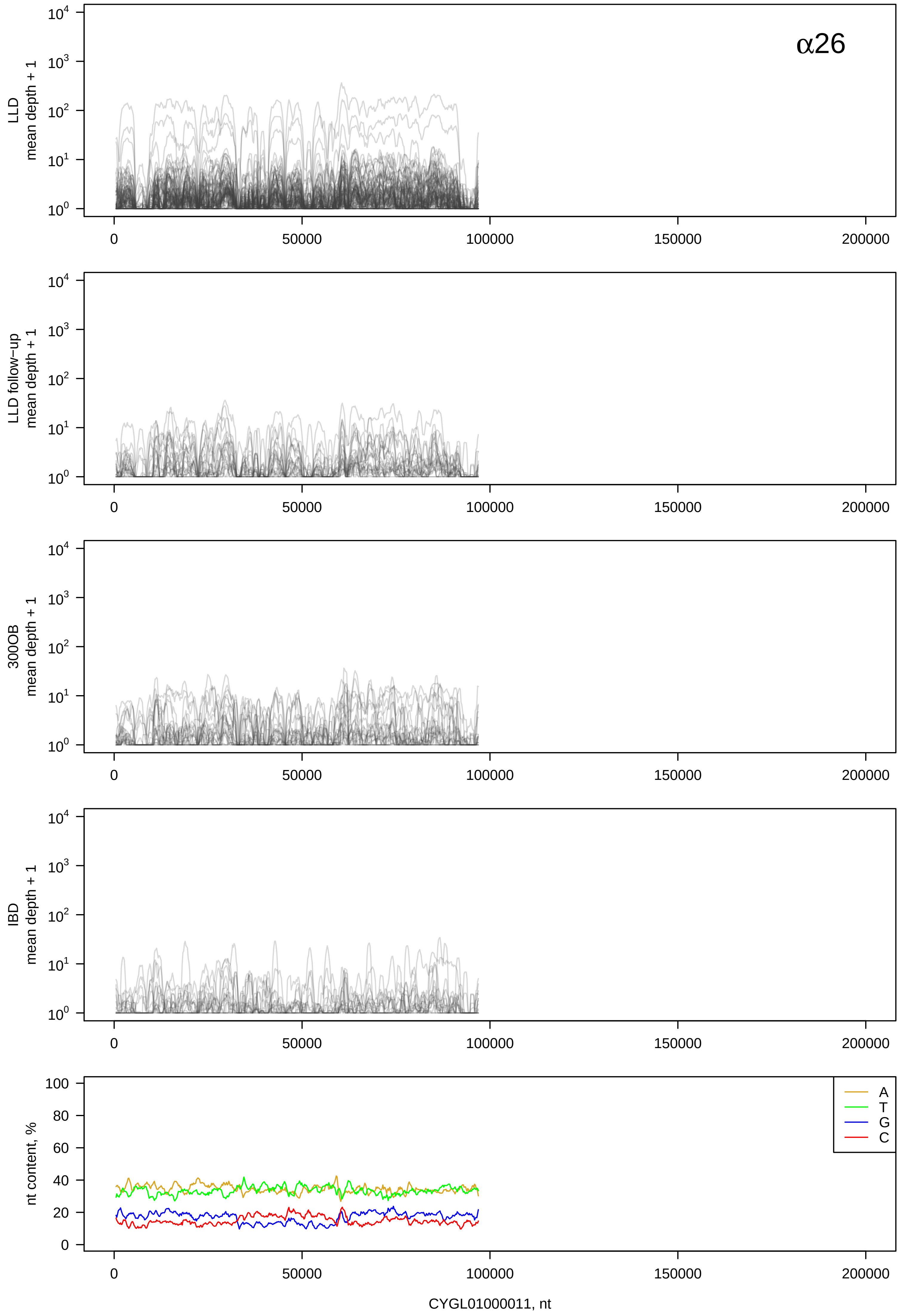

α26
LLD mean depth + 1
LLD follow-up mean depth + 1
300OB mean depth + 1
IBD mean depth + 1
nt content, %
A
T
G
C
CYGL01000011, nt

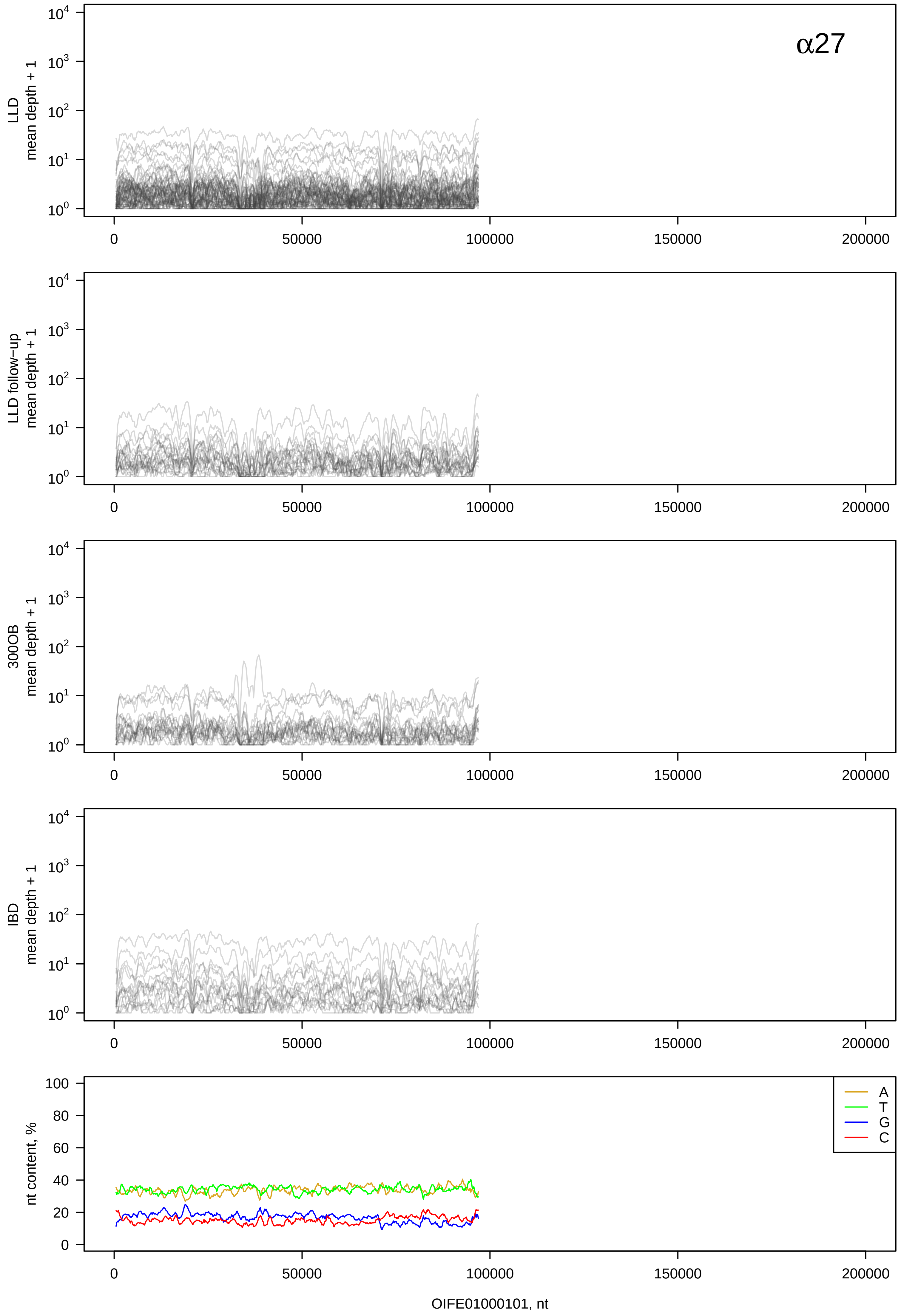

α27
LLD
mean depth + 1
LLD follow−up
mean depth + 1
300OB
mean depth + 1
IBD
mean depth + 1
nt content, %
A
T
G
C
OIFE01000101, nt

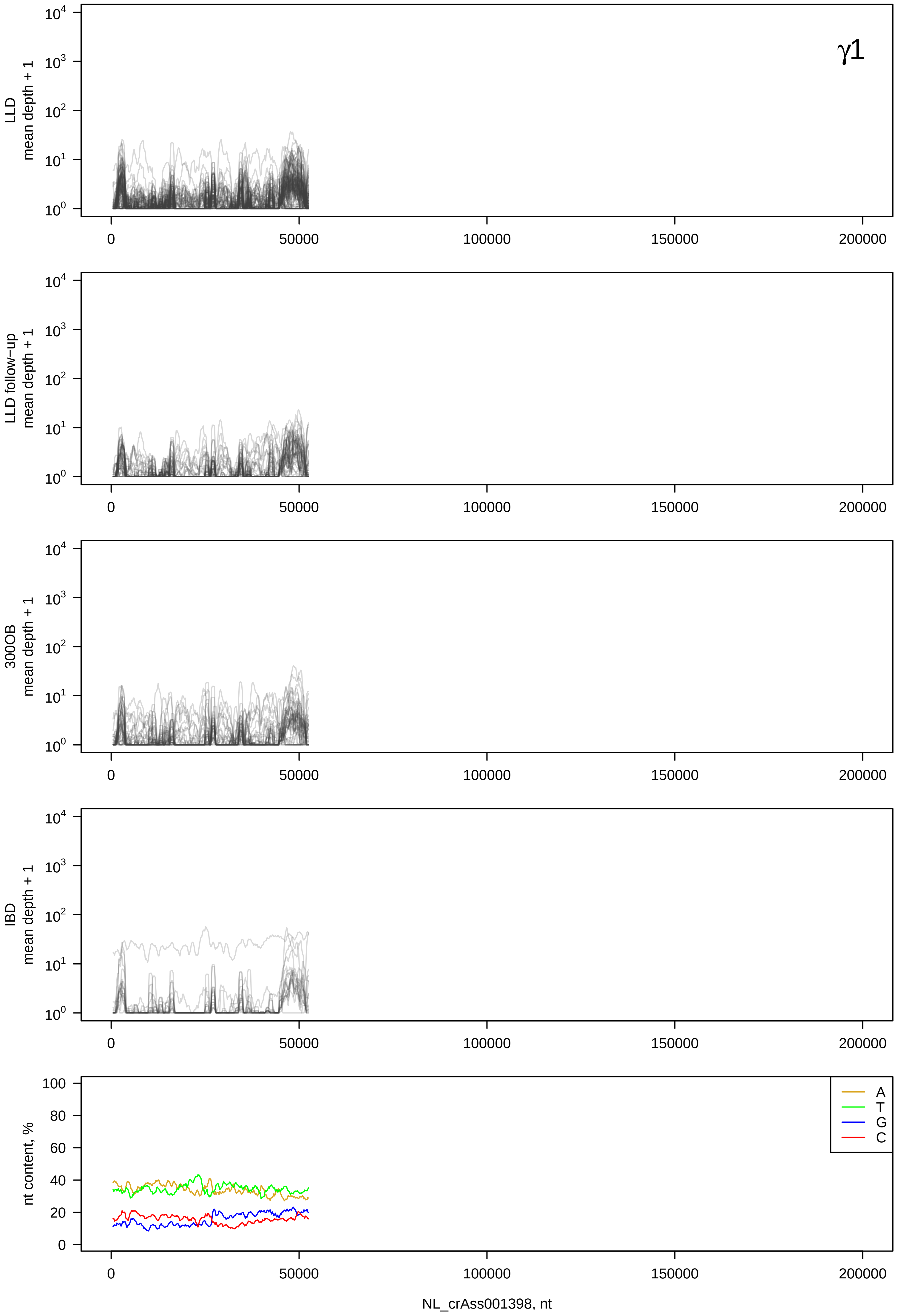
γ1
LLD mean depth + 1
LLD follow−up mean depth + 1
300OB mean depth + 1
IBD mean depth + 1
nt content, %
A
T
G
C
0
50000
100000
150000
200000
NL_crAss001398, nt

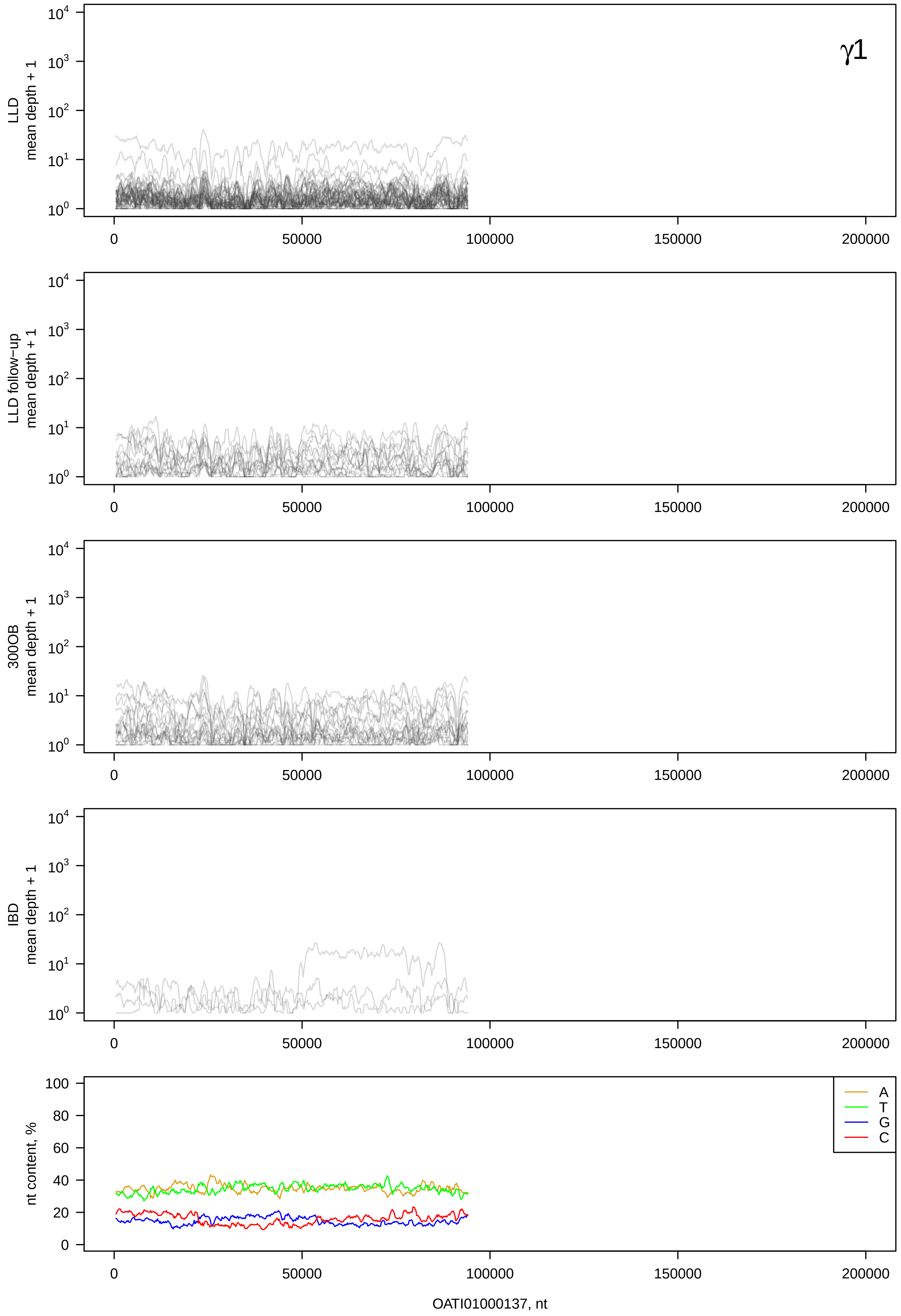
γ1
LLD mean depth + 1
LLD follow−up mean depth + 1
300OB mean depth + 1
IBD mean depth + 1
nt content, %
A
T
G
C
0
50000
100000
150000
200000
OATI01000137, nt

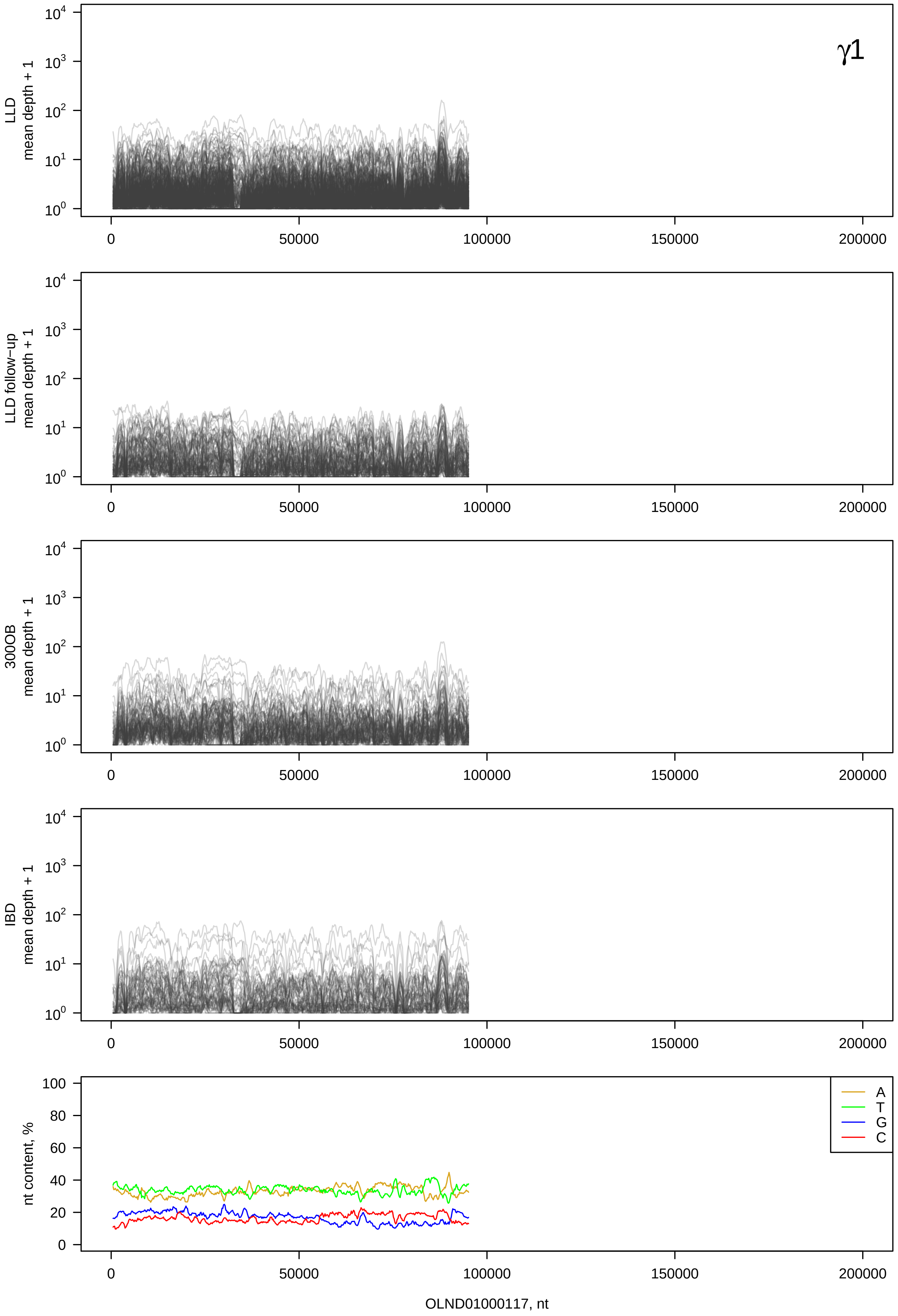
γ1
LLD mean depth + 1
LLD follow-up mean depth + 1
300OB mean depth + 1
IBD mean depth + 1
nt content, %
A
T
G
C
0
50000
100000
150000
200000
OLND01000117, nt

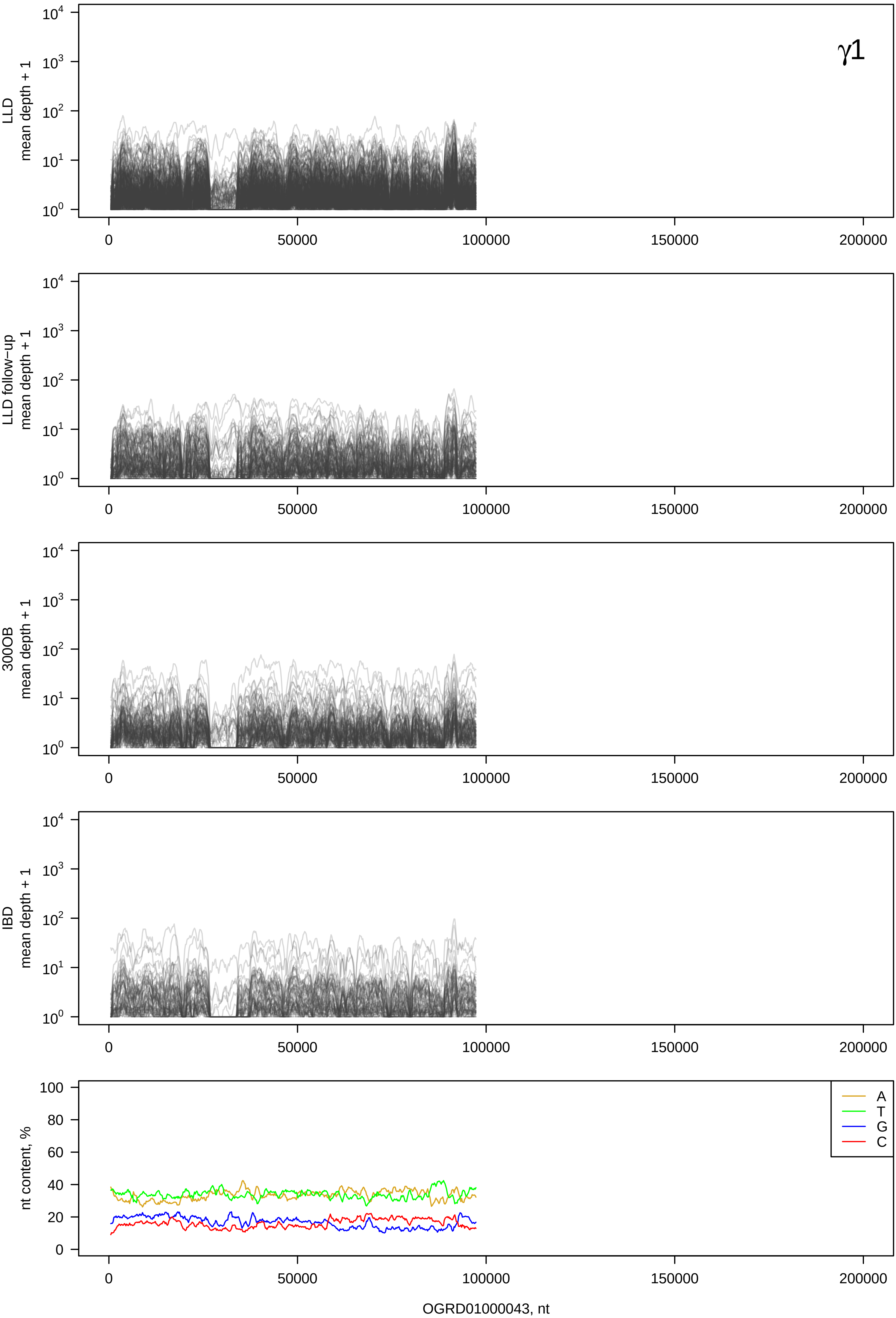

γ1
LLD mean depth + 1
LLD follow−up mean depth + 1
300OB mean depth + 1
IBD mean depth + 1
nt content, %
A
T
G
C
OGRD01000043, nt

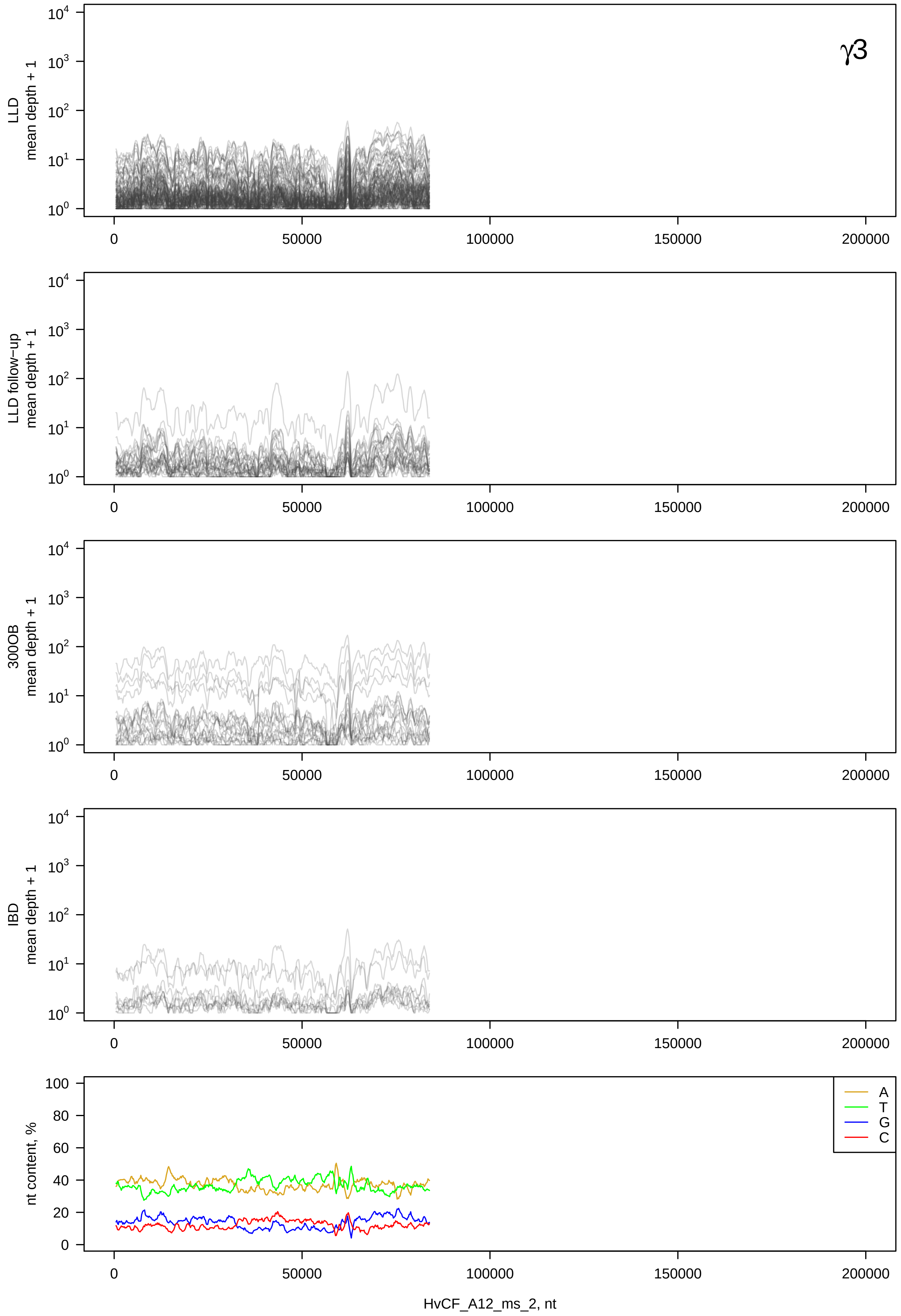
γ3
LLD mean depth + 1
LLD follow−up mean depth + 1
300OB mean depth + 1
IBD mean depth + 1
nt content, %
A
T
G
C
HvCF_A12_ms_2, nt

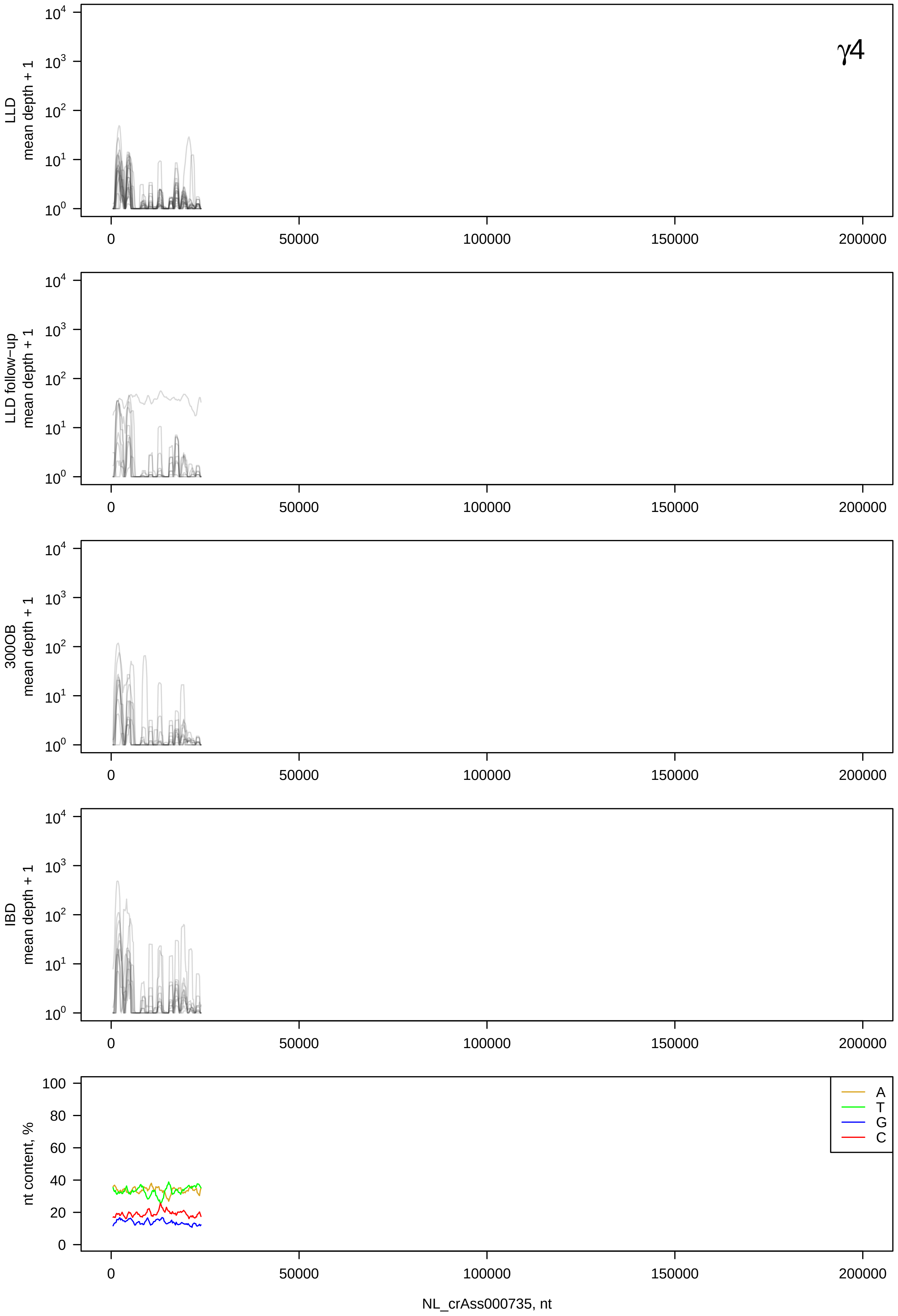

γ4
LLD mean depth + 1
LLD follow−up mean depth + 1
300OB mean depth + 1
IBD mean depth + 1
nt content, %
A
T
G
C
0
50000
100000
150000
200000
NL_crAss000735, nt

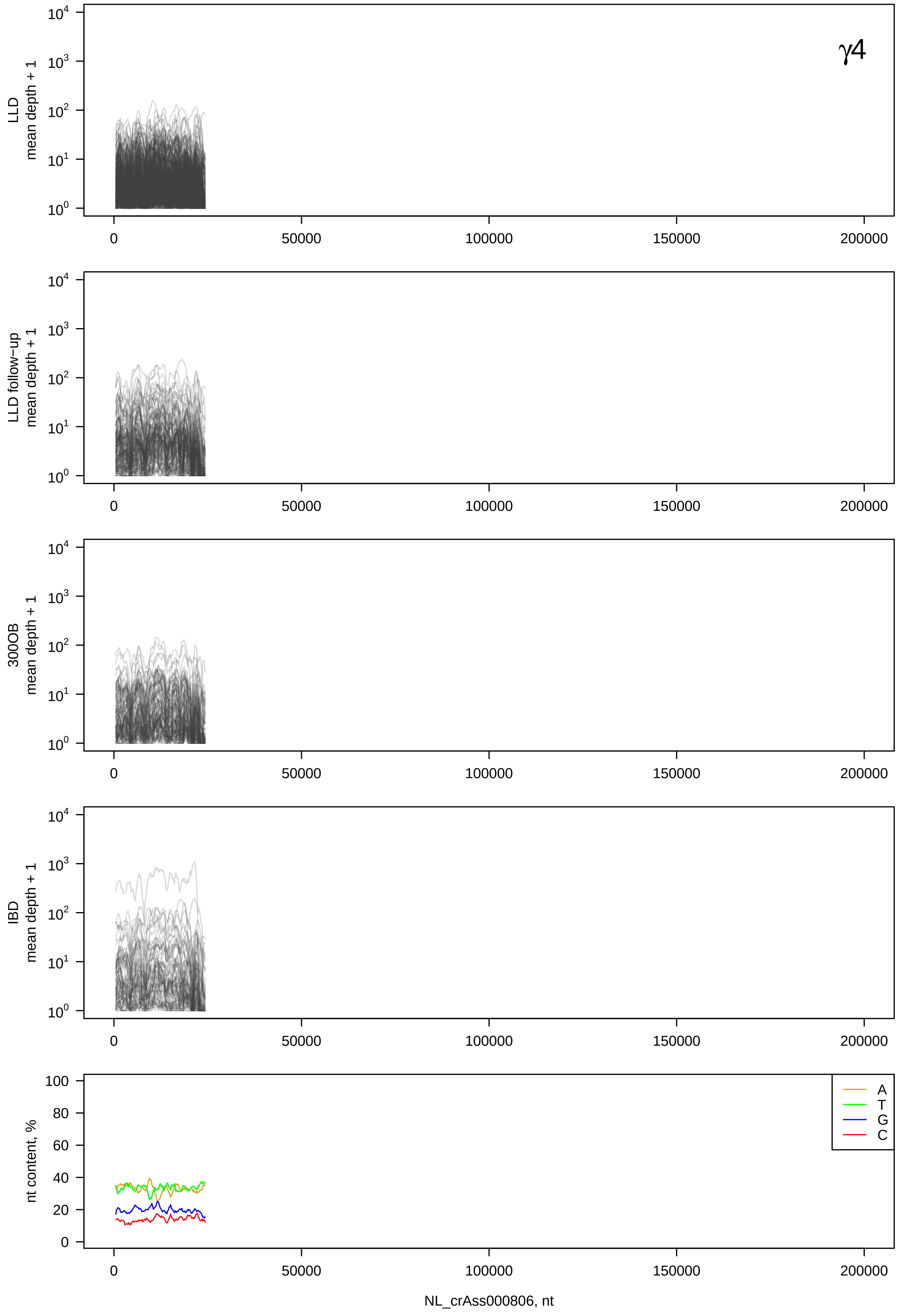

γ4
LLD
mean depth + 1
LLD follow−up
mean depth + 1
300OB
mean depth + 1
IBD
mean depth + 1
nt content, %
A
T
G
C
0
50000
100000
150000
200000
NL_crAss000806, nt

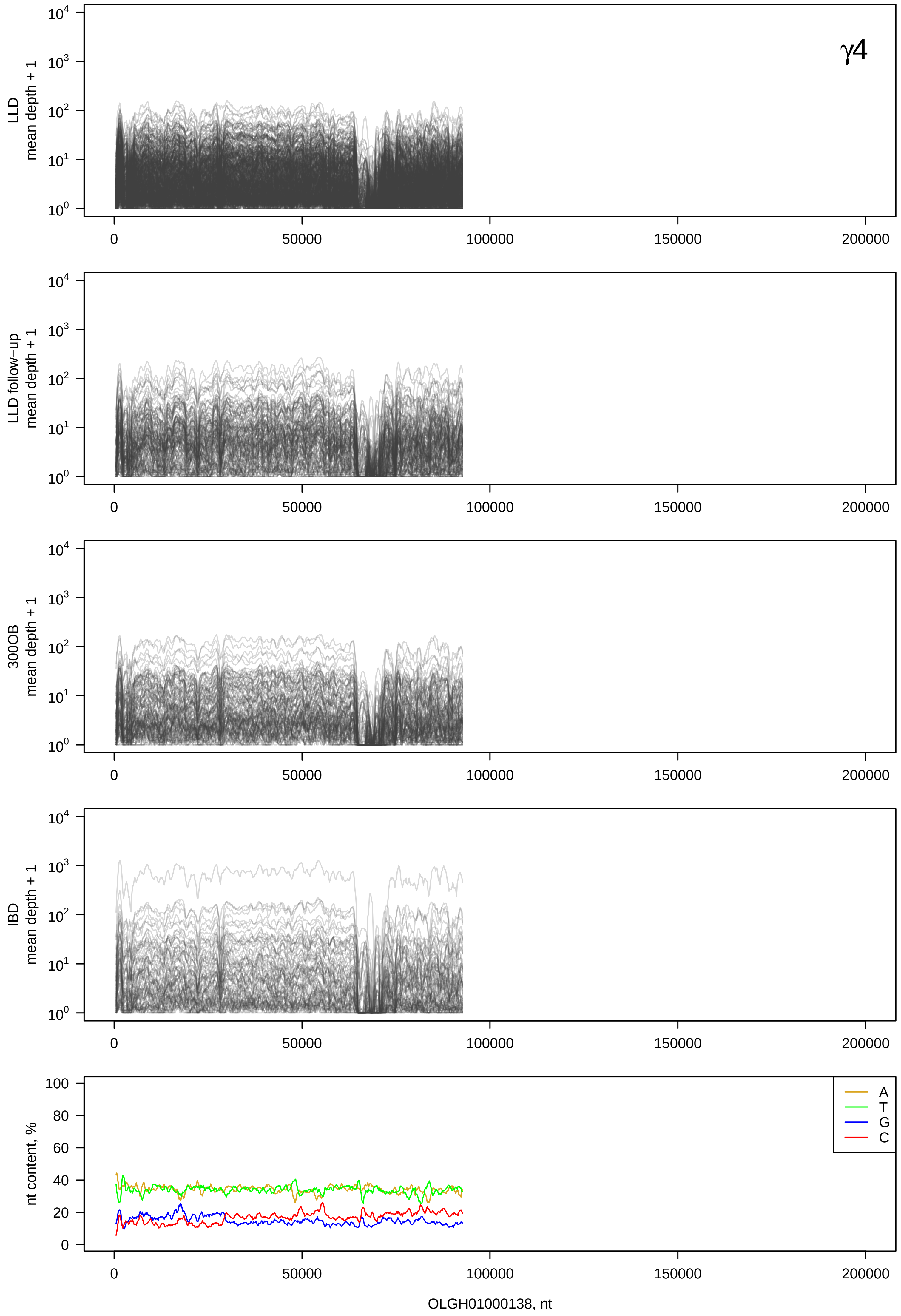
γ4
LLD mean depth + 1
LLD follow-up mean depth + 1
300OB mean depth + 1
IBD mean depth + 1
nt content, %
A
T
G
C
0
50000
100000
150000
200000
OLGH01000138, nt

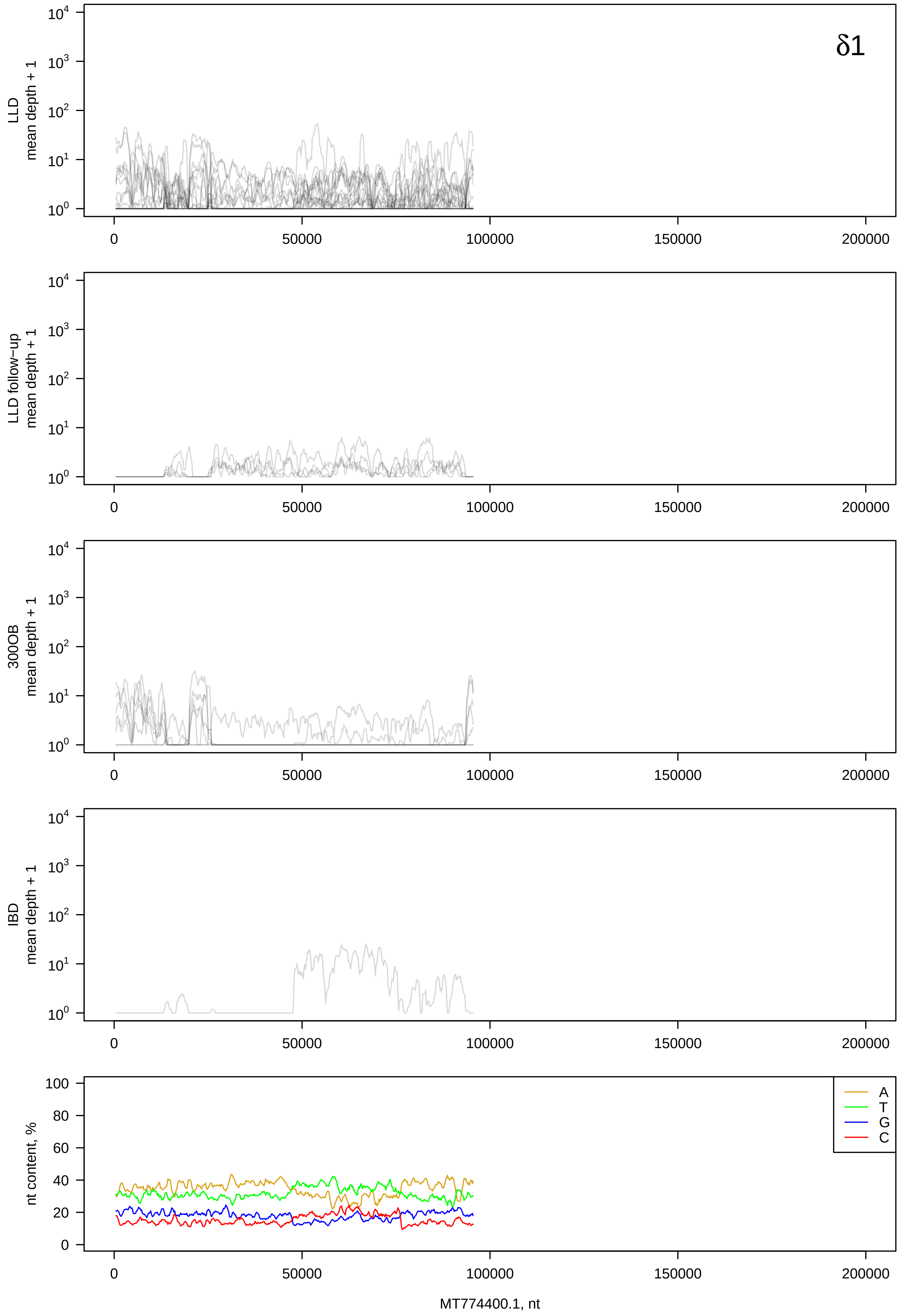

δ1
LLD
mean depth + 1
LLD follow−up
mean depth + 1
300OB
mean depth + 1
IBD
mean depth + 1
nt content, %
A
T
G
C
MT774400.1, nt

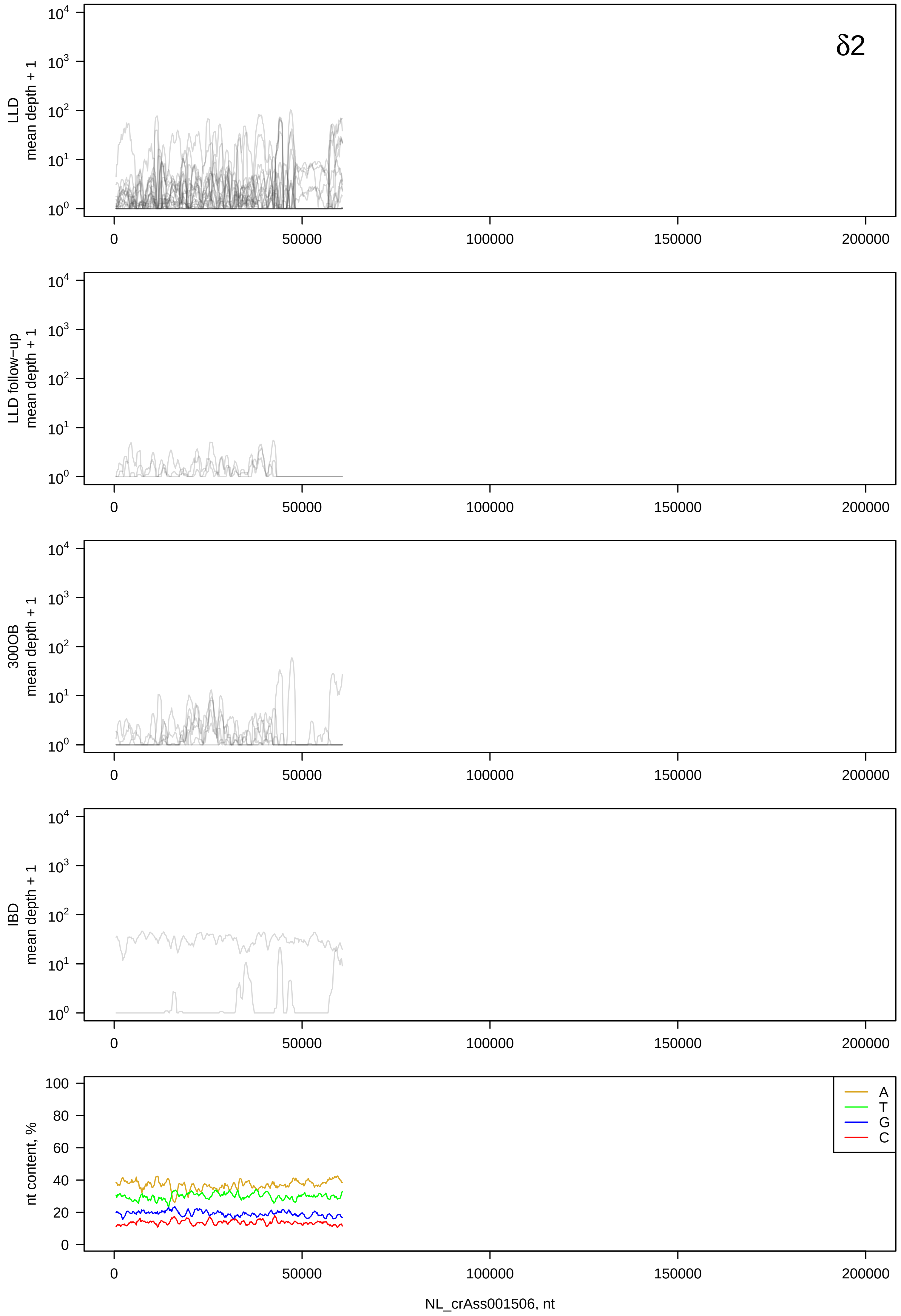
δ2
LLD
mean depth + 1
LLD follow−up
mean depth + 1
300OB
mean depth + 1
IBD
mean depth + 1
nt content, %
A
T
G
C
0
50000
100000
150000
200000
NL_crAss001506, nt

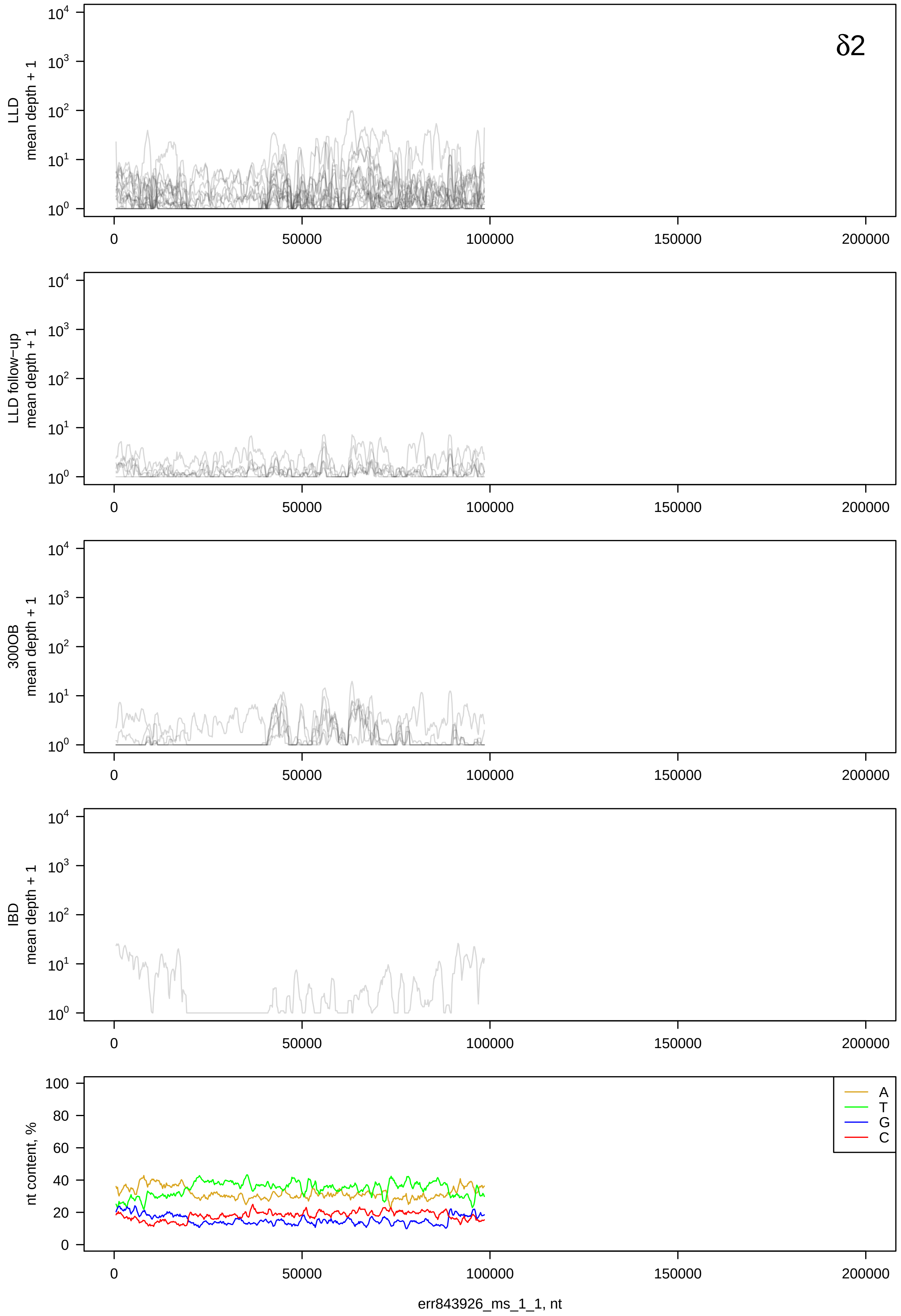

δ2
LLD
mean depth + 1
LLD follow-up
mean depth + 1
300OB
mean depth + 1
IBD
mean depth + 1
nt content, %
A
T
G
C
err843926_ms_1_1, nt

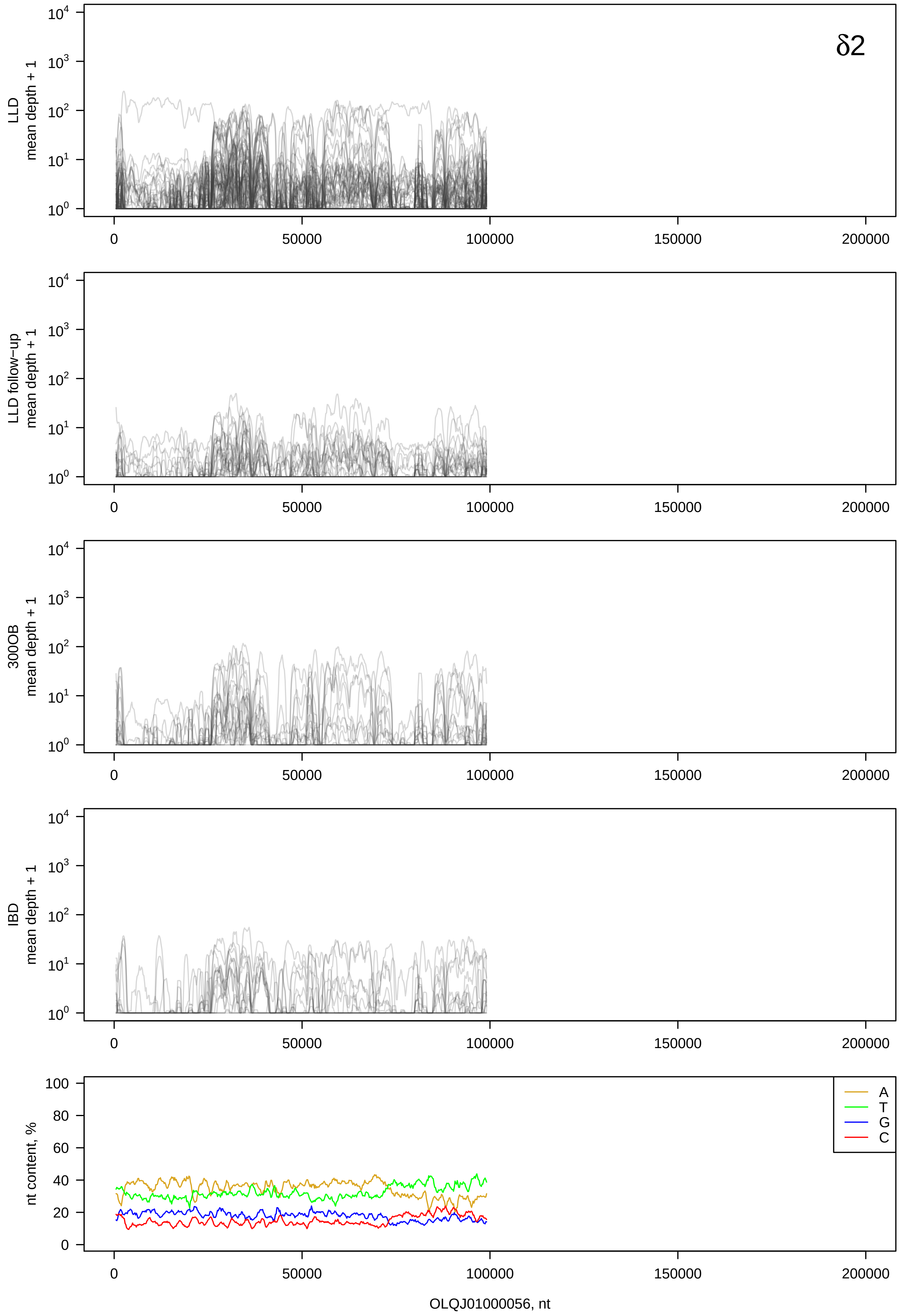

δ2
LLD
mean depth + 1
LLD follow−up
mean depth + 1
300OB
mean depth + 1
IBD
mean depth + 1
nt content, %
A
T
G
C
0
50000
100000
150000
200000
OLQJ01000056, nt

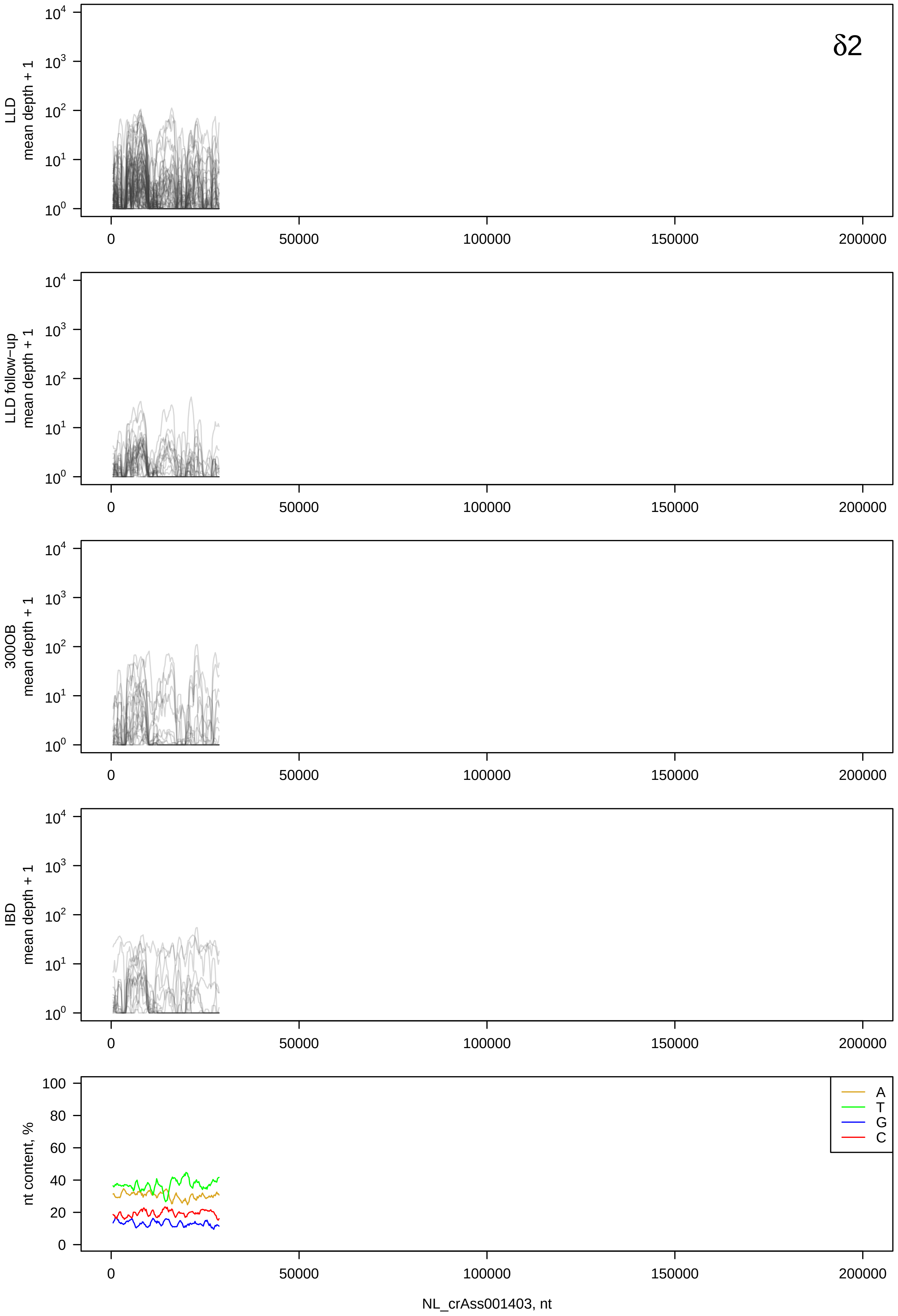
δ2
LLD mean depth + 1
LLD follow-up mean depth + 1
300OB mean depth + 1
IBD mean depth + 1
nt content, %
A
T
G
C
0
50000
100000
150000
200000
NL_crAss001403, nt

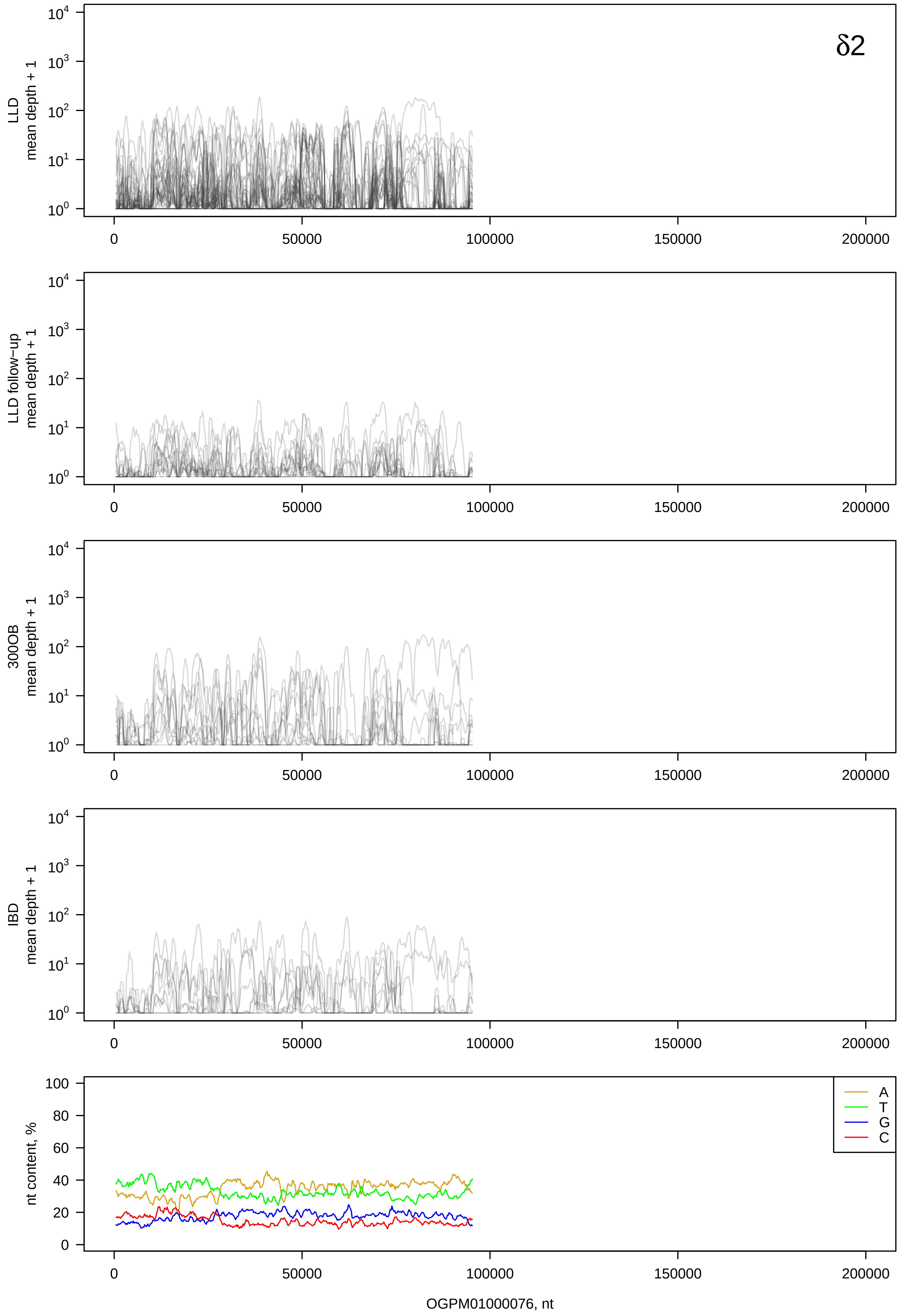

δ2
LLD mean depth + 1
LLD follow−up mean depth + 1
300OB mean depth + 1
IBD mean depth + 1
nt content, %
A
T
G
C
OGPM01000076, nt

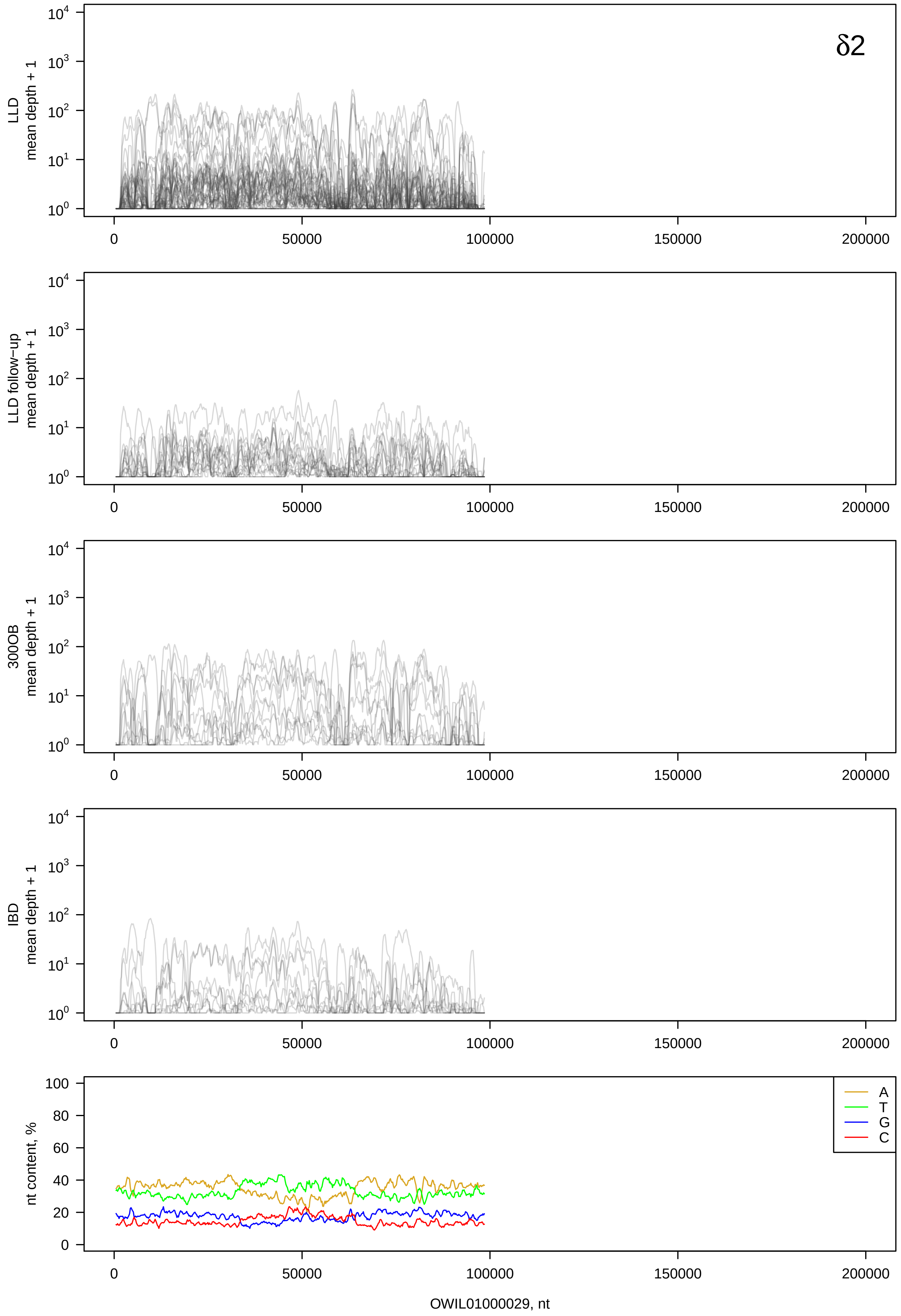

δ2
LLD mean depth + 1
LLD follow−up mean depth + 1
300OB mean depth + 1
IBD mean depth + 1
nt content, %
A
T
G
C
OWIL01000029, nt

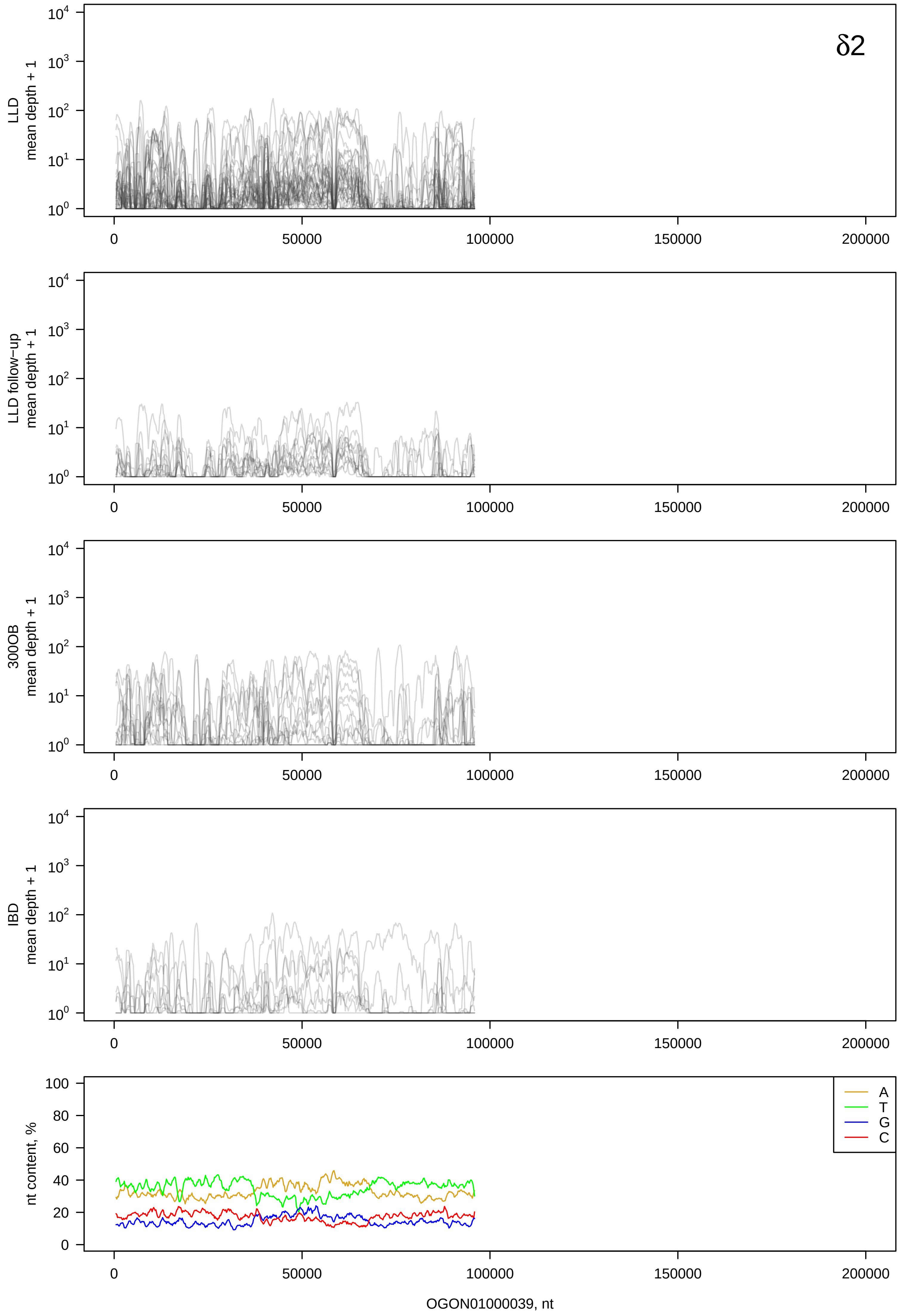

δ2
LLD
mean depth + 1
LLD follow−up
mean depth + 1
300OB
mean depth + 1
IBD
mean depth + 1
nt content, %
A
T
G
C
0
50000
100000
150000
200000
OGON01000039, nt

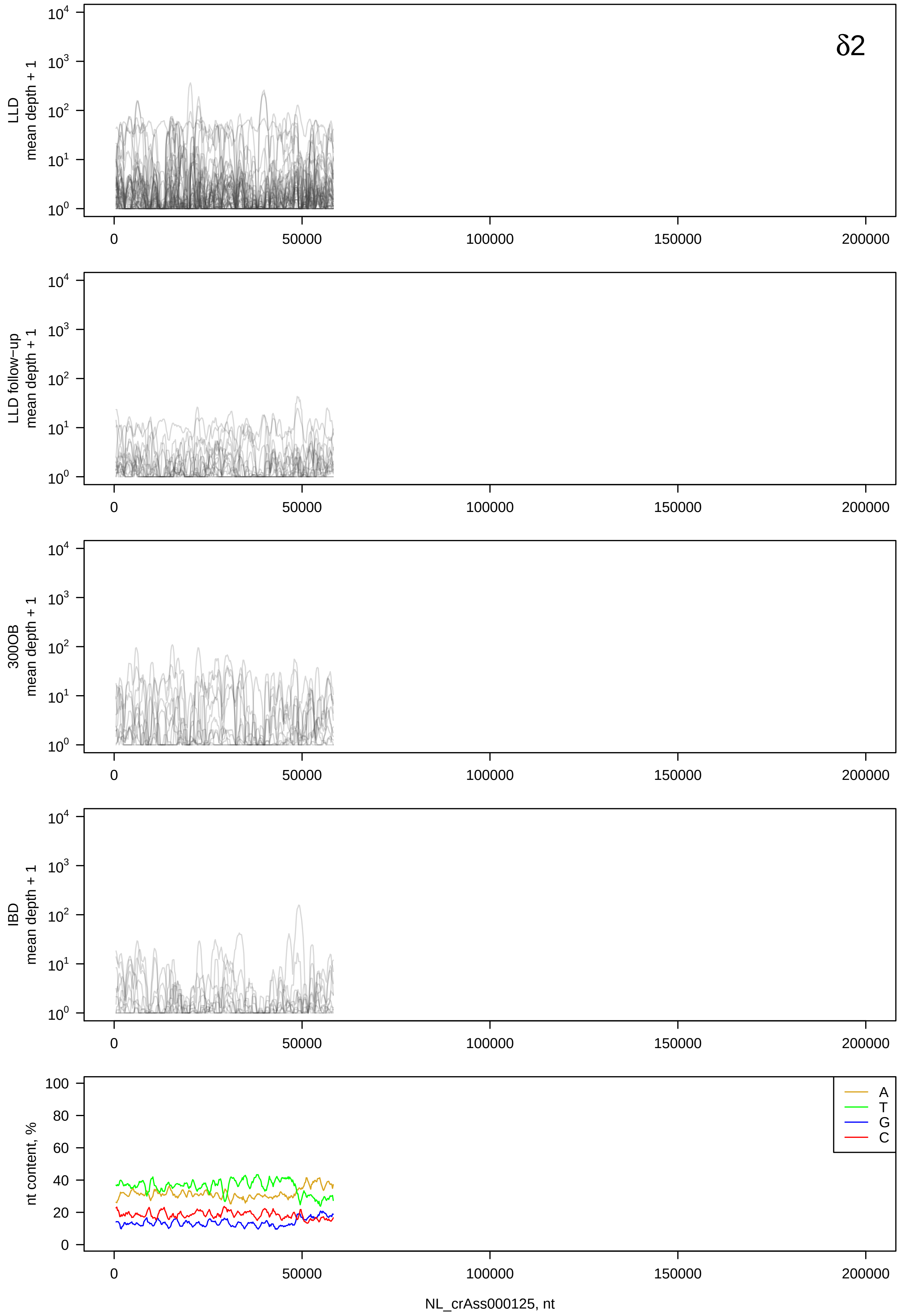
δ2
LLD
mean depth + 1
LLD follow-up
mean depth + 1
300OB
mean depth + 1
IBD
mean depth + 1
nt content, %
A
T
G
C
0
50000
100000
150000
200000
NL_crAss000125, nt

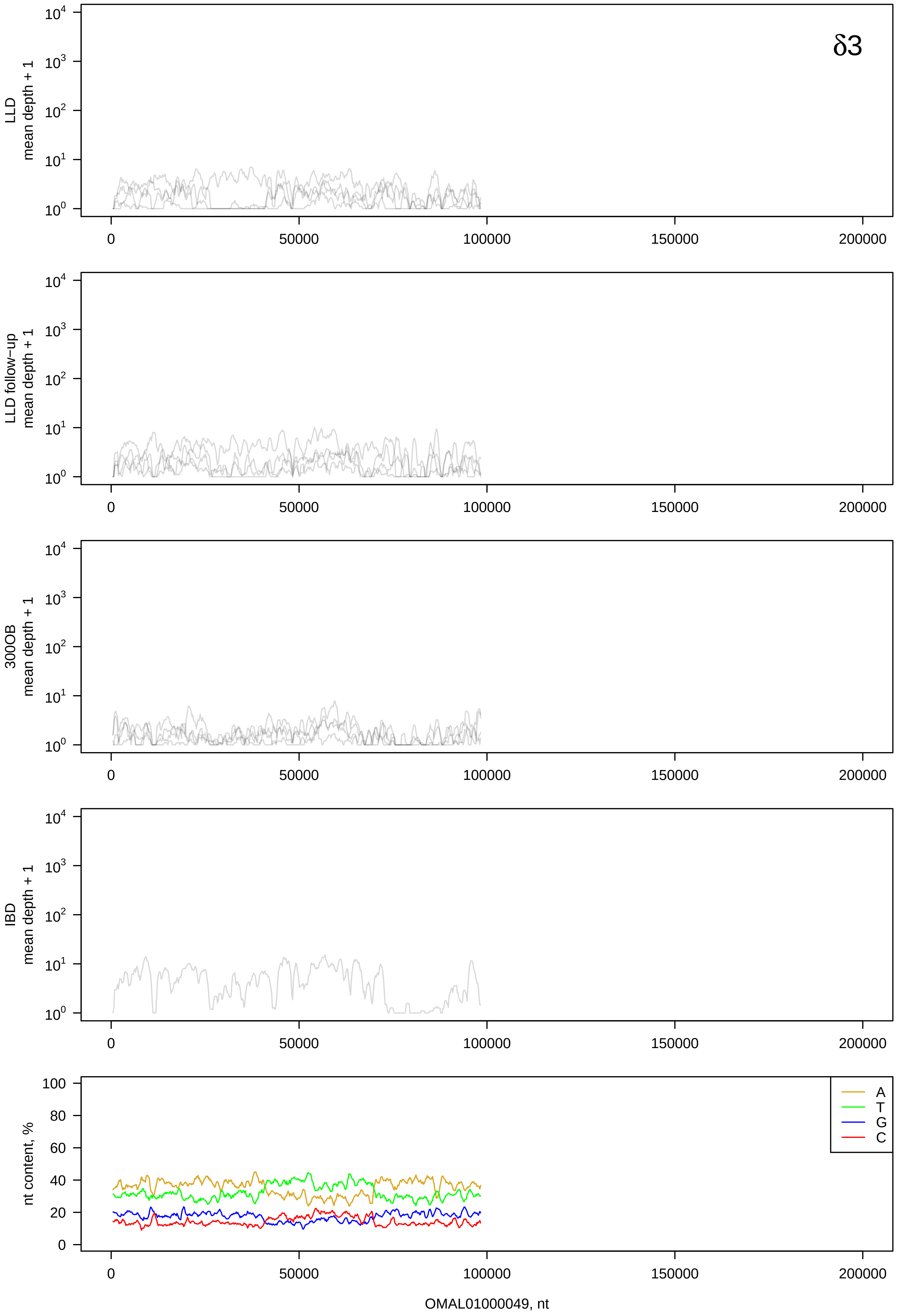
δ3
LLD
mean depth + 1
LLD follow−up
mean depth + 1
300OB
mean depth + 1
IBD
mean depth + 1
nt content, %
A
T
G
C
0
50000
100000
150000
200000
OMAL01000049, nt

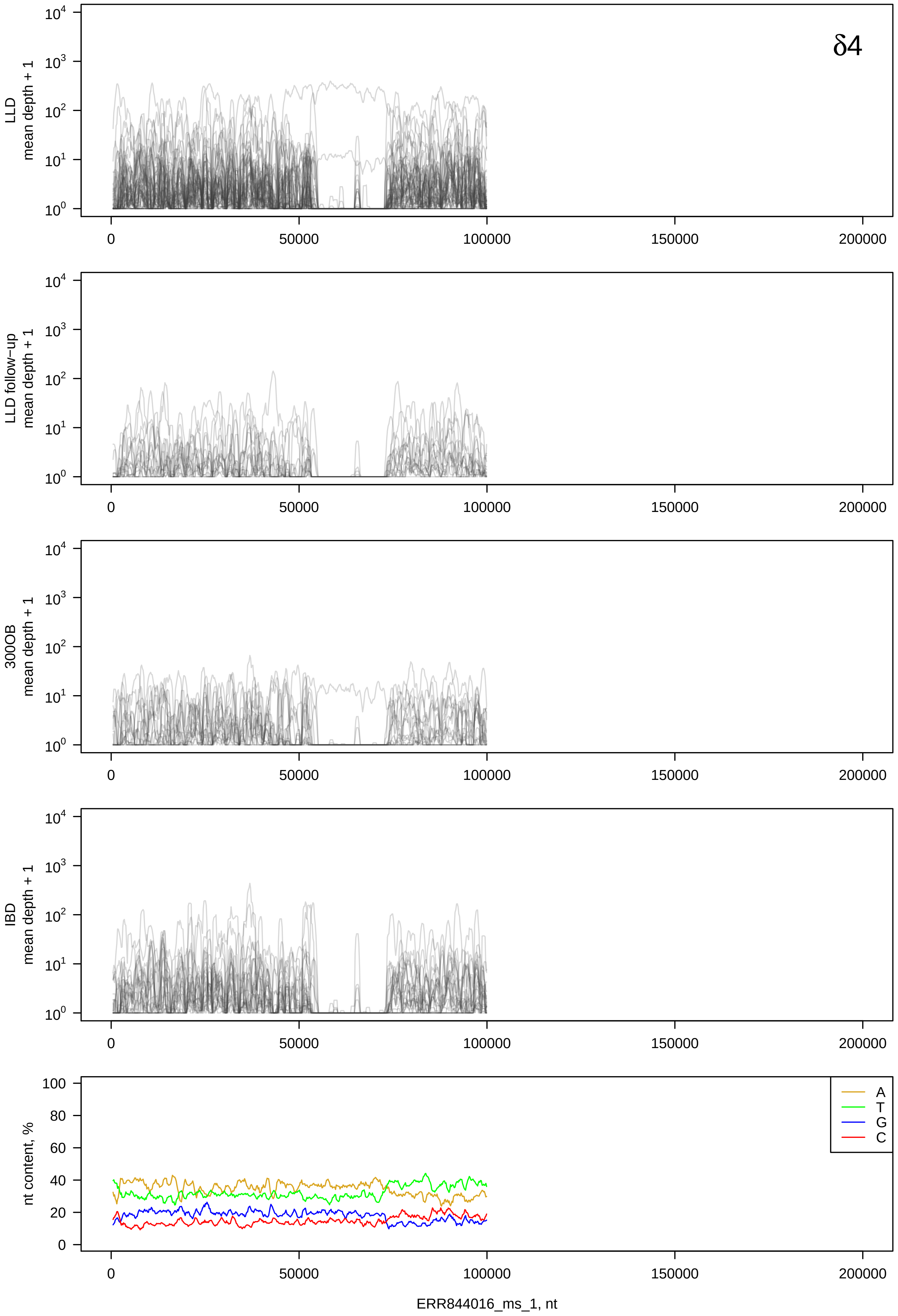
δ4
LLD mean depth + 1
LLD follow−up mean depth + 1
300OB mean depth + 1
IBD mean depth + 1
nt content, %
A
T
G
C
ERR844016_ms_1, nt

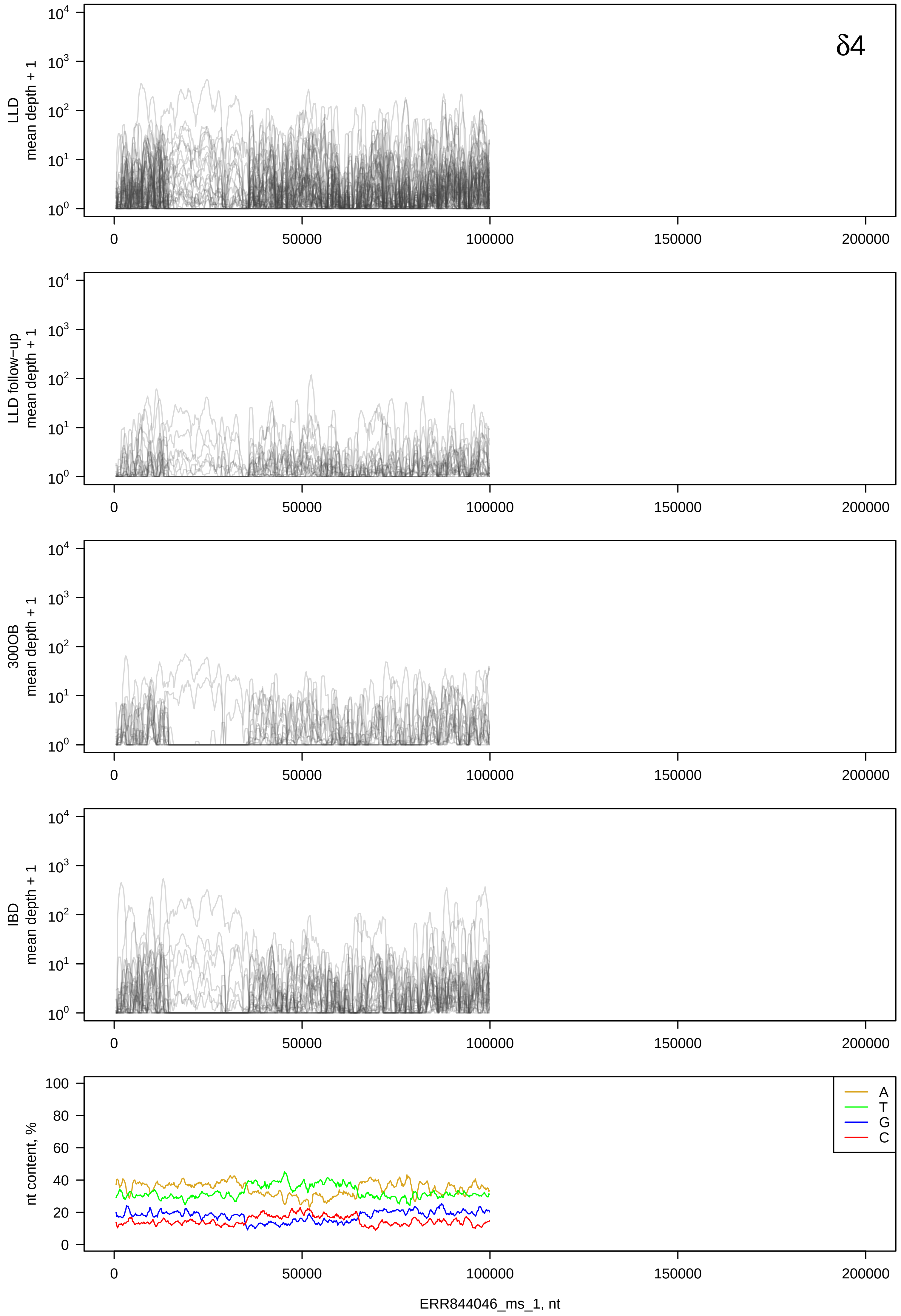
δ4
LLD mean depth + 1
LLD follow-up mean depth + 1
300OB mean depth + 1
IBD mean depth + 1
nt content, %
A
T
G
C
ERR844046_ms_1, nt

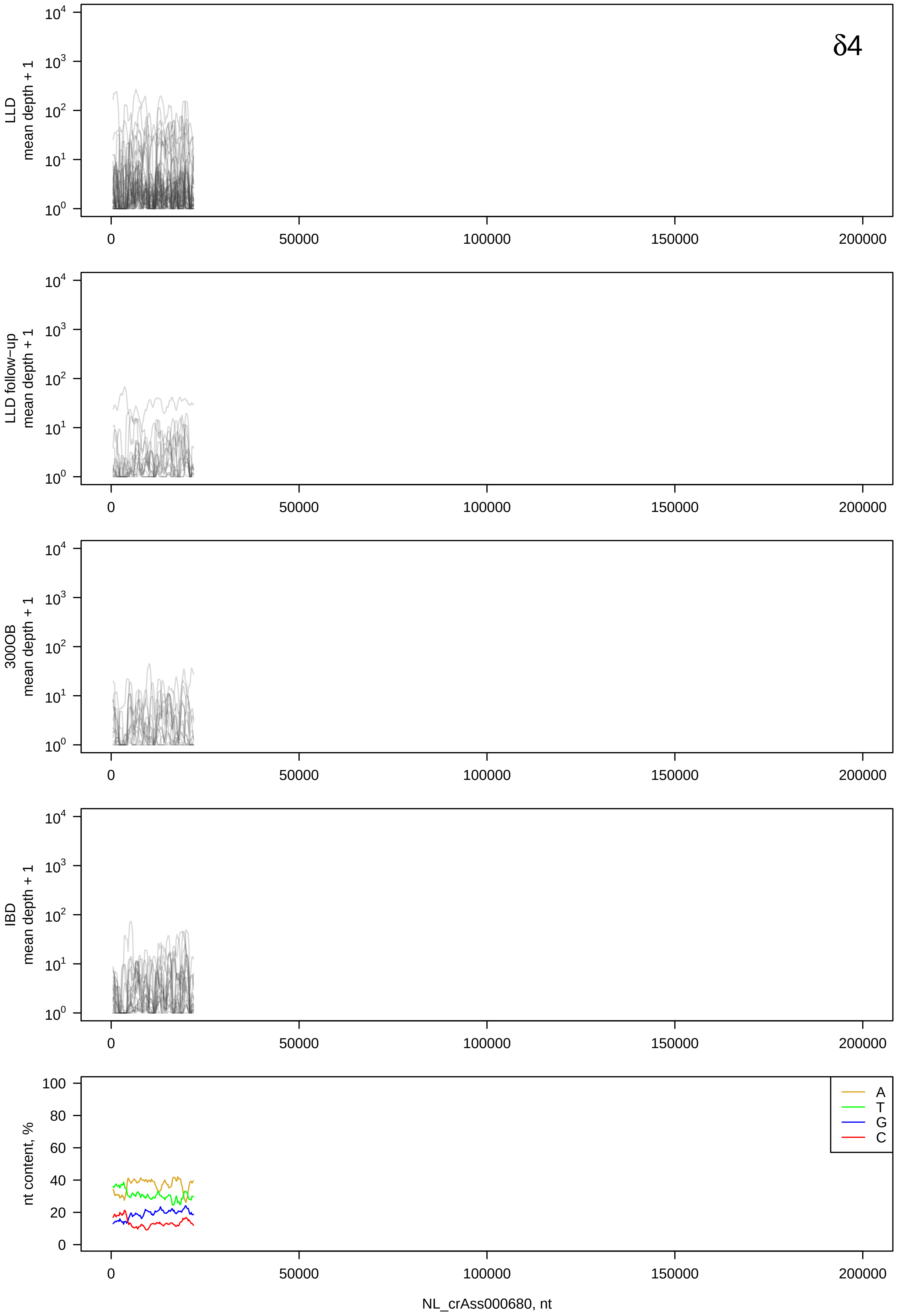

δ4
LLD mean depth + 1
LLD follow-up mean depth + 1
300OB mean depth + 1
IBD mean depth + 1
nt content, %
A
T
G
C
0
50000
100000
150000
200000
NL_crAss000680, nt

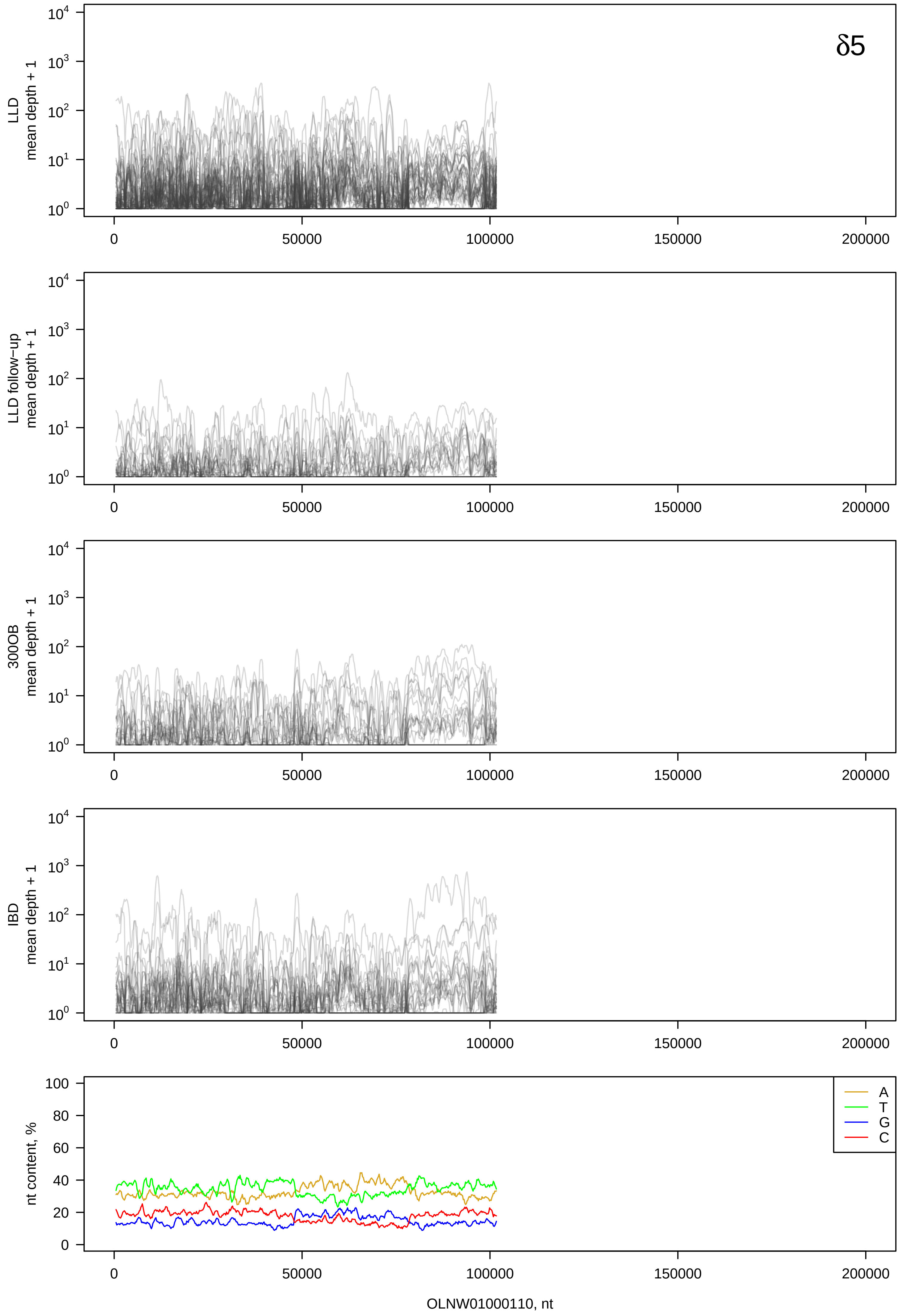
δ5
LLD mean depth + 1
LLD follow-up mean depth + 1
300OB mean depth + 1
IBD mean depth + 1
nt content, %
A
T
G
C
OLNW01000110, nt

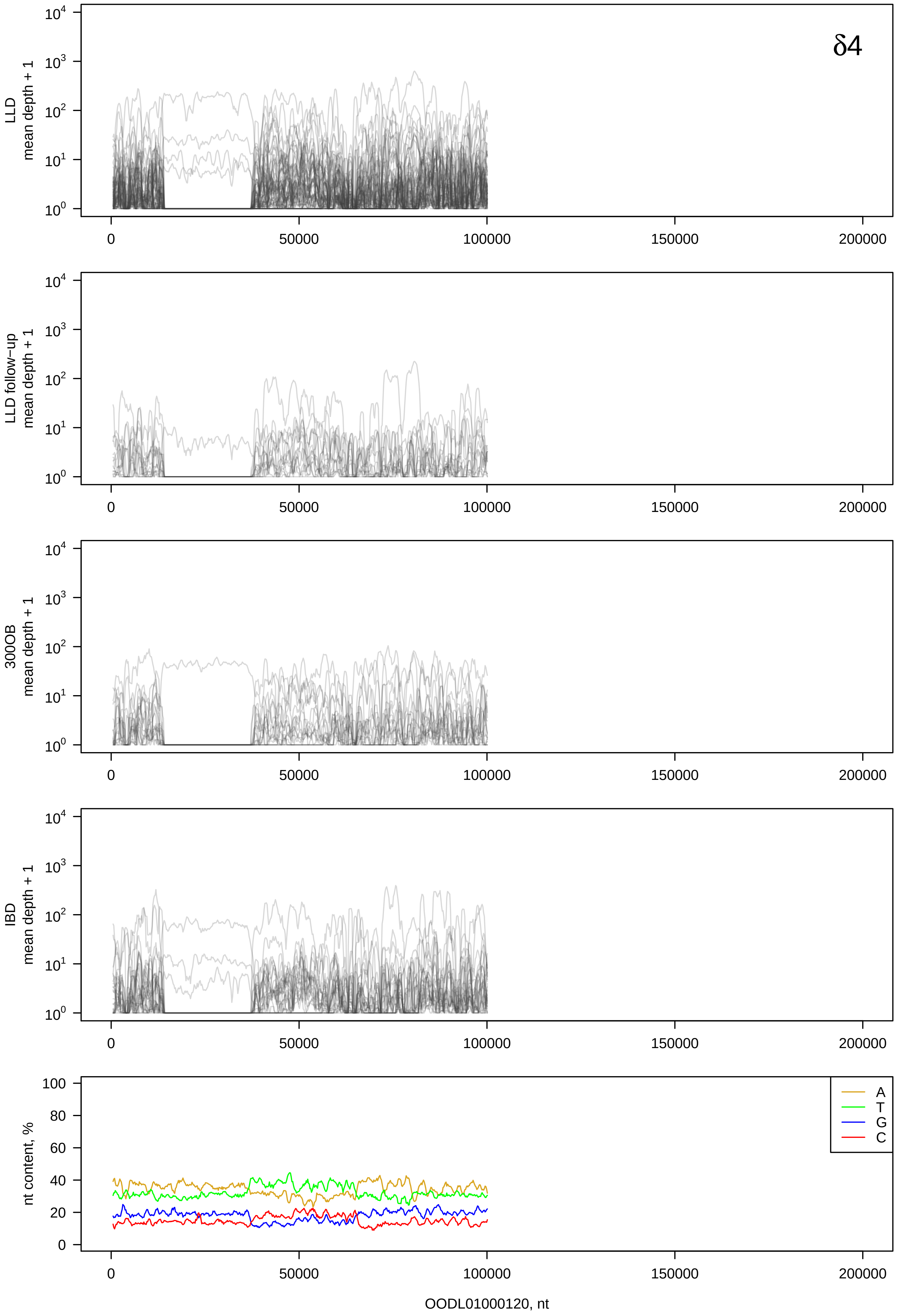

δ4
LLD mean depth + 1
LLD follow-up mean depth + 1
300OB mean depth + 1
IBD mean depth + 1
nt content, %
A
T
G
C
OODL01000120, nt

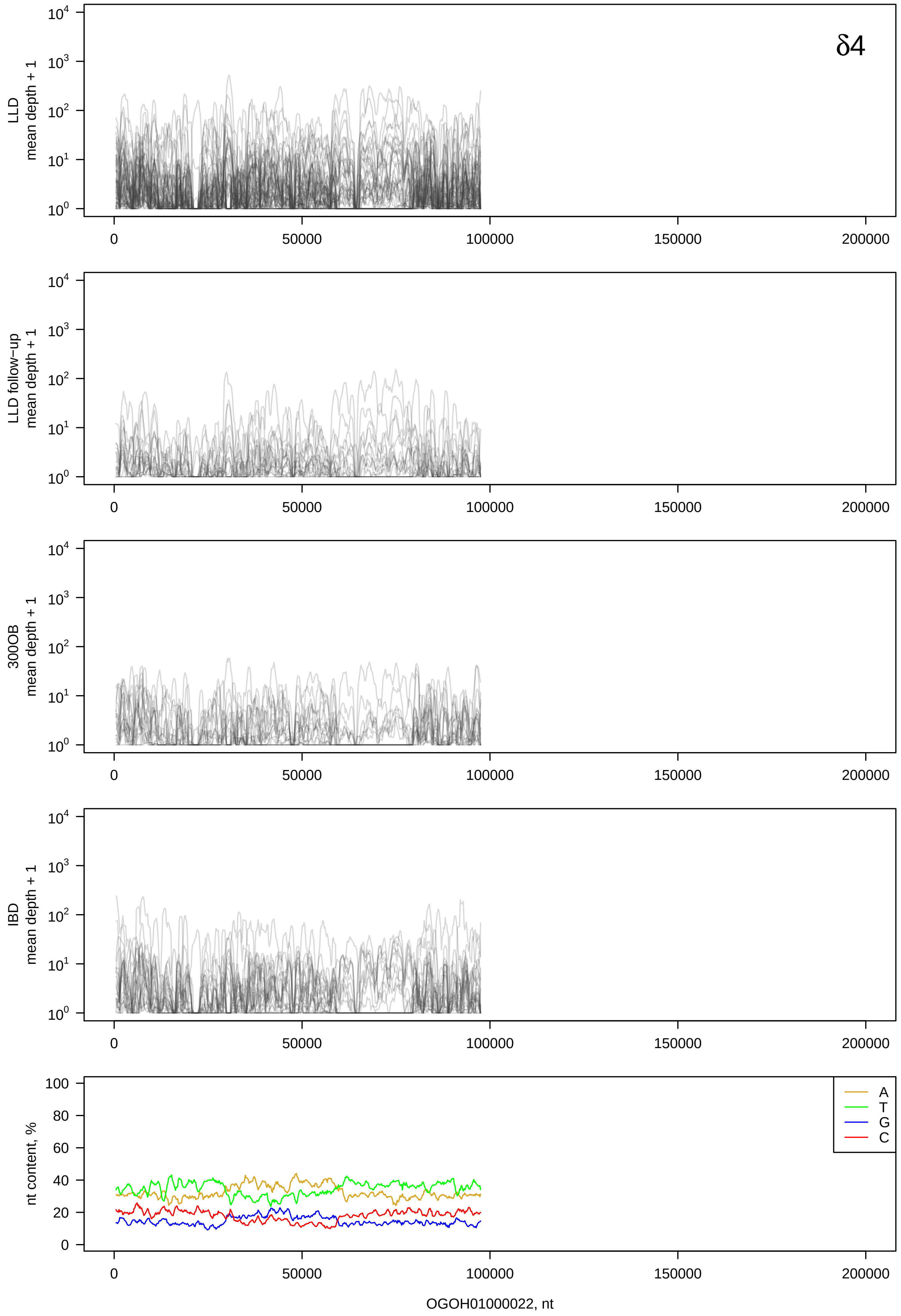

δ4
LLD mean depth + 1
LLD follow−up mean depth + 1
300OB mean depth + 1
IBD mean depth + 1
nt content, %
A
T
G
C
OGOH01000022, nt

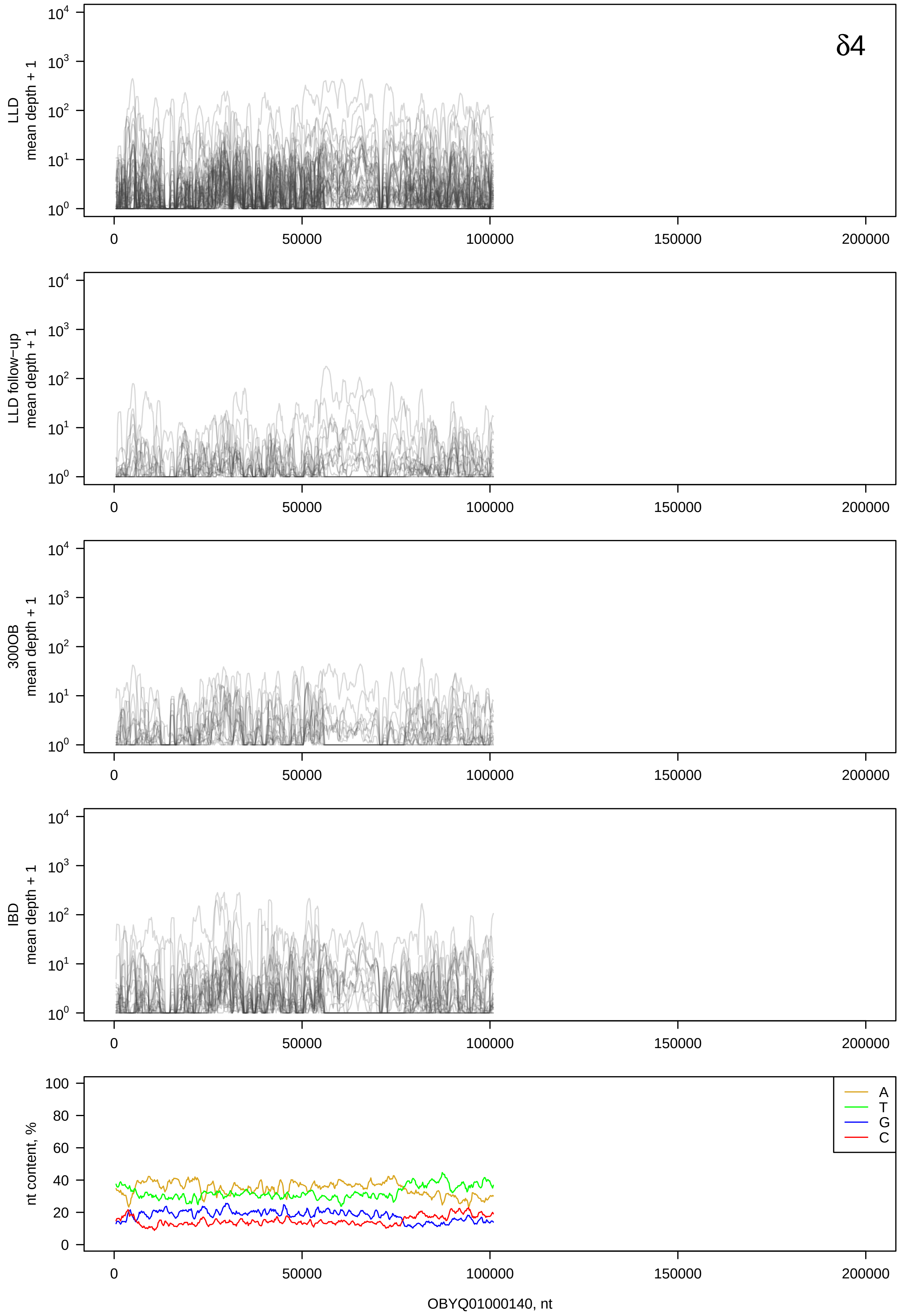
δ4
LLD
mean depth + 1
LLD follow−up
mean depth + 1
300OB
mean depth + 1
IBD
mean depth + 1
nt content, %
A
T
G
C
0
50000
100000
150000
200000
OBYQ01000140, nt

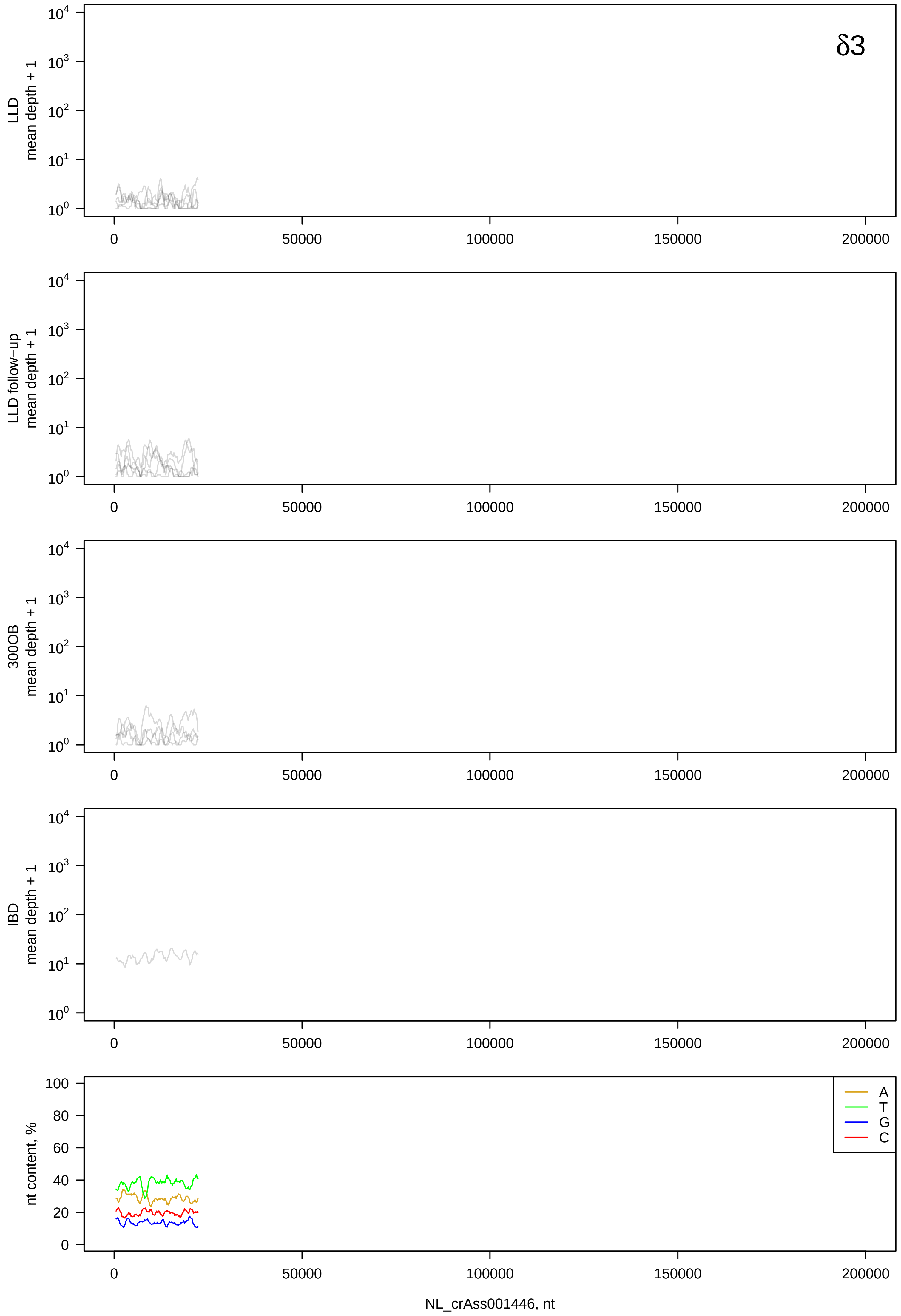

δ3
LLD mean depth + 1
LLD follow-up mean depth + 1
300OB mean depth + 1
IBD mean depth + 1
nt content, %
A
T
G
C
0
50000
100000
150000
200000
NL_crAss001446, nt

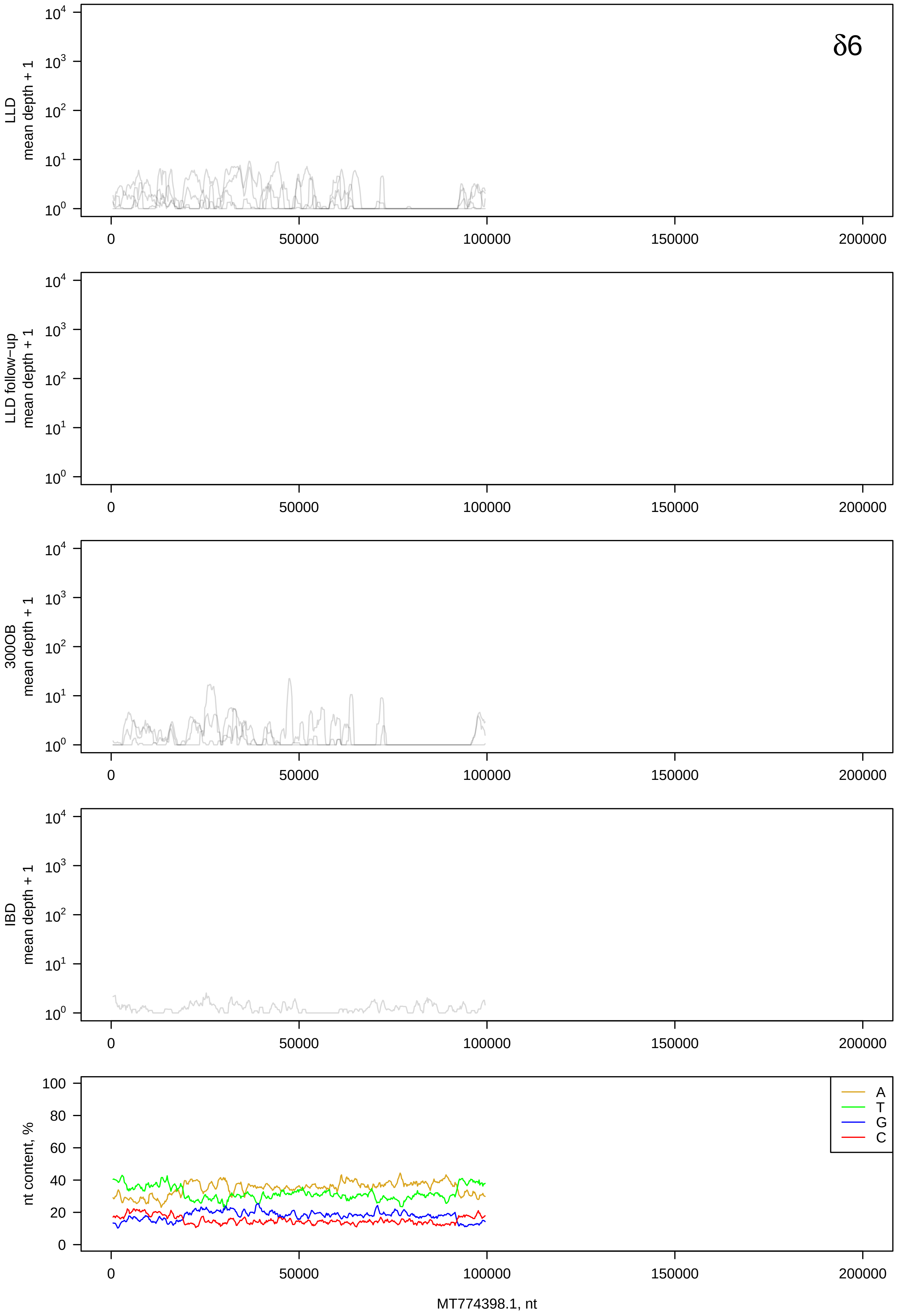

δ6
LLD mean depth + 1
LLD follow-up mean depth + 1
300OB mean depth + 1
IBD mean depth + 1
nt content, %
A
T
G
C
MT774398.1, nt

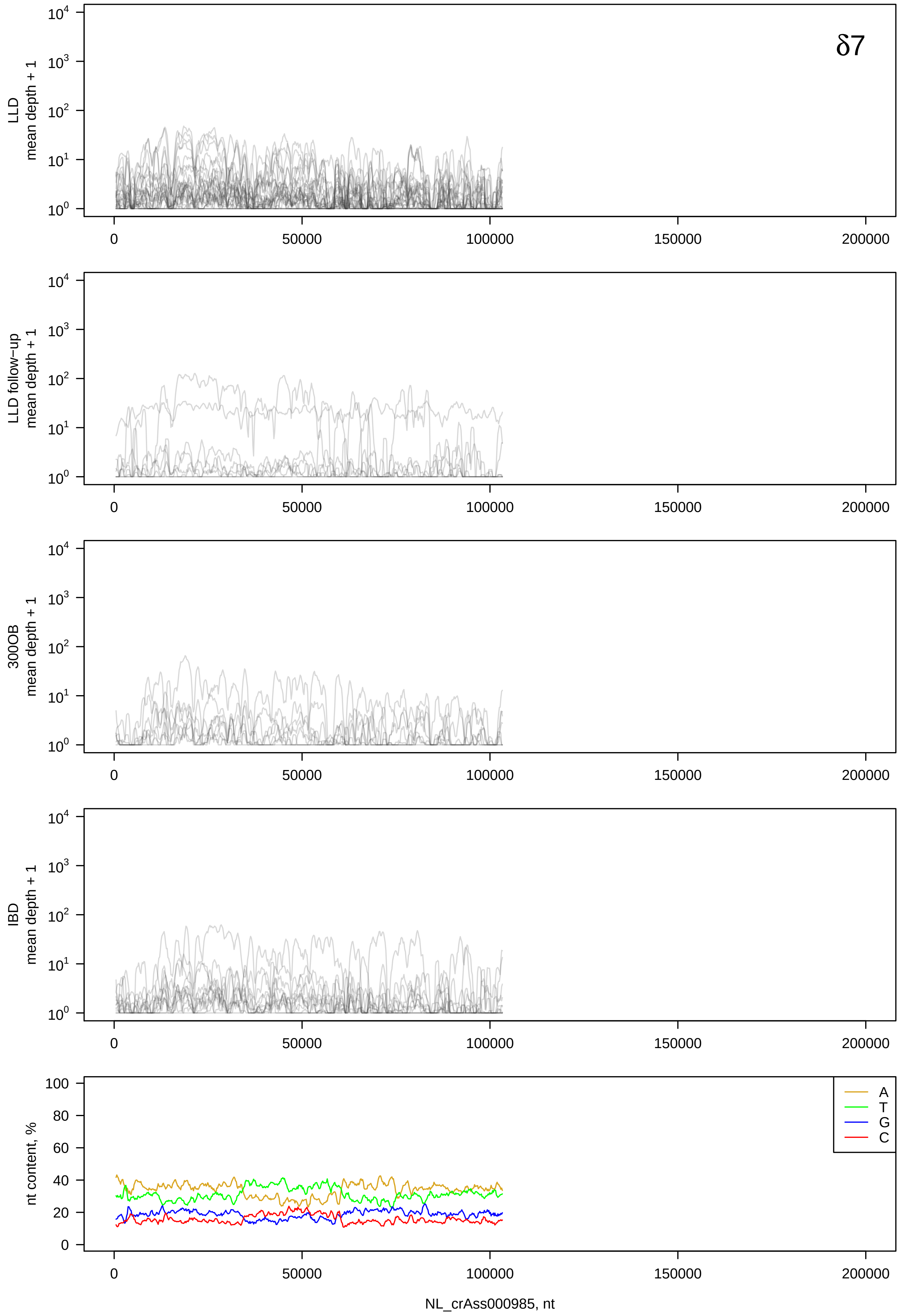
δ7
LLD mean depth + 1
LLD follow−up mean depth + 1
300OB mean depth + 1
IBD mean depth + 1
nt content, %
A
T
G
C
NL_crAss000985, nt

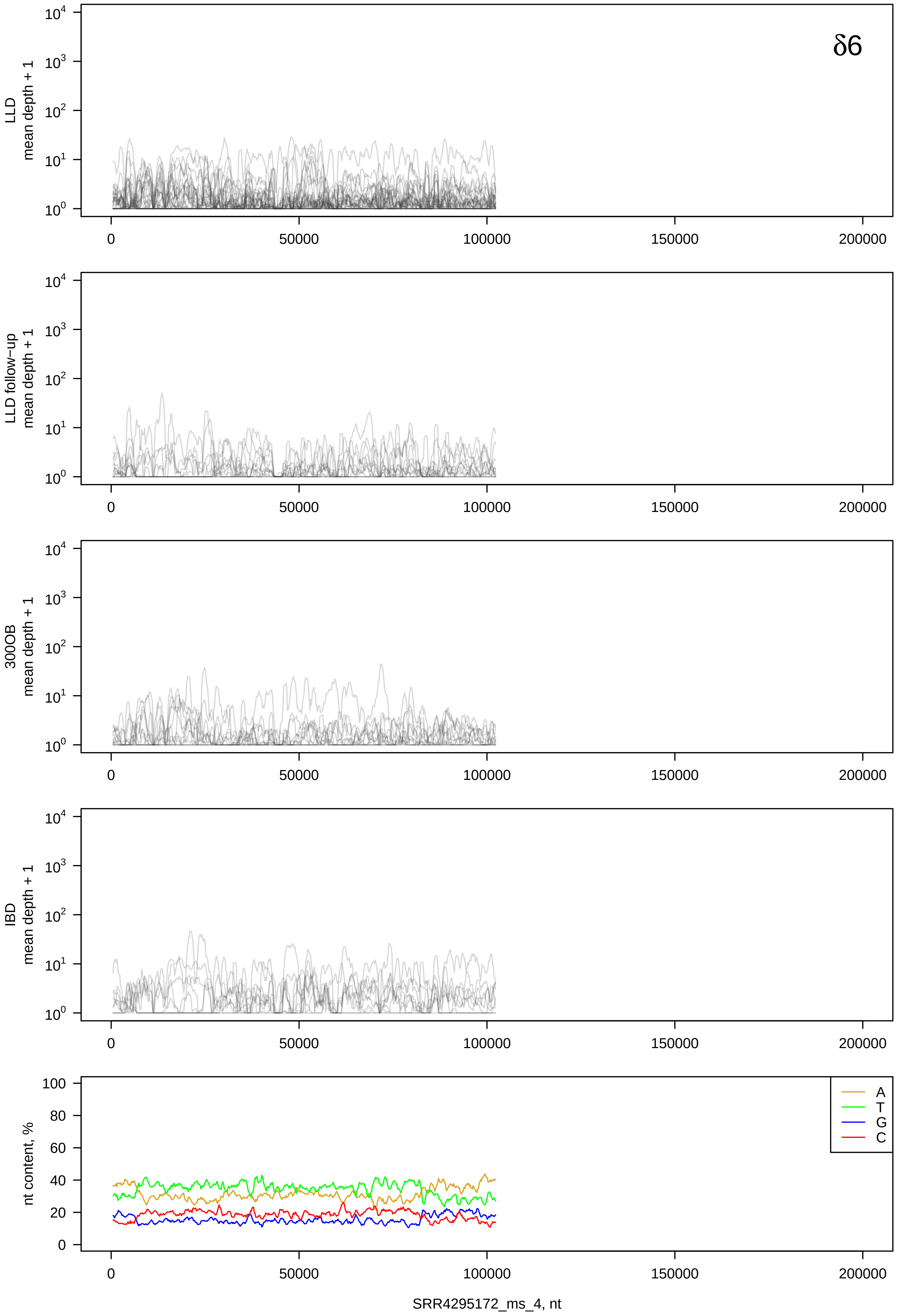

δ6
LLD
mean depth + 1
LLD follow−up
mean depth + 1
300OB
mean depth + 1
IBD
mean depth + 1
nt content, %
A
T
G
C
SRR4295172_ms_4, nt

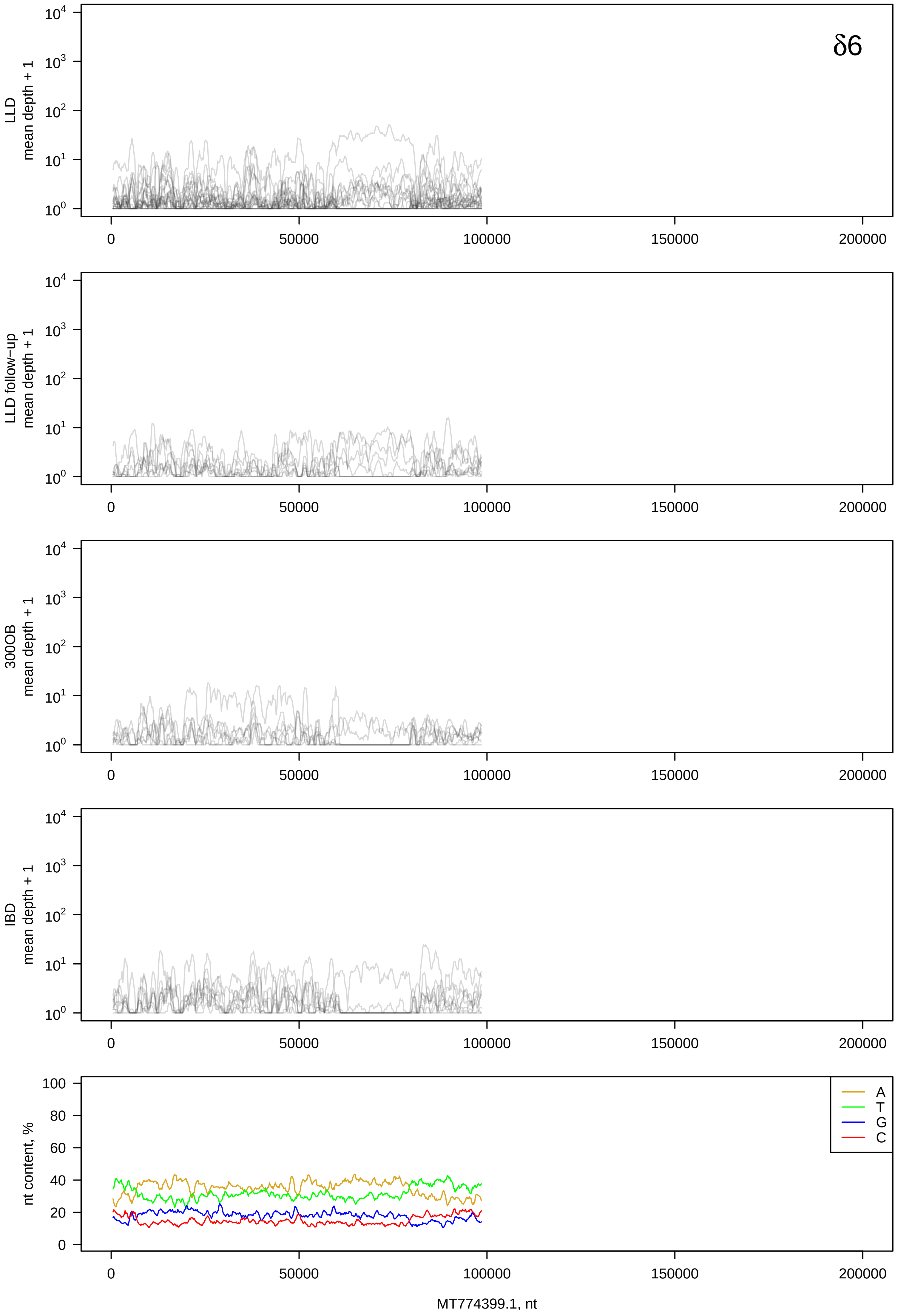

δ6
LLD
mean depth + 1
LLD follow−up
mean depth + 1
300OB
mean depth + 1
IBD
mean depth + 1
nt content, %
A
T
G
C
0
50000
100000
150000
200000
MT774399.1, nt

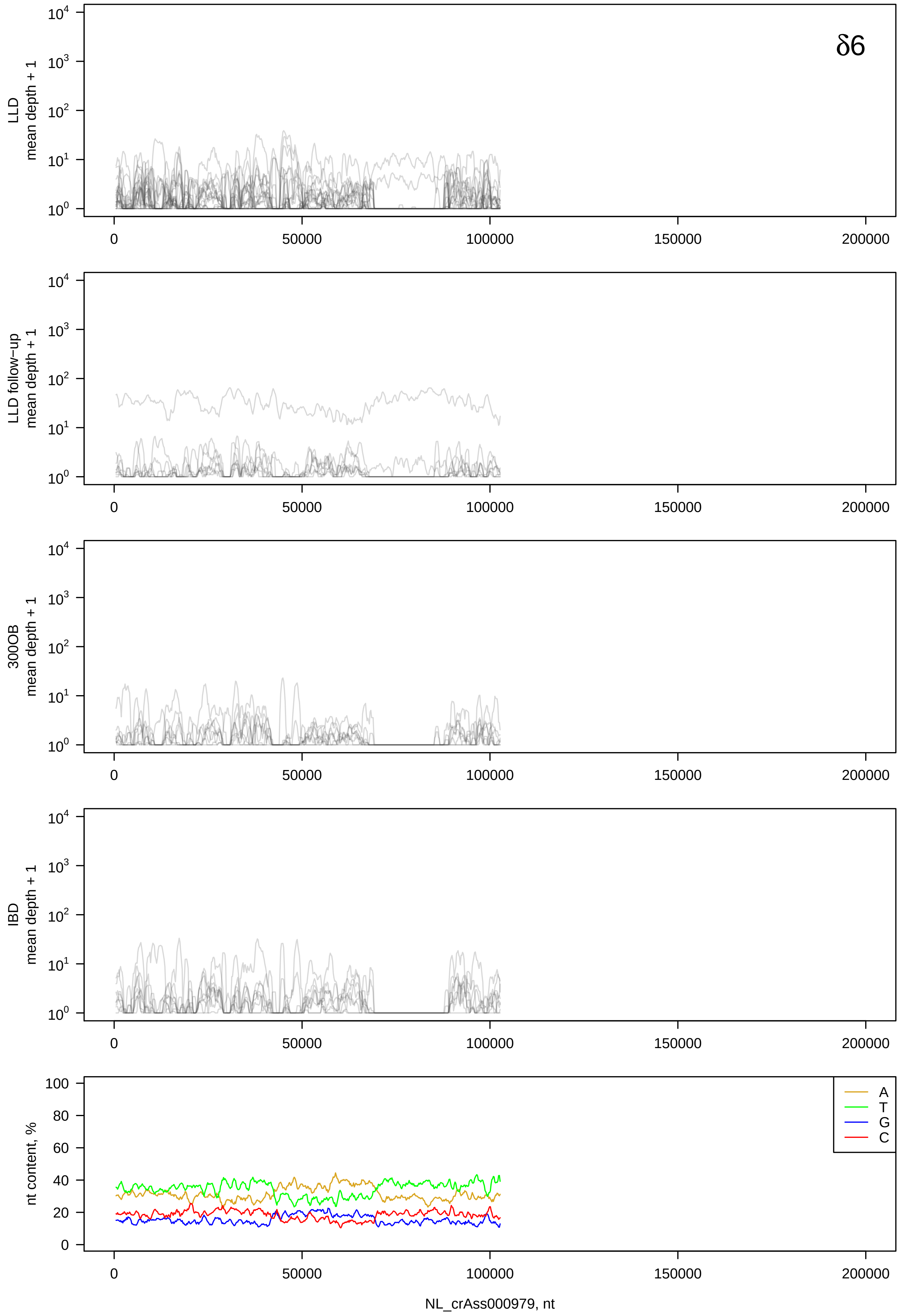

δ6
LLD
mean depth + 1
LLD follow-up
mean depth + 1
300OB
mean depth + 1
IBD
mean depth + 1
nt content, %
A
T
G
C
NL_crAss000979, nt

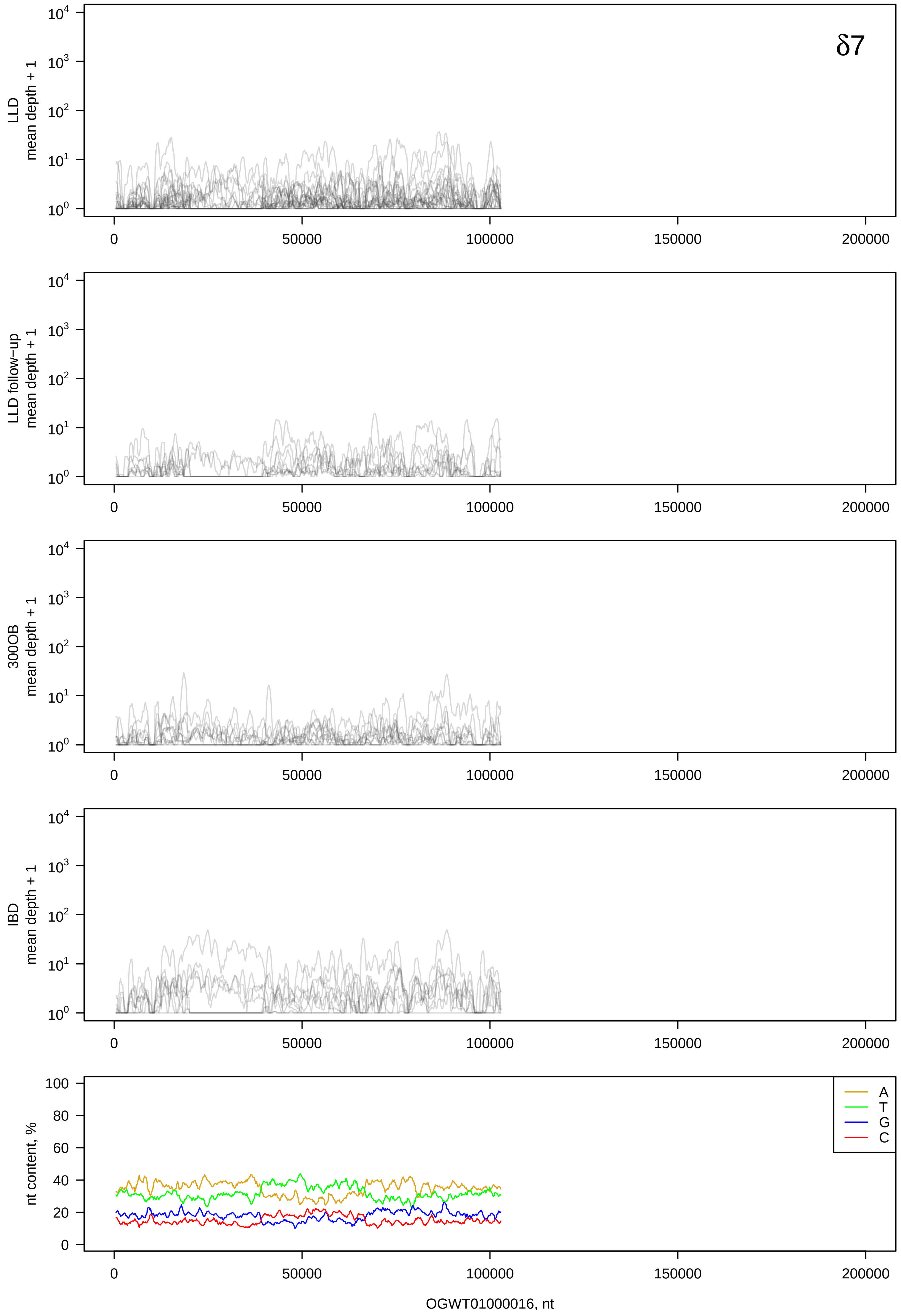
δ7
LLD mean depth + 1
LLD follow−up mean depth + 1
300OB mean depth + 1
IBD mean depth + 1
nt content, %
A
T
G
C
OGWT01000016, nt

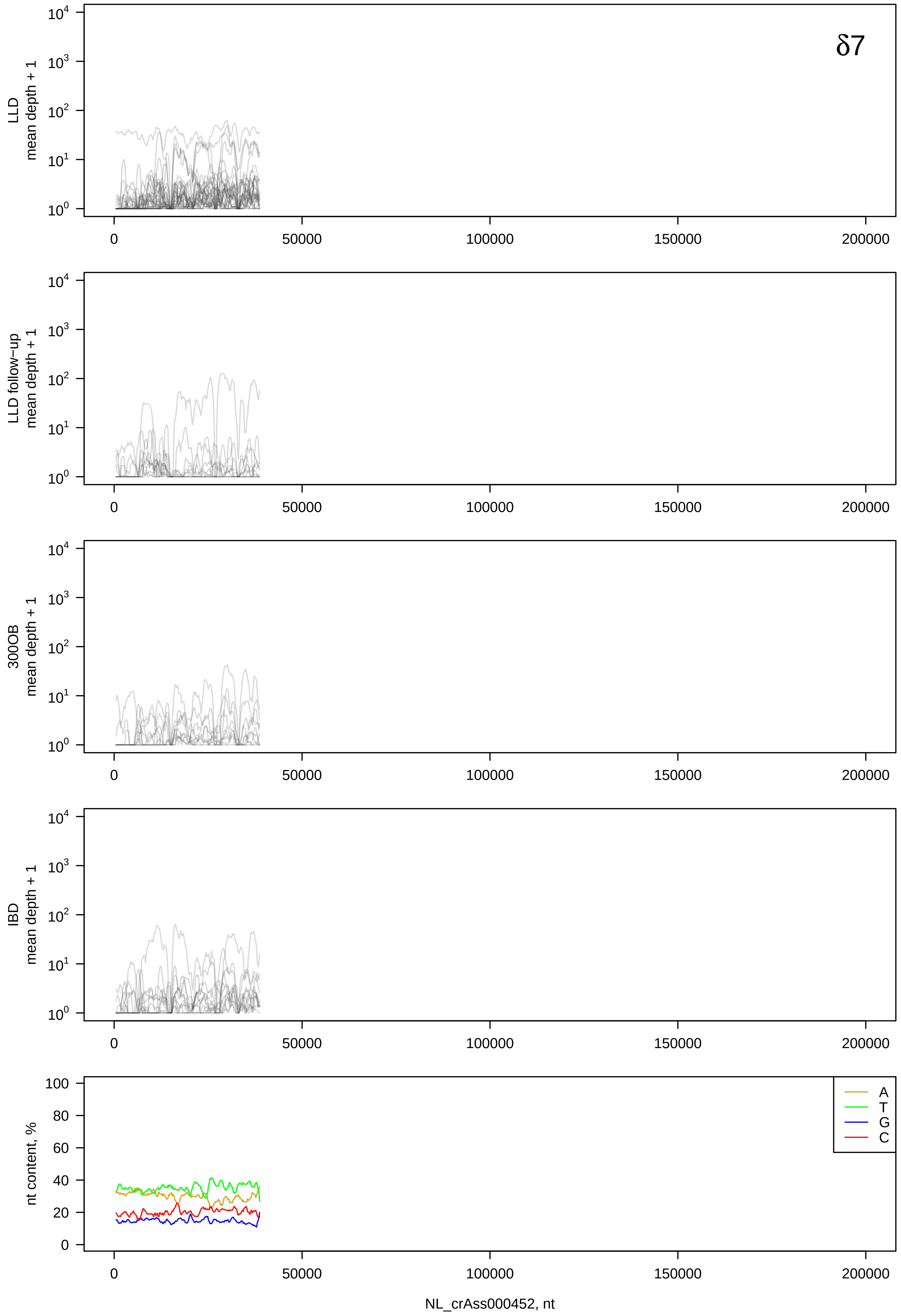

δ7
LLD
mean depth + 1
LLD follow−up
mean depth + 1
300OB
mean depth + 1
IBD
mean depth + 1
nt content, %
A
T
G
C
NL_crAss000452, nt

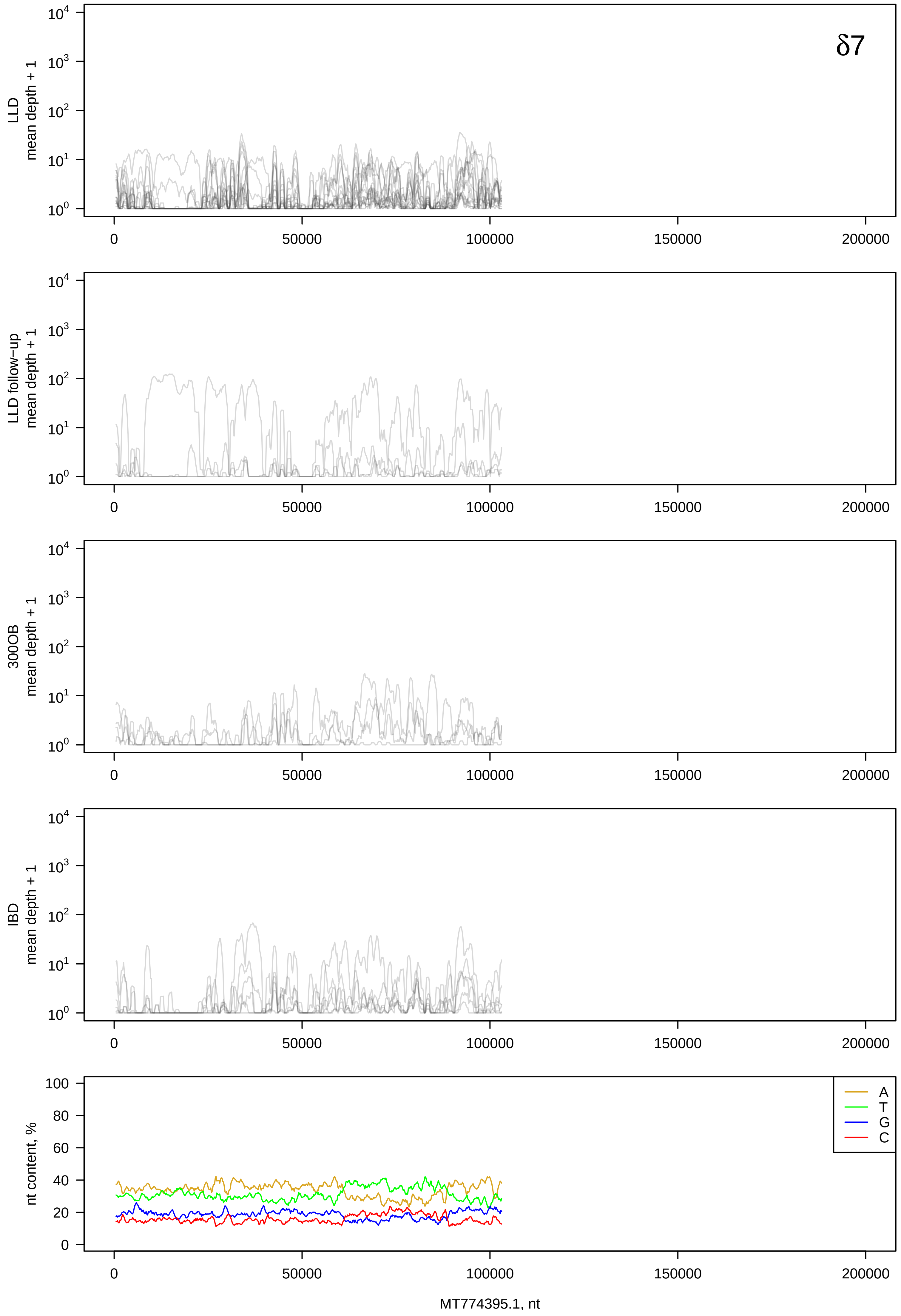

δ7
LLD
mean depth + 1
LLD follow−up
mean depth + 1
300OB
mean depth + 1
IBD
mean depth + 1
nt content, %
A
T
G
C
0
50000
100000
150000
200000
MT774395.1, nt

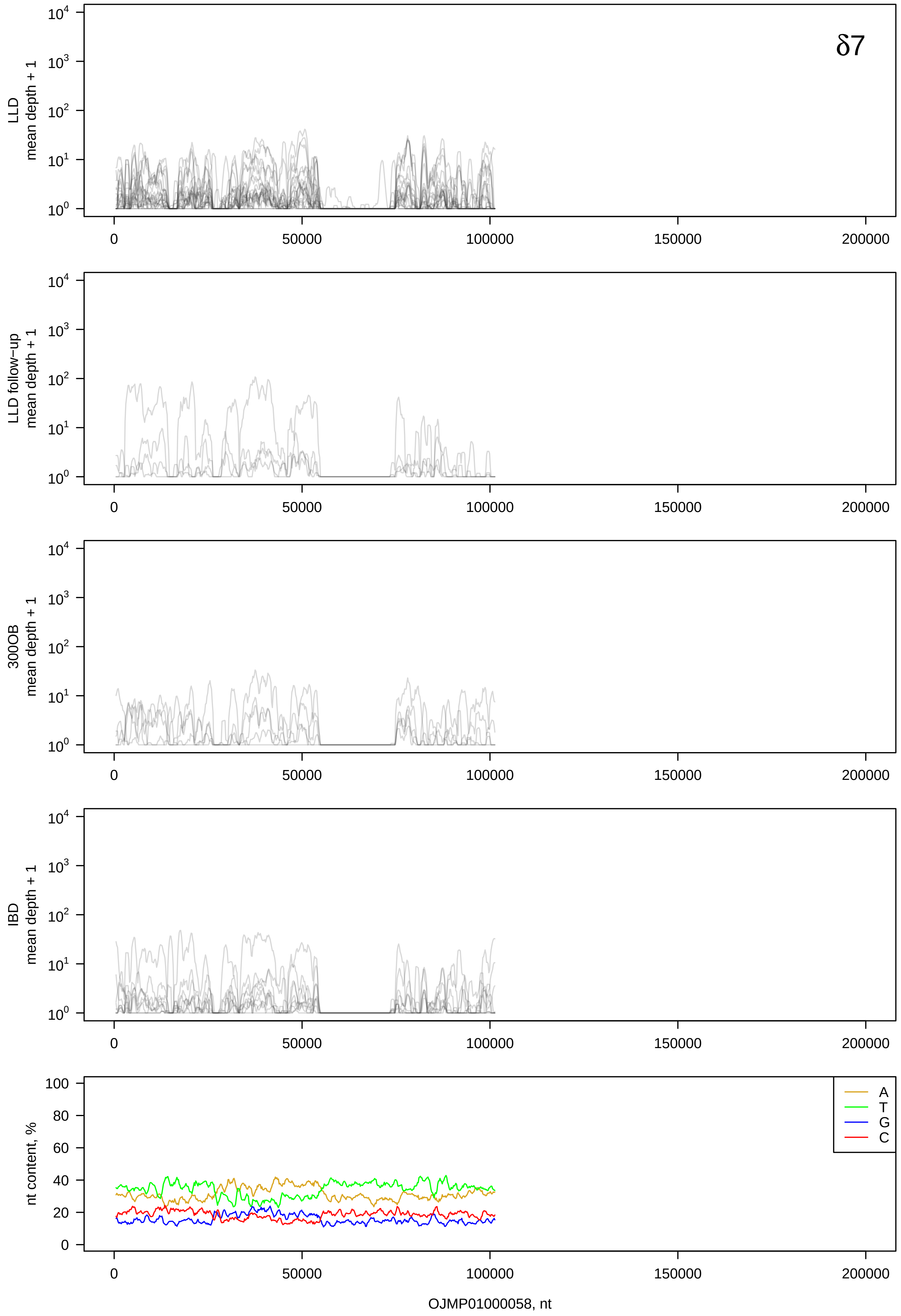
δ7
LLD
mean depth + 1
LLD follow−up
mean depth + 1
300OB
mean depth + 1
IBD
mean depth + 1
nt content, %
A
T
G
C
OJMP01000058, nt

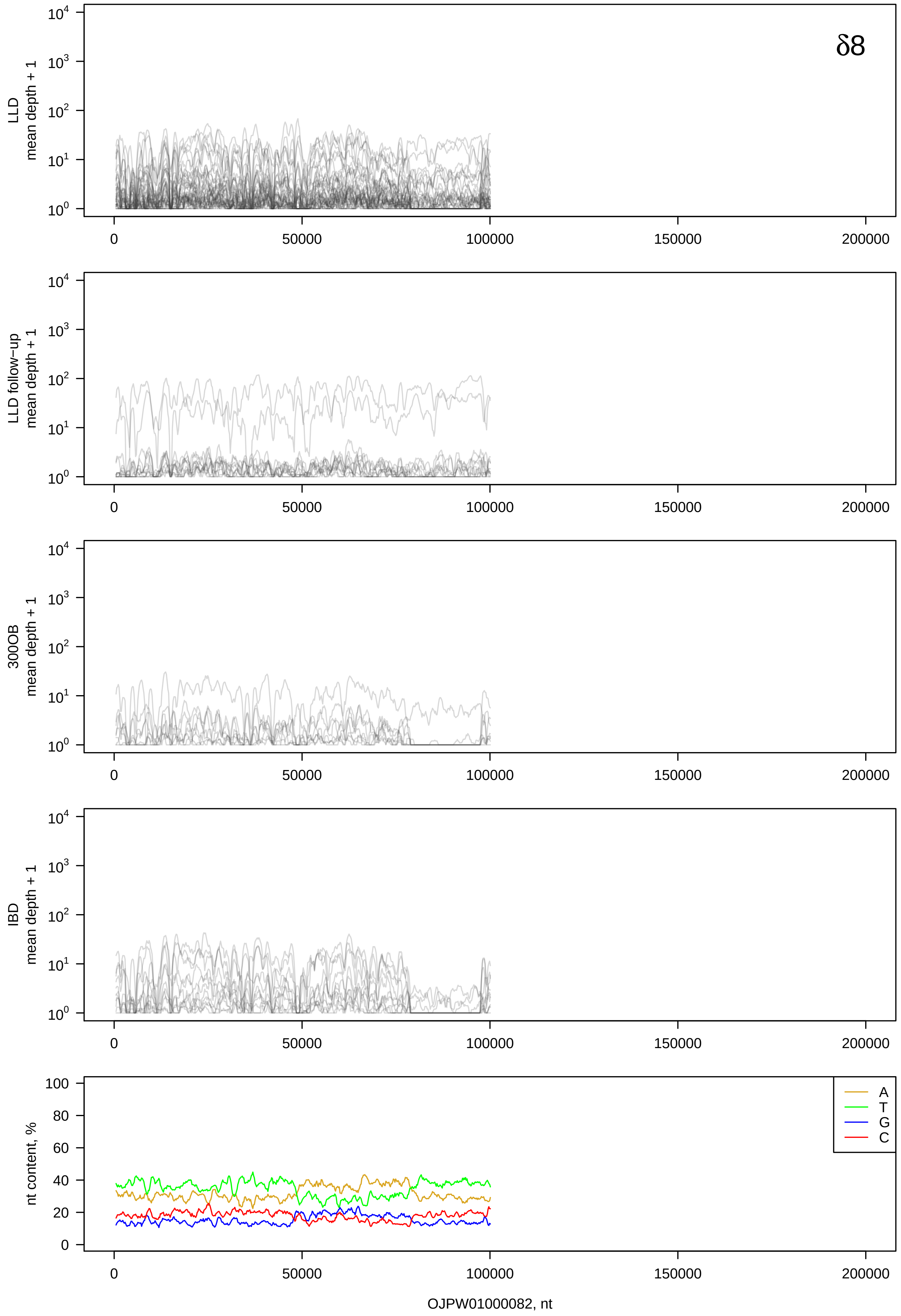
δ8
LLD
mean depth + 1
LLD follow-up
mean depth + 1
300OB
mean depth + 1
IBD
mean depth + 1
nt content, %
A
T
G
C
OJPW01000082, nt

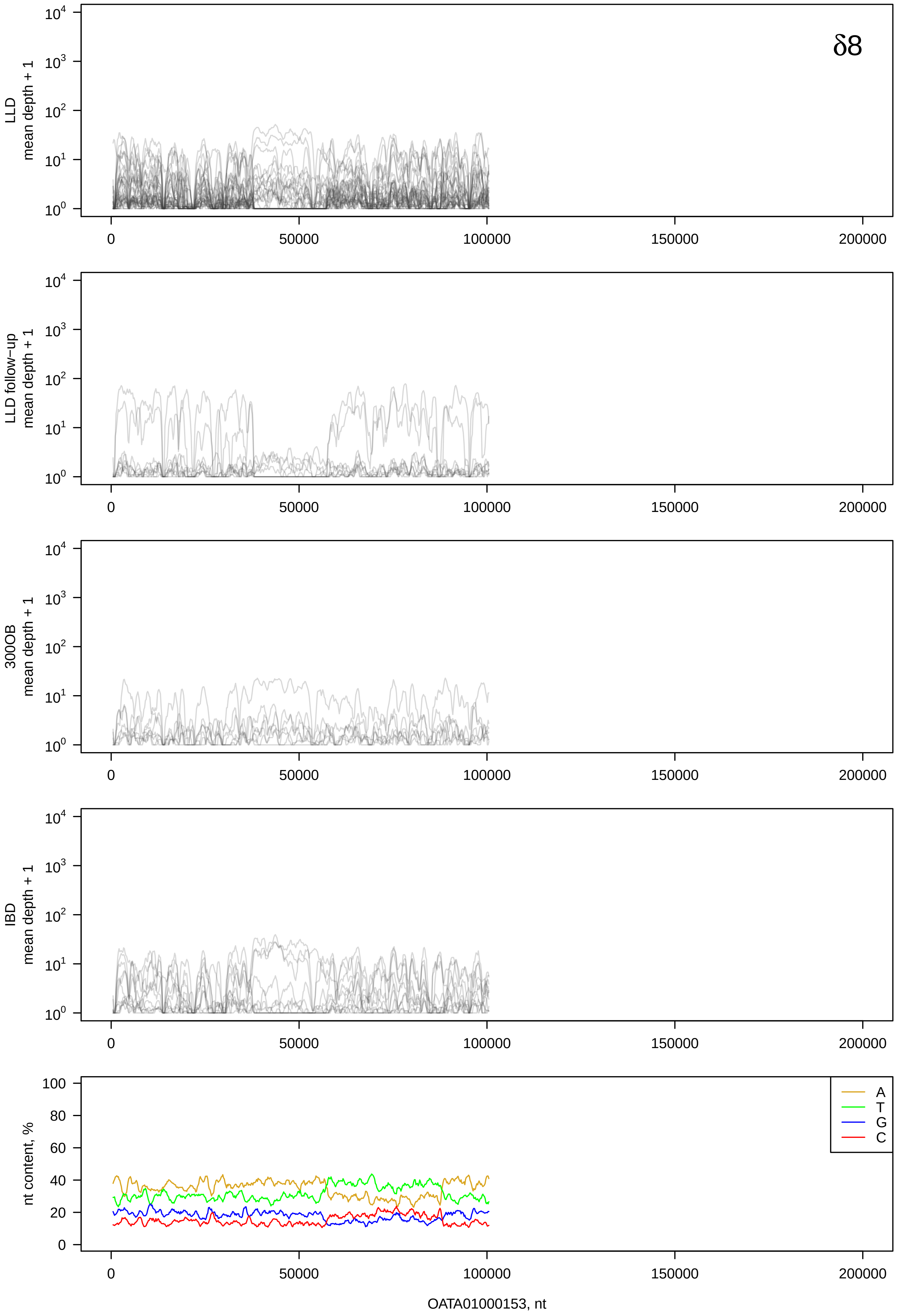
δ8
LLD mean depth + 1
LLD follow−up mean depth + 1
300OB mean depth + 1
IBD mean depth + 1
nt content, %
A
T
G
C
OATA01000153, nt

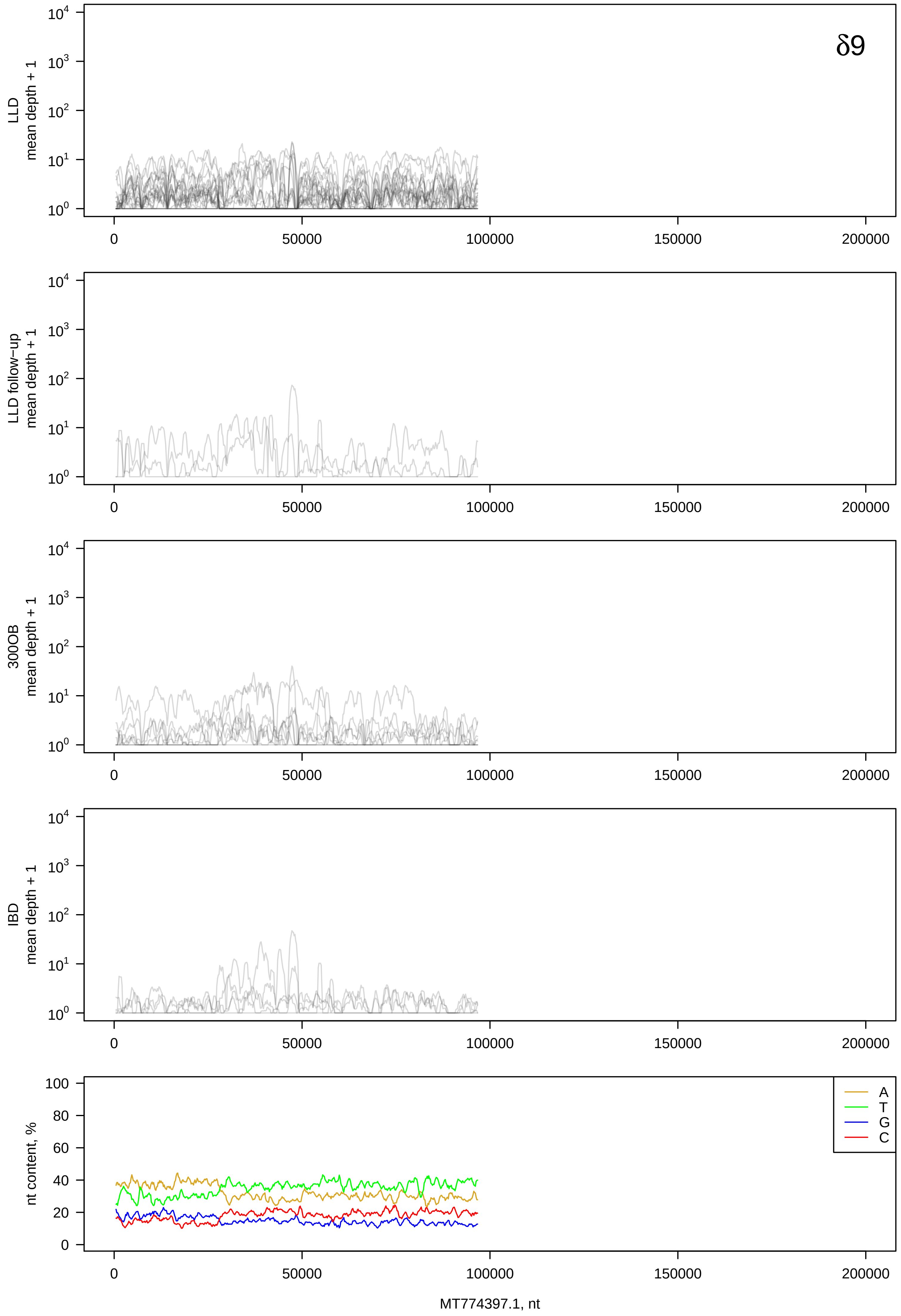

δ9
LLD mean depth + 1
LLD follow−up mean depth + 1
300OB mean depth + 1
IBD mean depth + 1
nt content, %
A
T
G
C
0
50000
100000
150000
200000
MT774397.1, nt

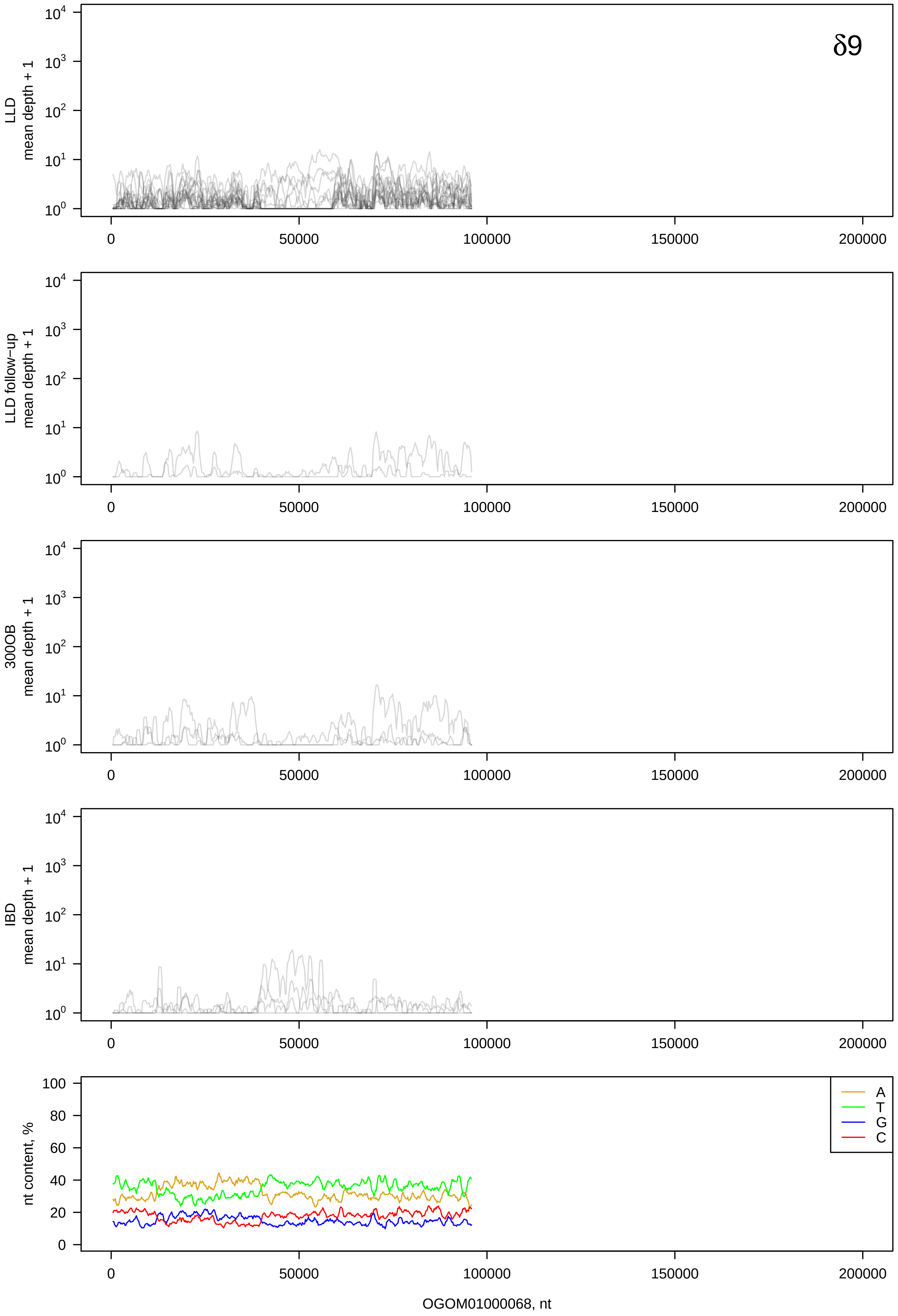
δ9
LLD mean depth + 1
LLD follow−up mean depth + 1
300OB mean depth + 1
IBD mean depth + 1
nt content, %
A
T
G
C
OGOM01000068, nt

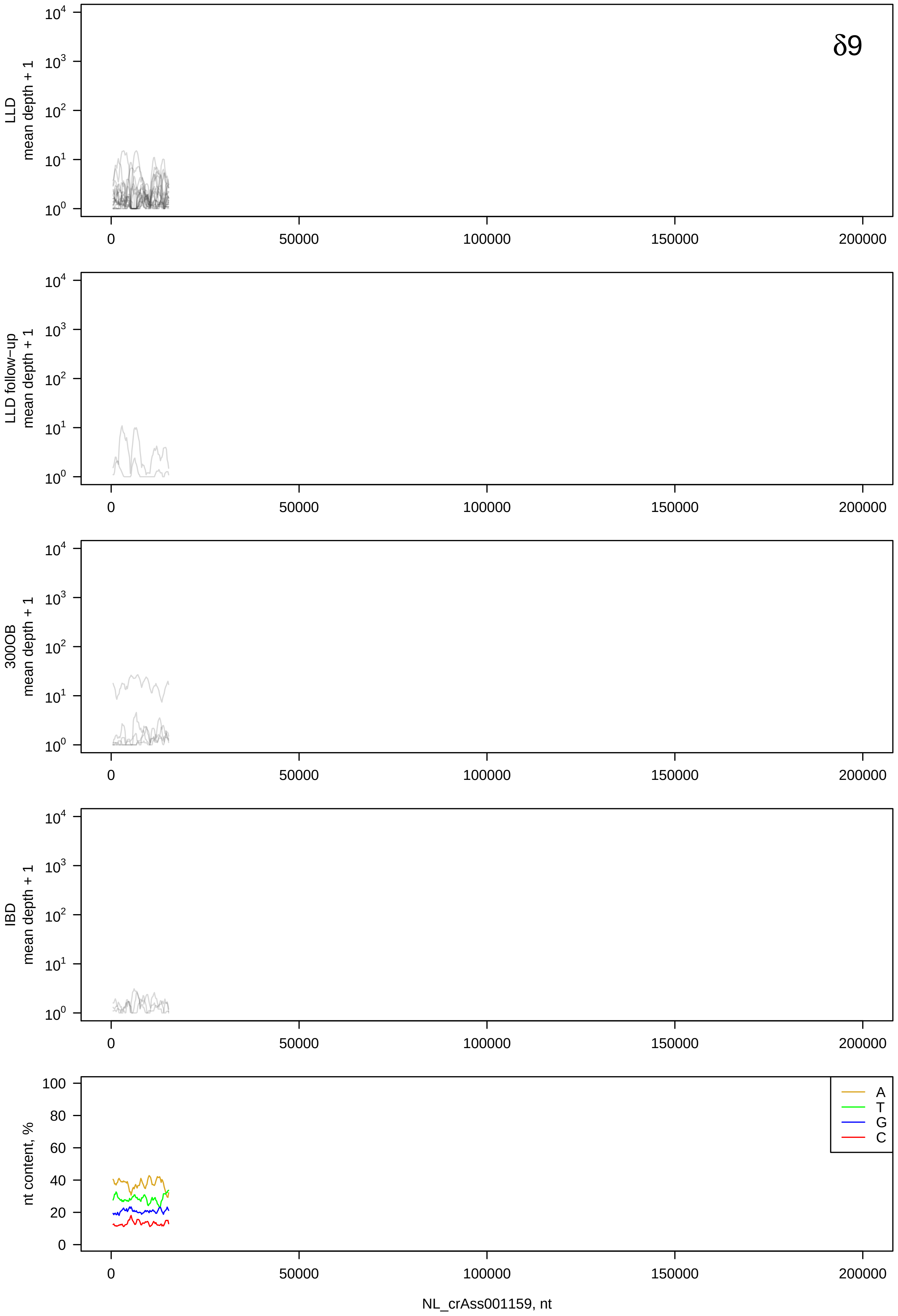
δ9
LLD mean depth + 1
LLD follow-up mean depth + 1
300OB mean depth + 1
IBD mean depth + 1
nt content, %
A
T
G
C
NL_crAss001159, nt

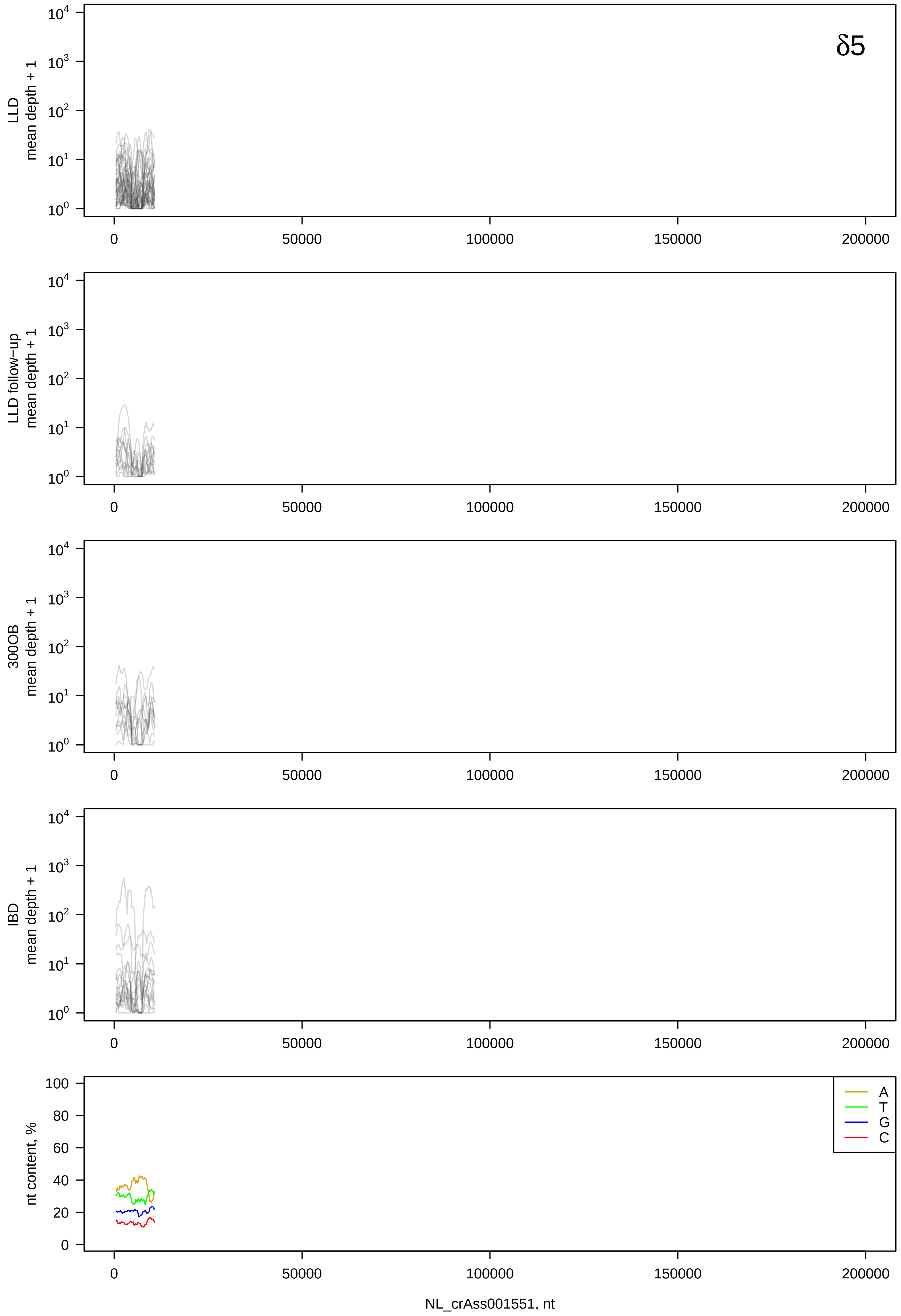

δ5
LLD mean depth + 1
LLD follow−up mean depth + 1
300OB mean depth + 1
IBD mean depth + 1
nt content, %
A
T
G
C
NL_crAss001551, nt

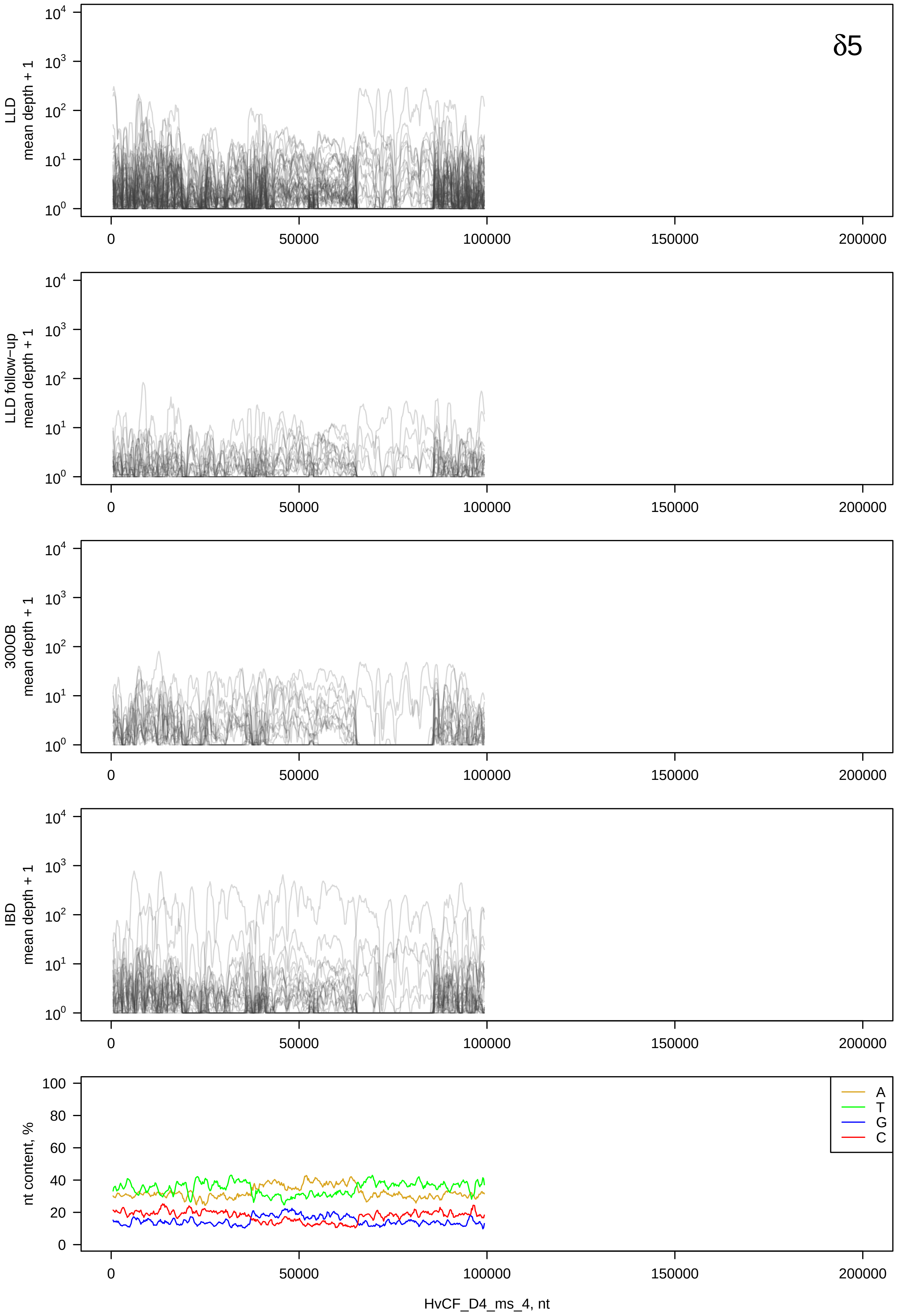

δ5
LLD
mean depth + 1
LLD follow−up
mean depth + 1
300OB
mean depth + 1
IBD
mean depth + 1
nt content, %
A
T
G
C
0
50000
100000
150000
200000
HvCF_D4_ms_4, nt

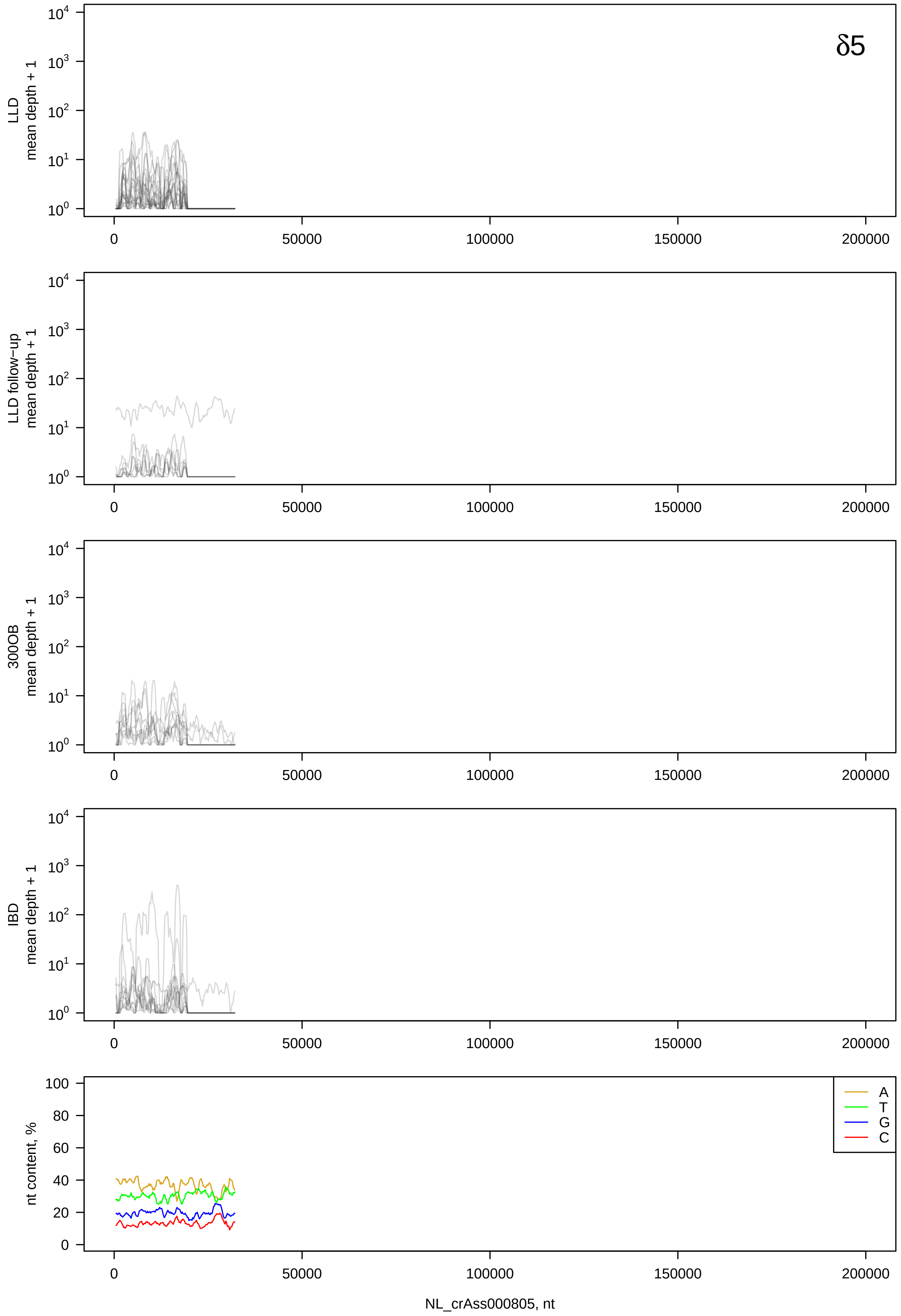

δ5
LLD mean depth + 1
LLD follow−up mean depth + 1
300OB mean depth + 1
IBD mean depth + 1
nt content, %
A
T
G
C
NL_crAss000805, nt

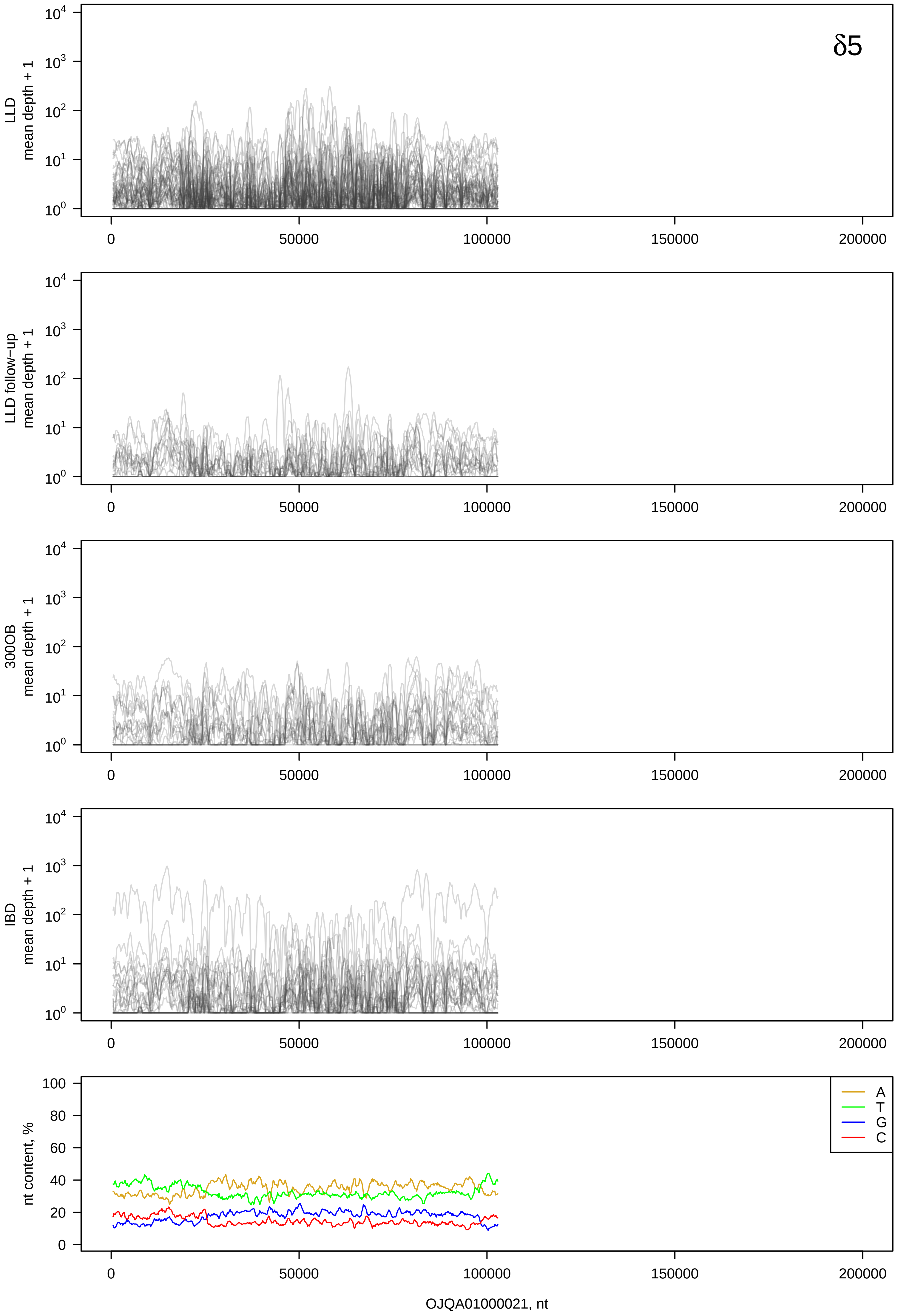
δ5
LLD mean depth + 1
LLD follow-up mean depth + 1
300OB mean depth + 1
IBD mean depth + 1
nt content, %
A
T
G
C
OJQA01000021, nt

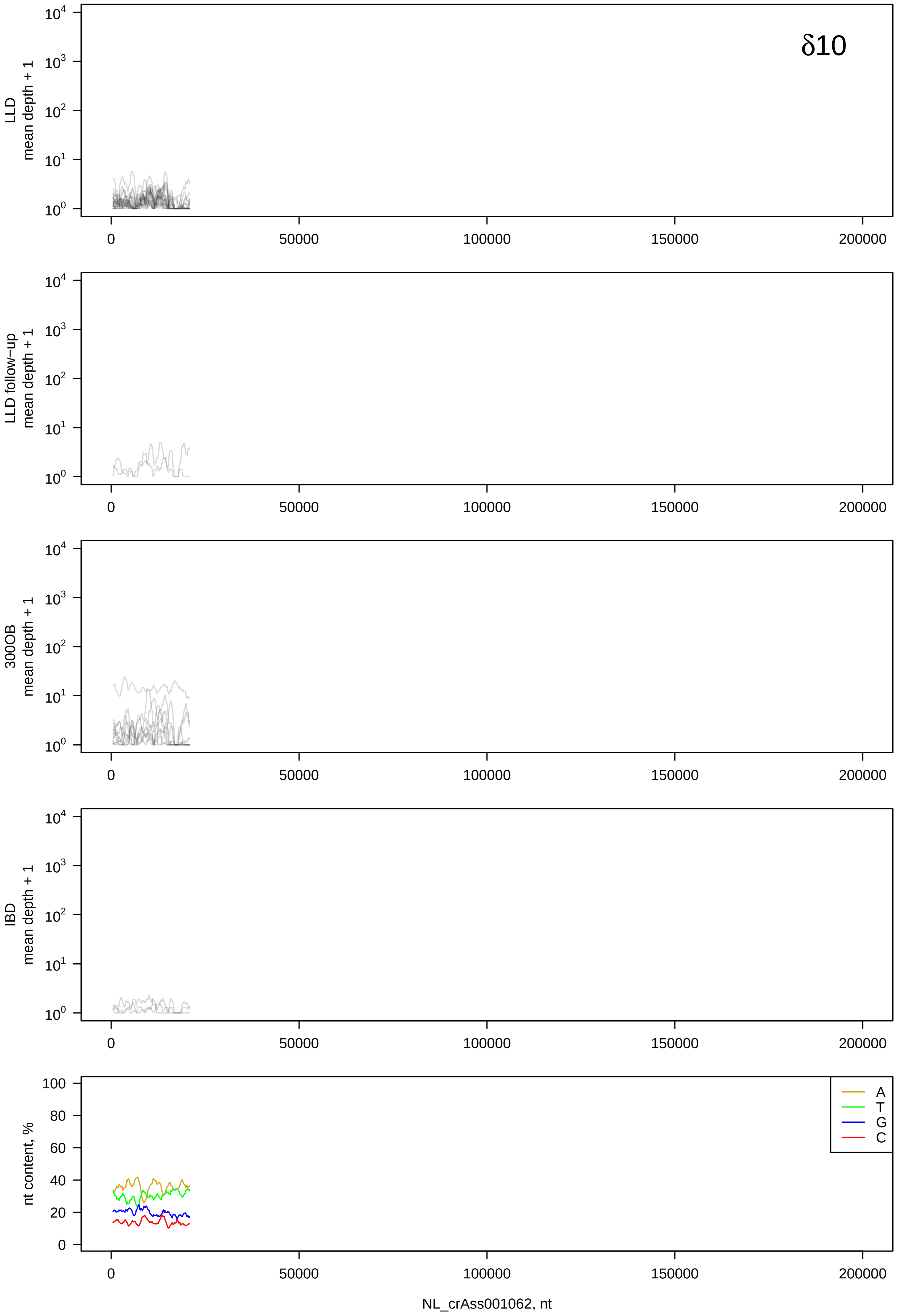

δ10
LLD
mean depth + 1
LLD follow−up
mean depth + 1
300OB
mean depth + 1
IBD
mean depth + 1
nt content, %
A
T
G
C
0
50000
100000
150000
200000
NL_crAss001062, nt

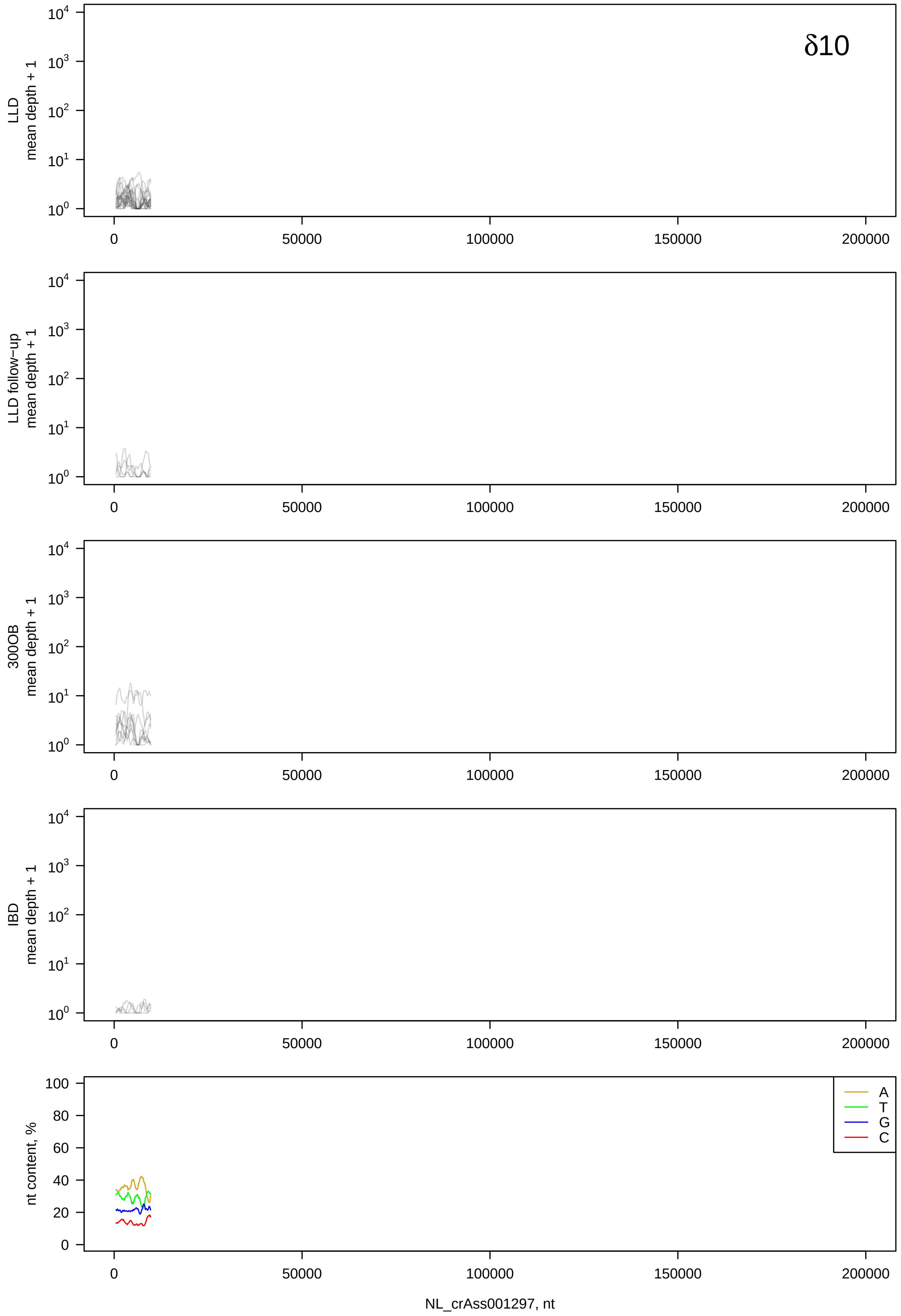
δ10
LLD
mean depth + 1
LLD follow-up
mean depth + 1
300OB
mean depth + 1
IBD
mean depth + 1
nt content, %
A
T
G
C
0
50000
100000
150000
200000
NL_crAss001297, nt

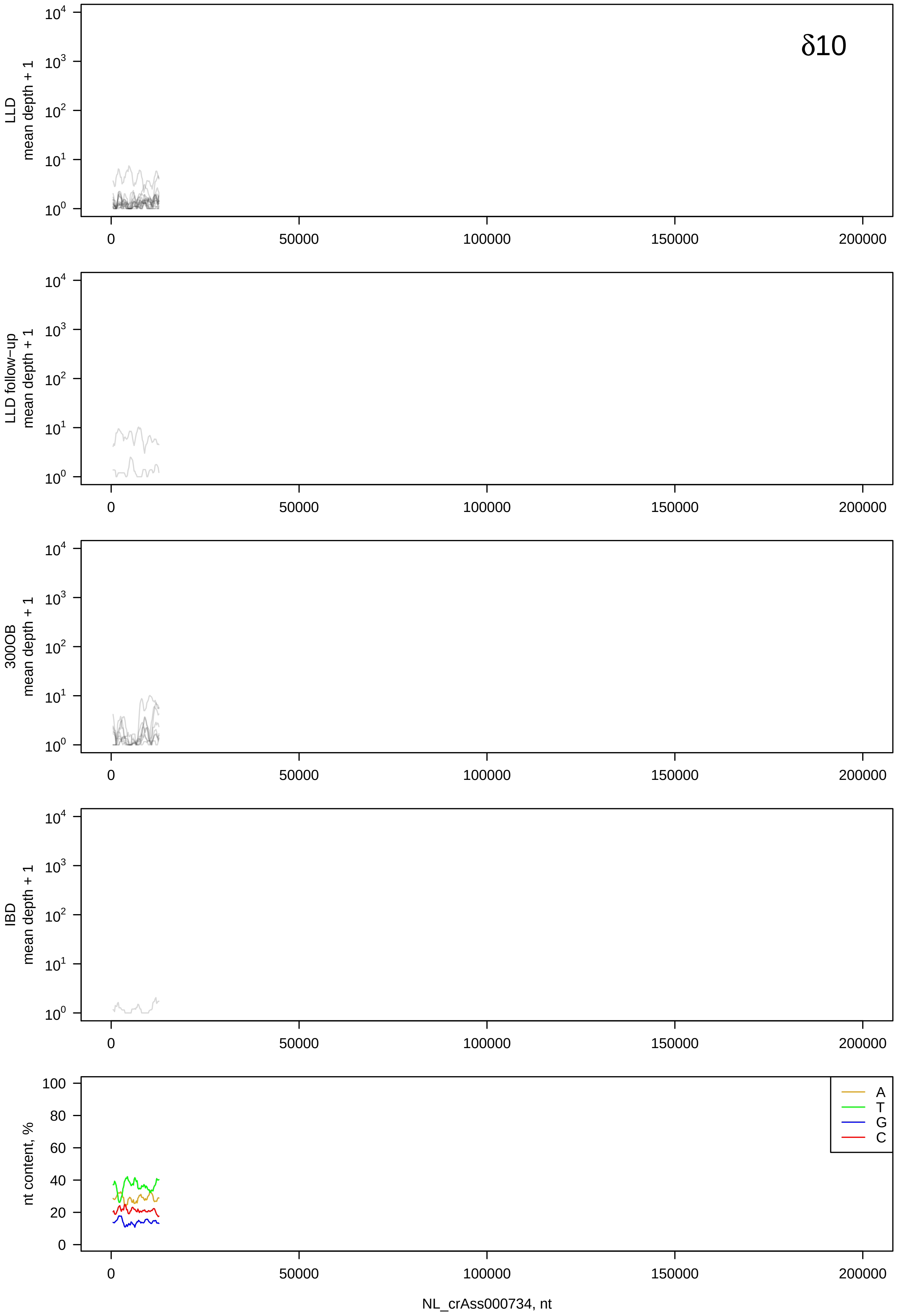
δ10
LLD
mean depth + 1
LLD follow-up
mean depth + 1
300OB
mean depth + 1
IBD
mean depth + 1
nt content, %
A
T
G
C
NL_crAss000734, nt

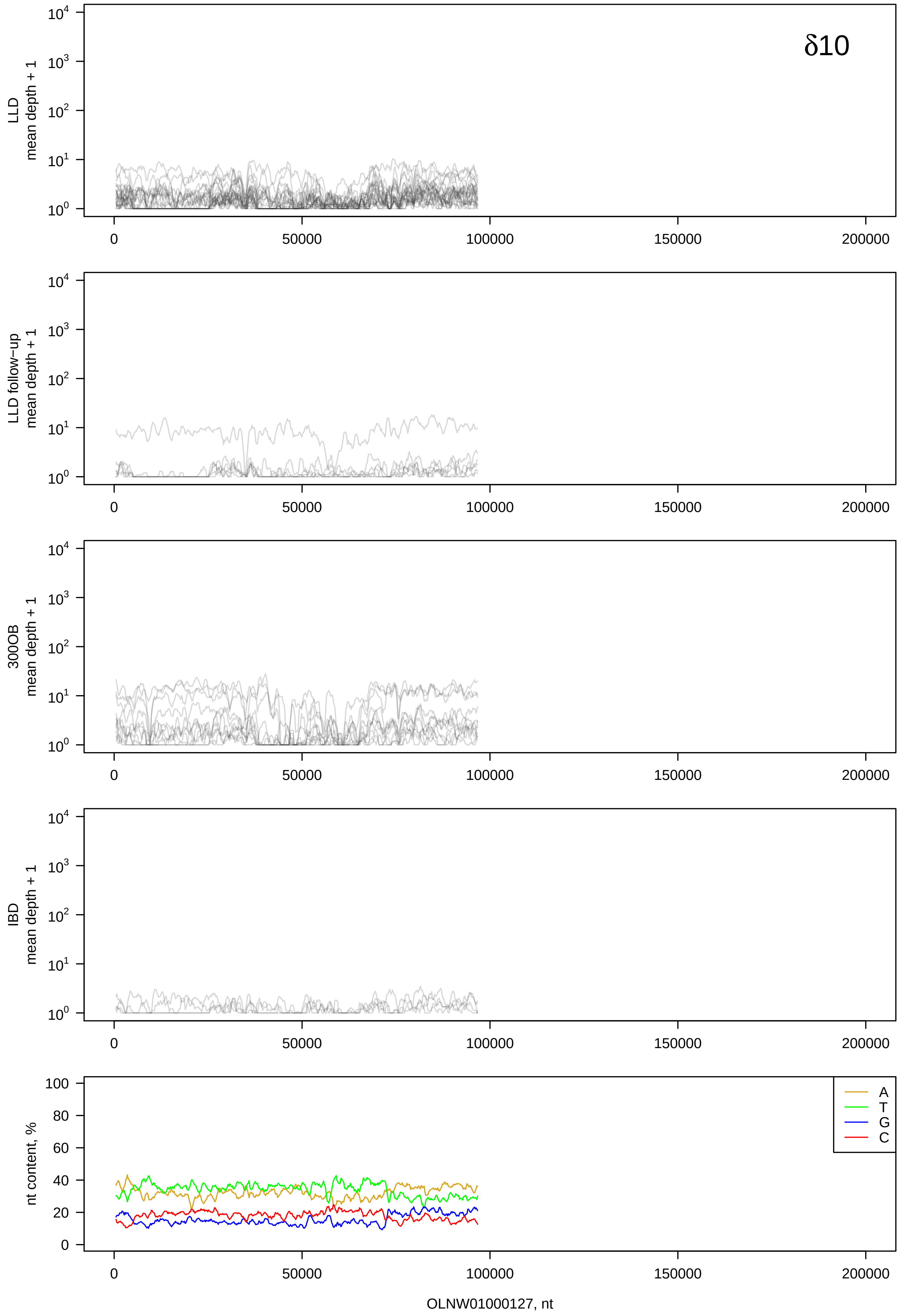
δ10
LLD
mean depth + 1
LLD follow−up
mean depth + 1
300OB
mean depth + 1
IBD
mean depth + 1
nt content, %
A
T
G
C
0
50000
100000
150000
200000
OLNW01000127, nt

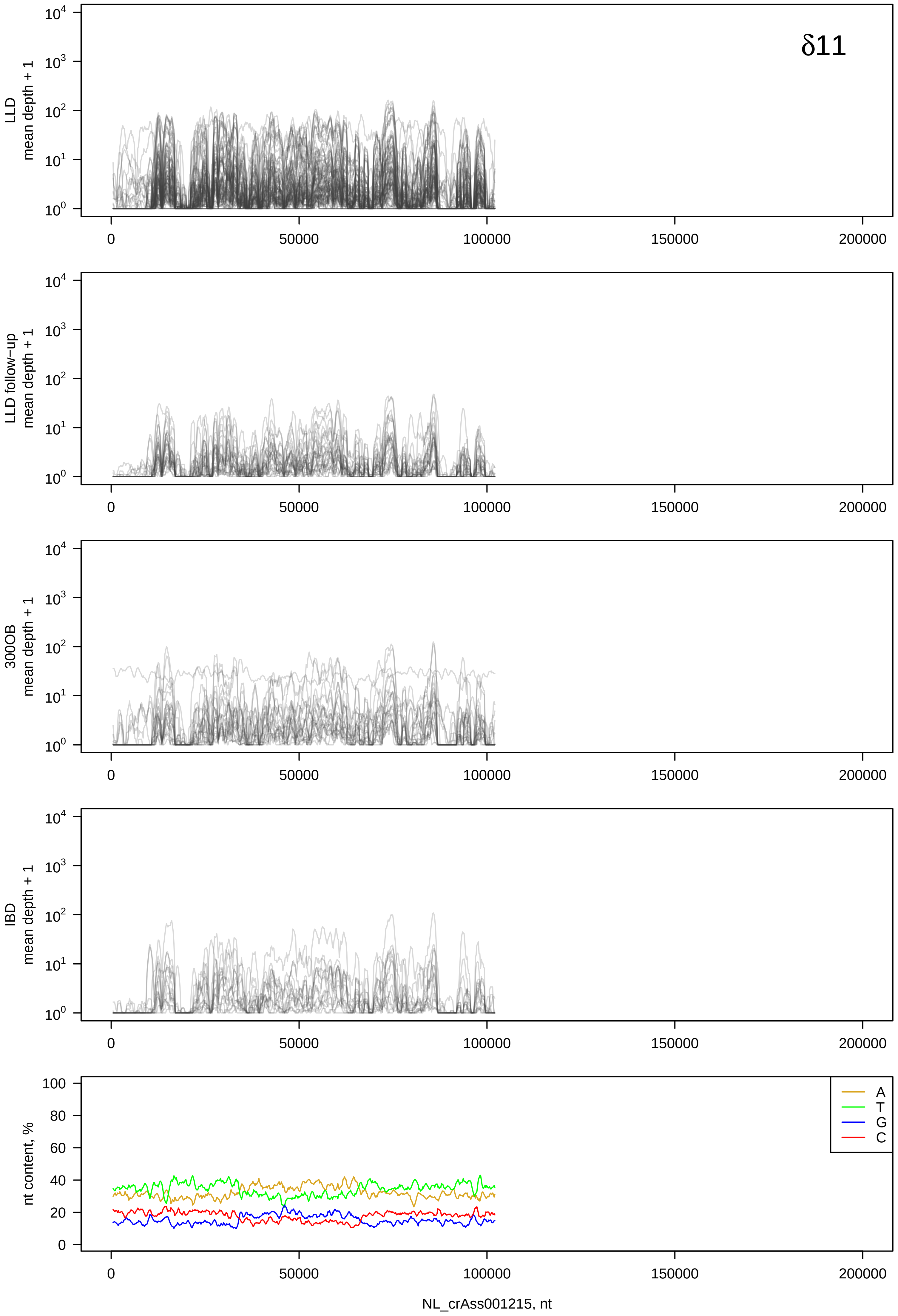

δ11
LLD
mean depth + 1
LLD follow−up
mean depth + 1
300OB
mean depth + 1
IBD
mean depth + 1
$10^0$
$10^1$
$10^2$
$10^3$
$10^4$
nt content, %
0
20
40
60
80
100
A
T
G
C
0
50000
100000
150000
200000
NL_crAss001215, nt

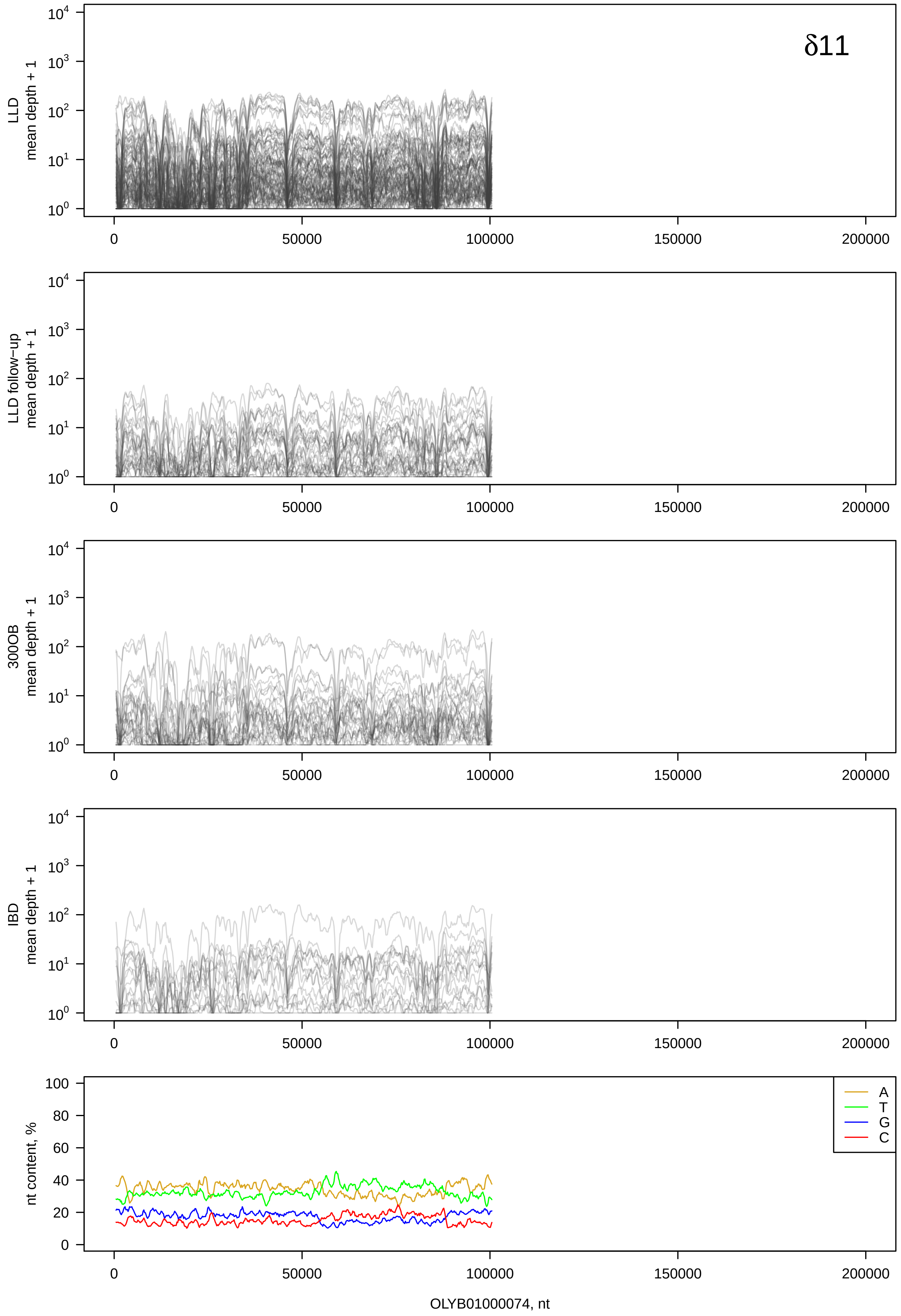
δ11
LLD
mean depth + 1
LLD follow−up
mean depth + 1
300OB
mean depth + 1
IBD
mean depth + 1
nt content, %
A
T
G
C
0
50000
100000
150000
200000
OLYB01000074, nt

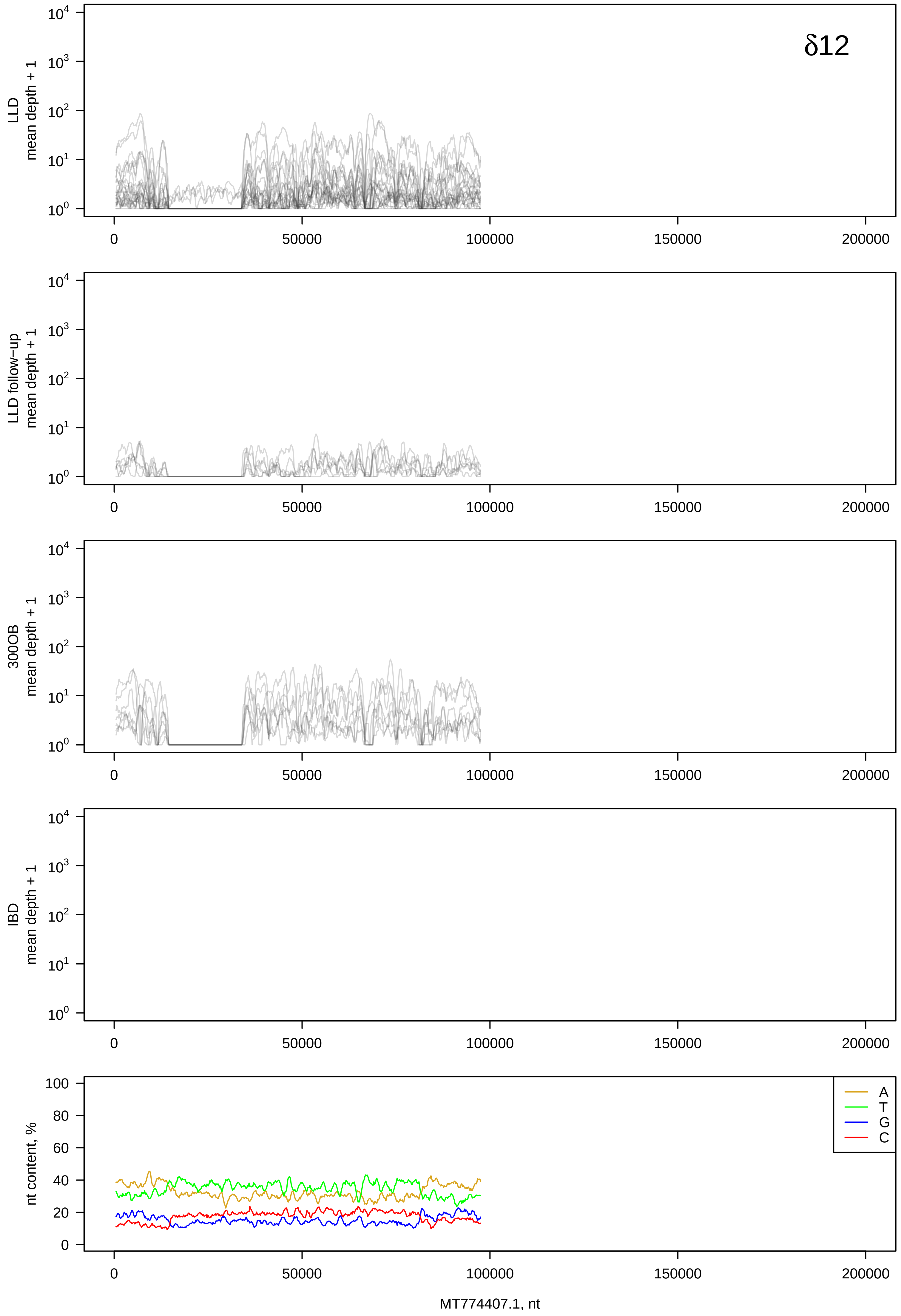
δ12
LLD
mean depth + 1
LLD follow−up
mean depth + 1
300OB
mean depth + 1
IBD
mean depth + 1
nt content, %
A
T
G
C
MT774407.1, nt

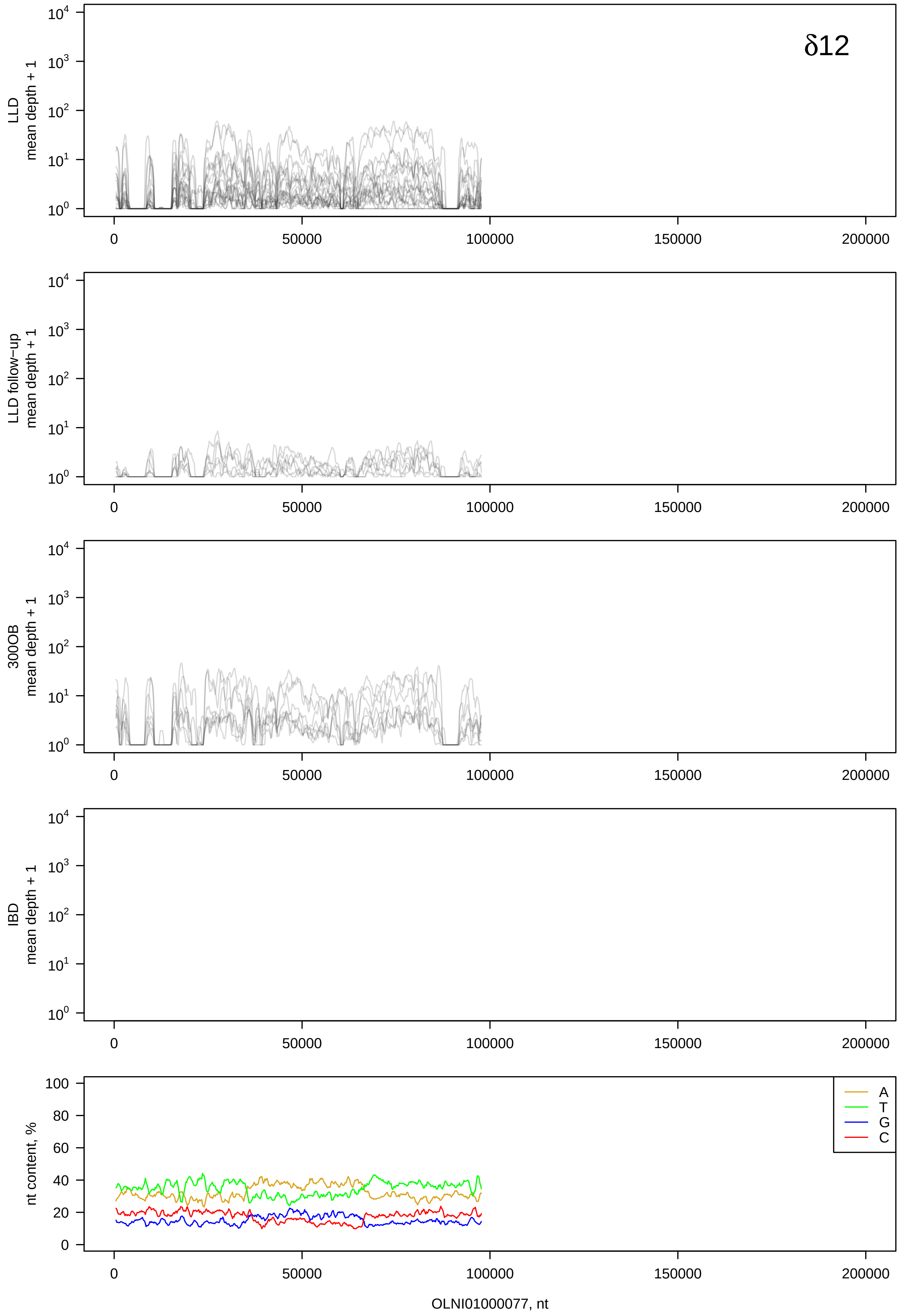
δ12
LLD mean depth + 1
LLD follow-up mean depth + 1
300OB mean depth + 1
IBD mean depth + 1
nt content, %
A
T
G
C
OLNI01000077, nt

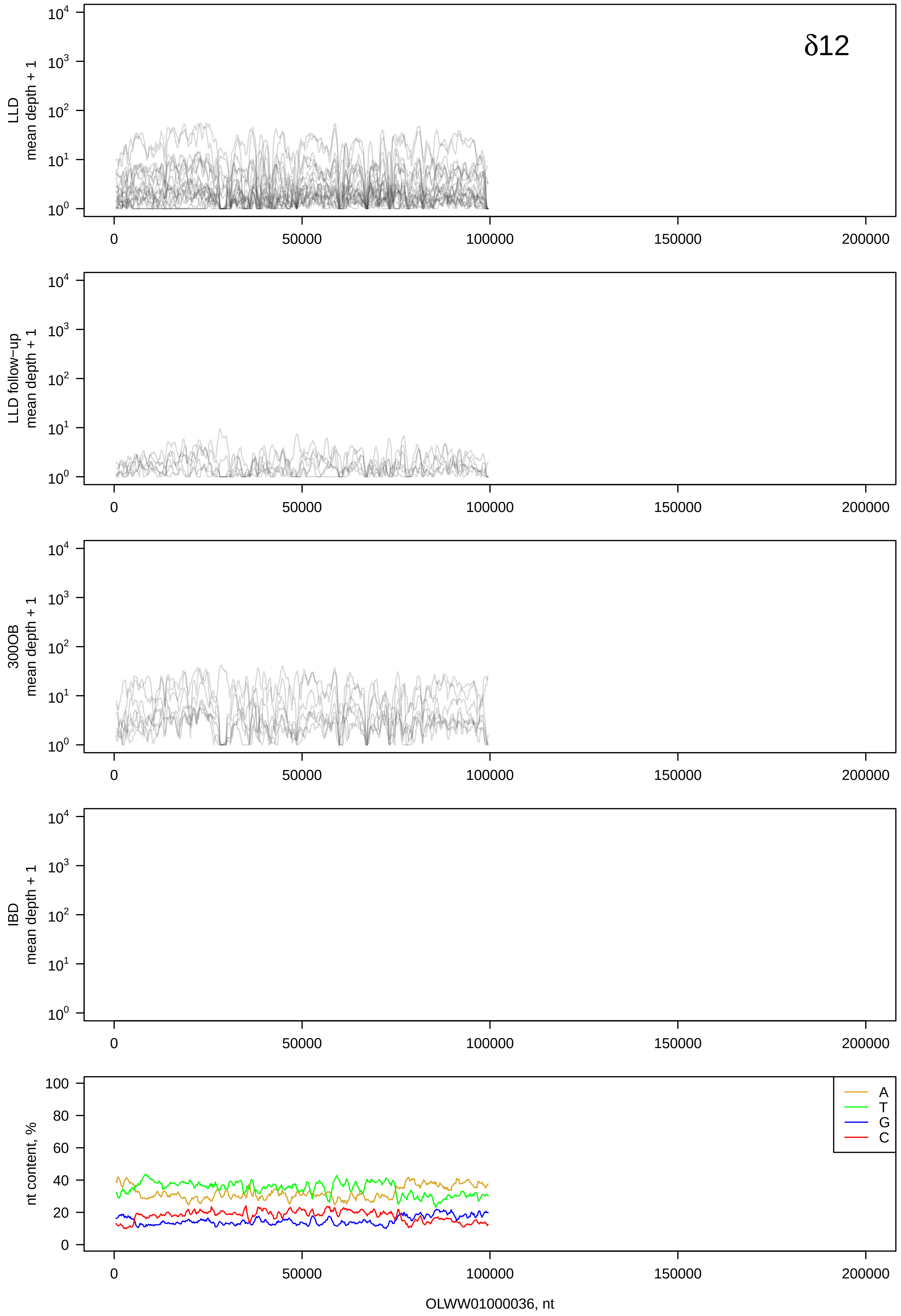
δ12
LLD mean depth + 1
LLD follow−up mean depth + 1
300OB mean depth + 1
IBD mean depth + 1
nt content, %
A
T
G
C
OLWW01000036, nt

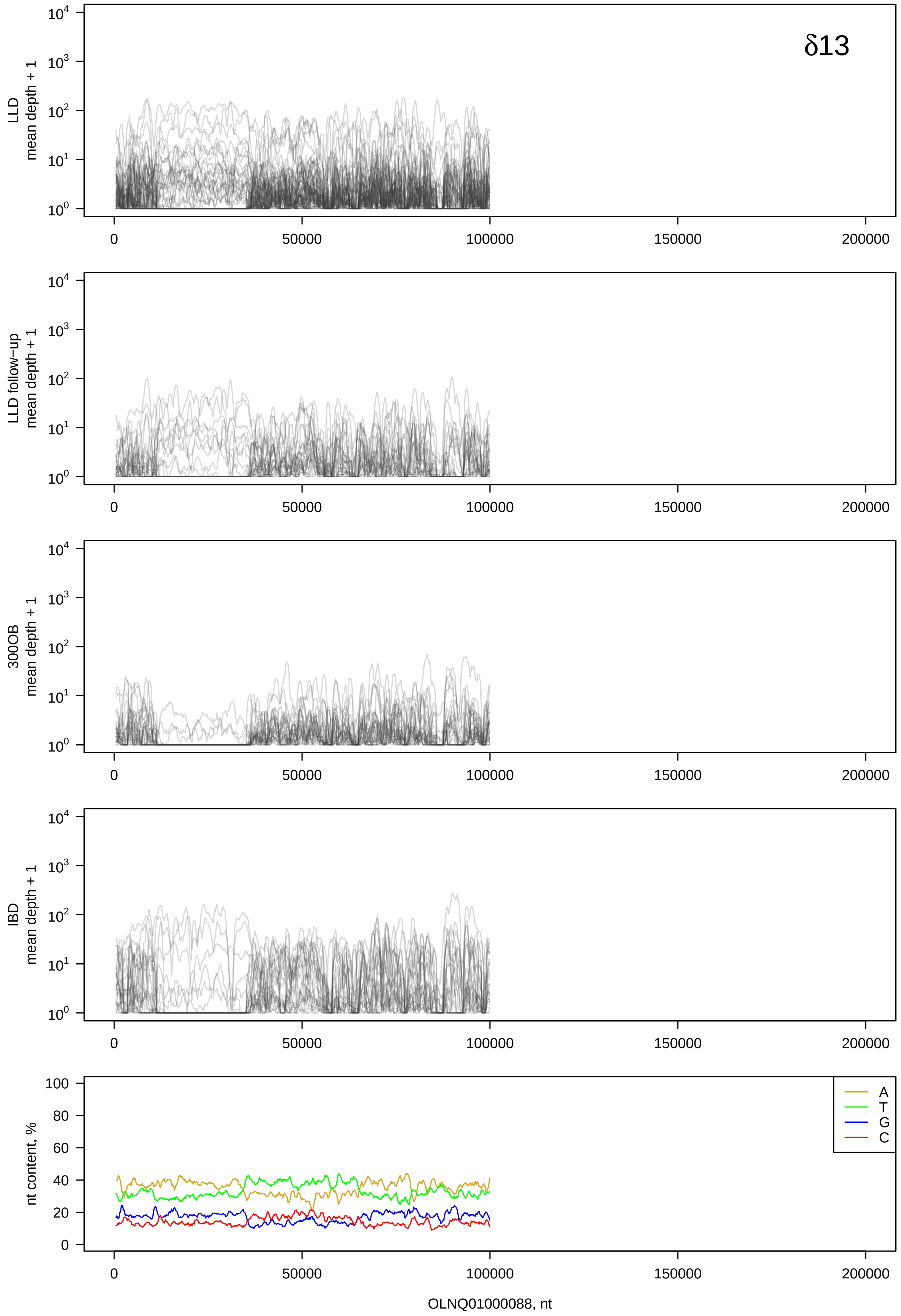

δ13
LLD
mean depth + 1
LLD follow−up
mean depth + 1
300OB
mean depth + 1
IBD
mean depth + 1
nt content, %
A
T
G
C
0
50000
100000
150000
200000
OLNQ01000088, nt

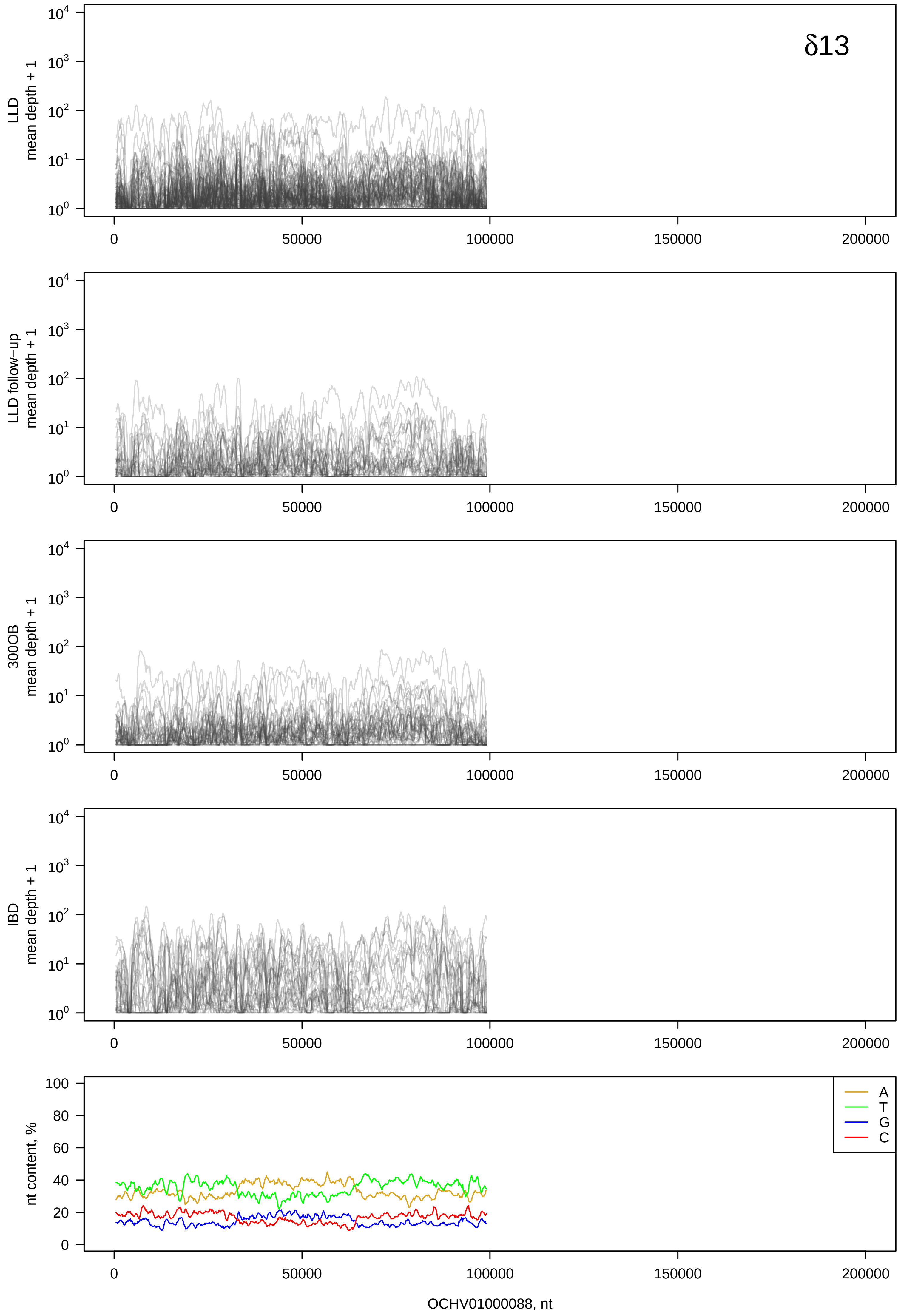
δ13
LLD
mean depth + 1
LLD follow−up
mean depth + 1
300OB
mean depth + 1
IBD
mean depth + 1
nt content, %
A
T
G
C
OCHV01000088, nt

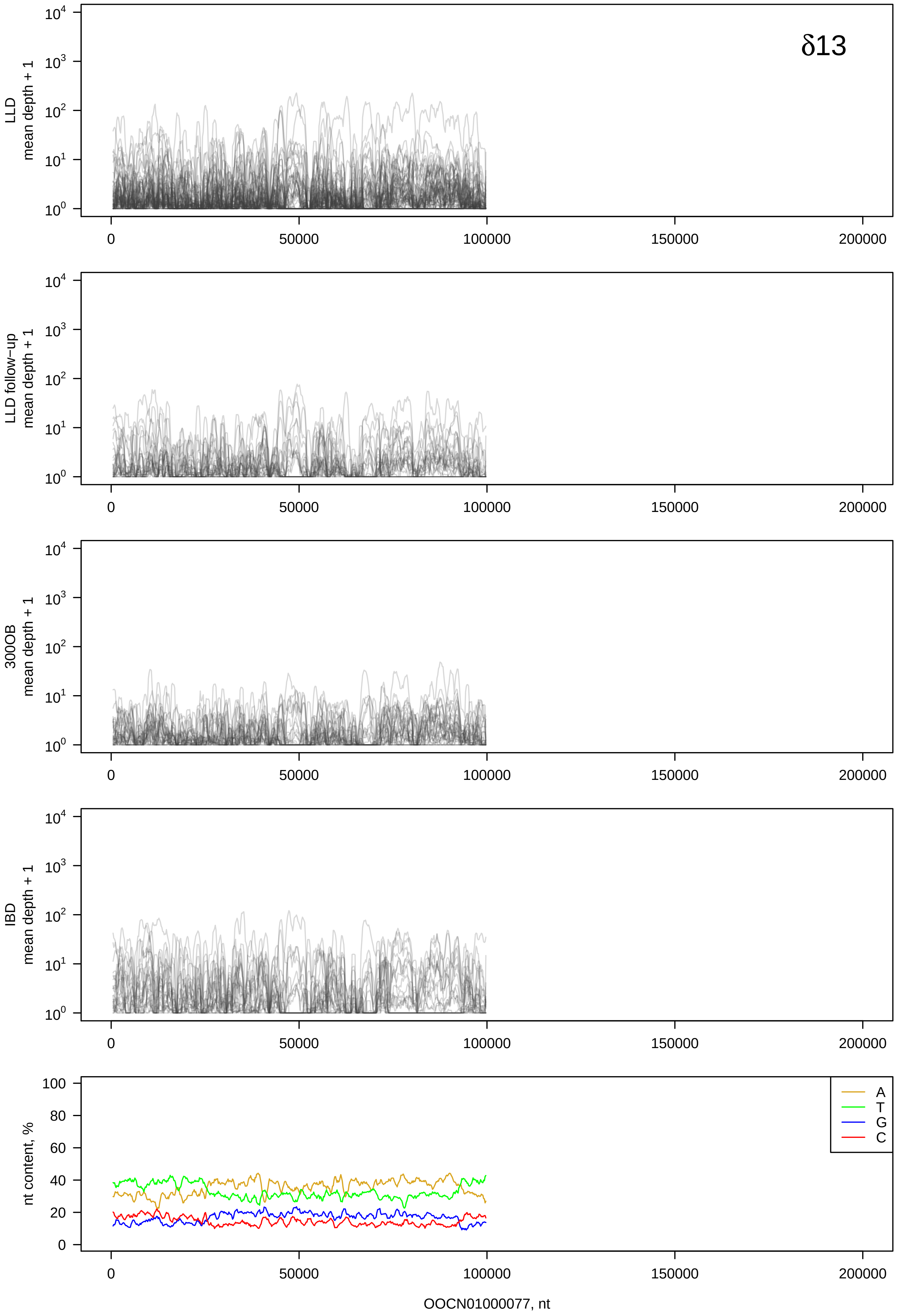
δ13
LLD
mean depth + 1
LLD follow−up
mean depth + 1
300OB
mean depth + 1
IBD
mean depth + 1
nt content, %
A
T
G
C
OOCN01000077, nt

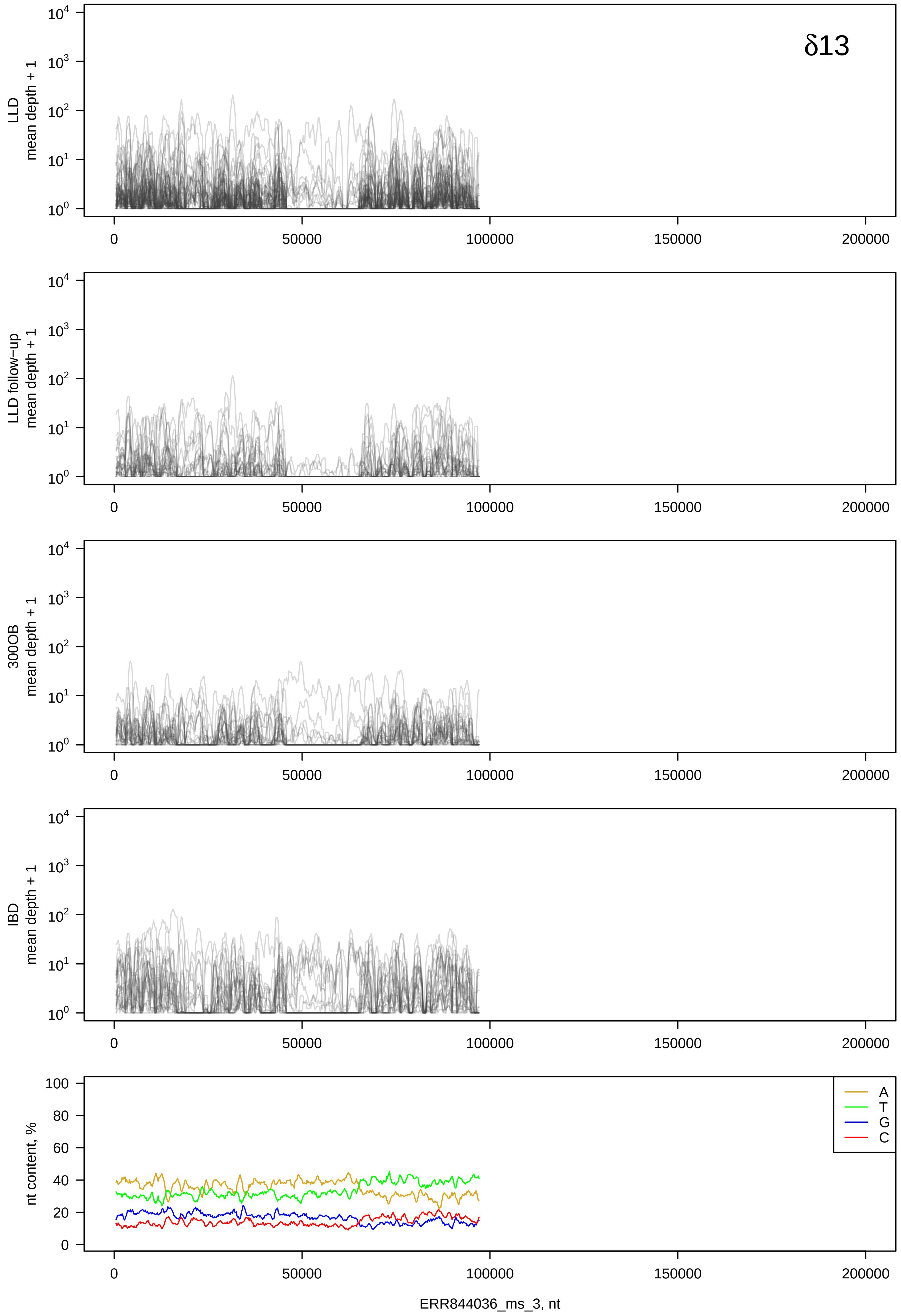

δ13
LLD mean depth + 1
LLD follow−up mean depth + 1
300OB mean depth + 1
IBD mean depth + 1
nt content, %
A
T
G
C
ERR844036_ms_3, nt

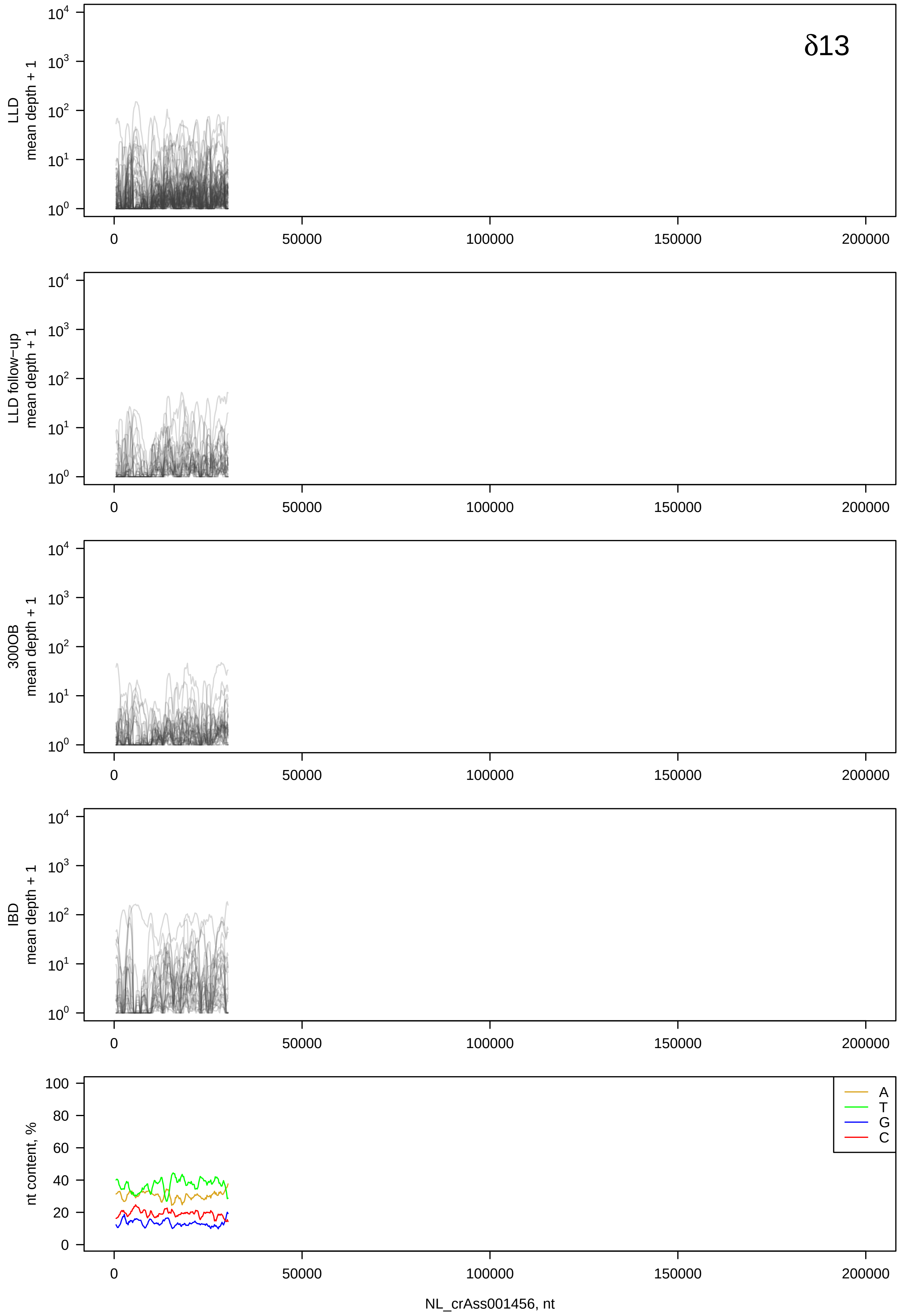
δ13
LLD
mean depth + 1
LLD follow−up
mean depth + 1
300OB
mean depth + 1
IBD
mean depth + 1
nt content, %
A
T
G
C
NL_crAss001456, nt

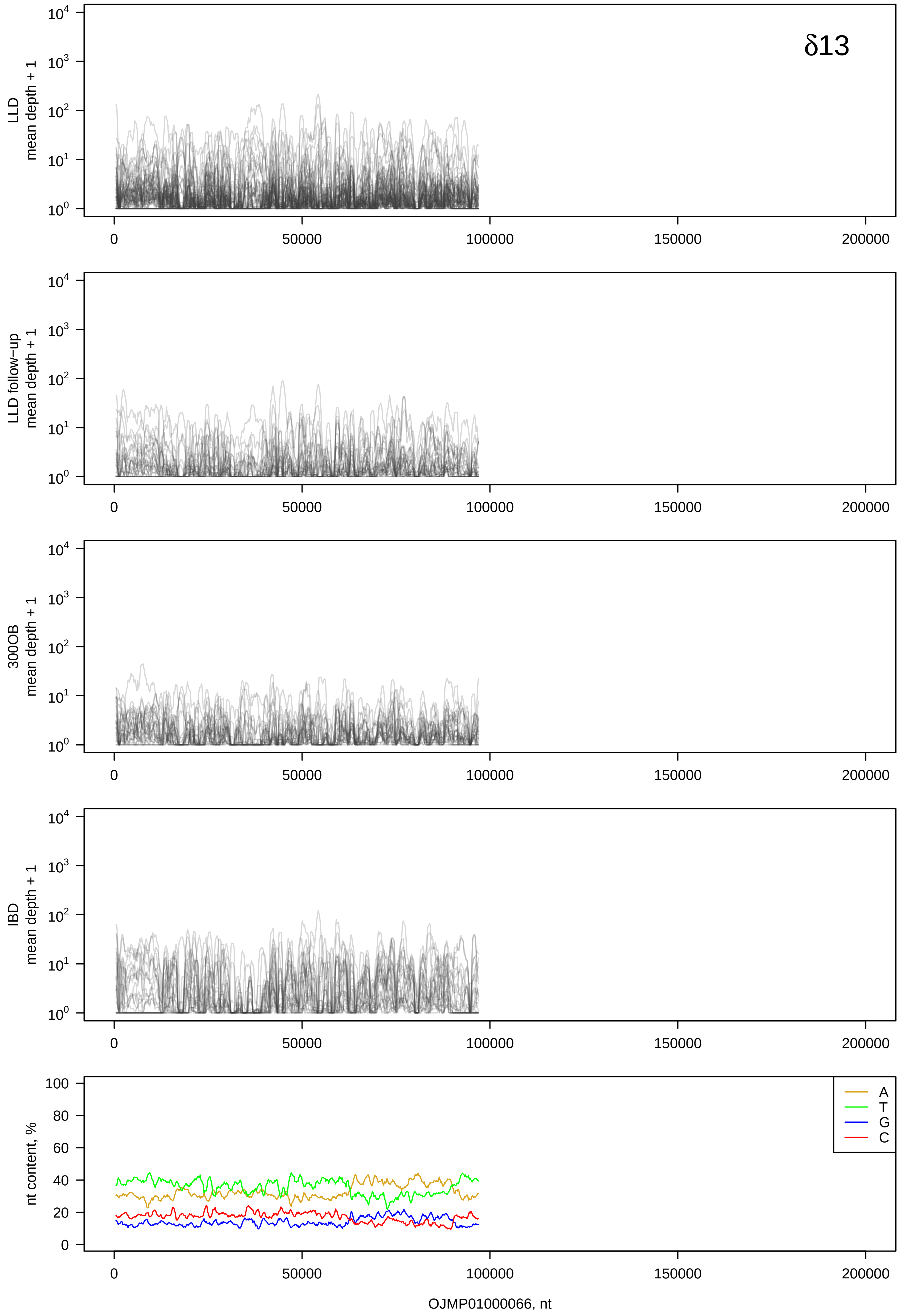

δ13
LLD mean depth + 1
LLD follow−up mean depth + 1
300OB mean depth + 1
IBD mean depth + 1
nt content, %
A
T
G
C
OJMP01000066, nt

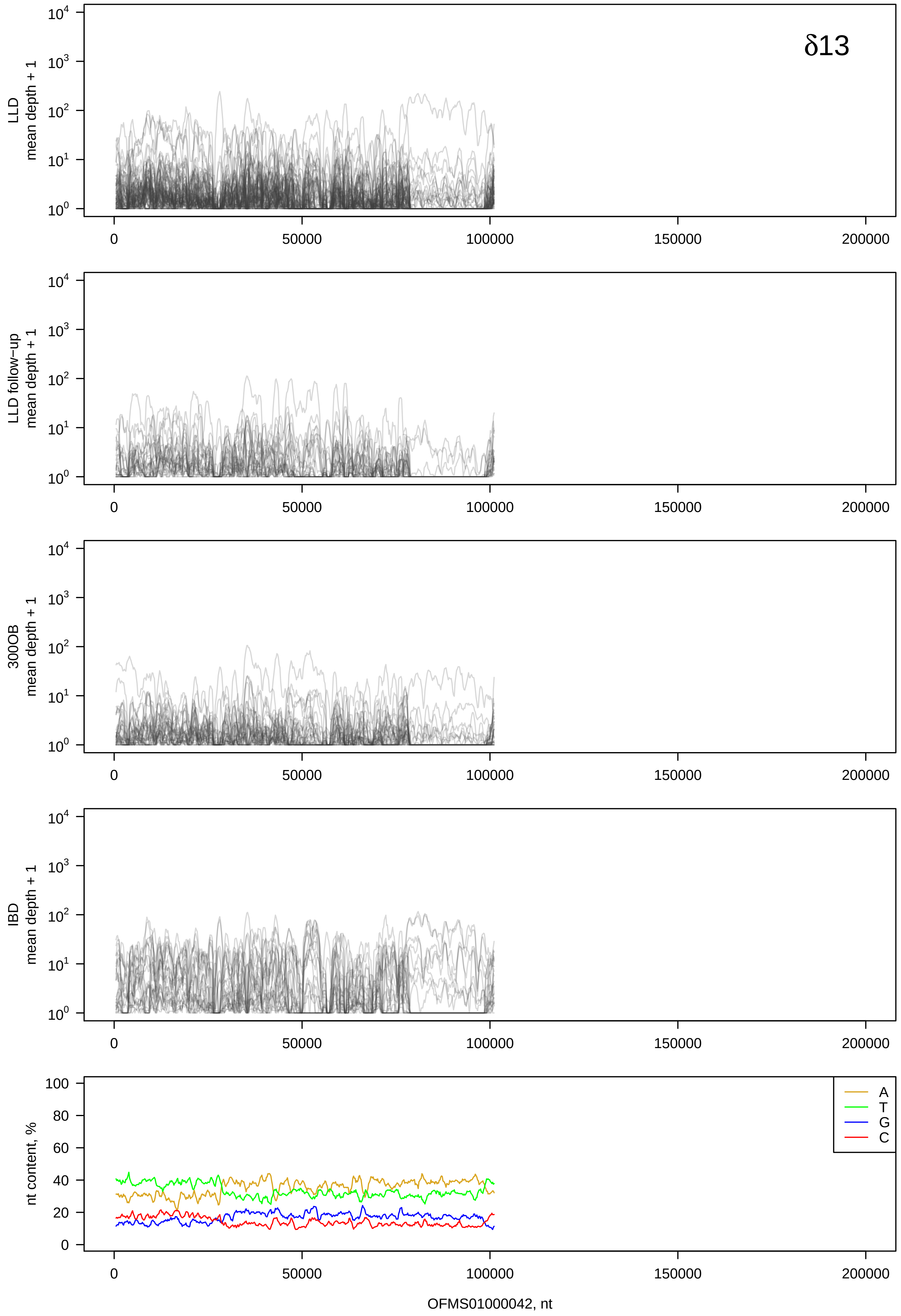

δ13
LLD mean depth + 1
LLD follow−up mean depth + 1
300OB mean depth + 1
IBD mean depth + 1
nt content, %
A
T
G
C
OFMS01000042, nt

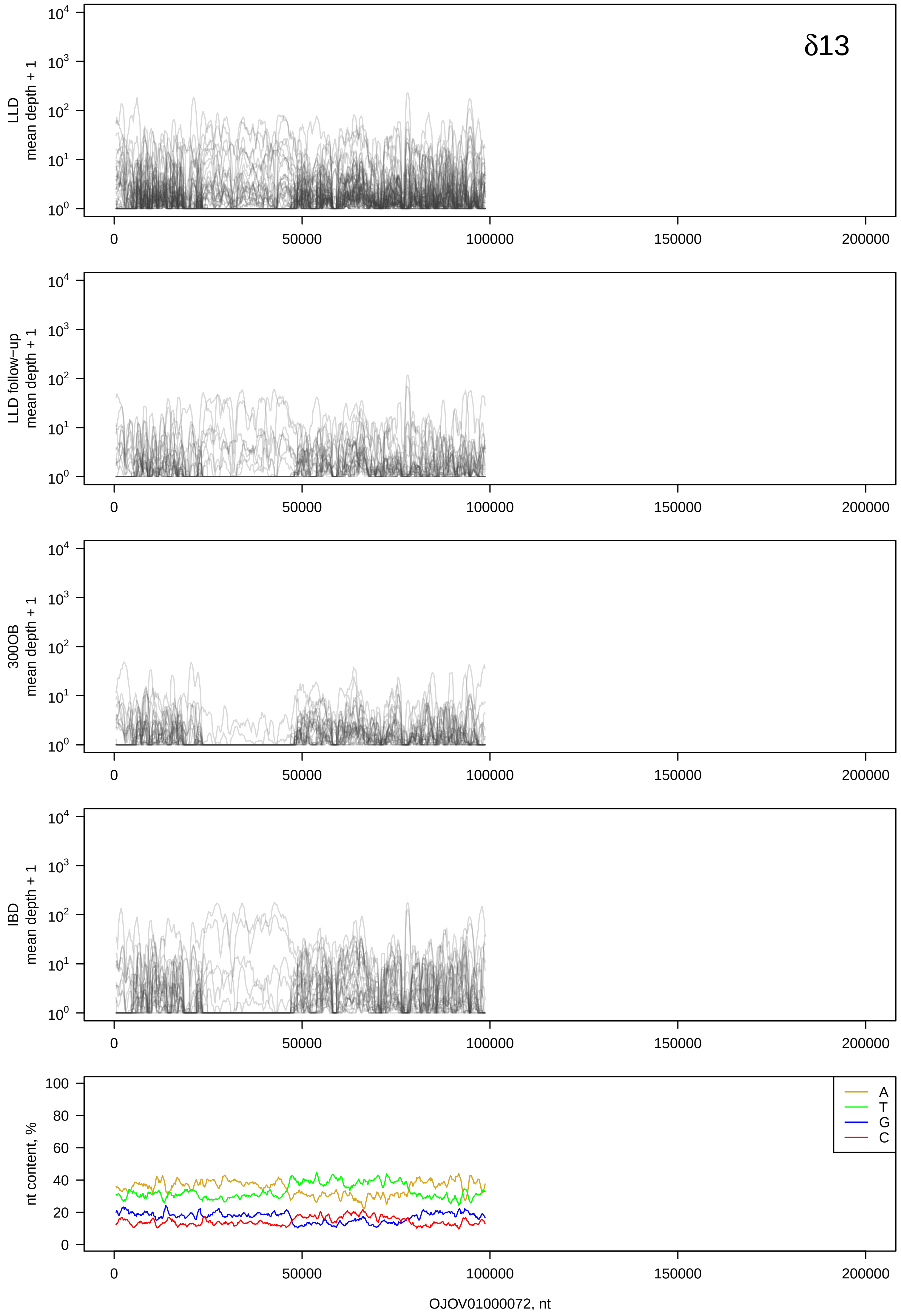

δ13
LLD
mean depth + 1
LLD follow−up
mean depth + 1
300OB
mean depth + 1
IBD
mean depth + 1
nt content, %
A
T
G
C
OJOV01000072, nt

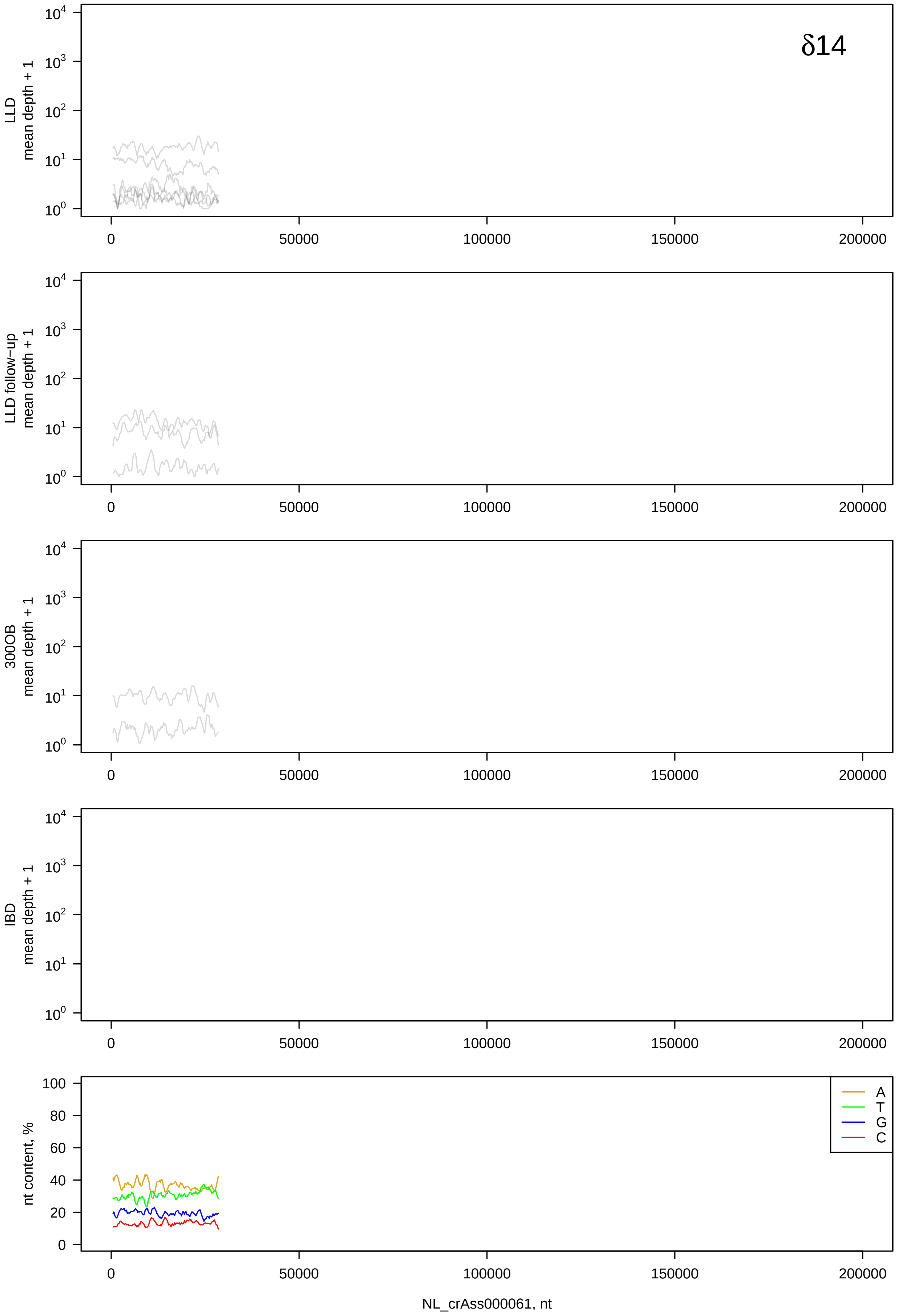
δ14
LLD
mean depth + 1
LLD follow-up
mean depth + 1
300OB
mean depth + 1
IBD
mean depth + 1
nt content, %
A
T
G
C
0
50000
100000
150000
200000
NL_crAss000061, nt

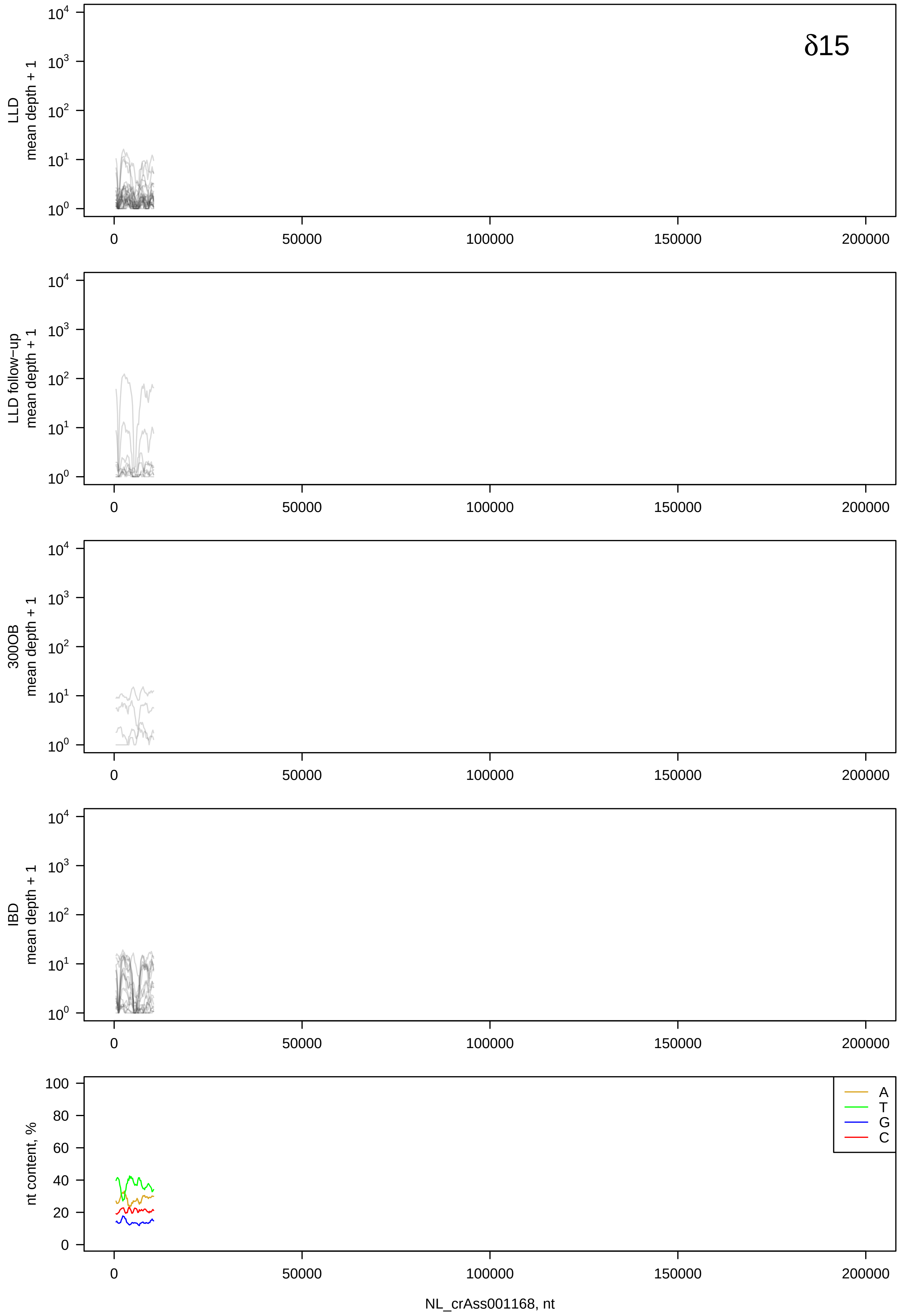
δ15
LLD
mean depth + 1
LLD follow−up
mean depth + 1
300OB
mean depth + 1
IBD
mean depth + 1
nt content, %
A
T
G
C
0
50000
100000
150000
200000
NL_crAss001168, nt

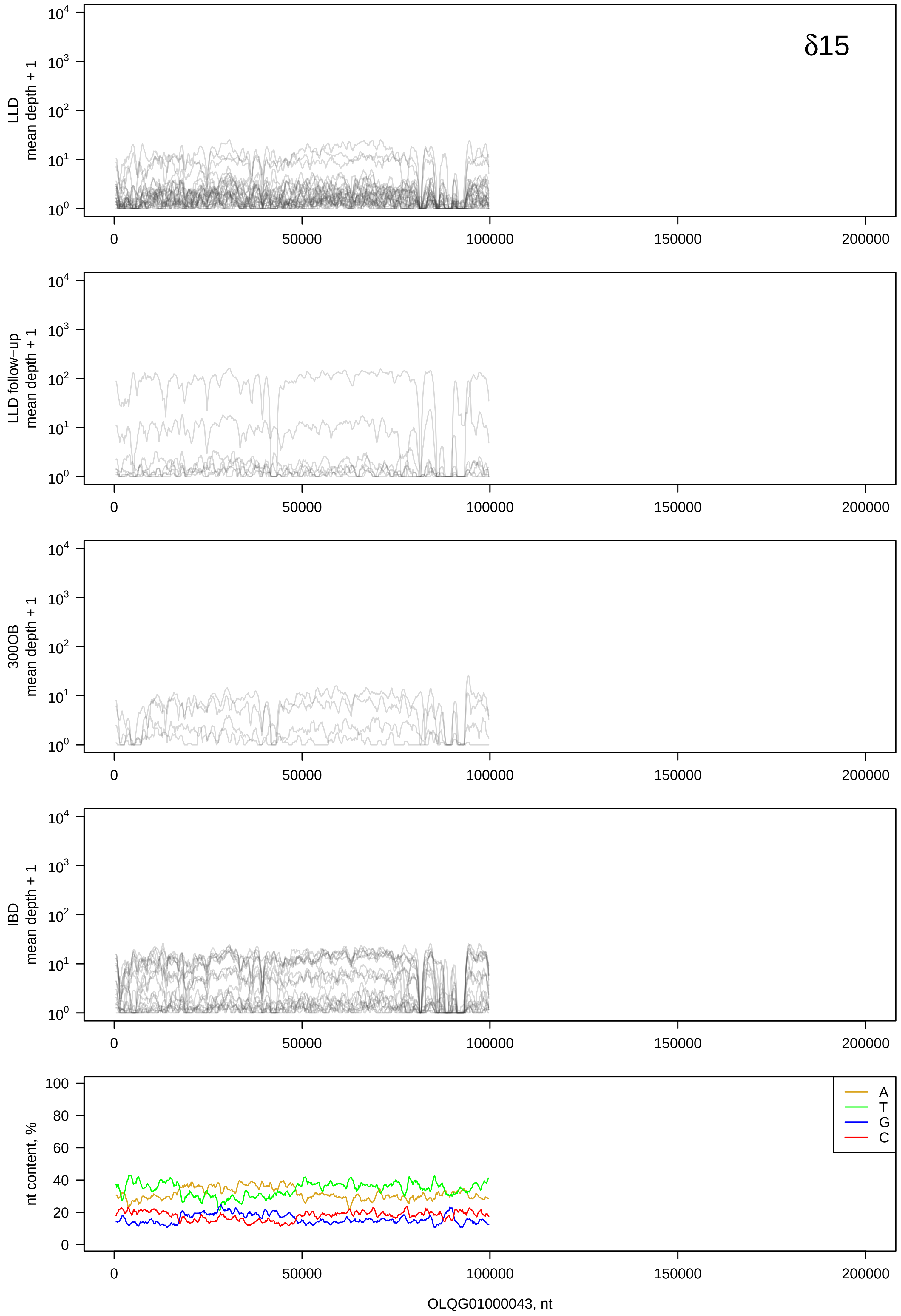
δ15
LLD
mean depth + 1
LLD follow-up
mean depth + 1
300OB
mean depth + 1
IBD
mean depth + 1
nt content, %
A
T
G
C
0
50000
100000
150000
200000
OLQG01000043, nt

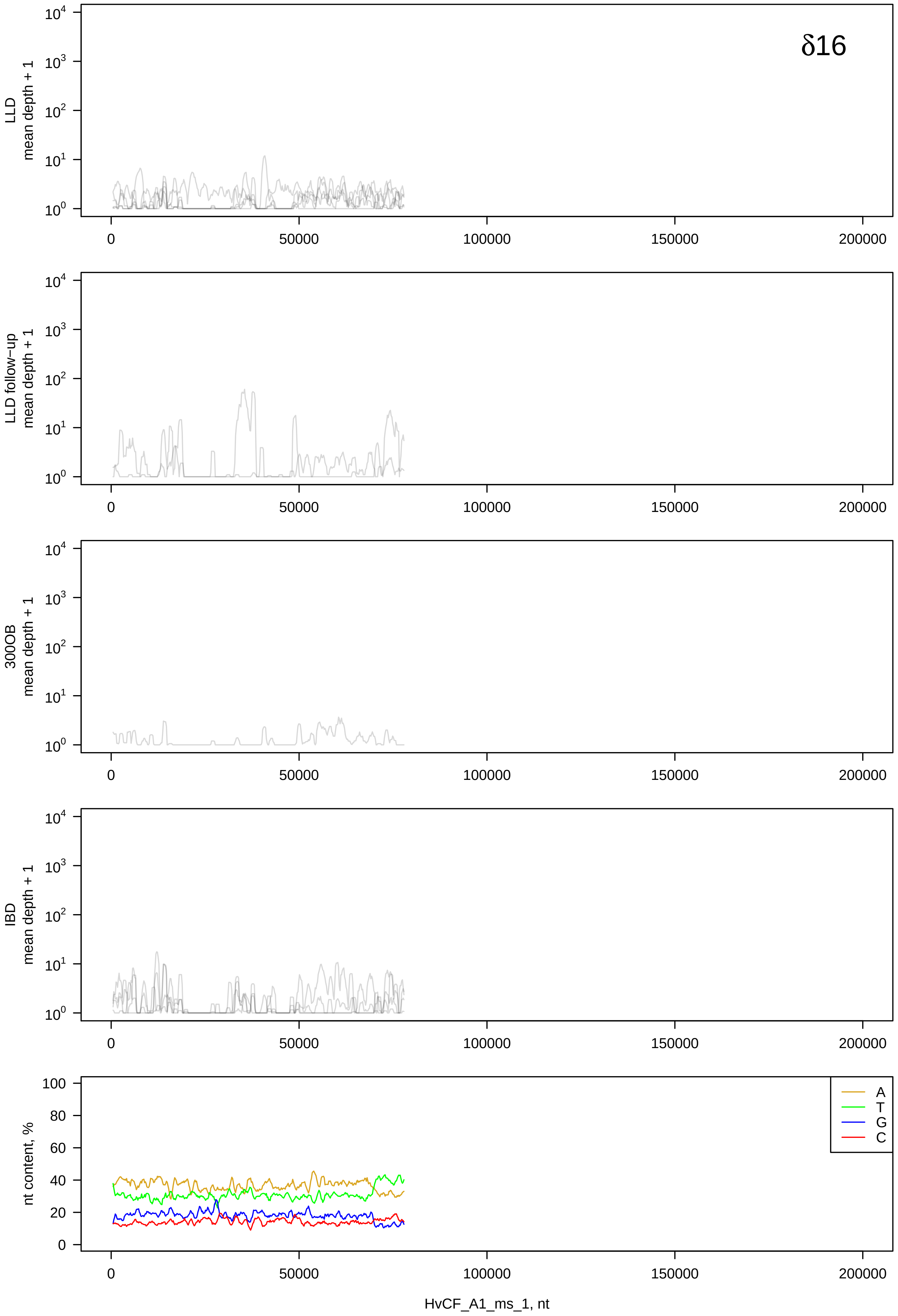
δ16
LLD mean depth + 1
LLD follow−up mean depth + 1
300OB mean depth + 1
IBD mean depth + 1
nt content, %
A
T
G
C
HvCF_A1_ms_1, nt

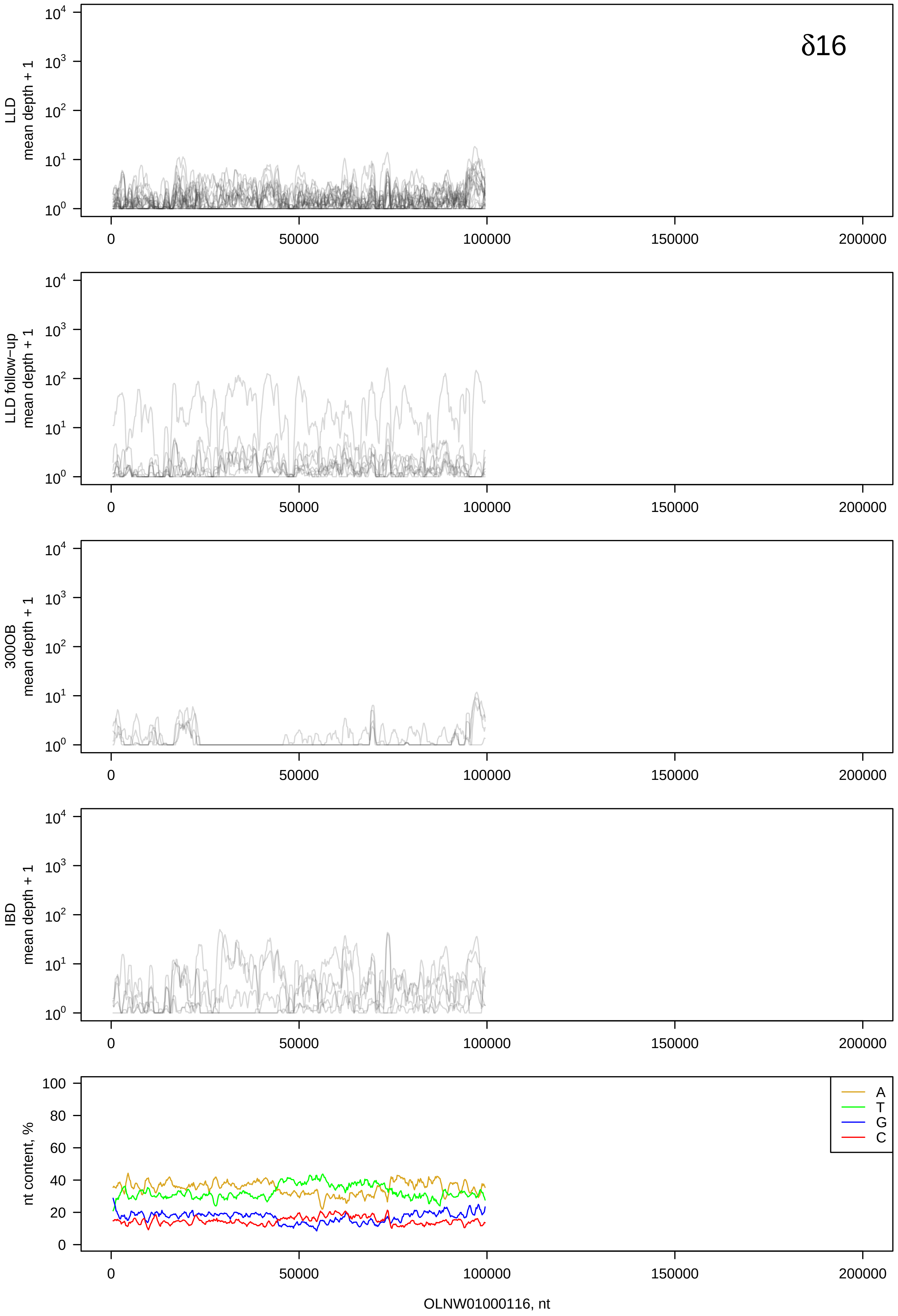

δ16
LLD
mean depth + 1
LLD follow−up
mean depth + 1
300OB
mean depth + 1
IBD
mean depth + 1
nt content, %
A
T
G
C
OLNW01000116, nt

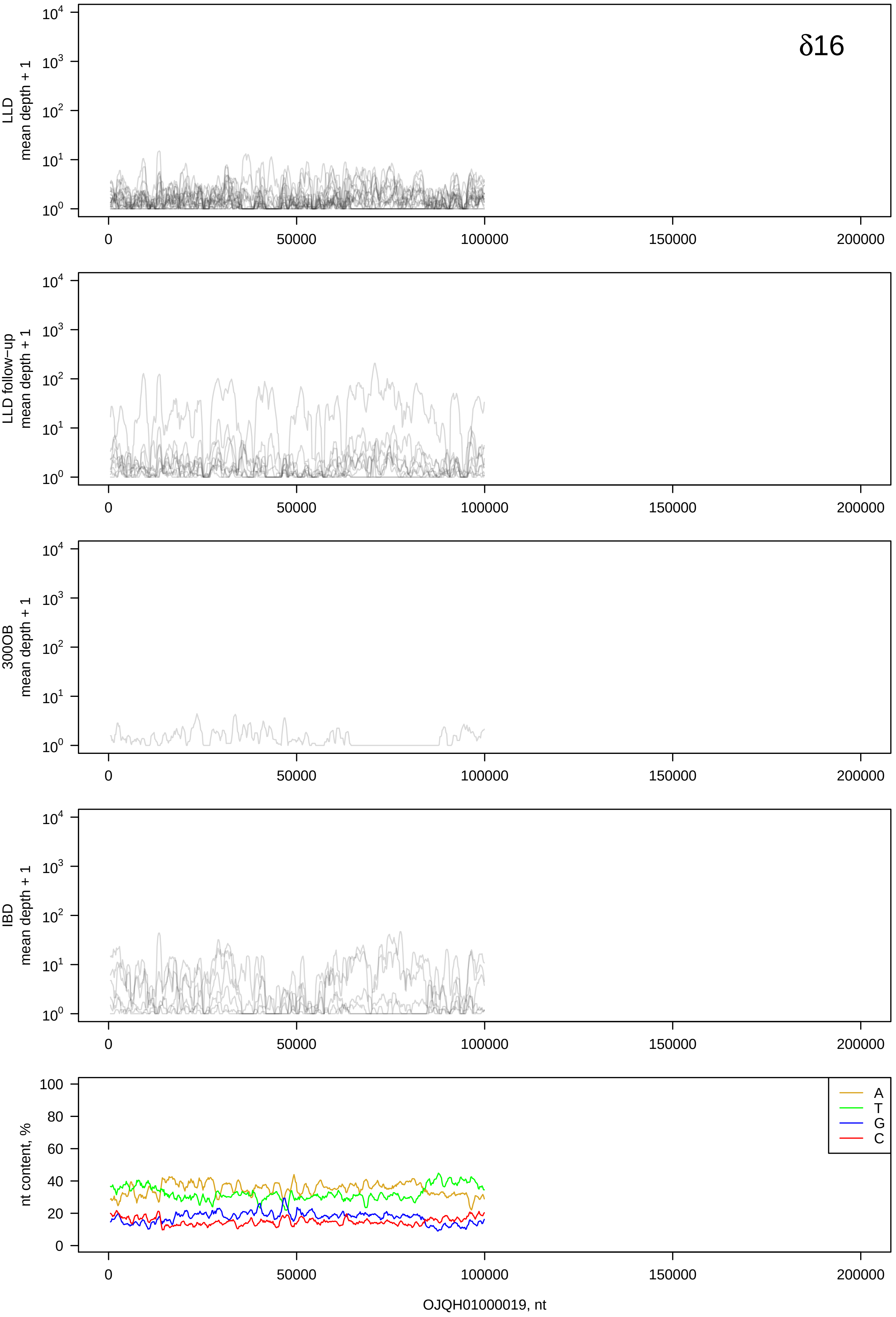

δ16
LLD
mean depth + 1
LLD follow−up
mean depth + 1
300OB
mean depth + 1
IBD
mean depth + 1
nt content, %
A
T
G
C
0
50000
100000
150000
200000
OJQH01000019, nt

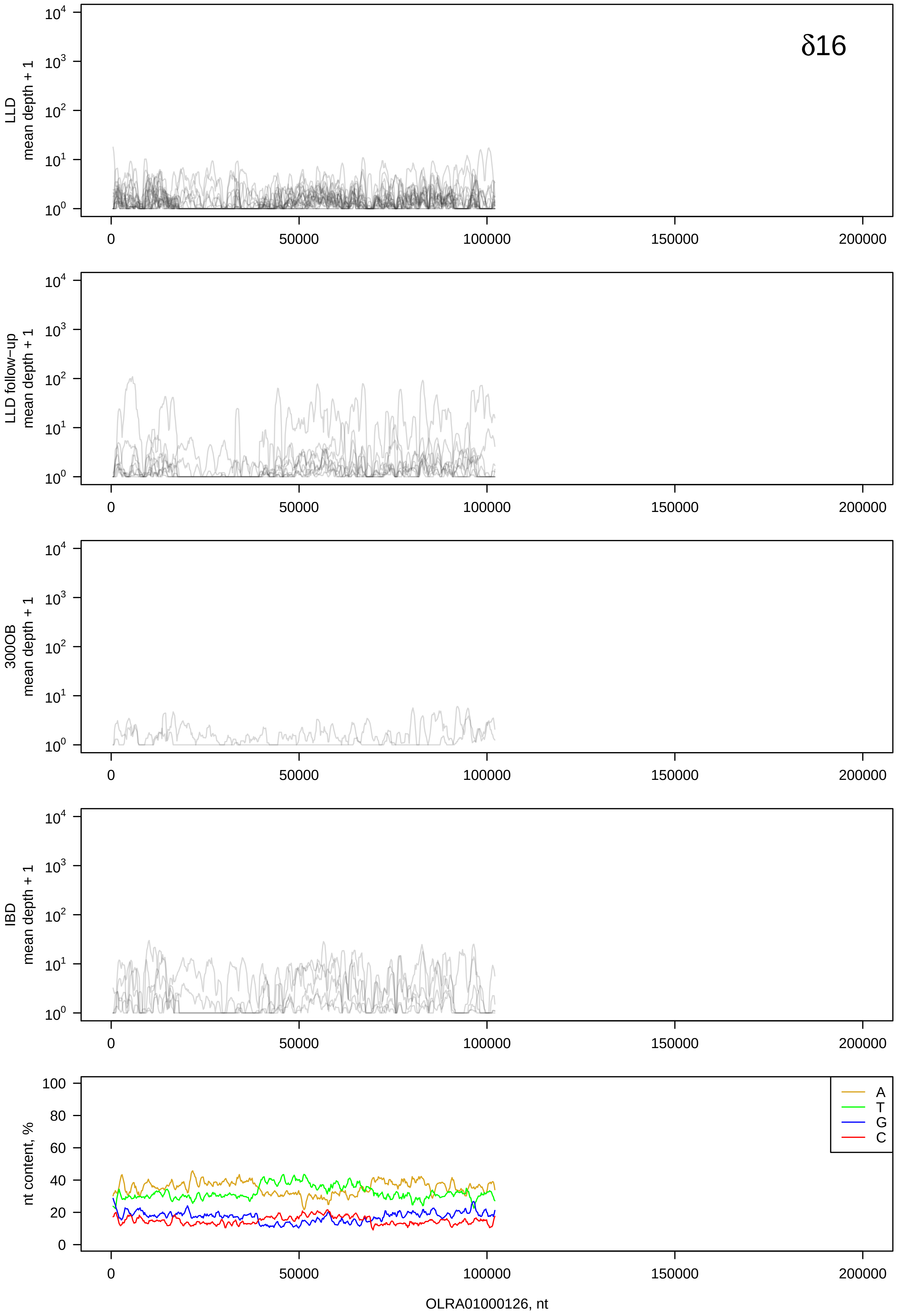
δ16
LLD
mean depth + 1
LLD follow−up
mean depth + 1
300OB
mean depth + 1
IBD
mean depth + 1
nt content, %
A
T
G
C
0
50000
100000
150000
200000
OLRA01000126, nt

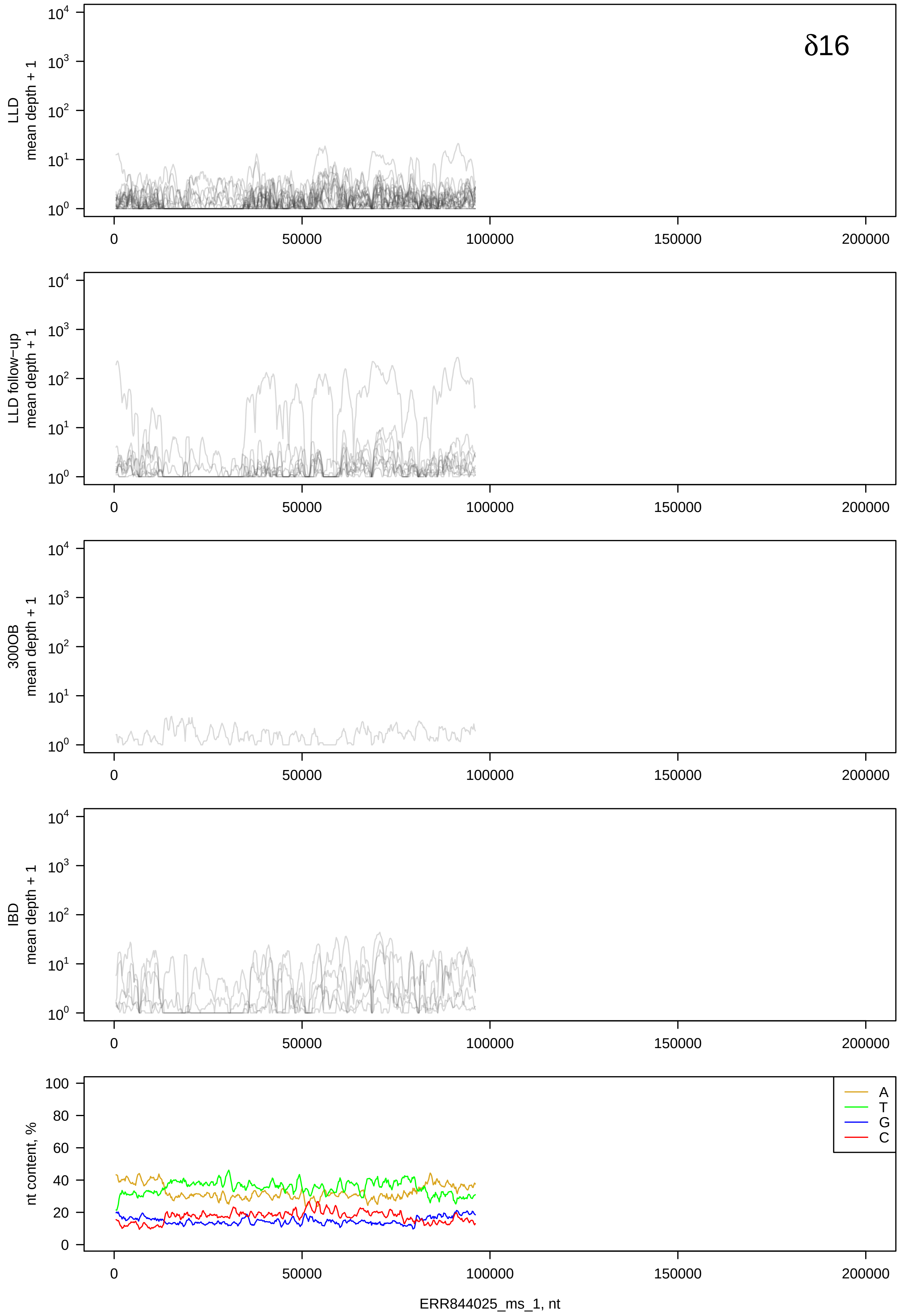

δ16
LLD
mean depth + 1
LLD follow-up
mean depth + 1
300OB
mean depth + 1
IBD
mean depth + 1
nt content, %
A
T
G
C
ERR844025_ms_1, nt

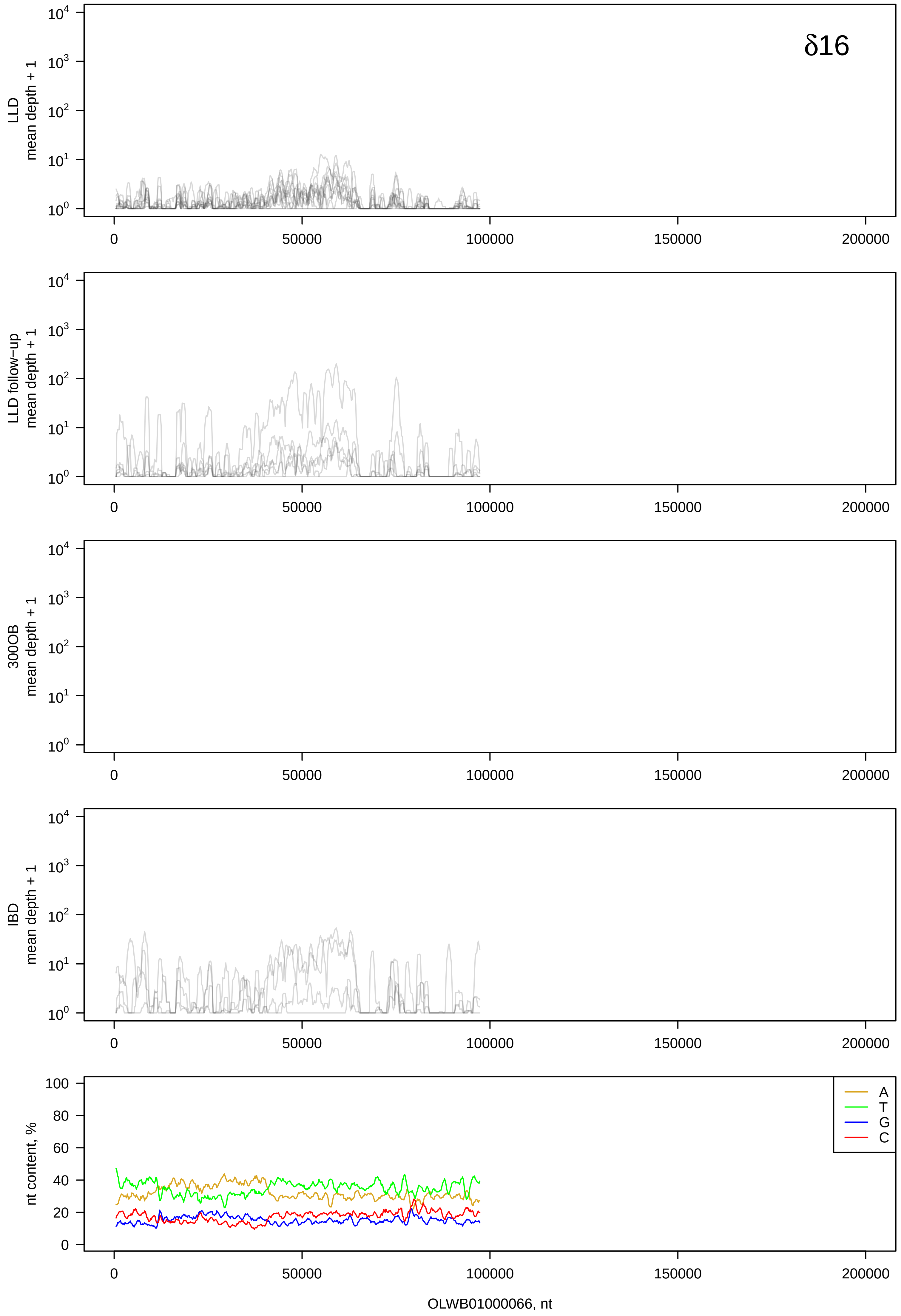

δ16
LLD
mean depth + 1
LLD follow−up
mean depth + 1
300OB
mean depth + 1
IBD
mean depth + 1
nt content, %
A
T
G
C
0
50000
100000
150000
200000
OLWB01000066, nt

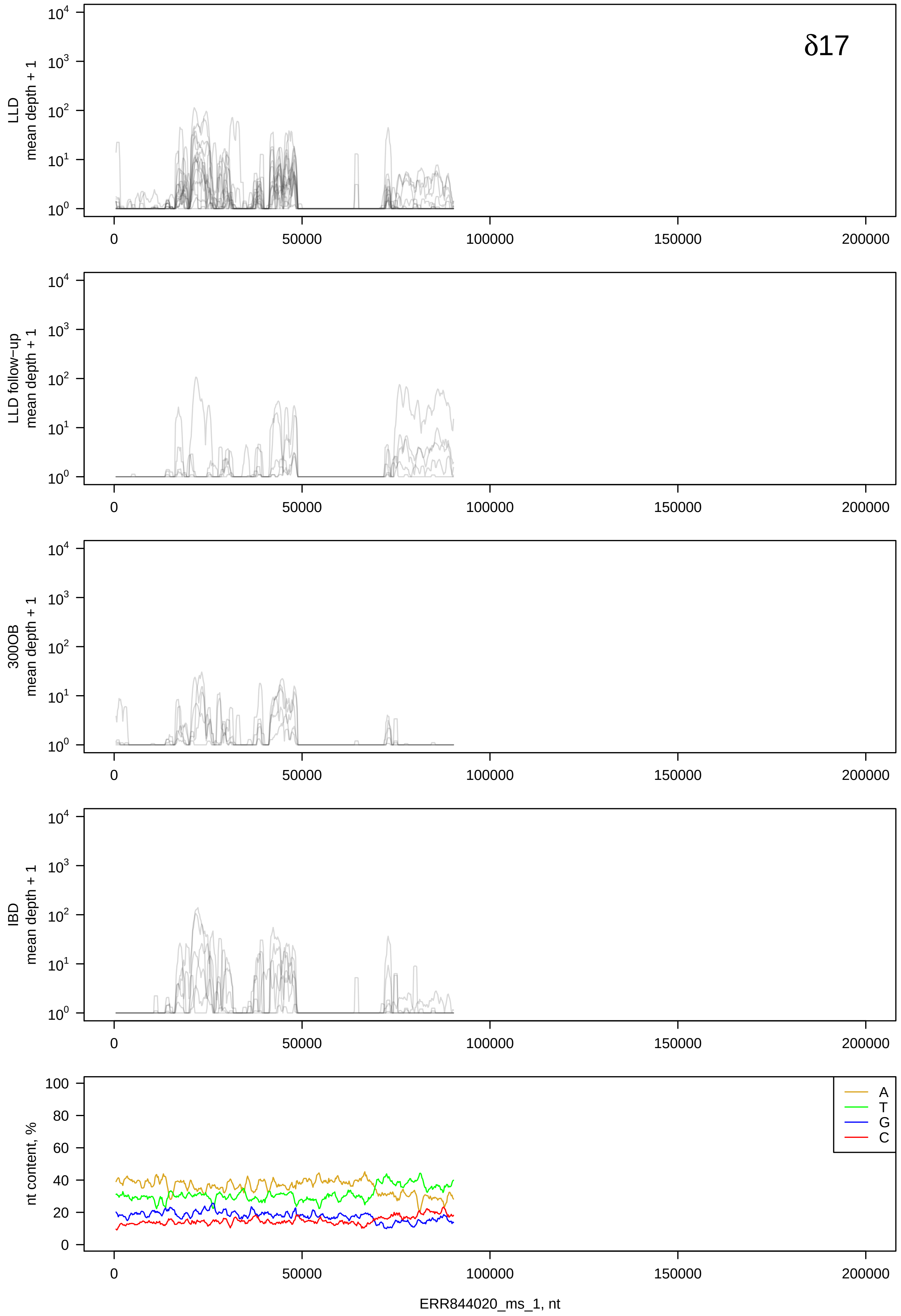

δ17
LLD mean depth + 1
LLD follow−up mean depth + 1
300OB mean depth + 1
IBD mean depth + 1
nt content, %
A
T
G
C
ERR844020_ms_1, nt

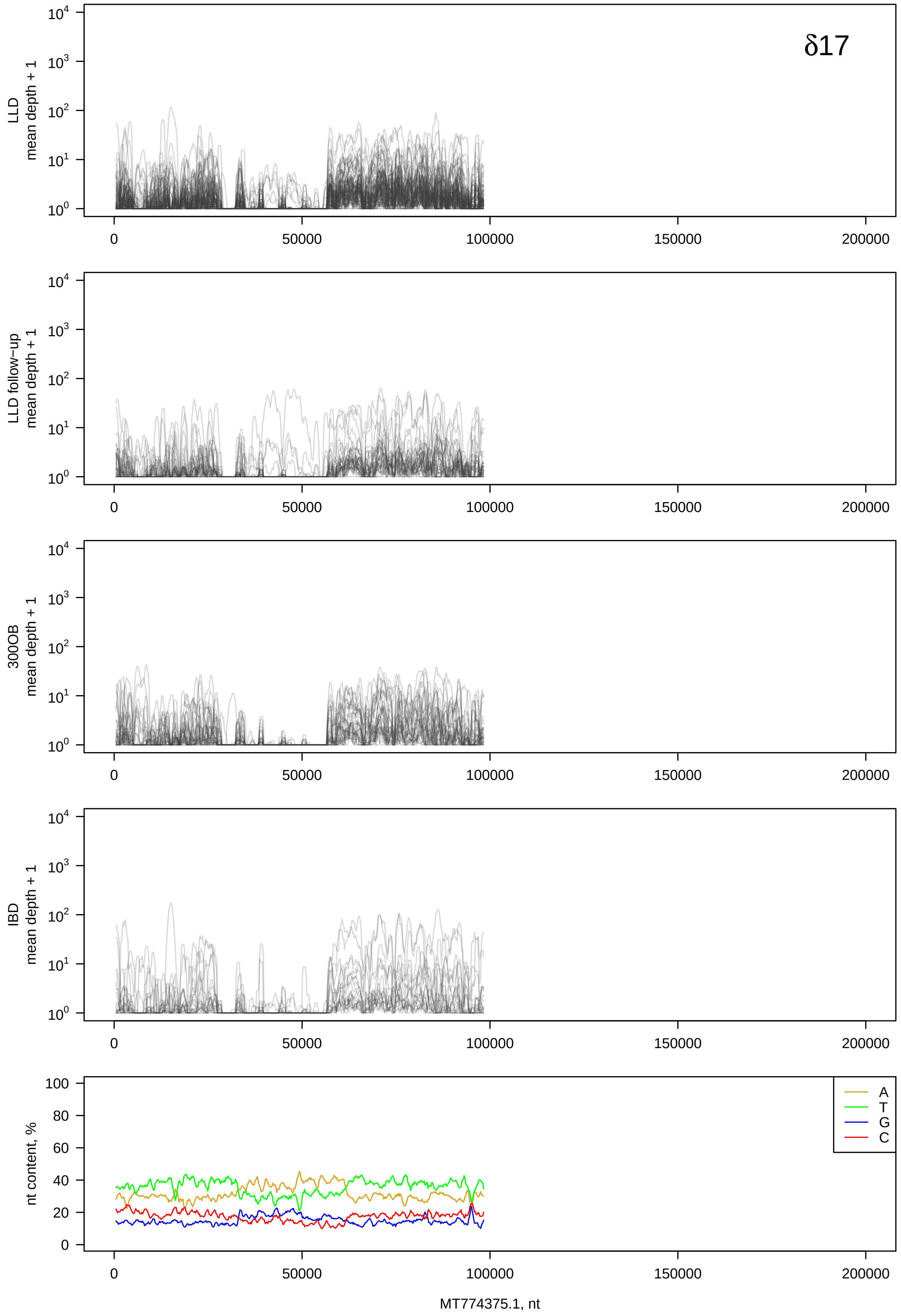

δ17
LLD
mean depth + 1
LLD follow-up
mean depth + 1
300OB
mean depth + 1
IBD
mean depth + 1
nt content, %
A
T
G
C
MT774375.1, nt

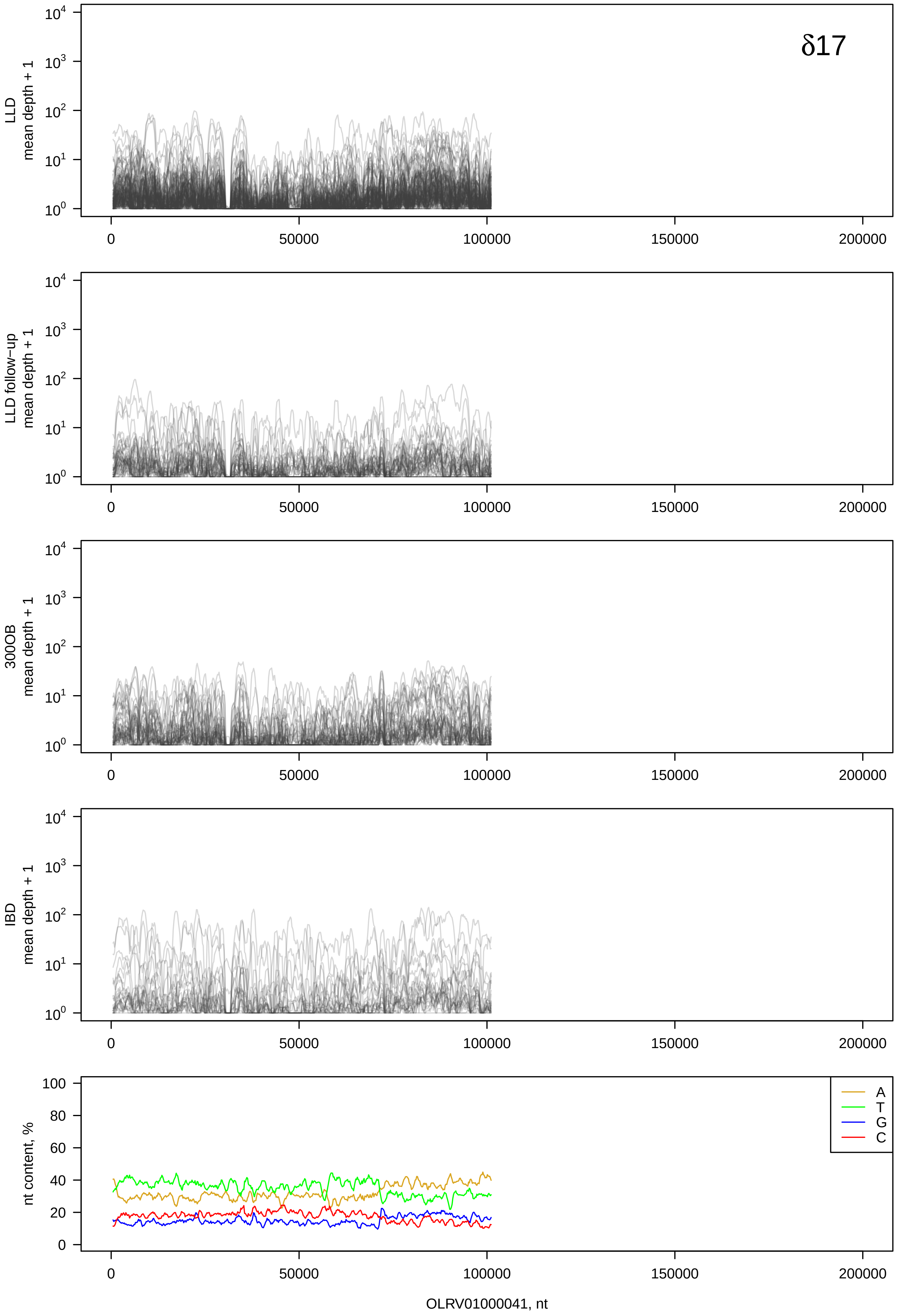
δ17
LLD
mean depth + 1
LLD follow−up
mean depth + 1
300OB
mean depth + 1
IBD
mean depth + 1
nt content, %
A
T
G
C
OLRV01000041, nt

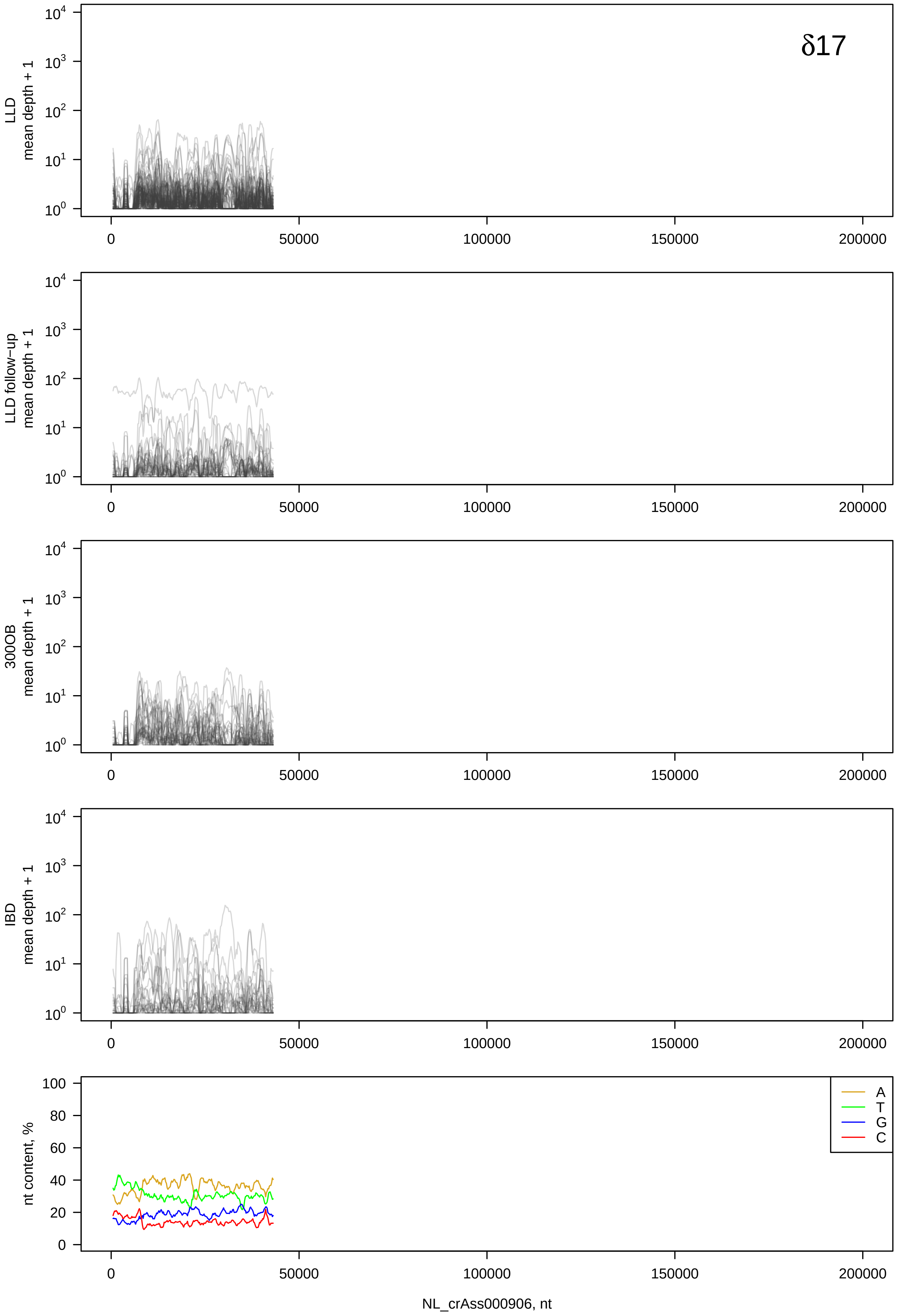

δ17
LLD mean depth + 1
LLD follow−up mean depth + 1
300OB mean depth + 1
IBD mean depth + 1
nt content, %
A
T
G
C
NL_crAss000906, nt

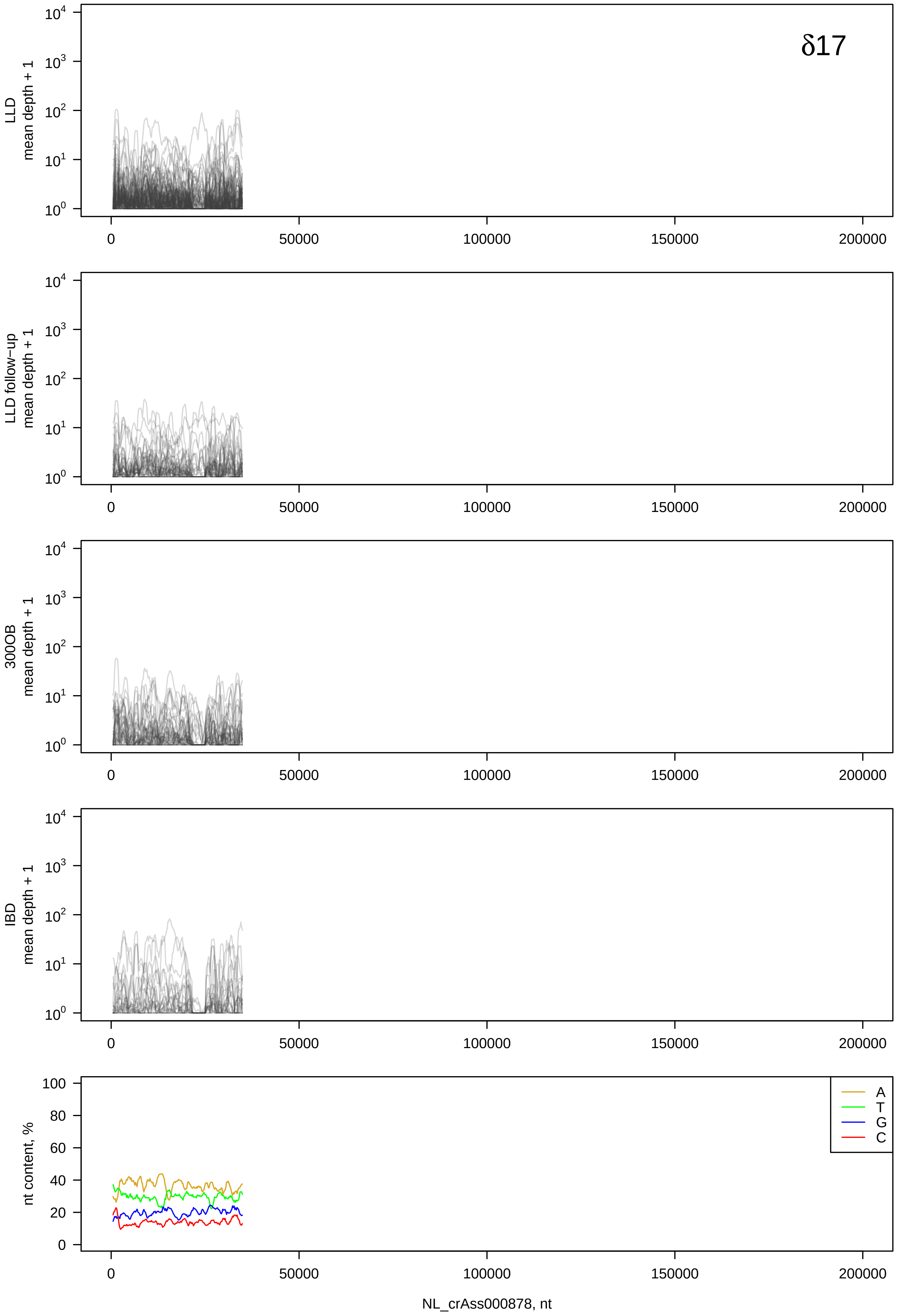
δ17
LLD
mean depth + 1
LLD follow-up
mean depth + 1
300OB
mean depth + 1
IBD
mean depth + 1
nt content, %
A
T
G
C
NL_crAss000878, nt

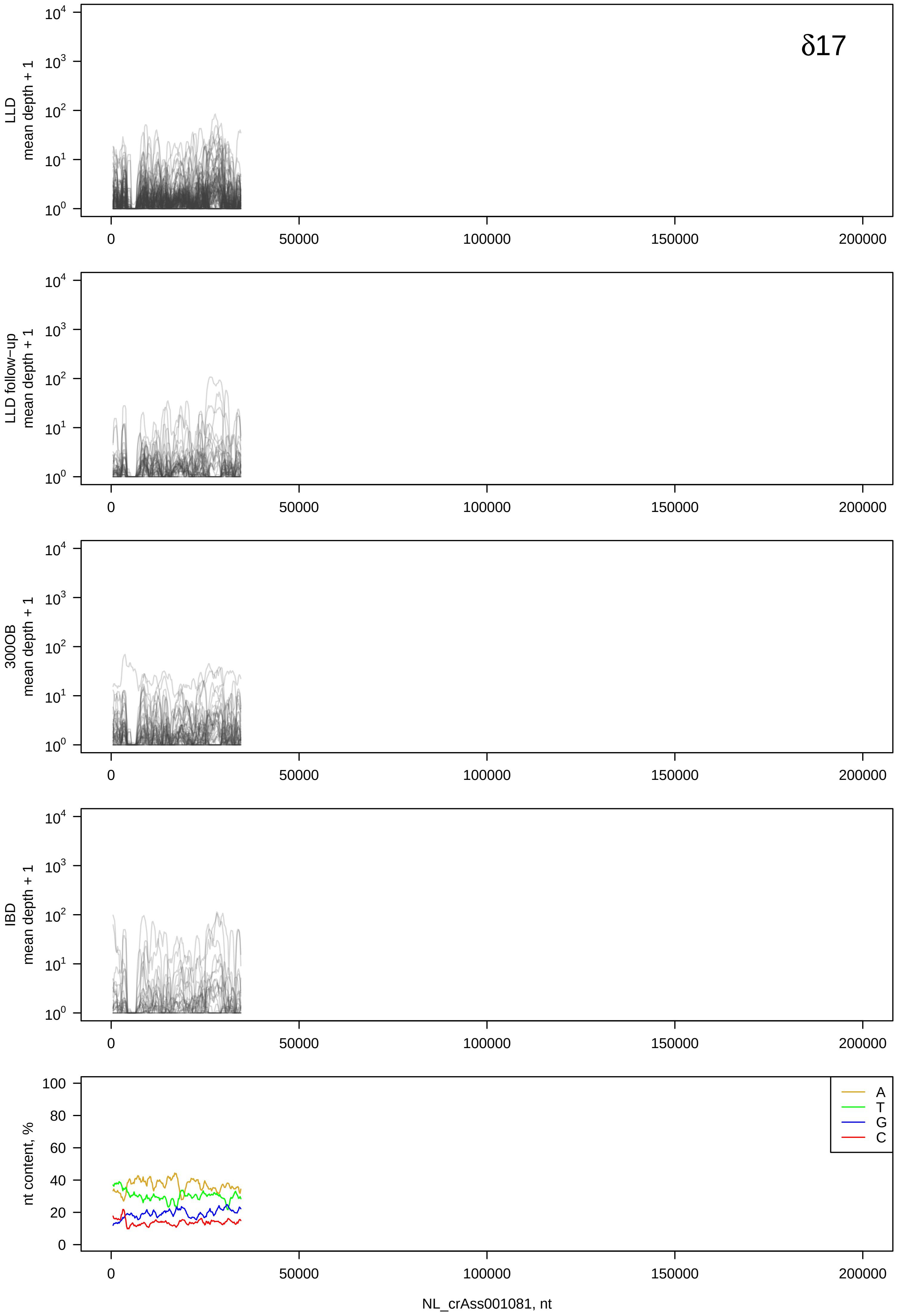

δ17
LLD
mean depth + 1
LLD follow−up
mean depth + 1
300OB
mean depth + 1
IBD
mean depth + 1
nt content, %
A
T
G
C
0
50000
100000
150000
200000
NL_crAss001081, nt

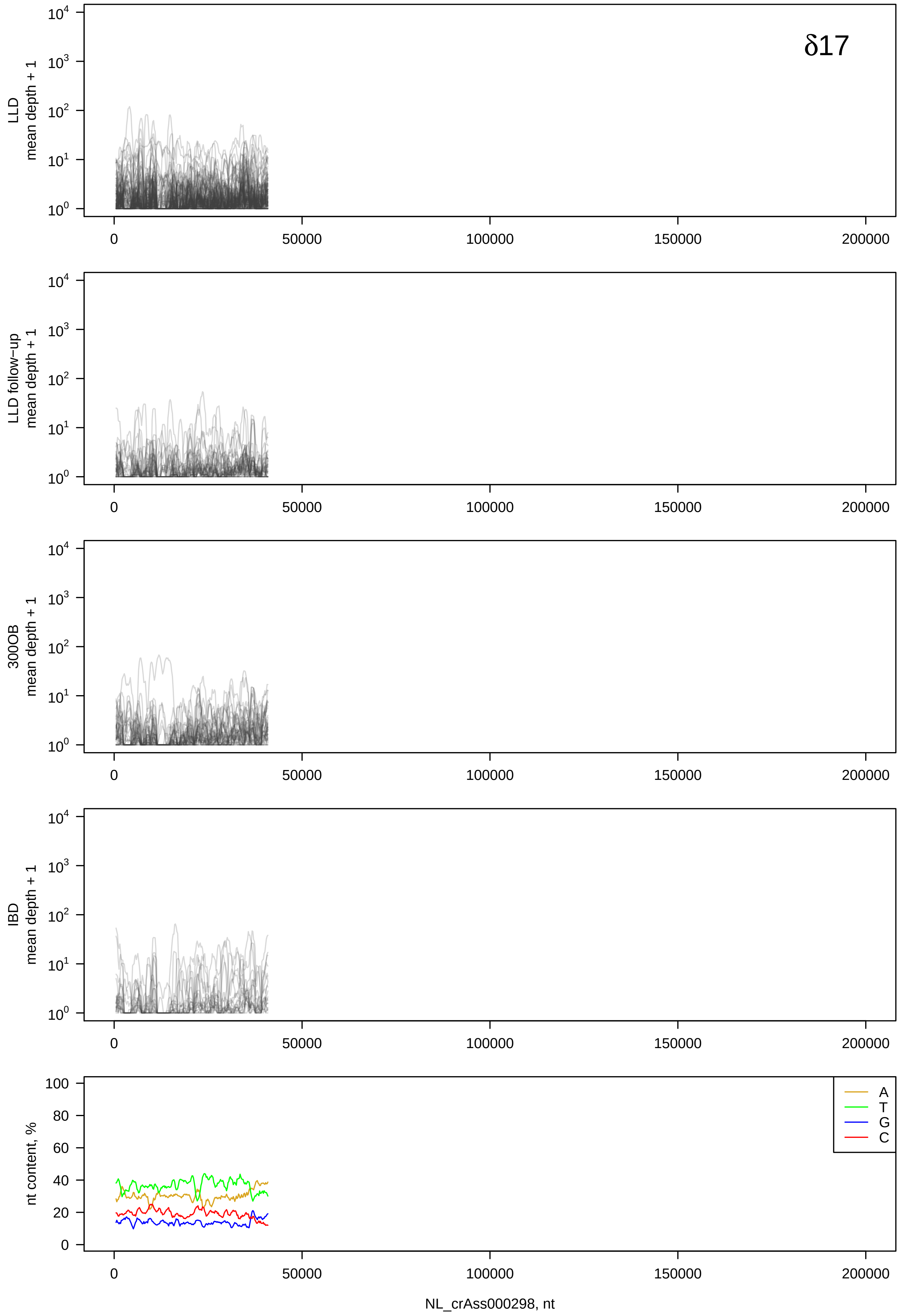

δ17
LLD
mean depth + 1
LLD follow-up
mean depth + 1
300OB
mean depth + 1
IBD
mean depth + 1
nt content, %
A
T
G
C
0
50000
100000
150000
200000
NL_crAss000298, nt

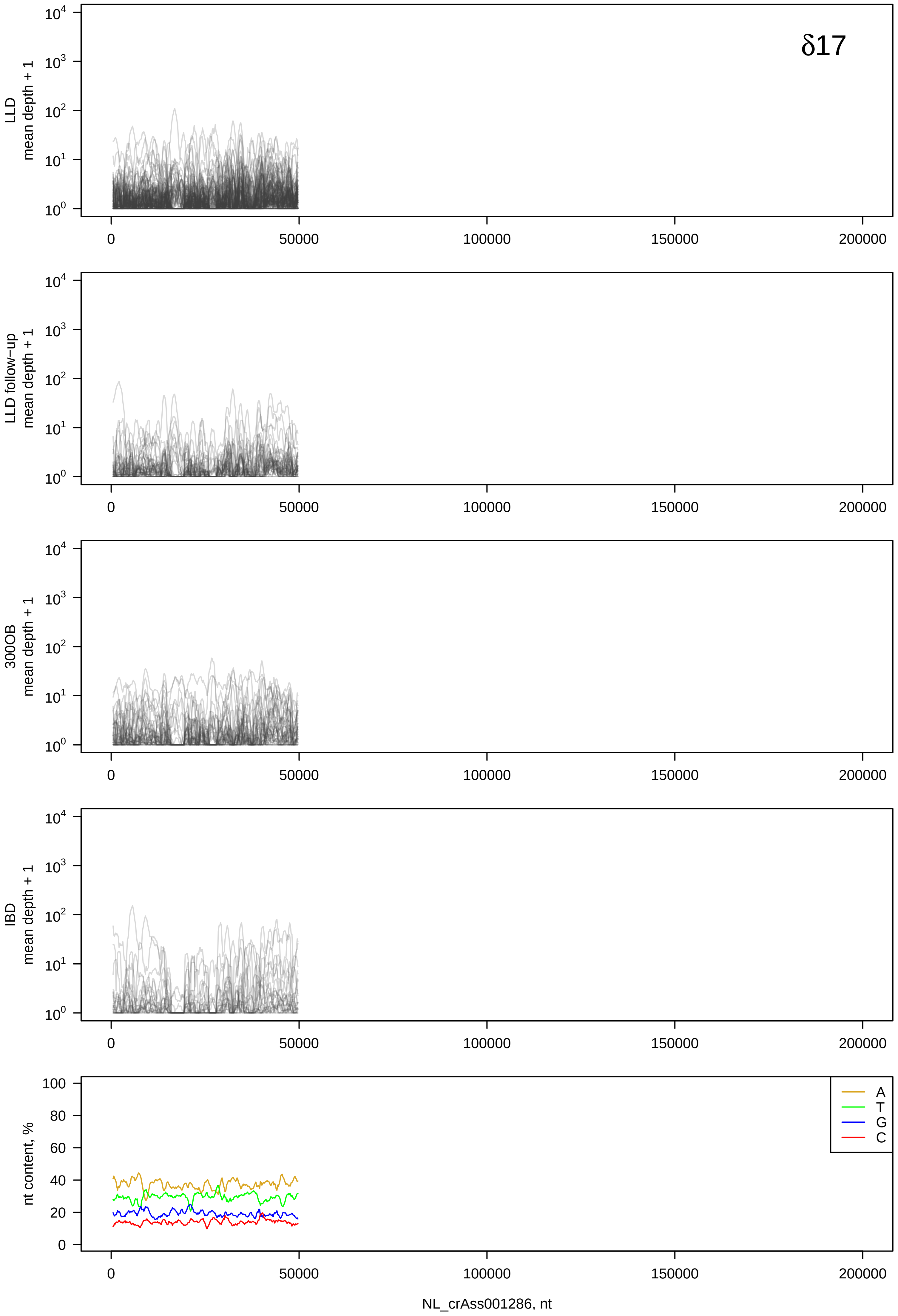
δ17
LLD mean depth + 1
LLD follow-up mean depth + 1
300OB mean depth + 1
IBD mean depth + 1
nt content, %
A
T
G
C
0
50000
100000
150000
200000
NL_crAss001286, nt

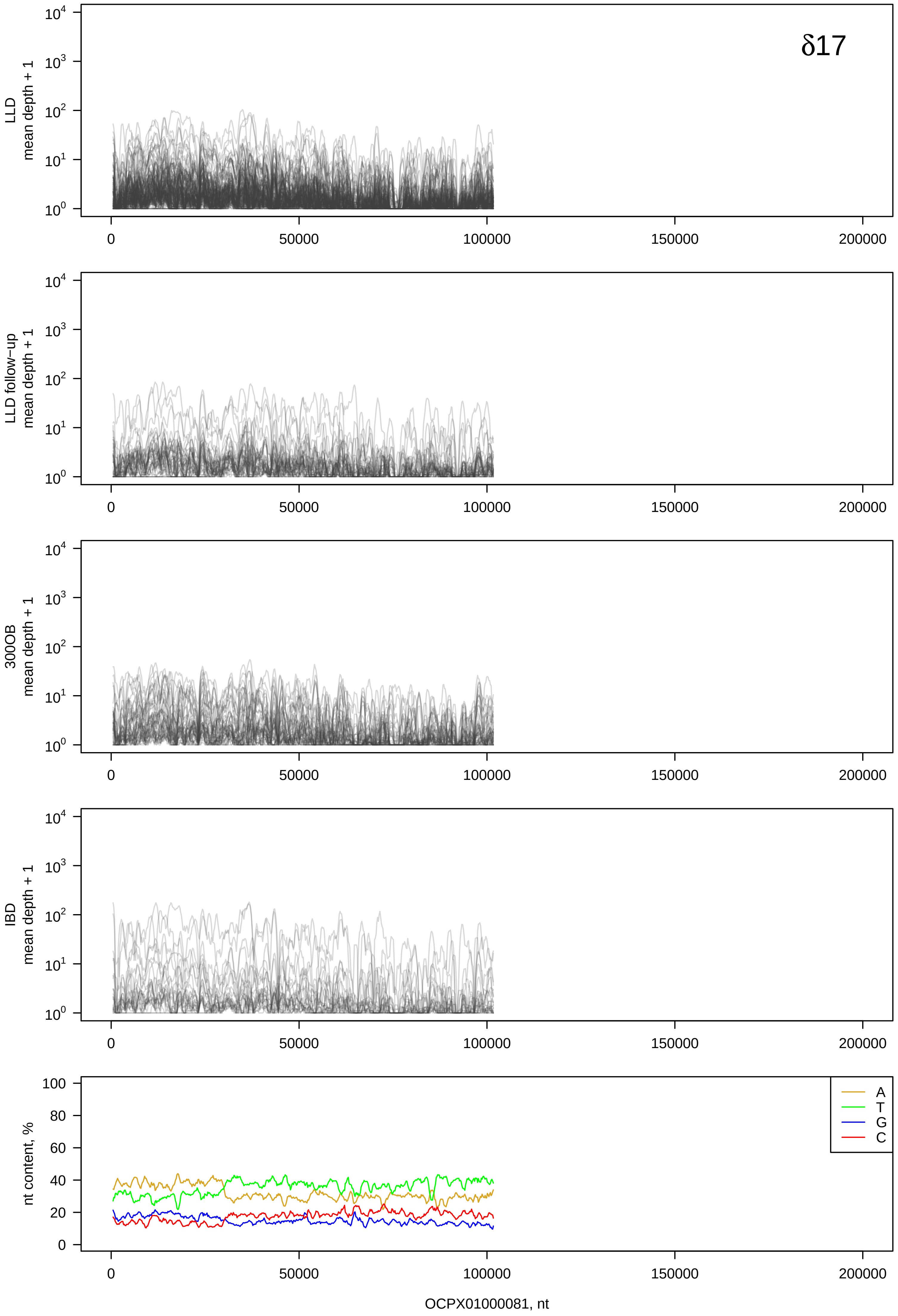
δ17
LLD mean depth + 1
LLD follow−up mean depth + 1
300OB mean depth + 1
IBD mean depth + 1
nt content, %
A
T
G
C
OCPX01000081, nt

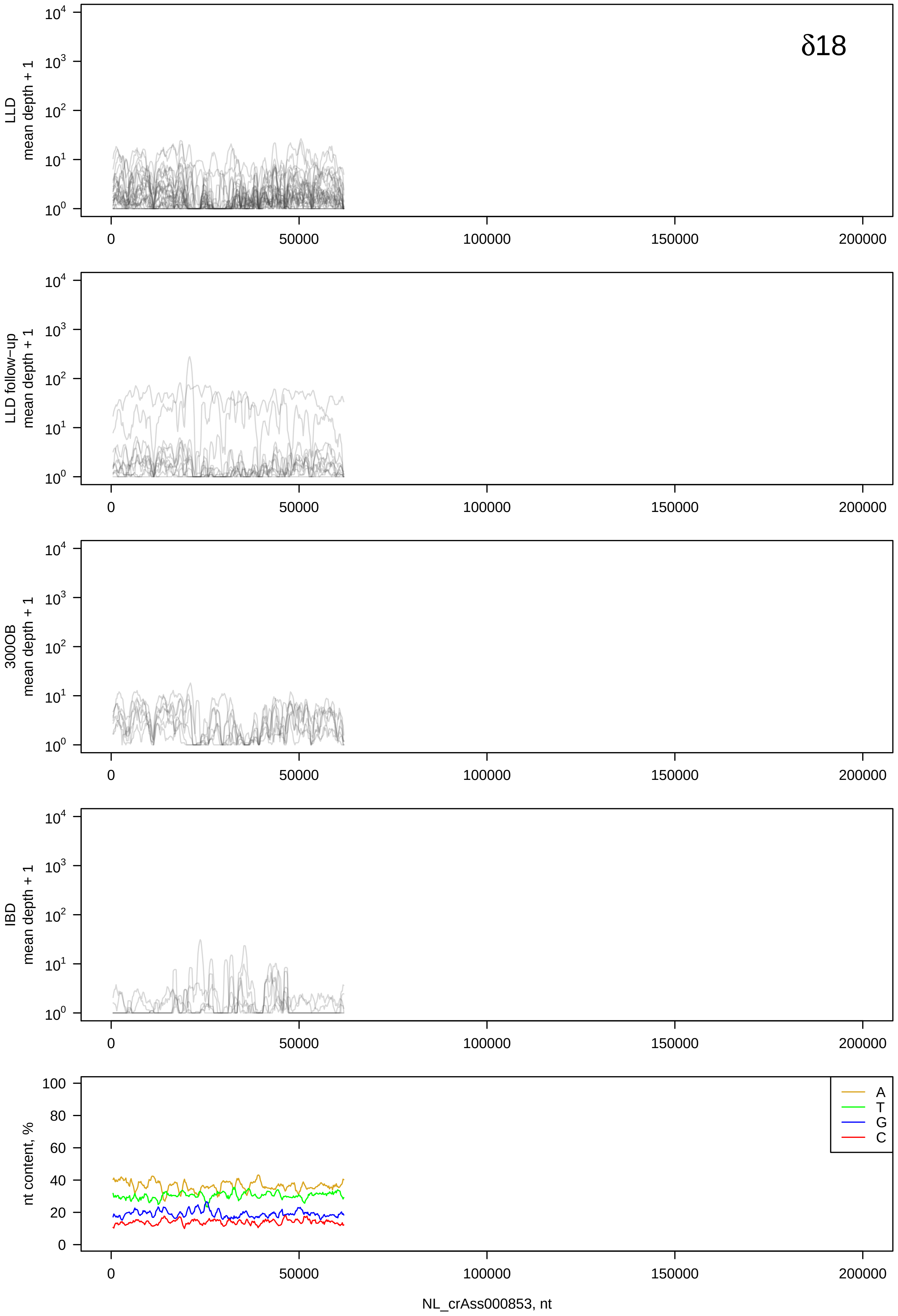

δ18
LLD
mean depth + 1
LLD follow-up
mean depth + 1
300OB
mean depth + 1
IBD
mean depth + 1
nt content, %
A
T
G
C
0
50000
100000
150000
200000
NL_crAss000853, nt

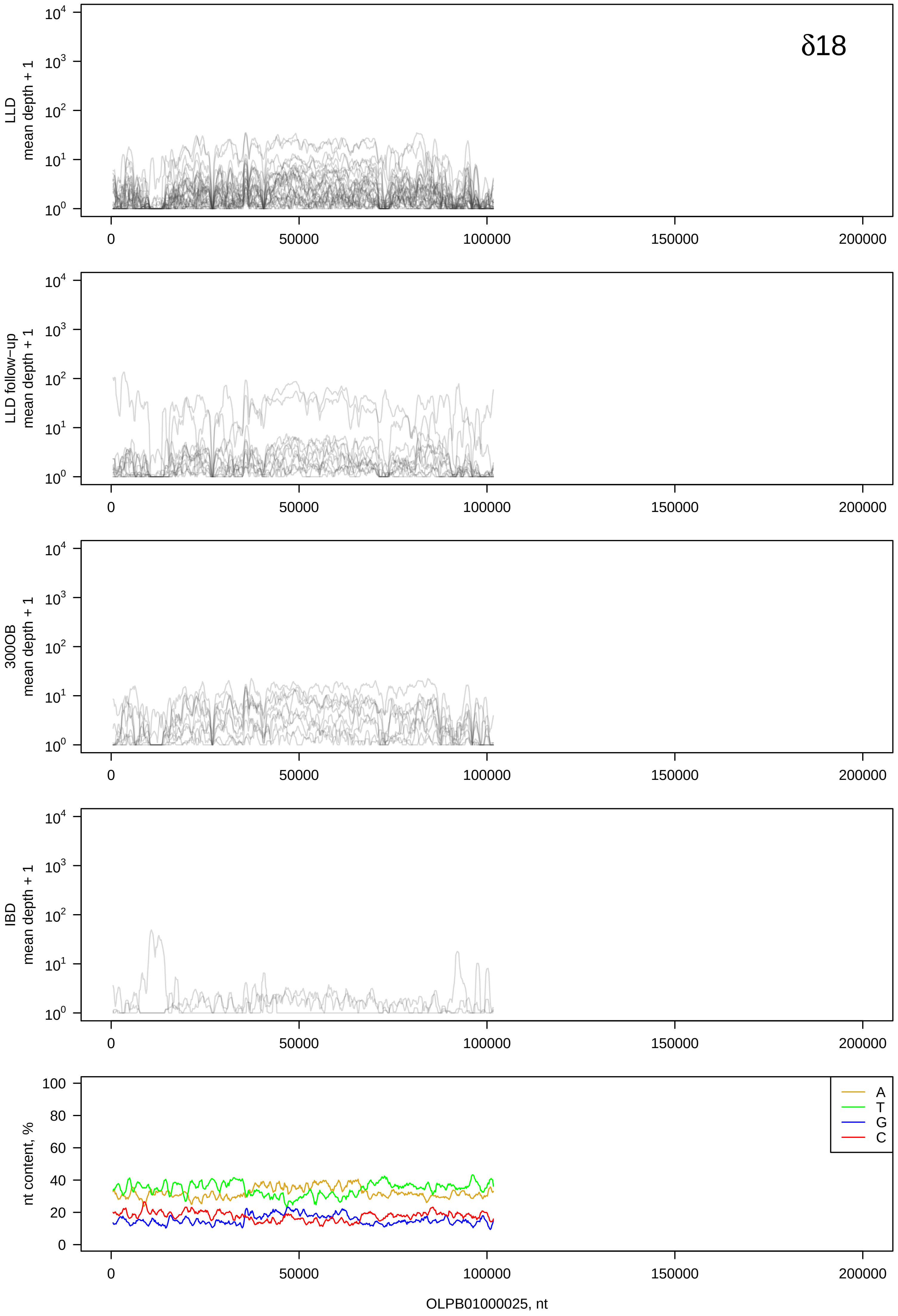

δ18
LLD
mean depth + 1
LLD follow−up
mean depth + 1
300OB
mean depth + 1
IBD
mean depth + 1
nt content, %
A
T
G
C
OLPB01000025, nt

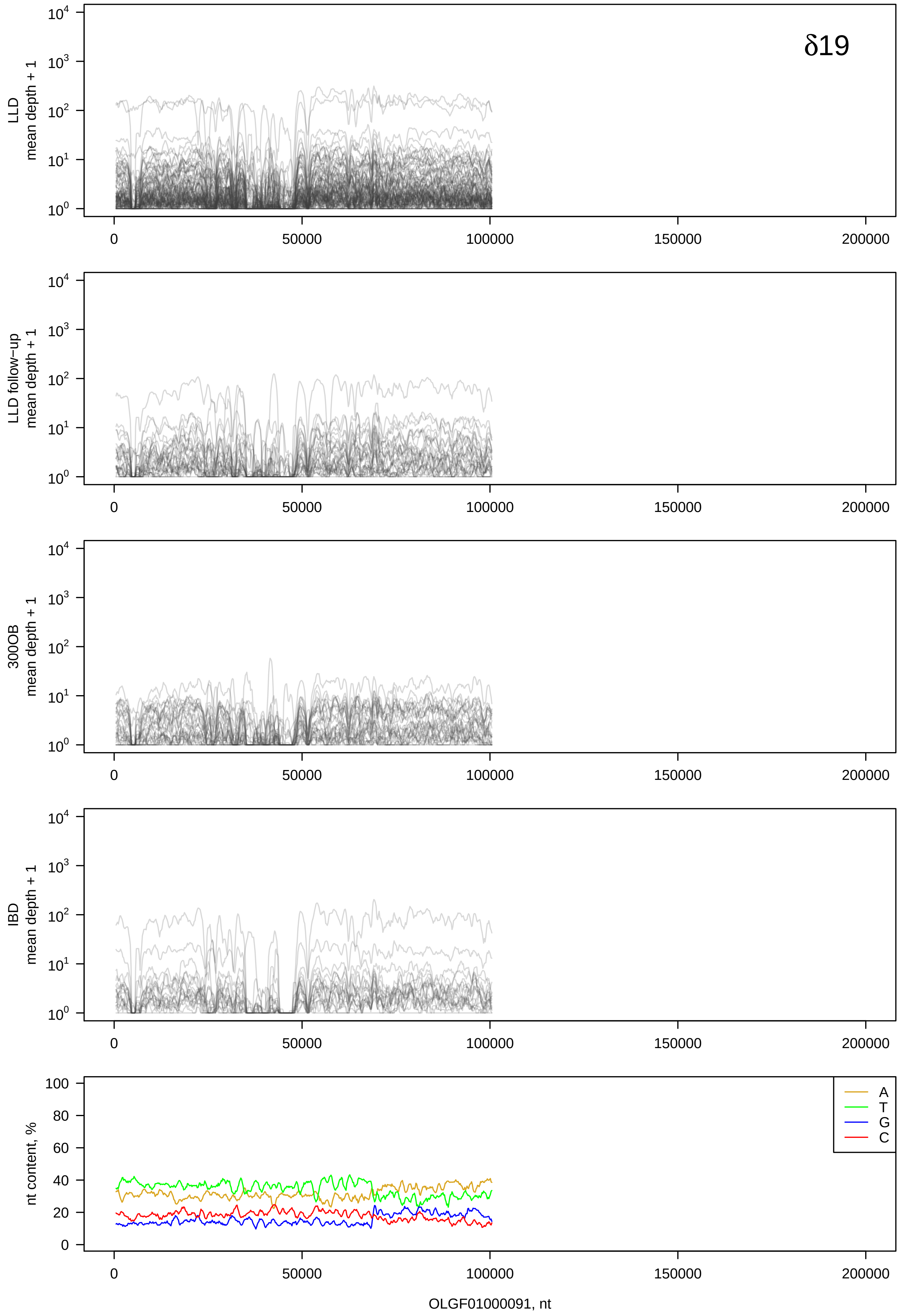
δ19
LLD
mean depth + 1
LLD follow−up
mean depth + 1
300OB
mean depth + 1
IBD
mean depth + 1
nt content, %
A
T
G
C
OLGF01000091, nt

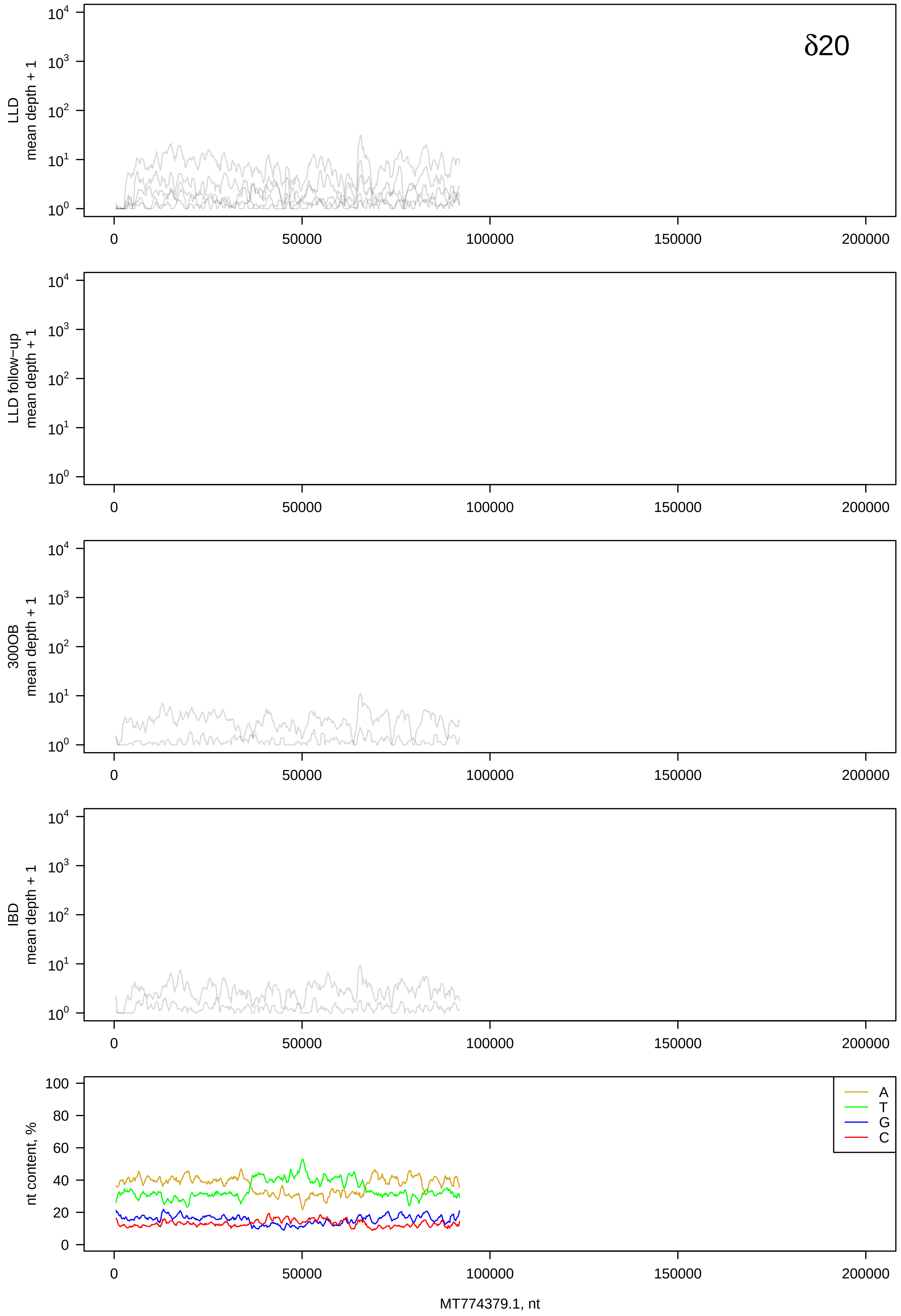

δ20
LLD
mean depth + 1
LLD follow−up
mean depth + 1
300OB
mean depth + 1
IBD
mean depth + 1
nt content, %
A
T
G
C
0
50000
100000
150000
200000
MT774379.1, nt

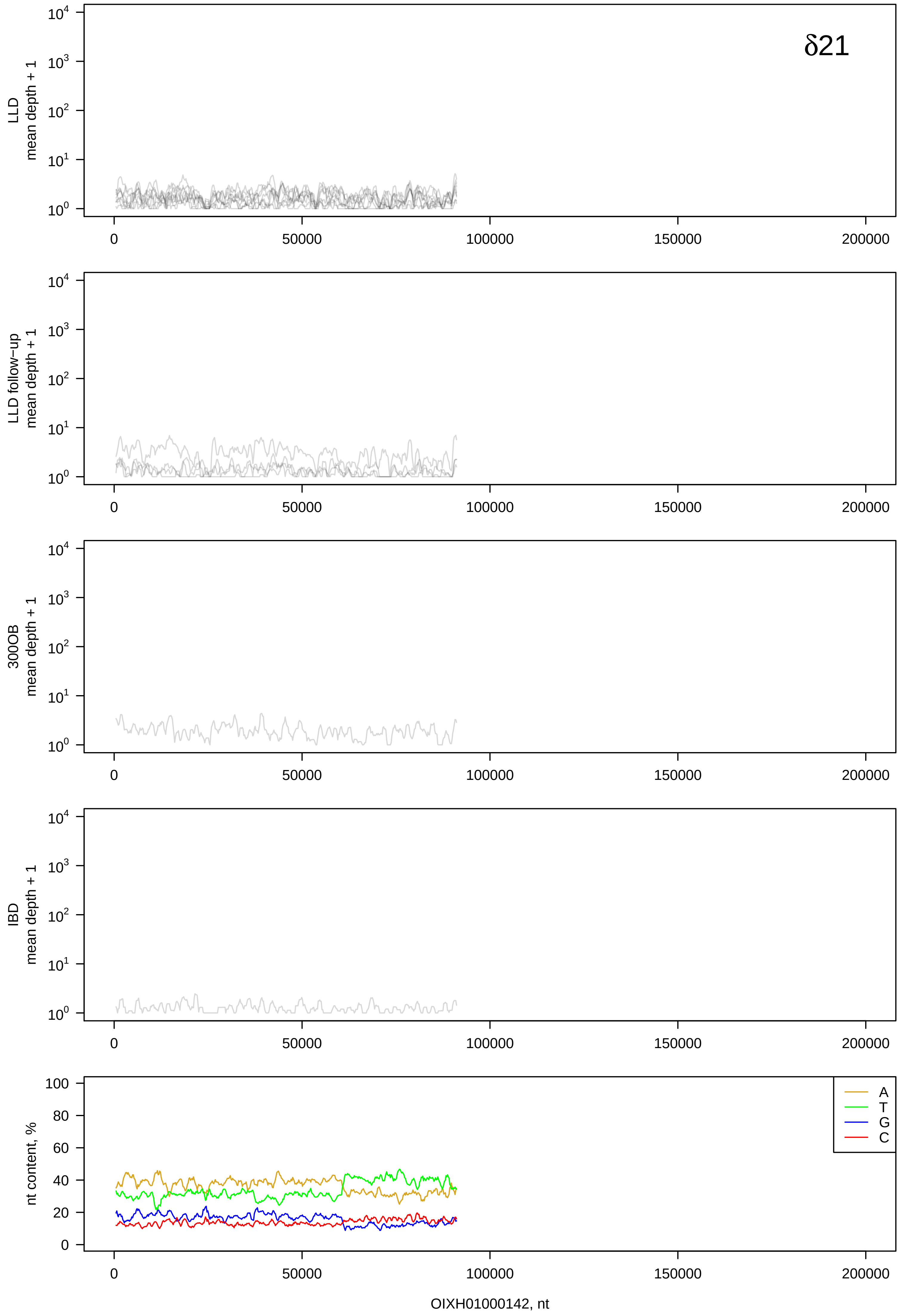
δ21
LLD
mean depth + 1
LLD follow−up
mean depth + 1
300OB
mean depth + 1
IBD
mean depth + 1
nt content, %
A
T
G
C
0
50000
100000
150000
200000
OIXH01000142, nt

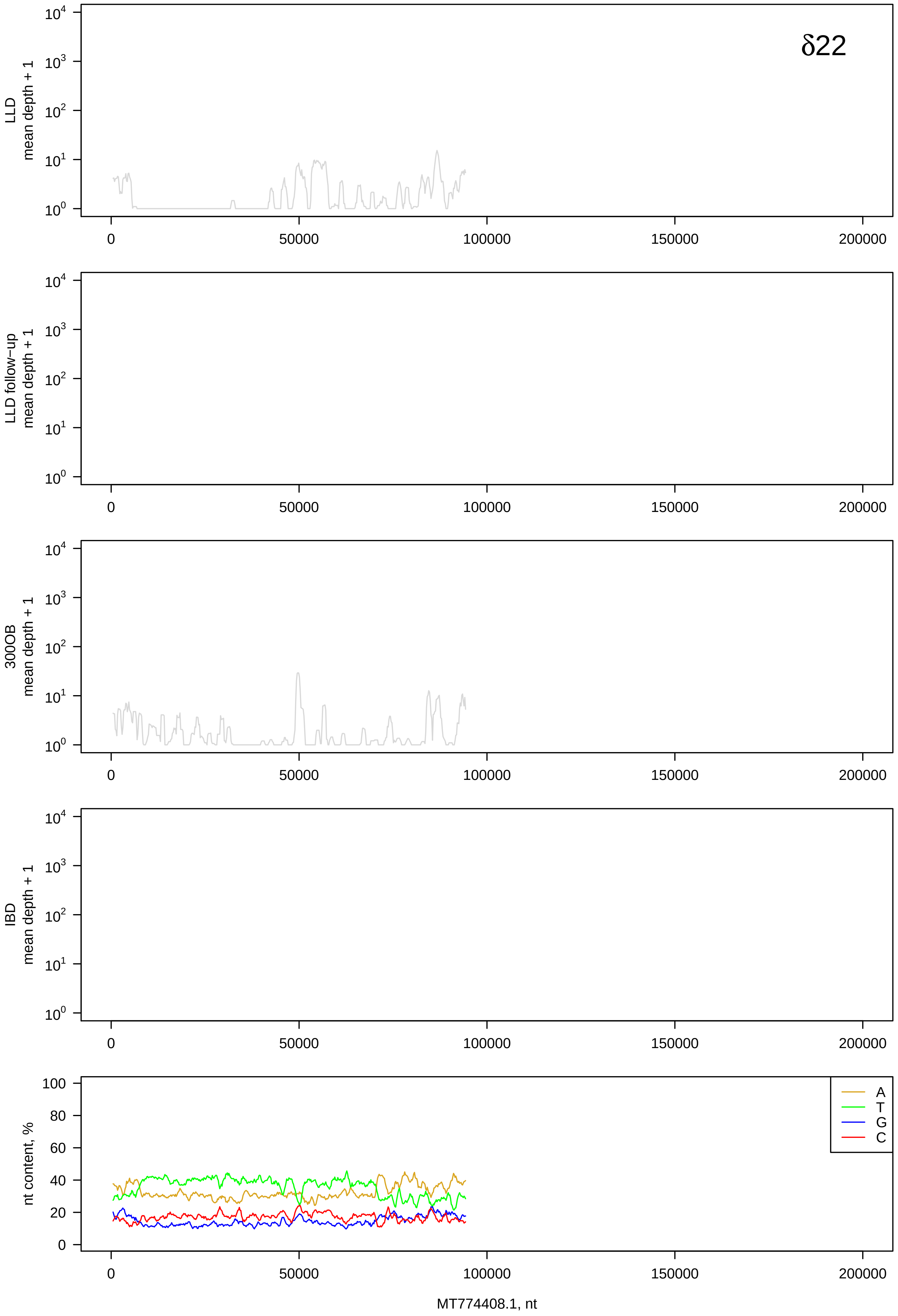
δ22
LLD mean depth + 1
LLD follow-up mean depth + 1
300OB mean depth + 1
IBD mean depth + 1
nt content, %
A
T
G
C
MT774408.1, nt

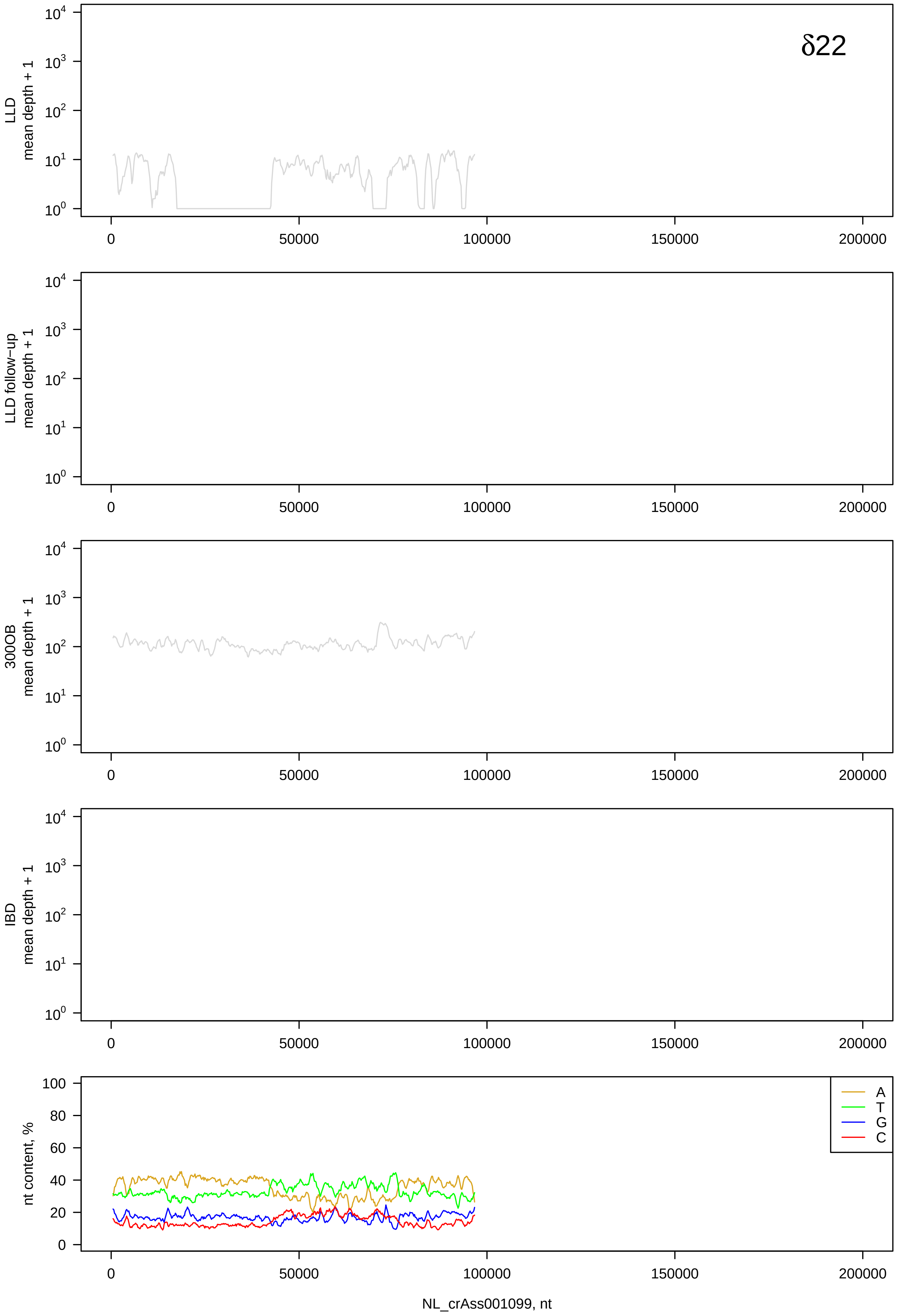

δ22
LLD
mean depth + 1
LLD follow-up
mean depth + 1
300OB
mean depth + 1
IBD
mean depth + 1
$10^0$
$10^1$
$10^2$
$10^3$
$10^4$
0
50000
100000
150000
200000
nt content, %
0
20
40
60
80
100
A
T
G
C
NL_crAss001099, nt

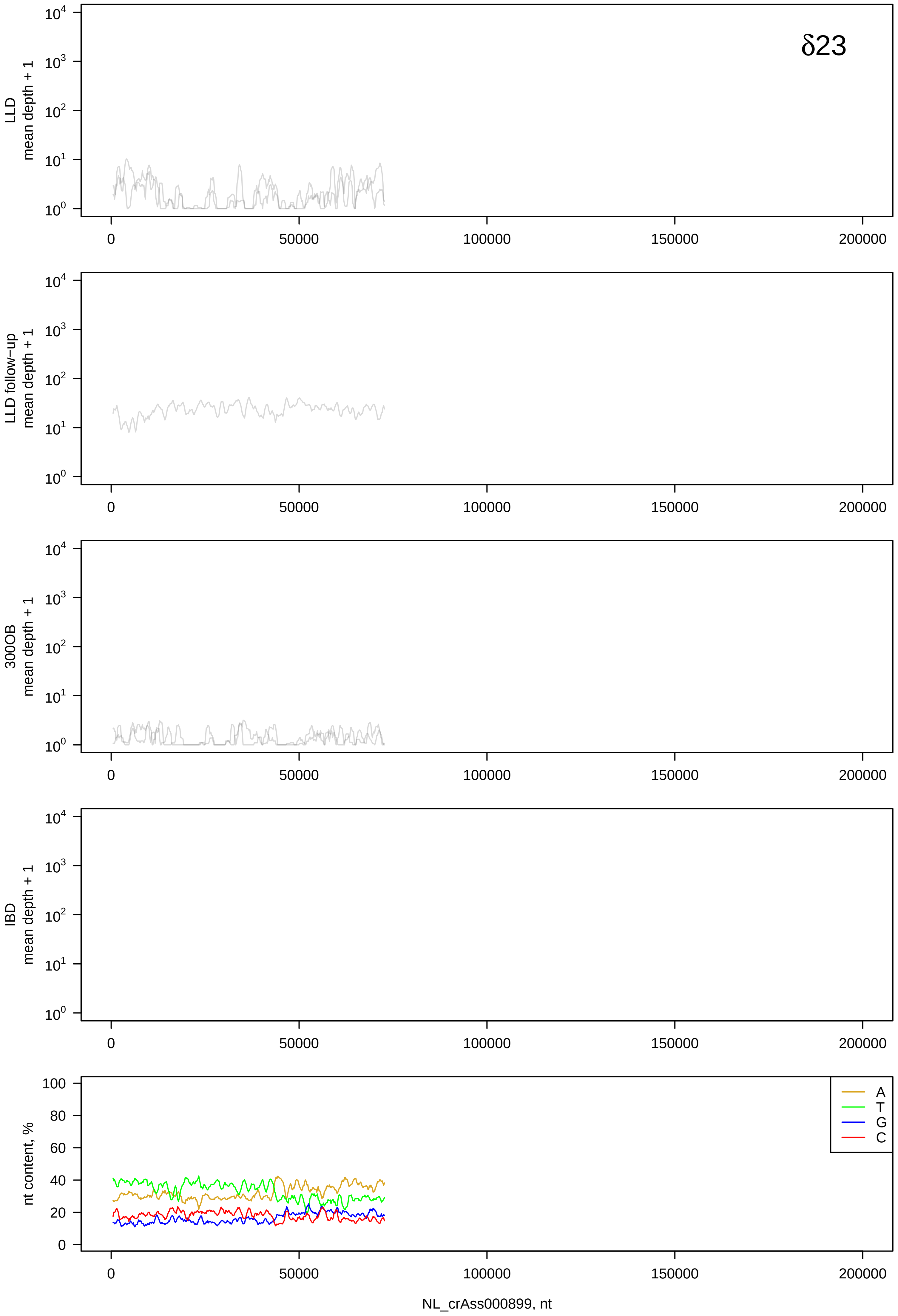
δ23
LLD mean depth + 1
LLD follow-up mean depth + 1
300OB mean depth + 1
IBD mean depth + 1
nt content, %
A
T
G
C
NL_crAss000899, nt

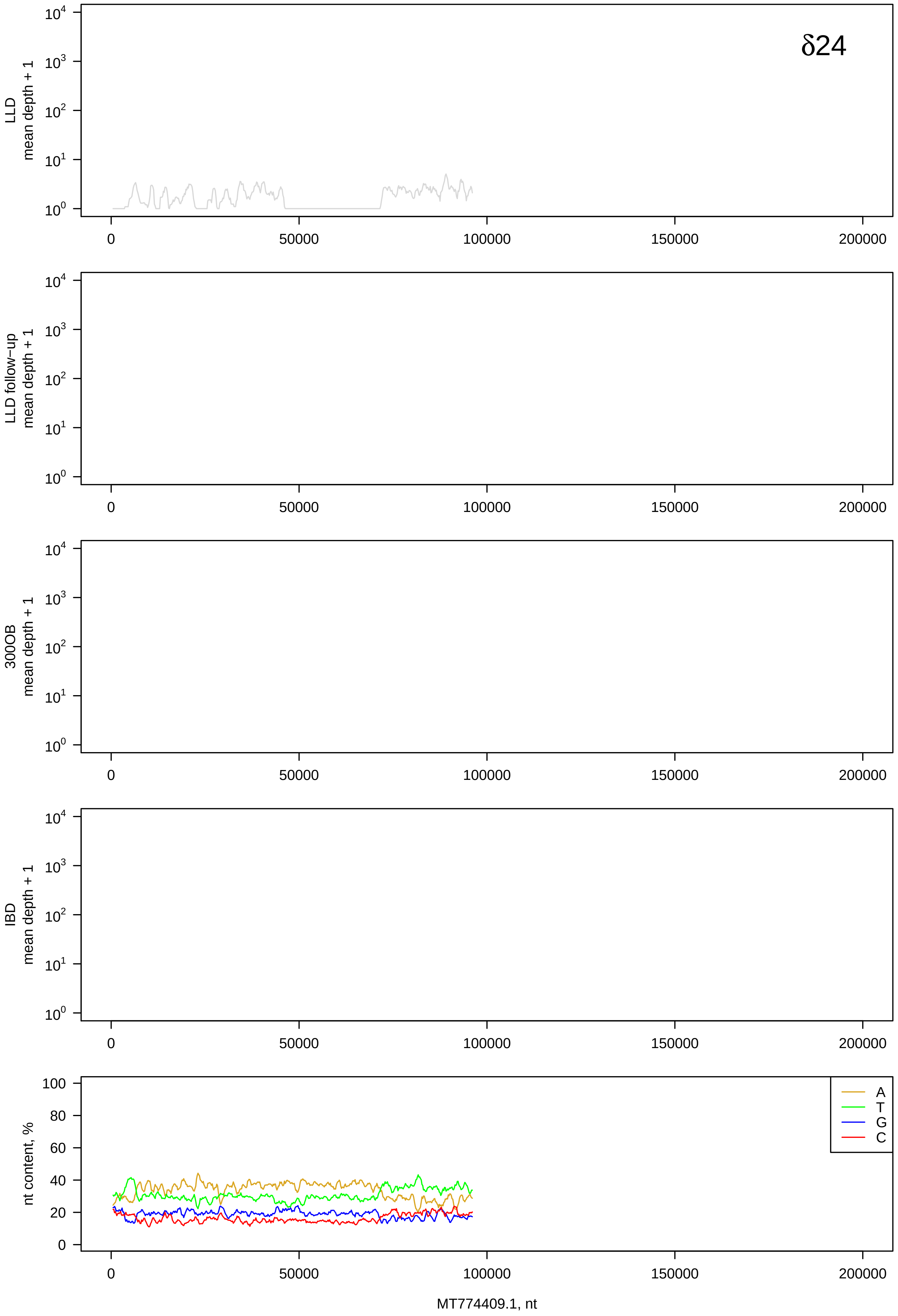
δ24
LLD
mean depth + 1
LLD follow−up
mean depth + 1
300OB
mean depth + 1
IBD
mean depth + 1
nt content, %
A
T
G
C
MT774409.1, nt

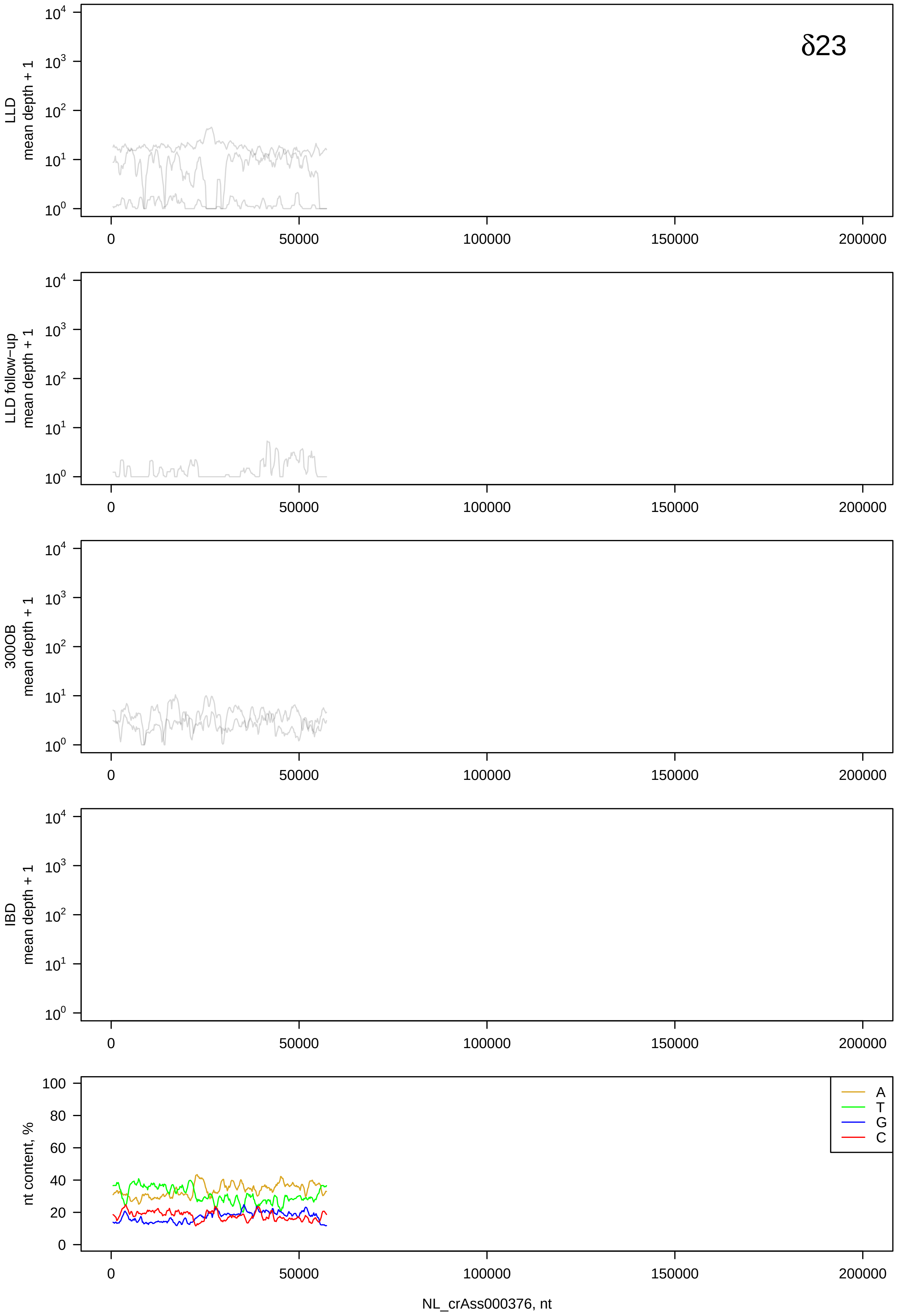

δ23
LLD
mean depth + 1
LLD follow−up
mean depth + 1
300OB
mean depth + 1
IBD
mean depth + 1
nt content, %
A
T
G
C
0
50000
100000
150000
200000
NL_crAss000376, nt

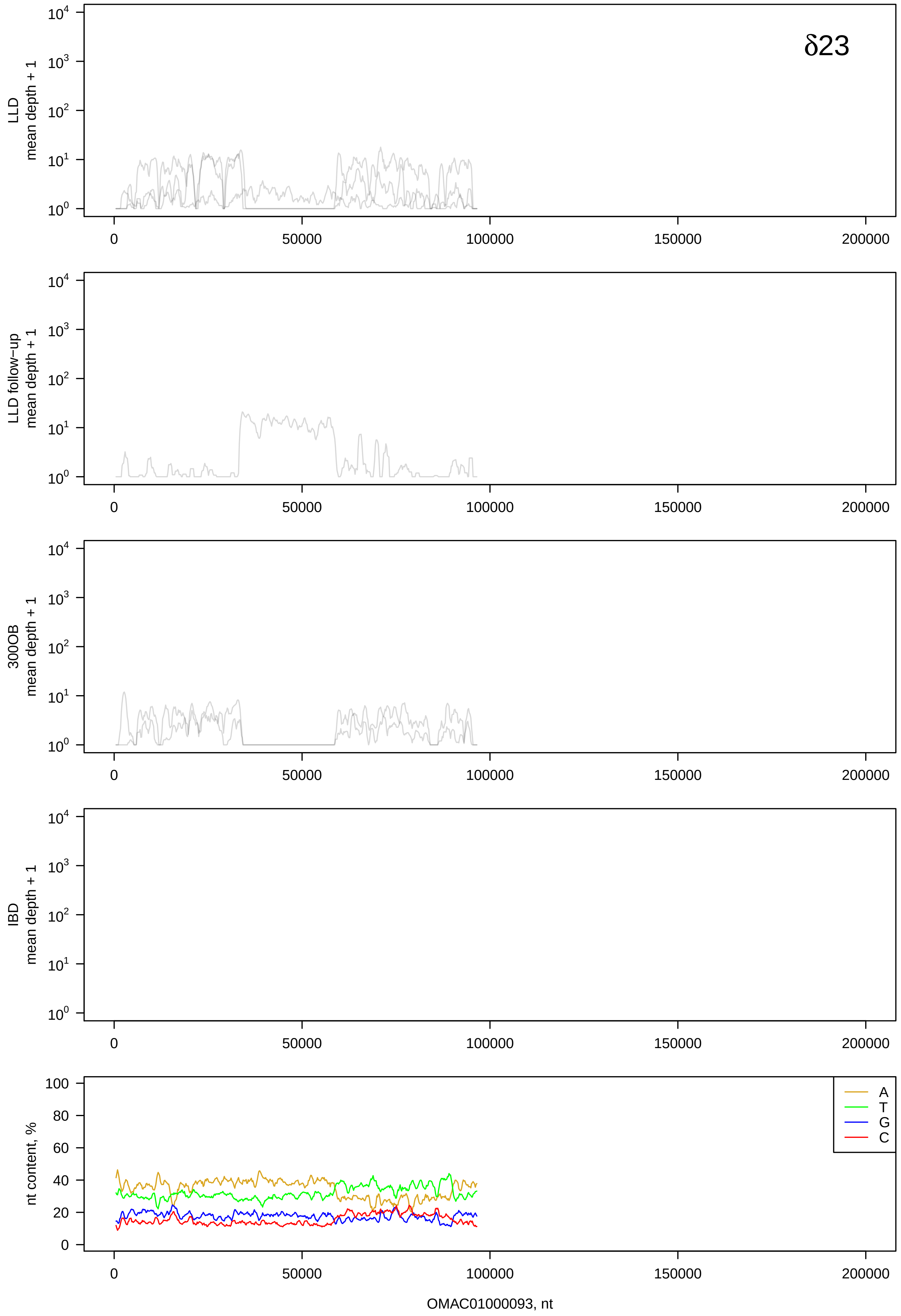
δ23
LLD mean depth + 1
LLD follow-up mean depth + 1
300OB mean depth + 1
IBD mean depth + 1
nt content, %
A
T
G
C
OMAC01000093, nt

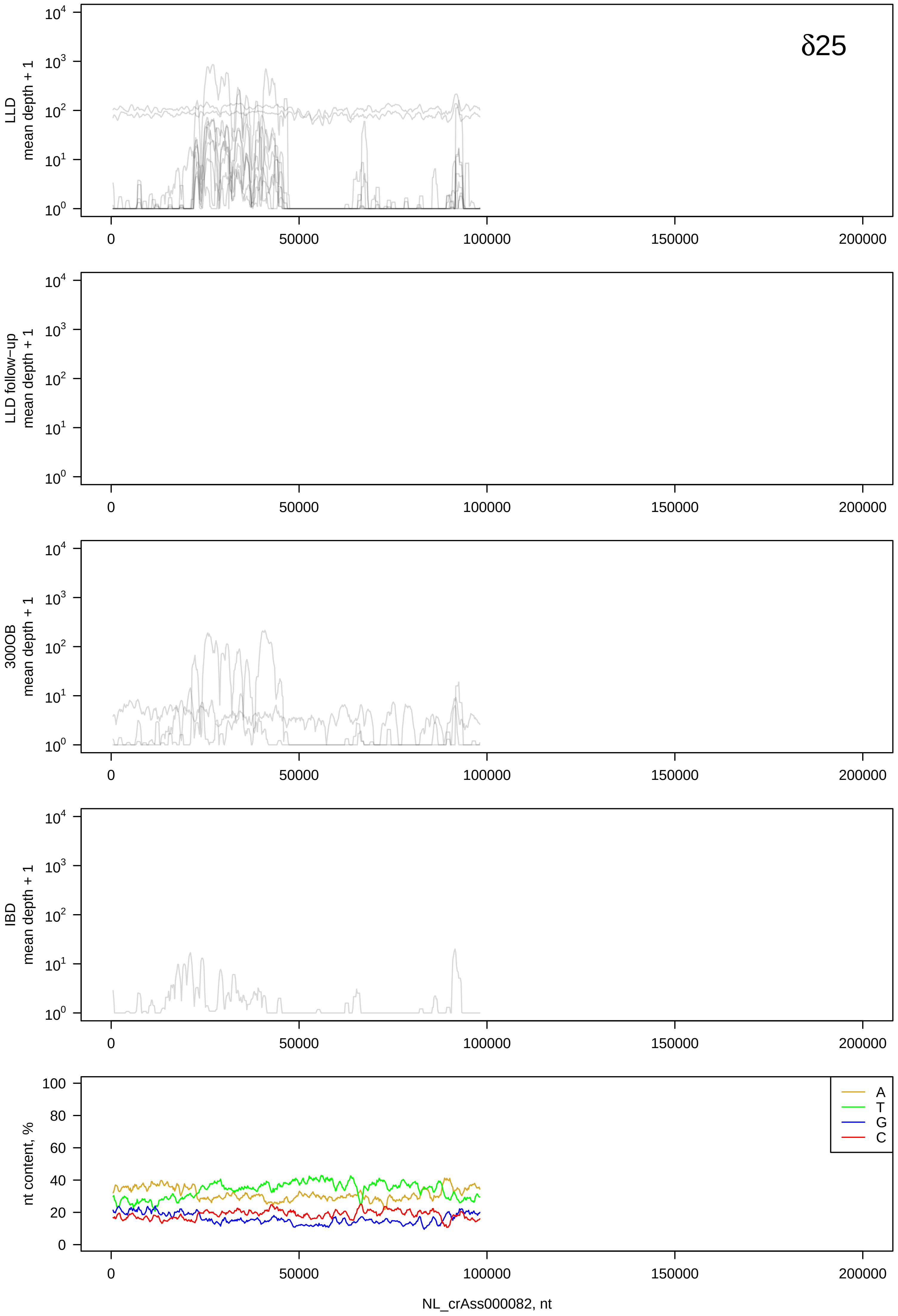

δ25
LLD
mean depth + 1
LLD follow−up
mean depth + 1
300OB
mean depth + 1
IBD
mean depth + 1
nt content, %
A
T
G
C
NL_crAss000082, nt

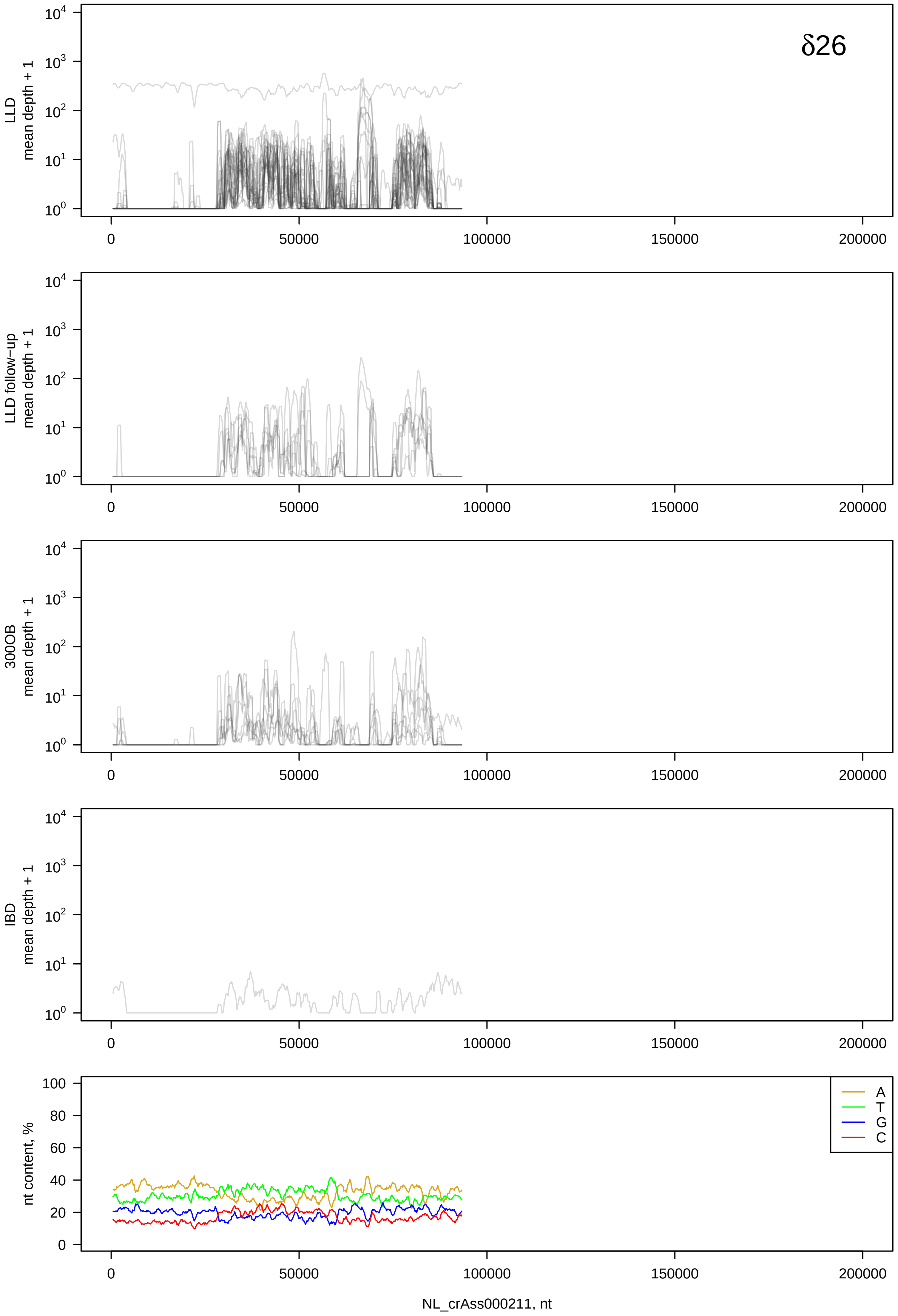

δ26
LLD
mean depth + 1
LLD follow−up
mean depth + 1
300OB
mean depth + 1
IBD
mean depth + 1
nt content, %
A
T
G
C
0
50000
100000
150000
200000
NL_crAss000211, nt

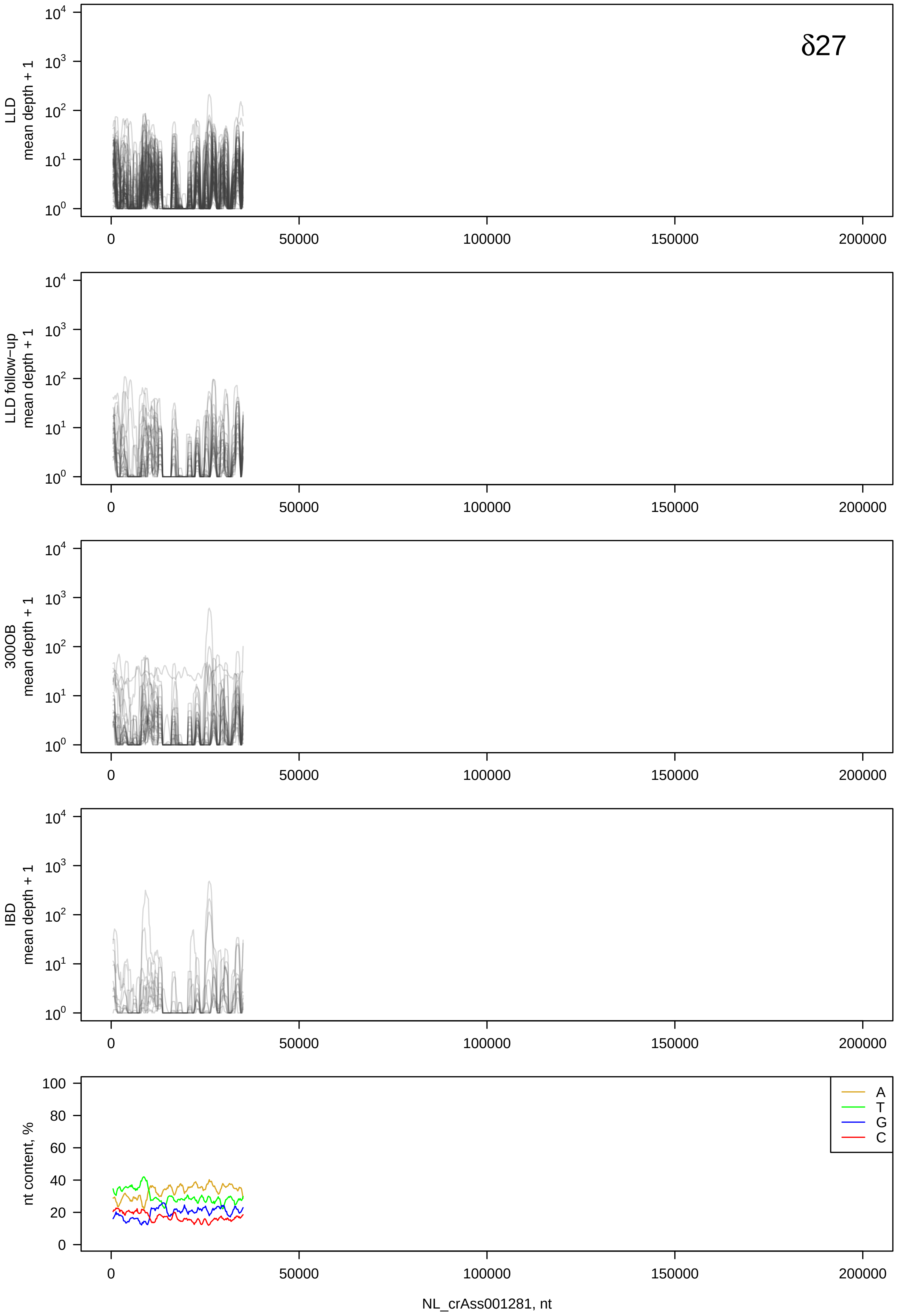
δ27
LLD mean depth + 1
LLD follow-up mean depth + 1
300OB mean depth + 1
IBD mean depth + 1
nt content, %
A
T
G
C
0
50000
100000
150000
200000
NL_crAss001281, nt

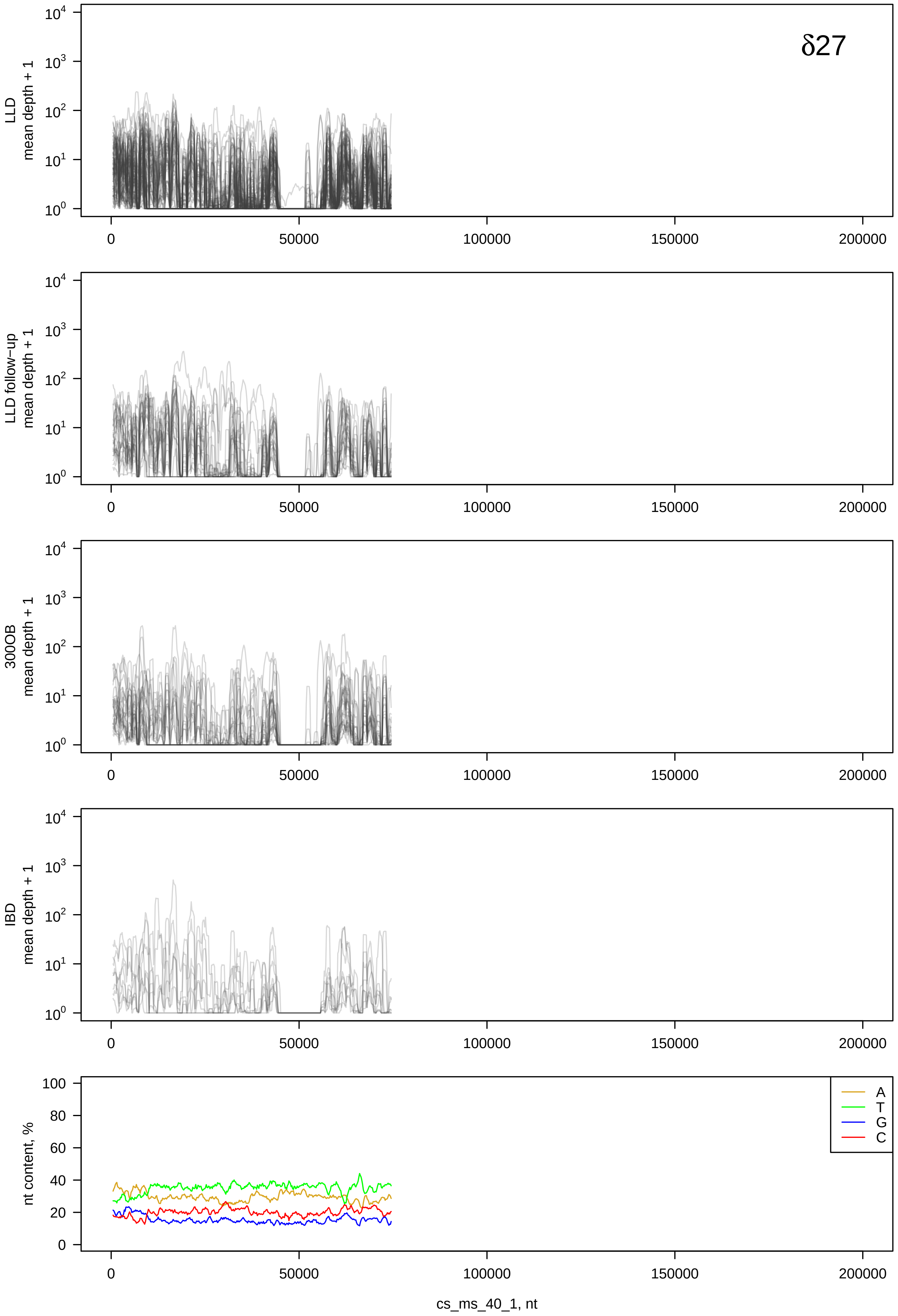
δ27
LLD
mean depth + 1
LLD follow-up
mean depth + 1
300OB
mean depth + 1
IBD
mean depth + 1
nt content, %
A
T
G
C
0
50000
100000
150000
200000
cs_ms_40_1, nt

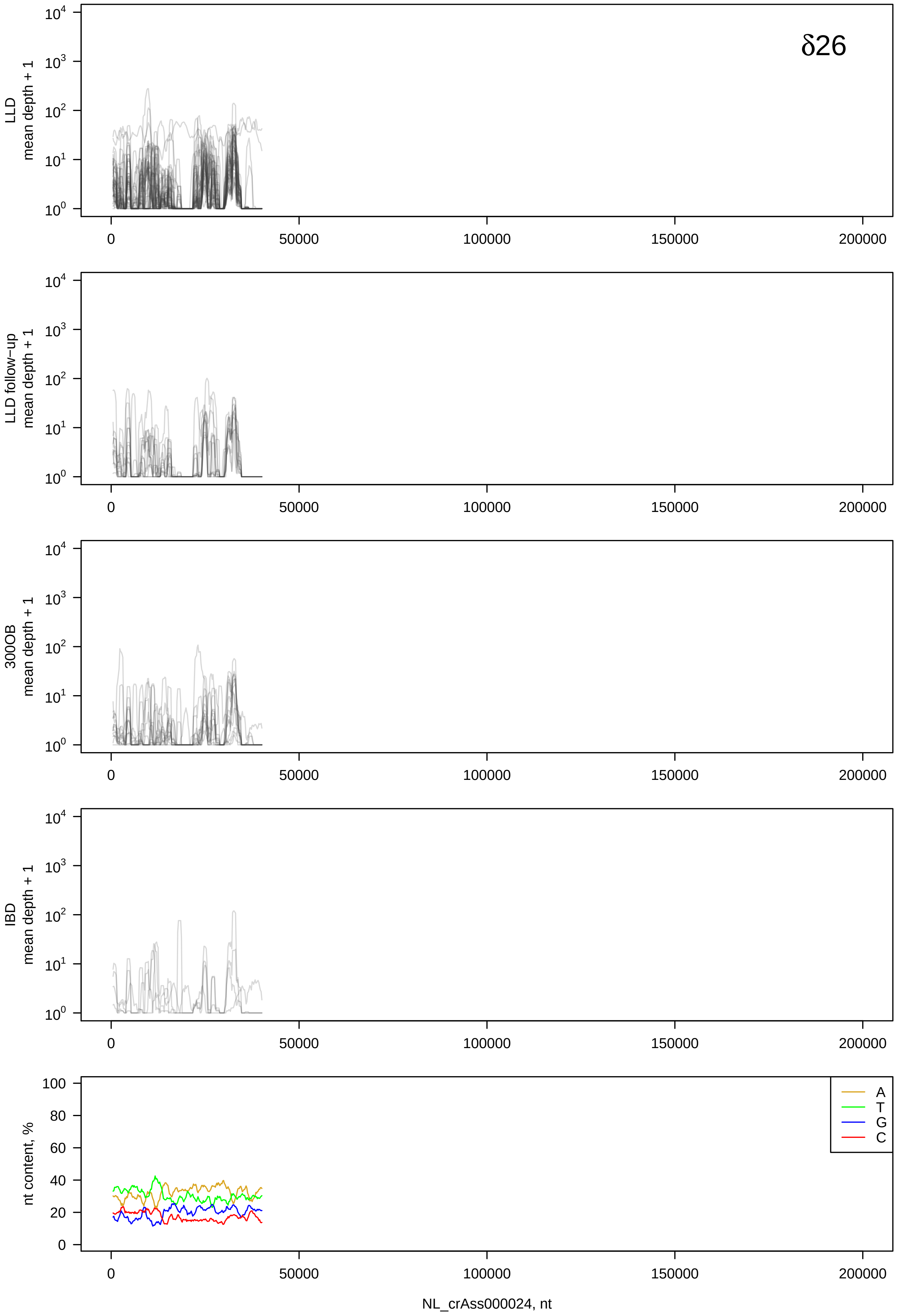

δ26
LLD mean depth + 1
LLD follow−up mean depth + 1
300OB mean depth + 1
IBD mean depth + 1
nt content, %
A
T
G
C
NL_crAss000024, nt

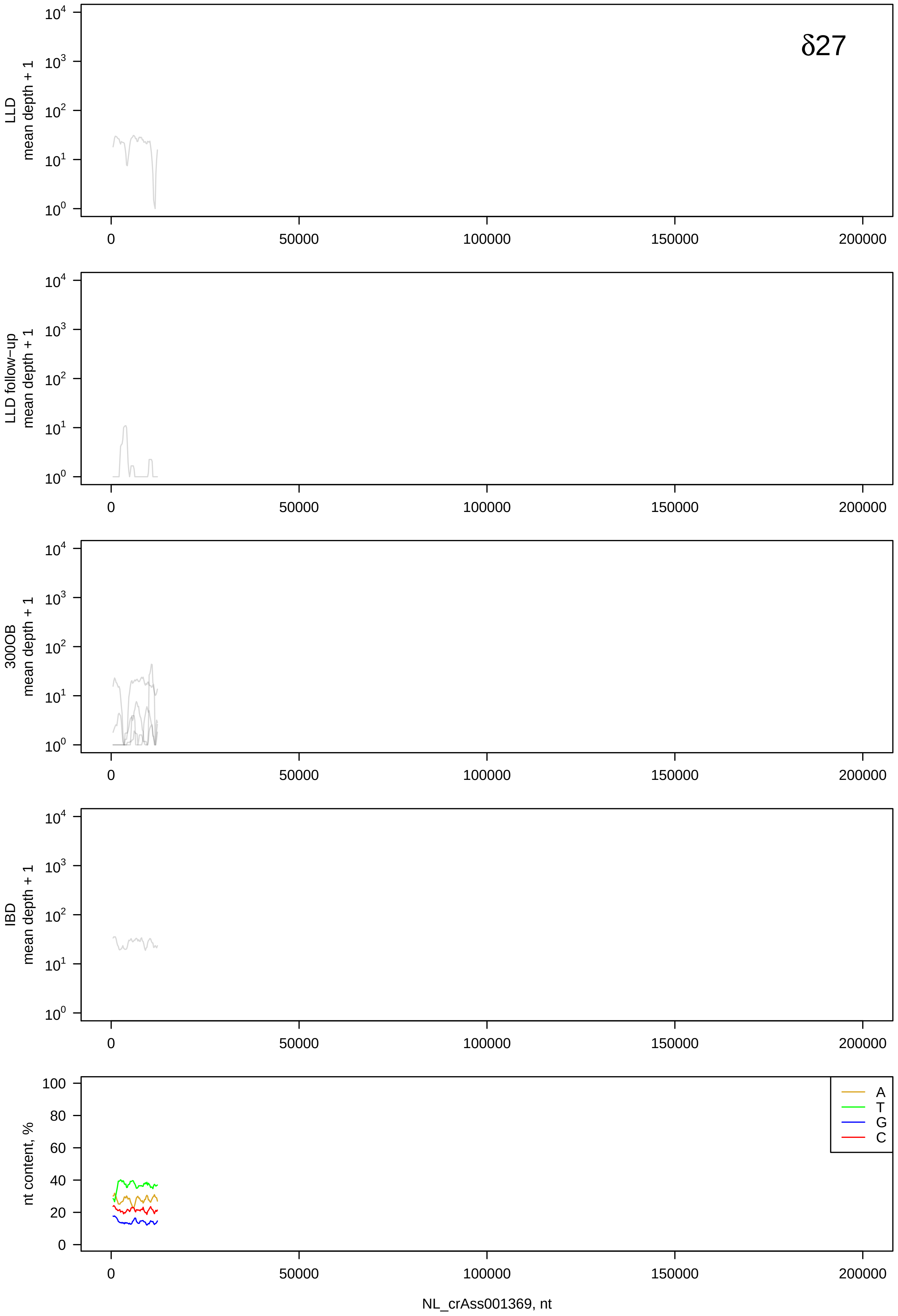
δ27
LLD
mean depth + 1
LLD follow−up
mean depth + 1
300OB
mean depth + 1
IBD
mean depth + 1
nt content, %
A
T
G
C
0
50000
100000
150000
200000
NL_crAss001369, nt

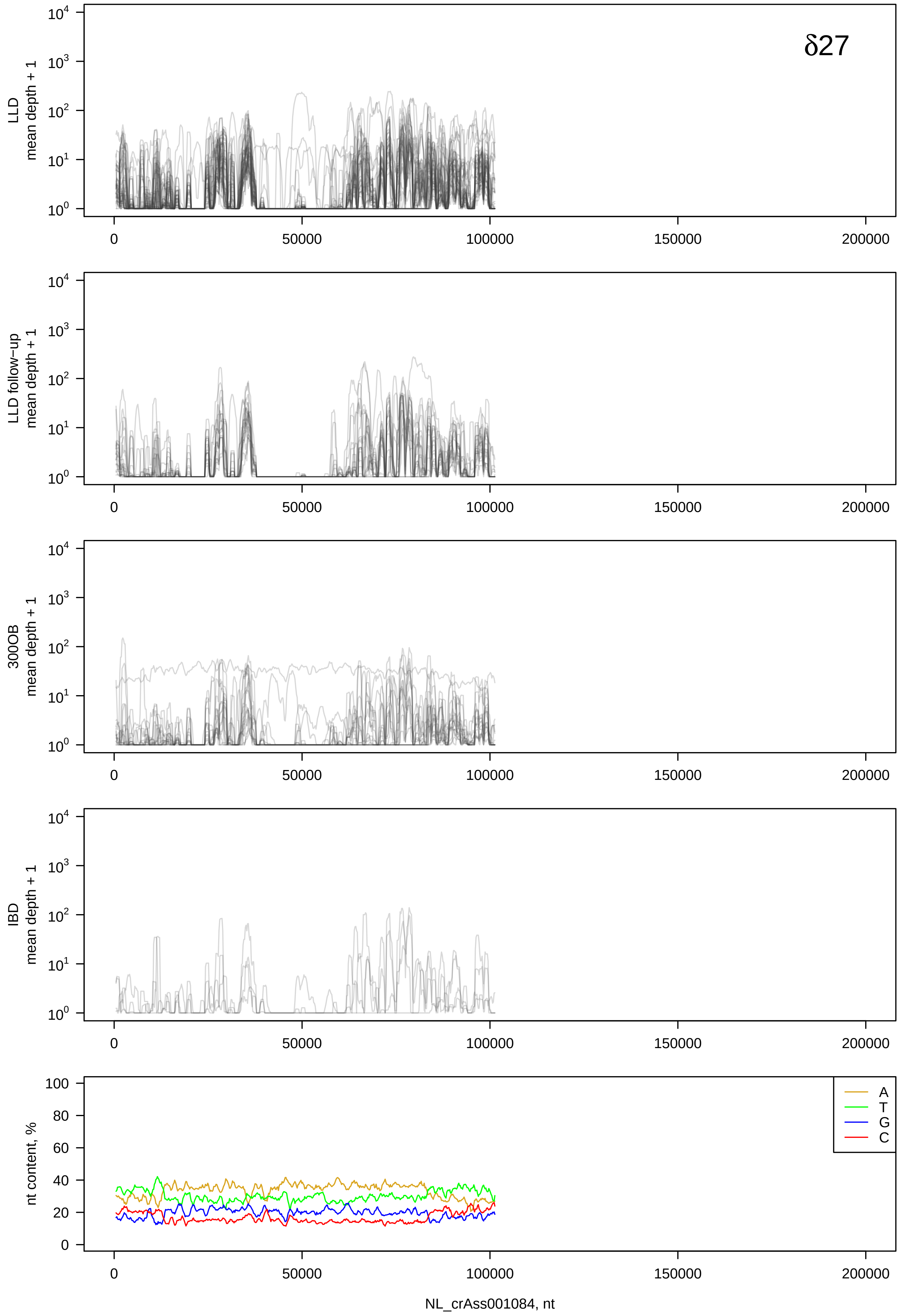

δ27
LLD mean depth + 1
LLD follow−up mean depth + 1
300OB mean depth + 1
IBD mean depth + 1
nt content, %
A
T
G
C
NL_crAss001084, nt

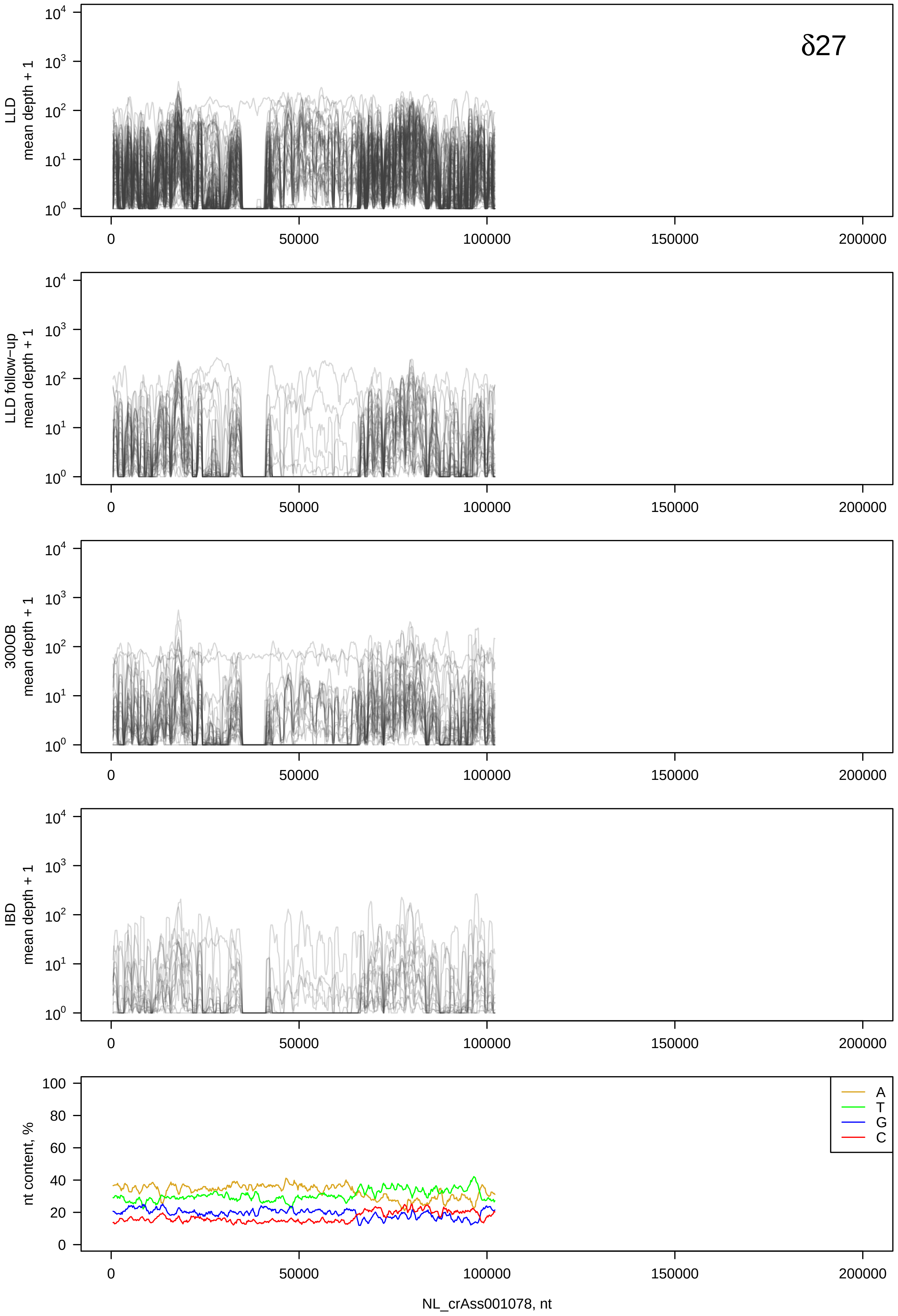

δ27
LLD mean depth + 1
LLD follow−up mean depth + 1
300OB mean depth + 1
IBD mean depth + 1
nt content, %
A
T
G
C
NL_crAss001078, nt

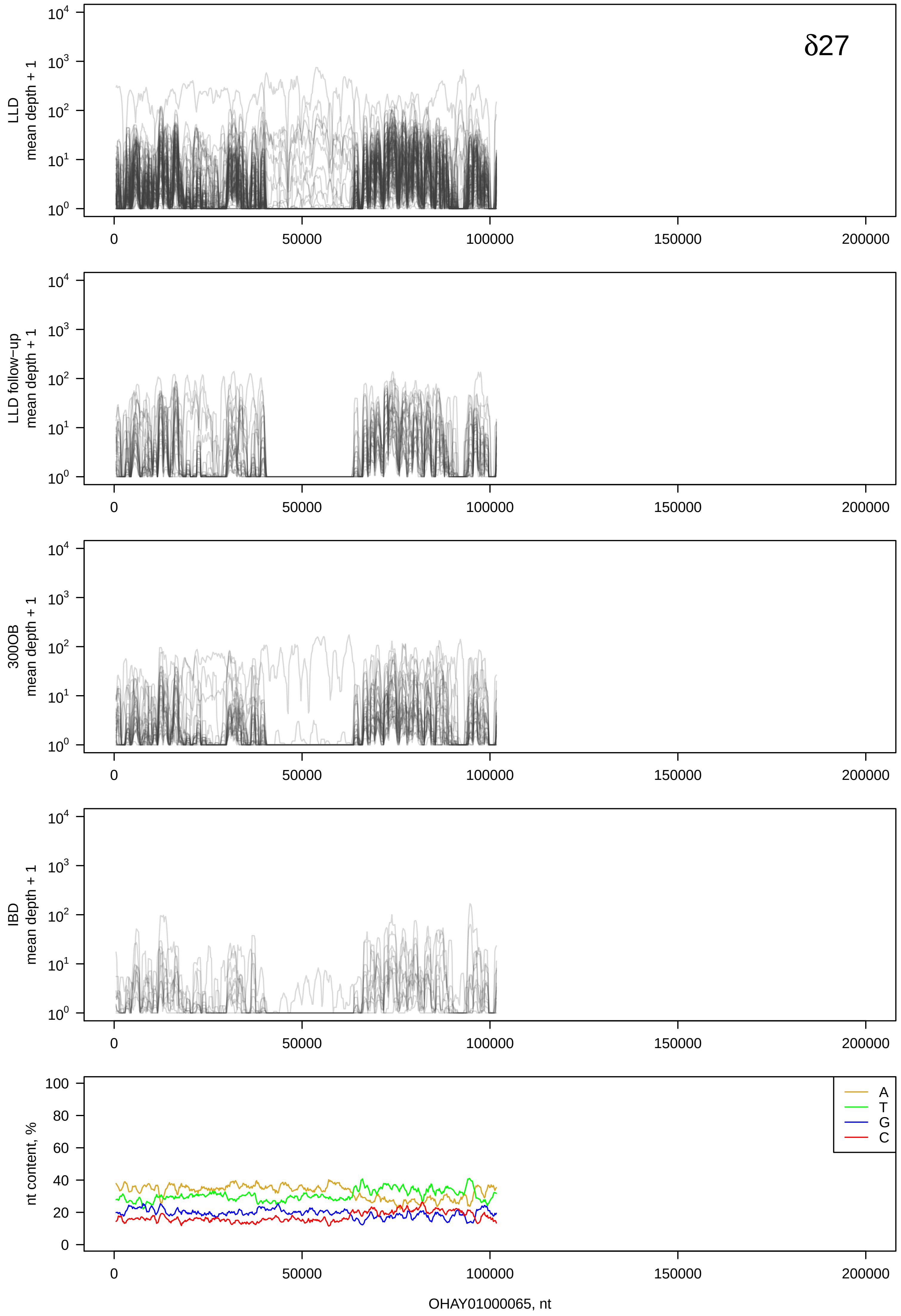
δ27
LLD
mean depth + 1
LLD follow−up
mean depth + 1
300OB
mean depth + 1
IBD
mean depth + 1
nt content, %
A
T
G
C
0
50000
100000
150000
200000
OHAY01000065, nt

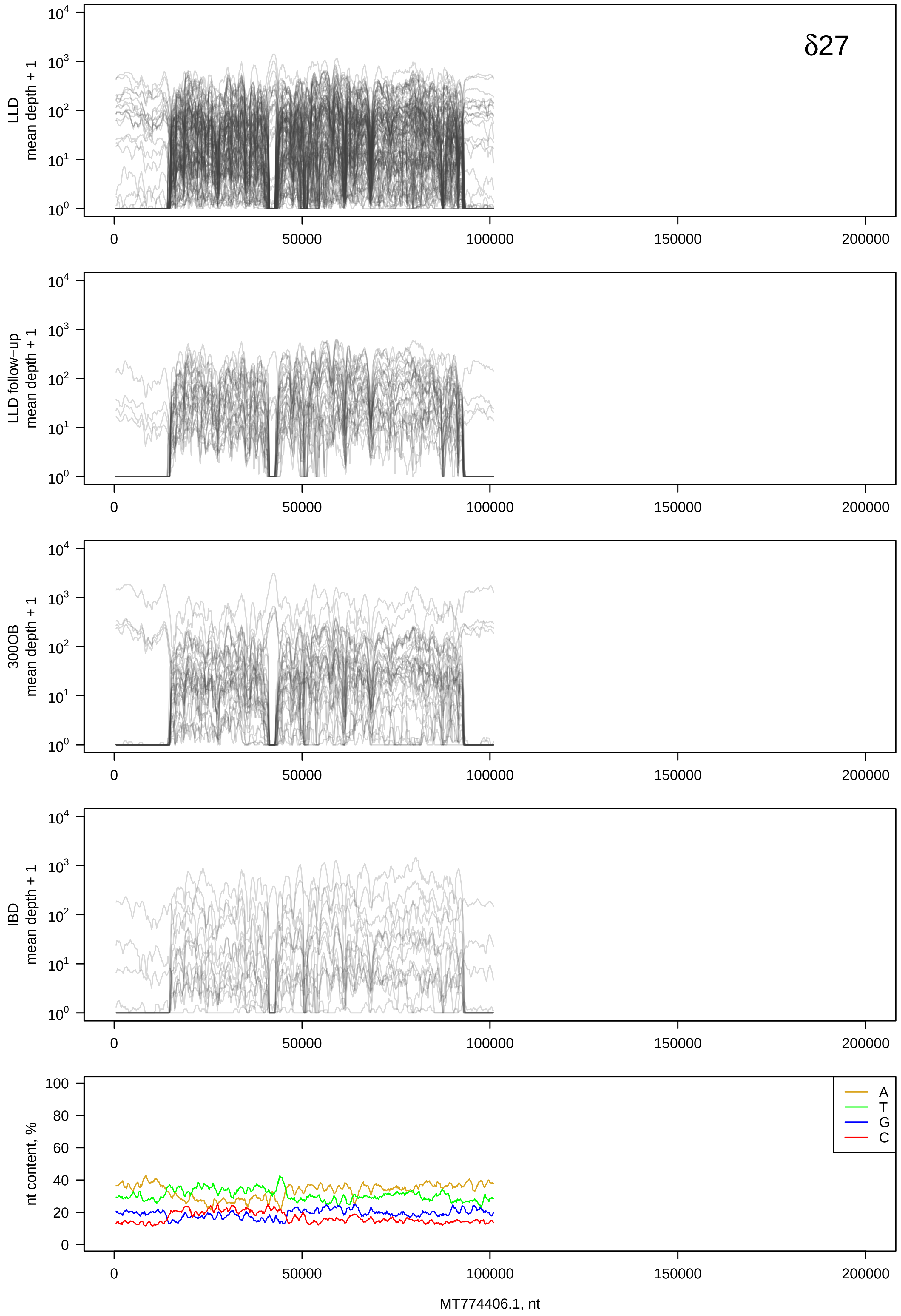
δ27
LLD
mean depth + 1
LLD follow−up
mean depth + 1
300OB
mean depth + 1
IBD
mean depth + 1
nt content, %
A
T
G
C
0
50000
100000
150000
200000
MT774406.1, nt

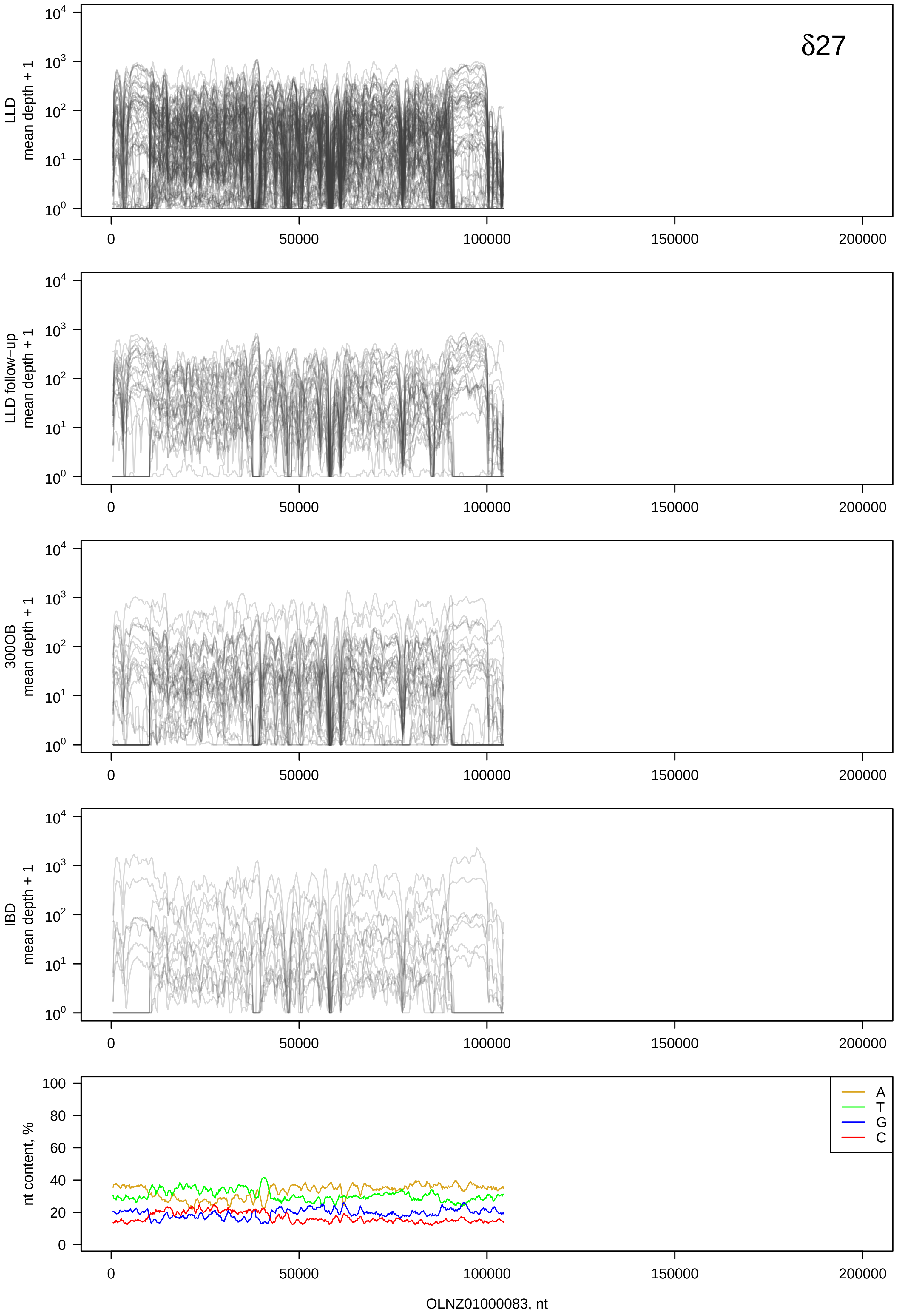

δ27
LLD mean depth + 1
LLD follow-up mean depth + 1
300OB mean depth + 1
IBD mean depth + 1
nt content, %
A
T
G
C
OLNZ01000083, nt

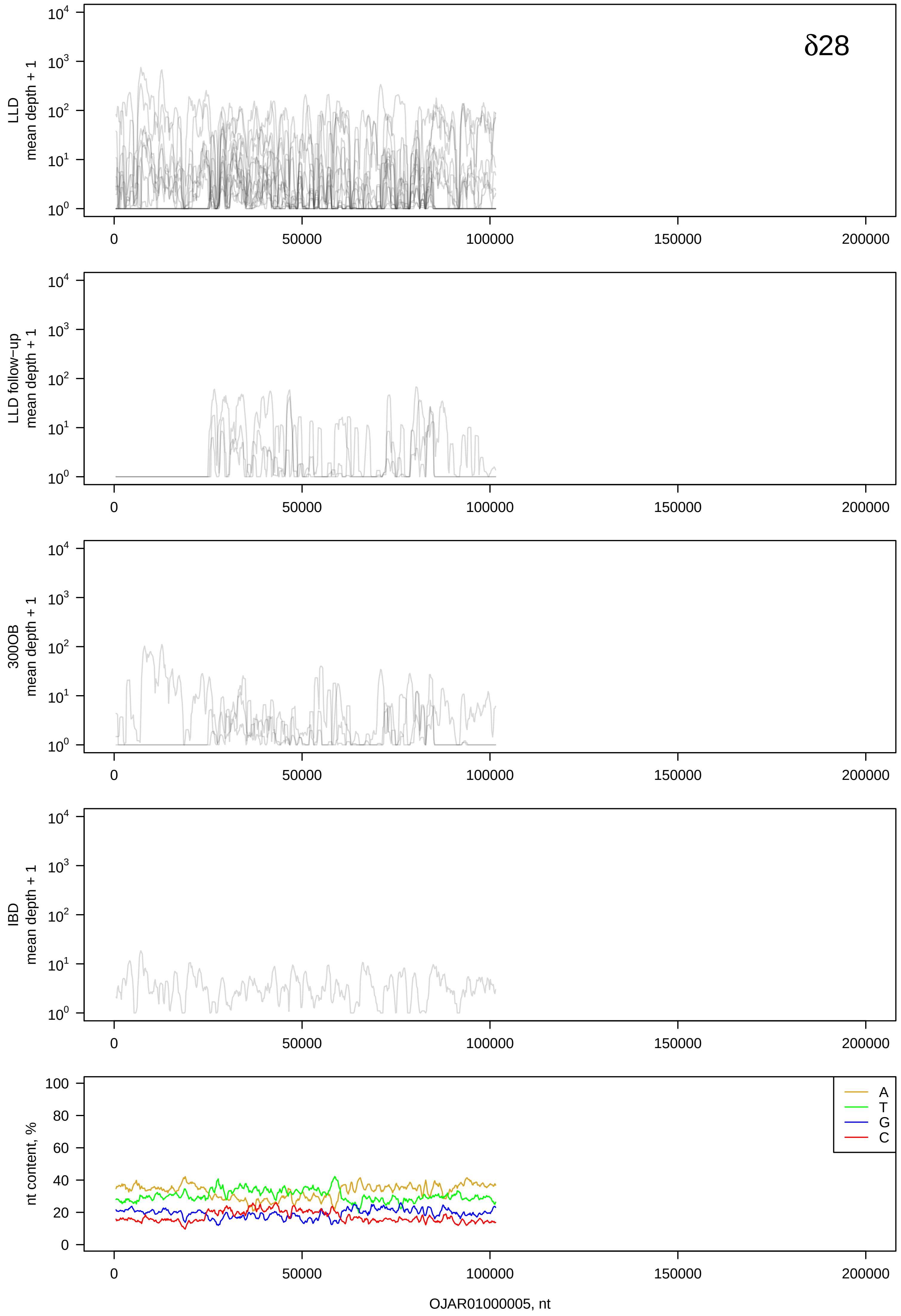
δ28
LLD mean depth + 1
LLD follow-up mean depth + 1
300OB mean depth + 1
IBD mean depth + 1
nt content, %
A
T
G
C
0
50000
100000
150000
200000
OJAR01000005, nt

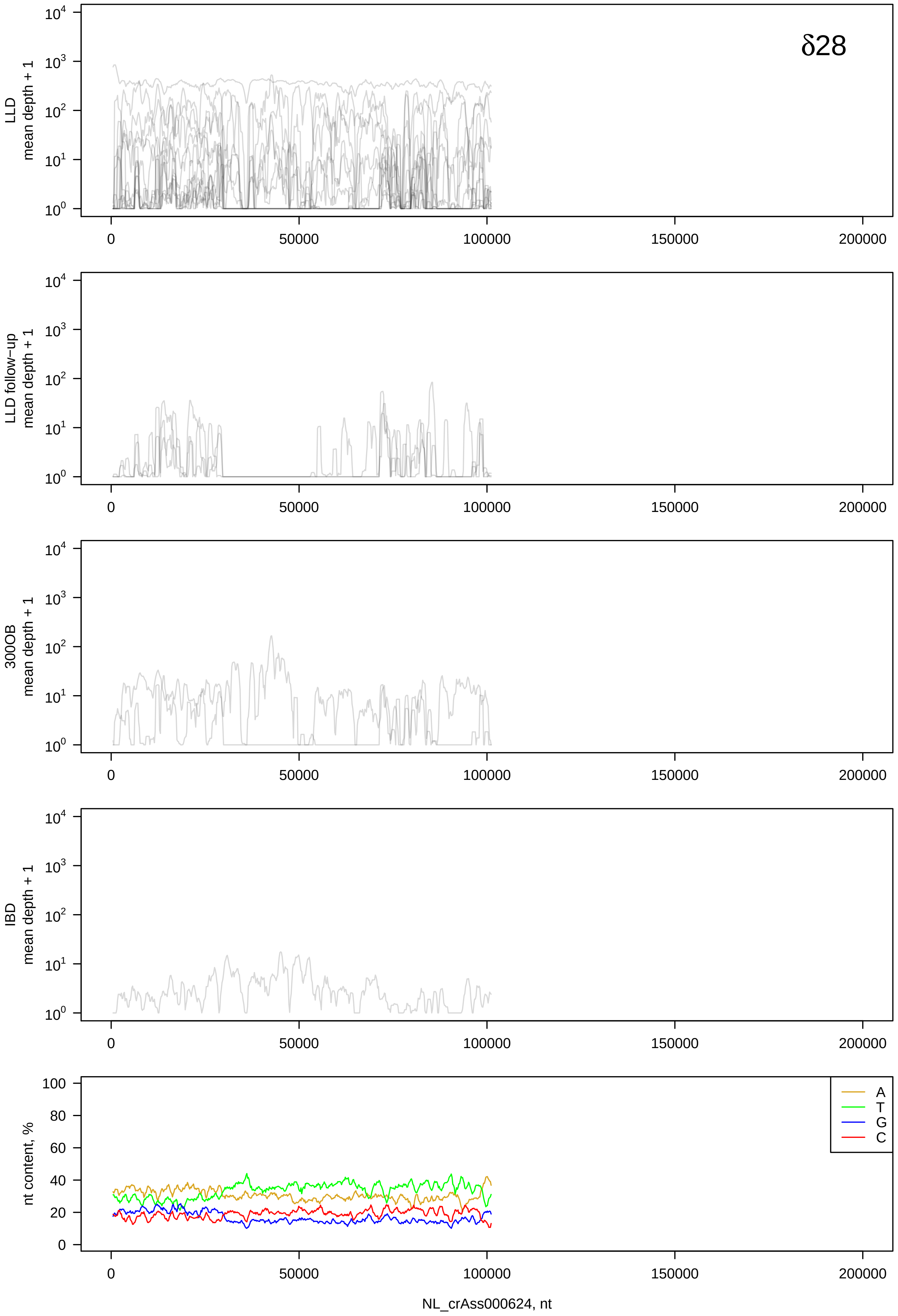
δ28
LLD
mean depth + 1
LLD follow−up
mean depth + 1
300OB
mean depth + 1
IBD
mean depth + 1
nt content, %
A
T
G
C
NL_crAss000624, nt

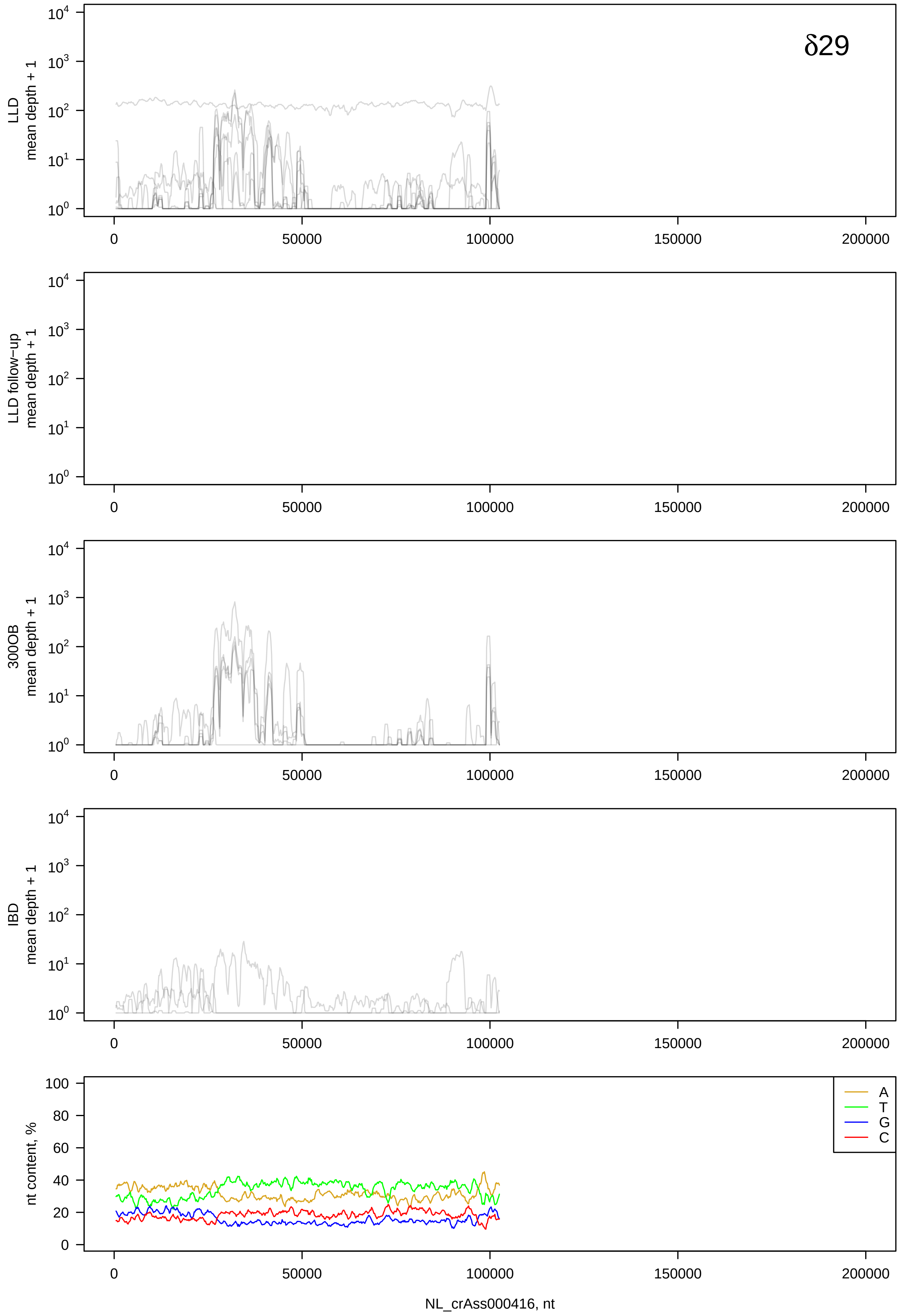

δ29
LLD
mean depth + 1
LLD follow−up
mean depth + 1
300OB
mean depth + 1
IBD
mean depth + 1
nt content, %
A
T
G
C
NL_crAss000416, nt

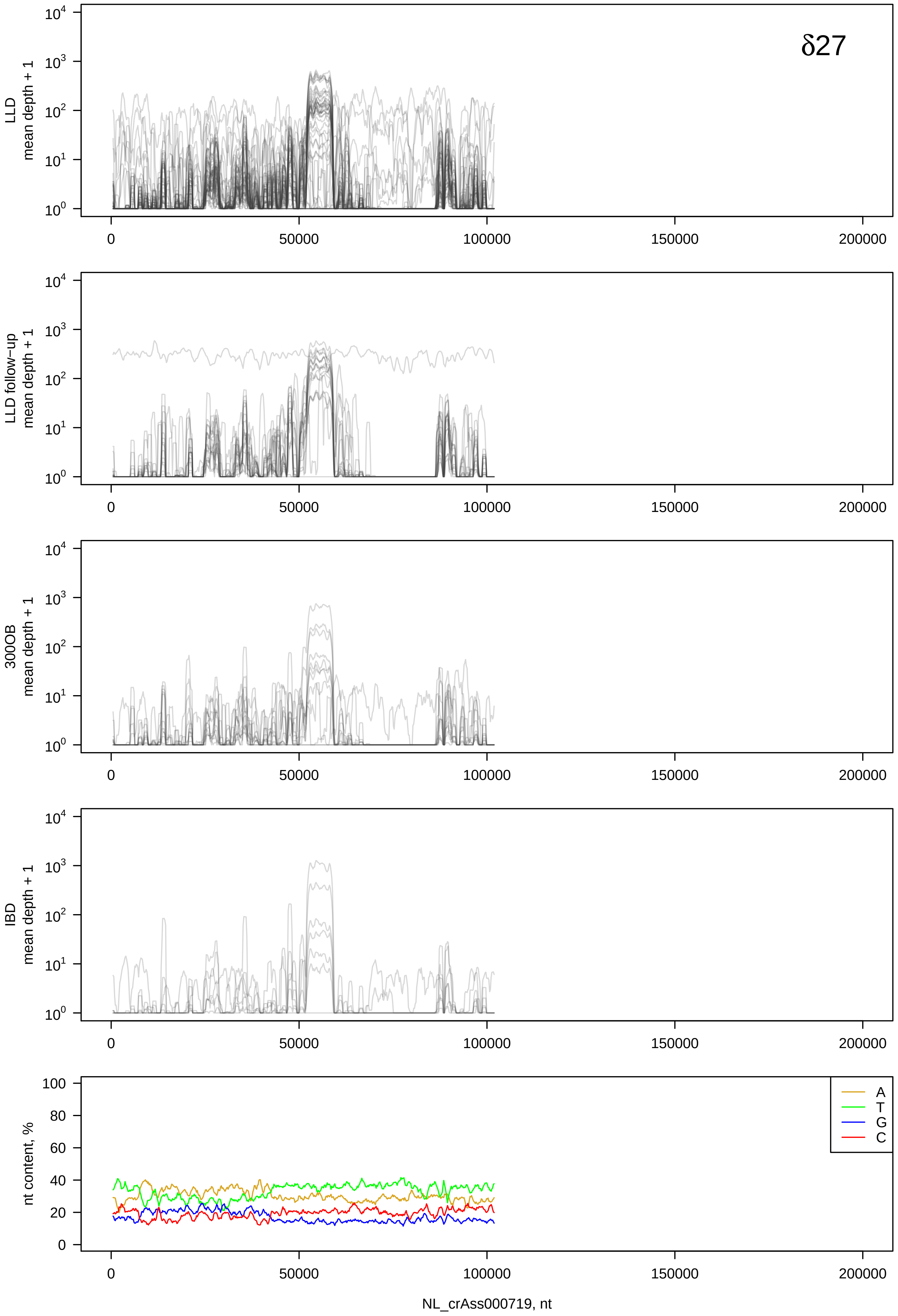

δ27
LLD
mean depth + 1
LLD follow−up
mean depth + 1
300OB
mean depth + 1
IBD
mean depth + 1
nt content, %
A
T
G
C
NL_crAss000719, nt

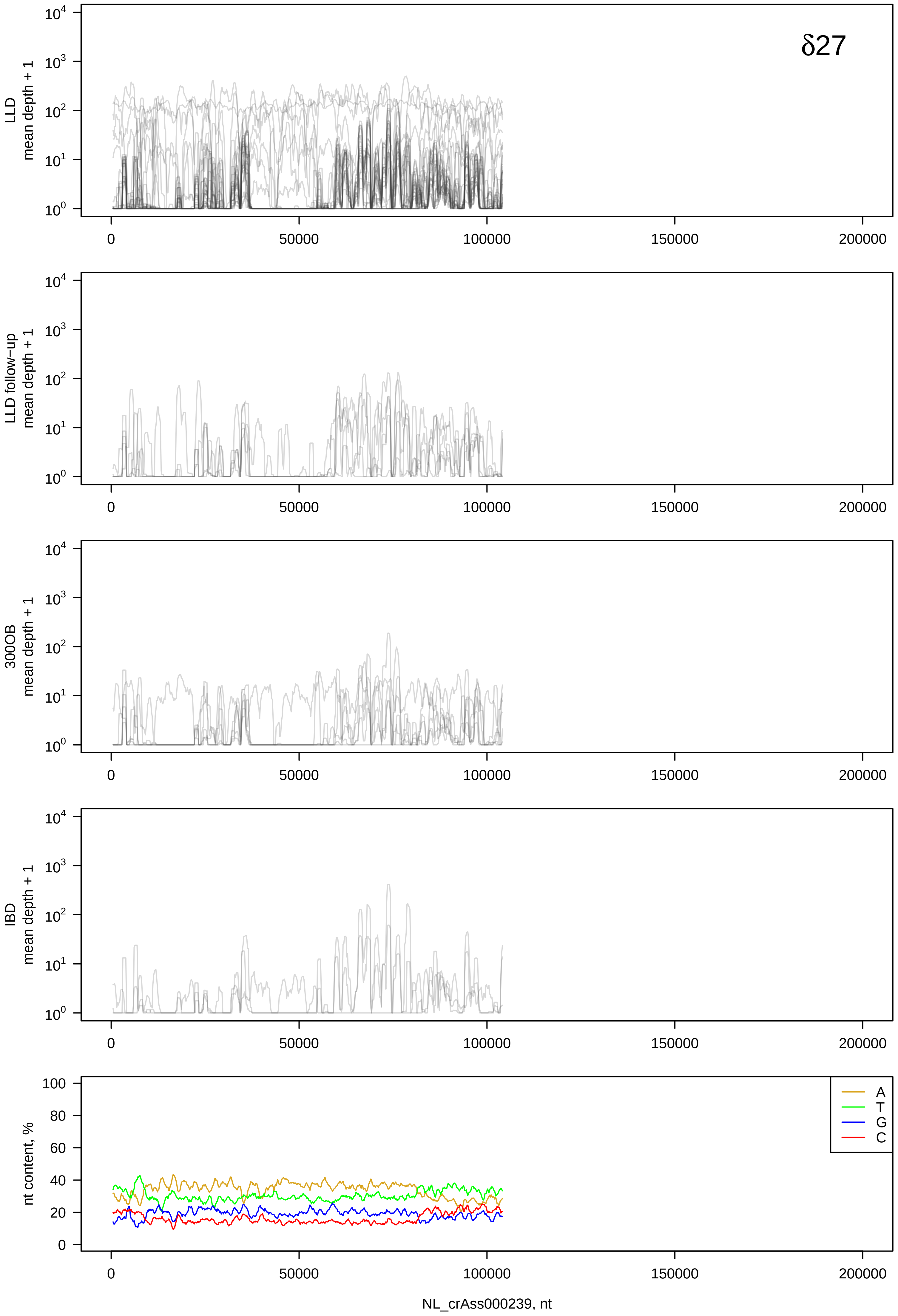
δ27
LLD mean depth + 1
LLD follow−up mean depth + 1
300OB mean depth + 1
IBD mean depth + 1
nt content, %
A
T
G
C
NL_crAss000239, nt

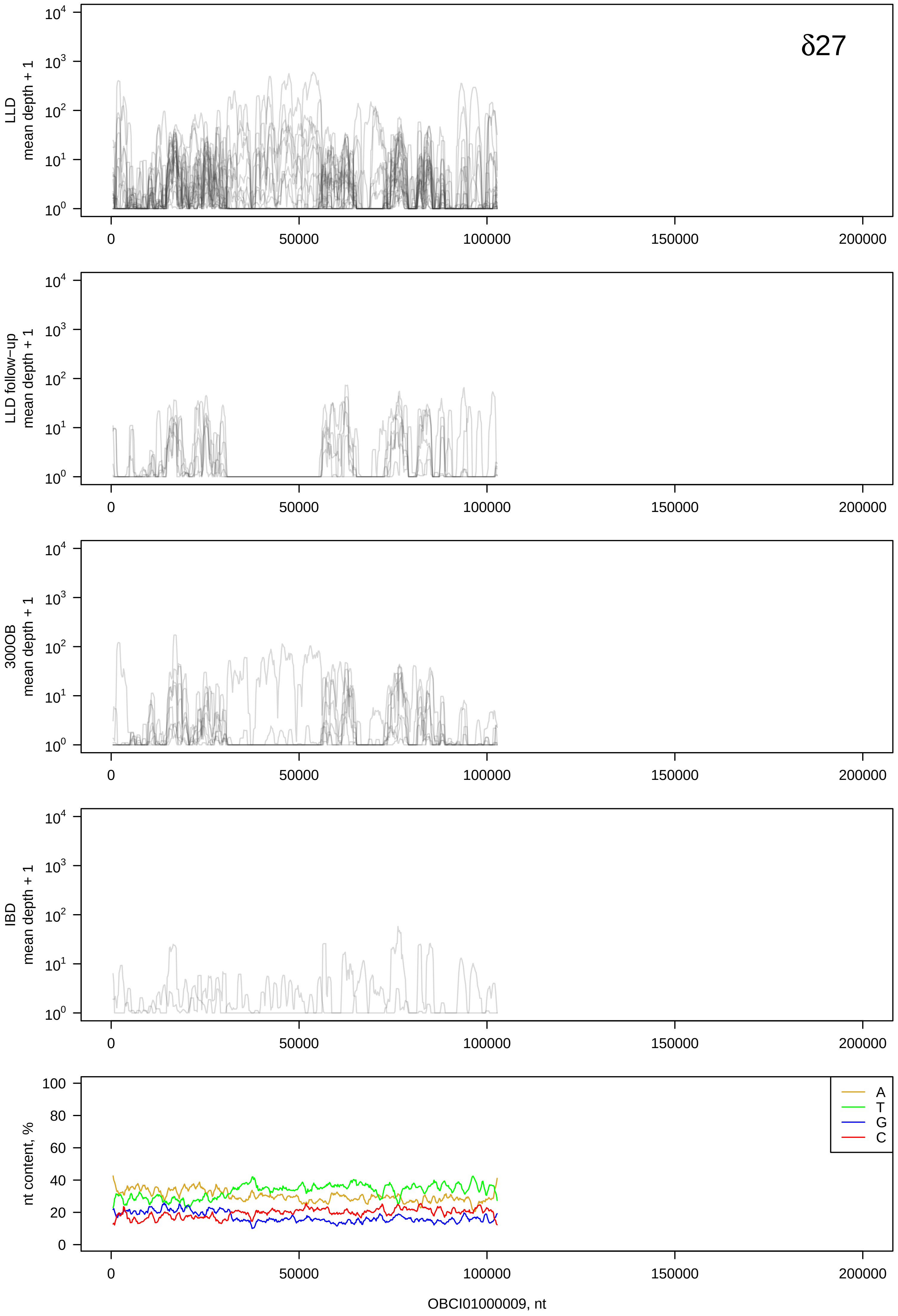

δ27
LLD
mean depth + 1
LLD follow−up
mean depth + 1
300OB
mean depth + 1
IBD
mean depth + 1
nt content, %
A
T
G
C
0
50000
100000
150000
200000
OBCI01000009, nt

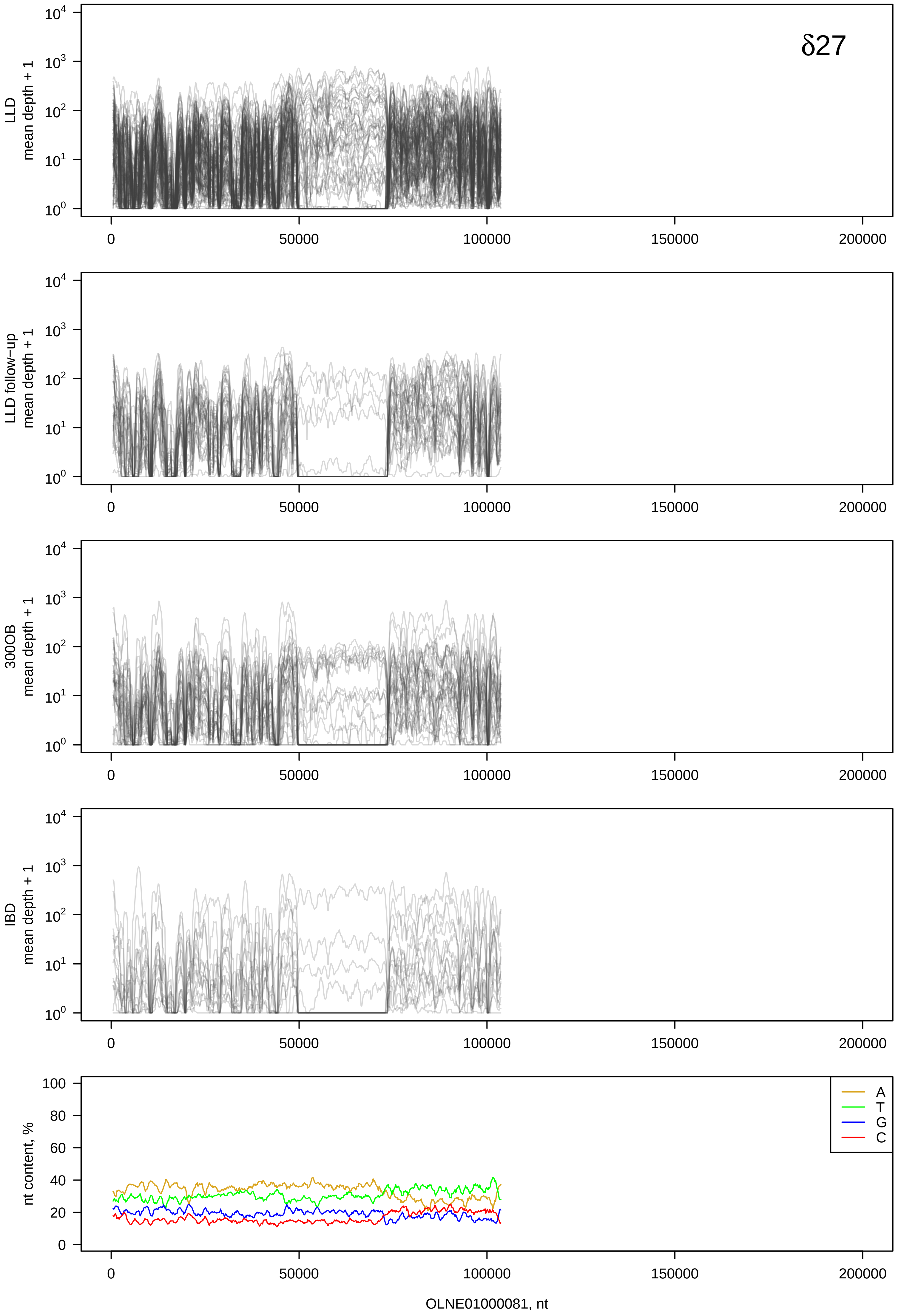
δ27
LLD
mean depth + 1
LLD follow-up
mean depth + 1
300OB
mean depth + 1
IBD
mean depth + 1
nt content, %
A
T
G
C
OLNE01000081, nt

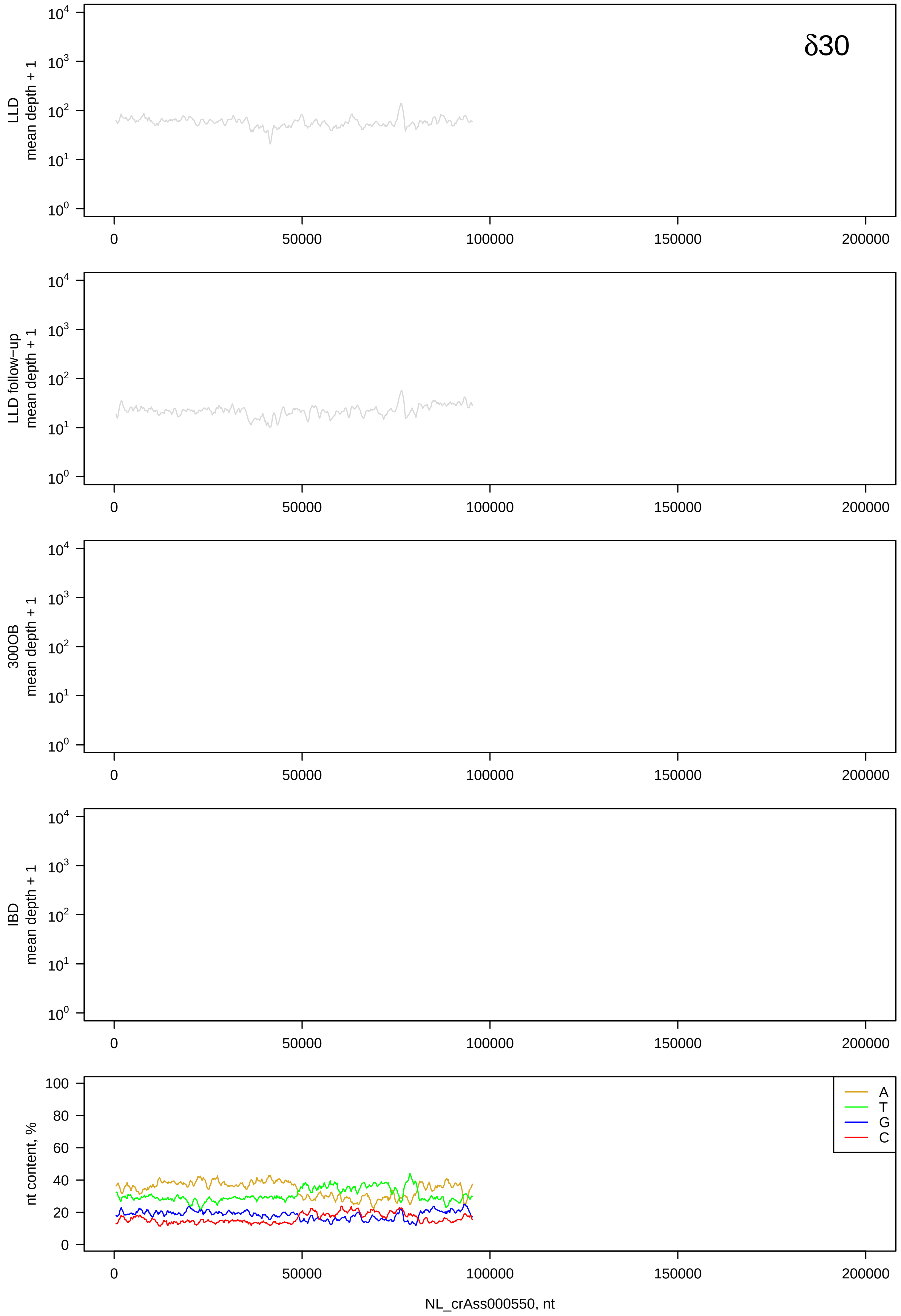
δ30
LLD
mean depth + 1
LLD follow−up
mean depth + 1
300OB
mean depth + 1
IBD
mean depth + 1
nt content, %
A
T
G
C
NL_crAss000550, nt

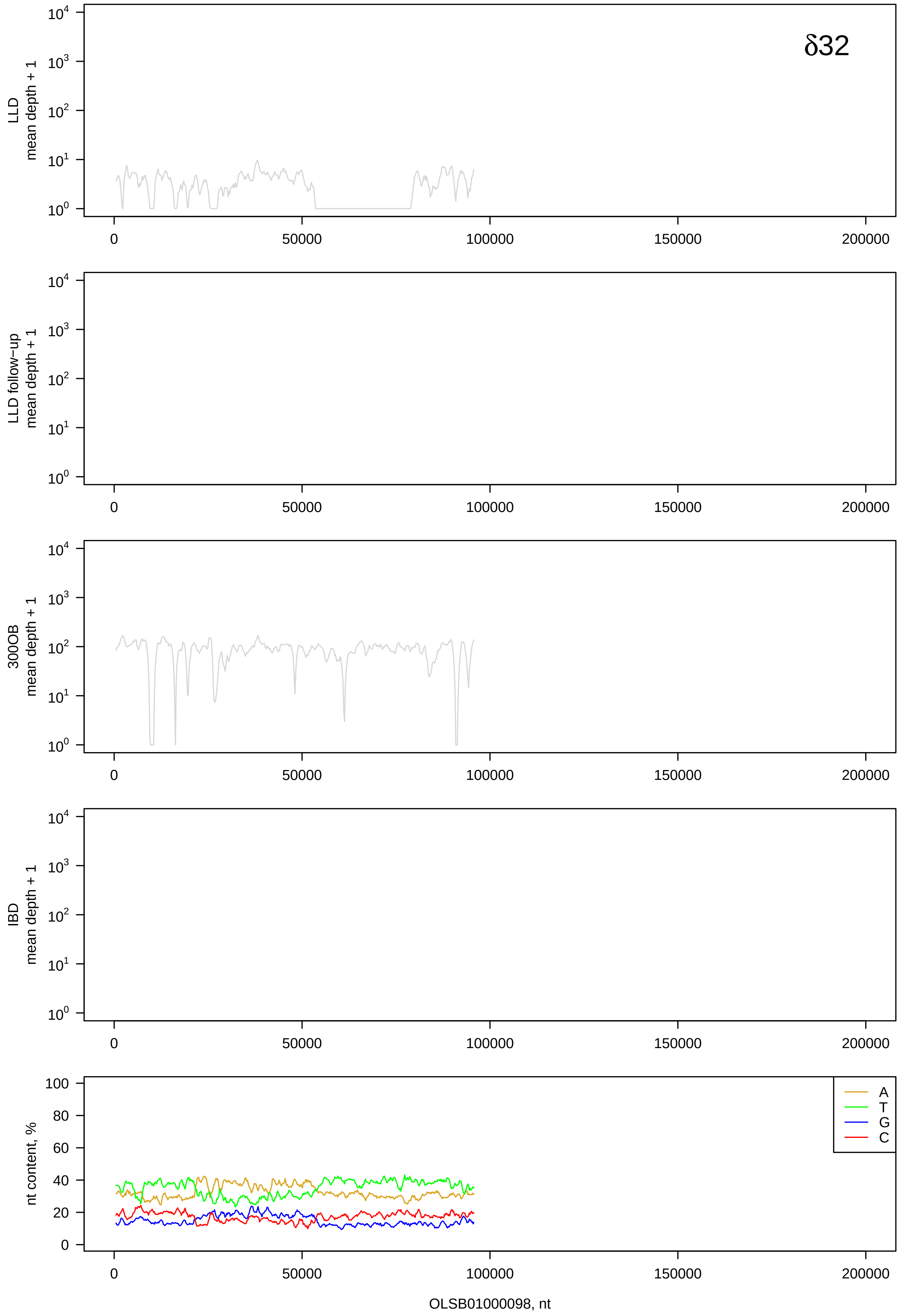

δ32
LLD mean depth + 1
LLD follow−up mean depth + 1
300OB mean depth + 1
IBD mean depth + 1
nt content, %
A
T
G
C
0
50000
100000
150000
200000
OLSB01000098, nt

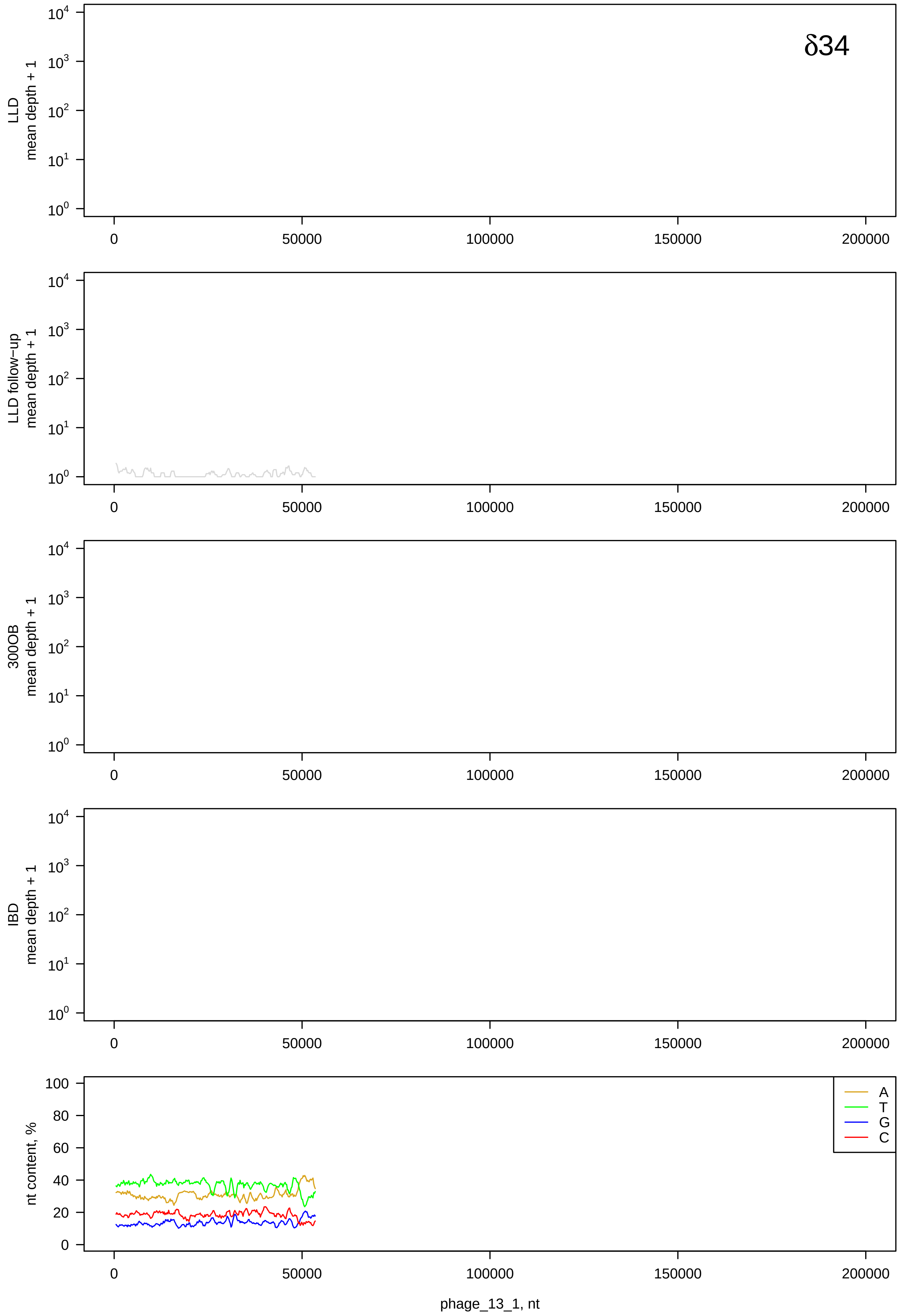
δ34
LLD
mean depth + 1
LLD follow−up
mean depth + 1
300OB
mean depth + 1
IBD
mean depth + 1
nt content, %
A
T
G
C
phage_13_1, nt

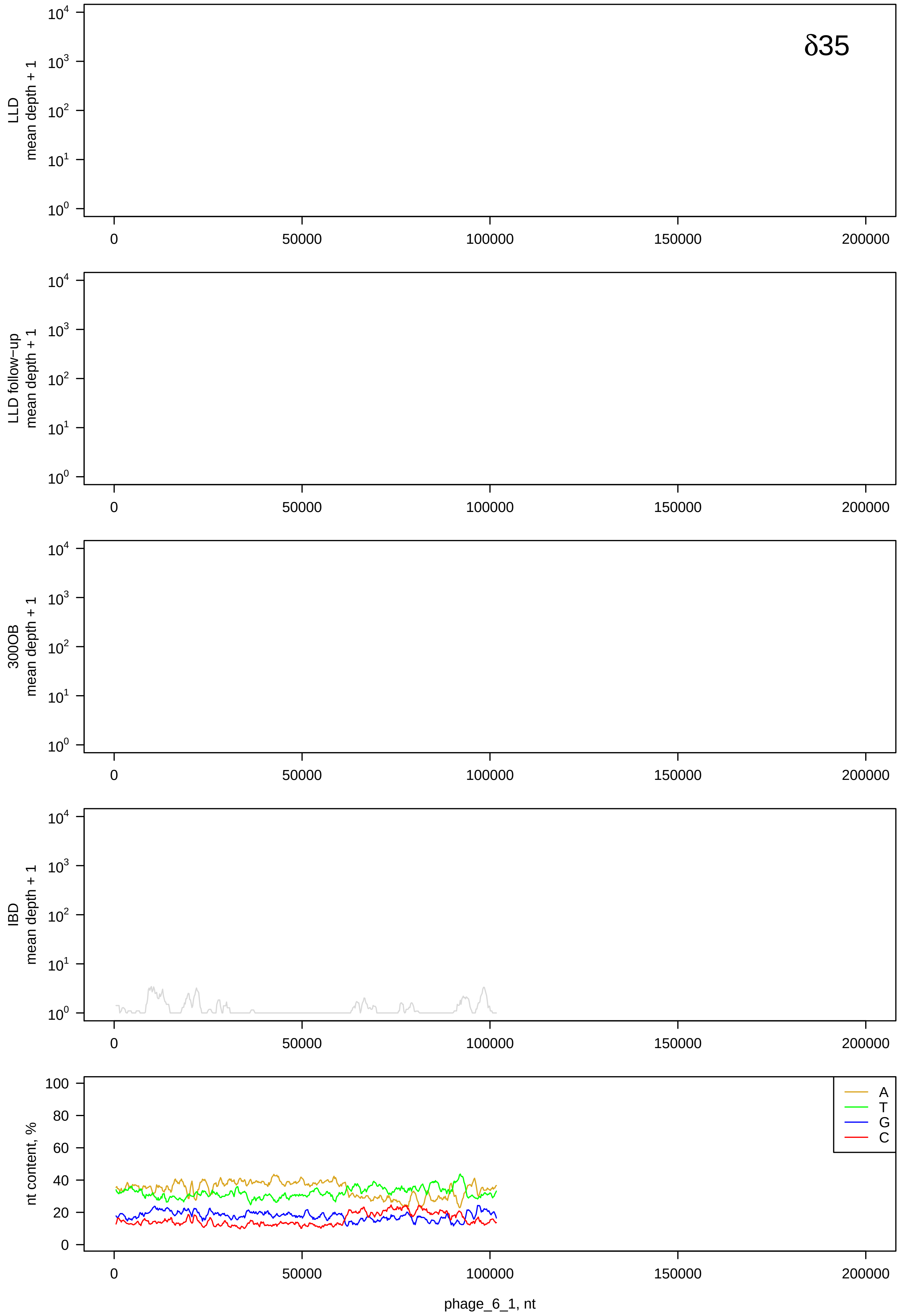

δ35
LLD
mean depth + 1
LLD follow−up
mean depth + 1
300OB
mean depth + 1
IBD
mean depth + 1
nt content, %
A
T
G
C
0
50000
100000
150000
200000
phage_6_1, nt

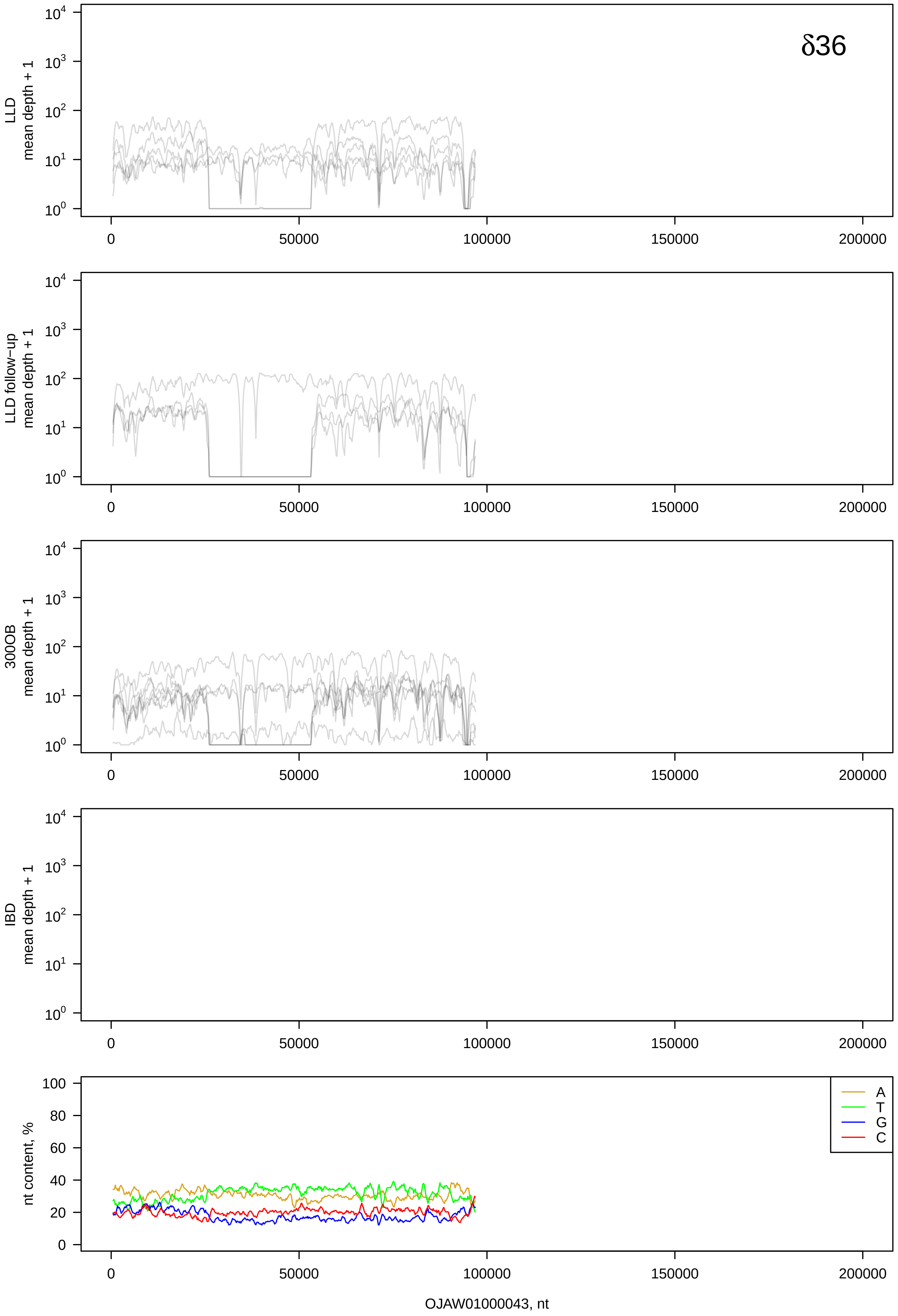
δ36
LLD
mean depth + 1
LLD follow-up
mean depth + 1
300OB
mean depth + 1
IBD
mean depth + 1
$10^0$
$10^1$
$10^2$
$10^3$
$10^4$
0
50000
100000
150000
200000
nt content, %
0
20
40
60
80
100
A
T
G
C
OJAW01000043, nt

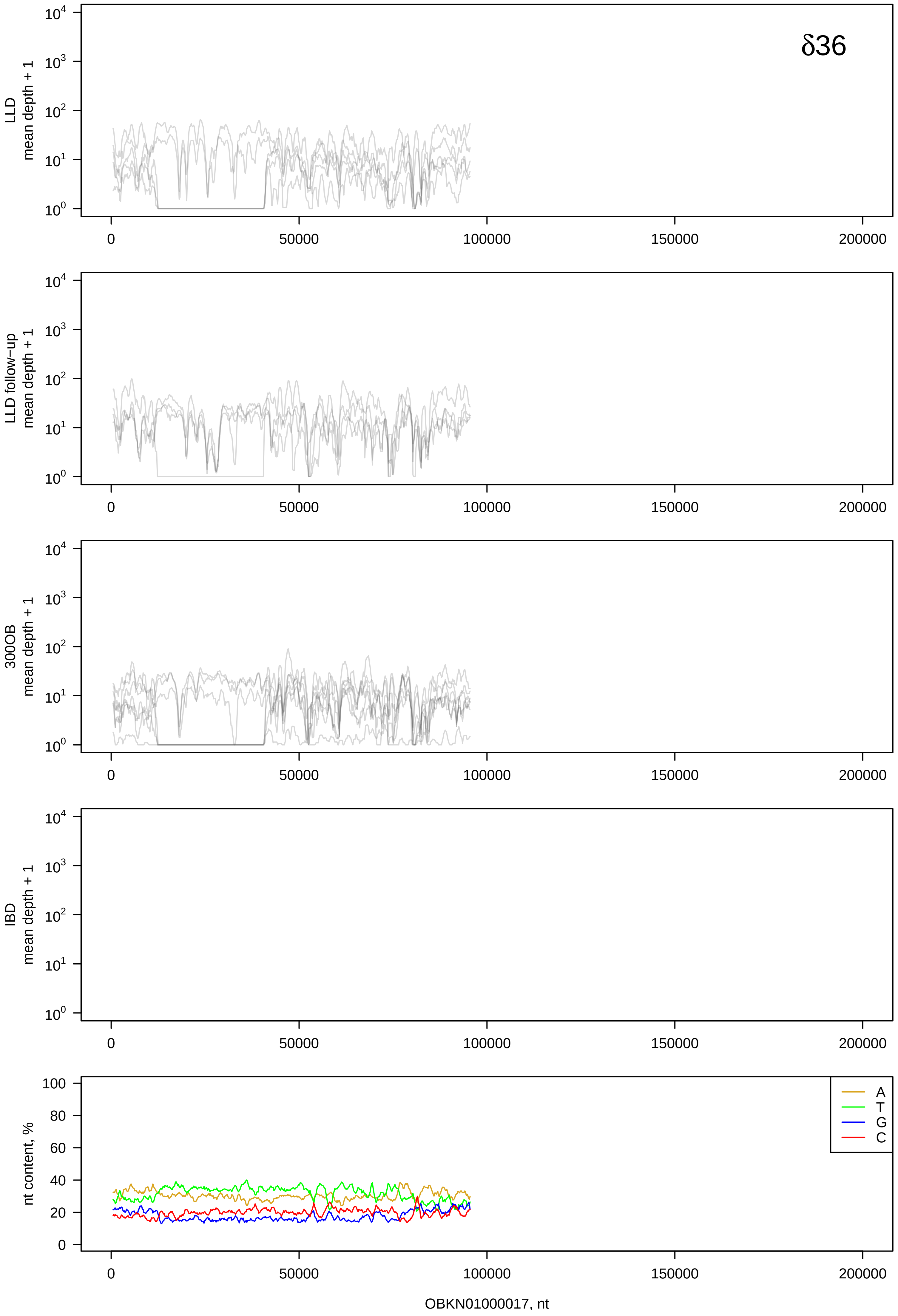
δ36
LLD
mean depth + 1
LLD follow-up
mean depth + 1
300OB
mean depth + 1
IBD
mean depth + 1
nt content, %
A
T
G
C
OBKN01000017, nt

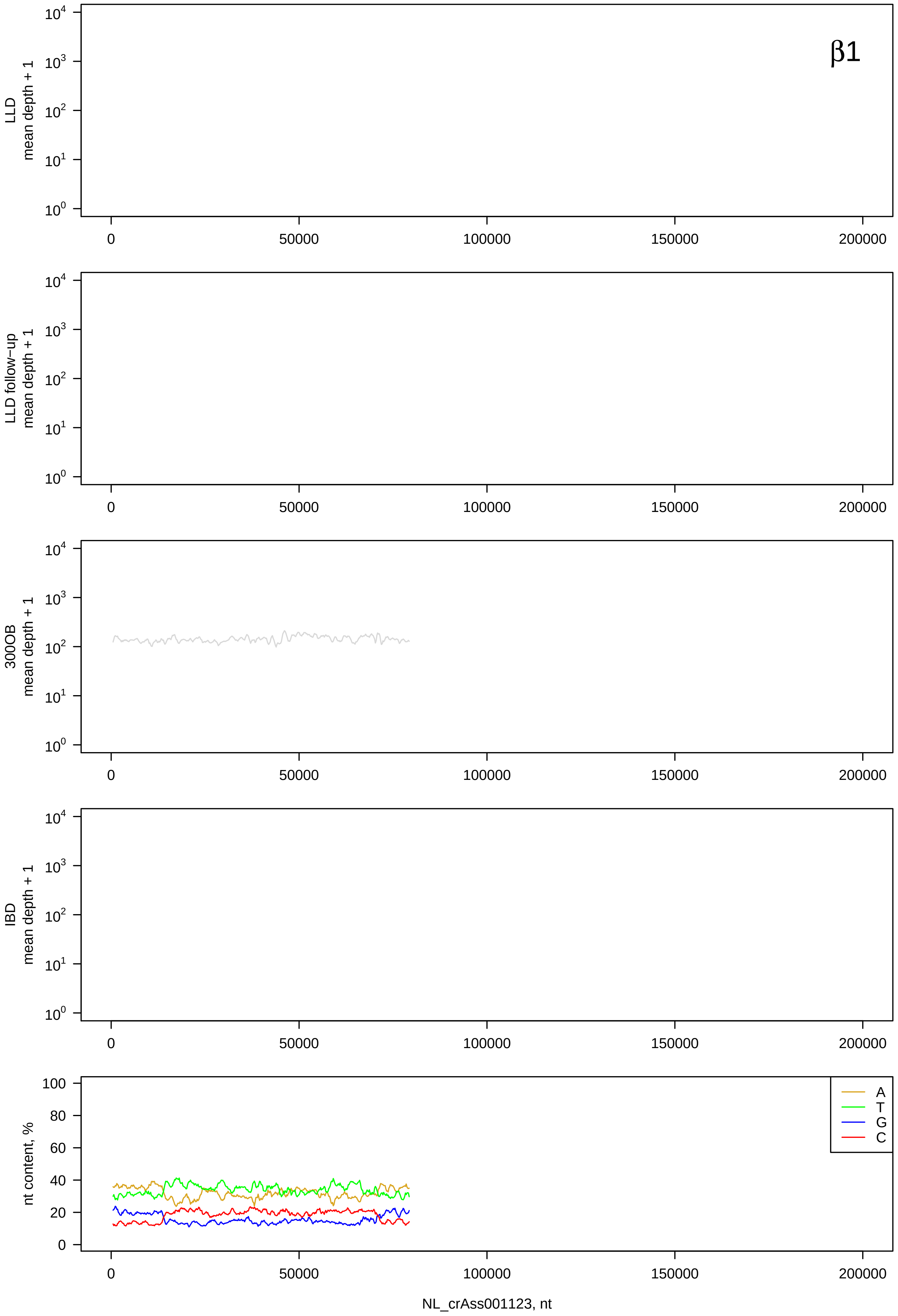

β1
LLD mean depth + 1
LLD follow−up mean depth + 1
300OB mean depth + 1
IBD mean depth + 1
nt content, %
A
T
G
C
NL_crAss001123, nt

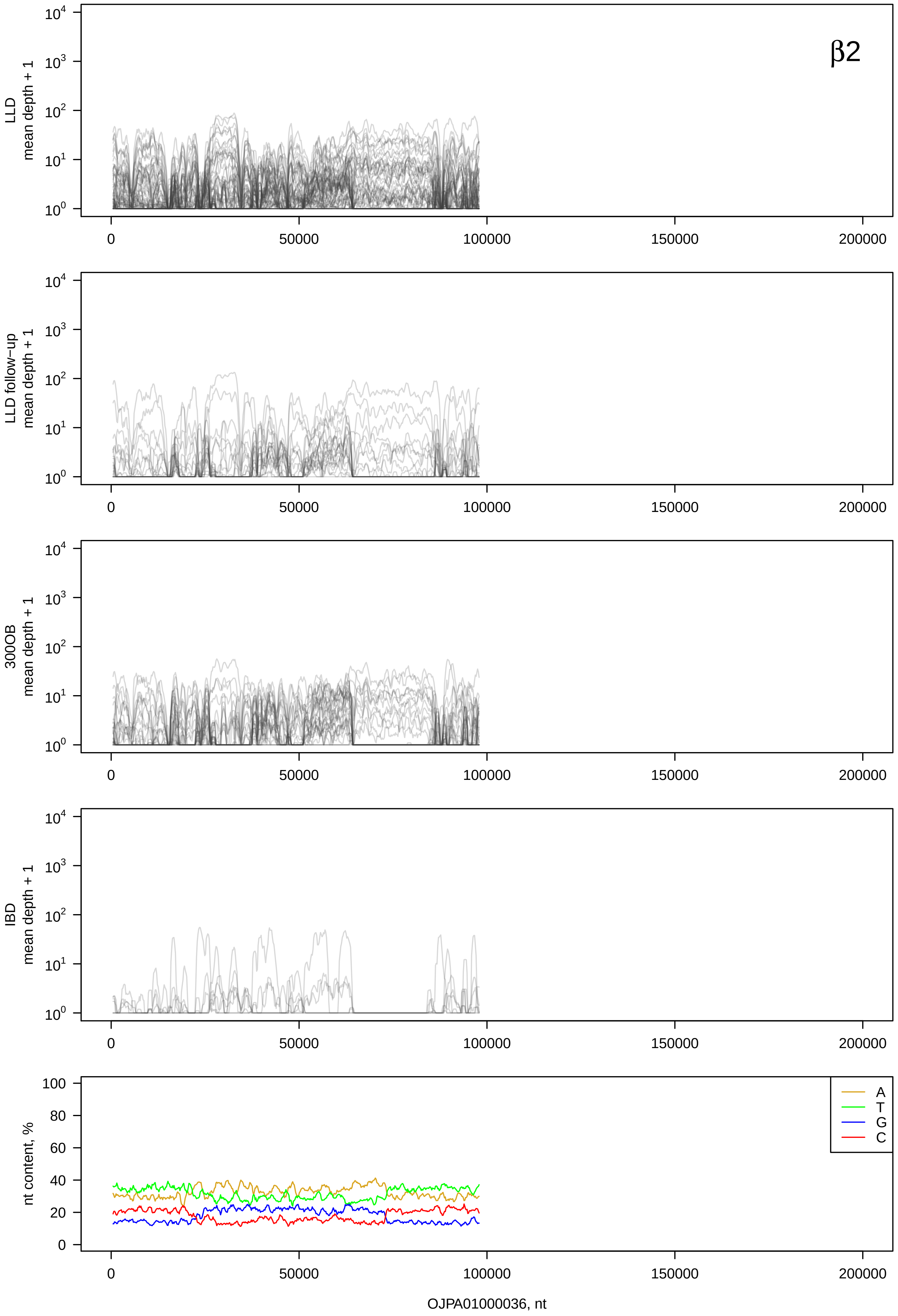
β2
LLD mean depth + 1
LLD follow-up mean depth + 1
300OB mean depth + 1
IBD mean depth + 1
nt content, %
A
T
G
C
OJPA01000036, nt

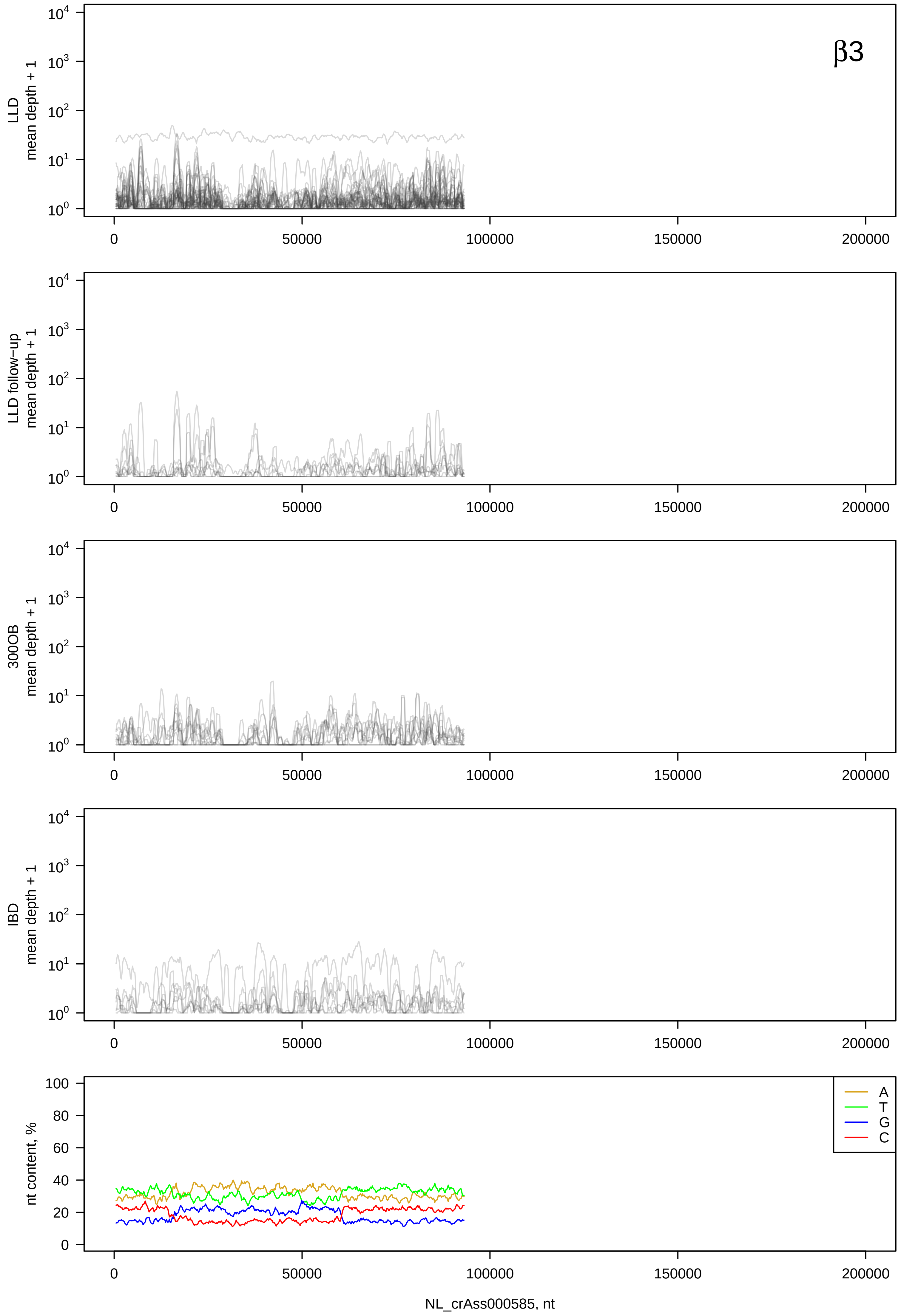
β3
LLD mean depth + 1
LLD follow-up mean depth + 1
300OB mean depth + 1
IBD mean depth + 1
nt content, %
A
T
G
C
NL_crAss000585, nt

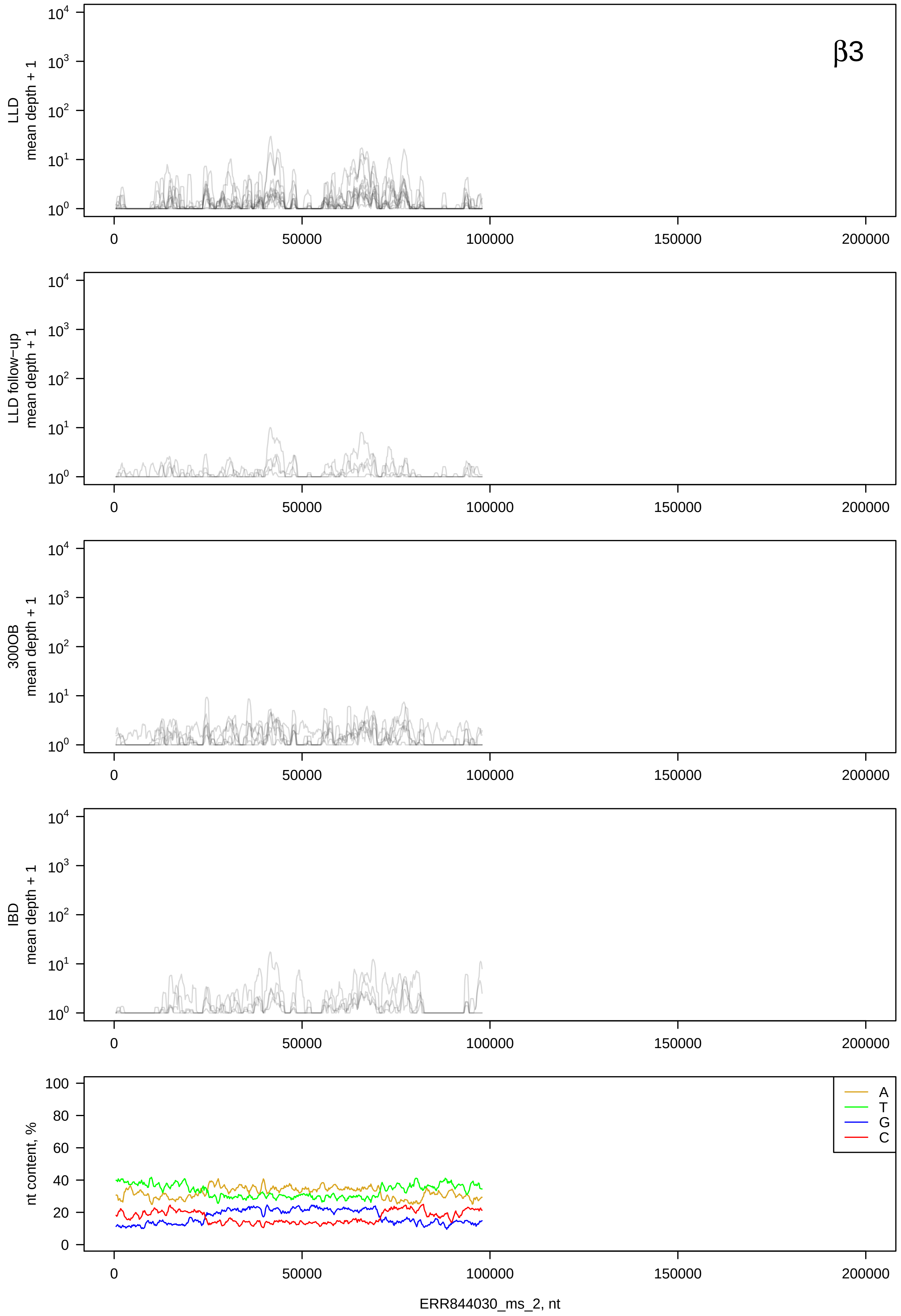

β3
LLD mean depth + 1
LLD follow−up mean depth + 1
300OB mean depth + 1
IBD mean depth + 1
nt content, %
A
T
G
C
0
50000
100000
150000
200000
ERR844030_ms_2, nt

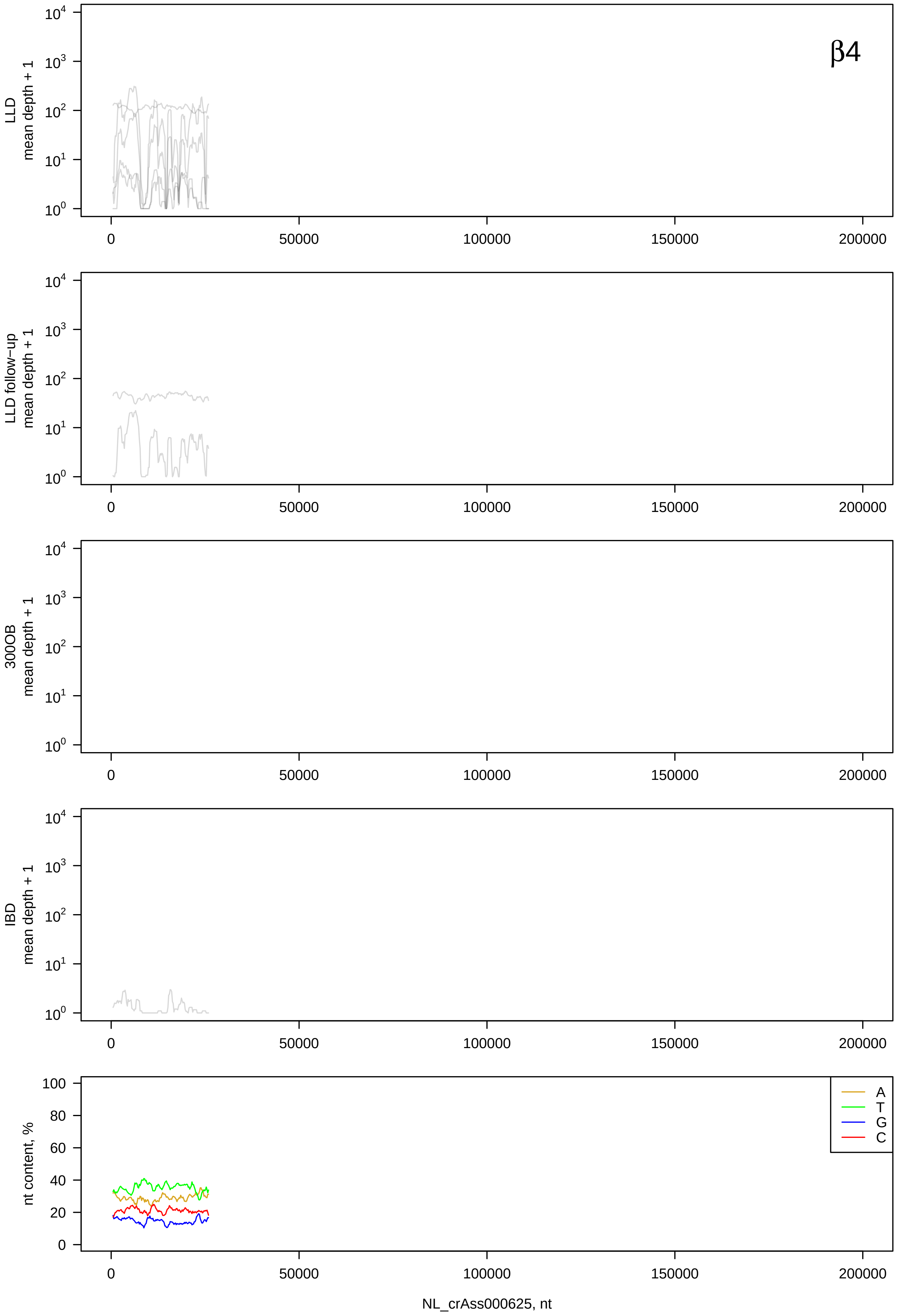
β4
LLD mean depth + 1
LLD follow−up mean depth + 1
300OB mean depth + 1
IBD mean depth + 1
nt content, %
A
T
G
C
NL_crAss000625, nt

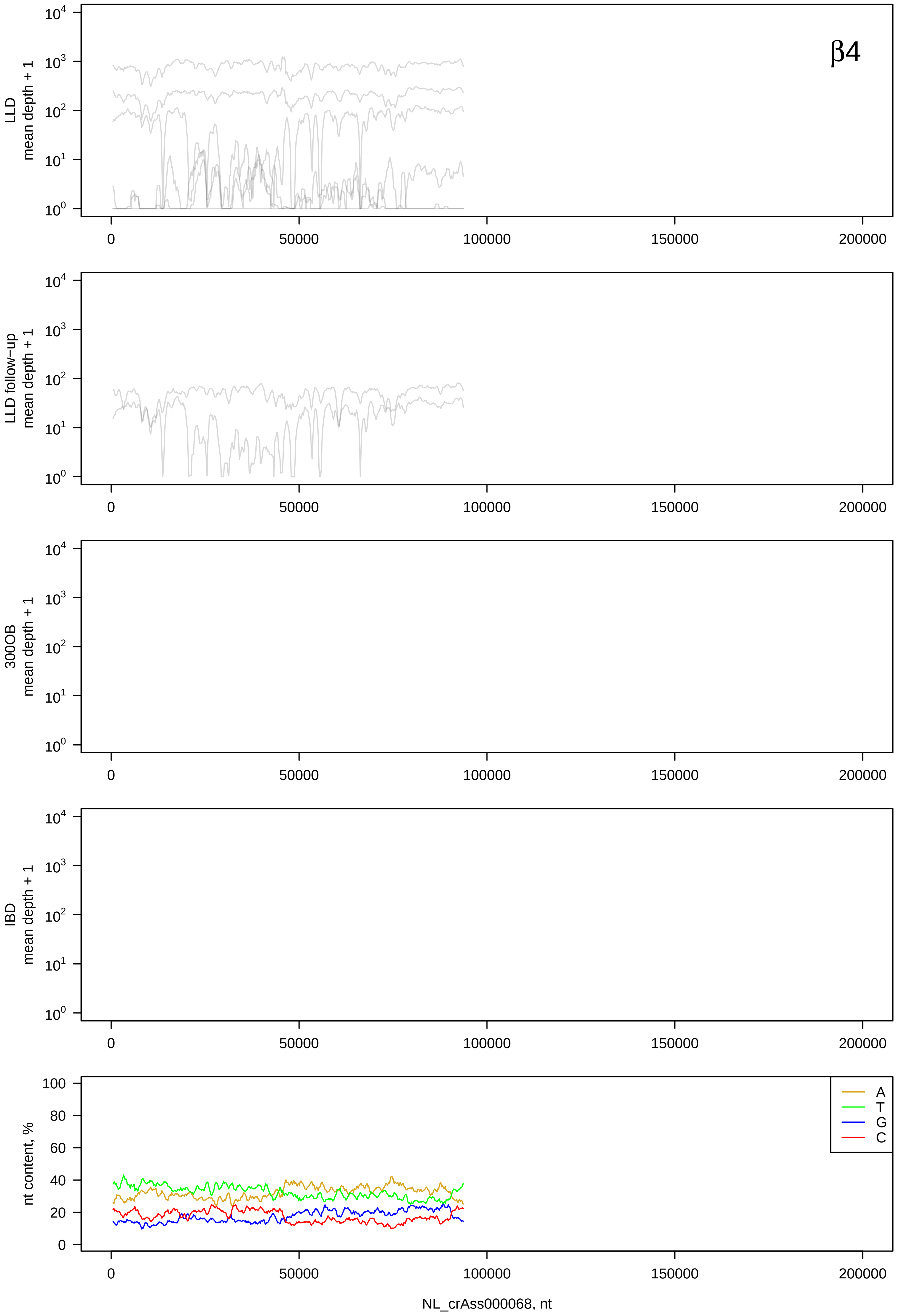

β4
LLD mean depth + 1
LLD follow−up mean depth + 1
300OB mean depth + 1
IBD mean depth + 1
nt content, %
A
T
G
C
0
50000
100000
150000
200000
NL_crAss000068, nt

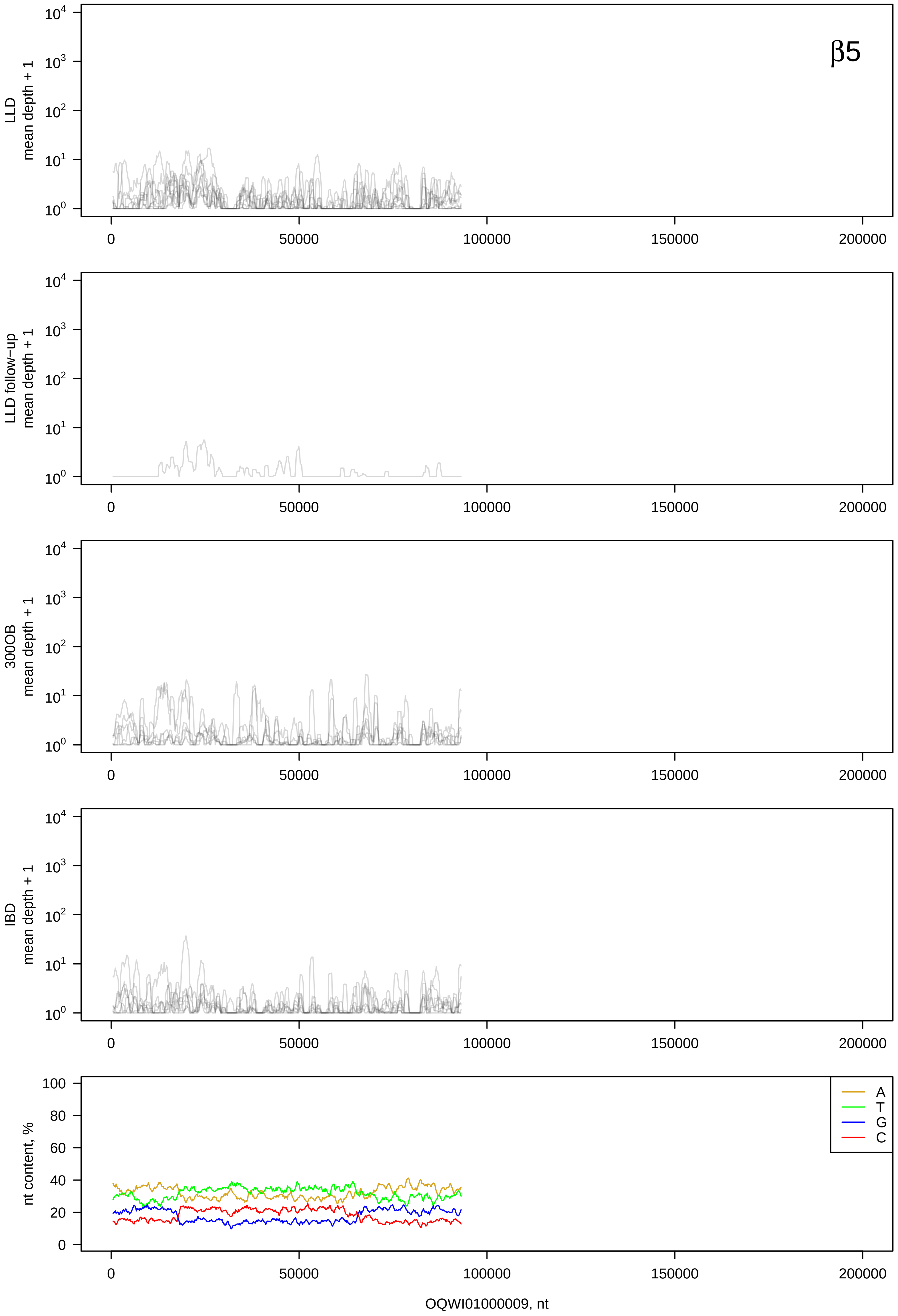

β5
LLD mean depth + 1
LLD follow−up mean depth + 1
300OB mean depth + 1
IBD mean depth + 1
nt content, %
A
T
G
C
0
50000
100000
150000
200000
OQWI01000009, nt

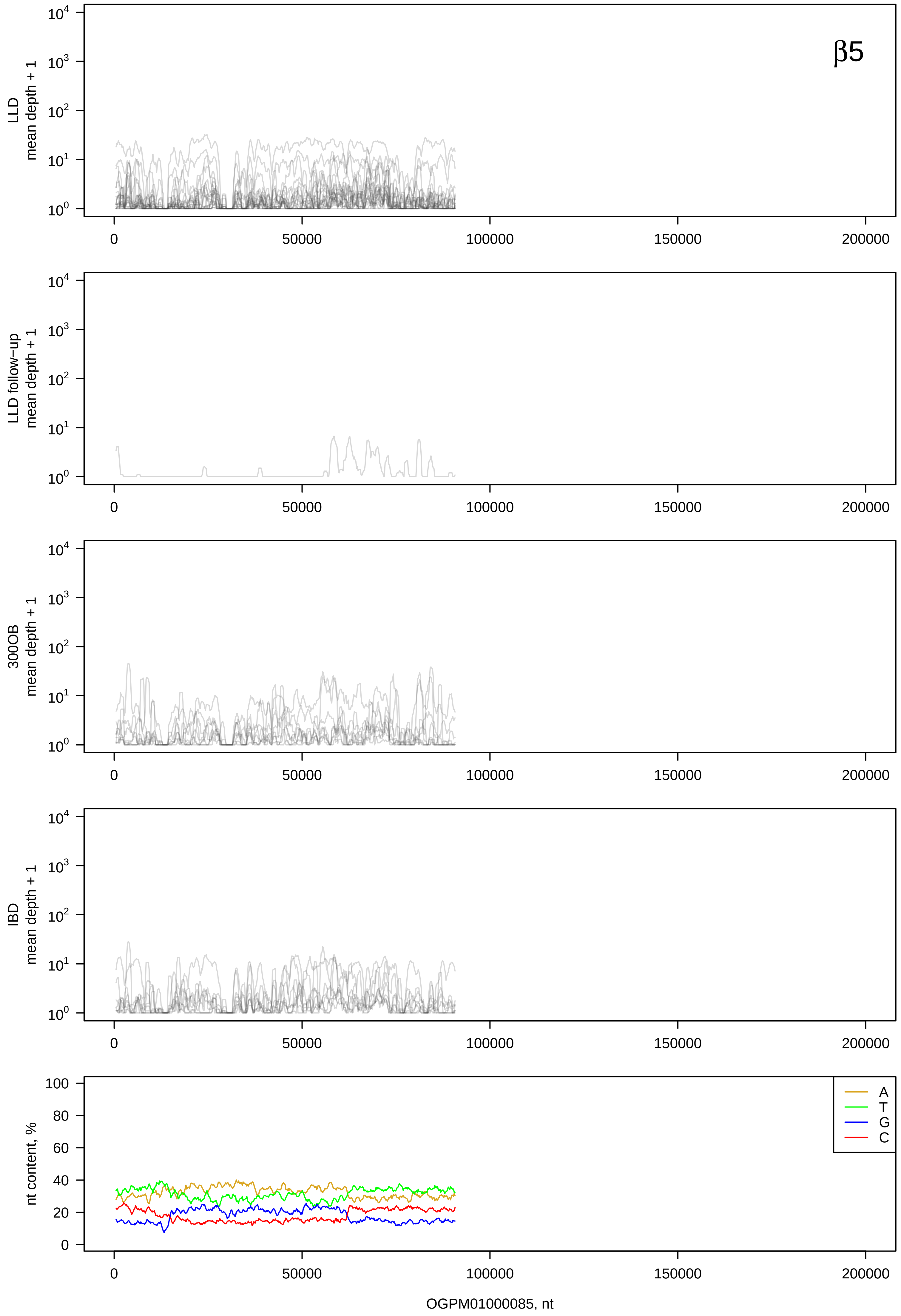

β5
LLD mean depth + 1
LLD follow−up mean depth + 1
300OB mean depth + 1
IBD mean depth + 1
nt content, %
A
T
G
C
0
50000
100000
150000
200000
OGPM01000085, nt

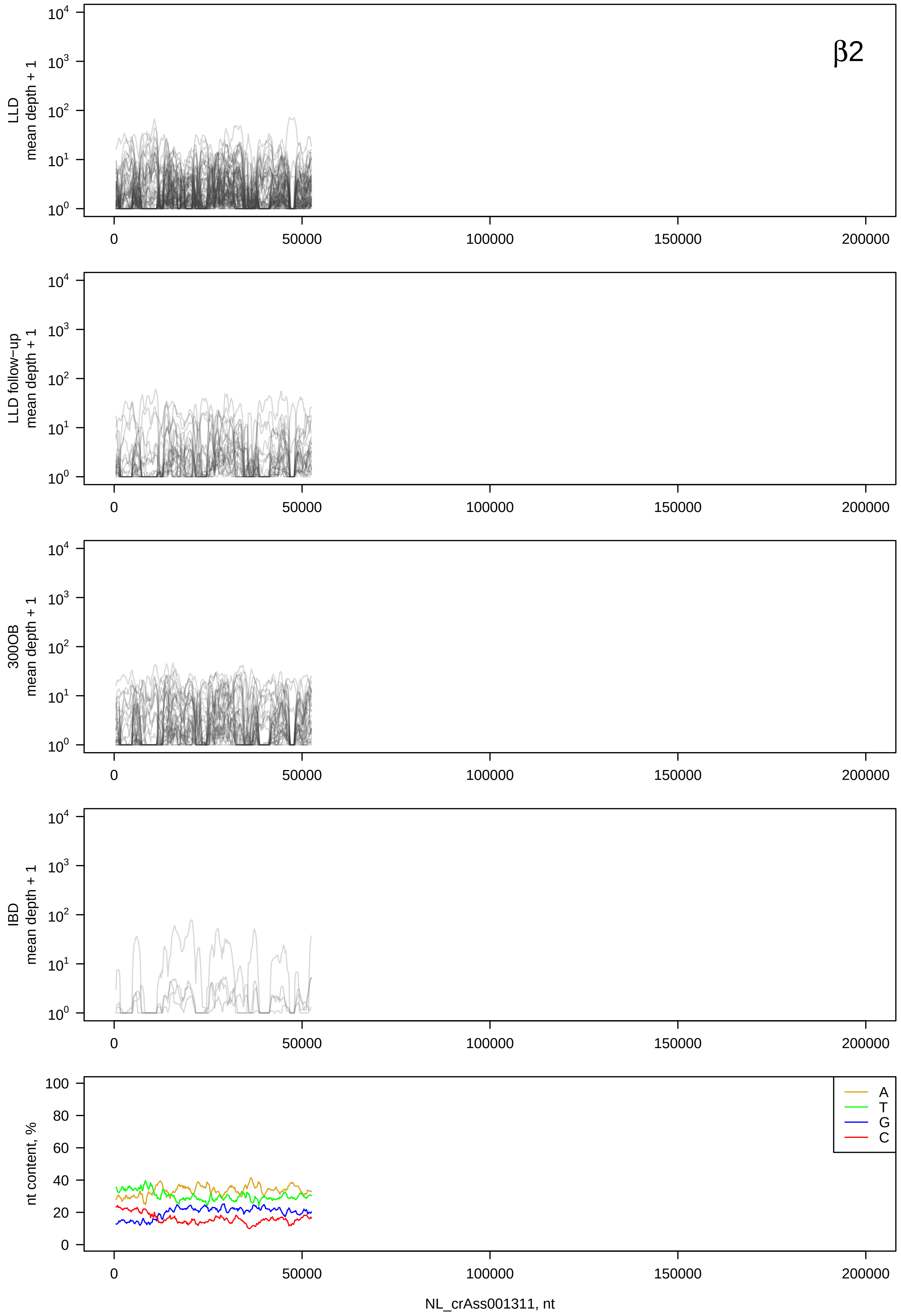
β2
LLD
mean depth + 1
LLD follow−up
mean depth + 1
300OB
mean depth + 1
IBD
mean depth + 1
nt content, %
A
T
G
C
NL_crAss001311, nt

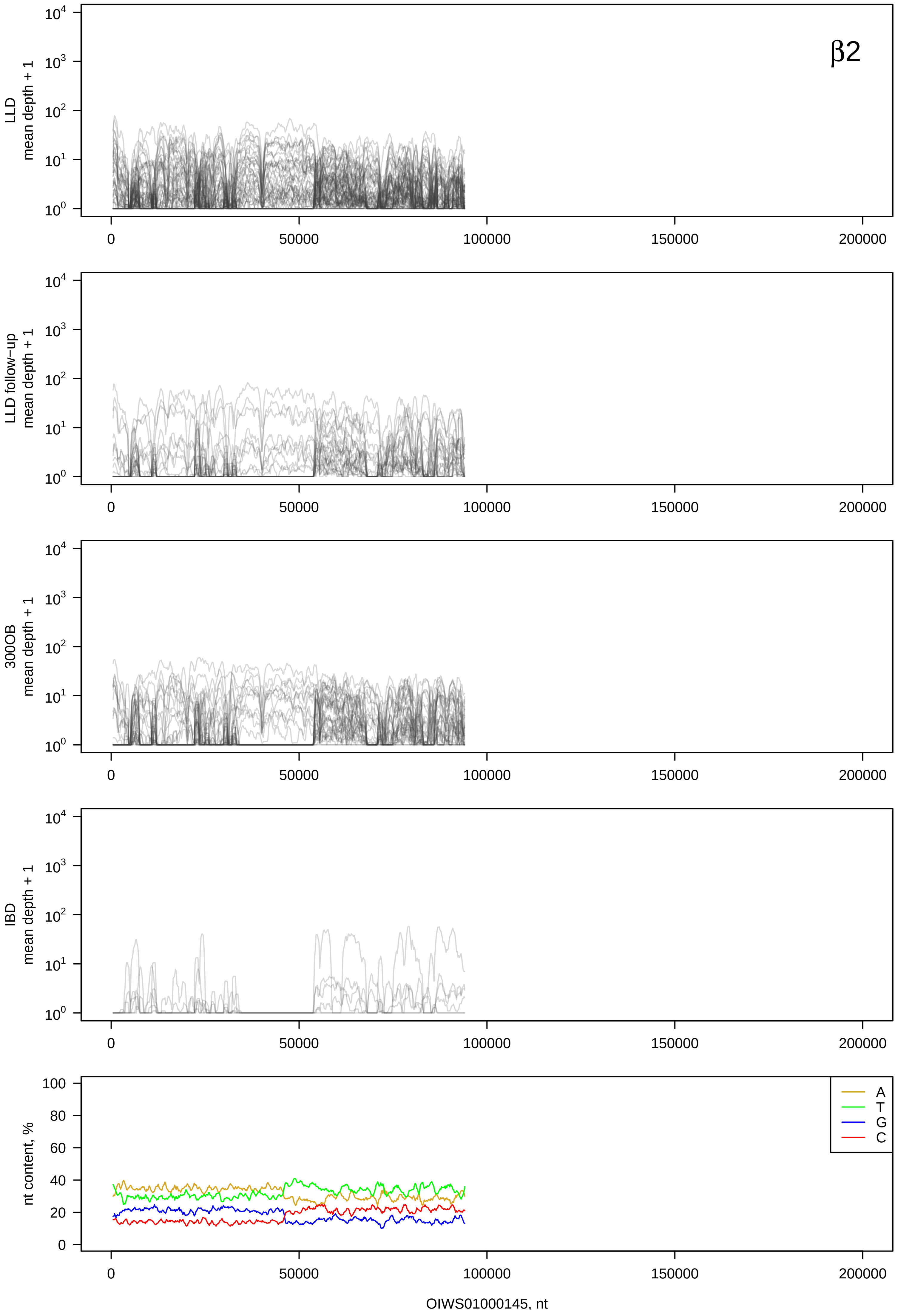

β2
LLD mean depth + 1
LLD follow-up mean depth + 1
300OB mean depth + 1
IBD mean depth + 1
nt content, %
A
T
G
C
OIWS01000145, nt

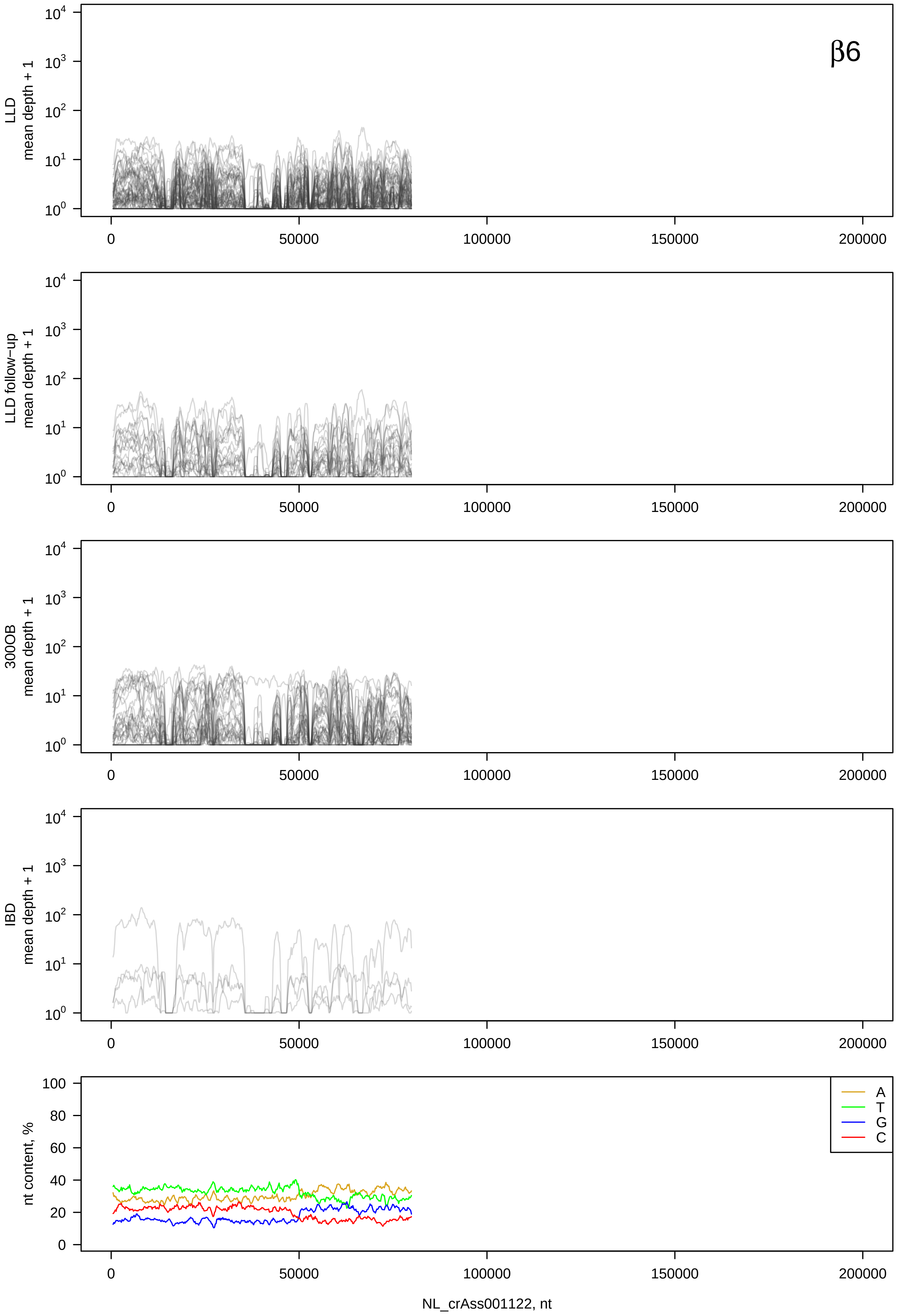

β6
LLD mean depth + 1
LLD follow-up mean depth + 1
300OB mean depth + 1
IBD mean depth + 1
nt content, %
A
T
G
C
0
50000
100000
150000
200000
NL_crAss001122, nt

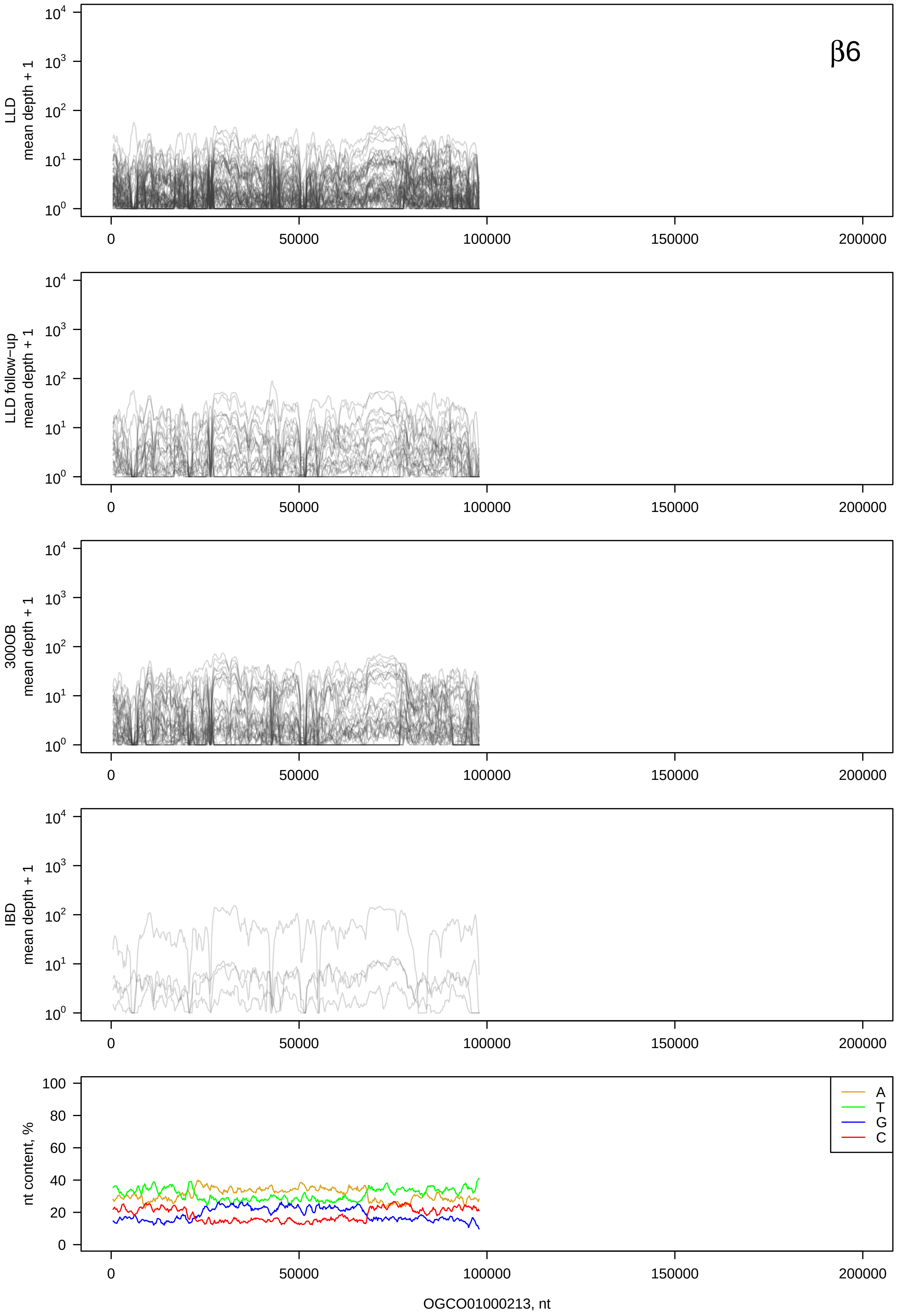
β6
LLD mean depth + 1
LLD follow-up mean depth + 1
300OB mean depth + 1
IBD mean depth + 1
nt content, %
A
T
G
C
0
50000
100000
150000
200000
OGCO01000213, nt

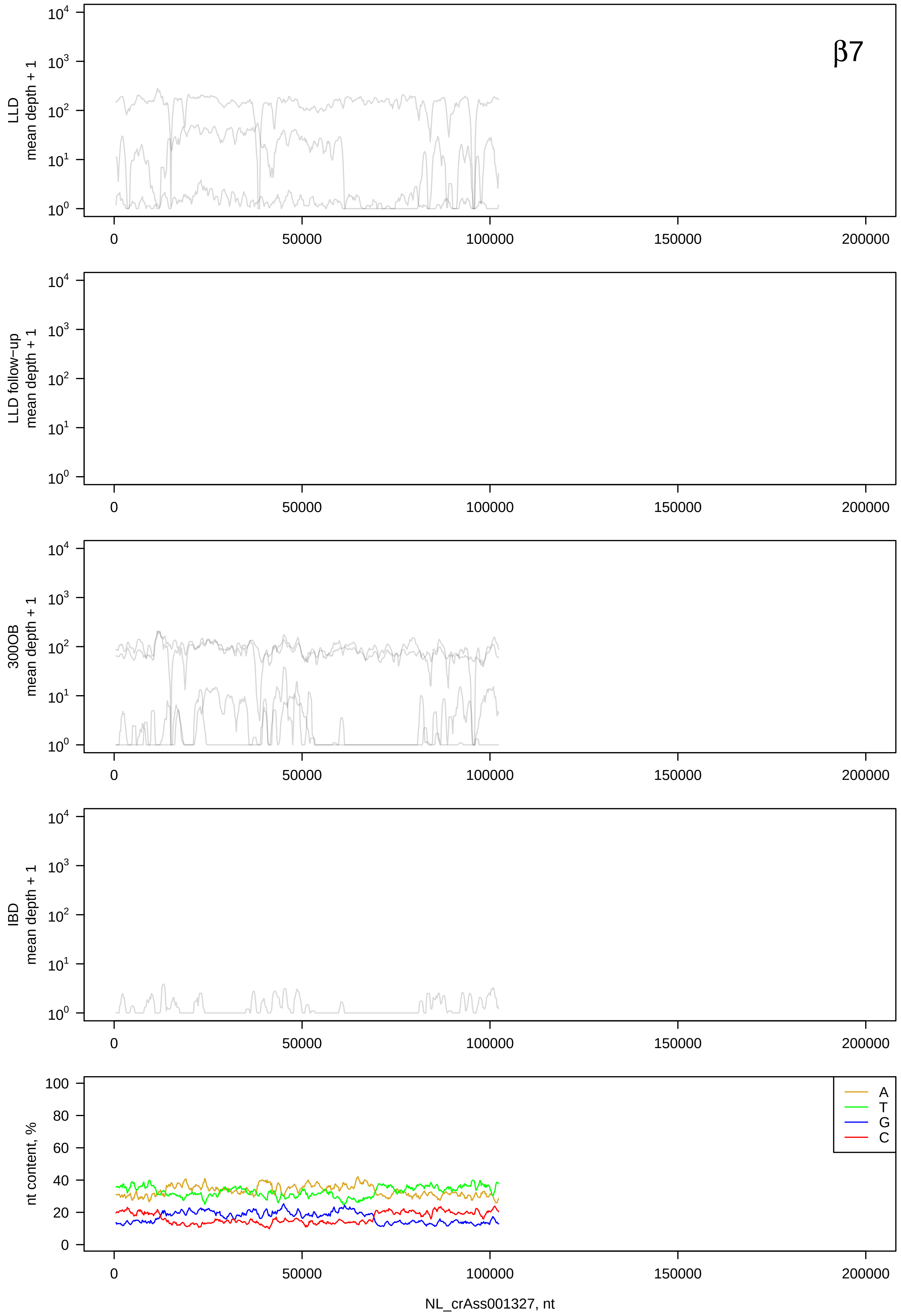

β7
LLD mean depth + 1
LLD follow−up mean depth + 1
300OB mean depth + 1
IBD mean depth + 1
nt content, %
A
T
G
C
NL_crAss001327, nt

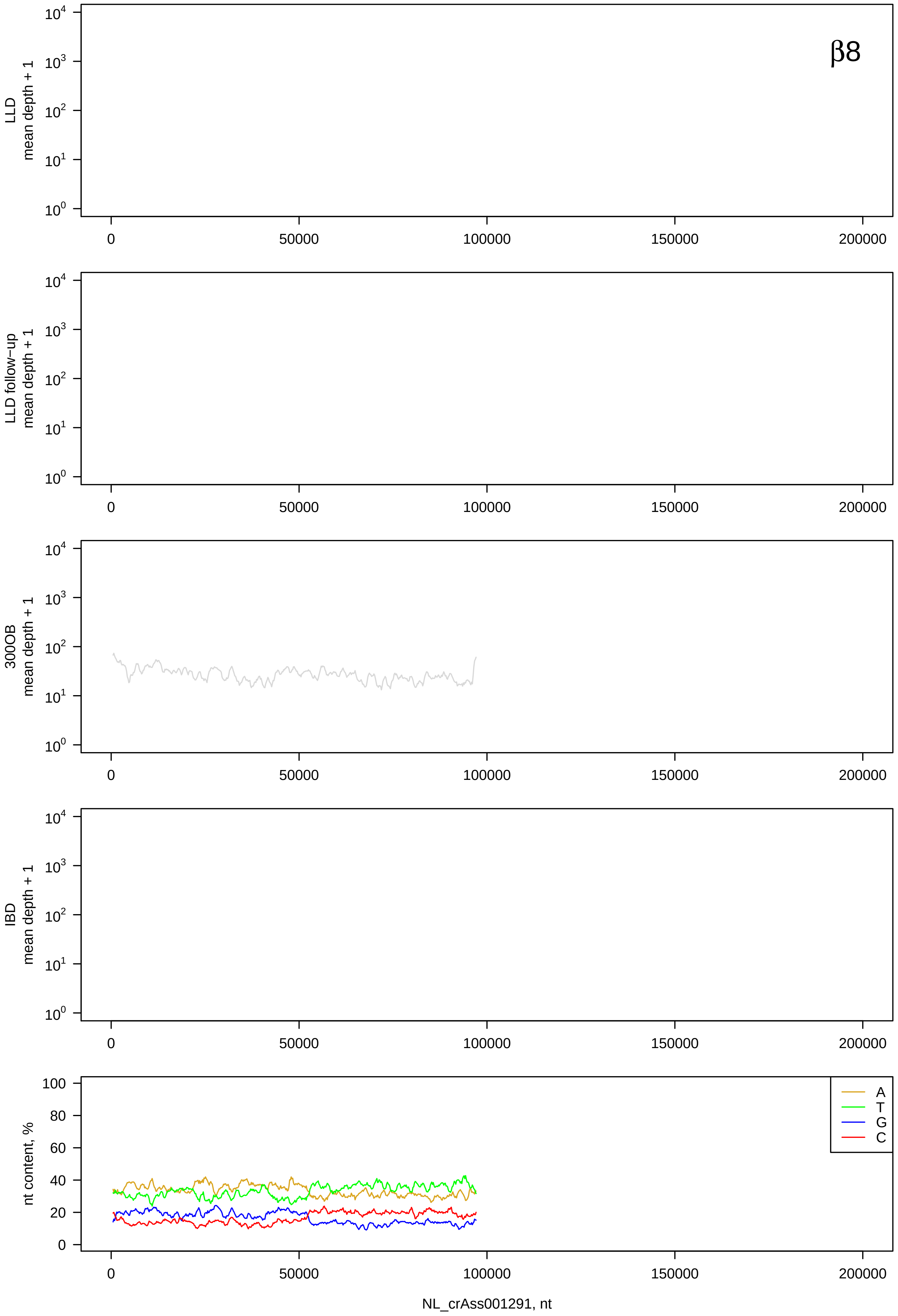
β8
LLD mean depth + 1
LLD follow−up mean depth + 1
300OB mean depth + 1
IBD mean depth + 1
nt content, %
A
T
G
C
NL_crAss001291, nt

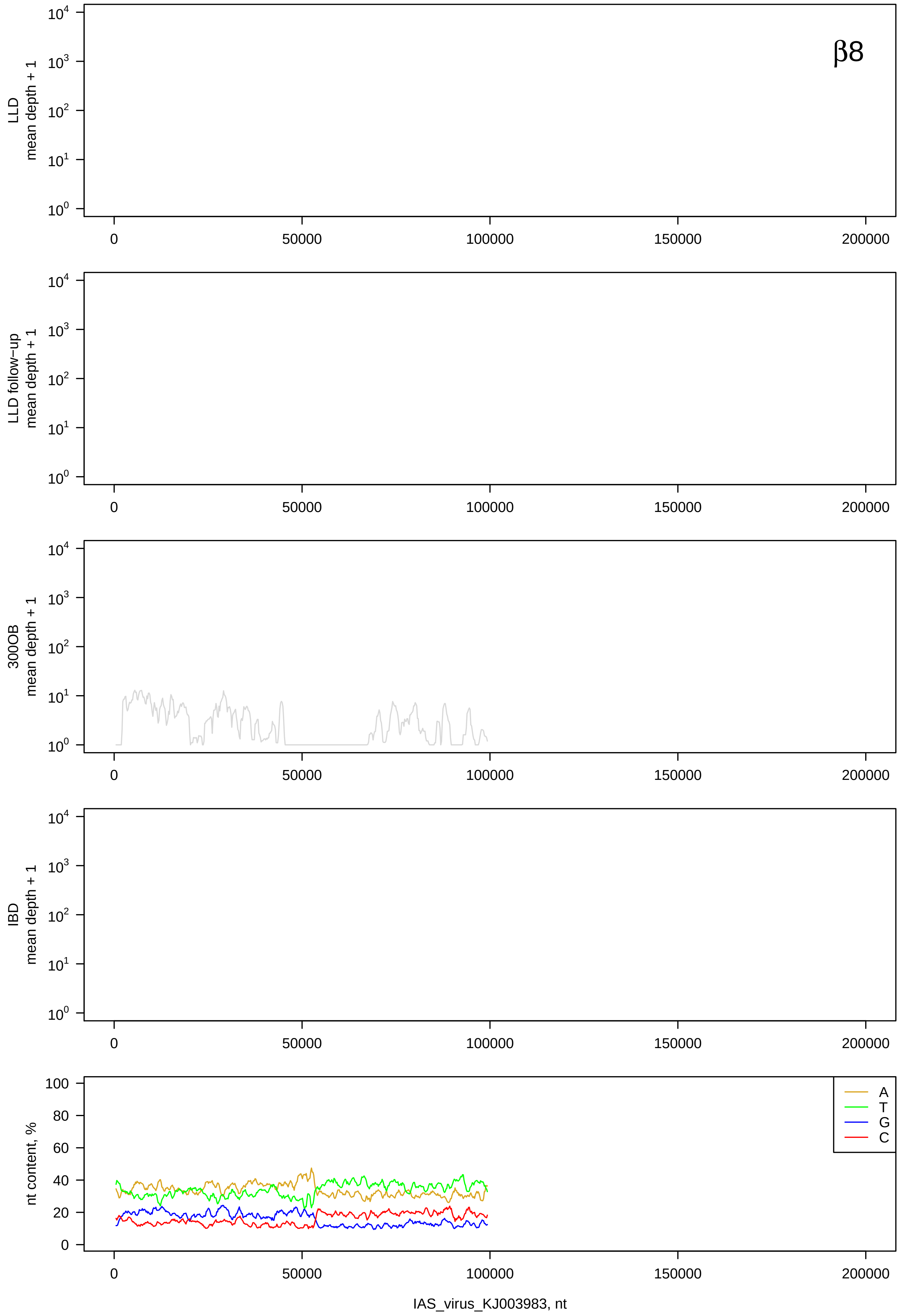
β8
LLD mean depth + 1
LLD follow-up mean depth + 1
300OB mean depth + 1
IBD mean depth + 1
nt content, %
A
T
G
C
IAS_virus_KJ003983, nt

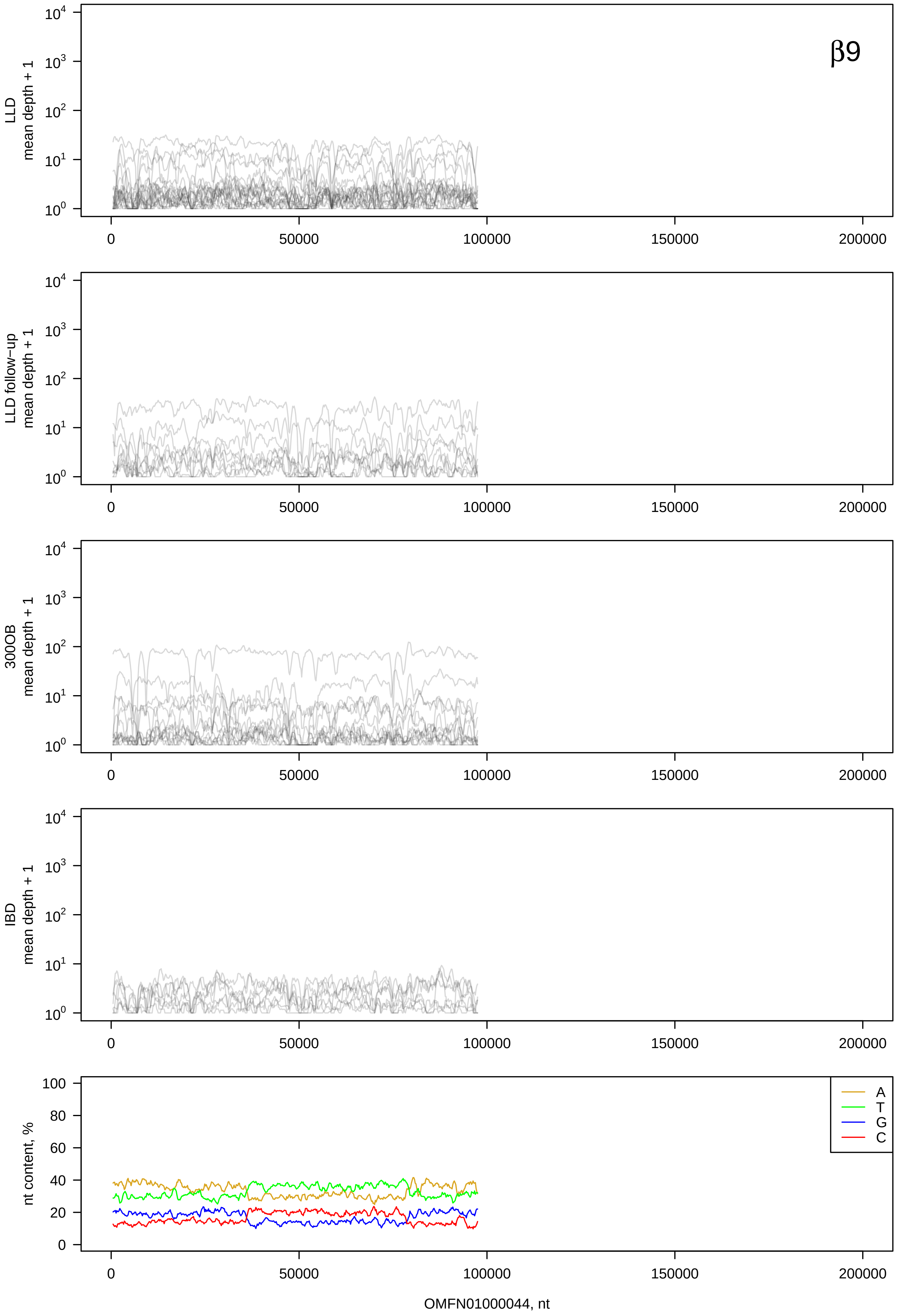

β9
LLD
mean depth + 1
LLD follow−up
mean depth + 1
300OB
mean depth + 1
IBD
mean depth + 1
nt content, %
A
T
G
C
OMFN01000044, nt

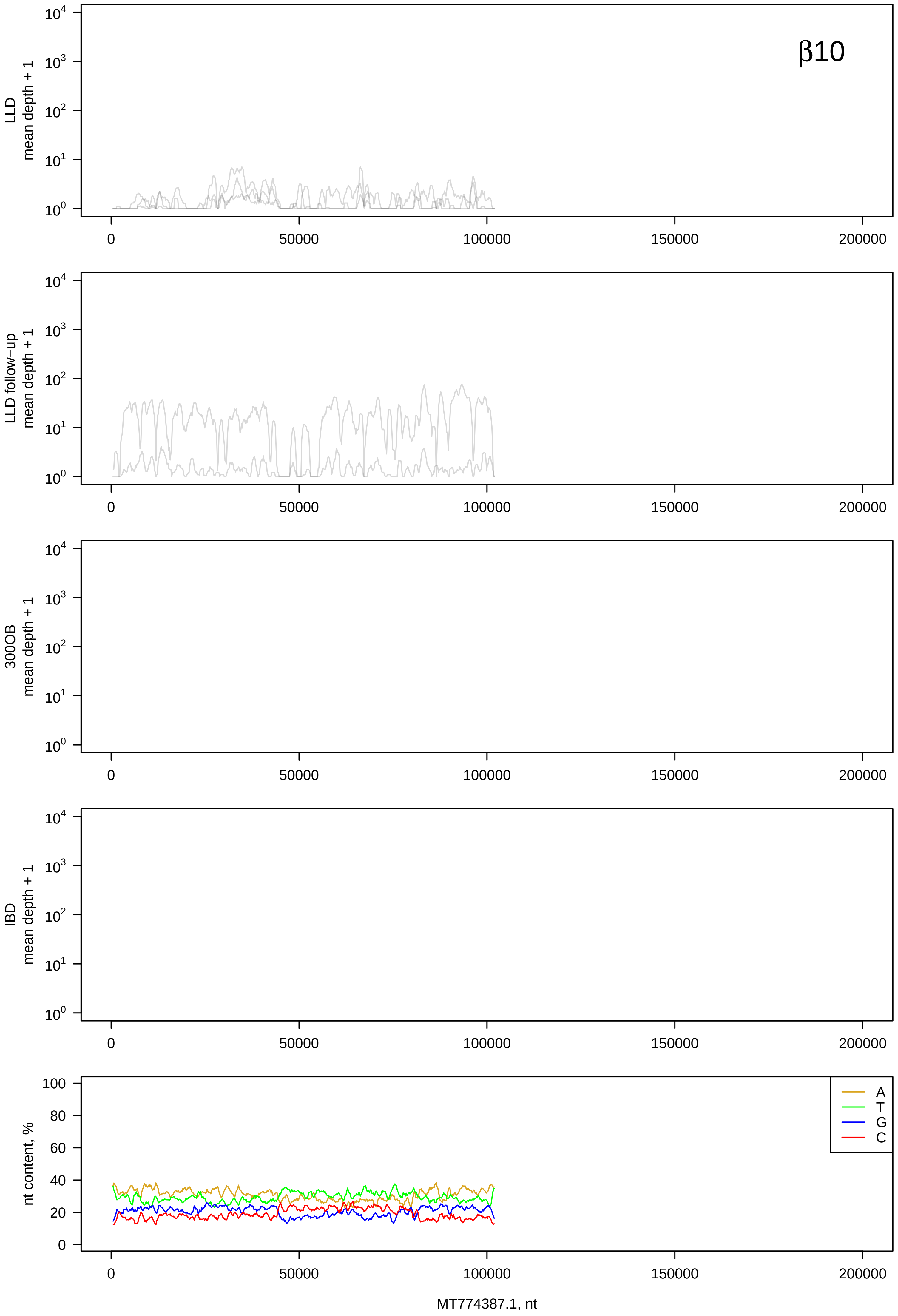

β10
LLD mean depth + 1
LLD follow-up mean depth + 1
300OB mean depth + 1
IBD mean depth + 1
nt content, %
A
T
G
C
MT774387.1, nt

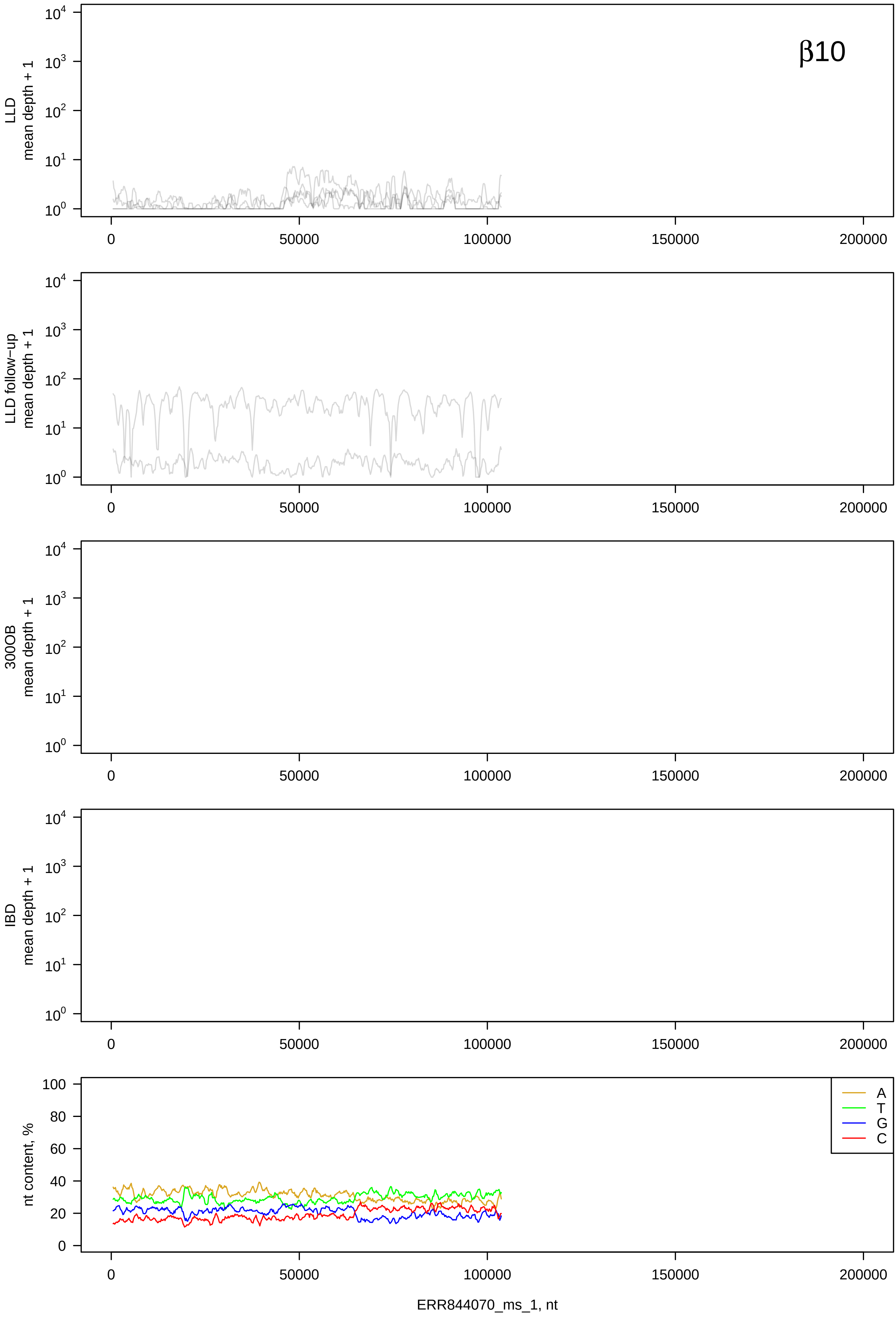
β10
LLD mean depth + 1
LLD follow-up mean depth + 1
300OB mean depth + 1
IBD mean depth + 1
nt content, %
A
T
G
C
ERR844070_ms_1, nt

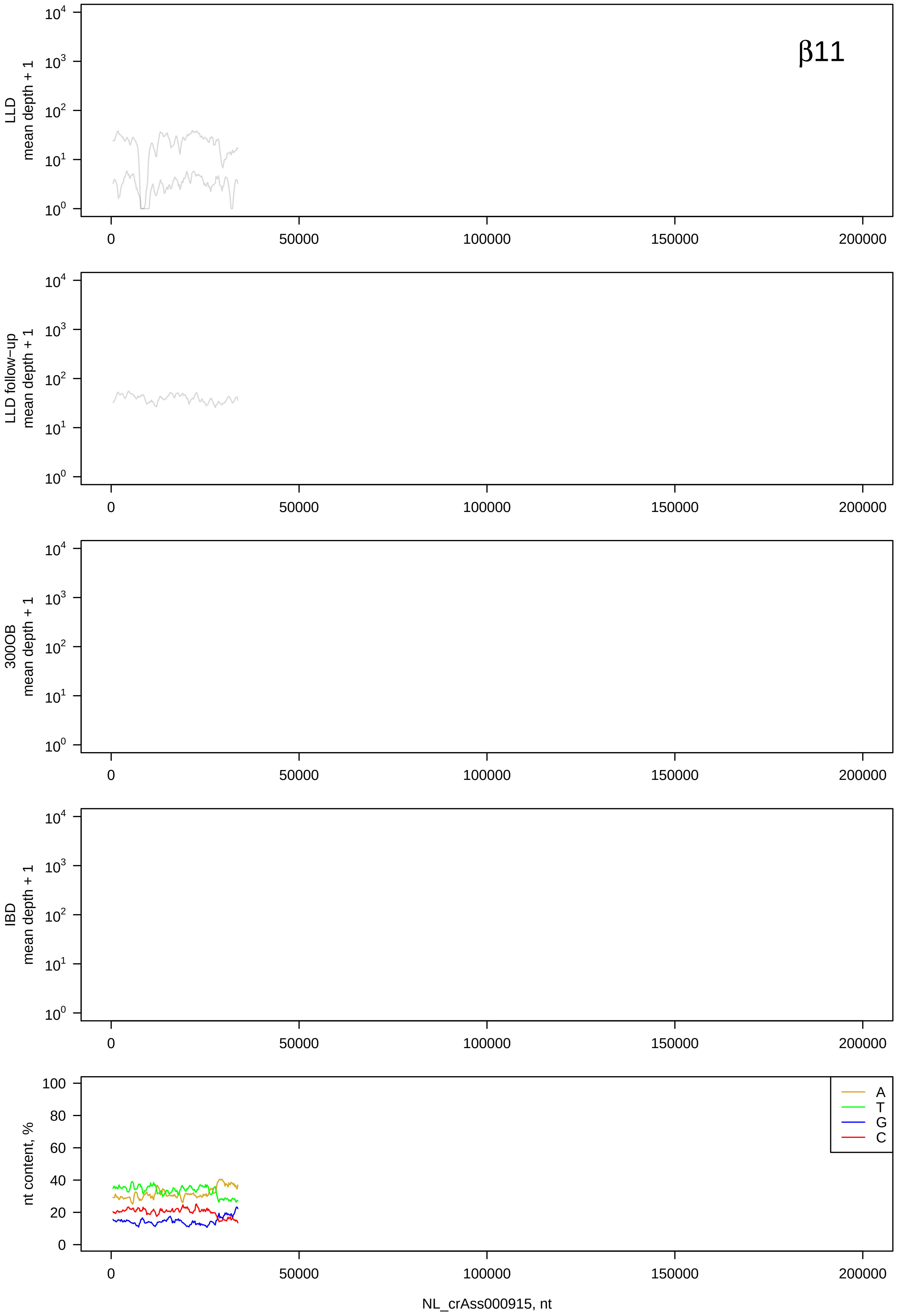
β11
LLD
mean depth + 1
LLD follow-up
mean depth + 1
300OB
mean depth + 1
IBD
mean depth + 1
nt content, %
A
T
G
C
NL_crAss000915, nt

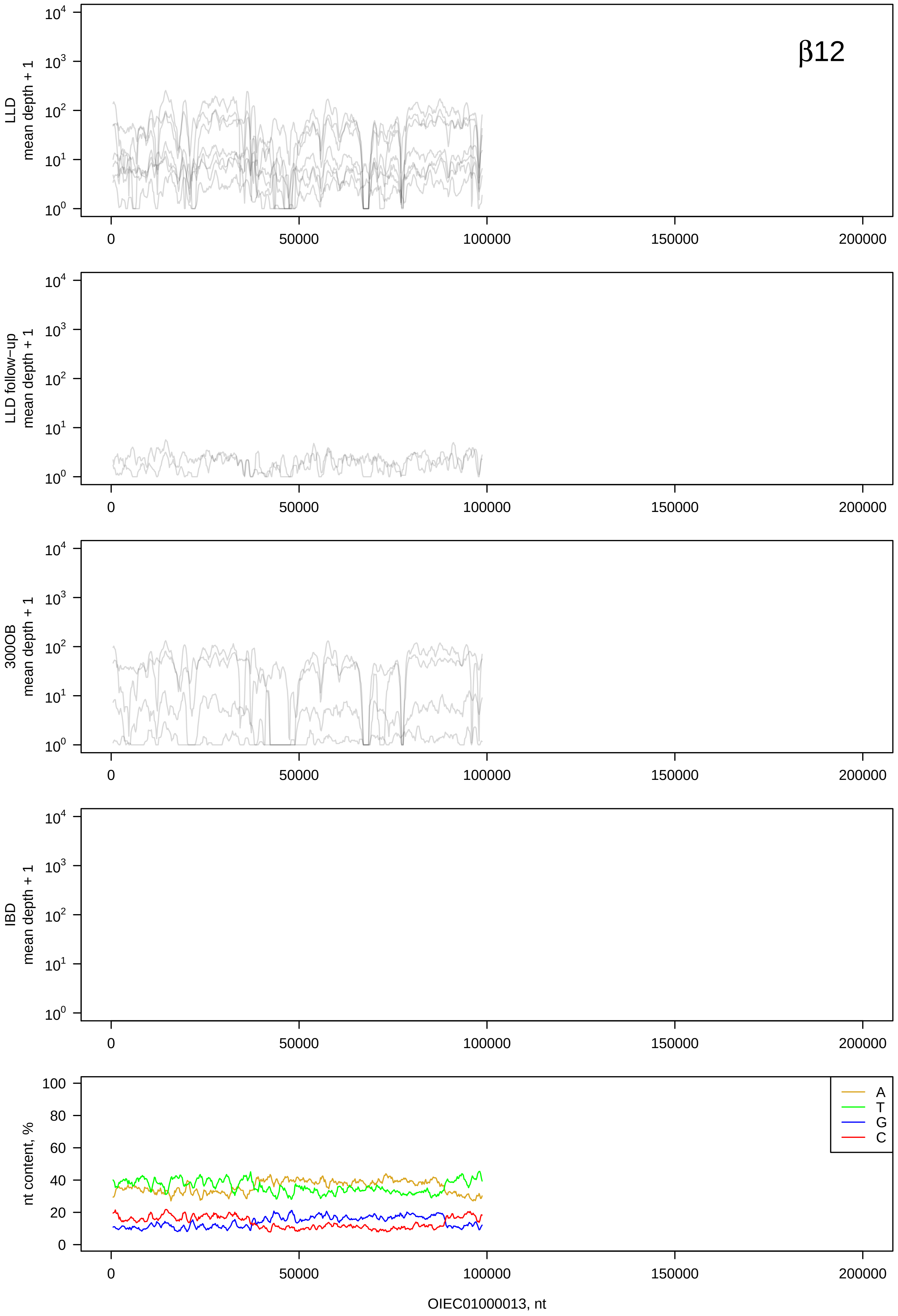

β12
LLD
mean depth + 1
LLD follow−up
mean depth + 1
300OB
mean depth + 1
IBD
mean depth + 1
nt content, %
A
T
G
C
0
50000
100000
150000
200000
OIEC01000013, nt

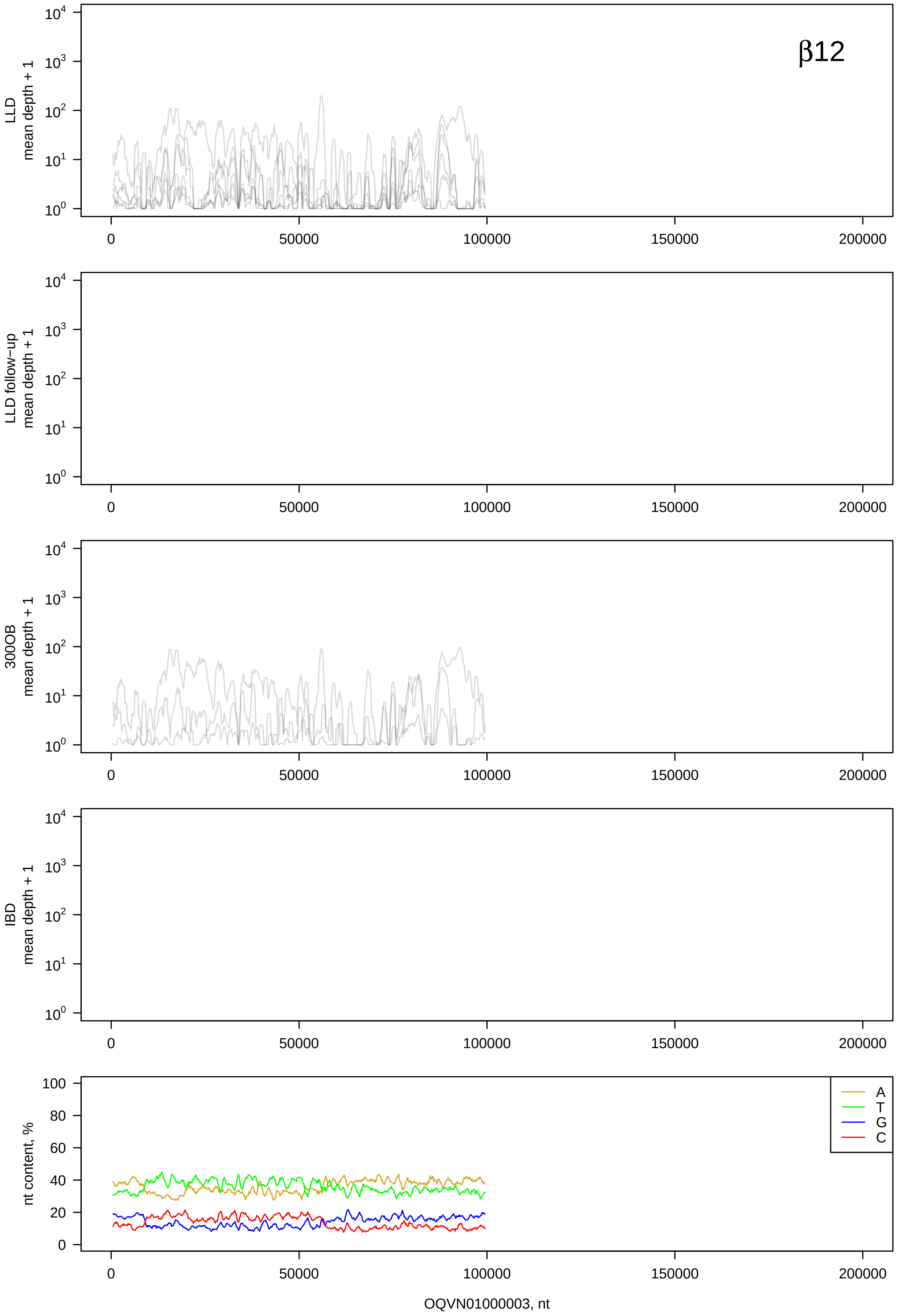
β12
LLD mean depth + 1
LLD follow−up mean depth + 1
300OB mean depth + 1
IBD mean depth + 1
nt content, %
A
T
G
C
OQVN01000003, nt

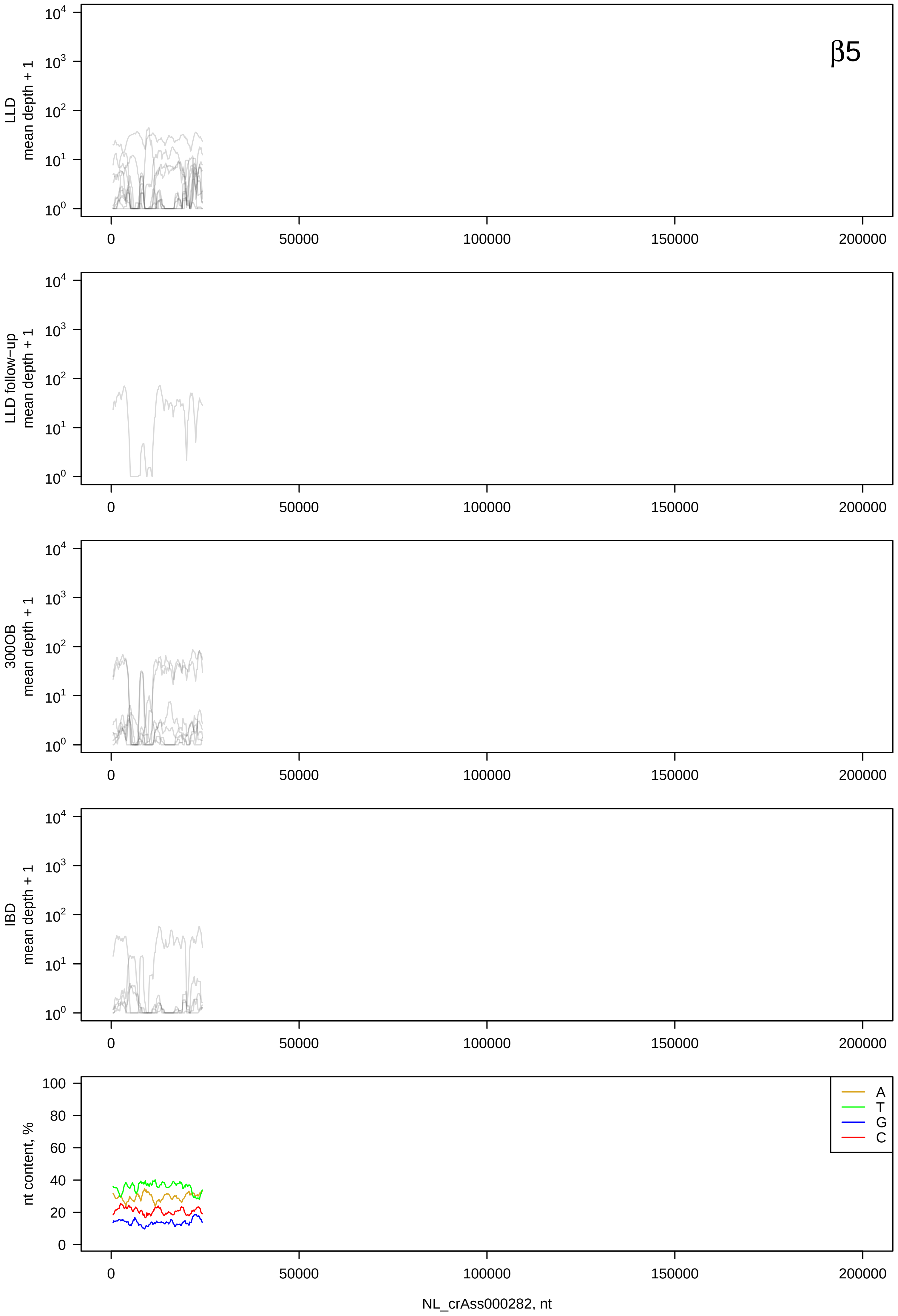
β5
LLD mean depth + 1
LLD follow-up mean depth + 1
300OB mean depth + 1
IBD mean depth + 1
nt content, %
A
T
G
C
NL_crAss000282, nt

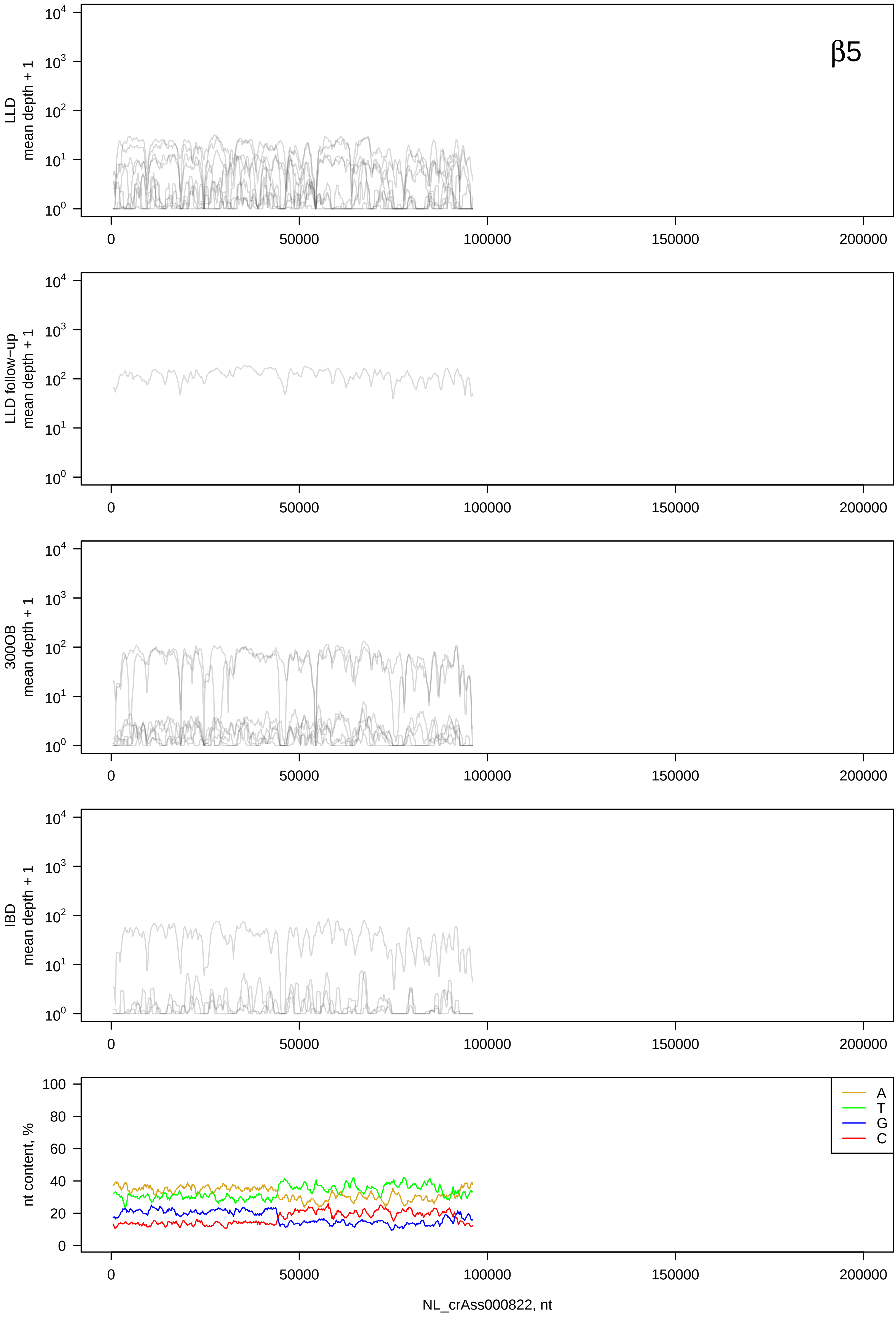
β5
LLD
mean depth + 1
LLD follow−up
mean depth + 1
300OB
mean depth + 1
IBD
mean depth + 1
nt content, %
A
T
G
C
NL_crAss000822, nt

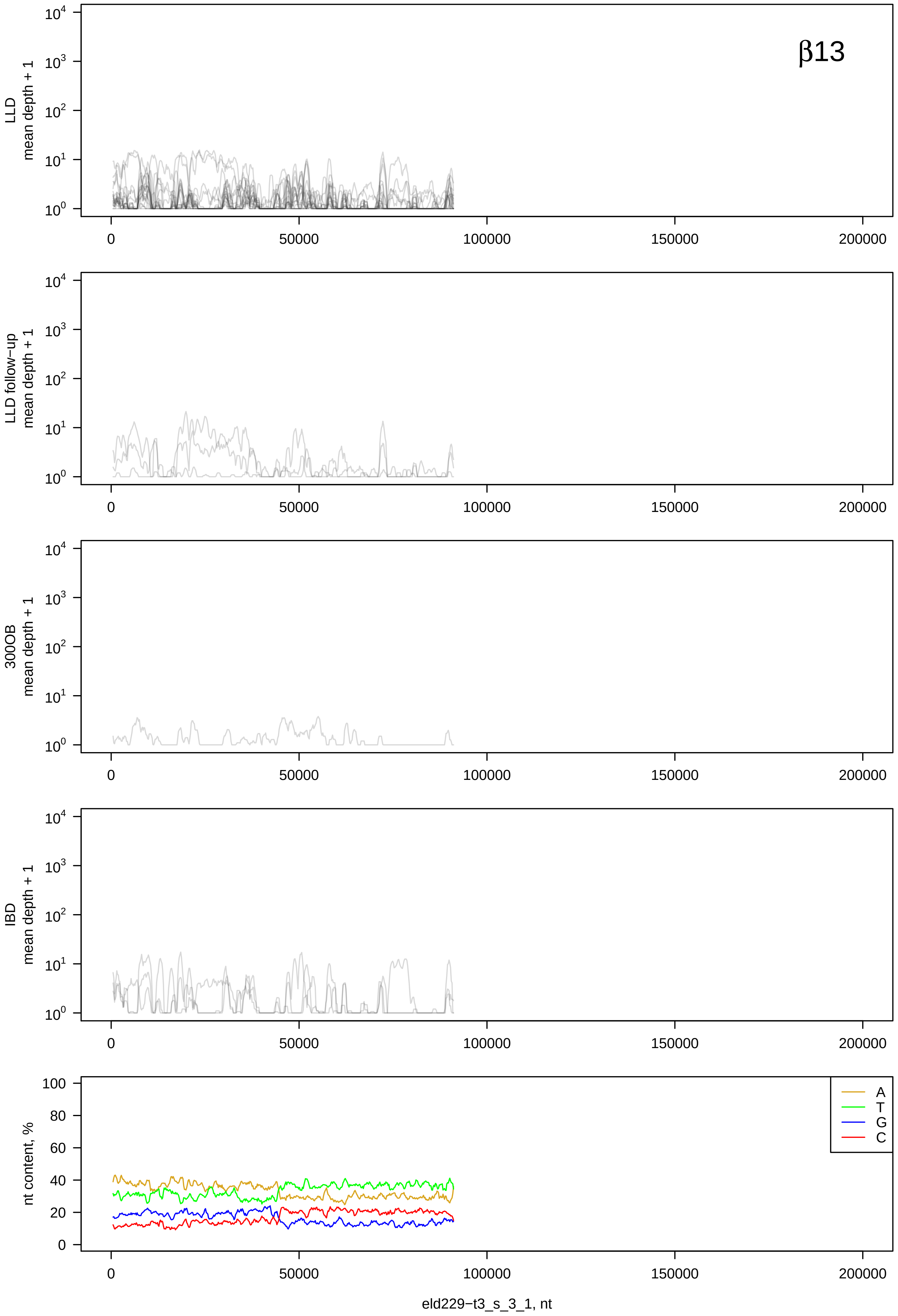
β13
LLD
mean depth + 1
LLD follow−up
mean depth + 1
300OB
mean depth + 1
IBD
mean depth + 1
nt content, %
A
T
G
C
0
50000
100000
150000
200000
eld229−t3_s_3_1, nt

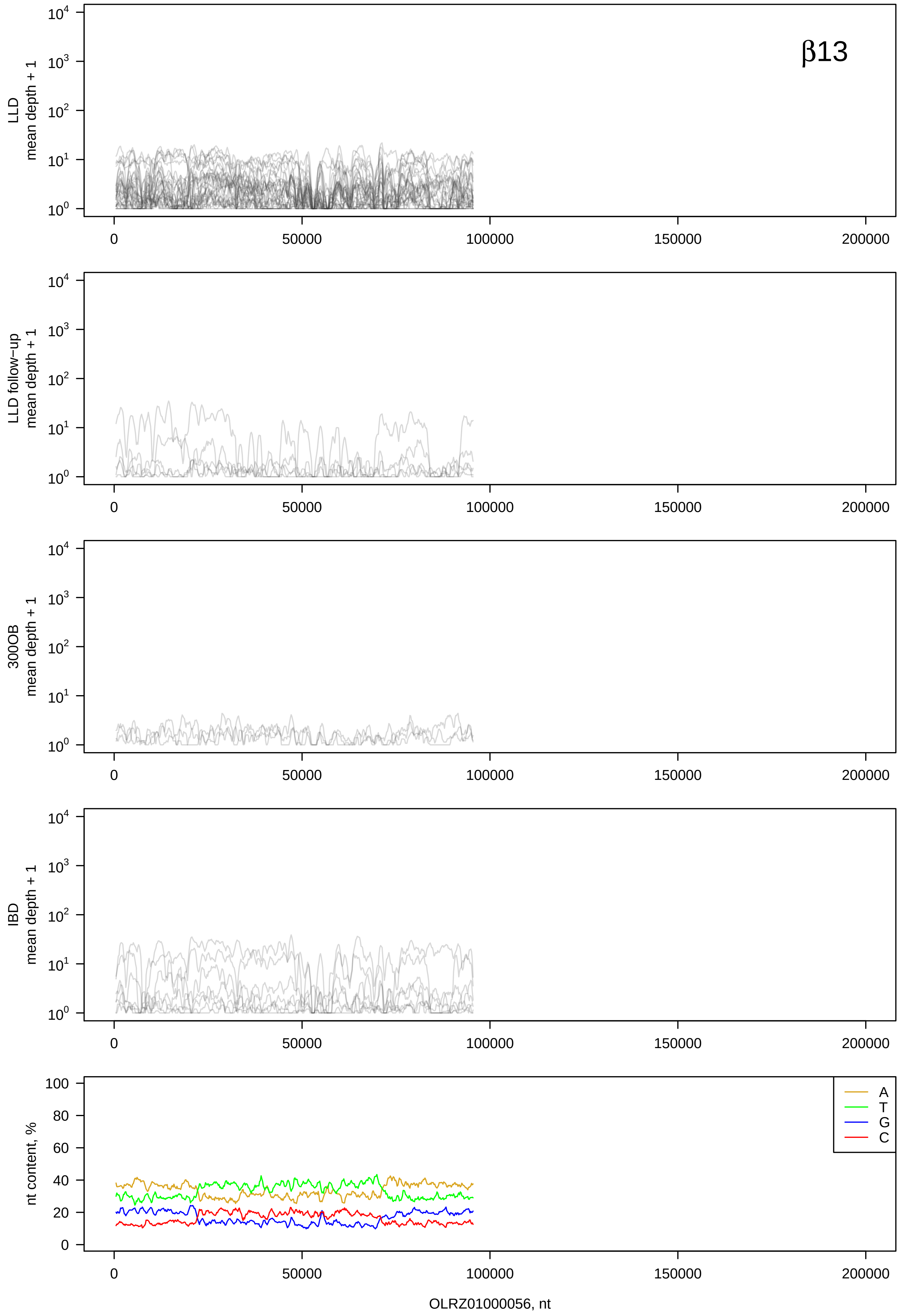

β13
LLD mean depth + 1
LLD follow−up mean depth + 1
300OB mean depth + 1
IBD mean depth + 1
nt content, %
A
T
G
C
0
50000
100000
150000
200000
OLRZ01000056, nt

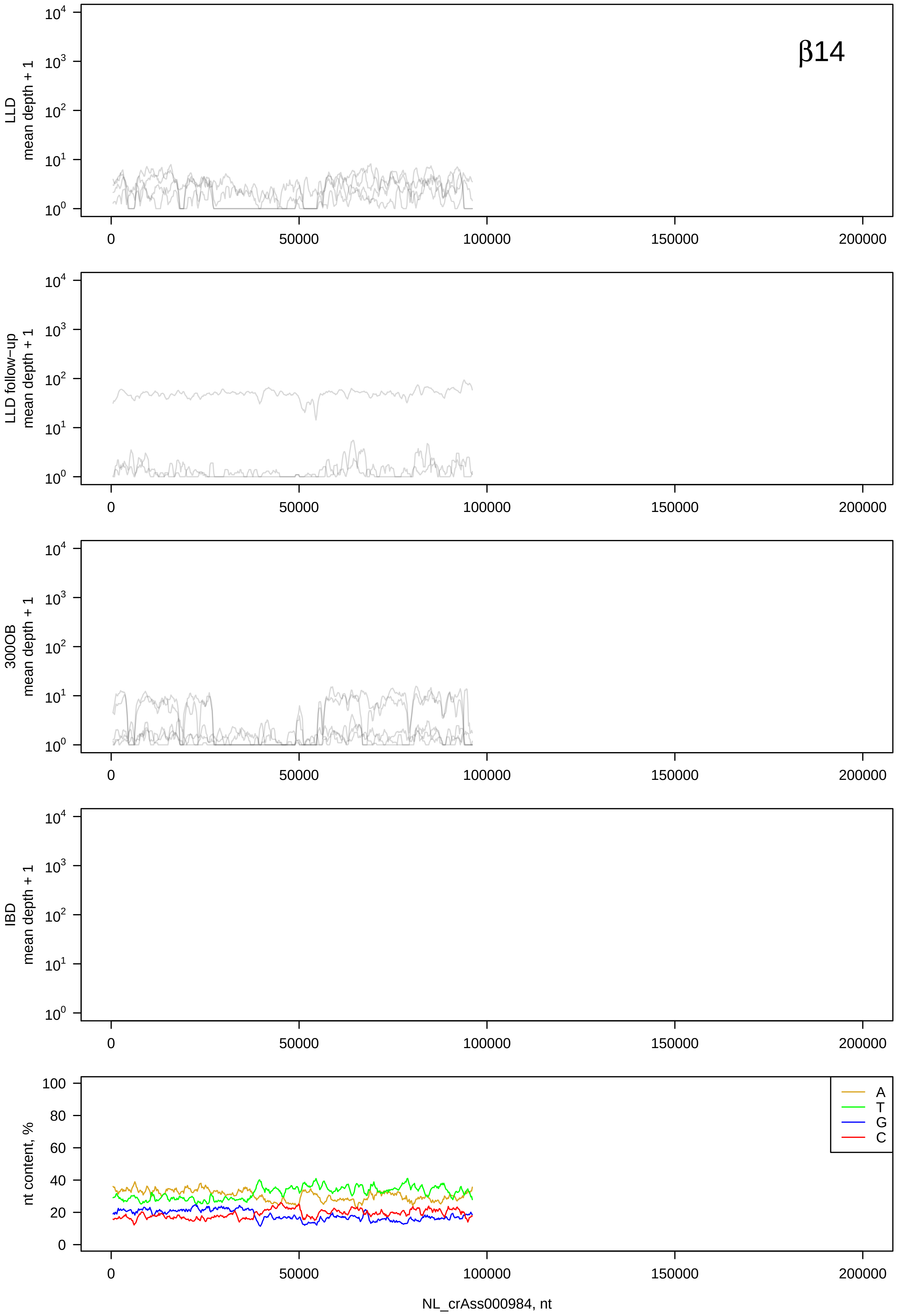
β14
LLD mean depth + 1
LLD follow-up mean depth + 1
300OB mean depth + 1
IBD mean depth + 1
nt content, %
A
T
G
C
NL_crAss000984, nt

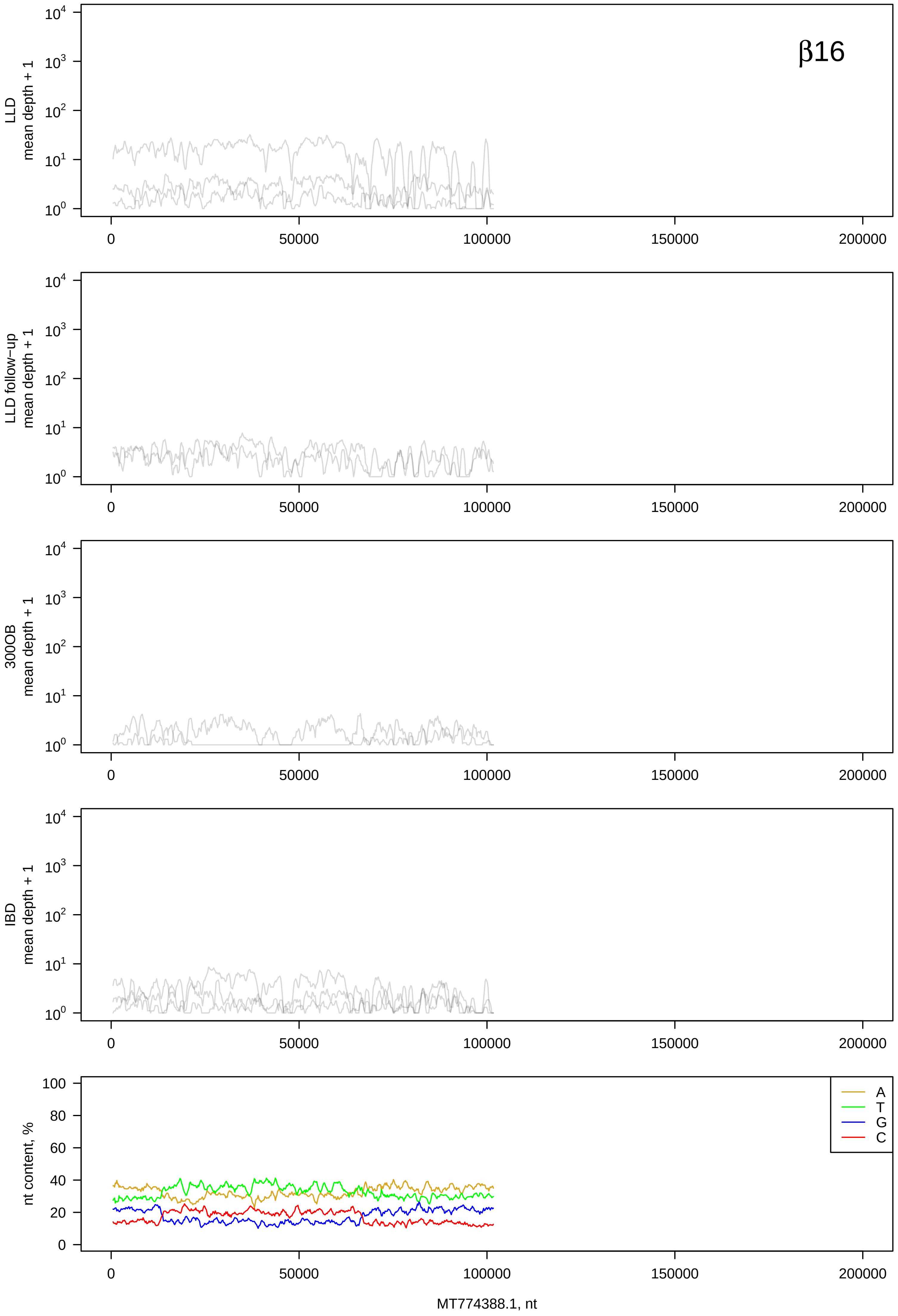
β16
LLD
mean depth + 1
LLD follow−up
mean depth + 1
300OB
mean depth + 1
IBD
mean depth + 1
nt content, %
A
T
G
C
MT774388.1, nt

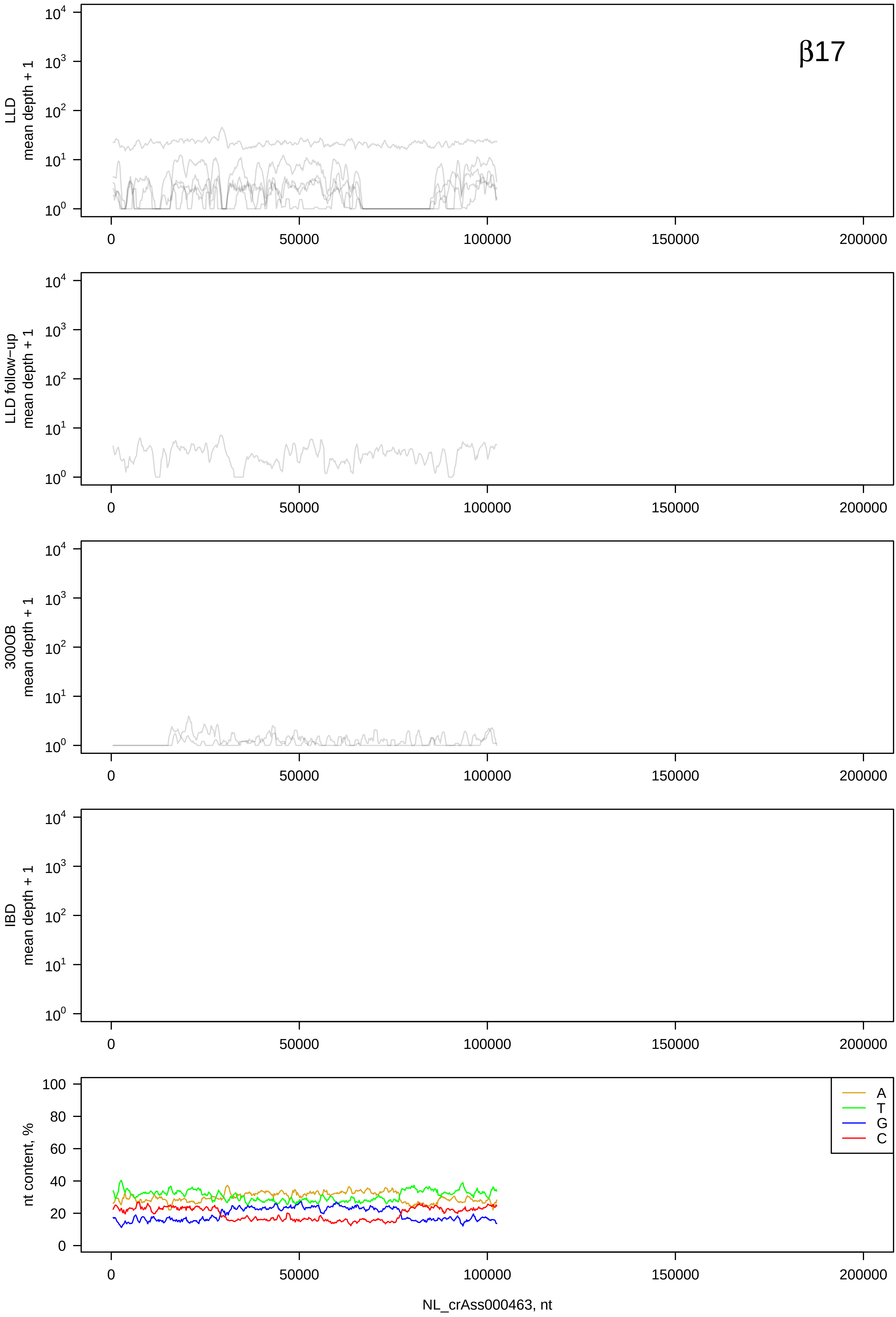

β17
LLD mean depth + 1
LLD follow-up mean depth + 1
300OB mean depth + 1
IBD mean depth + 1
nt content, %
A
T
G
C
0
50000
100000
150000
200000
NL_crAss000463, nt

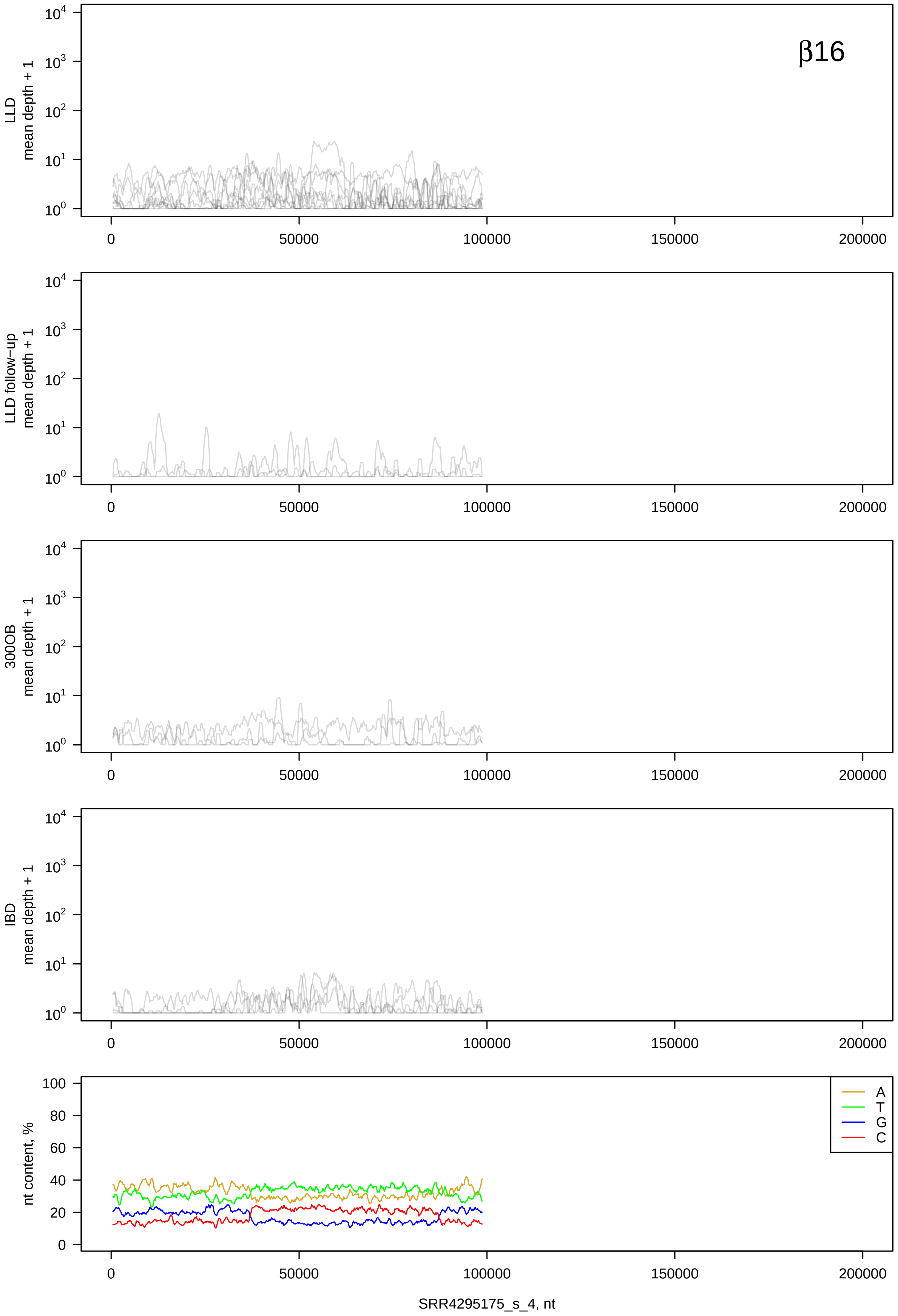

β16
LLD mean depth + 1
LLD follow-up mean depth + 1
300OB mean depth + 1
IBD mean depth + 1
nt content, %
A
T
G
C
0
50000
100000
150000
200000
SRR4295175_s_4, nt

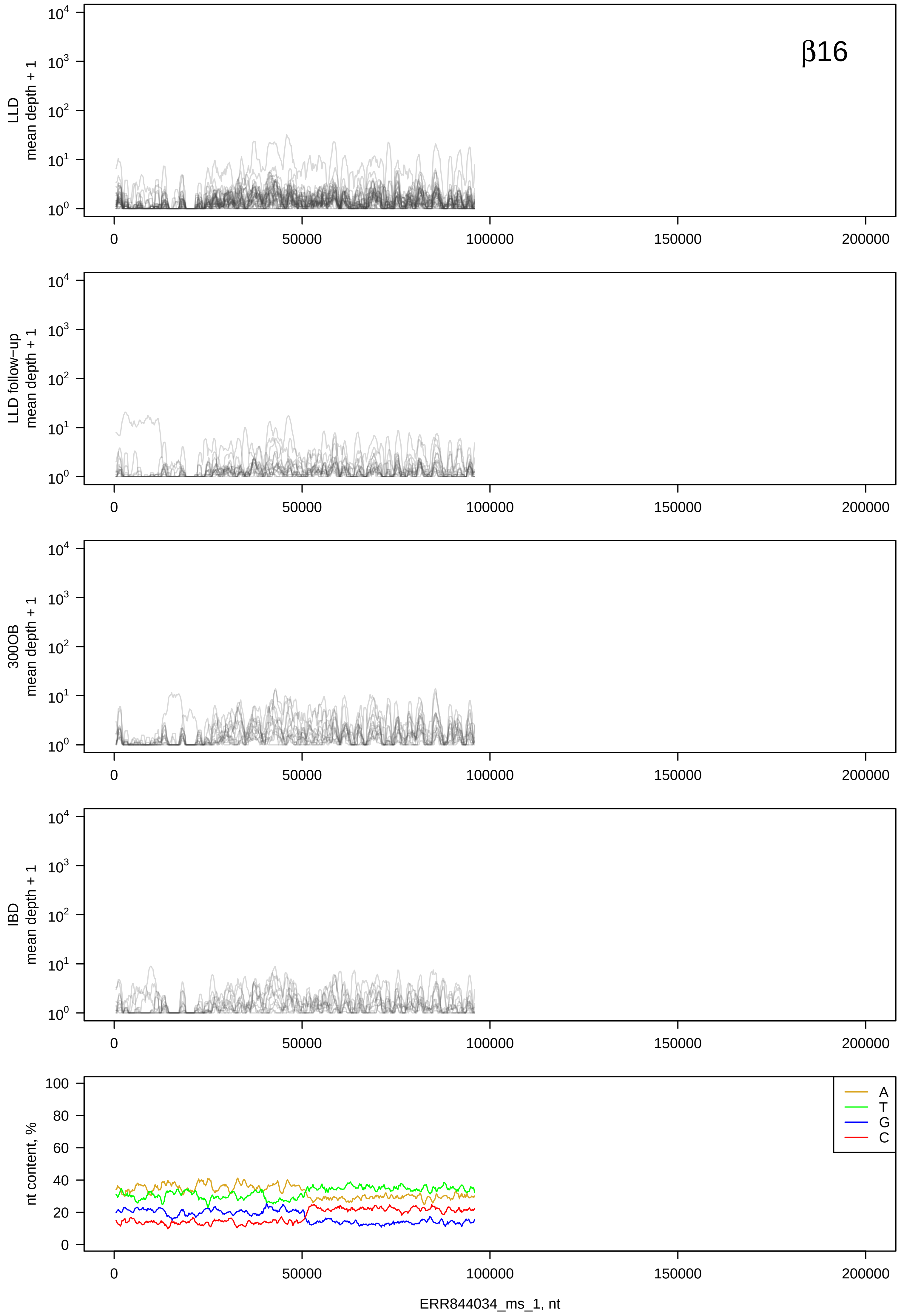
β16
LLD
mean depth + 1
LLD follow−up
mean depth + 1
300OB
mean depth + 1
IBD
mean depth + 1
nt content, %
A
T
G
C
0
50000
100000
150000
200000
ERR844034_ms_1, nt

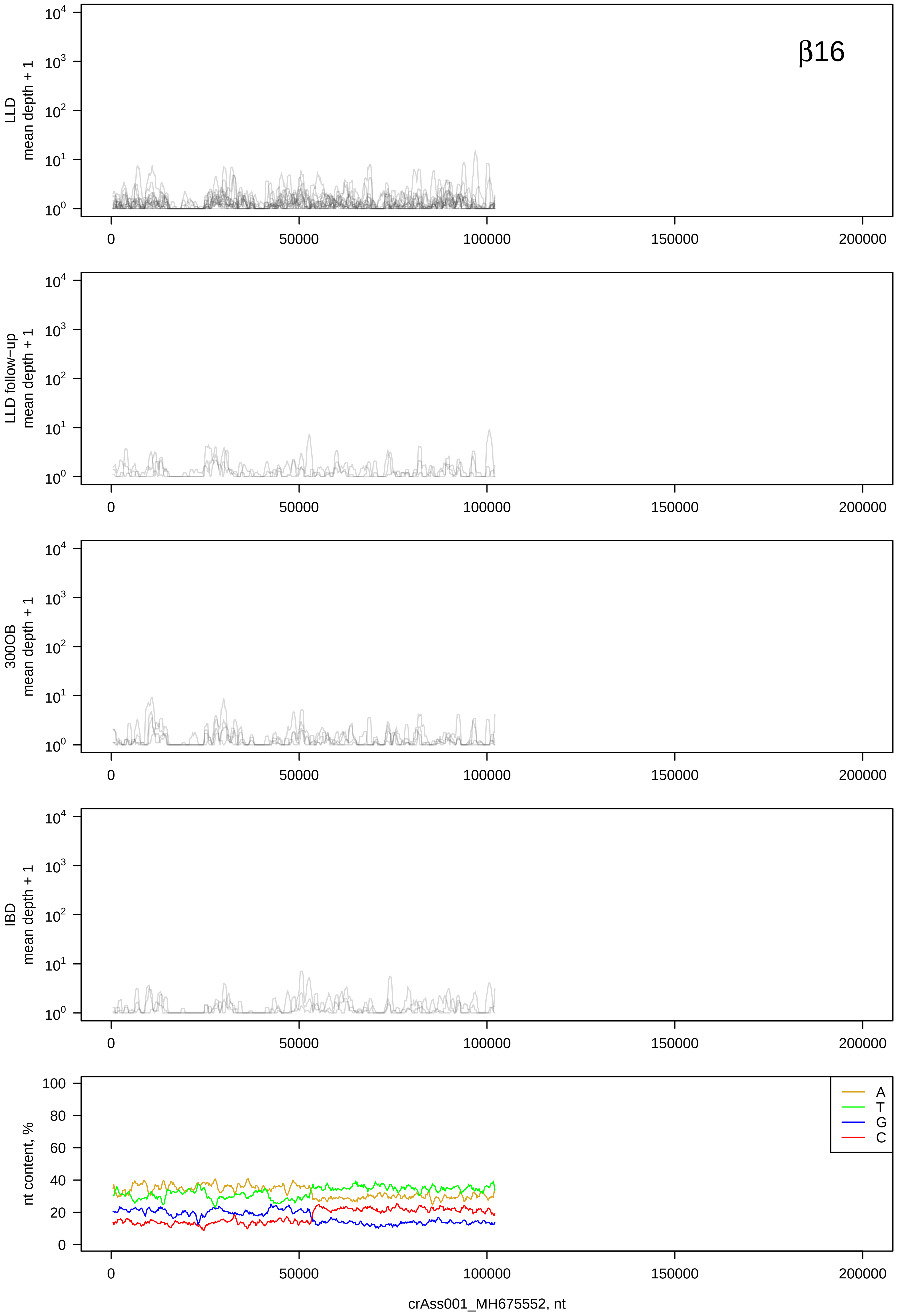

β16
LLD mean depth + 1
LLD follow-up mean depth + 1
300OB mean depth + 1
IBD mean depth + 1
nt content, %
A
T
G
C
0
50000
100000
150000
200000
crAss001_MH675552, nt

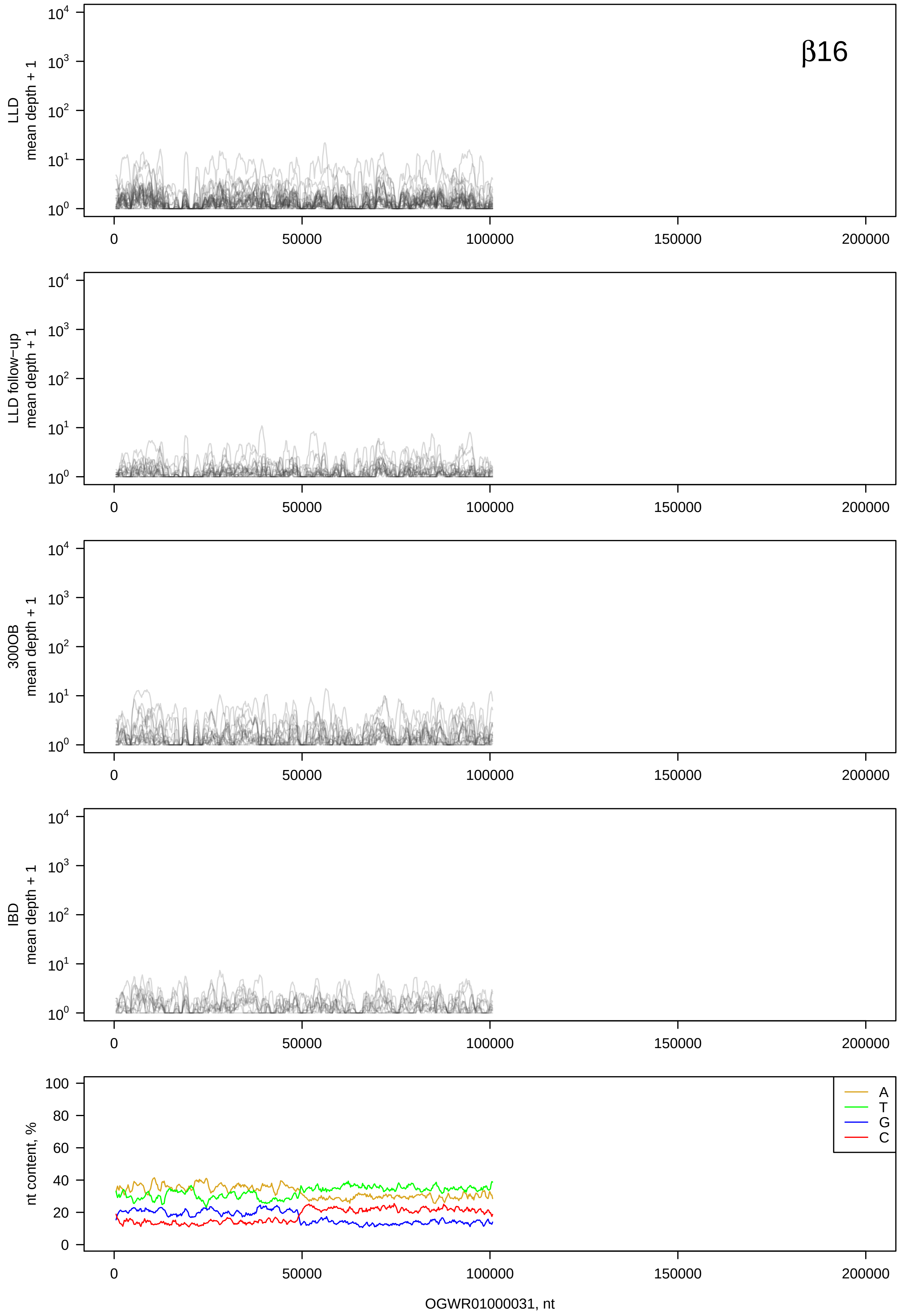
β16
LLD mean depth + 1
LLD follow-up mean depth + 1
300OB mean depth + 1
IBD mean depth + 1
nt content, %
A
T
G
C
OGWR01000031, nt

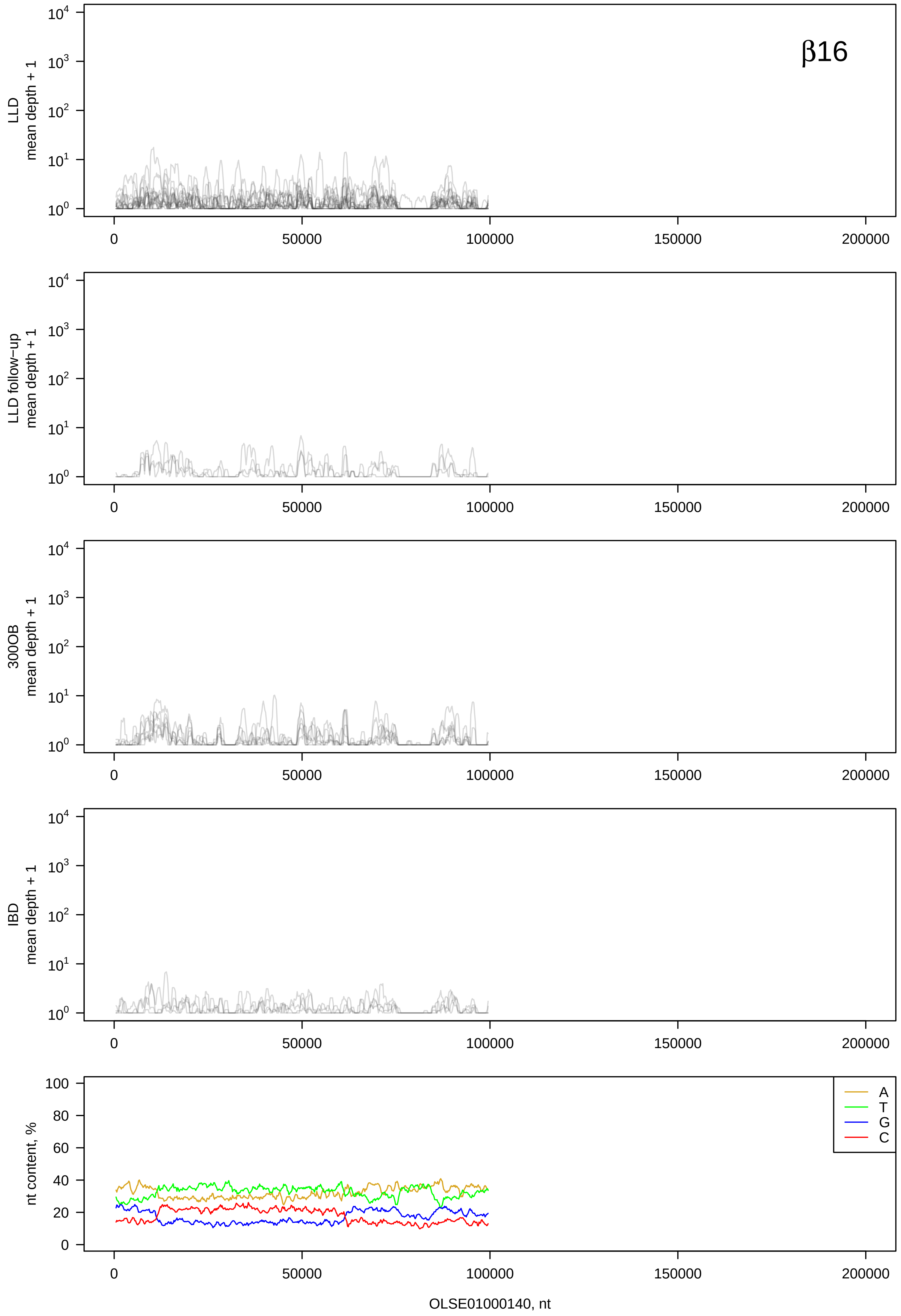

β16
LLD mean depth + 1
LLD follow−up mean depth + 1
300OB mean depth + 1
IBD mean depth + 1
nt content, %
A
T
G
C
0
50000
100000
150000
200000
OLSE01000140, nt

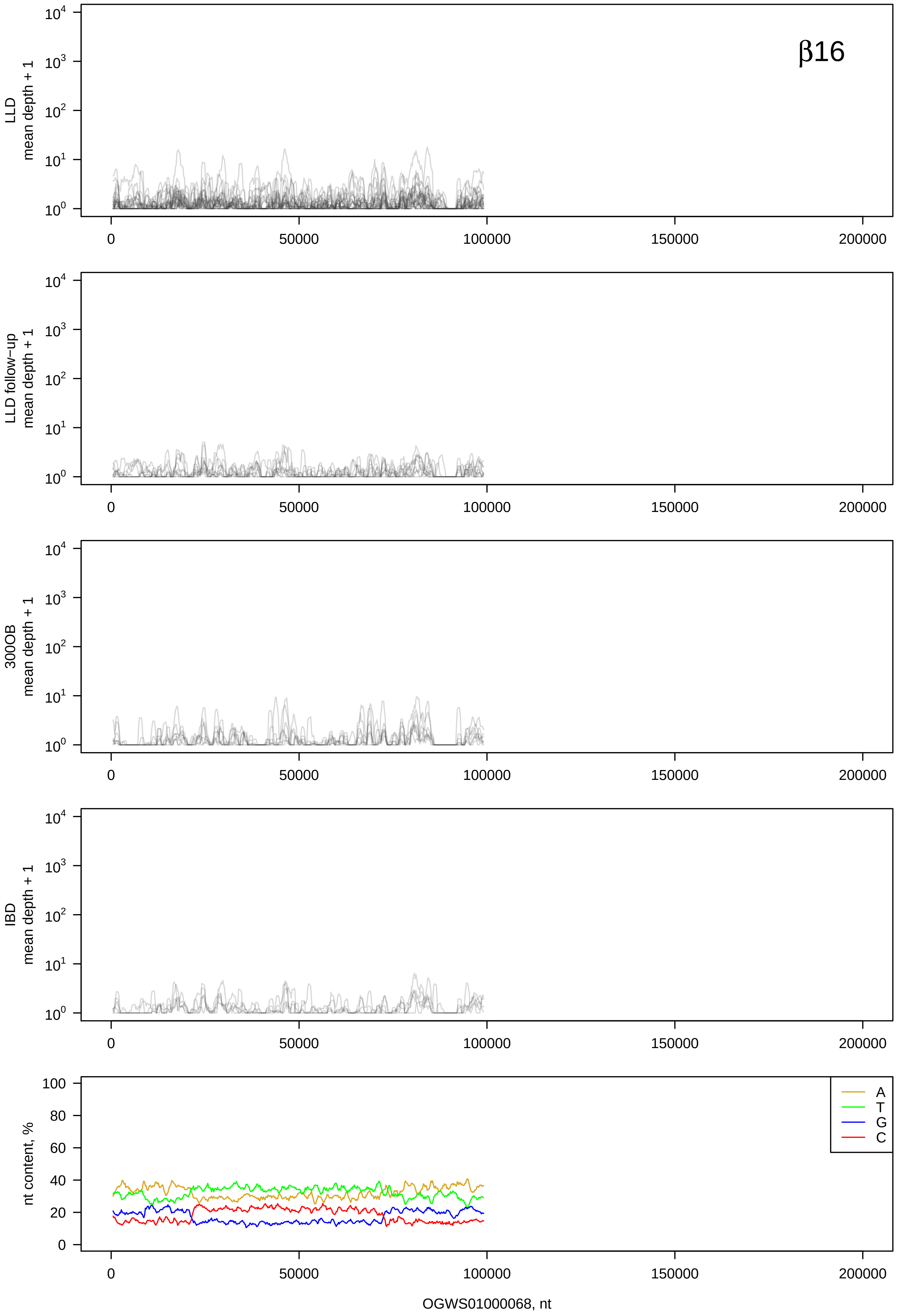
β16
LLD mean depth + 1
LLD follow−up mean depth + 1
300OB mean depth + 1
IBD mean depth + 1
nt content, %
A
T
G
C
0
50000
100000
150000
200000
OGWS01000068, nt

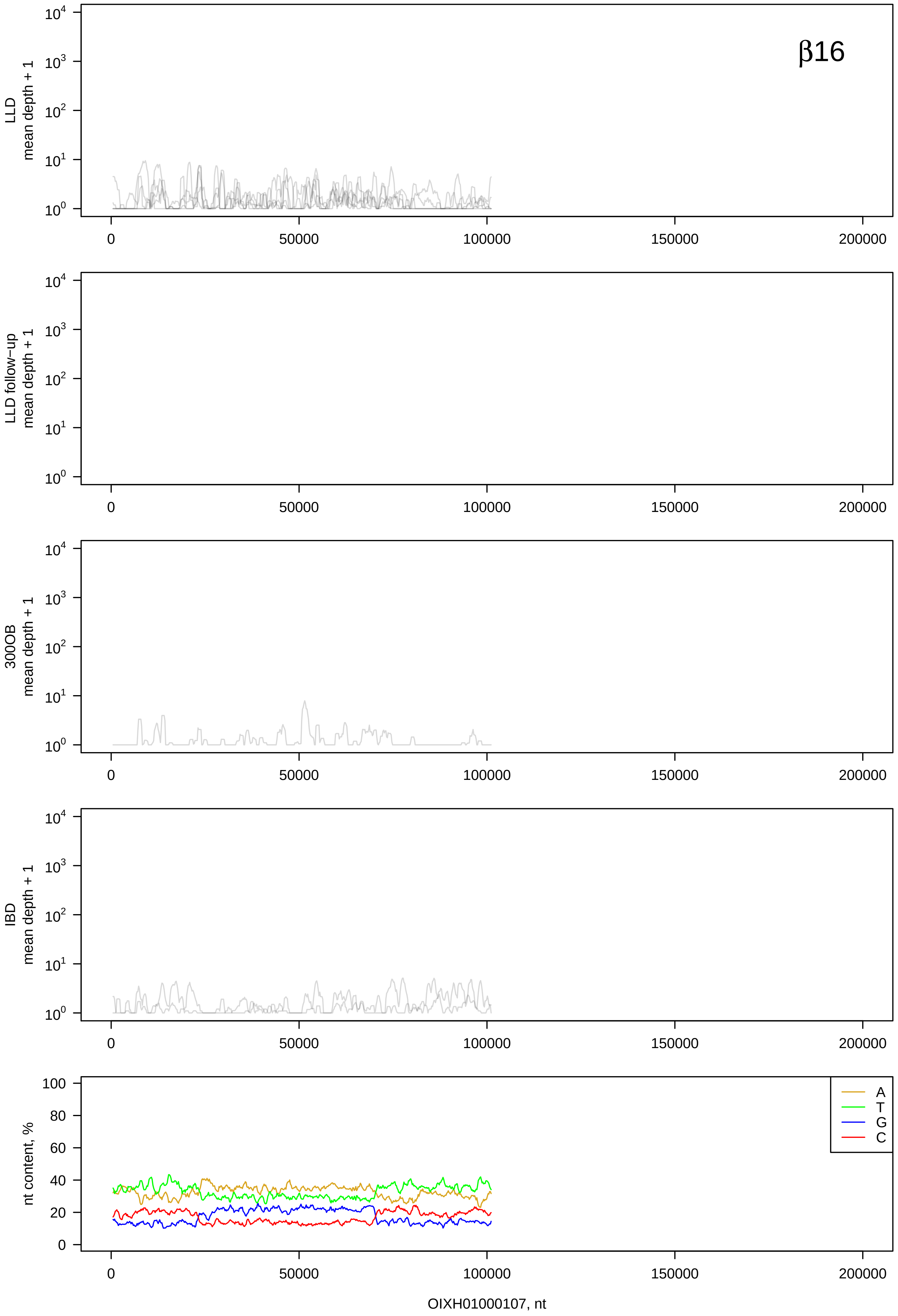

β16
LLD mean depth + 1
LLD follow−up mean depth + 1
300OB mean depth + 1
IBD mean depth + 1
nt content, %
A
T
G
C
0
50000
100000
150000
200000
OIXH01000107, nt

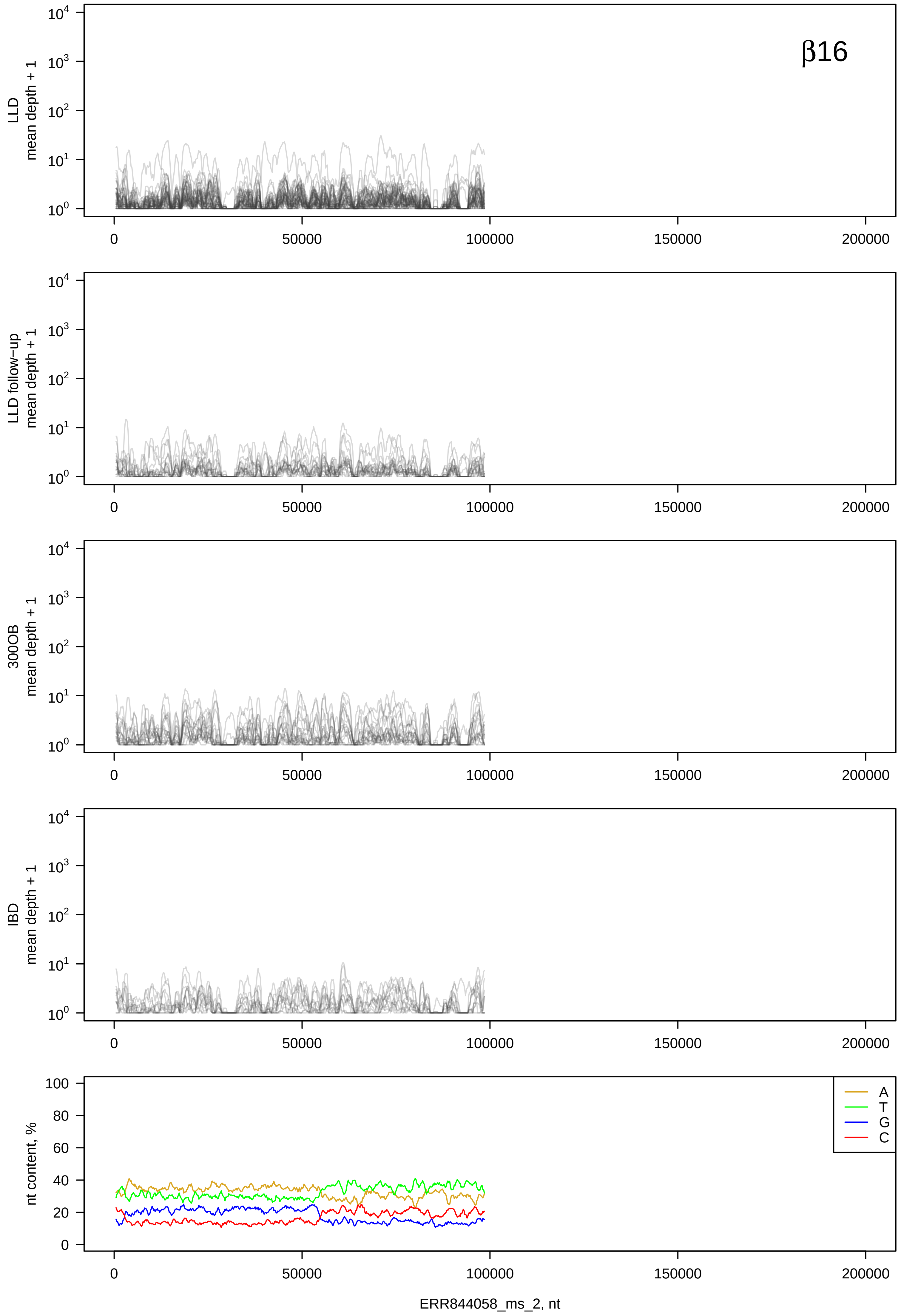

β16
LLD mean depth + 1
LLD follow−up mean depth + 1
300OB mean depth + 1
IBD mean depth + 1
nt content, %
A
T
G
C
0
50000
100000
150000
200000
ERR844058_ms_2, nt

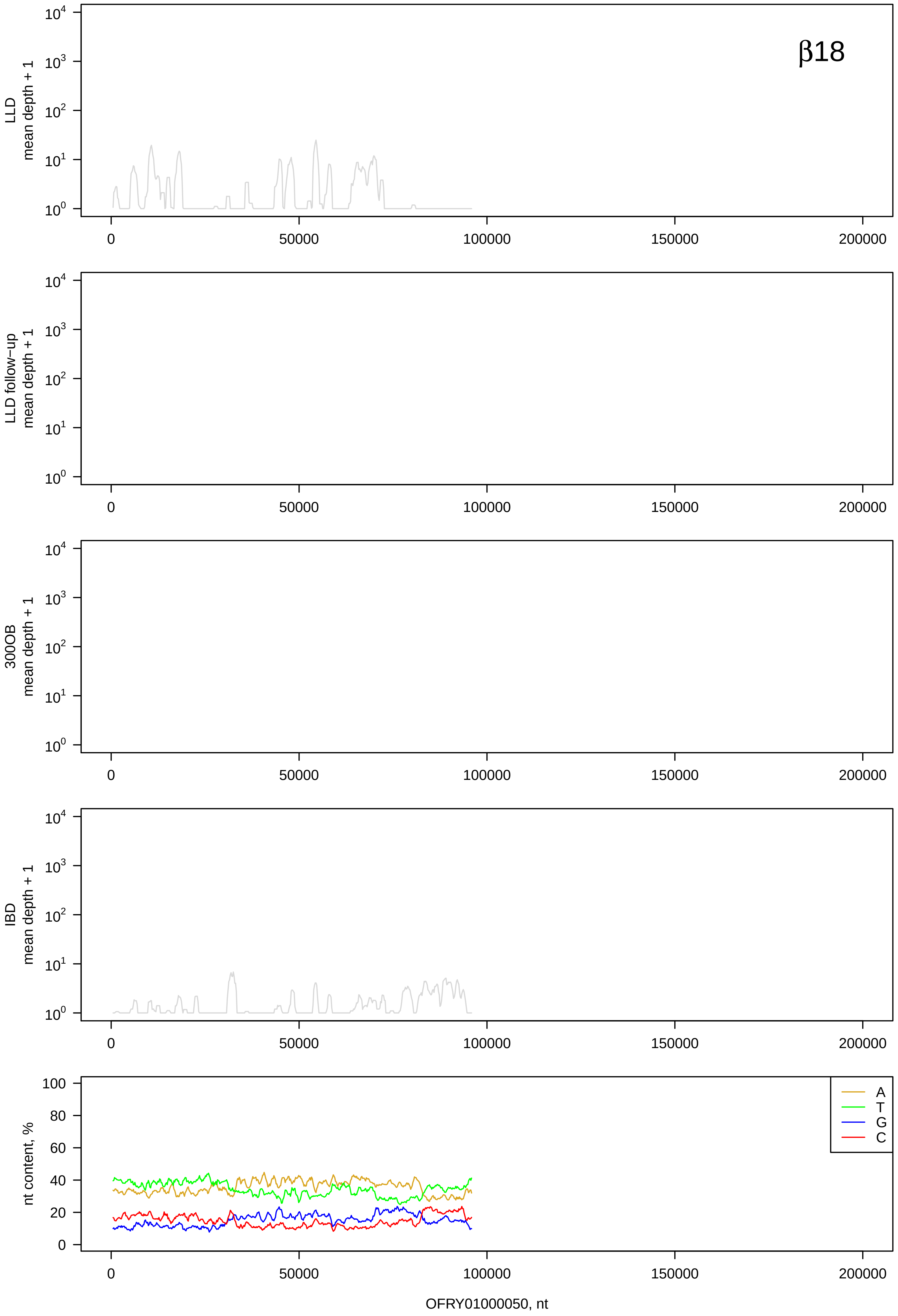
β18
LLD
mean depth + 1
LLD follow-up
mean depth + 1
300OB
mean depth + 1
IBD
mean depth + 1
nt content, %
A
T
G
C
0
50000
100000
150000
200000
OFRY01000050, nt

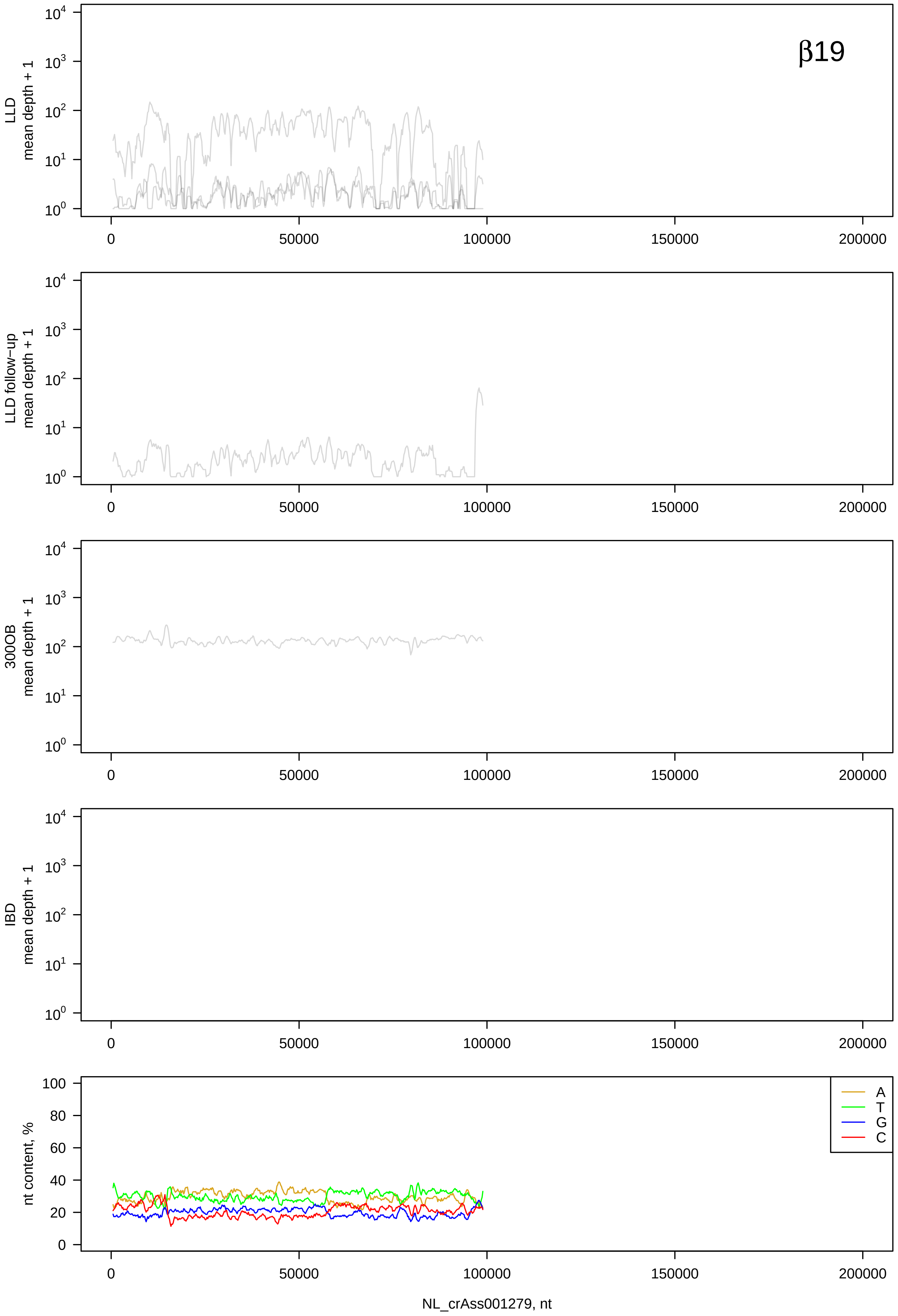

β19
LLD mean depth + 1
LLD follow−up mean depth + 1
300OB mean depth + 1
IBD mean depth + 1
nt content, %
A
T
G
C
0
50000
100000
150000
200000
NL_crAss001279, nt

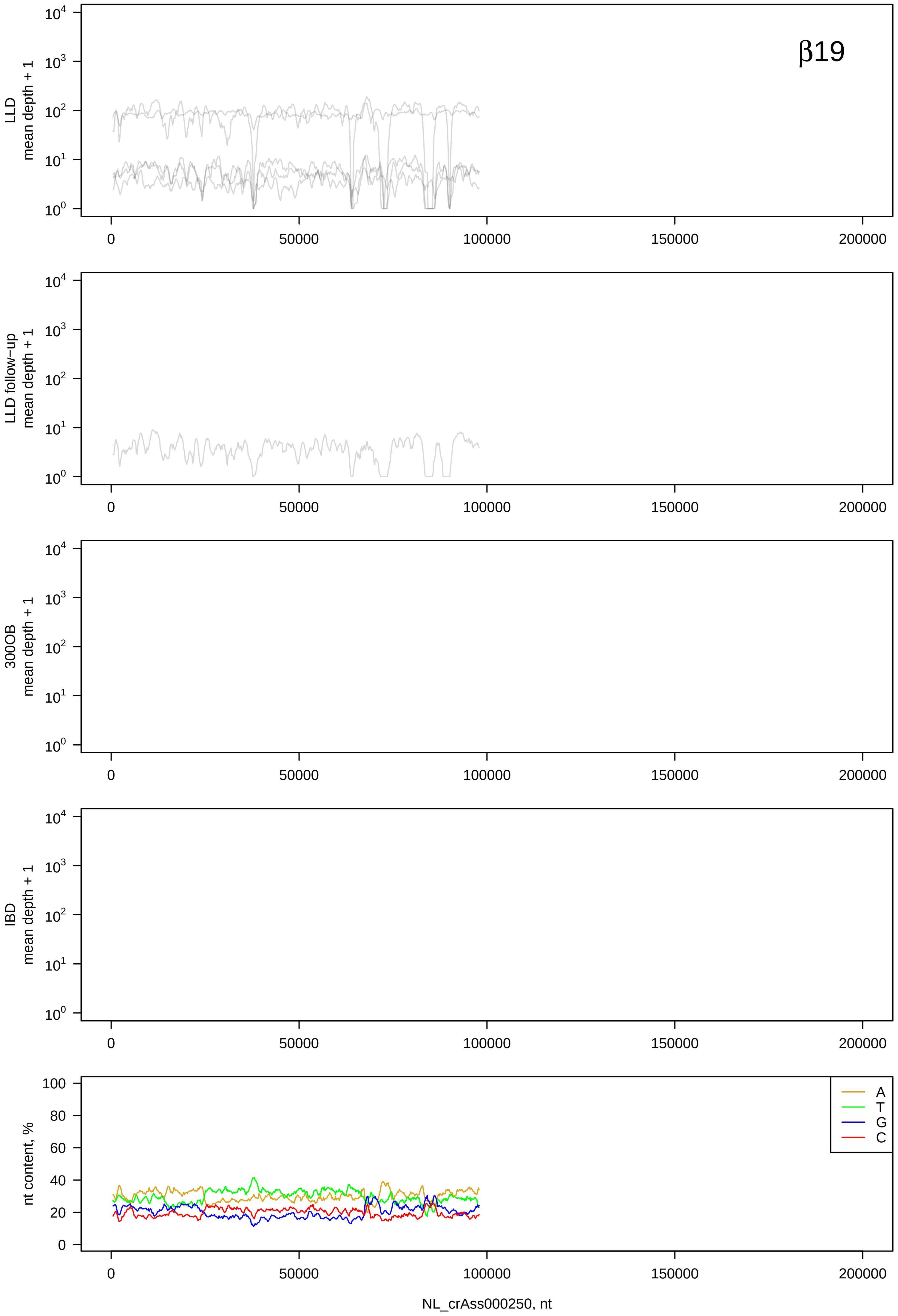

β19
LLD mean depth + 1
LLD follow−up mean depth + 1
300OB mean depth + 1
IBD mean depth + 1
nt content, %
A
T
G
C
NL_crAss000250, nt

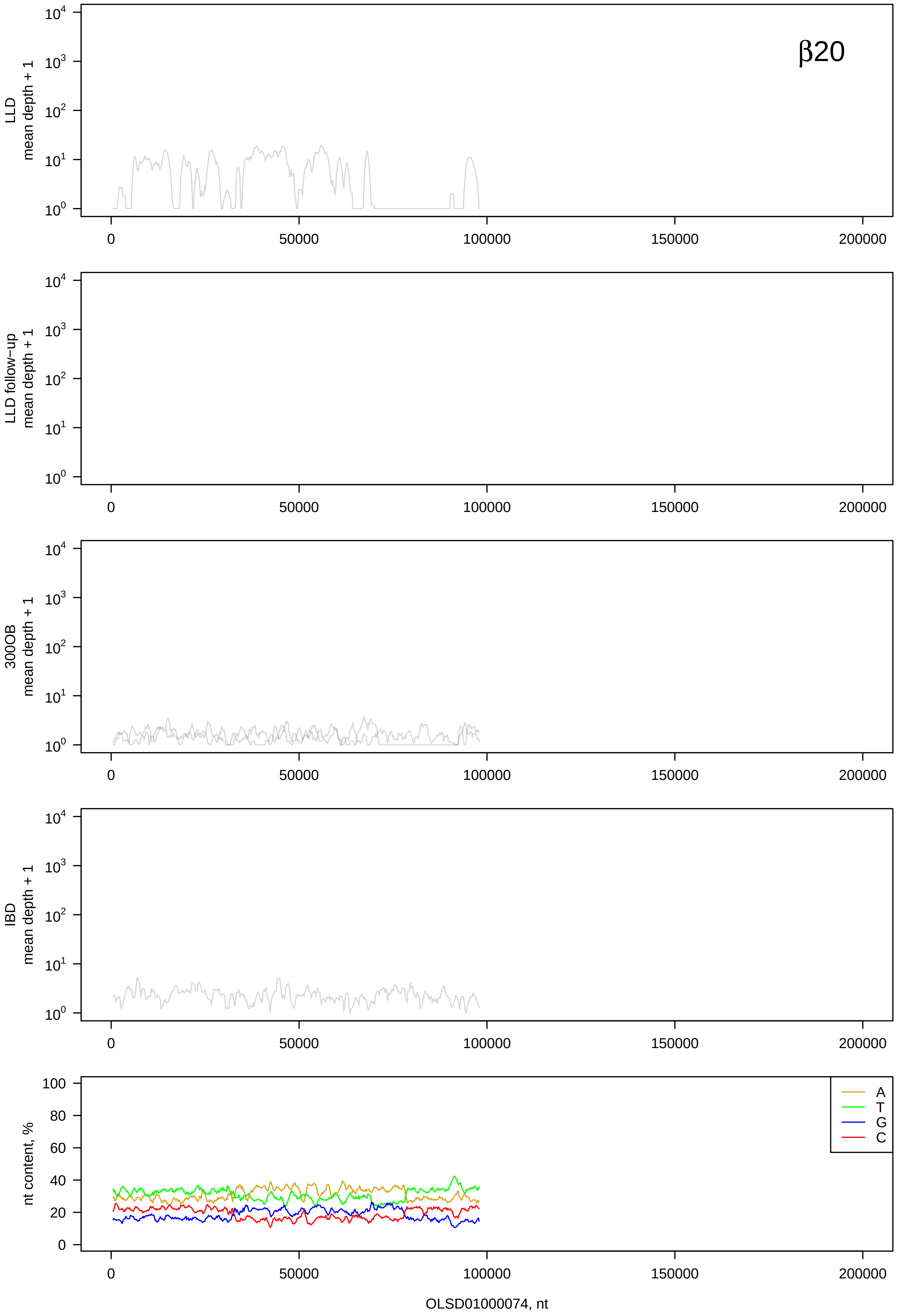
β20
LLD
mean depth + 1
LLD follow−up
mean depth + 1
300OB
mean depth + 1
IBD
mean depth + 1
nt content, %
A
T
G
C
0
50000
100000
150000
200000
OLSD01000074, nt

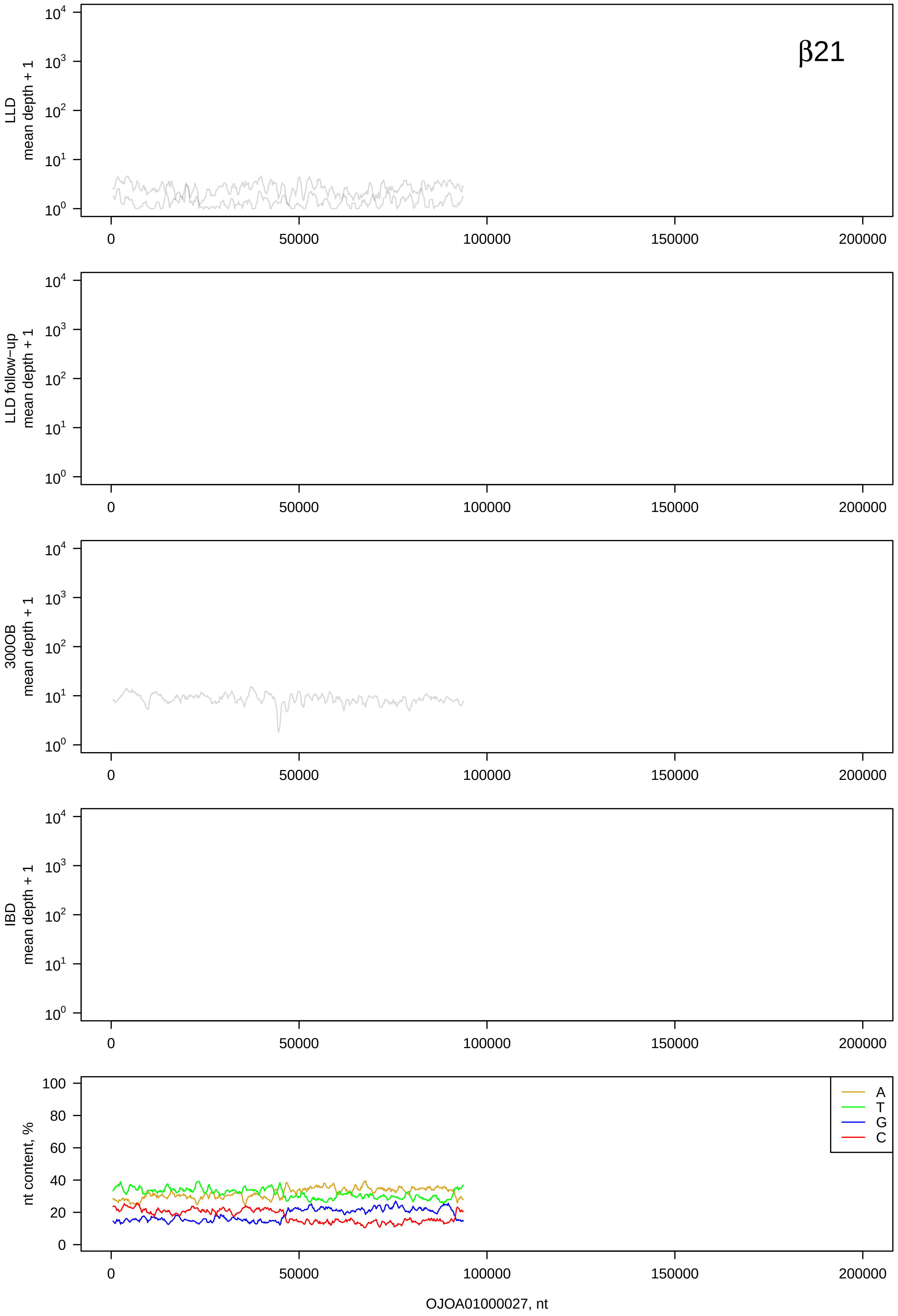
β21
LLD
mean depth + 1
LLD follow−up
mean depth + 1
300OB
mean depth + 1
IBD
mean depth + 1
nt content, %
A
T
G
C
0
50000
100000
150000
200000
OJOA01000027, nt

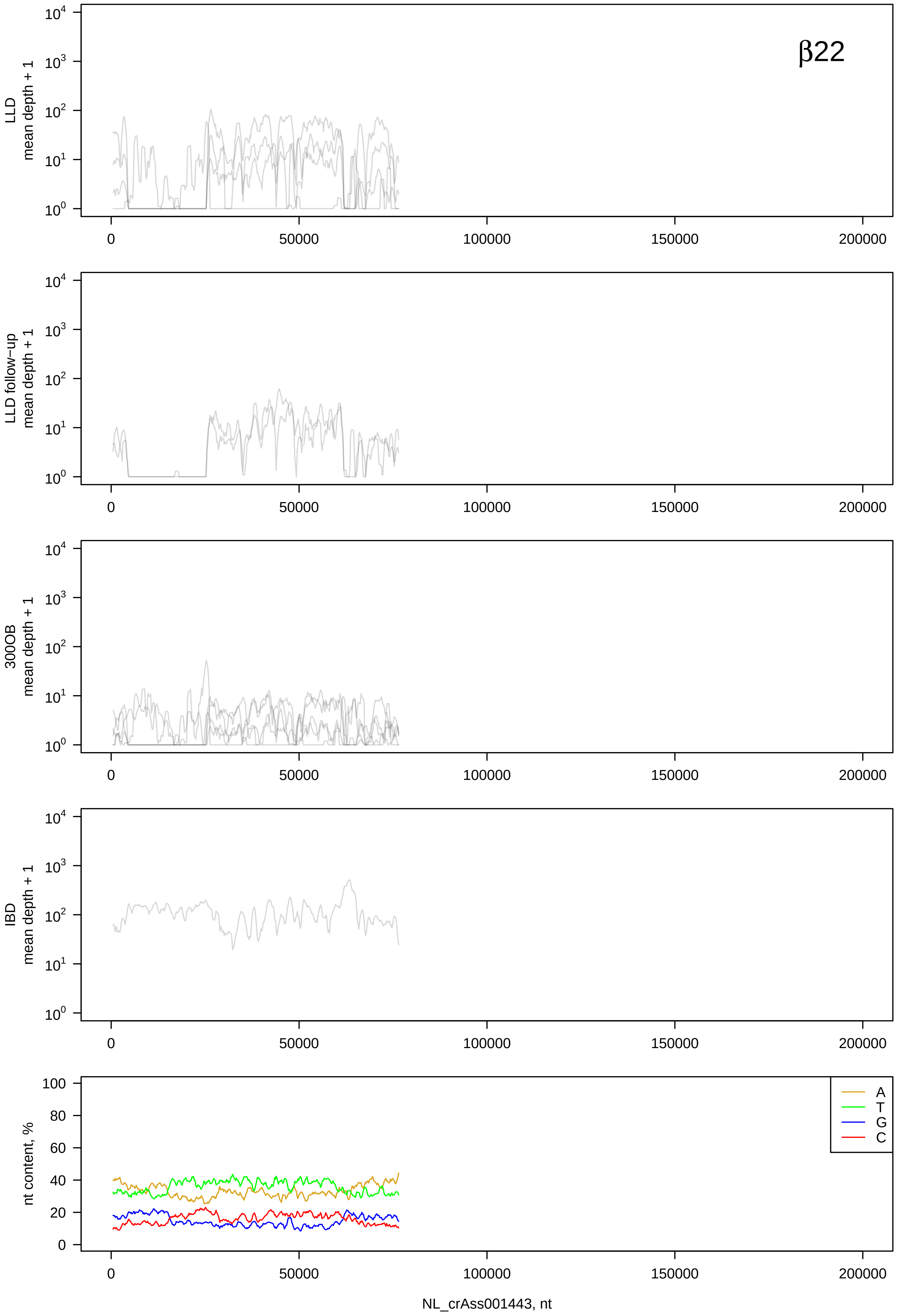

β22
LLD mean depth + 1
LLD follow-up mean depth + 1
300OB mean depth + 1
IBD mean depth + 1
nt content, %
A
T
G
C
0
50000
100000
150000
200000
NL_crAss001443, nt

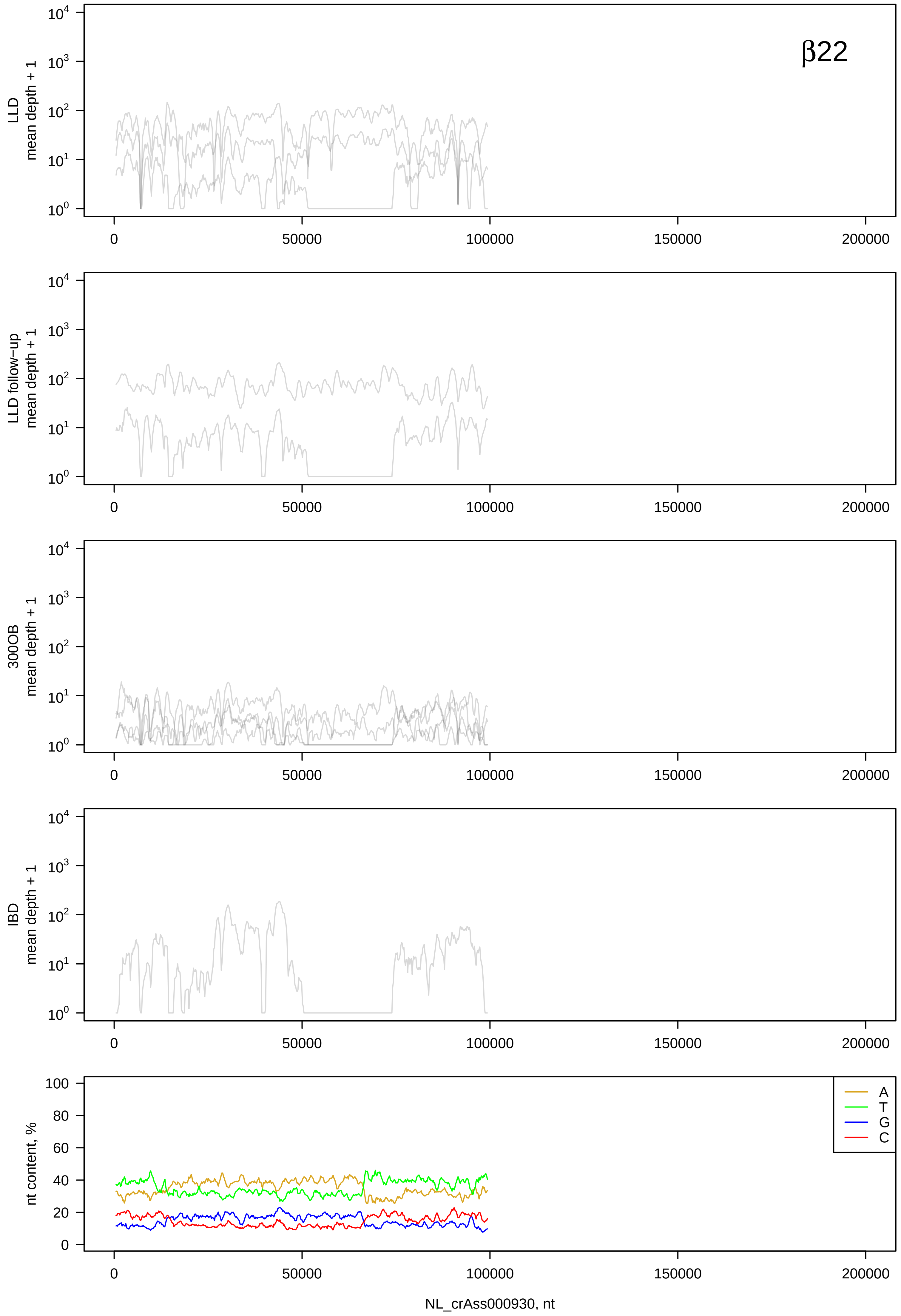
β22
LLD
mean depth + 1
LLD follow−up
mean depth + 1
300OB
mean depth + 1
IBD
mean depth + 1
nt content, %
A
T
G
C
NL_crAss000930, nt

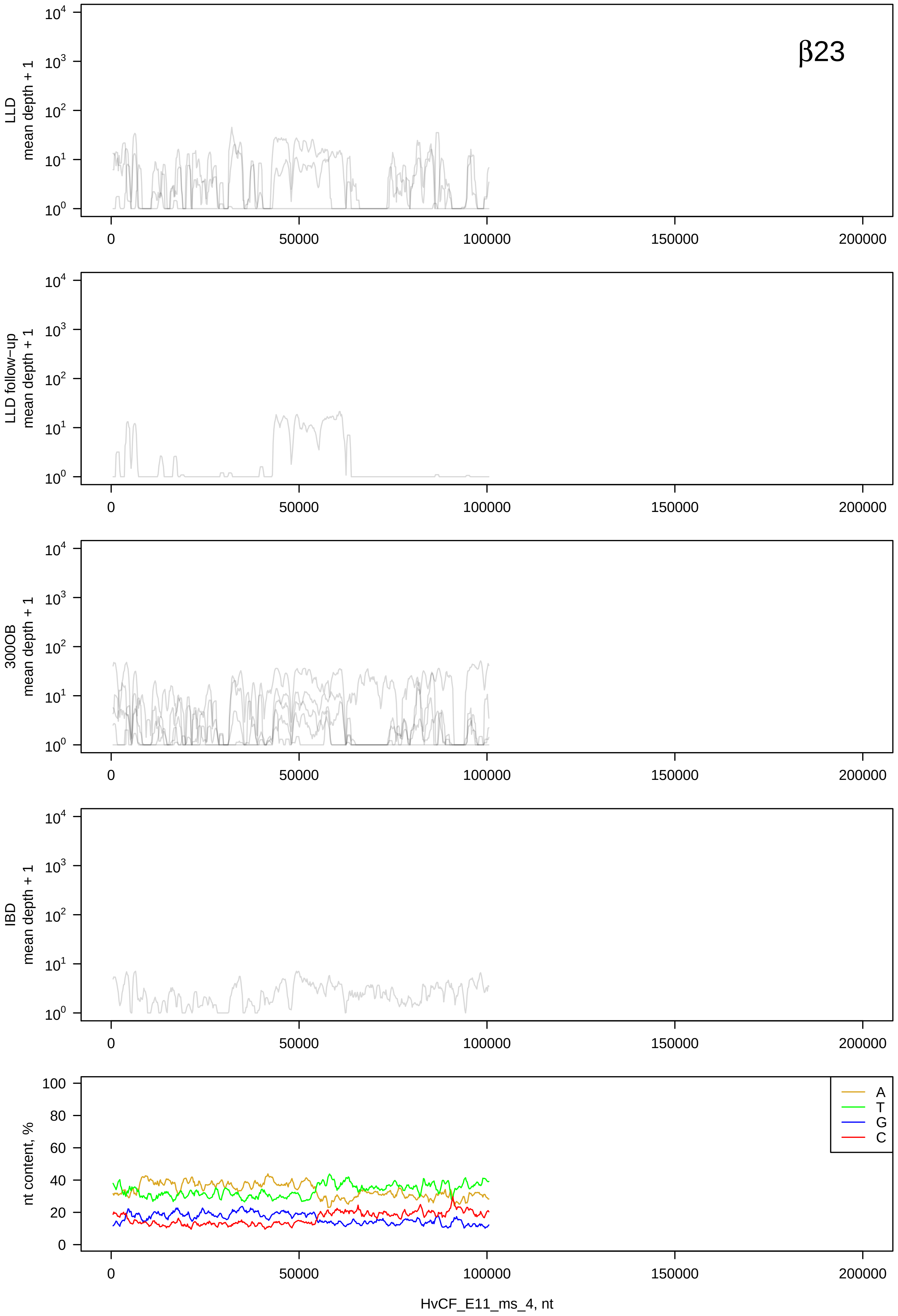
β23
LLD mean depth + 1
LLD follow-up mean depth + 1
300OB mean depth + 1
IBD mean depth + 1
nt content, %
A
T
G
C
HvCF_E11_ms_4, nt
0
50000
100000
150000
200000

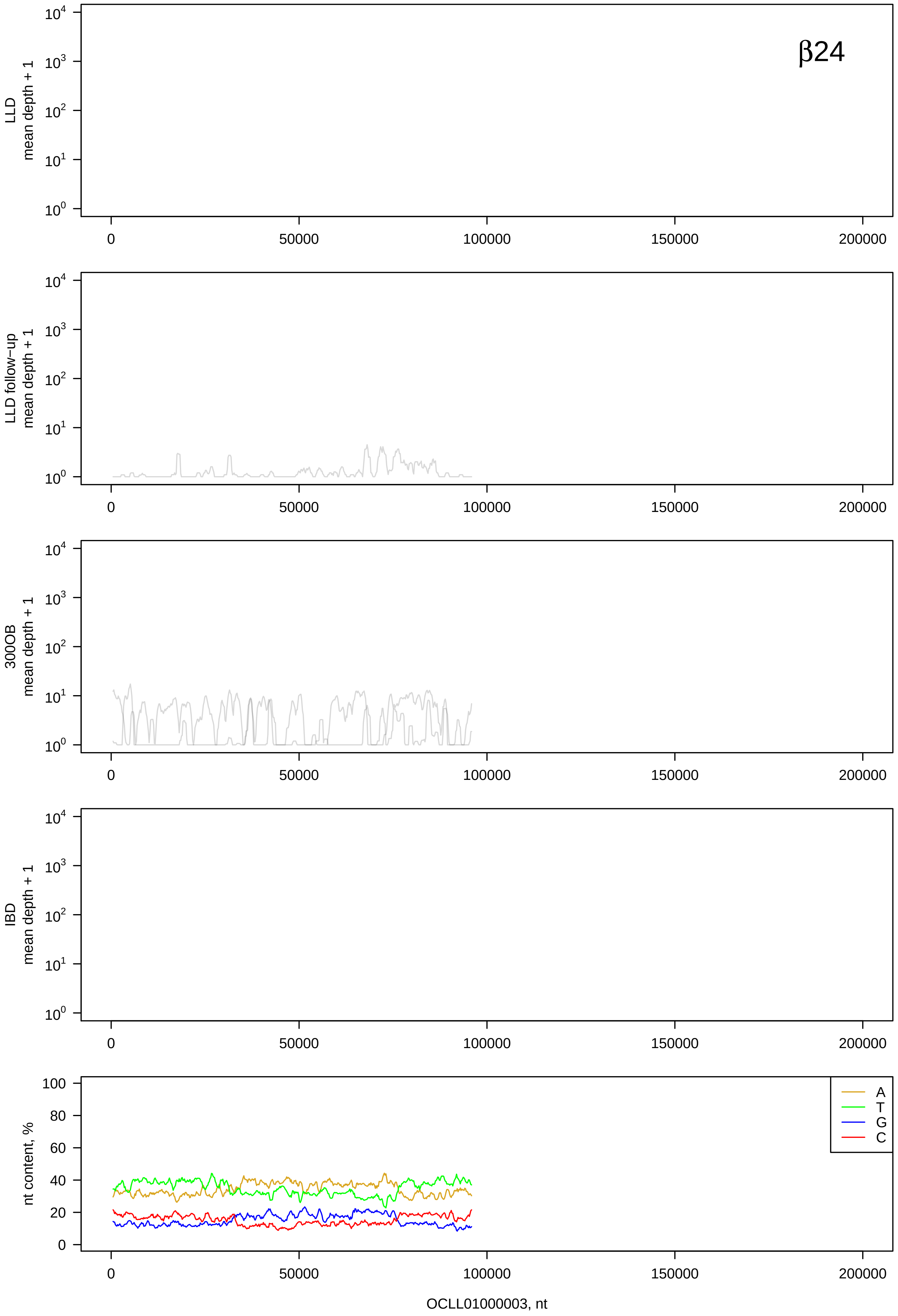

β24
LLD mean depth + 1
LLD follow−up mean depth + 1
300OB mean depth + 1
IBD mean depth + 1
nt content, %
A
T
G
C
0
50000
100000
150000
200000
OCLL01000003, nt

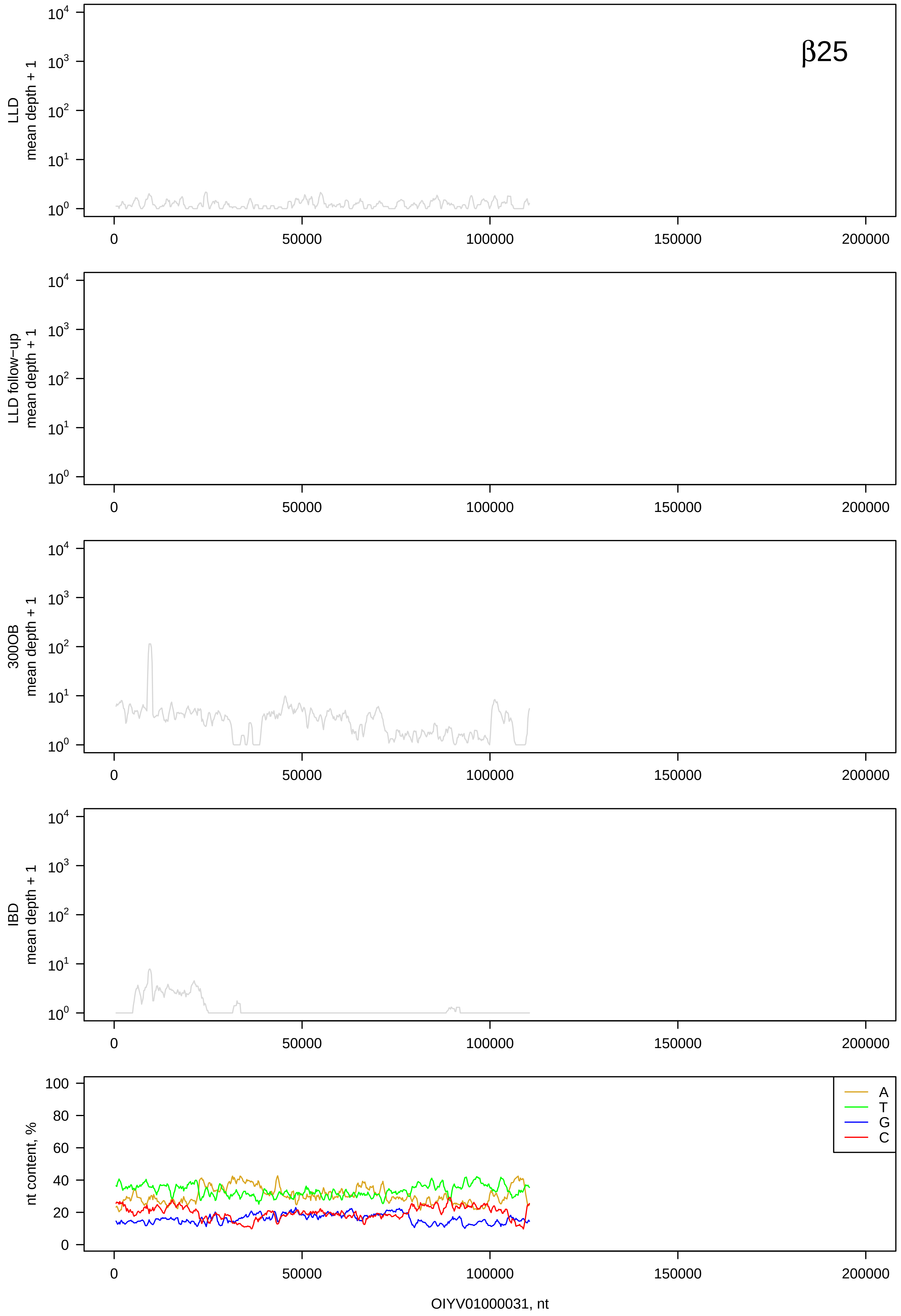
β25
LLD
mean depth + 1
LLD follow−up
mean depth + 1
300OB
mean depth + 1
IBD
mean depth + 1
nt content, %
A
T
G
C
0
50000
100000
150000
200000
OIYV01000031, nt

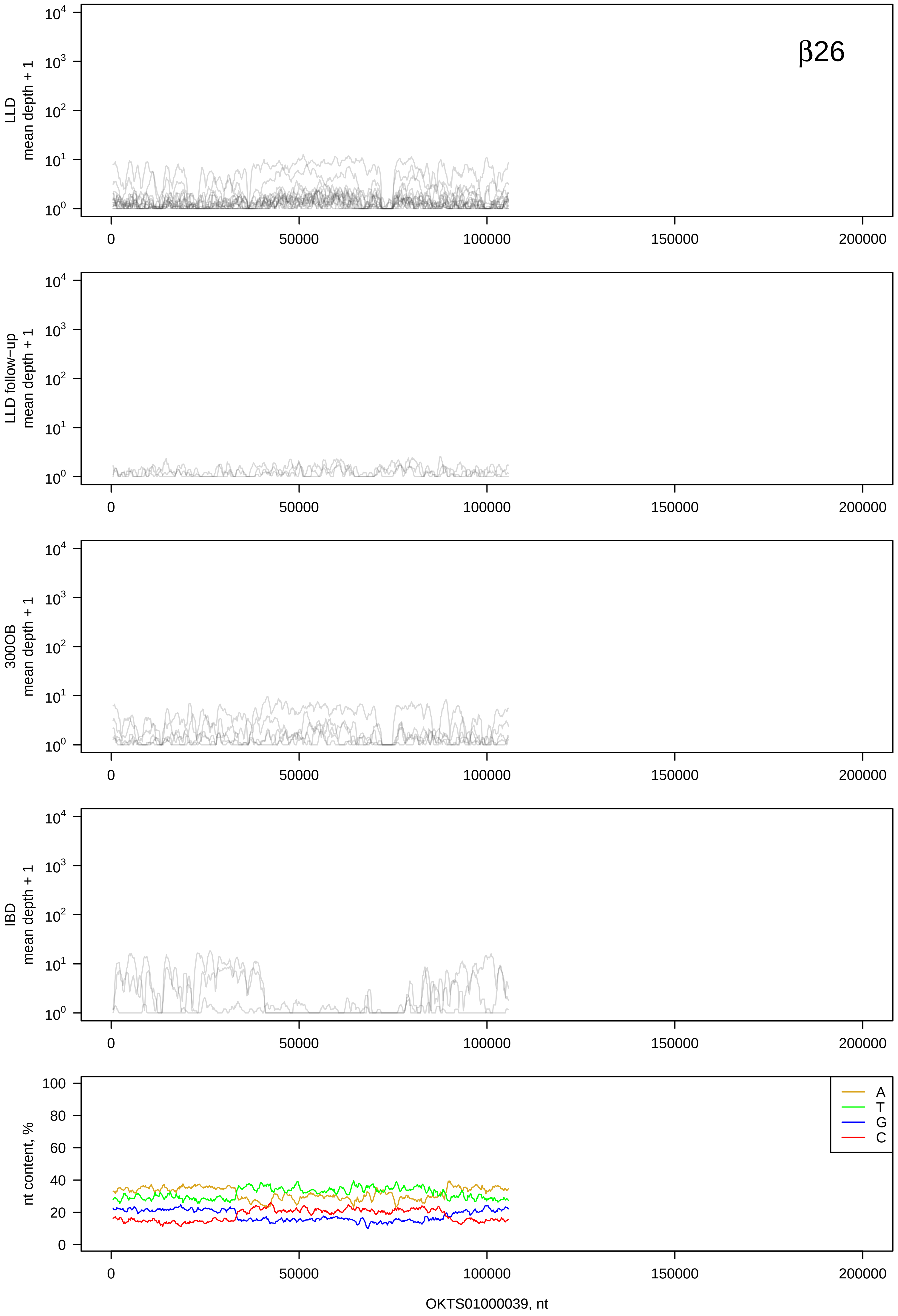

β26
LLD mean depth + 1
LLD follow−up mean depth + 1
300OB mean depth + 1
IBD mean depth + 1
nt content, %
A
T
G
C
OKTS01000039, nt

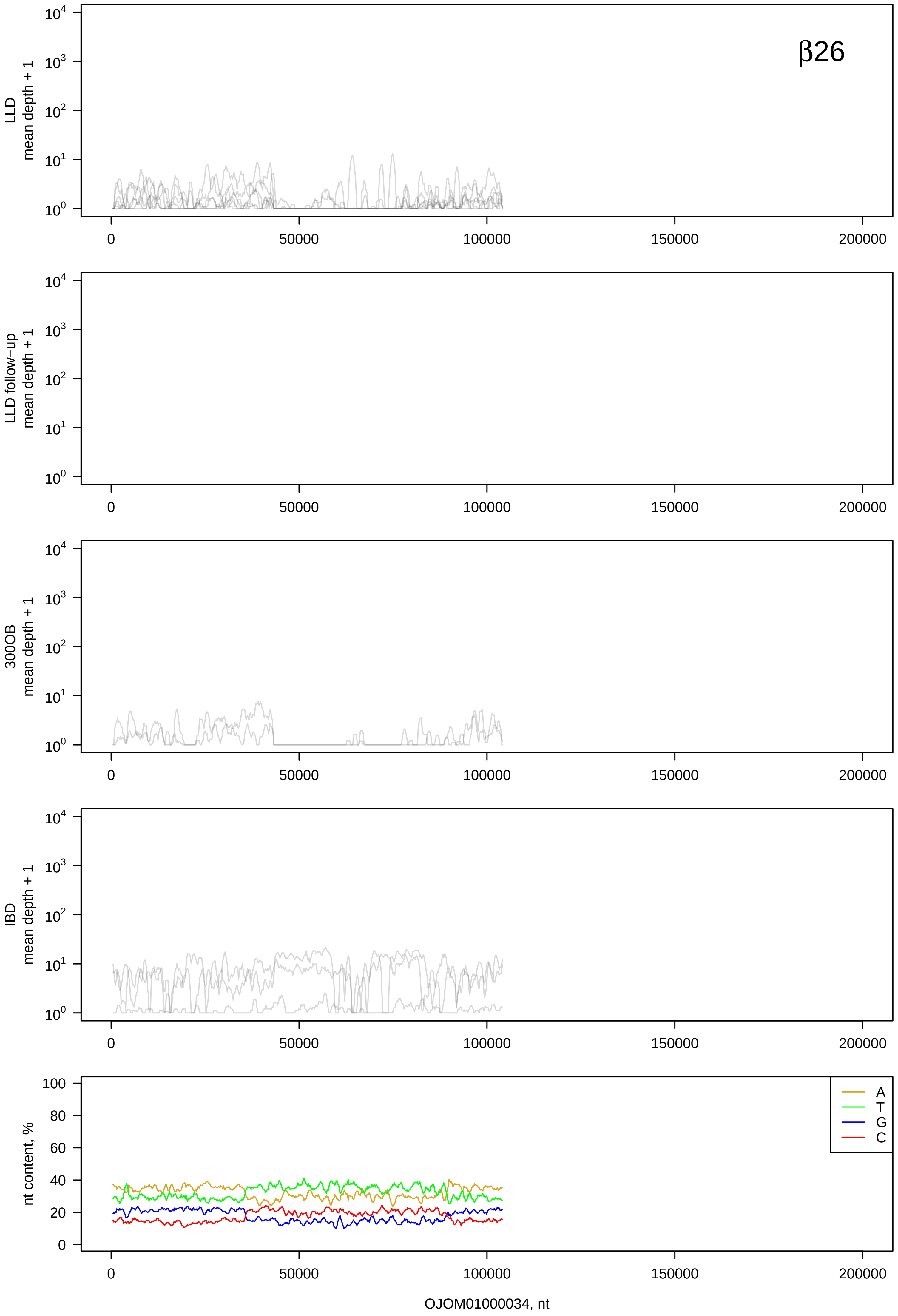
β26
LLD
mean depth + 1
LLD follow−up
mean depth + 1
300OB
mean depth + 1
IBD
mean depth + 1
nt content, %
A
T
G
C
OJOM01000034, nt

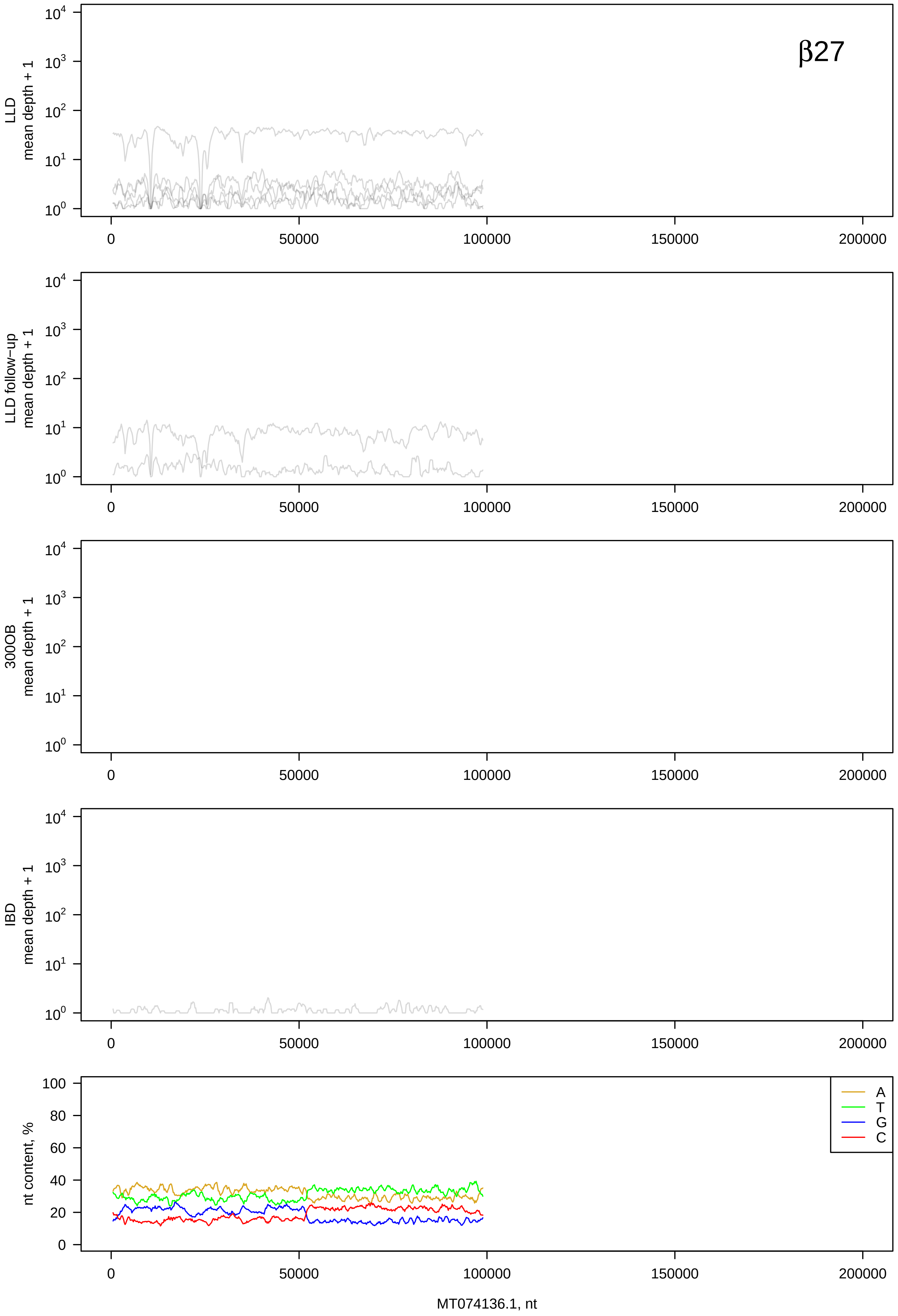

β27
LLD mean depth + 1
LLD follow-up mean depth + 1
300OB mean depth + 1
IBD mean depth + 1
nt content, %
A
T
G
C
MT074136.1, nt

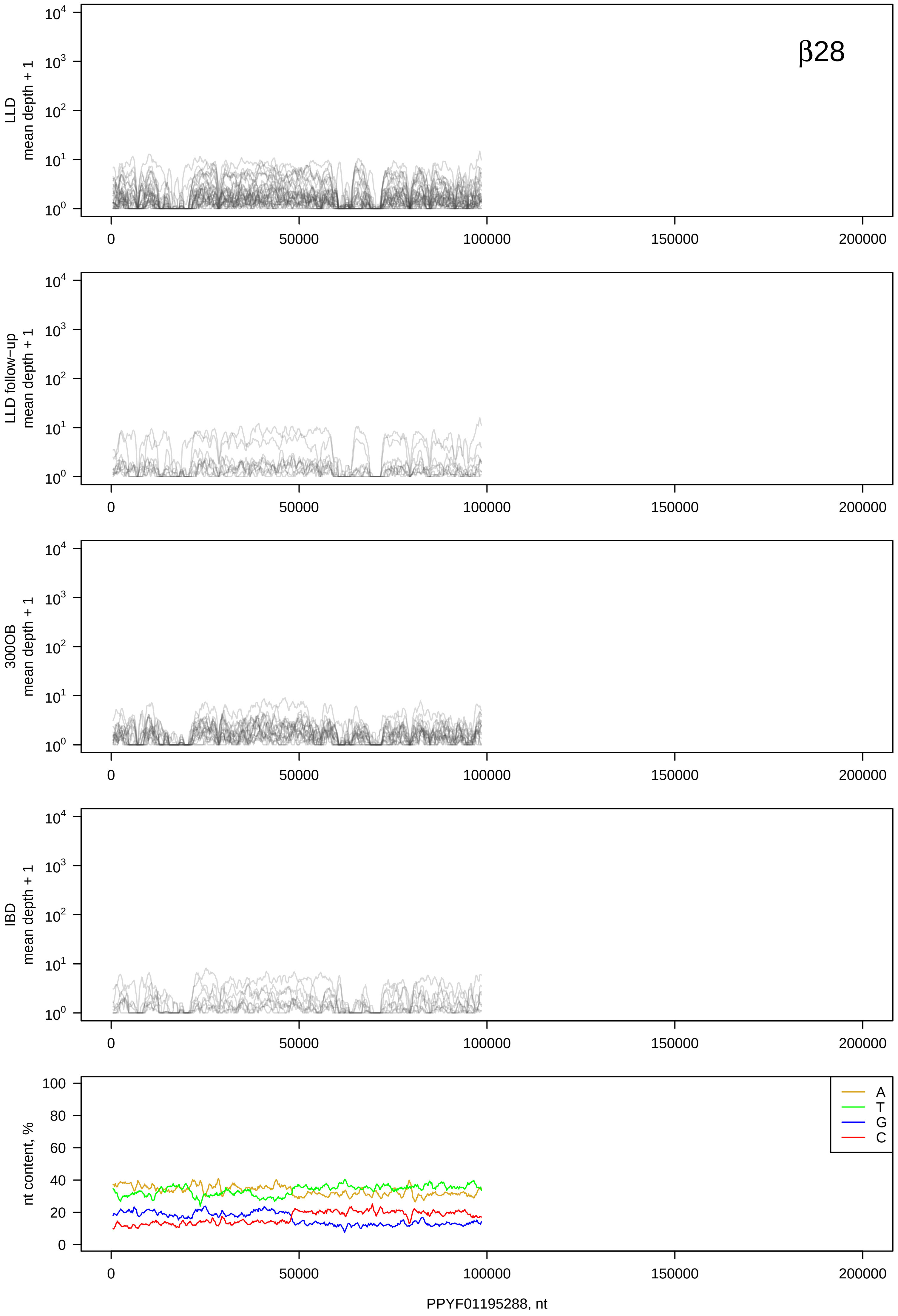
β28
LLD
mean depth + 1
LLD follow−up
mean depth + 1
300OB
mean depth + 1
IBD
mean depth + 1
nt content, %
A
T
G
C
0
50000
100000
150000
200000
PPYF01195288, nt

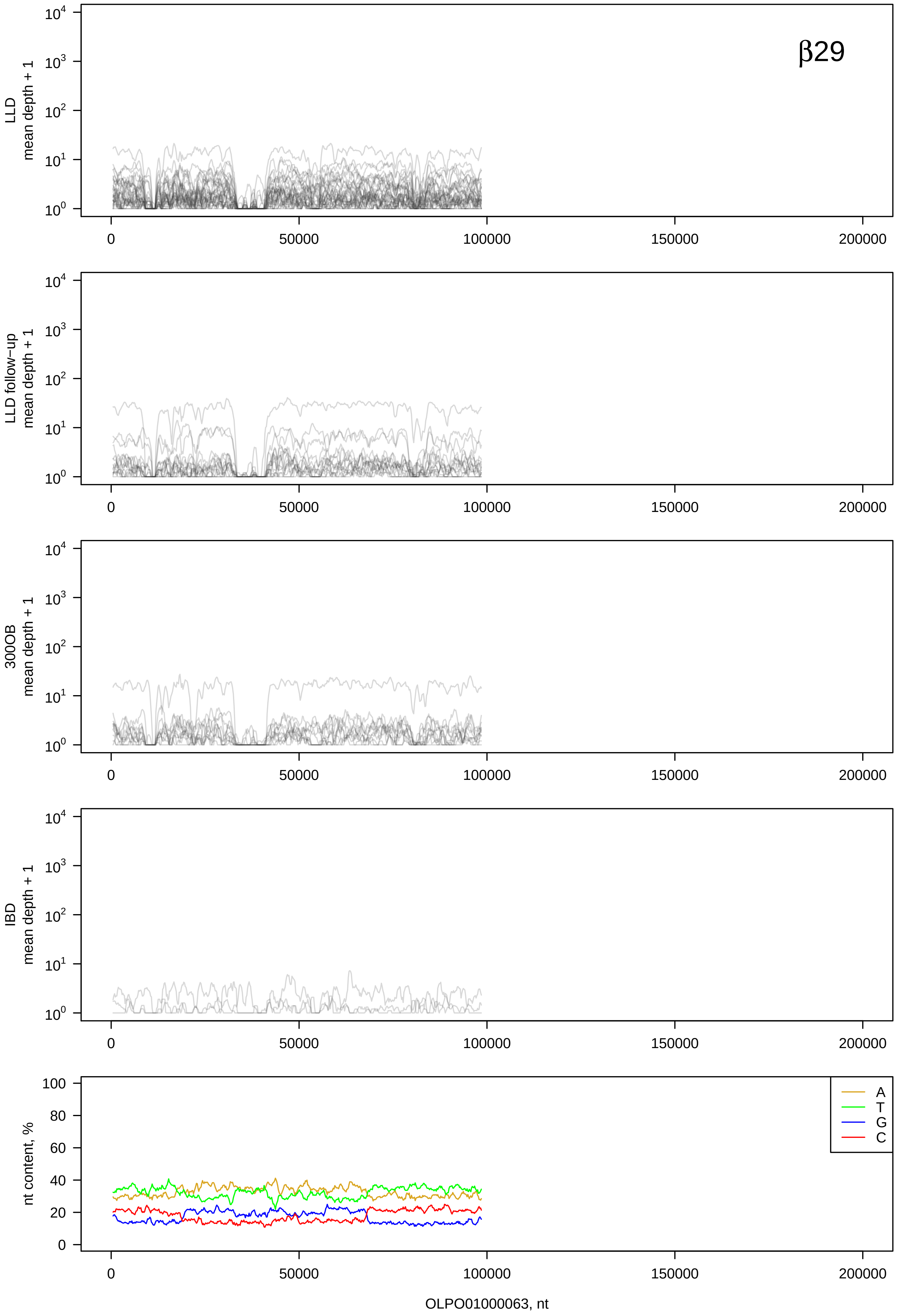

β29
LLD mean depth + 1
LLD follow−up mean depth + 1
300OB mean depth + 1
IBD mean depth + 1
nt content, %
A
T
G
C
0
50000
100000
150000
200000
OLPO01000063, nt

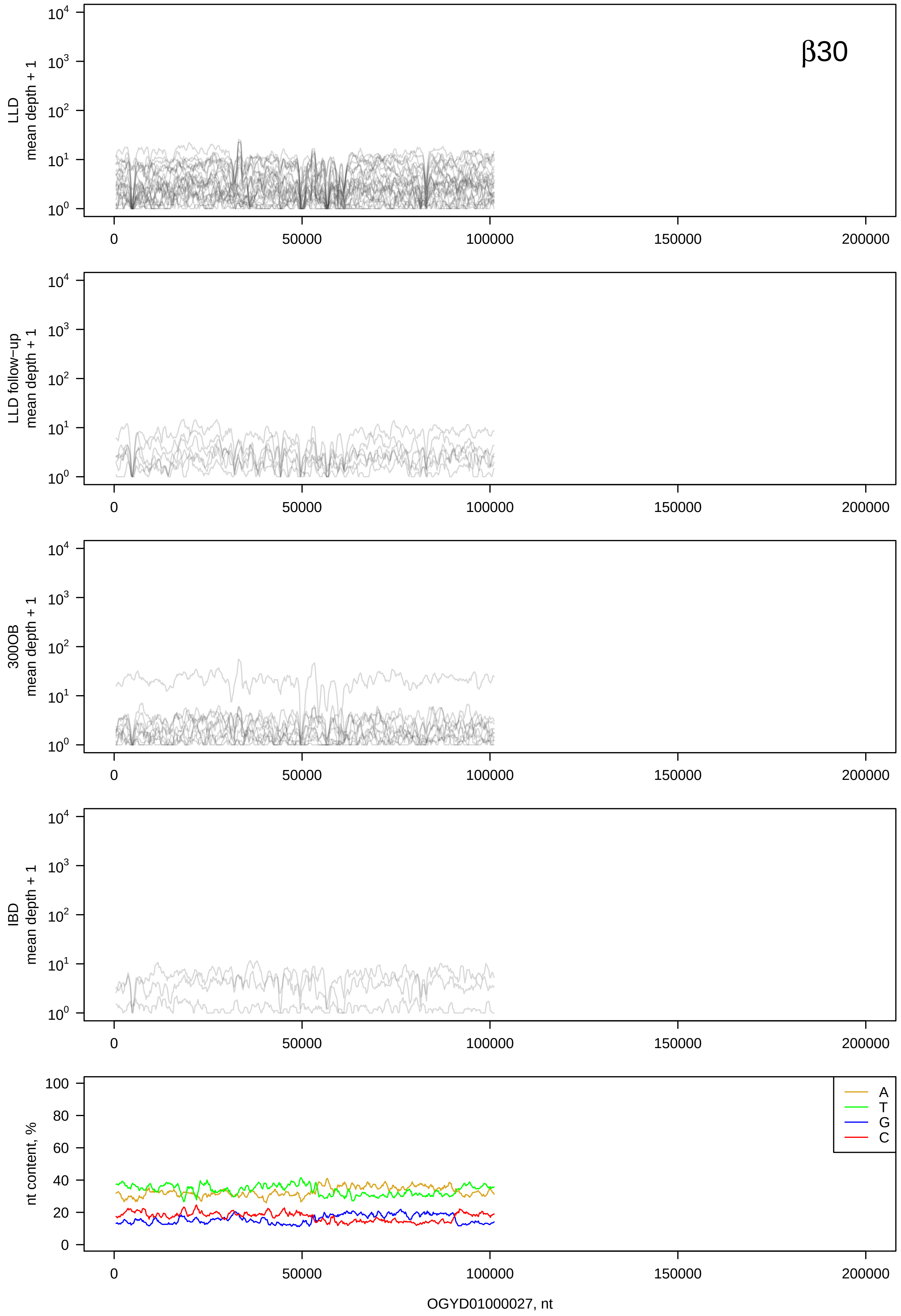
β30
LLD mean depth + 1
LLD follow−up mean depth + 1
300OB mean depth + 1
IBD mean depth + 1
nt content, %
A
T
G
C
0
50000
100000
150000
200000
OGYD01000027, nt

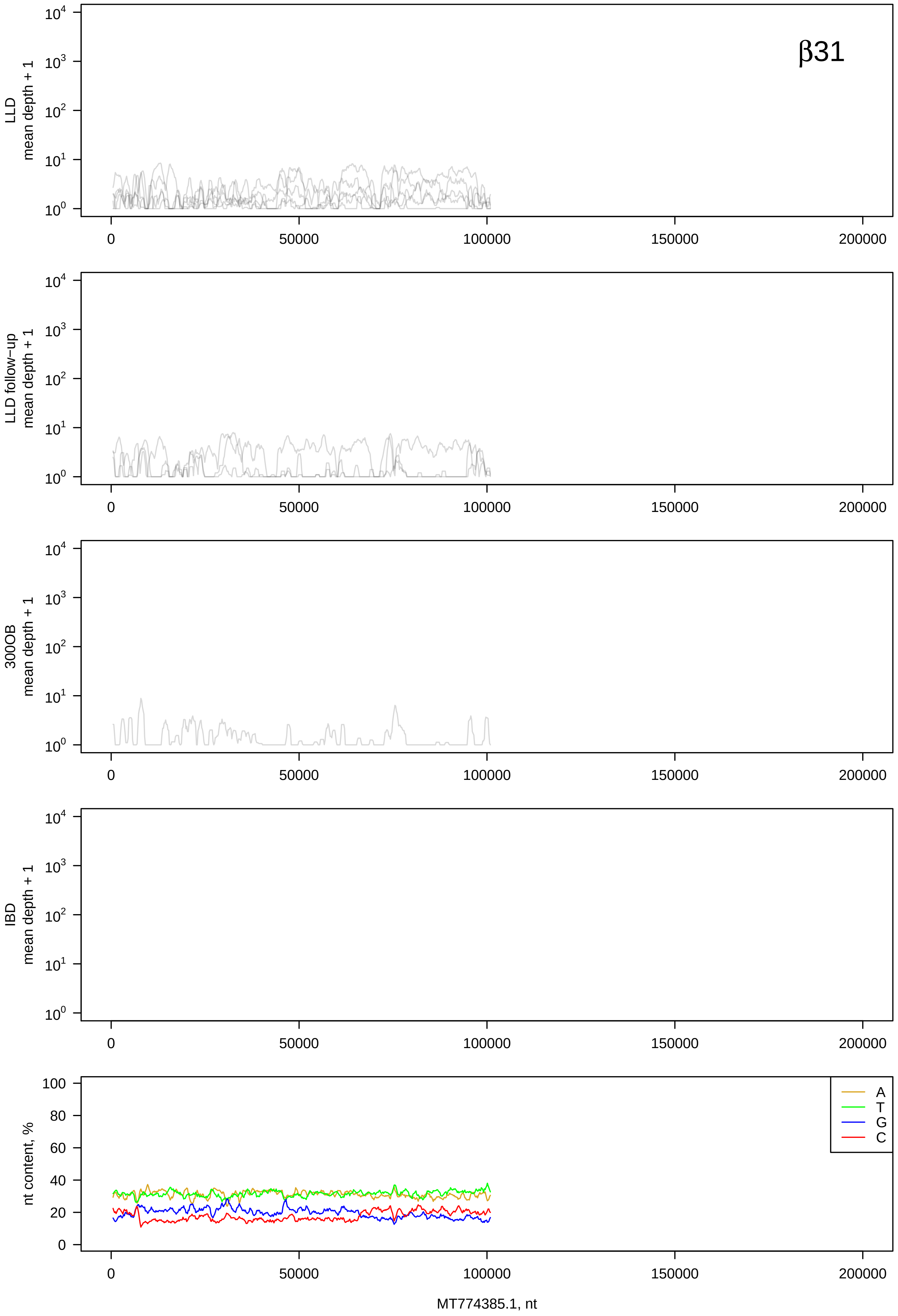
β31
LLD
mean depth + 1
LLD follow−up
mean depth + 1
300OB
mean depth + 1
IBD
mean depth + 1
nt content, %
A
T
G
C
MT774385.1, nt

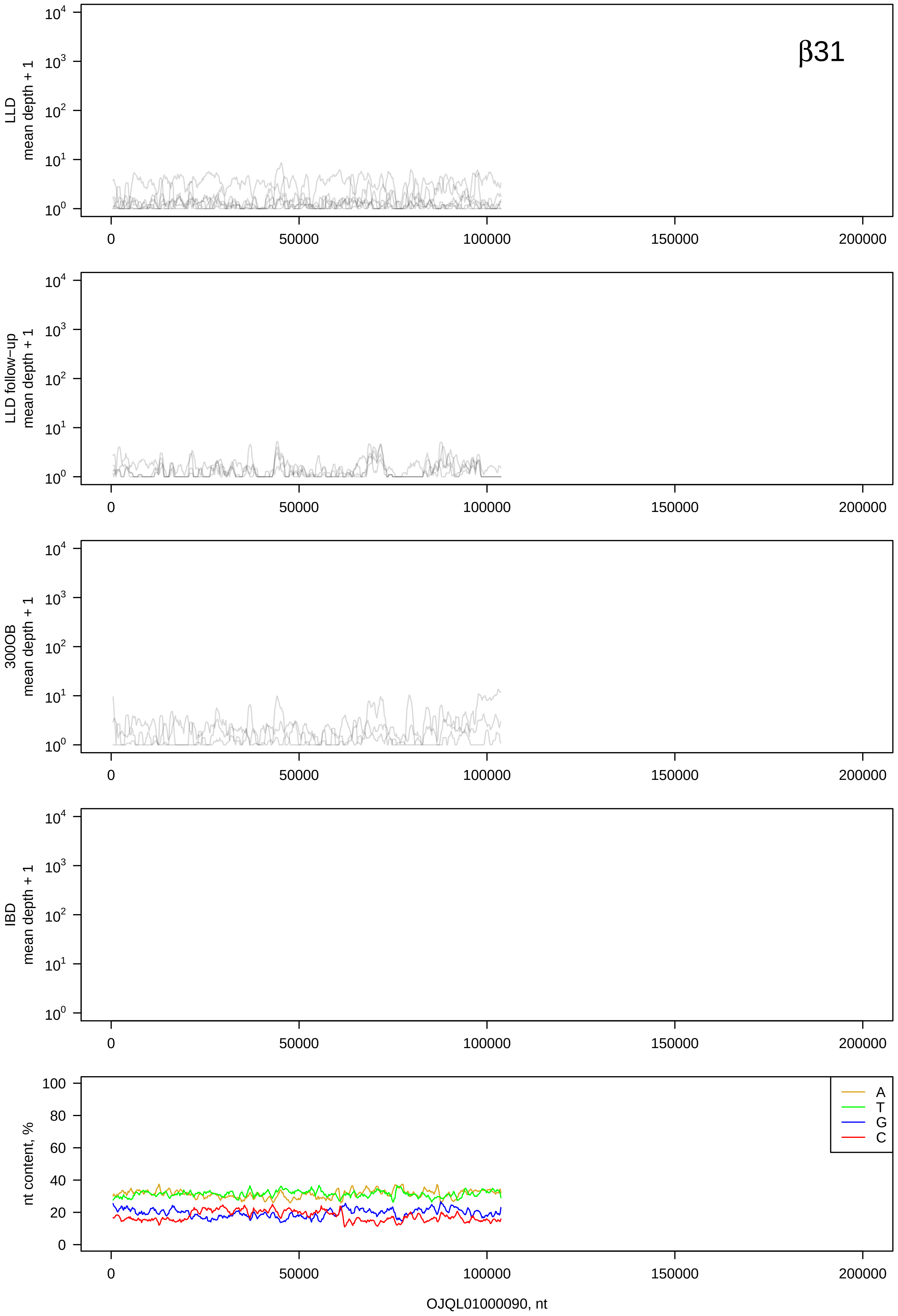

β31
LLD mean depth + 1
LLD follow−up mean depth + 1
300OB mean depth + 1
IBD mean depth + 1
nt content, %
A
T
G
C
0
50000
100000
150000
200000
OJQL01000090, nt

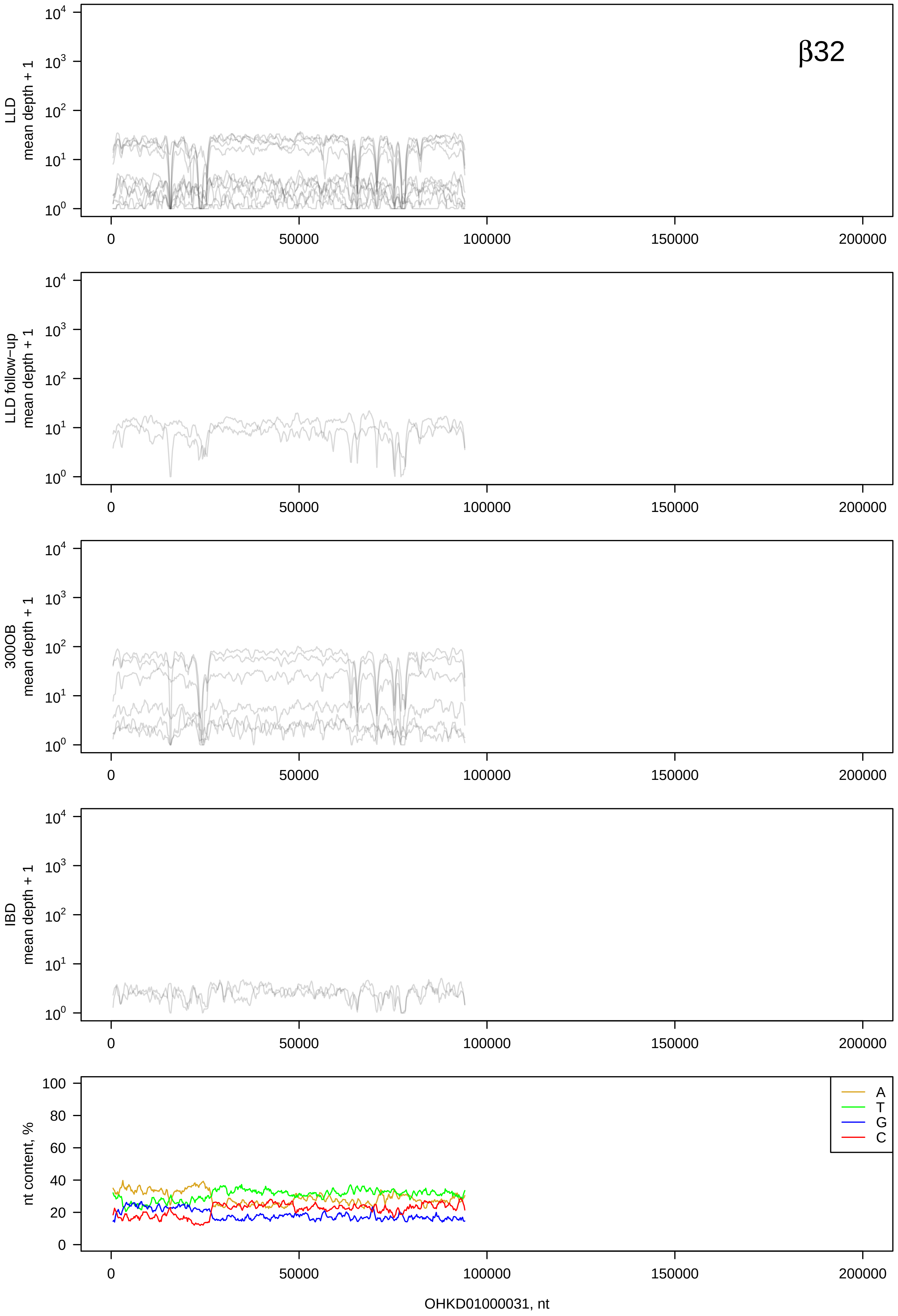
β32
LLD mean depth + 1
LLD follow−up mean depth + 1
300OB mean depth + 1
IBD mean depth + 1
nt content, %
A
T
G
C
0
50000
100000
150000
200000
OHKD01000031, nt

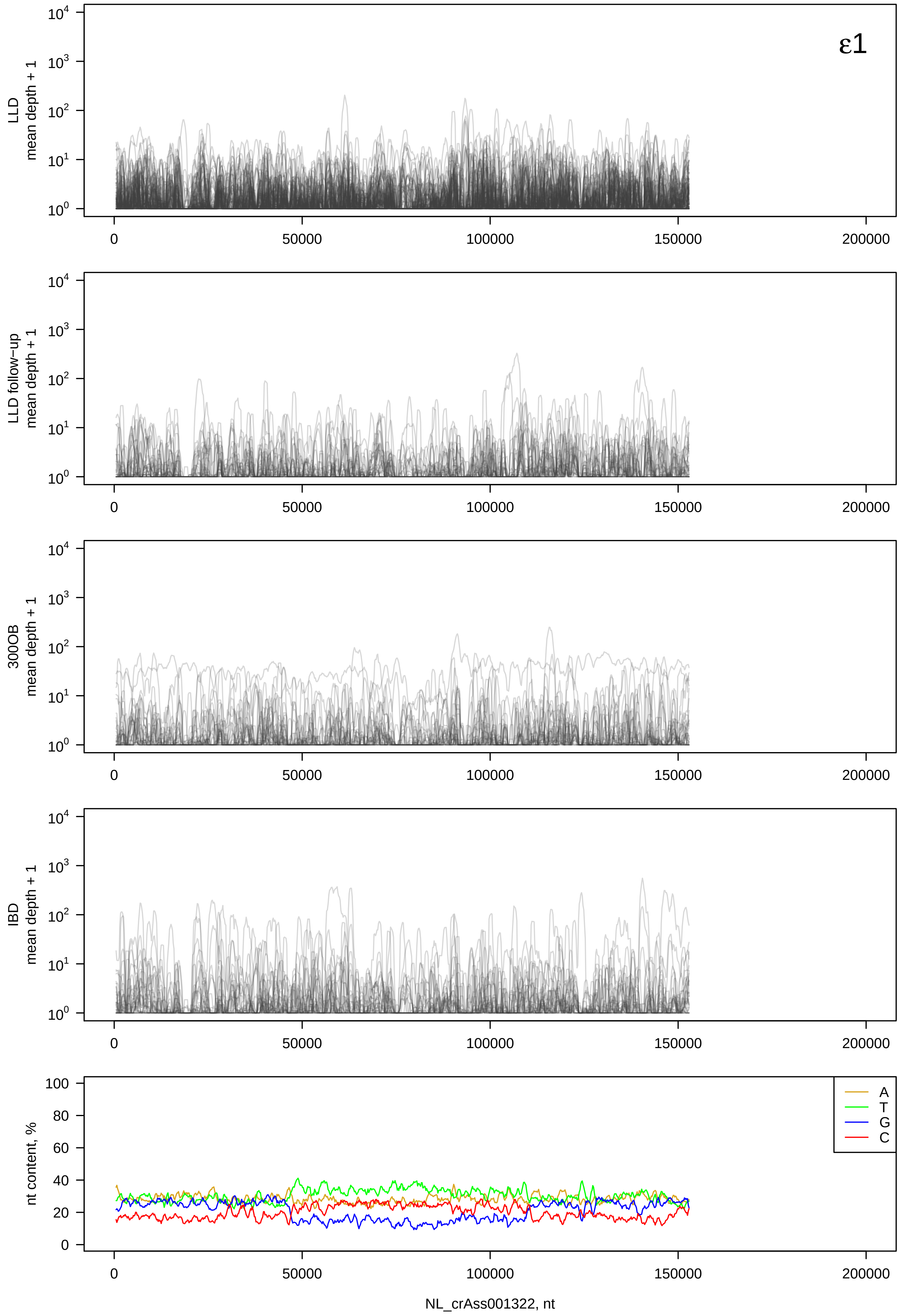

ε1
LLD mean depth + 1
LLD follow−up mean depth + 1
300OB mean depth + 1
IBD mean depth + 1
nt content, %
A
T
G
C
0
50000
100000
150000
200000
NL_crAss001322, nt

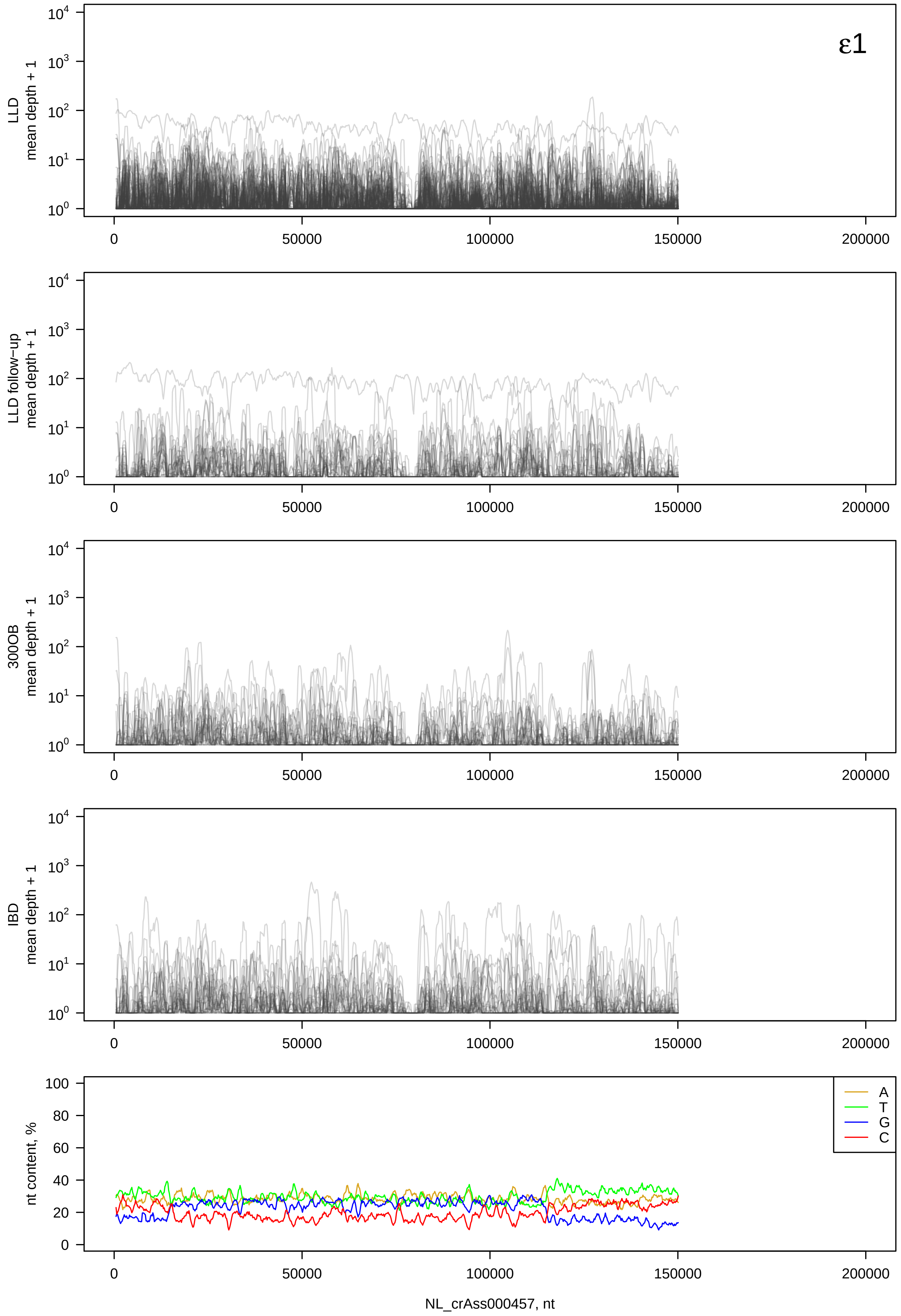

ε1
LLD mean depth + 1
LLD follow−up mean depth + 1
300OB mean depth + 1
IBD mean depth + 1
nt content, %
A
T
G
C
0
50000
100000
150000
200000
NL_crAss000457, nt

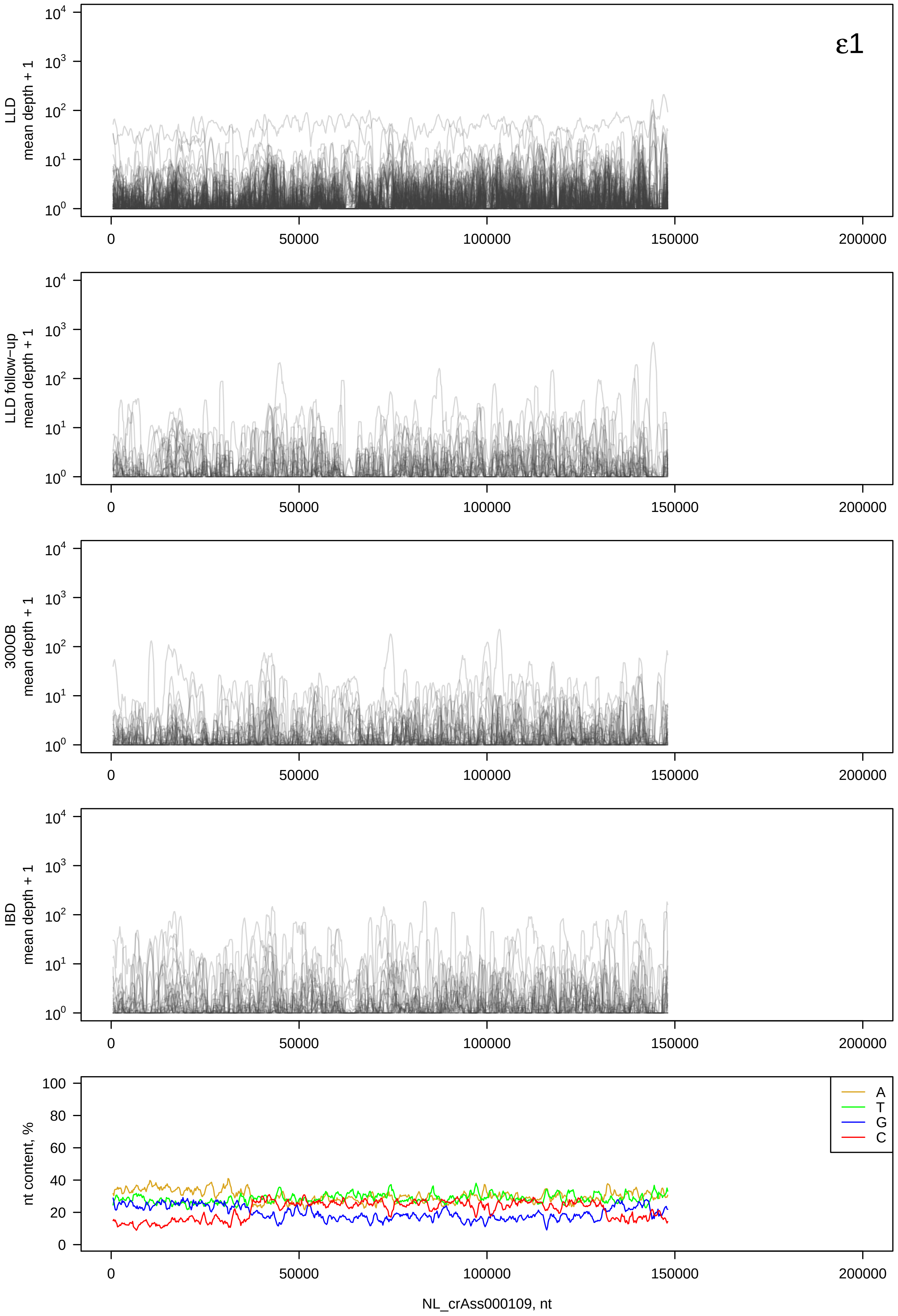
ε1
LLD
mean depth + 1
LLD follow-up
mean depth + 1
300OB
mean depth + 1
IBD
mean depth + 1
nt content, %
A
T
G
C
0
50000
100000
150000
200000
NL_crAss000109, nt

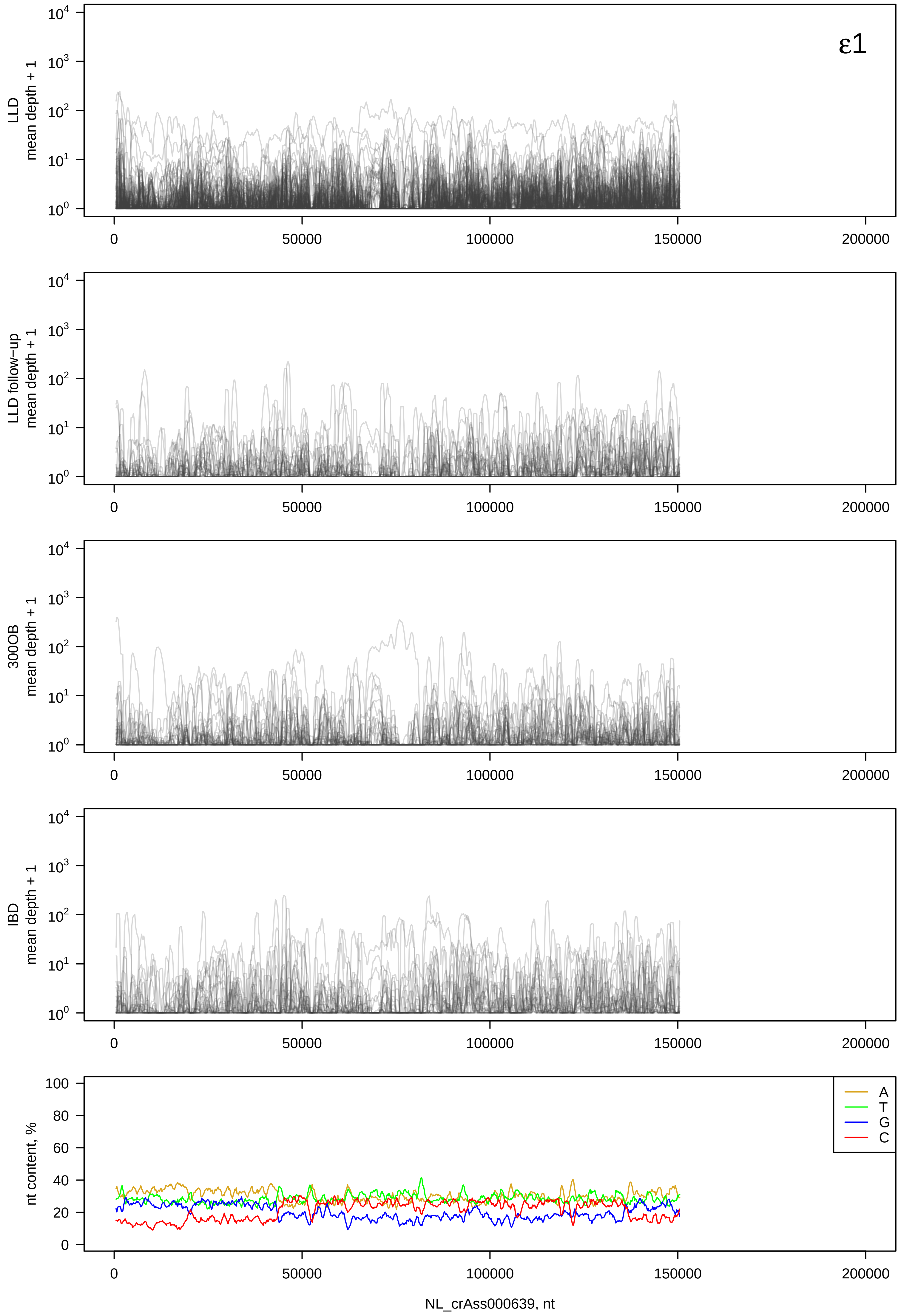
ε1
LLD mean depth + 1
LLD follow-up mean depth + 1
300OB mean depth + 1
IBD mean depth + 1
nt content, %
A
T
G
C
0
50000
100000
150000
200000
NL_crAss000639, nt

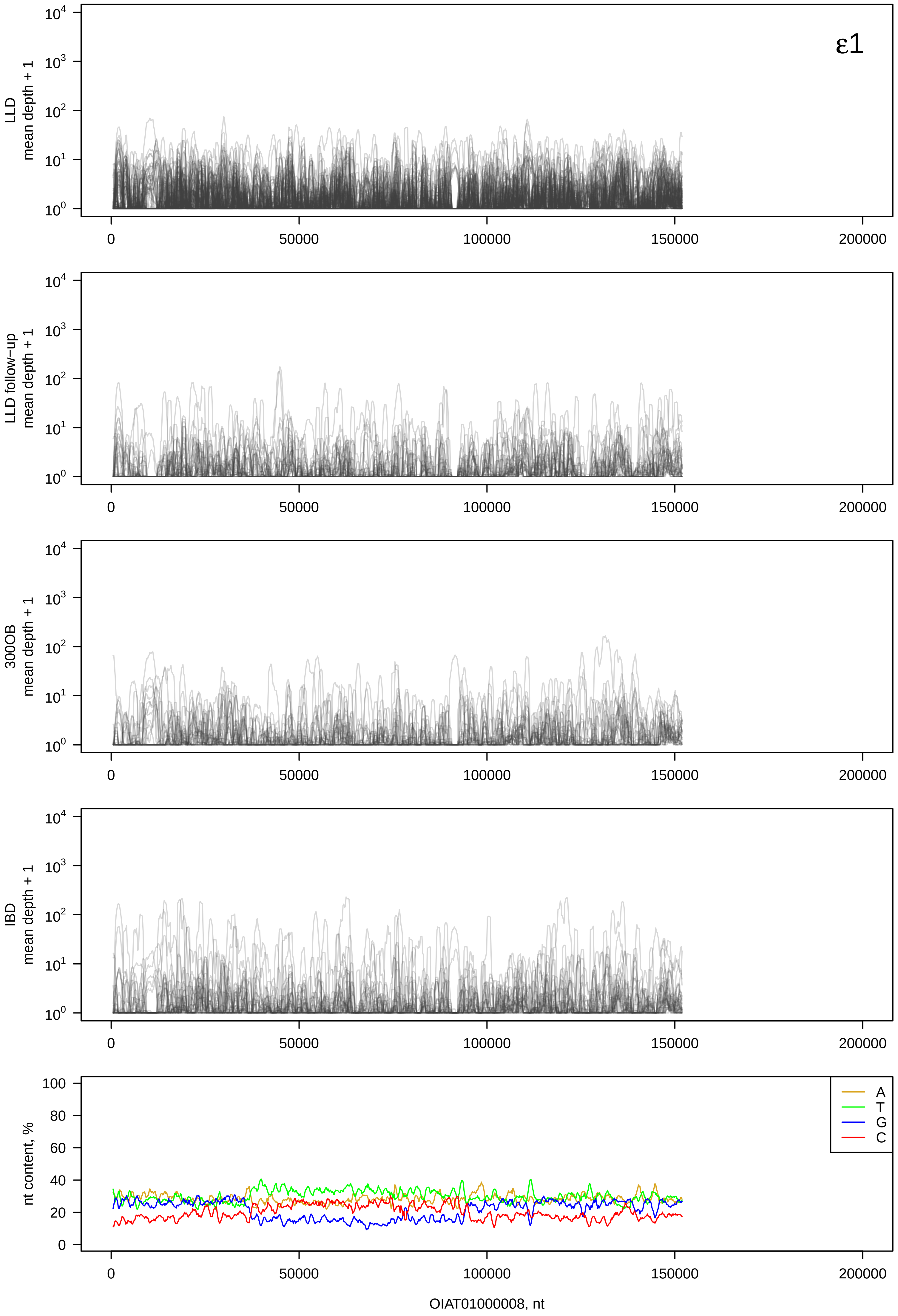

ε1
LLD mean depth + 1
LLD follow−up mean depth + 1
300OB mean depth + 1
IBD mean depth + 1
nt content, %
A
T
G
C
0
50000
100000
150000
200000
OIAT01000008, nt

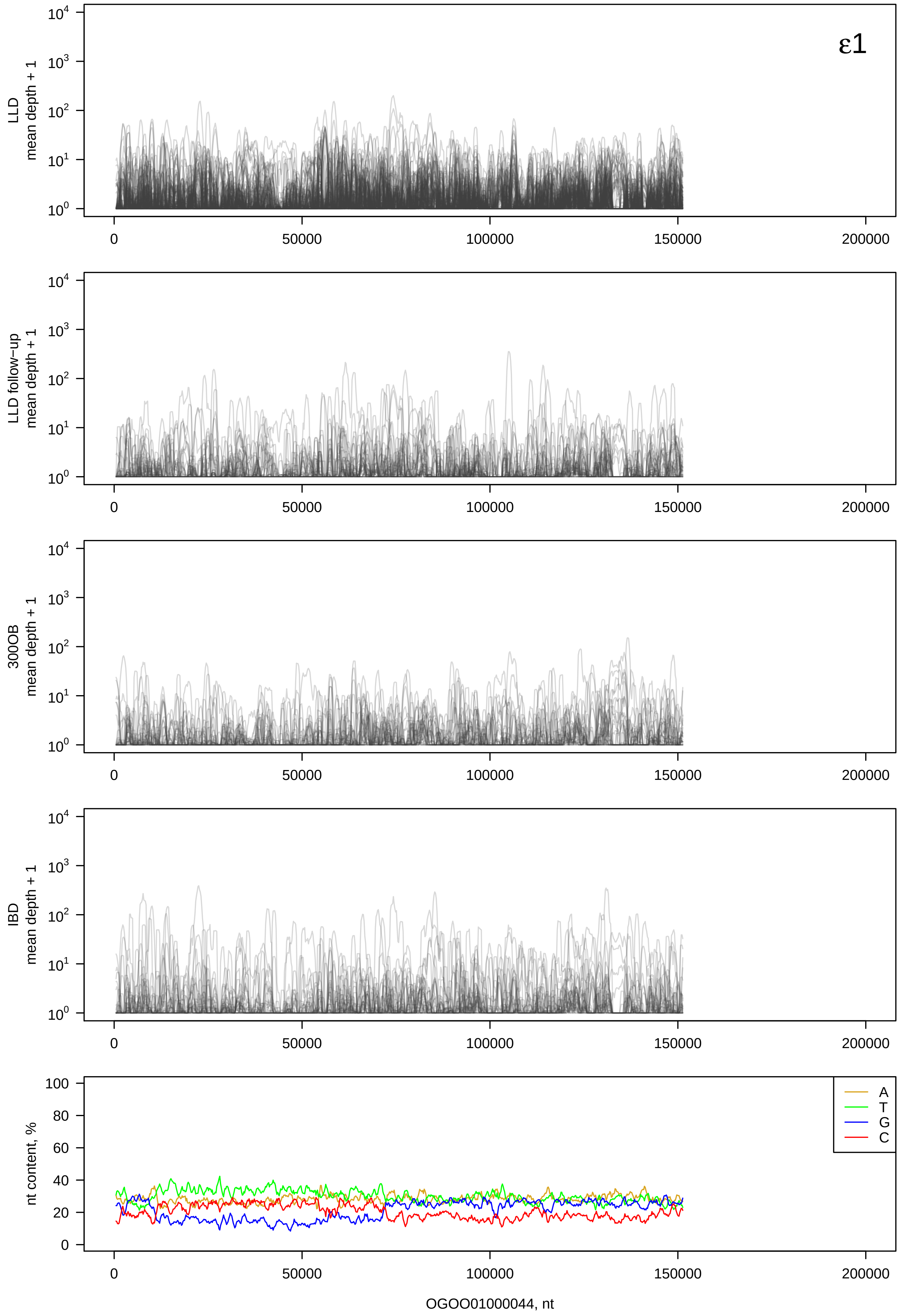
ε1
LLD mean depth + 1
LLD follow-up mean depth + 1
300OB mean depth + 1
IBD mean depth + 1
nt content, %
A
T
G
C
OGOO01000044, nt

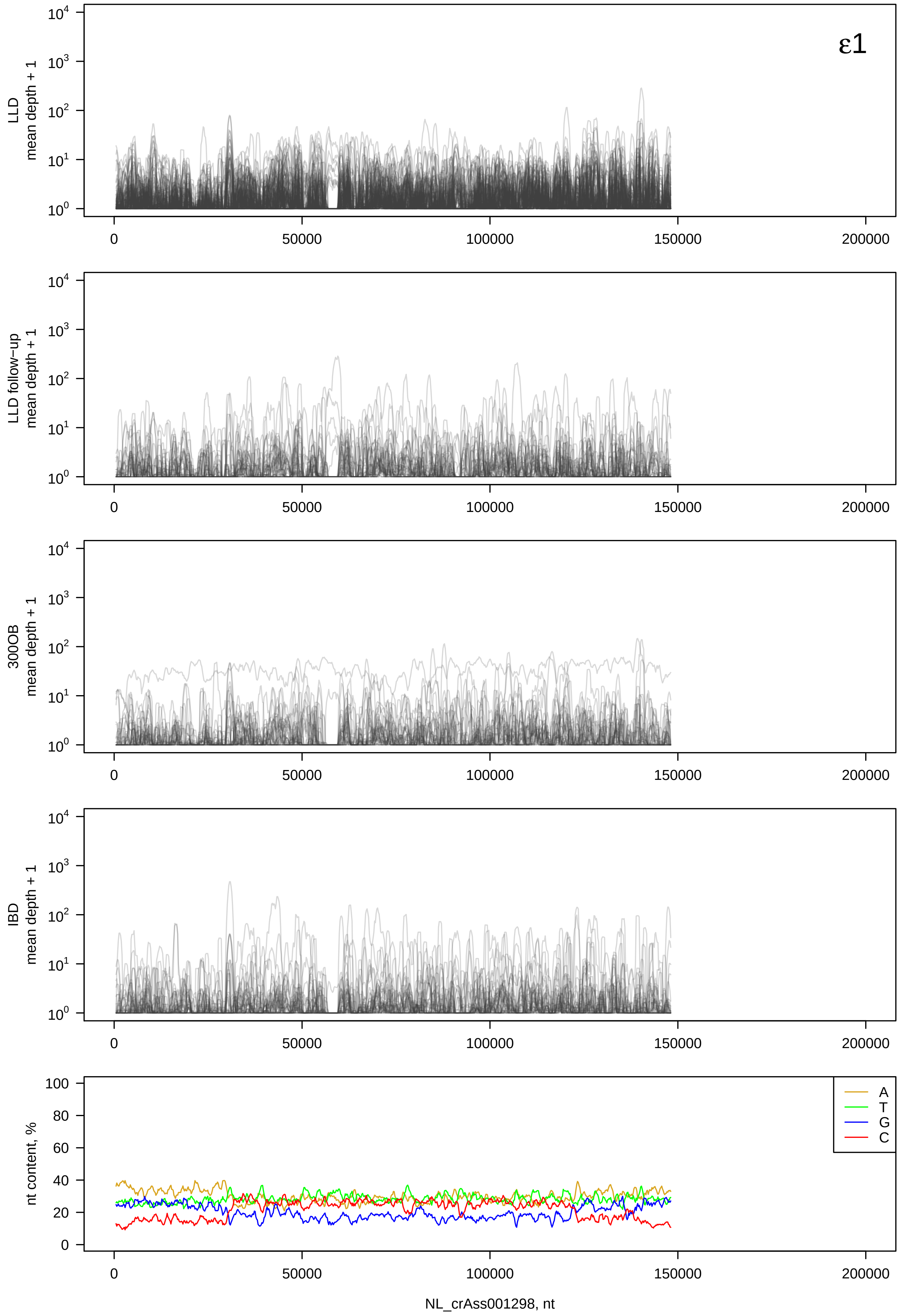

ε1
LLD mean depth + 1
LLD follow−up mean depth + 1
300OB mean depth + 1
IBD mean depth + 1
nt content, %
A
T
G
C
0
50000
100000
150000
200000
NL_crAss001298, nt

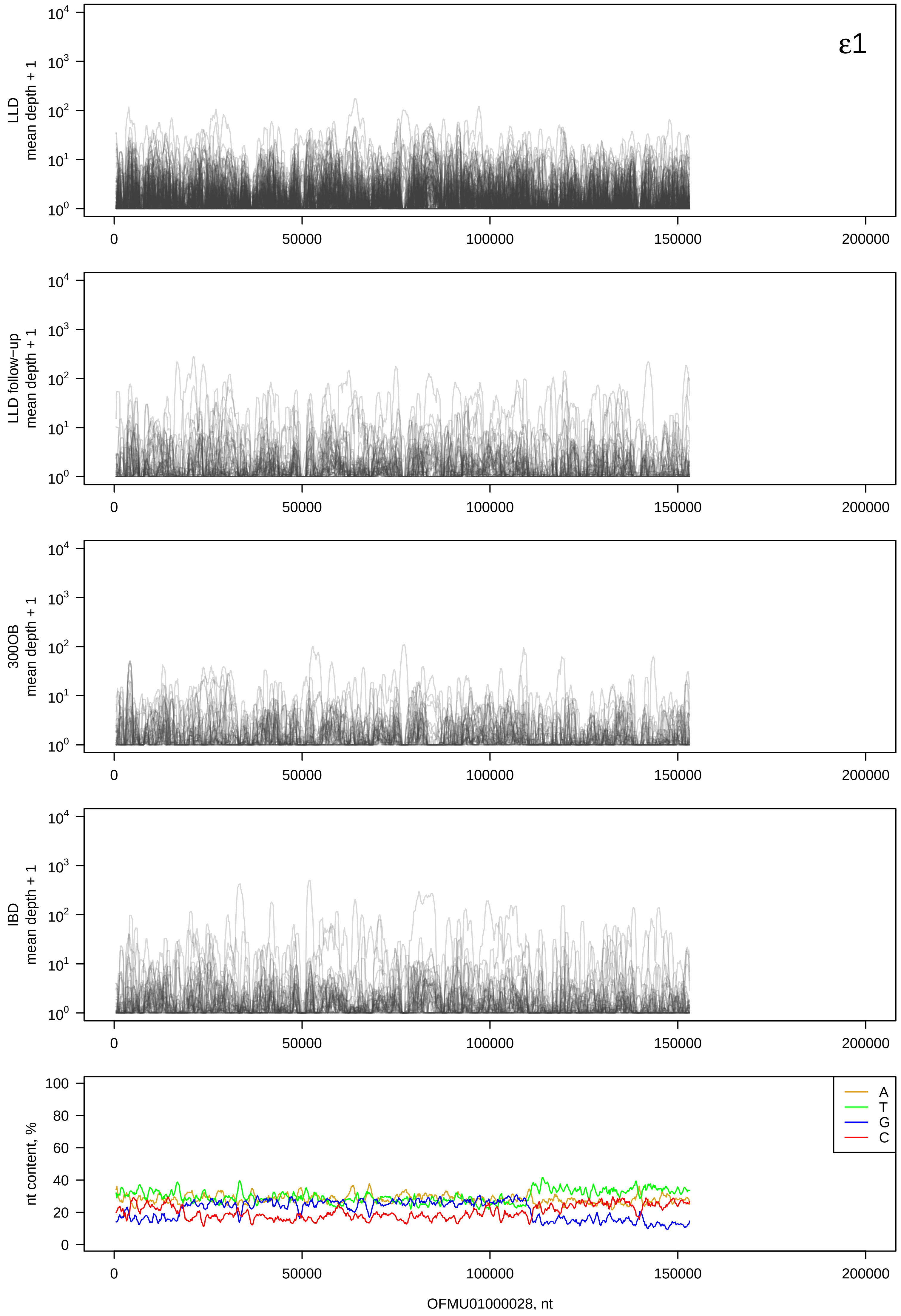

ε1
LLD mean depth + 1
LLD follow-up mean depth + 1
300OB mean depth + 1
IBD mean depth + 1
nt content, %
A
T
G
C
0
50000
100000
150000
200000
OFMU01000028, nt

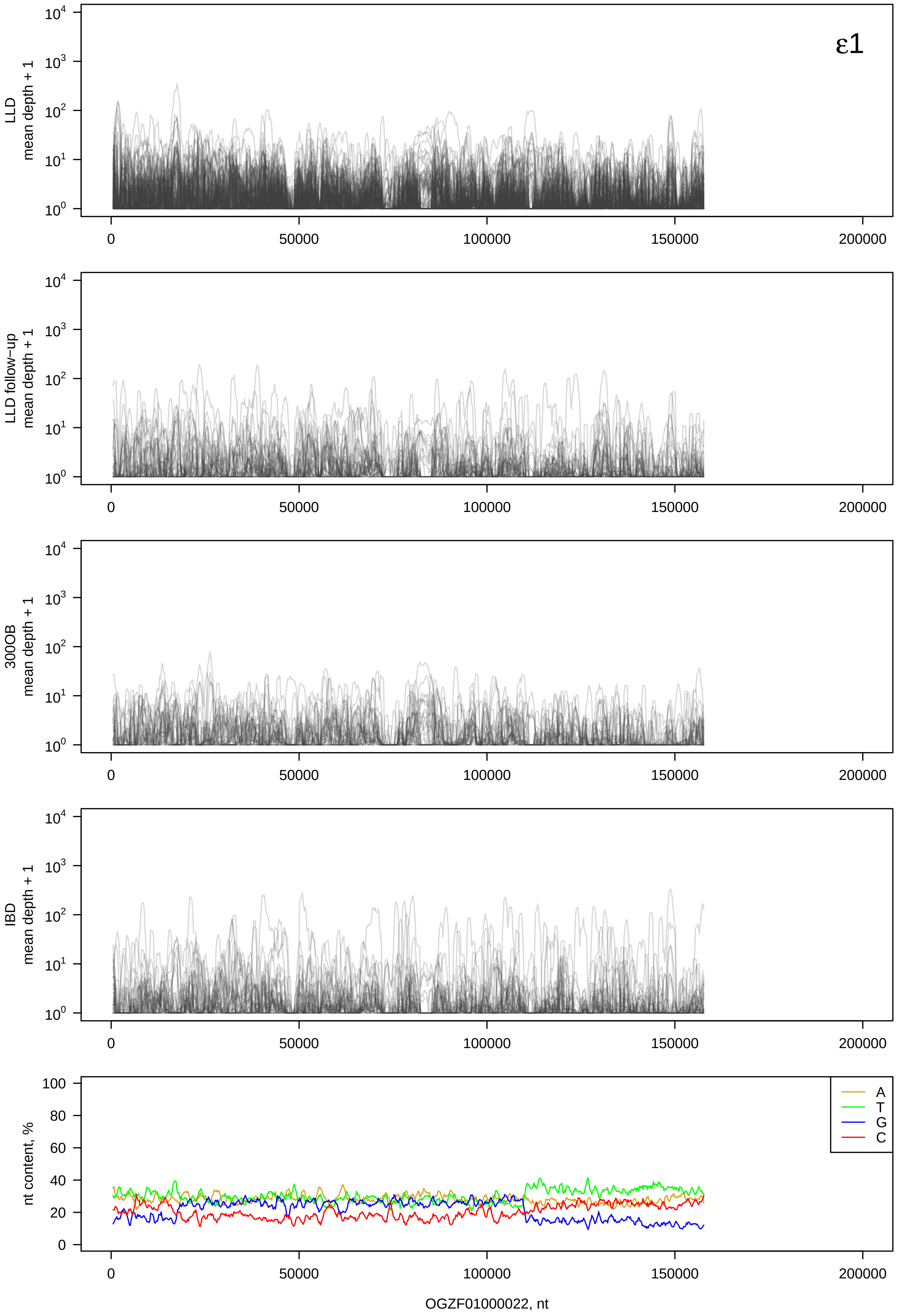

ε1
LLD mean depth + 1
LLD follow−up mean depth + 1
300OB mean depth + 1
IBD mean depth + 1
nt content, %
A
T
G
C
OGZF01000022, nt

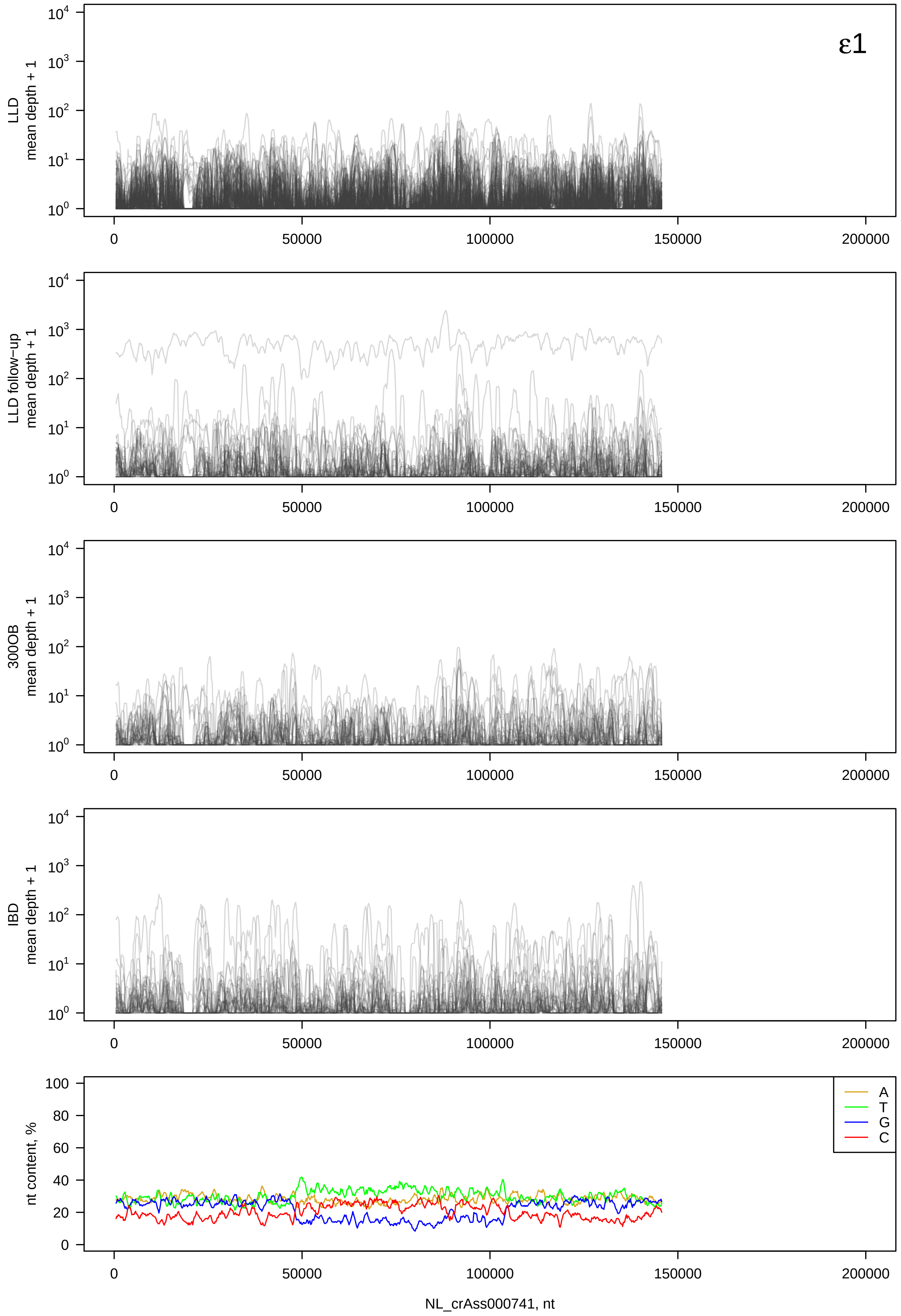
ε1
LLD mean depth + 1
LLD follow−up mean depth + 1
300OB mean depth + 1
IBD mean depth + 1
nt content, %
A
T
G
C
0
50000
100000
150000
200000
NL_crAss000741, nt

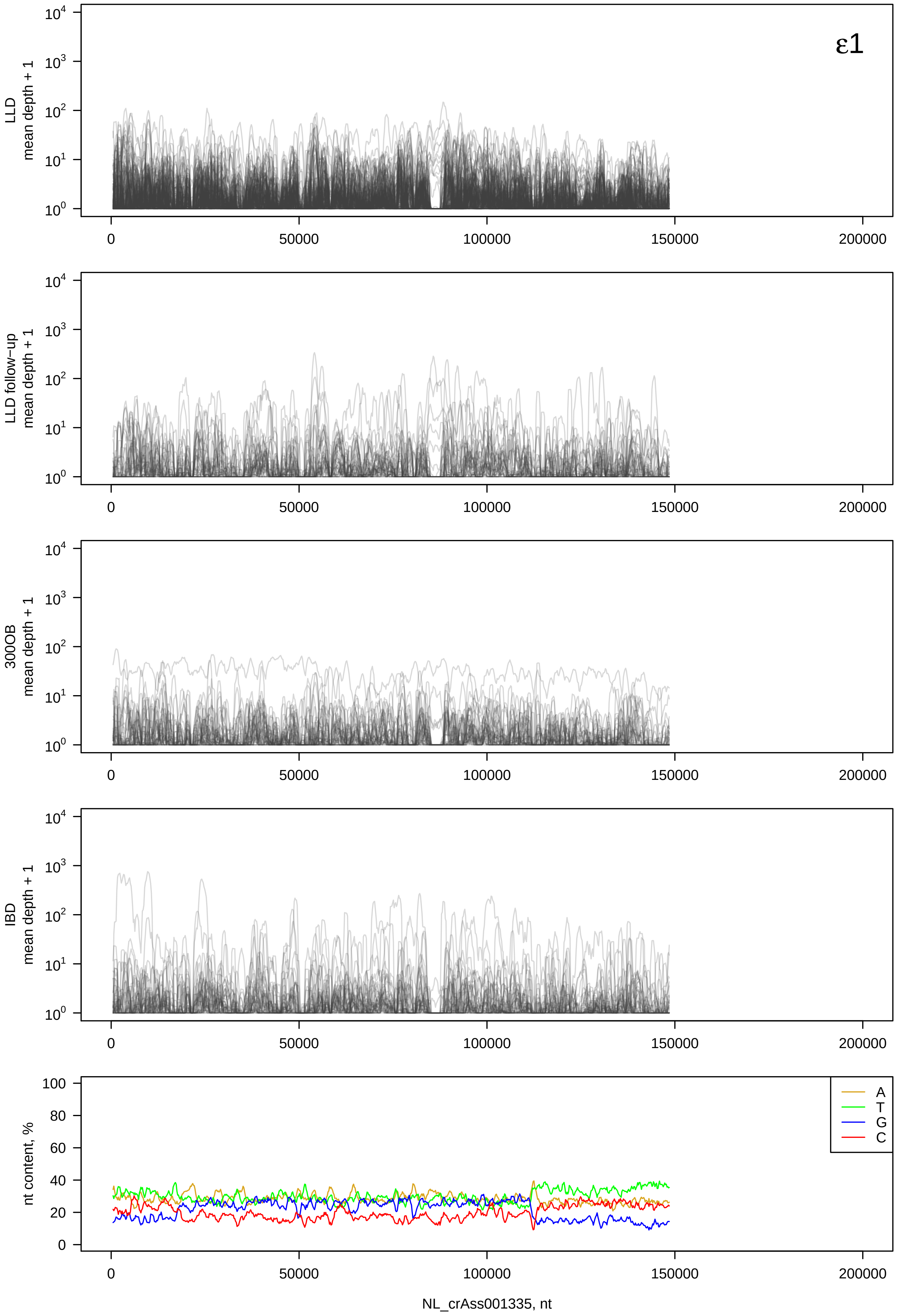

ε1
LLD mean depth + 1
LLD follow-up mean depth + 1
300OB mean depth + 1
IBD mean depth + 1
nt content, %
A
T
G
C
NL_crAss001335, nt

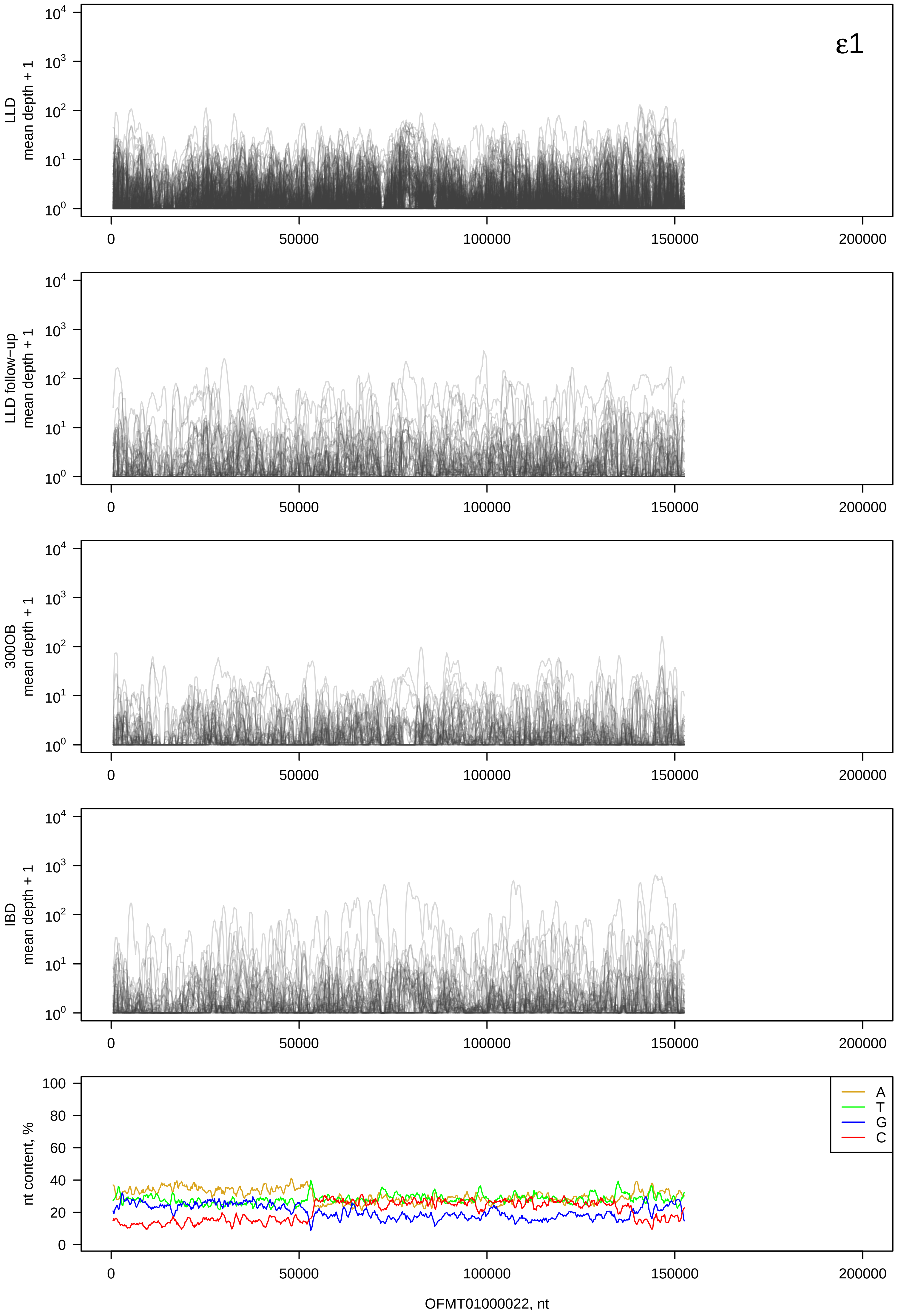

ε1
LLD mean depth + 1
LLD follow−up mean depth + 1
300OB mean depth + 1
IBD mean depth + 1
nt content, %
A
T
G
C
0
50000
100000
150000
200000
OFMT01000022, nt

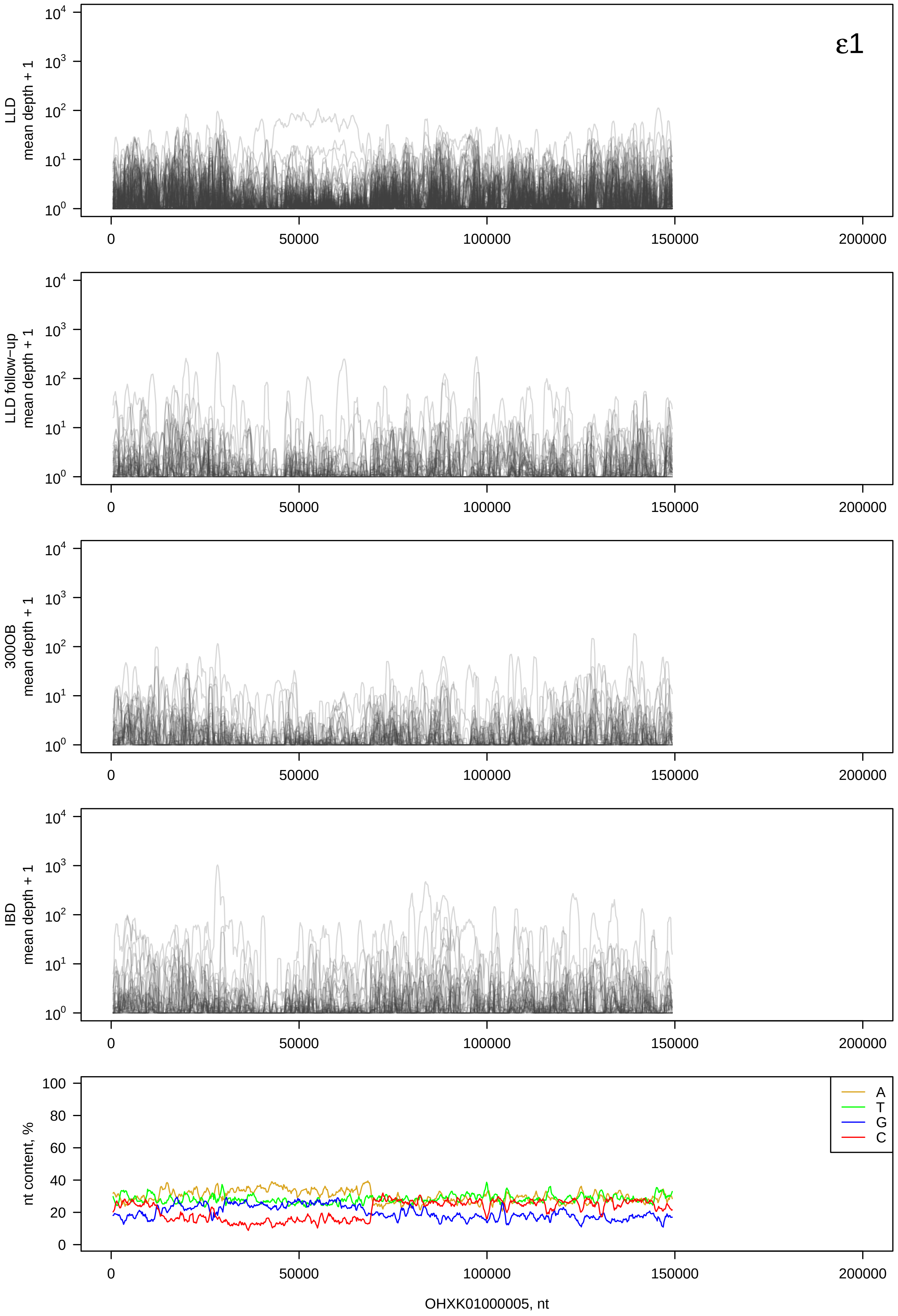

ε1
LLD mean depth + 1
LLD follow−up mean depth + 1
300OB mean depth + 1
IBD mean depth + 1
nt content, %
A
T
G
C
0
50000
100000
150000
200000
OHXK01000005, nt

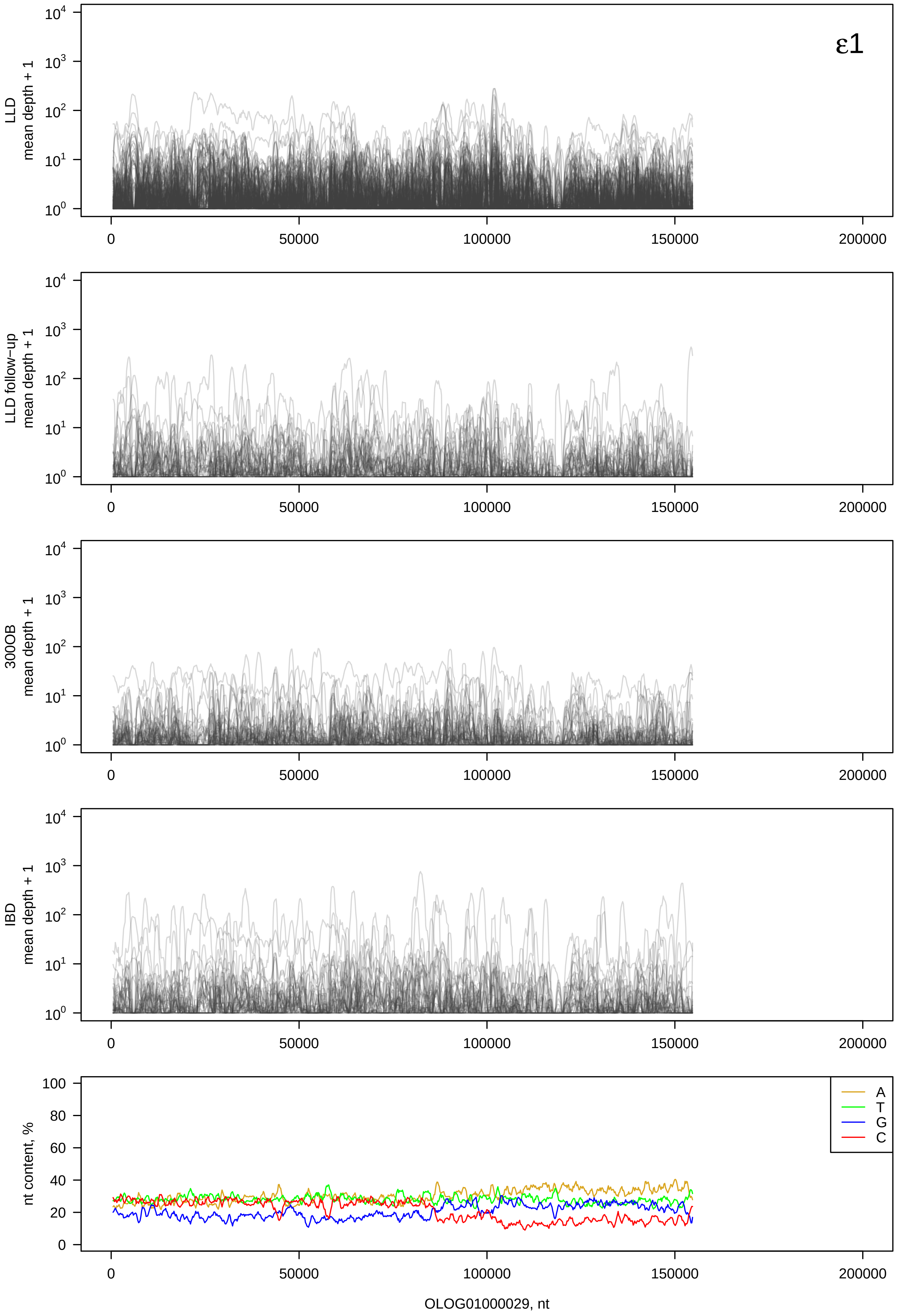

ε1
LLD mean depth + 1
LLD follow-up mean depth + 1
300OB mean depth + 1
IBD mean depth + 1
nt content, %
A
T
G
C
OLOG01000029, nt

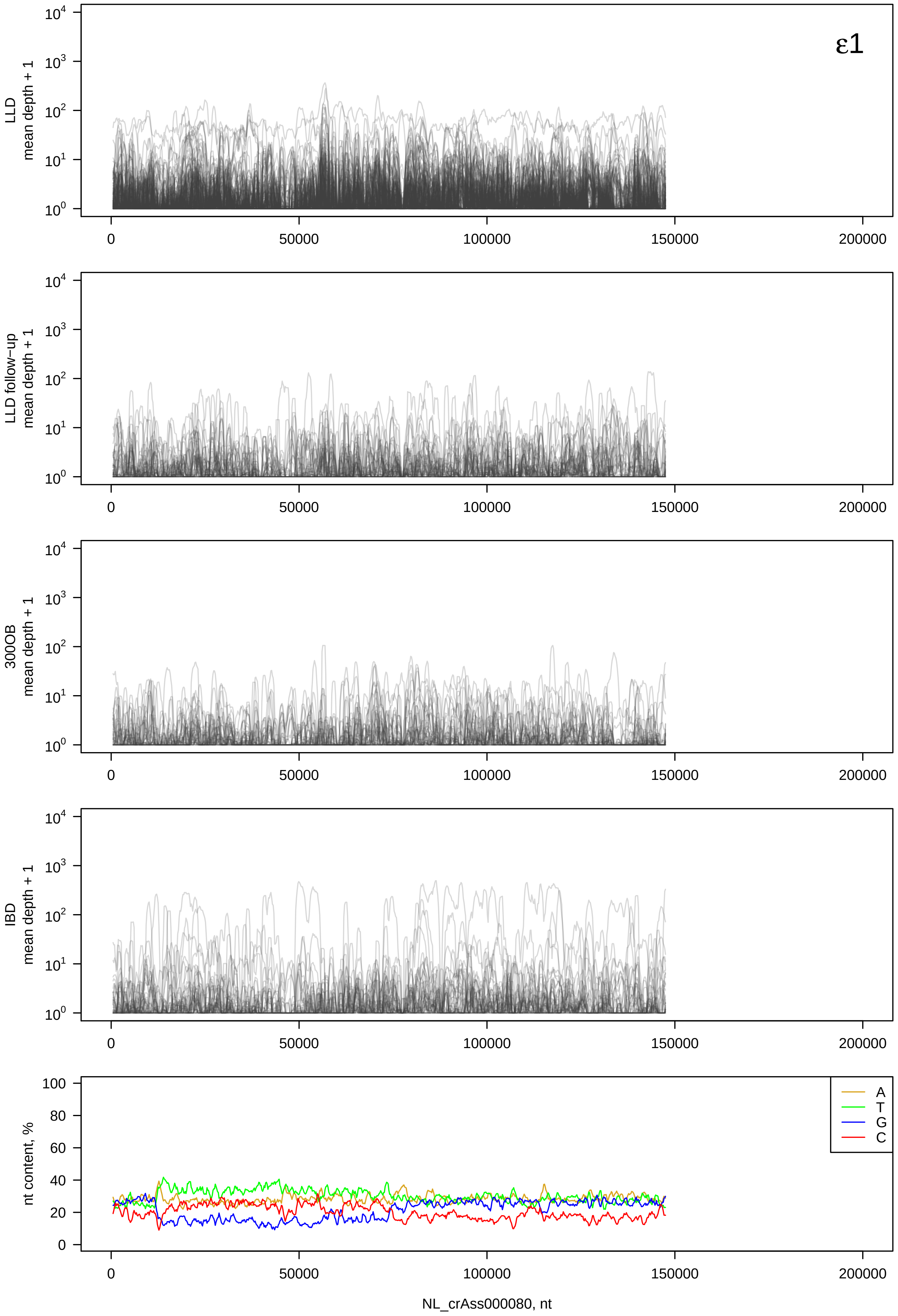

ε1
LLD mean depth + 1
LLD follow-up mean depth + 1
300OB mean depth + 1
IBD mean depth + 1
nt content, %
A
T
G
C
NL_crAss000080, nt

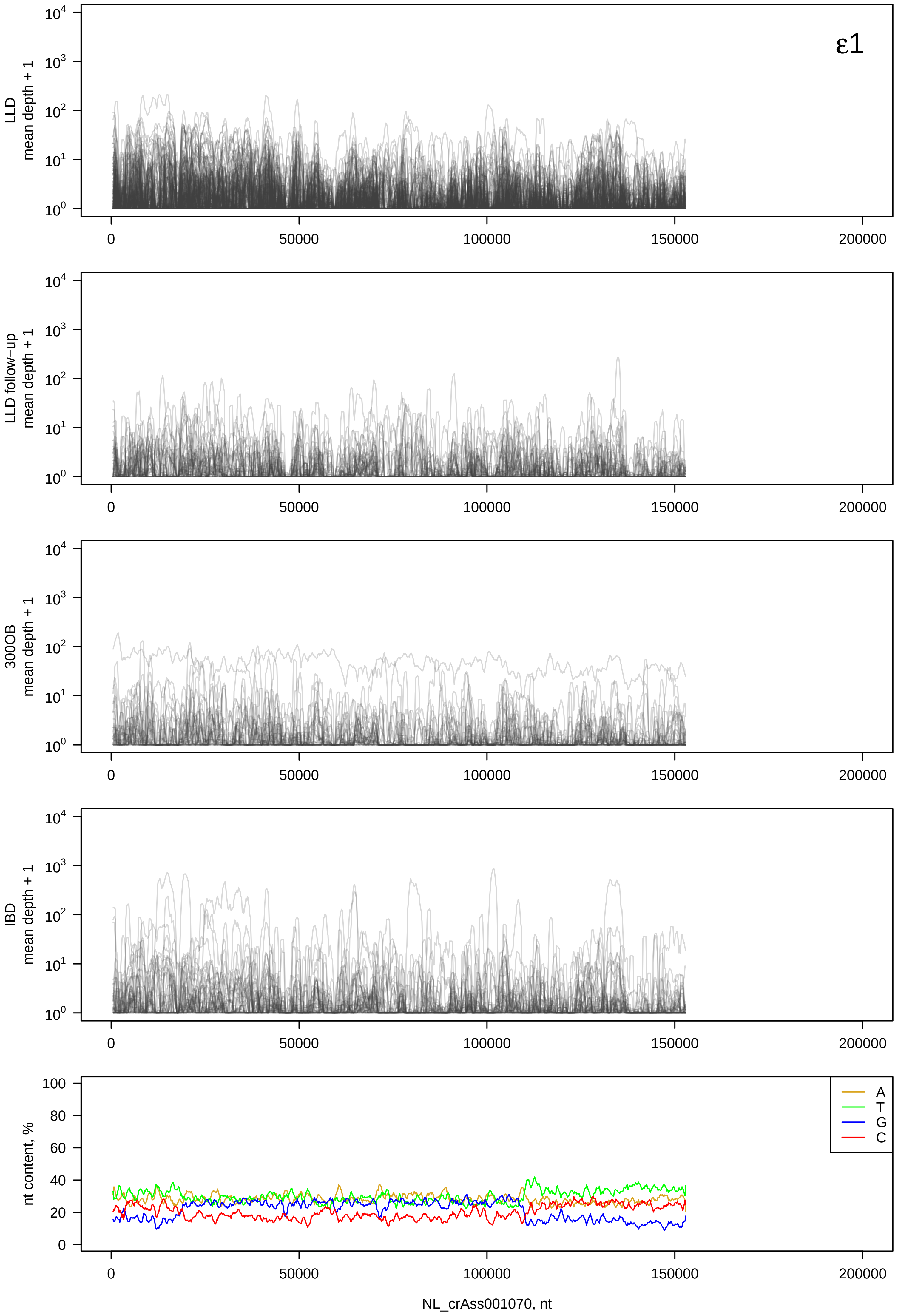

ε1
LLD mean depth + 1
LLD follow-up mean depth + 1
300OB mean depth + 1
IBD mean depth + 1
nt content, %
A
T
G
C
0
50000
100000
150000
200000
NL_crAss001070, nt

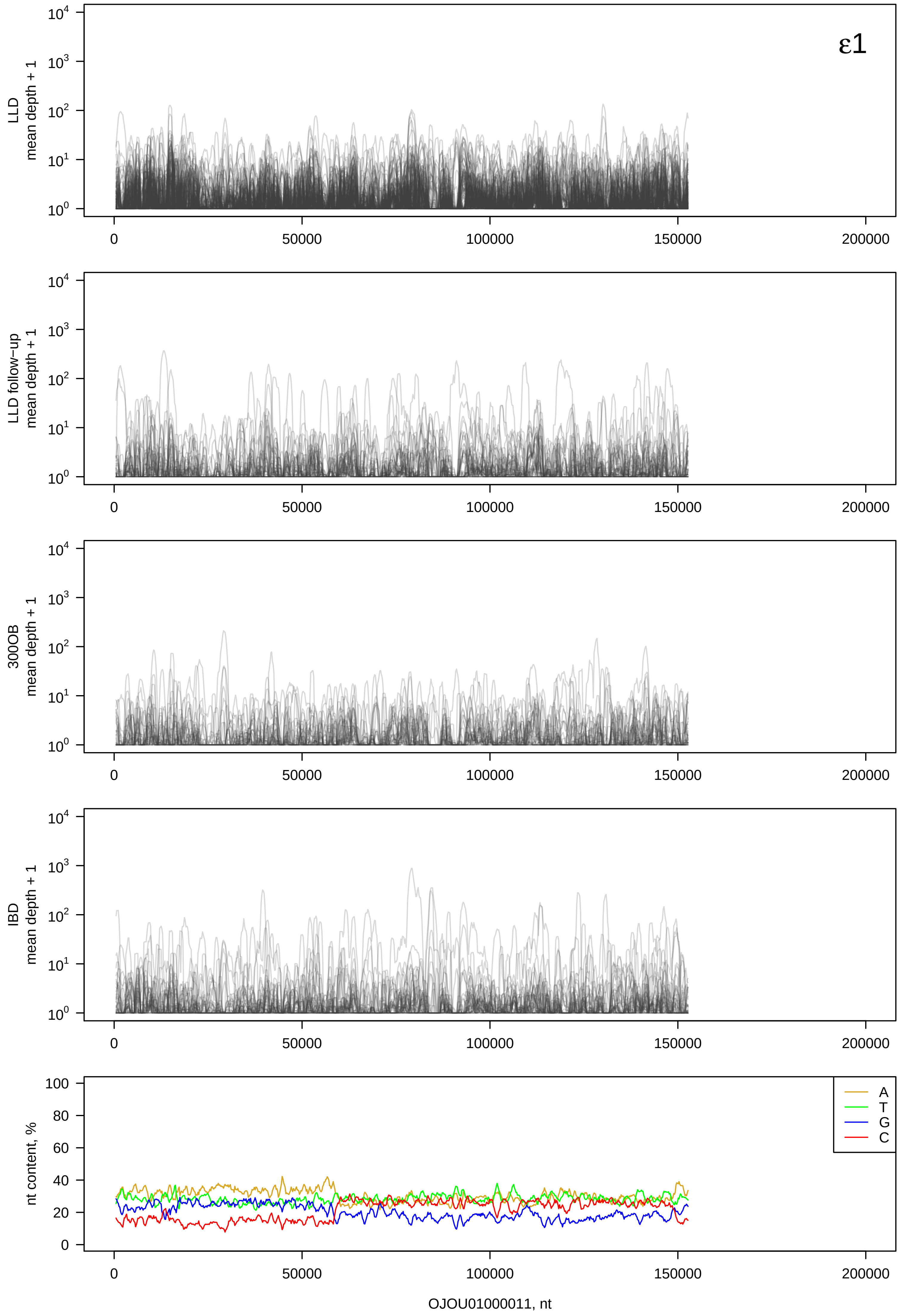

ε1
LLD mean depth + 1
LLD follow-up mean depth + 1
300OB mean depth + 1
IBD mean depth + 1
nt content, %
A
T
G
C
OJOU01000011, nt

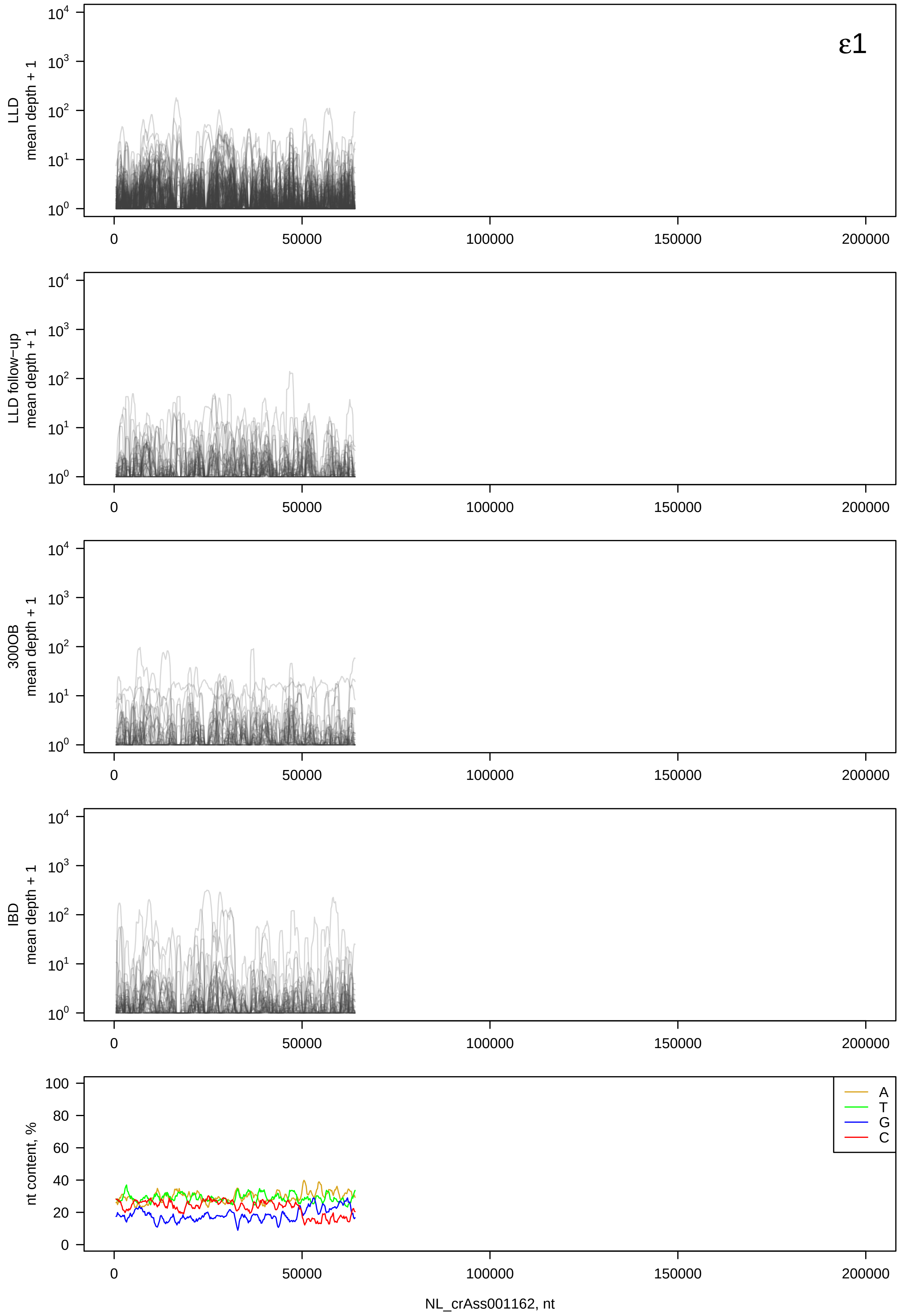

ε1
LLD mean depth + 1
LLD follow−up mean depth + 1
300OB mean depth + 1
IBD mean depth + 1
nt content, %
A
T
G
C
0
50000
100000
150000
200000
NL_crAss001162, nt

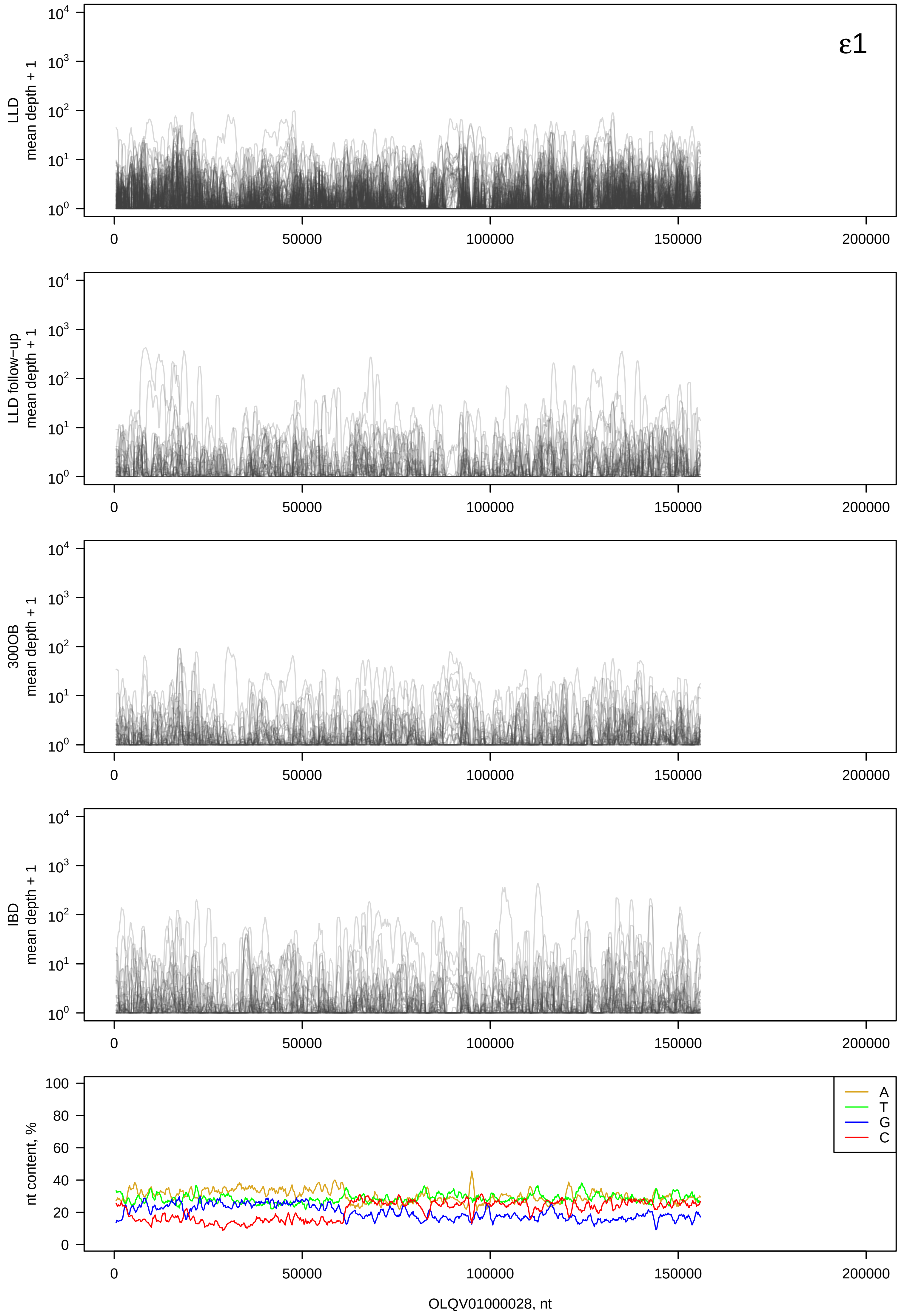

ε1
LLD mean depth + 1
LLD follow−up mean depth + 1
300OB mean depth + 1
IBD mean depth + 1
nt content, %
A
T
G
C
OLQV01000028, nt

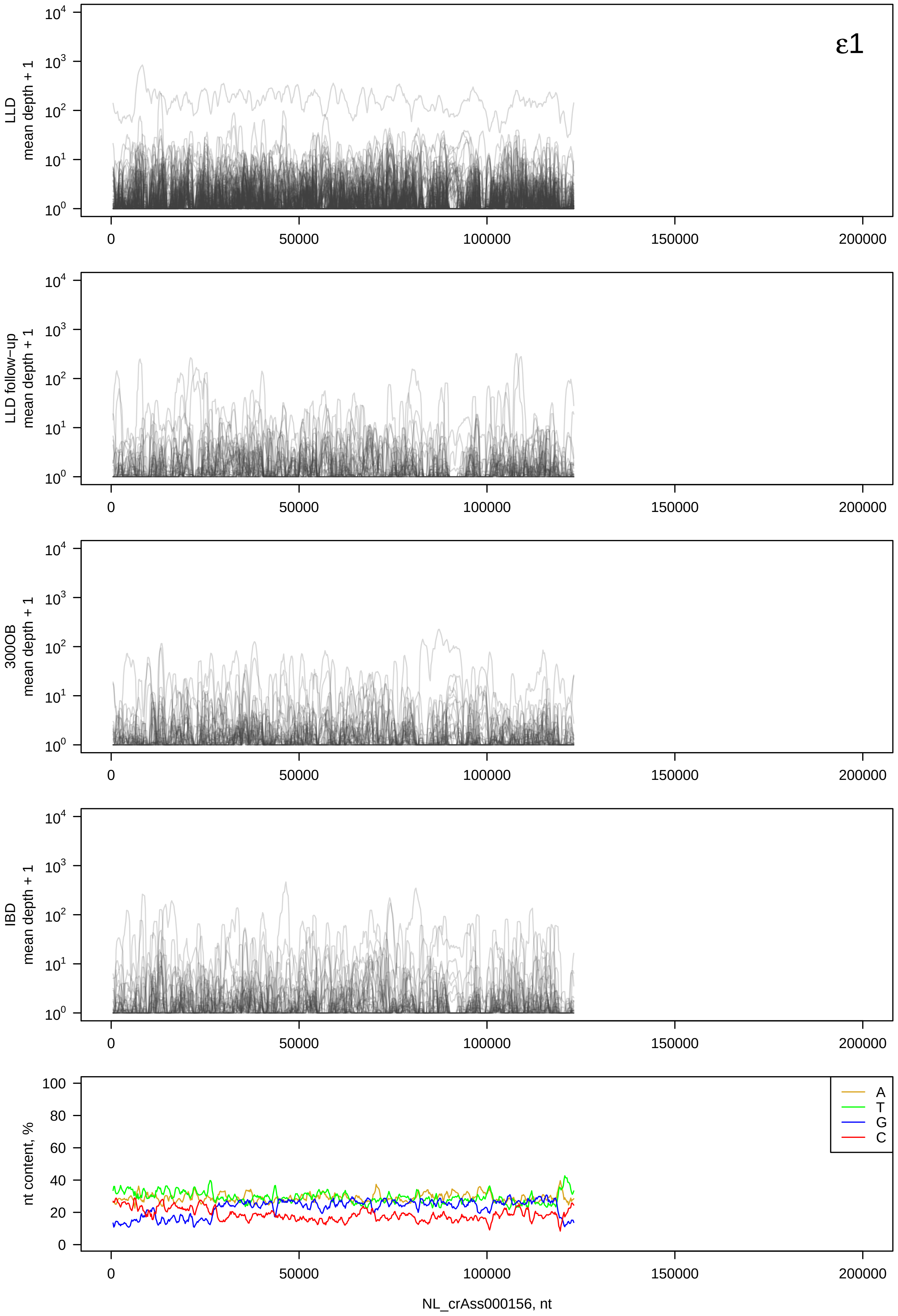

ε1
LLD mean depth + 1
LLD follow-up mean depth + 1
300OB mean depth + 1
IBD mean depth + 1
nt content, %
A
T
G
C
0
50000
100000
150000
200000
NL_crAss000156, nt

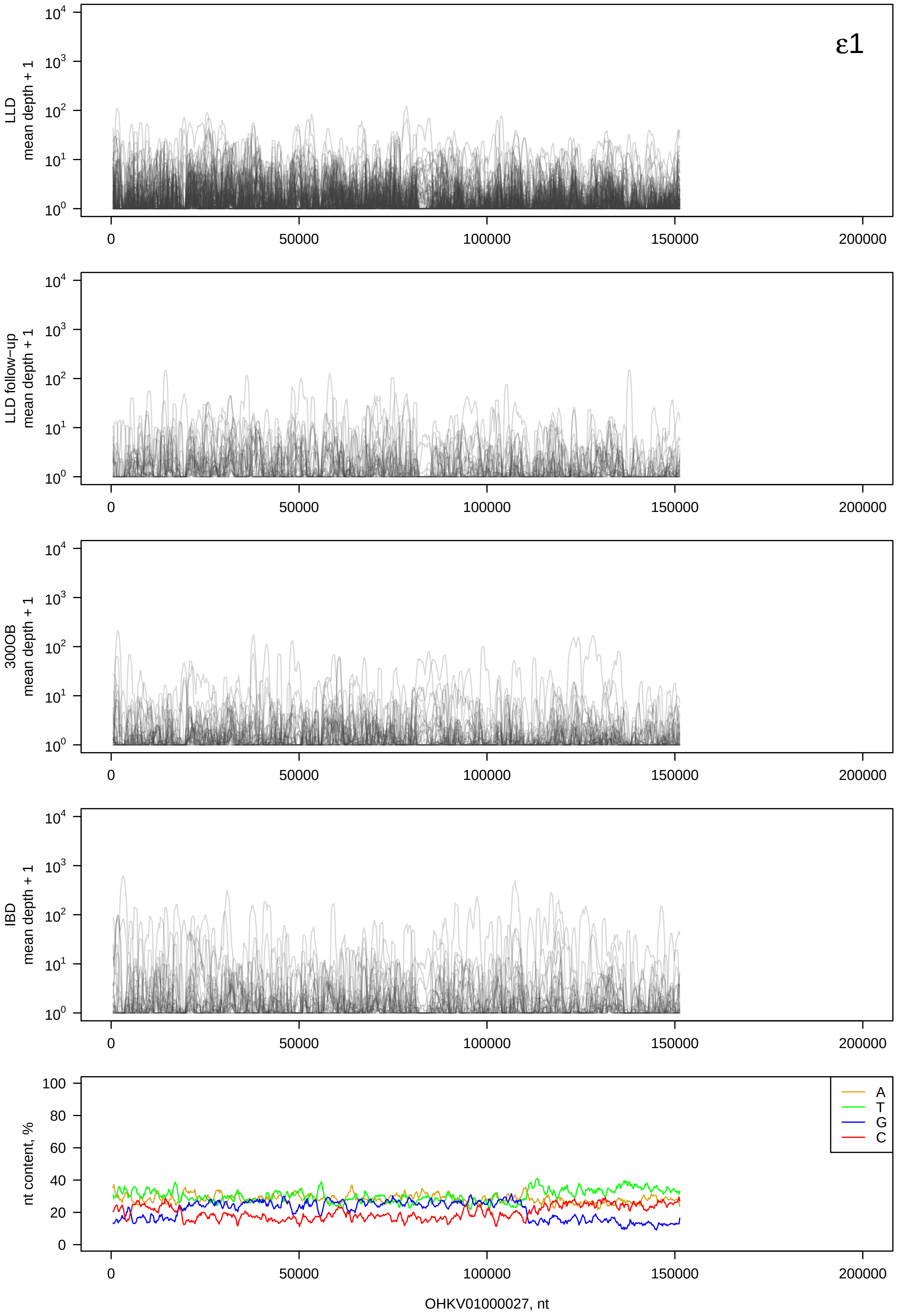

ε1
LLD mean depth + 1
LLD follow−up mean depth + 1
300OB mean depth + 1
IBD mean depth + 1
nt content, %
A
T
G
C
OHKV01000027, nt

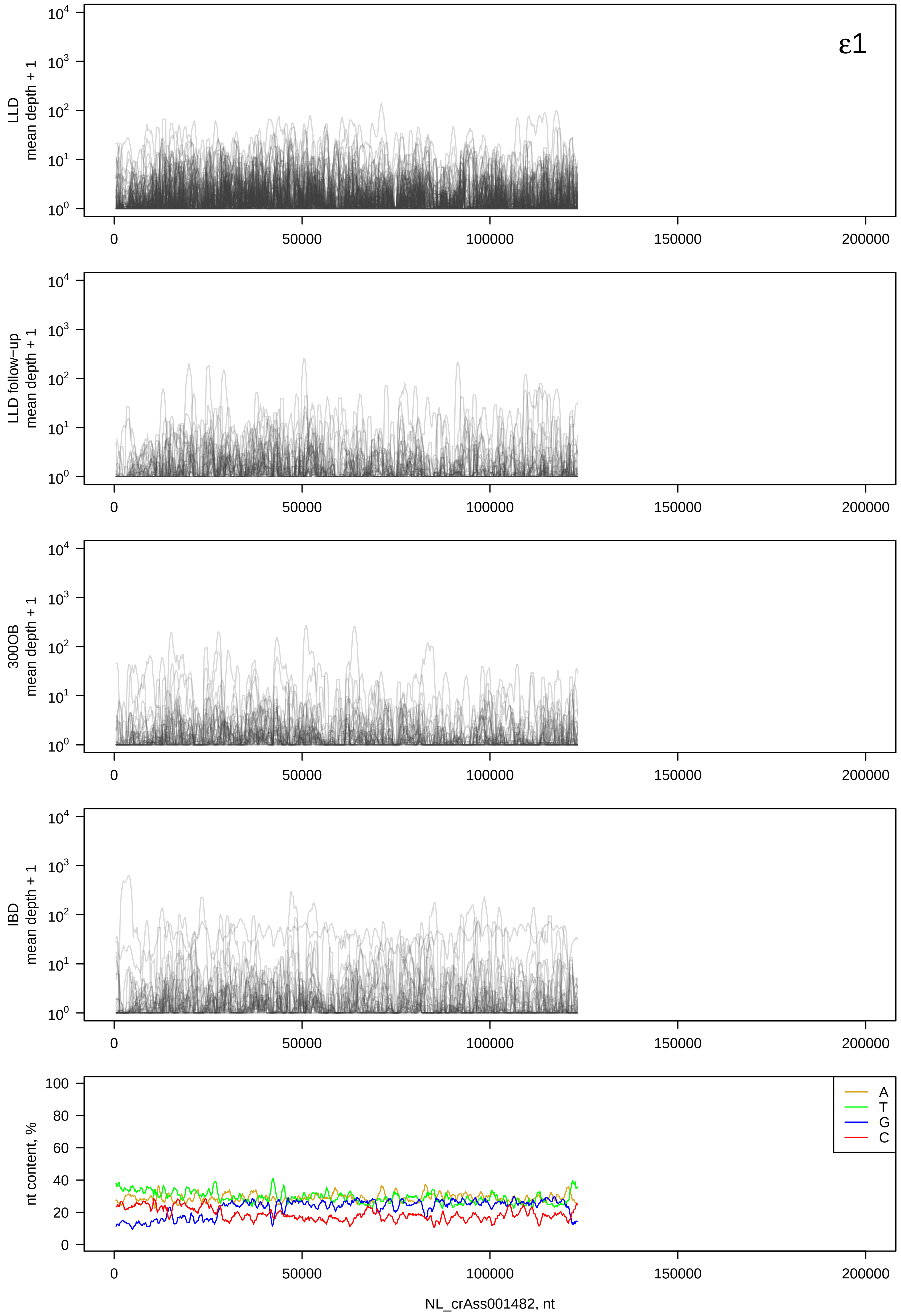

ε1
LLD mean depth + 1
LLD follow−up mean depth + 1
300OB mean depth + 1
IBD mean depth + 1
nt content, %
A
T
G
C
0
50000
100000
150000
200000
NL_crAss001482, nt

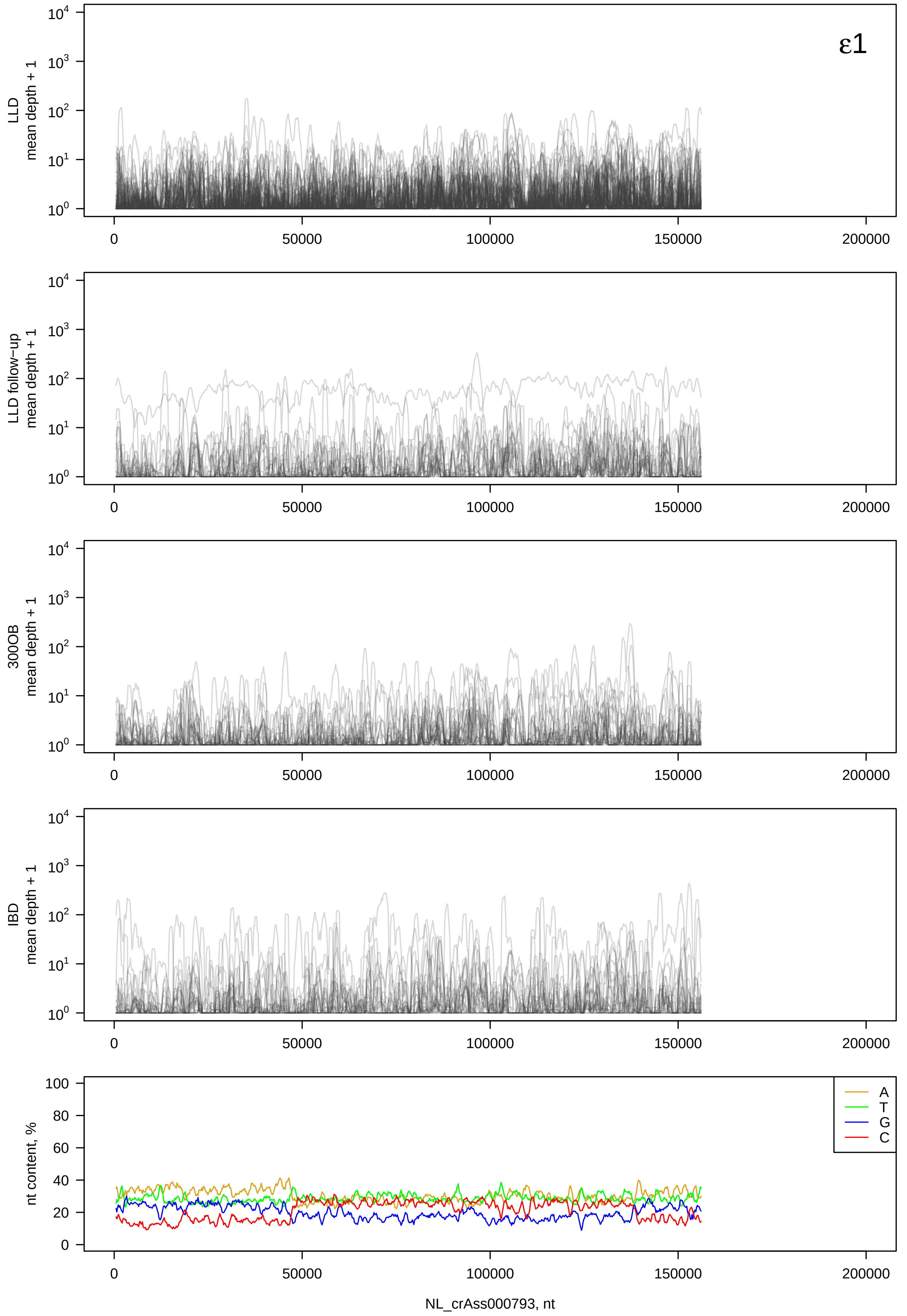

ε1
LLD mean depth + 1
LLD follow−up mean depth + 1
300OB mean depth + 1
IBD mean depth + 1
nt content, %
A
T
G
C
0
50000
100000
150000
200000
NL_crAss000793, nt

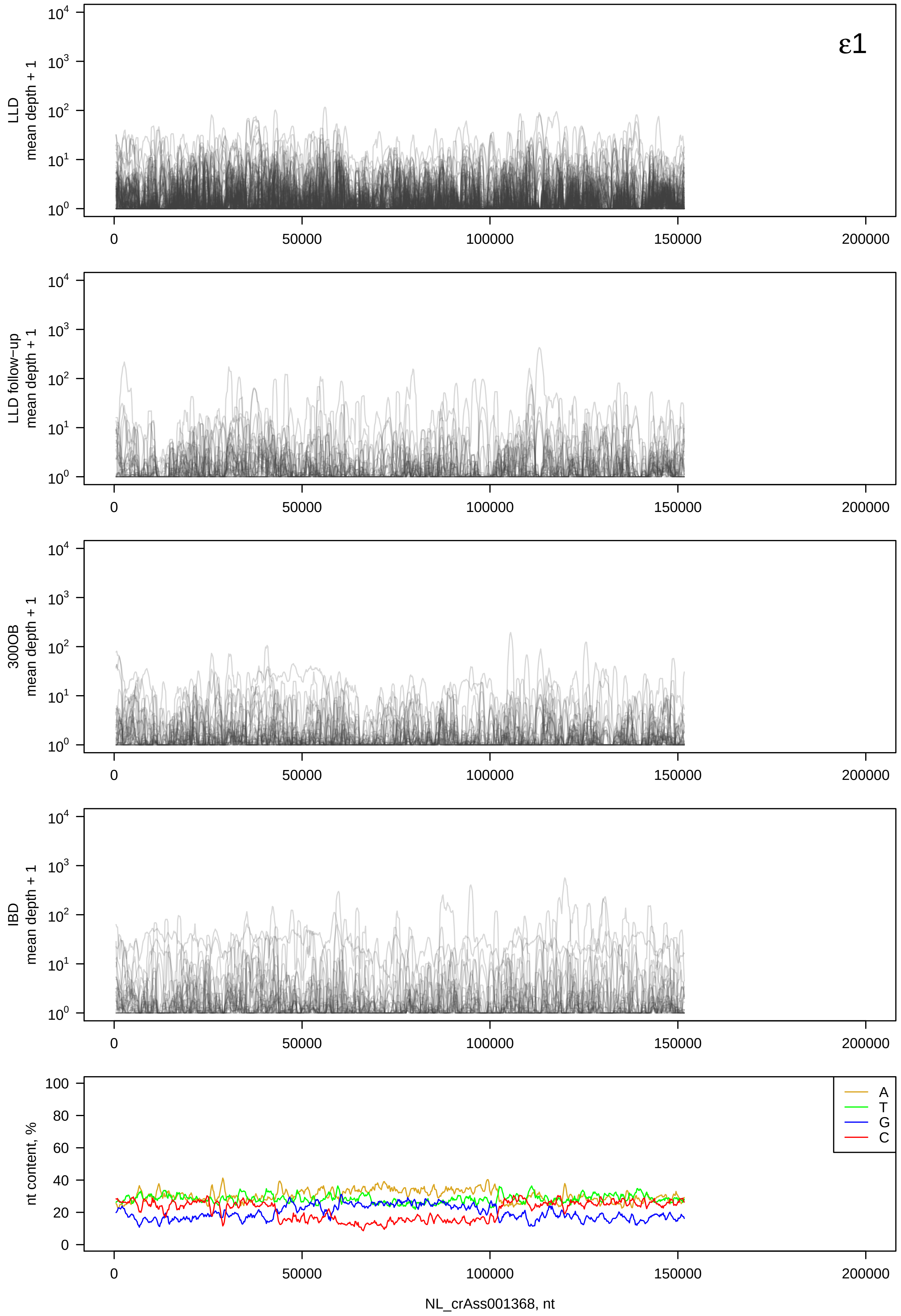

ε1
LLD mean depth + 1
LLD follow-up mean depth + 1
300OB mean depth + 1
IBD mean depth + 1
nt content, %
A
T
G
C
NL_crAss001368, nt

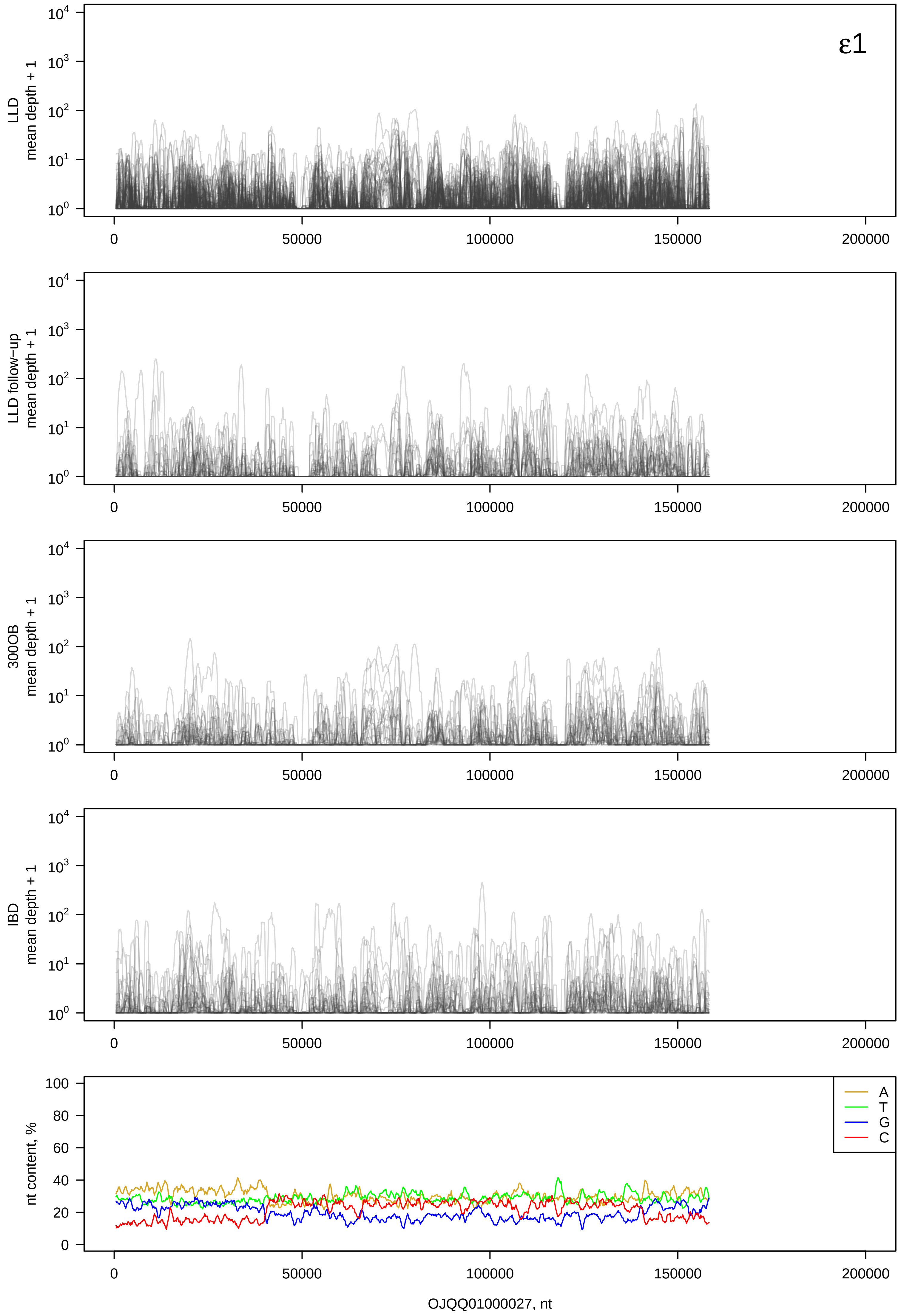

ε1
LLD mean depth + 1
LLD follow-up mean depth + 1
300OB mean depth + 1
IBD mean depth + 1
nt content, %
A
T
G
C
0
50000
100000
150000
200000
OJQQ01000027, nt

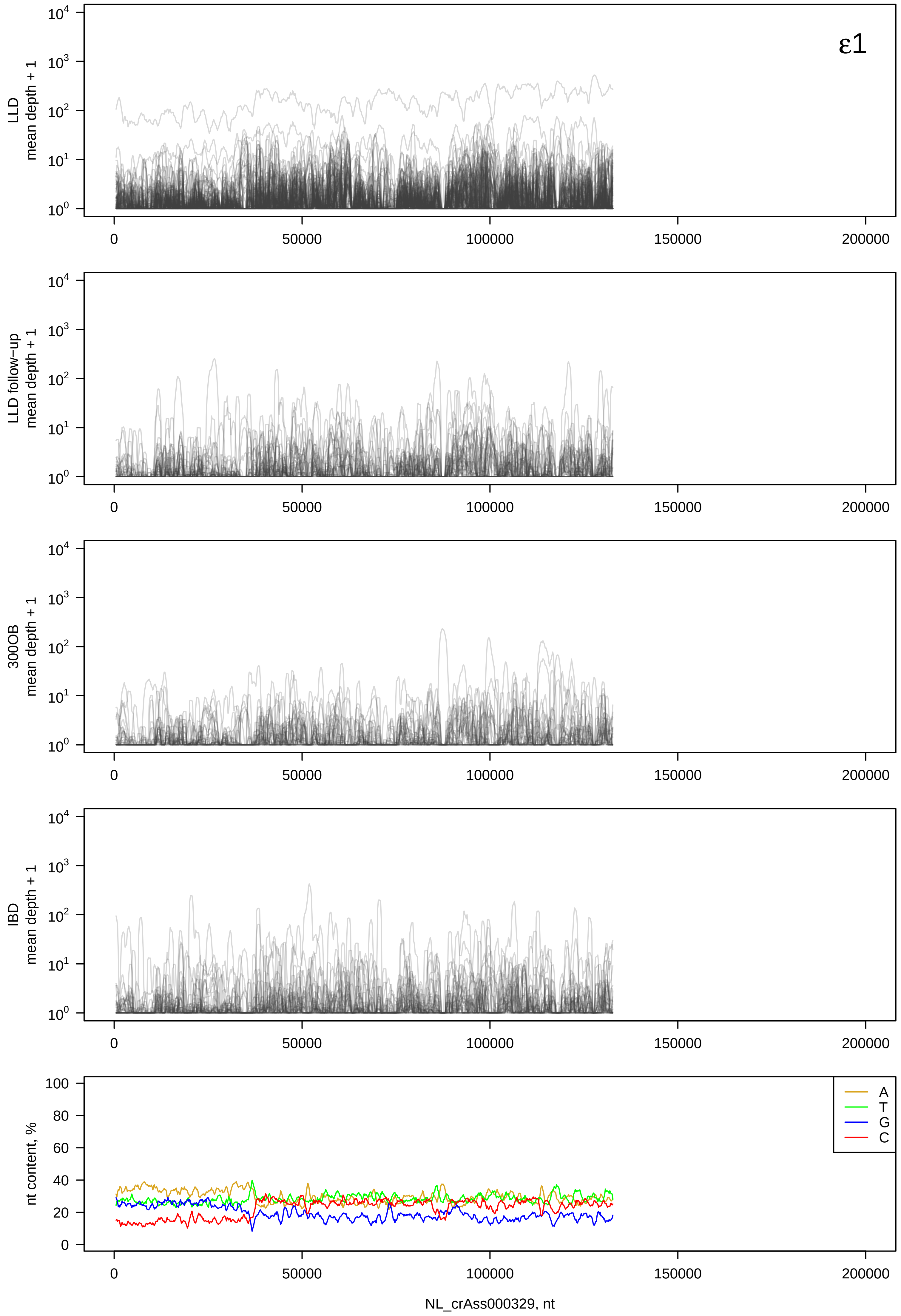
ε1
LLD mean depth + 1
LLD follow-up mean depth + 1
300OB mean depth + 1
IBD mean depth + 1
nt content, %
A
T
G
C
NL_crAss000329, nt
0
50000
100000
150000
200000

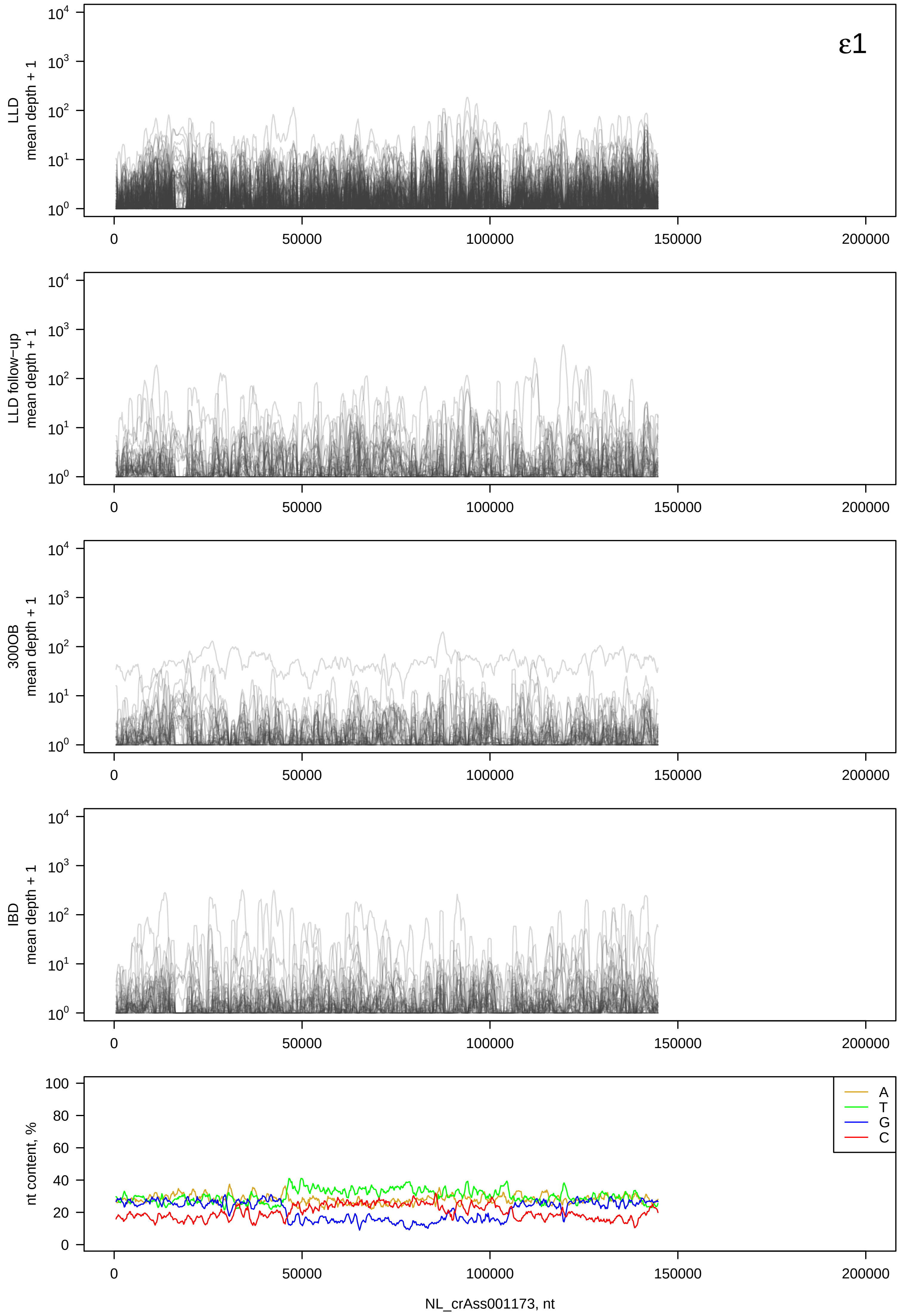
ε1
LLD
mean depth + 1
LLD follow−up
mean depth + 1
300OB
mean depth + 1
IBD
mean depth + 1
nt content, %
A
T
G
C
NL_crAss001173, nt

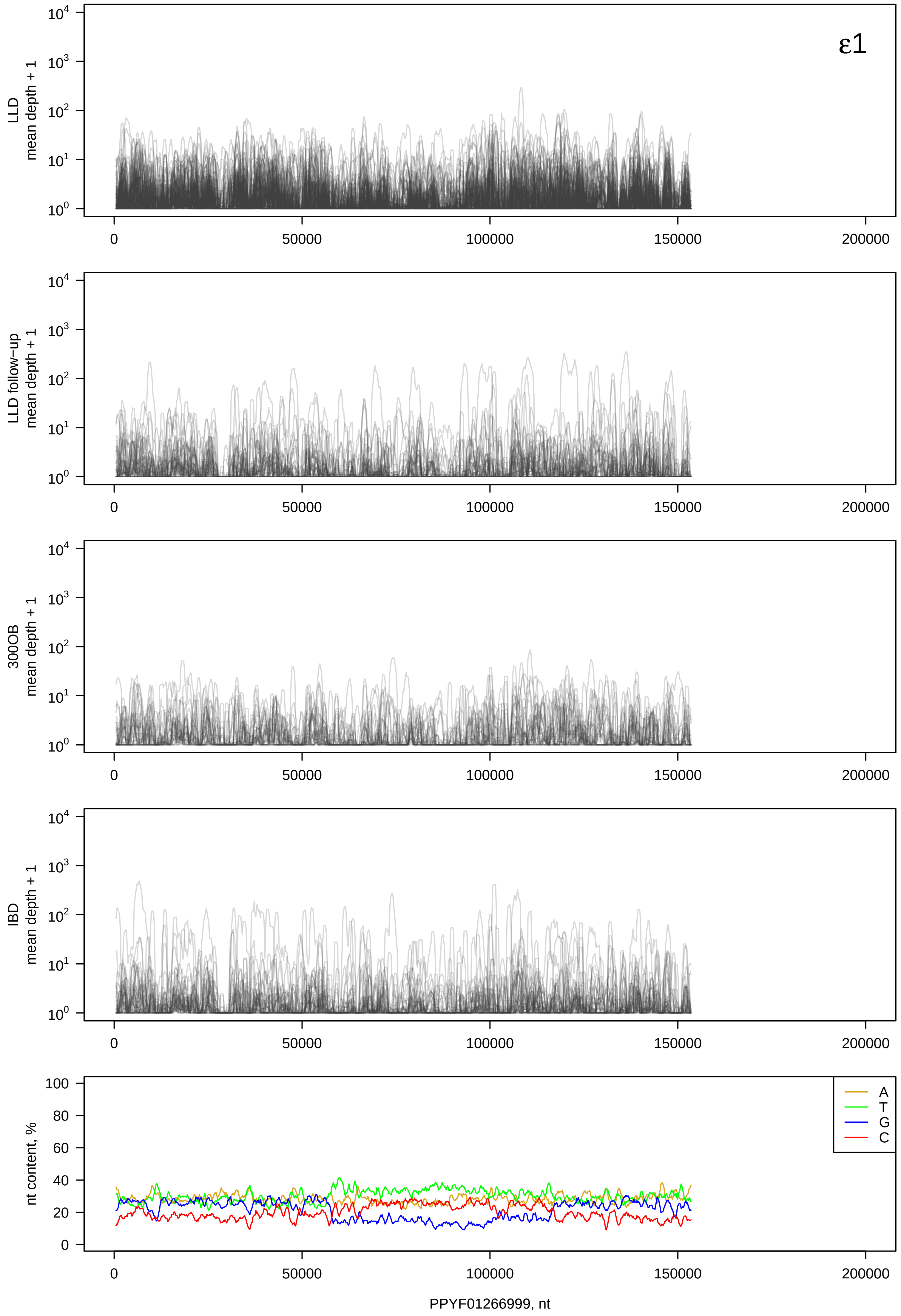

ε1
LLD mean depth + 1
LLD follow−up mean depth + 1
300OB mean depth + 1
IBD mean depth + 1
nt content, %
A
T
G
C
PPYF01266999, nt

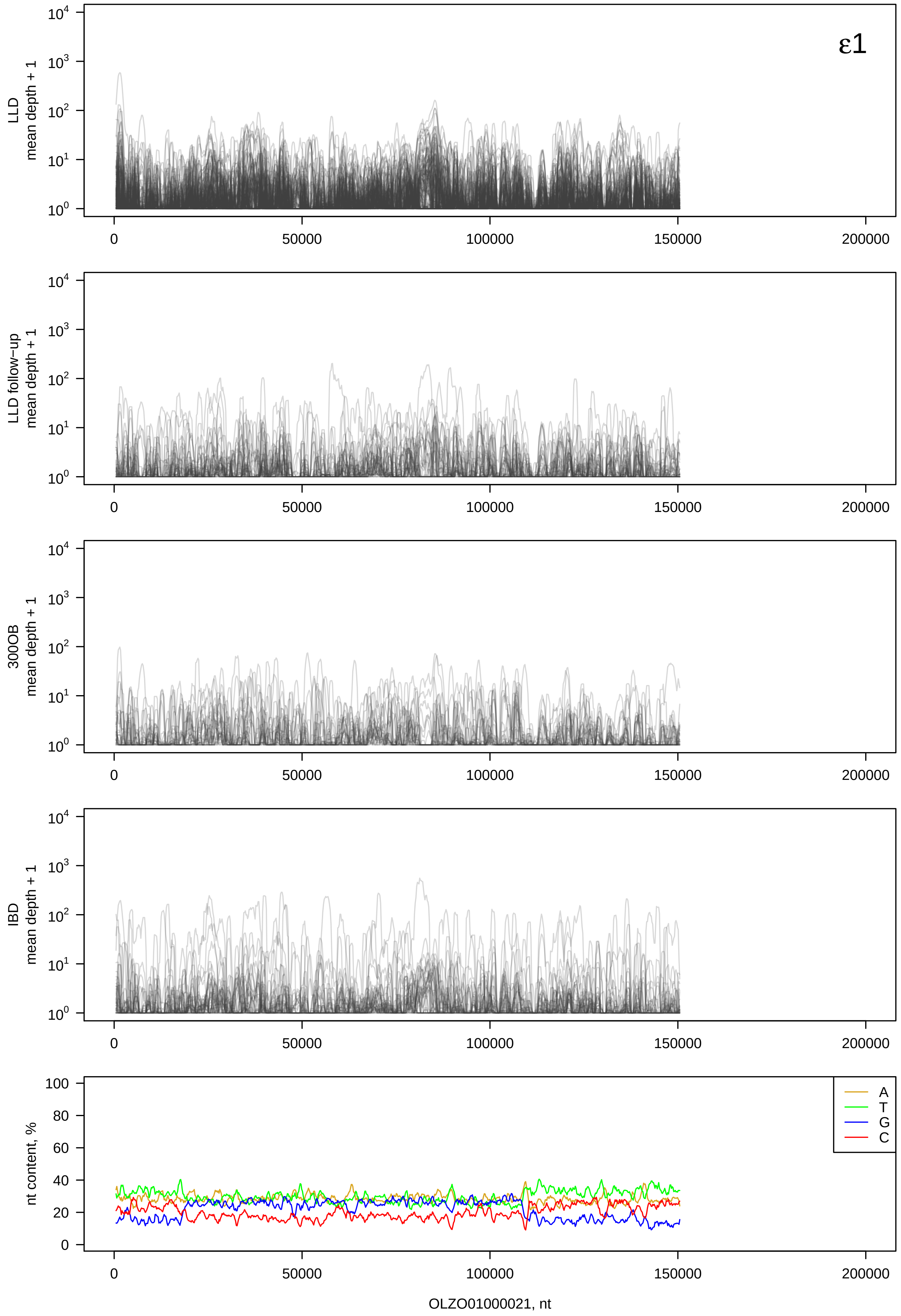
ε1
LLD mean depth + 1
LLD follow-up mean depth + 1
300OB mean depth + 1
IBD mean depth + 1
nt content, %
A
T
G
C
OLZO01000021, nt

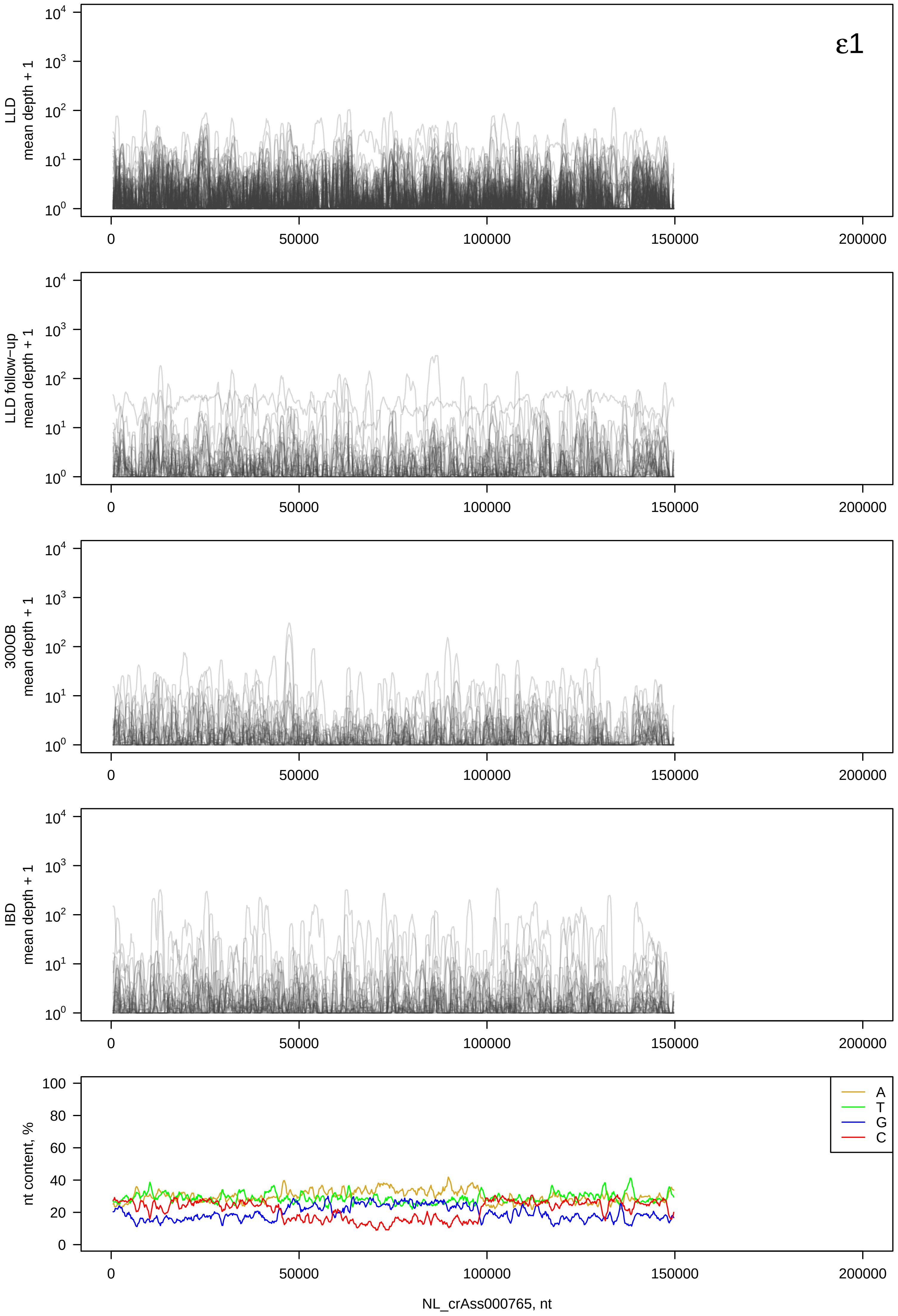

ε1
LLD mean depth + 1
LLD follow−up mean depth + 1
300OB mean depth + 1
IBD mean depth + 1
nt content, %
A
T
G
C
0
50000
100000
150000
200000
NL_crAss000765, nt

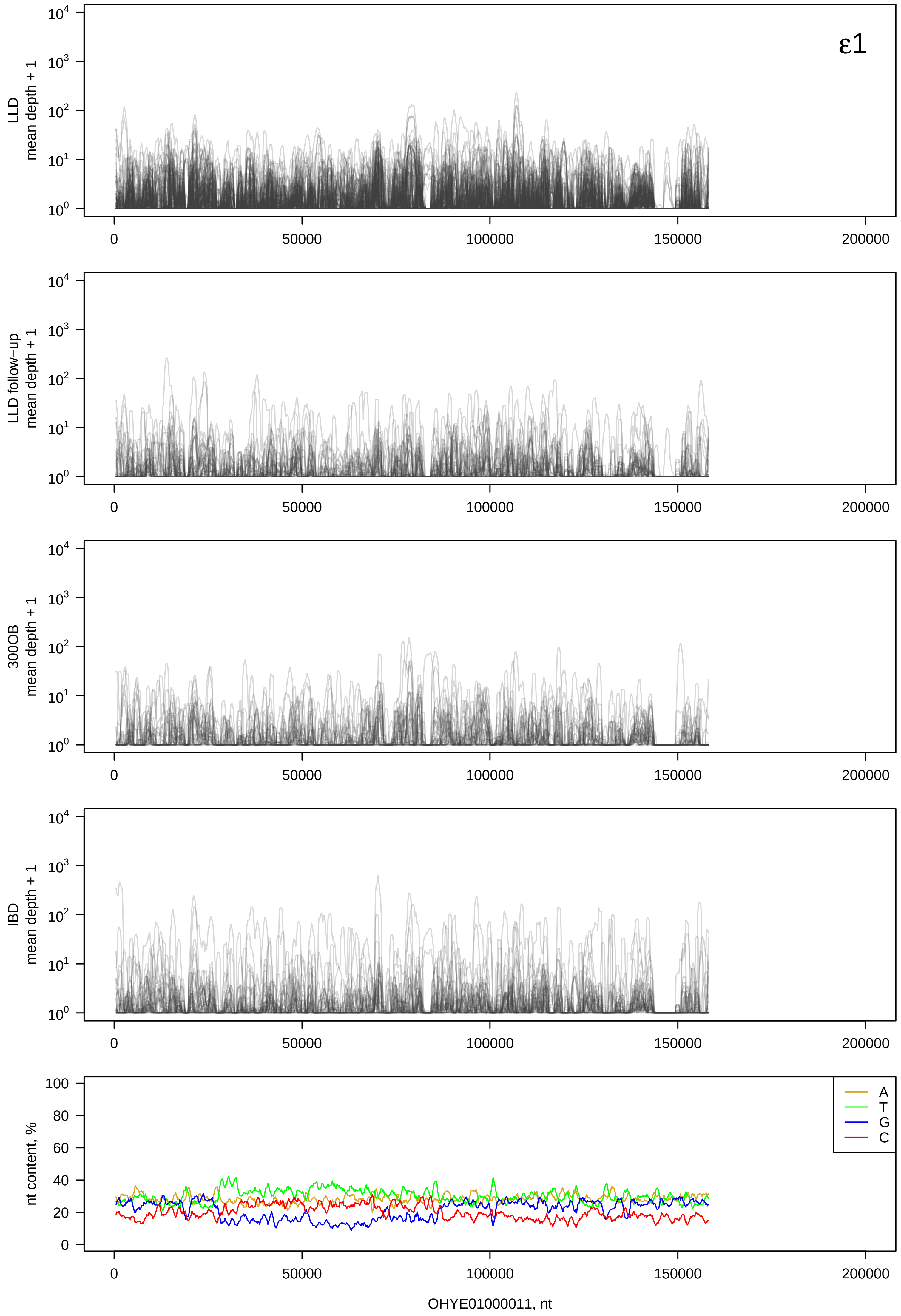
ε1
LLD mean depth + 1
LLD follow-up mean depth + 1
300OB mean depth + 1
IBD mean depth + 1
nt content, %
A
T
G
C
OHYE01000011, nt

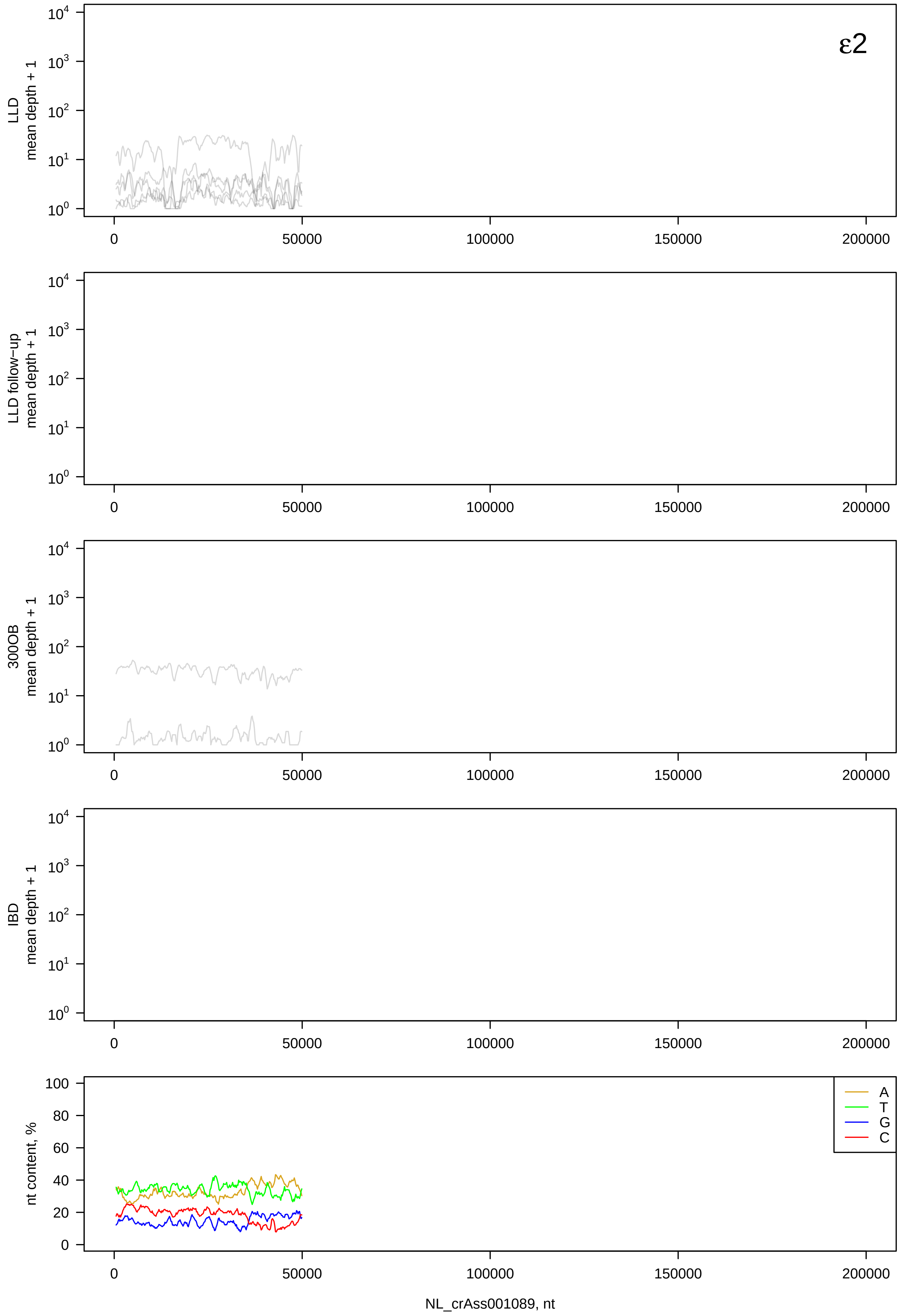

ε2
LLD mean depth + 1
LLD follow-up mean depth + 1
300OB mean depth + 1
IBD mean depth + 1
nt content, %
A
T
G
C
NL_crAss001089, nt

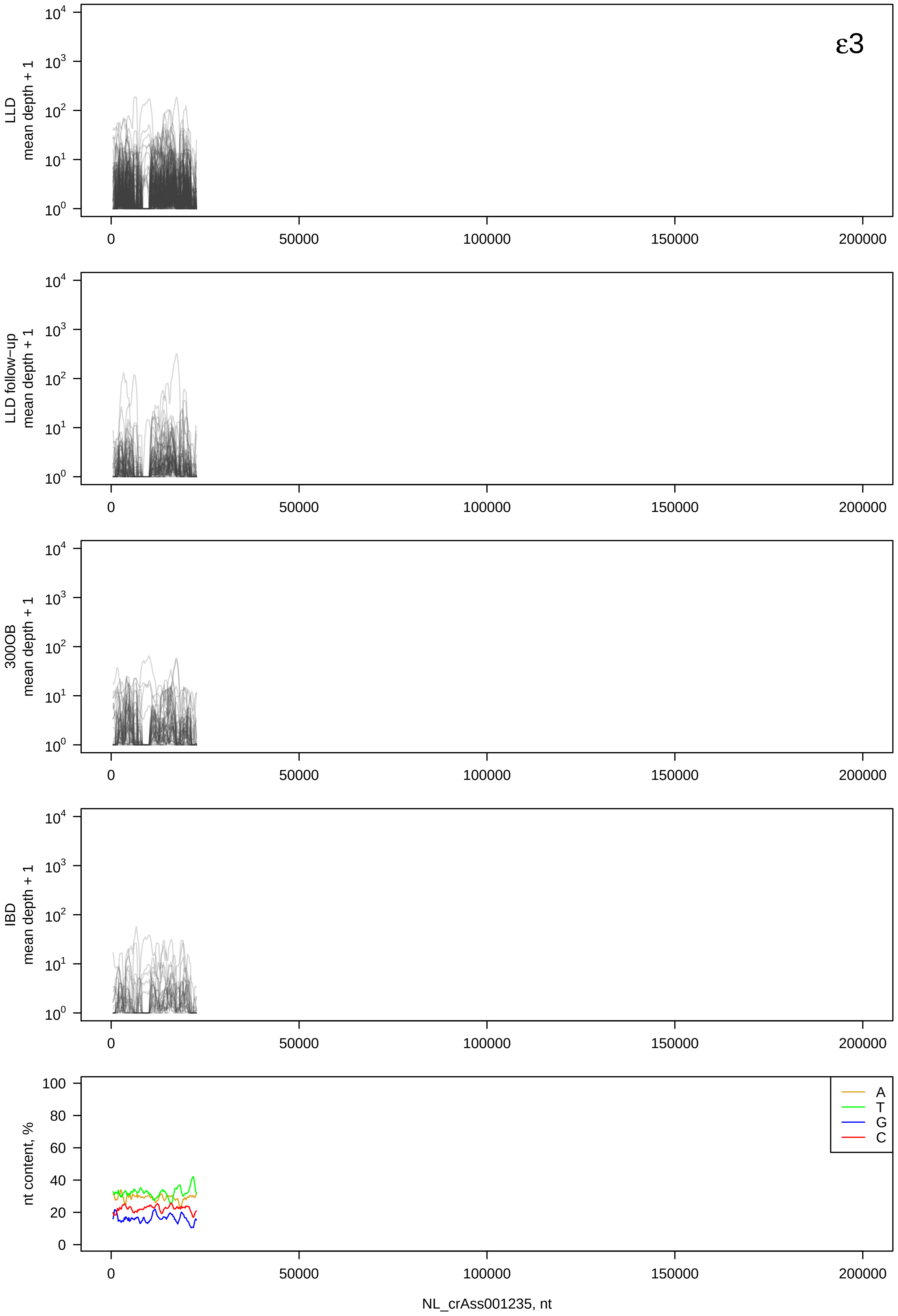

ε3
LLD
mean depth + 1
LLD follow−up
mean depth + 1
300OB
mean depth + 1
IBD
mean depth + 1
$10^0$
$10^1$
$10^2$
$10^3$
$10^4$
0
50000
100000
150000
200000
nt content, %
0
20
40
60
80
100
A
T
G
C
NL_crAss001235, nt

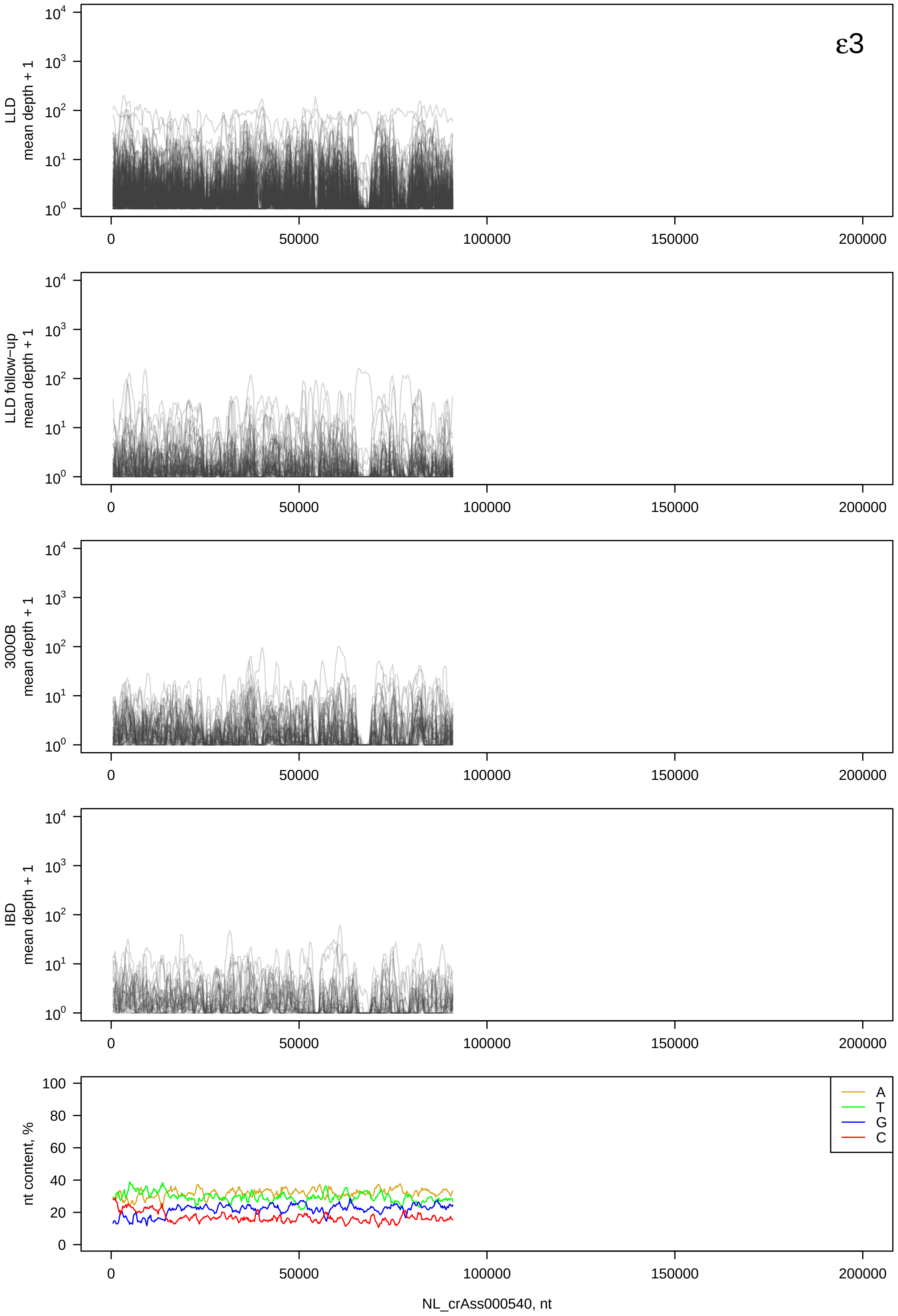

ε3
LLD mean depth + 1
LLD follow−up mean depth + 1
300OB mean depth + 1
IBD mean depth + 1
nt content, %
A
T
G
C
0
50000
100000
150000
200000
NL_crAss000540, nt

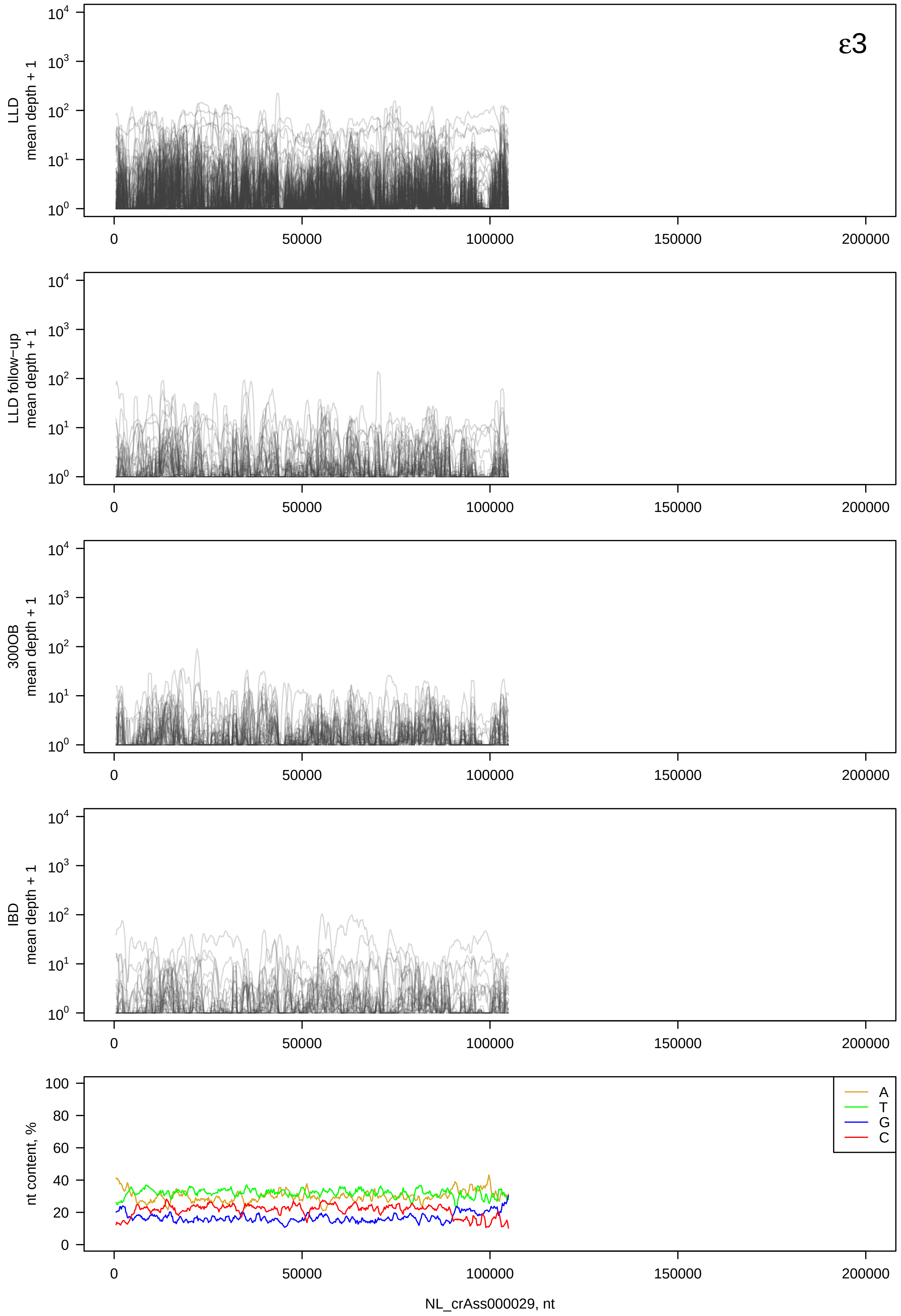

ε3
LLD
mean depth + 1
LLD follow−up
mean depth + 1
300OB
mean depth + 1
IBD
mean depth + 1
nt content, %
A
T
G
C
0
50000
100000
150000
200000
NL_crAss000029, nt

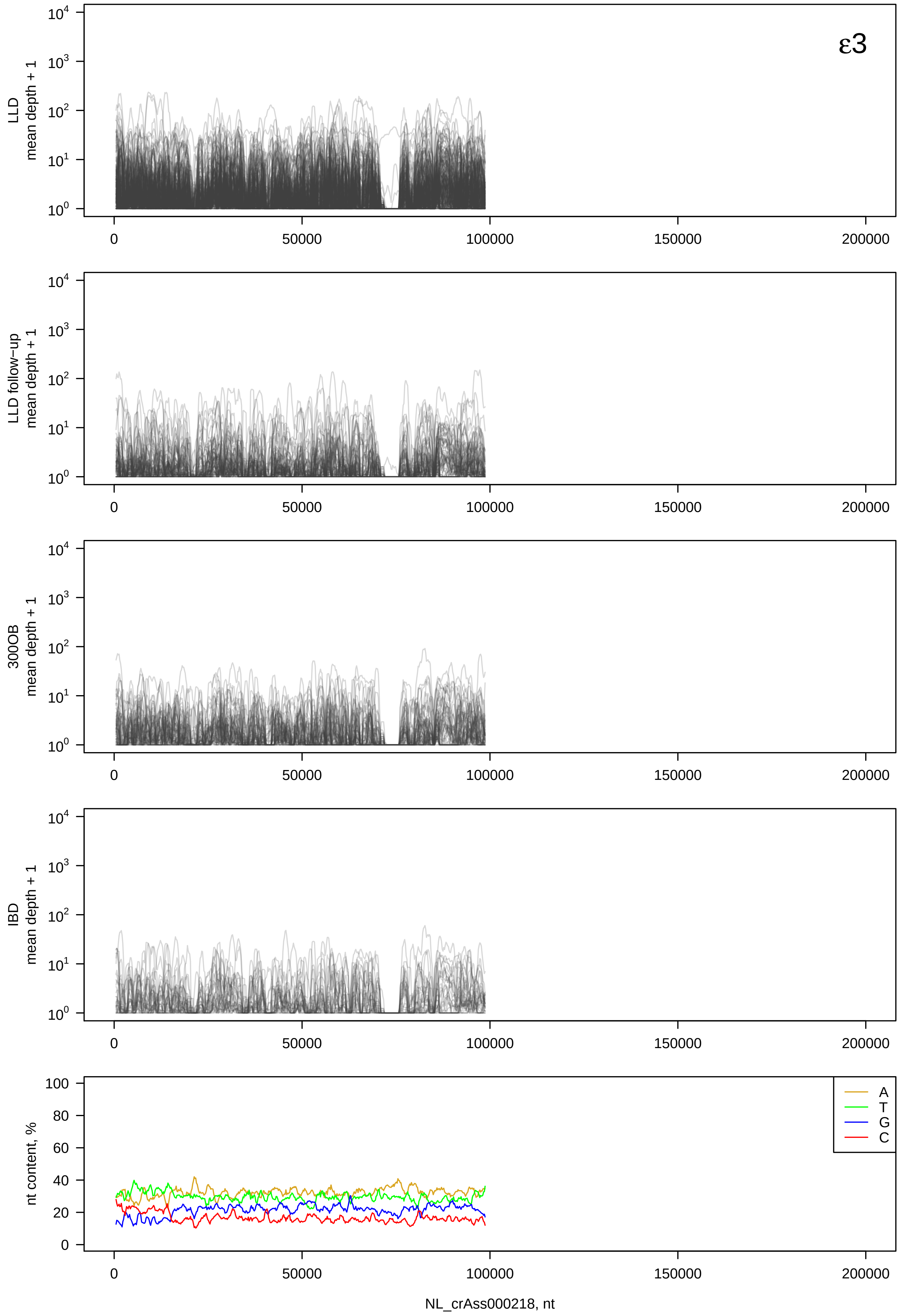

ε3
LLD
mean depth + 1
LLD follow−up
mean depth + 1
300OB
mean depth + 1
IBD
mean depth + 1
nt content, %
A
T
G
C
NL_crAss000218, nt

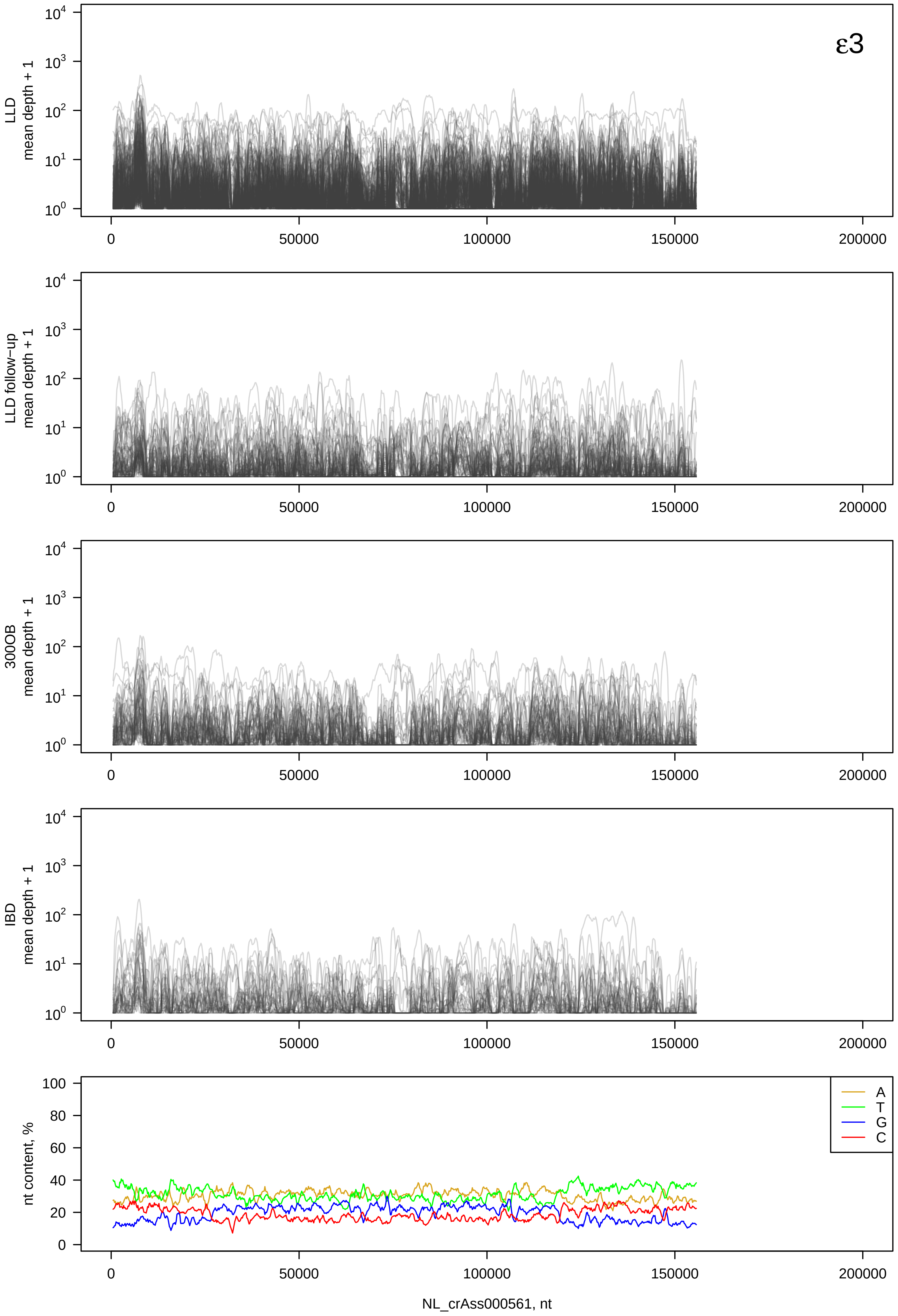

ε3
LLD
mean depth + 1
LLD follow−up
mean depth + 1
300OB
mean depth + 1
IBD
mean depth + 1
nt content, %
A
T
G
C
0
50000
100000
150000
200000
NL_crAss000561, nt

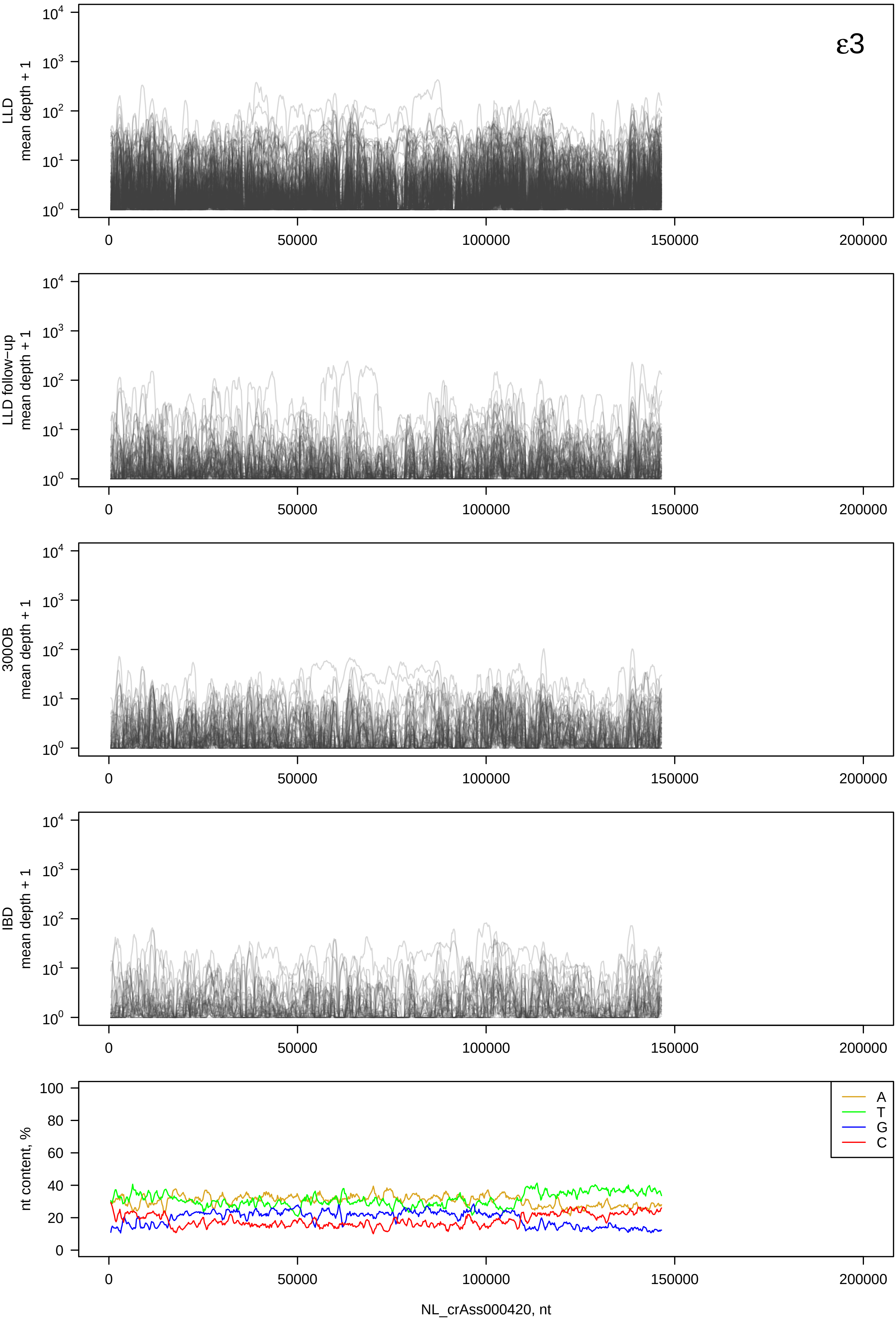

ε3
LLD mean depth + 1
LLD follow−up mean depth + 1
300OB mean depth + 1
IBD mean depth + 1
nt content, %
A
T
G
C
0
50000
100000
150000
200000
NL_crAss000420, nt

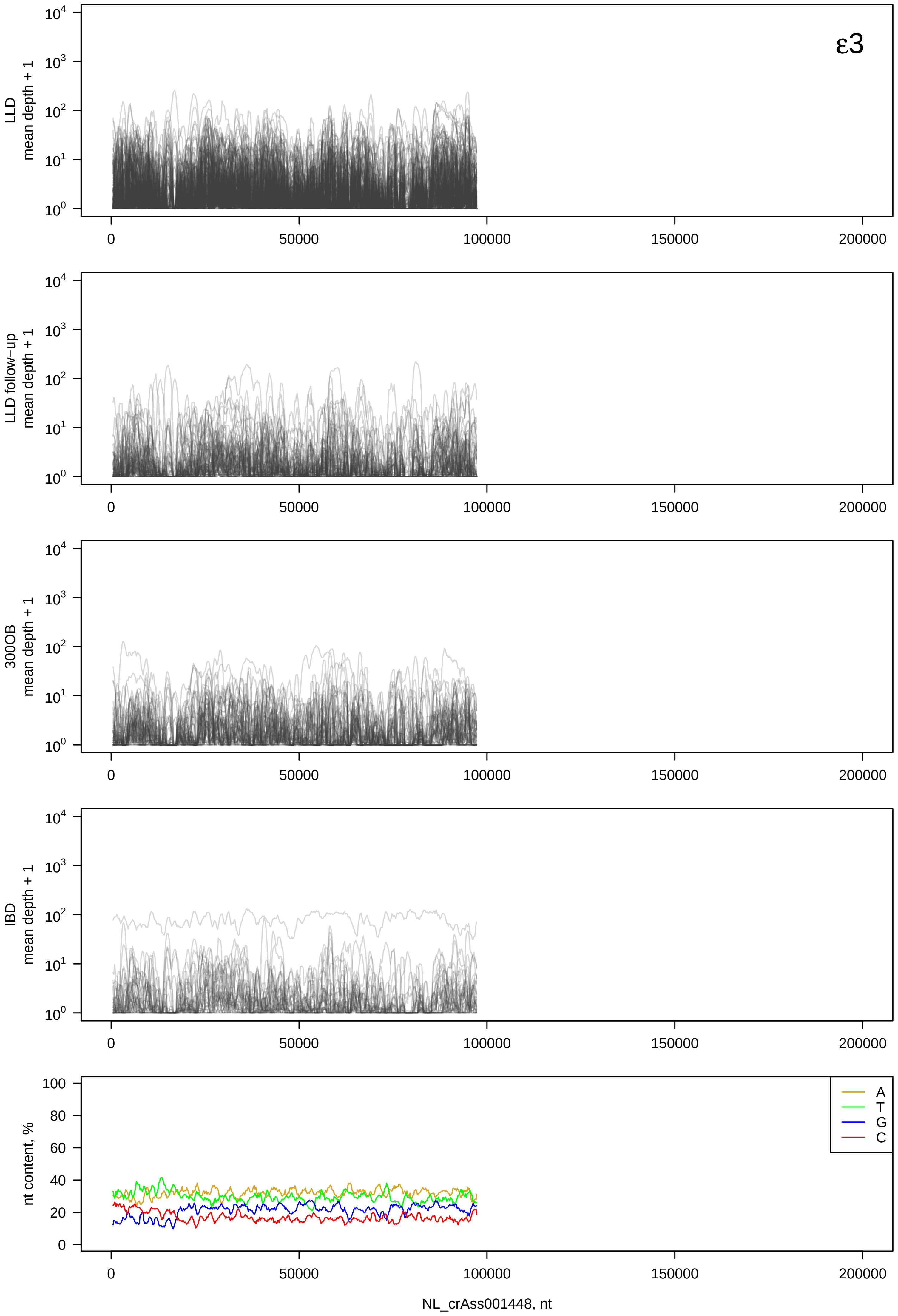

ε3
LLD mean depth + 1
LLD follow−up mean depth + 1
300OB mean depth + 1
IBD mean depth + 1
nt content, %
A
T
G
C
NL_crAss001448, nt

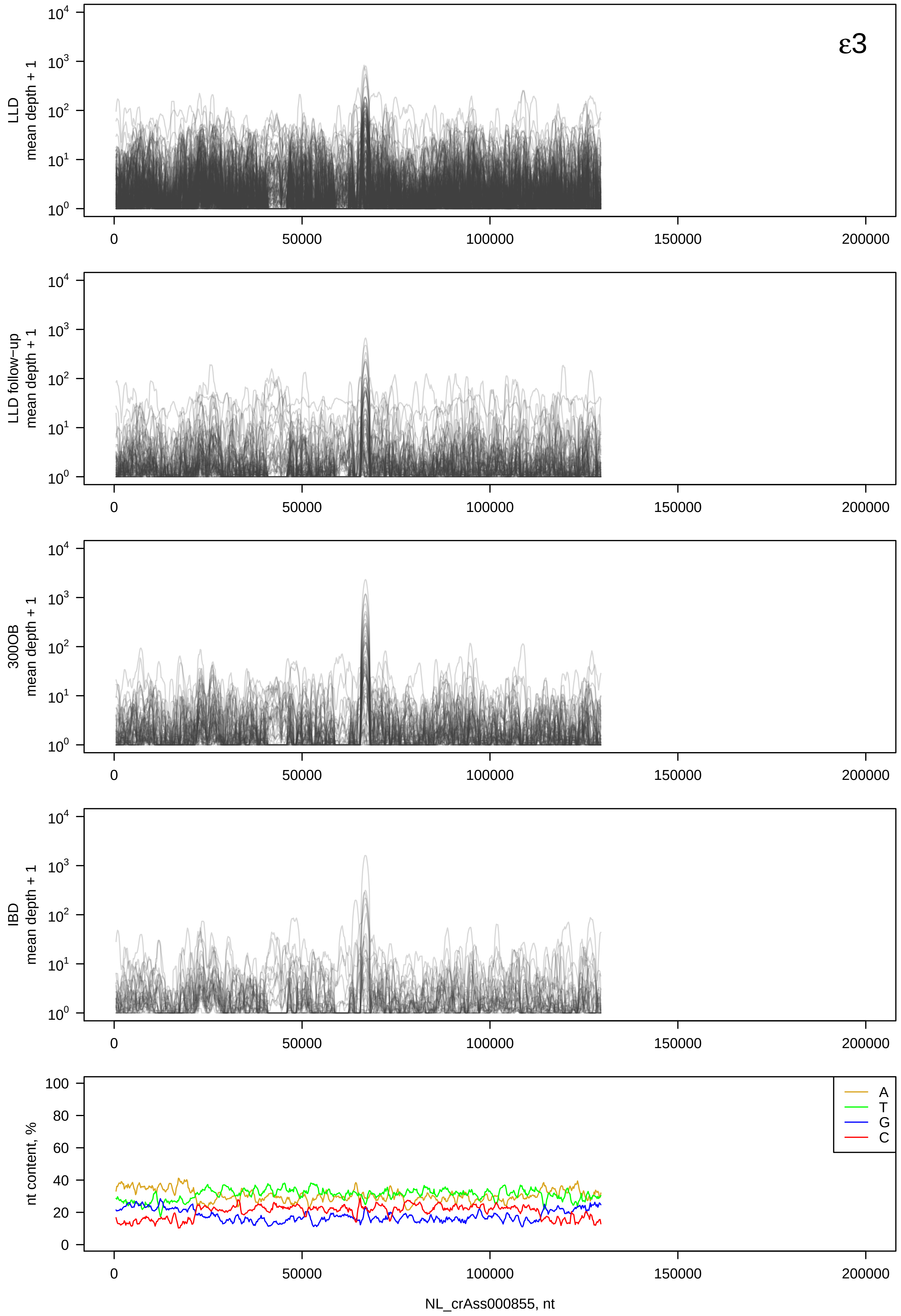

ε3
LLD mean depth + 1
LLD follow−up mean depth + 1
300OB mean depth + 1
IBD mean depth + 1
nt content, %
A
T
G
C
NL_crAss000855, nt

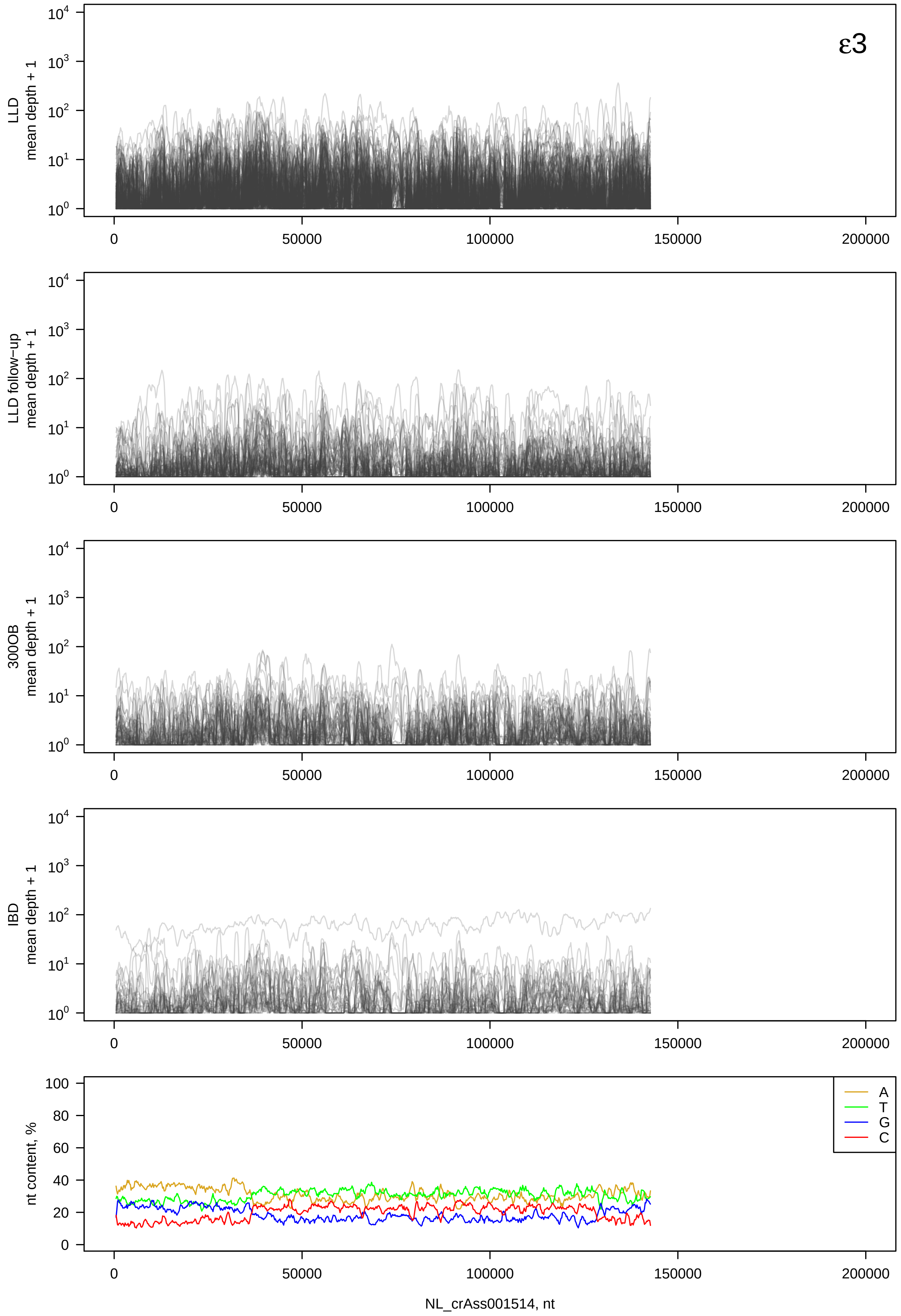
ε3
LLD
mean depth + 1
LLD follow−up
mean depth + 1
300OB
mean depth + 1
IBD
mean depth + 1
nt content, %
A
T
G
C
0
50000
100000
150000
200000
NL_crAss001514, nt

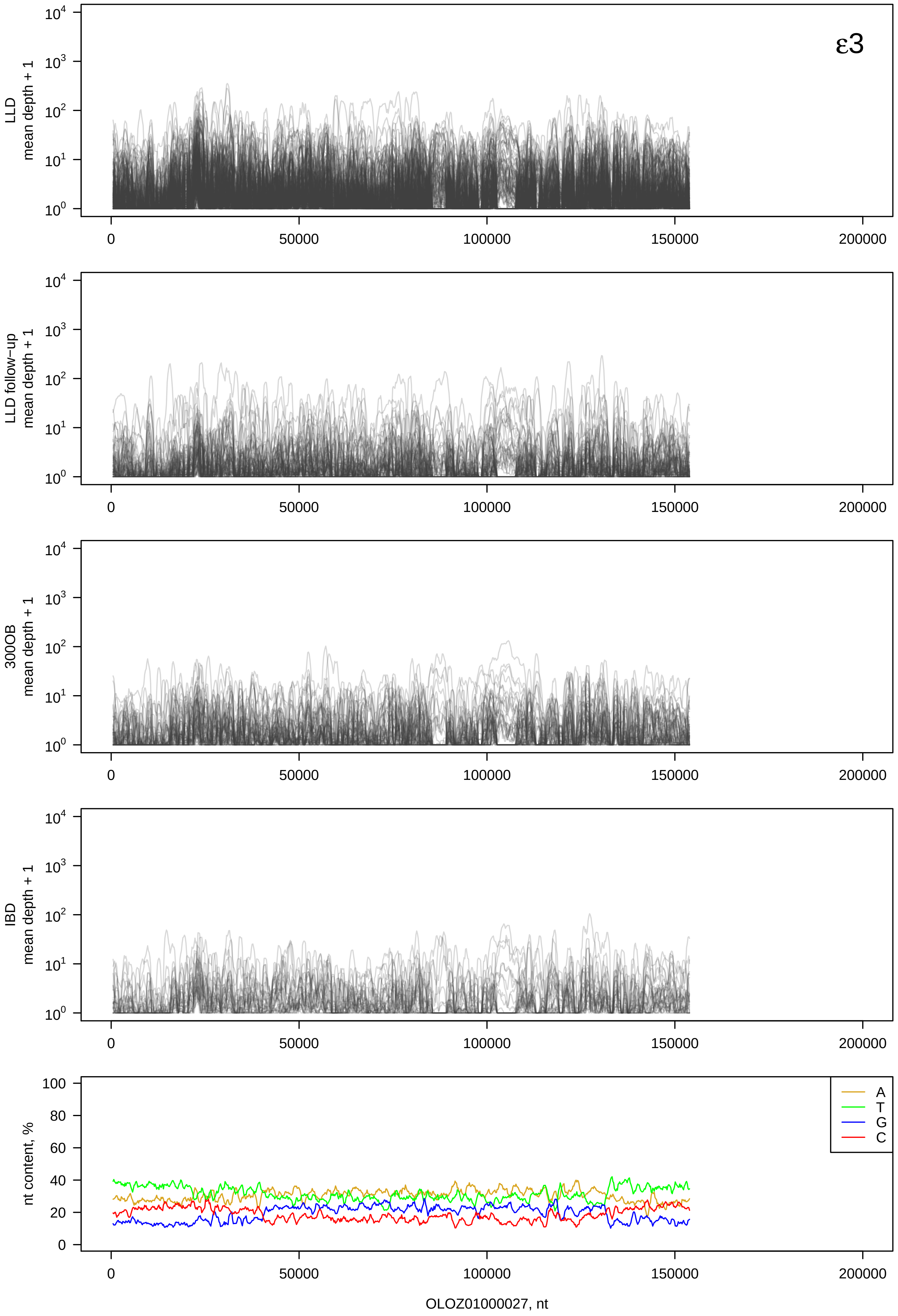

ε3
LLD
mean depth + 1
LLD follow−up
mean depth + 1
300OB
mean depth + 1
IBD
mean depth + 1
$10^0$
$10^1$
$10^2$
$10^3$
$10^4$
0
50000
100000
150000
200000
nt content, %
0
20
40
60
80
100
A
T
G
C
OLOZ01000027, nt

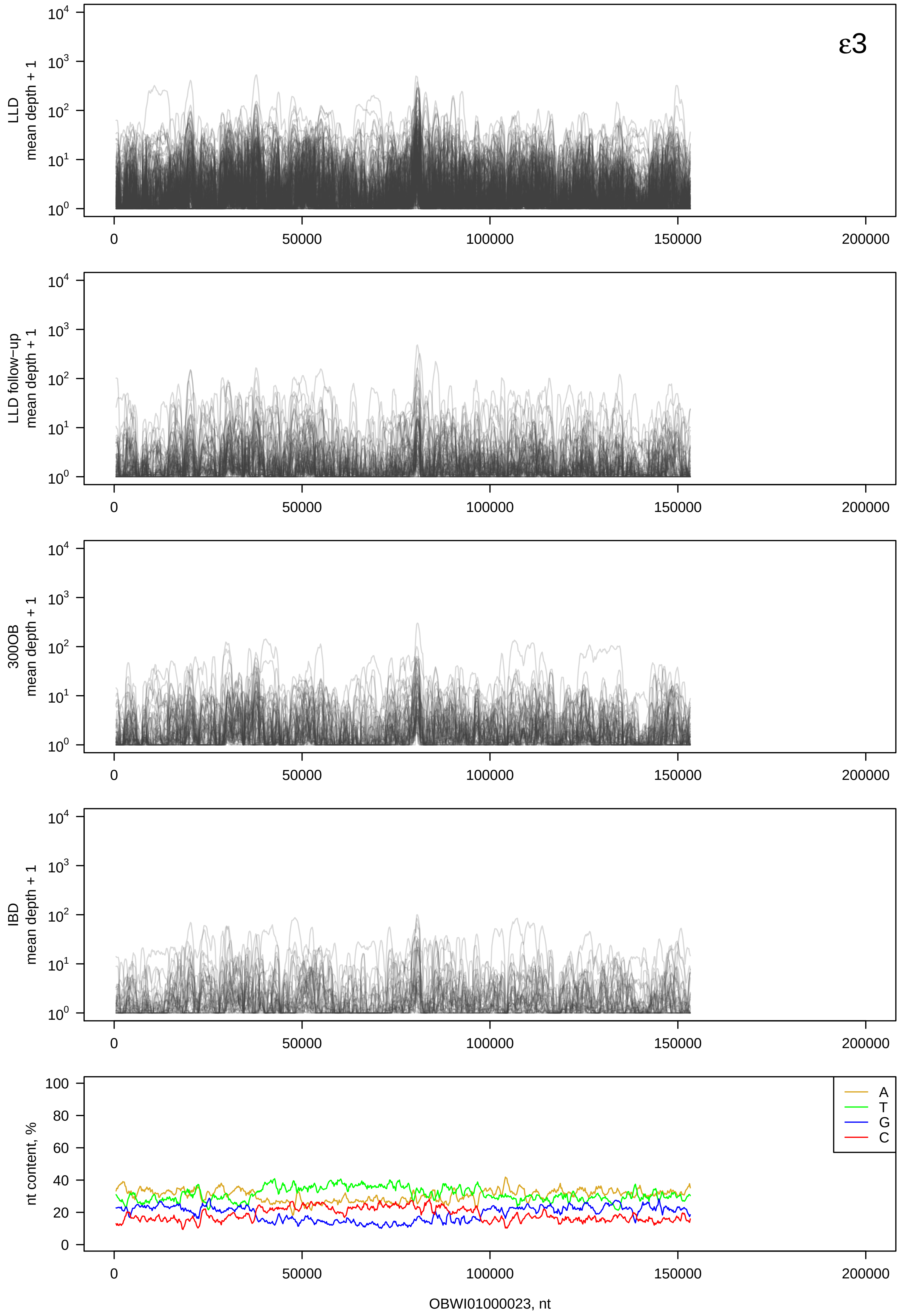
ε3
LLD
mean depth + 1
LLD follow-up
mean depth + 1
300OB
mean depth + 1
IBD
mean depth + 1
nt content, %
A
T
G
C
0
50000
100000
150000
200000
OBWI01000023, nt

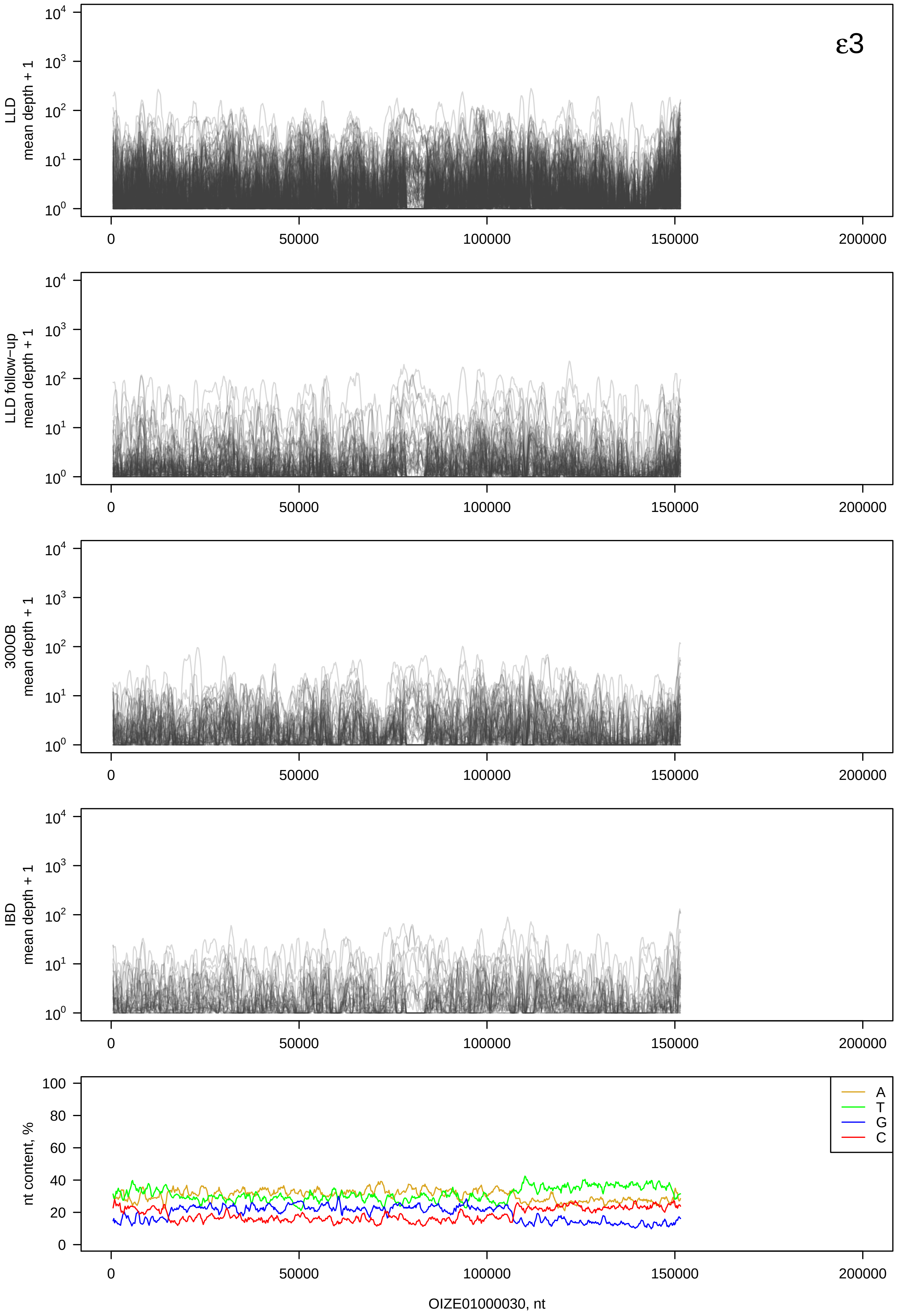

ε3
LLD mean depth + 1
LLD follow−up mean depth + 1
300OB mean depth + 1
IBD mean depth + 1
nt content, %
A
T
G
C
OIZE01000030, nt

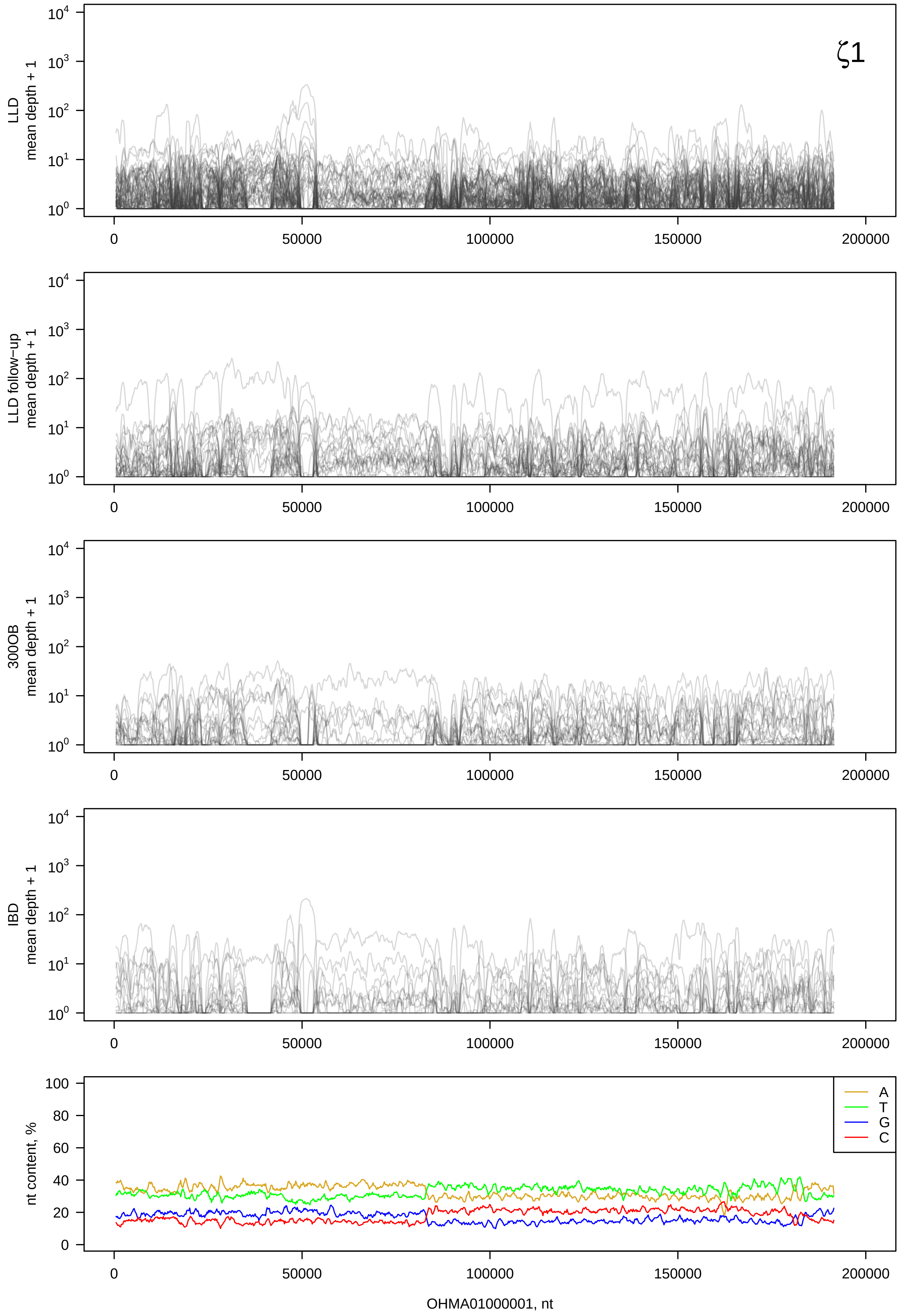
ζ1
LLD mean depth + 1
LLD follow−up mean depth + 1
300OB mean depth + 1
IBD mean depth + 1
nt content, %
A
T
G
C
OHMA01000001, nt

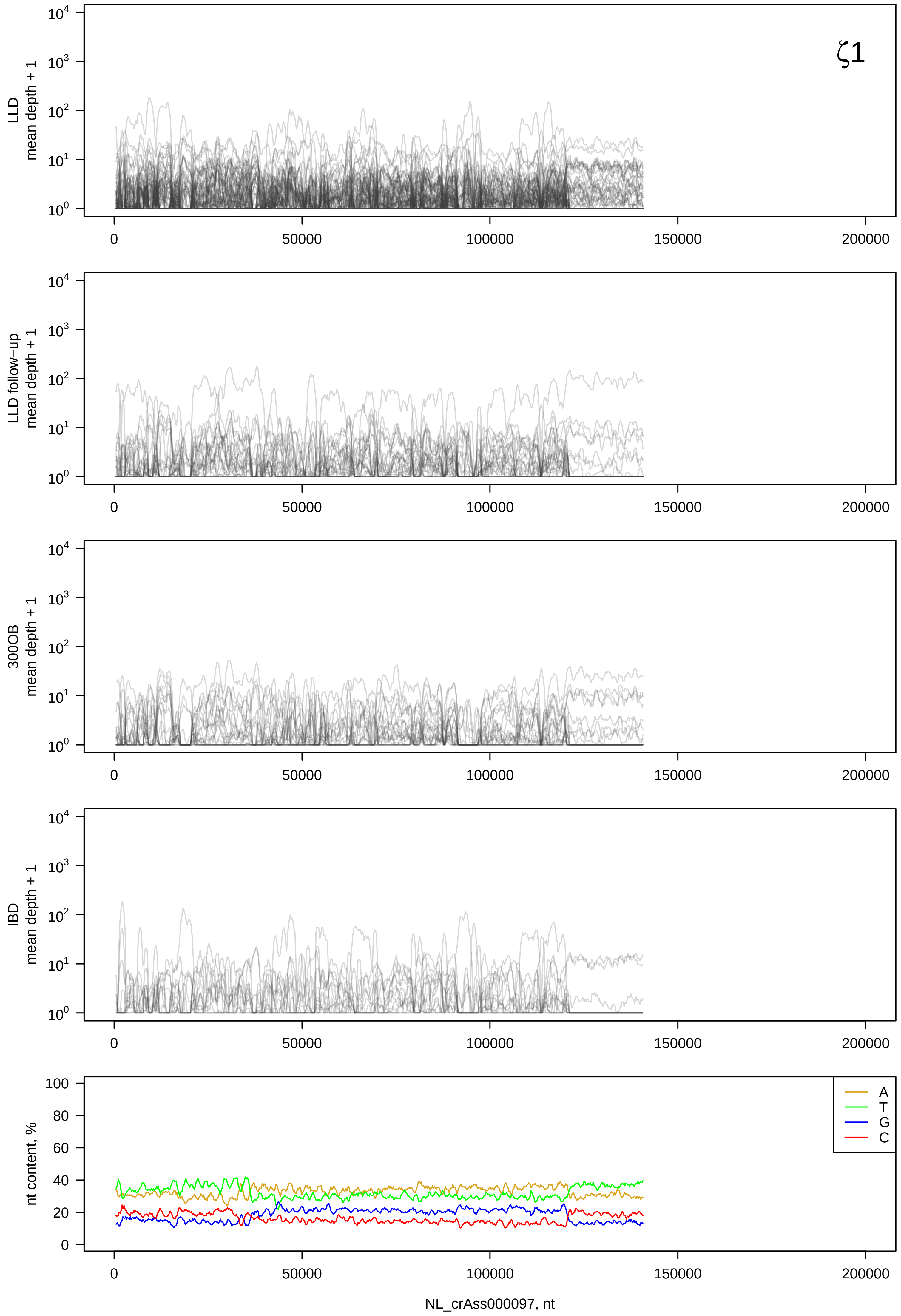

ζ1
LLD mean depth + 1
LLD follow-up mean depth + 1
300OB mean depth + 1
IBD mean depth + 1
nt content, %
A
T
G
C
0
50000
100000
150000
200000
NL_crAss000097, nt

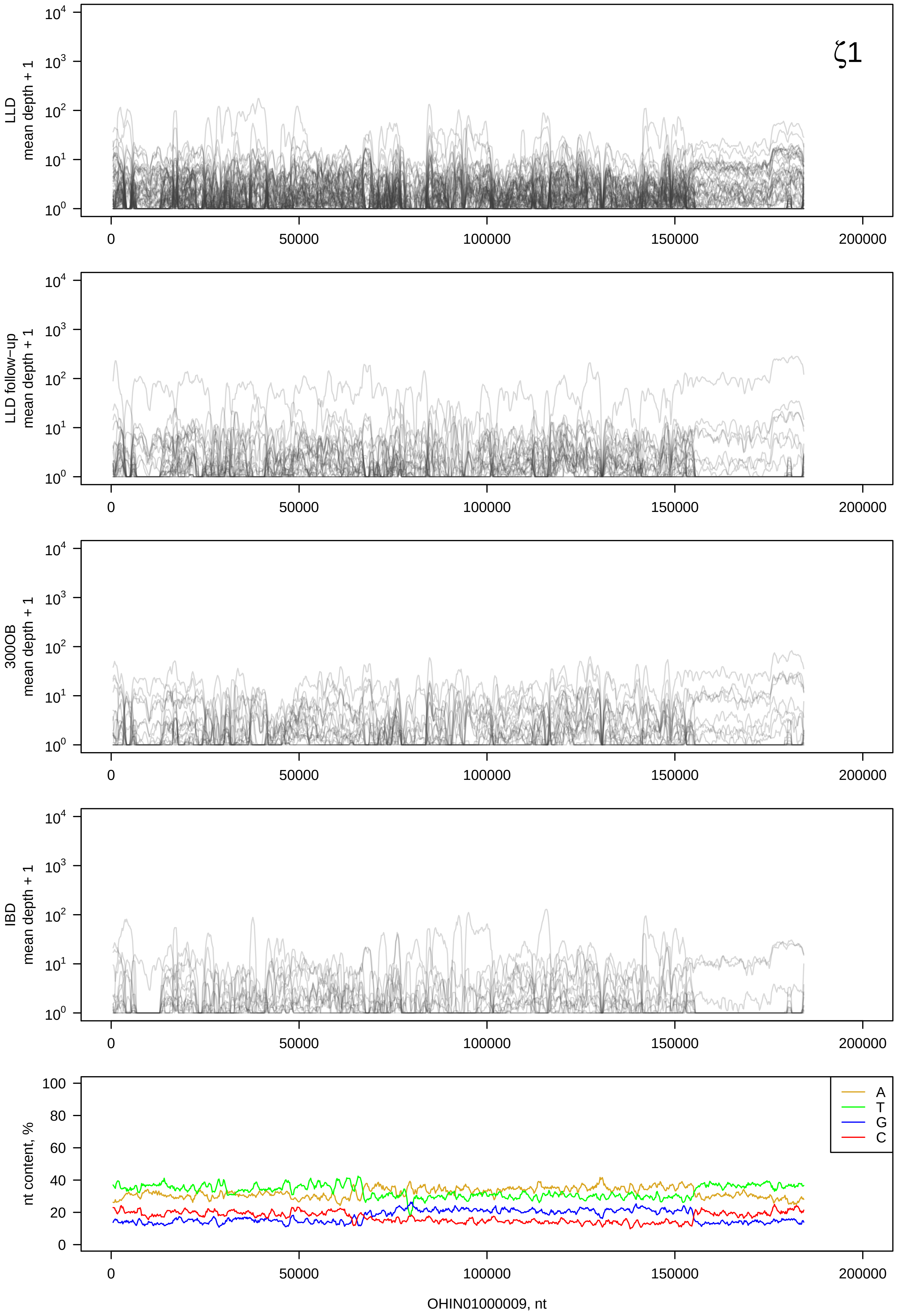

ζ1
LLD mean depth + 1
LLD follow−up mean depth + 1
300OB mean depth + 1
IBD mean depth + 1
nt content, %
A
T
G
C
OHIN01000009, nt

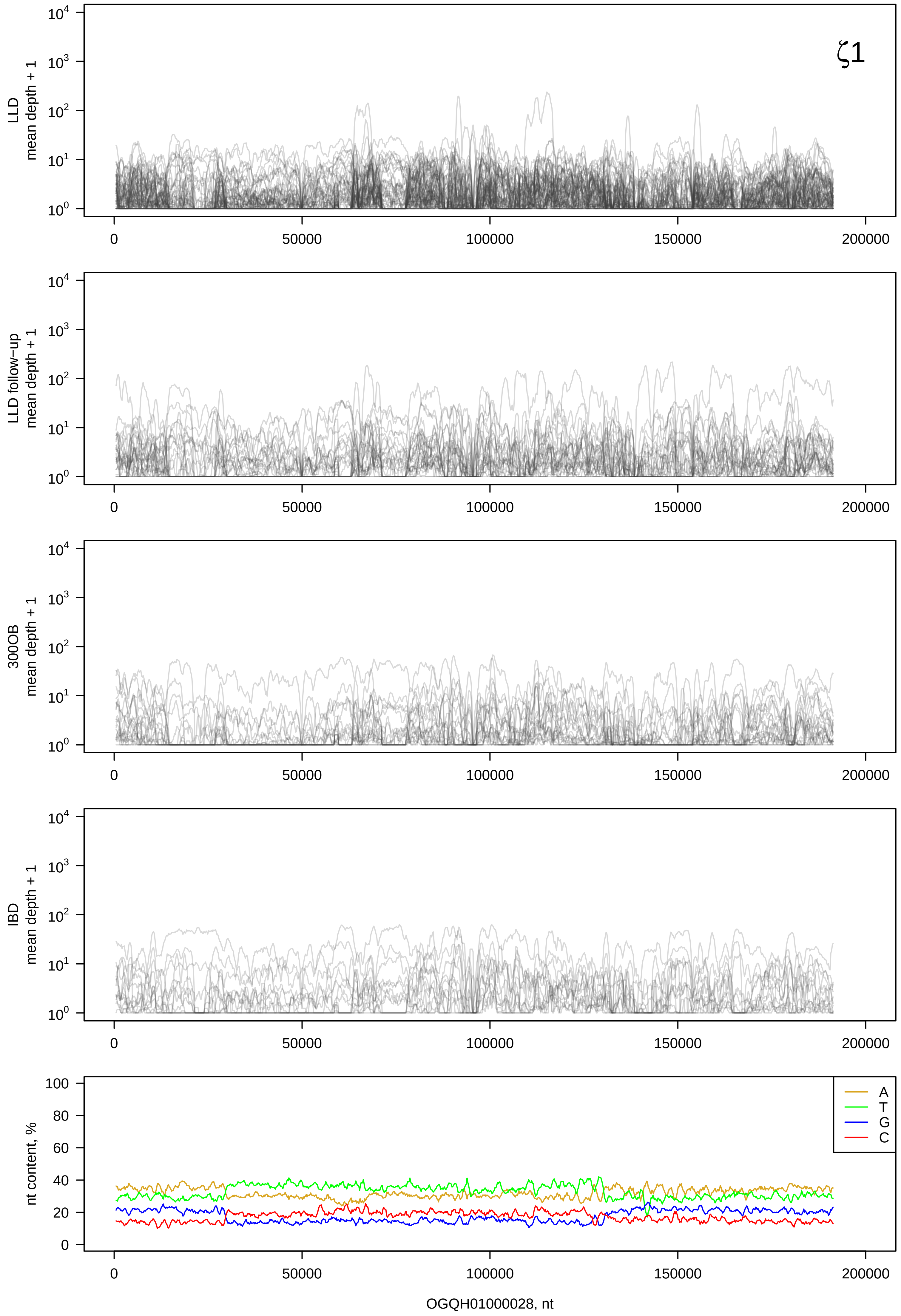

ζ1
LLD mean depth + 1
LLD follow−up mean depth + 1
300OB mean depth + 1
IBD mean depth + 1
nt content, %
A
T
G
C
OGQH01000028, nt

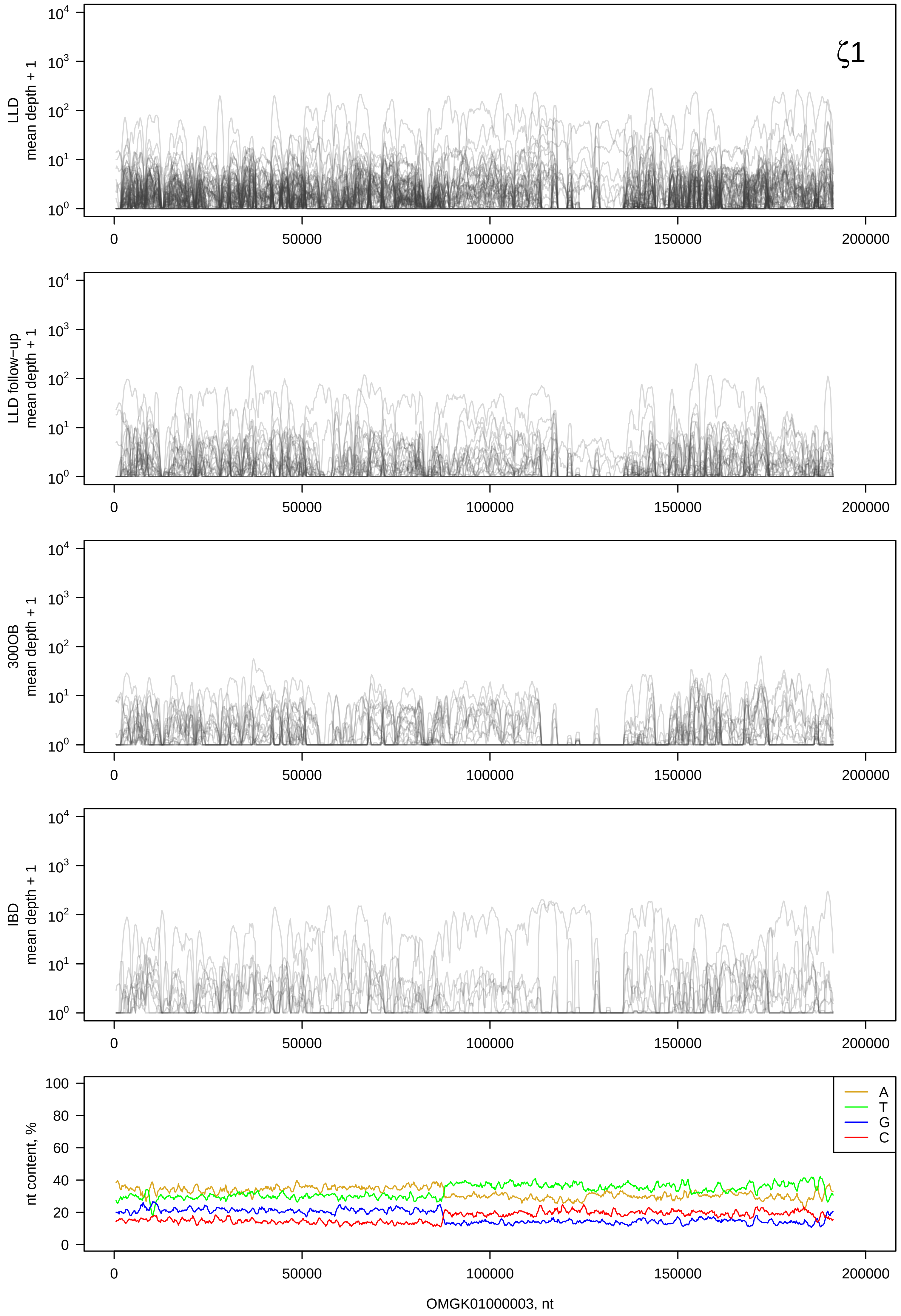
ζ1
LLD mean depth + 1
LLD follow-up mean depth + 1
300OB mean depth + 1
IBD mean depth + 1
nt content, %
A
T
G
C
OMGK01000003, nt

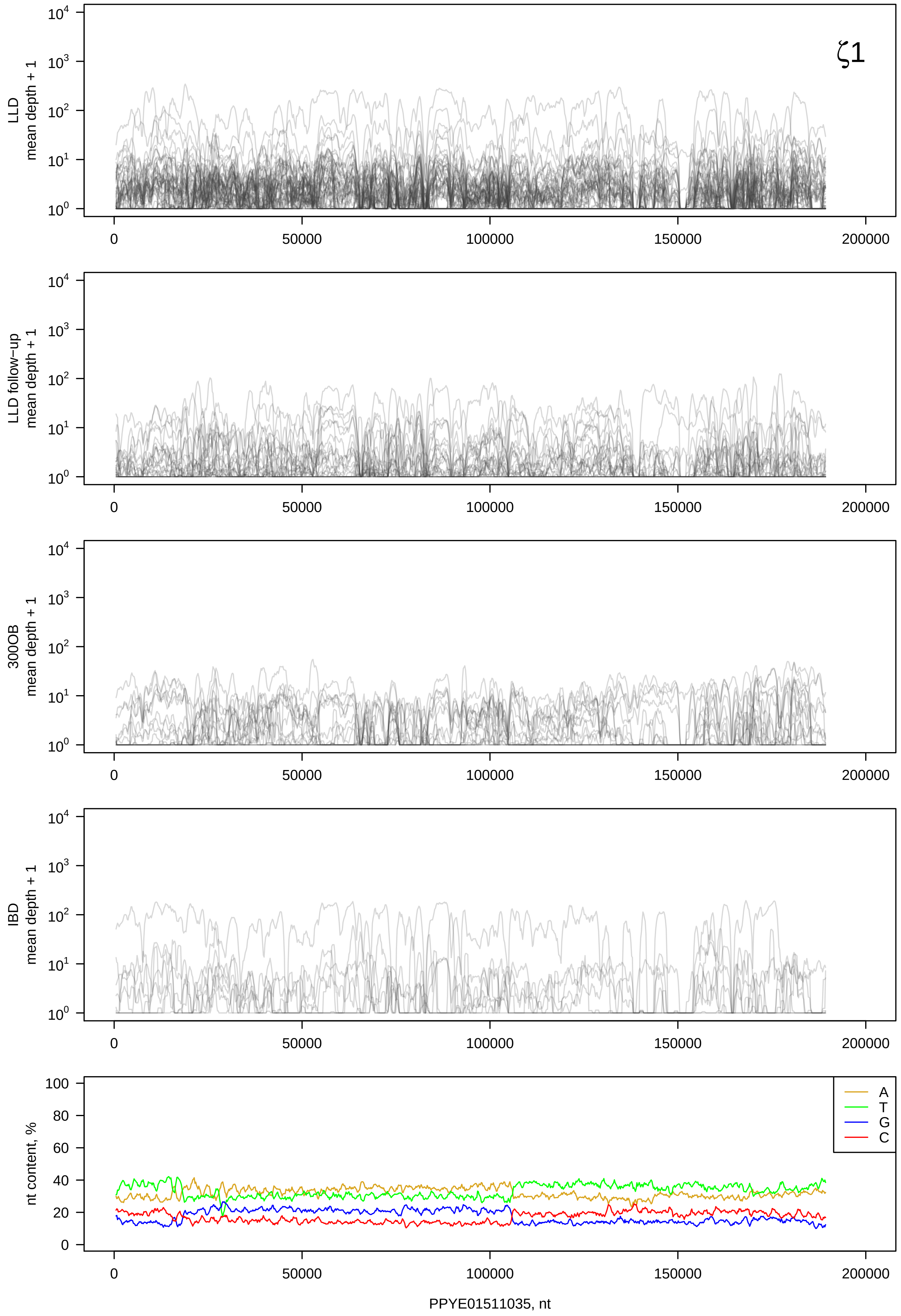

ζ1
LLD mean depth + 1
LLD follow−up mean depth + 1
300OB mean depth + 1
IBD mean depth + 1
nt content, %
A
T
G
C
0
50000
100000
150000
200000
PPYE01511035, nt

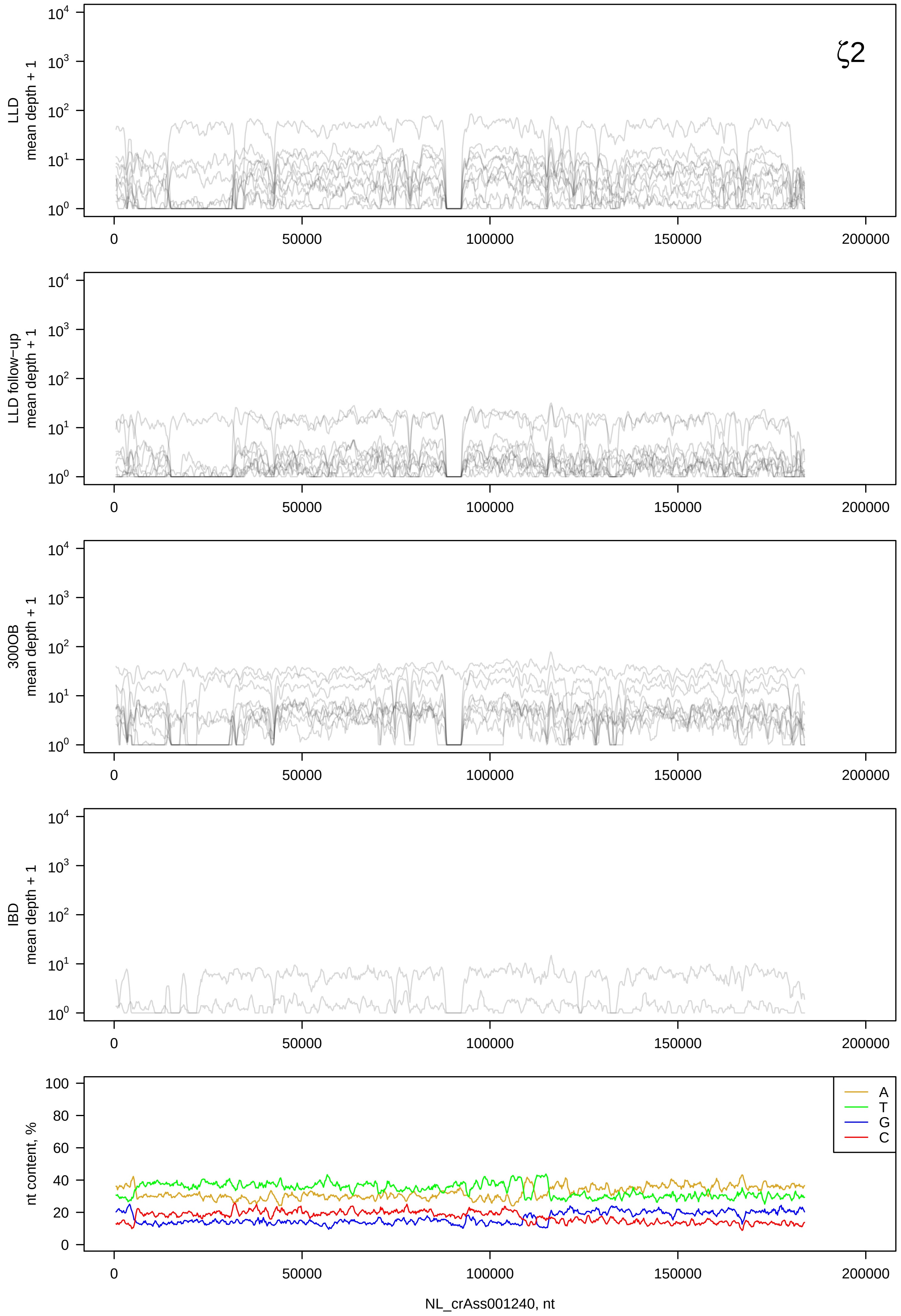

ζ2
LLD
mean depth + 1
LLD follow−up
mean depth + 1
300OB
mean depth + 1
IBD
mean depth + 1
nt content, %
A
T
G
C
NL_crAss001240, nt

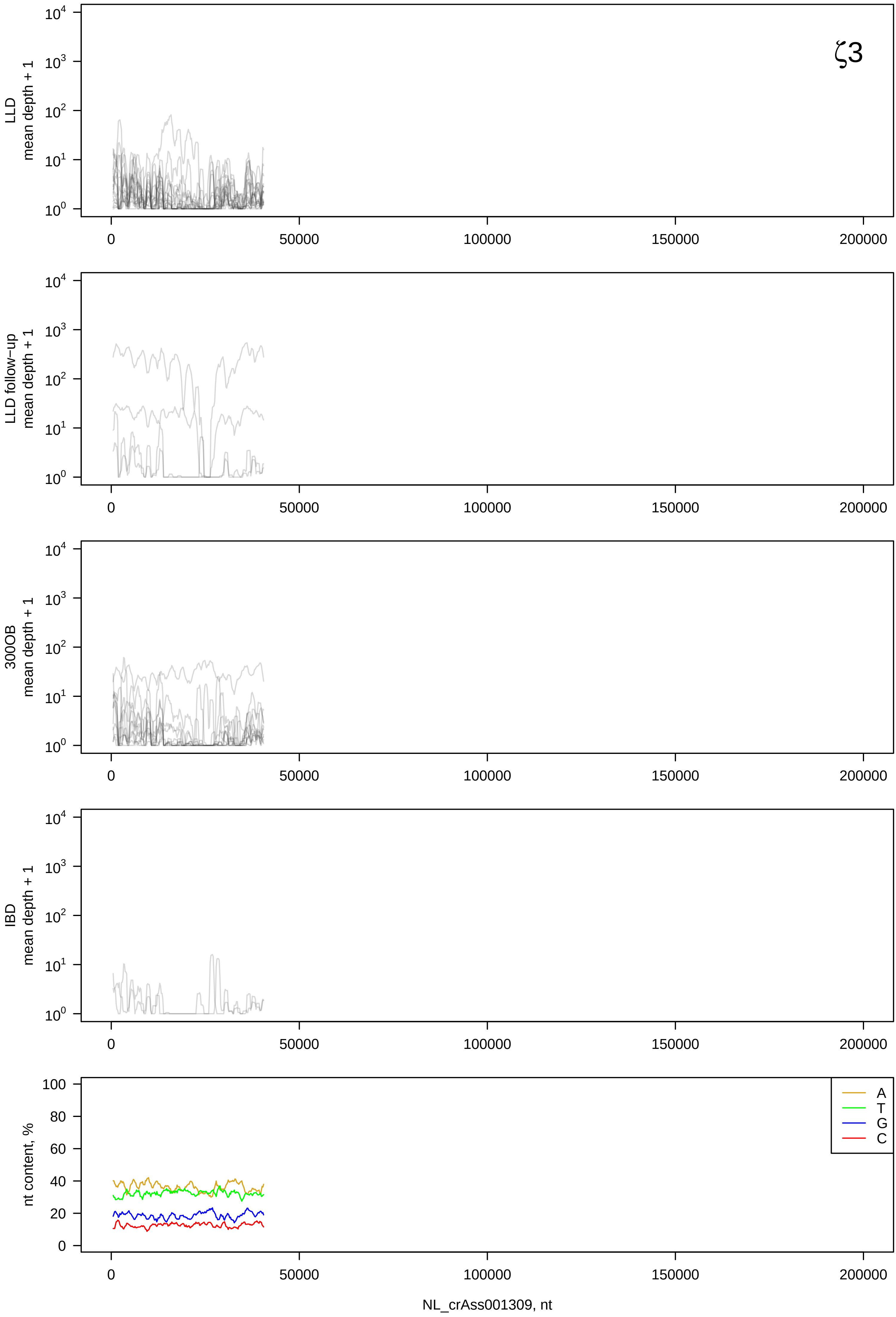

ζ3
LLD
mean depth + 1
LLD follow−up
mean depth + 1
300OB
mean depth + 1
IBD
mean depth + 1
nt content, %
A
T
G
C
0
50000
100000
150000
200000
NL_crAss001309, nt

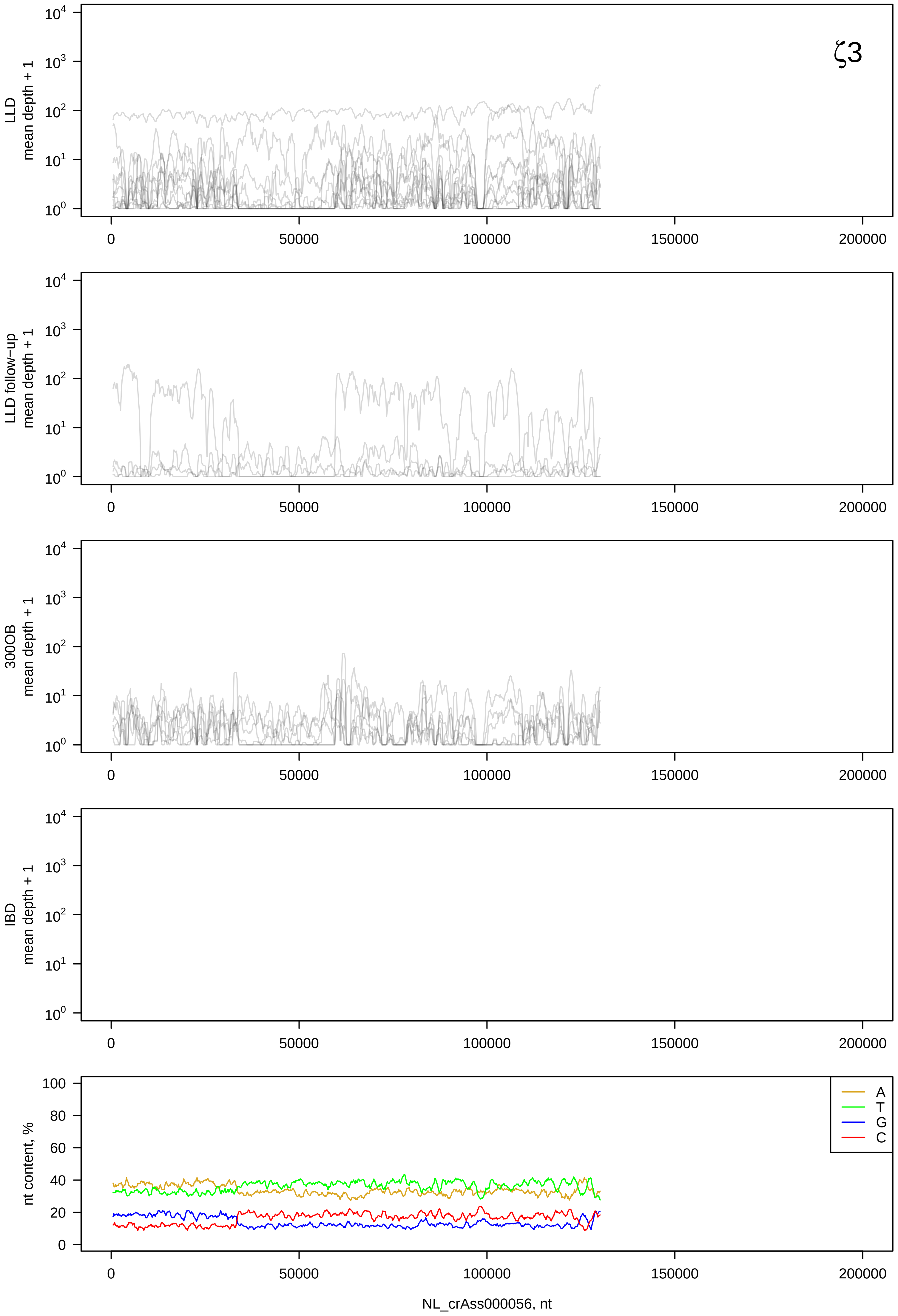
ζ3
LLD mean depth + 1
LLD follow−up mean depth + 1
300OB mean depth + 1
IBD mean depth + 1
nt content, %
A
T
G
C
NL_crAss000056, nt

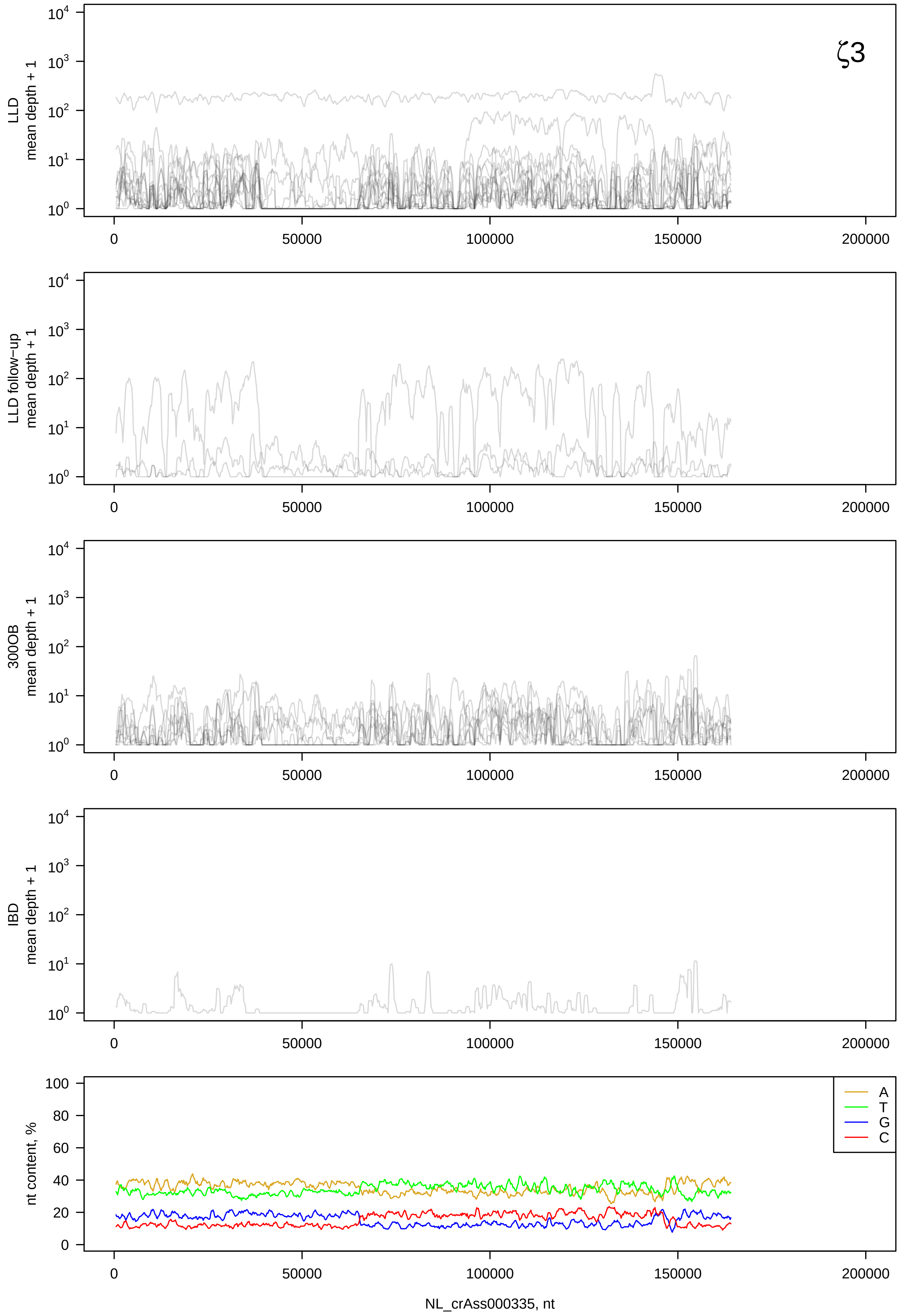

ζ3
LLD mean depth + 1
LLD follow−up mean depth + 1
300OB mean depth + 1
IBD mean depth + 1
nt content, %
A
T
G
C
NL_crAss000335, nt

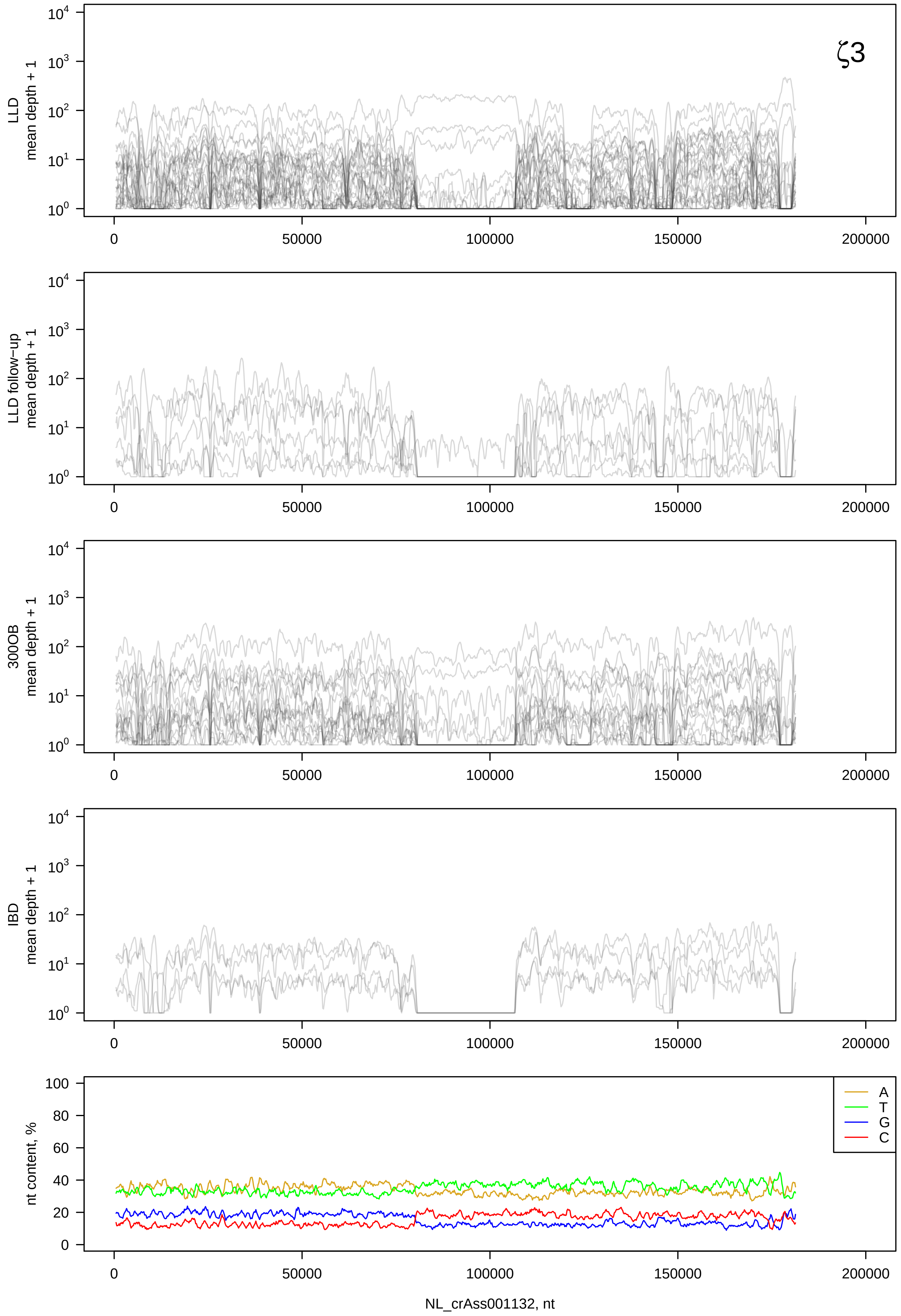

ζ3
LLD mean depth + 1
LLD follow-up mean depth + 1
300OB mean depth + 1
IBD mean depth + 1
nt content, %
A
T
G
C
$10^0$ $10^1$ $10^2$ $10^3$ $10^4$
0 20 40 60 80 100
0 50000 100000 150000 200000
NL_crAss001132, nt

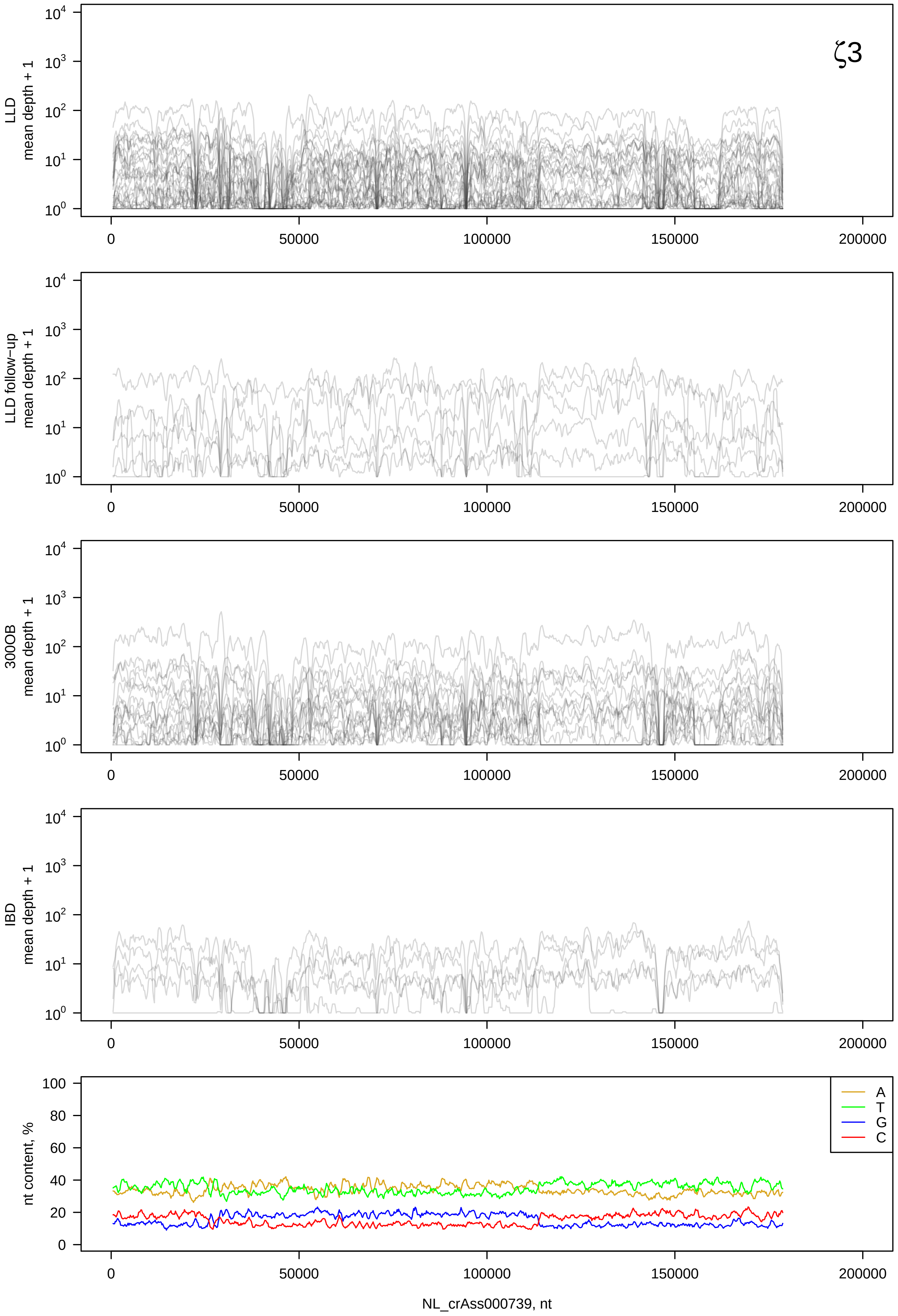

ζ3
LLD mean depth + 1
LLD follow-up mean depth + 1
300OB mean depth + 1
IBD mean depth + 1
nt content, %
A
T
G
C
NL_crAss000739, nt

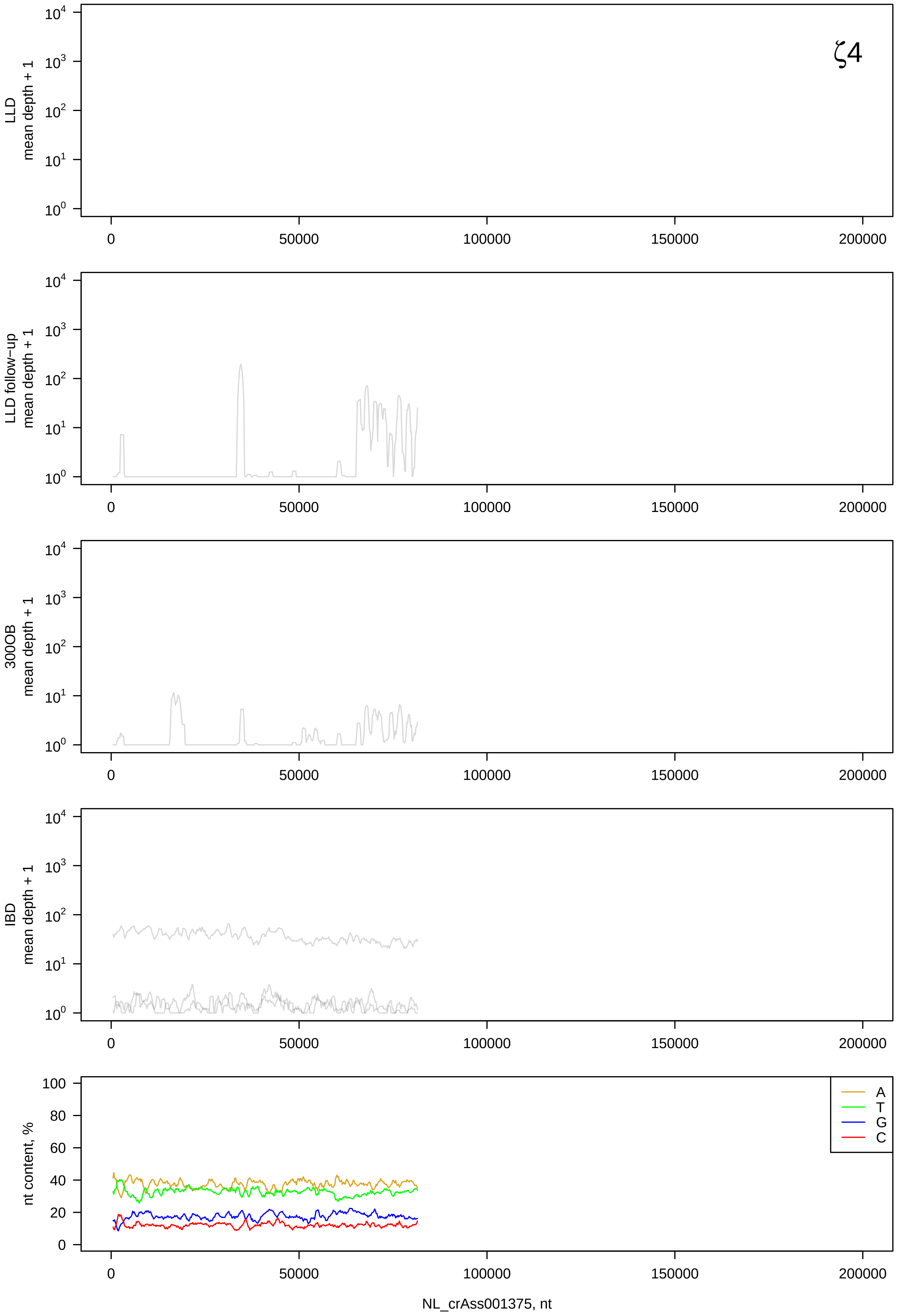

ζ4
LLD mean depth + 1
LLD follow-up mean depth + 1
300OB mean depth + 1
IBD mean depth + 1
nt content, %
A
T
G
C
0
50000
100000
150000
200000
NL_crAss001375, nt

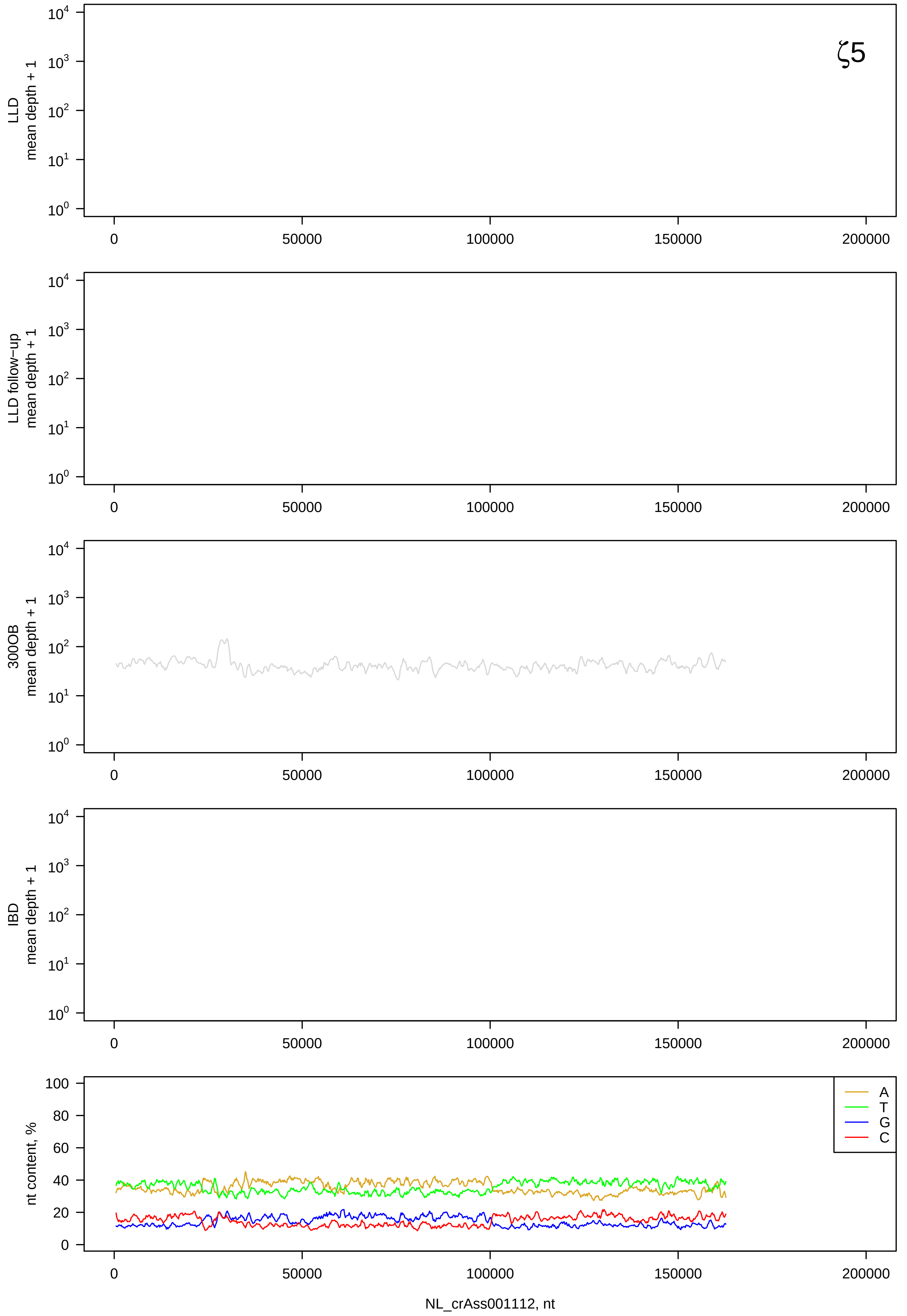

ζ5
LLD mean depth + 1
LLD follow−up mean depth + 1
300OB mean depth + 1
IBD mean depth + 1
nt content, %
A
T
G
C
NL_crAss001112, nt

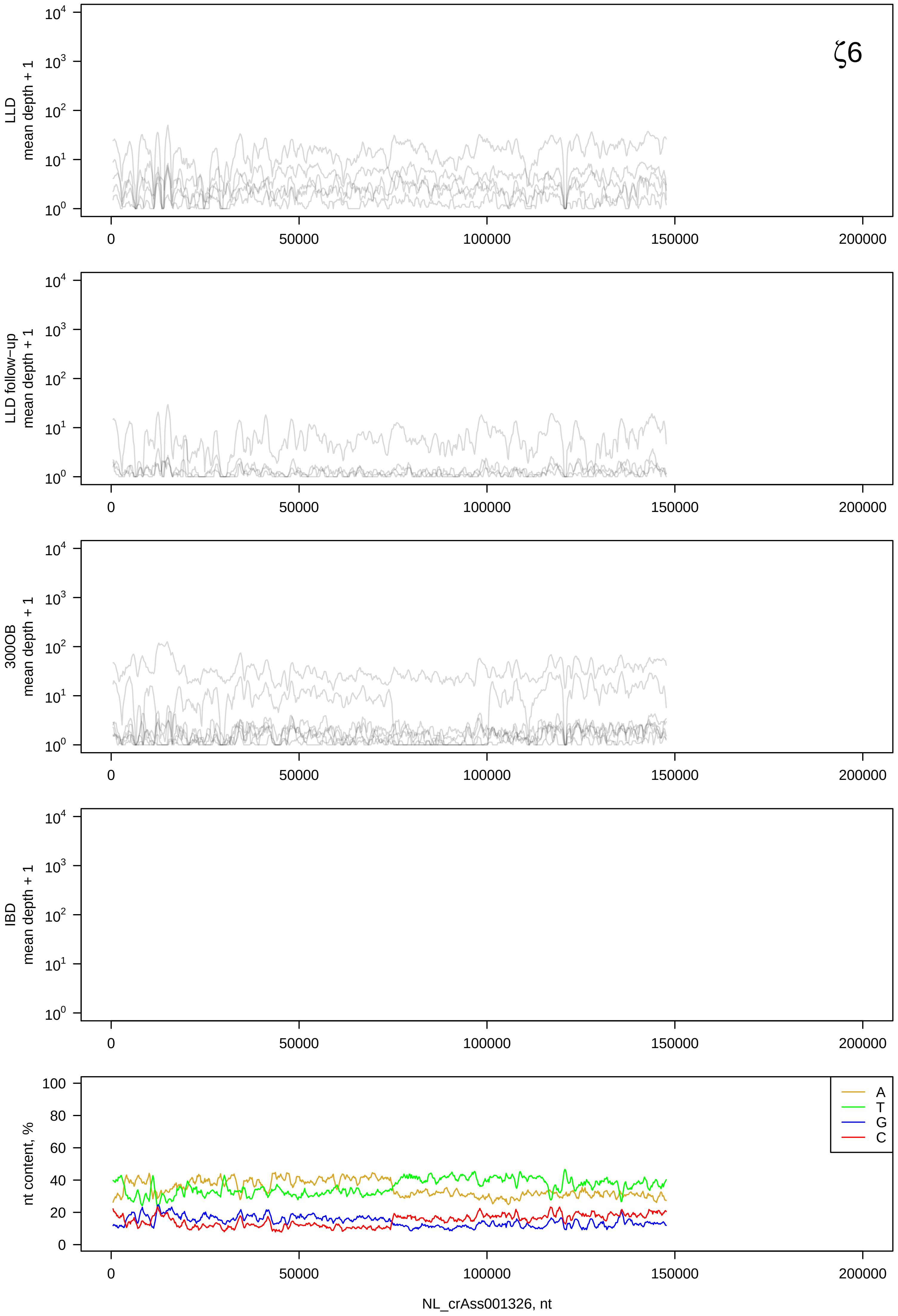
ζ6
LLD mean depth + 1
LLD follow−up mean depth + 1
300OB mean depth + 1
IBD mean depth + 1
nt content, %
A
T
G
C
0
50000
100000
150000
200000
NL_crAss001326, nt

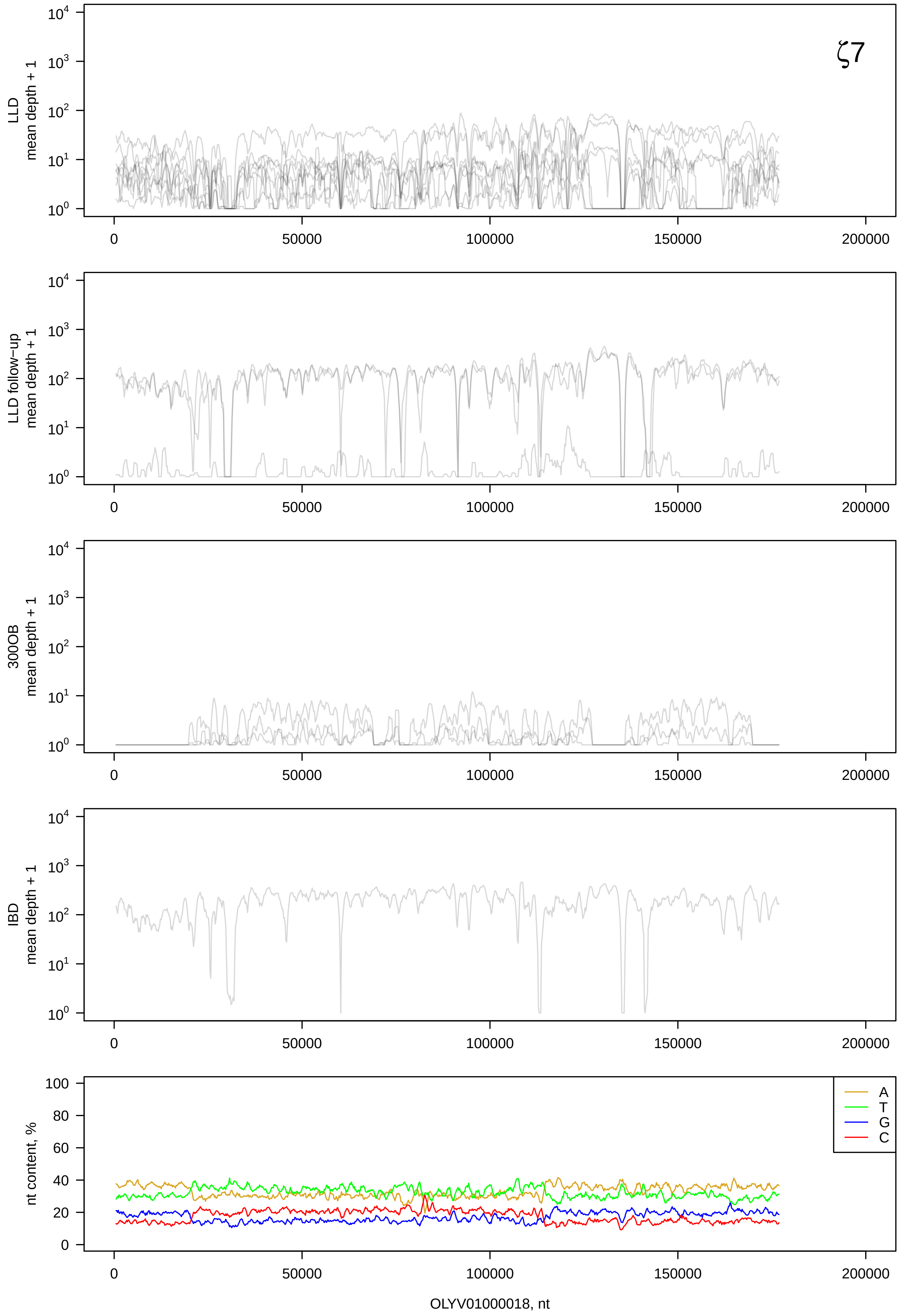

ζ7
LLD mean depth + 1
LLD follow−up mean depth + 1
300OB mean depth + 1
IBD mean depth + 1
nt content, %
A
T
G
C
0
50000
100000
150000
200000
OLYV01000018, nt

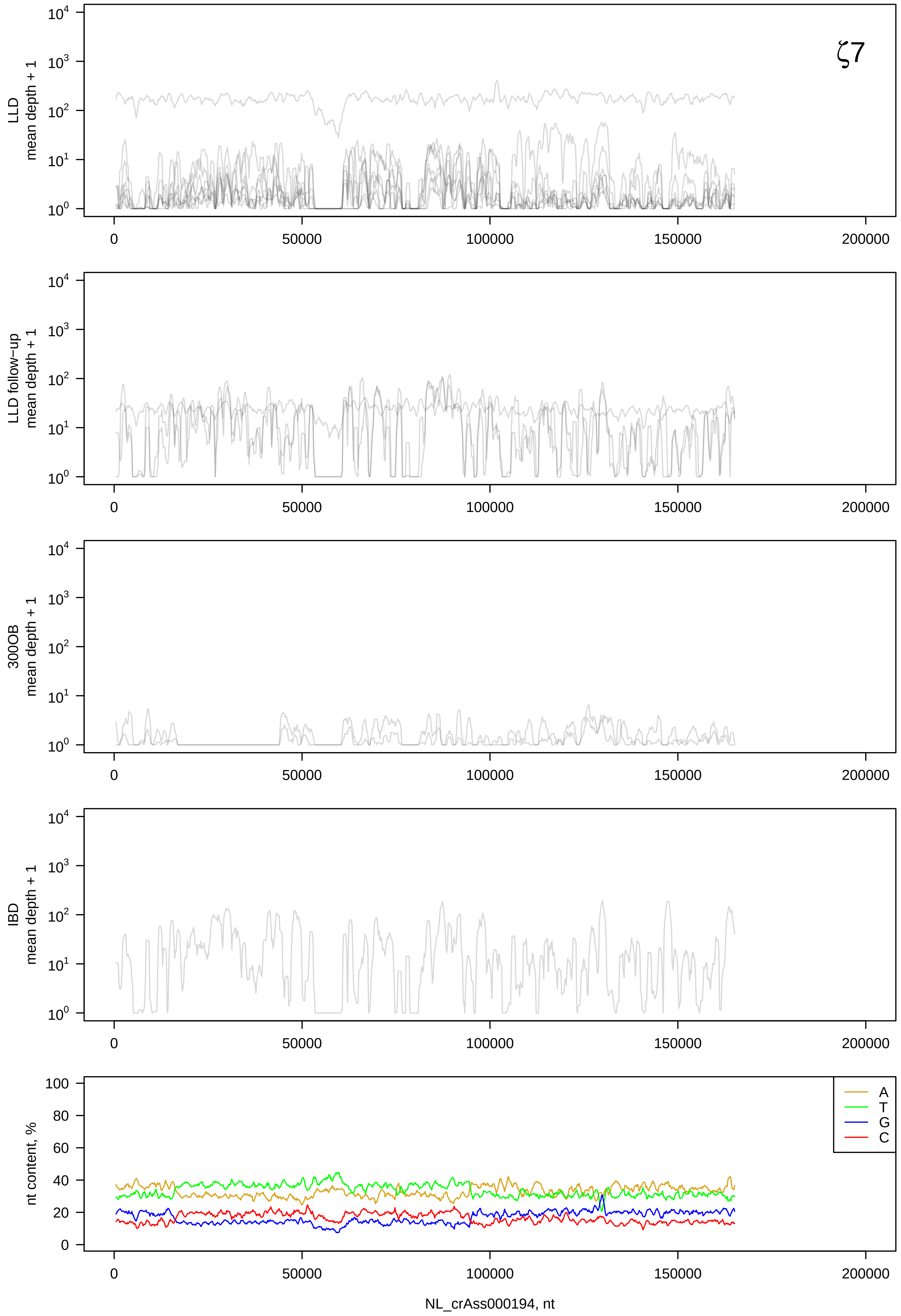

ζ7
LLD mean depth + 1
LLD follow−up mean depth + 1
300OB mean depth + 1
IBD mean depth + 1
nt content, %
A
T
G
C
0
50000
100000
150000
200000
NL_crAss000194, nt

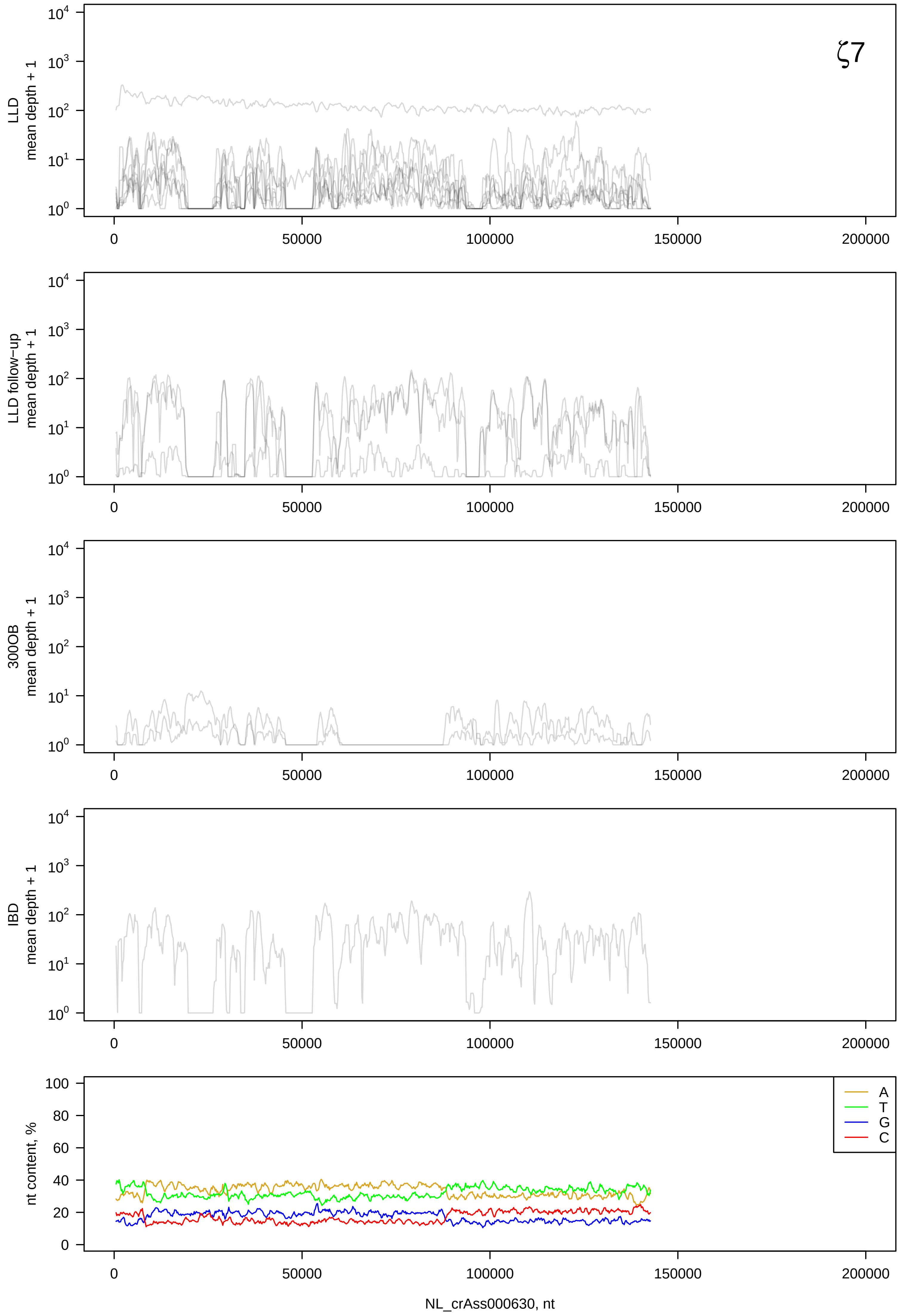

ζ7
LLD mean depth + 1
LLD follow−up mean depth + 1
300OB mean depth + 1
IBD mean depth + 1
nt content, %
A
T
G
C
NL_crAss000630, nt

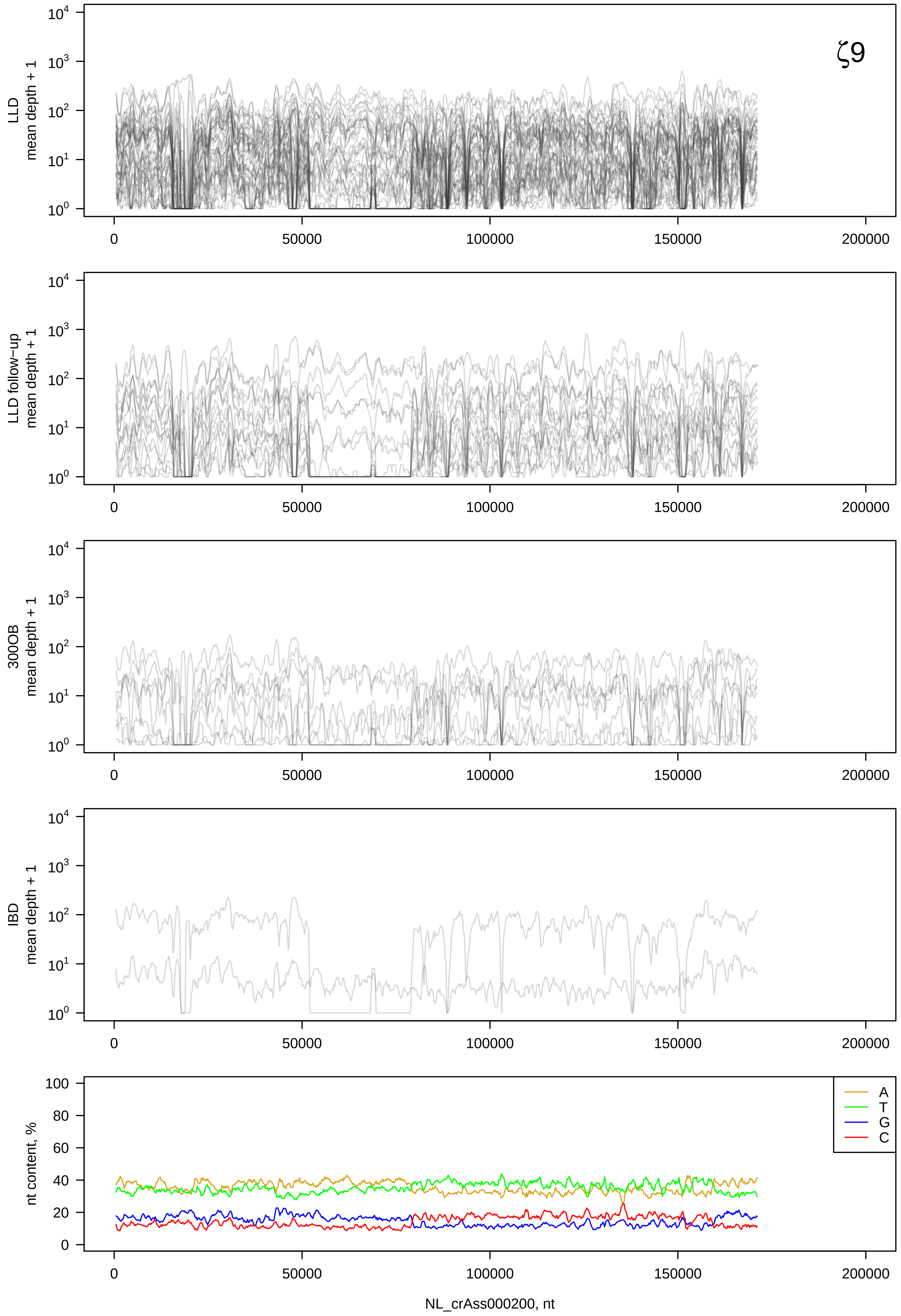

ζ9
LLD mean depth + 1
LLD follow-up mean depth + 1
300OB mean depth + 1
IBD mean depth + 1
nt content, %
A
T
G
C
NL_crAss000200, nt

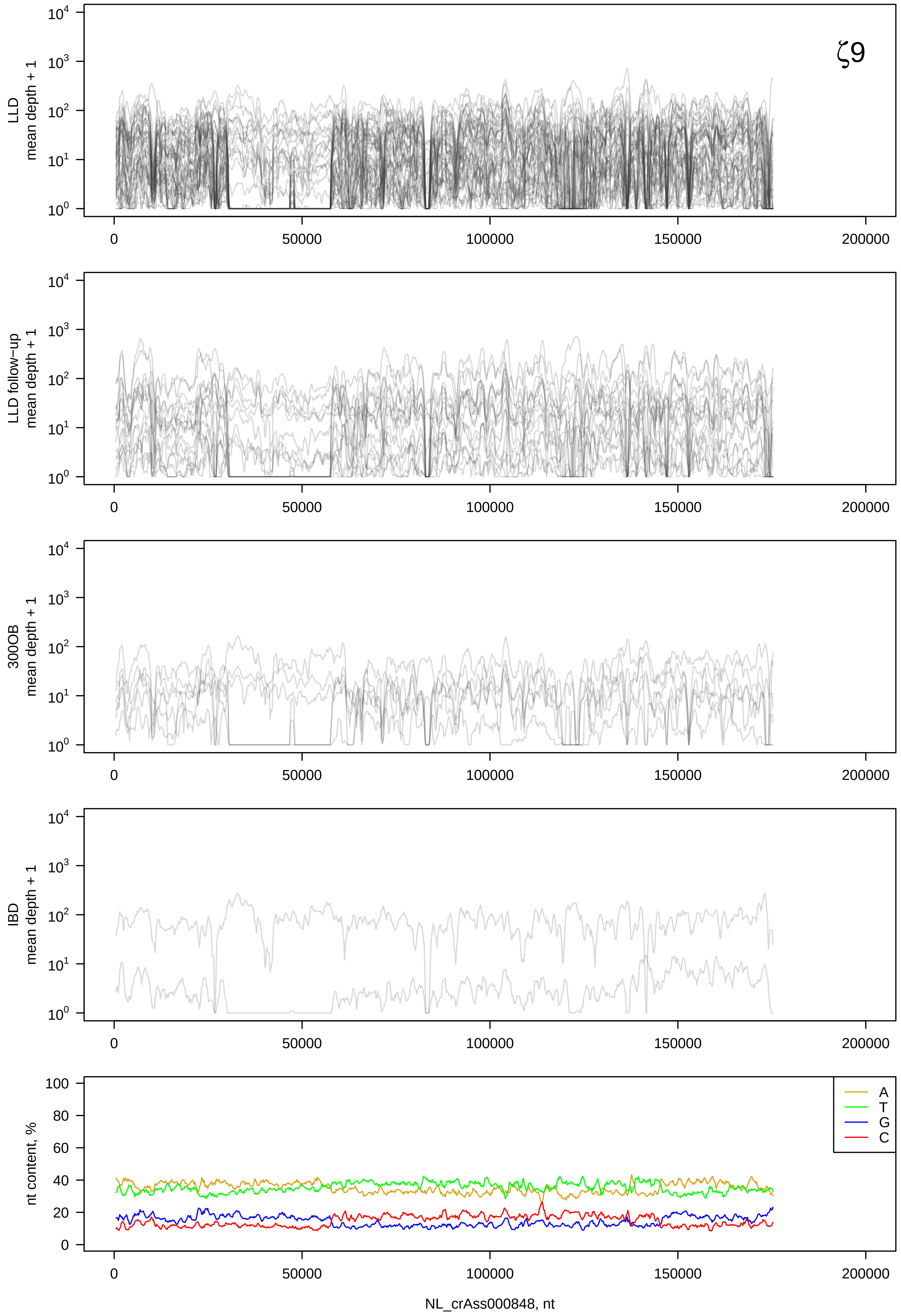

ζ9
LLD mean depth + 1
LLD follow−up mean depth + 1
300OB mean depth + 1
IBD mean depth + 1
nt content, %
A
T
G
C
NL_crAss000848, nt

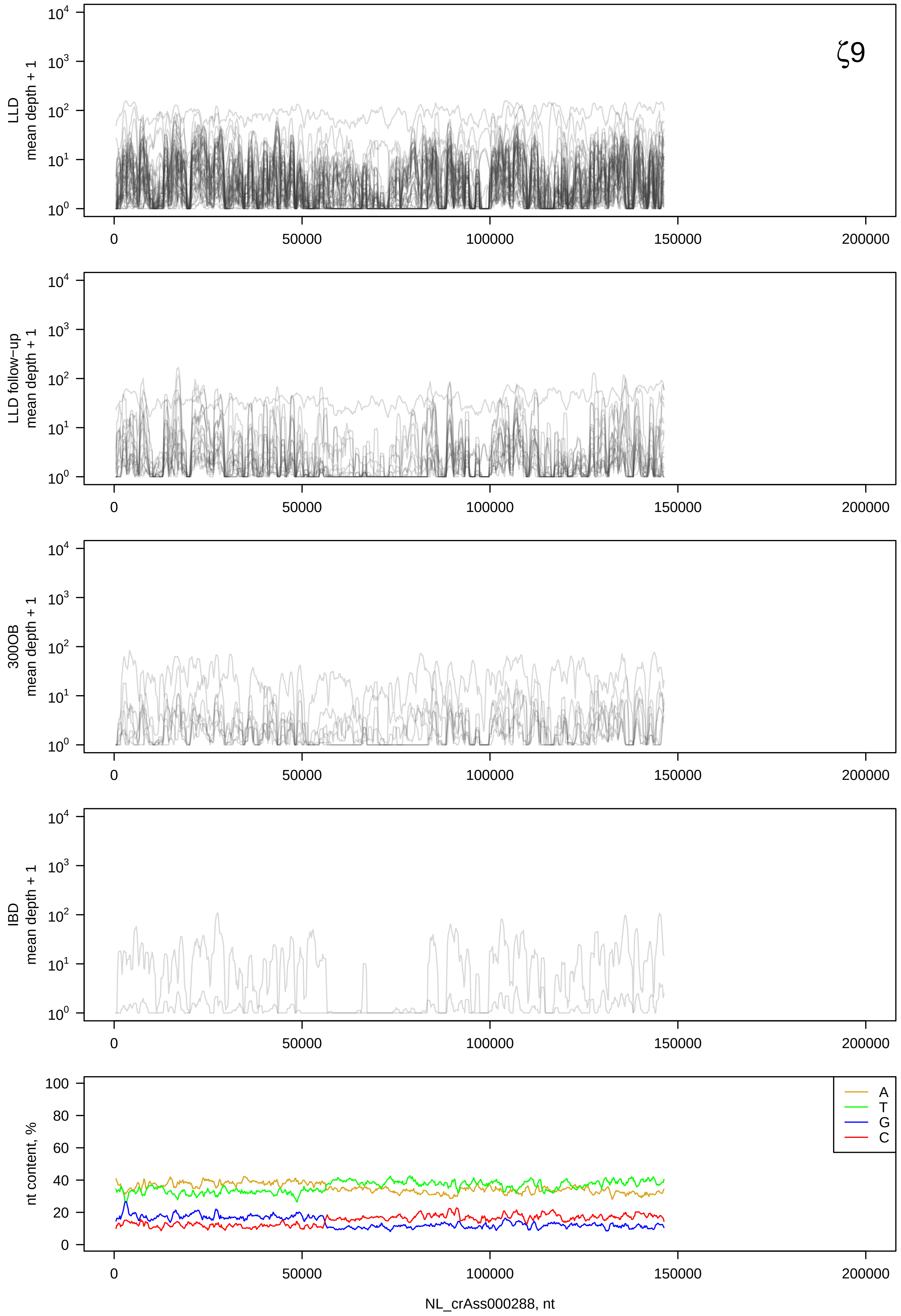
ζ9
LLD mean depth + 1
LLD follow-up mean depth + 1
300OB mean depth + 1
IBD mean depth + 1
nt content, %
A
T
G
C
0
50000
100000
150000
200000
NL_crAss000288, nt

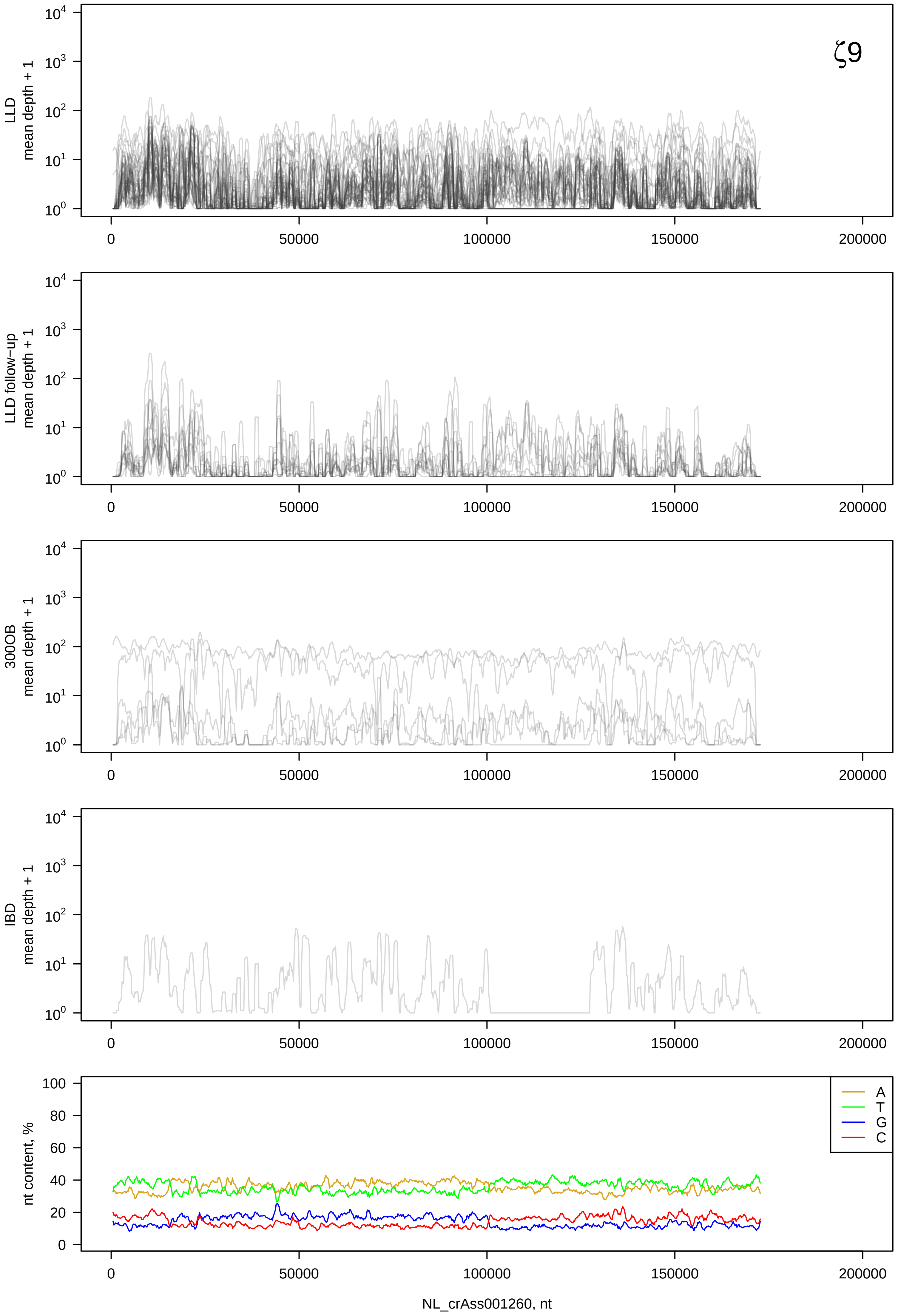

ζ9
LLD mean depth + 1
LLD follow−up mean depth + 1
300OB mean depth + 1
IBD mean depth + 1
nt content, %
A
T
G
C
NL_crAss001260, nt

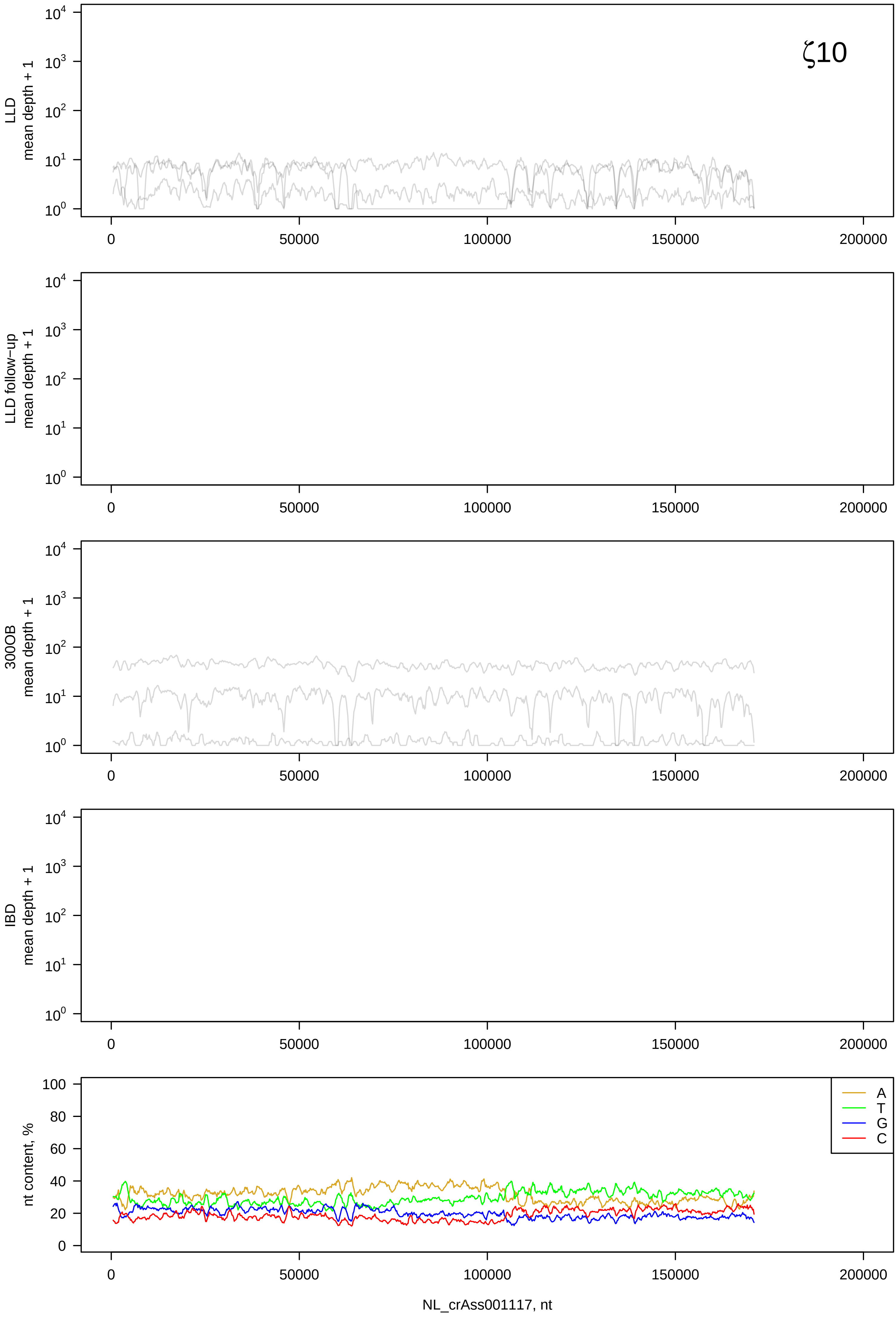

ζ10
LLD mean depth + 1
LLD follow-up mean depth + 1
300OB mean depth + 1
IBD mean depth + 1
nt content, %
A
T
G
C
NL_crAss001117, nt

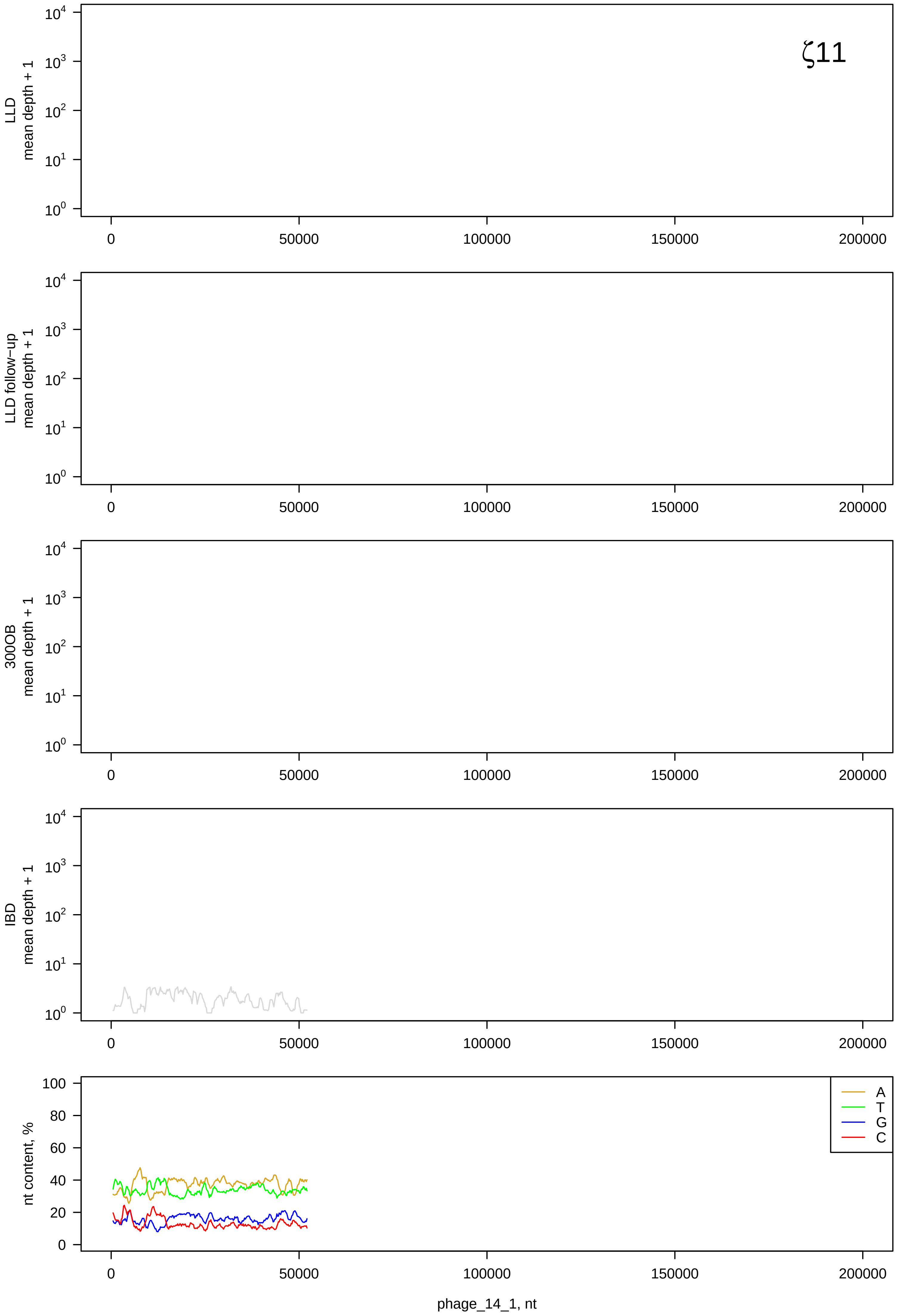

ζ11
LLD mean depth + 1
LLD follow−up mean depth + 1
300OB mean depth + 1
IBD mean depth + 1
nt content, %
A
T
G
C
0
50000
100000
150000
200000
phage_14_1, nt

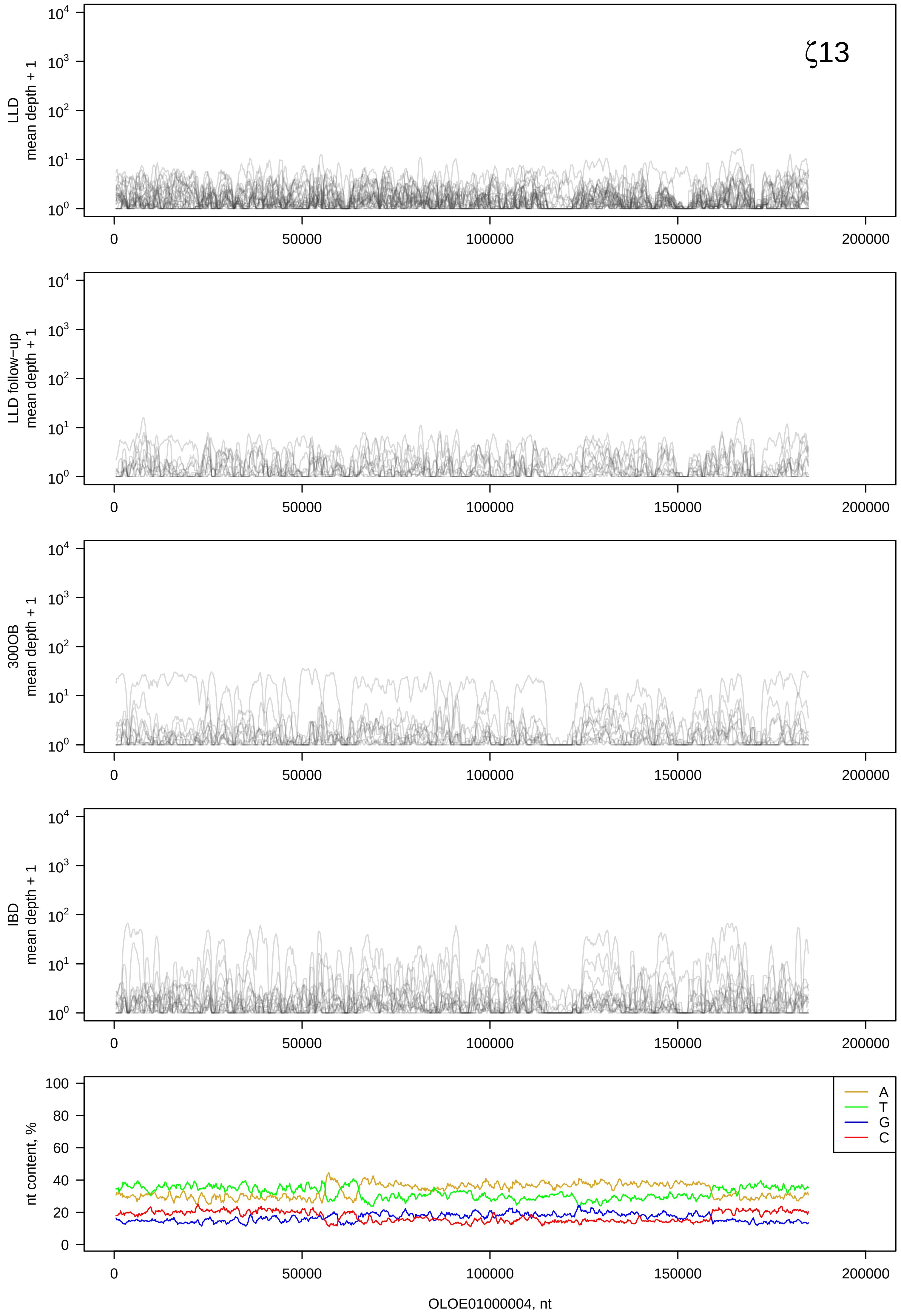

ζ13
LLD
mean depth + 1
LLD follow−up
mean depth + 1
300OB
mean depth + 1
IBD
mean depth + 1
nt content, %
A
T
G
C
0
50000
100000
150000
200000
OLOE01000004, nt

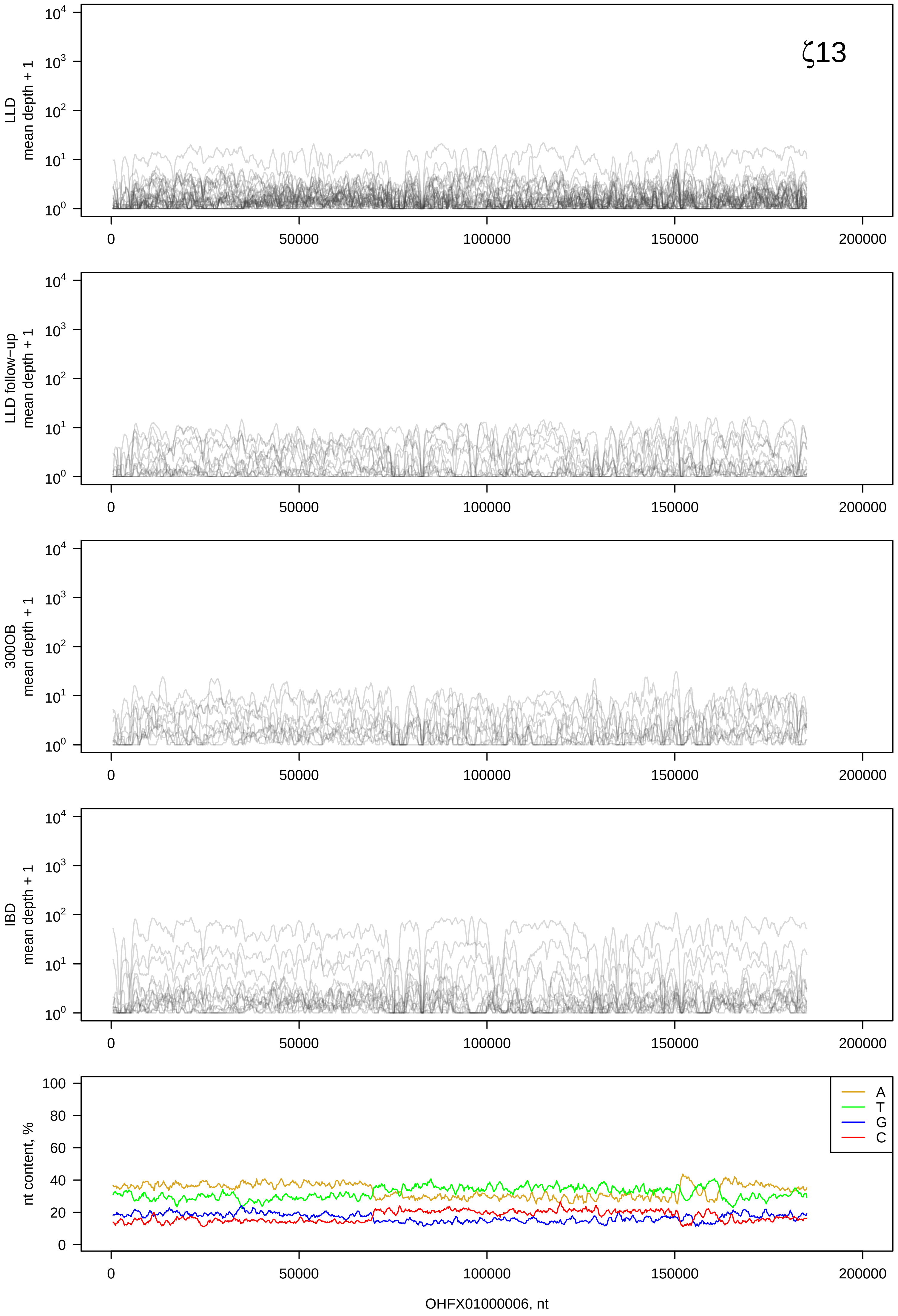

ζ13
LLD mean depth + 1
LLD follow-up mean depth + 1
300OB mean depth + 1
IBD mean depth + 1
nt content, %
A
T
G
C
0
50000
100000
150000
200000
OHFX01000006, nt

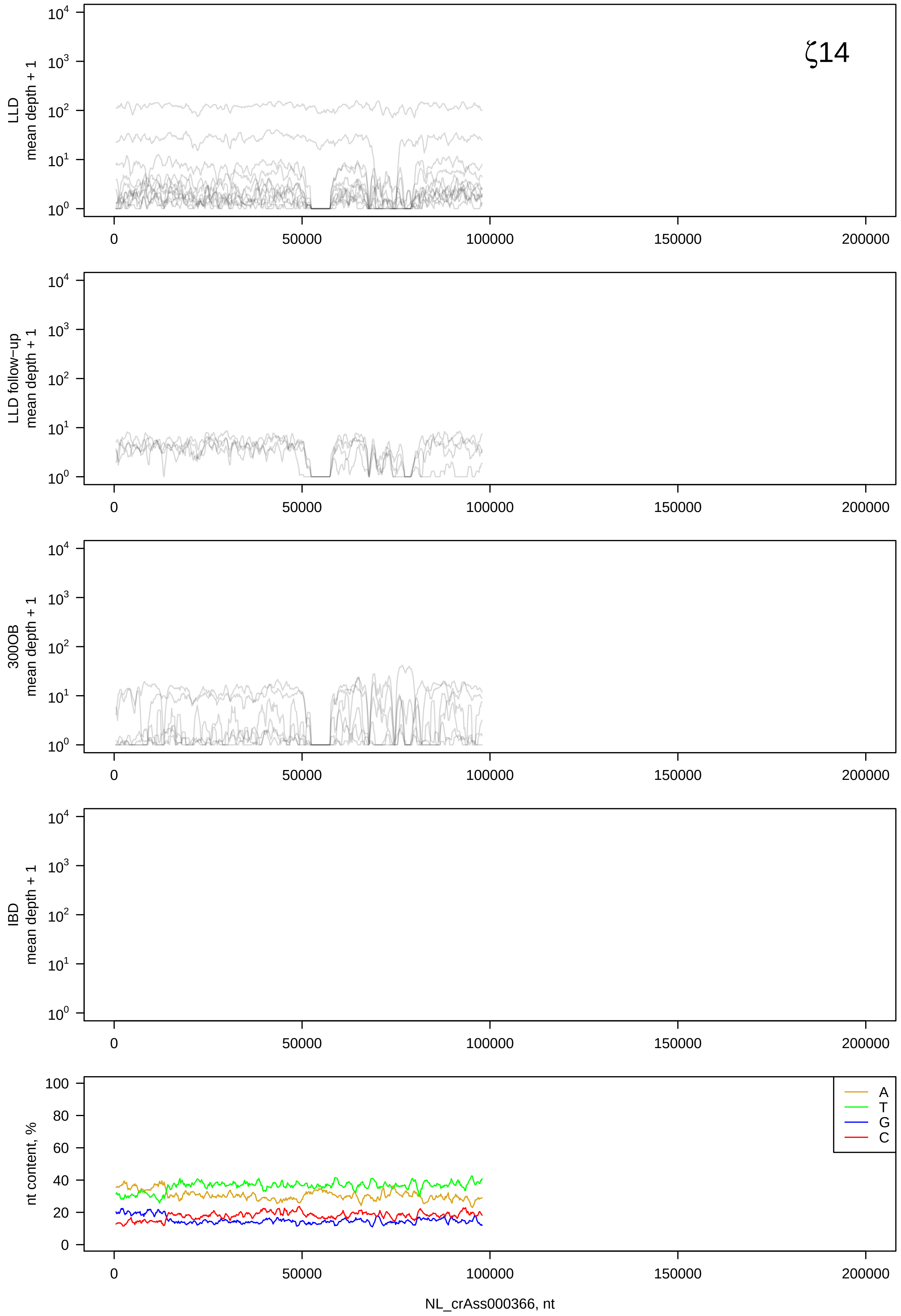

ζ14
LLD mean depth + 1
LLD follow−up mean depth + 1
300OB mean depth + 1
IBD mean depth + 1
nt content, %
A
T
G
C
NL_crAss000366, nt

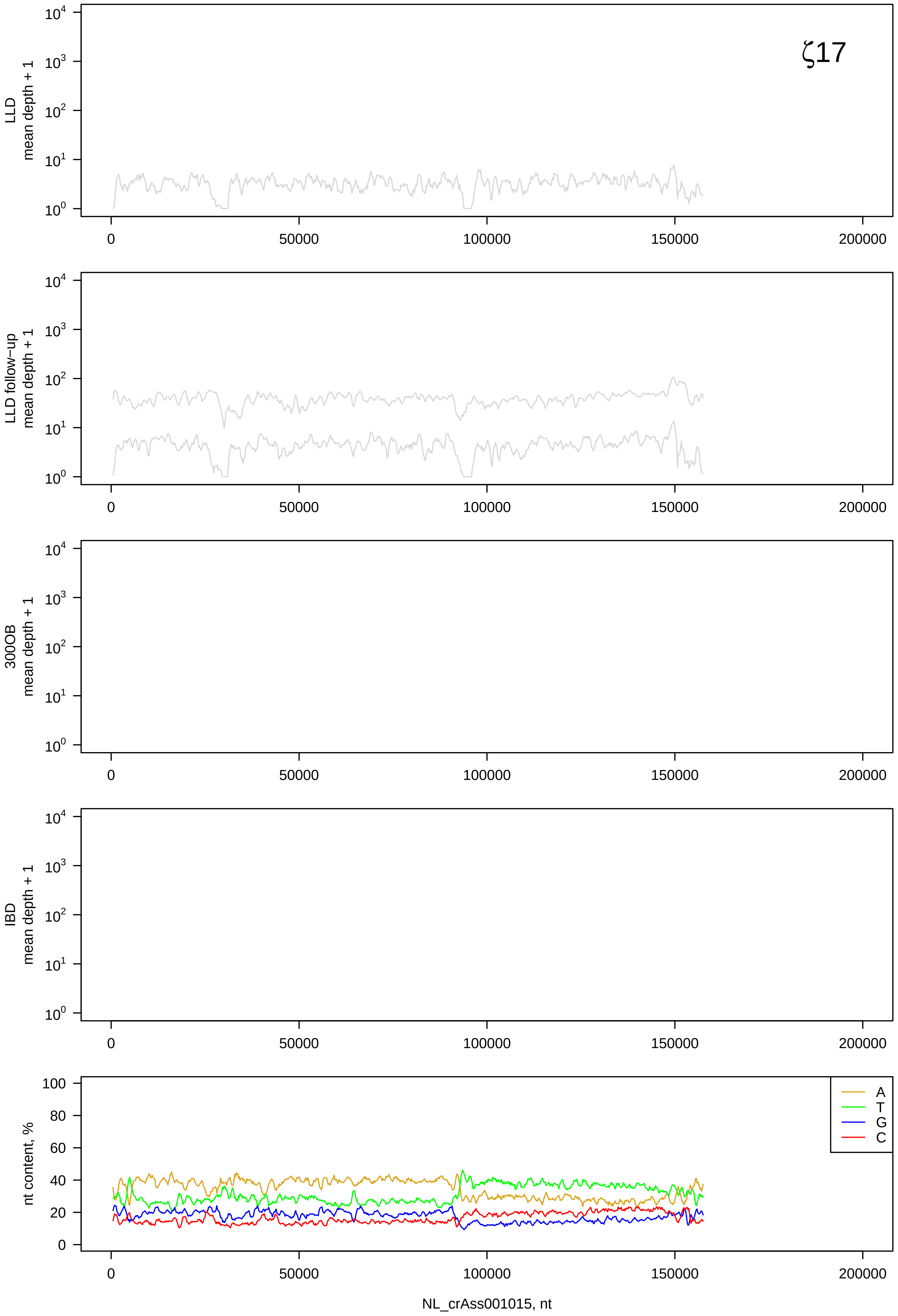

ζ17
LLD
mean depth + 1
LLD follow−up
mean depth + 1
300OB
mean depth + 1
IBD
mean depth + 1
nt content, %
A
T
G
C
NL_crAss001015, nt

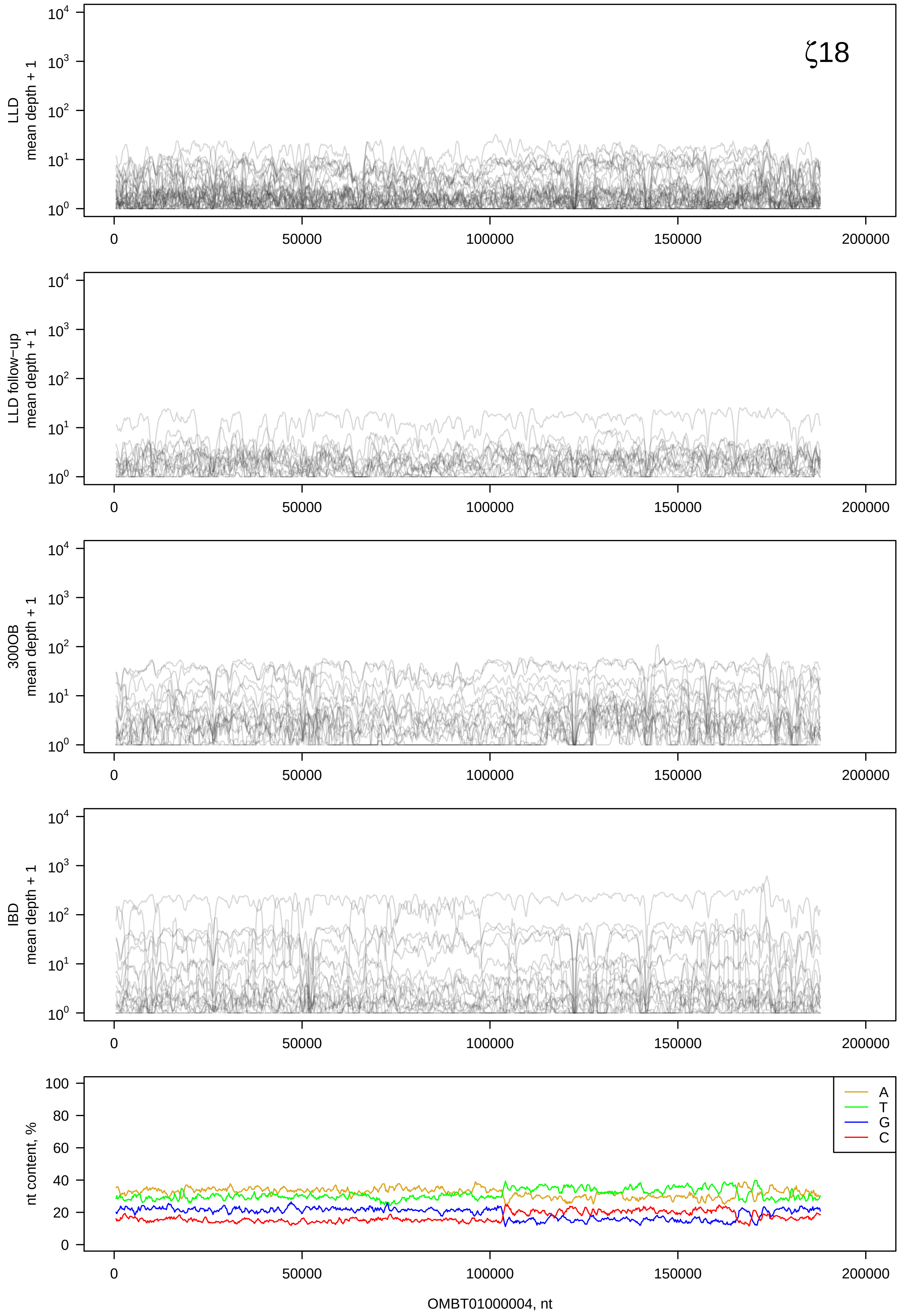

ζ18
LLD
mean depth + 1
LLD follow-up
mean depth + 1
300OB
mean depth + 1
IBD
mean depth + 1
nt content, %
A
T
G
C
0
50000
100000
150000
200000
OMBT01000004, nt

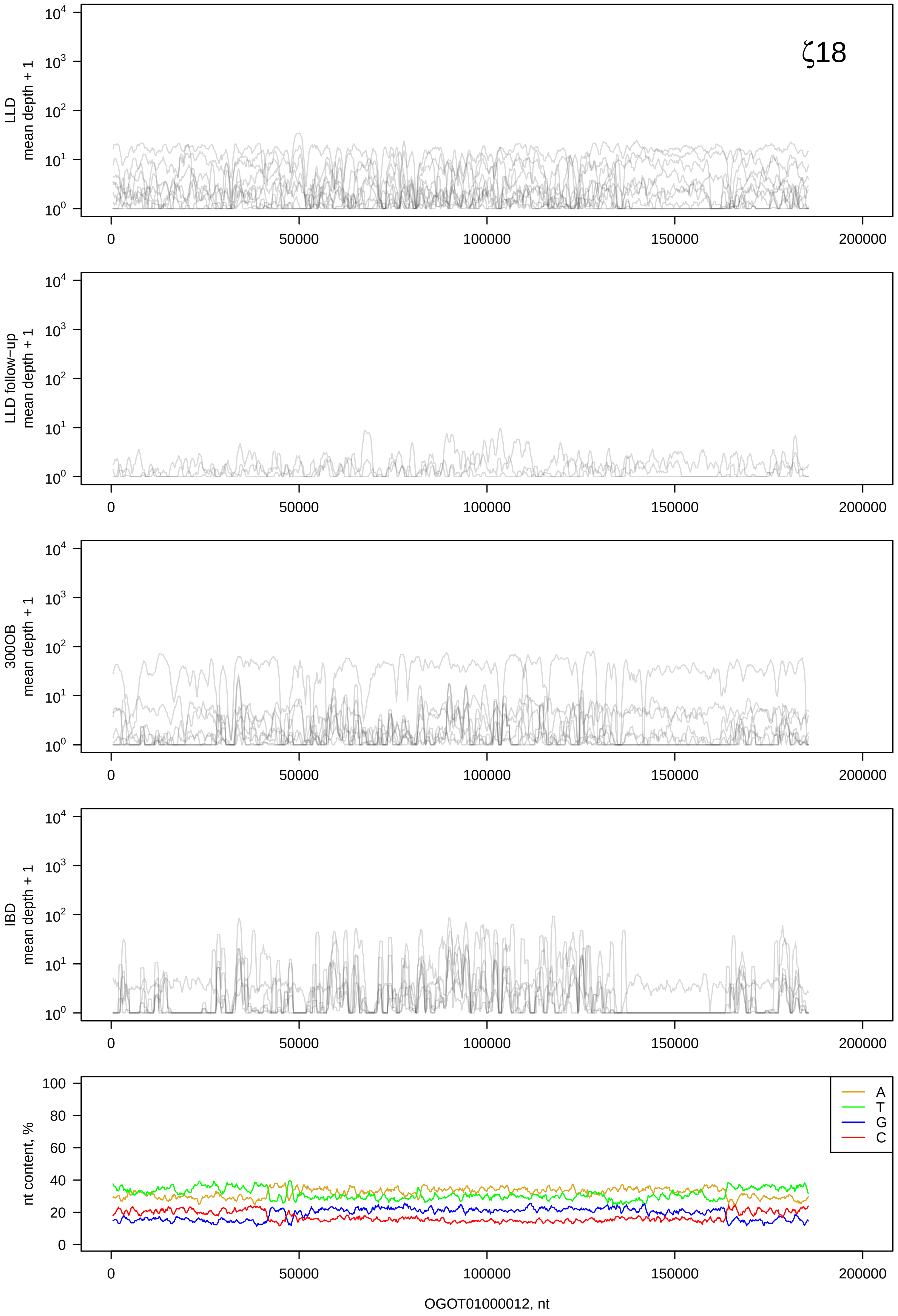

ζ18
LLD mean depth + 1
LLD follow-up mean depth + 1
300OB mean depth + 1
IBD mean depth + 1
nt content, %
A
T
G
C
0
50000
100000
150000
200000
OGOT01000012, nt

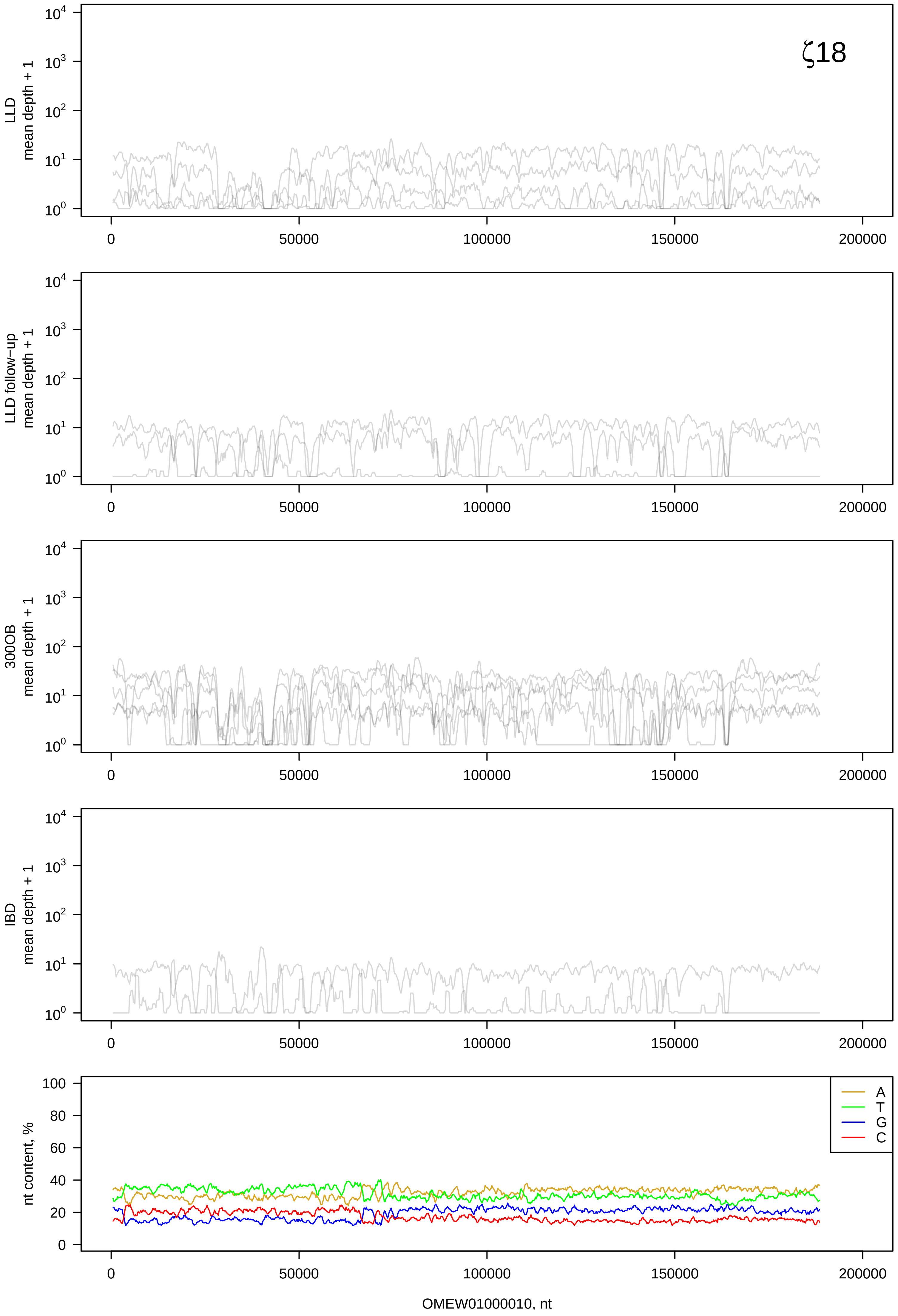
ζ18
LLD
mean depth + 1
LLD follow−up
mean depth + 1
300OB
mean depth + 1
IBD
mean depth + 1
nt content, %
A
T
G
C
OMEW01000010, nt

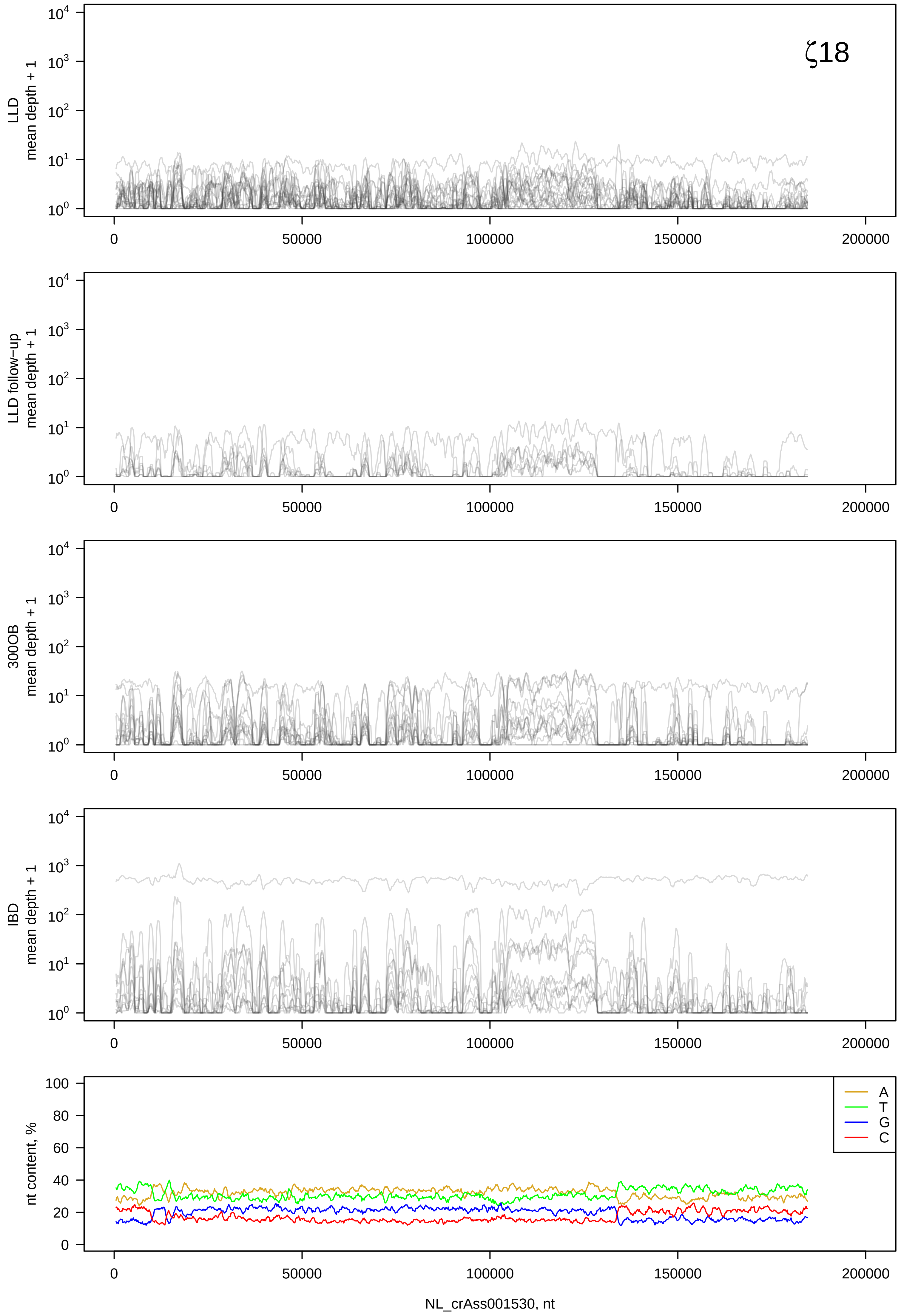
ζ18
LLD mean depth + 1
LLD follow-up mean depth + 1
300OB mean depth + 1
IBD mean depth + 1
nt content, %
A
T
G
C
NL_crAss001530, nt

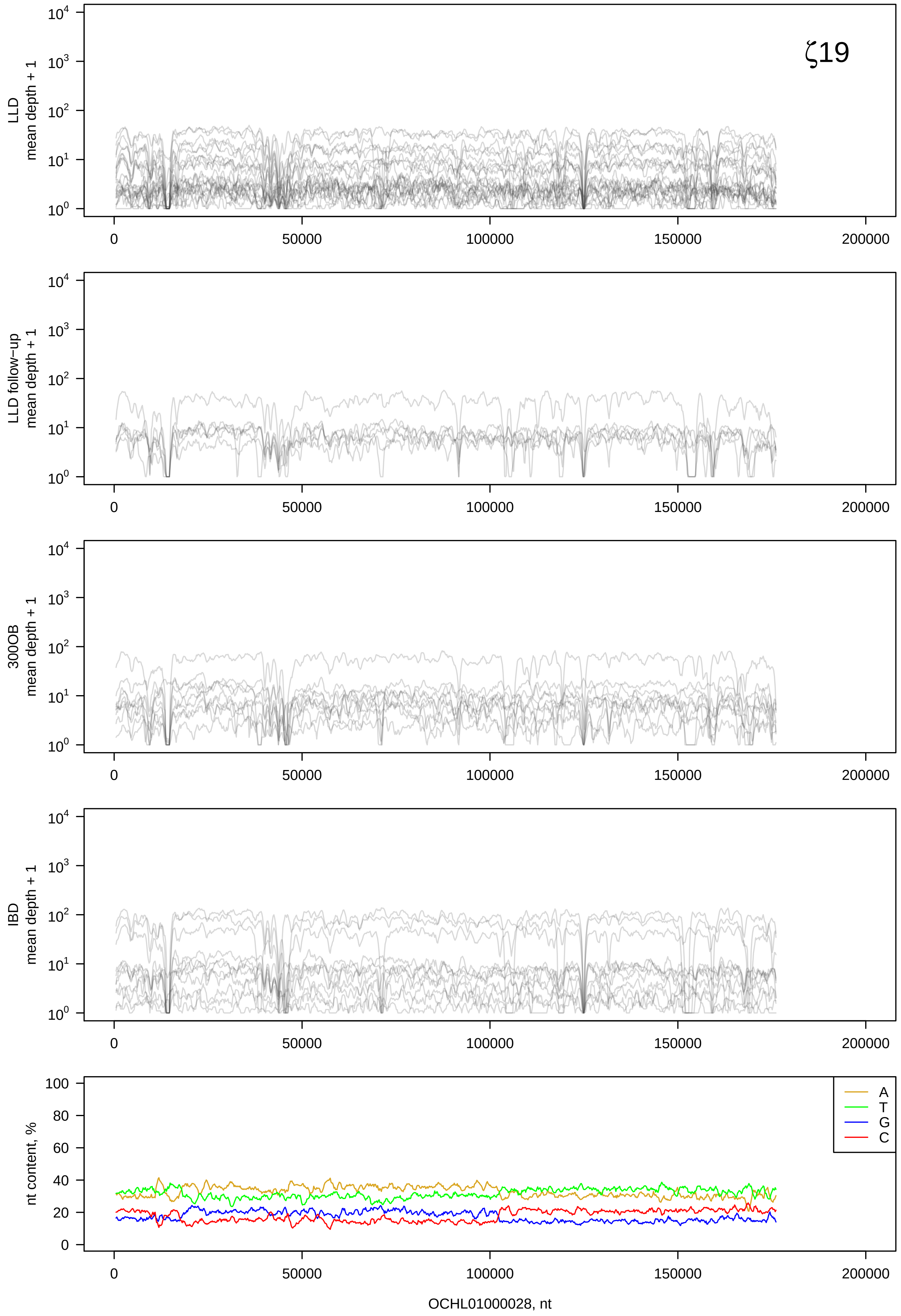

ζ19
LLD mean depth + 1
LLD follow−up mean depth + 1
300OB mean depth + 1
IBD mean depth + 1
nt content, %
A
T
G
C
0
50000
100000
150000
200000
OCHL01000028, nt

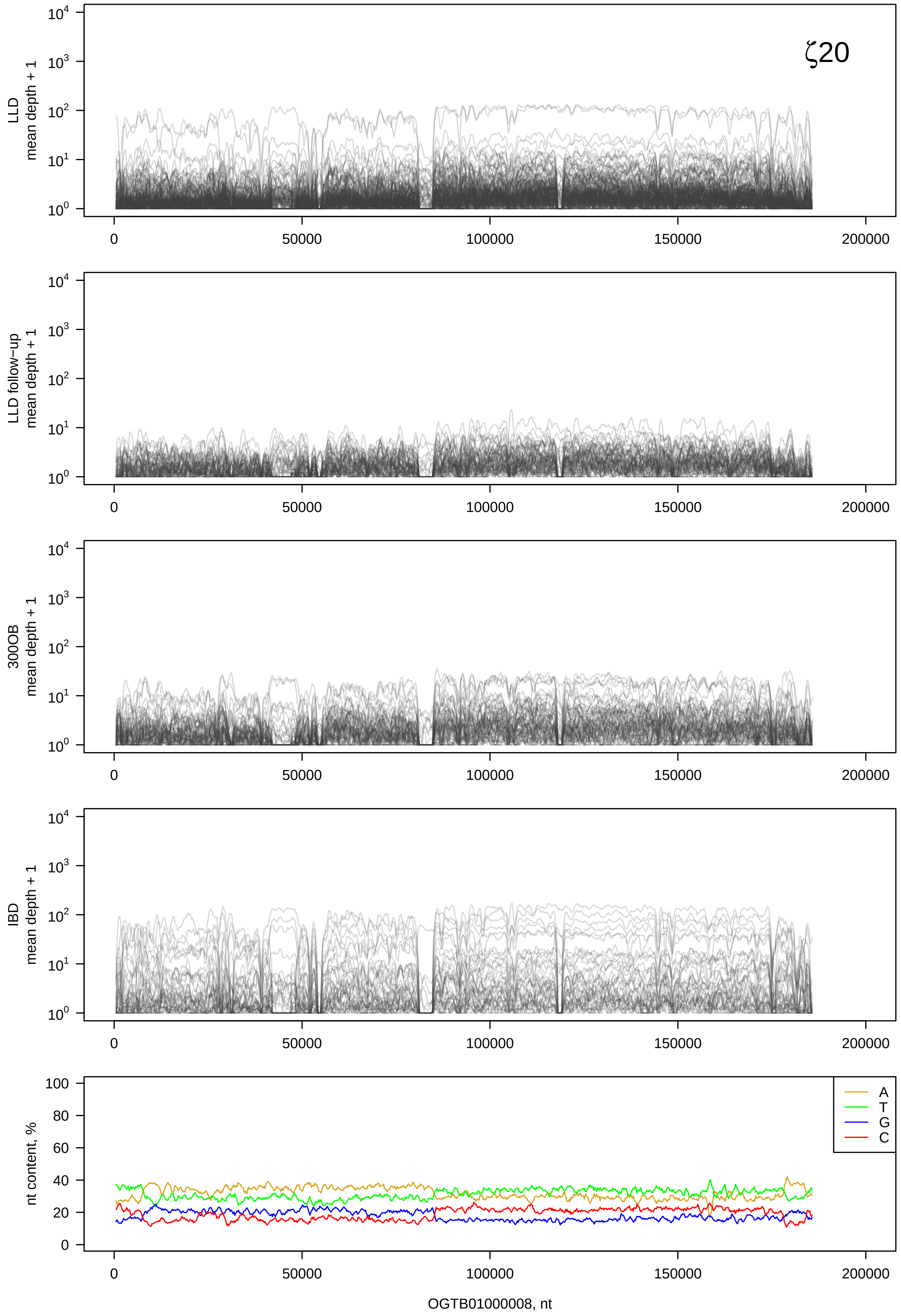

ζ20
LLD
mean depth + 1
LLD follow-up
mean depth + 1
300OB
mean depth + 1
IBD
mean depth + 1
nt content, %
A
T
G
C
OGTB01000008, nt

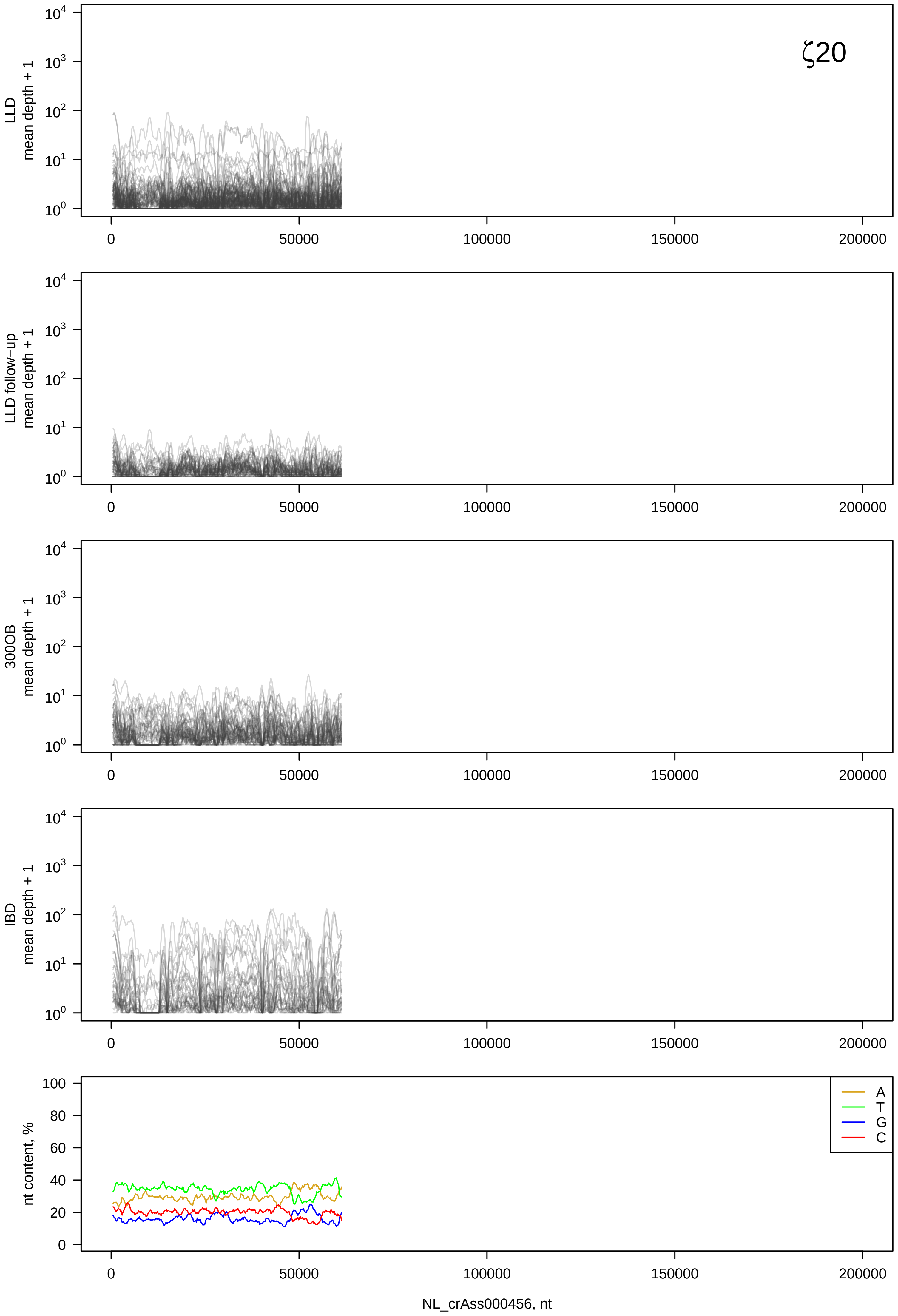

ζ20
LLD mean depth + 1
LLD follow-up mean depth + 1
300OB mean depth + 1
IBD mean depth + 1
nt content, %
A
T
G
C
0
50000
100000
150000
200000
NL_crAss000456, nt

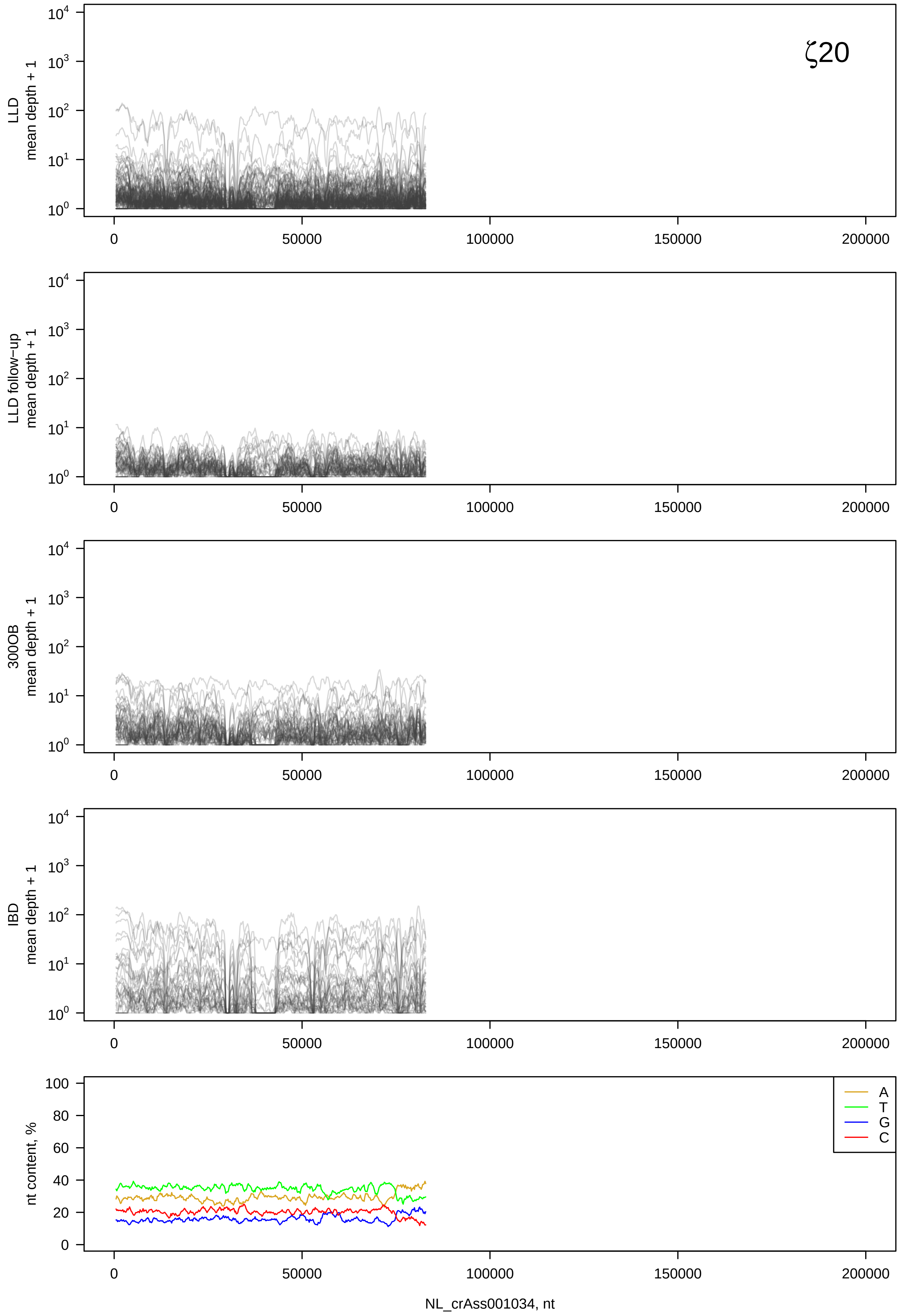
ζ20
LLD
mean depth + 1
LLD follow−up
mean depth + 1
300OB
mean depth + 1
IBD
mean depth + 1
nt content, %
A
T
G
C
NL_crAss001034, nt

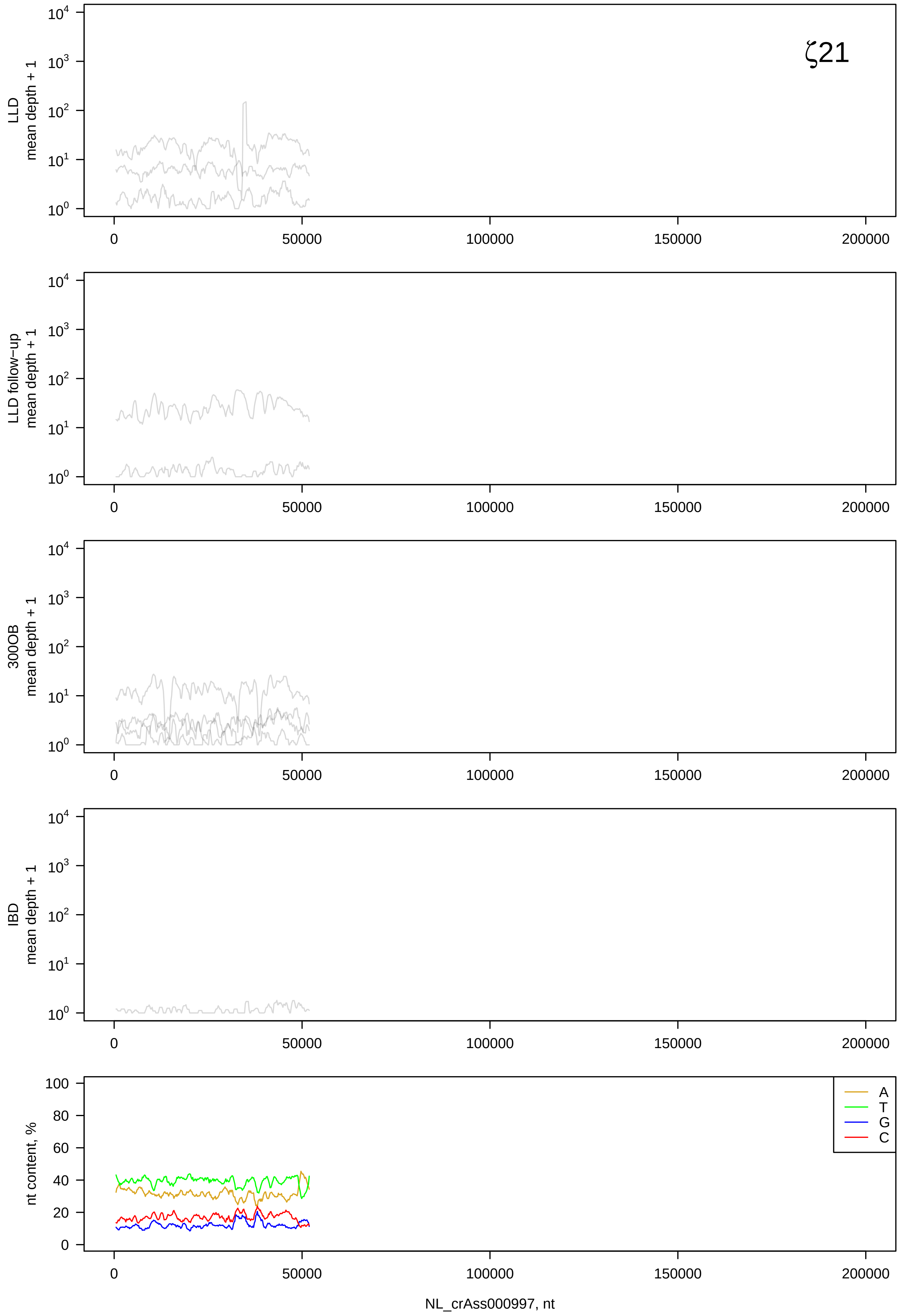
ζ21
LLD mean depth + 1
LLD follow-up mean depth + 1
300OB mean depth + 1
IBD mean depth + 1
nt content, %
A
T
G
C
NL_crAss000997, nt

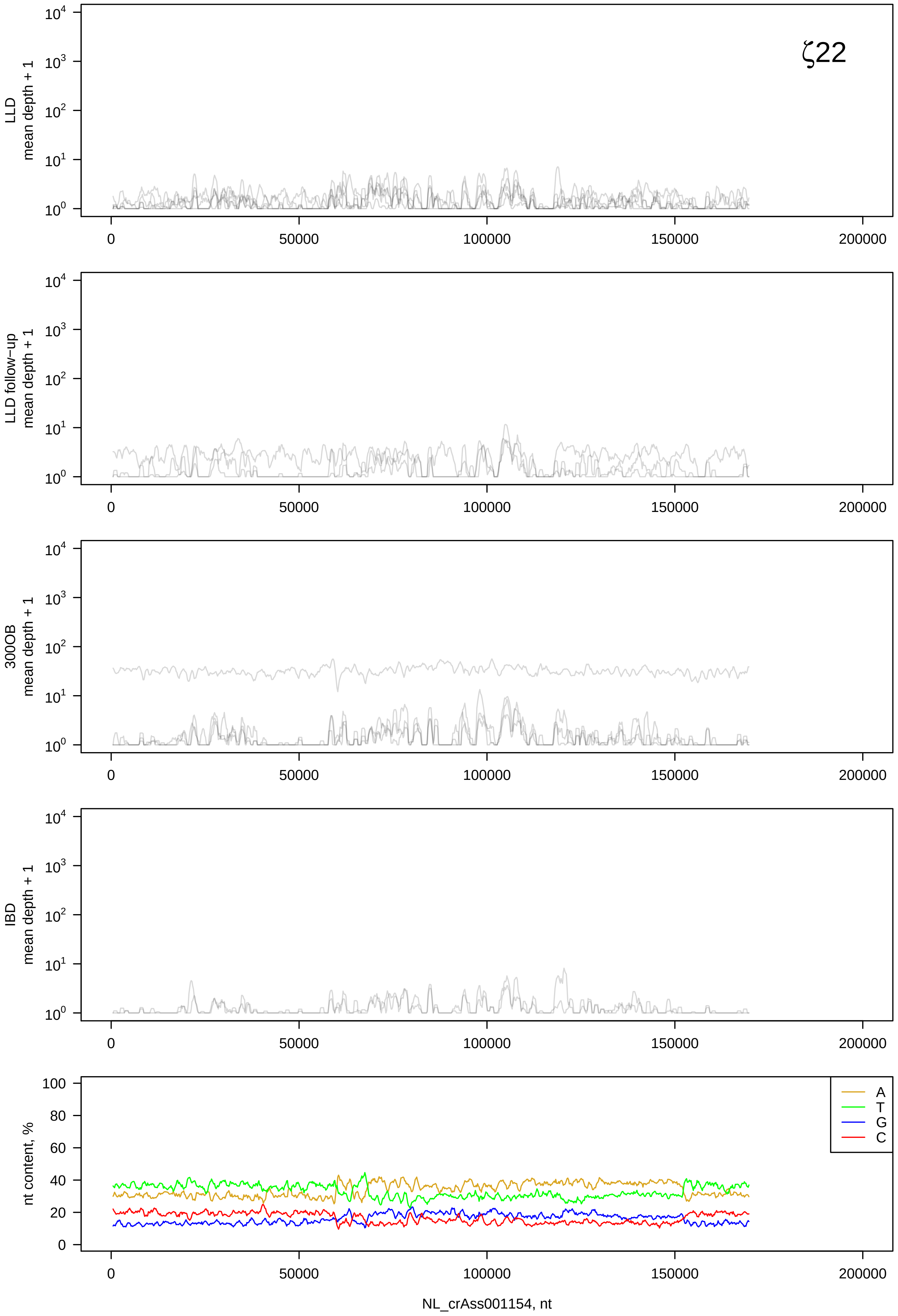

ζ22
LLD
mean depth + 1
LLD follow−up
mean depth + 1
300OB
mean depth + 1
IBD
mean depth + 1
nt content, %
A
T
G
C
0
50000
100000
150000
200000
NL_crAss001154, nt

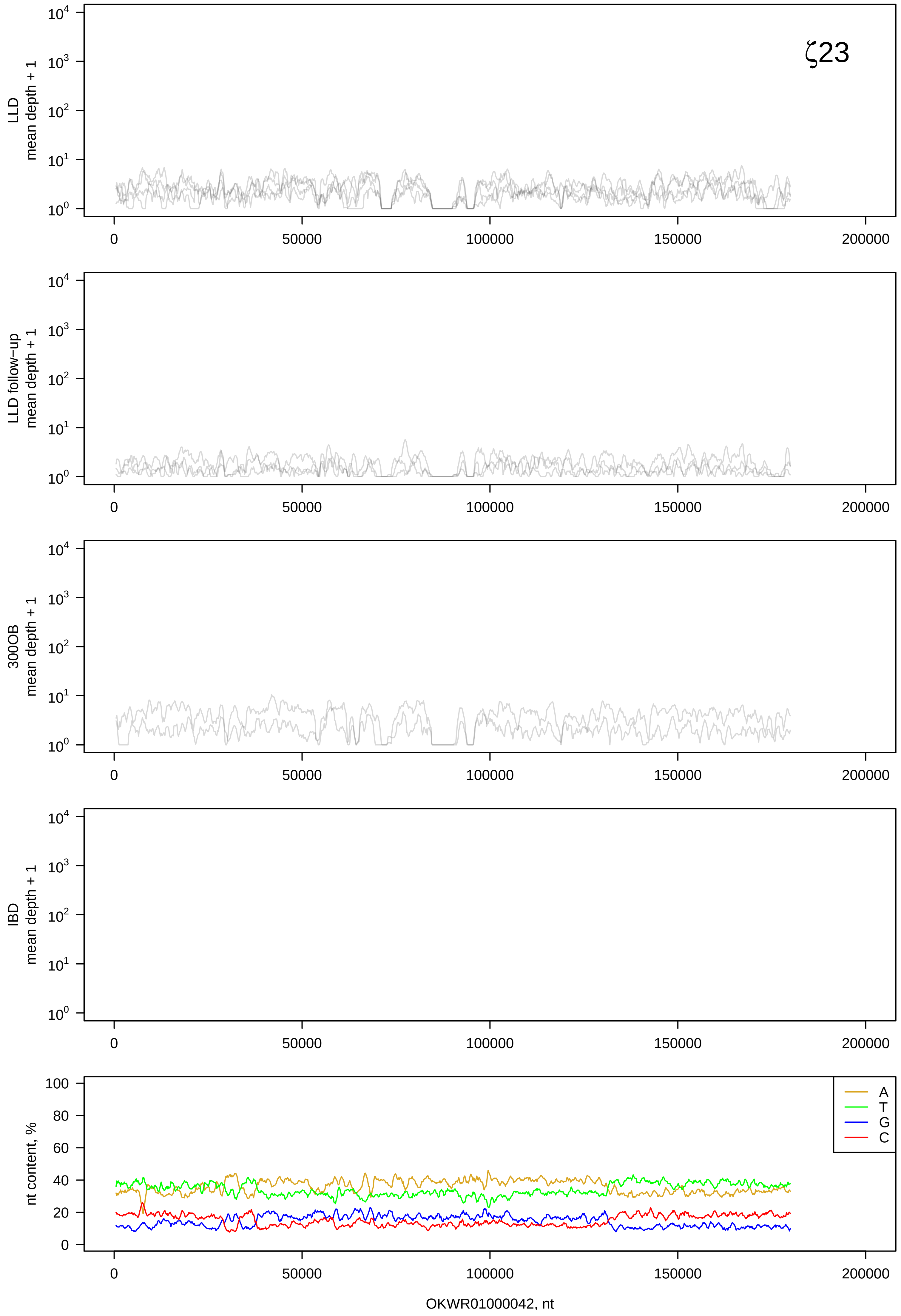

ζ23
LLD mean depth + 1
LLD follow-up mean depth + 1
300OB mean depth + 1
IBD mean depth + 1
nt content, %
A
T
G
C
0
50000
100000
150000
200000
OKWR01000042, nt

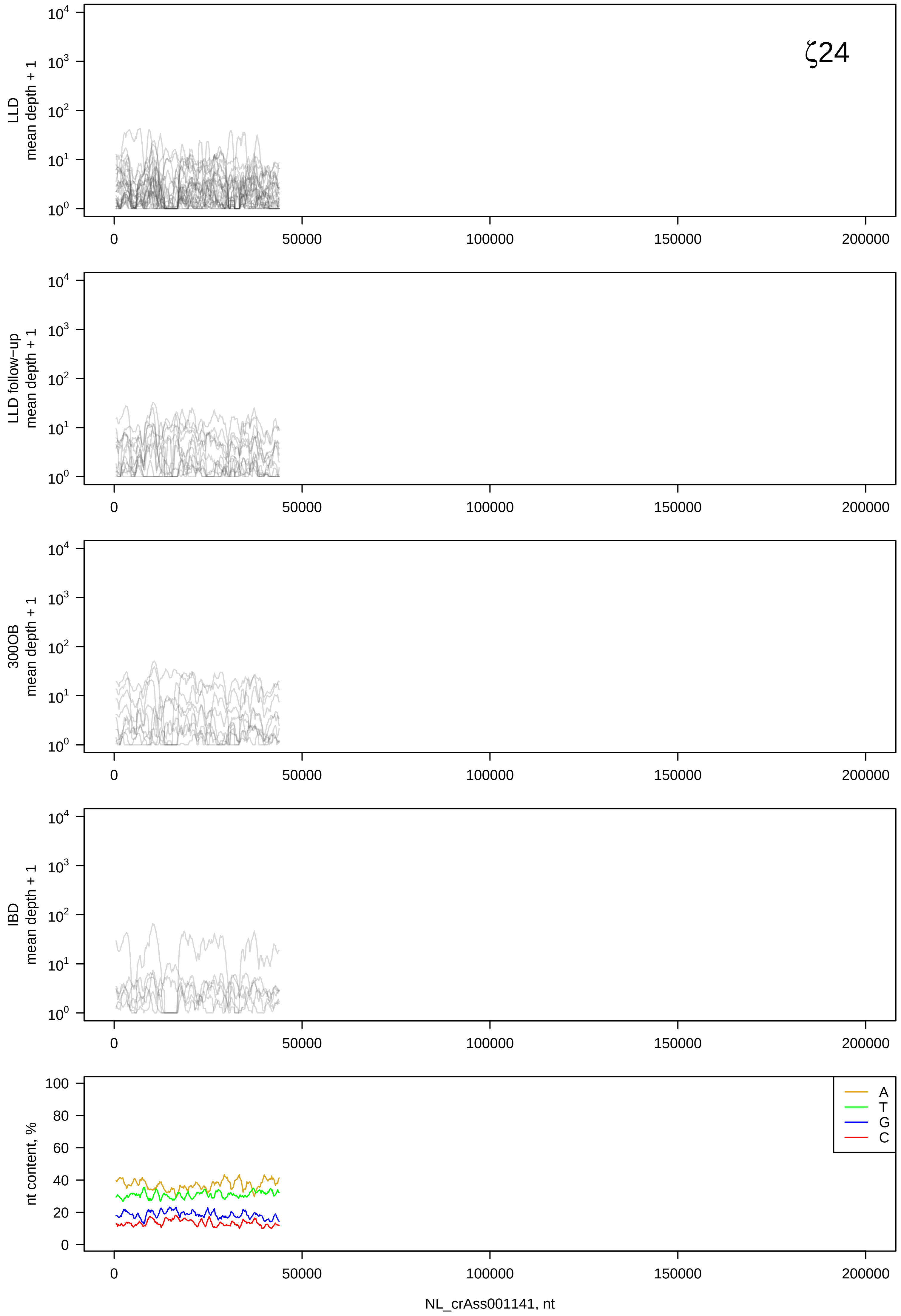

ζ24
LLD mean depth + 1
LLD follow−up mean depth + 1
300OB mean depth + 1
IBD mean depth + 1
nt content, %
A
T
G
C
0
50000
100000
150000
200000
NL_crAss001141, nt

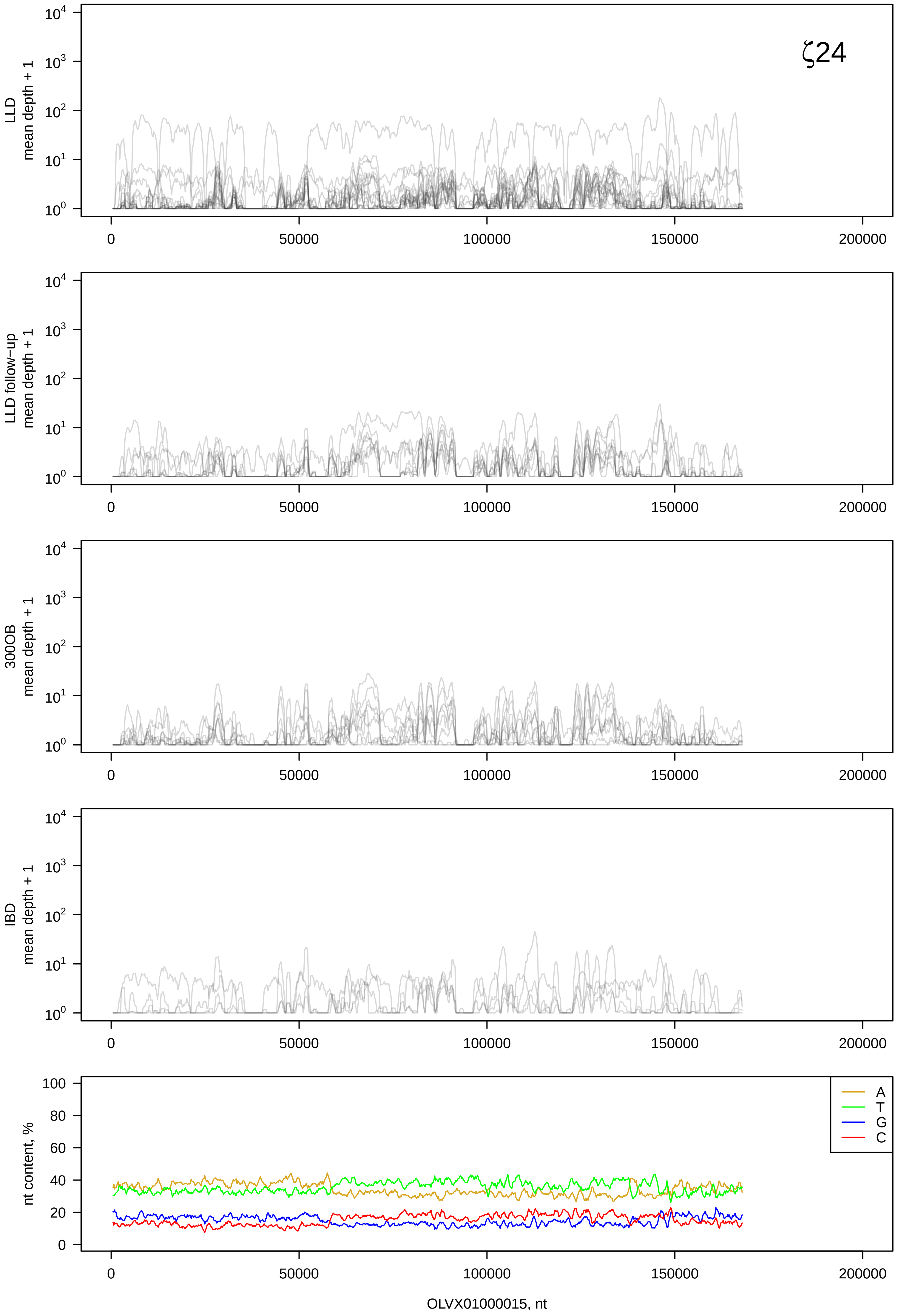

ζ24
LLD mean depth + 1
LLD follow-up mean depth + 1
300OB mean depth + 1
IBD mean depth + 1
nt content, %
A
T
G
C
OLVX01000015, nt

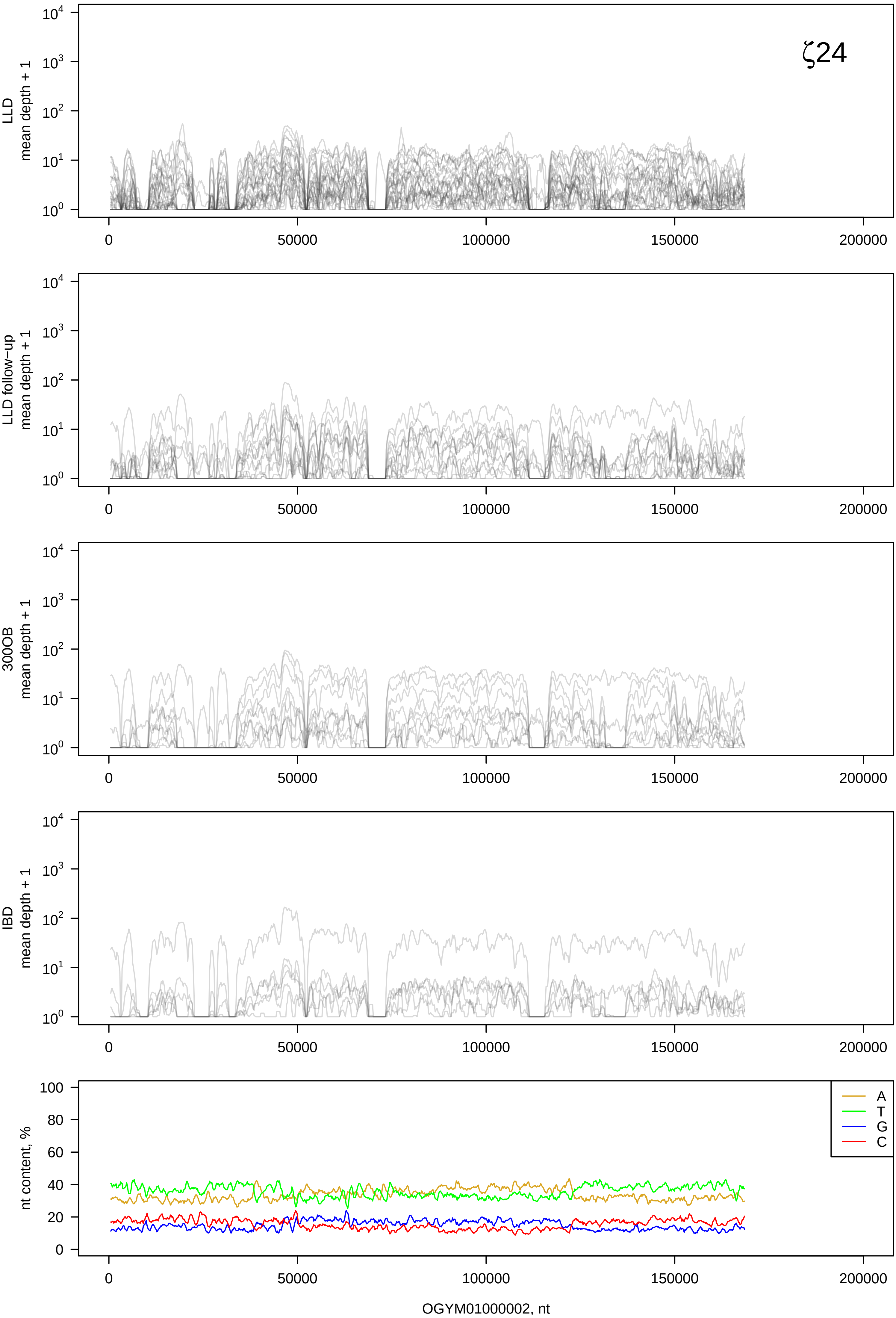

ζ24
LLD mean depth + 1
LLD follow−up mean depth + 1
300OB mean depth + 1
IBD mean depth + 1
nt content, %
A
T
G
C
0
50000
100000
150000
200000
OGYM01000002, nt

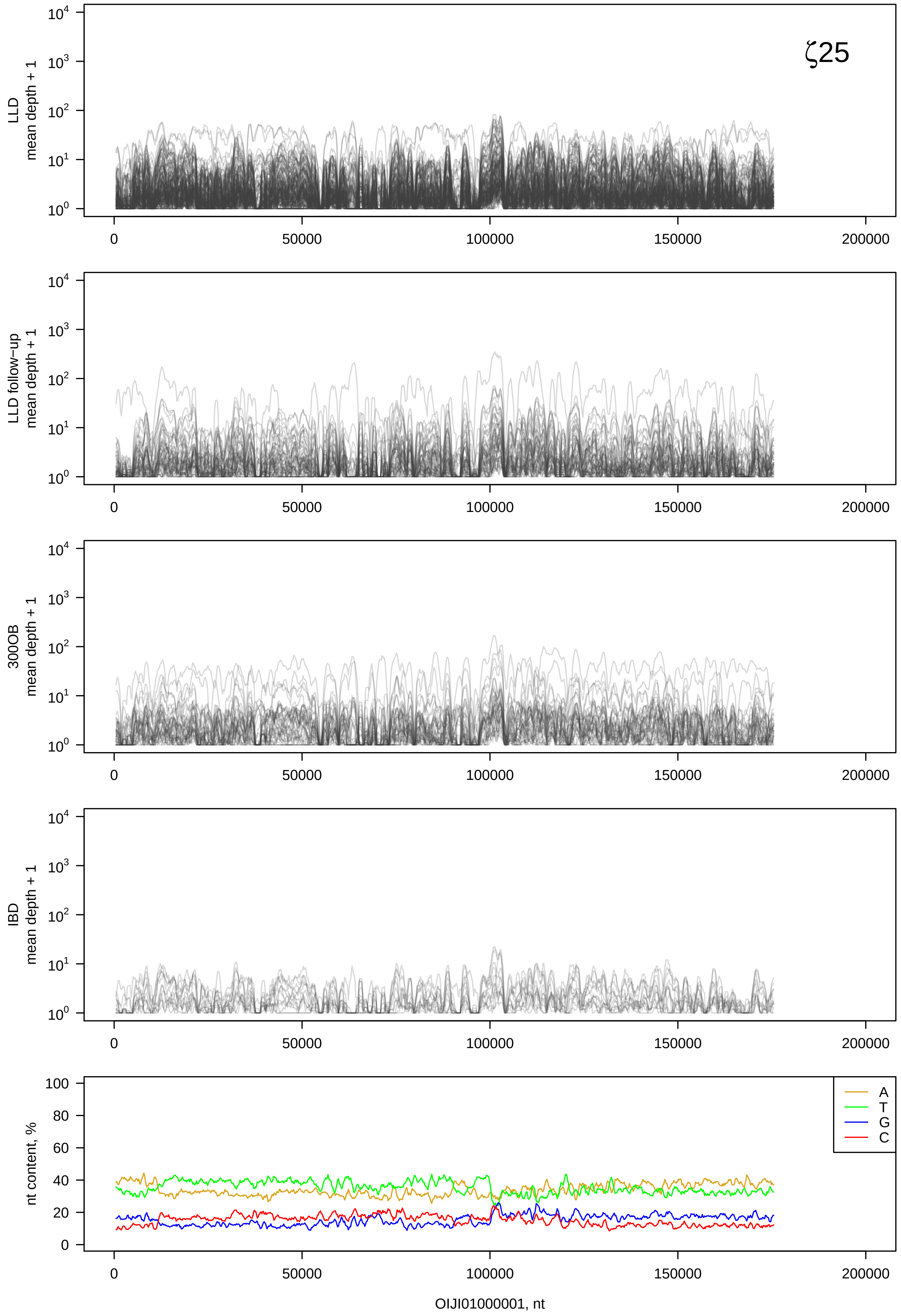
ζ25
LLD mean depth + 1
LLD follow−up mean depth + 1
300OB mean depth + 1
IBD mean depth + 1
nt content, %
A
T
G
C
OIJI01000001, nt

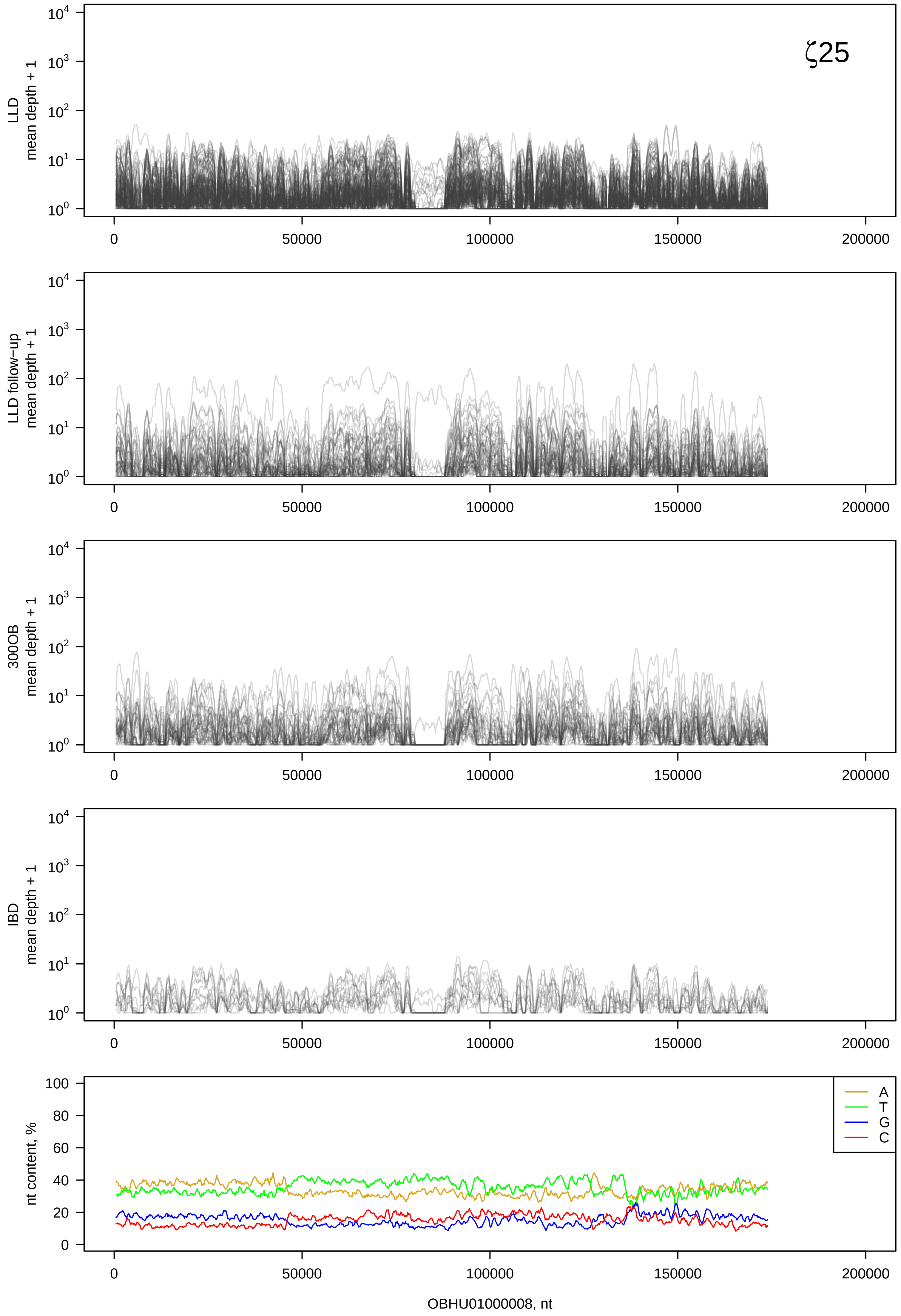
ζ25
LLD
mean depth + 1
LLD follow-up
mean depth + 1
300OB
mean depth + 1
IBD
mean depth + 1
nt content, %
A
T
G
C
0
50000
100000
150000
200000
OBHU01000008, nt

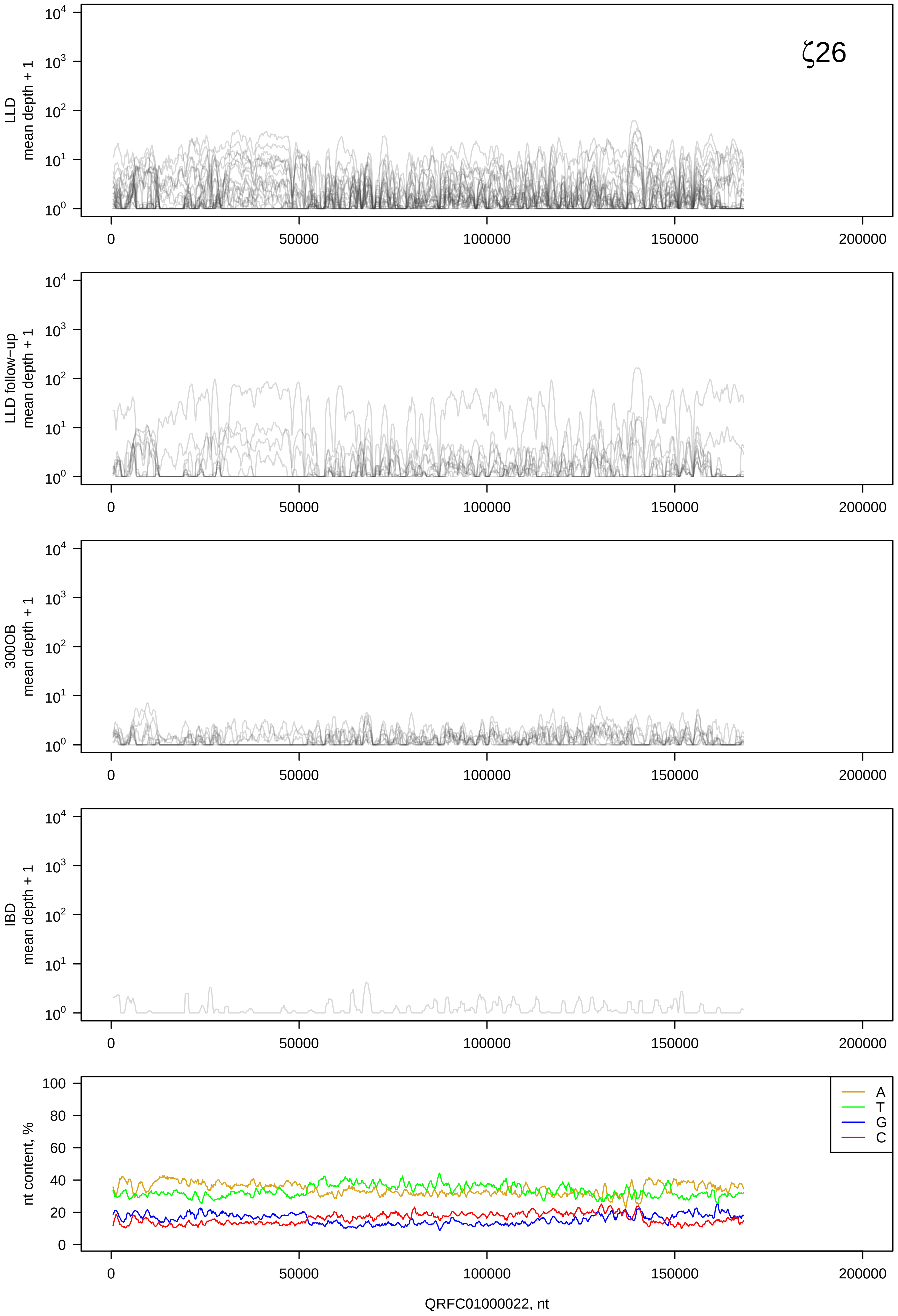

ζ26
LLD
mean depth + 1
LLD follow-up
mean depth + 1
300OB
mean depth + 1
IBD
mean depth + 1
nt content, %
A
T
G
C
0
50000
100000
150000
200000
QRFC01000022, nt

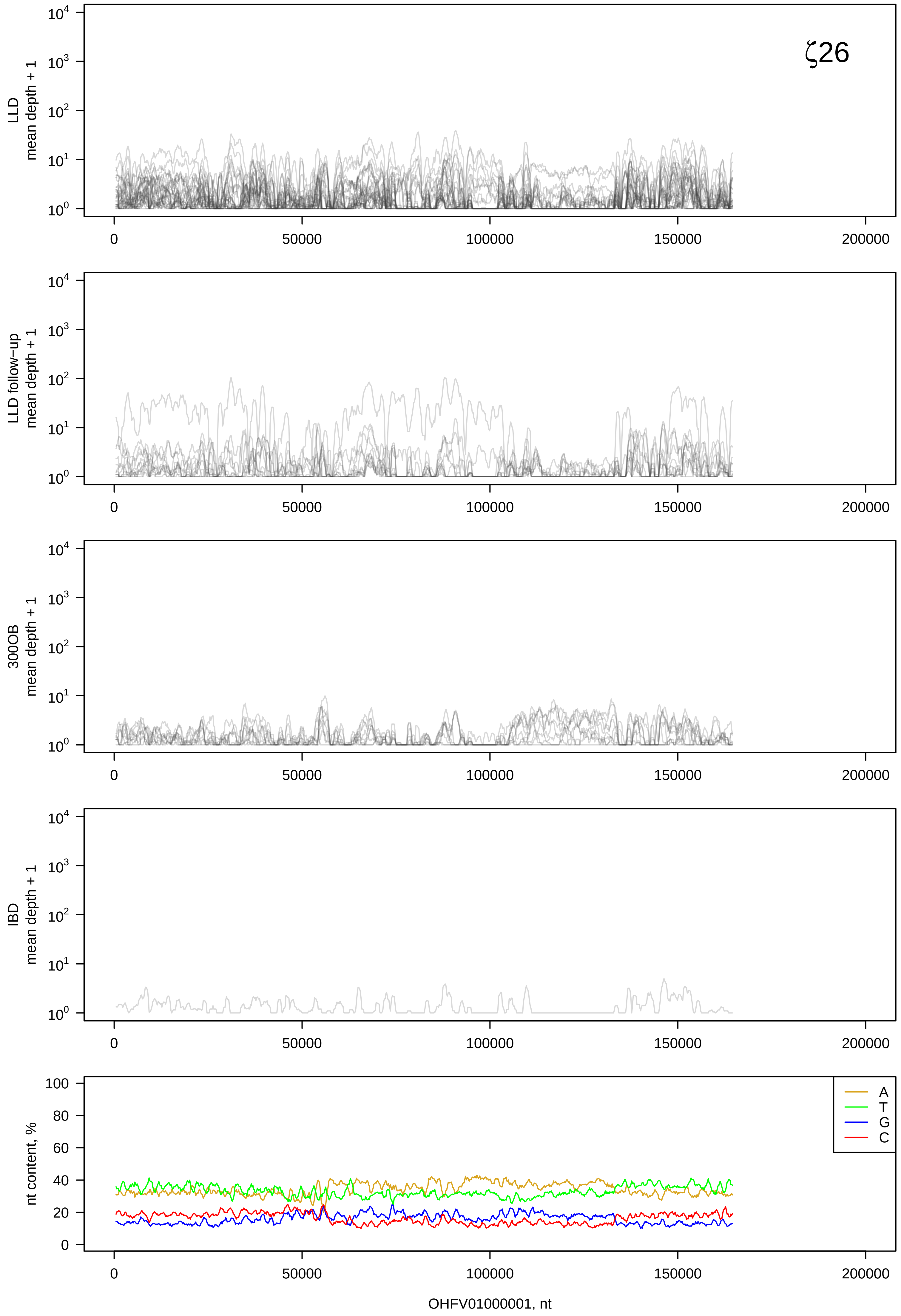
ζ26
LLD mean depth + 1
LLD follow-up mean depth + 1
300OB mean depth + 1
IBD mean depth + 1
nt content, %
A
T
G
C
OHFV01000001, nt

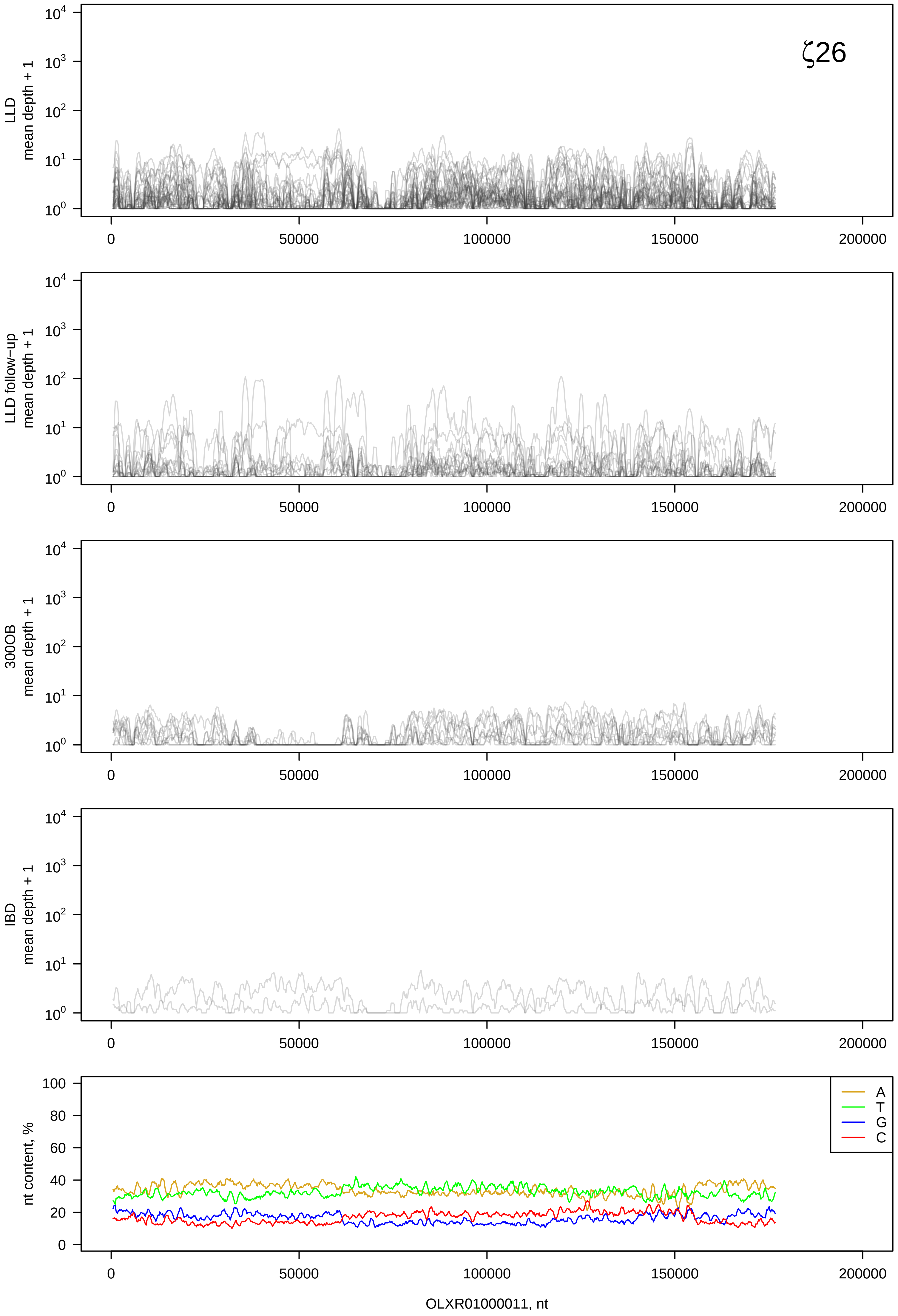

ζ26
LLD mean depth + 1
LLD follow-up mean depth + 1
300OB mean depth + 1
IBD mean depth + 1
nt content, %
A
T
G
C
0
50000
100000
150000
200000
OLXR01000011, nt

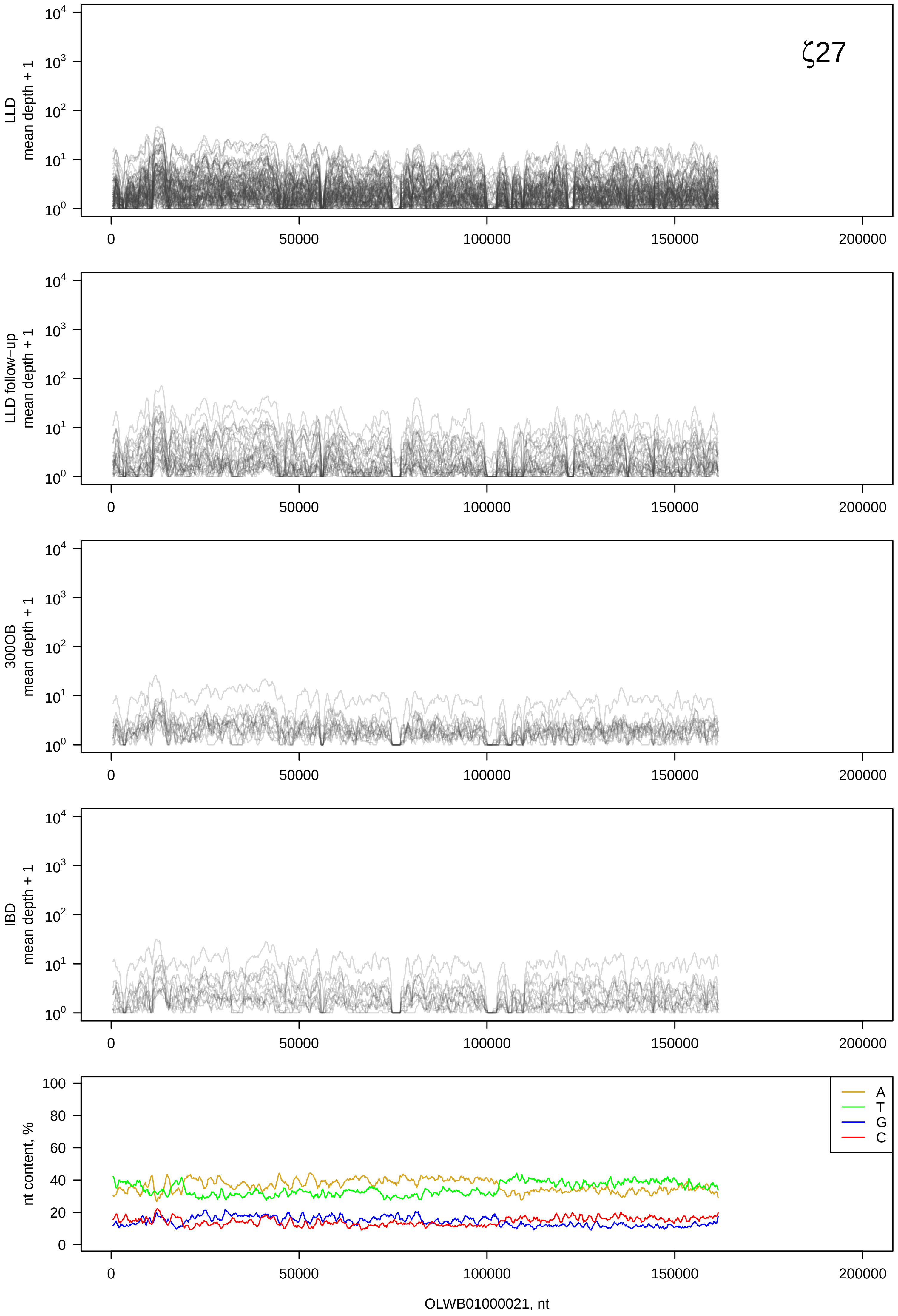

ζ27
LLD mean depth + 1
LLD follow−up mean depth + 1
300OB mean depth + 1
IBD mean depth + 1
nt content, %
A
T
G
C
OLWB01000021, nt

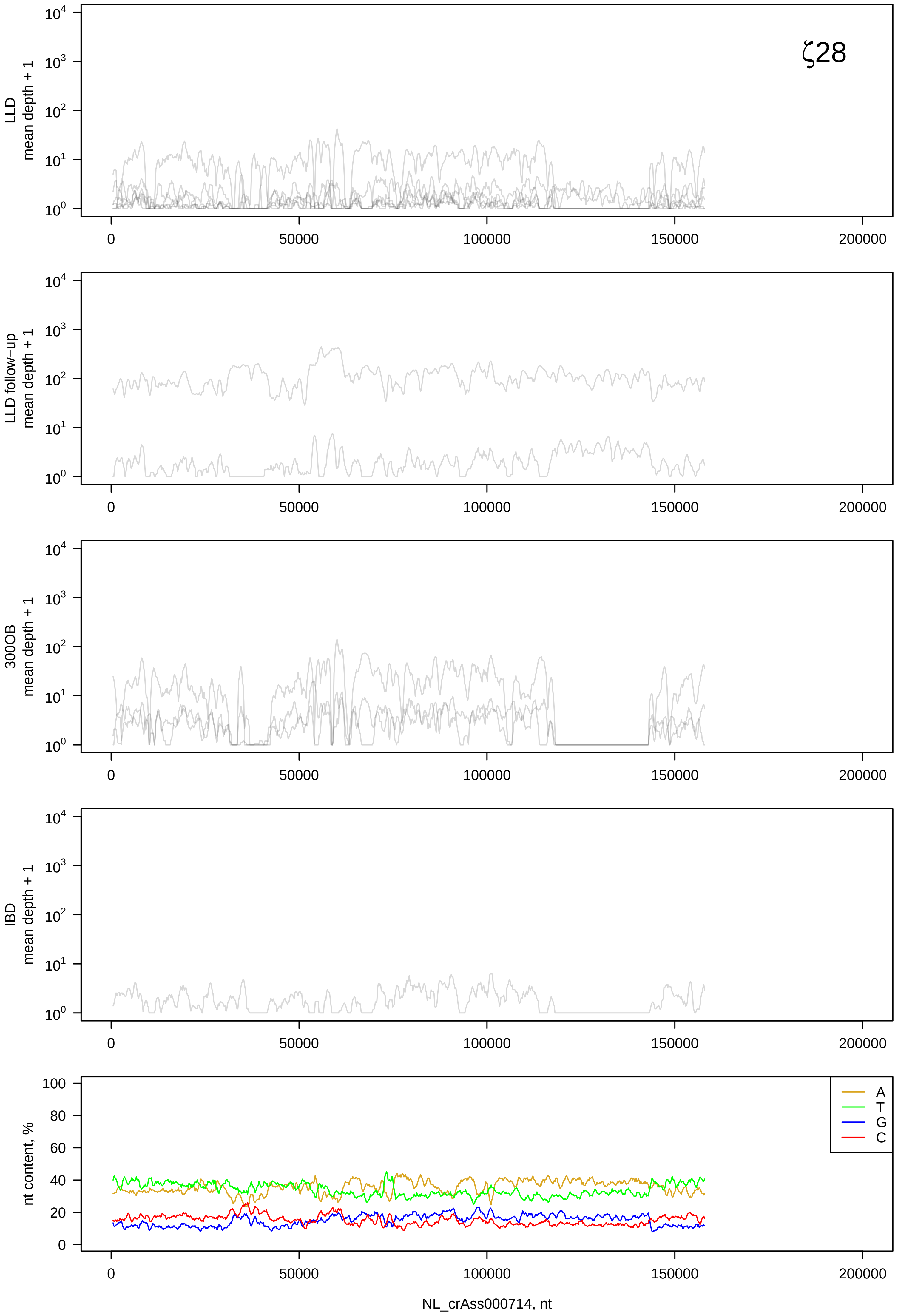

ζ28
LLD
mean depth + 1
LLD follow−up
mean depth + 1
300OB
mean depth + 1
IBD
mean depth + 1
nt content, %
A
T
G
C
NL_crAss000714, nt
0
50000
100000
150000
200000

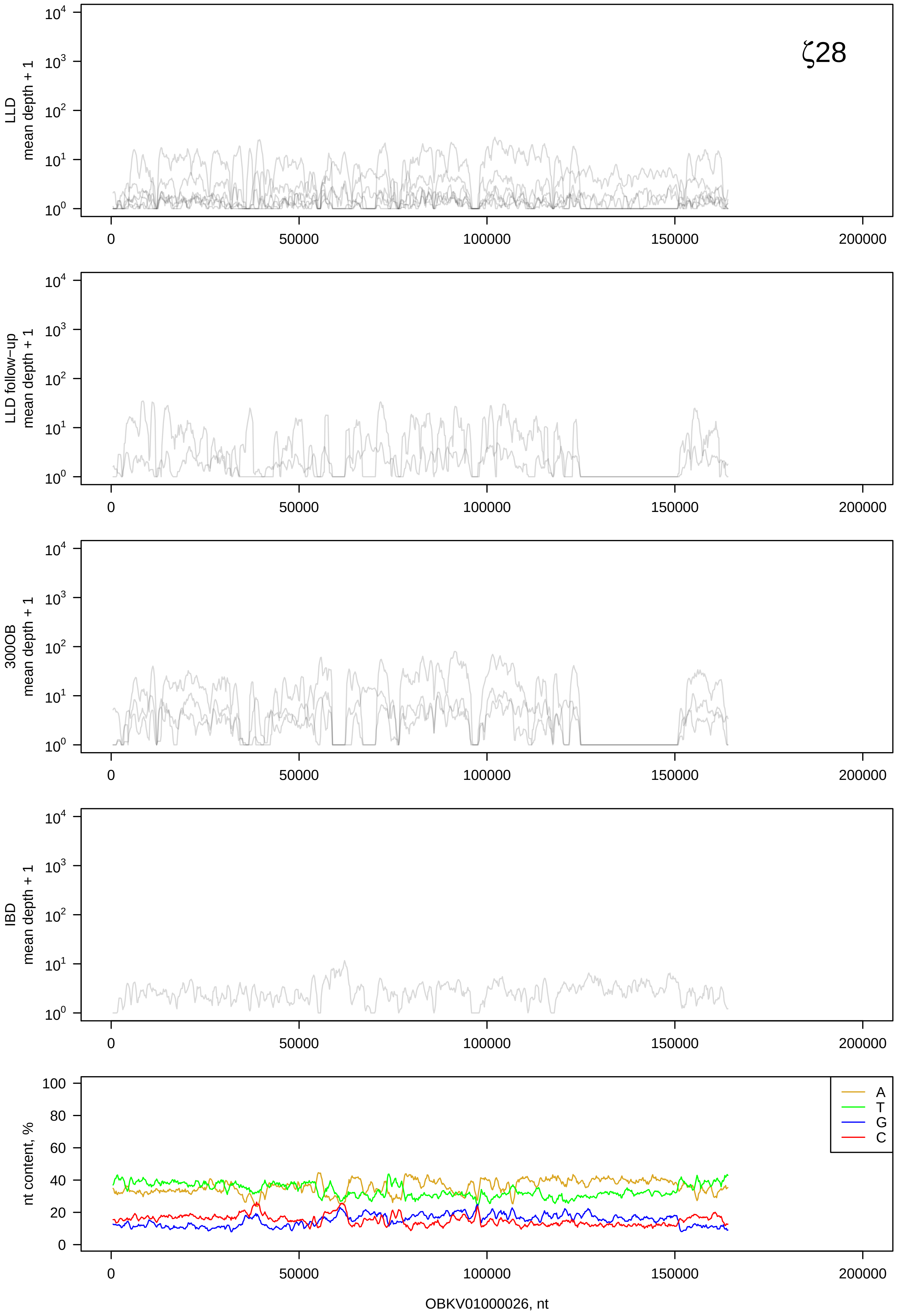

ζ28
LLD
mean depth + 1
LLD follow−up
mean depth + 1
300OB
mean depth + 1
IBD
mean depth + 1
$10^0$
$10^1$
$10^2$
$10^3$
$10^4$
0
50000
100000
150000
200000
nt content, %
100
80
60
40
20
0
A
T
G
C
OBKV01000026, nt

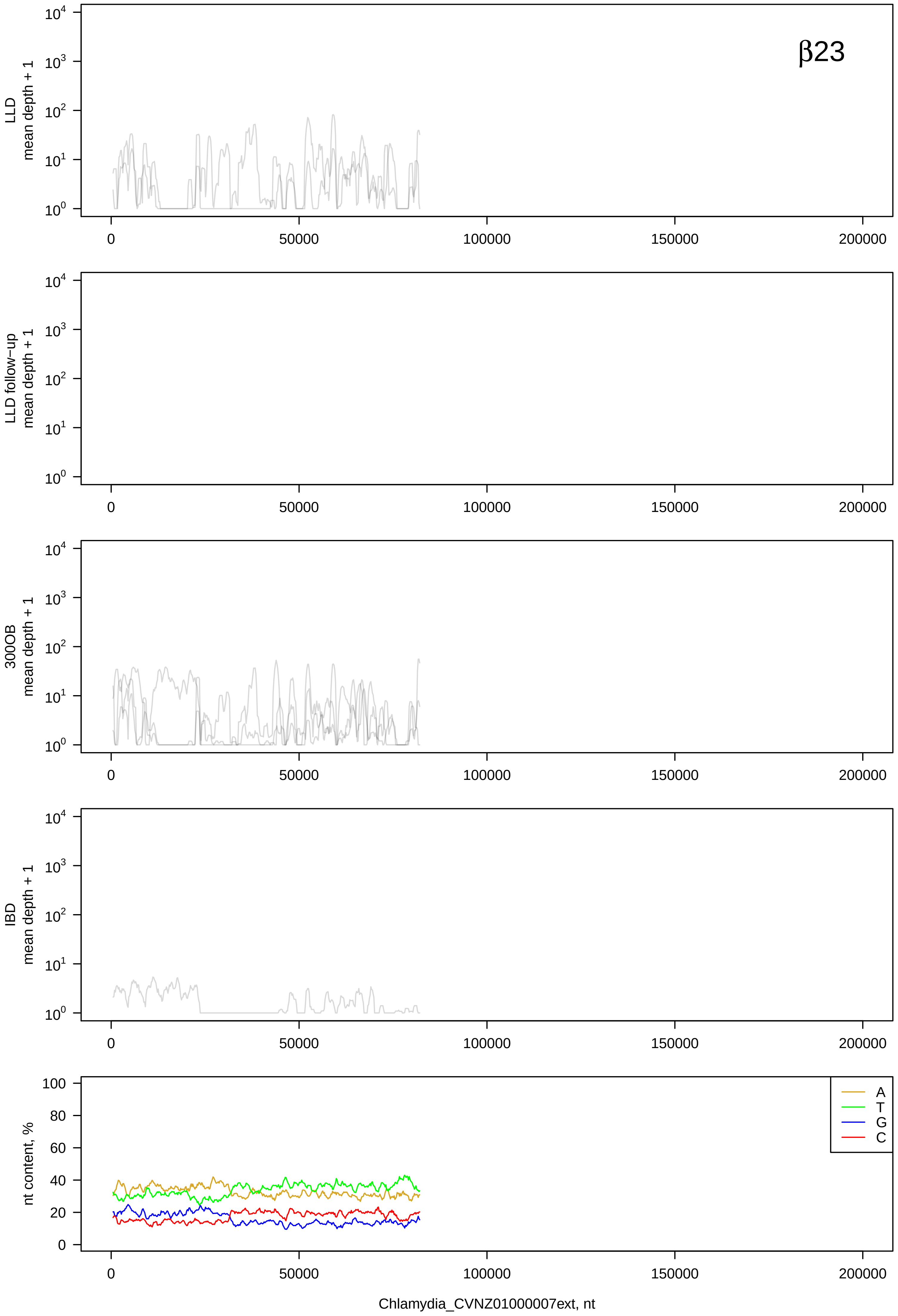

β23
LLD mean depth + 1
LLD follow−up mean depth + 1
300OB mean depth + 1
IBD mean depth + 1
nt content, %
A
T
G
C
0
50000
100000
150000
200000
Chlamydia_CVNZ01000007ext, nt

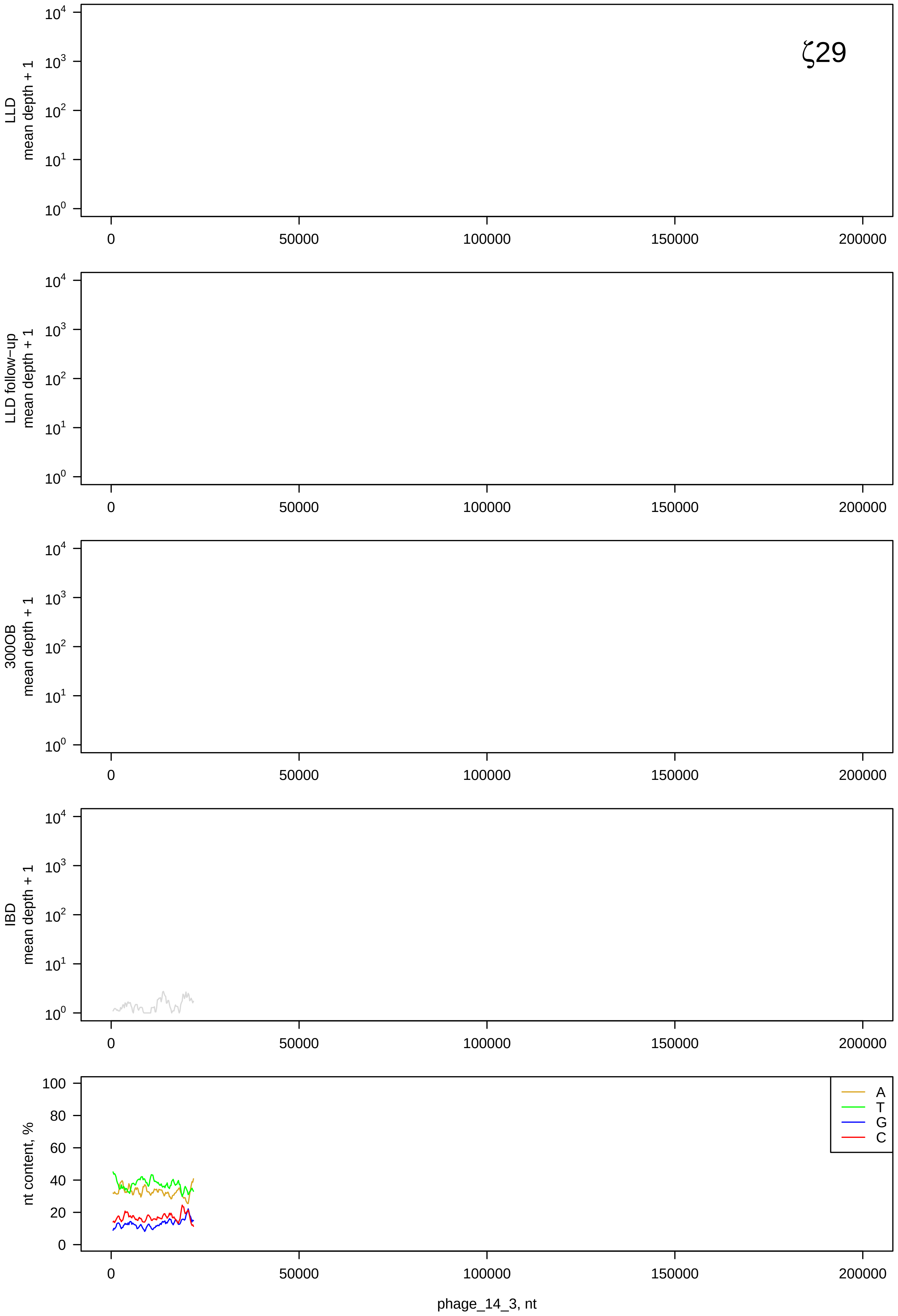
ζ29
LLD
mean depth + 1
LLD follow−up
mean depth + 1
300OB
mean depth + 1
IBD
mean depth + 1
nt content, %
A
T
G
C
0
50000
100000
150000
200000
phage_14_3, nt

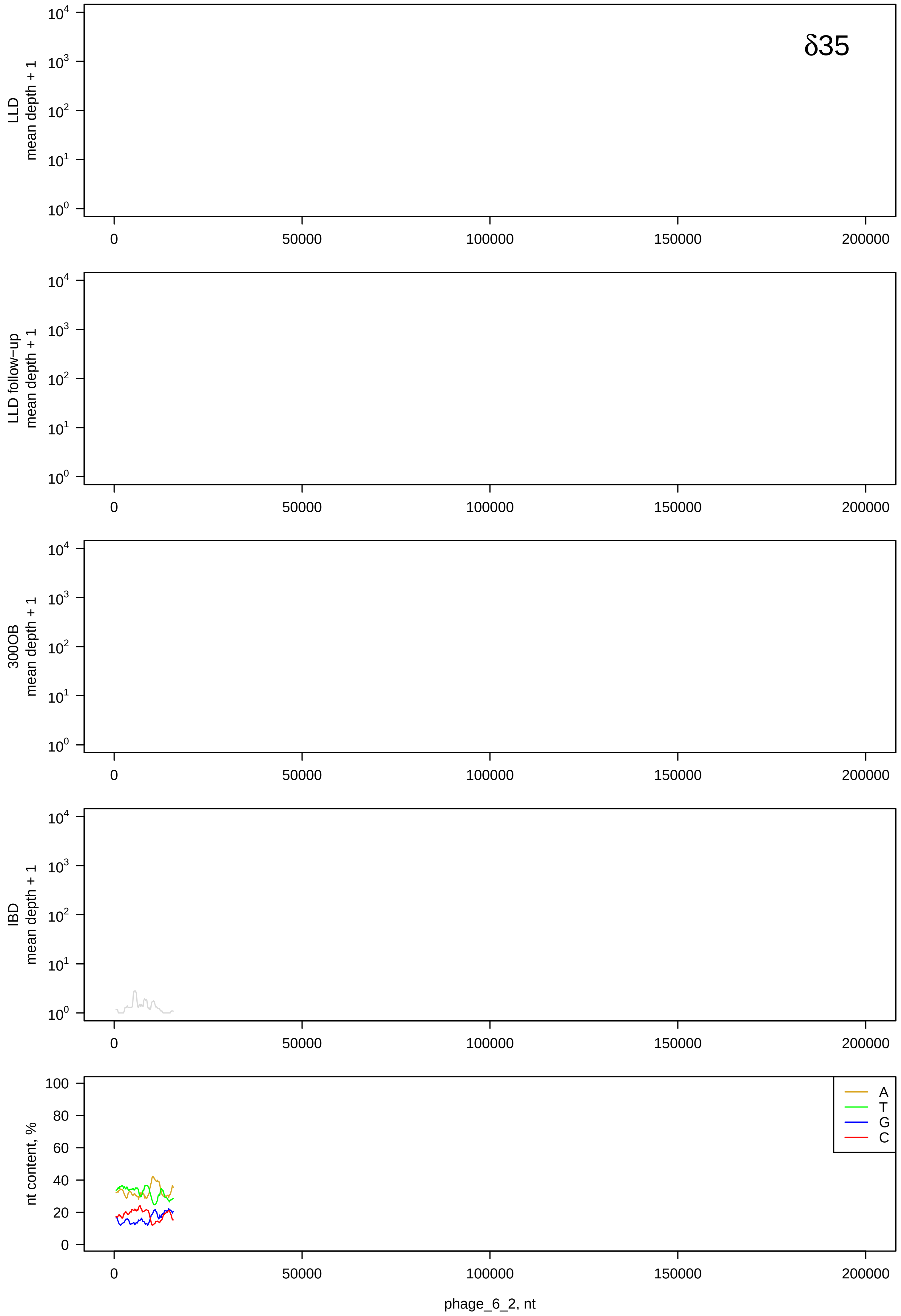

δ35
LLD mean depth + 1
LLD follow−up mean depth + 1
300OB mean depth + 1
IBD mean depth + 1
nt content, %
A
T
G
C
0
50000
100000
150000
200000
phage_6_2, nt

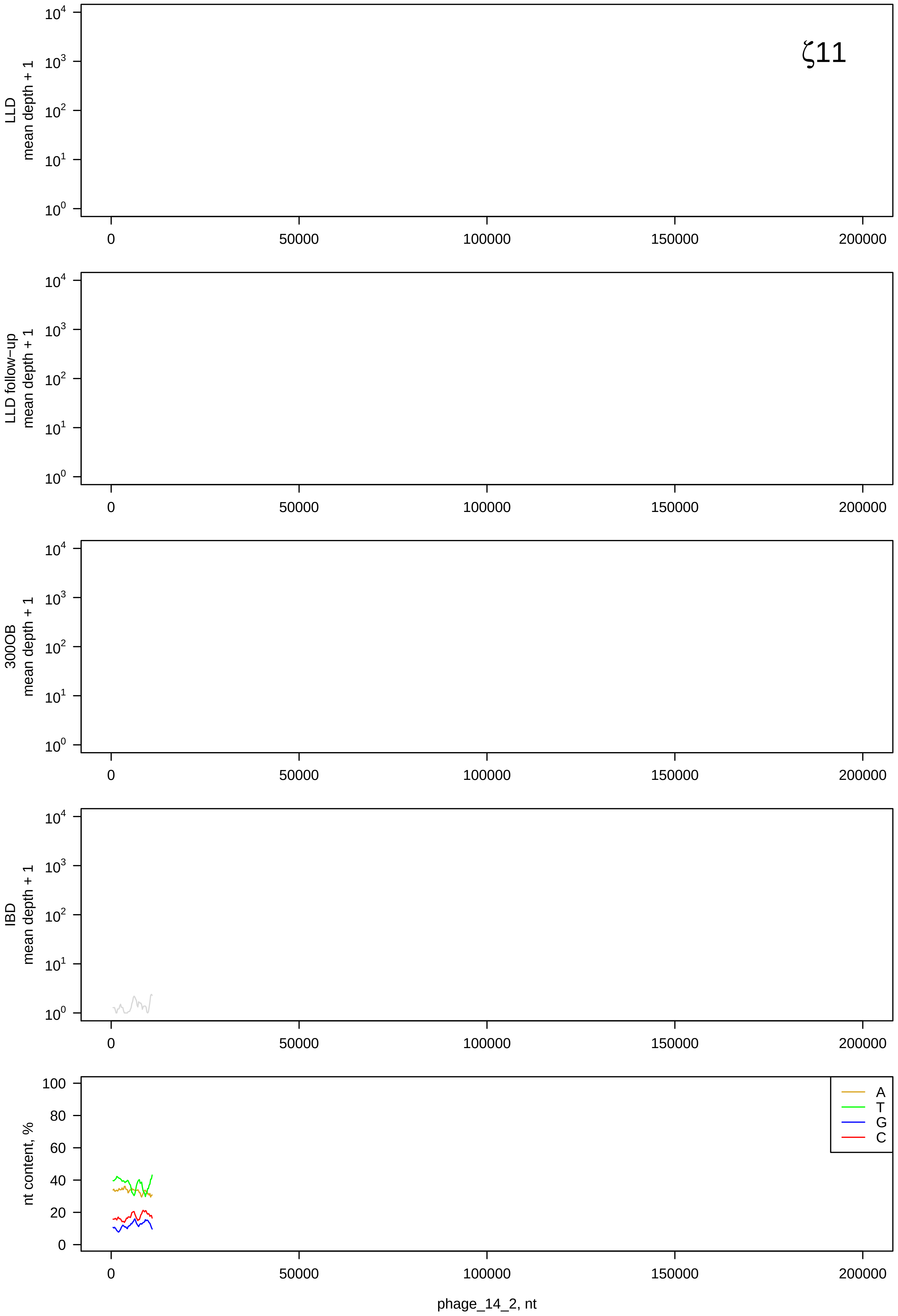
ζ11
LLD
mean depth + 1
LLD follow−up
mean depth + 1
300OB
mean depth + 1
IBD
mean depth + 1
nt content, %
A
T
G
C
phage_14_2, nt
